# Comparative pathway enrichment analysis in gastrointestinal cell lines Caco-2, HT-29, HEPG2, and colon fibroblasts using a custom expression panel for tight-junction and cytoskeletal regulatory genes

**DOI:** 10.1101/747105

**Authors:** JM Robinson

## Abstract

This brief report details results from a comparative analysis of Nanostring expression data between cell lines HEPG2, Caco-2, HT-29, and colon fibroblasts. Raw and normalized data are available publicly in the NCBI GEO/Bioproject databases. Results identify cell-line specific variations in gene expression relevant to intestinal epithelial function.

---

Cell lines provide invaluable *in vitro* research models for preclinical drug development and basic research into mechanisms of gastrointestinal and liver function. Standard intestinal epithelial models include two colorectal adenocarcinoma-derived lines, Caco-2 (ATCC® HTB-37™) and HT-29 (ATCC® HTB-38™), and the hepatocellular adenocarcinoma-derived HEPG2 (ATCC® HB-8065) cell line. Despite their shared origin from epithelial adenocarcinomatous tumors, each cell line possesses biological characteristics of its tissue of origin. Each cell line also possesses its own unique characteristics, therefore comparative data on cell line-specific molecular characteristics are important for generalizing experimental results to *in vivo* biology.

Some unique characteristics of these cell lines of interest include: Caco-2 forms polarized epithelia with a brush-border without specific induction; Caco-2 polarized monolayers have physiological barrier function as measured by trans-epithelial electrical resistance (TEER) [1]. HT-29 is characteristically known for mucous secretion and is therefore bears similarities with epithelial goblet cells. HT-29 polarization can be induced, and these cells have often also been used for differentiation studies [2]. Caco-2 and HT-29 have also been developed as a co-culture system, providing a more complex research model of intestinal epithelia [3]. HEPG2 possesses many characteristics of differentiated hepatocytes, and is a standard *in vitro* liver model for the biology of hepatitis and metabolism in the liver. [4]

A significant volume of transcriptomic data (predominantly microarray, some RNA-seq datasets are also available) have been reported for these lines in NCBI’s Gene Expression Omnibus (GEO) database [5], which contains (as of July 2019) for **HEPG2**: 1,253 series/11,178 samples; for **HT-29**: 224 series/1,466 samples; for **Caco-2**: 190 series/2,156 samples. Although other fibroblast lines are widely utilized by comparison, the colon fibroblast line **CCD-18Co** (ATCC® CRL-1459™) line contains 6 series/37 samples.

Robinson et al. (2019) recently reported results from a gene expression system utilizing a custom designed, 250-plex Nanostring codeset [6], performing expression profiling Caco-2 cultures treated with glucocorticoid hormone Dexamethasone (DEX) on a 30-day timecourse, and established DEX as having effects on epithelial-mesenchymal transition (EMT)-associated genes. Functional-morphological shifts between epithelial and mesenchymal cellular phenotypes underlie many important contexts of cellular differentiation, including embryonic development, organogenesis, tumorigenesis, and molecular mechanisms of EMT deeply conserved in the Metazoa [7].

For the analysis reported here, the full expression data and metadata for this experiment is publicly available in the NCBI Bioproject (accession#: PRJNA525237) and NCBI GEO (accession#: GSE132501) databases. Differential expression and and pathway enrichment analyses were performed using the Nanostring nSolver Advanced Analysis Module 2.0, available free of charge: (https://www.nanostring.com/products/analysis-software/advanced-analysis?jumpto=nsolver_advanced_analysis_getting_started) (Nanostring MAN-10030-03).

Previous report of this comparative expression data for Caco-2, HT-29, HEPG2, and CCD-18Co was reported in a conference proceeding [8]. For this analysis, expression data and metadata used are publicly available under the accessions listed above. In this analysis, days-post-seeding was used as a as a continuous co-variate in a generalized linear model (GLM), with the categorical “Cell Line” as a co-variate. Pathway scoring, differential expression (DE), and gene set enrichment (GSE) analyses of the data were performed using the Nanostring nSolver 4.0 with Advanced Analysis 2.0 plugin. ‘Pathway scores’ are determined relative to the normalized linear global average. DE and GSE results are relative to fibroblasts used as a reference category. Results presented here for each cell type should therefore to be interpreted as relative to expression in the fibroblast line, fibroblast effectively serve as an ‘outgroup’ for comparison to the cell lines with epithelial origin. The full results are provided as supplementary material. (**Sup. Results 1: HTML-formatted complete Nanostring Advanced Analysis Module results**).

Unsupervised, principle components analysis (PCA) results show that cell-type specific signal within the data represents the strongest DE signals. Hierarchical clustering analysis of normalized counts perfectly resolves cell lines, regardless of early or later timepoints (Fig. 1A,B). PCA results show that differences between cell types *and* time-associated effects explain 93% of variation in the overall expression data (Fig. 1B). *Generally*, **PC1** resolves between the 4 cell lines while highlighting <u>similarities between HEPG2 and Caco-2</u>. **PC2** resolves between the 4 lines but highlights <u>divergent features of Caco-2</u> vs. the other three cell lines. **PC3**, although determining less overall similarity between cell lines, highlights <u>divergent features of the CCD-18Co fibroblast line</u>. **PC4** resolves effects of earlier vs. later stages on the timecourse (Caco-2 line has several replicate samples at Day 1 and Day 10, while samples from other lines are from culture days 2-4 only).

**Figure 1.**
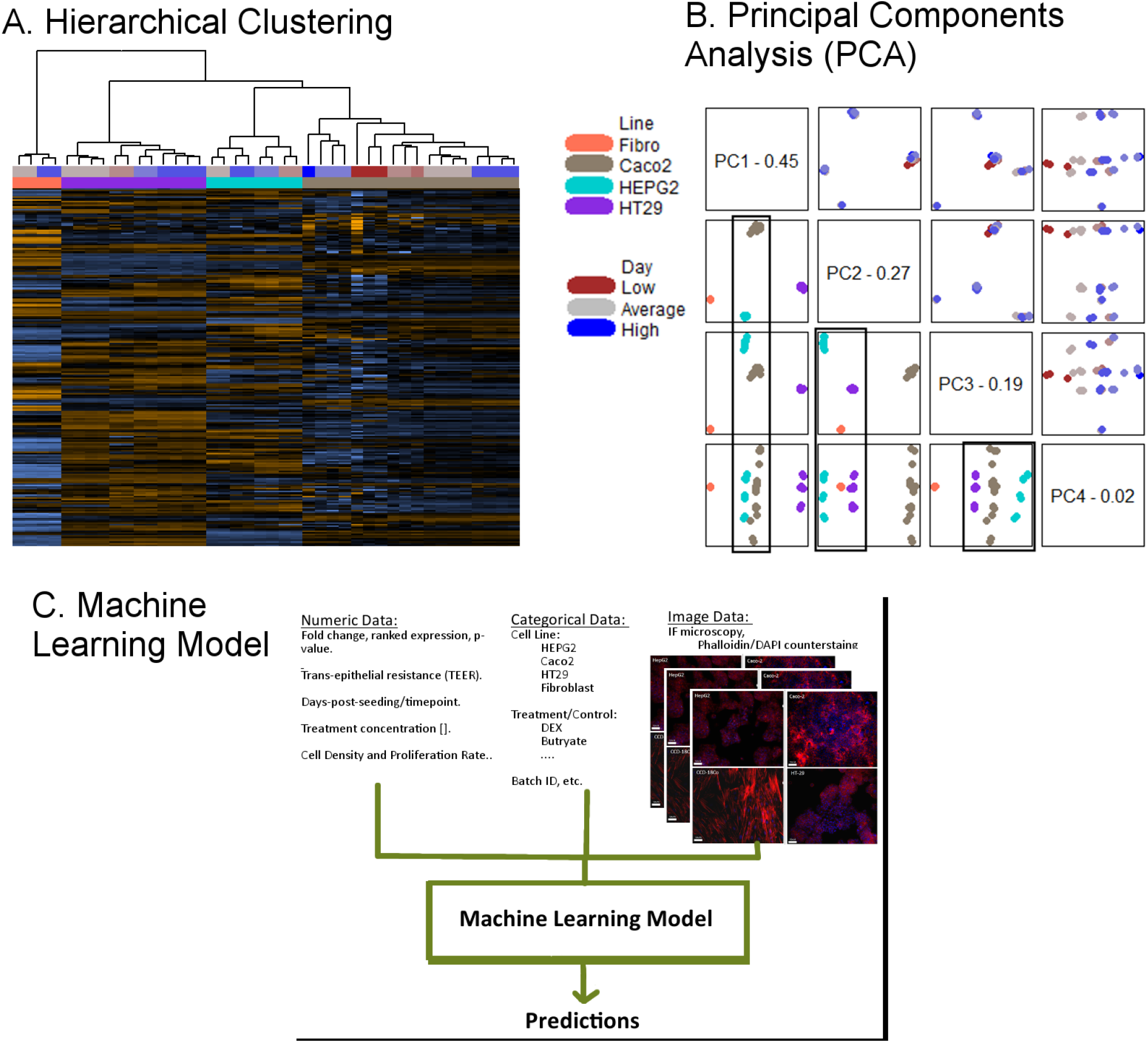
Comparative Results by Cell Type

Observed enrichment of pathway scores in the fibroblast line were consistent with the well-known mechanisms for regulating cytoskeletal dynamics in establishing epithelial identity [9]. For example, in this dataset, fibroblasts showed high scores for “RHO GTPases Activate Formins” and “TGF-beta receptor signaling activates SMADs”, and had low scores for “Tight junction interactions” and “RHO GTPase Activate ROCKs” compared with the other cell lines (Fig.2A).

**Figure 2.**
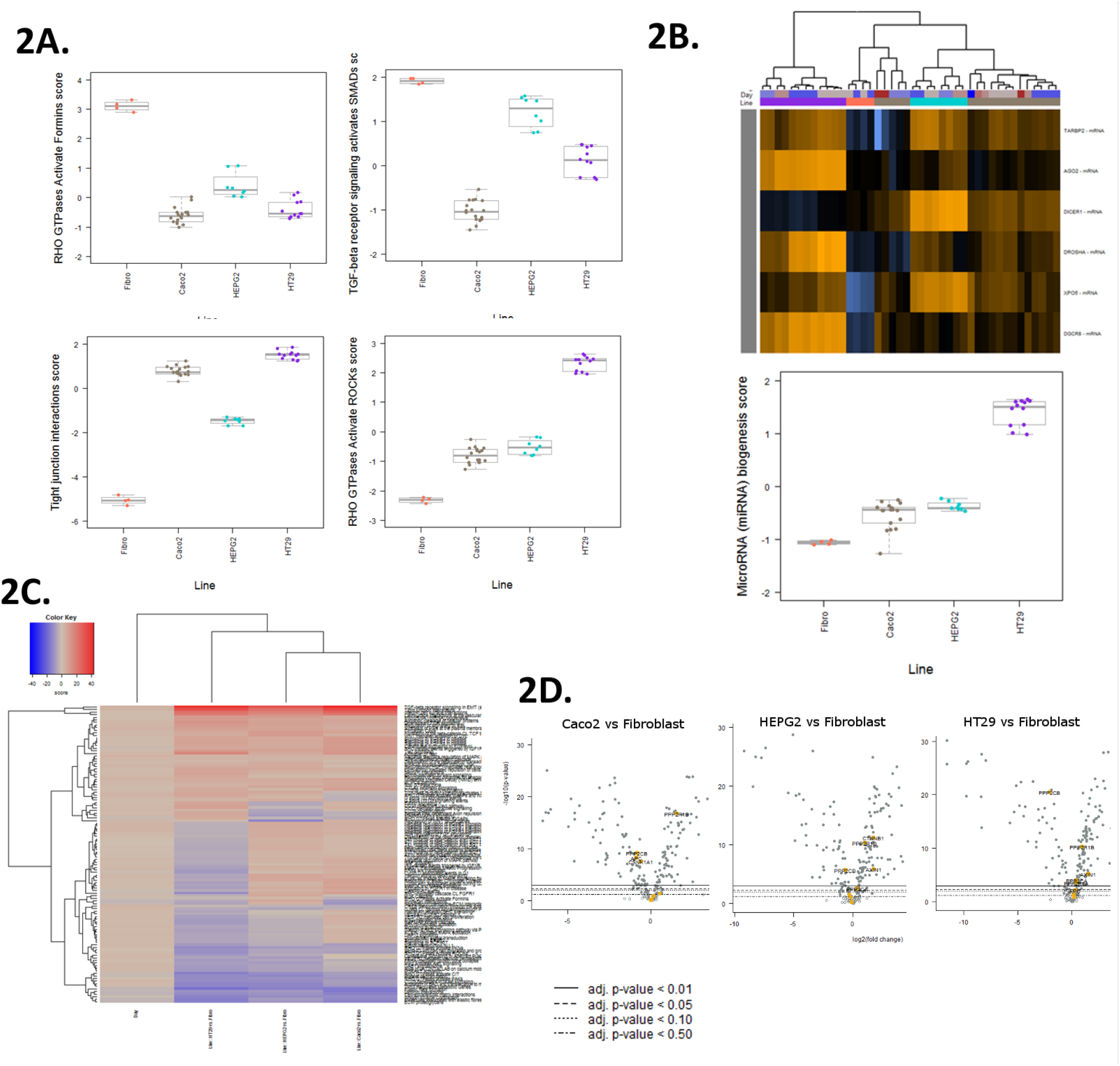
Cell Type-Associated Pathway Enrichment

Cell line-specific pathway enrichments were observed for each respective cell line. HT-29 often had divergent pathway scores vs. other cell lines, reflective of its “separateness” in PC1. “microRNA (miRNA) biogenesis” is another such pathway of interest, where HT-29 showed increased expression of Argonaute 2 (AGO2), DROSHA, and DGCR8 (Pasha), three of the key genes in the microRNA biogenesis pathway (Fig. 2B).

GSE analysis reveals relative enrichment of pathways associated with each “Cell Line vs. Fibroblast” comparison (Fig. 2C). GSE is a tools which helps to readily identify pathway-relevant genes by selecting variations in expression underlying significant GSE scores. For example, the “Beta-catenin phosphorylation cascade” shows −5.3 ratio in Caco-2 vs. fibroblasts, compared with both HT-29 and HEPG2, which have positive scores vs. fibroblasts (**Sup. Data 2: GSE Scores**), highlighting putative Caco2-specific responsiveness. The source of these differential scores are can be immediately traced to the significantly decreased expression of APC and CSNK1A1 in Caco-2, but not in HT-29 or HEPG2 (Fig. 2D).

Advanced machine learning data models have been proposed, which utilize diverse phenotypic data types such as immunofluorescent phalloidin staining [10]. Machine-learning methods for integrative data analysis have been proposed; an adaptation of such a model for this data, could include expression data, with a companion datastream of immunofluorescence microcopy images. In this model, system would include DE and DE-derived pathway quantifications, and immunoflurescent images. (Fig. 1C, after Rosebrock (2019) (https://www.pyimagesearch.com/2019/02/04/keras-multiple-inputs-and-mixed-data/)).

## Supporting information

Sup. Results 1

