## Supplementary material for "Comparative pathway enrichment analysis in gastrointestinal cell lines Caco-2, HT-29, HEPG2, and colon fibroblasts using a custom expression panel for tight-junction and cytoskeletal regulatory genes": Sup. Results 1: AdvAnalysisReport.html

          
   
     
     
     
     nCounter Advanced Analysis Report 
     
     
     
     
     
     
     
     
     
     
     
     
     
     
     
     
     
     
     
     
     
     
     
     
     
     
     
     
     
     
     
     
     
     
     
     
     
     
     
   
   
     
       
         
           
             
               
             
           
         
       
       
         
      
          Overview  
          Normalization  
          Pathway Score  
          Diff Expr  
          GSA  
          PathView  
          Analysis Parameters  
      
      Share   
       
     
     
      
        
          
                This module generates a series of high level plots that describe the data overall and may be useful for identifying anomalous data and/or covariates.     
              
                   Gene sets: 
                  
                      Summary  
                      Activation of BAD and translocation to mitochondria   
                      Activation of Rac  
                      Apoptotic cleavage of cellular proteins  
                      ARMS-mediated activation  
                      AXIN missense mutants destabilize the destruction complex  
                      Beta-catenin phosphorylation cascade  
                      CD209 (DC-SIGN) signaling  
                      CD28 co-stimulation  
                      CD28 dependent PI3KAkt signaling  
                      CD28 dependent Vav1 pathway  
                      Cell-extracellular matrix interactions  
                      Cell surface interactions at the vascular wall  
                      Circadian Clock  
                      Constitutive Signaling by Aberrant PI3K in Cancer  
                      Constitutive Signaling by EGFRvIII  
                      CREB phosphorylation through the activation of Ras  
                      CTLA4 inhibitory signaling  
                      Cyclin AB1 associated events during G2M transition  
                      Cyclin D associated events in G1  
                      DAP12 signaling  
                      DARPP-32 events  
                      DCC mediated attractive signaling  
                      Deactivation of the beta-catenin transactivating complex  
                      Disassembly of the destruction complex and recruitment of AXIN t...  
                      Downstream signal transduction  
                      Downstream TCR signaling  
                      ECM proteoglycans  
                      Elastic fibre formation  
                      EPH-ephrin mediated repulsion of cells  
                      EPHA-mediated growth cone collapse  
                      EPHB-mediated forward signaling  
                      Ephrin signaling  
                      ERKs are inactivated  
                      FCERI mediated MAPK activation  
                      Formation of the beta-catenin.CL.TCF transactivating complex  
                      Frs2-mediated activation  
                      G alpha (1213) signalling events  
                      G alpha (q) signalling events  
                      G beta.CL.gamma signalling through PI3Kgamma  
                      GAB1 signalosome  
                      Gastrin-CREB signalling pathway via PKC and MAPK  
                      GPVI-mediated activation cascade  
                      Integrin alphaIIb beta3 signaling  
                      Integrin cell surface interactions  
                      Interleukin receptor SHC signaling  
                      IRS-related events triggered by IGF1R  
                      Laminin interactions  
                      MAP2K and MAPK activation  
                      MAPK6MAPK4 signaling  
                      MASTL Facilitates Mitotic Progression  
                      MicroRNA (miRNA) biogenesis  
                      Misspliced GSK3beta mutants stabilize beta-catenin  
                      Molecules associated with elastic fibres  
                      Negative feedback regulation of MAPK pathway  
                      Negative regulation of FGFR1 signaling  
                      Negative regulation of FGFR2 signaling  
                      Negative regulation of FGFR3 signaling  
                      Negative regulation of FGFR4 signaling  
                      Negative regulation of MAPK pathway  
                      Nephrin interactions  
                      Non-integrin membrane-ECM interactions  
                      Nonsense Mediated Decay (NMD) enhanced by the Exon Junction Comp...  
                      NRAGE signals death through JNK  
                      p130Cas linkage to MAPK signaling for integrins  
                      PCPCE pathway  
                      PI3K Cascade  
                      PI3KAKT activation  
                      PIP3 activates AKT signaling  
                      Platelet degranulation   
                      RAF activation  
                      RAFMAP kinase cascade  
                      Regulation of actin dynamics for phagocytic cup formation  
                      Regulation of HSF1-mediated heat shock response  
                      Regulation of signaling by CBL  
                      Repression of WNT target genes  
                      Resolution of Sister Chromatid Cohesion  
                      Rho GTPase cycle  
                      RHO GTPases activate CIT  
                      RHO GTPases Activate Formins  
                      RHO GTPases activate IQGAPs  
                      RHO GTPases activate PAKs  
                      RHO GTPases activate PKNs  
                      RHO GTPases Activate ROCKs  
                      RHO GTPases Activate WASPs and WAVEs  
                      Role of LAT2NTALLAB on calcium mobilization  
                      S33 mutants of beta-catenin aren.SQT.t phosphorylated  
                      S37 mutants of beta-catenin aren.SQT.t phosphorylated  
                      S45 mutants of beta-catenin aren.SQT.t phosphorylated  
                      Sema3A PAK dependent Axon repulsion  
                      Sema4D induced cell migration and growth-cone collapse  
                      SHC-mediated cascade.CL.FGFR1  
                      SHC-related events triggered by IGF1R  
                      Signal transduction by L1  
                      Signaling by ERBB2  
                      Signaling by FGFR1 in disease  
                      Signaling by FGFR2 in disease  
                      Signaling by FGFR3 in disease  
                      Signaling by FGFR4 in disease  
                      Signaling by PDGF  
                      Signaling by SCF-KIT  
                      Smooth Muscle Contraction  
                      Spry regulation of FGF signaling  
                      Syndecan interactions  
                      Synthesis of PIPs at the plasma membrane  
                      T41 mutants of beta-catenin aren.SQT.t phosphorylated  
                      TGF-beta receptor signaling activates SMADs  
                      TGF-beta receptor signaling in EMT (epithelial to mesenchymal tr...  
                      Tie2 Signaling  
                      Tight junction interactions  
                      TP53 Regulates Metabolic Genes  
                      Truncations of AMER1 destabilize the destruction complex  
                      VEGFA-VEGFR2 Pathway  
                      VEGFR2 mediated cell proliferation  
                      VEGFR2 mediated vascular permeability  
                      WNT mediated activation of DVL  
                  
                   
                  
                    
                      
                           
                          
                               Heatmaps    PCA    Study design    Other QC     
                              
                                  Heatmaps  
                              
                               
                              
                                  Line  
                                  Day  
                              
                               
                              
                                  Covariates  
                              
                               
                              
                                  Other QC  
                              
                              
                            
                              
                                
                                  
                                       
                                      
                                           Heatmap of Raw Data   More Plot Information      Download detected/undetected calls data                   Heatmap of Raw Data  Heatmap of the raw counts. The plot is meant to provide an overview of how robust the raw expression levels are across samples and gene sets. Datasets that entirely lack higher level expressions (e.g. counts &gt; 100) may indicate experimental failure or low input. The detected/undetected calls links to a .csv file stating whether each probe is above background, with 0/1 indicating below/above background. If the user has not specified a detection threshold, probes are called detected if they have more than double the counts of the median negative control.   
                                           Heatmap of All Data   More Plot Information                   Heatmap of All Data  Heatmap of the normalized data, scaled to give all genes equal variance, generated via unsupervised clustering. Orange indicates high expression; blue indicates low expression. This plot is meant to provide a high level exploratory view of the data.   
                                      
                                       
                                  
                                
                              
                            
                            
                              
                                
                                  
                                       
                                      
                                           Principal Components of All Data   More Plot Information                      Principal Components of All Data  Principal component analysis maps high-dimensional datasets onto a smaller number of highly informative dimensions. Here, the first four principal components of the gene expression data are plotted against each other and colored by the values of the selected covariate. This plot may be used to identify clusters in the data and to identify variables associated with prominent signal in the data. Variables that are associated with these leading principal components should be considered in downstream analyses.   
                                      
                                       
                                  
                                
                                
                                  
                                       
                                      
                                           Principal Components of All Data   More Plot Information                      Principal Components of All Data  Principal component analysis maps high-dimensional datasets onto a smaller number of highly informative dimensions. Here, the first four principal components of the gene expression data are plotted against each other and colored by the values of the selected covariate. This plot may be used to identify clusters in the data and to identify variables associated with prominent signal in the data. Variables that are associated with these leading principal components should be considered in downstream analyses.   
                                      
                                       
                                  
                                
                              
                            
                            
                              
                                
                                  
                                       
                                      
                                              More Plot Information                  Pairwise comparisons of all covariates in the analysis. The type of plot is dependent on the types of variables compared; A categorical vs. categorical covariate plot is shown as a bar chart of counts (Y axis). Continuous vs. categorical covariates generate a boxplot with whiskers denoting 1.5 IQR. Continuous vs. continuous covariates are compared via a scatter plot. Variables that are correlated with a biological variable of interest are potential confounders that may influence downstream analyses. Additionally, bar plots and histograms show the distributions of categorical and continuous variables, respectively.   
                                              More Plot Information                  Pairwise comparisons of all covariates in the analysis. The type of plot is dependent on the types of variables compared; A categorical vs. categorical covariate plot is shown as a bar chart of counts (Y axis). Continuous vs. categorical covariates generate a boxplot with whiskers denoting 1.5 IQR. Continuous vs. continuous covariates are compared via a scatter plot. Variables that are correlated with a biological variable of interest are potential confounders that may influence downstream analyses. Additionally, bar plots and histograms show the distributions of categorical and continuous variables, respectively.   
                                              More Plot Information                  Pairwise comparisons of all covariates in the analysis. The type of plot is dependent on the types of variables compared; A categorical vs. categorical covariate plot is shown as a bar chart of counts (Y axis). Continuous vs. categorical covariates generate a boxplot with whiskers denoting 1.5 IQR. Continuous vs. continuous covariates are compared via a scatter plot. Variables that are correlated with a biological variable of interest are potential confounders that may influence downstream analyses. Additionally, bar plots and histograms show the distributions of categorical and continuous variables, respectively.   
                                      
                                       
                                  
                                
                              
                            
                            
                              
                                
                                  
                                       
                                      
                                           Variance vs. Mean normalized signal plot across all targets/probes   More Plot Information      Mean and Variance statistics across all genes                Variance vs. Mean normalized signal plot across all targets/probes  Each gene's variance in the log-scaled, normalized data is plotted against its mean value across all samples. Highly variable genes are indicated by gene name. Housekeeping genes are color coded according to their use in (or omission from) normalization.   
                                           p-value distribution plots   More Plot Information                p-value distribution plots  For each covariate included in the analysis, a histogram of p-values testing each gene's univariate association with the chosen covariate is displayed. Covariates with largely flat histograms have minimal association with gene expression; covariates with histograms with significantly more mass on the left are either associated with the expression of many genes or are confounded with a covariate that is associated with the expression. Low p-values indicate strong evidence for an association.   
                                      
                                       
                                  
                                
                              
                            
                           
                            
                            
                            
                          
                           
                      
                    
                    
                      
                           
                          
                               Heatmaps    PCA     
                              
                                  Heatmaps  
                              
                               
                              
                                  Line  
                                  Day  
                              
                              
                            
                              
                                
                                  
                                       
                                      
                                           Heatmap of Activation of BAD and translocation to mitochondria  Data   More Plot Information                      Heatmap of Activation of BAD and translocation to mitochondria  Data  Heatmap of the normalized data, scaled to give all genes equal variance, generated via unsupervised clustering. Orange indicates high expression; blue indicates low expression. This plot is meant to provide a high level exploratory view of the data.   
                                      
                                       
                                  
                                
                              
                            
                            
                              
                                
                                  
                                       
                                      
                                           Principal Components of Activation of BAD and translocation to mitochondria  Data   More Plot Information                      Principal Components of Activation of BAD and translocation to mitochondria  Data  Principal component analysis maps high-dimensional datasets onto a smaller number of highly informative dimensions. Here, the first four principal components of the gene expression data are plotted against each other and colored by the values of the selected covariate. This plot may be used to identify clusters in the data and to identify variables associated with prominent signal in the data. Variables that are associated with these leading principal components should be considered in downstream analyses.   
                                      
                                       
                                  
                                
                                
                                  
                                       
                                      
                                           Principal Components of Activation of BAD and translocation to mitochondria  Data   More Plot Information                      Principal Components of Activation of BAD and translocation to mitochondria  Data  Principal component analysis maps high-dimensional datasets onto a smaller number of highly informative dimensions. Here, the first four principal components of the gene expression data are plotted against each other and colored by the values of the selected covariate. This plot may be used to identify clusters in the data and to identify variables associated with prominent signal in the data. Variables that are associated with these leading principal components should be considered in downstream analyses.   
                                      
                                       
                                  
                                
                              
                            
                           
                            
                          
                           
                      
                    
                    
                      
                           
                          
                               Heatmaps    PCA     
                              
                                  Heatmaps  
                              
                               
                              
                                  Line  
                                  Day  
                              
                              
                            
                              
                                
                                  
                                       
                                      
                                           Heatmap of Activation of Rac Data   More Plot Information                      Heatmap of Activation of Rac Data  Heatmap of the normalized data, scaled to give all genes equal variance, generated via unsupervised clustering. Orange indicates high expression; blue indicates low expression. This plot is meant to provide a high level exploratory view of the data.   
                                      
                                       
                                  
                                
                              
                            
                            
                              
                                
                                  
                                       
                                      
                                           Principal Components of Activation of Rac Data   More Plot Information                      Principal Components of Activation of Rac Data  Principal component analysis maps high-dimensional datasets onto a smaller number of highly informative dimensions. Here, the first four principal components of the gene expression data are plotted against each other and colored by the values of the selected covariate. This plot may be used to identify clusters in the data and to identify variables associated with prominent signal in the data. Variables that are associated with these leading principal components should be considered in downstream analyses.   
                                      
                                       
                                  
                                
                                
                                  
                                       
                                      
                                           Principal Components of Activation of Rac Data   More Plot Information                      Principal Components of Activation of Rac Data  Principal component analysis maps high-dimensional datasets onto a smaller number of highly informative dimensions. Here, the first four principal components of the gene expression data are plotted against each other and colored by the values of the selected covariate. This plot may be used to identify clusters in the data and to identify variables associated with prominent signal in the data. Variables that are associated with these leading principal components should be considered in downstream analyses.   
                                      
                                       
                                  
                                
                              
                            
                           
                            
                          
                           
                      
                    
                    
                      
                           
                          
                               Heatmaps    PCA     
                              
                                  Heatmaps  
                              
                               
                              
                                  Line  
                                  Day  
                              
                              
                            
                              
                                
                                  
                                       
                                      
                                           Heatmap of Apoptotic cleavage of cellular proteins Data   More Plot Information                      Heatmap of Apoptotic cleavage of cellular proteins Data  Heatmap of the normalized data, scaled to give all genes equal variance, generated via unsupervised clustering. Orange indicates high expression; blue indicates low expression. This plot is meant to provide a high level exploratory view of the data.   
                                      
                                       
                                  
                                
                              
                            
                            
                              
                                
                                  
                                       
                                      
                                           Principal Components of Apoptotic cleavage of cellular proteins Data   More Plot Information                      Principal Components of Apoptotic cleavage of cellular proteins Data  Principal component analysis maps high-dimensional datasets onto a smaller number of highly informative dimensions. Here, the first four principal components of the gene expression data are plotted against each other and colored by the values of the selected covariate. This plot may be used to identify clusters in the data and to identify variables associated with prominent signal in the data. Variables that are associated with these leading principal components should be considered in downstream analyses.   
                                      
                                       
                                  
                                
                                
                                  
                                       
                                      
                                           Principal Components of Apoptotic cleavage of cellular proteins Data   More Plot Information                      Principal Components of Apoptotic cleavage of cellular proteins Data  Principal component analysis maps high-dimensional datasets onto a smaller number of highly informative dimensions. Here, the first four principal components of the gene expression data are plotted against each other and colored by the values of the selected covariate. This plot may be used to identify clusters in the data and to identify variables associated with prominent signal in the data. Variables that are associated with these leading principal components should be considered in downstream analyses.   
                                      
                                       
                                  
                                
                              
                            
                           
                            
                          
                           
                      
                    
                    
                      
                           
                          
                               Heatmaps    PCA     
                              
                                  Heatmaps  
                              
                               
                              
                                  Line  
                                  Day  
                              
                              
                            
                              
                                
                                  
                                       
                                      
                                           Heatmap of ARMS-mediated activation Data   More Plot Information                      Heatmap of ARMS-mediated activation Data  Heatmap of the normalized data, scaled to give all genes equal variance, generated via unsupervised clustering. Orange indicates high expression; blue indicates low expression. This plot is meant to provide a high level exploratory view of the data.   
                                      
                                       
                                  
                                
                              
                            
                            
                              
                                
                                  
                                       
                                      
                                           Principal Components of ARMS-mediated activation Data   More Plot Information                      Principal Components of ARMS-mediated activation Data  Principal component analysis maps high-dimensional datasets onto a smaller number of highly informative dimensions. Here, the first four principal components of the gene expression data are plotted against each other and colored by the values of the selected covariate. This plot may be used to identify clusters in the data and to identify variables associated with prominent signal in the data. Variables that are associated with these leading principal components should be considered in downstream analyses.   
                                      
                                       
                                  
                                
                                
                                  
                                       
                                      
                                           Principal Components of ARMS-mediated activation Data   More Plot Information                      Principal Components of ARMS-mediated activation Data  Principal component analysis maps high-dimensional datasets onto a smaller number of highly informative dimensions. Here, the first four principal components of the gene expression data are plotted against each other and colored by the values of the selected covariate. This plot may be used to identify clusters in the data and to identify variables associated with prominent signal in the data. Variables that are associated with these leading principal components should be considered in downstream analyses.   
                                      
                                       
                                  
                                
                              
                            
                           
                            
                          
                           
                      
                    
                    
                      
                           
                          
                               Heatmaps    PCA     
                              
                                  Heatmaps  
                              
                               
                              
                                  Line  
                                  Day  
                              
                              
                            
                              
                                
                                  
                                       
                                      
                                           Heatmap of AXIN missense mutants destabilize the destruction complex Data   More Plot Information                      Heatmap of AXIN missense mutants destabilize the destruction complex Data  Heatmap of the normalized data, scaled to give all genes equal variance, generated via unsupervised clustering. Orange indicates high expression; blue indicates low expression. This plot is meant to provide a high level exploratory view of the data.   
                                      
                                       
                                  
                                
                              
                            
                            
                              
                                
                                  
                                       
                                      
                                           Principal Components of AXIN missense mutants destabilize the destruction complex Data   More Plot Information                      Principal Components of AXIN missense mutants destabilize the destruction complex Data  Principal component analysis maps high-dimensional datasets onto a smaller number of highly informative dimensions. Here, the first four principal components of the gene expression data are plotted against each other and colored by the values of the selected covariate. This plot may be used to identify clusters in the data and to identify variables associated with prominent signal in the data. Variables that are associated with these leading principal components should be considered in downstream analyses.   
                                      
                                       
                                  
                                
                                
                                  
                                       
                                      
                                           Principal Components of AXIN missense mutants destabilize the destruction complex Data   More Plot Information                      Principal Components of AXIN missense mutants destabilize the destruction complex Data  Principal component analysis maps high-dimensional datasets onto a smaller number of highly informative dimensions. Here, the first four principal components of the gene expression data are plotted against each other and colored by the values of the selected covariate. This plot may be used to identify clusters in the data and to identify variables associated with prominent signal in the data. Variables that are associated with these leading principal components should be considered in downstream analyses.   
                                      
                                       
                                  
                                
                              
                            
                           
                            
                          
                           
                      
                    
                    
                      
                           
                          
                               Heatmaps    PCA     
                              
                                  Heatmaps  
                              
                               
                              
                                  Line  
                                  Day  
                              
                              
                            
                              
                                
                                  
                                       
                                      
                                           Heatmap of Beta-catenin phosphorylation cascade Data   More Plot Information                      Heatmap of Beta-catenin phosphorylation cascade Data  Heatmap of the normalized data, scaled to give all genes equal variance, generated via unsupervised clustering. Orange indicates high expression; blue indicates low expression. This plot is meant to provide a high level exploratory view of the data.   
                                      
                                       
                                  
                                
                              
                            
                            
                              
                                
                                  
                                       
                                      
                                           Principal Components of Beta-catenin phosphorylation cascade Data   More Plot Information                      Principal Components of Beta-catenin phosphorylation cascade Data  Principal component analysis maps high-dimensional datasets onto a smaller number of highly informative dimensions. Here, the first four principal components of the gene expression data are plotted against each other and colored by the values of the selected covariate. This plot may be used to identify clusters in the data and to identify variables associated with prominent signal in the data. Variables that are associated with these leading principal components should be considered in downstream analyses.   
                                      
                                       
                                  
                                
                                
                                  
                                       
                                      
                                           Principal Components of Beta-catenin phosphorylation cascade Data   More Plot Information                      Principal Components of Beta-catenin phosphorylation cascade Data  Principal component analysis maps high-dimensional datasets onto a smaller number of highly informative dimensions. Here, the first four principal components of the gene expression data are plotted against each other and colored by the values of the selected covariate. This plot may be used to identify clusters in the data and to identify variables associated with prominent signal in the data. Variables that are associated with these leading principal components should be considered in downstream analyses.   
                                      
                                       
                                  
                                
                              
                            
                           
                            
                          
                           
                      
                    
                    
                      
                           
                          
                               Heatmaps    PCA     
                              
                                  Heatmaps  
                              
                               
                              
                                  Line  
                                  Day  
                              
                              
                            
                              
                                
                                  
                                       
                                      
                                           Heatmap of CD209 (DC-SIGN) signaling Data   More Plot Information                      Heatmap of CD209 (DC-SIGN) signaling Data  Heatmap of the normalized data, scaled to give all genes equal variance, generated via unsupervised clustering. Orange indicates high expression; blue indicates low expression. This plot is meant to provide a high level exploratory view of the data.   
                                      
                                       
                                  
                                
                              
                            
                            
                              
                                
                                  
                                       
                                      
                                           Principal Components of CD209 (DC-SIGN) signaling Data   More Plot Information                      Principal Components of CD209 (DC-SIGN) signaling Data  Principal component analysis maps high-dimensional datasets onto a smaller number of highly informative dimensions. Here, the first four principal components of the gene expression data are plotted against each other and colored by the values of the selected covariate. This plot may be used to identify clusters in the data and to identify variables associated with prominent signal in the data. Variables that are associated with these leading principal components should be considered in downstream analyses.   
                                      
                                       
                                  
                                
                                
                                  
                                       
                                      
                                           Principal Components of CD209 (DC-SIGN) signaling Data   More Plot Information                      Principal Components of CD209 (DC-SIGN) signaling Data  Principal component analysis maps high-dimensional datasets onto a smaller number of highly informative dimensions. Here, the first four principal components of the gene expression data are plotted against each other and colored by the values of the selected covariate. This plot may be used to identify clusters in the data and to identify variables associated with prominent signal in the data. Variables that are associated with these leading principal components should be considered in downstream analyses.   
                                      
                                       
                                  
                                
                              
                            
                           
                            
                          
                           
                      
                    
                    
                      
                           
                          
                               Heatmaps    PCA     
                              
                                  Heatmaps  
                              
                               
                              
                                  Line  
                                  Day  
                              
                              
                            
                              
                                
                                  
                                       
                                      
                                           Heatmap of CD28 co-stimulation Data   More Plot Information                      Heatmap of CD28 co-stimulation Data  Heatmap of the normalized data, scaled to give all genes equal variance, generated via unsupervised clustering. Orange indicates high expression; blue indicates low expression. This plot is meant to provide a high level exploratory view of the data.   
                                      
                                       
                                  
                                
                              
                            
                            
                              
                                
                                  
                                       
                                      
                                           Principal Components of CD28 co-stimulation Data   More Plot Information                      Principal Components of CD28 co-stimulation Data  Principal component analysis maps high-dimensional datasets onto a smaller number of highly informative dimensions. Here, the first four principal components of the gene expression data are plotted against each other and colored by the values of the selected covariate. This plot may be used to identify clusters in the data and to identify variables associated with prominent signal in the data. Variables that are associated with these leading principal components should be considered in downstream analyses.   
                                      
                                       
                                  
                                
                                
                                  
                                       
                                      
                                           Principal Components of CD28 co-stimulation Data   More Plot Information                      Principal Components of CD28 co-stimulation Data  Principal component analysis maps high-dimensional datasets onto a smaller number of highly informative dimensions. Here, the first four principal components of the gene expression data are plotted against each other and colored by the values of the selected covariate. This plot may be used to identify clusters in the data and to identify variables associated with prominent signal in the data. Variables that are associated with these leading principal components should be considered in downstream analyses.   
                                      
                                       
                                  
                                
                              
                            
                           
                            
                          
                           
                      
                    
                    
                      
                           
                          
                               Heatmaps    PCA     
                              
                                  Heatmaps  
                              
                               
                              
                                  Line  
                                  Day  
                              
                              
                            
                              
                                
                                  
                                       
                                      
                                           Heatmap of CD28 dependent PI3KAkt signaling Data   More Plot Information                      Heatmap of CD28 dependent PI3KAkt signaling Data  Heatmap of the normalized data, scaled to give all genes equal variance, generated via unsupervised clustering. Orange indicates high expression; blue indicates low expression. This plot is meant to provide a high level exploratory view of the data.   
                                      
                                       
                                  
                                
                              
                            
                            
                              
                                
                                  
                                       
                                      
                                           Principal Components of CD28 dependent PI3KAkt signaling Data   More Plot Information                      Principal Components of CD28 dependent PI3KAkt signaling Data  Principal component analysis maps high-dimensional datasets onto a smaller number of highly informative dimensions. Here, the first four principal components of the gene expression data are plotted against each other and colored by the values of the selected covariate. This plot may be used to identify clusters in the data and to identify variables associated with prominent signal in the data. Variables that are associated with these leading principal components should be considered in downstream analyses.   
                                      
                                       
                                  
                                
                                
                                  
                                       
                                      
                                           Principal Components of CD28 dependent PI3KAkt signaling Data   More Plot Information                      Principal Components of CD28 dependent PI3KAkt signaling Data  Principal component analysis maps high-dimensional datasets onto a smaller number of highly informative dimensions. Here, the first four principal components of the gene expression data are plotted against each other and colored by the values of the selected covariate. This plot may be used to identify clusters in the data and to identify variables associated with prominent signal in the data. Variables that are associated with these leading principal components should be considered in downstream analyses.   
                                      
                                       
                                  
                                
                              
                            
                           
                            
                          
                           
                      
                    
                    
                      
                           
                          
                               Heatmaps    PCA     
                              
                                  Heatmaps  
                              
                               
                              
                                  Line  
                                  Day  
                              
                              
                            
                              
                                
                                  
                                       
                                      
                                           Heatmap of CD28 dependent Vav1 pathway Data   More Plot Information                      Heatmap of CD28 dependent Vav1 pathway Data  Heatmap of the normalized data, scaled to give all genes equal variance, generated via unsupervised clustering. Orange indicates high expression; blue indicates low expression. This plot is meant to provide a high level exploratory view of the data.   
                                      
                                       
                                  
                                
                              
                            
                            
                              
                                
                                  
                                       
                                      
                                           Principal Components of CD28 dependent Vav1 pathway Data   More Plot Information                      Principal Components of CD28 dependent Vav1 pathway Data  Principal component analysis maps high-dimensional datasets onto a smaller number of highly informative dimensions. Here, the first four principal components of the gene expression data are plotted against each other and colored by the values of the selected covariate. This plot may be used to identify clusters in the data and to identify variables associated with prominent signal in the data. Variables that are associated with these leading principal components should be considered in downstream analyses.   
                                      
                                       
                                  
                                
                                
                                  
                                       
                                      
                                           Principal Components of CD28 dependent Vav1 pathway Data   More Plot Information                      Principal Components of CD28 dependent Vav1 pathway Data  Principal component analysis maps high-dimensional datasets onto a smaller number of highly informative dimensions. Here, the first four principal components of the gene expression data are plotted against each other and colored by the values of the selected covariate. This plot may be used to identify clusters in the data and to identify variables associated with prominent signal in the data. Variables that are associated with these leading principal components should be considered in downstream analyses.   
                                      
                                       
                                  
                                
                              
                            
                           
                            
                          
                           
                      
                    
                    
                      
                           
                          
                               Heatmaps    PCA     
                              
                                  Heatmaps  
                              
                               
                              
                                  Line  
                                  Day  
                              
                              
                            
                              
                                
                                  
                                       
                                      
                                           Heatmap of Cell-extracellular matrix interactions Data   More Plot Information                      Heatmap of Cell-extracellular matrix interactions Data  Heatmap of the normalized data, scaled to give all genes equal variance, generated via unsupervised clustering. Orange indicates high expression; blue indicates low expression. This plot is meant to provide a high level exploratory view of the data.   
                                      
                                       
                                  
                                
                              
                            
                            
                              
                                
                                  
                                       
                                      
                                           Principal Components of Cell-extracellular matrix interactions Data   More Plot Information                      Principal Components of Cell-extracellular matrix interactions Data  Principal component analysis maps high-dimensional datasets onto a smaller number of highly informative dimensions. Here, the first four principal components of the gene expression data are plotted against each other and colored by the values of the selected covariate. This plot may be used to identify clusters in the data and to identify variables associated with prominent signal in the data. Variables that are associated with these leading principal components should be considered in downstream analyses.   
                                      
                                       
                                  
                                
                                
                                  
                                       
                                      
                                           Principal Components of Cell-extracellular matrix interactions Data   More Plot Information                      Principal Components of Cell-extracellular matrix interactions Data  Principal component analysis maps high-dimensional datasets onto a smaller number of highly informative dimensions. Here, the first four principal components of the gene expression data are plotted against each other and colored by the values of the selected covariate. This plot may be used to identify clusters in the data and to identify variables associated with prominent signal in the data. Variables that are associated with these leading principal components should be considered in downstream analyses.   
                                      
                                       
                                  
                                
                              
                            
                           
                            
                          
                           
                      
                    
                    
                      
                           
                          
                               Heatmaps    PCA     
                              
                                  Heatmaps  
                              
                               
                              
                                  Line  
                                  Day  
                              
                              
                            
                              
                                
                                  
                                       
                                      
                                           Heatmap of Cell surface interactions at the vascular wall Data   More Plot Information                      Heatmap of Cell surface interactions at the vascular wall Data  Heatmap of the normalized data, scaled to give all genes equal variance, generated via unsupervised clustering. Orange indicates high expression; blue indicates low expression. This plot is meant to provide a high level exploratory view of the data.   
                                      
                                       
                                  
                                
                              
                            
                            
                              
                                
                                  
                                       
                                      
                                           Principal Components of Cell surface interactions at the vascular wall Data   More Plot Information                      Principal Components of Cell surface interactions at the vascular wall Data  Principal component analysis maps high-dimensional datasets onto a smaller number of highly informative dimensions. Here, the first four principal components of the gene expression data are plotted against each other and colored by the values of the selected covariate. This plot may be used to identify clusters in the data and to identify variables associated with prominent signal in the data. Variables that are associated with these leading principal components should be considered in downstream analyses.   
                                      
                                       
                                  
                                
                                
                                  
                                       
                                      
                                           Principal Components of Cell surface interactions at the vascular wall Data   More Plot Information                      Principal Components of Cell surface interactions at the vascular wall Data  Principal component analysis maps high-dimensional datasets onto a smaller number of highly informative dimensions. Here, the first four principal components of the gene expression data are plotted against each other and colored by the values of the selected covariate. This plot may be used to identify clusters in the data and to identify variables associated with prominent signal in the data. Variables that are associated with these leading principal components should be considered in downstream analyses.   
                                      
                                       
                                  
                                
                              
                            
                           
                            
                          
                           
                      
                    
                    
                      
                           
                          
                               Heatmaps    PCA     
                              
                                  Heatmaps  
                              
                               
                              
                                  Line  
                                  Day  
                              
                              
                            
                              
                                
                                  
                                       
                                      
                                           Heatmap of Circadian Clock Data   More Plot Information                      Heatmap of Circadian Clock Data  Heatmap of the normalized data, scaled to give all genes equal variance, generated via unsupervised clustering. Orange indicates high expression; blue indicates low expression. This plot is meant to provide a high level exploratory view of the data.   
                                      
                                       
                                  
                                
                              
                            
                            
                              
                                
                                  
                                       
                                      
                                           Principal Components of Circadian Clock Data   More Plot Information                      Principal Components of Circadian Clock Data  Principal component analysis maps high-dimensional datasets onto a smaller number of highly informative dimensions. Here, the first four principal components of the gene expression data are plotted against each other and colored by the values of the selected covariate. This plot may be used to identify clusters in the data and to identify variables associated with prominent signal in the data. Variables that are associated with these leading principal components should be considered in downstream analyses.   
                                      
                                       
                                  
                                
                                
                                  
                                       
                                      
                                           Principal Components of Circadian Clock Data   More Plot Information                      Principal Components of Circadian Clock Data  Principal component analysis maps high-dimensional datasets onto a smaller number of highly informative dimensions. Here, the first four principal components of the gene expression data are plotted against each other and colored by the values of the selected covariate. This plot may be used to identify clusters in the data and to identify variables associated with prominent signal in the data. Variables that are associated with these leading principal components should be considered in downstream analyses.   
                                      
                                       
                                  
                                
                              
                            
                           
                            
                          
                           
                      
                    
                    
                      
                           
                          
                               Heatmaps    PCA     
                              
                                  Heatmaps  
                              
                               
                              
                                  Line  
                                  Day  
                              
                              
                            
                              
                                
                                  
                                       
                                      
                                           Heatmap of Constitutive Signaling by Aberrant PI3K in Cancer Data   More Plot Information                      Heatmap of Constitutive Signaling by Aberrant PI3K in Cancer Data  Heatmap of the normalized data, scaled to give all genes equal variance, generated via unsupervised clustering. Orange indicates high expression; blue indicates low expression. This plot is meant to provide a high level exploratory view of the data.   
                                      
                                       
                                  
                                
                              
                            
                            
                              
                                
                                  
                                       
                                      
                                           Principal Components of Constitutive Signaling by Aberrant PI3K in Cancer Data   More Plot Information                      Principal Components of Constitutive Signaling by Aberrant PI3K in Cancer Data  Principal component analysis maps high-dimensional datasets onto a smaller number of highly informative dimensions. Here, the first four principal components of the gene expression data are plotted against each other and colored by the values of the selected covariate. This plot may be used to identify clusters in the data and to identify variables associated with prominent signal in the data. Variables that are associated with these leading principal components should be considered in downstream analyses.   
                                      
                                       
                                  
                                
                                
                                  
                                       
                                      
                                           Principal Components of Constitutive Signaling by Aberrant PI3K in Cancer Data   More Plot Information                      Principal Components of Constitutive Signaling by Aberrant PI3K in Cancer Data  Principal component analysis maps high-dimensional datasets onto a smaller number of highly informative dimensions. Here, the first four principal components of the gene expression data are plotted against each other and colored by the values of the selected covariate. This plot may be used to identify clusters in the data and to identify variables associated with prominent signal in the data. Variables that are associated with these leading principal components should be considered in downstream analyses.   
                                      
                                       
                                  
                                
                              
                            
                           
                            
                          
                           
                      
                    
                    
                      
                           
                          
                               Heatmaps    PCA     
                              
                                  Heatmaps  
                              
                               
                              
                                  Line  
                                  Day  
                              
                              
                            
                              
                                
                                  
                                       
                                      
                                           Heatmap of Constitutive Signaling by EGFRvIII Data   More Plot Information                      Heatmap of Constitutive Signaling by EGFRvIII Data  Heatmap of the normalized data, scaled to give all genes equal variance, generated via unsupervised clustering. Orange indicates high expression; blue indicates low expression. This plot is meant to provide a high level exploratory view of the data.   
                                      
                                       
                                  
                                
                              
                            
                            
                              
                                
                                  
                                       
                                      
                                           Principal Components of Constitutive Signaling by EGFRvIII Data   More Plot Information                      Principal Components of Constitutive Signaling by EGFRvIII Data  Principal component analysis maps high-dimensional datasets onto a smaller number of highly informative dimensions. Here, the first four principal components of the gene expression data are plotted against each other and colored by the values of the selected covariate. This plot may be used to identify clusters in the data and to identify variables associated with prominent signal in the data. Variables that are associated with these leading principal components should be considered in downstream analyses.   
                                      
                                       
                                  
                                
                                
                                  
                                       
                                      
                                           Principal Components of Constitutive Signaling by EGFRvIII Data   More Plot Information                      Principal Components of Constitutive Signaling by EGFRvIII Data  Principal component analysis maps high-dimensional datasets onto a smaller number of highly informative dimensions. Here, the first four principal components of the gene expression data are plotted against each other and colored by the values of the selected covariate. This plot may be used to identify clusters in the data and to identify variables associated with prominent signal in the data. Variables that are associated with these leading principal components should be considered in downstream analyses.   
                                      
                                       
                                  
                                
                              
                            
                           
                            
                          
                           
                      
                    
                    
                      
                           
                          
                               Heatmaps    PCA     
                              
                                  Heatmaps  
                              
                               
                              
                                  Line  
                                  Day  
                              
                              
                            
                              
                                
                                  
                                       
                                      
                                           Heatmap of CREB phosphorylation through the activation of Ras Data   More Plot Information                      Heatmap of CREB phosphorylation through the activation of Ras Data  Heatmap of the normalized data, scaled to give all genes equal variance, generated via unsupervised clustering. Orange indicates high expression; blue indicates low expression. This plot is meant to provide a high level exploratory view of the data.   
                                      
                                       
                                  
                                
                              
                            
                            
                              
                                
                                  
                                       
                                      
                                           Principal Components of CREB phosphorylation through the activation of Ras Data   More Plot Information                      Principal Components of CREB phosphorylation through the activation of Ras Data  Principal component analysis maps high-dimensional datasets onto a smaller number of highly informative dimensions. Here, the first four principal components of the gene expression data are plotted against each other and colored by the values of the selected covariate. This plot may be used to identify clusters in the data and to identify variables associated with prominent signal in the data. Variables that are associated with these leading principal components should be considered in downstream analyses.   
                                      
                                       
                                  
                                
                                
                                  
                                       
                                      
                                           Principal Components of CREB phosphorylation through the activation of Ras Data   More Plot Information                      Principal Components of CREB phosphorylation through the activation of Ras Data  Principal component analysis maps high-dimensional datasets onto a smaller number of highly informative dimensions. Here, the first four principal components of the gene expression data are plotted against each other and colored by the values of the selected covariate. This plot may be used to identify clusters in the data and to identify variables associated with prominent signal in the data. Variables that are associated with these leading principal components should be considered in downstream analyses.   
                                      
                                       
                                  
                                
                              
                            
                           
                            
                          
                           
                      
                    
                    
                      
                           
                          
                               Heatmaps    PCA     
                              
                                  Heatmaps  
                              
                               
                              
                                  Line  
                                  Day  
                              
                              
                            
                              
                                
                                  
                                       
                                      
                                           Heatmap of CTLA4 inhibitory signaling Data   More Plot Information                      Heatmap of CTLA4 inhibitory signaling Data  Heatmap of the normalized data, scaled to give all genes equal variance, generated via unsupervised clustering. Orange indicates high expression; blue indicates low expression. This plot is meant to provide a high level exploratory view of the data.   
                                      
                                       
                                  
                                
                              
                            
                            
                              
                                
                                  
                                       
                                      
                                           Principal Components of CTLA4 inhibitory signaling Data   More Plot Information                      Principal Components of CTLA4 inhibitory signaling Data  Principal component analysis maps high-dimensional datasets onto a smaller number of highly informative dimensions. Here, the first four principal components of the gene expression data are plotted against each other and colored by the values of the selected covariate. This plot may be used to identify clusters in the data and to identify variables associated with prominent signal in the data. Variables that are associated with these leading principal components should be considered in downstream analyses.   
                                      
                                       
                                  
                                
                                
                                  
                                       
                                      
                                           Principal Components of CTLA4 inhibitory signaling Data   More Plot Information                      Principal Components of CTLA4 inhibitory signaling Data  Principal component analysis maps high-dimensional datasets onto a smaller number of highly informative dimensions. Here, the first four principal components of the gene expression data are plotted against each other and colored by the values of the selected covariate. This plot may be used to identify clusters in the data and to identify variables associated with prominent signal in the data. Variables that are associated with these leading principal components should be considered in downstream analyses.   
                                      
                                       
                                  
                                
                              
                            
                           
                            
                          
                           
                      
                    
                    
                      
                           
                          
                               Heatmaps    PCA     
                              
                                  Heatmaps  
                              
                               
                              
                                  Line  
                                  Day  
                              
                              
                            
                              
                                
                                  
                                       
                                      
                                           Heatmap of Cyclin AB1 associated events during G2M transition Data   More Plot Information                      Heatmap of Cyclin AB1 associated events during G2M transition Data  Heatmap of the normalized data, scaled to give all genes equal variance, generated via unsupervised clustering. Orange indicates high expression; blue indicates low expression. This plot is meant to provide a high level exploratory view of the data.   
                                      
                                       
                                  
                                
                              
                            
                            
                              
                                
                                  
                                       
                                      
                                           Principal Components of Cyclin AB1 associated events during G2M transition Data   More Plot Information                      Principal Components of Cyclin AB1 associated events during G2M transition Data  Principal component analysis maps high-dimensional datasets onto a smaller number of highly informative dimensions. Here, the first four principal components of the gene expression data are plotted against each other and colored by the values of the selected covariate. This plot may be used to identify clusters in the data and to identify variables associated with prominent signal in the data. Variables that are associated with these leading principal components should be considered in downstream analyses.   
                                      
                                       
                                  
                                
                                
                                  
                                       
                                      
                                           Principal Components of Cyclin AB1 associated events during G2M transition Data   More Plot Information                      Principal Components of Cyclin AB1 associated events during G2M transition Data  Principal component analysis maps high-dimensional datasets onto a smaller number of highly informative dimensions. Here, the first four principal components of the gene expression data are plotted against each other and colored by the values of the selected covariate. This plot may be used to identify clusters in the data and to identify variables associated with prominent signal in the data. Variables that are associated with these leading principal components should be considered in downstream analyses.   
                                      
                                       
                                  
                                
                              
                            
                           
                            
                          
                           
                      
                    
                    
                      
                           
                          
                               Heatmaps    PCA     
                              
                                  Heatmaps  
                              
                               
                              
                                  Line  
                                  Day  
                              
                              
                            
                              
                                
                                  
                                       
                                      
                                           Heatmap of Cyclin D associated events in G1 Data   More Plot Information                      Heatmap of Cyclin D associated events in G1 Data  Heatmap of the normalized data, scaled to give all genes equal variance, generated via unsupervised clustering. Orange indicates high expression; blue indicates low expression. This plot is meant to provide a high level exploratory view of the data.   
                                      
                                       
                                  
                                
                              
                            
                            
                              
                                
                                  
                                       
                                      
                                           Principal Components of Cyclin D associated events in G1 Data   More Plot Information                      Principal Components of Cyclin D associated events in G1 Data  Principal component analysis maps high-dimensional datasets onto a smaller number of highly informative dimensions. Here, the first four principal components of the gene expression data are plotted against each other and colored by the values of the selected covariate. This plot may be used to identify clusters in the data and to identify variables associated with prominent signal in the data. Variables that are associated with these leading principal components should be considered in downstream analyses.   
                                      
                                       
                                  
                                
                                
                                  
                                       
                                      
                                           Principal Components of Cyclin D associated events in G1 Data   More Plot Information                      Principal Components of Cyclin D associated events in G1 Data  Principal component analysis maps high-dimensional datasets onto a smaller number of highly informative dimensions. Here, the first four principal components of the gene expression data are plotted against each other and colored by the values of the selected covariate. This plot may be used to identify clusters in the data and to identify variables associated with prominent signal in the data. Variables that are associated with these leading principal components should be considered in downstream analyses.   
                                      
                                       
                                  
                                
                              
                            
                           
                            
                          
                           
                      
                    
                    
                      
                           
                          
                               Heatmaps    PCA     
                              
                                  Heatmaps  
                              
                               
                              
                                  Line  
                                  Day  
                              
                              
                            
                              
                                
                                  
                                       
                                      
                                           Heatmap of DAP12 signaling Data   More Plot Information                      Heatmap of DAP12 signaling Data  Heatmap of the normalized data, scaled to give all genes equal variance, generated via unsupervised clustering. Orange indicates high expression; blue indicates low expression. This plot is meant to provide a high level exploratory view of the data.   
                                      
                                       
                                  
                                
                              
                            
                            
                              
                                
                                  
                                       
                                      
                                           Principal Components of DAP12 signaling Data   More Plot Information                      Principal Components of DAP12 signaling Data  Principal component analysis maps high-dimensional datasets onto a smaller number of highly informative dimensions. Here, the first four principal components of the gene expression data are plotted against each other and colored by the values of the selected covariate. This plot may be used to identify clusters in the data and to identify variables associated with prominent signal in the data. Variables that are associated with these leading principal components should be considered in downstream analyses.   
                                      
                                       
                                  
                                
                                
                                  
                                       
                                      
                                           Principal Components of DAP12 signaling Data   More Plot Information                      Principal Components of DAP12 signaling Data  Principal component analysis maps high-dimensional datasets onto a smaller number of highly informative dimensions. Here, the first four principal components of the gene expression data are plotted against each other and colored by the values of the selected covariate. This plot may be used to identify clusters in the data and to identify variables associated with prominent signal in the data. Variables that are associated with these leading principal components should be considered in downstream analyses.   
                                      
                                       
                                  
                                
                              
                            
                           
                            
                          
                           
                      
                    
                    
                      
                           
                          
                               Heatmaps    PCA     
                              
                                  Heatmaps  
                              
                               
                              
                                  Line  
                                  Day  
                              
                              
                            
                              
                                
                                  
                                       
                                      
                                           Heatmap of DARPP-32 events Data   More Plot Information                      Heatmap of DARPP-32 events Data  Heatmap of the normalized data, scaled to give all genes equal variance, generated via unsupervised clustering. Orange indicates high expression; blue indicates low expression. This plot is meant to provide a high level exploratory view of the data.   
                                      
                                       
                                  
                                
                              
                            
                            
                              
                                
                                  
                                       
                                      
                                           Principal Components of DARPP-32 events Data   More Plot Information                      Principal Components of DARPP-32 events Data  Principal component analysis maps high-dimensional datasets onto a smaller number of highly informative dimensions. Here, the first four principal components of the gene expression data are plotted against each other and colored by the values of the selected covariate. This plot may be used to identify clusters in the data and to identify variables associated with prominent signal in the data. Variables that are associated with these leading principal components should be considered in downstream analyses.   
                                      
                                       
                                  
                                
                                
                                  
                                       
                                      
                                           Principal Components of DARPP-32 events Data   More Plot Information                      Principal Components of DARPP-32 events Data  Principal component analysis maps high-dimensional datasets onto a smaller number of highly informative dimensions. Here, the first four principal components of the gene expression data are plotted against each other and colored by the values of the selected covariate. This plot may be used to identify clusters in the data and to identify variables associated with prominent signal in the data. Variables that are associated with these leading principal components should be considered in downstream analyses.   
                                      
                                       
                                  
                                
                              
                            
                           
                            
                          
                           
                      
                    
                    
                      
                           
                          
                               Heatmaps    PCA     
                              
                                  Heatmaps  
                              
                               
                              
                                  Line  
                                  Day  
                              
                              
                            
                              
                                
                                  
                                       
                                      
                                           Heatmap of DCC mediated attractive signaling Data   More Plot Information                      Heatmap of DCC mediated attractive signaling Data  Heatmap of the normalized data, scaled to give all genes equal variance, generated via unsupervised clustering. Orange indicates high expression; blue indicates low expression. This plot is meant to provide a high level exploratory view of the data.   
                                      
                                       
                                  
                                
                              
                            
                            
                              
                                
                                  
                                       
                                      
                                           Principal Components of DCC mediated attractive signaling Data   More Plot Information                      Principal Components of DCC mediated attractive signaling Data  Principal component analysis maps high-dimensional datasets onto a smaller number of highly informative dimensions. Here, the first four principal components of the gene expression data are plotted against each other and colored by the values of the selected covariate. This plot may be used to identify clusters in the data and to identify variables associated with prominent signal in the data. Variables that are associated with these leading principal components should be considered in downstream analyses.   
                                      
                                       
                                  
                                
                                
                                  
                                       
                                      
                                           Principal Components of DCC mediated attractive signaling Data   More Plot Information                      Principal Components of DCC mediated attractive signaling Data  Principal component analysis maps high-dimensional datasets onto a smaller number of highly informative dimensions. Here, the first four principal components of the gene expression data are plotted against each other and colored by the values of the selected covariate. This plot may be used to identify clusters in the data and to identify variables associated with prominent signal in the data. Variables that are associated with these leading principal components should be considered in downstream analyses.   
                                      
                                       
                                  
                                
                              
                            
                           
                            
                          
                           
                      
                    
                    
                      
                           
                          
                               Heatmaps    PCA     
                              
                                  Heatmaps  
                              
                               
                              
                                  Line  
                                  Day  
                              
                              
                            
                              
                                
                                  
                                       
                                      
                                           Heatmap of Deactivation of the beta-catenin transactivating complex Data   More Plot Information                      Heatmap of Deactivation of the beta-catenin transactivating complex Data  Heatmap of the normalized data, scaled to give all genes equal variance, generated via unsupervised clustering. Orange indicates high expression; blue indicates low expression. This plot is meant to provide a high level exploratory view of the data.   
                                      
                                       
                                  
                                
                              
                            
                            
                              
                                
                                  
                                       
                                      
                                           Principal Components of Deactivation of the beta-catenin transactivating complex Data   More Plot Information                      Principal Components of Deactivation of the beta-catenin transactivating complex Data  Principal component analysis maps high-dimensional datasets onto a smaller number of highly informative dimensions. Here, the first four principal components of the gene expression data are plotted against each other and colored by the values of the selected covariate. This plot may be used to identify clusters in the data and to identify variables associated with prominent signal in the data. Variables that are associated with these leading principal components should be considered in downstream analyses.   
                                      
                                       
                                  
                                
                                
                                  
                                       
                                      
                                           Principal Components of Deactivation of the beta-catenin transactivating complex Data   More Plot Information                      Principal Components of Deactivation of the beta-catenin transactivating complex Data  Principal component analysis maps high-dimensional datasets onto a smaller number of highly informative dimensions. Here, the first four principal components of the gene expression data are plotted against each other and colored by the values of the selected covariate. This plot may be used to identify clusters in the data and to identify variables associated with prominent signal in the data. Variables that are associated with these leading principal components should be considered in downstream analyses.   
                                      
                                       
                                  
                                
                              
                            
                           
                            
                          
                           
                      
                    
                    
                      
                           
                          
                               Heatmaps    PCA     
                              
                                  Heatmaps  
                              
                               
                              
                                  Line  
                                  Day  
                              
                              
                            
                              
                                
                                  
                                       
                                      
                                           Heatmap of Disassembly of the destruction complex and recruitment of AXIN to the membrane Data   More Plot Information                      Heatmap of Disassembly of the destruction complex and recruitment of AXIN to the membrane Data  Heatmap of the normalized data, scaled to give all genes equal variance, generated via unsupervised clustering. Orange indicates high expression; blue indicates low expression. This plot is meant to provide a high level exploratory view of the data.   
                                      
                                       
                                  
                                
                              
                            
                            
                              
                                
                                  
                                       
                                      
                                           Principal Components of Disassembly of the destruction complex and recruitment of AXIN to the membrane Data   More Plot Information                      Principal Components of Disassembly of the destruction complex and recruitment of AXIN to the membrane Data  Principal component analysis maps high-dimensional datasets onto a smaller number of highly informative dimensions. Here, the first four principal components of the gene expression data are plotted against each other and colored by the values of the selected covariate. This plot may be used to identify clusters in the data and to identify variables associated with prominent signal in the data. Variables that are associated with these leading principal components should be considered in downstream analyses.   
                                      
                                       
                                  
                                
                                
                                  
                                       
                                      
                                           Principal Components of Disassembly of the destruction complex and recruitment of AXIN to the membrane Data   More Plot Information                      Principal Components of Disassembly of the destruction complex and recruitment of AXIN to the membrane Data  Principal component analysis maps high-dimensional datasets onto a smaller number of highly informative dimensions. Here, the first four principal components of the gene expression data are plotted against each other and colored by the values of the selected covariate. This plot may be used to identify clusters in the data and to identify variables associated with prominent signal in the data. Variables that are associated with these leading principal components should be considered in downstream analyses.   
                                      
                                       
                                  
                                
                              
                            
                           
                            
                          
                           
                      
                    
                    
                      
                           
                          
                               Heatmaps    PCA     
                              
                                  Heatmaps  
                              
                               
                              
                                  Line  
                                  Day  
                              
                              
                            
                              
                                
                                  
                                       
                                      
                                           Heatmap of Downstream signal transduction Data   More Plot Information                      Heatmap of Downstream signal transduction Data  Heatmap of the normalized data, scaled to give all genes equal variance, generated via unsupervised clustering. Orange indicates high expression; blue indicates low expression. This plot is meant to provide a high level exploratory view of the data.   
                                      
                                       
                                  
                                
                              
                            
                            
                              
                                
                                  
                                       
                                      
                                           Principal Components of Downstream signal transduction Data   More Plot Information                      Principal Components of Downstream signal transduction Data  Principal component analysis maps high-dimensional datasets onto a smaller number of highly informative dimensions. Here, the first four principal components of the gene expression data are plotted against each other and colored by the values of the selected covariate. This plot may be used to identify clusters in the data and to identify variables associated with prominent signal in the data. Variables that are associated with these leading principal components should be considered in downstream analyses.   
                                      
                                       
                                  
                                
                                
                                  
                                       
                                      
                                           Principal Components of Downstream signal transduction Data   More Plot Information                      Principal Components of Downstream signal transduction Data  Principal component analysis maps high-dimensional datasets onto a smaller number of highly informative dimensions. Here, the first four principal components of the gene expression data are plotted against each other and colored by the values of the selected covariate. This plot may be used to identify clusters in the data and to identify variables associated with prominent signal in the data. Variables that are associated with these leading principal components should be considered in downstream analyses.   
                                      
                                       
                                  
                                
                              
                            
                           
                            
                          
                           
                      
                    
                    
                      
                           
                          
                               Heatmaps    PCA     
                              
                                  Heatmaps  
                              
                               
                              
                                  Line  
                                  Day  
                              
                              
                            
                              
                                
                                  
                                       
                                      
                                           Heatmap of Downstream TCR signaling Data   More Plot Information                      Heatmap of Downstream TCR signaling Data  Heatmap of the normalized data, scaled to give all genes equal variance, generated via unsupervised clustering. Orange indicates high expression; blue indicates low expression. This plot is meant to provide a high level exploratory view of the data.   
                                      
                                       
                                  
                                
                              
                            
                            
                              
                                
                                  
                                       
                                      
                                           Principal Components of Downstream TCR signaling Data   More Plot Information                      Principal Components of Downstream TCR signaling Data  Principal component analysis maps high-dimensional datasets onto a smaller number of highly informative dimensions. Here, the first four principal components of the gene expression data are plotted against each other and colored by the values of the selected covariate. This plot may be used to identify clusters in the data and to identify variables associated with prominent signal in the data. Variables that are associated with these leading principal components should be considered in downstream analyses.   
                                      
                                       
                                  
                                
                                
                                  
                                       
                                      
                                           Principal Components of Downstream TCR signaling Data   More Plot Information                      Principal Components of Downstream TCR signaling Data  Principal component analysis maps high-dimensional datasets onto a smaller number of highly informative dimensions. Here, the first four principal components of the gene expression data are plotted against each other and colored by the values of the selected covariate. This plot may be used to identify clusters in the data and to identify variables associated with prominent signal in the data. Variables that are associated with these leading principal components should be considered in downstream analyses.   
                                      
                                       
                                  
                                
                              
                            
                           
                            
                          
                           
                      
                    
                    
                      
                           
                          
                               Heatmaps    PCA     
                              
                                  Heatmaps  
                              
                               
                              
                                  Line  
                                  Day  
                              
                              
                            
                              
                                
                                  
                                       
                                      
                                           Heatmap of ECM proteoglycans Data   More Plot Information                      Heatmap of ECM proteoglycans Data  Heatmap of the normalized data, scaled to give all genes equal variance, generated via unsupervised clustering. Orange indicates high expression; blue indicates low expression. This plot is meant to provide a high level exploratory view of the data.   
                                      
                                       
                                  
                                
                              
                            
                            
                              
                                
                                  
                                       
                                      
                                           Principal Components of ECM proteoglycans Data   More Plot Information                      Principal Components of ECM proteoglycans Data  Principal component analysis maps high-dimensional datasets onto a smaller number of highly informative dimensions. Here, the first four principal components of the gene expression data are plotted against each other and colored by the values of the selected covariate. This plot may be used to identify clusters in the data and to identify variables associated with prominent signal in the data. Variables that are associated with these leading principal components should be considered in downstream analyses.   
                                      
                                       
                                  
                                
                                
                                  
                                       
                                      
                                           Principal Components of ECM proteoglycans Data   More Plot Information                      Principal Components of ECM proteoglycans Data  Principal component analysis maps high-dimensional datasets onto a smaller number of highly informative dimensions. Here, the first four principal components of the gene expression data are plotted against each other and colored by the values of the selected covariate. This plot may be used to identify clusters in the data and to identify variables associated with prominent signal in the data. Variables that are associated with these leading principal components should be considered in downstream analyses.   
                                      
                                       
                                  
                                
                              
                            
                           
                            
                          
                           
                      
                    
                    
                      
                           
                          
                               Heatmaps    PCA     
                              
                                  Heatmaps  
                              
                               
                              
                                  Line  
                                  Day  
                              
                              
                            
                              
                                
                                  
                                       
                                      
                                           Heatmap of Elastic fibre formation Data   More Plot Information                      Heatmap of Elastic fibre formation Data  Heatmap of the normalized data, scaled to give all genes equal variance, generated via unsupervised clustering. Orange indicates high expression; blue indicates low expression. This plot is meant to provide a high level exploratory view of the data.   
                                      
                                       
                                  
                                
                              
                            
                            
                              
                                
                                  
                                       
                                      
                                           Principal Components of Elastic fibre formation Data   More Plot Information                      Principal Components of Elastic fibre formation Data  Principal component analysis maps high-dimensional datasets onto a smaller number of highly informative dimensions. Here, the first four principal components of the gene expression data are plotted against each other and colored by the values of the selected covariate. This plot may be used to identify clusters in the data and to identify variables associated with prominent signal in the data. Variables that are associated with these leading principal components should be considered in downstream analyses.   
                                      
                                       
                                  
                                
                                
                                  
                                       
                                      
                                           Principal Components of Elastic fibre formation Data   More Plot Information                      Principal Components of Elastic fibre formation Data  Principal component analysis maps high-dimensional datasets onto a smaller number of highly informative dimensions. Here, the first four principal components of the gene expression data are plotted against each other and colored by the values of the selected covariate. This plot may be used to identify clusters in the data and to identify variables associated with prominent signal in the data. Variables that are associated with these leading principal components should be considered in downstream analyses.   
                                      
                                       
                                  
                                
                              
                            
                           
                            
                          
                           
                      
                    
                    
                      
                           
                          
                               Heatmaps    PCA     
                              
                                  Heatmaps  
                              
                               
                              
                                  Line  
                                  Day  
                              
                              
                            
                              
                                
                                  
                                       
                                      
                                           Heatmap of EPH-ephrin mediated repulsion of cells Data   More Plot Information                      Heatmap of EPH-ephrin mediated repulsion of cells Data  Heatmap of the normalized data, scaled to give all genes equal variance, generated via unsupervised clustering. Orange indicates high expression; blue indicates low expression. This plot is meant to provide a high level exploratory view of the data.   
                                      
                                       
                                  
                                
                              
                            
                            
                              
                                
                                  
                                       
                                      
                                           Principal Components of EPH-ephrin mediated repulsion of cells Data   More Plot Information                      Principal Components of EPH-ephrin mediated repulsion of cells Data  Principal component analysis maps high-dimensional datasets onto a smaller number of highly informative dimensions. Here, the first four principal components of the gene expression data are plotted against each other and colored by the values of the selected covariate. This plot may be used to identify clusters in the data and to identify variables associated with prominent signal in the data. Variables that are associated with these leading principal components should be considered in downstream analyses.   
                                      
                                       
                                  
                                
                                
                                  
                                       
                                      
                                           Principal Components of EPH-ephrin mediated repulsion of cells Data   More Plot Information                      Principal Components of EPH-ephrin mediated repulsion of cells Data  Principal component analysis maps high-dimensional datasets onto a smaller number of highly informative dimensions. Here, the first four principal components of the gene expression data are plotted against each other and colored by the values of the selected covariate. This plot may be used to identify clusters in the data and to identify variables associated with prominent signal in the data. Variables that are associated with these leading principal components should be considered in downstream analyses.   
                                      
                                       
                                  
                                
                              
                            
                           
                            
                          
                           
                      
                    
                    
                      
                           
                          
                               Heatmaps    PCA     
                              
                                  Heatmaps  
                              
                               
                              
                                  Line  
                                  Day  
                              
                              
                            
                              
                                
                                  
                                       
                                      
                                           Heatmap of EPHA-mediated growth cone collapse Data   More Plot Information                      Heatmap of EPHA-mediated growth cone collapse Data  Heatmap of the normalized data, scaled to give all genes equal variance, generated via unsupervised clustering. Orange indicates high expression; blue indicates low expression. This plot is meant to provide a high level exploratory view of the data.   
                                      
                                       
                                  
                                
                              
                            
                            
                              
                                
                                  
                                       
                                      
                                           Principal Components of EPHA-mediated growth cone collapse Data   More Plot Information                      Principal Components of EPHA-mediated growth cone collapse Data  Principal component analysis maps high-dimensional datasets onto a smaller number of highly informative dimensions. Here, the first four principal components of the gene expression data are plotted against each other and colored by the values of the selected covariate. This plot may be used to identify clusters in the data and to identify variables associated with prominent signal in the data. Variables that are associated with these leading principal components should be considered in downstream analyses.   
                                      
                                       
                                  
                                
                                
                                  
                                       
                                      
                                           Principal Components of EPHA-mediated growth cone collapse Data   More Plot Information                      Principal Components of EPHA-mediated growth cone collapse Data  Principal component analysis maps high-dimensional datasets onto a smaller number of highly informative dimensions. Here, the first four principal components of the gene expression data are plotted against each other and colored by the values of the selected covariate. This plot may be used to identify clusters in the data and to identify variables associated with prominent signal in the data. Variables that are associated with these leading principal components should be considered in downstream analyses.   
                                      
                                       
                                  
                                
                              
                            
                           
                            
                          
                           
                      
                    
                    
                      
                           
                          
                               Heatmaps    PCA     
                              
                                  Heatmaps  
                              
                               
                              
                                  Line  
                                  Day  
                              
                              
                            
                              
                                
                                  
                                       
                                      
                                           Heatmap of EPHB-mediated forward signaling Data   More Plot Information                      Heatmap of EPHB-mediated forward signaling Data  Heatmap of the normalized data, scaled to give all genes equal variance, generated via unsupervised clustering. Orange indicates high expression; blue indicates low expression. This plot is meant to provide a high level exploratory view of the data.   
                                      
                                       
                                  
                                
                              
                            
                            
                              
                                
                                  
                                       
                                      
                                           Principal Components of EPHB-mediated forward signaling Data   More Plot Information                      Principal Components of EPHB-mediated forward signaling Data  Principal component analysis maps high-dimensional datasets onto a smaller number of highly informative dimensions. Here, the first four principal components of the gene expression data are plotted against each other and colored by the values of the selected covariate. This plot may be used to identify clusters in the data and to identify variables associated with prominent signal in the data. Variables that are associated with these leading principal components should be considered in downstream analyses.   
                                      
                                       
                                  
                                
                                
                                  
                                       
                                      
                                           Principal Components of EPHB-mediated forward signaling Data   More Plot Information                      Principal Components of EPHB-mediated forward signaling Data  Principal component analysis maps high-dimensional datasets onto a smaller number of highly informative dimensions. Here, the first four principal components of the gene expression data are plotted against each other and colored by the values of the selected covariate. This plot may be used to identify clusters in the data and to identify variables associated with prominent signal in the data. Variables that are associated with these leading principal components should be considered in downstream analyses.   
                                      
                                       
                                  
                                
                              
                            
                           
                            
                          
                           
                      
                    
                    
                      
                           
                          
                               Heatmaps    PCA     
                              
                                  Heatmaps  
                              
                               
                              
                                  Line  
                                  Day  
                              
                              
                            
                              
                                
                                  
                                       
                                      
                                           Heatmap of Ephrin signaling Data   More Plot Information                      Heatmap of Ephrin signaling Data  Heatmap of the normalized data, scaled to give all genes equal variance, generated via unsupervised clustering. Orange indicates high expression; blue indicates low expression. This plot is meant to provide a high level exploratory view of the data.   
                                      
                                       
                                  
                                
                              
                            
                            
                              
                                
                                  
                                       
                                      
                                           Principal Components of Ephrin signaling Data   More Plot Information                      Principal Components of Ephrin signaling Data  Principal component analysis maps high-dimensional datasets onto a smaller number of highly informative dimensions. Here, the first four principal components of the gene expression data are plotted against each other and colored by the values of the selected covariate. This plot may be used to identify clusters in the data and to identify variables associated with prominent signal in the data. Variables that are associated with these leading principal components should be considered in downstream analyses.   
                                      
                                       
                                  
                                
                                
                                  
                                       
                                      
                                           Principal Components of Ephrin signaling Data   More Plot Information                      Principal Components of Ephrin signaling Data  Principal component analysis maps high-dimensional datasets onto a smaller number of highly informative dimensions. Here, the first four principal components of the gene expression data are plotted against each other and colored by the values of the selected covariate. This plot may be used to identify clusters in the data and to identify variables associated with prominent signal in the data. Variables that are associated with these leading principal components should be considered in downstream analyses.   
                                      
                                       
                                  
                                
                              
                            
                           
                            
                          
                           
                      
                    
                    
                      
                           
                          
                               Heatmaps    PCA     
                              
                                  Heatmaps  
                              
                               
                              
                                  Line  
                                  Day  
                              
                              
                            
                              
                                
                                  
                                       
                                      
                                           Heatmap of ERKs are inactivated Data   More Plot Information                      Heatmap of ERKs are inactivated Data  Heatmap of the normalized data, scaled to give all genes equal variance, generated via unsupervised clustering. Orange indicates high expression; blue indicates low expression. This plot is meant to provide a high level exploratory view of the data.   
                                      
                                       
                                  
                                
                              
                            
                            
                              
                                
                                  
                                       
                                      
                                           Principal Components of ERKs are inactivated Data   More Plot Information                      Principal Components of ERKs are inactivated Data  Principal component analysis maps high-dimensional datasets onto a smaller number of highly informative dimensions. Here, the first four principal components of the gene expression data are plotted against each other and colored by the values of the selected covariate. This plot may be used to identify clusters in the data and to identify variables associated with prominent signal in the data. Variables that are associated with these leading principal components should be considered in downstream analyses.   
                                      
                                       
                                  
                                
                                
                                  
                                       
                                      
                                           Principal Components of ERKs are inactivated Data   More Plot Information                      Principal Components of ERKs are inactivated Data  Principal component analysis maps high-dimensional datasets onto a smaller number of highly informative dimensions. Here, the first four principal components of the gene expression data are plotted against each other and colored by the values of the selected covariate. This plot may be used to identify clusters in the data and to identify variables associated with prominent signal in the data. Variables that are associated with these leading principal components should be considered in downstream analyses.   
                                      
                                       
                                  
                                
                              
                            
                           
                            
                          
                           
                      
                    
                    
                      
                           
                          
                               Heatmaps    PCA     
                              
                                  Heatmaps  
                              
                               
                              
                                  Line  
                                  Day  
                              
                              
                            
                              
                                
                                  
                                       
                                      
                                           Heatmap of FCERI mediated MAPK activation Data   More Plot Information                      Heatmap of FCERI mediated MAPK activation Data  Heatmap of the normalized data, scaled to give all genes equal variance, generated via unsupervised clustering. Orange indicates high expression; blue indicates low expression. This plot is meant to provide a high level exploratory view of the data.   
                                      
                                       
                                  
                                
                              
                            
                            
                              
                                
                                  
                                       
                                      
                                           Principal Components of FCERI mediated MAPK activation Data   More Plot Information                      Principal Components of FCERI mediated MAPK activation Data  Principal component analysis maps high-dimensional datasets onto a smaller number of highly informative dimensions. Here, the first four principal components of the gene expression data are plotted against each other and colored by the values of the selected covariate. This plot may be used to identify clusters in the data and to identify variables associated with prominent signal in the data. Variables that are associated with these leading principal components should be considered in downstream analyses.   
                                      
                                       
                                  
                                
                                
                                  
                                       
                                      
                                           Principal Components of FCERI mediated MAPK activation Data   More Plot Information                      Principal Components of FCERI mediated MAPK activation Data  Principal component analysis maps high-dimensional datasets onto a smaller number of highly informative dimensions. Here, the first four principal components of the gene expression data are plotted against each other and colored by the values of the selected covariate. This plot may be used to identify clusters in the data and to identify variables associated with prominent signal in the data. Variables that are associated with these leading principal components should be considered in downstream analyses.   
                                      
                                       
                                  
                                
                              
                            
                           
                            
                          
                           
                      
                    
                    
                      
                           
                          
                               Heatmaps    PCA     
                              
                                  Heatmaps  
                              
                               
                              
                                  Line  
                                  Day  
                              
                              
                            
                              
                                
                                  
                                       
                                      
                                           Heatmap of Formation of the beta-catenin.CL.TCF transactivating complex Data   More Plot Information                      Heatmap of Formation of the beta-catenin.CL.TCF transactivating complex Data  Heatmap of the normalized data, scaled to give all genes equal variance, generated via unsupervised clustering. Orange indicates high expression; blue indicates low expression. This plot is meant to provide a high level exploratory view of the data.   
                                      
                                       
                                  
                                
                              
                            
                            
                              
                                
                                  
                                       
                                      
                                           Principal Components of Formation of the beta-catenin.CL.TCF transactivating complex Data   More Plot Information                      Principal Components of Formation of the beta-catenin.CL.TCF transactivating complex Data  Principal component analysis maps high-dimensional datasets onto a smaller number of highly informative dimensions. Here, the first four principal components of the gene expression data are plotted against each other and colored by the values of the selected covariate. This plot may be used to identify clusters in the data and to identify variables associated with prominent signal in the data. Variables that are associated with these leading principal components should be considered in downstream analyses.   
                                      
                                       
                                  
                                
                                
                                  
                                       
                                      
                                           Principal Components of Formation of the beta-catenin.CL.TCF transactivating complex Data   More Plot Information                      Principal Components of Formation of the beta-catenin.CL.TCF transactivating complex Data  Principal component analysis maps high-dimensional datasets onto a smaller number of highly informative dimensions. Here, the first four principal components of the gene expression data are plotted against each other and colored by the values of the selected covariate. This plot may be used to identify clusters in the data and to identify variables associated with prominent signal in the data. Variables that are associated with these leading principal components should be considered in downstream analyses.   
                                      
                                       
                                  
                                
                              
                            
                           
                            
                          
                           
                      
                    
                    
                      
                           
                          
                               Heatmaps    PCA     
                              
                                  Heatmaps  
                              
                               
                              
                                  Line  
                                  Day  
                              
                              
                            
                              
                                
                                  
                                       
                                      
                                           Heatmap of Frs2-mediated activation Data   More Plot Information                      Heatmap of Frs2-mediated activation Data  Heatmap of the normalized data, scaled to give all genes equal variance, generated via unsupervised clustering. Orange indicates high expression; blue indicates low expression. This plot is meant to provide a high level exploratory view of the data.   
                                      
                                       
                                  
                                
                              
                            
                            
                              
                                
                                  
                                       
                                      
                                           Principal Components of Frs2-mediated activation Data   More Plot Information                      Principal Components of Frs2-mediated activation Data  Principal component analysis maps high-dimensional datasets onto a smaller number of highly informative dimensions. Here, the first four principal components of the gene expression data are plotted against each other and colored by the values of the selected covariate. This plot may be used to identify clusters in the data and to identify variables associated with prominent signal in the data. Variables that are associated with these leading principal components should be considered in downstream analyses.   
                                      
                                       
                                  
                                
                                
                                  
                                       
                                      
                                           Principal Components of Frs2-mediated activation Data   More Plot Information                      Principal Components of Frs2-mediated activation Data  Principal component analysis maps high-dimensional datasets onto a smaller number of highly informative dimensions. Here, the first four principal components of the gene expression data are plotted against each other and colored by the values of the selected covariate. This plot may be used to identify clusters in the data and to identify variables associated with prominent signal in the data. Variables that are associated with these leading principal components should be considered in downstream analyses.   
                                      
                                       
                                  
                                
                              
                            
                           
                            
                          
                           
                      
                    
                    
                      
                           
                          
                               Heatmaps    PCA     
                              
                                  Heatmaps  
                              
                               
                              
                                  Line  
                                  Day  
                              
                              
                            
                              
                                
                                  
                                       
                                      
                                           Heatmap of G alpha (1213) signalling events Data   More Plot Information                      Heatmap of G alpha (1213) signalling events Data  Heatmap of the normalized data, scaled to give all genes equal variance, generated via unsupervised clustering. Orange indicates high expression; blue indicates low expression. This plot is meant to provide a high level exploratory view of the data.   
                                      
                                       
                                  
                                
                              
                            
                            
                              
                                
                                  
                                       
                                      
                                           Principal Components of G alpha (1213) signalling events Data   More Plot Information                      Principal Components of G alpha (1213) signalling events Data  Principal component analysis maps high-dimensional datasets onto a smaller number of highly informative dimensions. Here, the first four principal components of the gene expression data are plotted against each other and colored by the values of the selected covariate. This plot may be used to identify clusters in the data and to identify variables associated with prominent signal in the data. Variables that are associated with these leading principal components should be considered in downstream analyses.   
                                      
                                       
                                  
                                
                                
                                  
                                       
                                      
                                           Principal Components of G alpha (1213) signalling events Data   More Plot Information                      Principal Components of G alpha (1213) signalling events Data  Principal component analysis maps high-dimensional datasets onto a smaller number of highly informative dimensions. Here, the first four principal components of the gene expression data are plotted against each other and colored by the values of the selected covariate. This plot may be used to identify clusters in the data and to identify variables associated with prominent signal in the data. Variables that are associated with these leading principal components should be considered in downstream analyses.   
                                      
                                       
                                  
                                
                              
                            
                           
                            
                          
                           
                      
                    
                    
                      
                           
                          
                               Heatmaps    PCA     
                              
                                  Heatmaps  
                              
                               
                              
                                  Line  
                                  Day  
                              
                              
                            
                              
                                
                                  
                                       
                                      
                                           Heatmap of G alpha (q) signalling events Data   More Plot Information                      Heatmap of G alpha (q) signalling events Data  Heatmap of the normalized data, scaled to give all genes equal variance, generated via unsupervised clustering. Orange indicates high expression; blue indicates low expression. This plot is meant to provide a high level exploratory view of the data.   
                                      
                                       
                                  
                                
                              
                            
                            
                              
                                
                                  
                                       
                                      
                                           Principal Components of G alpha (q) signalling events Data   More Plot Information                      Principal Components of G alpha (q) signalling events Data  Principal component analysis maps high-dimensional datasets onto a smaller number of highly informative dimensions. Here, the first four principal components of the gene expression data are plotted against each other and colored by the values of the selected covariate. This plot may be used to identify clusters in the data and to identify variables associated with prominent signal in the data. Variables that are associated with these leading principal components should be considered in downstream analyses.   
                                      
                                       
                                  
                                
                                
                                  
                                       
                                      
                                           Principal Components of G alpha (q) signalling events Data   More Plot Information                      Principal Components of G alpha (q) signalling events Data  Principal component analysis maps high-dimensional datasets onto a smaller number of highly informative dimensions. Here, the first four principal components of the gene expression data are plotted against each other and colored by the values of the selected covariate. This plot may be used to identify clusters in the data and to identify variables associated with prominent signal in the data. Variables that are associated with these leading principal components should be considered in downstream analyses.   
                                      
                                       
                                  
                                
                              
                            
                           
                            
                          
                           
                      
                    
                    
                      
                           
                          
                               Heatmaps    PCA     
                              
                                  Heatmaps  
                              
                               
                              
                                  Line  
                                  Day  
                              
                              
                            
                              
                                
                                  
                                       
                                      
                                           Heatmap of G beta.CL.gamma signalling through PI3Kgamma Data   More Plot Information                      Heatmap of G beta.CL.gamma signalling through PI3Kgamma Data  Heatmap of the normalized data, scaled to give all genes equal variance, generated via unsupervised clustering. Orange indicates high expression; blue indicates low expression. This plot is meant to provide a high level exploratory view of the data.   
                                      
                                       
                                  
                                
                              
                            
                            
                              
                                
                                  
                                       
                                      
                                           Principal Components of G beta.CL.gamma signalling through PI3Kgamma Data   More Plot Information                      Principal Components of G beta.CL.gamma signalling through PI3Kgamma Data  Principal component analysis maps high-dimensional datasets onto a smaller number of highly informative dimensions. Here, the first four principal components of the gene expression data are plotted against each other and colored by the values of the selected covariate. This plot may be used to identify clusters in the data and to identify variables associated with prominent signal in the data. Variables that are associated with these leading principal components should be considered in downstream analyses.   
                                      
                                       
                                  
                                
                                
                                  
                                       
                                      
                                           Principal Components of G beta.CL.gamma signalling through PI3Kgamma Data   More Plot Information                      Principal Components of G beta.CL.gamma signalling through PI3Kgamma Data  Principal component analysis maps high-dimensional datasets onto a smaller number of highly informative dimensions. Here, the first four principal components of the gene expression data are plotted against each other and colored by the values of the selected covariate. This plot may be used to identify clusters in the data and to identify variables associated with prominent signal in the data. Variables that are associated with these leading principal components should be considered in downstream analyses.   
                                      
                                       
                                  
                                
                              
                            
                           
                            
                          
                           
                      
                    
                    
                      
                           
                          
                               Heatmaps    PCA     
                              
                                  Heatmaps  
                              
                               
                              
                                  Line  
                                  Day  
                              
                              
                            
                              
                                
                                  
                                       
                                      
                                           Heatmap of GAB1 signalosome Data   More Plot Information                      Heatmap of GAB1 signalosome Data  Heatmap of the normalized data, scaled to give all genes equal variance, generated via unsupervised clustering. Orange indicates high expression; blue indicates low expression. This plot is meant to provide a high level exploratory view of the data.   
                                      
                                       
                                  
                                
                              
                            
                            
                              
                                
                                  
                                       
                                      
                                           Principal Components of GAB1 signalosome Data   More Plot Information                      Principal Components of GAB1 signalosome Data  Principal component analysis maps high-dimensional datasets onto a smaller number of highly informative dimensions. Here, the first four principal components of the gene expression data are plotted against each other and colored by the values of the selected covariate. This plot may be used to identify clusters in the data and to identify variables associated with prominent signal in the data. Variables that are associated with these leading principal components should be considered in downstream analyses.   
                                      
                                       
                                  
                                
                                
                                  
                                       
                                      
                                           Principal Components of GAB1 signalosome Data   More Plot Information                      Principal Components of GAB1 signalosome Data  Principal component analysis maps high-dimensional datasets onto a smaller number of highly informative dimensions. Here, the first four principal components of the gene expression data are plotted against each other and colored by the values of the selected covariate. This plot may be used to identify clusters in the data and to identify variables associated with prominent signal in the data. Variables that are associated with these leading principal components should be considered in downstream analyses.   
                                      
                                       
                                  
                                
                              
                            
                           
                            
                          
                           
                      
                    
                    
                      
                           
                          
                               Heatmaps    PCA     
                              
                                  Heatmaps  
                              
                               
                              
                                  Line  
                                  Day  
                              
                              
                            
                              
                                
                                  
                                       
                                      
                                           Heatmap of Gastrin-CREB signalling pathway via PKC and MAPK Data   More Plot Information                      Heatmap of Gastrin-CREB signalling pathway via PKC and MAPK Data  Heatmap of the normalized data, scaled to give all genes equal variance, generated via unsupervised clustering. Orange indicates high expression; blue indicates low expression. This plot is meant to provide a high level exploratory view of the data.   
                                      
                                       
                                  
                                
                              
                            
                            
                              
                                
                                  
                                       
                                      
                                           Principal Components of Gastrin-CREB signalling pathway via PKC and MAPK Data   More Plot Information                      Principal Components of Gastrin-CREB signalling pathway via PKC and MAPK Data  Principal component analysis maps high-dimensional datasets onto a smaller number of highly informative dimensions. Here, the first four principal components of the gene expression data are plotted against each other and colored by the values of the selected covariate. This plot may be used to identify clusters in the data and to identify variables associated with prominent signal in the data. Variables that are associated with these leading principal components should be considered in downstream analyses.   
                                      
                                       
                                  
                                
                                
                                  
                                       
                                      
                                           Principal Components of Gastrin-CREB signalling pathway via PKC and MAPK Data   More Plot Information                      Principal Components of Gastrin-CREB signalling pathway via PKC and MAPK Data  Principal component analysis maps high-dimensional datasets onto a smaller number of highly informative dimensions. Here, the first four principal components of the gene expression data are plotted against each other and colored by the values of the selected covariate. This plot may be used to identify clusters in the data and to identify variables associated with prominent signal in the data. Variables that are associated with these leading principal components should be considered in downstream analyses.   
                                      
                                       
                                  
                                
                              
                            
                           
                            
                          
                           
                      
                    
                    
                      
                           
                          
                               Heatmaps    PCA     
                              
                                  Heatmaps  
                              
                               
                              
                                  Line  
                                  Day  
                              
                              
                            
                              
                                
                                  
                                       
                                      
                                           Heatmap of GPVI-mediated activation cascade Data   More Plot Information                      Heatmap of GPVI-mediated activation cascade Data  Heatmap of the normalized data, scaled to give all genes equal variance, generated via unsupervised clustering. Orange indicates high expression; blue indicates low expression. This plot is meant to provide a high level exploratory view of the data.   
                                      
                                       
                                  
                                
                              
                            
                            
                              
                                
                                  
                                       
                                      
                                           Principal Components of GPVI-mediated activation cascade Data   More Plot Information                      Principal Components of GPVI-mediated activation cascade Data  Principal component analysis maps high-dimensional datasets onto a smaller number of highly informative dimensions. Here, the first four principal components of the gene expression data are plotted against each other and colored by the values of the selected covariate. This plot may be used to identify clusters in the data and to identify variables associated with prominent signal in the data. Variables that are associated with these leading principal components should be considered in downstream analyses.   
                                      
                                       
                                  
                                
                                
                                  
                                       
                                      
                                           Principal Components of GPVI-mediated activation cascade Data   More Plot Information                      Principal Components of GPVI-mediated activation cascade Data  Principal component analysis maps high-dimensional datasets onto a smaller number of highly informative dimensions. Here, the first four principal components of the gene expression data are plotted against each other and colored by the values of the selected covariate. This plot may be used to identify clusters in the data and to identify variables associated with prominent signal in the data. Variables that are associated with these leading principal components should be considered in downstream analyses.   
                                      
                                       
                                  
                                
                              
                            
                           
                            
                          
                           
                      
                    
                    
                      
                           
                          
                               Heatmaps    PCA     
                              
                                  Heatmaps  
                              
                               
                              
                                  Line  
                                  Day  
                              
                              
                            
                              
                                
                                  
                                       
                                      
                                           Heatmap of Integrin alphaIIb beta3 signaling Data   More Plot Information                      Heatmap of Integrin alphaIIb beta3 signaling Data  Heatmap of the normalized data, scaled to give all genes equal variance, generated via unsupervised clustering. Orange indicates high expression; blue indicates low expression. This plot is meant to provide a high level exploratory view of the data.   
                                      
                                       
                                  
                                
                              
                            
                            
                              
                                
                                  
                                       
                                      
                                           Principal Components of Integrin alphaIIb beta3 signaling Data   More Plot Information                      Principal Components of Integrin alphaIIb beta3 signaling Data  Principal component analysis maps high-dimensional datasets onto a smaller number of highly informative dimensions. Here, the first four principal components of the gene expression data are plotted against each other and colored by the values of the selected covariate. This plot may be used to identify clusters in the data and to identify variables associated with prominent signal in the data. Variables that are associated with these leading principal components should be considered in downstream analyses.   
                                      
                                       
                                  
                                
                                
                                  
                                       
                                      
                                           Principal Components of Integrin alphaIIb beta3 signaling Data   More Plot Information                      Principal Components of Integrin alphaIIb beta3 signaling Data  Principal component analysis maps high-dimensional datasets onto a smaller number of highly informative dimensions. Here, the first four principal components of the gene expression data are plotted against each other and colored by the values of the selected covariate. This plot may be used to identify clusters in the data and to identify variables associated with prominent signal in the data. Variables that are associated with these leading principal components should be considered in downstream analyses.   
                                      
                                       
                                  
                                
                              
                            
                           
                            
                          
                           
                      
                    
                    
                      
                           
                          
                               Heatmaps    PCA     
                              
                                  Heatmaps  
                              
                               
                              
                                  Line  
                                  Day  
                              
                              
                            
                              
                                
                                  
                                       
                                      
                                           Heatmap of Integrin cell surface interactions Data   More Plot Information                      Heatmap of Integrin cell surface interactions Data  Heatmap of the normalized data, scaled to give all genes equal variance, generated via unsupervised clustering. Orange indicates high expression; blue indicates low expression. This plot is meant to provide a high level exploratory view of the data.   
                                      
                                       
                                  
                                
                              
                            
                            
                              
                                
                                  
                                       
                                      
                                           Principal Components of Integrin cell surface interactions Data   More Plot Information                      Principal Components of Integrin cell surface interactions Data  Principal component analysis maps high-dimensional datasets onto a smaller number of highly informative dimensions. Here, the first four principal components of the gene expression data are plotted against each other and colored by the values of the selected covariate. This plot may be used to identify clusters in the data and to identify variables associated with prominent signal in the data. Variables that are associated with these leading principal components should be considered in downstream analyses.   
                                      
                                       
                                  
                                
                                
                                  
                                       
                                      
                                           Principal Components of Integrin cell surface interactions Data   More Plot Information                      Principal Components of Integrin cell surface interactions Data  Principal component analysis maps high-dimensional datasets onto a smaller number of highly informative dimensions. Here, the first four principal components of the gene expression data are plotted against each other and colored by the values of the selected covariate. This plot may be used to identify clusters in the data and to identify variables associated with prominent signal in the data. Variables that are associated with these leading principal components should be considered in downstream analyses.   
                                      
                                       
                                  
                                
                              
                            
                           
                            
                          
                           
                      
                    
                    
                      
                           
                          
                               Heatmaps    PCA     
                              
                                  Heatmaps  
                              
                               
                              
                                  Line  
                                  Day  
                              
                              
                            
                              
                                
                                  
                                       
                                      
                                           Heatmap of Interleukin receptor SHC signaling Data   More Plot Information                      Heatmap of Interleukin receptor SHC signaling Data  Heatmap of the normalized data, scaled to give all genes equal variance, generated via unsupervised clustering. Orange indicates high expression; blue indicates low expression. This plot is meant to provide a high level exploratory view of the data.   
                                      
                                       
                                  
                                
                              
                            
                            
                              
                                
                                  
                                       
                                      
                                           Principal Components of Interleukin receptor SHC signaling Data   More Plot Information                      Principal Components of Interleukin receptor SHC signaling Data  Principal component analysis maps high-dimensional datasets onto a smaller number of highly informative dimensions. Here, the first four principal components of the gene expression data are plotted against each other and colored by the values of the selected covariate. This plot may be used to identify clusters in the data and to identify variables associated with prominent signal in the data. Variables that are associated with these leading principal components should be considered in downstream analyses.   
                                      
                                       
                                  
                                
                                
                                  
                                       
                                      
                                           Principal Components of Interleukin receptor SHC signaling Data   More Plot Information                      Principal Components of Interleukin receptor SHC signaling Data  Principal component analysis maps high-dimensional datasets onto a smaller number of highly informative dimensions. Here, the first four principal components of the gene expression data are plotted against each other and colored by the values of the selected covariate. This plot may be used to identify clusters in the data and to identify variables associated with prominent signal in the data. Variables that are associated with these leading principal components should be considered in downstream analyses.   
                                      
                                       
                                  
                                
                              
                            
                           
                            
                          
                           
                      
                    
                    
                      
                           
                          
                               Heatmaps    PCA     
                              
                                  Heatmaps  
                              
                               
                              
                                  Line  
                                  Day  
                              
                              
                            
                              
                                
                                  
                                       
                                      
                                           Heatmap of IRS-related events triggered by IGF1R Data   More Plot Information                      Heatmap of IRS-related events triggered by IGF1R Data  Heatmap of the normalized data, scaled to give all genes equal variance, generated via unsupervised clustering. Orange indicates high expression; blue indicates low expression. This plot is meant to provide a high level exploratory view of the data.   
                                      
                                       
                                  
                                
                              
                            
                            
                              
                                
                                  
                                       
                                      
                                           Principal Components of IRS-related events triggered by IGF1R Data   More Plot Information                      Principal Components of IRS-related events triggered by IGF1R Data  Principal component analysis maps high-dimensional datasets onto a smaller number of highly informative dimensions. Here, the first four principal components of the gene expression data are plotted against each other and colored by the values of the selected covariate. This plot may be used to identify clusters in the data and to identify variables associated with prominent signal in the data. Variables that are associated with these leading principal components should be considered in downstream analyses.   
                                      
                                       
                                  
                                
                                
                                  
                                       
                                      
                                           Principal Components of IRS-related events triggered by IGF1R Data   More Plot Information                      Principal Components of IRS-related events triggered by IGF1R Data  Principal component analysis maps high-dimensional datasets onto a smaller number of highly informative dimensions. Here, the first four principal components of the gene expression data are plotted against each other and colored by the values of the selected covariate. This plot may be used to identify clusters in the data and to identify variables associated with prominent signal in the data. Variables that are associated with these leading principal components should be considered in downstream analyses.   
                                      
                                       
                                  
                                
                              
                            
                           
                            
                          
                           
                      
                    
                    
                      
                           
                          
                               Heatmaps    PCA     
                              
                                  Heatmaps  
                              
                               
                              
                                  Line  
                                  Day  
                              
                              
                            
                              
                                
                                  
                                       
                                      
                                           Heatmap of Laminin interactions Data   More Plot Information                      Heatmap of Laminin interactions Data  Heatmap of the normalized data, scaled to give all genes equal variance, generated via unsupervised clustering. Orange indicates high expression; blue indicates low expression. This plot is meant to provide a high level exploratory view of the data.   
                                      
                                       
                                  
                                
                              
                            
                            
                              
                                
                                  
                                       
                                      
                                           Principal Components of Laminin interactions Data   More Plot Information                      Principal Components of Laminin interactions Data  Principal component analysis maps high-dimensional datasets onto a smaller number of highly informative dimensions. Here, the first four principal components of the gene expression data are plotted against each other and colored by the values of the selected covariate. This plot may be used to identify clusters in the data and to identify variables associated with prominent signal in the data. Variables that are associated with these leading principal components should be considered in downstream analyses.   
                                      
                                       
                                  
                                
                                
                                  
                                       
                                      
                                           Principal Components of Laminin interactions Data   More Plot Information                      Principal Components of Laminin interactions Data  Principal component analysis maps high-dimensional datasets onto a smaller number of highly informative dimensions. Here, the first four principal components of the gene expression data are plotted against each other and colored by the values of the selected covariate. This plot may be used to identify clusters in the data and to identify variables associated with prominent signal in the data. Variables that are associated with these leading principal components should be considered in downstream analyses.   
                                      
                                       
                                  
                                
                              
                            
                           
                            
                          
                           
                      
                    
                    
                      
                           
                          
                               Heatmaps    PCA     
                              
                                  Heatmaps  
                              
                               
                              
                                  Line  
                                  Day  
                              
                              
                            
                              
                                
                                  
                                       
                                      
                                           Heatmap of MAP2K and MAPK activation Data   More Plot Information                      Heatmap of MAP2K and MAPK activation Data  Heatmap of the normalized data, scaled to give all genes equal variance, generated via unsupervised clustering. Orange indicates high expression; blue indicates low expression. This plot is meant to provide a high level exploratory view of the data.   
                                      
                                       
                                  
                                
                              
                            
                            
                              
                                
                                  
                                       
                                      
                                           Principal Components of MAP2K and MAPK activation Data   More Plot Information                      Principal Components of MAP2K and MAPK activation Data  Principal component analysis maps high-dimensional datasets onto a smaller number of highly informative dimensions. Here, the first four principal components of the gene expression data are plotted against each other and colored by the values of the selected covariate. This plot may be used to identify clusters in the data and to identify variables associated with prominent signal in the data. Variables that are associated with these leading principal components should be considered in downstream analyses.   
                                      
                                       
                                  
                                
                                
                                  
                                       
                                      
                                           Principal Components of MAP2K and MAPK activation Data   More Plot Information                      Principal Components of MAP2K and MAPK activation Data  Principal component analysis maps high-dimensional datasets onto a smaller number of highly informative dimensions. Here, the first four principal components of the gene expression data are plotted against each other and colored by the values of the selected covariate. This plot may be used to identify clusters in the data and to identify variables associated with prominent signal in the data. Variables that are associated with these leading principal components should be considered in downstream analyses.   
                                      
                                       
                                  
                                
                              
                            
                           
                            
                          
                           
                      
                    
                    
                      
                           
                          
                               Heatmaps    PCA     
                              
                                  Heatmaps  
                              
                               
                              
                                  Line  
                                  Day  
                              
                              
                            
                              
                                
                                  
                                       
                                      
                                           Heatmap of MAPK6MAPK4 signaling Data   More Plot Information                      Heatmap of MAPK6MAPK4 signaling Data  Heatmap of the normalized data, scaled to give all genes equal variance, generated via unsupervised clustering. Orange indicates high expression; blue indicates low expression. This plot is meant to provide a high level exploratory view of the data.   
                                      
                                       
                                  
                                
                              
                            
                            
                              
                                
                                  
                                       
                                      
                                           Principal Components of MAPK6MAPK4 signaling Data   More Plot Information                      Principal Components of MAPK6MAPK4 signaling Data  Principal component analysis maps high-dimensional datasets onto a smaller number of highly informative dimensions. Here, the first four principal components of the gene expression data are plotted against each other and colored by the values of the selected covariate. This plot may be used to identify clusters in the data and to identify variables associated with prominent signal in the data. Variables that are associated with these leading principal components should be considered in downstream analyses.   
                                      
                                       
                                  
                                
                                
                                  
                                       
                                      
                                           Principal Components of MAPK6MAPK4 signaling Data   More Plot Information                      Principal Components of MAPK6MAPK4 signaling Data  Principal component analysis maps high-dimensional datasets onto a smaller number of highly informative dimensions. Here, the first four principal components of the gene expression data are plotted against each other and colored by the values of the selected covariate. This plot may be used to identify clusters in the data and to identify variables associated with prominent signal in the data. Variables that are associated with these leading principal components should be considered in downstream analyses.   
                                      
                                       
                                  
                                
                              
                            
                           
                            
                          
                           
                      
                    
                    
                      
                           
                          
                               Heatmaps    PCA     
                              
                                  Heatmaps  
                              
                               
                              
                                  Line  
                                  Day  
                              
                              
                            
                              
                                
                                  
                                       
                                      
                                           Heatmap of MASTL Facilitates Mitotic Progression Data   More Plot Information                      Heatmap of MASTL Facilitates Mitotic Progression Data  Heatmap of the normalized data, scaled to give all genes equal variance, generated via unsupervised clustering. Orange indicates high expression; blue indicates low expression. This plot is meant to provide a high level exploratory view of the data.   
                                      
                                       
                                  
                                
                              
                            
                            
                              
                                
                                  
                                       
                                      
                                           Principal Components of MASTL Facilitates Mitotic Progression Data   More Plot Information                      Principal Components of MASTL Facilitates Mitotic Progression Data  Principal component analysis maps high-dimensional datasets onto a smaller number of highly informative dimensions. Here, the first four principal components of the gene expression data are plotted against each other and colored by the values of the selected covariate. This plot may be used to identify clusters in the data and to identify variables associated with prominent signal in the data. Variables that are associated with these leading principal components should be considered in downstream analyses.   
                                      
                                       
                                  
                                
                                
                                  
                                       
                                      
                                           Principal Components of MASTL Facilitates Mitotic Progression Data   More Plot Information                      Principal Components of MASTL Facilitates Mitotic Progression Data  Principal component analysis maps high-dimensional datasets onto a smaller number of highly informative dimensions. Here, the first four principal components of the gene expression data are plotted against each other and colored by the values of the selected covariate. This plot may be used to identify clusters in the data and to identify variables associated with prominent signal in the data. Variables that are associated with these leading principal components should be considered in downstream analyses.   
                                      
                                       
                                  
                                
                              
                            
                           
                            
                          
                           
                      
                    
                    
                      
                           
                          
                               Heatmaps    PCA     
                              
                                  Heatmaps  
                              
                               
                              
                                  Line  
                                  Day  
                              
                              
                            
                              
                                
                                  
                                       
                                      
                                           Heatmap of MicroRNA (miRNA) biogenesis Data   More Plot Information                      Heatmap of MicroRNA (miRNA) biogenesis Data  Heatmap of the normalized data, scaled to give all genes equal variance, generated via unsupervised clustering. Orange indicates high expression; blue indicates low expression. This plot is meant to provide a high level exploratory view of the data.   
                                      
                                       
                                  
                                
                              
                            
                            
                              
                                
                                  
                                       
                                      
                                           Principal Components of MicroRNA (miRNA) biogenesis Data   More Plot Information                      Principal Components of MicroRNA (miRNA) biogenesis Data  Principal component analysis maps high-dimensional datasets onto a smaller number of highly informative dimensions. Here, the first four principal components of the gene expression data are plotted against each other and colored by the values of the selected covariate. This plot may be used to identify clusters in the data and to identify variables associated with prominent signal in the data. Variables that are associated with these leading principal components should be considered in downstream analyses.   
                                      
                                       
                                  
                                
                                
                                  
                                       
                                      
                                           Principal Components of MicroRNA (miRNA) biogenesis Data   More Plot Information                      Principal Components of MicroRNA (miRNA) biogenesis Data  Principal component analysis maps high-dimensional datasets onto a smaller number of highly informative dimensions. Here, the first four principal components of the gene expression data are plotted against each other and colored by the values of the selected covariate. This plot may be used to identify clusters in the data and to identify variables associated with prominent signal in the data. Variables that are associated with these leading principal components should be considered in downstream analyses.   
                                      
                                       
                                  
                                
                              
                            
                           
                            
                          
                           
                      
                    
                    
                      
                           
                          
                               Heatmaps    PCA     
                              
                                  Heatmaps  
                              
                               
                              
                                  Line  
                                  Day  
                              
                              
                            
                              
                                
                                  
                                       
                                      
                                           Heatmap of Misspliced GSK3beta mutants stabilize beta-catenin Data   More Plot Information                      Heatmap of Misspliced GSK3beta mutants stabilize beta-catenin Data  Heatmap of the normalized data, scaled to give all genes equal variance, generated via unsupervised clustering. Orange indicates high expression; blue indicates low expression. This plot is meant to provide a high level exploratory view of the data.   
                                      
                                       
                                  
                                
                              
                            
                            
                              
                                
                                  
                                       
                                      
                                           Principal Components of Misspliced GSK3beta mutants stabilize beta-catenin Data   More Plot Information                      Principal Components of Misspliced GSK3beta mutants stabilize beta-catenin Data  Principal component analysis maps high-dimensional datasets onto a smaller number of highly informative dimensions. Here, the first four principal components of the gene expression data are plotted against each other and colored by the values of the selected covariate. This plot may be used to identify clusters in the data and to identify variables associated with prominent signal in the data. Variables that are associated with these leading principal components should be considered in downstream analyses.   
                                      
                                       
                                  
                                
                                
                                  
                                       
                                      
                                           Principal Components of Misspliced GSK3beta mutants stabilize beta-catenin Data   More Plot Information                      Principal Components of Misspliced GSK3beta mutants stabilize beta-catenin Data  Principal component analysis maps high-dimensional datasets onto a smaller number of highly informative dimensions. Here, the first four principal components of the gene expression data are plotted against each other and colored by the values of the selected covariate. This plot may be used to identify clusters in the data and to identify variables associated with prominent signal in the data. Variables that are associated with these leading principal components should be considered in downstream analyses.   
                                      
                                       
                                  
                                
                              
                            
                           
                            
                          
                           
                      
                    
                    
                      
                           
                          
                               Heatmaps    PCA     
                              
                                  Heatmaps  
                              
                               
                              
                                  Line  
                                  Day  
                              
                              
                            
                              
                                
                                  
                                       
                                      
                                           Heatmap of Molecules associated with elastic fibres Data   More Plot Information                      Heatmap of Molecules associated with elastic fibres Data  Heatmap of the normalized data, scaled to give all genes equal variance, generated via unsupervised clustering. Orange indicates high expression; blue indicates low expression. This plot is meant to provide a high level exploratory view of the data.   
                                      
                                       
                                  
                                
                              
                            
                            
                              
                                
                                  
                                       
                                      
                                           Principal Components of Molecules associated with elastic fibres Data   More Plot Information                      Principal Components of Molecules associated with elastic fibres Data  Principal component analysis maps high-dimensional datasets onto a smaller number of highly informative dimensions. Here, the first four principal components of the gene expression data are plotted against each other and colored by the values of the selected covariate. This plot may be used to identify clusters in the data and to identify variables associated with prominent signal in the data. Variables that are associated with these leading principal components should be considered in downstream analyses.   
                                      
                                       
                                  
                                
                                
                                  
                                       
                                      
                                           Principal Components of Molecules associated with elastic fibres Data   More Plot Information                      Principal Components of Molecules associated with elastic fibres Data  Principal component analysis maps high-dimensional datasets onto a smaller number of highly informative dimensions. Here, the first four principal components of the gene expression data are plotted against each other and colored by the values of the selected covariate. This plot may be used to identify clusters in the data and to identify variables associated with prominent signal in the data. Variables that are associated with these leading principal components should be considered in downstream analyses.   
                                      
                                       
                                  
                                
                              
                            
                           
                            
                          
                           
                      
                    
                    
                      
                           
                          
                               Heatmaps    PCA     
                              
                                  Heatmaps  
                              
                               
                              
                                  Line  
                                  Day  
                              
                              
                            
                              
                                
                                  
                                       
                                      
                                           Heatmap of Negative feedback regulation of MAPK pathway Data   More Plot Information                      Heatmap of Negative feedback regulation of MAPK pathway Data  Heatmap of the normalized data, scaled to give all genes equal variance, generated via unsupervised clustering. Orange indicates high expression; blue indicates low expression. This plot is meant to provide a high level exploratory view of the data.   
                                      
                                       
                                  
                                
                              
                            
                            
                              
                                
                                  
                                       
                                      
                                           Principal Components of Negative feedback regulation of MAPK pathway Data   More Plot Information                      Principal Components of Negative feedback regulation of MAPK pathway Data  Principal component analysis maps high-dimensional datasets onto a smaller number of highly informative dimensions. Here, the first four principal components of the gene expression data are plotted against each other and colored by the values of the selected covariate. This plot may be used to identify clusters in the data and to identify variables associated with prominent signal in the data. Variables that are associated with these leading principal components should be considered in downstream analyses.   
                                      
                                       
                                  
                                
                                
                                  
                                       
                                      
                                           Principal Components of Negative feedback regulation of MAPK pathway Data   More Plot Information                      Principal Components of Negative feedback regulation of MAPK pathway Data  Principal component analysis maps high-dimensional datasets onto a smaller number of highly informative dimensions. Here, the first four principal components of the gene expression data are plotted against each other and colored by the values of the selected covariate. This plot may be used to identify clusters in the data and to identify variables associated with prominent signal in the data. Variables that are associated with these leading principal components should be considered in downstream analyses.   
                                      
                                       
                                  
                                
                              
                            
                           
                            
                          
                           
                      
                    
                    
                      
                           
                          
                               Heatmaps    PCA     
                              
                                  Heatmaps  
                              
                               
                              
                                  Line  
                                  Day  
                              
                              
                            
                              
                                
                                  
                                       
                                      
                                           Heatmap of Negative regulation of FGFR1 signaling Data   More Plot Information                      Heatmap of Negative regulation of FGFR1 signaling Data  Heatmap of the normalized data, scaled to give all genes equal variance, generated via unsupervised clustering. Orange indicates high expression; blue indicates low expression. This plot is meant to provide a high level exploratory view of the data.   
                                      
                                       
                                  
                                
                              
                            
                            
                              
                                
                                  
                                       
                                      
                                           Principal Components of Negative regulation of FGFR1 signaling Data   More Plot Information                      Principal Components of Negative regulation of FGFR1 signaling Data  Principal component analysis maps high-dimensional datasets onto a smaller number of highly informative dimensions. Here, the first four principal components of the gene expression data are plotted against each other and colored by the values of the selected covariate. This plot may be used to identify clusters in the data and to identify variables associated with prominent signal in the data. Variables that are associated with these leading principal components should be considered in downstream analyses.   
                                      
                                       
                                  
                                
                                
                                  
                                       
                                      
                                           Principal Components of Negative regulation of FGFR1 signaling Data   More Plot Information                      Principal Components of Negative regulation of FGFR1 signaling Data  Principal component analysis maps high-dimensional datasets onto a smaller number of highly informative dimensions. Here, the first four principal components of the gene expression data are plotted against each other and colored by the values of the selected covariate. This plot may be used to identify clusters in the data and to identify variables associated with prominent signal in the data. Variables that are associated with these leading principal components should be considered in downstream analyses.   
                                      
                                       
                                  
                                
                              
                            
                           
                            
                          
                           
                      
                    
                    
                      
                           
                          
                               Heatmaps    PCA     
                              
                                  Heatmaps  
                              
                               
                              
                                  Line  
                                  Day  
                              
                              
                            
                              
                                
                                  
                                       
                                      
                                           Heatmap of Negative regulation of FGFR2 signaling Data   More Plot Information                      Heatmap of Negative regulation of FGFR2 signaling Data  Heatmap of the normalized data, scaled to give all genes equal variance, generated via unsupervised clustering. Orange indicates high expression; blue indicates low expression. This plot is meant to provide a high level exploratory view of the data.   
                                      
                                       
                                  
                                
                              
                            
                            
                              
                                
                                  
                                       
                                      
                                           Principal Components of Negative regulation of FGFR2 signaling Data   More Plot Information                      Principal Components of Negative regulation of FGFR2 signaling Data  Principal component analysis maps high-dimensional datasets onto a smaller number of highly informative dimensions. Here, the first four principal components of the gene expression data are plotted against each other and colored by the values of the selected covariate. This plot may be used to identify clusters in the data and to identify variables associated with prominent signal in the data. Variables that are associated with these leading principal components should be considered in downstream analyses.   
                                      
                                       
                                  
                                
                                
                                  
                                       
                                      
                                           Principal Components of Negative regulation of FGFR2 signaling Data   More Plot Information                      Principal Components of Negative regulation of FGFR2 signaling Data  Principal component analysis maps high-dimensional datasets onto a smaller number of highly informative dimensions. Here, the first four principal components of the gene expression data are plotted against each other and colored by the values of the selected covariate. This plot may be used to identify clusters in the data and to identify variables associated with prominent signal in the data. Variables that are associated with these leading principal components should be considered in downstream analyses.   
                                      
                                       
                                  
                                
                              
                            
                           
                            
                          
                           
                      
                    
                    
                      
                           
                          
                               Heatmaps    PCA     
                              
                                  Heatmaps  
                              
                               
                              
                                  Line  
                                  Day  
                              
                              
                            
                              
                                
                                  
                                       
                                      
                                           Heatmap of Negative regulation of FGFR3 signaling Data   More Plot Information                      Heatmap of Negative regulation of FGFR3 signaling Data  Heatmap of the normalized data, scaled to give all genes equal variance, generated via unsupervised clustering. Orange indicates high expression; blue indicates low expression. This plot is meant to provide a high level exploratory view of the data.   
                                      
                                       
                                  
                                
                              
                            
                            
                              
                                
                                  
                                       
                                      
                                           Principal Components of Negative regulation of FGFR3 signaling Data   More Plot Information                      Principal Components of Negative regulation of FGFR3 signaling Data  Principal component analysis maps high-dimensional datasets onto a smaller number of highly informative dimensions. Here, the first four principal components of the gene expression data are plotted against each other and colored by the values of the selected covariate. This plot may be used to identify clusters in the data and to identify variables associated with prominent signal in the data. Variables that are associated with these leading principal components should be considered in downstream analyses.   
                                      
                                       
                                  
                                
                                
                                  
                                       
                                      
                                           Principal Components of Negative regulation of FGFR3 signaling Data   More Plot Information                      Principal Components of Negative regulation of FGFR3 signaling Data  Principal component analysis maps high-dimensional datasets onto a smaller number of highly informative dimensions. Here, the first four principal components of the gene expression data are plotted against each other and colored by the values of the selected covariate. This plot may be used to identify clusters in the data and to identify variables associated with prominent signal in the data. Variables that are associated with these leading principal components should be considered in downstream analyses.   
                                      
                                       
                                  
                                
                              
                            
                           
                            
                          
                           
                      
                    
                    
                      
                           
                          
                               Heatmaps    PCA     
                              
                                  Heatmaps  
                              
                               
                              
                                  Line  
                                  Day  
                              
                              
                            
                              
                                
                                  
                                       
                                      
                                           Heatmap of Negative regulation of FGFR4 signaling Data   More Plot Information                      Heatmap of Negative regulation of FGFR4 signaling Data  Heatmap of the normalized data, scaled to give all genes equal variance, generated via unsupervised clustering. Orange indicates high expression; blue indicates low expression. This plot is meant to provide a high level exploratory view of the data.   
                                      
                                       
                                  
                                
                              
                            
                            
                              
                                
                                  
                                       
                                      
                                           Principal Components of Negative regulation of FGFR4 signaling Data   More Plot Information                      Principal Components of Negative regulation of FGFR4 signaling Data  Principal component analysis maps high-dimensional datasets onto a smaller number of highly informative dimensions. Here, the first four principal components of the gene expression data are plotted against each other and colored by the values of the selected covariate. This plot may be used to identify clusters in the data and to identify variables associated with prominent signal in the data. Variables that are associated with these leading principal components should be considered in downstream analyses.   
                                      
                                       
                                  
                                
                                
                                  
                                       
                                      
                                           Principal Components of Negative regulation of FGFR4 signaling Data   More Plot Information                      Principal Components of Negative regulation of FGFR4 signaling Data  Principal component analysis maps high-dimensional datasets onto a smaller number of highly informative dimensions. Here, the first four principal components of the gene expression data are plotted against each other and colored by the values of the selected covariate. This plot may be used to identify clusters in the data and to identify variables associated with prominent signal in the data. Variables that are associated with these leading principal components should be considered in downstream analyses.   
                                      
                                       
                                  
                                
                              
                            
                           
                            
                          
                           
                      
                    
                    
                      
                           
                          
                               Heatmaps    PCA     
                              
                                  Heatmaps  
                              
                               
                              
                                  Line  
                                  Day  
                              
                              
                            
                              
                                
                                  
                                       
                                      
                                           Heatmap of Negative regulation of MAPK pathway Data   More Plot Information                      Heatmap of Negative regulation of MAPK pathway Data  Heatmap of the normalized data, scaled to give all genes equal variance, generated via unsupervised clustering. Orange indicates high expression; blue indicates low expression. This plot is meant to provide a high level exploratory view of the data.   
                                      
                                       
                                  
                                
                              
                            
                            
                              
                                
                                  
                                       
                                      
                                           Principal Components of Negative regulation of MAPK pathway Data   More Plot Information                      Principal Components of Negative regulation of MAPK pathway Data  Principal component analysis maps high-dimensional datasets onto a smaller number of highly informative dimensions. Here, the first four principal components of the gene expression data are plotted against each other and colored by the values of the selected covariate. This plot may be used to identify clusters in the data and to identify variables associated with prominent signal in the data. Variables that are associated with these leading principal components should be considered in downstream analyses.   
                                      
                                       
                                  
                                
                                
                                  
                                       
                                      
                                           Principal Components of Negative regulation of MAPK pathway Data   More Plot Information                      Principal Components of Negative regulation of MAPK pathway Data  Principal component analysis maps high-dimensional datasets onto a smaller number of highly informative dimensions. Here, the first four principal components of the gene expression data are plotted against each other and colored by the values of the selected covariate. This plot may be used to identify clusters in the data and to identify variables associated with prominent signal in the data. Variables that are associated with these leading principal components should be considered in downstream analyses.   
                                      
                                       
                                  
                                
                              
                            
                           
                            
                          
                           
                      
                    
                    
                      
                           
                          
                               Heatmaps    PCA     
                              
                                  Heatmaps  
                              
                               
                              
                                  Line  
                                  Day  
                              
                              
                            
                              
                                
                                  
                                       
                                      
                                           Heatmap of Nephrin interactions Data   More Plot Information                      Heatmap of Nephrin interactions Data  Heatmap of the normalized data, scaled to give all genes equal variance, generated via unsupervised clustering. Orange indicates high expression; blue indicates low expression. This plot is meant to provide a high level exploratory view of the data.   
                                      
                                       
                                  
                                
                              
                            
                            
                              
                                
                                  
                                       
                                      
                                           Principal Components of Nephrin interactions Data   More Plot Information                      Principal Components of Nephrin interactions Data  Principal component analysis maps high-dimensional datasets onto a smaller number of highly informative dimensions. Here, the first four principal components of the gene expression data are plotted against each other and colored by the values of the selected covariate. This plot may be used to identify clusters in the data and to identify variables associated with prominent signal in the data. Variables that are associated with these leading principal components should be considered in downstream analyses.   
                                      
                                       
                                  
                                
                                
                                  
                                       
                                      
                                           Principal Components of Nephrin interactions Data   More Plot Information                      Principal Components of Nephrin interactions Data  Principal component analysis maps high-dimensional datasets onto a smaller number of highly informative dimensions. Here, the first four principal components of the gene expression data are plotted against each other and colored by the values of the selected covariate. This plot may be used to identify clusters in the data and to identify variables associated with prominent signal in the data. Variables that are associated with these leading principal components should be considered in downstream analyses.   
                                      
                                       
                                  
                                
                              
                            
                           
                            
                          
                           
                      
                    
                    
                      
                           
                          
                               Heatmaps    PCA     
                              
                                  Heatmaps  
                              
                               
                              
                                  Line  
                                  Day  
                              
                              
                            
                              
                                
                                  
                                       
                                      
                                           Heatmap of Non-integrin membrane-ECM interactions Data   More Plot Information                      Heatmap of Non-integrin membrane-ECM interactions Data  Heatmap of the normalized data, scaled to give all genes equal variance, generated via unsupervised clustering. Orange indicates high expression; blue indicates low expression. This plot is meant to provide a high level exploratory view of the data.   
                                      
                                       
                                  
                                
                              
                            
                            
                              
                                
                                  
                                       
                                      
                                           Principal Components of Non-integrin membrane-ECM interactions Data   More Plot Information                      Principal Components of Non-integrin membrane-ECM interactions Data  Principal component analysis maps high-dimensional datasets onto a smaller number of highly informative dimensions. Here, the first four principal components of the gene expression data are plotted against each other and colored by the values of the selected covariate. This plot may be used to identify clusters in the data and to identify variables associated with prominent signal in the data. Variables that are associated with these leading principal components should be considered in downstream analyses.   
                                      
                                       
                                  
                                
                                
                                  
                                       
                                      
                                           Principal Components of Non-integrin membrane-ECM interactions Data   More Plot Information                      Principal Components of Non-integrin membrane-ECM interactions Data  Principal component analysis maps high-dimensional datasets onto a smaller number of highly informative dimensions. Here, the first four principal components of the gene expression data are plotted against each other and colored by the values of the selected covariate. This plot may be used to identify clusters in the data and to identify variables associated with prominent signal in the data. Variables that are associated with these leading principal components should be considered in downstream analyses.   
                                      
                                       
                                  
                                
                              
                            
                           
                            
                          
                           
                      
                    
                    
                      
                           
                          
                               Heatmaps    PCA     
                              
                                  Heatmaps  
                              
                               
                              
                                  Line  
                                  Day  
                              
                              
                            
                              
                                
                                  
                                       
                                      
                                           Heatmap of Nonsense Mediated Decay (NMD) enhanced by the Exon Junction Complex (EJC) Data   More Plot Information                      Heatmap of Nonsense Mediated Decay (NMD) enhanced by the Exon Junction Complex (EJC) Data  Heatmap of the normalized data, scaled to give all genes equal variance, generated via unsupervised clustering. Orange indicates high expression; blue indicates low expression. This plot is meant to provide a high level exploratory view of the data.   
                                      
                                       
                                  
                                
                              
                            
                            
                              
                                
                                  
                                       
                                      
                                           Principal Components of Nonsense Mediated Decay (NMD) enhanced by the Exon Junction Complex (EJC) Data   More Plot Information                      Principal Components of Nonsense Mediated Decay (NMD) enhanced by the Exon Junction Complex (EJC) Data  Principal component analysis maps high-dimensional datasets onto a smaller number of highly informative dimensions. Here, the first four principal components of the gene expression data are plotted against each other and colored by the values of the selected covariate. This plot may be used to identify clusters in the data and to identify variables associated with prominent signal in the data. Variables that are associated with these leading principal components should be considered in downstream analyses.   
                                      
                                       
                                  
                                
                                
                                  
                                       
                                      
                                           Principal Components of Nonsense Mediated Decay (NMD) enhanced by the Exon Junction Complex (EJC) Data   More Plot Information                      Principal Components of Nonsense Mediated Decay (NMD) enhanced by the Exon Junction Complex (EJC) Data  Principal component analysis maps high-dimensional datasets onto a smaller number of highly informative dimensions. Here, the first four principal components of the gene expression data are plotted against each other and colored by the values of the selected covariate. This plot may be used to identify clusters in the data and to identify variables associated with prominent signal in the data. Variables that are associated with these leading principal components should be considered in downstream analyses.   
                                      
                                       
                                  
                                
                              
                            
                           
                            
                          
                           
                      
                    
                    
                      
                           
                          
                               Heatmaps    PCA     
                              
                                  Heatmaps  
                              
                               
                              
                                  Line  
                                  Day  
                              
                              
                            
                              
                                
                                  
                                       
                                      
                                           Heatmap of NRAGE signals death through JNK Data   More Plot Information                      Heatmap of NRAGE signals death through JNK Data  Heatmap of the normalized data, scaled to give all genes equal variance, generated via unsupervised clustering. Orange indicates high expression; blue indicates low expression. This plot is meant to provide a high level exploratory view of the data.   
                                      
                                       
                                  
                                
                              
                            
                            
                              
                                
                                  
                                       
                                      
                                           Principal Components of NRAGE signals death through JNK Data   More Plot Information                      Principal Components of NRAGE signals death through JNK Data  Principal component analysis maps high-dimensional datasets onto a smaller number of highly informative dimensions. Here, the first four principal components of the gene expression data are plotted against each other and colored by the values of the selected covariate. This plot may be used to identify clusters in the data and to identify variables associated with prominent signal in the data. Variables that are associated with these leading principal components should be considered in downstream analyses.   
                                      
                                       
                                  
                                
                                
                                  
                                       
                                      
                                           Principal Components of NRAGE signals death through JNK Data   More Plot Information                      Principal Components of NRAGE signals death through JNK Data  Principal component analysis maps high-dimensional datasets onto a smaller number of highly informative dimensions. Here, the first four principal components of the gene expression data are plotted against each other and colored by the values of the selected covariate. This plot may be used to identify clusters in the data and to identify variables associated with prominent signal in the data. Variables that are associated with these leading principal components should be considered in downstream analyses.   
                                      
                                       
                                  
                                
                              
                            
                           
                            
                          
                           
                      
                    
                    
                      
                           
                          
                               Heatmaps    PCA     
                              
                                  Heatmaps  
                              
                               
                              
                                  Line  
                                  Day  
                              
                              
                            
                              
                                
                                  
                                       
                                      
                                           Heatmap of p130Cas linkage to MAPK signaling for integrins Data   More Plot Information                      Heatmap of p130Cas linkage to MAPK signaling for integrins Data  Heatmap of the normalized data, scaled to give all genes equal variance, generated via unsupervised clustering. Orange indicates high expression; blue indicates low expression. This plot is meant to provide a high level exploratory view of the data.   
                                      
                                       
                                  
                                
                              
                            
                            
                              
                                
                                  
                                       
                                      
                                           Principal Components of p130Cas linkage to MAPK signaling for integrins Data   More Plot Information                      Principal Components of p130Cas linkage to MAPK signaling for integrins Data  Principal component analysis maps high-dimensional datasets onto a smaller number of highly informative dimensions. Here, the first four principal components of the gene expression data are plotted against each other and colored by the values of the selected covariate. This plot may be used to identify clusters in the data and to identify variables associated with prominent signal in the data. Variables that are associated with these leading principal components should be considered in downstream analyses.   
                                      
                                       
                                  
                                
                                
                                  
                                       
                                      
                                           Principal Components of p130Cas linkage to MAPK signaling for integrins Data   More Plot Information                      Principal Components of p130Cas linkage to MAPK signaling for integrins Data  Principal component analysis maps high-dimensional datasets onto a smaller number of highly informative dimensions. Here, the first four principal components of the gene expression data are plotted against each other and colored by the values of the selected covariate. This plot may be used to identify clusters in the data and to identify variables associated with prominent signal in the data. Variables that are associated with these leading principal components should be considered in downstream analyses.   
                                      
                                       
                                  
                                
                              
                            
                           
                            
                          
                           
                      
                    
                    
                      
                           
                          
                               Heatmaps    PCA     
                              
                                  Heatmaps  
                              
                               
                              
                                  Line  
                                  Day  
                              
                              
                            
                              
                                
                                  
                                       
                                      
                                           Heatmap of PCPCE pathway Data   More Plot Information                      Heatmap of PCPCE pathway Data  Heatmap of the normalized data, scaled to give all genes equal variance, generated via unsupervised clustering. Orange indicates high expression; blue indicates low expression. This plot is meant to provide a high level exploratory view of the data.   
                                      
                                       
                                  
                                
                              
                            
                            
                              
                                
                                  
                                       
                                      
                                           Principal Components of PCPCE pathway Data   More Plot Information                      Principal Components of PCPCE pathway Data  Principal component analysis maps high-dimensional datasets onto a smaller number of highly informative dimensions. Here, the first four principal components of the gene expression data are plotted against each other and colored by the values of the selected covariate. This plot may be used to identify clusters in the data and to identify variables associated with prominent signal in the data. Variables that are associated with these leading principal components should be considered in downstream analyses.   
                                      
                                       
                                  
                                
                                
                                  
                                       
                                      
                                           Principal Components of PCPCE pathway Data   More Plot Information                      Principal Components of PCPCE pathway Data  Principal component analysis maps high-dimensional datasets onto a smaller number of highly informative dimensions. Here, the first four principal components of the gene expression data are plotted against each other and colored by the values of the selected covariate. This plot may be used to identify clusters in the data and to identify variables associated with prominent signal in the data. Variables that are associated with these leading principal components should be considered in downstream analyses.   
                                      
                                       
                                  
                                
                              
                            
                           
                            
                          
                           
                      
                    
                    
                      
                           
                          
                               Heatmaps    PCA     
                              
                                  Heatmaps  
                              
                               
                              
                                  Line  
                                  Day  
                              
                              
                            
                              
                                
                                  
                                       
                                      
                                           Heatmap of PI3K Cascade Data   More Plot Information                      Heatmap of PI3K Cascade Data  Heatmap of the normalized data, scaled to give all genes equal variance, generated via unsupervised clustering. Orange indicates high expression; blue indicates low expression. This plot is meant to provide a high level exploratory view of the data.   
                                      
                                       
                                  
                                
                              
                            
                            
                              
                                
                                  
                                       
                                      
                                           Principal Components of PI3K Cascade Data   More Plot Information                      Principal Components of PI3K Cascade Data  Principal component analysis maps high-dimensional datasets onto a smaller number of highly informative dimensions. Here, the first four principal components of the gene expression data are plotted against each other and colored by the values of the selected covariate. This plot may be used to identify clusters in the data and to identify variables associated with prominent signal in the data. Variables that are associated with these leading principal components should be considered in downstream analyses.   
                                      
                                       
                                  
                                
                                
                                  
                                       
                                      
                                           Principal Components of PI3K Cascade Data   More Plot Information                      Principal Components of PI3K Cascade Data  Principal component analysis maps high-dimensional datasets onto a smaller number of highly informative dimensions. Here, the first four principal components of the gene expression data are plotted against each other and colored by the values of the selected covariate. This plot may be used to identify clusters in the data and to identify variables associated with prominent signal in the data. Variables that are associated with these leading principal components should be considered in downstream analyses.   
                                      
                                       
                                  
                                
                              
                            
                           
                            
                          
                           
                      
                    
                    
                      
                           
                          
                               Heatmaps    PCA     
                              
                                  Heatmaps  
                              
                               
                              
                                  Line  
                                  Day  
                              
                              
                            
                              
                                
                                  
                                       
                                      
                                           Heatmap of PI3KAKT activation Data   More Plot Information                      Heatmap of PI3KAKT activation Data  Heatmap of the normalized data, scaled to give all genes equal variance, generated via unsupervised clustering. Orange indicates high expression; blue indicates low expression. This plot is meant to provide a high level exploratory view of the data.   
                                      
                                       
                                  
                                
                              
                            
                            
                              
                                
                                  
                                       
                                      
                                           Principal Components of PI3KAKT activation Data   More Plot Information                      Principal Components of PI3KAKT activation Data  Principal component analysis maps high-dimensional datasets onto a smaller number of highly informative dimensions. Here, the first four principal components of the gene expression data are plotted against each other and colored by the values of the selected covariate. This plot may be used to identify clusters in the data and to identify variables associated with prominent signal in the data. Variables that are associated with these leading principal components should be considered in downstream analyses.   
                                      
                                       
                                  
                                
                                
                                  
                                       
                                      
                                           Principal Components of PI3KAKT activation Data   More Plot Information                      Principal Components of PI3KAKT activation Data  Principal component analysis maps high-dimensional datasets onto a smaller number of highly informative dimensions. Here, the first four principal components of the gene expression data are plotted against each other and colored by the values of the selected covariate. This plot may be used to identify clusters in the data and to identify variables associated with prominent signal in the data. Variables that are associated with these leading principal components should be considered in downstream analyses.   
                                      
                                       
                                  
                                
                              
                            
                           
                            
                          
                           
                      
                    
                    
                      
                           
                          
                               Heatmaps    PCA     
                              
                                  Heatmaps  
                              
                               
                              
                                  Line  
                                  Day  
                              
                              
                            
                              
                                
                                  
                                       
                                      
                                           Heatmap of PIP3 activates AKT signaling Data   More Plot Information                      Heatmap of PIP3 activates AKT signaling Data  Heatmap of the normalized data, scaled to give all genes equal variance, generated via unsupervised clustering. Orange indicates high expression; blue indicates low expression. This plot is meant to provide a high level exploratory view of the data.   
                                      
                                       
                                  
                                
                              
                            
                            
                              
                                
                                  
                                       
                                      
                                           Principal Components of PIP3 activates AKT signaling Data   More Plot Information                      Principal Components of PIP3 activates AKT signaling Data  Principal component analysis maps high-dimensional datasets onto a smaller number of highly informative dimensions. Here, the first four principal components of the gene expression data are plotted against each other and colored by the values of the selected covariate. This plot may be used to identify clusters in the data and to identify variables associated with prominent signal in the data. Variables that are associated with these leading principal components should be considered in downstream analyses.   
                                      
                                       
                                  
                                
                                
                                  
                                       
                                      
                                           Principal Components of PIP3 activates AKT signaling Data   More Plot Information                      Principal Components of PIP3 activates AKT signaling Data  Principal component analysis maps high-dimensional datasets onto a smaller number of highly informative dimensions. Here, the first four principal components of the gene expression data are plotted against each other and colored by the values of the selected covariate. This plot may be used to identify clusters in the data and to identify variables associated with prominent signal in the data. Variables that are associated with these leading principal components should be considered in downstream analyses.   
                                      
                                       
                                  
                                
                              
                            
                           
                            
                          
                           
                      
                    
                    
                      
                           
                          
                               Heatmaps    PCA     
                              
                                  Heatmaps  
                              
                               
                              
                                  Line  
                                  Day  
                              
                              
                            
                              
                                
                                  
                                       
                                      
                                           Heatmap of Platelet degranulation  Data   More Plot Information                      Heatmap of Platelet degranulation  Data  Heatmap of the normalized data, scaled to give all genes equal variance, generated via unsupervised clustering. Orange indicates high expression; blue indicates low expression. This plot is meant to provide a high level exploratory view of the data.   
                                      
                                       
                                  
                                
                              
                            
                            
                              
                                
                                  
                                       
                                      
                                           Principal Components of Platelet degranulation  Data   More Plot Information                      Principal Components of Platelet degranulation  Data  Principal component analysis maps high-dimensional datasets onto a smaller number of highly informative dimensions. Here, the first four principal components of the gene expression data are plotted against each other and colored by the values of the selected covariate. This plot may be used to identify clusters in the data and to identify variables associated with prominent signal in the data. Variables that are associated with these leading principal components should be considered in downstream analyses.   
                                      
                                       
                                  
                                
                                
                                  
                                       
                                      
                                           Principal Components of Platelet degranulation  Data   More Plot Information                      Principal Components of Platelet degranulation  Data  Principal component analysis maps high-dimensional datasets onto a smaller number of highly informative dimensions. Here, the first four principal components of the gene expression data are plotted against each other and colored by the values of the selected covariate. This plot may be used to identify clusters in the data and to identify variables associated with prominent signal in the data. Variables that are associated with these leading principal components should be considered in downstream analyses.   
                                      
                                       
                                  
                                
                              
                            
                           
                            
                          
                           
                      
                    
                    
                      
                           
                          
                               Heatmaps    PCA     
                              
                                  Heatmaps  
                              
                               
                              
                                  Line  
                                  Day  
                              
                              
                            
                              
                                
                                  
                                       
                                      
                                           Heatmap of RAF activation Data   More Plot Information                      Heatmap of RAF activation Data  Heatmap of the normalized data, scaled to give all genes equal variance, generated via unsupervised clustering. Orange indicates high expression; blue indicates low expression. This plot is meant to provide a high level exploratory view of the data.   
                                      
                                       
                                  
                                
                              
                            
                            
                              
                                
                                  
                                       
                                      
                                           Principal Components of RAF activation Data   More Plot Information                      Principal Components of RAF activation Data  Principal component analysis maps high-dimensional datasets onto a smaller number of highly informative dimensions. Here, the first four principal components of the gene expression data are plotted against each other and colored by the values of the selected covariate. This plot may be used to identify clusters in the data and to identify variables associated with prominent signal in the data. Variables that are associated with these leading principal components should be considered in downstream analyses.   
                                      
                                       
                                  
                                
                                
                                  
                                       
                                      
                                           Principal Components of RAF activation Data   More Plot Information                      Principal Components of RAF activation Data  Principal component analysis maps high-dimensional datasets onto a smaller number of highly informative dimensions. Here, the first four principal components of the gene expression data are plotted against each other and colored by the values of the selected covariate. This plot may be used to identify clusters in the data and to identify variables associated with prominent signal in the data. Variables that are associated with these leading principal components should be considered in downstream analyses.   
                                      
                                       
                                  
                                
                              
                            
                           
                            
                          
                           
                      
                    
                    
                      
                           
                          
                               Heatmaps    PCA     
                              
                                  Heatmaps  
                              
                               
                              
                                  Line  
                                  Day  
                              
                              
                            
                              
                                
                                  
                                       
                                      
                                           Heatmap of RAFMAP kinase cascade Data   More Plot Information                      Heatmap of RAFMAP kinase cascade Data  Heatmap of the normalized data, scaled to give all genes equal variance, generated via unsupervised clustering. Orange indicates high expression; blue indicates low expression. This plot is meant to provide a high level exploratory view of the data.   
                                      
                                       
                                  
                                
                              
                            
                            
                              
                                
                                  
                                       
                                      
                                           Principal Components of RAFMAP kinase cascade Data   More Plot Information                      Principal Components of RAFMAP kinase cascade Data  Principal component analysis maps high-dimensional datasets onto a smaller number of highly informative dimensions. Here, the first four principal components of the gene expression data are plotted against each other and colored by the values of the selected covariate. This plot may be used to identify clusters in the data and to identify variables associated with prominent signal in the data. Variables that are associated with these leading principal components should be considered in downstream analyses.   
                                      
                                       
                                  
                                
                                
                                  
                                       
                                      
                                           Principal Components of RAFMAP kinase cascade Data   More Plot Information                      Principal Components of RAFMAP kinase cascade Data  Principal component analysis maps high-dimensional datasets onto a smaller number of highly informative dimensions. Here, the first four principal components of the gene expression data are plotted against each other and colored by the values of the selected covariate. This plot may be used to identify clusters in the data and to identify variables associated with prominent signal in the data. Variables that are associated with these leading principal components should be considered in downstream analyses.   
                                      
                                       
                                  
                                
                              
                            
                           
                            
                          
                           
                      
                    
                    
                      
                           
                          
                               Heatmaps    PCA     
                              
                                  Heatmaps  
                              
                               
                              
                                  Line  
                                  Day  
                              
                              
                            
                              
                                
                                  
                                       
                                      
                                           Heatmap of Regulation of actin dynamics for phagocytic cup formation Data   More Plot Information                      Heatmap of Regulation of actin dynamics for phagocytic cup formation Data  Heatmap of the normalized data, scaled to give all genes equal variance, generated via unsupervised clustering. Orange indicates high expression; blue indicates low expression. This plot is meant to provide a high level exploratory view of the data.   
                                      
                                       
                                  
                                
                              
                            
                            
                              
                                
                                  
                                       
                                      
                                           Principal Components of Regulation of actin dynamics for phagocytic cup formation Data   More Plot Information                      Principal Components of Regulation of actin dynamics for phagocytic cup formation Data  Principal component analysis maps high-dimensional datasets onto a smaller number of highly informative dimensions. Here, the first four principal components of the gene expression data are plotted against each other and colored by the values of the selected covariate. This plot may be used to identify clusters in the data and to identify variables associated with prominent signal in the data. Variables that are associated with these leading principal components should be considered in downstream analyses.   
                                      
                                       
                                  
                                
                                
                                  
                                       
                                      
                                           Principal Components of Regulation of actin dynamics for phagocytic cup formation Data   More Plot Information                      Principal Components of Regulation of actin dynamics for phagocytic cup formation Data  Principal component analysis maps high-dimensional datasets onto a smaller number of highly informative dimensions. Here, the first four principal components of the gene expression data are plotted against each other and colored by the values of the selected covariate. This plot may be used to identify clusters in the data and to identify variables associated with prominent signal in the data. Variables that are associated with these leading principal components should be considered in downstream analyses.   
                                      
                                       
                                  
                                
                              
                            
                           
                            
                          
                           
                      
                    
                    
                      
                           
                          
                               Heatmaps    PCA     
                              
                                  Heatmaps  
                              
                               
                              
                                  Line  
                                  Day  
                              
                              
                            
                              
                                
                                  
                                       
                                      
                                           Heatmap of Regulation of HSF1-mediated heat shock response Data   More Plot Information                      Heatmap of Regulation of HSF1-mediated heat shock response Data  Heatmap of the normalized data, scaled to give all genes equal variance, generated via unsupervised clustering. Orange indicates high expression; blue indicates low expression. This plot is meant to provide a high level exploratory view of the data.   
                                      
                                       
                                  
                                
                              
                            
                            
                              
                                
                                  
                                       
                                      
                                           Principal Components of Regulation of HSF1-mediated heat shock response Data   More Plot Information                      Principal Components of Regulation of HSF1-mediated heat shock response Data  Principal component analysis maps high-dimensional datasets onto a smaller number of highly informative dimensions. Here, the first four principal components of the gene expression data are plotted against each other and colored by the values of the selected covariate. This plot may be used to identify clusters in the data and to identify variables associated with prominent signal in the data. Variables that are associated with these leading principal components should be considered in downstream analyses.   
                                      
                                       
                                  
                                
                                
                                  
                                       
                                      
                                           Principal Components of Regulation of HSF1-mediated heat shock response Data   More Plot Information                      Principal Components of Regulation of HSF1-mediated heat shock response Data  Principal component analysis maps high-dimensional datasets onto a smaller number of highly informative dimensions. Here, the first four principal components of the gene expression data are plotted against each other and colored by the values of the selected covariate. This plot may be used to identify clusters in the data and to identify variables associated with prominent signal in the data. Variables that are associated with these leading principal components should be considered in downstream analyses.   
                                      
                                       
                                  
                                
                              
                            
                           
                            
                          
                           
                      
                    
                    
                      
                           
                          
                               Heatmaps    PCA     
                              
                                  Heatmaps  
                              
                               
                              
                                  Line  
                                  Day  
                              
                              
                            
                              
                                
                                  
                                       
                                      
                                           Heatmap of Regulation of signaling by CBL Data   More Plot Information                      Heatmap of Regulation of signaling by CBL Data  Heatmap of the normalized data, scaled to give all genes equal variance, generated via unsupervised clustering. Orange indicates high expression; blue indicates low expression. This plot is meant to provide a high level exploratory view of the data.   
                                      
                                       
                                  
                                
                              
                            
                            
                              
                                
                                  
                                       
                                      
                                           Principal Components of Regulation of signaling by CBL Data   More Plot Information                      Principal Components of Regulation of signaling by CBL Data  Principal component analysis maps high-dimensional datasets onto a smaller number of highly informative dimensions. Here, the first four principal components of the gene expression data are plotted against each other and colored by the values of the selected covariate. This plot may be used to identify clusters in the data and to identify variables associated with prominent signal in the data. Variables that are associated with these leading principal components should be considered in downstream analyses.   
                                      
                                       
                                  
                                
                                
                                  
                                       
                                      
                                           Principal Components of Regulation of signaling by CBL Data   More Plot Information                      Principal Components of Regulation of signaling by CBL Data  Principal component analysis maps high-dimensional datasets onto a smaller number of highly informative dimensions. Here, the first four principal components of the gene expression data are plotted against each other and colored by the values of the selected covariate. This plot may be used to identify clusters in the data and to identify variables associated with prominent signal in the data. Variables that are associated with these leading principal components should be considered in downstream analyses.   
                                      
                                       
                                  
                                
                              
                            
                           
                            
                          
                           
                      
                    
                    
                      
                           
                          
                               Heatmaps    PCA     
                              
                                  Heatmaps  
                              
                               
                              
                                  Line  
                                  Day  
                              
                              
                            
                              
                                
                                  
                                       
                                      
                                           Heatmap of Repression of WNT target genes Data   More Plot Information                      Heatmap of Repression of WNT target genes Data  Heatmap of the normalized data, scaled to give all genes equal variance, generated via unsupervised clustering. Orange indicates high expression; blue indicates low expression. This plot is meant to provide a high level exploratory view of the data.   
                                      
                                       
                                  
                                
                              
                            
                            
                              
                                
                                  
                                       
                                      
                                           Principal Components of Repression of WNT target genes Data   More Plot Information                      Principal Components of Repression of WNT target genes Data  Principal component analysis maps high-dimensional datasets onto a smaller number of highly informative dimensions. Here, the first four principal components of the gene expression data are plotted against each other and colored by the values of the selected covariate. This plot may be used to identify clusters in the data and to identify variables associated with prominent signal in the data. Variables that are associated with these leading principal components should be considered in downstream analyses.   
                                      
                                       
                                  
                                
                                
                                  
                                       
                                      
                                           Principal Components of Repression of WNT target genes Data   More Plot Information                      Principal Components of Repression of WNT target genes Data  Principal component analysis maps high-dimensional datasets onto a smaller number of highly informative dimensions. Here, the first four principal components of the gene expression data are plotted against each other and colored by the values of the selected covariate. This plot may be used to identify clusters in the data and to identify variables associated with prominent signal in the data. Variables that are associated with these leading principal components should be considered in downstream analyses.   
                                      
                                       
                                  
                                
                              
                            
                           
                            
                          
                           
                      
                    
                    
                      
                           
                          
                               Heatmaps    PCA     
                              
                                  Heatmaps  
                              
                               
                              
                                  Line  
                                  Day  
                              
                              
                            
                              
                                
                                  
                                       
                                      
                                           Heatmap of Resolution of Sister Chromatid Cohesion Data   More Plot Information                      Heatmap of Resolution of Sister Chromatid Cohesion Data  Heatmap of the normalized data, scaled to give all genes equal variance, generated via unsupervised clustering. Orange indicates high expression; blue indicates low expression. This plot is meant to provide a high level exploratory view of the data.   
                                      
                                       
                                  
                                
                              
                            
                            
                              
                                
                                  
                                       
                                      
                                           Principal Components of Resolution of Sister Chromatid Cohesion Data   More Plot Information                      Principal Components of Resolution of Sister Chromatid Cohesion Data  Principal component analysis maps high-dimensional datasets onto a smaller number of highly informative dimensions. Here, the first four principal components of the gene expression data are plotted against each other and colored by the values of the selected covariate. This plot may be used to identify clusters in the data and to identify variables associated with prominent signal in the data. Variables that are associated with these leading principal components should be considered in downstream analyses.   
                                      
                                       
                                  
                                
                                
                                  
                                       
                                      
                                           Principal Components of Resolution of Sister Chromatid Cohesion Data   More Plot Information                      Principal Components of Resolution of Sister Chromatid Cohesion Data  Principal component analysis maps high-dimensional datasets onto a smaller number of highly informative dimensions. Here, the first four principal components of the gene expression data are plotted against each other and colored by the values of the selected covariate. This plot may be used to identify clusters in the data and to identify variables associated with prominent signal in the data. Variables that are associated with these leading principal components should be considered in downstream analyses.   
                                      
                                       
                                  
                                
                              
                            
                           
                            
                          
                           
                      
                    
                    
                      
                           
                          
                               Heatmaps    PCA     
                              
                                  Heatmaps  
                              
                               
                              
                                  Line  
                                  Day  
                              
                              
                            
                              
                                
                                  
                                       
                                      
                                           Heatmap of Rho GTPase cycle Data   More Plot Information                      Heatmap of Rho GTPase cycle Data  Heatmap of the normalized data, scaled to give all genes equal variance, generated via unsupervised clustering. Orange indicates high expression; blue indicates low expression. This plot is meant to provide a high level exploratory view of the data.   
                                      
                                       
                                  
                                
                              
                            
                            
                              
                                
                                  
                                       
                                      
                                           Principal Components of Rho GTPase cycle Data   More Plot Information                      Principal Components of Rho GTPase cycle Data  Principal component analysis maps high-dimensional datasets onto a smaller number of highly informative dimensions. Here, the first four principal components of the gene expression data are plotted against each other and colored by the values of the selected covariate. This plot may be used to identify clusters in the data and to identify variables associated with prominent signal in the data. Variables that are associated with these leading principal components should be considered in downstream analyses.   
                                      
                                       
                                  
                                
                                
                                  
                                       
                                      
                                           Principal Components of Rho GTPase cycle Data   More Plot Information                      Principal Components of Rho GTPase cycle Data  Principal component analysis maps high-dimensional datasets onto a smaller number of highly informative dimensions. Here, the first four principal components of the gene expression data are plotted against each other and colored by the values of the selected covariate. This plot may be used to identify clusters in the data and to identify variables associated with prominent signal in the data. Variables that are associated with these leading principal components should be considered in downstream analyses.   
                                      
                                       
                                  
                                
                              
                            
                           
                            
                          
                           
                      
                    
                    
                      
                           
                          
                               Heatmaps    PCA     
                              
                                  Heatmaps  
                              
                               
                              
                                  Line  
                                  Day  
                              
                              
                            
                              
                                
                                  
                                       
                                      
                                           Heatmap of RHO GTPases activate CIT Data   More Plot Information                      Heatmap of RHO GTPases activate CIT Data  Heatmap of the normalized data, scaled to give all genes equal variance, generated via unsupervised clustering. Orange indicates high expression; blue indicates low expression. This plot is meant to provide a high level exploratory view of the data.   
                                      
                                       
                                  
                                
                              
                            
                            
                              
                                
                                  
                                       
                                      
                                           Principal Components of RHO GTPases activate CIT Data   More Plot Information                      Principal Components of RHO GTPases activate CIT Data  Principal component analysis maps high-dimensional datasets onto a smaller number of highly informative dimensions. Here, the first four principal components of the gene expression data are plotted against each other and colored by the values of the selected covariate. This plot may be used to identify clusters in the data and to identify variables associated with prominent signal in the data. Variables that are associated with these leading principal components should be considered in downstream analyses.   
                                      
                                       
                                  
                                
                                
                                  
                                       
                                      
                                           Principal Components of RHO GTPases activate CIT Data   More Plot Information                      Principal Components of RHO GTPases activate CIT Data  Principal component analysis maps high-dimensional datasets onto a smaller number of highly informative dimensions. Here, the first four principal components of the gene expression data are plotted against each other and colored by the values of the selected covariate. This plot may be used to identify clusters in the data and to identify variables associated with prominent signal in the data. Variables that are associated with these leading principal components should be considered in downstream analyses.   
                                      
                                       
                                  
                                
                              
                            
                           
                            
                          
                           
                      
                    
                    
                      
                           
                          
                               Heatmaps    PCA     
                              
                                  Heatmaps  
                              
                               
                              
                                  Line  
                                  Day  
                              
                              
                            
                              
                                
                                  
                                       
                                      
                                           Heatmap of RHO GTPases Activate Formins Data   More Plot Information                      Heatmap of RHO GTPases Activate Formins Data  Heatmap of the normalized data, scaled to give all genes equal variance, generated via unsupervised clustering. Orange indicates high expression; blue indicates low expression. This plot is meant to provide a high level exploratory view of the data.   
                                      
                                       
                                  
                                
                              
                            
                            
                              
                                
                                  
                                       
                                      
                                           Principal Components of RHO GTPases Activate Formins Data   More Plot Information                      Principal Components of RHO GTPases Activate Formins Data  Principal component analysis maps high-dimensional datasets onto a smaller number of highly informative dimensions. Here, the first four principal components of the gene expression data are plotted against each other and colored by the values of the selected covariate. This plot may be used to identify clusters in the data and to identify variables associated with prominent signal in the data. Variables that are associated with these leading principal components should be considered in downstream analyses.   
                                      
                                       
                                  
                                
                                
                                  
                                       
                                      
                                           Principal Components of RHO GTPases Activate Formins Data   More Plot Information                      Principal Components of RHO GTPases Activate Formins Data  Principal component analysis maps high-dimensional datasets onto a smaller number of highly informative dimensions. Here, the first four principal components of the gene expression data are plotted against each other and colored by the values of the selected covariate. This plot may be used to identify clusters in the data and to identify variables associated with prominent signal in the data. Variables that are associated with these leading principal components should be considered in downstream analyses.   
                                      
                                       
                                  
                                
                              
                            
                           
                            
                          
                           
                      
                    
                    
                      
                           
                          
                               Heatmaps    PCA     
                              
                                  Heatmaps  
                              
                               
                              
                                  Line  
                                  Day  
                              
                              
                            
                              
                                
                                  
                                       
                                      
                                           Heatmap of RHO GTPases activate IQGAPs Data   More Plot Information                      Heatmap of RHO GTPases activate IQGAPs Data  Heatmap of the normalized data, scaled to give all genes equal variance, generated via unsupervised clustering. Orange indicates high expression; blue indicates low expression. This plot is meant to provide a high level exploratory view of the data.   
                                      
                                       
                                  
                                
                              
                            
                            
                              
                                
                                  
                                       
                                      
                                           Principal Components of RHO GTPases activate IQGAPs Data   More Plot Information                      Principal Components of RHO GTPases activate IQGAPs Data  Principal component analysis maps high-dimensional datasets onto a smaller number of highly informative dimensions. Here, the first four principal components of the gene expression data are plotted against each other and colored by the values of the selected covariate. This plot may be used to identify clusters in the data and to identify variables associated with prominent signal in the data. Variables that are associated with these leading principal components should be considered in downstream analyses.   
                                      
                                       
                                  
                                
                                
                                  
                                       
                                      
                                           Principal Components of RHO GTPases activate IQGAPs Data   More Plot Information                      Principal Components of RHO GTPases activate IQGAPs Data  Principal component analysis maps high-dimensional datasets onto a smaller number of highly informative dimensions. Here, the first four principal components of the gene expression data are plotted against each other and colored by the values of the selected covariate. This plot may be used to identify clusters in the data and to identify variables associated with prominent signal in the data. Variables that are associated with these leading principal components should be considered in downstream analyses.   
                                      
                                       
                                  
                                
                              
                            
                           
                            
                          
                           
                      
                    
                    
                      
                           
                          
                               Heatmaps    PCA     
                              
                                  Heatmaps  
                              
                               
                              
                                  Line  
                                  Day  
                              
                              
                            
                              
                                
                                  
                                       
                                      
                                           Heatmap of RHO GTPases activate PAKs Data   More Plot Information                      Heatmap of RHO GTPases activate PAKs Data  Heatmap of the normalized data, scaled to give all genes equal variance, generated via unsupervised clustering. Orange indicates high expression; blue indicates low expression. This plot is meant to provide a high level exploratory view of the data.   
                                      
                                       
                                  
                                
                              
                            
                            
                              
                                
                                  
                                       
                                      
                                           Principal Components of RHO GTPases activate PAKs Data   More Plot Information                      Principal Components of RHO GTPases activate PAKs Data  Principal component analysis maps high-dimensional datasets onto a smaller number of highly informative dimensions. Here, the first four principal components of the gene expression data are plotted against each other and colored by the values of the selected covariate. This plot may be used to identify clusters in the data and to identify variables associated with prominent signal in the data. Variables that are associated with these leading principal components should be considered in downstream analyses.   
                                      
                                       
                                  
                                
                                
                                  
                                       
                                      
                                           Principal Components of RHO GTPases activate PAKs Data   More Plot Information                      Principal Components of RHO GTPases activate PAKs Data  Principal component analysis maps high-dimensional datasets onto a smaller number of highly informative dimensions. Here, the first four principal components of the gene expression data are plotted against each other and colored by the values of the selected covariate. This plot may be used to identify clusters in the data and to identify variables associated with prominent signal in the data. Variables that are associated with these leading principal components should be considered in downstream analyses.   
                                      
                                       
                                  
                                
                              
                            
                           
                            
                          
                           
                      
                    
                    
                      
                           
                          
                               Heatmaps    PCA     
                              
                                  Heatmaps  
                              
                               
                              
                                  Line  
                                  Day  
                              
                              
                            
                              
                                
                                  
                                       
                                      
                                           Heatmap of RHO GTPases activate PKNs Data   More Plot Information                      Heatmap of RHO GTPases activate PKNs Data  Heatmap of the normalized data, scaled to give all genes equal variance, generated via unsupervised clustering. Orange indicates high expression; blue indicates low expression. This plot is meant to provide a high level exploratory view of the data.   
                                      
                                       
                                  
                                
                              
                            
                            
                              
                                
                                  
                                       
                                      
                                           Principal Components of RHO GTPases activate PKNs Data   More Plot Information                      Principal Components of RHO GTPases activate PKNs Data  Principal component analysis maps high-dimensional datasets onto a smaller number of highly informative dimensions. Here, the first four principal components of the gene expression data are plotted against each other and colored by the values of the selected covariate. This plot may be used to identify clusters in the data and to identify variables associated with prominent signal in the data. Variables that are associated with these leading principal components should be considered in downstream analyses.   
                                      
                                       
                                  
                                
                                
                                  
                                       
                                      
                                           Principal Components of RHO GTPases activate PKNs Data   More Plot Information                      Principal Components of RHO GTPases activate PKNs Data  Principal component analysis maps high-dimensional datasets onto a smaller number of highly informative dimensions. Here, the first four principal components of the gene expression data are plotted against each other and colored by the values of the selected covariate. This plot may be used to identify clusters in the data and to identify variables associated with prominent signal in the data. Variables that are associated with these leading principal components should be considered in downstream analyses.   
                                      
                                       
                                  
                                
                              
                            
                           
                            
                          
                           
                      
                    
                    
                      
                           
                          
                               Heatmaps    PCA     
                              
                                  Heatmaps  
                              
                               
                              
                                  Line  
                                  Day  
                              
                              
                            
                              
                                
                                  
                                       
                                      
                                           Heatmap of RHO GTPases Activate ROCKs Data   More Plot Information                      Heatmap of RHO GTPases Activate ROCKs Data  Heatmap of the normalized data, scaled to give all genes equal variance, generated via unsupervised clustering. Orange indicates high expression; blue indicates low expression. This plot is meant to provide a high level exploratory view of the data.   
                                      
                                       
                                  
                                
                              
                            
                            
                              
                                
                                  
                                       
                                      
                                           Principal Components of RHO GTPases Activate ROCKs Data   More Plot Information                      Principal Components of RHO GTPases Activate ROCKs Data  Principal component analysis maps high-dimensional datasets onto a smaller number of highly informative dimensions. Here, the first four principal components of the gene expression data are plotted against each other and colored by the values of the selected covariate. This plot may be used to identify clusters in the data and to identify variables associated with prominent signal in the data. Variables that are associated with these leading principal components should be considered in downstream analyses.   
                                      
                                       
                                  
                                
                                
                                  
                                       
                                      
                                           Principal Components of RHO GTPases Activate ROCKs Data   More Plot Information                      Principal Components of RHO GTPases Activate ROCKs Data  Principal component analysis maps high-dimensional datasets onto a smaller number of highly informative dimensions. Here, the first four principal components of the gene expression data are plotted against each other and colored by the values of the selected covariate. This plot may be used to identify clusters in the data and to identify variables associated with prominent signal in the data. Variables that are associated with these leading principal components should be considered in downstream analyses.   
                                      
                                       
                                  
                                
                              
                            
                           
                            
                          
                           
                      
                    
                    
                      
                           
                          
                               Heatmaps    PCA     
                              
                                  Heatmaps  
                              
                               
                              
                                  Line  
                                  Day  
                              
                              
                            
                              
                                
                                  
                                       
                                      
                                           Heatmap of RHO GTPases Activate WASPs and WAVEs Data   More Plot Information                      Heatmap of RHO GTPases Activate WASPs and WAVEs Data  Heatmap of the normalized data, scaled to give all genes equal variance, generated via unsupervised clustering. Orange indicates high expression; blue indicates low expression. This plot is meant to provide a high level exploratory view of the data.   
                                      
                                       
                                  
                                
                              
                            
                            
                              
                                
                                  
                                       
                                      
                                           Principal Components of RHO GTPases Activate WASPs and WAVEs Data   More Plot Information                      Principal Components of RHO GTPases Activate WASPs and WAVEs Data  Principal component analysis maps high-dimensional datasets onto a smaller number of highly informative dimensions. Here, the first four principal components of the gene expression data are plotted against each other and colored by the values of the selected covariate. This plot may be used to identify clusters in the data and to identify variables associated with prominent signal in the data. Variables that are associated with these leading principal components should be considered in downstream analyses.   
                                      
                                       
                                  
                                
                                
                                  
                                       
                                      
                                           Principal Components of RHO GTPases Activate WASPs and WAVEs Data   More Plot Information                      Principal Components of RHO GTPases Activate WASPs and WAVEs Data  Principal component analysis maps high-dimensional datasets onto a smaller number of highly informative dimensions. Here, the first four principal components of the gene expression data are plotted against each other and colored by the values of the selected covariate. This plot may be used to identify clusters in the data and to identify variables associated with prominent signal in the data. Variables that are associated with these leading principal components should be considered in downstream analyses.   
                                      
                                       
                                  
                                
                              
                            
                           
                            
                          
                           
                      
                    
                    
                      
                           
                          
                               Heatmaps    PCA     
                              
                                  Heatmaps  
                              
                               
                              
                                  Line  
                                  Day  
                              
                              
                            
                              
                                
                                  
                                       
                                      
                                           Heatmap of Role of LAT2NTALLAB on calcium mobilization Data   More Plot Information                      Heatmap of Role of LAT2NTALLAB on calcium mobilization Data  Heatmap of the normalized data, scaled to give all genes equal variance, generated via unsupervised clustering. Orange indicates high expression; blue indicates low expression. This plot is meant to provide a high level exploratory view of the data.   
                                      
                                       
                                  
                                
                              
                            
                            
                              
                                
                                  
                                       
                                      
                                           Principal Components of Role of LAT2NTALLAB on calcium mobilization Data   More Plot Information                      Principal Components of Role of LAT2NTALLAB on calcium mobilization Data  Principal component analysis maps high-dimensional datasets onto a smaller number of highly informative dimensions. Here, the first four principal components of the gene expression data are plotted against each other and colored by the values of the selected covariate. This plot may be used to identify clusters in the data and to identify variables associated with prominent signal in the data. Variables that are associated with these leading principal components should be considered in downstream analyses.   
                                      
                                       
                                  
                                
                                
                                  
                                       
                                      
                                           Principal Components of Role of LAT2NTALLAB on calcium mobilization Data   More Plot Information                      Principal Components of Role of LAT2NTALLAB on calcium mobilization Data  Principal component analysis maps high-dimensional datasets onto a smaller number of highly informative dimensions. Here, the first four principal components of the gene expression data are plotted against each other and colored by the values of the selected covariate. This plot may be used to identify clusters in the data and to identify variables associated with prominent signal in the data. Variables that are associated with these leading principal components should be considered in downstream analyses.   
                                      
                                       
                                  
                                
                              
                            
                           
                            
                          
                           
                      
                    
                    
                      
                           
                          
                               Heatmaps    PCA     
                              
                                  Heatmaps  
                              
                               
                              
                                  Line  
                                  Day  
                              
                              
                            
                              
                                
                                  
                                       
                                      
                                           Heatmap of S33 mutants of beta-catenin aren.SQT.t phosphorylated Data   More Plot Information                      Heatmap of S33 mutants of beta-catenin aren.SQT.t phosphorylated Data  Heatmap of the normalized data, scaled to give all genes equal variance, generated via unsupervised clustering. Orange indicates high expression; blue indicates low expression. This plot is meant to provide a high level exploratory view of the data.   
                                      
                                       
                                  
                                
                              
                            
                            
                              
                                
                                  
                                       
                                      
                                           Principal Components of S33 mutants of beta-catenin aren.SQT.t phosphorylated Data   More Plot Information                      Principal Components of S33 mutants of beta-catenin aren.SQT.t phosphorylated Data  Principal component analysis maps high-dimensional datasets onto a smaller number of highly informative dimensions. Here, the first four principal components of the gene expression data are plotted against each other and colored by the values of the selected covariate. This plot may be used to identify clusters in the data and to identify variables associated with prominent signal in the data. Variables that are associated with these leading principal components should be considered in downstream analyses.   
                                      
                                       
                                  
                                
                                
                                  
                                       
                                      
                                           Principal Components of S33 mutants of beta-catenin aren.SQT.t phosphorylated Data   More Plot Information                      Principal Components of S33 mutants of beta-catenin aren.SQT.t phosphorylated Data  Principal component analysis maps high-dimensional datasets onto a smaller number of highly informative dimensions. Here, the first four principal components of the gene expression data are plotted against each other and colored by the values of the selected covariate. This plot may be used to identify clusters in the data and to identify variables associated with prominent signal in the data. Variables that are associated with these leading principal components should be considered in downstream analyses.   
                                      
                                       
                                  
                                
                              
                            
                           
                            
                          
                           
                      
                    
                    
                      
                           
                          
                               Heatmaps    PCA     
                              
                                  Heatmaps  
                              
                               
                              
                                  Line  
                                  Day  
                              
                              
                            
                              
                                
                                  
                                       
                                      
                                           Heatmap of S37 mutants of beta-catenin aren.SQT.t phosphorylated Data   More Plot Information                      Heatmap of S37 mutants of beta-catenin aren.SQT.t phosphorylated Data  Heatmap of the normalized data, scaled to give all genes equal variance, generated via unsupervised clustering. Orange indicates high expression; blue indicates low expression. This plot is meant to provide a high level exploratory view of the data.   
                                      
                                       
                                  
                                
                              
                            
                            
                              
                                
                                  
                                       
                                      
                                           Principal Components of S37 mutants of beta-catenin aren.SQT.t phosphorylated Data   More Plot Information                      Principal Components of S37 mutants of beta-catenin aren.SQT.t phosphorylated Data  Principal component analysis maps high-dimensional datasets onto a smaller number of highly informative dimensions. Here, the first four principal components of the gene expression data are plotted against each other and colored by the values of the selected covariate. This plot may be used to identify clusters in the data and to identify variables associated with prominent signal in the data. Variables that are associated with these leading principal components should be considered in downstream analyses.   
                                      
                                       
                                  
                                
                                
                                  
                                       
                                      
                                           Principal Components of S37 mutants of beta-catenin aren.SQT.t phosphorylated Data   More Plot Information                      Principal Components of S37 mutants of beta-catenin aren.SQT.t phosphorylated Data  Principal component analysis maps high-dimensional datasets onto a smaller number of highly informative dimensions. Here, the first four principal components of the gene expression data are plotted against each other and colored by the values of the selected covariate. This plot may be used to identify clusters in the data and to identify variables associated with prominent signal in the data. Variables that are associated with these leading principal components should be considered in downstream analyses.   
                                      
                                       
                                  
                                
                              
                            
                           
                            
                          
                           
                      
                    
                    
                      
                           
                          
                               Heatmaps    PCA     
                              
                                  Heatmaps  
                              
                               
                              
                                  Line  
                                  Day  
                              
                              
                            
                              
                                
                                  
                                       
                                      
                                           Heatmap of S45 mutants of beta-catenin aren.SQT.t phosphorylated Data   More Plot Information                      Heatmap of S45 mutants of beta-catenin aren.SQT.t phosphorylated Data  Heatmap of the normalized data, scaled to give all genes equal variance, generated via unsupervised clustering. Orange indicates high expression; blue indicates low expression. This plot is meant to provide a high level exploratory view of the data.   
                                      
                                       
                                  
                                
                              
                            
                            
                              
                                
                                  
                                       
                                      
                                           Principal Components of S45 mutants of beta-catenin aren.SQT.t phosphorylated Data   More Plot Information                      Principal Components of S45 mutants of beta-catenin aren.SQT.t phosphorylated Data  Principal component analysis maps high-dimensional datasets onto a smaller number of highly informative dimensions. Here, the first four principal components of the gene expression data are plotted against each other and colored by the values of the selected covariate. This plot may be used to identify clusters in the data and to identify variables associated with prominent signal in the data. Variables that are associated with these leading principal components should be considered in downstream analyses.   
                                      
                                       
                                  
                                
                                
                                  
                                       
                                      
                                           Principal Components of S45 mutants of beta-catenin aren.SQT.t phosphorylated Data   More Plot Information                      Principal Components of S45 mutants of beta-catenin aren.SQT.t phosphorylated Data  Principal component analysis maps high-dimensional datasets onto a smaller number of highly informative dimensions. Here, the first four principal components of the gene expression data are plotted against each other and colored by the values of the selected covariate. This plot may be used to identify clusters in the data and to identify variables associated with prominent signal in the data. Variables that are associated with these leading principal components should be considered in downstream analyses.   
                                      
                                       
                                  
                                
                              
                            
                           
                            
                          
                           
                      
                    
                    
                      
                           
                          
                               Heatmaps    PCA     
                              
                                  Heatmaps  
                              
                               
                              
                                  Line  
                                  Day  
                              
                              
                            
                              
                                
                                  
                                       
                                      
                                           Heatmap of Sema3A PAK dependent Axon repulsion Data   More Plot Information                      Heatmap of Sema3A PAK dependent Axon repulsion Data  Heatmap of the normalized data, scaled to give all genes equal variance, generated via unsupervised clustering. Orange indicates high expression; blue indicates low expression. This plot is meant to provide a high level exploratory view of the data.   
                                      
                                       
                                  
                                
                              
                            
                            
                              
                                
                                  
                                       
                                      
                                           Principal Components of Sema3A PAK dependent Axon repulsion Data   More Plot Information                      Principal Components of Sema3A PAK dependent Axon repulsion Data  Principal component analysis maps high-dimensional datasets onto a smaller number of highly informative dimensions. Here, the first four principal components of the gene expression data are plotted against each other and colored by the values of the selected covariate. This plot may be used to identify clusters in the data and to identify variables associated with prominent signal in the data. Variables that are associated with these leading principal components should be considered in downstream analyses.   
                                      
                                       
                                  
                                
                                
                                  
                                       
                                      
                                           Principal Components of Sema3A PAK dependent Axon repulsion Data   More Plot Information                      Principal Components of Sema3A PAK dependent Axon repulsion Data  Principal component analysis maps high-dimensional datasets onto a smaller number of highly informative dimensions. Here, the first four principal components of the gene expression data are plotted against each other and colored by the values of the selected covariate. This plot may be used to identify clusters in the data and to identify variables associated with prominent signal in the data. Variables that are associated with these leading principal components should be considered in downstream analyses.   
                                      
                                       
                                  
                                
                              
                            
                           
                            
                          
                           
                      
                    
                    
                      
                           
                          
                               Heatmaps    PCA     
                              
                                  Heatmaps  
                              
                               
                              
                                  Line  
                                  Day  
                              
                              
                            
                              
                                
                                  
                                       
                                      
                                           Heatmap of Sema4D induced cell migration and growth-cone collapse Data   More Plot Information                      Heatmap of Sema4D induced cell migration and growth-cone collapse Data  Heatmap of the normalized data, scaled to give all genes equal variance, generated via unsupervised clustering. Orange indicates high expression; blue indicates low expression. This plot is meant to provide a high level exploratory view of the data.   
                                      
                                       
                                  
                                
                              
                            
                            
                              
                                
                                  
                                       
                                      
                                           Principal Components of Sema4D induced cell migration and growth-cone collapse Data   More Plot Information                      Principal Components of Sema4D induced cell migration and growth-cone collapse Data  Principal component analysis maps high-dimensional datasets onto a smaller number of highly informative dimensions. Here, the first four principal components of the gene expression data are plotted against each other and colored by the values of the selected covariate. This plot may be used to identify clusters in the data and to identify variables associated with prominent signal in the data. Variables that are associated with these leading principal components should be considered in downstream analyses.   
                                      
                                       
                                  
                                
                                
                                  
                                       
                                      
                                           Principal Components of Sema4D induced cell migration and growth-cone collapse Data   More Plot Information                      Principal Components of Sema4D induced cell migration and growth-cone collapse Data  Principal component analysis maps high-dimensional datasets onto a smaller number of highly informative dimensions. Here, the first four principal components of the gene expression data are plotted against each other and colored by the values of the selected covariate. This plot may be used to identify clusters in the data and to identify variables associated with prominent signal in the data. Variables that are associated with these leading principal components should be considered in downstream analyses.   
                                      
                                       
                                  
                                
                              
                            
                           
                            
                          
                           
                      
                    
                    
                      
                           
                          
                               Heatmaps    PCA     
                              
                                  Heatmaps  
                              
                               
                              
                                  Line  
                                  Day  
                              
                              
                            
                              
                                
                                  
                                       
                                      
                                           Heatmap of SHC-mediated cascade.CL.FGFR1 Data   More Plot Information                      Heatmap of SHC-mediated cascade.CL.FGFR1 Data  Heatmap of the normalized data, scaled to give all genes equal variance, generated via unsupervised clustering. Orange indicates high expression; blue indicates low expression. This plot is meant to provide a high level exploratory view of the data.   
                                      
                                       
                                  
                                
                              
                            
                            
                              
                                
                                  
                                       
                                      
                                           Principal Components of SHC-mediated cascade.CL.FGFR1 Data   More Plot Information                      Principal Components of SHC-mediated cascade.CL.FGFR1 Data  Principal component analysis maps high-dimensional datasets onto a smaller number of highly informative dimensions. Here, the first four principal components of the gene expression data are plotted against each other and colored by the values of the selected covariate. This plot may be used to identify clusters in the data and to identify variables associated with prominent signal in the data. Variables that are associated with these leading principal components should be considered in downstream analyses.   
                                      
                                       
                                  
                                
                                
                                  
                                       
                                      
                                           Principal Components of SHC-mediated cascade.CL.FGFR1 Data   More Plot Information                      Principal Components of SHC-mediated cascade.CL.FGFR1 Data  Principal component analysis maps high-dimensional datasets onto a smaller number of highly informative dimensions. Here, the first four principal components of the gene expression data are plotted against each other and colored by the values of the selected covariate. This plot may be used to identify clusters in the data and to identify variables associated with prominent signal in the data. Variables that are associated with these leading principal components should be considered in downstream analyses.   
                                      
                                       
                                  
                                
                              
                            
                           
                            
                          
                           
                      
                    
                    
                      
                           
                          
                               Heatmaps    PCA     
                              
                                  Heatmaps  
                              
                               
                              
                                  Line  
                                  Day  
                              
                              
                            
                              
                                
                                  
                                       
                                      
                                           Heatmap of SHC-related events triggered by IGF1R Data   More Plot Information                      Heatmap of SHC-related events triggered by IGF1R Data  Heatmap of the normalized data, scaled to give all genes equal variance, generated via unsupervised clustering. Orange indicates high expression; blue indicates low expression. This plot is meant to provide a high level exploratory view of the data.   
                                      
                                       
                                  
                                
                              
                            
                            
                              
                                
                                  
                                       
                                      
                                           Principal Components of SHC-related events triggered by IGF1R Data   More Plot Information                      Principal Components of SHC-related events triggered by IGF1R Data  Principal component analysis maps high-dimensional datasets onto a smaller number of highly informative dimensions. Here, the first four principal components of the gene expression data are plotted against each other and colored by the values of the selected covariate. This plot may be used to identify clusters in the data and to identify variables associated with prominent signal in the data. Variables that are associated with these leading principal components should be considered in downstream analyses.   
                                      
                                       
                                  
                                
                                
                                  
                                       
                                      
                                           Principal Components of SHC-related events triggered by IGF1R Data   More Plot Information                      Principal Components of SHC-related events triggered by IGF1R Data  Principal component analysis maps high-dimensional datasets onto a smaller number of highly informative dimensions. Here, the first four principal components of the gene expression data are plotted against each other and colored by the values of the selected covariate. This plot may be used to identify clusters in the data and to identify variables associated with prominent signal in the data. Variables that are associated with these leading principal components should be considered in downstream analyses.   
                                      
                                       
                                  
                                
                              
                            
                           
                            
                          
                           
                      
                    
                    
                      
                           
                          
                               Heatmaps    PCA     
                              
                                  Heatmaps  
                              
                               
                              
                                  Line  
                                  Day  
                              
                              
                            
                              
                                
                                  
                                       
                                      
                                           Heatmap of Signal transduction by L1 Data   More Plot Information                      Heatmap of Signal transduction by L1 Data  Heatmap of the normalized data, scaled to give all genes equal variance, generated via unsupervised clustering. Orange indicates high expression; blue indicates low expression. This plot is meant to provide a high level exploratory view of the data.   
                                      
                                       
                                  
                                
                              
                            
                            
                              
                                
                                  
                                       
                                      
                                           Principal Components of Signal transduction by L1 Data   More Plot Information                      Principal Components of Signal transduction by L1 Data  Principal component analysis maps high-dimensional datasets onto a smaller number of highly informative dimensions. Here, the first four principal components of the gene expression data are plotted against each other and colored by the values of the selected covariate. This plot may be used to identify clusters in the data and to identify variables associated with prominent signal in the data. Variables that are associated with these leading principal components should be considered in downstream analyses.   
                                      
                                       
                                  
                                
                                
                                  
                                       
                                      
                                           Principal Components of Signal transduction by L1 Data   More Plot Information                      Principal Components of Signal transduction by L1 Data  Principal component analysis maps high-dimensional datasets onto a smaller number of highly informative dimensions. Here, the first four principal components of the gene expression data are plotted against each other and colored by the values of the selected covariate. This plot may be used to identify clusters in the data and to identify variables associated with prominent signal in the data. Variables that are associated with these leading principal components should be considered in downstream analyses.   
                                      
                                       
                                  
                                
                              
                            
                           
                            
                          
                           
                      
                    
                    
                      
                           
                          
                               Heatmaps    PCA     
                              
                                  Heatmaps  
                              
                               
                              
                                  Line  
                                  Day  
                              
                              
                            
                              
                                
                                  
                                       
                                      
                                           Heatmap of Signaling by ERBB2 Data   More Plot Information                      Heatmap of Signaling by ERBB2 Data  Heatmap of the normalized data, scaled to give all genes equal variance, generated via unsupervised clustering. Orange indicates high expression; blue indicates low expression. This plot is meant to provide a high level exploratory view of the data.   
                                      
                                       
                                  
                                
                              
                            
                            
                              
                                
                                  
                                       
                                      
                                           Principal Components of Signaling by ERBB2 Data   More Plot Information                      Principal Components of Signaling by ERBB2 Data  Principal component analysis maps high-dimensional datasets onto a smaller number of highly informative dimensions. Here, the first four principal components of the gene expression data are plotted against each other and colored by the values of the selected covariate. This plot may be used to identify clusters in the data and to identify variables associated with prominent signal in the data. Variables that are associated with these leading principal components should be considered in downstream analyses.   
                                      
                                       
                                  
                                
                                
                                  
                                       
                                      
                                           Principal Components of Signaling by ERBB2 Data   More Plot Information                      Principal Components of Signaling by ERBB2 Data  Principal component analysis maps high-dimensional datasets onto a smaller number of highly informative dimensions. Here, the first four principal components of the gene expression data are plotted against each other and colored by the values of the selected covariate. This plot may be used to identify clusters in the data and to identify variables associated with prominent signal in the data. Variables that are associated with these leading principal components should be considered in downstream analyses.   
                                      
                                       
                                  
                                
                              
                            
                           
                            
                          
                           
                      
                    
                    
                      
                           
                          
                               Heatmaps    PCA     
                              
                                  Heatmaps  
                              
                               
                              
                                  Line  
                                  Day  
                              
                              
                            
                              
                                
                                  
                                       
                                      
                                           Heatmap of Signaling by FGFR1 in disease Data   More Plot Information                      Heatmap of Signaling by FGFR1 in disease Data  Heatmap of the normalized data, scaled to give all genes equal variance, generated via unsupervised clustering. Orange indicates high expression; blue indicates low expression. This plot is meant to provide a high level exploratory view of the data.   
                                      
                                       
                                  
                                
                              
                            
                            
                              
                                
                                  
                                       
                                      
                                           Principal Components of Signaling by FGFR1 in disease Data   More Plot Information                      Principal Components of Signaling by FGFR1 in disease Data  Principal component analysis maps high-dimensional datasets onto a smaller number of highly informative dimensions. Here, the first four principal components of the gene expression data are plotted against each other and colored by the values of the selected covariate. This plot may be used to identify clusters in the data and to identify variables associated with prominent signal in the data. Variables that are associated with these leading principal components should be considered in downstream analyses.   
                                      
                                       
                                  
                                
                                
                                  
                                       
                                      
                                           Principal Components of Signaling by FGFR1 in disease Data   More Plot Information                      Principal Components of Signaling by FGFR1 in disease Data  Principal component analysis maps high-dimensional datasets onto a smaller number of highly informative dimensions. Here, the first four principal components of the gene expression data are plotted against each other and colored by the values of the selected covariate. This plot may be used to identify clusters in the data and to identify variables associated with prominent signal in the data. Variables that are associated with these leading principal components should be considered in downstream analyses.   
                                      
                                       
                                  
                                
                              
                            
                           
                            
                          
                           
                      
                    
                    
                      
                           
                          
                               Heatmaps    PCA     
                              
                                  Heatmaps  
                              
                               
                              
                                  Line  
                                  Day  
                              
                              
                            
                              
                                
                                  
                                       
                                      
                                           Heatmap of Signaling by FGFR2 in disease Data   More Plot Information                      Heatmap of Signaling by FGFR2 in disease Data  Heatmap of the normalized data, scaled to give all genes equal variance, generated via unsupervised clustering. Orange indicates high expression; blue indicates low expression. This plot is meant to provide a high level exploratory view of the data.   
                                      
                                       
                                  
                                
                              
                            
                            
                              
                                
                                  
                                       
                                      
                                           Principal Components of Signaling by FGFR2 in disease Data   More Plot Information                      Principal Components of Signaling by FGFR2 in disease Data  Principal component analysis maps high-dimensional datasets onto a smaller number of highly informative dimensions. Here, the first four principal components of the gene expression data are plotted against each other and colored by the values of the selected covariate. This plot may be used to identify clusters in the data and to identify variables associated with prominent signal in the data. Variables that are associated with these leading principal components should be considered in downstream analyses.   
                                      
                                       
                                  
                                
                                
                                  
                                       
                                      
                                           Principal Components of Signaling by FGFR2 in disease Data   More Plot Information                      Principal Components of Signaling by FGFR2 in disease Data  Principal component analysis maps high-dimensional datasets onto a smaller number of highly informative dimensions. Here, the first four principal components of the gene expression data are plotted against each other and colored by the values of the selected covariate. This plot may be used to identify clusters in the data and to identify variables associated with prominent signal in the data. Variables that are associated with these leading principal components should be considered in downstream analyses.   
                                      
                                       
                                  
                                
                              
                            
                           
                            
                          
                           
                      
                    
                    
                      
                           
                          
                               Heatmaps    PCA     
                              
                                  Heatmaps  
                              
                               
                              
                                  Line  
                                  Day  
                              
                              
                            
                              
                                
                                  
                                       
                                      
                                           Heatmap of Signaling by FGFR3 in disease Data   More Plot Information                      Heatmap of Signaling by FGFR3 in disease Data  Heatmap of the normalized data, scaled to give all genes equal variance, generated via unsupervised clustering. Orange indicates high expression; blue indicates low expression. This plot is meant to provide a high level exploratory view of the data.   
                                      
                                       
                                  
                                
                              
                            
                            
                              
                                
                                  
                                       
                                      
                                           Principal Components of Signaling by FGFR3 in disease Data   More Plot Information                      Principal Components of Signaling by FGFR3 in disease Data  Principal component analysis maps high-dimensional datasets onto a smaller number of highly informative dimensions. Here, the first four principal components of the gene expression data are plotted against each other and colored by the values of the selected covariate. This plot may be used to identify clusters in the data and to identify variables associated with prominent signal in the data. Variables that are associated with these leading principal components should be considered in downstream analyses.   
                                      
                                       
                                  
                                
                                
                                  
                                       
                                      
                                           Principal Components of Signaling by FGFR3 in disease Data   More Plot Information                      Principal Components of Signaling by FGFR3 in disease Data  Principal component analysis maps high-dimensional datasets onto a smaller number of highly informative dimensions. Here, the first four principal components of the gene expression data are plotted against each other and colored by the values of the selected covariate. This plot may be used to identify clusters in the data and to identify variables associated with prominent signal in the data. Variables that are associated with these leading principal components should be considered in downstream analyses.   
                                      
                                       
                                  
                                
                              
                            
                           
                            
                          
                           
                      
                    
                    
                      
                           
                          
                               Heatmaps    PCA     
                              
                                  Heatmaps  
                              
                               
                              
                                  Line  
                                  Day  
                              
                              
                            
                              
                                
                                  
                                       
                                      
                                           Heatmap of Signaling by FGFR4 in disease Data   More Plot Information                      Heatmap of Signaling by FGFR4 in disease Data  Heatmap of the normalized data, scaled to give all genes equal variance, generated via unsupervised clustering. Orange indicates high expression; blue indicates low expression. This plot is meant to provide a high level exploratory view of the data.   
                                      
                                       
                                  
                                
                              
                            
                            
                              
                                
                                  
                                       
                                      
                                           Principal Components of Signaling by FGFR4 in disease Data   More Plot Information                      Principal Components of Signaling by FGFR4 in disease Data  Principal component analysis maps high-dimensional datasets onto a smaller number of highly informative dimensions. Here, the first four principal components of the gene expression data are plotted against each other and colored by the values of the selected covariate. This plot may be used to identify clusters in the data and to identify variables associated with prominent signal in the data. Variables that are associated with these leading principal components should be considered in downstream analyses.   
                                      
                                       
                                  
                                
                                
                                  
                                       
                                      
                                           Principal Components of Signaling by FGFR4 in disease Data   More Plot Information                      Principal Components of Signaling by FGFR4 in disease Data  Principal component analysis maps high-dimensional datasets onto a smaller number of highly informative dimensions. Here, the first four principal components of the gene expression data are plotted against each other and colored by the values of the selected covariate. This plot may be used to identify clusters in the data and to identify variables associated with prominent signal in the data. Variables that are associated with these leading principal components should be considered in downstream analyses.   
                                      
                                       
                                  
                                
                              
                            
                           
                            
                          
                           
                      
                    
                    
                      
                           
                          
                               Heatmaps    PCA     
                              
                                  Heatmaps  
                              
                               
                              
                                  Line  
                                  Day  
                              
                              
                            
                              
                                
                                  
                                       
                                      
                                           Heatmap of Signaling by PDGF Data   More Plot Information                      Heatmap of Signaling by PDGF Data  Heatmap of the normalized data, scaled to give all genes equal variance, generated via unsupervised clustering. Orange indicates high expression; blue indicates low expression. This plot is meant to provide a high level exploratory view of the data.   
                                      
                                       
                                  
                                
                              
                            
                            
                              
                                
                                  
                                       
                                      
                                           Principal Components of Signaling by PDGF Data   More Plot Information                      Principal Components of Signaling by PDGF Data  Principal component analysis maps high-dimensional datasets onto a smaller number of highly informative dimensions. Here, the first four principal components of the gene expression data are plotted against each other and colored by the values of the selected covariate. This plot may be used to identify clusters in the data and to identify variables associated with prominent signal in the data. Variables that are associated with these leading principal components should be considered in downstream analyses.   
                                      
                                       
                                  
                                
                                
                                  
                                       
                                      
                                           Principal Components of Signaling by PDGF Data   More Plot Information                      Principal Components of Signaling by PDGF Data  Principal component analysis maps high-dimensional datasets onto a smaller number of highly informative dimensions. Here, the first four principal components of the gene expression data are plotted against each other and colored by the values of the selected covariate. This plot may be used to identify clusters in the data and to identify variables associated with prominent signal in the data. Variables that are associated with these leading principal components should be considered in downstream analyses.   
                                      
                                       
                                  
                                
                              
                            
                           
                            
                          
                           
                      
                    
                    
                      
                           
                          
                               Heatmaps    PCA     
                              
                                  Heatmaps  
                              
                               
                              
                                  Line  
                                  Day  
                              
                              
                            
                              
                                
                                  
                                       
                                      
                                           Heatmap of Signaling by SCF-KIT Data   More Plot Information                      Heatmap of Signaling by SCF-KIT Data  Heatmap of the normalized data, scaled to give all genes equal variance, generated via unsupervised clustering. Orange indicates high expression; blue indicates low expression. This plot is meant to provide a high level exploratory view of the data.   
                                      
                                       
                                  
                                
                              
                            
                            
                              
                                
                                  
                                       
                                      
                                           Principal Components of Signaling by SCF-KIT Data   More Plot Information                      Principal Components of Signaling by SCF-KIT Data  Principal component analysis maps high-dimensional datasets onto a smaller number of highly informative dimensions. Here, the first four principal components of the gene expression data are plotted against each other and colored by the values of the selected covariate. This plot may be used to identify clusters in the data and to identify variables associated with prominent signal in the data. Variables that are associated with these leading principal components should be considered in downstream analyses.   
                                      
                                       
                                  
                                
                                
                                  
                                       
                                      
                                           Principal Components of Signaling by SCF-KIT Data   More Plot Information                      Principal Components of Signaling by SCF-KIT Data  Principal component analysis maps high-dimensional datasets onto a smaller number of highly informative dimensions. Here, the first four principal components of the gene expression data are plotted against each other and colored by the values of the selected covariate. This plot may be used to identify clusters in the data and to identify variables associated with prominent signal in the data. Variables that are associated with these leading principal components should be considered in downstream analyses.   
                                      
                                       
                                  
                                
                              
                            
                           
                            
                          
                           
                      
                    
                    
                      
                           
                          
                               Heatmaps    PCA     
                              
                                  Heatmaps  
                              
                               
                              
                                  Line  
                                  Day  
                              
                              
                            
                              
                                
                                  
                                       
                                      
                                           Heatmap of Smooth Muscle Contraction Data   More Plot Information                      Heatmap of Smooth Muscle Contraction Data  Heatmap of the normalized data, scaled to give all genes equal variance, generated via unsupervised clustering. Orange indicates high expression; blue indicates low expression. This plot is meant to provide a high level exploratory view of the data.   
                                      
                                       
                                  
                                
                              
                            
                            
                              
                                
                                  
                                       
                                      
                                           Principal Components of Smooth Muscle Contraction Data   More Plot Information                      Principal Components of Smooth Muscle Contraction Data  Principal component analysis maps high-dimensional datasets onto a smaller number of highly informative dimensions. Here, the first four principal components of the gene expression data are plotted against each other and colored by the values of the selected covariate. This plot may be used to identify clusters in the data and to identify variables associated with prominent signal in the data. Variables that are associated with these leading principal components should be considered in downstream analyses.   
                                      
                                       
                                  
                                
                                
                                  
                                       
                                      
                                           Principal Components of Smooth Muscle Contraction Data   More Plot Information                      Principal Components of Smooth Muscle Contraction Data  Principal component analysis maps high-dimensional datasets onto a smaller number of highly informative dimensions. Here, the first four principal components of the gene expression data are plotted against each other and colored by the values of the selected covariate. This plot may be used to identify clusters in the data and to identify variables associated with prominent signal in the data. Variables that are associated with these leading principal components should be considered in downstream analyses.   
                                      
                                       
                                  
                                
                              
                            
                           
                            
                          
                           
                      
                    
                    
                      
                           
                          
                               Heatmaps    PCA     
                              
                                  Heatmaps  
                              
                               
                              
                                  Line  
                                  Day  
                              
                              
                            
                              
                                
                                  
                                       
                                      
                                           Heatmap of Spry regulation of FGF signaling Data   More Plot Information                      Heatmap of Spry regulation of FGF signaling Data  Heatmap of the normalized data, scaled to give all genes equal variance, generated via unsupervised clustering. Orange indicates high expression; blue indicates low expression. This plot is meant to provide a high level exploratory view of the data.   
                                      
                                       
                                  
                                
                              
                            
                            
                              
                                
                                  
                                       
                                      
                                           Principal Components of Spry regulation of FGF signaling Data   More Plot Information                      Principal Components of Spry regulation of FGF signaling Data  Principal component analysis maps high-dimensional datasets onto a smaller number of highly informative dimensions. Here, the first four principal components of the gene expression data are plotted against each other and colored by the values of the selected covariate. This plot may be used to identify clusters in the data and to identify variables associated with prominent signal in the data. Variables that are associated with these leading principal components should be considered in downstream analyses.   
                                      
                                       
                                  
                                
                                
                                  
                                       
                                      
                                           Principal Components of Spry regulation of FGF signaling Data   More Plot Information                      Principal Components of Spry regulation of FGF signaling Data  Principal component analysis maps high-dimensional datasets onto a smaller number of highly informative dimensions. Here, the first four principal components of the gene expression data are plotted against each other and colored by the values of the selected covariate. This plot may be used to identify clusters in the data and to identify variables associated with prominent signal in the data. Variables that are associated with these leading principal components should be considered in downstream analyses.   
                                      
                                       
                                  
                                
                              
                            
                           
                            
                          
                           
                      
                    
                    
                      
                           
                          
                               Heatmaps    PCA     
                              
                                  Heatmaps  
                              
                               
                              
                                  Line  
                                  Day  
                              
                              
                            
                              
                                
                                  
                                       
                                      
                                           Heatmap of Syndecan interactions Data   More Plot Information                      Heatmap of Syndecan interactions Data  Heatmap of the normalized data, scaled to give all genes equal variance, generated via unsupervised clustering. Orange indicates high expression; blue indicates low expression. This plot is meant to provide a high level exploratory view of the data.   
                                      
                                       
                                  
                                
                              
                            
                            
                              
                                
                                  
                                       
                                      
                                           Principal Components of Syndecan interactions Data   More Plot Information                      Principal Components of Syndecan interactions Data  Principal component analysis maps high-dimensional datasets onto a smaller number of highly informative dimensions. Here, the first four principal components of the gene expression data are plotted against each other and colored by the values of the selected covariate. This plot may be used to identify clusters in the data and to identify variables associated with prominent signal in the data. Variables that are associated with these leading principal components should be considered in downstream analyses.   
                                      
                                       
                                  
                                
                                
                                  
                                       
                                      
                                           Principal Components of Syndecan interactions Data   More Plot Information                      Principal Components of Syndecan interactions Data  Principal component analysis maps high-dimensional datasets onto a smaller number of highly informative dimensions. Here, the first four principal components of the gene expression data are plotted against each other and colored by the values of the selected covariate. This plot may be used to identify clusters in the data and to identify variables associated with prominent signal in the data. Variables that are associated with these leading principal components should be considered in downstream analyses.   
                                      
                                       
                                  
                                
                              
                            
                           
                            
                          
                           
                      
                    
                    
                      
                           
                          
                               Heatmaps    PCA     
                              
                                  Heatmaps  
                              
                               
                              
                                  Line  
                                  Day  
                              
                              
                            
                              
                                
                                  
                                       
                                      
                                           Heatmap of Synthesis of PIPs at the plasma membrane Data   More Plot Information                      Heatmap of Synthesis of PIPs at the plasma membrane Data  Heatmap of the normalized data, scaled to give all genes equal variance, generated via unsupervised clustering. Orange indicates high expression; blue indicates low expression. This plot is meant to provide a high level exploratory view of the data.   
                                      
                                       
                                  
                                
                              
                            
                            
                              
                                
                                  
                                       
                                      
                                           Principal Components of Synthesis of PIPs at the plasma membrane Data   More Plot Information                      Principal Components of Synthesis of PIPs at the plasma membrane Data  Principal component analysis maps high-dimensional datasets onto a smaller number of highly informative dimensions. Here, the first four principal components of the gene expression data are plotted against each other and colored by the values of the selected covariate. This plot may be used to identify clusters in the data and to identify variables associated with prominent signal in the data. Variables that are associated with these leading principal components should be considered in downstream analyses.   
                                      
                                       
                                  
                                
                                
                                  
                                       
                                      
                                           Principal Components of Synthesis of PIPs at the plasma membrane Data   More Plot Information                      Principal Components of Synthesis of PIPs at the plasma membrane Data  Principal component analysis maps high-dimensional datasets onto a smaller number of highly informative dimensions. Here, the first four principal components of the gene expression data are plotted against each other and colored by the values of the selected covariate. This plot may be used to identify clusters in the data and to identify variables associated with prominent signal in the data. Variables that are associated with these leading principal components should be considered in downstream analyses.   
                                      
                                       
                                  
                                
                              
                            
                           
                            
                          
                           
                      
                    
                    
                      
                           
                          
                               Heatmaps    PCA     
                              
                                  Heatmaps  
                              
                               
                              
                                  Line  
                                  Day  
                              
                              
                            
                              
                                
                                  
                                       
                                      
                                           Heatmap of T41 mutants of beta-catenin aren.SQT.t phosphorylated Data   More Plot Information                      Heatmap of T41 mutants of beta-catenin aren.SQT.t phosphorylated Data  Heatmap of the normalized data, scaled to give all genes equal variance, generated via unsupervised clustering. Orange indicates high expression; blue indicates low expression. This plot is meant to provide a high level exploratory view of the data.   
                                      
                                       
                                  
                                
                              
                            
                            
                              
                                
                                  
                                       
                                      
                                           Principal Components of T41 mutants of beta-catenin aren.SQT.t phosphorylated Data   More Plot Information                      Principal Components of T41 mutants of beta-catenin aren.SQT.t phosphorylated Data  Principal component analysis maps high-dimensional datasets onto a smaller number of highly informative dimensions. Here, the first four principal components of the gene expression data are plotted against each other and colored by the values of the selected covariate. This plot may be used to identify clusters in the data and to identify variables associated with prominent signal in the data. Variables that are associated with these leading principal components should be considered in downstream analyses.   
                                      
                                       
                                  
                                
                                
                                  
                                       
                                      
                                           Principal Components of T41 mutants of beta-catenin aren.SQT.t phosphorylated Data   More Plot Information                      Principal Components of T41 mutants of beta-catenin aren.SQT.t phosphorylated Data  Principal component analysis maps high-dimensional datasets onto a smaller number of highly informative dimensions. Here, the first four principal components of the gene expression data are plotted against each other and colored by the values of the selected covariate. This plot may be used to identify clusters in the data and to identify variables associated with prominent signal in the data. Variables that are associated with these leading principal components should be considered in downstream analyses.   
                                      
                                       
                                  
                                
                              
                            
                           
                            
                          
                           
                      
                    
                    
                      
                           
                          
                               Heatmaps    PCA     
                              
                                  Heatmaps  
                              
                               
                              
                                  Line  
                                  Day  
                              
                              
                            
                              
                                
                                  
                                       
                                      
                                           Heatmap of TGF-beta receptor signaling activates SMADs Data   More Plot Information                      Heatmap of TGF-beta receptor signaling activates SMADs Data  Heatmap of the normalized data, scaled to give all genes equal variance, generated via unsupervised clustering. Orange indicates high expression; blue indicates low expression. This plot is meant to provide a high level exploratory view of the data.   
                                      
                                       
                                  
                                
                              
                            
                            
                              
                                
                                  
                                       
                                      
                                           Principal Components of TGF-beta receptor signaling activates SMADs Data   More Plot Information                      Principal Components of TGF-beta receptor signaling activates SMADs Data  Principal component analysis maps high-dimensional datasets onto a smaller number of highly informative dimensions. Here, the first four principal components of the gene expression data are plotted against each other and colored by the values of the selected covariate. This plot may be used to identify clusters in the data and to identify variables associated with prominent signal in the data. Variables that are associated with these leading principal components should be considered in downstream analyses.   
                                      
                                       
                                  
                                
                                
                                  
                                       
                                      
                                           Principal Components of TGF-beta receptor signaling activates SMADs Data   More Plot Information                      Principal Components of TGF-beta receptor signaling activates SMADs Data  Principal component analysis maps high-dimensional datasets onto a smaller number of highly informative dimensions. Here, the first four principal components of the gene expression data are plotted against each other and colored by the values of the selected covariate. This plot may be used to identify clusters in the data and to identify variables associated with prominent signal in the data. Variables that are associated with these leading principal components should be considered in downstream analyses.   
                                      
                                       
                                  
                                
                              
                            
                           
                            
                          
                           
                      
                    
                    
                      
                           
                          
                               Heatmaps    PCA     
                              
                                  Heatmaps  
                              
                               
                              
                                  Line  
                                  Day  
                              
                              
                            
                              
                                
                                  
                                       
                                      
                                           Heatmap of TGF-beta receptor signaling in EMT (epithelial to mesenchymal transition) Data   More Plot Information                      Heatmap of TGF-beta receptor signaling in EMT (epithelial to mesenchymal transition) Data  Heatmap of the normalized data, scaled to give all genes equal variance, generated via unsupervised clustering. Orange indicates high expression; blue indicates low expression. This plot is meant to provide a high level exploratory view of the data.   
                                      
                                       
                                  
                                
                              
                            
                            
                              
                                
                                  
                                       
                                      
                                           Principal Components of TGF-beta receptor signaling in EMT (epithelial to mesenchymal transition) Data   More Plot Information                      Principal Components of TGF-beta receptor signaling in EMT (epithelial to mesenchymal transition) Data  Principal component analysis maps high-dimensional datasets onto a smaller number of highly informative dimensions. Here, the first four principal components of the gene expression data are plotted against each other and colored by the values of the selected covariate. This plot may be used to identify clusters in the data and to identify variables associated with prominent signal in the data. Variables that are associated with these leading principal components should be considered in downstream analyses.   
                                      
                                       
                                  
                                
                                
                                  
                                       
                                      
                                           Principal Components of TGF-beta receptor signaling in EMT (epithelial to mesenchymal transition) Data   More Plot Information                      Principal Components of TGF-beta receptor signaling in EMT (epithelial to mesenchymal transition) Data  Principal component analysis maps high-dimensional datasets onto a smaller number of highly informative dimensions. Here, the first four principal components of the gene expression data are plotted against each other and colored by the values of the selected covariate. This plot may be used to identify clusters in the data and to identify variables associated with prominent signal in the data. Variables that are associated with these leading principal components should be considered in downstream analyses.   
                                      
                                       
                                  
                                
                              
                            
                           
                            
                          
                           
                      
                    
                    
                      
                           
                          
                               Heatmaps    PCA     
                              
                                  Heatmaps  
                              
                               
                              
                                  Line  
                                  Day  
                              
                              
                            
                              
                                
                                  
                                       
                                      
                                           Heatmap of Tie2 Signaling Data   More Plot Information                      Heatmap of Tie2 Signaling Data  Heatmap of the normalized data, scaled to give all genes equal variance, generated via unsupervised clustering. Orange indicates high expression; blue indicates low expression. This plot is meant to provide a high level exploratory view of the data.   
                                      
                                       
                                  
                                
                              
                            
                            
                              
                                
                                  
                                       
                                      
                                           Principal Components of Tie2 Signaling Data   More Plot Information                      Principal Components of Tie2 Signaling Data  Principal component analysis maps high-dimensional datasets onto a smaller number of highly informative dimensions. Here, the first four principal components of the gene expression data are plotted against each other and colored by the values of the selected covariate. This plot may be used to identify clusters in the data and to identify variables associated with prominent signal in the data. Variables that are associated with these leading principal components should be considered in downstream analyses.   
                                      
                                       
                                  
                                
                                
                                  
                                       
                                      
                                           Principal Components of Tie2 Signaling Data   More Plot Information                      Principal Components of Tie2 Signaling Data  Principal component analysis maps high-dimensional datasets onto a smaller number of highly informative dimensions. Here, the first four principal components of the gene expression data are plotted against each other and colored by the values of the selected covariate. This plot may be used to identify clusters in the data and to identify variables associated with prominent signal in the data. Variables that are associated with these leading principal components should be considered in downstream analyses.   
                                      
                                       
                                  
                                
                              
                            
                           
                            
                          
                           
                      
                    
                    
                      
                           
                          
                               Heatmaps    PCA     
                              
                                  Heatmaps  
                              
                               
                              
                                  Line  
                                  Day  
                              
                              
                            
                              
                                
                                  
                                       
                                      
                                           Heatmap of Tight junction interactions Data   More Plot Information                      Heatmap of Tight junction interactions Data  Heatmap of the normalized data, scaled to give all genes equal variance, generated via unsupervised clustering. Orange indicates high expression; blue indicates low expression. This plot is meant to provide a high level exploratory view of the data.   
                                      
                                       
                                  
                                
                              
                            
                            
                              
                                
                                  
                                       
                                      
                                           Principal Components of Tight junction interactions Data   More Plot Information                      Principal Components of Tight junction interactions Data  Principal component analysis maps high-dimensional datasets onto a smaller number of highly informative dimensions. Here, the first four principal components of the gene expression data are plotted against each other and colored by the values of the selected covariate. This plot may be used to identify clusters in the data and to identify variables associated with prominent signal in the data. Variables that are associated with these leading principal components should be considered in downstream analyses.   
                                      
                                       
                                  
                                
                                
                                  
                                       
                                      
                                           Principal Components of Tight junction interactions Data   More Plot Information                      Principal Components of Tight junction interactions Data  Principal component analysis maps high-dimensional datasets onto a smaller number of highly informative dimensions. Here, the first four principal components of the gene expression data are plotted against each other and colored by the values of the selected covariate. This plot may be used to identify clusters in the data and to identify variables associated with prominent signal in the data. Variables that are associated with these leading principal components should be considered in downstream analyses.   
                                      
                                       
                                  
                                
                              
                            
                           
                            
                          
                           
                      
                    
                    
                      
                           
                          
                               Heatmaps    PCA     
                              
                                  Heatmaps  
                              
                               
                              
                                  Line  
                                  Day  
                              
                              
                            
                              
                                
                                  
                                       
                                      
                                           Heatmap of TP53 Regulates Metabolic Genes Data   More Plot Information                      Heatmap of TP53 Regulates Metabolic Genes Data  Heatmap of the normalized data, scaled to give all genes equal variance, generated via unsupervised clustering. Orange indicates high expression; blue indicates low expression. This plot is meant to provide a high level exploratory view of the data.   
                                      
                                       
                                  
                                
                              
                            
                            
                              
                                
                                  
                                       
                                      
                                           Principal Components of TP53 Regulates Metabolic Genes Data   More Plot Information                      Principal Components of TP53 Regulates Metabolic Genes Data  Principal component analysis maps high-dimensional datasets onto a smaller number of highly informative dimensions. Here, the first four principal components of the gene expression data are plotted against each other and colored by the values of the selected covariate. This plot may be used to identify clusters in the data and to identify variables associated with prominent signal in the data. Variables that are associated with these leading principal components should be considered in downstream analyses.   
                                      
                                       
                                  
                                
                                
                                  
                                       
                                      
                                           Principal Components of TP53 Regulates Metabolic Genes Data   More Plot Information                      Principal Components of TP53 Regulates Metabolic Genes Data  Principal component analysis maps high-dimensional datasets onto a smaller number of highly informative dimensions. Here, the first four principal components of the gene expression data are plotted against each other and colored by the values of the selected covariate. This plot may be used to identify clusters in the data and to identify variables associated with prominent signal in the data. Variables that are associated with these leading principal components should be considered in downstream analyses.   
                                      
                                       
                                  
                                
                              
                            
                           
                            
                          
                           
                      
                    
                    
                      
                           
                          
                               Heatmaps    PCA     
                              
                                  Heatmaps  
                              
                               
                              
                                  Line  
                                  Day  
                              
                              
                            
                              
                                
                                  
                                       
                                      
                                           Heatmap of Truncations of AMER1 destabilize the destruction complex Data   More Plot Information                      Heatmap of Truncations of AMER1 destabilize the destruction complex Data  Heatmap of the normalized data, scaled to give all genes equal variance, generated via unsupervised clustering. Orange indicates high expression; blue indicates low expression. This plot is meant to provide a high level exploratory view of the data.   
                                      
                                       
                                  
                                
                              
                            
                            
                              
                                
                                  
                                       
                                      
                                           Principal Components of Truncations of AMER1 destabilize the destruction complex Data   More Plot Information                      Principal Components of Truncations of AMER1 destabilize the destruction complex Data  Principal component analysis maps high-dimensional datasets onto a smaller number of highly informative dimensions. Here, the first four principal components of the gene expression data are plotted against each other and colored by the values of the selected covariate. This plot may be used to identify clusters in the data and to identify variables associated with prominent signal in the data. Variables that are associated with these leading principal components should be considered in downstream analyses.   
                                      
                                       
                                  
                                
                                
                                  
                                       
                                      
                                           Principal Components of Truncations of AMER1 destabilize the destruction complex Data   More Plot Information                      Principal Components of Truncations of AMER1 destabilize the destruction complex Data  Principal component analysis maps high-dimensional datasets onto a smaller number of highly informative dimensions. Here, the first four principal components of the gene expression data are plotted against each other and colored by the values of the selected covariate. This plot may be used to identify clusters in the data and to identify variables associated with prominent signal in the data. Variables that are associated with these leading principal components should be considered in downstream analyses.   
                                      
                                       
                                  
                                
                              
                            
                           
                            
                          
                           
                      
                    
                    
                      
                           
                          
                               Heatmaps    PCA     
                              
                                  Heatmaps  
                              
                               
                              
                                  Line  
                                  Day  
                              
                              
                            
                              
                                
                                  
                                       
                                      
                                           Heatmap of VEGFA-VEGFR2 Pathway Data   More Plot Information                      Heatmap of VEGFA-VEGFR2 Pathway Data  Heatmap of the normalized data, scaled to give all genes equal variance, generated via unsupervised clustering. Orange indicates high expression; blue indicates low expression. This plot is meant to provide a high level exploratory view of the data.   
                                      
                                       
                                  
                                
                              
                            
                            
                              
                                
                                  
                                       
                                      
                                           Principal Components of VEGFA-VEGFR2 Pathway Data   More Plot Information                      Principal Components of VEGFA-VEGFR2 Pathway Data  Principal component analysis maps high-dimensional datasets onto a smaller number of highly informative dimensions. Here, the first four principal components of the gene expression data are plotted against each other and colored by the values of the selected covariate. This plot may be used to identify clusters in the data and to identify variables associated with prominent signal in the data. Variables that are associated with these leading principal components should be considered in downstream analyses.   
                                      
                                       
                                  
                                
                                
                                  
                                       
                                      
                                           Principal Components of VEGFA-VEGFR2 Pathway Data   More Plot Information                      Principal Components of VEGFA-VEGFR2 Pathway Data  Principal component analysis maps high-dimensional datasets onto a smaller number of highly informative dimensions. Here, the first four principal components of the gene expression data are plotted against each other and colored by the values of the selected covariate. This plot may be used to identify clusters in the data and to identify variables associated with prominent signal in the data. Variables that are associated with these leading principal components should be considered in downstream analyses.   
                                      
                                       
                                  
                                
                              
                            
                           
                            
                          
                           
                      
                    
                    
                      
                           
                          
                               Heatmaps    PCA     
                              
                                  Heatmaps  
                              
                               
                              
                                  Line  
                                  Day  
                              
                              
                            
                              
                                
                                  
                                       
                                      
                                           Heatmap of VEGFR2 mediated cell proliferation Data   More Plot Information                      Heatmap of VEGFR2 mediated cell proliferation Data  Heatmap of the normalized data, scaled to give all genes equal variance, generated via unsupervised clustering. Orange indicates high expression; blue indicates low expression. This plot is meant to provide a high level exploratory view of the data.   
                                      
                                       
                                  
                                
                              
                            
                            
                              
                                
                                  
                                       
                                      
                                           Principal Components of VEGFR2 mediated cell proliferation Data   More Plot Information                      Principal Components of VEGFR2 mediated cell proliferation Data  Principal component analysis maps high-dimensional datasets onto a smaller number of highly informative dimensions. Here, the first four principal components of the gene expression data are plotted against each other and colored by the values of the selected covariate. This plot may be used to identify clusters in the data and to identify variables associated with prominent signal in the data. Variables that are associated with these leading principal components should be considered in downstream analyses.   
                                      
                                       
                                  
                                
                                
                                  
                                       
                                      
                                           Principal Components of VEGFR2 mediated cell proliferation Data   More Plot Information                      Principal Components of VEGFR2 mediated cell proliferation Data  Principal component analysis maps high-dimensional datasets onto a smaller number of highly informative dimensions. Here, the first four principal components of the gene expression data are plotted against each other and colored by the values of the selected covariate. This plot may be used to identify clusters in the data and to identify variables associated with prominent signal in the data. Variables that are associated with these leading principal components should be considered in downstream analyses.   
                                      
                                       
                                  
                                
                              
                            
                           
                            
                          
                           
                      
                    
                    
                      
                           
                          
                               Heatmaps    PCA     
                              
                                  Heatmaps  
                              
                               
                              
                                  Line  
                                  Day  
                              
                              
                            
                              
                                
                                  
                                       
                                      
                                           Heatmap of VEGFR2 mediated vascular permeability Data   More Plot Information                      Heatmap of VEGFR2 mediated vascular permeability Data  Heatmap of the normalized data, scaled to give all genes equal variance, generated via unsupervised clustering. Orange indicates high expression; blue indicates low expression. This plot is meant to provide a high level exploratory view of the data.   
                                      
                                       
                                  
                                
                              
                            
                            
                              
                                
                                  
                                       
                                      
                                           Principal Components of VEGFR2 mediated vascular permeability Data   More Plot Information                      Principal Components of VEGFR2 mediated vascular permeability Data  Principal component analysis maps high-dimensional datasets onto a smaller number of highly informative dimensions. Here, the first four principal components of the gene expression data are plotted against each other and colored by the values of the selected covariate. This plot may be used to identify clusters in the data and to identify variables associated with prominent signal in the data. Variables that are associated with these leading principal components should be considered in downstream analyses.   
                                      
                                       
                                  
                                
                                
                                  
                                       
                                      
                                           Principal Components of VEGFR2 mediated vascular permeability Data   More Plot Information                      Principal Components of VEGFR2 mediated vascular permeability Data  Principal component analysis maps high-dimensional datasets onto a smaller number of highly informative dimensions. Here, the first four principal components of the gene expression data are plotted against each other and colored by the values of the selected covariate. This plot may be used to identify clusters in the data and to identify variables associated with prominent signal in the data. Variables that are associated with these leading principal components should be considered in downstream analyses.   
                                      
                                       
                                  
                                
                              
                            
                           
                            
                          
                           
                      
                    
                    
                      
                           
                          
                               Heatmaps    PCA     
                              
                                  Heatmaps  
                              
                               
                              
                                  Line  
                                  Day  
                              
                              
                            
                              
                                
                                  
                                       
                                      
                                           Heatmap of WNT mediated activation of DVL Data   More Plot Information                      Heatmap of WNT mediated activation of DVL Data  Heatmap of the normalized data, scaled to give all genes equal variance, generated via unsupervised clustering. Orange indicates high expression; blue indicates low expression. This plot is meant to provide a high level exploratory view of the data.   
                                      
                                       
                                  
                                
                              
                            
                            
                              
                                
                                  
                                       
                                      
                                           Principal Components of WNT mediated activation of DVL Data   More Plot Information                      Principal Components of WNT mediated activation of DVL Data  Principal component analysis maps high-dimensional datasets onto a smaller number of highly informative dimensions. Here, the first four principal components of the gene expression data are plotted against each other and colored by the values of the selected covariate. This plot may be used to identify clusters in the data and to identify variables associated with prominent signal in the data. Variables that are associated with these leading principal components should be considered in downstream analyses.   
                                      
                                       
                                  
                                
                                
                                  
                                       
                                      
                                           Principal Components of WNT mediated activation of DVL Data   More Plot Information                      Principal Components of WNT mediated activation of DVL Data  Principal component analysis maps high-dimensional datasets onto a smaller number of highly informative dimensions. Here, the first four principal components of the gene expression data are plotted against each other and colored by the values of the selected covariate. This plot may be used to identify clusters in the data and to identify variables associated with prominent signal in the data. Variables that are associated with these leading principal components should be considered in downstream analyses.   
                                      
                                       
                                  
                                
                              
                            
                           
                            
                          
                           
                      
                    
                  
                 
              
               
          
        
        
          
                Displays plots that detail the impact of normalization on the data.      
              
                  Analyte: 
                  
                      mRNA  
                  
                  
                  
                    
                      
                           
                          
                               Pairwise Variance during HK Selection   More Plot Information                Pairwise Variance during HK Selection  Displays the geNorm pairwise variation statistic after successive genes are removed. This statistic cannot be computed for the final two genes, which are therefore not displayed. The ideal normalization gene set will minimize the pairwise variation statistic.   
                               Normalization Summary: mRNA   More Plot Information      mRNA Normalization Summary Data      All Normalized Data      mRNA Normalized Data      Download HK Genes                Normalization Summary: mRNA  The lower the MSE of reference profile the higher quality of normalization. The more positive normalization factor, the higher counts were registered for the sample and vice versa   
                          
                           
                      
                    
                  
                 
              
                
          
        
        
          
                Pathway scores condense each sample's gene expression profile into a small set of pathway scores. An experiment can then be explored through the lens of pathway scores instead of in the much higher-dimension lens of gene expression values. Pathway scores are fit using the first principal component of each gene set's data. They are oriented such that increasing score corresponds to mostly increasing expression (specifically, each pathway score has positive weights for at least half its genes).
        Summary plots explore the joint behavior of pathways, and Covariates plots compare pathway scores to covariates.      
              
                   Summary    Covariates     
                  
                      score  
                  
                   
                  
                      score  
                  
                  
                
                  
                    
                      
                           
                          
                               Choose a Pathway: 
                              
                                  Summary  
                                  Activation of BAD and translocation to mitochondria   
                                  Activation of Rac  
                                  Apoptotic cleavage of cellular proteins  
                                  ARMS-mediated activation  
                                  AXIN missense mutants destabilize the destruction complex  
                                  Beta-catenin phosphorylation cascade  
                                  CD209 (DC-SIGN) signaling  
                                  CD28 co-stimulation  
                                  CD28 dependent PI3KAkt signaling  
                                  CD28 dependent Vav1 pathway  
                                  Cell-extracellular matrix interactions  
                                  Cell surface interactions at the vascular wall  
                                  Circadian Clock  
                                  Constitutive Signaling by Aberrant PI3K in Cancer  
                                  Constitutive Signaling by EGFRvIII  
                                  CTLA4 inhibitory signaling  
                                  Cyclin D associated events in G1  
                                  DAP12 signaling  
                                  Deactivation of the beta-catenin transactivating complex  
                                  Disassembly of the destruction complex and recruitment of AXIN t...  
                                  Downstream signal transduction  
                                  ECM proteoglycans  
                                  Elastic fibre formation  
                                  EPH-ephrin mediated repulsion of cells  
                                  EPHA-mediated growth cone collapse  
                                  EPHB-mediated forward signaling  
                                  Ephrin signaling  
                                  FCERI mediated MAPK activation  
                                  Formation of the beta-catenin.CL.TCF transactivating complex  
                                  Frs2-mediated activation  
                                  G alpha (1213) signalling events  
                                  G alpha (q) signalling events  
                                  G beta.CL.gamma signalling through PI3Kgamma  
                                  GAB1 signalosome  
                                  Gastrin-CREB signalling pathway via PKC and MAPK  
                                  GPVI-mediated activation cascade  
                                  Integrin alphaIIb beta3 signaling  
                                  Integrin cell surface interactions  
                                  Interleukin receptor SHC signaling  
                                  IRS-related events triggered by IGF1R  
                                  Laminin interactions  
                                  MAP2K and MAPK activation  
                                  MAPK6MAPK4 signaling  
                                  MicroRNA (miRNA) biogenesis  
                                  Misspliced GSK3beta mutants stabilize beta-catenin  
                                  Molecules associated with elastic fibres  
                                  Negative regulation of FGFR1 signaling  
                                  Negative regulation of FGFR2 signaling  
                                  Negative regulation of FGFR3 signaling  
                                  Negative regulation of FGFR4 signaling  
                                  Negative regulation of MAPK pathway  
                                  Nephrin interactions  
                                  Non-integrin membrane-ECM interactions  
                                  NRAGE signals death through JNK  
                                  PCPCE pathway  
                                  PI3K Cascade  
                                  PI3KAKT activation  
                                  PIP3 activates AKT signaling  
                                  Platelet degranulation   
                                  RAF activation  
                                  RAFMAP kinase cascade  
                                  Regulation of actin dynamics for phagocytic cup formation  
                                  Regulation of signaling by CBL  
                                  Repression of WNT target genes  
                                  Rho GTPase cycle  
                                  RHO GTPases activate CIT  
                                  RHO GTPases Activate Formins  
                                  RHO GTPases activate IQGAPs  
                                  RHO GTPases activate PAKs  
                                  RHO GTPases activate PKNs  
                                  RHO GTPases Activate ROCKs  
                                  RHO GTPases Activate WASPs and WAVEs  
                                  Role of LAT2NTALLAB on calcium mobilization  
                                  S33 mutants of beta-catenin aren.SQT.t phosphorylated  
                                  S37 mutants of beta-catenin aren.SQT.t phosphorylated  
                                  S45 mutants of beta-catenin aren.SQT.t phosphorylated  
                                  Sema3A PAK dependent Axon repulsion  
                                  Sema4D induced cell migration and growth-cone collapse  
                                  SHC-mediated cascade.CL.FGFR1  
                                  SHC-related events triggered by IGF1R  
                                  Signal transduction by L1  
                                  Signaling by ERBB2  
                                  Signaling by FGFR1 in disease  
                                  Signaling by FGFR2 in disease  
                                  Signaling by FGFR3 in disease  
                                  Signaling by FGFR4 in disease  
                                  Signaling by PDGF  
                                  Signaling by SCF-KIT  
                                  Smooth Muscle Contraction  
                                  Spry regulation of FGF signaling  
                                  Syndecan interactions  
                                  Synthesis of PIPs at the plasma membrane  
                                  T41 mutants of beta-catenin aren.SQT.t phosphorylated  
                                  TGF-beta receptor signaling activates SMADs  
                                  Tie2 Signaling  
                                  Tight junction interactions  
                                  TP53 Regulates Metabolic Genes  
                                  Truncations of AMER1 destabilize the destruction complex  
                                  VEGFA-VEGFR2 Pathway  
                                  VEGFR2 mediated cell proliferation  
                                  VEGFR2 mediated vascular permeability  
                                  WNT mediated activation of DVL  
                              
                               
                              
                                
                                  
                                       
                                      
                                           Heatmap of Pathway scores   More Plot Information      Download Pathway scores                      Heatmap of Pathway scores  Heatmap showing pathway score. This plot is a high level overview of how the pathway scores change across samples. Use it to understand how pathway scores cluster together and which samples exhibit similar pathway score profiles. Orange indicates high scores; blue indicates low scores. Scores are displayed on the same scale via a Z-transformation.   
                                           Heatmap of Correlation Matrix of Pathway scores   More Plot Information                Heatmap of Correlation Matrix of Pathway scores  Heatmap showing the correlation matrix of pathway scores. Orange and blue indicate positive and negative correlation, respectively.   
                                      
                                       
                                  
                                
                                
                                  
                                       
                                      
                                           Activation of BAD and translocation to mitochondria  measurements vs. other pathway scores   More Plot Information                      Activation of BAD and translocation to mitochondria  measurements vs. other pathway scores  Each panel plots Activation of BAD and translocation to mitochondria  against another pathway score.  Points are colored by Day.   
                                           Activation of BAD and translocation to mitochondria  measurements vs. other pathway scores   More Plot Information                      Activation of BAD and translocation to mitochondria  measurements vs. other pathway scores  Each panel plots Activation of BAD and translocation to mitochondria  against another pathway score.  Points are colored by Line.   
                                      
                                       
                                  
                                
                                
                                  
                                       
                                      
                                           Activation of Rac measurements vs. other pathway scores   More Plot Information                      Activation of Rac measurements vs. other pathway scores  Each panel plots Activation of Rac against another pathway score.  Points are colored by Day.   
                                           Activation of Rac measurements vs. other pathway scores   More Plot Information                      Activation of Rac measurements vs. other pathway scores  Each panel plots Activation of Rac against another pathway score.  Points are colored by Line.   
                                      
                                       
                                  
                                
                                
                                  
                                       
                                      
                                           Apoptotic cleavage of cellular proteins measurements vs. other pathway scores   More Plot Information                      Apoptotic cleavage of cellular proteins measurements vs. other pathway scores  Each panel plots Apoptotic cleavage of cellular proteins against another pathway score.  Points are colored by Day.   
                                           Apoptotic cleavage of cellular proteins measurements vs. other pathway scores   More Plot Information                      Apoptotic cleavage of cellular proteins measurements vs. other pathway scores  Each panel plots Apoptotic cleavage of cellular proteins against another pathway score.  Points are colored by Line.   
                                      
                                       
                                  
                                
                                
                                  
                                       
                                      
                                           ARMS-mediated activation measurements vs. other pathway scores   More Plot Information                      ARMS-mediated activation measurements vs. other pathway scores  Each panel plots ARMS-mediated activation against another pathway score.  Points are colored by Day.   
                                           ARMS-mediated activation measurements vs. other pathway scores   More Plot Information                      ARMS-mediated activation measurements vs. other pathway scores  Each panel plots ARMS-mediated activation against another pathway score.  Points are colored by Line.   
                                      
                                       
                                  
                                
                                
                                  
                                       
                                      
                                           AXIN missense mutants destabilize the destruction complex measurements vs. other pathway scores   More Plot Information                      AXIN missense mutants destabilize the destruction complex measurements vs. other pathway scores  Each panel plots AXIN missense mutants destabilize the destruction complex against another pathway score.  Points are colored by Day.   
                                           AXIN missense mutants destabilize the destruction complex measurements vs. other pathway scores   More Plot Information                      AXIN missense mutants destabilize the destruction complex measurements vs. other pathway scores  Each panel plots AXIN missense mutants destabilize the destruction complex against another pathway score.  Points are colored by Line.   
                                      
                                       
                                  
                                
                                
                                  
                                       
                                      
                                           Beta-catenin phosphorylation cascade measurements vs. other pathway scores   More Plot Information                      Beta-catenin phosphorylation cascade measurements vs. other pathway scores  Each panel plots Beta-catenin phosphorylation cascade against another pathway score.  Points are colored by Day.   
                                           Beta-catenin phosphorylation cascade measurements vs. other pathway scores   More Plot Information                      Beta-catenin phosphorylation cascade measurements vs. other pathway scores  Each panel plots Beta-catenin phosphorylation cascade against another pathway score.  Points are colored by Line.   
                                      
                                       
                                  
                                
                                
                                  
                                       
                                      
                                           CD209 (DC-SIGN) signaling measurements vs. other pathway scores   More Plot Information                      CD209 (DC-SIGN) signaling measurements vs. other pathway scores  Each panel plots CD209 (DC-SIGN) signaling against another pathway score.  Points are colored by Day.   
                                           CD209 (DC-SIGN) signaling measurements vs. other pathway scores   More Plot Information                      CD209 (DC-SIGN) signaling measurements vs. other pathway scores  Each panel plots CD209 (DC-SIGN) signaling against another pathway score.  Points are colored by Line.   
                                      
                                       
                                  
                                
                                
                                  
                                       
                                      
                                           CD28 co-stimulation measurements vs. other pathway scores   More Plot Information                      CD28 co-stimulation measurements vs. other pathway scores  Each panel plots CD28 co-stimulation against another pathway score.  Points are colored by Day.   
                                           CD28 co-stimulation measurements vs. other pathway scores   More Plot Information                      CD28 co-stimulation measurements vs. other pathway scores  Each panel plots CD28 co-stimulation against another pathway score.  Points are colored by Line.   
                                      
                                       
                                  
                                
                                
                                  
                                       
                                      
                                           CD28 dependent PI3KAkt signaling measurements vs. other pathway scores   More Plot Information                      CD28 dependent PI3KAkt signaling measurements vs. other pathway scores  Each panel plots CD28 dependent PI3KAkt signaling against another pathway score.  Points are colored by Day.   
                                           CD28 dependent PI3KAkt signaling measurements vs. other pathway scores   More Plot Information                      CD28 dependent PI3KAkt signaling measurements vs. other pathway scores  Each panel plots CD28 dependent PI3KAkt signaling against another pathway score.  Points are colored by Line.   
                                      
                                       
                                  
                                
                                
                                  
                                       
                                      
                                           CD28 dependent Vav1 pathway measurements vs. other pathway scores   More Plot Information                      CD28 dependent Vav1 pathway measurements vs. other pathway scores  Each panel plots CD28 dependent Vav1 pathway against another pathway score.  Points are colored by Day.   
                                           CD28 dependent Vav1 pathway measurements vs. other pathway scores   More Plot Information                      CD28 dependent Vav1 pathway measurements vs. other pathway scores  Each panel plots CD28 dependent Vav1 pathway against another pathway score.  Points are colored by Line.   
                                      
                                       
                                  
                                
                                
                                  
                                       
                                      
                                           Cell-extracellular matrix interactions measurements vs. other pathway scores   More Plot Information                      Cell-extracellular matrix interactions measurements vs. other pathway scores  Each panel plots Cell-extracellular matrix interactions against another pathway score.  Points are colored by Day.   
                                           Cell-extracellular matrix interactions measurements vs. other pathway scores   More Plot Information                      Cell-extracellular matrix interactions measurements vs. other pathway scores  Each panel plots Cell-extracellular matrix interactions against another pathway score.  Points are colored by Line.   
                                      
                                       
                                  
                                
                                
                                  
                                       
                                      
                                           Cell surface interactions at the vascular wall measurements vs. other pathway scores   More Plot Information                      Cell surface interactions at the vascular wall measurements vs. other pathway scores  Each panel plots Cell surface interactions at the vascular wall against another pathway score.  Points are colored by Day.   
                                           Cell surface interactions at the vascular wall measurements vs. other pathway scores   More Plot Information                      Cell surface interactions at the vascular wall measurements vs. other pathway scores  Each panel plots Cell surface interactions at the vascular wall against another pathway score.  Points are colored by Line.   
                                      
                                       
                                  
                                
                                
                                  
                                       
                                      
                                           Circadian Clock measurements vs. other pathway scores   More Plot Information                      Circadian Clock measurements vs. other pathway scores  Each panel plots Circadian Clock against another pathway score.  Points are colored by Day.   
                                           Circadian Clock measurements vs. other pathway scores   More Plot Information                      Circadian Clock measurements vs. other pathway scores  Each panel plots Circadian Clock against another pathway score.  Points are colored by Line.   
                                      
                                       
                                  
                                
                                
                                  
                                       
                                      
                                           Constitutive Signaling by Aberrant PI3K in Cancer measurements vs. other pathway scores   More Plot Information                      Constitutive Signaling by Aberrant PI3K in Cancer measurements vs. other pathway scores  Each panel plots Constitutive Signaling by Aberrant PI3K in Cancer against another pathway score.  Points are colored by Day.   
                                           Constitutive Signaling by Aberrant PI3K in Cancer measurements vs. other pathway scores   More Plot Information                      Constitutive Signaling by Aberrant PI3K in Cancer measurements vs. other pathway scores  Each panel plots Constitutive Signaling by Aberrant PI3K in Cancer against another pathway score.  Points are colored by Line.   
                                      
                                       
                                  
                                
                                
                                  
                                       
                                      
                                           Constitutive Signaling by EGFRvIII measurements vs. other pathway scores   More Plot Information                      Constitutive Signaling by EGFRvIII measurements vs. other pathway scores  Each panel plots Constitutive Signaling by EGFRvIII against another pathway score.  Points are colored by Day.   
                                           Constitutive Signaling by EGFRvIII measurements vs. other pathway scores   More Plot Information                      Constitutive Signaling by EGFRvIII measurements vs. other pathway scores  Each panel plots Constitutive Signaling by EGFRvIII against another pathway score.  Points are colored by Line.   
                                      
                                       
                                  
                                
                                
                                  
                                       
                                      
                                           CTLA4 inhibitory signaling measurements vs. other pathway scores   More Plot Information                      CTLA4 inhibitory signaling measurements vs. other pathway scores  Each panel plots CTLA4 inhibitory signaling against another pathway score.  Points are colored by Day.   
                                           CTLA4 inhibitory signaling measurements vs. other pathway scores   More Plot Information                      CTLA4 inhibitory signaling measurements vs. other pathway scores  Each panel plots CTLA4 inhibitory signaling against another pathway score.  Points are colored by Line.   
                                      
                                       
                                  
                                
                                
                                  
                                       
                                      
                                           Cyclin D associated events in G1 measurements vs. other pathway scores   More Plot Information                      Cyclin D associated events in G1 measurements vs. other pathway scores  Each panel plots Cyclin D associated events in G1 against another pathway score.  Points are colored by Day.   
                                           Cyclin D associated events in G1 measurements vs. other pathway scores   More Plot Information                      Cyclin D associated events in G1 measurements vs. other pathway scores  Each panel plots Cyclin D associated events in G1 against another pathway score.  Points are colored by Line.   
                                      
                                       
                                  
                                
                                
                                  
                                       
                                      
                                           DAP12 signaling measurements vs. other pathway scores   More Plot Information                      DAP12 signaling measurements vs. other pathway scores  Each panel plots DAP12 signaling against another pathway score.  Points are colored by Day.   
                                           DAP12 signaling measurements vs. other pathway scores   More Plot Information                      DAP12 signaling measurements vs. other pathway scores  Each panel plots DAP12 signaling against another pathway score.  Points are colored by Line.   
                                      
                                       
                                  
                                
                                
                                  
                                       
                                      
                                           Deactivation of the beta-catenin transactivating complex measurements vs. other pathway scores   More Plot Information                      Deactivation of the beta-catenin transactivating complex measurements vs. other pathway scores  Each panel plots Deactivation of the beta-catenin transactivating complex against another pathway score.  Points are colored by Day.   
                                           Deactivation of the beta-catenin transactivating complex measurements vs. other pathway scores   More Plot Information                      Deactivation of the beta-catenin transactivating complex measurements vs. other pathway scores  Each panel plots Deactivation of the beta-catenin transactivating complex against another pathway score.  Points are colored by Line.   
                                      
                                       
                                  
                                
                                
                                  
                                       
                                      
                                           Disassembly of the destruction complex and recruitment of AXIN to the membrane measurements vs. other pathway scores   More Plot Information                      Disassembly of the destruction complex and recruitment of AXIN to the membrane measurements vs. other pathway scores  Each panel plots Disassembly of the destruction complex and recruitment of AXIN to the membrane against another pathway score.  Points are colored by Day.   
                                           Disassembly of the destruction complex and recruitment of AXIN to the membrane measurements vs. other pathway scores   More Plot Information                      Disassembly of the destruction complex and recruitment of AXIN to the membrane measurements vs. other pathway scores  Each panel plots Disassembly of the destruction complex and recruitment of AXIN to the membrane against another pathway score.  Points are colored by Line.   
                                      
                                       
                                  
                                
                                
                                  
                                       
                                      
                                           Downstream signal transduction measurements vs. other pathway scores   More Plot Information                      Downstream signal transduction measurements vs. other pathway scores  Each panel plots Downstream signal transduction against another pathway score.  Points are colored by Day.   
                                           Downstream signal transduction measurements vs. other pathway scores   More Plot Information                      Downstream signal transduction measurements vs. other pathway scores  Each panel plots Downstream signal transduction against another pathway score.  Points are colored by Line.   
                                      
                                       
                                  
                                
                                
                                  
                                       
                                      
                                           ECM proteoglycans measurements vs. other pathway scores   More Plot Information                      ECM proteoglycans measurements vs. other pathway scores  Each panel plots ECM proteoglycans against another pathway score.  Points are colored by Day.   
                                           ECM proteoglycans measurements vs. other pathway scores   More Plot Information                      ECM proteoglycans measurements vs. other pathway scores  Each panel plots ECM proteoglycans against another pathway score.  Points are colored by Line.   
                                      
                                       
                                  
                                
                                
                                  
                                       
                                      
                                           Elastic fibre formation measurements vs. other pathway scores   More Plot Information                      Elastic fibre formation measurements vs. other pathway scores  Each panel plots Elastic fibre formation against another pathway score.  Points are colored by Day.   
                                           Elastic fibre formation measurements vs. other pathway scores   More Plot Information                      Elastic fibre formation measurements vs. other pathway scores  Each panel plots Elastic fibre formation against another pathway score.  Points are colored by Line.   
                                      
                                       
                                  
                                
                                
                                  
                                       
                                      
                                           EPH-ephrin mediated repulsion of cells measurements vs. other pathway scores   More Plot Information                      EPH-ephrin mediated repulsion of cells measurements vs. other pathway scores  Each panel plots EPH-ephrin mediated repulsion of cells against another pathway score.  Points are colored by Day.   
                                           EPH-ephrin mediated repulsion of cells measurements vs. other pathway scores   More Plot Information                      EPH-ephrin mediated repulsion of cells measurements vs. other pathway scores  Each panel plots EPH-ephrin mediated repulsion of cells against another pathway score.  Points are colored by Line.   
                                      
                                       
                                  
                                
                                
                                  
                                       
                                      
                                           EPHA-mediated growth cone collapse measurements vs. other pathway scores   More Plot Information                      EPHA-mediated growth cone collapse measurements vs. other pathway scores  Each panel plots EPHA-mediated growth cone collapse against another pathway score.  Points are colored by Day.   
                                           EPHA-mediated growth cone collapse measurements vs. other pathway scores   More Plot Information                      EPHA-mediated growth cone collapse measurements vs. other pathway scores  Each panel plots EPHA-mediated growth cone collapse against another pathway score.  Points are colored by Line.   
                                      
                                       
                                  
                                
                                
                                  
                                       
                                      
                                           EPHB-mediated forward signaling measurements vs. other pathway scores   More Plot Information                      EPHB-mediated forward signaling measurements vs. other pathway scores  Each panel plots EPHB-mediated forward signaling against another pathway score.  Points are colored by Day.   
                                           EPHB-mediated forward signaling measurements vs. other pathway scores   More Plot Information                      EPHB-mediated forward signaling measurements vs. other pathway scores  Each panel plots EPHB-mediated forward signaling against another pathway score.  Points are colored by Line.   
                                      
                                       
                                  
                                
                                
                                  
                                       
                                      
                                           Ephrin signaling measurements vs. other pathway scores   More Plot Information                      Ephrin signaling measurements vs. other pathway scores  Each panel plots Ephrin signaling against another pathway score.  Points are colored by Day.   
                                           Ephrin signaling measurements vs. other pathway scores   More Plot Information                      Ephrin signaling measurements vs. other pathway scores  Each panel plots Ephrin signaling against another pathway score.  Points are colored by Line.   
                                      
                                       
                                  
                                
                                
                                  
                                       
                                      
                                           FCERI mediated MAPK activation measurements vs. other pathway scores   More Plot Information                      FCERI mediated MAPK activation measurements vs. other pathway scores  Each panel plots FCERI mediated MAPK activation against another pathway score.  Points are colored by Day.   
                                           FCERI mediated MAPK activation measurements vs. other pathway scores   More Plot Information                      FCERI mediated MAPK activation measurements vs. other pathway scores  Each panel plots FCERI mediated MAPK activation against another pathway score.  Points are colored by Line.   
                                      
                                       
                                  
                                
                                
                                  
                                       
                                      
                                           Formation of the beta-catenin.CL.TCF transactivating complex measurements vs. other pathway scores   More Plot Information                      Formation of the beta-catenin.CL.TCF transactivating complex measurements vs. other pathway scores  Each panel plots Formation of the beta-catenin.CL.TCF transactivating complex against another pathway score.  Points are colored by Day.   
                                           Formation of the beta-catenin.CL.TCF transactivating complex measurements vs. other pathway scores   More Plot Information                      Formation of the beta-catenin.CL.TCF transactivating complex measurements vs. other pathway scores  Each panel plots Formation of the beta-catenin.CL.TCF transactivating complex against another pathway score.  Points are colored by Line.   
                                      
                                       
                                  
                                
                                
                                  
                                       
                                      
                                           Frs2-mediated activation measurements vs. other pathway scores   More Plot Information                      Frs2-mediated activation measurements vs. other pathway scores  Each panel plots Frs2-mediated activation against another pathway score.  Points are colored by Day.   
                                           Frs2-mediated activation measurements vs. other pathway scores   More Plot Information                      Frs2-mediated activation measurements vs. other pathway scores  Each panel plots Frs2-mediated activation against another pathway score.  Points are colored by Line.   
                                      
                                       
                                  
                                
                                
                                  
                                       
                                      
                                           G alpha (1213) signalling events measurements vs. other pathway scores   More Plot Information                      G alpha (1213) signalling events measurements vs. other pathway scores  Each panel plots G alpha (1213) signalling events against another pathway score.  Points are colored by Day.   
                                           G alpha (1213) signalling events measurements vs. other pathway scores   More Plot Information                      G alpha (1213) signalling events measurements vs. other pathway scores  Each panel plots G alpha (1213) signalling events against another pathway score.  Points are colored by Line.   
                                      
                                       
                                  
                                
                                
                                  
                                       
                                      
                                           G alpha (q) signalling events measurements vs. other pathway scores   More Plot Information                      G alpha (q) signalling events measurements vs. other pathway scores  Each panel plots G alpha (q) signalling events against another pathway score.  Points are colored by Day.   
                                           G alpha (q) signalling events measurements vs. other pathway scores   More Plot Information                      G alpha (q) signalling events measurements vs. other pathway scores  Each panel plots G alpha (q) signalling events against another pathway score.  Points are colored by Line.   
                                      
                                       
                                  
                                
                                
                                  
                                       
                                      
                                           G beta.CL.gamma signalling through PI3Kgamma measurements vs. other pathway scores   More Plot Information                      G beta.CL.gamma signalling through PI3Kgamma measurements vs. other pathway scores  Each panel plots G beta.CL.gamma signalling through PI3Kgamma against another pathway score.  Points are colored by Day.   
                                           G beta.CL.gamma signalling through PI3Kgamma measurements vs. other pathway scores   More Plot Information                      G beta.CL.gamma signalling through PI3Kgamma measurements vs. other pathway scores  Each panel plots G beta.CL.gamma signalling through PI3Kgamma against another pathway score.  Points are colored by Line.   
                                      
                                       
                                  
                                
                                
                                  
                                       
                                      
                                           GAB1 signalosome measurements vs. other pathway scores   More Plot Information                      GAB1 signalosome measurements vs. other pathway scores  Each panel plots GAB1 signalosome against another pathway score.  Points are colored by Day.   
                                           GAB1 signalosome measurements vs. other pathway scores   More Plot Information                      GAB1 signalosome measurements vs. other pathway scores  Each panel plots GAB1 signalosome against another pathway score.  Points are colored by Line.   
                                      
                                       
                                  
                                
                                
                                  
                                       
                                      
                                           Gastrin-CREB signalling pathway via PKC and MAPK measurements vs. other pathway scores   More Plot Information                      Gastrin-CREB signalling pathway via PKC and MAPK measurements vs. other pathway scores  Each panel plots Gastrin-CREB signalling pathway via PKC and MAPK against another pathway score.  Points are colored by Day.   
                                           Gastrin-CREB signalling pathway via PKC and MAPK measurements vs. other pathway scores   More Plot Information                      Gastrin-CREB signalling pathway via PKC and MAPK measurements vs. other pathway scores  Each panel plots Gastrin-CREB signalling pathway via PKC and MAPK against another pathway score.  Points are colored by Line.   
                                      
                                       
                                  
                                
                                
                                  
                                       
                                      
                                           GPVI-mediated activation cascade measurements vs. other pathway scores   More Plot Information                      GPVI-mediated activation cascade measurements vs. other pathway scores  Each panel plots GPVI-mediated activation cascade against another pathway score.  Points are colored by Day.   
                                           GPVI-mediated activation cascade measurements vs. other pathway scores   More Plot Information                      GPVI-mediated activation cascade measurements vs. other pathway scores  Each panel plots GPVI-mediated activation cascade against another pathway score.  Points are colored by Line.   
                                      
                                       
                                  
                                
                                
                                  
                                       
                                      
                                           Integrin alphaIIb beta3 signaling measurements vs. other pathway scores   More Plot Information                      Integrin alphaIIb beta3 signaling measurements vs. other pathway scores  Each panel plots Integrin alphaIIb beta3 signaling against another pathway score.  Points are colored by Day.   
                                           Integrin alphaIIb beta3 signaling measurements vs. other pathway scores   More Plot Information                      Integrin alphaIIb beta3 signaling measurements vs. other pathway scores  Each panel plots Integrin alphaIIb beta3 signaling against another pathway score.  Points are colored by Line.   
                                      
                                       
                                  
                                
                                
                                  
                                       
                                      
                                           Integrin cell surface interactions measurements vs. other pathway scores   More Plot Information                      Integrin cell surface interactions measurements vs. other pathway scores  Each panel plots Integrin cell surface interactions against another pathway score.  Points are colored by Day.   
                                           Integrin cell surface interactions measurements vs. other pathway scores   More Plot Information                      Integrin cell surface interactions measurements vs. other pathway scores  Each panel plots Integrin cell surface interactions against another pathway score.  Points are colored by Line.   
                                      
                                       
                                  
                                
                                
                                  
                                       
                                      
                                           Interleukin receptor SHC signaling measurements vs. other pathway scores   More Plot Information                      Interleukin receptor SHC signaling measurements vs. other pathway scores  Each panel plots Interleukin receptor SHC signaling against another pathway score.  Points are colored by Day.   
                                           Interleukin receptor SHC signaling measurements vs. other pathway scores   More Plot Information                      Interleukin receptor SHC signaling measurements vs. other pathway scores  Each panel plots Interleukin receptor SHC signaling against another pathway score.  Points are colored by Line.   
                                      
                                       
                                  
                                
                                
                                  
                                       
                                      
                                           IRS-related events triggered by IGF1R measurements vs. other pathway scores   More Plot Information                      IRS-related events triggered by IGF1R measurements vs. other pathway scores  Each panel plots IRS-related events triggered by IGF1R against another pathway score.  Points are colored by Day.   
                                           IRS-related events triggered by IGF1R measurements vs. other pathway scores   More Plot Information                      IRS-related events triggered by IGF1R measurements vs. other pathway scores  Each panel plots IRS-related events triggered by IGF1R against another pathway score.  Points are colored by Line.   
                                      
                                       
                                  
                                
                                
                                  
                                       
                                      
                                           Laminin interactions measurements vs. other pathway scores   More Plot Information                      Laminin interactions measurements vs. other pathway scores  Each panel plots Laminin interactions against another pathway score.  Points are colored by Day.   
                                           Laminin interactions measurements vs. other pathway scores   More Plot Information                      Laminin interactions measurements vs. other pathway scores  Each panel plots Laminin interactions against another pathway score.  Points are colored by Line.   
                                      
                                       
                                  
                                
                                
                                  
                                       
                                      
                                           MAP2K and MAPK activation measurements vs. other pathway scores   More Plot Information                      MAP2K and MAPK activation measurements vs. other pathway scores  Each panel plots MAP2K and MAPK activation against another pathway score.  Points are colored by Day.   
                                           MAP2K and MAPK activation measurements vs. other pathway scores   More Plot Information                      MAP2K and MAPK activation measurements vs. other pathway scores  Each panel plots MAP2K and MAPK activation against another pathway score.  Points are colored by Line.   
                                      
                                       
                                  
                                
                                
                                  
                                       
                                      
                                           MAPK6MAPK4 signaling measurements vs. other pathway scores   More Plot Information                      MAPK6MAPK4 signaling measurements vs. other pathway scores  Each panel plots MAPK6MAPK4 signaling against another pathway score.  Points are colored by Day.   
                                           MAPK6MAPK4 signaling measurements vs. other pathway scores   More Plot Information                      MAPK6MAPK4 signaling measurements vs. other pathway scores  Each panel plots MAPK6MAPK4 signaling against another pathway score.  Points are colored by Line.   
                                      
                                       
                                  
                                
                                
                                  
                                       
                                      
                                           MicroRNA (miRNA) biogenesis measurements vs. other pathway scores   More Plot Information                      MicroRNA (miRNA) biogenesis measurements vs. other pathway scores  Each panel plots MicroRNA (miRNA) biogenesis against another pathway score.  Points are colored by Day.   
                                           MicroRNA (miRNA) biogenesis measurements vs. other pathway scores   More Plot Information                      MicroRNA (miRNA) biogenesis measurements vs. other pathway scores  Each panel plots MicroRNA (miRNA) biogenesis against another pathway score.  Points are colored by Line.   
                                      
                                       
                                  
                                
                                
                                  
                                       
                                      
                                           Misspliced GSK3beta mutants stabilize beta-catenin measurements vs. other pathway scores   More Plot Information                      Misspliced GSK3beta mutants stabilize beta-catenin measurements vs. other pathway scores  Each panel plots Misspliced GSK3beta mutants stabilize beta-catenin against another pathway score.  Points are colored by Day.   
                                           Misspliced GSK3beta mutants stabilize beta-catenin measurements vs. other pathway scores   More Plot Information                      Misspliced GSK3beta mutants stabilize beta-catenin measurements vs. other pathway scores  Each panel plots Misspliced GSK3beta mutants stabilize beta-catenin against another pathway score.  Points are colored by Line.   
                                      
                                       
                                  
                                
                                
                                  
                                       
                                      
                                           Molecules associated with elastic fibres measurements vs. other pathway scores   More Plot Information                      Molecules associated with elastic fibres measurements vs. other pathway scores  Each panel plots Molecules associated with elastic fibres against another pathway score.  Points are colored by Day.   
                                           Molecules associated with elastic fibres measurements vs. other pathway scores   More Plot Information                      Molecules associated with elastic fibres measurements vs. other pathway scores  Each panel plots Molecules associated with elastic fibres against another pathway score.  Points are colored by Line.   
                                      
                                       
                                  
                                
                                
                                  
                                       
                                      
                                           Negative regulation of FGFR1 signaling measurements vs. other pathway scores   More Plot Information                      Negative regulation of FGFR1 signaling measurements vs. other pathway scores  Each panel plots Negative regulation of FGFR1 signaling against another pathway score.  Points are colored by Day.   
                                           Negative regulation of FGFR1 signaling measurements vs. other pathway scores   More Plot Information                      Negative regulation of FGFR1 signaling measurements vs. other pathway scores  Each panel plots Negative regulation of FGFR1 signaling against another pathway score.  Points are colored by Line.   
                                      
                                       
                                  
                                
                                
                                  
                                       
                                      
                                           Negative regulation of FGFR2 signaling measurements vs. other pathway scores   More Plot Information                      Negative regulation of FGFR2 signaling measurements vs. other pathway scores  Each panel plots Negative regulation of FGFR2 signaling against another pathway score.  Points are colored by Day.   
                                           Negative regulation of FGFR2 signaling measurements vs. other pathway scores   More Plot Information                      Negative regulation of FGFR2 signaling measurements vs. other pathway scores  Each panel plots Negative regulation of FGFR2 signaling against another pathway score.  Points are colored by Line.   
                                      
                                       
                                  
                                
                                
                                  
                                       
                                      
                                           Negative regulation of FGFR3 signaling measurements vs. other pathway scores   More Plot Information                      Negative regulation of FGFR3 signaling measurements vs. other pathway scores  Each panel plots Negative regulation of FGFR3 signaling against another pathway score.  Points are colored by Day.   
                                           Negative regulation of FGFR3 signaling measurements vs. other pathway scores   More Plot Information                      Negative regulation of FGFR3 signaling measurements vs. other pathway scores  Each panel plots Negative regulation of FGFR3 signaling against another pathway score.  Points are colored by Line.   
                                      
                                       
                                  
                                
                                
                                  
                                       
                                      
                                           Negative regulation of FGFR4 signaling measurements vs. other pathway scores   More Plot Information                      Negative regulation of FGFR4 signaling measurements vs. other pathway scores  Each panel plots Negative regulation of FGFR4 signaling against another pathway score.  Points are colored by Day.   
                                           Negative regulation of FGFR4 signaling measurements vs. other pathway scores   More Plot Information                      Negative regulation of FGFR4 signaling measurements vs. other pathway scores  Each panel plots Negative regulation of FGFR4 signaling against another pathway score.  Points are colored by Line.   
                                      
                                       
                                  
                                
                                
                                  
                                       
                                      
                                           Negative regulation of MAPK pathway measurements vs. other pathway scores   More Plot Information                      Negative regulation of MAPK pathway measurements vs. other pathway scores  Each panel plots Negative regulation of MAPK pathway against another pathway score.  Points are colored by Day.   
                                           Negative regulation of MAPK pathway measurements vs. other pathway scores   More Plot Information                      Negative regulation of MAPK pathway measurements vs. other pathway scores  Each panel plots Negative regulation of MAPK pathway against another pathway score.  Points are colored by Line.   
                                      
                                       
                                  
                                
                                
                                  
                                       
                                      
                                           Nephrin interactions measurements vs. other pathway scores   More Plot Information                      Nephrin interactions measurements vs. other pathway scores  Each panel plots Nephrin interactions against another pathway score.  Points are colored by Day.   
                                           Nephrin interactions measurements vs. other pathway scores   More Plot Information                      Nephrin interactions measurements vs. other pathway scores  Each panel plots Nephrin interactions against another pathway score.  Points are colored by Line.   
                                      
                                       
                                  
                                
                                
                                  
                                       
                                      
                                           Non-integrin membrane-ECM interactions measurements vs. other pathway scores   More Plot Information                      Non-integrin membrane-ECM interactions measurements vs. other pathway scores  Each panel plots Non-integrin membrane-ECM interactions against another pathway score.  Points are colored by Day.   
                                           Non-integrin membrane-ECM interactions measurements vs. other pathway scores   More Plot Information                      Non-integrin membrane-ECM interactions measurements vs. other pathway scores  Each panel plots Non-integrin membrane-ECM interactions against another pathway score.  Points are colored by Line.   
                                      
                                       
                                  
                                
                                
                                  
                                       
                                      
                                           NRAGE signals death through JNK measurements vs. other pathway scores   More Plot Information                      NRAGE signals death through JNK measurements vs. other pathway scores  Each panel plots NRAGE signals death through JNK against another pathway score.  Points are colored by Day.   
                                           NRAGE signals death through JNK measurements vs. other pathway scores   More Plot Information                      NRAGE signals death through JNK measurements vs. other pathway scores  Each panel plots NRAGE signals death through JNK against another pathway score.  Points are colored by Line.   
                                      
                                       
                                  
                                
                                
                                  
                                       
                                      
                                           PCPCE pathway measurements vs. other pathway scores   More Plot Information                      PCPCE pathway measurements vs. other pathway scores  Each panel plots PCPCE pathway against another pathway score.  Points are colored by Day.   
                                           PCPCE pathway measurements vs. other pathway scores   More Plot Information                      PCPCE pathway measurements vs. other pathway scores  Each panel plots PCPCE pathway against another pathway score.  Points are colored by Line.   
                                      
                                       
                                  
                                
                                
                                  
                                       
                                      
                                           PI3K Cascade measurements vs. other pathway scores   More Plot Information                      PI3K Cascade measurements vs. other pathway scores  Each panel plots PI3K Cascade against another pathway score.  Points are colored by Day.   
                                           PI3K Cascade measurements vs. other pathway scores   More Plot Information                      PI3K Cascade measurements vs. other pathway scores  Each panel plots PI3K Cascade against another pathway score.  Points are colored by Line.   
                                      
                                       
                                  
                                
                                
                                  
                                       
                                      
                                           PI3KAKT activation measurements vs. other pathway scores   More Plot Information                      PI3KAKT activation measurements vs. other pathway scores  Each panel plots PI3KAKT activation against another pathway score.  Points are colored by Day.   
                                           PI3KAKT activation measurements vs. other pathway scores   More Plot Information                      PI3KAKT activation measurements vs. other pathway scores  Each panel plots PI3KAKT activation against another pathway score.  Points are colored by Line.   
                                      
                                       
                                  
                                
                                
                                  
                                       
                                      
                                           PIP3 activates AKT signaling measurements vs. other pathway scores   More Plot Information                      PIP3 activates AKT signaling measurements vs. other pathway scores  Each panel plots PIP3 activates AKT signaling against another pathway score.  Points are colored by Day.   
                                           PIP3 activates AKT signaling measurements vs. other pathway scores   More Plot Information                      PIP3 activates AKT signaling measurements vs. other pathway scores  Each panel plots PIP3 activates AKT signaling against another pathway score.  Points are colored by Line.   
                                      
                                       
                                  
                                
                                
                                  
                                       
                                      
                                           Platelet degranulation  measurements vs. other pathway scores   More Plot Information                      Platelet degranulation  measurements vs. other pathway scores  Each panel plots Platelet degranulation  against another pathway score.  Points are colored by Day.   
                                           Platelet degranulation  measurements vs. other pathway scores   More Plot Information                      Platelet degranulation  measurements vs. other pathway scores  Each panel plots Platelet degranulation  against another pathway score.  Points are colored by Line.   
                                      
                                       
                                  
                                
                                
                                  
                                       
                                      
                                           RAF activation measurements vs. other pathway scores   More Plot Information                      RAF activation measurements vs. other pathway scores  Each panel plots RAF activation against another pathway score.  Points are colored by Day.   
                                           RAF activation measurements vs. other pathway scores   More Plot Information                      RAF activation measurements vs. other pathway scores  Each panel plots RAF activation against another pathway score.  Points are colored by Line.   
                                      
                                       
                                  
                                
                                
                                  
                                       
                                      
                                           RAFMAP kinase cascade measurements vs. other pathway scores   More Plot Information                      RAFMAP kinase cascade measurements vs. other pathway scores  Each panel plots RAFMAP kinase cascade against another pathway score.  Points are colored by Day.   
                                           RAFMAP kinase cascade measurements vs. other pathway scores   More Plot Information                      RAFMAP kinase cascade measurements vs. other pathway scores  Each panel plots RAFMAP kinase cascade against another pathway score.  Points are colored by Line.   
                                      
                                       
                                  
                                
                                
                                  
                                       
                                      
                                           Regulation of actin dynamics for phagocytic cup formation measurements vs. other pathway scores   More Plot Information                      Regulation of actin dynamics for phagocytic cup formation measurements vs. other pathway scores  Each panel plots Regulation of actin dynamics for phagocytic cup formation against another pathway score.  Points are colored by Day.   
                                           Regulation of actin dynamics for phagocytic cup formation measurements vs. other pathway scores   More Plot Information                      Regulation of actin dynamics for phagocytic cup formation measurements vs. other pathway scores  Each panel plots Regulation of actin dynamics for phagocytic cup formation against another pathway score.  Points are colored by Line.   
                                      
                                       
                                  
                                
                                
                                  
                                       
                                      
                                           Regulation of signaling by CBL measurements vs. other pathway scores   More Plot Information                      Regulation of signaling by CBL measurements vs. other pathway scores  Each panel plots Regulation of signaling by CBL against another pathway score.  Points are colored by Day.   
                                           Regulation of signaling by CBL measurements vs. other pathway scores   More Plot Information                      Regulation of signaling by CBL measurements vs. other pathway scores  Each panel plots Regulation of signaling by CBL against another pathway score.  Points are colored by Line.   
                                      
                                       
                                  
                                
                                
                                  
                                       
                                      
                                           Repression of WNT target genes measurements vs. other pathway scores   More Plot Information                      Repression of WNT target genes measurements vs. other pathway scores  Each panel plots Repression of WNT target genes against another pathway score.  Points are colored by Day.   
                                           Repression of WNT target genes measurements vs. other pathway scores   More Plot Information                      Repression of WNT target genes measurements vs. other pathway scores  Each panel plots Repression of WNT target genes against another pathway score.  Points are colored by Line.   
                                      
                                       
                                  
                                
                                
                                  
                                       
                                      
                                           Rho GTPase cycle measurements vs. other pathway scores   More Plot Information                      Rho GTPase cycle measurements vs. other pathway scores  Each panel plots Rho GTPase cycle against another pathway score.  Points are colored by Day.   
                                           Rho GTPase cycle measurements vs. other pathway scores   More Plot Information                      Rho GTPase cycle measurements vs. other pathway scores  Each panel plots Rho GTPase cycle against another pathway score.  Points are colored by Line.   
                                      
                                       
                                  
                                
                                
                                  
                                       
                                      
                                           RHO GTPases activate CIT measurements vs. other pathway scores   More Plot Information                      RHO GTPases activate CIT measurements vs. other pathway scores  Each panel plots RHO GTPases activate CIT against another pathway score.  Points are colored by Day.   
                                           RHO GTPases activate CIT measurements vs. other pathway scores   More Plot Information                      RHO GTPases activate CIT measurements vs. other pathway scores  Each panel plots RHO GTPases activate CIT against another pathway score.  Points are colored by Line.   
                                      
                                       
                                  
                                
                                
                                  
                                       
                                      
                                           RHO GTPases Activate Formins measurements vs. other pathway scores   More Plot Information                      RHO GTPases Activate Formins measurements vs. other pathway scores  Each panel plots RHO GTPases Activate Formins against another pathway score.  Points are colored by Day.   
                                           RHO GTPases Activate Formins measurements vs. other pathway scores   More Plot Information                      RHO GTPases Activate Formins measurements vs. other pathway scores  Each panel plots RHO GTPases Activate Formins against another pathway score.  Points are colored by Line.   
                                      
                                       
                                  
                                
                                
                                  
                                       
                                      
                                           RHO GTPases activate IQGAPs measurements vs. other pathway scores   More Plot Information                      RHO GTPases activate IQGAPs measurements vs. other pathway scores  Each panel plots RHO GTPases activate IQGAPs against another pathway score.  Points are colored by Day.   
                                           RHO GTPases activate IQGAPs measurements vs. other pathway scores   More Plot Information                      RHO GTPases activate IQGAPs measurements vs. other pathway scores  Each panel plots RHO GTPases activate IQGAPs against another pathway score.  Points are colored by Line.   
                                      
                                       
                                  
                                
                                
                                  
                                       
                                      
                                           RHO GTPases activate PAKs measurements vs. other pathway scores   More Plot Information                      RHO GTPases activate PAKs measurements vs. other pathway scores  Each panel plots RHO GTPases activate PAKs against another pathway score.  Points are colored by Day.   
                                           RHO GTPases activate PAKs measurements vs. other pathway scores   More Plot Information                      RHO GTPases activate PAKs measurements vs. other pathway scores  Each panel plots RHO GTPases activate PAKs against another pathway score.  Points are colored by Line.   
                                      
                                       
                                  
                                
                                
                                  
                                       
                                      
                                           RHO GTPases activate PKNs measurements vs. other pathway scores   More Plot Information                      RHO GTPases activate PKNs measurements vs. other pathway scores  Each panel plots RHO GTPases activate PKNs against another pathway score.  Points are colored by Day.   
                                           RHO GTPases activate PKNs measurements vs. other pathway scores   More Plot Information                      RHO GTPases activate PKNs measurements vs. other pathway scores  Each panel plots RHO GTPases activate PKNs against another pathway score.  Points are colored by Line.   
                                      
                                       
                                  
                                
                                
                                  
                                       
                                      
                                           RHO GTPases Activate ROCKs measurements vs. other pathway scores   More Plot Information                      RHO GTPases Activate ROCKs measurements vs. other pathway scores  Each panel plots RHO GTPases Activate ROCKs against another pathway score.  Points are colored by Day.   
                                           RHO GTPases Activate ROCKs measurements vs. other pathway scores   More Plot Information                      RHO GTPases Activate ROCKs measurements vs. other pathway scores  Each panel plots RHO GTPases Activate ROCKs against another pathway score.  Points are colored by Line.   
                                      
                                       
                                  
                                
                                
                                  
                                       
                                      
                                           RHO GTPases Activate WASPs and WAVEs measurements vs. other pathway scores   More Plot Information                      RHO GTPases Activate WASPs and WAVEs measurements vs. other pathway scores  Each panel plots RHO GTPases Activate WASPs and WAVEs against another pathway score.  Points are colored by Day.   
                                           RHO GTPases Activate WASPs and WAVEs measurements vs. other pathway scores   More Plot Information                      RHO GTPases Activate WASPs and WAVEs measurements vs. other pathway scores  Each panel plots RHO GTPases Activate WASPs and WAVEs against another pathway score.  Points are colored by Line.   
                                      
                                       
                                  
                                
                                
                                  
                                       
                                      
                                           Role of LAT2NTALLAB on calcium mobilization measurements vs. other pathway scores   More Plot Information                      Role of LAT2NTALLAB on calcium mobilization measurements vs. other pathway scores  Each panel plots Role of LAT2NTALLAB on calcium mobilization against another pathway score.  Points are colored by Day.   
                                           Role of LAT2NTALLAB on calcium mobilization measurements vs. other pathway scores   More Plot Information                      Role of LAT2NTALLAB on calcium mobilization measurements vs. other pathway scores  Each panel plots Role of LAT2NTALLAB on calcium mobilization against another pathway score.  Points are colored by Line.   
                                      
                                       
                                  
                                
                                
                                  
                                       
                                      
                                           S33 mutants of beta-catenin aren.SQT.t phosphorylated measurements vs. other pathway scores   More Plot Information                      S33 mutants of beta-catenin aren.SQT.t phosphorylated measurements vs. other pathway scores  Each panel plots S33 mutants of beta-catenin aren.SQT.t phosphorylated against another pathway score.  Points are colored by Day.   
                                           S33 mutants of beta-catenin aren.SQT.t phosphorylated measurements vs. other pathway scores   More Plot Information                      S33 mutants of beta-catenin aren.SQT.t phosphorylated measurements vs. other pathway scores  Each panel plots S33 mutants of beta-catenin aren.SQT.t phosphorylated against another pathway score.  Points are colored by Line.   
                                      
                                       
                                  
                                
                                
                                  
                                       
                                      
                                           S37 mutants of beta-catenin aren.SQT.t phosphorylated measurements vs. other pathway scores   More Plot Information                      S37 mutants of beta-catenin aren.SQT.t phosphorylated measurements vs. other pathway scores  Each panel plots S37 mutants of beta-catenin aren.SQT.t phosphorylated against another pathway score.  Points are colored by Day.   
                                           S37 mutants of beta-catenin aren.SQT.t phosphorylated measurements vs. other pathway scores   More Plot Information                      S37 mutants of beta-catenin aren.SQT.t phosphorylated measurements vs. other pathway scores  Each panel plots S37 mutants of beta-catenin aren.SQT.t phosphorylated against another pathway score.  Points are colored by Line.   
                                      
                                       
                                  
                                
                                
                                  
                                       
                                      
                                           S45 mutants of beta-catenin aren.SQT.t phosphorylated measurements vs. other pathway scores   More Plot Information                      S45 mutants of beta-catenin aren.SQT.t phosphorylated measurements vs. other pathway scores  Each panel plots S45 mutants of beta-catenin aren.SQT.t phosphorylated against another pathway score.  Points are colored by Day.   
                                           S45 mutants of beta-catenin aren.SQT.t phosphorylated measurements vs. other pathway scores   More Plot Information                      S45 mutants of beta-catenin aren.SQT.t phosphorylated measurements vs. other pathway scores  Each panel plots S45 mutants of beta-catenin aren.SQT.t phosphorylated against another pathway score.  Points are colored by Line.   
                                      
                                       
                                  
                                
                                
                                  
                                       
                                      
                                           Sema3A PAK dependent Axon repulsion measurements vs. other pathway scores   More Plot Information                      Sema3A PAK dependent Axon repulsion measurements vs. other pathway scores  Each panel plots Sema3A PAK dependent Axon repulsion against another pathway score.  Points are colored by Day.   
                                           Sema3A PAK dependent Axon repulsion measurements vs. other pathway scores   More Plot Information                      Sema3A PAK dependent Axon repulsion measurements vs. other pathway scores  Each panel plots Sema3A PAK dependent Axon repulsion against another pathway score.  Points are colored by Line.   
                                      
                                       
                                  
                                
                                
                                  
                                       
                                      
                                           Sema4D induced cell migration and growth-cone collapse measurements vs. other pathway scores   More Plot Information                      Sema4D induced cell migration and growth-cone collapse measurements vs. other pathway scores  Each panel plots Sema4D induced cell migration and growth-cone collapse against another pathway score.  Points are colored by Day.   
                                           Sema4D induced cell migration and growth-cone collapse measurements vs. other pathway scores   More Plot Information                      Sema4D induced cell migration and growth-cone collapse measurements vs. other pathway scores  Each panel plots Sema4D induced cell migration and growth-cone collapse against another pathway score.  Points are colored by Line.   
                                      
                                       
                                  
                                
                                
                                  
                                       
                                      
                                           SHC-mediated cascade.CL.FGFR1 measurements vs. other pathway scores   More Plot Information                      SHC-mediated cascade.CL.FGFR1 measurements vs. other pathway scores  Each panel plots SHC-mediated cascade.CL.FGFR1 against another pathway score.  Points are colored by Day.   
                                           SHC-mediated cascade.CL.FGFR1 measurements vs. other pathway scores   More Plot Information                      SHC-mediated cascade.CL.FGFR1 measurements vs. other pathway scores  Each panel plots SHC-mediated cascade.CL.FGFR1 against another pathway score.  Points are colored by Line.   
                                      
                                       
                                  
                                
                                
                                  
                                       
                                      
                                           SHC-related events triggered by IGF1R measurements vs. other pathway scores   More Plot Information                      SHC-related events triggered by IGF1R measurements vs. other pathway scores  Each panel plots SHC-related events triggered by IGF1R against another pathway score.  Points are colored by Day.   
                                           SHC-related events triggered by IGF1R measurements vs. other pathway scores   More Plot Information                      SHC-related events triggered by IGF1R measurements vs. other pathway scores  Each panel plots SHC-related events triggered by IGF1R against another pathway score.  Points are colored by Line.   
                                      
                                       
                                  
                                
                                
                                  
                                       
                                      
                                           Signal transduction by L1 measurements vs. other pathway scores   More Plot Information                      Signal transduction by L1 measurements vs. other pathway scores  Each panel plots Signal transduction by L1 against another pathway score.  Points are colored by Day.   
                                           Signal transduction by L1 measurements vs. other pathway scores   More Plot Information                      Signal transduction by L1 measurements vs. other pathway scores  Each panel plots Signal transduction by L1 against another pathway score.  Points are colored by Line.   
                                      
                                       
                                  
                                
                                
                                  
                                       
                                      
                                           Signaling by ERBB2 measurements vs. other pathway scores   More Plot Information                      Signaling by ERBB2 measurements vs. other pathway scores  Each panel plots Signaling by ERBB2 against another pathway score.  Points are colored by Day.   
                                           Signaling by ERBB2 measurements vs. other pathway scores   More Plot Information                      Signaling by ERBB2 measurements vs. other pathway scores  Each panel plots Signaling by ERBB2 against another pathway score.  Points are colored by Line.   
                                      
                                       
                                  
                                
                                
                                  
                                       
                                      
                                           Signaling by FGFR1 in disease measurements vs. other pathway scores   More Plot Information                      Signaling by FGFR1 in disease measurements vs. other pathway scores  Each panel plots Signaling by FGFR1 in disease against another pathway score.  Points are colored by Day.   
                                           Signaling by FGFR1 in disease measurements vs. other pathway scores   More Plot Information                      Signaling by FGFR1 in disease measurements vs. other pathway scores  Each panel plots Signaling by FGFR1 in disease against another pathway score.  Points are colored by Line.   
                                      
                                       
                                  
                                
                                
                                  
                                       
                                      
                                           Signaling by FGFR2 in disease measurements vs. other pathway scores   More Plot Information                      Signaling by FGFR2 in disease measurements vs. other pathway scores  Each panel plots Signaling by FGFR2 in disease against another pathway score.  Points are colored by Day.   
                                           Signaling by FGFR2 in disease measurements vs. other pathway scores   More Plot Information                      Signaling by FGFR2 in disease measurements vs. other pathway scores  Each panel plots Signaling by FGFR2 in disease against another pathway score.  Points are colored by Line.   
                                      
                                       
                                  
                                
                                
                                  
                                       
                                      
                                           Signaling by FGFR3 in disease measurements vs. other pathway scores   More Plot Information                      Signaling by FGFR3 in disease measurements vs. other pathway scores  Each panel plots Signaling by FGFR3 in disease against another pathway score.  Points are colored by Day.   
                                           Signaling by FGFR3 in disease measurements vs. other pathway scores   More Plot Information                      Signaling by FGFR3 in disease measurements vs. other pathway scores  Each panel plots Signaling by FGFR3 in disease against another pathway score.  Points are colored by Line.   
                                      
                                       
                                  
                                
                                
                                  
                                       
                                      
                                           Signaling by FGFR4 in disease measurements vs. other pathway scores   More Plot Information                      Signaling by FGFR4 in disease measurements vs. other pathway scores  Each panel plots Signaling by FGFR4 in disease against another pathway score.  Points are colored by Day.   
                                           Signaling by FGFR4 in disease measurements vs. other pathway scores   More Plot Information                      Signaling by FGFR4 in disease measurements vs. other pathway scores  Each panel plots Signaling by FGFR4 in disease against another pathway score.  Points are colored by Line.   
                                      
                                       
                                  
                                
                                
                                  
                                       
                                      
                                           Signaling by PDGF measurements vs. other pathway scores   More Plot Information                      Signaling by PDGF measurements vs. other pathway scores  Each panel plots Signaling by PDGF against another pathway score.  Points are colored by Day.   
                                           Signaling by PDGF measurements vs. other pathway scores   More Plot Information                      Signaling by PDGF measurements vs. other pathway scores  Each panel plots Signaling by PDGF against another pathway score.  Points are colored by Line.   
                                      
                                       
                                  
                                
                                
                                  
                                       
                                      
                                           Signaling by SCF-KIT measurements vs. other pathway scores   More Plot Information                      Signaling by SCF-KIT measurements vs. other pathway scores  Each panel plots Signaling by SCF-KIT against another pathway score.  Points are colored by Day.   
                                           Signaling by SCF-KIT measurements vs. other pathway scores   More Plot Information                      Signaling by SCF-KIT measurements vs. other pathway scores  Each panel plots Signaling by SCF-KIT against another pathway score.  Points are colored by Line.   
                                      
                                       
                                  
                                
                                
                                  
                                       
                                      
                                           Smooth Muscle Contraction measurements vs. other pathway scores   More Plot Information                      Smooth Muscle Contraction measurements vs. other pathway scores  Each panel plots Smooth Muscle Contraction against another pathway score.  Points are colored by Day.   
                                           Smooth Muscle Contraction measurements vs. other pathway scores   More Plot Information                      Smooth Muscle Contraction measurements vs. other pathway scores  Each panel plots Smooth Muscle Contraction against another pathway score.  Points are colored by Line.   
                                      
                                       
                                  
                                
                                
                                  
                                       
                                      
                                           Spry regulation of FGF signaling measurements vs. other pathway scores   More Plot Information                      Spry regulation of FGF signaling measurements vs. other pathway scores  Each panel plots Spry regulation of FGF signaling against another pathway score.  Points are colored by Day.   
                                           Spry regulation of FGF signaling measurements vs. other pathway scores   More Plot Information                      Spry regulation of FGF signaling measurements vs. other pathway scores  Each panel plots Spry regulation of FGF signaling against another pathway score.  Points are colored by Line.   
                                      
                                       
                                  
                                
                                
                                  
                                       
                                      
                                           Syndecan interactions measurements vs. other pathway scores   More Plot Information                      Syndecan interactions measurements vs. other pathway scores  Each panel plots Syndecan interactions against another pathway score.  Points are colored by Day.   
                                           Syndecan interactions measurements vs. other pathway scores   More Plot Information                      Syndecan interactions measurements vs. other pathway scores  Each panel plots Syndecan interactions against another pathway score.  Points are colored by Line.   
                                      
                                       
                                  
                                
                                
                                  
                                       
                                      
                                           Synthesis of PIPs at the plasma membrane measurements vs. other pathway scores   More Plot Information                      Synthesis of PIPs at the plasma membrane measurements vs. other pathway scores  Each panel plots Synthesis of PIPs at the plasma membrane against another pathway score.  Points are colored by Day.   
                                           Synthesis of PIPs at the plasma membrane measurements vs. other pathway scores   More Plot Information                      Synthesis of PIPs at the plasma membrane measurements vs. other pathway scores  Each panel plots Synthesis of PIPs at the plasma membrane against another pathway score.  Points are colored by Line.   
                                      
                                       
                                  
                                
                                
                                  
                                       
                                      
                                           T41 mutants of beta-catenin aren.SQT.t phosphorylated measurements vs. other pathway scores   More Plot Information                      T41 mutants of beta-catenin aren.SQT.t phosphorylated measurements vs. other pathway scores  Each panel plots T41 mutants of beta-catenin aren.SQT.t phosphorylated against another pathway score.  Points are colored by Day.   
                                           T41 mutants of beta-catenin aren.SQT.t phosphorylated measurements vs. other pathway scores   More Plot Information                      T41 mutants of beta-catenin aren.SQT.t phosphorylated measurements vs. other pathway scores  Each panel plots T41 mutants of beta-catenin aren.SQT.t phosphorylated against another pathway score.  Points are colored by Line.   
                                      
                                       
                                  
                                
                                
                                  
                                       
                                      
                                           TGF-beta receptor signaling activates SMADs measurements vs. other pathway scores   More Plot Information                      TGF-beta receptor signaling activates SMADs measurements vs. other pathway scores  Each panel plots TGF-beta receptor signaling activates SMADs against another pathway score.  Points are colored by Day.   
                                           TGF-beta receptor signaling activates SMADs measurements vs. other pathway scores   More Plot Information                      TGF-beta receptor signaling activates SMADs measurements vs. other pathway scores  Each panel plots TGF-beta receptor signaling activates SMADs against another pathway score.  Points are colored by Line.   
                                      
                                       
                                  
                                
                                
                                  
                                       
                                      
                                           Tie2 Signaling measurements vs. other pathway scores   More Plot Information                      Tie2 Signaling measurements vs. other pathway scores  Each panel plots Tie2 Signaling against another pathway score.  Points are colored by Day.   
                                           Tie2 Signaling measurements vs. other pathway scores   More Plot Information                      Tie2 Signaling measurements vs. other pathway scores  Each panel plots Tie2 Signaling against another pathway score.  Points are colored by Line.   
                                      
                                       
                                  
                                
                                
                                  
                                       
                                      
                                           Tight junction interactions measurements vs. other pathway scores   More Plot Information                      Tight junction interactions measurements vs. other pathway scores  Each panel plots Tight junction interactions against another pathway score.  Points are colored by Day.   
                                           Tight junction interactions measurements vs. other pathway scores   More Plot Information                      Tight junction interactions measurements vs. other pathway scores  Each panel plots Tight junction interactions against another pathway score.  Points are colored by Line.   
                                      
                                       
                                  
                                
                                
                                  
                                       
                                      
                                           TP53 Regulates Metabolic Genes measurements vs. other pathway scores   More Plot Information                      TP53 Regulates Metabolic Genes measurements vs. other pathway scores  Each panel plots TP53 Regulates Metabolic Genes against another pathway score.  Points are colored by Day.   
                                           TP53 Regulates Metabolic Genes measurements vs. other pathway scores   More Plot Information                      TP53 Regulates Metabolic Genes measurements vs. other pathway scores  Each panel plots TP53 Regulates Metabolic Genes against another pathway score.  Points are colored by Line.   
                                      
                                       
                                  
                                
                                
                                  
                                       
                                      
                                           Truncations of AMER1 destabilize the destruction complex measurements vs. other pathway scores   More Plot Information                      Truncations of AMER1 destabilize the destruction complex measurements vs. other pathway scores  Each panel plots Truncations of AMER1 destabilize the destruction complex against another pathway score.  Points are colored by Day.   
                                           Truncations of AMER1 destabilize the destruction complex measurements vs. other pathway scores   More Plot Information                      Truncations of AMER1 destabilize the destruction complex measurements vs. other pathway scores  Each panel plots Truncations of AMER1 destabilize the destruction complex against another pathway score.  Points are colored by Line.   
                                      
                                       
                                  
                                
                                
                                  
                                       
                                      
                                           VEGFA-VEGFR2 Pathway measurements vs. other pathway scores   More Plot Information                      VEGFA-VEGFR2 Pathway measurements vs. other pathway scores  Each panel plots VEGFA-VEGFR2 Pathway against another pathway score.  Points are colored by Day.   
                                           VEGFA-VEGFR2 Pathway measurements vs. other pathway scores   More Plot Information                      VEGFA-VEGFR2 Pathway measurements vs. other pathway scores  Each panel plots VEGFA-VEGFR2 Pathway against another pathway score.  Points are colored by Line.   
                                      
                                       
                                  
                                
                                
                                  
                                       
                                      
                                           VEGFR2 mediated cell proliferation measurements vs. other pathway scores   More Plot Information                      VEGFR2 mediated cell proliferation measurements vs. other pathway scores  Each panel plots VEGFR2 mediated cell proliferation against another pathway score.  Points are colored by Day.   
                                           VEGFR2 mediated cell proliferation measurements vs. other pathway scores   More Plot Information                      VEGFR2 mediated cell proliferation measurements vs. other pathway scores  Each panel plots VEGFR2 mediated cell proliferation against another pathway score.  Points are colored by Line.   
                                      
                                       
                                  
                                
                                
                                  
                                       
                                      
                                           VEGFR2 mediated vascular permeability measurements vs. other pathway scores   More Plot Information                      VEGFR2 mediated vascular permeability measurements vs. other pathway scores  Each panel plots VEGFR2 mediated vascular permeability against another pathway score.  Points are colored by Day.   
                                           VEGFR2 mediated vascular permeability measurements vs. other pathway scores   More Plot Information                      VEGFR2 mediated vascular permeability measurements vs. other pathway scores  Each panel plots VEGFR2 mediated vascular permeability against another pathway score.  Points are colored by Line.   
                                      
                                       
                                  
                                
                                
                                  
                                       
                                      
                                           WNT mediated activation of DVL measurements vs. other pathway scores   More Plot Information                      WNT mediated activation of DVL measurements vs. other pathway scores  Each panel plots WNT mediated activation of DVL against another pathway score.  Points are colored by Day.   
                                           WNT mediated activation of DVL measurements vs. other pathway scores   More Plot Information                      WNT mediated activation of DVL measurements vs. other pathway scores  Each panel plots WNT mediated activation of DVL against another pathway score.  Points are colored by Line.   
                                      
                                       
                                  
                                
                              
                             
                          
                           
                      
                    
                  
                
                
                  
                    
                      
                           
                          
                               Choose a Pathway: 
                              
                                  Summary  
                                  Activation of BAD and translocation to mitochondria   
                                  Activation of Rac  
                                  Apoptotic cleavage of cellular proteins  
                                  ARMS-mediated activation  
                                  AXIN missense mutants destabilize the destruction complex  
                                  Beta-catenin phosphorylation cascade  
                                  CD209 (DC-SIGN) signaling  
                                  CD28 co-stimulation  
                                  CD28 dependent PI3KAkt signaling  
                                  CD28 dependent Vav1 pathway  
                                  Cell-extracellular matrix interactions  
                                  Cell surface interactions at the vascular wall  
                                  Circadian Clock  
                                  Constitutive Signaling by Aberrant PI3K in Cancer  
                                  Constitutive Signaling by EGFRvIII  
                                  CTLA4 inhibitory signaling  
                                  Cyclin D associated events in G1  
                                  DAP12 signaling  
                                  Deactivation of the beta-catenin transactivating complex  
                                  Disassembly of the destruction complex and recruitment of AXIN t...  
                                  Downstream signal transduction  
                                  ECM proteoglycans  
                                  Elastic fibre formation  
                                  EPH-ephrin mediated repulsion of cells  
                                  EPHA-mediated growth cone collapse  
                                  EPHB-mediated forward signaling  
                                  Ephrin signaling  
                                  FCERI mediated MAPK activation  
                                  Formation of the beta-catenin.CL.TCF transactivating complex  
                                  Frs2-mediated activation  
                                  G alpha (1213) signalling events  
                                  G alpha (q) signalling events  
                                  G beta.CL.gamma signalling through PI3Kgamma  
                                  GAB1 signalosome  
                                  Gastrin-CREB signalling pathway via PKC and MAPK  
                                  GPVI-mediated activation cascade  
                                  Integrin alphaIIb beta3 signaling  
                                  Integrin cell surface interactions  
                                  Interleukin receptor SHC signaling  
                                  IRS-related events triggered by IGF1R  
                                  Laminin interactions  
                                  MAP2K and MAPK activation  
                                  MAPK6MAPK4 signaling  
                                  MicroRNA (miRNA) biogenesis  
                                  Misspliced GSK3beta mutants stabilize beta-catenin  
                                  Molecules associated with elastic fibres  
                                  Negative regulation of FGFR1 signaling  
                                  Negative regulation of FGFR2 signaling  
                                  Negative regulation of FGFR3 signaling  
                                  Negative regulation of FGFR4 signaling  
                                  Negative regulation of MAPK pathway  
                                  Nephrin interactions  
                                  Non-integrin membrane-ECM interactions  
                                  NRAGE signals death through JNK  
                                  PCPCE pathway  
                                  PI3K Cascade  
                                  PI3KAKT activation  
                                  PIP3 activates AKT signaling  
                                  Platelet degranulation   
                                  RAF activation  
                                  RAFMAP kinase cascade  
                                  Regulation of actin dynamics for phagocytic cup formation  
                                  Regulation of signaling by CBL  
                                  Repression of WNT target genes  
                                  Rho GTPase cycle  
                                  RHO GTPases activate CIT  
                                  RHO GTPases Activate Formins  
                                  RHO GTPases activate IQGAPs  
                                  RHO GTPases activate PAKs  
                                  RHO GTPases activate PKNs  
                                  RHO GTPases Activate ROCKs  
                                  RHO GTPases Activate WASPs and WAVEs  
                                  Role of LAT2NTALLAB on calcium mobilization  
                                  S33 mutants of beta-catenin aren.SQT.t phosphorylated  
                                  S37 mutants of beta-catenin aren.SQT.t phosphorylated  
                                  S45 mutants of beta-catenin aren.SQT.t phosphorylated  
                                  Sema3A PAK dependent Axon repulsion  
                                  Sema4D induced cell migration and growth-cone collapse  
                                  SHC-mediated cascade.CL.FGFR1  
                                  SHC-related events triggered by IGF1R  
                                  Signal transduction by L1  
                                  Signaling by ERBB2  
                                  Signaling by FGFR1 in disease  
                                  Signaling by FGFR2 in disease  
                                  Signaling by FGFR3 in disease  
                                  Signaling by FGFR4 in disease  
                                  Signaling by PDGF  
                                  Signaling by SCF-KIT  
                                  Smooth Muscle Contraction  
                                  Spry regulation of FGF signaling  
                                  Syndecan interactions  
                                  Synthesis of PIPs at the plasma membrane  
                                  T41 mutants of beta-catenin aren.SQT.t phosphorylated  
                                  TGF-beta receptor signaling activates SMADs  
                                  Tie2 Signaling  
                                  Tight junction interactions  
                                  TP53 Regulates Metabolic Genes  
                                  Truncations of AMER1 destabilize the destruction complex  
                                  VEGFA-VEGFR2 Pathway  
                                  VEGFR2 mediated cell proliferation  
                                  VEGFR2 mediated vascular permeability  
                                  WNT mediated activation of DVL  
                              
                               
                              
                                
                                  
                                       
                                      
                                           Pathway scores vs. Day   More Plot Information                Pathway scores vs. Day  Plots the pathway scores against Day. The pathway scores are plotted to show how they vary across different values of Day conditions. Lines show each pathway's average score across values of Day.   
                                           Pathway scores vs. Line   More Plot Information                Pathway scores vs. Line  Plots the pathway scores against Line. The pathway scores are plotted to show how they vary across different values of Line conditions. Lines show each pathway's average score across values of Line.   
                                      
                                       
                                  
                                
                                
                                  
                                       
                                      
                                           Activation of BAD and translocation to mitochondria   pathway scores vs. Day   More Plot Information                Activation of BAD and translocation to mitochondria   pathway scores vs. Day  Plots Activation of BAD and translocation to mitochondria  pathway scores against Day.   
                                           Activation of BAD and translocation to mitochondria   pathway scores vs. Line   More Plot Information                Activation of BAD and translocation to mitochondria   pathway scores vs. Line  Plots Activation of BAD and translocation to mitochondria  pathway scores against Line.   
                                      
                                       
                                  
                                
                                
                                  
                                       
                                      
                                           Activation of Rac  pathway scores vs. Day   More Plot Information                Activation of Rac  pathway scores vs. Day  Plots Activation of Rac pathway scores against Day.   
                                           Activation of Rac  pathway scores vs. Line   More Plot Information                Activation of Rac  pathway scores vs. Line  Plots Activation of Rac pathway scores against Line.   
                                      
                                       
                                  
                                
                                
                                  
                                       
                                      
                                           Apoptotic cleavage of cellular proteins  pathway scores vs. Day   More Plot Information                Apoptotic cleavage of cellular proteins  pathway scores vs. Day  Plots Apoptotic cleavage of cellular proteins pathway scores against Day.   
                                           Apoptotic cleavage of cellular proteins  pathway scores vs. Line   More Plot Information                Apoptotic cleavage of cellular proteins  pathway scores vs. Line  Plots Apoptotic cleavage of cellular proteins pathway scores against Line.   
                                      
                                       
                                  
                                
                                
                                  
                                       
                                      
                                           ARMS-mediated activation  pathway scores vs. Day   More Plot Information                ARMS-mediated activation  pathway scores vs. Day  Plots ARMS-mediated activation pathway scores against Day.   
                                           ARMS-mediated activation  pathway scores vs. Line   More Plot Information                ARMS-mediated activation  pathway scores vs. Line  Plots ARMS-mediated activation pathway scores against Line.   
                                      
                                       
                                  
                                
                                
                                  
                                       
                                      
                                           AXIN missense mutants destabilize the destruction complex  pathway scores vs. Day   More Plot Information                AXIN missense mutants destabilize the destruction complex  pathway scores vs. Day  Plots AXIN missense mutants destabilize the destruction complex pathway scores against Day.   
                                           AXIN missense mutants destabilize the destruction complex  pathway scores vs. Line   More Plot Information                AXIN missense mutants destabilize the destruction complex  pathway scores vs. Line  Plots AXIN missense mutants destabilize the destruction complex pathway scores against Line.   
                                      
                                       
                                  
                                
                                
                                  
                                       
                                      
                                           Beta-catenin phosphorylation cascade  pathway scores vs. Day   More Plot Information                Beta-catenin phosphorylation cascade  pathway scores vs. Day  Plots Beta-catenin phosphorylation cascade pathway scores against Day.   
                                           Beta-catenin phosphorylation cascade  pathway scores vs. Line   More Plot Information                Beta-catenin phosphorylation cascade  pathway scores vs. Line  Plots Beta-catenin phosphorylation cascade pathway scores against Line.   
                                      
                                       
                                  
                                
                                
                                  
                                       
                                      
                                           CD209 (DC-SIGN) signaling  pathway scores vs. Day   More Plot Information                CD209 (DC-SIGN) signaling  pathway scores vs. Day  Plots CD209 (DC-SIGN) signaling pathway scores against Day.   
                                           CD209 (DC-SIGN) signaling  pathway scores vs. Line   More Plot Information                CD209 (DC-SIGN) signaling  pathway scores vs. Line  Plots CD209 (DC-SIGN) signaling pathway scores against Line.   
                                      
                                       
                                  
                                
                                
                                  
                                       
                                      
                                           CD28 co-stimulation  pathway scores vs. Day   More Plot Information                CD28 co-stimulation  pathway scores vs. Day  Plots CD28 co-stimulation pathway scores against Day.   
                                           CD28 co-stimulation  pathway scores vs. Line   More Plot Information                CD28 co-stimulation  pathway scores vs. Line  Plots CD28 co-stimulation pathway scores against Line.   
                                      
                                       
                                  
                                
                                
                                  
                                       
                                      
                                           CD28 dependent PI3KAkt signaling  pathway scores vs. Day   More Plot Information                CD28 dependent PI3KAkt signaling  pathway scores vs. Day  Plots CD28 dependent PI3KAkt signaling pathway scores against Day.   
                                           CD28 dependent PI3KAkt signaling  pathway scores vs. Line   More Plot Information                CD28 dependent PI3KAkt signaling  pathway scores vs. Line  Plots CD28 dependent PI3KAkt signaling pathway scores against Line.   
                                      
                                       
                                  
                                
                                
                                  
                                       
                                      
                                           CD28 dependent Vav1 pathway  pathway scores vs. Day   More Plot Information                CD28 dependent Vav1 pathway  pathway scores vs. Day  Plots CD28 dependent Vav1 pathway pathway scores against Day.   
                                           CD28 dependent Vav1 pathway  pathway scores vs. Line   More Plot Information                CD28 dependent Vav1 pathway  pathway scores vs. Line  Plots CD28 dependent Vav1 pathway pathway scores against Line.   
                                      
                                       
                                  
                                
                                
                                  
                                       
                                      
                                           Cell-extracellular matrix interactions  pathway scores vs. Day   More Plot Information                Cell-extracellular matrix interactions  pathway scores vs. Day  Plots Cell-extracellular matrix interactions pathway scores against Day.   
                                           Cell-extracellular matrix interactions  pathway scores vs. Line   More Plot Information                Cell-extracellular matrix interactions  pathway scores vs. Line  Plots Cell-extracellular matrix interactions pathway scores against Line.   
                                      
                                       
                                  
                                
                                
                                  
                                       
                                      
                                           Cell surface interactions at the vascular wall  pathway scores vs. Day   More Plot Information                Cell surface interactions at the vascular wall  pathway scores vs. Day  Plots Cell surface interactions at the vascular wall pathway scores against Day.   
                                           Cell surface interactions at the vascular wall  pathway scores vs. Line   More Plot Information                Cell surface interactions at the vascular wall  pathway scores vs. Line  Plots Cell surface interactions at the vascular wall pathway scores against Line.   
                                      
                                       
                                  
                                
                                
                                  
                                       
                                      
                                           Circadian Clock  pathway scores vs. Day   More Plot Information                Circadian Clock  pathway scores vs. Day  Plots Circadian Clock pathway scores against Day.   
                                           Circadian Clock  pathway scores vs. Line   More Plot Information                Circadian Clock  pathway scores vs. Line  Plots Circadian Clock pathway scores against Line.   
                                      
                                       
                                  
                                
                                
                                  
                                       
                                      
                                           Constitutive Signaling by Aberrant PI3K in Cancer  pathway scores vs. Day   More Plot Information                Constitutive Signaling by Aberrant PI3K in Cancer  pathway scores vs. Day  Plots Constitutive Signaling by Aberrant PI3K in Cancer pathway scores against Day.   
                                           Constitutive Signaling by Aberrant PI3K in Cancer  pathway scores vs. Line   More Plot Information                Constitutive Signaling by Aberrant PI3K in Cancer  pathway scores vs. Line  Plots Constitutive Signaling by Aberrant PI3K in Cancer pathway scores against Line.   
                                      
                                       
                                  
                                
                                
                                  
                                       
                                      
                                           Constitutive Signaling by EGFRvIII  pathway scores vs. Day   More Plot Information                Constitutive Signaling by EGFRvIII  pathway scores vs. Day  Plots Constitutive Signaling by EGFRvIII pathway scores against Day.   
                                           Constitutive Signaling by EGFRvIII  pathway scores vs. Line   More Plot Information                Constitutive Signaling by EGFRvIII  pathway scores vs. Line  Plots Constitutive Signaling by EGFRvIII pathway scores against Line.   
                                      
                                       
                                  
                                
                                
                                  
                                       
                                      
                                           CTLA4 inhibitory signaling  pathway scores vs. Day   More Plot Information                CTLA4 inhibitory signaling  pathway scores vs. Day  Plots CTLA4 inhibitory signaling pathway scores against Day.   
                                           CTLA4 inhibitory signaling  pathway scores vs. Line   More Plot Information                CTLA4 inhibitory signaling  pathway scores vs. Line  Plots CTLA4 inhibitory signaling pathway scores against Line.   
                                      
                                       
                                  
                                
                                
                                  
                                       
                                      
                                           Cyclin D associated events in G1  pathway scores vs. Day   More Plot Information                Cyclin D associated events in G1  pathway scores vs. Day  Plots Cyclin D associated events in G1 pathway scores against Day.   
                                           Cyclin D associated events in G1  pathway scores vs. Line   More Plot Information                Cyclin D associated events in G1  pathway scores vs. Line  Plots Cyclin D associated events in G1 pathway scores against Line.   
                                      
                                       
                                  
                                
                                
                                  
                                       
                                      
                                           DAP12 signaling  pathway scores vs. Day   More Plot Information                DAP12 signaling  pathway scores vs. Day  Plots DAP12 signaling pathway scores against Day.   
                                           DAP12 signaling  pathway scores vs. Line   More Plot Information                DAP12 signaling  pathway scores vs. Line  Plots DAP12 signaling pathway scores against Line.   
                                      
                                       
                                  
                                
                                
                                  
                                       
                                      
                                           Deactivation of the beta-catenin transactivating complex  pathway scores vs. Day   More Plot Information                Deactivation of the beta-catenin transactivating complex  pathway scores vs. Day  Plots Deactivation of the beta-catenin transactivating complex pathway scores against Day.   
                                           Deactivation of the beta-catenin transactivating complex  pathway scores vs. Line   More Plot Information                Deactivation of the beta-catenin transactivating complex  pathway scores vs. Line  Plots Deactivation of the beta-catenin transactivating complex pathway scores against Line.   
                                      
                                       
                                  
                                
                                
                                  
                                       
                                      
                                           Disassembly of the destruction complex and recruitment of AXIN to the membrane  pathway scores vs. Day   More Plot Information                Disassembly of the destruction complex and recruitment of AXIN to the membrane  pathway scores vs. Day  Plots Disassembly of the destruction complex and recruitment of AXIN to the membrane pathway scores against Day.   
                                           Disassembly of the destruction complex and recruitment of AXIN to the membrane  pathway scores vs. Line   More Plot Information                Disassembly of the destruction complex and recruitment of AXIN to the membrane  pathway scores vs. Line  Plots Disassembly of the destruction complex and recruitment of AXIN to the membrane pathway scores against Line.   
                                      
                                       
                                  
                                
                                
                                  
                                       
                                      
                                           Downstream signal transduction  pathway scores vs. Day   More Plot Information                Downstream signal transduction  pathway scores vs. Day  Plots Downstream signal transduction pathway scores against Day.   
                                           Downstream signal transduction  pathway scores vs. Line   More Plot Information                Downstream signal transduction  pathway scores vs. Line  Plots Downstream signal transduction pathway scores against Line.   
                                      
                                       
                                  
                                
                                
                                  
                                       
                                      
                                           ECM proteoglycans  pathway scores vs. Day   More Plot Information                ECM proteoglycans  pathway scores vs. Day  Plots ECM proteoglycans pathway scores against Day.   
                                           ECM proteoglycans  pathway scores vs. Line   More Plot Information                ECM proteoglycans  pathway scores vs. Line  Plots ECM proteoglycans pathway scores against Line.   
                                      
                                       
                                  
                                
                                
                                  
                                       
                                      
                                           Elastic fibre formation  pathway scores vs. Day   More Plot Information                Elastic fibre formation  pathway scores vs. Day  Plots Elastic fibre formation pathway scores against Day.   
                                           Elastic fibre formation  pathway scores vs. Line   More Plot Information                Elastic fibre formation  pathway scores vs. Line  Plots Elastic fibre formation pathway scores against Line.   
                                      
                                       
                                  
                                
                                
                                  
                                       
                                      
                                           EPH-ephrin mediated repulsion of cells  pathway scores vs. Day   More Plot Information                EPH-ephrin mediated repulsion of cells  pathway scores vs. Day  Plots EPH-ephrin mediated repulsion of cells pathway scores against Day.   
                                           EPH-ephrin mediated repulsion of cells  pathway scores vs. Line   More Plot Information                EPH-ephrin mediated repulsion of cells  pathway scores vs. Line  Plots EPH-ephrin mediated repulsion of cells pathway scores against Line.   
                                      
                                       
                                  
                                
                                
                                  
                                       
                                      
                                           EPHA-mediated growth cone collapse  pathway scores vs. Day   More Plot Information                EPHA-mediated growth cone collapse  pathway scores vs. Day  Plots EPHA-mediated growth cone collapse pathway scores against Day.   
                                           EPHA-mediated growth cone collapse  pathway scores vs. Line   More Plot Information                EPHA-mediated growth cone collapse  pathway scores vs. Line  Plots EPHA-mediated growth cone collapse pathway scores against Line.   
                                      
                                       
                                  
                                
                                
                                  
                                       
                                      
                                           EPHB-mediated forward signaling  pathway scores vs. Day   More Plot Information                EPHB-mediated forward signaling  pathway scores vs. Day  Plots EPHB-mediated forward signaling pathway scores against Day.   
                                           EPHB-mediated forward signaling  pathway scores vs. Line   More Plot Information                EPHB-mediated forward signaling  pathway scores vs. Line  Plots EPHB-mediated forward signaling pathway scores against Line.   
                                      
                                       
                                  
                                
                                
                                  
                                       
                                      
                                           Ephrin signaling  pathway scores vs. Day   More Plot Information                Ephrin signaling  pathway scores vs. Day  Plots Ephrin signaling pathway scores against Day.   
                                           Ephrin signaling  pathway scores vs. Line   More Plot Information                Ephrin signaling  pathway scores vs. Line  Plots Ephrin signaling pathway scores against Line.   
                                      
                                       
                                  
                                
                                
                                  
                                       
                                      
                                           FCERI mediated MAPK activation  pathway scores vs. Day   More Plot Information                FCERI mediated MAPK activation  pathway scores vs. Day  Plots FCERI mediated MAPK activation pathway scores against Day.   
                                           FCERI mediated MAPK activation  pathway scores vs. Line   More Plot Information                FCERI mediated MAPK activation  pathway scores vs. Line  Plots FCERI mediated MAPK activation pathway scores against Line.   
                                      
                                       
                                  
                                
                                
                                  
                                       
                                      
                                           Formation of the beta-catenin.CL.TCF transactivating complex  pathway scores vs. Day   More Plot Information                Formation of the beta-catenin.CL.TCF transactivating complex  pathway scores vs. Day  Plots Formation of the beta-catenin.CL.TCF transactivating complex pathway scores against Day.   
                                           Formation of the beta-catenin.CL.TCF transactivating complex  pathway scores vs. Line   More Plot Information                Formation of the beta-catenin.CL.TCF transactivating complex  pathway scores vs. Line  Plots Formation of the beta-catenin.CL.TCF transactivating complex pathway scores against Line.   
                                      
                                       
                                  
                                
                                
                                  
                                       
                                      
                                           Frs2-mediated activation  pathway scores vs. Day   More Plot Information                Frs2-mediated activation  pathway scores vs. Day  Plots Frs2-mediated activation pathway scores against Day.   
                                           Frs2-mediated activation  pathway scores vs. Line   More Plot Information                Frs2-mediated activation  pathway scores vs. Line  Plots Frs2-mediated activation pathway scores against Line.   
                                      
                                       
                                  
                                
                                
                                  
                                       
                                      
                                           G alpha (1213) signalling events  pathway scores vs. Day   More Plot Information                G alpha (1213) signalling events  pathway scores vs. Day  Plots G alpha (1213) signalling events pathway scores against Day.   
                                           G alpha (1213) signalling events  pathway scores vs. Line   More Plot Information                G alpha (1213) signalling events  pathway scores vs. Line  Plots G alpha (1213) signalling events pathway scores against Line.   
                                      
                                       
                                  
                                
                                
                                  
                                       
                                      
                                           G alpha (q) signalling events  pathway scores vs. Day   More Plot Information                G alpha (q) signalling events  pathway scores vs. Day  Plots G alpha (q) signalling events pathway scores against Day.   
                                           G alpha (q) signalling events  pathway scores vs. Line   More Plot Information                G alpha (q) signalling events  pathway scores vs. Line  Plots G alpha (q) signalling events pathway scores against Line.   
                                      
                                       
                                  
                                
                                
                                  
                                       
                                      
                                           G beta.CL.gamma signalling through PI3Kgamma  pathway scores vs. Day   More Plot Information                G beta.CL.gamma signalling through PI3Kgamma  pathway scores vs. Day  Plots G beta.CL.gamma signalling through PI3Kgamma pathway scores against Day.   
                                           G beta.CL.gamma signalling through PI3Kgamma  pathway scores vs. Line   More Plot Information                G beta.CL.gamma signalling through PI3Kgamma  pathway scores vs. Line  Plots G beta.CL.gamma signalling through PI3Kgamma pathway scores against Line.   
                                      
                                       
                                  
                                
                                
                                  
                                       
                                      
                                           GAB1 signalosome  pathway scores vs. Day   More Plot Information                GAB1 signalosome  pathway scores vs. Day  Plots GAB1 signalosome pathway scores against Day.   
                                           GAB1 signalosome  pathway scores vs. Line   More Plot Information                GAB1 signalosome  pathway scores vs. Line  Plots GAB1 signalosome pathway scores against Line.   
                                      
                                       
                                  
                                
                                
                                  
                                       
                                      
                                           Gastrin-CREB signalling pathway via PKC and MAPK  pathway scores vs. Day   More Plot Information                Gastrin-CREB signalling pathway via PKC and MAPK  pathway scores vs. Day  Plots Gastrin-CREB signalling pathway via PKC and MAPK pathway scores against Day.   
                                           Gastrin-CREB signalling pathway via PKC and MAPK  pathway scores vs. Line   More Plot Information                Gastrin-CREB signalling pathway via PKC and MAPK  pathway scores vs. Line  Plots Gastrin-CREB signalling pathway via PKC and MAPK pathway scores against Line.   
                                      
                                       
                                  
                                
                                
                                  
                                       
                                      
                                           GPVI-mediated activation cascade  pathway scores vs. Day   More Plot Information                GPVI-mediated activation cascade  pathway scores vs. Day  Plots GPVI-mediated activation cascade pathway scores against Day.   
                                           GPVI-mediated activation cascade  pathway scores vs. Line   More Plot Information                GPVI-mediated activation cascade  pathway scores vs. Line  Plots GPVI-mediated activation cascade pathway scores against Line.   
                                      
                                       
                                  
                                
                                
                                  
                                       
                                      
                                           Integrin alphaIIb beta3 signaling  pathway scores vs. Day   More Plot Information                Integrin alphaIIb beta3 signaling  pathway scores vs. Day  Plots Integrin alphaIIb beta3 signaling pathway scores against Day.   
                                           Integrin alphaIIb beta3 signaling  pathway scores vs. Line   More Plot Information                Integrin alphaIIb beta3 signaling  pathway scores vs. Line  Plots Integrin alphaIIb beta3 signaling pathway scores against Line.   
                                      
                                       
                                  
                                
                                
                                  
                                       
                                      
                                           Integrin cell surface interactions  pathway scores vs. Day   More Plot Information                Integrin cell surface interactions  pathway scores vs. Day  Plots Integrin cell surface interactions pathway scores against Day.   
                                           Integrin cell surface interactions  pathway scores vs. Line   More Plot Information                Integrin cell surface interactions  pathway scores vs. Line  Plots Integrin cell surface interactions pathway scores against Line.   
                                      
                                       
                                  
                                
                                
                                  
                                       
                                      
                                           Interleukin receptor SHC signaling  pathway scores vs. Day   More Plot Information                Interleukin receptor SHC signaling  pathway scores vs. Day  Plots Interleukin receptor SHC signaling pathway scores against Day.   
                                           Interleukin receptor SHC signaling  pathway scores vs. Line   More Plot Information                Interleukin receptor SHC signaling  pathway scores vs. Line  Plots Interleukin receptor SHC signaling pathway scores against Line.   
                                      
                                       
                                  
                                
                                
                                  
                                       
                                      
                                           IRS-related events triggered by IGF1R  pathway scores vs. Day   More Plot Information                IRS-related events triggered by IGF1R  pathway scores vs. Day  Plots IRS-related events triggered by IGF1R pathway scores against Day.   
                                           IRS-related events triggered by IGF1R  pathway scores vs. Line   More Plot Information                IRS-related events triggered by IGF1R  pathway scores vs. Line  Plots IRS-related events triggered by IGF1R pathway scores against Line.   
                                      
                                       
                                  
                                
                                
                                  
                                       
                                      
                                           Laminin interactions  pathway scores vs. Day   More Plot Information                Laminin interactions  pathway scores vs. Day  Plots Laminin interactions pathway scores against Day.   
                                           Laminin interactions  pathway scores vs. Line   More Plot Information                Laminin interactions  pathway scores vs. Line  Plots Laminin interactions pathway scores against Line.   
                                      
                                       
                                  
                                
                                
                                  
                                       
                                      
                                           MAP2K and MAPK activation  pathway scores vs. Day   More Plot Information                MAP2K and MAPK activation  pathway scores vs. Day  Plots MAP2K and MAPK activation pathway scores against Day.   
                                           MAP2K and MAPK activation  pathway scores vs. Line   More Plot Information                MAP2K and MAPK activation  pathway scores vs. Line  Plots MAP2K and MAPK activation pathway scores against Line.   
                                      
                                       
                                  
                                
                                
                                  
                                       
                                      
                                           MAPK6MAPK4 signaling  pathway scores vs. Day   More Plot Information                MAPK6MAPK4 signaling  pathway scores vs. Day  Plots MAPK6MAPK4 signaling pathway scores against Day.   
                                           MAPK6MAPK4 signaling  pathway scores vs. Line   More Plot Information                MAPK6MAPK4 signaling  pathway scores vs. Line  Plots MAPK6MAPK4 signaling pathway scores against Line.   
                                      
                                       
                                  
                                
                                
                                  
                                       
                                      
                                           MicroRNA (miRNA) biogenesis  pathway scores vs. Day   More Plot Information                MicroRNA (miRNA) biogenesis  pathway scores vs. Day  Plots MicroRNA (miRNA) biogenesis pathway scores against Day.   
                                           MicroRNA (miRNA) biogenesis  pathway scores vs. Line   More Plot Information                MicroRNA (miRNA) biogenesis  pathway scores vs. Line  Plots MicroRNA (miRNA) biogenesis pathway scores against Line.   
                                      
                                       
                                  
                                
                                
                                  
                                       
                                      
                                           Misspliced GSK3beta mutants stabilize beta-catenin  pathway scores vs. Day   More Plot Information                Misspliced GSK3beta mutants stabilize beta-catenin  pathway scores vs. Day  Plots Misspliced GSK3beta mutants stabilize beta-catenin pathway scores against Day.   
                                           Misspliced GSK3beta mutants stabilize beta-catenin  pathway scores vs. Line   More Plot Information                Misspliced GSK3beta mutants stabilize beta-catenin  pathway scores vs. Line  Plots Misspliced GSK3beta mutants stabilize beta-catenin pathway scores against Line.   
                                      
                                       
                                  
                                
                                
                                  
                                       
                                      
                                           Molecules associated with elastic fibres  pathway scores vs. Day   More Plot Information                Molecules associated with elastic fibres  pathway scores vs. Day  Plots Molecules associated with elastic fibres pathway scores against Day.   
                                           Molecules associated with elastic fibres  pathway scores vs. Line   More Plot Information                Molecules associated with elastic fibres  pathway scores vs. Line  Plots Molecules associated with elastic fibres pathway scores against Line.   
                                      
                                       
                                  
                                
                                
                                  
                                       
                                      
                                           Negative regulation of FGFR1 signaling  pathway scores vs. Day   More Plot Information                Negative regulation of FGFR1 signaling  pathway scores vs. Day  Plots Negative regulation of FGFR1 signaling pathway scores against Day.   
                                           Negative regulation of FGFR1 signaling  pathway scores vs. Line   More Plot Information                Negative regulation of FGFR1 signaling  pathway scores vs. Line  Plots Negative regulation of FGFR1 signaling pathway scores against Line.   
                                      
                                       
                                  
                                
                                
                                  
                                       
                                      
                                           Negative regulation of FGFR2 signaling  pathway scores vs. Day   More Plot Information                Negative regulation of FGFR2 signaling  pathway scores vs. Day  Plots Negative regulation of FGFR2 signaling pathway scores against Day.   
                                           Negative regulation of FGFR2 signaling  pathway scores vs. Line   More Plot Information                Negative regulation of FGFR2 signaling  pathway scores vs. Line  Plots Negative regulation of FGFR2 signaling pathway scores against Line.   
                                      
                                       
                                  
                                
                                
                                  
                                       
                                      
                                           Negative regulation of FGFR3 signaling  pathway scores vs. Day   More Plot Information                Negative regulation of FGFR3 signaling  pathway scores vs. Day  Plots Negative regulation of FGFR3 signaling pathway scores against Day.   
                                           Negative regulation of FGFR3 signaling  pathway scores vs. Line   More Plot Information                Negative regulation of FGFR3 signaling  pathway scores vs. Line  Plots Negative regulation of FGFR3 signaling pathway scores against Line.   
                                      
                                       
                                  
                                
                                
                                  
                                       
                                      
                                           Negative regulation of FGFR4 signaling  pathway scores vs. Day   More Plot Information                Negative regulation of FGFR4 signaling  pathway scores vs. Day  Plots Negative regulation of FGFR4 signaling pathway scores against Day.   
                                           Negative regulation of FGFR4 signaling  pathway scores vs. Line   More Plot Information                Negative regulation of FGFR4 signaling  pathway scores vs. Line  Plots Negative regulation of FGFR4 signaling pathway scores against Line.   
                                      
                                       
                                  
                                
                                
                                  
                                       
                                      
                                           Negative regulation of MAPK pathway  pathway scores vs. Day   More Plot Information                Negative regulation of MAPK pathway  pathway scores vs. Day  Plots Negative regulation of MAPK pathway pathway scores against Day.   
                                           Negative regulation of MAPK pathway  pathway scores vs. Line   More Plot Information                Negative regulation of MAPK pathway  pathway scores vs. Line  Plots Negative regulation of MAPK pathway pathway scores against Line.   
                                      
                                       
                                  
                                
                                
                                  
                                       
                                      
                                           Nephrin interactions  pathway scores vs. Day   More Plot Information                Nephrin interactions  pathway scores vs. Day  Plots Nephrin interactions pathway scores against Day.   
                                           Nephrin interactions  pathway scores vs. Line   More Plot Information                Nephrin interactions  pathway scores vs. Line  Plots Nephrin interactions pathway scores against Line.   
                                      
                                       
                                  
                                
                                
                                  
                                       
                                      
                                           Non-integrin membrane-ECM interactions  pathway scores vs. Day   More Plot Information                Non-integrin membrane-ECM interactions  pathway scores vs. Day  Plots Non-integrin membrane-ECM interactions pathway scores against Day.   
                                           Non-integrin membrane-ECM interactions  pathway scores vs. Line   More Plot Information                Non-integrin membrane-ECM interactions  pathway scores vs. Line  Plots Non-integrin membrane-ECM interactions pathway scores against Line.   
                                      
                                       
                                  
                                
                                
                                  
                                       
                                      
                                           NRAGE signals death through JNK  pathway scores vs. Day   More Plot Information                NRAGE signals death through JNK  pathway scores vs. Day  Plots NRAGE signals death through JNK pathway scores against Day.   
                                           NRAGE signals death through JNK  pathway scores vs. Line   More Plot Information                NRAGE signals death through JNK  pathway scores vs. Line  Plots NRAGE signals death through JNK pathway scores against Line.   
                                      
                                       
                                  
                                
                                
                                  
                                       
                                      
                                           PCPCE pathway  pathway scores vs. Day   More Plot Information                PCPCE pathway  pathway scores vs. Day  Plots PCPCE pathway pathway scores against Day.   
                                           PCPCE pathway  pathway scores vs. Line   More Plot Information                PCPCE pathway  pathway scores vs. Line  Plots PCPCE pathway pathway scores against Line.   
                                      
                                       
                                  
                                
                                
                                  
                                       
                                      
                                           PI3K Cascade  pathway scores vs. Day   More Plot Information                PI3K Cascade  pathway scores vs. Day  Plots PI3K Cascade pathway scores against Day.   
                                           PI3K Cascade  pathway scores vs. Line   More Plot Information                PI3K Cascade  pathway scores vs. Line  Plots PI3K Cascade pathway scores against Line.   
                                      
                                       
                                  
                                
                                
                                  
                                       
                                      
                                           PI3KAKT activation  pathway scores vs. Day   More Plot Information                PI3KAKT activation  pathway scores vs. Day  Plots PI3KAKT activation pathway scores against Day.   
                                           PI3KAKT activation  pathway scores vs. Line   More Plot Information                PI3KAKT activation  pathway scores vs. Line  Plots PI3KAKT activation pathway scores against Line.   
                                      
                                       
                                  
                                
                                
                                  
                                       
                                      
                                           PIP3 activates AKT signaling  pathway scores vs. Day   More Plot Information                PIP3 activates AKT signaling  pathway scores vs. Day  Plots PIP3 activates AKT signaling pathway scores against Day.   
                                           PIP3 activates AKT signaling  pathway scores vs. Line   More Plot Information                PIP3 activates AKT signaling  pathway scores vs. Line  Plots PIP3 activates AKT signaling pathway scores against Line.   
                                      
                                       
                                  
                                
                                
                                  
                                       
                                      
                                           Platelet degranulation   pathway scores vs. Day   More Plot Information                Platelet degranulation   pathway scores vs. Day  Plots Platelet degranulation  pathway scores against Day.   
                                           Platelet degranulation   pathway scores vs. Line   More Plot Information                Platelet degranulation   pathway scores vs. Line  Plots Platelet degranulation  pathway scores against Line.   
                                      
                                       
                                  
                                
                                
                                  
                                       
                                      
                                           RAF activation  pathway scores vs. Day   More Plot Information                RAF activation  pathway scores vs. Day  Plots RAF activation pathway scores against Day.   
                                           RAF activation  pathway scores vs. Line   More Plot Information                RAF activation  pathway scores vs. Line  Plots RAF activation pathway scores against Line.   
                                      
                                       
                                  
                                
                                
                                  
                                       
                                      
                                           RAFMAP kinase cascade  pathway scores vs. Day   More Plot Information                RAFMAP kinase cascade  pathway scores vs. Day  Plots RAFMAP kinase cascade pathway scores against Day.   
                                           RAFMAP kinase cascade  pathway scores vs. Line   More Plot Information                RAFMAP kinase cascade  pathway scores vs. Line  Plots RAFMAP kinase cascade pathway scores against Line.   
                                      
                                       
                                  
                                
                                
                                  
                                       
                                      
                                           Regulation of actin dynamics for phagocytic cup formation  pathway scores vs. Day   More Plot Information                Regulation of actin dynamics for phagocytic cup formation  pathway scores vs. Day  Plots Regulation of actin dynamics for phagocytic cup formation pathway scores against Day.   
                                           Regulation of actin dynamics for phagocytic cup formation  pathway scores vs. Line   More Plot Information                Regulation of actin dynamics for phagocytic cup formation  pathway scores vs. Line  Plots Regulation of actin dynamics for phagocytic cup formation pathway scores against Line.   
                                      
                                       
                                  
                                
                                
                                  
                                       
                                      
                                           Regulation of signaling by CBL  pathway scores vs. Day   More Plot Information                Regulation of signaling by CBL  pathway scores vs. Day  Plots Regulation of signaling by CBL pathway scores against Day.   
                                           Regulation of signaling by CBL  pathway scores vs. Line   More Plot Information                Regulation of signaling by CBL  pathway scores vs. Line  Plots Regulation of signaling by CBL pathway scores against Line.   
                                      
                                       
                                  
                                
                                
                                  
                                       
                                      
                                           Repression of WNT target genes  pathway scores vs. Day   More Plot Information                Repression of WNT target genes  pathway scores vs. Day  Plots Repression of WNT target genes pathway scores against Day.   
                                           Repression of WNT target genes  pathway scores vs. Line   More Plot Information                Repression of WNT target genes  pathway scores vs. Line  Plots Repression of WNT target genes pathway scores against Line.   
                                      
                                       
                                  
                                
                                
                                  
                                       
                                      
                                           Rho GTPase cycle  pathway scores vs. Day   More Plot Information                Rho GTPase cycle  pathway scores vs. Day  Plots Rho GTPase cycle pathway scores against Day.   
                                           Rho GTPase cycle  pathway scores vs. Line   More Plot Information                Rho GTPase cycle  pathway scores vs. Line  Plots Rho GTPase cycle pathway scores against Line.   
                                      
                                       
                                  
                                
                                
                                  
                                       
                                      
                                           RHO GTPases activate CIT  pathway scores vs. Day   More Plot Information                RHO GTPases activate CIT  pathway scores vs. Day  Plots RHO GTPases activate CIT pathway scores against Day.   
                                           RHO GTPases activate CIT  pathway scores vs. Line   More Plot Information                RHO GTPases activate CIT  pathway scores vs. Line  Plots RHO GTPases activate CIT pathway scores against Line.   
                                      
                                       
                                  
                                
                                
                                  
                                       
                                      
                                           RHO GTPases Activate Formins  pathway scores vs. Day   More Plot Information                RHO GTPases Activate Formins  pathway scores vs. Day  Plots RHO GTPases Activate Formins pathway scores against Day.   
                                           RHO GTPases Activate Formins  pathway scores vs. Line   More Plot Information                RHO GTPases Activate Formins  pathway scores vs. Line  Plots RHO GTPases Activate Formins pathway scores against Line.   
                                      
                                       
                                  
                                
                                
                                  
                                       
                                      
                                           RHO GTPases activate IQGAPs  pathway scores vs. Day   More Plot Information                RHO GTPases activate IQGAPs  pathway scores vs. Day  Plots RHO GTPases activate IQGAPs pathway scores against Day.   
                                           RHO GTPases activate IQGAPs  pathway scores vs. Line   More Plot Information                RHO GTPases activate IQGAPs  pathway scores vs. Line  Plots RHO GTPases activate IQGAPs pathway scores against Line.   
                                      
                                       
                                  
                                
                                
                                  
                                       
                                      
                                           RHO GTPases activate PAKs  pathway scores vs. Day   More Plot Information                RHO GTPases activate PAKs  pathway scores vs. Day  Plots RHO GTPases activate PAKs pathway scores against Day.   
                                           RHO GTPases activate PAKs  pathway scores vs. Line   More Plot Information                RHO GTPases activate PAKs  pathway scores vs. Line  Plots RHO GTPases activate PAKs pathway scores against Line.   
                                      
                                       
                                  
                                
                                
                                  
                                       
                                      
                                           RHO GTPases activate PKNs  pathway scores vs. Day   More Plot Information                RHO GTPases activate PKNs  pathway scores vs. Day  Plots RHO GTPases activate PKNs pathway scores against Day.   
                                           RHO GTPases activate PKNs  pathway scores vs. Line   More Plot Information                RHO GTPases activate PKNs  pathway scores vs. Line  Plots RHO GTPases activate PKNs pathway scores against Line.   
                                      
                                       
                                  
                                
                                
                                  
                                       
                                      
                                           RHO GTPases Activate ROCKs  pathway scores vs. Day   More Plot Information                RHO GTPases Activate ROCKs  pathway scores vs. Day  Plots RHO GTPases Activate ROCKs pathway scores against Day.   
                                           RHO GTPases Activate ROCKs  pathway scores vs. Line   More Plot Information                RHO GTPases Activate ROCKs  pathway scores vs. Line  Plots RHO GTPases Activate ROCKs pathway scores against Line.   
                                      
                                       
                                  
                                
                                
                                  
                                       
                                      
                                           RHO GTPases Activate WASPs and WAVEs  pathway scores vs. Day   More Plot Information                RHO GTPases Activate WASPs and WAVEs  pathway scores vs. Day  Plots RHO GTPases Activate WASPs and WAVEs pathway scores against Day.   
                                           RHO GTPases Activate WASPs and WAVEs  pathway scores vs. Line   More Plot Information                RHO GTPases Activate WASPs and WAVEs  pathway scores vs. Line  Plots RHO GTPases Activate WASPs and WAVEs pathway scores against Line.   
                                      
                                       
                                  
                                
                                
                                  
                                       
                                      
                                           Role of LAT2NTALLAB on calcium mobilization  pathway scores vs. Day   More Plot Information                Role of LAT2NTALLAB on calcium mobilization  pathway scores vs. Day  Plots Role of LAT2NTALLAB on calcium mobilization pathway scores against Day.   
                                           Role of LAT2NTALLAB on calcium mobilization  pathway scores vs. Line   More Plot Information                Role of LAT2NTALLAB on calcium mobilization  pathway scores vs. Line  Plots Role of LAT2NTALLAB on calcium mobilization pathway scores against Line.   
                                      
                                       
                                  
                                
                                
                                  
                                       
                                      
                                           S33 mutants of beta-catenin aren.SQT.t phosphorylated  pathway scores vs. Day   More Plot Information                S33 mutants of beta-catenin aren.SQT.t phosphorylated  pathway scores vs. Day  Plots S33 mutants of beta-catenin aren.SQT.t phosphorylated pathway scores against Day.   
                                           S33 mutants of beta-catenin aren.SQT.t phosphorylated  pathway scores vs. Line   More Plot Information                S33 mutants of beta-catenin aren.SQT.t phosphorylated  pathway scores vs. Line  Plots S33 mutants of beta-catenin aren.SQT.t phosphorylated pathway scores against Line.   
                                      
                                       
                                  
                                
                                
                                  
                                       
                                      
                                           S37 mutants of beta-catenin aren.SQT.t phosphorylated  pathway scores vs. Day   More Plot Information                S37 mutants of beta-catenin aren.SQT.t phosphorylated  pathway scores vs. Day  Plots S37 mutants of beta-catenin aren.SQT.t phosphorylated pathway scores against Day.   
                                           S37 mutants of beta-catenin aren.SQT.t phosphorylated  pathway scores vs. Line   More Plot Information                S37 mutants of beta-catenin aren.SQT.t phosphorylated  pathway scores vs. Line  Plots S37 mutants of beta-catenin aren.SQT.t phosphorylated pathway scores against Line.   
                                      
                                       
                                  
                                
                                
                                  
                                       
                                      
                                           S45 mutants of beta-catenin aren.SQT.t phosphorylated  pathway scores vs. Day   More Plot Information                S45 mutants of beta-catenin aren.SQT.t phosphorylated  pathway scores vs. Day  Plots S45 mutants of beta-catenin aren.SQT.t phosphorylated pathway scores against Day.   
                                           S45 mutants of beta-catenin aren.SQT.t phosphorylated  pathway scores vs. Line   More Plot Information                S45 mutants of beta-catenin aren.SQT.t phosphorylated  pathway scores vs. Line  Plots S45 mutants of beta-catenin aren.SQT.t phosphorylated pathway scores against Line.   
                                      
                                       
                                  
                                
                                
                                  
                                       
                                      
                                           Sema3A PAK dependent Axon repulsion  pathway scores vs. Day   More Plot Information                Sema3A PAK dependent Axon repulsion  pathway scores vs. Day  Plots Sema3A PAK dependent Axon repulsion pathway scores against Day.   
                                           Sema3A PAK dependent Axon repulsion  pathway scores vs. Line   More Plot Information                Sema3A PAK dependent Axon repulsion  pathway scores vs. Line  Plots Sema3A PAK dependent Axon repulsion pathway scores against Line.   
                                      
                                       
                                  
                                
                                
                                  
                                       
                                      
                                           Sema4D induced cell migration and growth-cone collapse  pathway scores vs. Day   More Plot Information                Sema4D induced cell migration and growth-cone collapse  pathway scores vs. Day  Plots Sema4D induced cell migration and growth-cone collapse pathway scores against Day.   
                                           Sema4D induced cell migration and growth-cone collapse  pathway scores vs. Line   More Plot Information                Sema4D induced cell migration and growth-cone collapse  pathway scores vs. Line  Plots Sema4D induced cell migration and growth-cone collapse pathway scores against Line.   
                                      
                                       
                                  
                                
                                
                                  
                                       
                                      
                                           SHC-mediated cascade.CL.FGFR1  pathway scores vs. Day   More Plot Information                SHC-mediated cascade.CL.FGFR1  pathway scores vs. Day  Plots SHC-mediated cascade.CL.FGFR1 pathway scores against Day.   
                                           SHC-mediated cascade.CL.FGFR1  pathway scores vs. Line   More Plot Information                SHC-mediated cascade.CL.FGFR1  pathway scores vs. Line  Plots SHC-mediated cascade.CL.FGFR1 pathway scores against Line.   
                                      
                                       
                                  
                                
                                
                                  
                                       
                                      
                                           SHC-related events triggered by IGF1R  pathway scores vs. Day   More Plot Information                SHC-related events triggered by IGF1R  pathway scores vs. Day  Plots SHC-related events triggered by IGF1R pathway scores against Day.   
                                           SHC-related events triggered by IGF1R  pathway scores vs. Line   More Plot Information                SHC-related events triggered by IGF1R  pathway scores vs. Line  Plots SHC-related events triggered by IGF1R pathway scores against Line.   
                                      
                                       
                                  
                                
                                
                                  
                                       
                                      
                                           Signal transduction by L1  pathway scores vs. Day   More Plot Information                Signal transduction by L1  pathway scores vs. Day  Plots Signal transduction by L1 pathway scores against Day.   
                                           Signal transduction by L1  pathway scores vs. Line   More Plot Information                Signal transduction by L1  pathway scores vs. Line  Plots Signal transduction by L1 pathway scores against Line.   
                                      
                                       
                                  
                                
                                
                                  
                                       
                                      
                                           Signaling by ERBB2  pathway scores vs. Day   More Plot Information                Signaling by ERBB2  pathway scores vs. Day  Plots Signaling by ERBB2 pathway scores against Day.   
                                           Signaling by ERBB2  pathway scores vs. Line   More Plot Information                Signaling by ERBB2  pathway scores vs. Line  Plots Signaling by ERBB2 pathway scores against Line.   
                                      
                                       
                                  
                                
                                
                                  
                                       
                                      
                                           Signaling by FGFR1 in disease  pathway scores vs. Day   More Plot Information                Signaling by FGFR1 in disease  pathway scores vs. Day  Plots Signaling by FGFR1 in disease pathway scores against Day.   
                                           Signaling by FGFR1 in disease  pathway scores vs. Line   More Plot Information                Signaling by FGFR1 in disease  pathway scores vs. Line  Plots Signaling by FGFR1 in disease pathway scores against Line.   
                                      
                                       
                                  
                                
                                
                                  
                                       
                                      
                                           Signaling by FGFR2 in disease  pathway scores vs. Day   More Plot Information                Signaling by FGFR2 in disease  pathway scores vs. Day  Plots Signaling by FGFR2 in disease pathway scores against Day.   
                                           Signaling by FGFR2 in disease  pathway scores vs. Line   More Plot Information                Signaling by FGFR2 in disease  pathway scores vs. Line  Plots Signaling by FGFR2 in disease pathway scores against Line.   
                                      
                                       
                                  
                                
                                
                                  
                                       
                                      
                                           Signaling by FGFR3 in disease  pathway scores vs. Day   More Plot Information                Signaling by FGFR3 in disease  pathway scores vs. Day  Plots Signaling by FGFR3 in disease pathway scores against Day.   
                                           Signaling by FGFR3 in disease  pathway scores vs. Line   More Plot Information                Signaling by FGFR3 in disease  pathway scores vs. Line  Plots Signaling by FGFR3 in disease pathway scores against Line.   
                                      
                                       
                                  
                                
                                
                                  
                                       
                                      
                                           Signaling by FGFR4 in disease  pathway scores vs. Day   More Plot Information                Signaling by FGFR4 in disease  pathway scores vs. Day  Plots Signaling by FGFR4 in disease pathway scores against Day.   
                                           Signaling by FGFR4 in disease  pathway scores vs. Line   More Plot Information                Signaling by FGFR4 in disease  pathway scores vs. Line  Plots Signaling by FGFR4 in disease pathway scores against Line.   
                                      
                                       
                                  
                                
                                
                                  
                                       
                                      
                                           Signaling by PDGF  pathway scores vs. Day   More Plot Information                Signaling by PDGF  pathway scores vs. Day  Plots Signaling by PDGF pathway scores against Day.   
                                           Signaling by PDGF  pathway scores vs. Line   More Plot Information                Signaling by PDGF  pathway scores vs. Line  Plots Signaling by PDGF pathway scores against Line.   
                                      
                                       
                                  
                                
                                
                                  
                                       
                                      
                                           Signaling by SCF-KIT  pathway scores vs. Day   More Plot Information                Signaling by SCF-KIT  pathway scores vs. Day  Plots Signaling by SCF-KIT pathway scores against Day.   
                                           Signaling by SCF-KIT  pathway scores vs. Line   More Plot Information                Signaling by SCF-KIT  pathway scores vs. Line  Plots Signaling by SCF-KIT pathway scores against Line.   
                                      
                                       
                                  
                                
                                
                                  
                                       
                                      
                                           Smooth Muscle Contraction  pathway scores vs. Day   More Plot Information                Smooth Muscle Contraction  pathway scores vs. Day  Plots Smooth Muscle Contraction pathway scores against Day.   
                                           Smooth Muscle Contraction  pathway scores vs. Line   More Plot Information                Smooth Muscle Contraction  pathway scores vs. Line  Plots Smooth Muscle Contraction pathway scores against Line.   
                                      
                                       
                                  
                                
                                
                                  
                                       
                                      
                                           Spry regulation of FGF signaling  pathway scores vs. Day   More Plot Information                Spry regulation of FGF signaling  pathway scores vs. Day  Plots Spry regulation of FGF signaling pathway scores against Day.   
                                           Spry regulation of FGF signaling  pathway scores vs. Line   More Plot Information                Spry regulation of FGF signaling  pathway scores vs. Line  Plots Spry regulation of FGF signaling pathway scores against Line.   
                                      
                                       
                                  
                                
                                
                                  
                                       
                                      
                                           Syndecan interactions  pathway scores vs. Day   More Plot Information                Syndecan interactions  pathway scores vs. Day  Plots Syndecan interactions pathway scores against Day.   
                                           Syndecan interactions  pathway scores vs. Line   More Plot Information                Syndecan interactions  pathway scores vs. Line  Plots Syndecan interactions pathway scores against Line.   
                                      
                                       
                                  
                                
                                
                                  
                                       
                                      
                                           Synthesis of PIPs at the plasma membrane  pathway scores vs. Day   More Plot Information                Synthesis of PIPs at the plasma membrane  pathway scores vs. Day  Plots Synthesis of PIPs at the plasma membrane pathway scores against Day.   
                                           Synthesis of PIPs at the plasma membrane  pathway scores vs. Line   More Plot Information                Synthesis of PIPs at the plasma membrane  pathway scores vs. Line  Plots Synthesis of PIPs at the plasma membrane pathway scores against Line.   
                                      
                                       
                                  
                                
                                
                                  
                                       
                                      
                                           T41 mutants of beta-catenin aren.SQT.t phosphorylated  pathway scores vs. Day   More Plot Information                T41 mutants of beta-catenin aren.SQT.t phosphorylated  pathway scores vs. Day  Plots T41 mutants of beta-catenin aren.SQT.t phosphorylated pathway scores against Day.   
                                           T41 mutants of beta-catenin aren.SQT.t phosphorylated  pathway scores vs. Line   More Plot Information                T41 mutants of beta-catenin aren.SQT.t phosphorylated  pathway scores vs. Line  Plots T41 mutants of beta-catenin aren.SQT.t phosphorylated pathway scores against Line.   
                                      
                                       
                                  
                                
                                
                                  
                                       
                                      
                                           TGF-beta receptor signaling activates SMADs  pathway scores vs. Day   More Plot Information                TGF-beta receptor signaling activates SMADs  pathway scores vs. Day  Plots TGF-beta receptor signaling activates SMADs pathway scores against Day.   
                                           TGF-beta receptor signaling activates SMADs  pathway scores vs. Line   More Plot Information                TGF-beta receptor signaling activates SMADs  pathway scores vs. Line  Plots TGF-beta receptor signaling activates SMADs pathway scores against Line.   
                                      
                                       
                                  
                                
                                
                                  
                                       
                                      
                                           Tie2 Signaling  pathway scores vs. Day   More Plot Information                Tie2 Signaling  pathway scores vs. Day  Plots Tie2 Signaling pathway scores against Day.   
                                           Tie2 Signaling  pathway scores vs. Line   More Plot Information                Tie2 Signaling  pathway scores vs. Line  Plots Tie2 Signaling pathway scores against Line.   
                                      
                                       
                                  
                                
                                
                                  
                                       
                                      
                                           Tight junction interactions  pathway scores vs. Day   More Plot Information                Tight junction interactions  pathway scores vs. Day  Plots Tight junction interactions pathway scores against Day.   
                                           Tight junction interactions  pathway scores vs. Line   More Plot Information                Tight junction interactions  pathway scores vs. Line  Plots Tight junction interactions pathway scores against Line.   
                                      
                                       
                                  
                                
                                
                                  
                                       
                                      
                                           TP53 Regulates Metabolic Genes  pathway scores vs. Day   More Plot Information                TP53 Regulates Metabolic Genes  pathway scores vs. Day  Plots TP53 Regulates Metabolic Genes pathway scores against Day.   
                                           TP53 Regulates Metabolic Genes  pathway scores vs. Line   More Plot Information                TP53 Regulates Metabolic Genes  pathway scores vs. Line  Plots TP53 Regulates Metabolic Genes pathway scores against Line.   
                                      
                                       
                                  
                                
                                
                                  
                                       
                                      
                                           Truncations of AMER1 destabilize the destruction complex  pathway scores vs. Day   More Plot Information                Truncations of AMER1 destabilize the destruction complex  pathway scores vs. Day  Plots Truncations of AMER1 destabilize the destruction complex pathway scores against Day.   
                                           Truncations of AMER1 destabilize the destruction complex  pathway scores vs. Line   More Plot Information                Truncations of AMER1 destabilize the destruction complex  pathway scores vs. Line  Plots Truncations of AMER1 destabilize the destruction complex pathway scores against Line.   
                                      
                                       
                                  
                                
                                
                                  
                                       
                                      
                                           VEGFA-VEGFR2 Pathway  pathway scores vs. Day   More Plot Information                VEGFA-VEGFR2 Pathway  pathway scores vs. Day  Plots VEGFA-VEGFR2 Pathway pathway scores against Day.   
                                           VEGFA-VEGFR2 Pathway  pathway scores vs. Line   More Plot Information                VEGFA-VEGFR2 Pathway  pathway scores vs. Line  Plots VEGFA-VEGFR2 Pathway pathway scores against Line.   
                                      
                                       
                                  
                                
                                
                                  
                                       
                                      
                                           VEGFR2 mediated cell proliferation  pathway scores vs. Day   More Plot Information                VEGFR2 mediated cell proliferation  pathway scores vs. Day  Plots VEGFR2 mediated cell proliferation pathway scores against Day.   
                                           VEGFR2 mediated cell proliferation  pathway scores vs. Line   More Plot Information                VEGFR2 mediated cell proliferation  pathway scores vs. Line  Plots VEGFR2 mediated cell proliferation pathway scores against Line.   
                                      
                                       
                                  
                                
                                
                                  
                                       
                                      
                                           VEGFR2 mediated vascular permeability  pathway scores vs. Day   More Plot Information                VEGFR2 mediated vascular permeability  pathway scores vs. Day  Plots VEGFR2 mediated vascular permeability pathway scores against Day.   
                                           VEGFR2 mediated vascular permeability  pathway scores vs. Line   More Plot Information                VEGFR2 mediated vascular permeability  pathway scores vs. Line  Plots VEGFR2 mediated vascular permeability pathway scores against Line.   
                                      
                                       
                                  
                                
                                
                                  
                                       
                                      
                                           WNT mediated activation of DVL  pathway scores vs. Day   More Plot Information                WNT mediated activation of DVL  pathway scores vs. Day  Plots WNT mediated activation of DVL pathway scores against Day.   
                                           WNT mediated activation of DVL  pathway scores vs. Line   More Plot Information                WNT mediated activation of DVL  pathway scores vs. Line  Plots WNT mediated activation of DVL pathway scores against Line.   
                                      
                                       
                                  
                                
                              
                             
                          
                           
                      
                    
                  
                
               
                
              
                
          
        
        
          
                Genes are tested for differential expression in response to each selected covariate. For each gene, a single linear 
  regression is fit using all selected covariates to predict expression. This approach eliminates confounding due to measured
  covariates and isolates the independent association of each covariate with gene expression, measuring each variable's
  association with a gene after holding all other variables constant.      
              
                   Day    Line     
                  
                      Day  
                  
                   
                  
                      Line: differential expression in Caco2 vs. baseline of Fibro  
                      Line: differential expression in HEPG2 vs. baseline of Fibro  
                      Line: differential expression in HT29 vs. baseline of Fibro  
                  
                  
                
                  
                    
                      
                           
                          
                               Volcano Plot: Day   More Plot Information                Volcano Plot: Day  Volcano plot displaying each gene's -log10(p-value) and log2 fold change with the selected covariate. Highly statistically significant genes fall at the top of the plot above the horizontal lines, and highly differentially expressed genes fall to either side. Horizontal lines indicate various False Discovery Rate (FDR) thresholds or p-value thresholds if there is no adjustment to the p-values. Genes are colored if the resulting p-value is below the given FDR or p-value threshold. The 40 most statistically significant genes are labeled in the plot.   
                               DE Results: Day   More Plot Information      Download CSV Data     
                               
                                 
                                 
                               
                               
                                 
                                   
                                   Log2 fold change 
                                   std error (log2) 
                                   Lower confidence limit (log2) 
                                   Upper confidence limit (log2) 
                                   Linear fold change 
                                   Lower confidence limit (linear) 
                                   Upper confidence limit (linear) 
                                   P-value 
                                   BY.p.value 
                                   method 
                                   Gene.sets 
                                   probe.ID 
                                 
                               
                               
                                 
                                   GUSB-mRNA 
                                   0.171 
                                   0.0188 
                                   0.134 
                                   0.208 
                                   1.13 
                                   1.1 
                                   1.15 
                                   6.09e-11 
                                   8.65e-08 
                                   lm.nb 
                                   
                                   NM_000181.1:1350 
                                 
                                 
                                   CCNG2-mRNA 
                                   0.281 
                                   0.0366 
                                   0.209 
                                   0.352 
                                   1.21 
                                   1.16 
                                   1.28 
                                   3.62e-09 
                                   2.57e-06 
                                   lm.nb 
                                   
                                   NM_004354.2:336 
                                 
                                 
                                   LDHA-mRNA 
                                   0.258 
                                   0.0354 
                                   0.189 
                                   0.327 
                                   1.2 
                                   1.14 
                                   1.25 
                                   1.16e-08 
                                   5.48e-06 
                                   lm.nb 
                                   
                                   NM_001165414.1:1690 
                                 
                                 
                                   F11R-mRNA 
                                   0.0662 
                                   0.0096 
                                   0.0473 
                                   0.085 
                                   1.05 
                                   1.03 
                                   1.06 
                                   3.95e-08 
                                   1.4e-05 
                                   loglinear 
                                   Cell surface interactions at the vascular wall, Integrin cell surface interactions, TGF-beta receptor signaling in EMT (epithelial to mesenchymal transition), Tight junction interactions 
                                   NM_144503.1:3650 
                                 
                                 
                                   EXOC4-mRNA 
                                   0.0851 
                                   0.0127 
                                   0.0603 
                                   0.11 
                                   1.06 
                                   1.04 
                                   1.08 
                                   6.67e-08 
                                   1.89e-05 
                                   lm.nb 
                                   
                                   NM_001037126.1:560 
                                 
                                 
                                   CLDN3-mRNA 
                                   0.173 
                                   0.0264 
                                   0.121 
                                   0.225 
                                   1.13 
                                   1.09 
                                   1.17 
                                   1.18e-07 
                                   2.79e-05 
                                   lm.nb 
                                   Tight junction interactions 
                                   NM_001306.3:606 
                                 
                                 
                                   ADAM9-mRNA 
                                   -0.0767 
                                   0.0118 
                                   -0.0999 
                                   -0.0535 
                                   0.948 
                                   0.933 
                                   0.964 
                                   1.39e-07 
                                   2.83e-05 
                                   loglinear 
                                   
                                   NM_001005845.1:695 
                                 
                                 
                                   MAPK1-mRNA 
                                   0.0599 
                                   0.00943 
                                   0.0414 
                                   0.0784 
                                   1.04 
                                   1.03 
                                   1.06 
                                   2.11e-07 
                                   3.74e-05 
                                   loglinear 
                                   ARMS-mediated activation, CREB phosphorylation through the activation of Ras, DAP12 signaling, Downstream signal transduction, FCERI mediated MAPK activation, Frs2-mediated activation, Gastrin-CREB signalling pathway via PKC and MAPK, Interleukin receptor SHC signaling, IRS-related events triggered by IGF1R, MAP2K and MAPK activation, Negative feedback regulation of MAPK pathway, Negative regulation of FGFR1 signaling, Negative regulation of FGFR2 signaling, Negative regulation of FGFR3 signaling, Negative regulation of FGFR4 signaling, Negative regulation of MAPK pathway, RAFMAP kinase cascade, Regulation of HSF1-mediated heat shock response, RHO GTPases Activate WASPs and WAVEs, Signal transduction by L1, Signaling by ERBB2, Signaling by PDGF, Signaling by SCF-KIT, VEGFA-VEGFR2 Pathway, VEGFR2 mediated cell proliferation 
                                   NM_138957.2:430 
                                 
                                 
                                   PDCD4-mRNA 
                                   0.239 
                                   0.0385 
                                   0.164 
                                   0.315 
                                   1.18 
                                   1.12 
                                   1.24 
                                   3.19e-07 
                                   5.02e-05 
                                   lm.nb 
                                   
                                   NM_014456.3:1115 
                                 
                                 
                                   WASF2-mRNA 
                                   0.0691 
                                   0.0112 
                                   0.0471 
                                   0.0911 
                                   1.05 
                                   1.03 
                                   1.07 
                                   3.81e-07 
                                   5.4e-05 
                                   loglinear 
                                   Regulation of actin dynamics for phagocytic cup formation, RHO GTPases Activate WASPs and WAVEs, VEGFA-VEGFR2 Pathway 
                                   NM_006990.2:485 
                                 
                                 
                                   PAK1-mRNA 
                                   0.128 
                                   0.0211 
                                   0.0869 
                                   0.169 
                                   1.09 
                                   1.06 
                                   1.12 
                                   4.84e-07 
                                   6.25e-05 
                                   lm.nb 
                                   Activation of Rac, CD209 (DC-SIGN) signaling, CD28 co-stimulation, CD28 dependent Vav1 pathway, EPHB-mediated forward signaling, Ephrin signaling, FCERI mediated MAPK activation, MAPK6MAPK4 signaling, Regulation of actin dynamics for phagocytic cup formation, RHO GTPases activate PAKs, Sema3A PAK dependent Axon repulsion, VEGFA-VEGFR2 Pathway, VEGFR2 mediated vascular permeability 
                                   NM_002576.3:975 
                                 
                                 
                                   TJP1-mRNA 
                                   -0.068 
                                   0.0114 
                                   -0.0903 
                                   -0.0458 
                                   0.954 
                                   0.939 
                                   0.969 
                                   6.41e-07 
                                   7.39e-05 
                                   lm.nb 
                                   Apoptotic cleavage of cellular proteins 
                                   NM_003257.3:6277 
                                 
                                 
                                   PPP1CB-mRNA 
                                   0.0489 
                                   0.00818 
                                   0.0328 
                                   0.0649 
                                   1.03 
                                   1.02 
                                   1.05 
                                   6.77e-07 
                                   7.39e-05 
                                   loglinear 
                                   Circadian Clock, RHO GTPases activate PKNs, RHO GTPases Activate ROCKs, TGF-beta receptor signaling activates SMADs 
                                   NM_002709.2:393 
                                 
                                 
                                   TBP-mRNA 
                                   0.0453 
                                   0.00766 
                                   0.0302 
                                   0.0603 
                                   1.03 
                                   1.02 
                                   1.04 
                                   8.4e-07 
                                   8.52e-05 
                                   loglinear 
                                   
                                   NM_001172085.1:587 
                                 
                                 
                                   GAPDH-mRNA 
                                   0.11 
                                   0.0195 
                                   0.0722 
                                   0.149 
                                   1.08 
                                   1.05 
                                   1.11 
                                   1.82e-06 
                                   0.000172 
                                   lm.nb 
                                   
                                   NM_002046.3:972 
                                 
                                 
                                   ERBB2-mRNA 
                                   0.185 
                                   0.0329 
                                   0.12 
                                   0.249 
                                   1.14 
                                   1.09 
                                   1.19 
                                   2.12e-06 
                                   0.000181 
                                   lm.nb 
                                   ARMS-mediated activation, DAP12 signaling, Downstream signal transduction, FCERI mediated MAPK activation, Frs2-mediated activation, GAB1 signalosome, Gastrin-CREB signalling pathway via PKC and MAPK, Interleukin receptor SHC signaling, IRS-related events triggered by IGF1R, PI3KAKT activation, RAFMAP kinase cascade, Role of LAT2NTALLAB on calcium mobilization, Sema4D induced cell migration and growth-cone collapse, Signaling by ERBB2, Signaling by PDGF, Signaling by SCF-KIT, VEGFA-VEGFR2 Pathway, VEGFR2 mediated cell proliferation 
                                   NM_001005862.1:1255 
                                 
                                 
                                   TP53-mRNA 
                                   0.113 
                                   0.0202 
                                   0.0733 
                                   0.152 
                                   1.08 
                                   1.05 
                                   1.11 
                                   2.25e-06 
                                   0.000181 
                                   lm.nb 
                                   
                                   NM_000546.2:1330 
                                 
                                 
                                   PTK2-mRNA 
                                   0.0792 
                                   0.0142 
                                   0.0514 
                                   0.107 
                                   1.06 
                                   1.04 
                                   1.08 
                                   2.29e-06 
                                   0.000181 
                                   lm.nb 
                                   Apoptotic cleavage of cellular proteins, ARMS-mediated activation, DAP12 signaling, DCC mediated attractive signaling, Downstream signal transduction, EPHA-mediated growth cone collapse, EPHB-mediated forward signaling, FCERI mediated MAPK activation, Frs2-mediated activation, Gastrin-CREB signalling pathway via PKC and MAPK, Integrin alphaIIb beta3 signaling, Interleukin receptor SHC signaling, IRS-related events triggered by IGF1R, p130Cas linkage to MAPK signaling for integrins, RAFMAP kinase cascade, Regulation of actin dynamics for phagocytic cup formation, RHO GTPases Activate WASPs and WAVEs, Signaling by ERBB2, Signaling by PDGF, Signaling by SCF-KIT, VEGFA-VEGFR2 Pathway, VEGFR2 mediated cell proliferation 
                                   NM_153831.2:965 
                                 
                                 
                                   CRKL-mRNA 
                                   0.0742 
                                   0.0136 
                                   0.0476 
                                   0.101 
                                   1.05 
                                   1.03 
                                   1.07 
                                   3.37e-06 
                                   0.000251 
                                   lm.nb 
                                   Downstream signal transduction, Frs2-mediated activation, Regulation of signaling by CBL, Signaling by PDGF 
                                   NM_005207.3:1372 
                                 
                                 
                                   CYFIP2-mRNA 
                                   0.387 
                                   0.0717 
                                   0.247 
                                   0.528 
                                   1.31 
                                   1.19 
                                   1.44 
                                   4.06e-06 
                                   0.000288 
                                   lm.nb 
                                   Regulation of actin dynamics for phagocytic cup formation, RHO GTPases Activate WASPs and WAVEs, VEGFA-VEGFR2 Pathway 
                                   NM_001037332.2:4990 
                                 
                               
                               DE Results: Day  Table presenting the most statistically significantly differentially expressed genes with the selected covariate. 'Estimated log fold-change' estimates a gene's differential expression. For categorical covariates, a gene is estimated to have 2^(log fold change) times its expression in baseline samples, holding all other variables in the analysis constant. If the covariate is continuous, for each unit increase in the selected covariate, a gene's expression is estimated to increase by 2^(log fold change)-fold, holding all other variables in the analysis constant. The 95% confidence interval for the log fold change is also presented, along with a p-value and an adjusted p-value or FDR if requested. Method column indicates the model used to estimate differential expression. Mixed negative binomial model uses the mle function to run the Wald test. The simplified negative binomial model uses the glm.nb function . The loglinear model uses the lm function.   
                                   
                          
                           
                      
                    
                  
                
                
                  
                    
                      
                           
                          
                               Volcano Plot: Line: Caco2 vs.Fibro   More Plot Information                Volcano Plot: Line: Caco2 vs.Fibro  Volcano plot displaying each gene's -log10(p-value) and log2 fold change with the selected covariate. Highly statistically significant genes fall at the top of the plot above the horizontal lines, and highly differentially expressed genes fall to either side. Horizontal lines indicate various False Discovery Rate (FDR) thresholds or p-value thresholds if there is no adjustment to the p-values. Genes are colored if the resulting p-value is below the given FDR or p-value threshold. The 40 most statistically significant genes are labeled in the plot.   
                               DE Results: Line: Caco2 vs.Fibro   More Plot Information      Download CSV Data     
                               
                                 
                                 
                               
                               
                                 
                                   
                                   Log2 fold change 
                                   std error (log2) 
                                   Lower confidence limit (log2) 
                                   Upper confidence limit (log2) 
                                   Linear fold change 
                                   Lower confidence limit (linear) 
                                   Upper confidence limit (linear) 
                                   P-value 
                                   BY.p.value 
                                   method 
                                   Gene.sets 
                                   probe.ID 
                                 
                               
                               
                                 
                                   F11R-mRNA 
                                   6.84 
                                   0.0726 
                                   6.7 
                                   6.98 
                                   114 
                                   104 
                                   126 
                                   1.12e-45 
                                   1.59e-42 
                                   loglinear 
                                   Cell surface interactions at the vascular wall, Integrin cell surface interactions, TGF-beta receptor signaling in EMT (epithelial to mesenchymal transition), Tight junction interactions 
                                   NM_144503.1:3650 
                                 
                                 
                                   CLDN4-mRNA 
                                   7.95 
                                   0.241 
                                   7.48 
                                   8.42 
                                   247 
                                   178 
                                   343 
                                   4.89e-29 
                                   3.47e-26 
                                   lm.nb 
                                   Tight junction interactions 
                                   NM_001305.3:1242 
                                 
                                 
                                   MYLK-mRNA 
                                   -6.27 
                                   0.235 
                                   -6.73 
                                   -5.81 
                                   0.013 
                                   0.00943 
                                   0.0178 
                                   8.55e-26 
                                   4.04e-23 
                                   lm.nb 
                                   RHO GTPases activate PAKs, Smooth Muscle Contraction 
                                   NM_053032.2:710 
                                 
                                 
                                   BAIAP2L1-mRNA 
                                   2.53 
                                   0.102 
                                   2.33 
                                   2.73 
                                   5.79 
                                   5.04 
                                   6.65 
                                   1.29e-24 
                                   4.25e-22 
                                   lm.nb 
                                   
                                   NM_018842.3:259 
                                 
                                 
                                   PAK4-mRNA 
                                   2.18 
                                   0.0884 
                                   2 
                                   2.35 
                                   4.52 
                                   4.01 
                                   5.1 
                                   1.5e-24 
                                   4.25e-22 
                                   loglinear 
                                   Activation of Rac 
                                   NM_005884.3:2130 
                                 
                                 
                                   KRAS-mRNA 
                                   2.3 
                                   0.0937 
                                   2.11 
                                   2.48 
                                   4.91 
                                   4.32 
                                   5.58 
                                   1.8e-24 
                                   4.27e-22 
                                   lm.nb 
                                   ARMS-mediated activation, CD209 (DC-SIGN) signaling, Cell surface interactions at the vascular wall, Constitutive Signaling by EGFRvIII, DAP12 signaling, Downstream signal transduction, FCERI mediated MAPK activation, Frs2-mediated activation, Gastrin-CREB signalling pathway via PKC and MAPK, Interleukin receptor SHC signaling, IRS-related events triggered by IGF1R, MAP2K and MAPK activation, RAFMAP kinase cascade, SHC-mediated cascade.CL.FGFR1, SHC-related events triggered by IGF1R, Signaling by ERBB2, Signaling by FGFR1 in disease, Signaling by FGFR2 in disease, Signaling by FGFR3 in disease, Signaling by FGFR4 in disease, Signaling by PDGF, Signaling by SCF-KIT, Tie2 Signaling, VEGFA-VEGFR2 Pathway, VEGFR2 mediated cell proliferation 
                                   NM_004985.3:327 
                                 
                                 
                                   ADAM9-mRNA 
                                   -2.17 
                                   0.0895 
                                   -2.35 
                                   -2 
                                   0.222 
                                   0.196 
                                   0.25 
                                   2.47e-24 
                                   5e-22 
                                   loglinear 
                                   
                                   NM_001005845.1:695 
                                 
                                 
                                   CRB3-mRNA 
                                   7.21 
                                   0.298 
                                   6.62 
                                   7.79 
                                   148 
                                   98.7 
                                   222 
                                   2.93e-24 
                                   5.2e-22 
                                   lm.nb 
                                   Tight junction interactions 
                                   NM_139161.3:300 
                                 
                                 
                                   ITGB1-mRNA 
                                   -2.14 
                                   0.0938 
                                   -2.32 
                                   -1.95 
                                   0.227 
                                   0.2 
                                   0.258 
                                   2.21e-23 
                                   3.48e-21 
                                   lm.nb 
                                   Cell-extracellular matrix interactions, Cell surface interactions at the vascular wall, ECM proteoglycans, Elastic fibre formation, Integrin cell surface interactions, Laminin interactions, Non-integrin membrane-ECM interactions, RHO GTPases Activate Formins 
                                   NM_033666.2:2000 
                                 
                                 
                                   ITGAV-mRNA 
                                   -2.46 
                                   0.111 
                                   -2.67 
                                   -2.24 
                                   0.182 
                                   0.157 
                                   0.212 
                                   5.51e-23 
                                   7.82e-21 
                                   lm.nb 
                                   Cell surface interactions at the vascular wall, ECM proteoglycans, Elastic fibre formation, Integrin cell surface interactions, Laminin interactions, Molecules associated with elastic fibres, Non-integrin membrane-ECM interactions, Signal transduction by L1, Syndecan interactions, VEGFA-VEGFR2 Pathway 
                                   NM_002210.2:2615 
                                 
                                 
                                   CLDN7-mRNA 
                                   8.5 
                                   0.393 
                                   7.73 
                                   9.27 
                                   362 
                                   212 
                                   617 
                                   1.32e-22 
                                   1.71e-20 
                                   lm.nb 
                                   Tight junction interactions 
                                   NM_001307.3:175 
                                 
                                 
                                   TP53-mRNA 
                                   -3.1 
                                   0.145 
                                   -3.39 
                                   -2.82 
                                   0.116 
                                   0.0956 
                                   0.142 
                                   1.85e-22 
                                   2.19e-20 
                                   lm.nb 
                                   
                                   NM_000546.2:1330 
                                 
                                 
                                   OCLN-mRNA 
                                   4.7 
                                   0.221 
                                   4.26 
                                   5.13 
                                   25.9 
                                   19.2 
                                   35 
                                   2.42e-22 
                                   2.64e-20 
                                   lm.nb 
                                   Apoptotic cleavage of cellular proteins 
                                   NM_002538.3:5130 
                                 
                                 
                                   LYZ-mRNA 
                                   7.51 
                                   0.357 
                                   6.81 
                                   8.21 
                                   182 
                                   112 
                                   296 
                                   3.52e-22 
                                   3.57e-20 
                                   lm.nb 
                                   
                                   NM_000239.2:305 
                                 
                                 
                                   RAC3-mRNA 
                                   3.64 
                                   0.179 
                                   3.29 
                                   3.99 
                                   12.5 
                                   9.77 
                                   15.9 
                                   1.18e-21 
                                   1.12e-19 
                                   lm.nb 
                                   PCPCE pathway, Rho GTPase cycle 
                                   NM_005052.2:702 
                                 
                                 
                                   MLLT4-mRNA 
                                   2.72 
                                   0.145 
                                   2.44 
                                   3 
                                   6.6 
                                   5.42 
                                   8.03 
                                   1.52e-20 
                                   1.35e-18 
                                   lm.nb 
                                   
                                   NM_005936.2:1560 
                                 
                                 
                                   INADL-mRNA 
                                   3.41 
                                   0.184 
                                   3.05 
                                   3.77 
                                   10.6 
                                   8.26 
                                   13.6 
                                   2.54e-20 
                                   2.12e-18 
                                   lm.nb 
                                   
                                   XM_005270347.1:343 
                                 
                                 
                                   MAPK10-mRNA 
                                   -4.81 
                                   0.26 
                                   -5.32 
                                   -4.3 
                                   0.0355 
                                   0.025 
                                   0.0506 
                                   2.84e-20 
                                   2.17e-18 
                                   lm.nb 
                                   FCERI mediated MAPK activation 
                                   NM_138981.2:450 
                                 
                                 
                                   GNAI2-mRNA 
                                   -2.14 
                                   0.116 
                                   -2.37 
                                   -1.92 
                                   0.226 
                                   0.193 
                                   0.265 
                                   2.9e-20 
                                   2.17e-18 
                                   lm.nb 
                                   
                                   NM_002070.2:1000 
                                 
                                 
                                   MET-mRNA 
                                   2.88 
                                   0.16 
                                   2.56 
                                   3.19 
                                   7.35 
                                   5.91 
                                   9.14 
                                   7.57e-20 
                                   5.37e-18 
                                   lm.nb 
                                   
                                   NM_001127500.1:1925 
                                 
                               
                               DE Results: Line: Caco2 vs.Fibro  Table presenting the most statistically significantly differentially expressed genes with the selected covariate. 'Estimated log fold-change' estimates a gene's differential expression. For categorical covariates, a gene is estimated to have 2^(log fold change) times its expression in baseline samples, holding all other variables in the analysis constant. If the covariate is continuous, for each unit increase in the selected covariate, a gene's expression is estimated to increase by 2^(log fold change)-fold, holding all other variables in the analysis constant. The 95% confidence interval for the log fold change is also presented, along with a p-value and an adjusted p-value or FDR if requested. Method column indicates the model used to estimate differential expression. Mixed negative binomial model uses the mle function to run the Wald test. The simplified negative binomial model uses the glm.nb function . The loglinear model uses the lm function.   
                                   
                          
                           
                      
                    
                    
                      
                           
                          
                               Volcano Plot: Line: HEPG2 vs.Fibro   More Plot Information                Volcano Plot: Line: HEPG2 vs.Fibro  Volcano plot displaying each gene's -log10(p-value) and log2 fold change with the selected covariate. Highly statistically significant genes fall at the top of the plot above the horizontal lines, and highly differentially expressed genes fall to either side. Horizontal lines indicate various False Discovery Rate (FDR) thresholds or p-value thresholds if there is no adjustment to the p-values. Genes are colored if the resulting p-value is below the given FDR or p-value threshold. The 40 most statistically significant genes are labeled in the plot.   
                               DE Results: Line: HEPG2 vs.Fibro   More Plot Information      Download CSV Data     
                               
                                 
                                 
                               
                               
                                 
                                   
                                   Log2 fold change 
                                   std error (log2) 
                                   Lower confidence limit (log2) 
                                   Upper confidence limit (log2) 
                                   Linear fold change 
                                   Lower confidence limit (linear) 
                                   Upper confidence limit (linear) 
                                   P-value 
                                   BY.p.value 
                                   method 
                                   Gene.sets 
                                   probe.ID 
                                 
                               
                               
                                 
                                   F11R-mRNA 
                                   5.4 
                                   0.0797 
                                   5.24 
                                   5.56 
                                   42.3 
                                   37.9 
                                   47.1 
                                   2.04e-40 
                                   2.89e-37 
                                   loglinear 
                                   Cell surface interactions at the vascular wall, Integrin cell surface interactions, TGF-beta receptor signaling in EMT (epithelial to mesenchymal transition), Tight junction interactions 
                                   NM_144503.1:3650 
                                 
                                 
                                   LYZ-mRNA 
                                   13.5 
                                   0.379 
                                   12.7 
                                   14.2 
                                   11400 
                                   6790 
                                   19000 
                                   3.38e-30 
                                   2.4e-27 
                                   lm.nb 
                                   
                                   NM_000239.2:305 
                                 
                                 
                                   CTBP2-mRNA 
                                   -5.05 
                                   0.149 
                                   -5.34 
                                   -4.75 
                                   0.0303 
                                   0.0247 
                                   0.0371 
                                   1.89e-29 
                                   8.93e-27 
                                   lm.nb 
                                   Repression of WNT target genes 
                                   NM_001329.2:1884 
                                 
                                 
                                   AKT3-mRNA 
                                   -7.38 
                                   0.252 
                                   -7.87 
                                   -6.89 
                                   0.006 
                                   0.00426 
                                   0.00845 
                                   3.23e-27 
                                   1.15e-24 
                                   lm.nb 
                                   Activation of BAD and translocation to mitochondria , CD28 co-stimulation, CD28 dependent PI3KAkt signaling, DAP12 signaling, Downstream signal transduction, G beta.CL.gamma signalling through PI3Kgamma, GAB1 signalosome, GPVI-mediated activation cascade, PI3KAKT activation, PIP3 activates AKT signaling, Role of LAT2NTALLAB on calcium mobilization, Signaling by ERBB2, Signaling by PDGF, Signaling by SCF-KIT, TP53 Regulates Metabolic Genes, VEGFA-VEGFR2 Pathway, VEGFR2 mediated vascular permeability 
                                   NM_005465.4:287 
                                 
                                 
                                   CD9-mRNA 
                                   -3.37 
                                   0.119 
                                   -3.61 
                                   -3.14 
                                   0.0965 
                                   0.0821 
                                   0.113 
                                   1.07e-26 
                                   2.78e-24 
                                   lm.nb 
                                   Platelet degranulation  
                                   NM_001769.2:405 
                                 
                                 
                                   ITGA5-mRNA 
                                   -9.22 
                                   0.326 
                                   -9.86 
                                   -8.58 
                                   0.00168 
                                   0.00108 
                                   0.00261 
                                   1.18e-26 
                                   2.78e-24 
                                   lm.nb 
                                   Cell surface interactions at the vascular wall, Elastic fibre formation, Integrin cell surface interactions, Signal transduction by L1 
                                   NM_002205.2:925 
                                 
                                 
                                   MYL9-mRNA 
                                   -8.2 
                                   0.308 
                                   -8.8 
                                   -7.59 
                                   0.00341 
                                   0.00224 
                                   0.00519 
                                   1.03e-25 
                                   2.1e-23 
                                   lm.nb 
                                   EPHA-mediated growth cone collapse, RHO GTPases activate CIT, RHO GTPases activate PAKs, RHO GTPases activate PKNs, RHO GTPases Activate ROCKs, Sema4D induced cell migration and growth-cone collapse, Smooth Muscle Contraction 
                                   NM_181526.1:133 
                                 
                                 
                                   TBP-mRNA 
                                   1.68 
                                   0.0636 
                                   1.56 
                                   1.81 
                                   3.21 
                                   2.94 
                                   3.5 
                                   1.27e-25 
                                   2.26e-23 
                                   loglinear 
                                   
                                   NM_001172085.1:587 
                                 
                                 
                                   PDGFRA-mRNA 
                                   -7.73 
                                   0.294 
                                   -8.31 
                                   -7.16 
                                   0.0047 
                                   0.00315 
                                   0.00701 
                                   1.54e-25 
                                   2.43e-23 
                                   lm.nb 
                                   ARMS-mediated activation, Constitutive Signaling by Aberrant PI3K in Cancer, DAP12 signaling, Downstream signal transduction, FCERI mediated MAPK activation, Frs2-mediated activation, GAB1 signalosome, Gastrin-CREB signalling pathway via PKC and MAPK, Interleukin receptor SHC signaling, IRS-related events triggered by IGF1R, PI3KAKT activation, PIP3 activates AKT signaling, RAFMAP kinase cascade, Role of LAT2NTALLAB on calcium mobilization, Signaling by ERBB2, Signaling by PDGF, Signaling by SCF-KIT, VEGFA-VEGFR2 Pathway, VEGFR2 mediated cell proliferation 
                                   NM_006206.3:1925 
                                 
                                 
                                   CLDN4-mRNA 
                                   6.14 
                                   0.25 
                                   5.65 
                                   6.63 
                                   70.6 
                                   50.3 
                                   99.2 
                                   1.67e-24 
                                   2.37e-22 
                                   lm.nb 
                                   Tight junction interactions 
                                   NM_001305.3:1242 
                                 
                                 
                                   MLLT4-mRNA 
                                   3.75 
                                   0.157 
                                   3.44 
                                   4.06 
                                   13.5 
                                   10.9 
                                   16.7 
                                   4.7e-24 
                                   6.06e-22 
                                   lm.nb 
                                   
                                   NM_005936.2:1560 
                                 
                                 
                                   CRB3-mRNA 
                                   6.88 
                                   0.302 
                                   6.29 
                                   7.47 
                                   118 
                                   78.2 
                                   177 
                                   2.18e-23 
                                   2.58e-21 
                                   lm.nb 
                                   Tight junction interactions 
                                   NM_139161.3:300 
                                 
                                 
                                   ITGA6-mRNA 
                                   3.36 
                                   0.153 
                                   3.06 
                                   3.66 
                                   10.3 
                                   8.36 
                                   12.7 
                                   7.67e-23 
                                   8.37e-21 
                                   lm.nb 
                                   Cell surface interactions at the vascular wall, Non-integrin membrane-ECM interactions, Syndecan interactions 
                                   NM_000210.1:3065 
                                 
                                 
                                   AXIN2-mRNA 
                                   5.52 
                                   0.278 
                                   4.97 
                                   6.06 
                                   45.8 
                                   31.4 
                                   66.8 
                                   2.49e-21 
                                   2.53e-19 
                                   lm.nb 
                                   Formation of the beta-catenin.CL.TCF transactivating complex 
                                   NM_004655.3:1035 
                                 
                                 
                                   PAK4-mRNA 
                                   1.84 
                                   0.097 
                                   1.65 
                                   2.03 
                                   3.58 
                                   3.13 
                                   4.08 
                                   1.24e-20 
                                   1.18e-18 
                                   loglinear 
                                   Activation of Rac 
                                   NM_005884.3:2130 
                                 
                                 
                                   CD44-mRNA 
                                   -8.44 
                                   0.447 
                                   -9.32 
                                   -7.57 
                                   0.00287 
                                   0.00156 
                                   0.00527 
                                   1.33e-20 
                                   1.18e-18 
                                   lm.nb 
                                   Cell surface interactions at the vascular wall 
                                   NM_001001392.1:429 
                                 
                                 
                                   EPB41-mRNA 
                                   3.9 
                                   0.208 
                                   3.5 
                                   4.31 
                                   15 
                                   11.3 
                                   19.8 
                                   1.73e-20 
                                   1.45e-18 
                                   lm.nb 
                                   
                                   NM_001166005.1:1330 
                                 
                                 
                                   PIK3CB-mRNA 
                                   1.85 
                                   0.0992 
                                   1.66 
                                   2.05 
                                   3.61 
                                   3.15 
                                   4.13 
                                   2.04e-20 
                                   1.61e-18 
                                   loglinear 
                                   Cell surface interactions at the vascular wall, Constitutive Signaling by Aberrant PI3K in Cancer, DAP12 signaling, Downstream signal transduction, Downstream TCR signaling, G beta.CL.gamma signalling through PI3Kgamma, GAB1 signalosome, GPVI-mediated activation cascade, Interleukin receptor SHC signaling, IRS-related events triggered by IGF1R, Nephrin interactions, PI3K Cascade, PI3KAKT activation, PIP3 activates AKT signaling, Regulation of signaling by CBL, Role of LAT2NTALLAB on calcium mobilization, Signaling by ERBB2, Signaling by PDGF, Signaling by SCF-KIT, Synthesis of PIPs at the plasma membrane, Tie2 Signaling, VEGFA-VEGFR2 Pathway 
                                   NM_006219.1:2945 
                                 
                                 
                                   BAIAP2L1-mRNA 
                                   2.03 
                                   0.111 
                                   1.81 
                                   2.25 
                                   4.09 
                                   3.52 
                                   4.75 
                                   3.7e-20 
                                   2.77e-18 
                                   lm.nb 
                                   
                                   NM_018842.3:259 
                                 
                                 
                                   MAPK10-mRNA 
                                   -5.48 
                                   0.311 
                                   -6.09 
                                   -4.87 
                                   0.0225 
                                   0.0147 
                                   0.0343 
                                   1.41e-19 
                                   9.99e-18 
                                   lm.nb 
                                   FCERI mediated MAPK activation 
                                   NM_138981.2:450 
                                 
                               
                               DE Results: Line: HEPG2 vs.Fibro  Table presenting the most statistically significantly differentially expressed genes with the selected covariate. 'Estimated log fold-change' estimates a gene's differential expression. For categorical covariates, a gene is estimated to have 2^(log fold change) times its expression in baseline samples, holding all other variables in the analysis constant. If the covariate is continuous, for each unit increase in the selected covariate, a gene's expression is estimated to increase by 2^(log fold change)-fold, holding all other variables in the analysis constant. The 95% confidence interval for the log fold change is also presented, along with a p-value and an adjusted p-value or FDR if requested. Method column indicates the model used to estimate differential expression. Mixed negative binomial model uses the mle function to run the Wald test. The simplified negative binomial model uses the glm.nb function . The loglinear model uses the lm function.   
                                   
                          
                           
                      
                    
                    
                      
                           
                          
                               Volcano Plot: Line: HT29 vs.Fibro   More Plot Information                Volcano Plot: Line: HT29 vs.Fibro  Volcano plot displaying each gene's -log10(p-value) and log2 fold change with the selected covariate. Highly statistically significant genes fall at the top of the plot above the horizontal lines, and highly differentially expressed genes fall to either side. Horizontal lines indicate various False Discovery Rate (FDR) thresholds or p-value thresholds if there is no adjustment to the p-values. Genes are colored if the resulting p-value is below the given FDR or p-value threshold. The 40 most statistically significant genes are labeled in the plot.   
                               DE Results: Line: HT29 vs.Fibro   More Plot Information      Download CSV Data     
                               
                                 
                                 
                               
                               
                                 
                                   
                                   Log2 fold change 
                                   std error (log2) 
                                   Lower confidence limit (log2) 
                                   Upper confidence limit (log2) 
                                   Linear fold change 
                                   Lower confidence limit (linear) 
                                   Upper confidence limit (linear) 
                                   P-value 
                                   BY.p.value 
                                   method 
                                   Gene.sets 
                                   probe.ID 
                                 
                               
                               
                                 
                                   F11R-mRNA 
                                   6.55 
                                   0.075 
                                   6.4 
                                   6.7 
                                   93.7 
                                   84.6 
                                   104 
                                   1.87e-44 
                                   2.66e-41 
                                   loglinear 
                                   Cell surface interactions at the vascular wall, Integrin cell surface interactions, TGF-beta receptor signaling in EMT (epithelial to mesenchymal transition), Tight junction interactions 
                                   NM_144503.1:3650 
                                 
                                 
                                   EPB41L2-mRNA 
                                   -5.22 
                                   0.119 
                                   -5.45 
                                   -4.98 
                                   0.0269 
                                   0.0229 
                                   0.0317 
                                   1.87e-33 
                                   1.33e-30 
                                   lm.nb 
                                   
                                   NM_001199389.1:1880 
                                 
                                 
                                   MYL9-mRNA 
                                   -11.6 
                                   0.313 
                                   -12.3 
                                   -11 
                                   0.000312 
                                   0.000204 
                                   0.000477 
                                   6.43e-31 
                                   3.04e-28 
                                   lm.nb 
                                   EPHA-mediated growth cone collapse, RHO GTPases activate CIT, RHO GTPases activate PAKs, RHO GTPases activate PKNs, RHO GTPases Activate ROCKs, Sema4D induced cell migration and growth-cone collapse, Smooth Muscle Contraction 
                                   NM_181526.1:133 
                                 
                                 
                                   LYZ-mRNA 
                                   13.3 
                                   0.365 
                                   12.6 
                                   14.1 
                                   10400 
                                   6350 
                                   17100 
                                   1.16e-30 
                                   4.1e-28 
                                   lm.nb 
                                   
                                   NM_000239.2:305 
                                 
                                 
                                   CLDN4-mRNA 
                                   8.58 
                                   0.244 
                                   8.1 
                                   9.06 
                                   382 
                                   274 
                                   533 
                                   4.85e-30 
                                   1.38e-27 
                                   lm.nb 
                                   Tight junction interactions 
                                   NM_001305.3:1242 
                                 
                                 
                                   BAIAP2L1-mRNA 
                                   3.38 
                                   0.105 
                                   3.18 
                                   3.59 
                                   10.4 
                                   9.06 
                                   12 
                                   1.05e-28 
                                   2.47e-26 
                                   lm.nb 
                                   
                                   NM_018842.3:259 
                                 
                                 
                                   PAK4-mRNA 
                                   2.93 
                                   0.0914 
                                   2.75 
                                   3.11 
                                   7.64 
                                   6.75 
                                   8.65 
                                   1.3e-28 
                                   2.64e-26 
                                   loglinear 
                                   Activation of Rac 
                                   NM_005884.3:2130 
                                 
                                 
                                   AKT3-mRNA 
                                   -8.38 
                                   0.265 
                                   -8.9 
                                   -7.86 
                                   0.003 
                                   0.00209 
                                   0.0043 
                                   2.3e-28 
                                   4.09e-26 
                                   lm.nb 
                                   Activation of BAD and translocation to mitochondria , CD28 co-stimulation, CD28 dependent PI3KAkt signaling, DAP12 signaling, Downstream signal transduction, G beta.CL.gamma signalling through PI3Kgamma, GAB1 signalosome, GPVI-mediated activation cascade, PI3KAKT activation, PIP3 activates AKT signaling, Role of LAT2NTALLAB on calcium mobilization, Signaling by ERBB2, Signaling by PDGF, Signaling by SCF-KIT, TP53 Regulates Metabolic Genes, VEGFA-VEGFR2 Pathway, VEGFR2 mediated vascular permeability 
                                   NM_005465.4:287 
                                 
                                 
                                   PDGFRA-mRNA 
                                   -7.96 
                                   0.273 
                                   -8.5 
                                   -7.42 
                                   0.00402 
                                   0.00277 
                                   0.00582 
                                   4.02e-27 
                                   6.33e-25 
                                   lm.nb 
                                   ARMS-mediated activation, Constitutive Signaling by Aberrant PI3K in Cancer, DAP12 signaling, Downstream signal transduction, FCERI mediated MAPK activation, Frs2-mediated activation, GAB1 signalosome, Gastrin-CREB signalling pathway via PKC and MAPK, Interleukin receptor SHC signaling, IRS-related events triggered by IGF1R, PI3KAKT activation, PIP3 activates AKT signaling, RAFMAP kinase cascade, Role of LAT2NTALLAB on calcium mobilization, Signaling by ERBB2, Signaling by PDGF, Signaling by SCF-KIT, VEGFA-VEGFR2 Pathway, VEGFR2 mediated cell proliferation 
                                   NM_006206.3:1925 
                                 
                                 
                                   FGFR1-mRNA 
                                   -9.61 
                                   0.332 
                                   -10.3 
                                   -8.96 
                                   0.00128 
                                   0.000816 
                                   0.00201 
                                   5.21e-27 
                                   7.39e-25 
                                   lm.nb 
                                   ARMS-mediated activation, Constitutive Signaling by Aberrant PI3K in Cancer, DAP12 signaling, Downstream signal transduction, FCERI mediated MAPK activation, Frs2-mediated activation, GAB1 signalosome, Gastrin-CREB signalling pathway via PKC and MAPK, Interleukin receptor SHC signaling, IRS-related events triggered by IGF1R, PI3K Cascade, PI3KAKT activation, RAFMAP kinase cascade, Role of LAT2NTALLAB on calcium mobilization, SHC-mediated cascade.CL.FGFR1, Signaling by ERBB2, Signaling by FGFR1 in disease, Signaling by PDGF, Signaling by SCF-KIT, VEGFA-VEGFR2 Pathway, VEGFR2 mediated cell proliferation 
                                   NM_015850.2:1335 
                                 
                                 
                                   FN1-mRNA 
                                   -9.78 
                                   0.348 
                                   -10.5 
                                   -9.1 
                                   0.00114 
                                   0.000708 
                                   0.00182 
                                   1.46e-26 
                                   1.89e-24 
                                   lm.nb 
                                   ARMS-mediated activation, DAP12 signaling, Downstream signal transduction, ECM proteoglycans, Elastic fibre formation, FCERI mediated MAPK activation, Frs2-mediated activation, Gastrin-CREB signalling pathway via PKC and MAPK, Integrin alphaIIb beta3 signaling, Integrin cell surface interactions, Interleukin receptor SHC signaling, IRS-related events triggered by IGF1R, MAP2K and MAPK activation, Molecules associated with elastic fibres, Non-integrin membrane-ECM interactions, p130Cas linkage to MAPK signaling for integrins, Platelet degranulation , RAFMAP kinase cascade, Signaling by ERBB2, Signaling by PDGF, Signaling by SCF-KIT, Syndecan interactions, VEGFA-VEGFR2 Pathway, VEGFR2 mediated cell proliferation 
                                   NM_212482.1:1776 
                                 
                                 
                                   ITGA5-mRNA 
                                   -11.6 
                                   0.414 
                                   -12.4 
                                   -10.8 
                                   0.000321 
                                   0.000183 
                                   0.000563 
                                   1.66e-26 
                                   1.96e-24 
                                   lm.nb 
                                   Cell surface interactions at the vascular wall, Elastic fibre formation, Integrin cell surface interactions, Signal transduction by L1 
                                   NM_002205.2:925 
                                 
                                 
                                   ITGA6-mRNA 
                                   3.72 
                                   0.144 
                                   3.44 
                                   4 
                                   13.2 
                                   10.8 
                                   16 
                                   3.2e-25 
                                   3.49e-23 
                                   lm.nb 
                                   Cell surface interactions at the vascular wall, Non-integrin membrane-ECM interactions, Syndecan interactions 
                                   NM_000210.1:3065 
                                 
                                 
                                   CRB3-mRNA 
                                   7.61 
                                   0.299 
                                   7.02 
                                   8.19 
                                   195 
                                   130 
                                   293 
                                   5.05e-25 
                                   5.12e-23 
                                   lm.nb 
                                   Tight junction interactions 
                                   NM_139161.3:300 
                                 
                                 
                                   CLDN7-mRNA 
                                   9.85 
                                   0.395 
                                   9.08 
                                   10.6 
                                   922 
                                   539 
                                   1580 
                                   9.49e-25 
                                   8.98e-23 
                                   lm.nb 
                                   Tight junction interactions 
                                   NM_001307.3:175 
                                 
                                 
                                   MET-mRNA 
                                   4.07 
                                   0.165 
                                   3.75 
                                   4.4 
                                   16.8 
                                   13.5 
                                   21.1 
                                   1.47e-24 
                                   1.3e-22 
                                   lm.nb 
                                   
                                   NM_001127500.1:1925 
                                 
                                 
                                   INADL-mRNA 
                                   4.54 
                                   0.189 
                                   4.17 
                                   4.91 
                                   23.2 
                                   18 
                                   30 
                                   3.62e-24 
                                   3.02e-22 
                                   lm.nb 
                                   
                                   XM_005270347.1:343 
                                 
                                 
                                   PIK3CB-mRNA 
                                   2.18 
                                   0.0935 
                                   2 
                                   2.36 
                                   4.53 
                                   3.99 
                                   5.14 
                                   1.02e-23 
                                   8.03e-22 
                                   loglinear 
                                   Cell surface interactions at the vascular wall, Constitutive Signaling by Aberrant PI3K in Cancer, DAP12 signaling, Downstream signal transduction, Downstream TCR signaling, G beta.CL.gamma signalling through PI3Kgamma, GAB1 signalosome, GPVI-mediated activation cascade, Interleukin receptor SHC signaling, IRS-related events triggered by IGF1R, Nephrin interactions, PI3K Cascade, PI3KAKT activation, PIP3 activates AKT signaling, Regulation of signaling by CBL, Role of LAT2NTALLAB on calcium mobilization, Signaling by ERBB2, Signaling by PDGF, Signaling by SCF-KIT, Synthesis of PIPs at the plasma membrane, Tie2 Signaling, VEGFA-VEGFR2 Pathway 
                                   NM_006219.1:2945 
                                 
                                 
                                   MAGI3-mRNA 
                                   2.93 
                                   0.129 
                                   2.67 
                                   3.18 
                                   7.61 
                                   6.38 
                                   9.07 
                                   2.77e-23 
                                   2.07e-21 
                                   lm.nb 
                                   
                                   NM_152900.2:6685 
                                 
                                 
                                   MYH10-mRNA 
                                   -3.59 
                                   0.16 
                                   -3.9 
                                   -3.28 
                                   0.083 
                                   0.0668 
                                   0.103 
                                   3.47e-23 
                                   2.46e-21 
                                   lm.nb 
                                   EPHA-mediated growth cone collapse, RHO GTPases activate CIT, RHO GTPases activate PAKs, RHO GTPases activate PKNs, RHO GTPases Activate ROCKs, Sema4D induced cell migration and growth-cone collapse 
                                   NM_005964.1:7092 
                                 
                               
                               DE Results: Line: HT29 vs.Fibro  Table presenting the most statistically significantly differentially expressed genes with the selected covariate. 'Estimated log fold-change' estimates a gene's differential expression. For categorical covariates, a gene is estimated to have 2^(log fold change) times its expression in baseline samples, holding all other variables in the analysis constant. If the covariate is continuous, for each unit increase in the selected covariate, a gene's expression is estimated to increase by 2^(log fold change)-fold, holding all other variables in the analysis constant. The 95% confidence interval for the log fold change is also presented, along with a p-value and an adjusted p-value or FDR if requested. Method column indicates the model used to estimate differential expression. Mixed negative binomial model uses the mle function to run the Wald test. The simplified negative binomial model uses the glm.nb function . The loglinear model uses the lm function.   
                                   
                          
                           
                      
                    
                  
                
               
                
              
                
          
        
        
          
                The results of differential expression testing are summarized at the gene set level. Each gene set's most differentially expressed genes are identified, and the extent of differential expression in each gene set is summarized using a 'global significance score'.     
              
                   Gene sets: 
                  
                      Summary  
                      Activation of BAD and translocation to mitochondria   
                      Activation of Rac  
                      Apoptotic cleavage of cellular proteins  
                      ARMS-mediated activation  
                      AXIN missense mutants destabilize the destruction complex  
                      Beta-catenin phosphorylation cascade  
                      CD209 (DC-SIGN) signaling  
                      CD28 co-stimulation  
                      CD28 dependent PI3KAkt signaling  
                      CD28 dependent Vav1 pathway  
                      Cell-extracellular matrix interactions  
                      Cell surface interactions at the vascular wall  
                      Circadian Clock  
                      Constitutive Signaling by Aberrant PI3K in Cancer  
                      Constitutive Signaling by EGFRvIII  
                      CREB phosphorylation through the activation of Ras  
                      CTLA4 inhibitory signaling  
                      Cyclin AB1 associated events during G2M transition  
                      Cyclin D associated events in G1  
                      DAP12 signaling  
                      DARPP-32 events  
                      DCC mediated attractive signaling  
                      Deactivation of the beta-catenin transactivating complex  
                      Disassembly of the destruction complex and recruitment of AXIN t...  
                      Downstream signal transduction  
                      Downstream TCR signaling  
                      ECM proteoglycans  
                      Elastic fibre formation  
                      EPH-ephrin mediated repulsion of cells  
                      EPHA-mediated growth cone collapse  
                      EPHB-mediated forward signaling  
                      Ephrin signaling  
                      ERKs are inactivated  
                      FCERI mediated MAPK activation  
                      Formation of the beta-catenin.CL.TCF transactivating complex  
                      Frs2-mediated activation  
                      G alpha (1213) signalling events  
                      G alpha (q) signalling events  
                      G beta.CL.gamma signalling through PI3Kgamma  
                      GAB1 signalosome  
                      Gastrin-CREB signalling pathway via PKC and MAPK  
                      GPVI-mediated activation cascade  
                      Integrin alphaIIb beta3 signaling  
                      Integrin cell surface interactions  
                      Interleukin receptor SHC signaling  
                      IRS-related events triggered by IGF1R  
                      Laminin interactions  
                      MAP2K and MAPK activation  
                      MAPK6MAPK4 signaling  
                      MASTL Facilitates Mitotic Progression  
                      MicroRNA (miRNA) biogenesis  
                      Misspliced GSK3beta mutants stabilize beta-catenin  
                      Molecules associated with elastic fibres  
                      Negative feedback regulation of MAPK pathway  
                      Negative regulation of FGFR1 signaling  
                      Negative regulation of FGFR2 signaling  
                      Negative regulation of FGFR3 signaling  
                      Negative regulation of FGFR4 signaling  
                      Negative regulation of MAPK pathway  
                      Nephrin interactions  
                      Non-integrin membrane-ECM interactions  
                      Nonsense Mediated Decay (NMD) enhanced by the Exon Junction Comp...  
                      NRAGE signals death through JNK  
                      p130Cas linkage to MAPK signaling for integrins  
                      PCPCE pathway  
                      PI3K Cascade  
                      PI3KAKT activation  
                      PIP3 activates AKT signaling  
                      Platelet degranulation   
                      RAF activation  
                      RAFMAP kinase cascade  
                      Regulation of actin dynamics for phagocytic cup formation  
                      Regulation of HSF1-mediated heat shock response  
                      Regulation of signaling by CBL  
                      Repression of WNT target genes  
                      Resolution of Sister Chromatid Cohesion  
                      Rho GTPase cycle  
                      RHO GTPases activate CIT  
                      RHO GTPases Activate Formins  
                      RHO GTPases activate IQGAPs  
                      RHO GTPases activate PAKs  
                      RHO GTPases activate PKNs  
                      RHO GTPases Activate ROCKs  
                      RHO GTPases Activate WASPs and WAVEs  
                      Role of LAT2NTALLAB on calcium mobilization  
                      S33 mutants of beta-catenin aren.SQT.t phosphorylated  
                      S37 mutants of beta-catenin aren.SQT.t phosphorylated  
                      S45 mutants of beta-catenin aren.SQT.t phosphorylated  
                      Sema3A PAK dependent Axon repulsion  
                      Sema4D induced cell migration and growth-cone collapse  
                      SHC-mediated cascade.CL.FGFR1  
                      SHC-related events triggered by IGF1R  
                      Signal transduction by L1  
                      Signaling by ERBB2  
                      Signaling by FGFR1 in disease  
                      Signaling by FGFR2 in disease  
                      Signaling by FGFR3 in disease  
                      Signaling by FGFR4 in disease  
                      Signaling by PDGF  
                      Signaling by SCF-KIT  
                      Smooth Muscle Contraction  
                      Spry regulation of FGF signaling  
                      Syndecan interactions  
                      Synthesis of PIPs at the plasma membrane  
                      T41 mutants of beta-catenin aren.SQT.t phosphorylated  
                      TGF-beta receptor signaling activates SMADs  
                      TGF-beta receptor signaling in EMT (epithelial to mesenchymal tr...  
                      Tie2 Signaling  
                      Tight junction interactions  
                      TP53 Regulates Metabolic Genes  
                      Truncations of AMER1 destabilize the destruction complex  
                      VEGFA-VEGFR2 Pathway  
                      VEGFR2 mediated cell proliferation  
                      VEGFR2 mediated vascular permeability  
                      WNT mediated activation of DVL  
                  
                   
                  
                    
                      
                           
                          
                               Global Significance Scores   More Plot Information                Global Significance Scores  Heatmap displaying each sample's global significance scores. Global significance statistics measure the extent of differential expression of a gene set's genes with a covariate, ignoring whether each gene within the set is up- or down-regulated. Orange denotes gene sets whose genes exhibit extensive differential expression with the covariate, blue denotes gene sets with less differential expression.   
                               Directed Global Significance Scores   More Plot Information                Directed Global Significance Scores  Heatmap displaying each sample's directed global significance scores. Directed global significance statistics measure the extent to which a gene set's genes are up- or down-regulated with the variable. Red denotes gene sets whose genes exhibit extensive over-expression with the covariate, blue denotes gene sets with extensive under-expression.   
                          
                           
                           
                          
                               Undirected and Directed global significance scores table   More Plot Information      Download CSV Data     
                               
                                 
                                 
                               
                               
                                 
                                   
                                   Undirected Day 
                                   Undirected Line: differential expression in Caco2 vs. baseline of Fibro 
                                   Undirected Line: differential expression in HT29 vs. baseline of Fibro 
                                   Undirected Line: differential expression in HEPG2 vs. baseline of Fibro 
                                   Directed Day 
                                   Directed Line: differential expression in Caco2 vs. baseline of Fibro 
                                   Directed Line: differential expression in HT29 vs. baseline of Fibro 
                                   Directed Line: differential expression in HEPG2 vs. baseline of Fibro 
                                 
                               
                               
                                 
                                   Activation of BAD and translocation to mitochondria  
                                   1.877 
                                   8.356 
                                   14.977 
                                   13.505 
                                   1.865 
                                   -7.966 
                                   -13.266 
                                   -12.826 
                                 
                                 
                                   Activation of Rac 
                                   3.703 
                                   12.485 
                                   16.53 
                                   10.735 
                                   2.701 
                                   12.071 
                                   16.524 
                                   6.378 
                                 
                                 
                                   Apoptotic cleavage of cellular proteins 
                                   3.999 
                                   8.61 
                                   11.021 
                                   8.472 
                                   1.56 
                                   7.417 
                                   10.606 
                                   7.618 
                                 
                                 
                                   ARMS-mediated activation 
                                   3.012 
                                   8.286 
                                   12.312 
                                   7.722 
                                   2.056 
                                   4.061 
                                   -8.287 
                                   -3.877 
                                 
                                 
                                   AXIN missense mutants destabilize the destruction complex 
                                   1.874 
                                   7.133 
                                   8.659 
                                   4.572 
                                   1.043 
                                   2.777 
                                   -4.705 
                                   3.335 
                                 
                                 
                                   Beta-catenin phosphorylation cascade 
                                   1.823 
                                   7.133 
                                   8.278 
                                   5.748 
                                   1.129 
                                   2.777 
                                   -5.346 
                                   4.823 
                                 
                                 
                                   CD209 (DC-SIGN) signaling 
                                   2.931 
                                   9.97 
                                   7.189 
                                   7.213 
                                   1.082 
                                   9.058 
                                   6.215 
                                   2.762 
                                 
                                 
                                   CD28 co-stimulation 
                                   2.927 
                                   9.094 
                                   12.629 
                                   10.6 
                                   2.421 
                                   2.435 
                                   3.457 
                                   -6.353 
                                 
                                 
                                   CD28 dependent PI3KAkt signaling 
                                   2.858 
                                   10.325 
                                   14.581 
                                   12.744 
                                   2.858 
                                   -7.585 
                                   -11.473 
                                   -11.445 
                                 
                                 
                                   CD28 dependent Vav1 pathway 
                                   3.34 
                                   6.892 
                                   8.618 
                                   6.644 
                                   1.893 
                                   3.291 
                                   8.328 
                                   -5.602 
                                 
                                 
                                   Cell-extracellular matrix interactions 
                                   2.981 
                                   13.714 
                                   11.595 
                                   9.843 
                                   -2.795 
                                   -13.714 
                                   -11.595 
                                   -9.843 
                                 
                                 
                                   Cell surface interactions at the vascular wall 
                                   3.1 
                                   25.39 
                                   24.515 
                                   20.089 
                                   1.511 
                                   20.922 
                                   20.626 
                                   15.264 
                                 
                                 
                                   Circadian Clock 
                                   3.368 
                                   7.386 
                                   8.172 
                                   8.431 
                                   3.199 
                                   7.064 
                                   8.149 
                                   8.431 
                                 
                                 
                                   Constitutive Signaling by Aberrant PI3K in Cancer 
                                   2.716 
                                   8.885 
                                   14.666 
                                   9.82 
                                   2.455 
                                   0.773 
                                   -9.387 
                                   -4.899 
                                 
                                 
                                   Constitutive Signaling by EGFRvIII 
                                   2.84 
                                   11.261 
                                   7.965 
                                   6.775 
                                   1.762 
                                   10.104 
                                   7.799 
                                   5.828 
                                 
                                 
                                   CREB phosphorylation through the activation of Ras 
                                   3.642 
                                   7.864 
                                   4.395 
                                   7.021 
                                   3.295 
                                   -4.627 
                                   -1.748 
                                   -2.778 
                                 
                                 
                                   CTLA4 inhibitory signaling 
                                   2.124 
                                   9.795 
                                   11.931 
                                   8.176 
                                   1.473 
                                   7.673 
                                   5.278 
                                   7.518 
                                 
                                 
                                   Cyclin AB1 associated events during G2M transition 
                                   2.748 
                                   10.619 
                                   10.068 
                                   5.494 
                                   2.067 
                                   9.258 
                                   -7.355 
                                   3.966 
                                 
                                 
                                   Cyclin D associated events in G1 
                                   2.663 
                                   9.528 
                                   10.477 
                                   5.734 
                                   -0.69 
                                   4.961 
                                   -8.509 
                                   1.47 
                                 
                                 
                                   DAP12 signaling 
                                   2.966 
                                   8.309 
                                   12.26 
                                   8.652 
                                   2.407 
                                   3.545 
                                   -6.103 
                                   -3.22 
                                 
                               
                               Undirected and Directed global significance scores table  Table displaying each sample's global significance scores and directed global significance scores as defined in the heatmaps above. The global significance score is calculated as the square root of the mean squared t-statistic for the genes in a gene set, with t-statistics coming from the linear regression underlying our differential expression analysis. The directed global significance score is calculated as the square root of the mean signed squared t-statistic for the genes in a gene set, with t-statistics coming from the linear regression underlying our differential expression analysis.   
                          
                           
                      
                    
                    
                      
                           
                          
                               Day    Line     
                              
                                  Day  
                              
                               
                              
                                  Line: differential expression in Caco2 vs. baseline of Fibro  
                                  Line: differential expression in HEPG2 vs. baseline of Fibro  
                                  Line: differential expression in HT29 vs. baseline of Fibro  
                              
                              
                            
                              
                                
                                  
                                       
                                      
                                           Volcano Plot: Day   More Plot Information                Volcano Plot: Day  Volcano plot displaying each gene's -log10(p-value) and log2 fold change for the selected covariate. Highly statistically significant genes fall at the top of the plot, and highly differentially expressed genes fall to either side. Genes within the selected gene set are highlighted in orange. Horizontal lines indicate various False Discovery Rate (FDR) thresholds.   
                                           DE Results - Activation of BAD and translocation to mitochondria  genes - Day   More Plot Information      Download CSV Data     
                                           
                                             
                                             
                                           
                                           
                                             
                                               Probe Label 
                                               Log2 fold change 
                                               std error (log2) 
                                               Lower confidence limit (log2) 
                                               Upper confidence limit (log2) 
                                               Linear fold change 
                                               Lower confidence limit (linear) 
                                               Upper confidence limit (linear) 
                                               P-value 
                                               BY.p.value 
                                               method 
                                               Gene.sets 
                                               Codeset.Name 
                                               Probe.Label 
                                               Analyte.Type 
                                               Is.Control 
                                               Control.Type 
                                               Related.Probes 
                                               Probe.Annotation 
                                               KEGG.Pathways 
                                               Official.Gene.Name 
                                               Control.Conc 
                                               volcanocol 
                                               volcanopch 
                                               tstats.all 
                                             
                                           
                                           
                                             
                                               AKT3-mRNA 
                                               0.0985 
                                               0.0254 
                                               0.0488 
                                               0.148 
                                               1.07 
                                               1.03 
                                               1.11 
                                               0.000409 
                                               0.0111 
                                               lm.nb 
                                               Activation of BAD and translocation to mitochondria , CD28 co-stimulation, CD28 dependent PI3KAkt signaling, DAP12 signaling, Downstream signal transduction, G beta.CL.gamma signalling through PI3Kgamma, GAB1 signalosome, GPVI-mediated activation cascade, PI3KAKT activation, PIP3 activates AKT signaling, Role of LAT2NTALLAB on calcium mobilization, Signaling by ERBB2, Signaling by PDGF, Signaling by SCF-KIT, TP53 Regulates Metabolic Genes, VEGFA-VEGFR2 Pathway, VEGFR2 mediated vascular permeability 
                                               SEBELIUS_2_C4636 
                                               AKT3 
                                               mRNA 
                                               0 
                                               
                                               
                                               TP53 Regulates Metabolic Genes;CD28 dependent PI3KAkt signaling;AKT phosphorylates targets in the cytosol;Negative regulation of the PI3KAKT network;Activation of BAD and translocation to mitochondria ;G beta.CL.gamma signalling through PI3Kgamma;AKT phosphorylates targets in the nucleus;VEGFR2 mediated vascular permeability;GPVI-mediated activation cascade;Signaling by ERBB2;PIP3 activates AKT signaling;Downstream signal transduction;GAB1 signalosome;Signaling by SCF-KIT;PI3KAKT activation;Role of LAT2NTALLAB on calcium mobilization;DAP12 signaling;Signaling by PDGF;CD28 co-stimulation;VEGFA-VEGFR2 Pathway 
                                               hsa04010;hsa04012;hsa04014;hsa04015;hsa04022;hsa04024;hsa04062;hsa04066;hsa04068;hsa04071;hsa04072;hsa04150;hsa04151;hsa04152;hsa04210;hsa04211;hsa04213;hsa04261;hsa04370;hsa04380;hsa04510;hsa04530;hsa04550;hsa04611;hsa04620;hsa04630;hsa04660;hsa04662;hsa04664;hsa04666;hsa04668;hsa04722;hsa04725;hsa04728;hsa04910;hsa04914;hsa04915;hsa04917;hsa04919;hsa04920;hsa04922;hsa04923;hsa04931;hsa04932;hsa04933;hsa04973;hsa05142;hsa05145;hsa05152;hsa05160;hsa05161;hsa05162;hsa05164;hsa05166;hsa05169;hsa05200;hsa05205;hsa05210;hsa05211;hsa05212;hsa05213;hsa05214;hsa05215;hsa05218;hsa05220;hsa05221;hsa05222;hsa05223;hsa05230;hsa05231 
                                               AKT3 
                                               NA 
                                               slateblue2 
                                               16 
                                               3.87795275590551 
                                             
                                             
                                               BAD-mRNA 
                                               0.0496 
                                               0.0392 
                                               -0.0273 
                                               0.126 
                                               1.03 
                                               0.981 
                                               1.09 
                                               0.214 
                                               1 
                                               lm.nb 
                                               Activation of BAD and translocation to mitochondria , DAP12 signaling, Downstream signal transduction, GAB1 signalosome, NRAGE signals death through JNK, PI3KAKT activation, PIP3 activates AKT signaling, Role of LAT2NTALLAB on calcium mobilization, Signaling by ERBB2, Signaling by PDGF, Signaling by SCF-KIT 
                                               SEBELIUS_2_C4636 
                                               BAD 
                                               mRNA 
                                               0 
                                               
                                               
                                               Activation of BAD and translocation to mitochondria ;NRAGE signals death through JNK;Constitutive Signaling by AKT1 E17K in Cancer;AKT phosphorylates targets in the cytosol;PIP3 activates AKT signaling;Downstream signal transduction;GAB1 signalosome;Signaling by SCF-KIT;PI3KAKT activation;Role of LAT2NTALLAB on calcium mobilization;DAP12 signaling;Signaling by PDGF;Signaling by ERBB2 
                                               hsa04012;hsa04014;hsa04022;hsa04024;hsa04151;hsa04210;hsa04370;hsa04510;hsa04722;hsa04910;hsa04919;hsa05010;hsa05145;hsa05152;hsa05160;hsa05161;hsa05200;hsa05203;hsa05210;hsa05212;hsa05213;hsa05215;hsa05218;hsa05220;hsa05221;hsa05223 
                                               BAD 
                                               NA 
                                               azure4 
                                               1 
                                               1.26530612244898 
                                             
                                             
                                               AKT1-mRNA 
                                               0.0219 
                                               0.031 
                                               -0.0388 
                                               0.0827 
                                               1.02 
                                               0.973 
                                               1.06 
                                               0.484 
                                               1 
                                               lm.nb 
                                               Activation of BAD and translocation to mitochondria , CD28 co-stimulation, DAP12 signaling, Deactivation of the beta-catenin transactivating complex, Downstream signal transduction, G beta.CL.gamma signalling through PI3Kgamma, GAB1 signalosome, GPVI-mediated activation cascade, IRS-related events triggered by IGF1R, PI3K Cascade, PI3KAKT activation, PIP3 activates AKT signaling, Role of LAT2NTALLAB on calcium mobilization, Signaling by ERBB2, Signaling by PDGF, Signaling by SCF-KIT, TP53 Regulates Metabolic Genes, VEGFA-VEGFR2 Pathway, VEGFR2 mediated vascular permeability 
                                               SEBELIUS_2_C4636 
                                               AKT1 
                                               mRNA 
                                               0 
                                               
                                               
                                               TP53 Regulates Metabolic Genes;Deactivation of the beta-catenin transactivating complex;Activation of BAD and translocation to mitochondria ;eNOS activation;mTOR signalling;G beta.CL.gamma signalling through PI3Kgamma;KSRP (KHSRP) binds and destabilizes mRNA;AKT phosphorylates targets in the nucleus;VEGFR2 mediated vascular permeability;GPVI-mediated activation cascade;Tetrahydrobiopterin (BH4) synthesis# recycling# salvage and regulation;Butyrate Response Factor 1 (BRF1) binds and destabilizes mRNA;Signaling by ERBB2;PIP3 activates AKT signaling;Downstream signal transduction;GAB1 signalosome;Signaling by SCF-KIT;PI3KAKT activation;Role of LAT2NTALLAB on calcium mobilization;DAP12 signaling;Signaling by PDGF;CD28 co-stimulation;PI3K Cascade;IRS-related events triggered by IGF1R;VEGFA-VEGFR2 Pathway 
                                               hsa04010;hsa04012;hsa04014;hsa04015;hsa04022;hsa04024;hsa04062;hsa04066;hsa04068;hsa04071;hsa04072;hsa04150;hsa04151;hsa04152;hsa04210;hsa04211;hsa04213;hsa04261;hsa04370;hsa04380;hsa04510;hsa04530;hsa04550;hsa04611;hsa04620;hsa04630;hsa04660;hsa04662;hsa04664;hsa04666;hsa04668;hsa04722;hsa04725;hsa04728;hsa04910;hsa04914;hsa04915;hsa04917;hsa04919;hsa04920;hsa04922;hsa04923;hsa04931;hsa04932;hsa04933;hsa04973;hsa05142;hsa05145;hsa05152;hsa05160;hsa05161;hsa05162;hsa05164;hsa05166;hsa05169;hsa05200;hsa05205;hsa05210;hsa05211;hsa05212;hsa05213;hsa05214;hsa05215;hsa05218;hsa05220;hsa05221;hsa05222;hsa05223;hsa05230;hsa05231 
                                               AKT1 
                                               NA 
                                               azure4 
                                               1 
                                               0.706451612903226 
                                             
                                             
                                               AKT2-mRNA 
                                               0.00957 
                                               0.0157 
                                               -0.0212 
                                               0.0403 
                                               1.01 
                                               0.985 
                                               1.03 
                                               0.546 
                                               1 
                                               lm.nb 
                                               Activation of BAD and translocation to mitochondria , CD28 co-stimulation, DAP12 signaling, Deactivation of the beta-catenin transactivating complex, Downstream signal transduction, G beta.CL.gamma signalling through PI3Kgamma, GAB1 signalosome, GPVI-mediated activation cascade, IRS-related events triggered by IGF1R, PI3K Cascade, PI3KAKT activation, PIP3 activates AKT signaling, Role of LAT2NTALLAB on calcium mobilization, Signaling by ERBB2, Signaling by PDGF, Signaling by SCF-KIT, TP53 Regulates Metabolic Genes, VEGFA-VEGFR2 Pathway, VEGFR2 mediated vascular permeability 
                                               SEBELIUS_2_C4636 
                                               AKT2 
                                               mRNA 
                                               0 
                                               
                                               
                                               TP53 Regulates Metabolic Genes;PDE3B signalling;AKT phosphorylates targets in the cytosol;Negative regulation of the PI3KAKT network;Deactivation of the beta-catenin transactivating complex;Activation of BAD and translocation to mitochondria ;G beta.CL.gamma signalling through PI3Kgamma;AKT phosphorylates targets in the nucleus;VEGFR2 mediated vascular permeability;GPVI-mediated activation cascade;Signaling by ERBB2;PI3K Cascade;IRS-related events triggered by IGF1R;mTOR signalling;PIP3 activates AKT signaling;Downstream signal transduction;GAB1 signalosome;Signaling by SCF-KIT;PI3KAKT activation;Role of LAT2NTALLAB on calcium mobilization;DAP12 signaling;Signaling by PDGF;CD28 co-stimulation;VEGFA-VEGFR2 Pathway 
                                               hsa04010;hsa04012;hsa04014;hsa04015;hsa04022;hsa04024;hsa04062;hsa04066;hsa04068;hsa04071;hsa04072;hsa04150;hsa04151;hsa04152;hsa04210;hsa04211;hsa04213;hsa04261;hsa04370;hsa04380;hsa04510;hsa04530;hsa04550;hsa04611;hsa04620;hsa04630;hsa04660;hsa04662;hsa04664;hsa04666;hsa04668;hsa04722;hsa04725;hsa04728;hsa04910;hsa04914;hsa04915;hsa04917;hsa04919;hsa04920;hsa04922;hsa04923;hsa04931;hsa04932;hsa04933;hsa04973;hsa05142;hsa05145;hsa05152;hsa05160;hsa05161;hsa05162;hsa05164;hsa05166;hsa05169;hsa05200;hsa05205;hsa05210;hsa05211;hsa05212;hsa05213;hsa05214;hsa05215;hsa05218;hsa05220;hsa05221;hsa05222;hsa05223;hsa05230;hsa05231 
                                               AKT2 
                                               NA 
                                               azure4 
                                               1 
                                               0.609554140127389 
                                             
                                             
                                               BCL2-mRNA 
                                               -0.102 
                                               0.305 
                                               -0.701 
                                               0.496 
                                               0.931 
                                               0.615 
                                               1.41 
                                               0.739 
                                               1 
                                               Wald 
                                               Activation of BAD and translocation to mitochondria  
                                               SEBELIUS_2_C4636 
                                               BCL2 
                                               mRNA 
                                               0 
                                               
                                               
                                               Activation of BAD and translocation to mitochondria  
                                               hsa04064;hsa04066;hsa04071;hsa04141;hsa04151;hsa04210;hsa04261;hsa04510;hsa04630;hsa04722;hsa04725;hsa04933;hsa05145;hsa05152;hsa05161;hsa05169;hsa05200;hsa05206;hsa05210;hsa05215;hsa05222 
                                               BCL2 
                                               NA 
                                               azure4 
                                               1 
                                               -0.334426229508197 
                                             
                                           
                                           DE Results - Activation of BAD and translocation to mitochondria  genes - Day  Table displaying each sample's global significance scores and directed global significance scores as defined in the heatmaps above. The global significance score is calculated as the square root of the mean squared t-statistic for the genes in a gene set, with t-statistics coming from the linear regression underlying our differential expression analysis. The directed global significance score is calculated as the square root of the mean signed squared t-statistic for the genes in a gene set, with t-statistics coming from the linear regression underlying our differential expression analysis.   
                                      
                                       
                                  
                                
                              
                            
                            
                              
                                
                                  
                                       
                                      
                                           Volcano Plot: Line: Caco2 vs.Fibro   More Plot Information                Volcano Plot: Line: Caco2 vs.Fibro  Volcano plot displaying each gene's -log10(p-value) and log2 fold change for the selected covariate. Highly statistically significant genes fall at the top of the plot, and highly differentially expressed genes fall to either side. Genes within the selected gene set are highlighted in orange. Horizontal lines indicate various False Discovery Rate (FDR) thresholds.   
                                           DE Results - Activation of BAD and translocation to mitochondria  genes - Line: Caco2 vs.Fibro   More Plot Information      Download CSV Data     
                                           
                                             
                                             
                                           
                                           
                                             
                                               Probe Label 
                                               Log2 fold change 
                                               std error (log2) 
                                               Lower confidence limit (log2) 
                                               Upper confidence limit (log2) 
                                               Linear fold change 
                                               Lower confidence limit (linear) 
                                               Upper confidence limit (linear) 
                                               P-value 
                                               BY.p.value 
                                               method 
                                               Gene.sets 
                                               Codeset.Name 
                                               Probe.Label 
                                               Analyte.Type 
                                               Is.Control 
                                               Control.Type 
                                               Related.Probes 
                                               Probe.Annotation 
                                               KEGG.Pathways 
                                               Official.Gene.Name 
                                               Control.Conc 
                                               volcanocol 
                                               volcanopch 
                                               tstats.all 
                                             
                                           
                                           
                                             
                                               AKT3-mRNA 
                                               -2.88 
                                               0.162 
                                               -3.2 
                                               -2.57 
                                               0.135 
                                               0.109 
                                               0.169 
                                               9.07e-20 
                                               6.13e-18 
                                               lm.nb 
                                               Activation of BAD and translocation to mitochondria , CD28 co-stimulation, CD28 dependent PI3KAkt signaling, DAP12 signaling, Downstream signal transduction, G beta.CL.gamma signalling through PI3Kgamma, GAB1 signalosome, GPVI-mediated activation cascade, PI3KAKT activation, PIP3 activates AKT signaling, Role of LAT2NTALLAB on calcium mobilization, Signaling by ERBB2, Signaling by PDGF, Signaling by SCF-KIT, TP53 Regulates Metabolic Genes, VEGFA-VEGFR2 Pathway, VEGFR2 mediated vascular permeability 
                                               SEBELIUS_2_C4636 
                                               AKT3 
                                               mRNA 
                                               0 
                                               
                                               
                                               TP53 Regulates Metabolic Genes;CD28 dependent PI3KAkt signaling;AKT phosphorylates targets in the cytosol;Negative regulation of the PI3KAKT network;Activation of BAD and translocation to mitochondria ;G beta.CL.gamma signalling through PI3Kgamma;AKT phosphorylates targets in the nucleus;VEGFR2 mediated vascular permeability;GPVI-mediated activation cascade;Signaling by ERBB2;PIP3 activates AKT signaling;Downstream signal transduction;GAB1 signalosome;Signaling by SCF-KIT;PI3KAKT activation;Role of LAT2NTALLAB on calcium mobilization;DAP12 signaling;Signaling by PDGF;CD28 co-stimulation;VEGFA-VEGFR2 Pathway 
                                               hsa04010;hsa04012;hsa04014;hsa04015;hsa04022;hsa04024;hsa04062;hsa04066;hsa04068;hsa04071;hsa04072;hsa04150;hsa04151;hsa04152;hsa04210;hsa04211;hsa04213;hsa04261;hsa04370;hsa04380;hsa04510;hsa04530;hsa04550;hsa04611;hsa04620;hsa04630;hsa04660;hsa04662;hsa04664;hsa04666;hsa04668;hsa04722;hsa04725;hsa04728;hsa04910;hsa04914;hsa04915;hsa04917;hsa04919;hsa04920;hsa04922;hsa04923;hsa04931;hsa04932;hsa04933;hsa04973;hsa05142;hsa05145;hsa05152;hsa05160;hsa05161;hsa05162;hsa05164;hsa05166;hsa05169;hsa05200;hsa05205;hsa05210;hsa05211;hsa05212;hsa05213;hsa05214;hsa05215;hsa05218;hsa05220;hsa05221;hsa05222;hsa05223;hsa05230;hsa05231 
                                               AKT3 
                                               NA 
                                               slateblue1 
                                               16 
                                               -17.7777777777778 
                                             
                                             
                                               AKT2-mRNA 
                                               0.479 
                                               0.12 
                                               0.245 
                                               0.713 
                                               1.39 
                                               1.18 
                                               1.64 
                                               0.000285 
                                               0.00263 
                                               lm.nb 
                                               Activation of BAD and translocation to mitochondria , CD28 co-stimulation, DAP12 signaling, Deactivation of the beta-catenin transactivating complex, Downstream signal transduction, G beta.CL.gamma signalling through PI3Kgamma, GAB1 signalosome, GPVI-mediated activation cascade, IRS-related events triggered by IGF1R, PI3K Cascade, PI3KAKT activation, PIP3 activates AKT signaling, Role of LAT2NTALLAB on calcium mobilization, Signaling by ERBB2, Signaling by PDGF, Signaling by SCF-KIT, TP53 Regulates Metabolic Genes, VEGFA-VEGFR2 Pathway, VEGFR2 mediated vascular permeability 
                                               SEBELIUS_2_C4636 
                                               AKT2 
                                               mRNA 
                                               0 
                                               
                                               
                                               TP53 Regulates Metabolic Genes;PDE3B signalling;AKT phosphorylates targets in the cytosol;Negative regulation of the PI3KAKT network;Deactivation of the beta-catenin transactivating complex;Activation of BAD and translocation to mitochondria ;G beta.CL.gamma signalling through PI3Kgamma;AKT phosphorylates targets in the nucleus;VEGFR2 mediated vascular permeability;GPVI-mediated activation cascade;Signaling by ERBB2;PI3K Cascade;IRS-related events triggered by IGF1R;mTOR signalling;PIP3 activates AKT signaling;Downstream signal transduction;GAB1 signalosome;Signaling by SCF-KIT;PI3KAKT activation;Role of LAT2NTALLAB on calcium mobilization;DAP12 signaling;Signaling by PDGF;CD28 co-stimulation;VEGFA-VEGFR2 Pathway 
                                               hsa04010;hsa04012;hsa04014;hsa04015;hsa04022;hsa04024;hsa04062;hsa04066;hsa04068;hsa04071;hsa04072;hsa04150;hsa04151;hsa04152;hsa04210;hsa04211;hsa04213;hsa04261;hsa04370;hsa04380;hsa04510;hsa04530;hsa04550;hsa04611;hsa04620;hsa04630;hsa04660;hsa04662;hsa04664;hsa04666;hsa04668;hsa04722;hsa04725;hsa04728;hsa04910;hsa04914;hsa04915;hsa04917;hsa04919;hsa04920;hsa04922;hsa04923;hsa04931;hsa04932;hsa04933;hsa04973;hsa05142;hsa05145;hsa05152;hsa05160;hsa05161;hsa05162;hsa05164;hsa05166;hsa05169;hsa05200;hsa05205;hsa05210;hsa05211;hsa05212;hsa05213;hsa05214;hsa05215;hsa05218;hsa05220;hsa05221;hsa05222;hsa05223;hsa05230;hsa05231 
                                               AKT2 
                                               NA 
                                               slateblue1 
                                               16 
                                               3.99166666666667 
                                             
                                             
                                               AKT1-mRNA 
                                               -0.919 
                                               0.234 
                                               -1.38 
                                               -0.461 
                                               0.529 
                                               0.385 
                                               0.726 
                                               0.000354 
                                               0.00322 
                                               lm.nb 
                                               Activation of BAD and translocation to mitochondria , CD28 co-stimulation, DAP12 signaling, Deactivation of the beta-catenin transactivating complex, Downstream signal transduction, G beta.CL.gamma signalling through PI3Kgamma, GAB1 signalosome, GPVI-mediated activation cascade, IRS-related events triggered by IGF1R, PI3K Cascade, PI3KAKT activation, PIP3 activates AKT signaling, Role of LAT2NTALLAB on calcium mobilization, Signaling by ERBB2, Signaling by PDGF, Signaling by SCF-KIT, TP53 Regulates Metabolic Genes, VEGFA-VEGFR2 Pathway, VEGFR2 mediated vascular permeability 
                                               SEBELIUS_2_C4636 
                                               AKT1 
                                               mRNA 
                                               0 
                                               
                                               
                                               TP53 Regulates Metabolic Genes;Deactivation of the beta-catenin transactivating complex;Activation of BAD and translocation to mitochondria ;eNOS activation;mTOR signalling;G beta.CL.gamma signalling through PI3Kgamma;KSRP (KHSRP) binds and destabilizes mRNA;AKT phosphorylates targets in the nucleus;VEGFR2 mediated vascular permeability;GPVI-mediated activation cascade;Tetrahydrobiopterin (BH4) synthesis# recycling# salvage and regulation;Butyrate Response Factor 1 (BRF1) binds and destabilizes mRNA;Signaling by ERBB2;PIP3 activates AKT signaling;Downstream signal transduction;GAB1 signalosome;Signaling by SCF-KIT;PI3KAKT activation;Role of LAT2NTALLAB on calcium mobilization;DAP12 signaling;Signaling by PDGF;CD28 co-stimulation;PI3K Cascade;IRS-related events triggered by IGF1R;VEGFA-VEGFR2 Pathway 
                                               hsa04010;hsa04012;hsa04014;hsa04015;hsa04022;hsa04024;hsa04062;hsa04066;hsa04068;hsa04071;hsa04072;hsa04150;hsa04151;hsa04152;hsa04210;hsa04211;hsa04213;hsa04261;hsa04370;hsa04380;hsa04510;hsa04530;hsa04550;hsa04611;hsa04620;hsa04630;hsa04660;hsa04662;hsa04664;hsa04666;hsa04668;hsa04722;hsa04725;hsa04728;hsa04910;hsa04914;hsa04915;hsa04917;hsa04919;hsa04920;hsa04922;hsa04923;hsa04931;hsa04932;hsa04933;hsa04973;hsa05142;hsa05145;hsa05152;hsa05160;hsa05161;hsa05162;hsa05164;hsa05166;hsa05169;hsa05200;hsa05205;hsa05210;hsa05211;hsa05212;hsa05213;hsa05214;hsa05215;hsa05218;hsa05220;hsa05221;hsa05222;hsa05223;hsa05230;hsa05231 
                                               AKT1 
                                               NA 
                                               slateblue1 
                                               16 
                                               -3.92735042735043 
                                             
                                             
                                               BAD-mRNA 
                                               -0.386 
                                               0.294 
                                               -0.963 
                                               0.192 
                                               0.765 
                                               0.513 
                                               1.14 
                                               0.198 
                                               1 
                                               lm.nb 
                                               Activation of BAD and translocation to mitochondria , DAP12 signaling, Downstream signal transduction, GAB1 signalosome, NRAGE signals death through JNK, PI3KAKT activation, PIP3 activates AKT signaling, Role of LAT2NTALLAB on calcium mobilization, Signaling by ERBB2, Signaling by PDGF, Signaling by SCF-KIT 
                                               SEBELIUS_2_C4636 
                                               BAD 
                                               mRNA 
                                               0 
                                               
                                               
                                               Activation of BAD and translocation to mitochondria ;NRAGE signals death through JNK;Constitutive Signaling by AKT1 E17K in Cancer;AKT phosphorylates targets in the cytosol;PIP3 activates AKT signaling;Downstream signal transduction;GAB1 signalosome;Signaling by SCF-KIT;PI3KAKT activation;Role of LAT2NTALLAB on calcium mobilization;DAP12 signaling;Signaling by PDGF;Signaling by ERBB2 
                                               hsa04012;hsa04014;hsa04022;hsa04024;hsa04151;hsa04210;hsa04370;hsa04510;hsa04722;hsa04910;hsa04919;hsa05010;hsa05145;hsa05152;hsa05160;hsa05161;hsa05200;hsa05203;hsa05210;hsa05212;hsa05213;hsa05215;hsa05218;hsa05220;hsa05221;hsa05223 
                                               BAD 
                                               NA 
                                               azure4 
                                               1 
                                               -1.31292517006803 
                                             
                                             
                                               BCL2-mRNA 
                                               0.0304 
                                               2.3 
                                               -4.48 
                                               4.54 
                                               1.02 
                                               0.0447 
                                               23.3 
                                               0.99 
                                               1 
                                               Wald 
                                               Activation of BAD and translocation to mitochondria  
                                               SEBELIUS_2_C4636 
                                               BCL2 
                                               mRNA 
                                               0 
                                               
                                               
                                               Activation of BAD and translocation to mitochondria  
                                               hsa04064;hsa04066;hsa04071;hsa04141;hsa04151;hsa04210;hsa04261;hsa04510;hsa04630;hsa04722;hsa04725;hsa04933;hsa05145;hsa05152;hsa05161;hsa05169;hsa05200;hsa05206;hsa05210;hsa05215;hsa05222 
                                               BCL2 
                                               NA 
                                               azure4 
                                               1 
                                               0.0132173913043478 
                                             
                                           
                                           DE Results - Activation of BAD and translocation to mitochondria  genes - Line: Caco2 vs.Fibro  Table displaying each sample's global significance scores and directed global significance scores as defined in the heatmaps above. The global significance score is calculated as the square root of the mean squared t-statistic for the genes in a gene set, with t-statistics coming from the linear regression underlying our differential expression analysis. The directed global significance score is calculated as the square root of the mean signed squared t-statistic for the genes in a gene set, with t-statistics coming from the linear regression underlying our differential expression analysis.   
                                      
                                       
                                  
                                
                                
                                  
                                       
                                      
                                           Volcano Plot: Line: HEPG2 vs.Fibro   More Plot Information                Volcano Plot: Line: HEPG2 vs.Fibro  Volcano plot displaying each gene's -log10(p-value) and log2 fold change for the selected covariate. Highly statistically significant genes fall at the top of the plot, and highly differentially expressed genes fall to either side. Genes within the selected gene set are highlighted in orange. Horizontal lines indicate various False Discovery Rate (FDR) thresholds.   
                                           DE Results - Activation of BAD and translocation to mitochondria  genes - Line: HEPG2 vs.Fibro   More Plot Information      Download CSV Data     
                                           
                                             
                                             
                                           
                                           
                                             
                                               Probe Label 
                                               Log2 fold change 
                                               std error (log2) 
                                               Lower confidence limit (log2) 
                                               Upper confidence limit (log2) 
                                               Linear fold change 
                                               Lower confidence limit (linear) 
                                               Upper confidence limit (linear) 
                                               P-value 
                                               BY.p.value 
                                               method 
                                               Gene.sets 
                                               Codeset.Name 
                                               Probe.Label 
                                               Analyte.Type 
                                               Is.Control 
                                               Control.Type 
                                               Related.Probes 
                                               Probe.Annotation 
                                               KEGG.Pathways 
                                               Official.Gene.Name 
                                               Control.Conc 
                                               volcanocol 
                                               volcanopch 
                                               tstats.all 
                                             
                                           
                                           
                                             
                                               AKT3-mRNA 
                                               -7.38 
                                               0.252 
                                               -7.87 
                                               -6.89 
                                               0.006 
                                               0.00426 
                                               0.00845 
                                               3.23e-27 
                                               1.15e-24 
                                               lm.nb 
                                               Activation of BAD and translocation to mitochondria , CD28 co-stimulation, CD28 dependent PI3KAkt signaling, DAP12 signaling, Downstream signal transduction, G beta.CL.gamma signalling through PI3Kgamma, GAB1 signalosome, GPVI-mediated activation cascade, PI3KAKT activation, PIP3 activates AKT signaling, Role of LAT2NTALLAB on calcium mobilization, Signaling by ERBB2, Signaling by PDGF, Signaling by SCF-KIT, TP53 Regulates Metabolic Genes, VEGFA-VEGFR2 Pathway, VEGFR2 mediated vascular permeability 
                                               SEBELIUS_2_C4636 
                                               AKT3 
                                               mRNA 
                                               0 
                                               
                                               
                                               TP53 Regulates Metabolic Genes;CD28 dependent PI3KAkt signaling;AKT phosphorylates targets in the cytosol;Negative regulation of the PI3KAKT network;Activation of BAD and translocation to mitochondria ;G beta.CL.gamma signalling through PI3Kgamma;AKT phosphorylates targets in the nucleus;VEGFR2 mediated vascular permeability;GPVI-mediated activation cascade;Signaling by ERBB2;PIP3 activates AKT signaling;Downstream signal transduction;GAB1 signalosome;Signaling by SCF-KIT;PI3KAKT activation;Role of LAT2NTALLAB on calcium mobilization;DAP12 signaling;Signaling by PDGF;CD28 co-stimulation;VEGFA-VEGFR2 Pathway 
                                               hsa04010;hsa04012;hsa04014;hsa04015;hsa04022;hsa04024;hsa04062;hsa04066;hsa04068;hsa04071;hsa04072;hsa04150;hsa04151;hsa04152;hsa04210;hsa04211;hsa04213;hsa04261;hsa04370;hsa04380;hsa04510;hsa04530;hsa04550;hsa04611;hsa04620;hsa04630;hsa04660;hsa04662;hsa04664;hsa04666;hsa04668;hsa04722;hsa04725;hsa04728;hsa04910;hsa04914;hsa04915;hsa04917;hsa04919;hsa04920;hsa04922;hsa04923;hsa04931;hsa04932;hsa04933;hsa04973;hsa05142;hsa05145;hsa05152;hsa05160;hsa05161;hsa05162;hsa05164;hsa05166;hsa05169;hsa05200;hsa05205;hsa05210;hsa05211;hsa05212;hsa05213;hsa05214;hsa05215;hsa05218;hsa05220;hsa05221;hsa05222;hsa05223;hsa05230;hsa05231 
                                               AKT3 
                                               NA 
                                               slateblue1 
                                               16 
                                               -29.2857142857143 
                                             
                                             
                                               AKT2-mRNA 
                                               0.873 
                                               0.131 
                                               0.617 
                                               1.13 
                                               1.83 
                                               1.53 
                                               2.19 
                                               7.75e-08 
                                               1.06e-06 
                                               lm.nb 
                                               Activation of BAD and translocation to mitochondria , CD28 co-stimulation, DAP12 signaling, Deactivation of the beta-catenin transactivating complex, Downstream signal transduction, G beta.CL.gamma signalling through PI3Kgamma, GAB1 signalosome, GPVI-mediated activation cascade, IRS-related events triggered by IGF1R, PI3K Cascade, PI3KAKT activation, PIP3 activates AKT signaling, Role of LAT2NTALLAB on calcium mobilization, Signaling by ERBB2, Signaling by PDGF, Signaling by SCF-KIT, TP53 Regulates Metabolic Genes, VEGFA-VEGFR2 Pathway, VEGFR2 mediated vascular permeability 
                                               SEBELIUS_2_C4636 
                                               AKT2 
                                               mRNA 
                                               0 
                                               
                                               
                                               TP53 Regulates Metabolic Genes;PDE3B signalling;AKT phosphorylates targets in the cytosol;Negative regulation of the PI3KAKT network;Deactivation of the beta-catenin transactivating complex;Activation of BAD and translocation to mitochondria ;G beta.CL.gamma signalling through PI3Kgamma;AKT phosphorylates targets in the nucleus;VEGFR2 mediated vascular permeability;GPVI-mediated activation cascade;Signaling by ERBB2;PI3K Cascade;IRS-related events triggered by IGF1R;mTOR signalling;PIP3 activates AKT signaling;Downstream signal transduction;GAB1 signalosome;Signaling by SCF-KIT;PI3KAKT activation;Role of LAT2NTALLAB on calcium mobilization;DAP12 signaling;Signaling by PDGF;CD28 co-stimulation;VEGFA-VEGFR2 Pathway 
                                               hsa04010;hsa04012;hsa04014;hsa04015;hsa04022;hsa04024;hsa04062;hsa04066;hsa04068;hsa04071;hsa04072;hsa04150;hsa04151;hsa04152;hsa04210;hsa04211;hsa04213;hsa04261;hsa04370;hsa04380;hsa04510;hsa04530;hsa04550;hsa04611;hsa04620;hsa04630;hsa04660;hsa04662;hsa04664;hsa04666;hsa04668;hsa04722;hsa04725;hsa04728;hsa04910;hsa04914;hsa04915;hsa04917;hsa04919;hsa04920;hsa04922;hsa04923;hsa04931;hsa04932;hsa04933;hsa04973;hsa05142;hsa05145;hsa05152;hsa05160;hsa05161;hsa05162;hsa05164;hsa05166;hsa05169;hsa05200;hsa05205;hsa05210;hsa05211;hsa05212;hsa05213;hsa05214;hsa05215;hsa05218;hsa05220;hsa05221;hsa05222;hsa05223;hsa05230;hsa05231 
                                               AKT2 
                                               NA 
                                               slateblue1 
                                               16 
                                               6.66412213740458 
                                             
                                             
                                               BAD-mRNA 
                                               -0.997 
                                               0.325 
                                               -1.63 
                                               -0.359 
                                               0.501 
                                               0.322 
                                               0.779 
                                               0.00404 
                                               0.0352 
                                               lm.nb 
                                               Activation of BAD and translocation to mitochondria , DAP12 signaling, Downstream signal transduction, GAB1 signalosome, NRAGE signals death through JNK, PI3KAKT activation, PIP3 activates AKT signaling, Role of LAT2NTALLAB on calcium mobilization, Signaling by ERBB2, Signaling by PDGF, Signaling by SCF-KIT 
                                               SEBELIUS_2_C4636 
                                               BAD 
                                               mRNA 
                                               0 
                                               
                                               
                                               Activation of BAD and translocation to mitochondria ;NRAGE signals death through JNK;Constitutive Signaling by AKT1 E17K in Cancer;AKT phosphorylates targets in the cytosol;PIP3 activates AKT signaling;Downstream signal transduction;GAB1 signalosome;Signaling by SCF-KIT;PI3KAKT activation;Role of LAT2NTALLAB on calcium mobilization;DAP12 signaling;Signaling by PDGF;Signaling by ERBB2 
                                               hsa04012;hsa04014;hsa04022;hsa04024;hsa04151;hsa04210;hsa04370;hsa04510;hsa04722;hsa04910;hsa04919;hsa05010;hsa05145;hsa05152;hsa05160;hsa05161;hsa05200;hsa05203;hsa05210;hsa05212;hsa05213;hsa05215;hsa05218;hsa05220;hsa05221;hsa05223 
                                               BAD 
                                               NA 
                                               slateblue2 
                                               16 
                                               -3.06769230769231 
                                             
                                             
                                               AKT1-mRNA 
                                               0.141 
                                               0.256 
                                               -0.361 
                                               0.643 
                                               1.1 
                                               0.779 
                                               1.56 
                                               0.585 
                                               1 
                                               lm.nb 
                                               Activation of BAD and translocation to mitochondria , CD28 co-stimulation, DAP12 signaling, Deactivation of the beta-catenin transactivating complex, Downstream signal transduction, G beta.CL.gamma signalling through PI3Kgamma, GAB1 signalosome, GPVI-mediated activation cascade, IRS-related events triggered by IGF1R, PI3K Cascade, PI3KAKT activation, PIP3 activates AKT signaling, Role of LAT2NTALLAB on calcium mobilization, Signaling by ERBB2, Signaling by PDGF, Signaling by SCF-KIT, TP53 Regulates Metabolic Genes, VEGFA-VEGFR2 Pathway, VEGFR2 mediated vascular permeability 
                                               SEBELIUS_2_C4636 
                                               AKT1 
                                               mRNA 
                                               0 
                                               
                                               
                                               TP53 Regulates Metabolic Genes;Deactivation of the beta-catenin transactivating complex;Activation of BAD and translocation to mitochondria ;eNOS activation;mTOR signalling;G beta.CL.gamma signalling through PI3Kgamma;KSRP (KHSRP) binds and destabilizes mRNA;AKT phosphorylates targets in the nucleus;VEGFR2 mediated vascular permeability;GPVI-mediated activation cascade;Tetrahydrobiopterin (BH4) synthesis# recycling# salvage and regulation;Butyrate Response Factor 1 (BRF1) binds and destabilizes mRNA;Signaling by ERBB2;PIP3 activates AKT signaling;Downstream signal transduction;GAB1 signalosome;Signaling by SCF-KIT;PI3KAKT activation;Role of LAT2NTALLAB on calcium mobilization;DAP12 signaling;Signaling by PDGF;CD28 co-stimulation;PI3K Cascade;IRS-related events triggered by IGF1R;VEGFA-VEGFR2 Pathway 
                                               hsa04010;hsa04012;hsa04014;hsa04015;hsa04022;hsa04024;hsa04062;hsa04066;hsa04068;hsa04071;hsa04072;hsa04150;hsa04151;hsa04152;hsa04210;hsa04211;hsa04213;hsa04261;hsa04370;hsa04380;hsa04510;hsa04530;hsa04550;hsa04611;hsa04620;hsa04630;hsa04660;hsa04662;hsa04664;hsa04666;hsa04668;hsa04722;hsa04725;hsa04728;hsa04910;hsa04914;hsa04915;hsa04917;hsa04919;hsa04920;hsa04922;hsa04923;hsa04931;hsa04932;hsa04933;hsa04973;hsa05142;hsa05145;hsa05152;hsa05160;hsa05161;hsa05162;hsa05164;hsa05166;hsa05169;hsa05200;hsa05205;hsa05210;hsa05211;hsa05212;hsa05213;hsa05214;hsa05215;hsa05218;hsa05220;hsa05221;hsa05222;hsa05223;hsa05230;hsa05231 
                                               AKT1 
                                               NA 
                                               azure4 
                                               1 
                                               0.55078125 
                                             
                                             
                                               BCL2-mRNA 
                                               -0.895 
                                               2.49 
                                               -5.77 
                                               3.98 
                                               0.538 
                                               0.0183 
                                               15.8 
                                               0.721 
                                               1 
                                               Wald 
                                               Activation of BAD and translocation to mitochondria  
                                               SEBELIUS_2_C4636 
                                               BCL2 
                                               mRNA 
                                               0 
                                               
                                               
                                               Activation of BAD and translocation to mitochondria  
                                               hsa04064;hsa04066;hsa04071;hsa04141;hsa04151;hsa04210;hsa04261;hsa04510;hsa04630;hsa04722;hsa04725;hsa04933;hsa05145;hsa05152;hsa05161;hsa05169;hsa05200;hsa05206;hsa05210;hsa05215;hsa05222 
                                               BCL2 
                                               NA 
                                               azure4 
                                               1 
                                               -0.359437751004016 
                                             
                                           
                                           DE Results - Activation of BAD and translocation to mitochondria  genes - Line: HEPG2 vs.Fibro  Table displaying each sample's global significance scores and directed global significance scores as defined in the heatmaps above. The global significance score is calculated as the square root of the mean squared t-statistic for the genes in a gene set, with t-statistics coming from the linear regression underlying our differential expression analysis. The directed global significance score is calculated as the square root of the mean signed squared t-statistic for the genes in a gene set, with t-statistics coming from the linear regression underlying our differential expression analysis.   
                                      
                                       
                                  
                                
                                
                                  
                                       
                                      
                                           Volcano Plot: Line: HT29 vs.Fibro   More Plot Information                Volcano Plot: Line: HT29 vs.Fibro  Volcano plot displaying each gene's -log10(p-value) and log2 fold change for the selected covariate. Highly statistically significant genes fall at the top of the plot, and highly differentially expressed genes fall to either side. Genes within the selected gene set are highlighted in orange. Horizontal lines indicate various False Discovery Rate (FDR) thresholds.   
                                           DE Results - Activation of BAD and translocation to mitochondria  genes - Line: HT29 vs.Fibro   More Plot Information      Download CSV Data     
                                           
                                             
                                             
                                           
                                           
                                             
                                               Probe Label 
                                               Log2 fold change 
                                               std error (log2) 
                                               Lower confidence limit (log2) 
                                               Upper confidence limit (log2) 
                                               Linear fold change 
                                               Lower confidence limit (linear) 
                                               Upper confidence limit (linear) 
                                               P-value 
                                               BY.p.value 
                                               method 
                                               Gene.sets 
                                               Codeset.Name 
                                               Probe.Label 
                                               Analyte.Type 
                                               Is.Control 
                                               Control.Type 
                                               Related.Probes 
                                               Probe.Annotation 
                                               KEGG.Pathways 
                                               Official.Gene.Name 
                                               Control.Conc 
                                               volcanocol 
                                               volcanopch 
                                               tstats.all 
                                             
                                           
                                           
                                             
                                               AKT3-mRNA 
                                               -8.38 
                                               0.265 
                                               -8.9 
                                               -7.86 
                                               0.003 
                                               0.00209 
                                               0.0043 
                                               2.3e-28 
                                               4.09e-26 
                                               lm.nb 
                                               Activation of BAD and translocation to mitochondria , CD28 co-stimulation, CD28 dependent PI3KAkt signaling, DAP12 signaling, Downstream signal transduction, G beta.CL.gamma signalling through PI3Kgamma, GAB1 signalosome, GPVI-mediated activation cascade, PI3KAKT activation, PIP3 activates AKT signaling, Role of LAT2NTALLAB on calcium mobilization, Signaling by ERBB2, Signaling by PDGF, Signaling by SCF-KIT, TP53 Regulates Metabolic Genes, VEGFA-VEGFR2 Pathway, VEGFR2 mediated vascular permeability 
                                               SEBELIUS_2_C4636 
                                               AKT3 
                                               mRNA 
                                               0 
                                               
                                               
                                               TP53 Regulates Metabolic Genes;CD28 dependent PI3KAkt signaling;AKT phosphorylates targets in the cytosol;Negative regulation of the PI3KAKT network;Activation of BAD and translocation to mitochondria ;G beta.CL.gamma signalling through PI3Kgamma;AKT phosphorylates targets in the nucleus;VEGFR2 mediated vascular permeability;GPVI-mediated activation cascade;Signaling by ERBB2;PIP3 activates AKT signaling;Downstream signal transduction;GAB1 signalosome;Signaling by SCF-KIT;PI3KAKT activation;Role of LAT2NTALLAB on calcium mobilization;DAP12 signaling;Signaling by PDGF;CD28 co-stimulation;VEGFA-VEGFR2 Pathway 
                                               hsa04010;hsa04012;hsa04014;hsa04015;hsa04022;hsa04024;hsa04062;hsa04066;hsa04068;hsa04071;hsa04072;hsa04150;hsa04151;hsa04152;hsa04210;hsa04211;hsa04213;hsa04261;hsa04370;hsa04380;hsa04510;hsa04530;hsa04550;hsa04611;hsa04620;hsa04630;hsa04660;hsa04662;hsa04664;hsa04666;hsa04668;hsa04722;hsa04725;hsa04728;hsa04910;hsa04914;hsa04915;hsa04917;hsa04919;hsa04920;hsa04922;hsa04923;hsa04931;hsa04932;hsa04933;hsa04973;hsa05142;hsa05145;hsa05152;hsa05160;hsa05161;hsa05162;hsa05164;hsa05166;hsa05169;hsa05200;hsa05205;hsa05210;hsa05211;hsa05212;hsa05213;hsa05214;hsa05215;hsa05218;hsa05220;hsa05221;hsa05222;hsa05223;hsa05230;hsa05231 
                                               AKT3 
                                               NA 
                                               slateblue1 
                                               16 
                                               -31.622641509434 
                                             
                                             
                                               AKT2-mRNA 
                                               1.35 
                                               0.123 
                                               1.11 
                                               1.59 
                                               2.55 
                                               2.15 
                                               3.01 
                                               3.72e-13 
                                               6.37e-12 
                                               lm.nb 
                                               Activation of BAD and translocation to mitochondria , CD28 co-stimulation, DAP12 signaling, Deactivation of the beta-catenin transactivating complex, Downstream signal transduction, G beta.CL.gamma signalling through PI3Kgamma, GAB1 signalosome, GPVI-mediated activation cascade, IRS-related events triggered by IGF1R, PI3K Cascade, PI3KAKT activation, PIP3 activates AKT signaling, Role of LAT2NTALLAB on calcium mobilization, Signaling by ERBB2, Signaling by PDGF, Signaling by SCF-KIT, TP53 Regulates Metabolic Genes, VEGFA-VEGFR2 Pathway, VEGFR2 mediated vascular permeability 
                                               SEBELIUS_2_C4636 
                                               AKT2 
                                               mRNA 
                                               0 
                                               
                                               
                                               TP53 Regulates Metabolic Genes;PDE3B signalling;AKT phosphorylates targets in the cytosol;Negative regulation of the PI3KAKT network;Deactivation of the beta-catenin transactivating complex;Activation of BAD and translocation to mitochondria ;G beta.CL.gamma signalling through PI3Kgamma;AKT phosphorylates targets in the nucleus;VEGFR2 mediated vascular permeability;GPVI-mediated activation cascade;Signaling by ERBB2;PI3K Cascade;IRS-related events triggered by IGF1R;mTOR signalling;PIP3 activates AKT signaling;Downstream signal transduction;GAB1 signalosome;Signaling by SCF-KIT;PI3KAKT activation;Role of LAT2NTALLAB on calcium mobilization;DAP12 signaling;Signaling by PDGF;CD28 co-stimulation;VEGFA-VEGFR2 Pathway 
                                               hsa04010;hsa04012;hsa04014;hsa04015;hsa04022;hsa04024;hsa04062;hsa04066;hsa04068;hsa04071;hsa04072;hsa04150;hsa04151;hsa04152;hsa04210;hsa04211;hsa04213;hsa04261;hsa04370;hsa04380;hsa04510;hsa04530;hsa04550;hsa04611;hsa04620;hsa04630;hsa04660;hsa04662;hsa04664;hsa04666;hsa04668;hsa04722;hsa04725;hsa04728;hsa04910;hsa04914;hsa04915;hsa04917;hsa04919;hsa04920;hsa04922;hsa04923;hsa04931;hsa04932;hsa04933;hsa04973;hsa05142;hsa05145;hsa05152;hsa05160;hsa05161;hsa05162;hsa05164;hsa05166;hsa05169;hsa05200;hsa05205;hsa05210;hsa05211;hsa05212;hsa05213;hsa05214;hsa05215;hsa05218;hsa05220;hsa05221;hsa05222;hsa05223;hsa05230;hsa05231 
                                               AKT2 
                                               NA 
                                               slateblue1 
                                               16 
                                               10.9756097560976 
                                             
                                             
                                               BCL2-mRNA 
                                               -1.98 
                                               2.36 
                                               -6.61 
                                               2.64 
                                               0.253 
                                               0.0102 
                                               6.24 
                                               0.406 
                                               1 
                                               Wald 
                                               Activation of BAD and translocation to mitochondria  
                                               SEBELIUS_2_C4636 
                                               BCL2 
                                               mRNA 
                                               0 
                                               
                                               
                                               Activation of BAD and translocation to mitochondria  
                                               hsa04064;hsa04066;hsa04071;hsa04141;hsa04151;hsa04210;hsa04261;hsa04510;hsa04630;hsa04722;hsa04725;hsa04933;hsa05145;hsa05152;hsa05161;hsa05169;hsa05200;hsa05206;hsa05210;hsa05215;hsa05222 
                                               BCL2 
                                               NA 
                                               azure4 
                                               1 
                                               -0.838983050847458 
                                             
                                             
                                               BAD-mRNA 
                                               0.175 
                                               0.304 
                                               -0.42 
                                               0.771 
                                               1.13 
                                               0.747 
                                               1.71 
                                               0.568 
                                               1 
                                               lm.nb 
                                               Activation of BAD and translocation to mitochondria , DAP12 signaling, Downstream signal transduction, GAB1 signalosome, NRAGE signals death through JNK, PI3KAKT activation, PIP3 activates AKT signaling, Role of LAT2NTALLAB on calcium mobilization, Signaling by ERBB2, Signaling by PDGF, Signaling by SCF-KIT 
                                               SEBELIUS_2_C4636 
                                               BAD 
                                               mRNA 
                                               0 
                                               
                                               
                                               Activation of BAD and translocation to mitochondria ;NRAGE signals death through JNK;Constitutive Signaling by AKT1 E17K in Cancer;AKT phosphorylates targets in the cytosol;PIP3 activates AKT signaling;Downstream signal transduction;GAB1 signalosome;Signaling by SCF-KIT;PI3KAKT activation;Role of LAT2NTALLAB on calcium mobilization;DAP12 signaling;Signaling by PDGF;Signaling by ERBB2 
                                               hsa04012;hsa04014;hsa04022;hsa04024;hsa04151;hsa04210;hsa04370;hsa04510;hsa04722;hsa04910;hsa04919;hsa05010;hsa05145;hsa05152;hsa05160;hsa05161;hsa05200;hsa05203;hsa05210;hsa05212;hsa05213;hsa05215;hsa05218;hsa05220;hsa05221;hsa05223 
                                               BAD 
                                               NA 
                                               azure4 
                                               1 
                                               0.575657894736842 
                                             
                                             
                                               AKT1-mRNA 
                                               0.00517 
                                               0.241 
                                               -0.468 
                                               0.478 
                                               1 
                                               0.723 
                                               1.39 
                                               0.983 
                                               1 
                                               lm.nb 
                                               Activation of BAD and translocation to mitochondria , CD28 co-stimulation, DAP12 signaling, Deactivation of the beta-catenin transactivating complex, Downstream signal transduction, G beta.CL.gamma signalling through PI3Kgamma, GAB1 signalosome, GPVI-mediated activation cascade, IRS-related events triggered by IGF1R, PI3K Cascade, PI3KAKT activation, PIP3 activates AKT signaling, Role of LAT2NTALLAB on calcium mobilization, Signaling by ERBB2, Signaling by PDGF, Signaling by SCF-KIT, TP53 Regulates Metabolic Genes, VEGFA-VEGFR2 Pathway, VEGFR2 mediated vascular permeability 
                                               SEBELIUS_2_C4636 
                                               AKT1 
                                               mRNA 
                                               0 
                                               
                                               
                                               TP53 Regulates Metabolic Genes;Deactivation of the beta-catenin transactivating complex;Activation of BAD and translocation to mitochondria ;eNOS activation;mTOR signalling;G beta.CL.gamma signalling through PI3Kgamma;KSRP (KHSRP) binds and destabilizes mRNA;AKT phosphorylates targets in the nucleus;VEGFR2 mediated vascular permeability;GPVI-mediated activation cascade;Tetrahydrobiopterin (BH4) synthesis# recycling# salvage and regulation;Butyrate Response Factor 1 (BRF1) binds and destabilizes mRNA;Signaling by ERBB2;PIP3 activates AKT signaling;Downstream signal transduction;GAB1 signalosome;Signaling by SCF-KIT;PI3KAKT activation;Role of LAT2NTALLAB on calcium mobilization;DAP12 signaling;Signaling by PDGF;CD28 co-stimulation;PI3K Cascade;IRS-related events triggered by IGF1R;VEGFA-VEGFR2 Pathway 
                                               hsa04010;hsa04012;hsa04014;hsa04015;hsa04022;hsa04024;hsa04062;hsa04066;hsa04068;hsa04071;hsa04072;hsa04150;hsa04151;hsa04152;hsa04210;hsa04211;hsa04213;hsa04261;hsa04370;hsa04380;hsa04510;hsa04530;hsa04550;hsa04611;hsa04620;hsa04630;hsa04660;hsa04662;hsa04664;hsa04666;hsa04668;hsa04722;hsa04725;hsa04728;hsa04910;hsa04914;hsa04915;hsa04917;hsa04919;hsa04920;hsa04922;hsa04923;hsa04931;hsa04932;hsa04933;hsa04973;hsa05142;hsa05145;hsa05152;hsa05160;hsa05161;hsa05162;hsa05164;hsa05166;hsa05169;hsa05200;hsa05205;hsa05210;hsa05211;hsa05212;hsa05213;hsa05214;hsa05215;hsa05218;hsa05220;hsa05221;hsa05222;hsa05223;hsa05230;hsa05231 
                                               AKT1 
                                               NA 
                                               azure4 
                                               1 
                                               0.0214522821576764 
                                             
                                           
                                           DE Results - Activation of BAD and translocation to mitochondria  genes - Line: HT29 vs.Fibro  Table displaying each sample's global significance scores and directed global significance scores as defined in the heatmaps above. The global significance score is calculated as the square root of the mean squared t-statistic for the genes in a gene set, with t-statistics coming from the linear regression underlying our differential expression analysis. The directed global significance score is calculated as the square root of the mean signed squared t-statistic for the genes in a gene set, with t-statistics coming from the linear regression underlying our differential expression analysis.   
                                      
                                       
                                  
                                
                              
                            
                           
                            
                          
                           
                      
                    
                    
                      
                           
                          
                               Day    Line     
                              
                                  Day  
                              
                               
                              
                                  Line: differential expression in Caco2 vs. baseline of Fibro  
                                  Line: differential expression in HEPG2 vs. baseline of Fibro  
                                  Line: differential expression in HT29 vs. baseline of Fibro  
                              
                              
                            
                              
                                
                                  
                                       
                                      
                                           Volcano Plot: Day   More Plot Information                Volcano Plot: Day  Volcano plot displaying each gene's -log10(p-value) and log2 fold change for the selected covariate. Highly statistically significant genes fall at the top of the plot, and highly differentially expressed genes fall to either side. Genes within the selected gene set are highlighted in orange. Horizontal lines indicate various False Discovery Rate (FDR) thresholds.   
                                           DE Results - Activation of Rac genes - Day   More Plot Information      Download CSV Data     
                                           
                                             
                                             
                                           
                                           
                                             
                                               Probe Label 
                                               Log2 fold change 
                                               std error (log2) 
                                               Lower confidence limit (log2) 
                                               Upper confidence limit (log2) 
                                               Linear fold change 
                                               Lower confidence limit (linear) 
                                               Upper confidence limit (linear) 
                                               P-value 
                                               BY.p.value 
                                               method 
                                               Gene.sets 
                                               Codeset.Name 
                                               Probe.Label 
                                               Analyte.Type 
                                               Is.Control 
                                               Control.Type 
                                               Related.Probes 
                                               Probe.Annotation 
                                               KEGG.Pathways 
                                               Official.Gene.Name 
                                               Control.Conc 
                                               volcanocol 
                                               volcanopch 
                                               tstats.all 
                                             
                                           
                                           
                                             
                                               PAK1-mRNA 
                                               0.128 
                                               0.0211 
                                               0.0869 
                                               0.169 
                                               1.09 
                                               1.06 
                                               1.12 
                                               4.84e-07 
                                               6.25e-05 
                                               lm.nb 
                                               Activation of Rac, CD209 (DC-SIGN) signaling, CD28 co-stimulation, CD28 dependent Vav1 pathway, EPHB-mediated forward signaling, Ephrin signaling, FCERI mediated MAPK activation, MAPK6MAPK4 signaling, Regulation of actin dynamics for phagocytic cup formation, RHO GTPases activate PAKs, Sema3A PAK dependent Axon repulsion, VEGFA-VEGFR2 Pathway, VEGFR2 mediated vascular permeability 
                                               SEBELIUS_2_C4636 
                                               PAK1 
                                               mRNA 
                                               0 
                                               
                                               
                                               EPHB-mediated forward signaling;Ephrin signaling;MAPK6MAPK4 signaling;DSCAM interactions;Activation of Rac;VEGFR2 mediated vascular permeability;RHO GTPases activate PAKs;Sema3A PAK dependent Axon repulsion;FCERI mediated MAPK activation;Regulation of actin dynamics for phagocytic cup formation;CD209 (DC-SIGN) signaling;CD28 dependent Vav1 pathway;VEGFA-VEGFR2 Pathway;CD28 co-stimulation 
                                               hsa04010;hsa04012;hsa04014;hsa04024;hsa04062;hsa04360;hsa04510;hsa04650;hsa04660;hsa04666;hsa04810;hsa05120;hsa05205;hsa05211 
                                               PAK1 
                                               NA 
                                               slateblue1 
                                               16 
                                               6.06635071090047 
                                             
                                             
                                               PAK2-mRNA 
                                               -0.0398 
                                               0.00994 
                                               -0.0593 
                                               -0.0203 
                                               0.973 
                                               0.96 
                                               0.986 
                                               0.000289 
                                               0.00836 
                                               loglinear 
                                               Activation of Rac, CD209 (DC-SIGN) signaling, CD28 co-stimulation, CD28 dependent Vav1 pathway, Ephrin signaling, RHO GTPases activate PAKs, Sema3A PAK dependent Axon repulsion, VEGFA-VEGFR2 Pathway, VEGFR2 mediated vascular permeability 
                                               SEBELIUS_2_C4636 
                                               PAK2 
                                               mRNA 
                                               0 
                                               
                                               
                                               Nef and signal transduction;CD28 dependent Vav1 pathway;Stimulation of the cell death response by PAK-2p34;Ephrin signaling;Activation of Rac;RHO GTPases activate PAKs;VEGFR2 mediated vascular permeability;Sema3A PAK dependent Axon repulsion;CD209 (DC-SIGN) signaling;VEGFA-VEGFR2 Pathway;CD28 co-stimulation 
                                               hsa04010;hsa04012;hsa04014;hsa04360;hsa04510;hsa04660;hsa04810;hsa05211 
                                               PAK2 
                                               NA 
                                               slateblue1 
                                               16 
                                               -4.00402414486922 
                                             
                                             
                                               SOS2-mRNA 
                                               0.0982 
                                               0.0316 
                                               0.0362 
                                               0.16 
                                               1.07 
                                               1.03 
                                               1.12 
                                               0.00365 
                                               0.0602 
                                               lm.nb 
                                               Activation of Rac, G alpha (1213) signalling events, NRAGE signals death through JNK, Rho GTPase cycle 
                                               SEBELIUS_2_C4636 
                                               SOS2 
                                               mRNA 
                                               0 
                                               
                                               
                                               Rho GTPase cycle;Activation of Rac;NRAGE signals death through JNK;G alpha (1213) signalling events 
                                               hsa04010;hsa04012;hsa04014;hsa04062;hsa04068;hsa04072;hsa04151;hsa04320;hsa04510;hsa04540;hsa04630;hsa04650;hsa04660;hsa04662;hsa04664;hsa04722;hsa04810;hsa04910;hsa04912;hsa04915;hsa04917;hsa05034;hsa05160;hsa05200;hsa05205;hsa05206;hsa05211;hsa05213;hsa05214;hsa05215;hsa05220;hsa05221;hsa05223;hsa05231 
                                               SOS2 
                                               NA 
                                               slateblue3 
                                               16 
                                               3.10759493670886 
                                             
                                             
                                               PAK4-mRNA 
                                               0.0288 
                                               0.0117 
                                               0.00588 
                                               0.0517 
                                               1.02 
                                               1 
                                               1.04 
                                               0.0186 
                                               0.237 
                                               loglinear 
                                               Activation of Rac 
                                               SEBELIUS_2_C4636 
                                               PAK4 
                                               mRNA 
                                               0 
                                               
                                               
                                               Activation of Rac 
                                               hsa04012;hsa04014;hsa04360;hsa04510;hsa04660;hsa04810;hsa05206;hsa05211 
                                               PAK4 
                                               NA 
                                               slateblue4 
                                               16 
                                               2.46153846153846 
                                             
                                             
                                               RAC1-mRNA 
                                               -0.000703 
                                               0.0123 
                                               -0.0248 
                                               0.0234 
                                               1 
                                               0.983 
                                               1.02 
                                               0.955 
                                               1 
                                               lm.nb 
                                               Activation of Rac, CD28 co-stimulation, CD28 dependent Vav1 pathway, DAP12 signaling, EPH-ephrin mediated repulsion of cells, Ephrin signaling, G alpha (1213) signalling events, GPVI-mediated activation cascade, MAPK6MAPK4 signaling, NRAGE signals death through JNK, PCPCE pathway, Regulation of actin dynamics for phagocytic cup formation, RHO GTPases activate CIT, RHO GTPases Activate Formins, RHO GTPases activate IQGAPs, RHO GTPases activate PAKs, RHO GTPases activate PKNs, RHO GTPases Activate WASPs and WAVEs, Sema3A PAK dependent Axon repulsion, Sema4D induced cell migration and growth-cone collapse, Signal transduction by L1, Signaling by SCF-KIT, VEGFA-VEGFR2 Pathway, VEGFR2 mediated vascular permeability 
                                               SEBELIUS_2_C4636 
                                               RAC1 
                                               mRNA 
                                               0 
                                               
                                               
                                               MAPK6MAPK4 signaling;Signaling by SCF-KIT;Inactivation of Cdc42 and Rac;DAP12 signaling;Ephrin signaling;EPH-ephrin mediated repulsion of cells;Activation of Rac;RHO GTPases Activate Formins;VEGFR2 mediated vascular permeability;RHO GTPases activate PAKs;Sema4D mediated inhibition of cell attachment and migration;G alpha (1213) signalling events;Regulation of actin dynamics for phagocytic cup formation;Sema4D induced cell migration and growth-cone collapse;Signal transduction by L1;RHO GTPases activate CIT;RHO GTPases Activate WASPs and WAVEs;Factors involved in megakaryocyte development and platelet production;RHO GTPases activate KTN1;CD28 dependent Vav1 pathway;DSCAM interactions;NRAGE signals death through JNK;GPVI-mediated activation cascade;Sema3A PAK dependent Axon repulsion;RHO GTPases activate PKNs;SEMA3A-Plexin repulsion signaling by inhibiting Integrin adhesion;RHO GTPases activate IQGAPs;PCPCE pathway;VEGFA-VEGFR2 Pathway;CD28 co-stimulation 
                                               hsa04010;hsa04014;hsa04015;hsa04024;hsa04062;hsa04071;hsa04145;hsa04151;hsa04310;hsa04360;hsa04370;hsa04380;hsa04510;hsa04520;hsa04620;hsa04650;hsa04662;hsa04664;hsa04666;hsa04670;hsa04722;hsa04810;hsa04932;hsa04933;hsa04972;hsa05100;hsa05120;hsa05131;hsa05132;hsa05200;hsa05203;hsa05205;hsa05210;hsa05211;hsa05212;hsa05231;hsa05416 
                                               RAC1 
                                               NA 
                                               azure4 
                                               1 
                                               -0.0571544715447154 
                                             
                                           
                                           DE Results - Activation of Rac genes - Day  Table displaying each sample's global significance scores and directed global significance scores as defined in the heatmaps above. The global significance score is calculated as the square root of the mean squared t-statistic for the genes in a gene set, with t-statistics coming from the linear regression underlying our differential expression analysis. The directed global significance score is calculated as the square root of the mean signed squared t-statistic for the genes in a gene set, with t-statistics coming from the linear regression underlying our differential expression analysis.   
                                      
                                       
                                  
                                
                              
                            
                            
                              
                                
                                  
                                       
                                      
                                           Volcano Plot: Line: Caco2 vs.Fibro   More Plot Information                Volcano Plot: Line: Caco2 vs.Fibro  Volcano plot displaying each gene's -log10(p-value) and log2 fold change for the selected covariate. Highly statistically significant genes fall at the top of the plot, and highly differentially expressed genes fall to either side. Genes within the selected gene set are highlighted in orange. Horizontal lines indicate various False Discovery Rate (FDR) thresholds.   
                                           DE Results - Activation of Rac genes - Line: Caco2 vs.Fibro   More Plot Information      Download CSV Data     
                                           
                                             
                                             
                                           
                                           
                                             
                                               Probe Label 
                                               Log2 fold change 
                                               std error (log2) 
                                               Lower confidence limit (log2) 
                                               Upper confidence limit (log2) 
                                               Linear fold change 
                                               Lower confidence limit (linear) 
                                               Upper confidence limit (linear) 
                                               P-value 
                                               BY.p.value 
                                               method 
                                               Gene.sets 
                                               Codeset.Name 
                                               Probe.Label 
                                               Analyte.Type 
                                               Is.Control 
                                               Control.Type 
                                               Related.Probes 
                                               Probe.Annotation 
                                               KEGG.Pathways 
                                               Official.Gene.Name 
                                               Control.Conc 
                                               volcanocol 
                                               volcanopch 
                                               tstats.all 
                                             
                                           
                                           
                                             
                                               PAK4-mRNA 
                                               2.18 
                                               0.0884 
                                               2 
                                               2.35 
                                               4.52 
                                               4.01 
                                               5.1 
                                               1.5e-24 
                                               4.25e-22 
                                               loglinear 
                                               Activation of Rac 
                                               SEBELIUS_2_C4636 
                                               PAK4 
                                               mRNA 
                                               0 
                                               
                                               
                                               Activation of Rac 
                                               hsa04012;hsa04014;hsa04360;hsa04510;hsa04660;hsa04810;hsa05206;hsa05211 
                                               PAK4 
                                               NA 
                                               slateblue1 
                                               16 
                                               24.6606334841629 
                                             
                                             
                                               PAK1-mRNA 
                                               1.92 
                                               0.159 
                                               1.61 
                                               2.23 
                                               3.79 
                                               3.05 
                                               4.7 
                                               2.23e-14 
                                               5.02e-13 
                                               lm.nb 
                                               Activation of Rac, CD209 (DC-SIGN) signaling, CD28 co-stimulation, CD28 dependent Vav1 pathway, EPHB-mediated forward signaling, Ephrin signaling, FCERI mediated MAPK activation, MAPK6MAPK4 signaling, Regulation of actin dynamics for phagocytic cup formation, RHO GTPases activate PAKs, Sema3A PAK dependent Axon repulsion, VEGFA-VEGFR2 Pathway, VEGFR2 mediated vascular permeability 
                                               SEBELIUS_2_C4636 
                                               PAK1 
                                               mRNA 
                                               0 
                                               
                                               
                                               EPHB-mediated forward signaling;Ephrin signaling;MAPK6MAPK4 signaling;DSCAM interactions;Activation of Rac;VEGFR2 mediated vascular permeability;RHO GTPases activate PAKs;Sema3A PAK dependent Axon repulsion;FCERI mediated MAPK activation;Regulation of actin dynamics for phagocytic cup formation;CD209 (DC-SIGN) signaling;CD28 dependent Vav1 pathway;VEGFA-VEGFR2 Pathway;CD28 co-stimulation 
                                               hsa04010;hsa04012;hsa04014;hsa04024;hsa04062;hsa04360;hsa04510;hsa04650;hsa04660;hsa04666;hsa04810;hsa05120;hsa05205;hsa05211 
                                               PAK1 
                                               NA 
                                               slateblue1 
                                               16 
                                               12.0754716981132 
                                             
                                             
                                               RAC1-mRNA 
                                               -0.326 
                                               0.093 
                                               -0.508 
                                               -0.144 
                                               0.798 
                                               0.703 
                                               0.905 
                                               0.00121 
                                               0.0104 
                                               lm.nb 
                                               Activation of Rac, CD28 co-stimulation, CD28 dependent Vav1 pathway, DAP12 signaling, EPH-ephrin mediated repulsion of cells, Ephrin signaling, G alpha (1213) signalling events, GPVI-mediated activation cascade, MAPK6MAPK4 signaling, NRAGE signals death through JNK, PCPCE pathway, Regulation of actin dynamics for phagocytic cup formation, RHO GTPases activate CIT, RHO GTPases Activate Formins, RHO GTPases activate IQGAPs, RHO GTPases activate PAKs, RHO GTPases activate PKNs, RHO GTPases Activate WASPs and WAVEs, Sema3A PAK dependent Axon repulsion, Sema4D induced cell migration and growth-cone collapse, Signal transduction by L1, Signaling by SCF-KIT, VEGFA-VEGFR2 Pathway, VEGFR2 mediated vascular permeability 
                                               SEBELIUS_2_C4636 
                                               RAC1 
                                               mRNA 
                                               0 
                                               
                                               
                                               MAPK6MAPK4 signaling;Signaling by SCF-KIT;Inactivation of Cdc42 and Rac;DAP12 signaling;Ephrin signaling;EPH-ephrin mediated repulsion of cells;Activation of Rac;RHO GTPases Activate Formins;VEGFR2 mediated vascular permeability;RHO GTPases activate PAKs;Sema4D mediated inhibition of cell attachment and migration;G alpha (1213) signalling events;Regulation of actin dynamics for phagocytic cup formation;Sema4D induced cell migration and growth-cone collapse;Signal transduction by L1;RHO GTPases activate CIT;RHO GTPases Activate WASPs and WAVEs;Factors involved in megakaryocyte development and platelet production;RHO GTPases activate KTN1;CD28 dependent Vav1 pathway;DSCAM interactions;NRAGE signals death through JNK;GPVI-mediated activation cascade;Sema3A PAK dependent Axon repulsion;RHO GTPases activate PKNs;SEMA3A-Plexin repulsion signaling by inhibiting Integrin adhesion;RHO GTPases activate IQGAPs;PCPCE pathway;VEGFA-VEGFR2 Pathway;CD28 co-stimulation 
                                               hsa04010;hsa04014;hsa04015;hsa04024;hsa04062;hsa04071;hsa04145;hsa04151;hsa04310;hsa04360;hsa04370;hsa04380;hsa04510;hsa04520;hsa04620;hsa04650;hsa04662;hsa04664;hsa04666;hsa04670;hsa04722;hsa04810;hsa04932;hsa04933;hsa04972;hsa05100;hsa05120;hsa05131;hsa05132;hsa05200;hsa05203;hsa05205;hsa05210;hsa05211;hsa05212;hsa05231;hsa05416 
                                               RAC1 
                                               NA 
                                               slateblue2 
                                               16 
                                               -3.50537634408602 
                                             
                                             
                                               SOS2-mRNA 
                                               -0.739 
                                               0.237 
                                               -1.2 
                                               -0.274 
                                               0.599 
                                               0.434 
                                               0.827 
                                               0.00355 
                                               0.0294 
                                               lm.nb 
                                               Activation of Rac, G alpha (1213) signalling events, NRAGE signals death through JNK, Rho GTPase cycle 
                                               SEBELIUS_2_C4636 
                                               SOS2 
                                               mRNA 
                                               0 
                                               
                                               
                                               Rho GTPase cycle;Activation of Rac;NRAGE signals death through JNK;G alpha (1213) signalling events 
                                               hsa04010;hsa04012;hsa04014;hsa04062;hsa04068;hsa04072;hsa04151;hsa04320;hsa04510;hsa04540;hsa04630;hsa04650;hsa04660;hsa04662;hsa04664;hsa04722;hsa04810;hsa04910;hsa04912;hsa04915;hsa04917;hsa05034;hsa05160;hsa05200;hsa05205;hsa05206;hsa05211;hsa05213;hsa05214;hsa05215;hsa05220;hsa05221;hsa05223;hsa05231 
                                               SOS2 
                                               NA 
                                               slateblue2 
                                               16 
                                               -3.11814345991561 
                                             
                                             
                                               PAK2-mRNA 
                                               -0.138 
                                               0.0752 
                                               -0.285 
                                               0.00968 
                                               0.909 
                                               0.821 
                                               1.01 
                                               0.0751 
                                               0.552 
                                               loglinear 
                                               Activation of Rac, CD209 (DC-SIGN) signaling, CD28 co-stimulation, CD28 dependent Vav1 pathway, Ephrin signaling, RHO GTPases activate PAKs, Sema3A PAK dependent Axon repulsion, VEGFA-VEGFR2 Pathway, VEGFR2 mediated vascular permeability 
                                               SEBELIUS_2_C4636 
                                               PAK2 
                                               mRNA 
                                               0 
                                               
                                               
                                               Nef and signal transduction;CD28 dependent Vav1 pathway;Stimulation of the cell death response by PAK-2p34;Ephrin signaling;Activation of Rac;RHO GTPases activate PAKs;VEGFR2 mediated vascular permeability;Sema3A PAK dependent Axon repulsion;CD209 (DC-SIGN) signaling;VEGFA-VEGFR2 Pathway;CD28 co-stimulation 
                                               hsa04010;hsa04012;hsa04014;hsa04360;hsa04510;hsa04660;hsa04810;hsa05211 
                                               PAK2 
                                               NA 
                                               azure4 
                                               1 
                                               -1.83510638297872 
                                             
                                           
                                           DE Results - Activation of Rac genes - Line: Caco2 vs.Fibro  Table displaying each sample's global significance scores and directed global significance scores as defined in the heatmaps above. The global significance score is calculated as the square root of the mean squared t-statistic for the genes in a gene set, with t-statistics coming from the linear regression underlying our differential expression analysis. The directed global significance score is calculated as the square root of the mean signed squared t-statistic for the genes in a gene set, with t-statistics coming from the linear regression underlying our differential expression analysis.   
                                      
                                       
                                  
                                
                                
                                  
                                       
                                      
                                           Volcano Plot: Line: HEPG2 vs.Fibro   More Plot Information                Volcano Plot: Line: HEPG2 vs.Fibro  Volcano plot displaying each gene's -log10(p-value) and log2 fold change for the selected covariate. Highly statistically significant genes fall at the top of the plot, and highly differentially expressed genes fall to either side. Genes within the selected gene set are highlighted in orange. Horizontal lines indicate various False Discovery Rate (FDR) thresholds.   
                                           DE Results - Activation of Rac genes - Line: HEPG2 vs.Fibro   More Plot Information      Download CSV Data     
                                           
                                             
                                             
                                           
                                           
                                             
                                               Probe Label 
                                               Log2 fold change 
                                               std error (log2) 
                                               Lower confidence limit (log2) 
                                               Upper confidence limit (log2) 
                                               Linear fold change 
                                               Lower confidence limit (linear) 
                                               Upper confidence limit (linear) 
                                               P-value 
                                               BY.p.value 
                                               method 
                                               Gene.sets 
                                               Codeset.Name 
                                               Probe.Label 
                                               Analyte.Type 
                                               Is.Control 
                                               Control.Type 
                                               Related.Probes 
                                               Probe.Annotation 
                                               KEGG.Pathways 
                                               Official.Gene.Name 
                                               Control.Conc 
                                               volcanocol 
                                               volcanopch 
                                               tstats.all 
                                             
                                           
                                           
                                             
                                               PAK4-mRNA 
                                               1.84 
                                               0.097 
                                               1.65 
                                               2.03 
                                               3.58 
                                               3.13 
                                               4.08 
                                               1.24e-20 
                                               1.18e-18 
                                               loglinear 
                                               Activation of Rac 
                                               SEBELIUS_2_C4636 
                                               PAK4 
                                               mRNA 
                                               0 
                                               
                                               
                                               Activation of Rac 
                                               hsa04012;hsa04014;hsa04360;hsa04510;hsa04660;hsa04810;hsa05206;hsa05211 
                                               PAK4 
                                               NA 
                                               slateblue1 
                                               16 
                                               18.9690721649485 
                                             
                                             
                                               PAK1-mRNA 
                                               -2.44 
                                               0.179 
                                               -2.79 
                                               -2.09 
                                               0.184 
                                               0.145 
                                               0.235 
                                               5.34e-16 
                                               1.72e-14 
                                               lm.nb 
                                               Activation of Rac, CD209 (DC-SIGN) signaling, CD28 co-stimulation, CD28 dependent Vav1 pathway, EPHB-mediated forward signaling, Ephrin signaling, FCERI mediated MAPK activation, MAPK6MAPK4 signaling, Regulation of actin dynamics for phagocytic cup formation, RHO GTPases activate PAKs, Sema3A PAK dependent Axon repulsion, VEGFA-VEGFR2 Pathway, VEGFR2 mediated vascular permeability 
                                               SEBELIUS_2_C4636 
                                               PAK1 
                                               mRNA 
                                               0 
                                               
                                               
                                               EPHB-mediated forward signaling;Ephrin signaling;MAPK6MAPK4 signaling;DSCAM interactions;Activation of Rac;VEGFR2 mediated vascular permeability;RHO GTPases activate PAKs;Sema3A PAK dependent Axon repulsion;FCERI mediated MAPK activation;Regulation of actin dynamics for phagocytic cup formation;CD209 (DC-SIGN) signaling;CD28 dependent Vav1 pathway;VEGFA-VEGFR2 Pathway;CD28 co-stimulation 
                                               hsa04010;hsa04012;hsa04014;hsa04024;hsa04062;hsa04360;hsa04510;hsa04650;hsa04660;hsa04666;hsa04810;hsa05120;hsa05205;hsa05211 
                                               PAK1 
                                               NA 
                                               slateblue1 
                                               16 
                                               -13.6312849162011 
                                             
                                             
                                               PAK2-mRNA 
                                               0.446 
                                               0.0825 
                                               0.284 
                                               0.608 
                                               1.36 
                                               1.22 
                                               1.52 
                                               4.03e-06 
                                               4.65e-05 
                                               loglinear 
                                               Activation of Rac, CD209 (DC-SIGN) signaling, CD28 co-stimulation, CD28 dependent Vav1 pathway, Ephrin signaling, RHO GTPases activate PAKs, Sema3A PAK dependent Axon repulsion, VEGFA-VEGFR2 Pathway, VEGFR2 mediated vascular permeability 
                                               SEBELIUS_2_C4636 
                                               PAK2 
                                               mRNA 
                                               0 
                                               
                                               
                                               Nef and signal transduction;CD28 dependent Vav1 pathway;Stimulation of the cell death response by PAK-2p34;Ephrin signaling;Activation of Rac;RHO GTPases activate PAKs;VEGFR2 mediated vascular permeability;Sema3A PAK dependent Axon repulsion;CD209 (DC-SIGN) signaling;VEGFA-VEGFR2 Pathway;CD28 co-stimulation 
                                               hsa04010;hsa04012;hsa04014;hsa04360;hsa04510;hsa04660;hsa04810;hsa05211 
                                               PAK2 
                                               NA 
                                               slateblue1 
                                               16 
                                               5.40606060606061 
                                             
                                             
                                               SOS2-mRNA 
                                               0.228 
                                               0.26 
                                               -0.281 
                                               0.737 
                                               1.17 
                                               0.823 
                                               1.67 
                                               0.386 
                                               1 
                                               lm.nb 
                                               Activation of Rac, G alpha (1213) signalling events, NRAGE signals death through JNK, Rho GTPase cycle 
                                               SEBELIUS_2_C4636 
                                               SOS2 
                                               mRNA 
                                               0 
                                               
                                               
                                               Rho GTPase cycle;Activation of Rac;NRAGE signals death through JNK;G alpha (1213) signalling events 
                                               hsa04010;hsa04012;hsa04014;hsa04062;hsa04068;hsa04072;hsa04151;hsa04320;hsa04510;hsa04540;hsa04630;hsa04650;hsa04660;hsa04662;hsa04664;hsa04722;hsa04810;hsa04910;hsa04912;hsa04915;hsa04917;hsa05034;hsa05160;hsa05200;hsa05205;hsa05206;hsa05211;hsa05213;hsa05214;hsa05215;hsa05220;hsa05221;hsa05223;hsa05231 
                                               SOS2 
                                               NA 
                                               azure4 
                                               1 
                                               0.876923076923077 
                                             
                                             
                                               RAC1-mRNA 
                                               -0.0792 
                                               0.102 
                                               -0.279 
                                               0.121 
                                               0.947 
                                               0.824 
                                               1.09 
                                               0.442 
                                               1 
                                               lm.nb 
                                               Activation of Rac, CD28 co-stimulation, CD28 dependent Vav1 pathway, DAP12 signaling, EPH-ephrin mediated repulsion of cells, Ephrin signaling, G alpha (1213) signalling events, GPVI-mediated activation cascade, MAPK6MAPK4 signaling, NRAGE signals death through JNK, PCPCE pathway, Regulation of actin dynamics for phagocytic cup formation, RHO GTPases activate CIT, RHO GTPases Activate Formins, RHO GTPases activate IQGAPs, RHO GTPases activate PAKs, RHO GTPases activate PKNs, RHO GTPases Activate WASPs and WAVEs, Sema3A PAK dependent Axon repulsion, Sema4D induced cell migration and growth-cone collapse, Signal transduction by L1, Signaling by SCF-KIT, VEGFA-VEGFR2 Pathway, VEGFR2 mediated vascular permeability 
                                               SEBELIUS_2_C4636 
                                               RAC1 
                                               mRNA 
                                               0 
                                               
                                               
                                               MAPK6MAPK4 signaling;Signaling by SCF-KIT;Inactivation of Cdc42 and Rac;DAP12 signaling;Ephrin signaling;EPH-ephrin mediated repulsion of cells;Activation of Rac;RHO GTPases Activate Formins;VEGFR2 mediated vascular permeability;RHO GTPases activate PAKs;Sema4D mediated inhibition of cell attachment and migration;G alpha (1213) signalling events;Regulation of actin dynamics for phagocytic cup formation;Sema4D induced cell migration and growth-cone collapse;Signal transduction by L1;RHO GTPases activate CIT;RHO GTPases Activate WASPs and WAVEs;Factors involved in megakaryocyte development and platelet production;RHO GTPases activate KTN1;CD28 dependent Vav1 pathway;DSCAM interactions;NRAGE signals death through JNK;GPVI-mediated activation cascade;Sema3A PAK dependent Axon repulsion;RHO GTPases activate PKNs;SEMA3A-Plexin repulsion signaling by inhibiting Integrin adhesion;RHO GTPases activate IQGAPs;PCPCE pathway;VEGFA-VEGFR2 Pathway;CD28 co-stimulation 
                                               hsa04010;hsa04014;hsa04015;hsa04024;hsa04062;hsa04071;hsa04145;hsa04151;hsa04310;hsa04360;hsa04370;hsa04380;hsa04510;hsa04520;hsa04620;hsa04650;hsa04662;hsa04664;hsa04666;hsa04670;hsa04722;hsa04810;hsa04932;hsa04933;hsa04972;hsa05100;hsa05120;hsa05131;hsa05132;hsa05200;hsa05203;hsa05205;hsa05210;hsa05211;hsa05212;hsa05231;hsa05416 
                                               RAC1 
                                               NA 
                                               azure4 
                                               1 
                                               -0.776470588235294 
                                             
                                           
                                           DE Results - Activation of Rac genes - Line: HEPG2 vs.Fibro  Table displaying each sample's global significance scores and directed global significance scores as defined in the heatmaps above. The global significance score is calculated as the square root of the mean squared t-statistic for the genes in a gene set, with t-statistics coming from the linear regression underlying our differential expression analysis. The directed global significance score is calculated as the square root of the mean signed squared t-statistic for the genes in a gene set, with t-statistics coming from the linear regression underlying our differential expression analysis.   
                                      
                                       
                                  
                                
                                
                                  
                                       
                                      
                                           Volcano Plot: Line: HT29 vs.Fibro   More Plot Information                Volcano Plot: Line: HT29 vs.Fibro  Volcano plot displaying each gene's -log10(p-value) and log2 fold change for the selected covariate. Highly statistically significant genes fall at the top of the plot, and highly differentially expressed genes fall to either side. Genes within the selected gene set are highlighted in orange. Horizontal lines indicate various False Discovery Rate (FDR) thresholds.   
                                           DE Results - Activation of Rac genes - Line: HT29 vs.Fibro   More Plot Information      Download CSV Data     
                                           
                                             
                                             
                                           
                                           
                                             
                                               Probe Label 
                                               Log2 fold change 
                                               std error (log2) 
                                               Lower confidence limit (log2) 
                                               Upper confidence limit (log2) 
                                               Linear fold change 
                                               Lower confidence limit (linear) 
                                               Upper confidence limit (linear) 
                                               P-value 
                                               BY.p.value 
                                               method 
                                               Gene.sets 
                                               Codeset.Name 
                                               Probe.Label 
                                               Analyte.Type 
                                               Is.Control 
                                               Control.Type 
                                               Related.Probes 
                                               Probe.Annotation 
                                               KEGG.Pathways 
                                               Official.Gene.Name 
                                               Control.Conc 
                                               volcanocol 
                                               volcanopch 
                                               tstats.all 
                                             
                                           
                                           
                                             
                                               PAK4-mRNA 
                                               2.93 
                                               0.0914 
                                               2.75 
                                               3.11 
                                               7.64 
                                               6.75 
                                               8.65 
                                               1.3e-28 
                                               2.64e-26 
                                               loglinear 
                                               Activation of Rac 
                                               SEBELIUS_2_C4636 
                                               PAK4 
                                               mRNA 
                                               0 
                                               
                                               
                                               Activation of Rac 
                                               hsa04012;hsa04014;hsa04360;hsa04510;hsa04660;hsa04810;hsa05206;hsa05211 
                                               PAK4 
                                               NA 
                                               slateblue1 
                                               16 
                                               32.0568927789934 
                                             
                                             
                                               PAK1-mRNA 
                                               2.31 
                                               0.165 
                                               1.99 
                                               2.63 
                                               4.96 
                                               3.96 
                                               6.2 
                                               2.18e-16 
                                               5.51e-15 
                                               lm.nb 
                                               Activation of Rac, CD209 (DC-SIGN) signaling, CD28 co-stimulation, CD28 dependent Vav1 pathway, EPHB-mediated forward signaling, Ephrin signaling, FCERI mediated MAPK activation, MAPK6MAPK4 signaling, Regulation of actin dynamics for phagocytic cup formation, RHO GTPases activate PAKs, Sema3A PAK dependent Axon repulsion, VEGFA-VEGFR2 Pathway, VEGFR2 mediated vascular permeability 
                                               SEBELIUS_2_C4636 
                                               PAK1 
                                               mRNA 
                                               0 
                                               
                                               
                                               EPHB-mediated forward signaling;Ephrin signaling;MAPK6MAPK4 signaling;DSCAM interactions;Activation of Rac;VEGFR2 mediated vascular permeability;RHO GTPases activate PAKs;Sema3A PAK dependent Axon repulsion;FCERI mediated MAPK activation;Regulation of actin dynamics for phagocytic cup formation;CD209 (DC-SIGN) signaling;CD28 dependent Vav1 pathway;VEGFA-VEGFR2 Pathway;CD28 co-stimulation 
                                               hsa04010;hsa04012;hsa04014;hsa04024;hsa04062;hsa04360;hsa04510;hsa04650;hsa04660;hsa04666;hsa04810;hsa05120;hsa05205;hsa05211 
                                               PAK1 
                                               NA 
                                               slateblue1 
                                               16 
                                               14 
                                             
                                             
                                               RAC1-mRNA 
                                               1.11 
                                               0.0959 
                                               0.925 
                                               1.3 
                                               2.16 
                                               1.9 
                                               2.46 
                                               6.93e-14 
                                               1.24e-12 
                                               lm.nb 
                                               Activation of Rac, CD28 co-stimulation, CD28 dependent Vav1 pathway, DAP12 signaling, EPH-ephrin mediated repulsion of cells, Ephrin signaling, G alpha (1213) signalling events, GPVI-mediated activation cascade, MAPK6MAPK4 signaling, NRAGE signals death through JNK, PCPCE pathway, Regulation of actin dynamics for phagocytic cup formation, RHO GTPases activate CIT, RHO GTPases Activate Formins, RHO GTPases activate IQGAPs, RHO GTPases activate PAKs, RHO GTPases activate PKNs, RHO GTPases Activate WASPs and WAVEs, Sema3A PAK dependent Axon repulsion, Sema4D induced cell migration and growth-cone collapse, Signal transduction by L1, Signaling by SCF-KIT, VEGFA-VEGFR2 Pathway, VEGFR2 mediated vascular permeability 
                                               SEBELIUS_2_C4636 
                                               RAC1 
                                               mRNA 
                                               0 
                                               
                                               
                                               MAPK6MAPK4 signaling;Signaling by SCF-KIT;Inactivation of Cdc42 and Rac;DAP12 signaling;Ephrin signaling;EPH-ephrin mediated repulsion of cells;Activation of Rac;RHO GTPases Activate Formins;VEGFR2 mediated vascular permeability;RHO GTPases activate PAKs;Sema4D mediated inhibition of cell attachment and migration;G alpha (1213) signalling events;Regulation of actin dynamics for phagocytic cup formation;Sema4D induced cell migration and growth-cone collapse;Signal transduction by L1;RHO GTPases activate CIT;RHO GTPases Activate WASPs and WAVEs;Factors involved in megakaryocyte development and platelet production;RHO GTPases activate KTN1;CD28 dependent Vav1 pathway;DSCAM interactions;NRAGE signals death through JNK;GPVI-mediated activation cascade;Sema3A PAK dependent Axon repulsion;RHO GTPases activate PKNs;SEMA3A-Plexin repulsion signaling by inhibiting Integrin adhesion;RHO GTPases activate IQGAPs;PCPCE pathway;VEGFA-VEGFR2 Pathway;CD28 co-stimulation 
                                               hsa04010;hsa04014;hsa04015;hsa04024;hsa04062;hsa04071;hsa04145;hsa04151;hsa04310;hsa04360;hsa04370;hsa04380;hsa04510;hsa04520;hsa04620;hsa04650;hsa04662;hsa04664;hsa04666;hsa04670;hsa04722;hsa04810;hsa04932;hsa04933;hsa04972;hsa05100;hsa05120;hsa05131;hsa05132;hsa05200;hsa05203;hsa05205;hsa05210;hsa05211;hsa05212;hsa05231;hsa05416 
                                               RAC1 
                                               NA 
                                               slateblue1 
                                               16 
                                               11.5745568300313 
                                             
                                             
                                               PAK2-mRNA 
                                               0.221 
                                               0.0777 
                                               0.0684 
                                               0.373 
                                               1.17 
                                               1.05 
                                               1.3 
                                               0.00728 
                                               0.0571 
                                               loglinear 
                                               Activation of Rac, CD209 (DC-SIGN) signaling, CD28 co-stimulation, CD28 dependent Vav1 pathway, Ephrin signaling, RHO GTPases activate PAKs, Sema3A PAK dependent Axon repulsion, VEGFA-VEGFR2 Pathway, VEGFR2 mediated vascular permeability 
                                               SEBELIUS_2_C4636 
                                               PAK2 
                                               mRNA 
                                               0 
                                               
                                               
                                               Nef and signal transduction;CD28 dependent Vav1 pathway;Stimulation of the cell death response by PAK-2p34;Ephrin signaling;Activation of Rac;RHO GTPases activate PAKs;VEGFR2 mediated vascular permeability;Sema3A PAK dependent Axon repulsion;CD209 (DC-SIGN) signaling;VEGFA-VEGFR2 Pathway;CD28 co-stimulation 
                                               hsa04010;hsa04012;hsa04014;hsa04360;hsa04510;hsa04660;hsa04810;hsa05211 
                                               PAK2 
                                               NA 
                                               slateblue3 
                                               16 
                                               2.84427284427284 
                                             
                                             
                                               SOS2-mRNA 
                                               -0.16 
                                               0.245 
                                               -0.64 
                                               0.321 
                                               0.895 
                                               0.642 
                                               1.25 
                                               0.519 
                                               1 
                                               lm.nb 
                                               Activation of Rac, G alpha (1213) signalling events, NRAGE signals death through JNK, Rho GTPase cycle 
                                               SEBELIUS_2_C4636 
                                               SOS2 
                                               mRNA 
                                               0 
                                               
                                               
                                               Rho GTPase cycle;Activation of Rac;NRAGE signals death through JNK;G alpha (1213) signalling events 
                                               hsa04010;hsa04012;hsa04014;hsa04062;hsa04068;hsa04072;hsa04151;hsa04320;hsa04510;hsa04540;hsa04630;hsa04650;hsa04660;hsa04662;hsa04664;hsa04722;hsa04810;hsa04910;hsa04912;hsa04915;hsa04917;hsa05034;hsa05160;hsa05200;hsa05205;hsa05206;hsa05211;hsa05213;hsa05214;hsa05215;hsa05220;hsa05221;hsa05223;hsa05231 
                                               SOS2 
                                               NA 
                                               azure4 
                                               1 
                                               -0.653061224489796 
                                             
                                           
                                           DE Results - Activation of Rac genes - Line: HT29 vs.Fibro  Table displaying each sample's global significance scores and directed global significance scores as defined in the heatmaps above. The global significance score is calculated as the square root of the mean squared t-statistic for the genes in a gene set, with t-statistics coming from the linear regression underlying our differential expression analysis. The directed global significance score is calculated as the square root of the mean signed squared t-statistic for the genes in a gene set, with t-statistics coming from the linear regression underlying our differential expression analysis.   
                                      
                                       
                                  
                                
                              
                            
                           
                            
                          
                           
                      
                    
                    
                      
                           
                          
                               Day    Line     
                              
                                  Day  
                              
                               
                              
                                  Line: differential expression in Caco2 vs. baseline of Fibro  
                                  Line: differential expression in HEPG2 vs. baseline of Fibro  
                                  Line: differential expression in HT29 vs. baseline of Fibro  
                              
                              
                            
                              
                                
                                  
                                       
                                      
                                           Volcano Plot: Day   More Plot Information                Volcano Plot: Day  Volcano plot displaying each gene's -log10(p-value) and log2 fold change for the selected covariate. Highly statistically significant genes fall at the top of the plot, and highly differentially expressed genes fall to either side. Genes within the selected gene set are highlighted in orange. Horizontal lines indicate various False Discovery Rate (FDR) thresholds.   
                                           DE Results - Apoptotic cleavage of cellular proteins genes - Day   More Plot Information      Download CSV Data     
                                           
                                             
                                             
                                           
                                           
                                             
                                               Probe Label 
                                               Log2 fold change 
                                               std error (log2) 
                                               Lower confidence limit (log2) 
                                               Upper confidence limit (log2) 
                                               Linear fold change 
                                               Lower confidence limit (linear) 
                                               Upper confidence limit (linear) 
                                               P-value 
                                               BY.p.value 
                                               method 
                                               Gene.sets 
                                               Codeset.Name 
                                               Probe.Label 
                                               Analyte.Type 
                                               Is.Control 
                                               Control.Type 
                                               Related.Probes 
                                               Probe.Annotation 
                                               KEGG.Pathways 
                                               Official.Gene.Name 
                                               Control.Conc 
                                               volcanocol 
                                               volcanopch 
                                               tstats.all 
                                             
                                           
                                           
                                             
                                               TJP1-mRNA 
                                               -0.068 
                                               0.0114 
                                               -0.0903 
                                               -0.0458 
                                               0.954 
                                               0.939 
                                               0.969 
                                               6.41e-07 
                                               7.39e-05 
                                               lm.nb 
                                               Apoptotic cleavage of cellular proteins 
                                               SEBELIUS_2_C4636 
                                               TJP1 
                                               mRNA 
                                               0 
                                               
                                               
                                               Apoptotic cleavage of cellular proteins 
                                               hsa04520;hsa04530;hsa04540;hsa05110;hsa05120;hsa05132 
                                               TJP1 
                                               NA 
                                               slateblue1 
                                               16 
                                               -5.96491228070175 
                                             
                                             
                                               PTK2-mRNA 
                                               0.0792 
                                               0.0142 
                                               0.0514 
                                               0.107 
                                               1.06 
                                               1.04 
                                               1.08 
                                               2.29e-06 
                                               0.000181 
                                               lm.nb 
                                               Apoptotic cleavage of cellular proteins, ARMS-mediated activation, DAP12 signaling, DCC mediated attractive signaling, Downstream signal transduction, EPHA-mediated growth cone collapse, EPHB-mediated forward signaling, FCERI mediated MAPK activation, Frs2-mediated activation, Gastrin-CREB signalling pathway via PKC and MAPK, Integrin alphaIIb beta3 signaling, Interleukin receptor SHC signaling, IRS-related events triggered by IGF1R, p130Cas linkage to MAPK signaling for integrins, RAFMAP kinase cascade, Regulation of actin dynamics for phagocytic cup formation, RHO GTPases Activate WASPs and WAVEs, Signaling by ERBB2, Signaling by PDGF, Signaling by SCF-KIT, VEGFA-VEGFR2 Pathway, VEGFR2 mediated cell proliferation 
                                               SEBELIUS_2_C4636 
                                               PTK2 
                                               mRNA 
                                               0 
                                               
                                               
                                               Apoptotic cleavage of cellular proteins;RAFMAP kinase cascade;RHO GTPases Activate WASPs and WAVEs;DCC mediated attractive signaling;p130Cas linkage to MAPK signaling for integrins;EPHB-mediated forward signaling;EPHA-mediated growth cone collapse;Regulation of actin dynamics for phagocytic cup formation;Downstream signal transduction;VEGFR2 mediated cell proliferation;Signaling by SCF-KIT;Frs2-mediated activation;ARMS-mediated activation;Gastrin-CREB signalling pathway via PKC and MAPK;Interleukin receptor SHC signaling;FCERI mediated MAPK activation;DAP12 signaling;Signaling by PDGF;VEGFA-VEGFR2 Pathway;Signaling by ERBB2;IRS-related events triggered by IGF1R;Integrin alphaIIb beta3 signaling 
                                               hsa04012;hsa04062;hsa04151;hsa04360;hsa04370;hsa04510;hsa04670;hsa04810;hsa05100;hsa05146;hsa05200;hsa05202;hsa05205;hsa05222 
                                               PTK2 
                                               NA 
                                               slateblue1 
                                               16 
                                               5.57746478873239 
                                             
                                             
                                               TJP2-mRNA 
                                               0.0982 
                                               0.0202 
                                               0.0586 
                                               0.138 
                                               1.07 
                                               1.04 
                                               1.1 
                                               2.22e-05 
                                               0.00105 
                                               lm.nb 
                                               Apoptotic cleavage of cellular proteins 
                                               SEBELIUS_2_C4636 
                                               TJP2 
                                               mRNA 
                                               0 
                                               
                                               
                                               Apoptotic cleavage of cellular proteins 
                                               hsa04530;hsa05110 
                                               TJP2 
                                               NA 
                                               slateblue1 
                                               16 
                                               4.86138613861386 
                                             
                                             
                                               OCLN-mRNA 
                                               0.0986 
                                               0.0249 
                                               0.0498 
                                               0.147 
                                               1.07 
                                               1.04 
                                               1.11 
                                               0.000325 
                                               0.00903 
                                               lm.nb 
                                               Apoptotic cleavage of cellular proteins 
                                               SEBELIUS_2_C4636 
                                               OCLN 
                                               mRNA 
                                               0 
                                               
                                               
                                               Apoptotic cleavage of cellular proteins 
                                               hsa04514;hsa04530;hsa04670;hsa05130;hsa05160 
                                               OCLN 
                                               NA 
                                               slateblue2 
                                               16 
                                               3.95983935742972 
                                             
                                             
                                               SPTAN1-mRNA 
                                               -0.0399 
                                               0.0123 
                                               -0.0641 
                                               -0.0157 
                                               0.973 
                                               0.957 
                                               0.989 
                                               0.00257 
                                               0.0486 
                                               lm.nb 
                                               Apoptotic cleavage of cellular proteins, ARMS-mediated activation, DAP12 signaling, Downstream signal transduction, FCERI mediated MAPK activation, Frs2-mediated activation, Gastrin-CREB signalling pathway via PKC and MAPK, Interleukin receptor SHC signaling, IRS-related events triggered by IGF1R, Nephrin interactions, Signaling by ERBB2, Signaling by PDGF, Signaling by SCF-KIT, VEGFA-VEGFR2 Pathway, VEGFR2 mediated cell proliferation 
                                               SEBELIUS_2_C4636 
                                               SPTAN1 
                                               mRNA 
                                               0 
                                               
                                               
                                               Nephrin interactions;Caspase-mediated cleavage of cytoskeletal proteins;Downstream signal transduction;VEGFR2 mediated cell proliferation;Signaling by SCF-KIT;Frs2-mediated activation;ARMS-mediated activation;Gastrin-CREB signalling pathway via PKC and MAPK;Interleukin receptor SHC signaling;FCERI mediated MAPK activation;DAP12 signaling;Signaling by PDGF;VEGFA-VEGFR2 Pathway;Signaling by ERBB2;IRS-related events triggered by IGF1R;Apoptotic cleavage of cellular proteins 
                                               hsa04530 
                                               SPTAN1 
                                               NA 
                                               slateblue2 
                                               16 
                                               -3.24390243902439 
                                             
                                             
                                               ROCK1-mRNA 
                                               -0.0239 
                                               0.00838 
                                               -0.0403 
                                               -0.00747 
                                               0.984 
                                               0.972 
                                               0.995 
                                               0.00709 
                                               0.103 
                                               loglinear 
                                               Apoptotic cleavage of cellular proteins, EPHA-mediated growth cone collapse, EPHB-mediated forward signaling, G alpha (1213) signalling events, RHO GTPases Activate ROCKs, Sema4D induced cell migration and growth-cone collapse, VEGFA-VEGFR2 Pathway 
                                               SEBELIUS_2_C4636 
                                               ROCK1 
                                               mRNA 
                                               0 
                                               
                                               
                                               EPHB-mediated forward signaling;EPHA-mediated growth cone collapse;RHO GTPases Activate ROCKs;Apoptotic cleavage of cellular proteins;G alpha (1213) signalling events;VEGFA-VEGFR2 Pathway;Sema4D induced cell migration and growth-cone collapse 
                                               hsa04022;hsa04024;hsa04062;hsa04071;hsa04270;hsa04350;hsa04360;hsa04510;hsa04611;hsa04670;hsa04810;hsa04921;hsa05130;hsa05131;hsa05132;hsa05200;hsa05205;hsa05206 
                                               ROCK1 
                                               NA 
                                               slateblue4 
                                               16 
                                               -2.85202863961814 
                                             
                                             
                                               APC-mRNA 
                                               0.0285 
                                               0.0157 
                                               -0.0023 
                                               0.0592 
                                               1.02 
                                               0.998 
                                               1.04 
                                               0.0779 
                                               0.762 
                                               lm.nb 
                                               Apoptotic cleavage of cellular proteins, AXIN missense mutants destabilize the destruction complex, Beta-catenin phosphorylation cascade, Deactivation of the beta-catenin transactivating complex, Disassembly of the destruction complex and recruitment of AXIN to the membrane, Misspliced GSK3beta mutants stabilize beta-catenin, S33 mutants of beta-catenin aren.SQT.t phosphorylated, S37 mutants of beta-catenin aren.SQT.t phosphorylated, S45 mutants of beta-catenin aren.SQT.t phosphorylated, T41 mutants of beta-catenin aren.SQT.t phosphorylated, Truncations of AMER1 destabilize the destruction complex 
                                               SEBELIUS_2_C4636 
                                               APC 
                                               mRNA 
                                               0 
                                               
                                               
                                               Apoptotic cleavage of cellular proteins;Misspliced GSK3beta mutants stabilize beta-catenin;APC truncation mutants have impaired AXIN binding;T41 mutants of beta-catenin aren.SQT.t phosphorylated;AXIN missense mutants destabilize the destruction complex;S45 mutants of beta-catenin aren.SQT.t phosphorylated;Deactivation of the beta-catenin transactivating complex;Disassembly of the destruction complex and recruitment of AXIN to the membrane;S37 mutants of beta-catenin aren.SQT.t phosphorylated;S33 mutants of beta-catenin aren.SQT.t phosphorylated;Beta-catenin phosphorylation cascade;Truncations of AMER1 destabilize the destruction complex 
                                               hsa04310;hsa04390;hsa04550;hsa04810;hsa05166;hsa05200;hsa05206;hsa05210;hsa05213;hsa05217 
                                               APC 
                                               NA 
                                               azure4 
                                               1 
                                               1.81528662420382 
                                             
                                             
                                               CTNNB1-mRNA 
                                               -0.000519 
                                               0.0215 
                                               -0.0427 
                                               0.0416 
                                               1 
                                               0.971 
                                               1.03 
                                               0.981 
                                               1 
                                               lm.nb 
                                               Apoptotic cleavage of cellular proteins, Beta-catenin phosphorylation cascade, Deactivation of the beta-catenin transactivating complex, Disassembly of the destruction complex and recruitment of AXIN to the membrane, Formation of the beta-catenin.CL.TCF transactivating complex, Misspliced GSK3beta mutants stabilize beta-catenin, Repression of WNT target genes, RHO GTPases activate IQGAPs, S37 mutants of beta-catenin aren.SQT.t phosphorylated, S45 mutants of beta-catenin aren.SQT.t phosphorylated, T41 mutants of beta-catenin aren.SQT.t phosphorylated, VEGFA-VEGFR2 Pathway, VEGFR2 mediated vascular permeability 
                                               SEBELIUS_2_C4636 
                                               CTNNB1 
                                               mRNA 
                                               0 
                                               
                                               
                                               Disassembly of the destruction complex and recruitment of AXIN to the membrane;S37 mutants of beta-catenin aren.SQT.t phosphorylated;T41 mutants of beta-catenin aren.SQT.t phosphorylated;S45 mutants of beta-catenin aren.SQT.t phosphorylated;Binding of TCFLEF.CL.CTNNB1 to target gene promoters;RHO GTPases activate IQGAPs;Deactivation of the beta-catenin transactivating complex;Repression of WNT target genes;Apoptotic cleavage of cell adhesion  proteins;Adherens junctions interactions;Misspliced GSK3beta mutants stabilize beta-catenin;VEGFR2 mediated vascular permeability;Beta-catenin phosphorylation cascade;Synthesis# secretion# and inactivation of Glucagon-like Peptide-1 (GLP-1);Formation of the beta-catenin.CL.TCF transactivating complex;Apoptotic cleavage of cellular proteins;VEGFA-VEGFR2 Pathway 
                                               hsa04015;hsa04310;hsa04390;hsa04510;hsa04520;hsa04530;hsa04550;hsa04670;hsa04916;hsa04919;hsa05100;hsa05130;hsa05166;hsa05200;hsa05205;hsa05210;hsa05213;hsa05215;hsa05216;hsa05217;hsa05412 
                                               CTNNB1 
                                               NA 
                                               azure4 
                                               1 
                                               -0.0241395348837209 
                                             
                                           
                                           DE Results - Apoptotic cleavage of cellular proteins genes - Day  Table displaying each sample's global significance scores and directed global significance scores as defined in the heatmaps above. The global significance score is calculated as the square root of the mean squared t-statistic for the genes in a gene set, with t-statistics coming from the linear regression underlying our differential expression analysis. The directed global significance score is calculated as the square root of the mean signed squared t-statistic for the genes in a gene set, with t-statistics coming from the linear regression underlying our differential expression analysis.   
                                      
                                       
                                  
                                
                              
                            
                            
                              
                                
                                  
                                       
                                      
                                           Volcano Plot: Line: Caco2 vs.Fibro   More Plot Information                Volcano Plot: Line: Caco2 vs.Fibro  Volcano plot displaying each gene's -log10(p-value) and log2 fold change for the selected covariate. Highly statistically significant genes fall at the top of the plot, and highly differentially expressed genes fall to either side. Genes within the selected gene set are highlighted in orange. Horizontal lines indicate various False Discovery Rate (FDR) thresholds.   
                                           DE Results - Apoptotic cleavage of cellular proteins genes - Line: Caco2 vs.Fibro   More Plot Information      Download CSV Data     
                                           
                                             
                                             
                                           
                                           
                                             
                                               Probe Label 
                                               Log2 fold change 
                                               std error (log2) 
                                               Lower confidence limit (log2) 
                                               Upper confidence limit (log2) 
                                               Linear fold change 
                                               Lower confidence limit (linear) 
                                               Upper confidence limit (linear) 
                                               P-value 
                                               BY.p.value 
                                               method 
                                               Gene.sets 
                                               Codeset.Name 
                                               Probe.Label 
                                               Analyte.Type 
                                               Is.Control 
                                               Control.Type 
                                               Related.Probes 
                                               Probe.Annotation 
                                               KEGG.Pathways 
                                               Official.Gene.Name 
                                               Control.Conc 
                                               volcanocol 
                                               volcanopch 
                                               tstats.all 
                                             
                                           
                                           
                                             
                                               OCLN-mRNA 
                                               4.7 
                                               0.221 
                                               4.26 
                                               5.13 
                                               25.9 
                                               19.2 
                                               35 
                                               2.42e-22 
                                               2.64e-20 
                                               lm.nb 
                                               Apoptotic cleavage of cellular proteins 
                                               SEBELIUS_2_C4636 
                                               OCLN 
                                               mRNA 
                                               0 
                                               
                                               
                                               Apoptotic cleavage of cellular proteins 
                                               hsa04514;hsa04530;hsa04670;hsa05130;hsa05160 
                                               OCLN 
                                               NA 
                                               slateblue1 
                                               16 
                                               21.2669683257919 
                                             
                                             
                                               PTK2-mRNA 
                                               0.866 
                                               0.108 
                                               0.654 
                                               1.08 
                                               1.82 
                                               1.57 
                                               2.11 
                                               1.35e-09 
                                               2e-08 
                                               lm.nb 
                                               Apoptotic cleavage of cellular proteins, ARMS-mediated activation, DAP12 signaling, DCC mediated attractive signaling, Downstream signal transduction, EPHA-mediated growth cone collapse, EPHB-mediated forward signaling, FCERI mediated MAPK activation, Frs2-mediated activation, Gastrin-CREB signalling pathway via PKC and MAPK, Integrin alphaIIb beta3 signaling, Interleukin receptor SHC signaling, IRS-related events triggered by IGF1R, p130Cas linkage to MAPK signaling for integrins, RAFMAP kinase cascade, Regulation of actin dynamics for phagocytic cup formation, RHO GTPases Activate WASPs and WAVEs, Signaling by ERBB2, Signaling by PDGF, Signaling by SCF-KIT, VEGFA-VEGFR2 Pathway, VEGFR2 mediated cell proliferation 
                                               SEBELIUS_2_C4636 
                                               PTK2 
                                               mRNA 
                                               0 
                                               
                                               
                                               Apoptotic cleavage of cellular proteins;RAFMAP kinase cascade;RHO GTPases Activate WASPs and WAVEs;DCC mediated attractive signaling;p130Cas linkage to MAPK signaling for integrins;EPHB-mediated forward signaling;EPHA-mediated growth cone collapse;Regulation of actin dynamics for phagocytic cup formation;Downstream signal transduction;VEGFR2 mediated cell proliferation;Signaling by SCF-KIT;Frs2-mediated activation;ARMS-mediated activation;Gastrin-CREB signalling pathway via PKC and MAPK;Interleukin receptor SHC signaling;FCERI mediated MAPK activation;DAP12 signaling;Signaling by PDGF;VEGFA-VEGFR2 Pathway;Signaling by ERBB2;IRS-related events triggered by IGF1R;Integrin alphaIIb beta3 signaling 
                                               hsa04012;hsa04062;hsa04151;hsa04360;hsa04370;hsa04510;hsa04670;hsa04810;hsa05100;hsa05146;hsa05200;hsa05202;hsa05205;hsa05222 
                                               PTK2 
                                               NA 
                                               slateblue1 
                                               16 
                                               8.01851851851852 
                                             
                                             
                                               APC-mRNA 
                                               -0.862 
                                               0.115 
                                               -1.09 
                                               -0.637 
                                               0.55 
                                               0.471 
                                               0.643 
                                               6.14e-09 
                                               8.54e-08 
                                               lm.nb 
                                               Apoptotic cleavage of cellular proteins, AXIN missense mutants destabilize the destruction complex, Beta-catenin phosphorylation cascade, Deactivation of the beta-catenin transactivating complex, Disassembly of the destruction complex and recruitment of AXIN to the membrane, Misspliced GSK3beta mutants stabilize beta-catenin, S33 mutants of beta-catenin aren.SQT.t phosphorylated, S37 mutants of beta-catenin aren.SQT.t phosphorylated, S45 mutants of beta-catenin aren.SQT.t phosphorylated, T41 mutants of beta-catenin aren.SQT.t phosphorylated, Truncations of AMER1 destabilize the destruction complex 
                                               SEBELIUS_2_C4636 
                                               APC 
                                               mRNA 
                                               0 
                                               
                                               
                                               Apoptotic cleavage of cellular proteins;Misspliced GSK3beta mutants stabilize beta-catenin;APC truncation mutants have impaired AXIN binding;T41 mutants of beta-catenin aren.SQT.t phosphorylated;AXIN missense mutants destabilize the destruction complex;S45 mutants of beta-catenin aren.SQT.t phosphorylated;Deactivation of the beta-catenin transactivating complex;Disassembly of the destruction complex and recruitment of AXIN to the membrane;S37 mutants of beta-catenin aren.SQT.t phosphorylated;S33 mutants of beta-catenin aren.SQT.t phosphorylated;Beta-catenin phosphorylation cascade;Truncations of AMER1 destabilize the destruction complex 
                                               hsa04310;hsa04390;hsa04550;hsa04810;hsa05166;hsa05200;hsa05206;hsa05210;hsa05213;hsa05217 
                                               APC 
                                               NA 
                                               slateblue1 
                                               16 
                                               -7.49565217391304 
                                             
                                             
                                               SPTAN1-mRNA 
                                               -0.389 
                                               0.093 
                                               -0.571 
                                               -0.206 
                                               0.764 
                                               0.673 
                                               0.867 
                                               0.000171 
                                               0.00162 
                                               lm.nb 
                                               Apoptotic cleavage of cellular proteins, ARMS-mediated activation, DAP12 signaling, Downstream signal transduction, FCERI mediated MAPK activation, Frs2-mediated activation, Gastrin-CREB signalling pathway via PKC and MAPK, Interleukin receptor SHC signaling, IRS-related events triggered by IGF1R, Nephrin interactions, Signaling by ERBB2, Signaling by PDGF, Signaling by SCF-KIT, VEGFA-VEGFR2 Pathway, VEGFR2 mediated cell proliferation 
                                               SEBELIUS_2_C4636 
                                               SPTAN1 
                                               mRNA 
                                               0 
                                               
                                               
                                               Nephrin interactions;Caspase-mediated cleavage of cytoskeletal proteins;Downstream signal transduction;VEGFR2 mediated cell proliferation;Signaling by SCF-KIT;Frs2-mediated activation;ARMS-mediated activation;Gastrin-CREB signalling pathway via PKC and MAPK;Interleukin receptor SHC signaling;FCERI mediated MAPK activation;DAP12 signaling;Signaling by PDGF;VEGFA-VEGFR2 Pathway;Signaling by ERBB2;IRS-related events triggered by IGF1R;Apoptotic cleavage of cellular proteins 
                                               hsa04530 
                                               SPTAN1 
                                               NA 
                                               slateblue1 
                                               16 
                                               -4.18279569892473 
                                             
                                             
                                               TJP1-mRNA 
                                               -0.137 
                                               0.0858 
                                               -0.305 
                                               0.0315 
                                               0.91 
                                               0.809 
                                               1.02 
                                               0.12 
                                               0.858 
                                               lm.nb 
                                               Apoptotic cleavage of cellular proteins 
                                               SEBELIUS_2_C4636 
                                               TJP1 
                                               mRNA 
                                               0 
                                               
                                               
                                               Apoptotic cleavage of cellular proteins 
                                               hsa04520;hsa04530;hsa04540;hsa05110;hsa05120;hsa05132 
                                               TJP1 
                                               NA 
                                               azure4 
                                               1 
                                               -1.5967365967366 
                                             
                                             
                                               ROCK1-mRNA 
                                               -0.032 
                                               0.0634 
                                               -0.156 
                                               0.0922 
                                               0.978 
                                               0.897 
                                               1.07 
                                               0.617 
                                               1 
                                               loglinear 
                                               Apoptotic cleavage of cellular proteins, EPHA-mediated growth cone collapse, EPHB-mediated forward signaling, G alpha (1213) signalling events, RHO GTPases Activate ROCKs, Sema4D induced cell migration and growth-cone collapse, VEGFA-VEGFR2 Pathway 
                                               SEBELIUS_2_C4636 
                                               ROCK1 
                                               mRNA 
                                               0 
                                               
                                               
                                               EPHB-mediated forward signaling;EPHA-mediated growth cone collapse;RHO GTPases Activate ROCKs;Apoptotic cleavage of cellular proteins;G alpha (1213) signalling events;VEGFA-VEGFR2 Pathway;Sema4D induced cell migration and growth-cone collapse 
                                               hsa04022;hsa04024;hsa04062;hsa04071;hsa04270;hsa04350;hsa04360;hsa04510;hsa04611;hsa04670;hsa04810;hsa04921;hsa05130;hsa05131;hsa05132;hsa05200;hsa05205;hsa05206 
                                               ROCK1 
                                               NA 
                                               azure4 
                                               1 
                                               -0.504731861198738 
                                             
                                             
                                               TJP2-mRNA 
                                               0.00744 
                                               0.155 
                                               -0.297 
                                               0.312 
                                               1.01 
                                               0.814 
                                               1.24 
                                               0.962 
                                               1 
                                               lm.nb 
                                               Apoptotic cleavage of cellular proteins 
                                               SEBELIUS_2_C4636 
                                               TJP2 
                                               mRNA 
                                               0 
                                               
                                               
                                               Apoptotic cleavage of cellular proteins 
                                               hsa04530;hsa05110 
                                               TJP2 
                                               NA 
                                               azure4 
                                               1 
                                               0.048 
                                             
                                             
                                               CTNNB1-mRNA 
                                               0.0018 
                                               0.163 
                                               -0.317 
                                               0.32 
                                               1 
                                               0.803 
                                               1.25 
                                               0.991 
                                               1 
                                               lm.nb 
                                               Apoptotic cleavage of cellular proteins, Beta-catenin phosphorylation cascade, Deactivation of the beta-catenin transactivating complex, Disassembly of the destruction complex and recruitment of AXIN to the membrane, Formation of the beta-catenin.CL.TCF transactivating complex, Misspliced GSK3beta mutants stabilize beta-catenin, Repression of WNT target genes, RHO GTPases activate IQGAPs, S37 mutants of beta-catenin aren.SQT.t phosphorylated, S45 mutants of beta-catenin aren.SQT.t phosphorylated, T41 mutants of beta-catenin aren.SQT.t phosphorylated, VEGFA-VEGFR2 Pathway, VEGFR2 mediated vascular permeability 
                                               SEBELIUS_2_C4636 
                                               CTNNB1 
                                               mRNA 
                                               0 
                                               
                                               
                                               Disassembly of the destruction complex and recruitment of AXIN to the membrane;S37 mutants of beta-catenin aren.SQT.t phosphorylated;T41 mutants of beta-catenin aren.SQT.t phosphorylated;S45 mutants of beta-catenin aren.SQT.t phosphorylated;Binding of TCFLEF.CL.CTNNB1 to target gene promoters;RHO GTPases activate IQGAPs;Deactivation of the beta-catenin transactivating complex;Repression of WNT target genes;Apoptotic cleavage of cell adhesion  proteins;Adherens junctions interactions;Misspliced GSK3beta mutants stabilize beta-catenin;VEGFR2 mediated vascular permeability;Beta-catenin phosphorylation cascade;Synthesis# secretion# and inactivation of Glucagon-like Peptide-1 (GLP-1);Formation of the beta-catenin.CL.TCF transactivating complex;Apoptotic cleavage of cellular proteins;VEGFA-VEGFR2 Pathway 
                                               hsa04015;hsa04310;hsa04390;hsa04510;hsa04520;hsa04530;hsa04550;hsa04670;hsa04916;hsa04919;hsa05100;hsa05130;hsa05166;hsa05200;hsa05205;hsa05210;hsa05213;hsa05215;hsa05216;hsa05217;hsa05412 
                                               CTNNB1 
                                               NA 
                                               azure4 
                                               1 
                                               0.0110429447852761 
                                             
                                           
                                           DE Results - Apoptotic cleavage of cellular proteins genes - Line: Caco2 vs.Fibro  Table displaying each sample's global significance scores and directed global significance scores as defined in the heatmaps above. The global significance score is calculated as the square root of the mean squared t-statistic for the genes in a gene set, with t-statistics coming from the linear regression underlying our differential expression analysis. The directed global significance score is calculated as the square root of the mean signed squared t-statistic for the genes in a gene set, with t-statistics coming from the linear regression underlying our differential expression analysis.   
                                      
                                       
                                  
                                
                                
                                  
                                       
                                      
                                           Volcano Plot: Line: HEPG2 vs.Fibro   More Plot Information                Volcano Plot: Line: HEPG2 vs.Fibro  Volcano plot displaying each gene's -log10(p-value) and log2 fold change for the selected covariate. Highly statistically significant genes fall at the top of the plot, and highly differentially expressed genes fall to either side. Genes within the selected gene set are highlighted in orange. Horizontal lines indicate various False Discovery Rate (FDR) thresholds.   
                                           DE Results - Apoptotic cleavage of cellular proteins genes - Line: HEPG2 vs.Fibro   More Plot Information      Download CSV Data     
                                           
                                             
                                             
                                           
                                           
                                             
                                               Probe Label 
                                               Log2 fold change 
                                               std error (log2) 
                                               Lower confidence limit (log2) 
                                               Upper confidence limit (log2) 
                                               Linear fold change 
                                               Lower confidence limit (linear) 
                                               Upper confidence limit (linear) 
                                               P-value 
                                               BY.p.value 
                                               method 
                                               Gene.sets 
                                               Codeset.Name 
                                               Probe.Label 
                                               Analyte.Type 
                                               Is.Control 
                                               Control.Type 
                                               Related.Probes 
                                               Probe.Annotation 
                                               KEGG.Pathways 
                                               Official.Gene.Name 
                                               Control.Conc 
                                               volcanocol 
                                               volcanopch 
                                               tstats.all 
                                             
                                           
                                           
                                             
                                               TJP2-mRNA 
                                               2.41 
                                               0.166 
                                               2.09 
                                               2.74 
                                               5.33 
                                               4.26 
                                               6.68 
                                               7.15e-17 
                                               2.67e-15 
                                               lm.nb 
                                               Apoptotic cleavage of cellular proteins 
                                               SEBELIUS_2_C4636 
                                               TJP2 
                                               mRNA 
                                               0 
                                               
                                               
                                               Apoptotic cleavage of cellular proteins 
                                               hsa04530;hsa05110 
                                               TJP2 
                                               NA 
                                               slateblue1 
                                               16 
                                               14.5180722891566 
                                             
                                             
                                               OCLN-mRNA 
                                               3.13 
                                               0.237 
                                               2.66 
                                               3.59 
                                               8.73 
                                               6.32 
                                               12 
                                               1.53e-15 
                                               4.62e-14 
                                               lm.nb 
                                               Apoptotic cleavage of cellular proteins 
                                               SEBELIUS_2_C4636 
                                               OCLN 
                                               mRNA 
                                               0 
                                               
                                               
                                               Apoptotic cleavage of cellular proteins 
                                               hsa04514;hsa04530;hsa04670;hsa05130;hsa05160 
                                               OCLN 
                                               NA 
                                               slateblue1 
                                               16 
                                               13.2067510548523 
                                             
                                             
                                               CTNNB1-mRNA 
                                               1.77 
                                               0.178 
                                               1.42 
                                               2.12 
                                               3.4 
                                               2.67 
                                               4.33 
                                               6.08e-12 
                                               1.41e-10 
                                               lm.nb 
                                               Apoptotic cleavage of cellular proteins, Beta-catenin phosphorylation cascade, Deactivation of the beta-catenin transactivating complex, Disassembly of the destruction complex and recruitment of AXIN to the membrane, Formation of the beta-catenin.CL.TCF transactivating complex, Misspliced GSK3beta mutants stabilize beta-catenin, Repression of WNT target genes, RHO GTPases activate IQGAPs, S37 mutants of beta-catenin aren.SQT.t phosphorylated, S45 mutants of beta-catenin aren.SQT.t phosphorylated, T41 mutants of beta-catenin aren.SQT.t phosphorylated, VEGFA-VEGFR2 Pathway, VEGFR2 mediated vascular permeability 
                                               SEBELIUS_2_C4636 
                                               CTNNB1 
                                               mRNA 
                                               0 
                                               
                                               
                                               Disassembly of the destruction complex and recruitment of AXIN to the membrane;S37 mutants of beta-catenin aren.SQT.t phosphorylated;T41 mutants of beta-catenin aren.SQT.t phosphorylated;S45 mutants of beta-catenin aren.SQT.t phosphorylated;Binding of TCFLEF.CL.CTNNB1 to target gene promoters;RHO GTPases activate IQGAPs;Deactivation of the beta-catenin transactivating complex;Repression of WNT target genes;Apoptotic cleavage of cell adhesion  proteins;Adherens junctions interactions;Misspliced GSK3beta mutants stabilize beta-catenin;VEGFR2 mediated vascular permeability;Beta-catenin phosphorylation cascade;Synthesis# secretion# and inactivation of Glucagon-like Peptide-1 (GLP-1);Formation of the beta-catenin.CL.TCF transactivating complex;Apoptotic cleavage of cellular proteins;VEGFA-VEGFR2 Pathway 
                                               hsa04015;hsa04310;hsa04390;hsa04510;hsa04520;hsa04530;hsa04550;hsa04670;hsa04916;hsa04919;hsa05100;hsa05130;hsa05166;hsa05200;hsa05205;hsa05210;hsa05213;hsa05215;hsa05216;hsa05217;hsa05412 
                                               CTNNB1 
                                               NA 
                                               slateblue1 
                                               16 
                                               9.9438202247191 
                                             
                                             
                                               TJP1-mRNA 
                                               -0.671 
                                               0.0945 
                                               -0.856 
                                               -0.486 
                                               0.628 
                                               0.552 
                                               0.714 
                                               2.07e-08 
                                               3.03e-07 
                                               lm.nb 
                                               Apoptotic cleavage of cellular proteins 
                                               SEBELIUS_2_C4636 
                                               TJP1 
                                               mRNA 
                                               0 
                                               
                                               
                                               Apoptotic cleavage of cellular proteins 
                                               hsa04520;hsa04530;hsa04540;hsa05110;hsa05120;hsa05132 
                                               TJP1 
                                               NA 
                                               slateblue1 
                                               16 
                                               -7.1005291005291 
                                             
                                             
                                               PTK2-mRNA 
                                               0.703 
                                               0.119 
                                               0.471 
                                               0.936 
                                               1.63 
                                               1.39 
                                               1.91 
                                               7.91e-07 
                                               1e-05 
                                               lm.nb 
                                               Apoptotic cleavage of cellular proteins, ARMS-mediated activation, DAP12 signaling, DCC mediated attractive signaling, Downstream signal transduction, EPHA-mediated growth cone collapse, EPHB-mediated forward signaling, FCERI mediated MAPK activation, Frs2-mediated activation, Gastrin-CREB signalling pathway via PKC and MAPK, Integrin alphaIIb beta3 signaling, Interleukin receptor SHC signaling, IRS-related events triggered by IGF1R, p130Cas linkage to MAPK signaling for integrins, RAFMAP kinase cascade, Regulation of actin dynamics for phagocytic cup formation, RHO GTPases Activate WASPs and WAVEs, Signaling by ERBB2, Signaling by PDGF, Signaling by SCF-KIT, VEGFA-VEGFR2 Pathway, VEGFR2 mediated cell proliferation 
                                               SEBELIUS_2_C4636 
                                               PTK2 
                                               mRNA 
                                               0 
                                               
                                               
                                               Apoptotic cleavage of cellular proteins;RAFMAP kinase cascade;RHO GTPases Activate WASPs and WAVEs;DCC mediated attractive signaling;p130Cas linkage to MAPK signaling for integrins;EPHB-mediated forward signaling;EPHA-mediated growth cone collapse;Regulation of actin dynamics for phagocytic cup formation;Downstream signal transduction;VEGFR2 mediated cell proliferation;Signaling by SCF-KIT;Frs2-mediated activation;ARMS-mediated activation;Gastrin-CREB signalling pathway via PKC and MAPK;Interleukin receptor SHC signaling;FCERI mediated MAPK activation;DAP12 signaling;Signaling by PDGF;VEGFA-VEGFR2 Pathway;Signaling by ERBB2;IRS-related events triggered by IGF1R;Integrin alphaIIb beta3 signaling 
                                               hsa04012;hsa04062;hsa04151;hsa04360;hsa04370;hsa04510;hsa04670;hsa04810;hsa05100;hsa05146;hsa05200;hsa05202;hsa05205;hsa05222 
                                               PTK2 
                                               NA 
                                               slateblue1 
                                               16 
                                               5.90756302521008 
                                             
                                             
                                               SPTAN1-mRNA 
                                               -0.211 
                                               0.102 
                                               -0.411 
                                               -0.0111 
                                               0.864 
                                               0.752 
                                               0.992 
                                               0.0456 
                                               0.35 
                                               lm.nb 
                                               Apoptotic cleavage of cellular proteins, ARMS-mediated activation, DAP12 signaling, Downstream signal transduction, FCERI mediated MAPK activation, Frs2-mediated activation, Gastrin-CREB signalling pathway via PKC and MAPK, Interleukin receptor SHC signaling, IRS-related events triggered by IGF1R, Nephrin interactions, Signaling by ERBB2, Signaling by PDGF, Signaling by SCF-KIT, VEGFA-VEGFR2 Pathway, VEGFR2 mediated cell proliferation 
                                               SEBELIUS_2_C4636 
                                               SPTAN1 
                                               mRNA 
                                               0 
                                               
                                               
                                               Nephrin interactions;Caspase-mediated cleavage of cytoskeletal proteins;Downstream signal transduction;VEGFR2 mediated cell proliferation;Signaling by SCF-KIT;Frs2-mediated activation;ARMS-mediated activation;Gastrin-CREB signalling pathway via PKC and MAPK;Interleukin receptor SHC signaling;FCERI mediated MAPK activation;DAP12 signaling;Signaling by PDGF;VEGFA-VEGFR2 Pathway;Signaling by ERBB2;IRS-related events triggered by IGF1R;Apoptotic cleavage of cellular proteins 
                                               hsa04530 
                                               SPTAN1 
                                               NA 
                                               slateblue4 
                                               16 
                                               -2.06862745098039 
                                             
                                             
                                               ROCK1-mRNA 
                                               0.0361 
                                               0.0696 
                                               -0.1 
                                               0.172 
                                               1.03 
                                               0.933 
                                               1.13 
                                               0.606 
                                               1 
                                               loglinear 
                                               Apoptotic cleavage of cellular proteins, EPHA-mediated growth cone collapse, EPHB-mediated forward signaling, G alpha (1213) signalling events, RHO GTPases Activate ROCKs, Sema4D induced cell migration and growth-cone collapse, VEGFA-VEGFR2 Pathway 
                                               SEBELIUS_2_C4636 
                                               ROCK1 
                                               mRNA 
                                               0 
                                               
                                               
                                               EPHB-mediated forward signaling;EPHA-mediated growth cone collapse;RHO GTPases Activate ROCKs;Apoptotic cleavage of cellular proteins;G alpha (1213) signalling events;VEGFA-VEGFR2 Pathway;Sema4D induced cell migration and growth-cone collapse 
                                               hsa04022;hsa04024;hsa04062;hsa04071;hsa04270;hsa04350;hsa04360;hsa04510;hsa04611;hsa04670;hsa04810;hsa04921;hsa05130;hsa05131;hsa05132;hsa05200;hsa05205;hsa05206 
                                               ROCK1 
                                               NA 
                                               azure4 
                                               1 
                                               0.51867816091954 
                                             
                                             
                                               APC-mRNA 
                                               -0.0616 
                                               0.125 
                                               -0.306 
                                               0.183 
                                               0.958 
                                               0.809 
                                               1.14 
                                               0.624 
                                               1 
                                               lm.nb 
                                               Apoptotic cleavage of cellular proteins, AXIN missense mutants destabilize the destruction complex, Beta-catenin phosphorylation cascade, Deactivation of the beta-catenin transactivating complex, Disassembly of the destruction complex and recruitment of AXIN to the membrane, Misspliced GSK3beta mutants stabilize beta-catenin, S33 mutants of beta-catenin aren.SQT.t phosphorylated, S37 mutants of beta-catenin aren.SQT.t phosphorylated, S45 mutants of beta-catenin aren.SQT.t phosphorylated, T41 mutants of beta-catenin aren.SQT.t phosphorylated, Truncations of AMER1 destabilize the destruction complex 
                                               SEBELIUS_2_C4636 
                                               APC 
                                               mRNA 
                                               0 
                                               
                                               
                                               Apoptotic cleavage of cellular proteins;Misspliced GSK3beta mutants stabilize beta-catenin;APC truncation mutants have impaired AXIN binding;T41 mutants of beta-catenin aren.SQT.t phosphorylated;AXIN missense mutants destabilize the destruction complex;S45 mutants of beta-catenin aren.SQT.t phosphorylated;Deactivation of the beta-catenin transactivating complex;Disassembly of the destruction complex and recruitment of AXIN to the membrane;S37 mutants of beta-catenin aren.SQT.t phosphorylated;S33 mutants of beta-catenin aren.SQT.t phosphorylated;Beta-catenin phosphorylation cascade;Truncations of AMER1 destabilize the destruction complex 
                                               hsa04310;hsa04390;hsa04550;hsa04810;hsa05166;hsa05200;hsa05206;hsa05210;hsa05213;hsa05217 
                                               APC 
                                               NA 
                                               azure4 
                                               1 
                                               -0.4928 
                                             
                                           
                                           DE Results - Apoptotic cleavage of cellular proteins genes - Line: HEPG2 vs.Fibro  Table displaying each sample's global significance scores and directed global significance scores as defined in the heatmaps above. The global significance score is calculated as the square root of the mean squared t-statistic for the genes in a gene set, with t-statistics coming from the linear regression underlying our differential expression analysis. The directed global significance score is calculated as the square root of the mean signed squared t-statistic for the genes in a gene set, with t-statistics coming from the linear regression underlying our differential expression analysis.   
                                      
                                       
                                  
                                
                                
                                  
                                       
                                      
                                           Volcano Plot: Line: HT29 vs.Fibro   More Plot Information                Volcano Plot: Line: HT29 vs.Fibro  Volcano plot displaying each gene's -log10(p-value) and log2 fold change for the selected covariate. Highly statistically significant genes fall at the top of the plot, and highly differentially expressed genes fall to either side. Genes within the selected gene set are highlighted in orange. Horizontal lines indicate various False Discovery Rate (FDR) thresholds.   
                                           DE Results - Apoptotic cleavage of cellular proteins genes - Line: HT29 vs.Fibro   More Plot Information      Download CSV Data     
                                           
                                             
                                             
                                           
                                           
                                             
                                               Probe Label 
                                               Log2 fold change 
                                               std error (log2) 
                                               Lower confidence limit (log2) 
                                               Upper confidence limit (log2) 
                                               Linear fold change 
                                               Lower confidence limit (linear) 
                                               Upper confidence limit (linear) 
                                               P-value 
                                               BY.p.value 
                                               method 
                                               Gene.sets 
                                               Codeset.Name 
                                               Probe.Label 
                                               Analyte.Type 
                                               Is.Control 
                                               Control.Type 
                                               Related.Probes 
                                               Probe.Annotation 
                                               KEGG.Pathways 
                                               Official.Gene.Name 
                                               Control.Conc 
                                               volcanocol 
                                               volcanopch 
                                               tstats.all 
                                             
                                           
                                           
                                             
                                               PTK2-mRNA 
                                               2.36 
                                               0.112 
                                               2.14 
                                               2.58 
                                               5.14 
                                               4.41 
                                               5.98 
                                               2.87e-22 
                                               1.7e-20 
                                               lm.nb 
                                               Apoptotic cleavage of cellular proteins, ARMS-mediated activation, DAP12 signaling, DCC mediated attractive signaling, Downstream signal transduction, EPHA-mediated growth cone collapse, EPHB-mediated forward signaling, FCERI mediated MAPK activation, Frs2-mediated activation, Gastrin-CREB signalling pathway via PKC and MAPK, Integrin alphaIIb beta3 signaling, Interleukin receptor SHC signaling, IRS-related events triggered by IGF1R, p130Cas linkage to MAPK signaling for integrins, RAFMAP kinase cascade, Regulation of actin dynamics for phagocytic cup formation, RHO GTPases Activate WASPs and WAVEs, Signaling by ERBB2, Signaling by PDGF, Signaling by SCF-KIT, VEGFA-VEGFR2 Pathway, VEGFR2 mediated cell proliferation 
                                               SEBELIUS_2_C4636 
                                               PTK2 
                                               mRNA 
                                               0 
                                               
                                               
                                               Apoptotic cleavage of cellular proteins;RAFMAP kinase cascade;RHO GTPases Activate WASPs and WAVEs;DCC mediated attractive signaling;p130Cas linkage to MAPK signaling for integrins;EPHB-mediated forward signaling;EPHA-mediated growth cone collapse;Regulation of actin dynamics for phagocytic cup formation;Downstream signal transduction;VEGFR2 mediated cell proliferation;Signaling by SCF-KIT;Frs2-mediated activation;ARMS-mediated activation;Gastrin-CREB signalling pathway via PKC and MAPK;Interleukin receptor SHC signaling;FCERI mediated MAPK activation;DAP12 signaling;Signaling by PDGF;VEGFA-VEGFR2 Pathway;Signaling by ERBB2;IRS-related events triggered by IGF1R;Integrin alphaIIb beta3 signaling 
                                               hsa04012;hsa04062;hsa04151;hsa04360;hsa04370;hsa04510;hsa04670;hsa04810;hsa05100;hsa05146;hsa05200;hsa05202;hsa05205;hsa05222 
                                               PTK2 
                                               NA 
                                               slateblue1 
                                               16 
                                               21.0714285714286 
                                             
                                             
                                               TJP2-mRNA 
                                               2.62 
                                               0.157 
                                               2.31 
                                               2.93 
                                               6.14 
                                               4.96 
                                               7.6 
                                               9.39e-19 
                                               3.6e-17 
                                               lm.nb 
                                               Apoptotic cleavage of cellular proteins 
                                               SEBELIUS_2_C4636 
                                               TJP2 
                                               mRNA 
                                               0 
                                               
                                               
                                               Apoptotic cleavage of cellular proteins 
                                               hsa04530;hsa05110 
                                               TJP2 
                                               NA 
                                               slateblue1 
                                               16 
                                               16.687898089172 
                                             
                                             
                                               OCLN-mRNA 
                                               2.89 
                                               0.227 
                                               2.44 
                                               3.33 
                                               7.4 
                                               5.44 
                                               10.1 
                                               4.43e-15 
                                               9.24e-14 
                                               lm.nb 
                                               Apoptotic cleavage of cellular proteins 
                                               SEBELIUS_2_C4636 
                                               OCLN 
                                               mRNA 
                                               0 
                                               
                                               
                                               Apoptotic cleavage of cellular proteins 
                                               hsa04514;hsa04530;hsa04670;hsa05130;hsa05160 
                                               OCLN 
                                               NA 
                                               slateblue1 
                                               16 
                                               12.7312775330396 
                                             
                                             
                                               TJP1-mRNA 
                                               0.569 
                                               0.0886 
                                               0.395 
                                               0.743 
                                               1.48 
                                               1.32 
                                               1.67 
                                               1.68e-07 
                                               1.87e-06 
                                               lm.nb 
                                               Apoptotic cleavage of cellular proteins 
                                               SEBELIUS_2_C4636 
                                               TJP1 
                                               mRNA 
                                               0 
                                               
                                               
                                               Apoptotic cleavage of cellular proteins 
                                               hsa04520;hsa04530;hsa04540;hsa05110;hsa05120;hsa05132 
                                               TJP1 
                                               NA 
                                               slateblue1 
                                               16 
                                               6.42212189616253 
                                             
                                             
                                               SPTAN1-mRNA 
                                               -0.527 
                                               0.0962 
                                               -0.716 
                                               -0.339 
                                               0.694 
                                               0.609 
                                               0.791 
                                               3.18e-06 
                                               3.24e-05 
                                               lm.nb 
                                               Apoptotic cleavage of cellular proteins, ARMS-mediated activation, DAP12 signaling, Downstream signal transduction, FCERI mediated MAPK activation, Frs2-mediated activation, Gastrin-CREB signalling pathway via PKC and MAPK, Interleukin receptor SHC signaling, IRS-related events triggered by IGF1R, Nephrin interactions, Signaling by ERBB2, Signaling by PDGF, Signaling by SCF-KIT, VEGFA-VEGFR2 Pathway, VEGFR2 mediated cell proliferation 
                                               SEBELIUS_2_C4636 
                                               SPTAN1 
                                               mRNA 
                                               0 
                                               
                                               
                                               Nephrin interactions;Caspase-mediated cleavage of cytoskeletal proteins;Downstream signal transduction;VEGFR2 mediated cell proliferation;Signaling by SCF-KIT;Frs2-mediated activation;ARMS-mediated activation;Gastrin-CREB signalling pathway via PKC and MAPK;Interleukin receptor SHC signaling;FCERI mediated MAPK activation;DAP12 signaling;Signaling by PDGF;VEGFA-VEGFR2 Pathway;Signaling by ERBB2;IRS-related events triggered by IGF1R;Apoptotic cleavage of cellular proteins 
                                               hsa04530 
                                               SPTAN1 
                                               NA 
                                               slateblue1 
                                               16 
                                               -5.47817047817048 
                                             
                                             
                                               APC-mRNA 
                                               0.329 
                                               0.117 
                                               0.1 
                                               0.558 
                                               1.26 
                                               1.07 
                                               1.47 
                                               0.00766 
                                               0.0598 
                                               lm.nb 
                                               Apoptotic cleavage of cellular proteins, AXIN missense mutants destabilize the destruction complex, Beta-catenin phosphorylation cascade, Deactivation of the beta-catenin transactivating complex, Disassembly of the destruction complex and recruitment of AXIN to the membrane, Misspliced GSK3beta mutants stabilize beta-catenin, S33 mutants of beta-catenin aren.SQT.t phosphorylated, S37 mutants of beta-catenin aren.SQT.t phosphorylated, S45 mutants of beta-catenin aren.SQT.t phosphorylated, T41 mutants of beta-catenin aren.SQT.t phosphorylated, Truncations of AMER1 destabilize the destruction complex 
                                               SEBELIUS_2_C4636 
                                               APC 
                                               mRNA 
                                               0 
                                               
                                               
                                               Apoptotic cleavage of cellular proteins;Misspliced GSK3beta mutants stabilize beta-catenin;APC truncation mutants have impaired AXIN binding;T41 mutants of beta-catenin aren.SQT.t phosphorylated;AXIN missense mutants destabilize the destruction complex;S45 mutants of beta-catenin aren.SQT.t phosphorylated;Deactivation of the beta-catenin transactivating complex;Disassembly of the destruction complex and recruitment of AXIN to the membrane;S37 mutants of beta-catenin aren.SQT.t phosphorylated;S33 mutants of beta-catenin aren.SQT.t phosphorylated;Beta-catenin phosphorylation cascade;Truncations of AMER1 destabilize the destruction complex 
                                               hsa04310;hsa04390;hsa04550;hsa04810;hsa05166;hsa05200;hsa05206;hsa05210;hsa05213;hsa05217 
                                               APC 
                                               NA 
                                               slateblue3 
                                               16 
                                               2.81196581196581 
                                             
                                             
                                               ROCK1-mRNA 
                                               -0.159 
                                               0.0655 
                                               -0.287 
                                               -0.0304 
                                               0.896 
                                               0.819 
                                               0.979 
                                               0.0204 
                                               0.155 
                                               loglinear 
                                               Apoptotic cleavage of cellular proteins, EPHA-mediated growth cone collapse, EPHB-mediated forward signaling, G alpha (1213) signalling events, RHO GTPases Activate ROCKs, Sema4D induced cell migration and growth-cone collapse, VEGFA-VEGFR2 Pathway 
                                               SEBELIUS_2_C4636 
                                               ROCK1 
                                               mRNA 
                                               0 
                                               
                                               
                                               EPHB-mediated forward signaling;EPHA-mediated growth cone collapse;RHO GTPases Activate ROCKs;Apoptotic cleavage of cellular proteins;G alpha (1213) signalling events;VEGFA-VEGFR2 Pathway;Sema4D induced cell migration and growth-cone collapse 
                                               hsa04022;hsa04024;hsa04062;hsa04071;hsa04270;hsa04350;hsa04360;hsa04510;hsa04611;hsa04670;hsa04810;hsa04921;hsa05130;hsa05131;hsa05132;hsa05200;hsa05205;hsa05206 
                                               ROCK1 
                                               NA 
                                               slateblue4 
                                               16 
                                               -2.42748091603053 
                                             
                                             
                                               CTNNB1-mRNA 
                                               0.244 
                                               0.168 
                                               -0.0856 
                                               0.573 
                                               1.18 
                                               0.942 
                                               1.49 
                                               0.155 
                                               1 
                                               lm.nb 
                                               Apoptotic cleavage of cellular proteins, Beta-catenin phosphorylation cascade, Deactivation of the beta-catenin transactivating complex, Disassembly of the destruction complex and recruitment of AXIN to the membrane, Formation of the beta-catenin.CL.TCF transactivating complex, Misspliced GSK3beta mutants stabilize beta-catenin, Repression of WNT target genes, RHO GTPases activate IQGAPs, S37 mutants of beta-catenin aren.SQT.t phosphorylated, S45 mutants of beta-catenin aren.SQT.t phosphorylated, T41 mutants of beta-catenin aren.SQT.t phosphorylated, VEGFA-VEGFR2 Pathway, VEGFR2 mediated vascular permeability 
                                               SEBELIUS_2_C4636 
                                               CTNNB1 
                                               mRNA 
                                               0 
                                               
                                               
                                               Disassembly of the destruction complex and recruitment of AXIN to the membrane;S37 mutants of beta-catenin aren.SQT.t phosphorylated;T41 mutants of beta-catenin aren.SQT.t phosphorylated;S45 mutants of beta-catenin aren.SQT.t phosphorylated;Binding of TCFLEF.CL.CTNNB1 to target gene promoters;RHO GTPases activate IQGAPs;Deactivation of the beta-catenin transactivating complex;Repression of WNT target genes;Apoptotic cleavage of cell adhesion  proteins;Adherens junctions interactions;Misspliced GSK3beta mutants stabilize beta-catenin;VEGFR2 mediated vascular permeability;Beta-catenin phosphorylation cascade;Synthesis# secretion# and inactivation of Glucagon-like Peptide-1 (GLP-1);Formation of the beta-catenin.CL.TCF transactivating complex;Apoptotic cleavage of cellular proteins;VEGFA-VEGFR2 Pathway 
                                               hsa04015;hsa04310;hsa04390;hsa04510;hsa04520;hsa04530;hsa04550;hsa04670;hsa04916;hsa04919;hsa05100;hsa05130;hsa05166;hsa05200;hsa05205;hsa05210;hsa05213;hsa05215;hsa05216;hsa05217;hsa05412 
                                               CTNNB1 
                                               NA 
                                               azure4 
                                               1 
                                               1.45238095238095 
                                             
                                           
                                           DE Results - Apoptotic cleavage of cellular proteins genes - Line: HT29 vs.Fibro  Table displaying each sample's global significance scores and directed global significance scores as defined in the heatmaps above. The global significance score is calculated as the square root of the mean squared t-statistic for the genes in a gene set, with t-statistics coming from the linear regression underlying our differential expression analysis. The directed global significance score is calculated as the square root of the mean signed squared t-statistic for the genes in a gene set, with t-statistics coming from the linear regression underlying our differential expression analysis.   
                                      
                                       
                                  
                                
                              
                            
                           
                            
                          
                           
                      
                    
                    
                      
                           
                          
                               Day    Line     
                              
                                  Day  
                              
                               
                              
                                  Line: differential expression in Caco2 vs. baseline of Fibro  
                                  Line: differential expression in HEPG2 vs. baseline of Fibro  
                                  Line: differential expression in HT29 vs. baseline of Fibro  
                              
                              
                            
                              
                                
                                  
                                       
                                      
                                           Volcano Plot: Day   More Plot Information                Volcano Plot: Day  Volcano plot displaying each gene's -log10(p-value) and log2 fold change for the selected covariate. Highly statistically significant genes fall at the top of the plot, and highly differentially expressed genes fall to either side. Genes within the selected gene set are highlighted in orange. Horizontal lines indicate various False Discovery Rate (FDR) thresholds.   
                                           DE Results - ARMS-mediated activation genes - Day   More Plot Information      Download CSV Data     
                                           
                                             
                                             
                                           
                                           
                                             
                                               Probe Label 
                                               Log2 fold change 
                                               std error (log2) 
                                               Lower confidence limit (log2) 
                                               Upper confidence limit (log2) 
                                               Linear fold change 
                                               Lower confidence limit (linear) 
                                               Upper confidence limit (linear) 
                                               P-value 
                                               BY.p.value 
                                               method 
                                               Gene.sets 
                                               Codeset.Name 
                                               Probe.Label 
                                               Analyte.Type 
                                               Is.Control 
                                               Control.Type 
                                               Related.Probes 
                                               Probe.Annotation 
                                               KEGG.Pathways 
                                               Official.Gene.Name 
                                               Control.Conc 
                                               volcanocol 
                                               volcanopch 
                                               tstats.all 
                                             
                                           
                                           
                                             
                                               MAPK1-mRNA 
                                               0.0599 
                                               0.00943 
                                               0.0414 
                                               0.0784 
                                               1.04 
                                               1.03 
                                               1.06 
                                               2.11e-07 
                                               3.74e-05 
                                               loglinear 
                                               ARMS-mediated activation, CREB phosphorylation through the activation of Ras, DAP12 signaling, Downstream signal transduction, FCERI mediated MAPK activation, Frs2-mediated activation, Gastrin-CREB signalling pathway via PKC and MAPK, Interleukin receptor SHC signaling, IRS-related events triggered by IGF1R, MAP2K and MAPK activation, Negative feedback regulation of MAPK pathway, Negative regulation of FGFR1 signaling, Negative regulation of FGFR2 signaling, Negative regulation of FGFR3 signaling, Negative regulation of FGFR4 signaling, Negative regulation of MAPK pathway, RAFMAP kinase cascade, Regulation of HSF1-mediated heat shock response, RHO GTPases Activate WASPs and WAVEs, Signal transduction by L1, Signaling by ERBB2, Signaling by PDGF, Signaling by SCF-KIT, VEGFA-VEGFR2 Pathway, VEGFR2 mediated cell proliferation 
                                               SEBELIUS_2_C4636 
                                               MAPK1 
                                               mRNA 
                                               0 
                                               
                                               
                                               Negative regulation of FGFR1 signaling;Signal transduction by L1;Negative regulation of FGFR2 signaling;MAP2K and MAPK activation;RHO GTPases Activate WASPs and WAVEs;Senescence-Associated Secretory Phenotype (SASP);Regulation of HSF1-mediated heat shock response;Thrombin signalling through proteinase activated receptors (PARs);Negative regulation of FGFR4 signaling;Oxidative Stress Induced Senescence;Signal attenuation;Negative regulation of FGFR3 signaling;Negative feedback regulation of MAPK pathway;Growth hormone receptor signaling;PLC beta mediated events;FCERI mediated MAPK activation;CREB phosphorylation through the activation of Ras;Negative regulation of MAPK pathway;RAFMAP kinase cascade;Downstream signal transduction;VEGFR2 mediated cell proliferation;Signaling by SCF-KIT;Frs2-mediated activation;ARMS-mediated activation;Gastrin-CREB signalling pathway via PKC and MAPK;Interleukin receptor SHC signaling;DAP12 signaling;Signaling by PDGF;VEGFA-VEGFR2 Pathway;Signaling by ERBB2;IRS-related events triggered by IGF1R 
                                               hsa04010;hsa04012;hsa04014;hsa04015;hsa04022;hsa04024;hsa04062;hsa04066;hsa04068;hsa04071;hsa04072;hsa04114;hsa04150;hsa04151;hsa04261;hsa04270;hsa04320;hsa04350;hsa04360;hsa04370;hsa04380;hsa04510;hsa04520;hsa04540;hsa04550;hsa04611;hsa04620;hsa04621;hsa04650;hsa04660;hsa04662;hsa04664;hsa04666;hsa04668;hsa04713;hsa04720;hsa04722;hsa04723;hsa04724;hsa04725;hsa04726;hsa04730;hsa04810;hsa04910;hsa04912;hsa04914;hsa04915;hsa04916;hsa04917;hsa04919;hsa04921;hsa04930;hsa04933;hsa04960;hsa05010;hsa05020;hsa05034;hsa05131;hsa05132;hsa05133;hsa05140;hsa05142;hsa05145;hsa05152;hsa05160;hsa05161;hsa05164;hsa05200;hsa05203;hsa05205;hsa05206;hsa05210;hsa05211;hsa05212;hsa05213;hsa05214;hsa05215;hsa05216;hsa05218;hsa05219;hsa05220;hsa05221;hsa05223;hsa05230;hsa05231 
                                               MAPK1 
                                               NA 
                                               slateblue1 
                                               16 
                                               6.35206786850477 
                                             
                                             
                                               ERBB2-mRNA 
                                               0.185 
                                               0.0329 
                                               0.12 
                                               0.249 
                                               1.14 
                                               1.09 
                                               1.19 
                                               2.12e-06 
                                               0.000181 
                                               lm.nb 
                                               ARMS-mediated activation, DAP12 signaling, Downstream signal transduction, FCERI mediated MAPK activation, Frs2-mediated activation, GAB1 signalosome, Gastrin-CREB signalling pathway via PKC and MAPK, Interleukin receptor SHC signaling, IRS-related events triggered by IGF1R, PI3KAKT activation, RAFMAP kinase cascade, Role of LAT2NTALLAB on calcium mobilization, Sema4D induced cell migration and growth-cone collapse, Signaling by ERBB2, Signaling by PDGF, Signaling by SCF-KIT, VEGFA-VEGFR2 Pathway, VEGFR2 mediated cell proliferation 
                                               SEBELIUS_2_C4636 
                                               ERBB2 
                                               mRNA 
                                               0 
                                               
                                               
                                               RAFMAP kinase cascade;GRB7 events in ERBB2 signaling;PLCG1 events in ERBB2 signaling;Sema4D induced cell migration and growth-cone collapse;Signaling by ERBB2;Downstream signal transduction;GAB1 signalosome;Signaling by SCF-KIT;PI3KAKT activation;Role of LAT2NTALLAB on calcium mobilization;DAP12 signaling;Signaling by PDGF;VEGFR2 mediated cell proliferation;Frs2-mediated activation;ARMS-mediated activation;Gastrin-CREB signalling pathway via PKC and MAPK;Interleukin receptor SHC signaling;FCERI mediated MAPK activation;VEGFA-VEGFR2 Pathway;IRS-related events triggered by IGF1R 
                                               hsa04012;hsa04020;hsa04066;hsa04510;hsa04520;hsa05200;hsa05205;hsa05206;hsa05212;hsa05213;hsa05215;hsa05219;hsa05223;hsa05230 
                                               ERBB2 
                                               NA 
                                               slateblue1 
                                               16 
                                               5.62310030395137 
                                             
                                             
                                               PTK2-mRNA 
                                               0.0792 
                                               0.0142 
                                               0.0514 
                                               0.107 
                                               1.06 
                                               1.04 
                                               1.08 
                                               2.29e-06 
                                               0.000181 
                                               lm.nb 
                                               Apoptotic cleavage of cellular proteins, ARMS-mediated activation, DAP12 signaling, DCC mediated attractive signaling, Downstream signal transduction, EPHA-mediated growth cone collapse, EPHB-mediated forward signaling, FCERI mediated MAPK activation, Frs2-mediated activation, Gastrin-CREB signalling pathway via PKC and MAPK, Integrin alphaIIb beta3 signaling, Interleukin receptor SHC signaling, IRS-related events triggered by IGF1R, p130Cas linkage to MAPK signaling for integrins, RAFMAP kinase cascade, Regulation of actin dynamics for phagocytic cup formation, RHO GTPases Activate WASPs and WAVEs, Signaling by ERBB2, Signaling by PDGF, Signaling by SCF-KIT, VEGFA-VEGFR2 Pathway, VEGFR2 mediated cell proliferation 
                                               SEBELIUS_2_C4636 
                                               PTK2 
                                               mRNA 
                                               0 
                                               
                                               
                                               Apoptotic cleavage of cellular proteins;RAFMAP kinase cascade;RHO GTPases Activate WASPs and WAVEs;DCC mediated attractive signaling;p130Cas linkage to MAPK signaling for integrins;EPHB-mediated forward signaling;EPHA-mediated growth cone collapse;Regulation of actin dynamics for phagocytic cup formation;Downstream signal transduction;VEGFR2 mediated cell proliferation;Signaling by SCF-KIT;Frs2-mediated activation;ARMS-mediated activation;Gastrin-CREB signalling pathway via PKC and MAPK;Interleukin receptor SHC signaling;FCERI mediated MAPK activation;DAP12 signaling;Signaling by PDGF;VEGFA-VEGFR2 Pathway;Signaling by ERBB2;IRS-related events triggered by IGF1R;Integrin alphaIIb beta3 signaling 
                                               hsa04012;hsa04062;hsa04151;hsa04360;hsa04370;hsa04510;hsa04670;hsa04810;hsa05100;hsa05146;hsa05200;hsa05202;hsa05205;hsa05222 
                                               PTK2 
                                               NA 
                                               slateblue1 
                                               16 
                                               5.57746478873239 
                                             
                                             
                                               MAPK3-mRNA 
                                               0.112 
                                               0.0225 
                                               0.0681 
                                               0.156 
                                               1.08 
                                               1.05 
                                               1.11 
                                               1.44e-05 
                                               0.000788 
                                               lm.nb 
                                               ARMS-mediated activation, DAP12 signaling, Downstream signal transduction, ERKs are inactivated, FCERI mediated MAPK activation, Frs2-mediated activation, Gastrin-CREB signalling pathway via PKC and MAPK, Interleukin receptor SHC signaling, IRS-related events triggered by IGF1R, Negative feedback regulation of MAPK pathway, Negative regulation of FGFR3 signaling, Negative regulation of FGFR4 signaling, Negative regulation of MAPK pathway, RAFMAP kinase cascade, Regulation of HSF1-mediated heat shock response, RHO GTPases Activate WASPs and WAVEs, Signaling by ERBB2, Signaling by PDGF, Signaling by SCF-KIT, VEGFA-VEGFR2 Pathway, VEGFR2 mediated cell proliferation 
                                               SEBELIUS_2_C4636 
                                               MAPK3 
                                               mRNA 
                                               0 
                                               
                                               
                                               ERKs are inactivated;ISG15 antiviral mechanism;RHO GTPases Activate WASPs and WAVEs;Senescence-Associated Secretory Phenotype (SASP);Regulation of HSF1-mediated heat shock response;Thrombin signalling through proteinase activated receptors (PARs);Negative regulation of FGFR4 signaling;Oxidative Stress Induced Senescence;Signal attenuation;RNA Polymerase I Promoter Opening;Negative regulation of FGFR3 signaling;Negative feedback regulation of MAPK pathway;Growth hormone receptor signaling;Golgi Cisternae Pericentriolar Stack Reorganization;FCERI mediated MAPK activation;Negative regulation of MAPK pathway;RAFMAP kinase cascade;Downstream signal transduction;VEGFR2 mediated cell proliferation;Signaling by SCF-KIT;Frs2-mediated activation;ARMS-mediated activation;Gastrin-CREB signalling pathway via PKC and MAPK;Interleukin receptor SHC signaling;DAP12 signaling;Signaling by PDGF;VEGFA-VEGFR2 Pathway;Signaling by ERBB2;IRS-related events triggered by IGF1R 
                                               hsa04010;hsa04012;hsa04014;hsa04015;hsa04022;hsa04024;hsa04062;hsa04066;hsa04068;hsa04071;hsa04072;hsa04114;hsa04150;hsa04151;hsa04261;hsa04270;hsa04320;hsa04350;hsa04360;hsa04370;hsa04380;hsa04510;hsa04520;hsa04540;hsa04550;hsa04611;hsa04620;hsa04621;hsa04650;hsa04660;hsa04662;hsa04664;hsa04666;hsa04668;hsa04713;hsa04720;hsa04722;hsa04723;hsa04724;hsa04725;hsa04726;hsa04730;hsa04810;hsa04910;hsa04912;hsa04914;hsa04915;hsa04916;hsa04917;hsa04919;hsa04921;hsa04930;hsa04933;hsa04960;hsa05010;hsa05020;hsa05034;hsa05131;hsa05132;hsa05133;hsa05140;hsa05142;hsa05145;hsa05152;hsa05160;hsa05161;hsa05164;hsa05200;hsa05203;hsa05205;hsa05210;hsa05211;hsa05212;hsa05213;hsa05214;hsa05215;hsa05216;hsa05218;hsa05219;hsa05220;hsa05221;hsa05223;hsa05230;hsa05231 
                                               MAPK3 
                                               NA 
                                               slateblue1 
                                               16 
                                               4.97777777777778 
                                             
                                             
                                               BRAF-mRNA 
                                               0.0805 
                                               0.0222 
                                               0.037 
                                               0.124 
                                               1.06 
                                               1.03 
                                               1.09 
                                               0.000864 
                                               0.0198 
                                               lm.nb 
                                               ARMS-mediated activation, CREB phosphorylation through the activation of Ras, DAP12 signaling, Downstream signal transduction, FCERI mediated MAPK activation, Frs2-mediated activation, Gastrin-CREB signalling pathway via PKC and MAPK, Interleukin receptor SHC signaling, IRS-related events triggered by IGF1R, MAP2K and MAPK activation, Negative feedback regulation of MAPK pathway, Negative regulation of FGFR1 signaling, Negative regulation of FGFR2 signaling, Negative regulation of FGFR3 signaling, Negative regulation of FGFR4 signaling, Negative regulation of MAPK pathway, RAF activation, RAFMAP kinase cascade, Signaling by ERBB2, Signaling by PDGF, Signaling by SCF-KIT, Spry regulation of FGF signaling, VEGFA-VEGFR2 Pathway, VEGFR2 mediated cell proliferation 
                                               SEBELIUS_2_C4636 
                                               BRAF 
                                               mRNA 
                                               0 
                                               
                                               
                                               Spry regulation of FGF signaling;MAP2K and MAPK activation;RAF activation;CREB phosphorylation through the activation of Ras;Negative feedback regulation of MAPK pathway;ARMS-mediated activation;Negative regulation of FGFR3 signaling;Negative regulation of FGFR4 signaling;Negative regulation of FGFR2 signaling;Negative regulation of FGFR1 signaling;RAFMAP kinase cascade;Downstream signal transduction;VEGFR2 mediated cell proliferation;Signaling by SCF-KIT;Frs2-mediated activation;Gastrin-CREB signalling pathway via PKC and MAPK;Interleukin receptor SHC signaling;FCERI mediated MAPK activation;DAP12 signaling;Signaling by PDGF;VEGFA-VEGFR2 Pathway;Signaling by ERBB2;IRS-related events triggered by IGF1R;Negative regulation of MAPK pathway 
                                               hsa04010;hsa04012;hsa04015;hsa04024;hsa04062;hsa04068;hsa04150;hsa04270;hsa04510;hsa04650;hsa04720;hsa04722;hsa04726;hsa04730;hsa04810;hsa04910;hsa04914;hsa05034;hsa05160;hsa05200;hsa05205;hsa05210;hsa05211;hsa05212;hsa05213;hsa05214;hsa05215;hsa05216;hsa05218;hsa05219;hsa05220;hsa05221;hsa05223 
                                               BRAF 
                                               NA 
                                               slateblue2 
                                               16 
                                               3.62612612612613 
                                             
                                             
                                               VCL-mRNA 
                                               -0.101 
                                               0.029 
                                               -0.158 
                                               -0.0442 
                                               0.932 
                                               0.896 
                                               0.97 
                                               0.00128 
                                               0.0268 
                                               lm.nb 
                                               ARMS-mediated activation, DAP12 signaling, Downstream signal transduction, FCERI mediated MAPK activation, Frs2-mediated activation, Gastrin-CREB signalling pathway via PKC and MAPK, Interleukin receptor SHC signaling, IRS-related events triggered by IGF1R, MAP2K and MAPK activation, Platelet degranulation , RAFMAP kinase cascade, Signaling by ERBB2, Signaling by PDGF, Signaling by SCF-KIT, Smooth Muscle Contraction, VEGFA-VEGFR2 Pathway, VEGFR2 mediated cell proliferation 
                                               SEBELIUS_2_C4636 
                                               VCL 
                                               mRNA 
                                               0 
                                               
                                               
                                               Smooth Muscle Contraction;MAP2K and MAPK activation;Platelet degranulation ;RAFMAP kinase cascade;Downstream signal transduction;VEGFR2 mediated cell proliferation;Signaling by SCF-KIT;Frs2-mediated activation;ARMS-mediated activation;Gastrin-CREB signalling pathway via PKC and MAPK;Interleukin receptor SHC signaling;FCERI mediated MAPK activation;DAP12 signaling;Signaling by PDGF;VEGFA-VEGFR2 Pathway;Signaling by ERBB2;IRS-related events triggered by IGF1R 
                                               hsa04510;hsa04520;hsa04670;hsa04810;hsa05100;hsa05131;hsa05146 
                                               VCL 
                                               NA 
                                               slateblue2 
                                               16 
                                               -3.48275862068966 
                                             
                                             
                                               SRC-mRNA 
                                               0.0696 
                                               0.02 
                                               0.0304 
                                               0.109 
                                               1.05 
                                               1.02 
                                               1.08 
                                               0.00132 
                                               0.0268 
                                               lm.nb 
                                               ARMS-mediated activation, CD28 co-stimulation, Cell surface interactions at the vascular wall, CTLA4 inhibitory signaling, DAP12 signaling, Downstream signal transduction, FCERI mediated MAPK activation, Frs2-mediated activation, GAB1 signalosome, Gastrin-CREB signalling pathway via PKC and MAPK, Integrin alphaIIb beta3 signaling, Interleukin receptor SHC signaling, IRS-related events triggered by IGF1R, MAP2K and MAPK activation, Negative regulation of FGFR1 signaling, Negative regulation of FGFR2 signaling, Negative regulation of FGFR3 signaling, Negative regulation of FGFR4 signaling, p130Cas linkage to MAPK signaling for integrins, RAFMAP kinase cascade, Signaling by ERBB2, Signaling by PDGF, Signaling by SCF-KIT, Spry regulation of FGF signaling, VEGFA-VEGFR2 Pathway, VEGFR2 mediated cell proliferation 
                                               SEBELIUS_2_C4636 
                                               SRC 
                                               mRNA 
                                               0 
                                               
                                               
                                               FCGR activation;Spry regulation of FGF signaling;GAB1 signalosome;MAP2K and MAPK activation;CLEC7A (Dectin-1) signaling;CD28 co-stimulation;p130Cas linkage to MAPK signaling for integrins;CTLA4 inhibitory signaling;GRB2.CL.SOS provides linkage to MAPK signaling for Integrins ;Signal regulatory protein (SIRP) family interactions;PECAM1 interactions;Thrombin signalling through proteinase activated receptors (PARs);GP1b-IX-V activation signalling;c-src mediated regulation of Cx43 function and closure of gap junctions;Signaling by ERBB2;VEGFA-VEGFR2 Pathway;Signaling by SCF-KIT;VEGFR2 mediated cell proliferation;RAFMAP kinase cascade;Downstream signal transduction;Frs2-mediated activation;ARMS-mediated activation;Gastrin-CREB signalling pathway via PKC and MAPK;Interleukin receptor SHC signaling;FCERI mediated MAPK activation;DAP12 signaling;Signaling by PDGF;IRS-related events triggered by IGF1R;Negative regulation of FGFR3 signaling;Negative regulation of FGFR4 signaling;Negative regulation of FGFR2 signaling;Negative regulation of FGFR1 signaling;Integrin alphaIIb beta3 signaling;Cell surface interactions at the vascular wall 
                                               hsa04012;hsa04015;hsa04062;hsa04144;hsa04370;hsa04510;hsa04520;hsa04530;hsa04540;hsa04611;hsa04727;hsa04750;hsa04810;hsa04912;hsa04915;hsa04917;hsa04919;hsa04921;hsa05100;hsa05120;hsa05131;hsa05152;hsa05161;hsa05203;hsa05205;hsa05219 
                                               SRC 
                                               NA 
                                               slateblue2 
                                               16 
                                               3.48 
                                             
                                             
                                               SPTAN1-mRNA 
                                               -0.0399 
                                               0.0123 
                                               -0.0641 
                                               -0.0157 
                                               0.973 
                                               0.957 
                                               0.989 
                                               0.00257 
                                               0.0486 
                                               lm.nb 
                                               Apoptotic cleavage of cellular proteins, ARMS-mediated activation, DAP12 signaling, Downstream signal transduction, FCERI mediated MAPK activation, Frs2-mediated activation, Gastrin-CREB signalling pathway via PKC and MAPK, Interleukin receptor SHC signaling, IRS-related events triggered by IGF1R, Nephrin interactions, Signaling by ERBB2, Signaling by PDGF, Signaling by SCF-KIT, VEGFA-VEGFR2 Pathway, VEGFR2 mediated cell proliferation 
                                               SEBELIUS_2_C4636 
                                               SPTAN1 
                                               mRNA 
                                               0 
                                               
                                               
                                               Nephrin interactions;Caspase-mediated cleavage of cytoskeletal proteins;Downstream signal transduction;VEGFR2 mediated cell proliferation;Signaling by SCF-KIT;Frs2-mediated activation;ARMS-mediated activation;Gastrin-CREB signalling pathway via PKC and MAPK;Interleukin receptor SHC signaling;FCERI mediated MAPK activation;DAP12 signaling;Signaling by PDGF;VEGFA-VEGFR2 Pathway;Signaling by ERBB2;IRS-related events triggered by IGF1R;Apoptotic cleavage of cellular proteins 
                                               hsa04530 
                                               SPTAN1 
                                               NA 
                                               slateblue2 
                                               16 
                                               -3.24390243902439 
                                             
                                             
                                               PDGFRB-mRNA 
                                               0.272 
                                               0.0874 
                                               0.101 
                                               0.443 
                                               1.21 
                                               1.07 
                                               1.36 
                                               0.00355 
                                               0.0593 
                                               loglinear 
                                               ARMS-mediated activation, Constitutive Signaling by Aberrant PI3K in Cancer, DAP12 signaling, Downstream signal transduction, FCERI mediated MAPK activation, Frs2-mediated activation, GAB1 signalosome, Gastrin-CREB signalling pathway via PKC and MAPK, Interleukin receptor SHC signaling, IRS-related events triggered by IGF1R, PI3KAKT activation, PIP3 activates AKT signaling, RAFMAP kinase cascade, Role of LAT2NTALLAB on calcium mobilization, Signaling by ERBB2, Signaling by PDGF, Signaling by SCF-KIT, VEGFA-VEGFR2 Pathway, VEGFR2 mediated cell proliferation 
                                               SEBELIUS_2_C4636 
                                               PDGFRB 
                                               mRNA 
                                               0 
                                               
                                               
                                               PIP3 activates AKT signaling;Constitutive Signaling by Aberrant PI3K in Cancer;RAFMAP kinase cascade;Signaling by PDGF;Downstream signal transduction;GAB1 signalosome;Signaling by SCF-KIT;PI3KAKT activation;Role of LAT2NTALLAB on calcium mobilization;DAP12 signaling;Signaling by ERBB2;VEGFR2 mediated cell proliferation;Frs2-mediated activation;ARMS-mediated activation;Gastrin-CREB signalling pathway via PKC and MAPK;Interleukin receptor SHC signaling;FCERI mediated MAPK activation;VEGFA-VEGFR2 Pathway;IRS-related events triggered by IGF1R 
                                               hsa04010;hsa04014;hsa04015;hsa04020;hsa04060;hsa04072;hsa04151;hsa04510;hsa04540;hsa04810;hsa05166;hsa05200;hsa05206;hsa05214;hsa05215;hsa05218;hsa05230;hsa05231 
                                               PDGFRB 
                                               NA 
                                               slateblue3 
                                               16 
                                               3.11212814645309 
                                             
                                             
                                               ACTB-mRNA 
                                               -0.0865 
                                               0.0282 
                                               -0.142 
                                               -0.0313 
                                               0.942 
                                               0.906 
                                               0.979 
                                               0.00397 
                                               0.0647 
                                               lm.nb 
                                               ARMS-mediated activation, Cell-extracellular matrix interactions, DAP12 signaling, Downstream signal transduction, EPH-ephrin mediated repulsion of cells, EPHB-mediated forward signaling, FCERI mediated MAPK activation, Frs2-mediated activation, Gastrin-CREB signalling pathway via PKC and MAPK, Interleukin receptor SHC signaling, IRS-related events triggered by IGF1R, RAFMAP kinase cascade, Regulation of actin dynamics for phagocytic cup formation, RHO GTPases Activate Formins, RHO GTPases activate IQGAPs, Signaling by ERBB2, Signaling by PDGF, Signaling by SCF-KIT, VEGFA-VEGFR2 Pathway, VEGFR2 mediated cell proliferation 
                                               SEBELIUS_2_C4636 
                                               ACTB 
                                               mRNA 
                                               0 
                                               
                                               
                                               EPHB-mediated forward signaling;Interaction between L1 and Ankyrins;DNA Damage Recognition in GG-NER;EPH-ephrin mediated repulsion of cells;Prefoldin mediated transfer of substrate  to CCTTriC;RHO GTPases Activate Formins;RHO GTPases activate IQGAPs;Cell-extracellular matrix interactions;Regulation of actin dynamics for phagocytic cup formation;Folding of actin by CCTTriC;B-WICH complex positively regulates rRNA expression;RAFMAP kinase cascade;Downstream signal transduction;VEGFR2 mediated cell proliferation;Signaling by SCF-KIT;Frs2-mediated activation;ARMS-mediated activation;Gastrin-CREB signalling pathway via PKC and MAPK;Interleukin receptor SHC signaling;FCERI mediated MAPK activation;DAP12 signaling;Signaling by PDGF;VEGFA-VEGFR2 Pathway;Signaling by ERBB2;IRS-related events triggered by IGF1R 
                                               hsa04015;hsa04145;hsa04390;hsa04510;hsa04520;hsa04530;hsa04611;hsa04670;hsa04810;hsa04919;hsa04921;hsa04971;hsa05100;hsa05110;hsa05130;hsa05131;hsa05132;hsa05164;hsa05205;hsa05410;hsa05412;hsa05414;hsa05416 
                                               ACTB 
                                               NA 
                                               slateblue3 
                                               16 
                                               -3.06737588652482 
                                             
                                             
                                               IRS1-mRNA 
                                               0.0979 
                                               0.0323 
                                               0.0346 
                                               0.161 
                                               1.07 
                                               1.02 
                                               1.12 
                                               0.00444 
                                               0.0717 
                                               lm.nb 
                                               ARMS-mediated activation, Constitutive Signaling by Aberrant PI3K in Cancer, DAP12 signaling, Downstream signal transduction, FCERI mediated MAPK activation, Frs2-mediated activation, GAB1 signalosome, Gastrin-CREB signalling pathway via PKC and MAPK, Interleukin receptor SHC signaling, IRS-related events triggered by IGF1R, PI3K Cascade, PI3KAKT activation, RAFMAP kinase cascade, Role of LAT2NTALLAB on calcium mobilization, Signaling by ERBB2, Signaling by PDGF, Signaling by SCF-KIT, VEGFA-VEGFR2 Pathway, VEGFR2 mediated cell proliferation 
                                               SEBELIUS_2_C4636 
                                               IRS1 
                                               mRNA 
                                               0 
                                               
                                               
                                               Constitutive Signaling by Aberrant PI3K in Cancer;RAFMAP kinase cascade;PI3K Cascade;Growth hormone receptor signaling;IRS-related events triggered by IGF1R;PI3KAKT activation;Downstream signal transduction;GAB1 signalosome;Signaling by SCF-KIT;Role of LAT2NTALLAB on calcium mobilization;DAP12 signaling;Signaling by PDGF;Signaling by ERBB2;VEGFR2 mediated cell proliferation;Frs2-mediated activation;ARMS-mediated activation;Gastrin-CREB signalling pathway via PKC and MAPK;Interleukin receptor SHC signaling;FCERI mediated MAPK activation;VEGFA-VEGFR2 Pathway 
                                               hsa04022;hsa04068;hsa04150;hsa04151;hsa04152;hsa04211;hsa04213;hsa04722;hsa04910;hsa04920;hsa04923;hsa04930;hsa04931;hsa04932;hsa04960;hsa05206 
                                               IRS1 
                                               NA 
                                               slateblue3 
                                               16 
                                               3.03095975232198 
                                             
                                             
                                               PDGFRA-mRNA 
                                               -0.0981 
                                               0.0332 
                                               -0.163 
                                               -0.033 
                                               0.934 
                                               0.893 
                                               0.977 
                                               0.00544 
                                               0.0836 
                                               lm.nb 
                                               ARMS-mediated activation, Constitutive Signaling by Aberrant PI3K in Cancer, DAP12 signaling, Downstream signal transduction, FCERI mediated MAPK activation, Frs2-mediated activation, GAB1 signalosome, Gastrin-CREB signalling pathway via PKC and MAPK, Interleukin receptor SHC signaling, IRS-related events triggered by IGF1R, PI3KAKT activation, PIP3 activates AKT signaling, RAFMAP kinase cascade, Role of LAT2NTALLAB on calcium mobilization, Signaling by ERBB2, Signaling by PDGF, Signaling by SCF-KIT, VEGFA-VEGFR2 Pathway, VEGFR2 mediated cell proliferation 
                                               SEBELIUS_2_C4636 
                                               PDGFRA 
                                               mRNA 
                                               0 
                                               
                                               
                                               PIP3 activates AKT signaling;Constitutive Signaling by Aberrant PI3K in Cancer;RAFMAP kinase cascade;Signaling by PDGF;Downstream signal transduction;GAB1 signalosome;Signaling by SCF-KIT;PI3KAKT activation;Role of LAT2NTALLAB on calcium mobilization;DAP12 signaling;Signaling by ERBB2;VEGFR2 mediated cell proliferation;Frs2-mediated activation;ARMS-mediated activation;Gastrin-CREB signalling pathway via PKC and MAPK;Interleukin receptor SHC signaling;FCERI mediated MAPK activation;VEGFA-VEGFR2 Pathway;IRS-related events triggered by IGF1R 
                                               hsa04010;hsa04014;hsa04015;hsa04020;hsa04060;hsa04072;hsa04144;hsa04151;hsa04510;hsa04540;hsa04810;hsa05166;hsa05200;hsa05206;hsa05214;hsa05215;hsa05218;hsa05230;hsa05231 
                                               PDGFRA 
                                               NA 
                                               slateblue3 
                                               16 
                                               -2.95481927710843 
                                             
                                             
                                               EGFR-mRNA 
                                               0.0995 
                                               0.0337 
                                               0.0334 
                                               0.166 
                                               1.07 
                                               1.02 
                                               1.12 
                                               0.00548 
                                               0.0836 
                                               lm.nb 
                                               ARMS-mediated activation, Constitutive Signaling by Aberrant PI3K in Cancer, Constitutive Signaling by EGFRvIII, DAP12 signaling, Downstream signal transduction, FCERI mediated MAPK activation, Frs2-mediated activation, GAB1 signalosome, Gastrin-CREB signalling pathway via PKC and MAPK, Interleukin receptor SHC signaling, IRS-related events triggered by IGF1R, PI3KAKT activation, Role of LAT2NTALLAB on calcium mobilization, Signaling by ERBB2, Signaling by PDGF, Signaling by SCF-KIT, VEGFA-VEGFR2 Pathway, VEGFR2 mediated cell proliferation 
                                               SEBELIUS_2_C4636 
                                               EGFR 
                                               mRNA 
                                               0 
                                               
                                               
                                               Constitutive Signaling by Aberrant PI3K in Cancer;Inhibition of Signaling by Overexpressed EGFR;PLCG1 events in ERBB2 signaling;EGFR interacts with phospholipase C-gamma;EGFR Transactivation by Gastrin;Constitutive Signaling by Ligand-Responsive EGFR Cancer Variants;Constitutive Signaling by EGFRvIII;GAB1 signalosome;Downstream signal transduction;Signaling by SCF-KIT;PI3KAKT activation;Role of LAT2NTALLAB on calcium mobilization;DAP12 signaling;Signaling by PDGF;Signaling by ERBB2;VEGFR2 mediated cell proliferation;Frs2-mediated activation;ARMS-mediated activation;Gastrin-CREB signalling pathway via PKC and MAPK;Interleukin receptor SHC signaling;FCERI mediated MAPK activation;VEGFA-VEGFR2 Pathway;IRS-related events triggered by IGF1R 
                                               hsa04010;hsa04012;hsa04014;hsa04015;hsa04020;hsa04060;hsa04066;hsa04068;hsa04072;hsa04144;hsa04151;hsa04320;hsa04510;hsa04520;hsa04540;hsa04810;hsa04912;hsa04915;hsa04921;hsa05120;hsa05160;hsa05200;hsa05205;hsa05206;hsa05212;hsa05213;hsa05214;hsa05215;hsa05218;hsa05219;hsa05223;hsa05230;hsa05231 
                                               EGFR 
                                               NA 
                                               slateblue3 
                                               16 
                                               2.95252225519288 
                                             
                                             
                                               SOS1-mRNA 
                                               0.0646 
                                               0.0221 
                                               0.0213 
                                               0.108 
                                               1.05 
                                               1.01 
                                               1.08 
                                               0.00587 
                                               0.0886 
                                               lm.nb 
                                               ARMS-mediated activation, Cell surface interactions at the vascular wall, Constitutive Signaling by EGFRvIII, DAP12 signaling, Downstream signal transduction, FCERI mediated MAPK activation, Frs2-mediated activation, Gastrin-CREB signalling pathway via PKC and MAPK, Integrin alphaIIb beta3 signaling, Interleukin receptor SHC signaling, IRS-related events triggered by IGF1R, NRAGE signals death through JNK, RAFMAP kinase cascade, Role of LAT2NTALLAB on calcium mobilization, SHC-mediated cascade.CL.FGFR1, SHC-related events triggered by IGF1R, Signaling by ERBB2, Signaling by FGFR1 in disease, Signaling by FGFR2 in disease, Signaling by FGFR3 in disease, Signaling by FGFR4 in disease, Signaling by PDGF, Signaling by SCF-KIT, Tie2 Signaling, VEGFA-VEGFR2 Pathway, VEGFR2 mediated cell proliferation 
                                               SEBELIUS_2_C4636 
                                               SOS1 
                                               mRNA 
                                               0 
                                               
                                               
                                               Signaling by FGFR2 in disease;Antigen activates B Cell Receptor (BCR) leading to generation of second messengers;SHC-mediated cascade.CL.FGFR3;Signaling by FGFR3 in disease;RAFMAP kinase cascade;SHC-related events triggered by IGF1R;Signaling by FGFR4 in disease;SHC-mediated cascade.CL.FGFR1;GRB2.CL.SOS provides linkage to MAPK signaling for Integrins ;Tie2 Signaling;Role of LAT2NTALLAB on calcium mobilization;Signaling by FGFR1 in disease;Signal attenuation;NRAGE signals death through JNK;SHC-mediated cascade.CL.FGFR4;FCERI mediated Ca.PLS.2 mobilization;Constitutive Signaling by EGFRvIII;FCERI mediated MAPK activation;Signaling by ERBB2;Signaling by SCF-KIT;DAP12 signaling;Gastrin-CREB signalling pathway via PKC and MAPK;Interleukin receptor SHC signaling;IRS-related events triggered by IGF1R;Downstream signal transduction;VEGFR2 mediated cell proliferation;Frs2-mediated activation;ARMS-mediated activation;Signaling by PDGF;VEGFA-VEGFR2 Pathway;Integrin alphaIIb beta3 signaling;Cell surface interactions at the vascular wall 
                                               hsa04010;hsa04012;hsa04014;hsa04062;hsa04068;hsa04072;hsa04151;hsa04320;hsa04510;hsa04540;hsa04630;hsa04650;hsa04660;hsa04662;hsa04664;hsa04722;hsa04810;hsa04910;hsa04912;hsa04915;hsa04917;hsa05034;hsa05160;hsa05200;hsa05205;hsa05206;hsa05211;hsa05213;hsa05214;hsa05215;hsa05220;hsa05221;hsa05223;hsa05231 
                                               SOS1 
                                               NA 
                                               slateblue3 
                                               16 
                                               2.92307692307692 
                                             
                                             
                                               PPP2R1A-mRNA 
                                               0.0412 
                                               0.0143 
                                               0.013 
                                               0.0693 
                                               1.03 
                                               1.01 
                                               1.05 
                                               0.00677 
                                               0.099 
                                               lm.nb 
                                               ARMS-mediated activation, AXIN missense mutants destabilize the destruction complex, Beta-catenin phosphorylation cascade, CTLA4 inhibitory signaling, Cyclin AB1 associated events during G2M transition, Cyclin D associated events in G1, DAP12 signaling, DARPP-32 events, Disassembly of the destruction complex and recruitment of AXIN to the membrane, Downstream signal transduction, FCERI mediated MAPK activation, Frs2-mediated activation, Gastrin-CREB signalling pathway via PKC and MAPK, Interleukin receptor SHC signaling, IRS-related events triggered by IGF1R, MASTL Facilitates Mitotic Progression, Negative regulation of FGFR1 signaling, Negative regulation of FGFR2 signaling, Negative regulation of FGFR3 signaling, Negative regulation of FGFR4 signaling, Nonsense Mediated Decay (NMD) enhanced by the Exon Junction Complex (EJC), RAF activation, RAFMAP kinase cascade, Resolution of Sister Chromatid Cohesion, RHO GTPases Activate Formins, S33 mutants of beta-catenin aren.SQT.t phosphorylated, S37 mutants of beta-catenin aren.SQT.t phosphorylated, S45 mutants of beta-catenin aren.SQT.t phosphorylated, Signaling by ERBB2, Signaling by PDGF, Signaling by SCF-KIT, Spry regulation of FGF signaling, Truncations of AMER1 destabilize the destruction complex, VEGFA-VEGFR2 Pathway, VEGFR2 mediated cell proliferation 
                                               SEBELIUS_2_C4636 
                                               PPP2R1A 
                                               mRNA 
                                               0 
                                               
                                               
                                               MASTL Facilitates Mitotic Progression;Regulation of PLK1 Activity at G2M Transition;Disassembly of the destruction complex and recruitment of AXIN to the membrane;S37 mutants of beta-catenin aren.SQT.t phosphorylated;Cyclin AB1 associated events during G2M transition;RAF activation;RHO GTPases Activate Formins;S33 mutants of beta-catenin aren.SQT.t phosphorylated;Loss of Nlp from mitotic centrosomes;DARPP-32 events;Beta-catenin phosphorylation cascade;Initiation of Nuclear Envelope Reformation;Recruitment of mitotic centrosome proteins and complexes;Spry regulation of FGF signaling;Truncations of AMER1 destabilize the destruction complex;CTLA4 inhibitory signaling;AXIN missense mutants destabilize the destruction complex;S45 mutants of beta-catenin aren.SQT.t phosphorylated;Anchoring of the basal body to the plasma membrane;Nonsense Mediated Decay (NMD) enhanced by the Exon Junction Complex (EJC);Cyclin D associated events in G1;Platelet sensitization by LDL;Resolution of Sister Chromatid Cohesion;Inhibition of replication initiation of damaged DNA by RB1E2F1;RAFMAP kinase cascade;Downstream signal transduction;VEGFR2 mediated cell proliferation;Signaling by SCF-KIT;Frs2-mediated activation;ARMS-mediated activation;Gastrin-CREB signalling pathway via PKC and MAPK;Interleukin receptor SHC signaling;FCERI mediated MAPK activation;DAP12 signaling;Signaling by PDGF;VEGFA-VEGFR2 Pathway;Signaling by ERBB2;IRS-related events triggered by IGF1R;Negative regulation of FGFR3 signaling;Negative regulation of FGFR4 signaling;Negative regulation of FGFR2 signaling;Negative regulation of FGFR1 signaling 
                                               hsa03015;hsa04071;hsa04114;hsa04151;hsa04152;hsa04261;hsa04350;hsa04390;hsa04530;hsa04728;hsa04730;hsa05142;hsa05160 
                                               PPP2R1A 
                                               NA 
                                               slateblue4 
                                               16 
                                               2.88111888111888 
                                             
                                             
                                               RAF1-mRNA 
                                               0.0287 
                                               0.011 
                                               0.00708 
                                               0.0503 
                                               1.02 
                                               1 
                                               1.04 
                                               0.0132 
                                               0.177 
                                               loglinear 
                                               ARMS-mediated activation, CD209 (DC-SIGN) signaling, CREB phosphorylation through the activation of Ras, DAP12 signaling, Downstream signal transduction, FCERI mediated MAPK activation, Frs2-mediated activation, Gastrin-CREB signalling pathway via PKC and MAPK, Interleukin receptor SHC signaling, IRS-related events triggered by IGF1R, Negative feedback regulation of MAPK pathway, Negative regulation of MAPK pathway, RAF activation, RAFMAP kinase cascade, Signaling by ERBB2, Signaling by PDGF, Signaling by SCF-KIT, VEGFA-VEGFR2 Pathway, VEGFR2 mediated cell proliferation 
                                               SEBELIUS_2_C4636 
                                               RAF1 
                                               mRNA 
                                               0 
                                               
                                               
                                               RAF activation;Stimuli-sensing channels;CREB phosphorylation through the activation of Ras;Negative feedback regulation of MAPK pathway;CD209 (DC-SIGN) signaling;RAFMAP kinase cascade;Downstream signal transduction;VEGFR2 mediated cell proliferation;Signaling by SCF-KIT;Frs2-mediated activation;ARMS-mediated activation;Gastrin-CREB signalling pathway via PKC and MAPK;Interleukin receptor SHC signaling;FCERI mediated MAPK activation;DAP12 signaling;Signaling by PDGF;VEGFA-VEGFR2 Pathway;Signaling by ERBB2;IRS-related events triggered by IGF1R;Negative regulation of MAPK pathway 
                                               hsa04010;hsa04012;hsa04014;hsa04015;hsa04022;hsa04024;hsa04062;hsa04068;hsa04071;hsa04072;hsa04151;hsa04270;hsa04370;hsa04510;hsa04540;hsa04550;hsa04630;hsa04650;hsa04660;hsa04662;hsa04664;hsa04666;hsa04720;hsa04722;hsa04726;hsa04730;hsa04810;hsa04910;hsa04912;hsa04914;hsa04915;hsa04916;hsa04917;hsa04919;hsa04921;hsa05034;hsa05152;hsa05160;hsa05161;hsa05164;hsa05200;hsa05205;hsa05206;hsa05210;hsa05211;hsa05212;hsa05213;hsa05214;hsa05215;hsa05218;hsa05219;hsa05220;hsa05221;hsa05223;hsa05230;hsa05231 
                                               RAF1 
                                               NA 
                                               slateblue4 
                                               16 
                                               2.60909090909091 
                                             
                                             
                                               NRAS-mRNA 
                                               -0.03 
                                               0.0116 
                                               -0.0527 
                                               -0.00738 
                                               0.979 
                                               0.964 
                                               0.995 
                                               0.0134 
                                               0.177 
                                               lm.nb 
                                               ARMS-mediated activation, CD209 (DC-SIGN) signaling, Cell surface interactions at the vascular wall, Constitutive Signaling by EGFRvIII, DAP12 signaling, Downstream signal transduction, FCERI mediated MAPK activation, Frs2-mediated activation, Gastrin-CREB signalling pathway via PKC and MAPK, Interleukin receptor SHC signaling, IRS-related events triggered by IGF1R, MAP2K and MAPK activation, RAFMAP kinase cascade, SHC-mediated cascade.CL.FGFR1, SHC-related events triggered by IGF1R, Signaling by ERBB2, Signaling by FGFR1 in disease, Signaling by FGFR2 in disease, Signaling by FGFR3 in disease, Signaling by FGFR4 in disease, Signaling by PDGF, Signaling by SCF-KIT, Tie2 Signaling, VEGFA-VEGFR2 Pathway, VEGFR2 mediated cell proliferation 
                                               SEBELIUS_2_C4636 
                                               NRAS 
                                               mRNA 
                                               0 
                                               
                                               
                                               Signaling by FGFR2 in disease;Regulation of RAS by GAPs;SHC-mediated cascade.CL.FGFR3;Signaling by FGFR3 in disease;MAP2K and MAPK activation;SHC-related events triggered by IGF1R;Signaling by FGFR4 in disease;SHC-mediated cascade.CL.FGFR1;Tie2 Signaling;Signaling by FGFR1 in disease;Activation of RAS in B cells;SHC-mediated cascade.CL.FGFR4;Constitutive Signaling by EGFRvIII;CD209 (DC-SIGN) signaling;FCERI mediated MAPK activation;RAFMAP kinase cascade;Downstream signal transduction;VEGFR2 mediated cell proliferation;Signaling by SCF-KIT;Frs2-mediated activation;ARMS-mediated activation;Gastrin-CREB signalling pathway via PKC and MAPK;Interleukin receptor SHC signaling;DAP12 signaling;Signaling by PDGF;VEGFA-VEGFR2 Pathway;Signaling by ERBB2;IRS-related events triggered by IGF1R;Cell surface interactions at the vascular wall 
                                               hsa04010;hsa04012;hsa04014;hsa04015;hsa04062;hsa04068;hsa04071;hsa04072;hsa04151;hsa04211;hsa04213;hsa04360;hsa04370;hsa04530;hsa04540;hsa04550;hsa04650;hsa04660;hsa04662;hsa04664;hsa04720;hsa04722;hsa04725;hsa04726;hsa04730;hsa04810;hsa04910;hsa04912;hsa04915;hsa04916;hsa04917;hsa04919;hsa04921;hsa04933;hsa05034;hsa05160;hsa05161;hsa05166;hsa05200;hsa05203;hsa05205;hsa05206;hsa05211;hsa05213;hsa05214;hsa05215;hsa05216;hsa05218;hsa05219;hsa05220;hsa05221;hsa05223;hsa05230;hsa05231 
                                               NRAS 
                                               NA 
                                               slateblue4 
                                               16 
                                               -2.58620689655172 
                                             
                                             
                                               HRAS-mRNA 
                                               -0.0795 
                                               0.0324 
                                               -0.143 
                                               -0.016 
                                               0.946 
                                               0.906 
                                               0.989 
                                               0.019 
                                               0.24 
                                               lm.nb 
                                               ARMS-mediated activation, CD209 (DC-SIGN) signaling, Cell surface interactions at the vascular wall, Constitutive Signaling by EGFRvIII, CREB phosphorylation through the activation of Ras, DAP12 signaling, Downstream signal transduction, FCERI mediated MAPK activation, Frs2-mediated activation, Gastrin-CREB signalling pathway via PKC and MAPK, Interleukin receptor SHC signaling, IRS-related events triggered by IGF1R, MAP2K and MAPK activation, RAFMAP kinase cascade, SHC-mediated cascade.CL.FGFR1, SHC-related events triggered by IGF1R, Signaling by ERBB2, Signaling by FGFR1 in disease, Signaling by FGFR2 in disease, Signaling by FGFR3 in disease, Signaling by FGFR4 in disease, Signaling by PDGF, Signaling by SCF-KIT, Tie2 Signaling, VEGFA-VEGFR2 Pathway, VEGFR2 mediated cell proliferation 
                                               SEBELIUS_2_C4636 
                                               HRAS 
                                               mRNA 
                                               0 
                                               
                                               
                                               Signaling by FGFR2 in disease;Regulation of RAS by GAPs;SHC-mediated cascade.CL.FGFR3;Signaling by FGFR3 in disease;MAP2K and MAPK activation;SHC-related events triggered by IGF1R;Signaling by FGFR4 in disease;SHC-mediated cascade.CL.FGFR1;Tie2 Signaling;Signaling by FGFR1 in disease;Activation of RAS in B cells;SHC-mediated cascade.CL.FGFR4;Constitutive Signaling by EGFRvIII;CD209 (DC-SIGN) signaling;Ras activation uopn Ca2.PLS. infux through NMDA receptor;FCERI mediated MAPK activation;RAFMAP kinase cascade;Downstream signal transduction;VEGFR2 mediated cell proliferation;Signaling by SCF-KIT;Frs2-mediated activation;ARMS-mediated activation;Gastrin-CREB signalling pathway via PKC and MAPK;Interleukin receptor SHC signaling;DAP12 signaling;Signaling by PDGF;VEGFA-VEGFR2 Pathway;Signaling by ERBB2;IRS-related events triggered by IGF1R;Cell surface interactions at the vascular wall;CREB phosphorylation through the activation of Ras 
                                               hsa04010;hsa04012;hsa04014;hsa04015;hsa04062;hsa04068;hsa04071;hsa04072;hsa04144;hsa04151;hsa04211;hsa04213;hsa04360;hsa04370;hsa04510;hsa04530;hsa04540;hsa04550;hsa04630;hsa04650;hsa04660;hsa04662;hsa04664;hsa04720;hsa04722;hsa04725;hsa04726;hsa04730;hsa04810;hsa04910;hsa04912;hsa04915;hsa04916;hsa04917;hsa04919;hsa04921;hsa04933;hsa05034;hsa05160;hsa05161;hsa05166;hsa05200;hsa05203;hsa05205;hsa05206;hsa05211;hsa05213;hsa05214;hsa05215;hsa05216;hsa05218;hsa05219;hsa05220;hsa05221;hsa05223;hsa05230;hsa05231 
                                               HRAS 
                                               NA 
                                               slateblue4 
                                               16 
                                               -2.4537037037037 
                                             
                                             
                                               CRK-mRNA 
                                               -0.0293 
                                               0.0128 
                                               -0.0543 
                                               -0.00418 
                                               0.98 
                                               0.963 
                                               0.997 
                                               0.0281 
                                               0.319 
                                               lm.nb 
                                               ARMS-mediated activation, Downstream signal transduction, Integrin alphaIIb beta3 signaling, p130Cas linkage to MAPK signaling for integrins, Regulation of actin dynamics for phagocytic cup formation, Regulation of signaling by CBL, Signaling by PDGF, VEGFA-VEGFR2 Pathway 
                                               SEBELIUS_2_C4636 
                                               CRK 
                                               mRNA 
                                               0 
                                               
                                               
                                               ARMS-mediated activation;Downstream signal transduction;Regulation of signaling by CBL;p130Cas linkage to MAPK signaling for integrins;VEGFA-VEGFR2 Pathway;Regulation of actin dynamics for phagocytic cup formation;Signaling by PDGF;Integrin alphaIIb beta3 signaling 
                                               hsa04010;hsa04012;hsa04015;hsa04062;hsa04510;hsa04666;hsa04722;hsa04810;hsa04910;hsa05100;hsa05131;hsa05200;hsa05206;hsa05211;hsa05220 
                                               CRK 
                                               NA 
                                               slateblue4 
                                               16 
                                               -2.2890625 
                                             
                                             
                                               PPP2CB-mRNA 
                                               -0.0296 
                                               0.0132 
                                               -0.0554 
                                               -0.0037 
                                               0.98 
                                               0.962 
                                               0.997 
                                               0.0311 
                                               0.348 
                                               lm.nb 
                                               ARMS-mediated activation, AXIN missense mutants destabilize the destruction complex, Beta-catenin phosphorylation cascade, CTLA4 inhibitory signaling, Cyclin AB1 associated events during G2M transition, Cyclin D associated events in G1, DAP12 signaling, DARPP-32 events, Disassembly of the destruction complex and recruitment of AXIN to the membrane, Downstream signal transduction, ERKs are inactivated, FCERI mediated MAPK activation, Frs2-mediated activation, Gastrin-CREB signalling pathway via PKC and MAPK, Interleukin receptor SHC signaling, IRS-related events triggered by IGF1R, MASTL Facilitates Mitotic Progression, Misspliced GSK3beta mutants stabilize beta-catenin, Negative regulation of FGFR1 signaling, Negative regulation of FGFR2 signaling, Negative regulation of FGFR3 signaling, Negative regulation of FGFR4 signaling, Negative regulation of MAPK pathway, RAF activation, RAFMAP kinase cascade, Resolution of Sister Chromatid Cohesion, RHO GTPases Activate Formins, S33 mutants of beta-catenin aren.SQT.t phosphorylated, S37 mutants of beta-catenin aren.SQT.t phosphorylated, S45 mutants of beta-catenin aren.SQT.t phosphorylated, Signaling by ERBB2, Signaling by PDGF, Signaling by SCF-KIT, Spry regulation of FGF signaling, T41 mutants of beta-catenin aren.SQT.t phosphorylated, Truncations of AMER1 destabilize the destruction complex, VEGFA-VEGFR2 Pathway, VEGFR2 mediated cell proliferation 
                                               SEBELIUS_2_C4636 
                                               PPP2CB 
                                               mRNA 
                                               0 
                                               
                                               
                                               MASTL Facilitates Mitotic Progression;Misspliced GSK3beta mutants stabilize beta-catenin;T41 mutants of beta-catenin aren.SQT.t phosphorylated;ERKs are inactivated;Glycolysis;Negative regulation of MAPK pathway;Disassembly of the destruction complex and recruitment of AXIN to the membrane;S37 mutants of beta-catenin aren.SQT.t phosphorylated;Cyclin AB1 associated events during G2M transition;RAF activation;RHO GTPases Activate Formins;S33 mutants of beta-catenin aren.SQT.t phosphorylated;DARPP-32 events;Beta-catenin phosphorylation cascade;Spry regulation of FGF signaling;Truncations of AMER1 destabilize the destruction complex;CTLA4 inhibitory signaling;AXIN missense mutants destabilize the destruction complex;S45 mutants of beta-catenin aren.SQT.t phosphorylated;Cyclin D associated events in G1;Platelet sensitization by LDL;Resolution of Sister Chromatid Cohesion;Inhibition of replication initiation of damaged DNA by RB1E2F1;RAFMAP kinase cascade;Downstream signal transduction;VEGFR2 mediated cell proliferation;Signaling by SCF-KIT;Frs2-mediated activation;ARMS-mediated activation;Gastrin-CREB signalling pathway via PKC and MAPK;Interleukin receptor SHC signaling;FCERI mediated MAPK activation;DAP12 signaling;Signaling by PDGF;VEGFA-VEGFR2 Pathway;Signaling by ERBB2;IRS-related events triggered by IGF1R;Negative regulation of FGFR3 signaling;Negative regulation of FGFR4 signaling;Negative regulation of FGFR2 signaling;Negative regulation of FGFR1 signaling 
                                               hsa03015;hsa04071;hsa04114;hsa04151;hsa04152;hsa04261;hsa04350;hsa04390;hsa04530;hsa04728;hsa04730;hsa05142;hsa05160 
                                               PPP2CB 
                                               NA 
                                               slateblue4 
                                               16 
                                               -2.24242424242424 
                                             
                                           
                                           DE Results - ARMS-mediated activation genes - Day  Table displaying each sample's global significance scores and directed global significance scores as defined in the heatmaps above. The global significance score is calculated as the square root of the mean squared t-statistic for the genes in a gene set, with t-statistics coming from the linear regression underlying our differential expression analysis. The directed global significance score is calculated as the square root of the mean signed squared t-statistic for the genes in a gene set, with t-statistics coming from the linear regression underlying our differential expression analysis.   
                                      
                                       
                                  
                                
                              
                            
                            
                              
                                
                                  
                                       
                                      
                                           Volcano Plot: Line: Caco2 vs.Fibro   More Plot Information                Volcano Plot: Line: Caco2 vs.Fibro  Volcano plot displaying each gene's -log10(p-value) and log2 fold change for the selected covariate. Highly statistically significant genes fall at the top of the plot, and highly differentially expressed genes fall to either side. Genes within the selected gene set are highlighted in orange. Horizontal lines indicate various False Discovery Rate (FDR) thresholds.   
                                           DE Results - ARMS-mediated activation genes - Line: Caco2 vs.Fibro   More Plot Information      Download CSV Data     
                                           
                                             
                                             
                                           
                                           
                                             
                                               Probe Label 
                                               Log2 fold change 
                                               std error (log2) 
                                               Lower confidence limit (log2) 
                                               Upper confidence limit (log2) 
                                               Linear fold change 
                                               Lower confidence limit (linear) 
                                               Upper confidence limit (linear) 
                                               P-value 
                                               BY.p.value 
                                               method 
                                               Gene.sets 
                                               Codeset.Name 
                                               Probe.Label 
                                               Analyte.Type 
                                               Is.Control 
                                               Control.Type 
                                               Related.Probes 
                                               Probe.Annotation 
                                               KEGG.Pathways 
                                               Official.Gene.Name 
                                               Control.Conc 
                                               volcanocol 
                                               volcanopch 
                                               tstats.all 
                                             
                                           
                                           
                                             
                                               KRAS-mRNA 
                                               2.3 
                                               0.0937 
                                               2.11 
                                               2.48 
                                               4.91 
                                               4.32 
                                               5.58 
                                               1.8e-24 
                                               4.27e-22 
                                               lm.nb 
                                               ARMS-mediated activation, CD209 (DC-SIGN) signaling, Cell surface interactions at the vascular wall, Constitutive Signaling by EGFRvIII, DAP12 signaling, Downstream signal transduction, FCERI mediated MAPK activation, Frs2-mediated activation, Gastrin-CREB signalling pathway via PKC and MAPK, Interleukin receptor SHC signaling, IRS-related events triggered by IGF1R, MAP2K and MAPK activation, RAFMAP kinase cascade, SHC-mediated cascade.CL.FGFR1, SHC-related events triggered by IGF1R, Signaling by ERBB2, Signaling by FGFR1 in disease, Signaling by FGFR2 in disease, Signaling by FGFR3 in disease, Signaling by FGFR4 in disease, Signaling by PDGF, Signaling by SCF-KIT, Tie2 Signaling, VEGFA-VEGFR2 Pathway, VEGFR2 mediated cell proliferation 
                                               SEBELIUS_2_C4636 
                                               KRAS 
                                               mRNA 
                                               0 
                                               
                                               
                                               Signaling by FGFR2 in disease;Regulation of RAS by GAPs;SHC-mediated cascade.CL.FGFR3;Signaling by FGFR3 in disease;MAP2K and MAPK activation;SHC-related events triggered by IGF1R;Signaling by FGFR4 in disease;SHC-mediated cascade.CL.FGFR1;Tie2 Signaling;Signaling by FGFR1 in disease;Activation of RAS in B cells;SHC-mediated cascade.CL.FGFR4;Constitutive Signaling by EGFRvIII;CD209 (DC-SIGN) signaling;FCERI mediated MAPK activation;RAFMAP kinase cascade;Downstream signal transduction;VEGFR2 mediated cell proliferation;Signaling by SCF-KIT;Frs2-mediated activation;ARMS-mediated activation;Gastrin-CREB signalling pathway via PKC and MAPK;Interleukin receptor SHC signaling;DAP12 signaling;Signaling by PDGF;VEGFA-VEGFR2 Pathway;Signaling by ERBB2;IRS-related events triggered by IGF1R;Cell surface interactions at the vascular wall 
                                               hsa04010;hsa04012;hsa04014;hsa04015;hsa04062;hsa04068;hsa04071;hsa04072;hsa04151;hsa04211;hsa04213;hsa04320;hsa04360;hsa04370;hsa04530;hsa04540;hsa04550;hsa04650;hsa04660;hsa04662;hsa04664;hsa04720;hsa04722;hsa04725;hsa04726;hsa04730;hsa04810;hsa04910;hsa04912;hsa04914;hsa04915;hsa04916;hsa04917;hsa04919;hsa04921;hsa04933;hsa04960;hsa05034;hsa05160;hsa05161;hsa05166;hsa05200;hsa05203;hsa05205;hsa05206;hsa05210;hsa05211;hsa05212;hsa05213;hsa05214;hsa05215;hsa05216;hsa05218;hsa05219;hsa05220;hsa05221;hsa05223;hsa05230;hsa05231 
                                               KRAS 
                                               NA 
                                               slateblue1 
                                               16 
                                               24.5464247598719 
                                             
                                             
                                               PPP2R1B-mRNA 
                                               1.57 
                                               0.104 
                                               1.37 
                                               1.78 
                                               2.98 
                                               2.59 
                                               3.43 
                                               1.73e-17 
                                               6.65e-16 
                                               lm.nb 
                                               ARMS-mediated activation, AXIN missense mutants destabilize the destruction complex, Beta-catenin phosphorylation cascade, CTLA4 inhibitory signaling, Cyclin AB1 associated events during G2M transition, Cyclin D associated events in G1, DAP12 signaling, DARPP-32 events, Disassembly of the destruction complex and recruitment of AXIN to the membrane, Downstream signal transduction, ERKs are inactivated, FCERI mediated MAPK activation, Frs2-mediated activation, Gastrin-CREB signalling pathway via PKC and MAPK, Interleukin receptor SHC signaling, IRS-related events triggered by IGF1R, MASTL Facilitates Mitotic Progression, Misspliced GSK3beta mutants stabilize beta-catenin, Negative regulation of MAPK pathway, RAF activation, RAFMAP kinase cascade, Resolution of Sister Chromatid Cohesion, RHO GTPases Activate Formins, S33 mutants of beta-catenin aren.SQT.t phosphorylated, S37 mutants of beta-catenin aren.SQT.t phosphorylated, S45 mutants of beta-catenin aren.SQT.t phosphorylated, Signaling by ERBB2, Signaling by PDGF, Signaling by SCF-KIT, T41 mutants of beta-catenin aren.SQT.t phosphorylated, Truncations of AMER1 destabilize the destruction complex, VEGFA-VEGFR2 Pathway, VEGFR2 mediated cell proliferation 
                                               SEBELIUS_2_C4636 
                                               PPP2R1B 
                                               mRNA 
                                               0 
                                               
                                               
                                               MASTL Facilitates Mitotic Progression;Misspliced GSK3beta mutants stabilize beta-catenin;T41 mutants of beta-catenin aren.SQT.t phosphorylated;ERKs are inactivated;Glycolysis;Negative regulation of MAPK pathway;Disassembly of the destruction complex and recruitment of AXIN to the membrane;S37 mutants of beta-catenin aren.SQT.t phosphorylated;Cyclin AB1 associated events during G2M transition;RAF activation;RHO GTPases Activate Formins;S33 mutants of beta-catenin aren.SQT.t phosphorylated;DARPP-32 events;Beta-catenin phosphorylation cascade;Truncations of AMER1 destabilize the destruction complex;CTLA4 inhibitory signaling;AXIN missense mutants destabilize the destruction complex;S45 mutants of beta-catenin aren.SQT.t phosphorylated;Cyclin D associated events in G1;Platelet sensitization by LDL;Resolution of Sister Chromatid Cohesion;Inhibition of replication initiation of damaged DNA by RB1E2F1;RAFMAP kinase cascade;Downstream signal transduction;VEGFR2 mediated cell proliferation;Signaling by SCF-KIT;Frs2-mediated activation;ARMS-mediated activation;Gastrin-CREB signalling pathway via PKC and MAPK;Interleukin receptor SHC signaling;FCERI mediated MAPK activation;DAP12 signaling;Signaling by PDGF;VEGFA-VEGFR2 Pathway;Signaling by ERBB2;IRS-related events triggered by IGF1R 
                                               hsa03015;hsa04071;hsa04114;hsa04151;hsa04152;hsa04261;hsa04350;hsa04390;hsa04530;hsa04728;hsa04730;hsa05142;hsa05160 
                                               PPP2R1B 
                                               NA 
                                               slateblue1 
                                               16 
                                               15.0961538461538 
                                             
                                             
                                               VCL-mRNA 
                                               -3.08 
                                               0.218 
                                               -3.51 
                                               -2.66 
                                               0.118 
                                               0.0878 
                                               0.159 
                                               1.66e-16 
                                               5.1e-15 
                                               lm.nb 
                                               ARMS-mediated activation, DAP12 signaling, Downstream signal transduction, FCERI mediated MAPK activation, Frs2-mediated activation, Gastrin-CREB signalling pathway via PKC and MAPK, Interleukin receptor SHC signaling, IRS-related events triggered by IGF1R, MAP2K and MAPK activation, Platelet degranulation , RAFMAP kinase cascade, Signaling by ERBB2, Signaling by PDGF, Signaling by SCF-KIT, Smooth Muscle Contraction, VEGFA-VEGFR2 Pathway, VEGFR2 mediated cell proliferation 
                                               SEBELIUS_2_C4636 
                                               VCL 
                                               mRNA 
                                               0 
                                               
                                               
                                               Smooth Muscle Contraction;MAP2K and MAPK activation;Platelet degranulation ;RAFMAP kinase cascade;Downstream signal transduction;VEGFR2 mediated cell proliferation;Signaling by SCF-KIT;Frs2-mediated activation;ARMS-mediated activation;Gastrin-CREB signalling pathway via PKC and MAPK;Interleukin receptor SHC signaling;FCERI mediated MAPK activation;DAP12 signaling;Signaling by PDGF;VEGFA-VEGFR2 Pathway;Signaling by ERBB2;IRS-related events triggered by IGF1R 
                                               hsa04510;hsa04520;hsa04670;hsa04810;hsa05100;hsa05131;hsa05146 
                                               VCL 
                                               NA 
                                               slateblue1 
                                               16 
                                               -14.1284403669725 
                                             
                                             
                                               FGFR1-mRNA 
                                               -2.92 
                                               0.244 
                                               -3.4 
                                               -2.45 
                                               0.132 
                                               0.0946 
                                               0.183 
                                               2.56e-14 
                                               5.68e-13 
                                               lm.nb 
                                               ARMS-mediated activation, Constitutive Signaling by Aberrant PI3K in Cancer, DAP12 signaling, Downstream signal transduction, FCERI mediated MAPK activation, Frs2-mediated activation, GAB1 signalosome, Gastrin-CREB signalling pathway via PKC and MAPK, Interleukin receptor SHC signaling, IRS-related events triggered by IGF1R, PI3K Cascade, PI3KAKT activation, RAFMAP kinase cascade, Role of LAT2NTALLAB on calcium mobilization, SHC-mediated cascade.CL.FGFR1, Signaling by ERBB2, Signaling by FGFR1 in disease, Signaling by PDGF, Signaling by SCF-KIT, VEGFA-VEGFR2 Pathway, VEGFR2 mediated cell proliferation 
                                               SEBELIUS_2_C4636 
                                               FGFR1 
                                               mRNA 
                                               0 
                                               
                                               
                                               RAFMAP kinase cascade;Constitutive Signaling by Aberrant PI3K in Cancer;SHC-mediated cascade.CL.FGFR1;Phospholipase C-mediated cascade.CL. FGFR1;Signaling by FGFR1 amplification mutants;PI3K Cascade;FGFR1b ligand binding and activation;Signaling by FGFR1 in disease;Downstream signal transduction;GAB1 signalosome;Signaling by SCF-KIT;PI3KAKT activation;Role of LAT2NTALLAB on calcium mobilization;DAP12 signaling;Signaling by PDGF;Signaling by ERBB2;VEGFR2 mediated cell proliferation;Frs2-mediated activation;ARMS-mediated activation;Gastrin-CREB signalling pathway via PKC and MAPK;Interleukin receptor SHC signaling;FCERI mediated MAPK activation;VEGFA-VEGFR2 Pathway;IRS-related events triggered by IGF1R 
                                               hsa04010;hsa04014;hsa04015;hsa04151;hsa04520;hsa04550;hsa04810;hsa05200;hsa05205;hsa05215;hsa05218;hsa05230 
                                               FGFR1 
                                               NA 
                                               slateblue1 
                                               16 
                                               -11.9672131147541 
                                             
                                             
                                               SRC-mRNA 
                                               1.77 
                                               0.154 
                                               1.47 
                                               2.07 
                                               3.41 
                                               2.76 
                                               4.2 
                                               8.97e-14 
                                               1.9e-12 
                                               lm.nb 
                                               ARMS-mediated activation, CD28 co-stimulation, Cell surface interactions at the vascular wall, CTLA4 inhibitory signaling, DAP12 signaling, Downstream signal transduction, FCERI mediated MAPK activation, Frs2-mediated activation, GAB1 signalosome, Gastrin-CREB signalling pathway via PKC and MAPK, Integrin alphaIIb beta3 signaling, Interleukin receptor SHC signaling, IRS-related events triggered by IGF1R, MAP2K and MAPK activation, Negative regulation of FGFR1 signaling, Negative regulation of FGFR2 signaling, Negative regulation of FGFR3 signaling, Negative regulation of FGFR4 signaling, p130Cas linkage to MAPK signaling for integrins, RAFMAP kinase cascade, Signaling by ERBB2, Signaling by PDGF, Signaling by SCF-KIT, Spry regulation of FGF signaling, VEGFA-VEGFR2 Pathway, VEGFR2 mediated cell proliferation 
                                               SEBELIUS_2_C4636 
                                               SRC 
                                               mRNA 
                                               0 
                                               
                                               
                                               FCGR activation;Spry regulation of FGF signaling;GAB1 signalosome;MAP2K and MAPK activation;CLEC7A (Dectin-1) signaling;CD28 co-stimulation;p130Cas linkage to MAPK signaling for integrins;CTLA4 inhibitory signaling;GRB2.CL.SOS provides linkage to MAPK signaling for Integrins ;Signal regulatory protein (SIRP) family interactions;PECAM1 interactions;Thrombin signalling through proteinase activated receptors (PARs);GP1b-IX-V activation signalling;c-src mediated regulation of Cx43 function and closure of gap junctions;Signaling by ERBB2;VEGFA-VEGFR2 Pathway;Signaling by SCF-KIT;VEGFR2 mediated cell proliferation;RAFMAP kinase cascade;Downstream signal transduction;Frs2-mediated activation;ARMS-mediated activation;Gastrin-CREB signalling pathway via PKC and MAPK;Interleukin receptor SHC signaling;FCERI mediated MAPK activation;DAP12 signaling;Signaling by PDGF;IRS-related events triggered by IGF1R;Negative regulation of FGFR3 signaling;Negative regulation of FGFR4 signaling;Negative regulation of FGFR2 signaling;Negative regulation of FGFR1 signaling;Integrin alphaIIb beta3 signaling;Cell surface interactions at the vascular wall 
                                               hsa04012;hsa04015;hsa04062;hsa04144;hsa04370;hsa04510;hsa04520;hsa04530;hsa04540;hsa04611;hsa04727;hsa04750;hsa04810;hsa04912;hsa04915;hsa04917;hsa04919;hsa04921;hsa05100;hsa05120;hsa05131;hsa05152;hsa05161;hsa05203;hsa05205;hsa05219 
                                               SRC 
                                               NA 
                                               slateblue1 
                                               16 
                                               11.4935064935065 
                                             
                                             
                                               PDGFA-mRNA 
                                               2.89 
                                               0.257 
                                               2.38 
                                               3.39 
                                               7.39 
                                               5.21 
                                               10.5 
                                               1.86e-13 
                                               3.83e-12 
                                               lm.nb 
                                               ARMS-mediated activation, Constitutive Signaling by Aberrant PI3K in Cancer, DAP12 signaling, Downstream signal transduction, FCERI mediated MAPK activation, Frs2-mediated activation, GAB1 signalosome, Gastrin-CREB signalling pathway via PKC and MAPK, Interleukin receptor SHC signaling, IRS-related events triggered by IGF1R, Non-integrin membrane-ECM interactions, PI3KAKT activation, PIP3 activates AKT signaling, Platelet degranulation , RAFMAP kinase cascade, Role of LAT2NTALLAB on calcium mobilization, Signaling by ERBB2, Signaling by PDGF, Signaling by SCF-KIT, VEGFA-VEGFR2 Pathway, VEGFR2 mediated cell proliferation 
                                               SEBELIUS_2_C4636 
                                               PDGFA 
                                               mRNA 
                                               0 
                                               
                                               
                                               Non-integrin membrane-ECM interactions;PIP3 activates AKT signaling;Constitutive Signaling by Aberrant PI3K in Cancer;RAFMAP kinase cascade;Platelet degranulation ;Signaling by PDGF;Downstream signal transduction;GAB1 signalosome;Signaling by SCF-KIT;PI3KAKT activation;Role of LAT2NTALLAB on calcium mobilization;DAP12 signaling;Signaling by ERBB2;VEGFR2 mediated cell proliferation;Frs2-mediated activation;ARMS-mediated activation;Gastrin-CREB signalling pathway via PKC and MAPK;Interleukin receptor SHC signaling;FCERI mediated MAPK activation;VEGFA-VEGFR2 Pathway;IRS-related events triggered by IGF1R 
                                               hsa04010;hsa04014;hsa04015;hsa04060;hsa04072;hsa04151;hsa04510;hsa04540;hsa04810;hsa05166;hsa05200;hsa05202;hsa05206;hsa05214;hsa05215;hsa05218;hsa05231 
                                               PDGFA 
                                               NA 
                                               slateblue1 
                                               16 
                                               11.2451361867704 
                                             
                                             
                                               BRAF-mRNA 
                                               1.58 
                                               0.17 
                                               1.25 
                                               1.92 
                                               2.99 
                                               2.38 
                                               3.77 
                                               3.06e-11 
                                               5.43e-10 
                                               lm.nb 
                                               ARMS-mediated activation, CREB phosphorylation through the activation of Ras, DAP12 signaling, Downstream signal transduction, FCERI mediated MAPK activation, Frs2-mediated activation, Gastrin-CREB signalling pathway via PKC and MAPK, Interleukin receptor SHC signaling, IRS-related events triggered by IGF1R, MAP2K and MAPK activation, Negative feedback regulation of MAPK pathway, Negative regulation of FGFR1 signaling, Negative regulation of FGFR2 signaling, Negative regulation of FGFR3 signaling, Negative regulation of FGFR4 signaling, Negative regulation of MAPK pathway, RAF activation, RAFMAP kinase cascade, Signaling by ERBB2, Signaling by PDGF, Signaling by SCF-KIT, Spry regulation of FGF signaling, VEGFA-VEGFR2 Pathway, VEGFR2 mediated cell proliferation 
                                               SEBELIUS_2_C4636 
                                               BRAF 
                                               mRNA 
                                               0 
                                               
                                               
                                               Spry regulation of FGF signaling;MAP2K and MAPK activation;RAF activation;CREB phosphorylation through the activation of Ras;Negative feedback regulation of MAPK pathway;ARMS-mediated activation;Negative regulation of FGFR3 signaling;Negative regulation of FGFR4 signaling;Negative regulation of FGFR2 signaling;Negative regulation of FGFR1 signaling;RAFMAP kinase cascade;Downstream signal transduction;VEGFR2 mediated cell proliferation;Signaling by SCF-KIT;Frs2-mediated activation;Gastrin-CREB signalling pathway via PKC and MAPK;Interleukin receptor SHC signaling;FCERI mediated MAPK activation;DAP12 signaling;Signaling by PDGF;VEGFA-VEGFR2 Pathway;Signaling by ERBB2;IRS-related events triggered by IGF1R;Negative regulation of MAPK pathway 
                                               hsa04010;hsa04012;hsa04015;hsa04024;hsa04062;hsa04068;hsa04150;hsa04270;hsa04510;hsa04650;hsa04720;hsa04722;hsa04726;hsa04730;hsa04810;hsa04910;hsa04914;hsa05034;hsa05160;hsa05200;hsa05205;hsa05210;hsa05211;hsa05212;hsa05213;hsa05214;hsa05215;hsa05216;hsa05218;hsa05219;hsa05220;hsa05221;hsa05223 
                                               BRAF 
                                               NA 
                                               slateblue1 
                                               16 
                                               9.29411764705882 
                                             
                                             
                                               ACTG1-mRNA 
                                               -1.29 
                                               0.141 
                                               -1.56 
                                               -1.01 
                                               0.41 
                                               0.339 
                                               0.496 
                                               4.96e-11 
                                               8.29e-10 
                                               lm.nb 
                                               ARMS-mediated activation, Cell-extracellular matrix interactions, DAP12 signaling, Downstream signal transduction, EPH-ephrin mediated repulsion of cells, EPHB-mediated forward signaling, FCERI mediated MAPK activation, Frs2-mediated activation, Gastrin-CREB signalling pathway via PKC and MAPK, Interleukin receptor SHC signaling, IRS-related events triggered by IGF1R, RAFMAP kinase cascade, Regulation of actin dynamics for phagocytic cup formation, RHO GTPases Activate Formins, RHO GTPases activate IQGAPs, RHO GTPases Activate WASPs and WAVEs, Signaling by ERBB2, Signaling by PDGF, Signaling by SCF-KIT, VEGFA-VEGFR2 Pathway, VEGFR2 mediated cell proliferation 
                                               SEBELIUS_2_C4636 
                                               ACTG1 
                                               mRNA 
                                               0 
                                               
                                               
                                               RHO GTPases Activate WASPs and WAVEs;Translocation of GLUT4 to the plasma membrane;Recycling pathway of L1;Interaction between L1 and Ankyrins;EPHB-mediated forward signaling;EPH-ephrin mediated repulsion of cells;RHO GTPases Activate Formins;RHO GTPases activate IQGAPs;Cell-extracellular matrix interactions;Regulation of actin dynamics for phagocytic cup formation;RAFMAP kinase cascade;Downstream signal transduction;VEGFR2 mediated cell proliferation;Signaling by SCF-KIT;Frs2-mediated activation;ARMS-mediated activation;Gastrin-CREB signalling pathway via PKC and MAPK;Interleukin receptor SHC signaling;FCERI mediated MAPK activation;DAP12 signaling;Signaling by PDGF;VEGFA-VEGFR2 Pathway;Signaling by ERBB2;IRS-related events triggered by IGF1R 
                                               hsa04015;hsa04145;hsa04390;hsa04510;hsa04520;hsa04530;hsa04611;hsa04670;hsa04810;hsa04919;hsa04921;hsa05100;hsa05110;hsa05130;hsa05131;hsa05132;hsa05164;hsa05205;hsa05410;hsa05412;hsa05414;hsa05416 
                                               ACTG1 
                                               NA 
                                               slateblue1 
                                               16 
                                               -9.14893617021277 
                                             
                                             
                                               PPP2CB-mRNA 
                                               -0.814 
                                               0.099 
                                               -1.01 
                                               -0.62 
                                               0.569 
                                               0.497 
                                               0.651 
                                               7.15e-10 
                                               1.11e-08 
                                               lm.nb 
                                               ARMS-mediated activation, AXIN missense mutants destabilize the destruction complex, Beta-catenin phosphorylation cascade, CTLA4 inhibitory signaling, Cyclin AB1 associated events during G2M transition, Cyclin D associated events in G1, DAP12 signaling, DARPP-32 events, Disassembly of the destruction complex and recruitment of AXIN to the membrane, Downstream signal transduction, ERKs are inactivated, FCERI mediated MAPK activation, Frs2-mediated activation, Gastrin-CREB signalling pathway via PKC and MAPK, Interleukin receptor SHC signaling, IRS-related events triggered by IGF1R, MASTL Facilitates Mitotic Progression, Misspliced GSK3beta mutants stabilize beta-catenin, Negative regulation of FGFR1 signaling, Negative regulation of FGFR2 signaling, Negative regulation of FGFR3 signaling, Negative regulation of FGFR4 signaling, Negative regulation of MAPK pathway, RAF activation, RAFMAP kinase cascade, Resolution of Sister Chromatid Cohesion, RHO GTPases Activate Formins, S33 mutants of beta-catenin aren.SQT.t phosphorylated, S37 mutants of beta-catenin aren.SQT.t phosphorylated, S45 mutants of beta-catenin aren.SQT.t phosphorylated, Signaling by ERBB2, Signaling by PDGF, Signaling by SCF-KIT, Spry regulation of FGF signaling, T41 mutants of beta-catenin aren.SQT.t phosphorylated, Truncations of AMER1 destabilize the destruction complex, VEGFA-VEGFR2 Pathway, VEGFR2 mediated cell proliferation 
                                               SEBELIUS_2_C4636 
                                               PPP2CB 
                                               mRNA 
                                               0 
                                               
                                               
                                               MASTL Facilitates Mitotic Progression;Misspliced GSK3beta mutants stabilize beta-catenin;T41 mutants of beta-catenin aren.SQT.t phosphorylated;ERKs are inactivated;Glycolysis;Negative regulation of MAPK pathway;Disassembly of the destruction complex and recruitment of AXIN to the membrane;S37 mutants of beta-catenin aren.SQT.t phosphorylated;Cyclin AB1 associated events during G2M transition;RAF activation;RHO GTPases Activate Formins;S33 mutants of beta-catenin aren.SQT.t phosphorylated;DARPP-32 events;Beta-catenin phosphorylation cascade;Spry regulation of FGF signaling;Truncations of AMER1 destabilize the destruction complex;CTLA4 inhibitory signaling;AXIN missense mutants destabilize the destruction complex;S45 mutants of beta-catenin aren.SQT.t phosphorylated;Cyclin D associated events in G1;Platelet sensitization by LDL;Resolution of Sister Chromatid Cohesion;Inhibition of replication initiation of damaged DNA by RB1E2F1;RAFMAP kinase cascade;Downstream signal transduction;VEGFR2 mediated cell proliferation;Signaling by SCF-KIT;Frs2-mediated activation;ARMS-mediated activation;Gastrin-CREB signalling pathway via PKC and MAPK;Interleukin receptor SHC signaling;FCERI mediated MAPK activation;DAP12 signaling;Signaling by PDGF;VEGFA-VEGFR2 Pathway;Signaling by ERBB2;IRS-related events triggered by IGF1R;Negative regulation of FGFR3 signaling;Negative regulation of FGFR4 signaling;Negative regulation of FGFR2 signaling;Negative regulation of FGFR1 signaling 
                                               hsa03015;hsa04071;hsa04114;hsa04151;hsa04152;hsa04261;hsa04350;hsa04390;hsa04530;hsa04728;hsa04730;hsa05142;hsa05160 
                                               PPP2CB 
                                               NA 
                                               slateblue1 
                                               16 
                                               -8.22222222222222 
                                             
                                             
                                               PDGFRA-mRNA 
                                               -1.9 
                                               0.232 
                                               -2.35 
                                               -1.44 
                                               0.268 
                                               0.196 
                                               0.368 
                                               7.84e-10 
                                               1.18e-08 
                                               lm.nb 
                                               ARMS-mediated activation, Constitutive Signaling by Aberrant PI3K in Cancer, DAP12 signaling, Downstream signal transduction, FCERI mediated MAPK activation, Frs2-mediated activation, GAB1 signalosome, Gastrin-CREB signalling pathway via PKC and MAPK, Interleukin receptor SHC signaling, IRS-related events triggered by IGF1R, PI3KAKT activation, PIP3 activates AKT signaling, RAFMAP kinase cascade, Role of LAT2NTALLAB on calcium mobilization, Signaling by ERBB2, Signaling by PDGF, Signaling by SCF-KIT, VEGFA-VEGFR2 Pathway, VEGFR2 mediated cell proliferation 
                                               SEBELIUS_2_C4636 
                                               PDGFRA 
                                               mRNA 
                                               0 
                                               
                                               
                                               PIP3 activates AKT signaling;Constitutive Signaling by Aberrant PI3K in Cancer;RAFMAP kinase cascade;Signaling by PDGF;Downstream signal transduction;GAB1 signalosome;Signaling by SCF-KIT;PI3KAKT activation;Role of LAT2NTALLAB on calcium mobilization;DAP12 signaling;Signaling by ERBB2;VEGFR2 mediated cell proliferation;Frs2-mediated activation;ARMS-mediated activation;Gastrin-CREB signalling pathway via PKC and MAPK;Interleukin receptor SHC signaling;FCERI mediated MAPK activation;VEGFA-VEGFR2 Pathway;IRS-related events triggered by IGF1R 
                                               hsa04010;hsa04014;hsa04015;hsa04020;hsa04060;hsa04072;hsa04144;hsa04151;hsa04510;hsa04540;hsa04810;hsa05166;hsa05200;hsa05206;hsa05214;hsa05215;hsa05218;hsa05230;hsa05231 
                                               PDGFRA 
                                               NA 
                                               slateblue1 
                                               16 
                                               -8.18965517241379 
                                             
                                             
                                               PTK2-mRNA 
                                               0.866 
                                               0.108 
                                               0.654 
                                               1.08 
                                               1.82 
                                               1.57 
                                               2.11 
                                               1.35e-09 
                                               2e-08 
                                               lm.nb 
                                               Apoptotic cleavage of cellular proteins, ARMS-mediated activation, DAP12 signaling, DCC mediated attractive signaling, Downstream signal transduction, EPHA-mediated growth cone collapse, EPHB-mediated forward signaling, FCERI mediated MAPK activation, Frs2-mediated activation, Gastrin-CREB signalling pathway via PKC and MAPK, Integrin alphaIIb beta3 signaling, Interleukin receptor SHC signaling, IRS-related events triggered by IGF1R, p130Cas linkage to MAPK signaling for integrins, RAFMAP kinase cascade, Regulation of actin dynamics for phagocytic cup formation, RHO GTPases Activate WASPs and WAVEs, Signaling by ERBB2, Signaling by PDGF, Signaling by SCF-KIT, VEGFA-VEGFR2 Pathway, VEGFR2 mediated cell proliferation 
                                               SEBELIUS_2_C4636 
                                               PTK2 
                                               mRNA 
                                               0 
                                               
                                               
                                               Apoptotic cleavage of cellular proteins;RAFMAP kinase cascade;RHO GTPases Activate WASPs and WAVEs;DCC mediated attractive signaling;p130Cas linkage to MAPK signaling for integrins;EPHB-mediated forward signaling;EPHA-mediated growth cone collapse;Regulation of actin dynamics for phagocytic cup formation;Downstream signal transduction;VEGFR2 mediated cell proliferation;Signaling by SCF-KIT;Frs2-mediated activation;ARMS-mediated activation;Gastrin-CREB signalling pathway via PKC and MAPK;Interleukin receptor SHC signaling;FCERI mediated MAPK activation;DAP12 signaling;Signaling by PDGF;VEGFA-VEGFR2 Pathway;Signaling by ERBB2;IRS-related events triggered by IGF1R;Integrin alphaIIb beta3 signaling 
                                               hsa04012;hsa04062;hsa04151;hsa04360;hsa04370;hsa04510;hsa04670;hsa04810;hsa05100;hsa05146;hsa05200;hsa05202;hsa05205;hsa05222 
                                               PTK2 
                                               NA 
                                               slateblue1 
                                               16 
                                               8.01851851851852 
                                             
                                             
                                               FYN-mRNA 
                                               -2.64 
                                               0.335 
                                               -3.3 
                                               -1.99 
                                               0.16 
                                               0.102 
                                               0.253 
                                               1.96e-09 
                                               2.84e-08 
                                               lm.nb 
                                               ARMS-mediated activation, CD209 (DC-SIGN) signaling, CD28 co-stimulation, CD28 dependent PI3KAkt signaling, CD28 dependent Vav1 pathway, Cell surface interactions at the vascular wall, Constitutive Signaling by Aberrant PI3K in Cancer, CTLA4 inhibitory signaling, DAP12 signaling, Downstream signal transduction, EPH-ephrin mediated repulsion of cells, FCERI mediated MAPK activation, Frs2-mediated activation, GAB1 signalosome, Gastrin-CREB signalling pathway via PKC and MAPK, GPVI-mediated activation cascade, Interleukin receptor SHC signaling, IRS-related events triggered by IGF1R, Nephrin interactions, PI3KAKT activation, RAFMAP kinase cascade, Regulation of signaling by CBL, Role of LAT2NTALLAB on calcium mobilization, Sema3A PAK dependent Axon repulsion, Signaling by ERBB2, Signaling by PDGF, Signaling by SCF-KIT, VEGFA-VEGFR2 Pathway, VEGFR2 mediated cell proliferation 
                                               SEBELIUS_2_C4636 
                                               FYN 
                                               mRNA 
                                               0 
                                               
                                               
                                               EPH-ephrin mediated repulsion of cells;Regulation of signaling by CBL;FCGR activation;Antigen activates B Cell Receptor (BCR) leading to generation of second messengers;Dectin-2 family;RAFMAP kinase cascade;Constitutive Signaling by Aberrant PI3K in Cancer;CTLA4 inhibitory signaling;CD28 dependent PI3KAkt signaling;PECAM1 interactions;CD28 dependent Vav1 pathway;GPVI-mediated activation cascade;Sema3A PAK dependent Axon repulsion;Nephrin interactions;SEMA3A-Plexin repulsion signaling by inhibiting Integrin adhesion;CRMPs in Sema3A signaling;CD209 (DC-SIGN) signaling;Downstream signal transduction;GAB1 signalosome;Signaling by SCF-KIT;PI3KAKT activation;Role of LAT2NTALLAB on calcium mobilization;DAP12 signaling;Signaling by PDGF;Signaling by ERBB2;VEGFA-VEGFR2 Pathway;VEGFR2 mediated cell proliferation;Frs2-mediated activation;ARMS-mediated activation;Gastrin-CREB signalling pathway via PKC and MAPK;Interleukin receptor SHC signaling;FCERI mediated MAPK activation;IRS-related events triggered by IGF1R;CD28 co-stimulation;Cell surface interactions at the vascular wall 
                                               hsa04071;hsa04072;hsa04360;hsa04380;hsa04510;hsa04520;hsa04611;hsa04650;hsa04660;hsa04664;hsa04725;hsa05020;hsa05130;hsa05162;hsa05416 
                                               FYN 
                                               NA 
                                               slateblue1 
                                               16 
                                               -7.88059701492537 
                                             
                                             
                                               PDGFRB-mRNA 
                                               -4.62 
                                               0.661 
                                               -5.91 
                                               -3.32 
                                               0.0407 
                                               0.0166 
                                               0.0998 
                                               2.89e-08 
                                               3.73e-07 
                                               loglinear 
                                               ARMS-mediated activation, Constitutive Signaling by Aberrant PI3K in Cancer, DAP12 signaling, Downstream signal transduction, FCERI mediated MAPK activation, Frs2-mediated activation, GAB1 signalosome, Gastrin-CREB signalling pathway via PKC and MAPK, Interleukin receptor SHC signaling, IRS-related events triggered by IGF1R, PI3KAKT activation, PIP3 activates AKT signaling, RAFMAP kinase cascade, Role of LAT2NTALLAB on calcium mobilization, Signaling by ERBB2, Signaling by PDGF, Signaling by SCF-KIT, VEGFA-VEGFR2 Pathway, VEGFR2 mediated cell proliferation 
                                               SEBELIUS_2_C4636 
                                               PDGFRB 
                                               mRNA 
                                               0 
                                               
                                               
                                               PIP3 activates AKT signaling;Constitutive Signaling by Aberrant PI3K in Cancer;RAFMAP kinase cascade;Signaling by PDGF;Downstream signal transduction;GAB1 signalosome;Signaling by SCF-KIT;PI3KAKT activation;Role of LAT2NTALLAB on calcium mobilization;DAP12 signaling;Signaling by ERBB2;VEGFR2 mediated cell proliferation;Frs2-mediated activation;ARMS-mediated activation;Gastrin-CREB signalling pathway via PKC and MAPK;Interleukin receptor SHC signaling;FCERI mediated MAPK activation;VEGFA-VEGFR2 Pathway;IRS-related events triggered by IGF1R 
                                               hsa04010;hsa04014;hsa04015;hsa04020;hsa04060;hsa04072;hsa04151;hsa04510;hsa04540;hsa04810;hsa05166;hsa05200;hsa05206;hsa05214;hsa05215;hsa05218;hsa05230;hsa05231 
                                               PDGFRB 
                                               NA 
                                               slateblue1 
                                               16 
                                               -6.98940998487141 
                                             
                                             
                                               ACTB-mRNA 
                                               -1.41 
                                               0.213 
                                               -1.83 
                                               -0.997 
                                               0.375 
                                               0.281 
                                               0.501 
                                               8.48e-08 
                                               1.07e-06 
                                               lm.nb 
                                               ARMS-mediated activation, Cell-extracellular matrix interactions, DAP12 signaling, Downstream signal transduction, EPH-ephrin mediated repulsion of cells, EPHB-mediated forward signaling, FCERI mediated MAPK activation, Frs2-mediated activation, Gastrin-CREB signalling pathway via PKC and MAPK, Interleukin receptor SHC signaling, IRS-related events triggered by IGF1R, RAFMAP kinase cascade, Regulation of actin dynamics for phagocytic cup formation, RHO GTPases Activate Formins, RHO GTPases activate IQGAPs, Signaling by ERBB2, Signaling by PDGF, Signaling by SCF-KIT, VEGFA-VEGFR2 Pathway, VEGFR2 mediated cell proliferation 
                                               SEBELIUS_2_C4636 
                                               ACTB 
                                               mRNA 
                                               0 
                                               
                                               
                                               EPHB-mediated forward signaling;Interaction between L1 and Ankyrins;DNA Damage Recognition in GG-NER;EPH-ephrin mediated repulsion of cells;Prefoldin mediated transfer of substrate  to CCTTriC;RHO GTPases Activate Formins;RHO GTPases activate IQGAPs;Cell-extracellular matrix interactions;Regulation of actin dynamics for phagocytic cup formation;Folding of actin by CCTTriC;B-WICH complex positively regulates rRNA expression;RAFMAP kinase cascade;Downstream signal transduction;VEGFR2 mediated cell proliferation;Signaling by SCF-KIT;Frs2-mediated activation;ARMS-mediated activation;Gastrin-CREB signalling pathway via PKC and MAPK;Interleukin receptor SHC signaling;FCERI mediated MAPK activation;DAP12 signaling;Signaling by PDGF;VEGFA-VEGFR2 Pathway;Signaling by ERBB2;IRS-related events triggered by IGF1R 
                                               hsa04015;hsa04145;hsa04390;hsa04510;hsa04520;hsa04530;hsa04611;hsa04670;hsa04810;hsa04919;hsa04921;hsa04971;hsa05100;hsa05110;hsa05130;hsa05131;hsa05132;hsa05164;hsa05205;hsa05410;hsa05412;hsa05414;hsa05416 
                                               ACTB 
                                               NA 
                                               slateblue1 
                                               16 
                                               -6.61971830985915 
                                             
                                             
                                               SOS1-mRNA 
                                               1.07 
                                               0.168 
                                               0.737 
                                               1.39 
                                               2.09 
                                               1.67 
                                               2.63 
                                               2.11e-07 
                                               2.5e-06 
                                               lm.nb 
                                               ARMS-mediated activation, Cell surface interactions at the vascular wall, Constitutive Signaling by EGFRvIII, DAP12 signaling, Downstream signal transduction, FCERI mediated MAPK activation, Frs2-mediated activation, Gastrin-CREB signalling pathway via PKC and MAPK, Integrin alphaIIb beta3 signaling, Interleukin receptor SHC signaling, IRS-related events triggered by IGF1R, NRAGE signals death through JNK, RAFMAP kinase cascade, Role of LAT2NTALLAB on calcium mobilization, SHC-mediated cascade.CL.FGFR1, SHC-related events triggered by IGF1R, Signaling by ERBB2, Signaling by FGFR1 in disease, Signaling by FGFR2 in disease, Signaling by FGFR3 in disease, Signaling by FGFR4 in disease, Signaling by PDGF, Signaling by SCF-KIT, Tie2 Signaling, VEGFA-VEGFR2 Pathway, VEGFR2 mediated cell proliferation 
                                               SEBELIUS_2_C4636 
                                               SOS1 
                                               mRNA 
                                               0 
                                               
                                               
                                               Signaling by FGFR2 in disease;Antigen activates B Cell Receptor (BCR) leading to generation of second messengers;SHC-mediated cascade.CL.FGFR3;Signaling by FGFR3 in disease;RAFMAP kinase cascade;SHC-related events triggered by IGF1R;Signaling by FGFR4 in disease;SHC-mediated cascade.CL.FGFR1;GRB2.CL.SOS provides linkage to MAPK signaling for Integrins ;Tie2 Signaling;Role of LAT2NTALLAB on calcium mobilization;Signaling by FGFR1 in disease;Signal attenuation;NRAGE signals death through JNK;SHC-mediated cascade.CL.FGFR4;FCERI mediated Ca.PLS.2 mobilization;Constitutive Signaling by EGFRvIII;FCERI mediated MAPK activation;Signaling by ERBB2;Signaling by SCF-KIT;DAP12 signaling;Gastrin-CREB signalling pathway via PKC and MAPK;Interleukin receptor SHC signaling;IRS-related events triggered by IGF1R;Downstream signal transduction;VEGFR2 mediated cell proliferation;Frs2-mediated activation;ARMS-mediated activation;Signaling by PDGF;VEGFA-VEGFR2 Pathway;Integrin alphaIIb beta3 signaling;Cell surface interactions at the vascular wall 
                                               hsa04010;hsa04012;hsa04014;hsa04062;hsa04068;hsa04072;hsa04151;hsa04320;hsa04510;hsa04540;hsa04630;hsa04650;hsa04660;hsa04662;hsa04664;hsa04722;hsa04810;hsa04910;hsa04912;hsa04915;hsa04917;hsa05034;hsa05160;hsa05200;hsa05205;hsa05206;hsa05211;hsa05213;hsa05214;hsa05215;hsa05220;hsa05221;hsa05223;hsa05231 
                                               SOS1 
                                               NA 
                                               slateblue1 
                                               16 
                                               6.36904761904762 
                                             
                                             
                                               MAPK3-mRNA 
                                               -0.769 
                                               0.169 
                                               -1.1 
                                               -0.437 
                                               0.587 
                                               0.466 
                                               0.738 
                                               5.67e-05 
                                               0.000559 
                                               lm.nb 
                                               ARMS-mediated activation, DAP12 signaling, Downstream signal transduction, ERKs are inactivated, FCERI mediated MAPK activation, Frs2-mediated activation, Gastrin-CREB signalling pathway via PKC and MAPK, Interleukin receptor SHC signaling, IRS-related events triggered by IGF1R, Negative feedback regulation of MAPK pathway, Negative regulation of FGFR3 signaling, Negative regulation of FGFR4 signaling, Negative regulation of MAPK pathway, RAFMAP kinase cascade, Regulation of HSF1-mediated heat shock response, RHO GTPases Activate WASPs and WAVEs, Signaling by ERBB2, Signaling by PDGF, Signaling by SCF-KIT, VEGFA-VEGFR2 Pathway, VEGFR2 mediated cell proliferation 
                                               SEBELIUS_2_C4636 
                                               MAPK3 
                                               mRNA 
                                               0 
                                               
                                               
                                               ERKs are inactivated;ISG15 antiviral mechanism;RHO GTPases Activate WASPs and WAVEs;Senescence-Associated Secretory Phenotype (SASP);Regulation of HSF1-mediated heat shock response;Thrombin signalling through proteinase activated receptors (PARs);Negative regulation of FGFR4 signaling;Oxidative Stress Induced Senescence;Signal attenuation;RNA Polymerase I Promoter Opening;Negative regulation of FGFR3 signaling;Negative feedback regulation of MAPK pathway;Growth hormone receptor signaling;Golgi Cisternae Pericentriolar Stack Reorganization;FCERI mediated MAPK activation;Negative regulation of MAPK pathway;RAFMAP kinase cascade;Downstream signal transduction;VEGFR2 mediated cell proliferation;Signaling by SCF-KIT;Frs2-mediated activation;ARMS-mediated activation;Gastrin-CREB signalling pathway via PKC and MAPK;Interleukin receptor SHC signaling;DAP12 signaling;Signaling by PDGF;VEGFA-VEGFR2 Pathway;Signaling by ERBB2;IRS-related events triggered by IGF1R 
                                               hsa04010;hsa04012;hsa04014;hsa04015;hsa04022;hsa04024;hsa04062;hsa04066;hsa04068;hsa04071;hsa04072;hsa04114;hsa04150;hsa04151;hsa04261;hsa04270;hsa04320;hsa04350;hsa04360;hsa04370;hsa04380;hsa04510;hsa04520;hsa04540;hsa04550;hsa04611;hsa04620;hsa04621;hsa04650;hsa04660;hsa04662;hsa04664;hsa04666;hsa04668;hsa04713;hsa04720;hsa04722;hsa04723;hsa04724;hsa04725;hsa04726;hsa04730;hsa04810;hsa04910;hsa04912;hsa04914;hsa04915;hsa04916;hsa04917;hsa04919;hsa04921;hsa04930;hsa04933;hsa04960;hsa05010;hsa05020;hsa05034;hsa05131;hsa05132;hsa05133;hsa05140;hsa05142;hsa05145;hsa05152;hsa05160;hsa05161;hsa05164;hsa05200;hsa05203;hsa05205;hsa05210;hsa05211;hsa05212;hsa05213;hsa05214;hsa05215;hsa05216;hsa05218;hsa05219;hsa05220;hsa05221;hsa05223;hsa05230;hsa05231 
                                               MAPK3 
                                               NA 
                                               slateblue1 
                                               16 
                                               -4.55029585798817 
                                             
                                             
                                               IQGAP1-mRNA 
                                               -0.453 
                                               0.104 
                                               -0.657 
                                               -0.249 
                                               0.73 
                                               0.634 
                                               0.841 
                                               9.92e-05 
                                               0.000958 
                                               lm.nb 
                                               ARMS-mediated activation, DAP12 signaling, Downstream signal transduction, FCERI mediated MAPK activation, Frs2-mediated activation, Gastrin-CREB signalling pathway via PKC and MAPK, Interleukin receptor SHC signaling, IRS-related events triggered by IGF1R, MAP2K and MAPK activation, Nephrin interactions, RAFMAP kinase cascade, RHO GTPases activate IQGAPs, Signaling by ERBB2, Signaling by PDGF, Signaling by SCF-KIT, VEGFA-VEGFR2 Pathway, VEGFR2 mediated cell proliferation 
                                               SEBELIUS_2_C4636 
                                               IQGAP1 
                                               mRNA 
                                               0 
                                               
                                               
                                               MAP2K and MAPK activation;Nephrin interactions;RHO GTPases activate IQGAPs;RAFMAP kinase cascade;Downstream signal transduction;VEGFR2 mediated cell proliferation;Signaling by SCF-KIT;Frs2-mediated activation;ARMS-mediated activation;Gastrin-CREB signalling pathway via PKC and MAPK;Interleukin receptor SHC signaling;FCERI mediated MAPK activation;DAP12 signaling;Signaling by PDGF;VEGFA-VEGFR2 Pathway;Signaling by ERBB2;IRS-related events triggered by IGF1R 
                                               hsa04520;hsa04810;hsa05205 
                                               IQGAP1 
                                               NA 
                                               slateblue1 
                                               16 
                                               -4.35576923076923 
                                             
                                             
                                               IRS1-mRNA 
                                               -1.02 
                                               0.242 
                                               -1.5 
                                               -0.547 
                                               0.492 
                                               0.354 
                                               0.684 
                                               0.000152 
                                               0.00146 
                                               lm.nb 
                                               ARMS-mediated activation, Constitutive Signaling by Aberrant PI3K in Cancer, DAP12 signaling, Downstream signal transduction, FCERI mediated MAPK activation, Frs2-mediated activation, GAB1 signalosome, Gastrin-CREB signalling pathway via PKC and MAPK, Interleukin receptor SHC signaling, IRS-related events triggered by IGF1R, PI3K Cascade, PI3KAKT activation, RAFMAP kinase cascade, Role of LAT2NTALLAB on calcium mobilization, Signaling by ERBB2, Signaling by PDGF, Signaling by SCF-KIT, VEGFA-VEGFR2 Pathway, VEGFR2 mediated cell proliferation 
                                               SEBELIUS_2_C4636 
                                               IRS1 
                                               mRNA 
                                               0 
                                               
                                               
                                               Constitutive Signaling by Aberrant PI3K in Cancer;RAFMAP kinase cascade;PI3K Cascade;Growth hormone receptor signaling;IRS-related events triggered by IGF1R;PI3KAKT activation;Downstream signal transduction;GAB1 signalosome;Signaling by SCF-KIT;Role of LAT2NTALLAB on calcium mobilization;DAP12 signaling;Signaling by PDGF;Signaling by ERBB2;VEGFR2 mediated cell proliferation;Frs2-mediated activation;ARMS-mediated activation;Gastrin-CREB signalling pathway via PKC and MAPK;Interleukin receptor SHC signaling;FCERI mediated MAPK activation;VEGFA-VEGFR2 Pathway 
                                               hsa04022;hsa04068;hsa04150;hsa04151;hsa04152;hsa04211;hsa04213;hsa04722;hsa04910;hsa04920;hsa04923;hsa04930;hsa04931;hsa04932;hsa04960;hsa05206 
                                               IRS1 
                                               NA 
                                               slateblue1 
                                               16 
                                               -4.21487603305785 
                                             
                                             
                                               SPTAN1-mRNA 
                                               -0.389 
                                               0.093 
                                               -0.571 
                                               -0.206 
                                               0.764 
                                               0.673 
                                               0.867 
                                               0.000171 
                                               0.00162 
                                               lm.nb 
                                               Apoptotic cleavage of cellular proteins, ARMS-mediated activation, DAP12 signaling, Downstream signal transduction, FCERI mediated MAPK activation, Frs2-mediated activation, Gastrin-CREB signalling pathway via PKC and MAPK, Interleukin receptor SHC signaling, IRS-related events triggered by IGF1R, Nephrin interactions, Signaling by ERBB2, Signaling by PDGF, Signaling by SCF-KIT, VEGFA-VEGFR2 Pathway, VEGFR2 mediated cell proliferation 
                                               SEBELIUS_2_C4636 
                                               SPTAN1 
                                               mRNA 
                                               0 
                                               
                                               
                                               Nephrin interactions;Caspase-mediated cleavage of cytoskeletal proteins;Downstream signal transduction;VEGFR2 mediated cell proliferation;Signaling by SCF-KIT;Frs2-mediated activation;ARMS-mediated activation;Gastrin-CREB signalling pathway via PKC and MAPK;Interleukin receptor SHC signaling;FCERI mediated MAPK activation;DAP12 signaling;Signaling by PDGF;VEGFA-VEGFR2 Pathway;Signaling by ERBB2;IRS-related events triggered by IGF1R;Apoptotic cleavage of cellular proteins 
                                               hsa04530 
                                               SPTAN1 
                                               NA 
                                               slateblue1 
                                               16 
                                               -4.18279569892473 
                                             
                                             
                                               HRAS-mRNA 
                                               -0.868 
                                               0.244 
                                               -1.35 
                                               -0.391 
                                               0.548 
                                               0.393 
                                               0.763 
                                               0.00102 
                                               0.00903 
                                               lm.nb 
                                               ARMS-mediated activation, CD209 (DC-SIGN) signaling, Cell surface interactions at the vascular wall, Constitutive Signaling by EGFRvIII, CREB phosphorylation through the activation of Ras, DAP12 signaling, Downstream signal transduction, FCERI mediated MAPK activation, Frs2-mediated activation, Gastrin-CREB signalling pathway via PKC and MAPK, Interleukin receptor SHC signaling, IRS-related events triggered by IGF1R, MAP2K and MAPK activation, RAFMAP kinase cascade, SHC-mediated cascade.CL.FGFR1, SHC-related events triggered by IGF1R, Signaling by ERBB2, Signaling by FGFR1 in disease, Signaling by FGFR2 in disease, Signaling by FGFR3 in disease, Signaling by FGFR4 in disease, Signaling by PDGF, Signaling by SCF-KIT, Tie2 Signaling, VEGFA-VEGFR2 Pathway, VEGFR2 mediated cell proliferation 
                                               SEBELIUS_2_C4636 
                                               HRAS 
                                               mRNA 
                                               0 
                                               
                                               
                                               Signaling by FGFR2 in disease;Regulation of RAS by GAPs;SHC-mediated cascade.CL.FGFR3;Signaling by FGFR3 in disease;MAP2K and MAPK activation;SHC-related events triggered by IGF1R;Signaling by FGFR4 in disease;SHC-mediated cascade.CL.FGFR1;Tie2 Signaling;Signaling by FGFR1 in disease;Activation of RAS in B cells;SHC-mediated cascade.CL.FGFR4;Constitutive Signaling by EGFRvIII;CD209 (DC-SIGN) signaling;Ras activation uopn Ca2.PLS. infux through NMDA receptor;FCERI mediated MAPK activation;RAFMAP kinase cascade;Downstream signal transduction;VEGFR2 mediated cell proliferation;Signaling by SCF-KIT;Frs2-mediated activation;ARMS-mediated activation;Gastrin-CREB signalling pathway via PKC and MAPK;Interleukin receptor SHC signaling;DAP12 signaling;Signaling by PDGF;VEGFA-VEGFR2 Pathway;Signaling by ERBB2;IRS-related events triggered by IGF1R;Cell surface interactions at the vascular wall;CREB phosphorylation through the activation of Ras 
                                               hsa04010;hsa04012;hsa04014;hsa04015;hsa04062;hsa04068;hsa04071;hsa04072;hsa04144;hsa04151;hsa04211;hsa04213;hsa04360;hsa04370;hsa04510;hsa04530;hsa04540;hsa04550;hsa04630;hsa04650;hsa04660;hsa04662;hsa04664;hsa04720;hsa04722;hsa04725;hsa04726;hsa04730;hsa04810;hsa04910;hsa04912;hsa04915;hsa04916;hsa04917;hsa04919;hsa04921;hsa04933;hsa05034;hsa05160;hsa05161;hsa05166;hsa05200;hsa05203;hsa05205;hsa05206;hsa05211;hsa05213;hsa05214;hsa05215;hsa05216;hsa05218;hsa05219;hsa05220;hsa05221;hsa05223;hsa05230;hsa05231 
                                               HRAS 
                                               NA 
                                               slateblue1 
                                               16 
                                               -3.55737704918033 
                                             
                                           
                                           DE Results - ARMS-mediated activation genes - Line: Caco2 vs.Fibro  Table displaying each sample's global significance scores and directed global significance scores as defined in the heatmaps above. The global significance score is calculated as the square root of the mean squared t-statistic for the genes in a gene set, with t-statistics coming from the linear regression underlying our differential expression analysis. The directed global significance score is calculated as the square root of the mean signed squared t-statistic for the genes in a gene set, with t-statistics coming from the linear regression underlying our differential expression analysis.   
                                      
                                       
                                  
                                
                                
                                  
                                       
                                      
                                           Volcano Plot: Line: HEPG2 vs.Fibro   More Plot Information                Volcano Plot: Line: HEPG2 vs.Fibro  Volcano plot displaying each gene's -log10(p-value) and log2 fold change for the selected covariate. Highly statistically significant genes fall at the top of the plot, and highly differentially expressed genes fall to either side. Genes within the selected gene set are highlighted in orange. Horizontal lines indicate various False Discovery Rate (FDR) thresholds.   
                                           DE Results - ARMS-mediated activation genes - Line: HEPG2 vs.Fibro   More Plot Information      Download CSV Data     
                                           
                                             
                                             
                                           
                                           
                                             
                                               Probe Label 
                                               Log2 fold change 
                                               std error (log2) 
                                               Lower confidence limit (log2) 
                                               Upper confidence limit (log2) 
                                               Linear fold change 
                                               Lower confidence limit (linear) 
                                               Upper confidence limit (linear) 
                                               P-value 
                                               BY.p.value 
                                               method 
                                               Gene.sets 
                                               Codeset.Name 
                                               Probe.Label 
                                               Analyte.Type 
                                               Is.Control 
                                               Control.Type 
                                               Related.Probes 
                                               Probe.Annotation 
                                               KEGG.Pathways 
                                               Official.Gene.Name 
                                               Control.Conc 
                                               volcanocol 
                                               volcanopch 
                                               tstats.all 
                                             
                                           
                                           
                                             
                                               PDGFRA-mRNA 
                                               -7.73 
                                               0.294 
                                               -8.31 
                                               -7.16 
                                               0.0047 
                                               0.00315 
                                               0.00701 
                                               1.54e-25 
                                               2.43e-23 
                                               lm.nb 
                                               ARMS-mediated activation, Constitutive Signaling by Aberrant PI3K in Cancer, DAP12 signaling, Downstream signal transduction, FCERI mediated MAPK activation, Frs2-mediated activation, GAB1 signalosome, Gastrin-CREB signalling pathway via PKC and MAPK, Interleukin receptor SHC signaling, IRS-related events triggered by IGF1R, PI3KAKT activation, PIP3 activates AKT signaling, RAFMAP kinase cascade, Role of LAT2NTALLAB on calcium mobilization, Signaling by ERBB2, Signaling by PDGF, Signaling by SCF-KIT, VEGFA-VEGFR2 Pathway, VEGFR2 mediated cell proliferation 
                                               SEBELIUS_2_C4636 
                                               PDGFRA 
                                               mRNA 
                                               0 
                                               
                                               
                                               PIP3 activates AKT signaling;Constitutive Signaling by Aberrant PI3K in Cancer;RAFMAP kinase cascade;Signaling by PDGF;Downstream signal transduction;GAB1 signalosome;Signaling by SCF-KIT;PI3KAKT activation;Role of LAT2NTALLAB on calcium mobilization;DAP12 signaling;Signaling by ERBB2;VEGFR2 mediated cell proliferation;Frs2-mediated activation;ARMS-mediated activation;Gastrin-CREB signalling pathway via PKC and MAPK;Interleukin receptor SHC signaling;FCERI mediated MAPK activation;VEGFA-VEGFR2 Pathway;IRS-related events triggered by IGF1R 
                                               hsa04010;hsa04014;hsa04015;hsa04020;hsa04060;hsa04072;hsa04144;hsa04151;hsa04510;hsa04540;hsa04810;hsa05166;hsa05200;hsa05206;hsa05214;hsa05215;hsa05218;hsa05230;hsa05231 
                                               PDGFRA 
                                               NA 
                                               slateblue1 
                                               16 
                                               -26.2925170068027 
                                             
                                             
                                               VCL-mRNA 
                                               -3.46 
                                               0.24 
                                               -3.93 
                                               -2.99 
                                               0.0909 
                                               0.0657 
                                               0.126 
                                               8.72e-17 
                                               3.17e-15 
                                               lm.nb 
                                               ARMS-mediated activation, DAP12 signaling, Downstream signal transduction, FCERI mediated MAPK activation, Frs2-mediated activation, Gastrin-CREB signalling pathway via PKC and MAPK, Interleukin receptor SHC signaling, IRS-related events triggered by IGF1R, MAP2K and MAPK activation, Platelet degranulation , RAFMAP kinase cascade, Signaling by ERBB2, Signaling by PDGF, Signaling by SCF-KIT, Smooth Muscle Contraction, VEGFA-VEGFR2 Pathway, VEGFR2 mediated cell proliferation 
                                               SEBELIUS_2_C4636 
                                               VCL 
                                               mRNA 
                                               0 
                                               
                                               
                                               Smooth Muscle Contraction;MAP2K and MAPK activation;Platelet degranulation ;RAFMAP kinase cascade;Downstream signal transduction;VEGFR2 mediated cell proliferation;Signaling by SCF-KIT;Frs2-mediated activation;ARMS-mediated activation;Gastrin-CREB signalling pathway via PKC and MAPK;Interleukin receptor SHC signaling;FCERI mediated MAPK activation;DAP12 signaling;Signaling by PDGF;VEGFA-VEGFR2 Pathway;Signaling by ERBB2;IRS-related events triggered by IGF1R 
                                               hsa04510;hsa04520;hsa04670;hsa04810;hsa05100;hsa05131;hsa05146 
                                               VCL 
                                               NA 
                                               slateblue1 
                                               16 
                                               -14.4166666666667 
                                             
                                             
                                               SRC-mRNA 
                                               2.38 
                                               0.168 
                                               2.05 
                                               2.71 
                                               5.2 
                                               4.14 
                                               6.53 
                                               1.71e-16 
                                               6.08e-15 
                                               lm.nb 
                                               ARMS-mediated activation, CD28 co-stimulation, Cell surface interactions at the vascular wall, CTLA4 inhibitory signaling, DAP12 signaling, Downstream signal transduction, FCERI mediated MAPK activation, Frs2-mediated activation, GAB1 signalosome, Gastrin-CREB signalling pathway via PKC and MAPK, Integrin alphaIIb beta3 signaling, Interleukin receptor SHC signaling, IRS-related events triggered by IGF1R, MAP2K and MAPK activation, Negative regulation of FGFR1 signaling, Negative regulation of FGFR2 signaling, Negative regulation of FGFR3 signaling, Negative regulation of FGFR4 signaling, p130Cas linkage to MAPK signaling for integrins, RAFMAP kinase cascade, Signaling by ERBB2, Signaling by PDGF, Signaling by SCF-KIT, Spry regulation of FGF signaling, VEGFA-VEGFR2 Pathway, VEGFR2 mediated cell proliferation 
                                               SEBELIUS_2_C4636 
                                               SRC 
                                               mRNA 
                                               0 
                                               
                                               
                                               FCGR activation;Spry regulation of FGF signaling;GAB1 signalosome;MAP2K and MAPK activation;CLEC7A (Dectin-1) signaling;CD28 co-stimulation;p130Cas linkage to MAPK signaling for integrins;CTLA4 inhibitory signaling;GRB2.CL.SOS provides linkage to MAPK signaling for Integrins ;Signal regulatory protein (SIRP) family interactions;PECAM1 interactions;Thrombin signalling through proteinase activated receptors (PARs);GP1b-IX-V activation signalling;c-src mediated regulation of Cx43 function and closure of gap junctions;Signaling by ERBB2;VEGFA-VEGFR2 Pathway;Signaling by SCF-KIT;VEGFR2 mediated cell proliferation;RAFMAP kinase cascade;Downstream signal transduction;Frs2-mediated activation;ARMS-mediated activation;Gastrin-CREB signalling pathway via PKC and MAPK;Interleukin receptor SHC signaling;FCERI mediated MAPK activation;DAP12 signaling;Signaling by PDGF;IRS-related events triggered by IGF1R;Negative regulation of FGFR3 signaling;Negative regulation of FGFR4 signaling;Negative regulation of FGFR2 signaling;Negative regulation of FGFR1 signaling;Integrin alphaIIb beta3 signaling;Cell surface interactions at the vascular wall 
                                               hsa04012;hsa04015;hsa04062;hsa04144;hsa04370;hsa04510;hsa04520;hsa04530;hsa04540;hsa04611;hsa04727;hsa04750;hsa04810;hsa04912;hsa04915;hsa04917;hsa04919;hsa04921;hsa05100;hsa05120;hsa05131;hsa05152;hsa05161;hsa05203;hsa05205;hsa05219 
                                               SRC 
                                               NA 
                                               slateblue1 
                                               16 
                                               14.1666666666667 
                                             
                                             
                                               NRAS-mRNA 
                                               1.09 
                                               0.0962 
                                               0.904 
                                               1.28 
                                               2.13 
                                               1.87 
                                               2.43 
                                               1.28e-13 
                                               3.29e-12 
                                               lm.nb 
                                               ARMS-mediated activation, CD209 (DC-SIGN) signaling, Cell surface interactions at the vascular wall, Constitutive Signaling by EGFRvIII, DAP12 signaling, Downstream signal transduction, FCERI mediated MAPK activation, Frs2-mediated activation, Gastrin-CREB signalling pathway via PKC and MAPK, Interleukin receptor SHC signaling, IRS-related events triggered by IGF1R, MAP2K and MAPK activation, RAFMAP kinase cascade, SHC-mediated cascade.CL.FGFR1, SHC-related events triggered by IGF1R, Signaling by ERBB2, Signaling by FGFR1 in disease, Signaling by FGFR2 in disease, Signaling by FGFR3 in disease, Signaling by FGFR4 in disease, Signaling by PDGF, Signaling by SCF-KIT, Tie2 Signaling, VEGFA-VEGFR2 Pathway, VEGFR2 mediated cell proliferation 
                                               SEBELIUS_2_C4636 
                                               NRAS 
                                               mRNA 
                                               0 
                                               
                                               
                                               Signaling by FGFR2 in disease;Regulation of RAS by GAPs;SHC-mediated cascade.CL.FGFR3;Signaling by FGFR3 in disease;MAP2K and MAPK activation;SHC-related events triggered by IGF1R;Signaling by FGFR4 in disease;SHC-mediated cascade.CL.FGFR1;Tie2 Signaling;Signaling by FGFR1 in disease;Activation of RAS in B cells;SHC-mediated cascade.CL.FGFR4;Constitutive Signaling by EGFRvIII;CD209 (DC-SIGN) signaling;FCERI mediated MAPK activation;RAFMAP kinase cascade;Downstream signal transduction;VEGFR2 mediated cell proliferation;Signaling by SCF-KIT;Frs2-mediated activation;ARMS-mediated activation;Gastrin-CREB signalling pathway via PKC and MAPK;Interleukin receptor SHC signaling;DAP12 signaling;Signaling by PDGF;VEGFA-VEGFR2 Pathway;Signaling by ERBB2;IRS-related events triggered by IGF1R;Cell surface interactions at the vascular wall 
                                               hsa04010;hsa04012;hsa04014;hsa04015;hsa04062;hsa04068;hsa04071;hsa04072;hsa04151;hsa04211;hsa04213;hsa04360;hsa04370;hsa04530;hsa04540;hsa04550;hsa04650;hsa04660;hsa04662;hsa04664;hsa04720;hsa04722;hsa04725;hsa04726;hsa04730;hsa04810;hsa04910;hsa04912;hsa04915;hsa04916;hsa04917;hsa04919;hsa04921;hsa04933;hsa05034;hsa05160;hsa05161;hsa05166;hsa05200;hsa05203;hsa05205;hsa05206;hsa05211;hsa05213;hsa05214;hsa05215;hsa05216;hsa05218;hsa05219;hsa05220;hsa05221;hsa05223;hsa05230;hsa05231 
                                               NRAS 
                                               NA 
                                               slateblue1 
                                               16 
                                               11.3305613305613 
                                             
                                             
                                               IQGAP1-mRNA 
                                               -1.08 
                                               0.114 
                                               -1.31 
                                               -0.858 
                                               0.472 
                                               0.404 
                                               0.552 
                                               2.04e-11 
                                               4.53e-10 
                                               lm.nb 
                                               ARMS-mediated activation, DAP12 signaling, Downstream signal transduction, FCERI mediated MAPK activation, Frs2-mediated activation, Gastrin-CREB signalling pathway via PKC and MAPK, Interleukin receptor SHC signaling, IRS-related events triggered by IGF1R, MAP2K and MAPK activation, Nephrin interactions, RAFMAP kinase cascade, RHO GTPases activate IQGAPs, Signaling by ERBB2, Signaling by PDGF, Signaling by SCF-KIT, VEGFA-VEGFR2 Pathway, VEGFR2 mediated cell proliferation 
                                               SEBELIUS_2_C4636 
                                               IQGAP1 
                                               mRNA 
                                               0 
                                               
                                               
                                               MAP2K and MAPK activation;Nephrin interactions;RHO GTPases activate IQGAPs;RAFMAP kinase cascade;Downstream signal transduction;VEGFR2 mediated cell proliferation;Signaling by SCF-KIT;Frs2-mediated activation;ARMS-mediated activation;Gastrin-CREB signalling pathway via PKC and MAPK;Interleukin receptor SHC signaling;FCERI mediated MAPK activation;DAP12 signaling;Signaling by PDGF;VEGFA-VEGFR2 Pathway;Signaling by ERBB2;IRS-related events triggered by IGF1R 
                                               hsa04520;hsa04810;hsa05205 
                                               IQGAP1 
                                               NA 
                                               slateblue1 
                                               16 
                                               -9.47368421052632 
                                             
                                             
                                               PPP2R1B-mRNA 
                                               1.04 
                                               0.114 
                                               0.818 
                                               1.26 
                                               2.06 
                                               1.76 
                                               2.4 
                                               4.71e-11 
                                               9.7e-10 
                                               lm.nb 
                                               ARMS-mediated activation, AXIN missense mutants destabilize the destruction complex, Beta-catenin phosphorylation cascade, CTLA4 inhibitory signaling, Cyclin AB1 associated events during G2M transition, Cyclin D associated events in G1, DAP12 signaling, DARPP-32 events, Disassembly of the destruction complex and recruitment of AXIN to the membrane, Downstream signal transduction, ERKs are inactivated, FCERI mediated MAPK activation, Frs2-mediated activation, Gastrin-CREB signalling pathway via PKC and MAPK, Interleukin receptor SHC signaling, IRS-related events triggered by IGF1R, MASTL Facilitates Mitotic Progression, Misspliced GSK3beta mutants stabilize beta-catenin, Negative regulation of MAPK pathway, RAF activation, RAFMAP kinase cascade, Resolution of Sister Chromatid Cohesion, RHO GTPases Activate Formins, S33 mutants of beta-catenin aren.SQT.t phosphorylated, S37 mutants of beta-catenin aren.SQT.t phosphorylated, S45 mutants of beta-catenin aren.SQT.t phosphorylated, Signaling by ERBB2, Signaling by PDGF, Signaling by SCF-KIT, T41 mutants of beta-catenin aren.SQT.t phosphorylated, Truncations of AMER1 destabilize the destruction complex, VEGFA-VEGFR2 Pathway, VEGFR2 mediated cell proliferation 
                                               SEBELIUS_2_C4636 
                                               PPP2R1B 
                                               mRNA 
                                               0 
                                               
                                               
                                               MASTL Facilitates Mitotic Progression;Misspliced GSK3beta mutants stabilize beta-catenin;T41 mutants of beta-catenin aren.SQT.t phosphorylated;ERKs are inactivated;Glycolysis;Negative regulation of MAPK pathway;Disassembly of the destruction complex and recruitment of AXIN to the membrane;S37 mutants of beta-catenin aren.SQT.t phosphorylated;Cyclin AB1 associated events during G2M transition;RAF activation;RHO GTPases Activate Formins;S33 mutants of beta-catenin aren.SQT.t phosphorylated;DARPP-32 events;Beta-catenin phosphorylation cascade;Truncations of AMER1 destabilize the destruction complex;CTLA4 inhibitory signaling;AXIN missense mutants destabilize the destruction complex;S45 mutants of beta-catenin aren.SQT.t phosphorylated;Cyclin D associated events in G1;Platelet sensitization by LDL;Resolution of Sister Chromatid Cohesion;Inhibition of replication initiation of damaged DNA by RB1E2F1;RAFMAP kinase cascade;Downstream signal transduction;VEGFR2 mediated cell proliferation;Signaling by SCF-KIT;Frs2-mediated activation;ARMS-mediated activation;Gastrin-CREB signalling pathway via PKC and MAPK;Interleukin receptor SHC signaling;FCERI mediated MAPK activation;DAP12 signaling;Signaling by PDGF;VEGFA-VEGFR2 Pathway;Signaling by ERBB2;IRS-related events triggered by IGF1R 
                                               hsa03015;hsa04071;hsa04114;hsa04151;hsa04152;hsa04261;hsa04350;hsa04390;hsa04530;hsa04728;hsa04730;hsa05142;hsa05160 
                                               PPP2R1B 
                                               NA 
                                               slateblue1 
                                               16 
                                               9.12280701754386 
                                             
                                             
                                               MAPK1-mRNA 
                                               0.704 
                                               0.0782 
                                               0.55 
                                               0.857 
                                               1.63 
                                               1.46 
                                               1.81 
                                               7.61e-11 
                                               1.52e-09 
                                               loglinear 
                                               ARMS-mediated activation, CREB phosphorylation through the activation of Ras, DAP12 signaling, Downstream signal transduction, FCERI mediated MAPK activation, Frs2-mediated activation, Gastrin-CREB signalling pathway via PKC and MAPK, Interleukin receptor SHC signaling, IRS-related events triggered by IGF1R, MAP2K and MAPK activation, Negative feedback regulation of MAPK pathway, Negative regulation of FGFR1 signaling, Negative regulation of FGFR2 signaling, Negative regulation of FGFR3 signaling, Negative regulation of FGFR4 signaling, Negative regulation of MAPK pathway, RAFMAP kinase cascade, Regulation of HSF1-mediated heat shock response, RHO GTPases Activate WASPs and WAVEs, Signal transduction by L1, Signaling by ERBB2, Signaling by PDGF, Signaling by SCF-KIT, VEGFA-VEGFR2 Pathway, VEGFR2 mediated cell proliferation 
                                               SEBELIUS_2_C4636 
                                               MAPK1 
                                               mRNA 
                                               0 
                                               
                                               
                                               Negative regulation of FGFR1 signaling;Signal transduction by L1;Negative regulation of FGFR2 signaling;MAP2K and MAPK activation;RHO GTPases Activate WASPs and WAVEs;Senescence-Associated Secretory Phenotype (SASP);Regulation of HSF1-mediated heat shock response;Thrombin signalling through proteinase activated receptors (PARs);Negative regulation of FGFR4 signaling;Oxidative Stress Induced Senescence;Signal attenuation;Negative regulation of FGFR3 signaling;Negative feedback regulation of MAPK pathway;Growth hormone receptor signaling;PLC beta mediated events;FCERI mediated MAPK activation;CREB phosphorylation through the activation of Ras;Negative regulation of MAPK pathway;RAFMAP kinase cascade;Downstream signal transduction;VEGFR2 mediated cell proliferation;Signaling by SCF-KIT;Frs2-mediated activation;ARMS-mediated activation;Gastrin-CREB signalling pathway via PKC and MAPK;Interleukin receptor SHC signaling;DAP12 signaling;Signaling by PDGF;VEGFA-VEGFR2 Pathway;Signaling by ERBB2;IRS-related events triggered by IGF1R 
                                               hsa04010;hsa04012;hsa04014;hsa04015;hsa04022;hsa04024;hsa04062;hsa04066;hsa04068;hsa04071;hsa04072;hsa04114;hsa04150;hsa04151;hsa04261;hsa04270;hsa04320;hsa04350;hsa04360;hsa04370;hsa04380;hsa04510;hsa04520;hsa04540;hsa04550;hsa04611;hsa04620;hsa04621;hsa04650;hsa04660;hsa04662;hsa04664;hsa04666;hsa04668;hsa04713;hsa04720;hsa04722;hsa04723;hsa04724;hsa04725;hsa04726;hsa04730;hsa04810;hsa04910;hsa04912;hsa04914;hsa04915;hsa04916;hsa04917;hsa04919;hsa04921;hsa04930;hsa04933;hsa04960;hsa05010;hsa05020;hsa05034;hsa05131;hsa05132;hsa05133;hsa05140;hsa05142;hsa05145;hsa05152;hsa05160;hsa05161;hsa05164;hsa05200;hsa05203;hsa05205;hsa05206;hsa05210;hsa05211;hsa05212;hsa05213;hsa05214;hsa05215;hsa05216;hsa05218;hsa05219;hsa05220;hsa05221;hsa05223;hsa05230;hsa05231 
                                               MAPK1 
                                               NA 
                                               slateblue1 
                                               16 
                                               9.00255754475703 
                                             
                                             
                                               SOS1-mRNA 
                                               1.52 
                                               0.184 
                                               1.16 
                                               1.88 
                                               2.87 
                                               2.23 
                                               3.68 
                                               6.41e-10 
                                               1.06e-08 
                                               lm.nb 
                                               ARMS-mediated activation, Cell surface interactions at the vascular wall, Constitutive Signaling by EGFRvIII, DAP12 signaling, Downstream signal transduction, FCERI mediated MAPK activation, Frs2-mediated activation, Gastrin-CREB signalling pathway via PKC and MAPK, Integrin alphaIIb beta3 signaling, Interleukin receptor SHC signaling, IRS-related events triggered by IGF1R, NRAGE signals death through JNK, RAFMAP kinase cascade, Role of LAT2NTALLAB on calcium mobilization, SHC-mediated cascade.CL.FGFR1, SHC-related events triggered by IGF1R, Signaling by ERBB2, Signaling by FGFR1 in disease, Signaling by FGFR2 in disease, Signaling by FGFR3 in disease, Signaling by FGFR4 in disease, Signaling by PDGF, Signaling by SCF-KIT, Tie2 Signaling, VEGFA-VEGFR2 Pathway, VEGFR2 mediated cell proliferation 
                                               SEBELIUS_2_C4636 
                                               SOS1 
                                               mRNA 
                                               0 
                                               
                                               
                                               Signaling by FGFR2 in disease;Antigen activates B Cell Receptor (BCR) leading to generation of second messengers;SHC-mediated cascade.CL.FGFR3;Signaling by FGFR3 in disease;RAFMAP kinase cascade;SHC-related events triggered by IGF1R;Signaling by FGFR4 in disease;SHC-mediated cascade.CL.FGFR1;GRB2.CL.SOS provides linkage to MAPK signaling for Integrins ;Tie2 Signaling;Role of LAT2NTALLAB on calcium mobilization;Signaling by FGFR1 in disease;Signal attenuation;NRAGE signals death through JNK;SHC-mediated cascade.CL.FGFR4;FCERI mediated Ca.PLS.2 mobilization;Constitutive Signaling by EGFRvIII;FCERI mediated MAPK activation;Signaling by ERBB2;Signaling by SCF-KIT;DAP12 signaling;Gastrin-CREB signalling pathway via PKC and MAPK;Interleukin receptor SHC signaling;IRS-related events triggered by IGF1R;Downstream signal transduction;VEGFR2 mediated cell proliferation;Frs2-mediated activation;ARMS-mediated activation;Signaling by PDGF;VEGFA-VEGFR2 Pathway;Integrin alphaIIb beta3 signaling;Cell surface interactions at the vascular wall 
                                               hsa04010;hsa04012;hsa04014;hsa04062;hsa04068;hsa04072;hsa04151;hsa04320;hsa04510;hsa04540;hsa04630;hsa04650;hsa04660;hsa04662;hsa04664;hsa04722;hsa04810;hsa04910;hsa04912;hsa04915;hsa04917;hsa05034;hsa05160;hsa05200;hsa05205;hsa05206;hsa05211;hsa05213;hsa05214;hsa05215;hsa05220;hsa05221;hsa05223;hsa05231 
                                               SOS1 
                                               NA 
                                               slateblue1 
                                               16 
                                               8.26086956521739 
                                             
                                             
                                               ERBB2-mRNA 
                                               1.71 
                                               0.273 
                                               1.17 
                                               2.24 
                                               3.27 
                                               2.25 
                                               4.74 
                                               2.86e-07 
                                               3.8e-06 
                                               lm.nb 
                                               ARMS-mediated activation, DAP12 signaling, Downstream signal transduction, FCERI mediated MAPK activation, Frs2-mediated activation, GAB1 signalosome, Gastrin-CREB signalling pathway via PKC and MAPK, Interleukin receptor SHC signaling, IRS-related events triggered by IGF1R, PI3KAKT activation, RAFMAP kinase cascade, Role of LAT2NTALLAB on calcium mobilization, Sema4D induced cell migration and growth-cone collapse, Signaling by ERBB2, Signaling by PDGF, Signaling by SCF-KIT, VEGFA-VEGFR2 Pathway, VEGFR2 mediated cell proliferation 
                                               SEBELIUS_2_C4636 
                                               ERBB2 
                                               mRNA 
                                               0 
                                               
                                               
                                               RAFMAP kinase cascade;GRB7 events in ERBB2 signaling;PLCG1 events in ERBB2 signaling;Sema4D induced cell migration and growth-cone collapse;Signaling by ERBB2;Downstream signal transduction;GAB1 signalosome;Signaling by SCF-KIT;PI3KAKT activation;Role of LAT2NTALLAB on calcium mobilization;DAP12 signaling;Signaling by PDGF;VEGFR2 mediated cell proliferation;Frs2-mediated activation;ARMS-mediated activation;Gastrin-CREB signalling pathway via PKC and MAPK;Interleukin receptor SHC signaling;FCERI mediated MAPK activation;VEGFA-VEGFR2 Pathway;IRS-related events triggered by IGF1R 
                                               hsa04012;hsa04020;hsa04066;hsa04510;hsa04520;hsa05200;hsa05205;hsa05206;hsa05212;hsa05213;hsa05215;hsa05219;hsa05223;hsa05230 
                                               ERBB2 
                                               NA 
                                               slateblue1 
                                               16 
                                               6.26373626373626 
                                             
                                             
                                               ACTB-mRNA 
                                               -1.44 
                                               0.234 
                                               -1.9 
                                               -0.984 
                                               0.368 
                                               0.268 
                                               0.506 
                                               3.69e-07 
                                               4.85e-06 
                                               lm.nb 
                                               ARMS-mediated activation, Cell-extracellular matrix interactions, DAP12 signaling, Downstream signal transduction, EPH-ephrin mediated repulsion of cells, EPHB-mediated forward signaling, FCERI mediated MAPK activation, Frs2-mediated activation, Gastrin-CREB signalling pathway via PKC and MAPK, Interleukin receptor SHC signaling, IRS-related events triggered by IGF1R, RAFMAP kinase cascade, Regulation of actin dynamics for phagocytic cup formation, RHO GTPases Activate Formins, RHO GTPases activate IQGAPs, Signaling by ERBB2, Signaling by PDGF, Signaling by SCF-KIT, VEGFA-VEGFR2 Pathway, VEGFR2 mediated cell proliferation 
                                               SEBELIUS_2_C4636 
                                               ACTB 
                                               mRNA 
                                               0 
                                               
                                               
                                               EPHB-mediated forward signaling;Interaction between L1 and Ankyrins;DNA Damage Recognition in GG-NER;EPH-ephrin mediated repulsion of cells;Prefoldin mediated transfer of substrate  to CCTTriC;RHO GTPases Activate Formins;RHO GTPases activate IQGAPs;Cell-extracellular matrix interactions;Regulation of actin dynamics for phagocytic cup formation;Folding of actin by CCTTriC;B-WICH complex positively regulates rRNA expression;RAFMAP kinase cascade;Downstream signal transduction;VEGFR2 mediated cell proliferation;Signaling by SCF-KIT;Frs2-mediated activation;ARMS-mediated activation;Gastrin-CREB signalling pathway via PKC and MAPK;Interleukin receptor SHC signaling;FCERI mediated MAPK activation;DAP12 signaling;Signaling by PDGF;VEGFA-VEGFR2 Pathway;Signaling by ERBB2;IRS-related events triggered by IGF1R 
                                               hsa04015;hsa04145;hsa04390;hsa04510;hsa04520;hsa04530;hsa04611;hsa04670;hsa04810;hsa04919;hsa04921;hsa04971;hsa05100;hsa05110;hsa05130;hsa05131;hsa05132;hsa05164;hsa05205;hsa05410;hsa05412;hsa05414;hsa05416 
                                               ACTB 
                                               NA 
                                               slateblue1 
                                               16 
                                               -6.15384615384615 
                                             
                                             
                                               PTK2-mRNA 
                                               0.703 
                                               0.119 
                                               0.471 
                                               0.936 
                                               1.63 
                                               1.39 
                                               1.91 
                                               7.91e-07 
                                               1e-05 
                                               lm.nb 
                                               Apoptotic cleavage of cellular proteins, ARMS-mediated activation, DAP12 signaling, DCC mediated attractive signaling, Downstream signal transduction, EPHA-mediated growth cone collapse, EPHB-mediated forward signaling, FCERI mediated MAPK activation, Frs2-mediated activation, Gastrin-CREB signalling pathway via PKC and MAPK, Integrin alphaIIb beta3 signaling, Interleukin receptor SHC signaling, IRS-related events triggered by IGF1R, p130Cas linkage to MAPK signaling for integrins, RAFMAP kinase cascade, Regulation of actin dynamics for phagocytic cup formation, RHO GTPases Activate WASPs and WAVEs, Signaling by ERBB2, Signaling by PDGF, Signaling by SCF-KIT, VEGFA-VEGFR2 Pathway, VEGFR2 mediated cell proliferation 
                                               SEBELIUS_2_C4636 
                                               PTK2 
                                               mRNA 
                                               0 
                                               
                                               
                                               Apoptotic cleavage of cellular proteins;RAFMAP kinase cascade;RHO GTPases Activate WASPs and WAVEs;DCC mediated attractive signaling;p130Cas linkage to MAPK signaling for integrins;EPHB-mediated forward signaling;EPHA-mediated growth cone collapse;Regulation of actin dynamics for phagocytic cup formation;Downstream signal transduction;VEGFR2 mediated cell proliferation;Signaling by SCF-KIT;Frs2-mediated activation;ARMS-mediated activation;Gastrin-CREB signalling pathway via PKC and MAPK;Interleukin receptor SHC signaling;FCERI mediated MAPK activation;DAP12 signaling;Signaling by PDGF;VEGFA-VEGFR2 Pathway;Signaling by ERBB2;IRS-related events triggered by IGF1R;Integrin alphaIIb beta3 signaling 
                                               hsa04012;hsa04062;hsa04151;hsa04360;hsa04370;hsa04510;hsa04670;hsa04810;hsa05100;hsa05146;hsa05200;hsa05202;hsa05205;hsa05222 
                                               PTK2 
                                               NA 
                                               slateblue1 
                                               16 
                                               5.90756302521008 
                                             
                                             
                                               PPP2CB-mRNA 
                                               -0.609 
                                               0.109 
                                               -0.822 
                                               -0.397 
                                               0.655 
                                               0.565 
                                               0.76 
                                               2.12e-06 
                                               2.55e-05 
                                               lm.nb 
                                               ARMS-mediated activation, AXIN missense mutants destabilize the destruction complex, Beta-catenin phosphorylation cascade, CTLA4 inhibitory signaling, Cyclin AB1 associated events during G2M transition, Cyclin D associated events in G1, DAP12 signaling, DARPP-32 events, Disassembly of the destruction complex and recruitment of AXIN to the membrane, Downstream signal transduction, ERKs are inactivated, FCERI mediated MAPK activation, Frs2-mediated activation, Gastrin-CREB signalling pathway via PKC and MAPK, Interleukin receptor SHC signaling, IRS-related events triggered by IGF1R, MASTL Facilitates Mitotic Progression, Misspliced GSK3beta mutants stabilize beta-catenin, Negative regulation of FGFR1 signaling, Negative regulation of FGFR2 signaling, Negative regulation of FGFR3 signaling, Negative regulation of FGFR4 signaling, Negative regulation of MAPK pathway, RAF activation, RAFMAP kinase cascade, Resolution of Sister Chromatid Cohesion, RHO GTPases Activate Formins, S33 mutants of beta-catenin aren.SQT.t phosphorylated, S37 mutants of beta-catenin aren.SQT.t phosphorylated, S45 mutants of beta-catenin aren.SQT.t phosphorylated, Signaling by ERBB2, Signaling by PDGF, Signaling by SCF-KIT, Spry regulation of FGF signaling, T41 mutants of beta-catenin aren.SQT.t phosphorylated, Truncations of AMER1 destabilize the destruction complex, VEGFA-VEGFR2 Pathway, VEGFR2 mediated cell proliferation 
                                               SEBELIUS_2_C4636 
                                               PPP2CB 
                                               mRNA 
                                               0 
                                               
                                               
                                               MASTL Facilitates Mitotic Progression;Misspliced GSK3beta mutants stabilize beta-catenin;T41 mutants of beta-catenin aren.SQT.t phosphorylated;ERKs are inactivated;Glycolysis;Negative regulation of MAPK pathway;Disassembly of the destruction complex and recruitment of AXIN to the membrane;S37 mutants of beta-catenin aren.SQT.t phosphorylated;Cyclin AB1 associated events during G2M transition;RAF activation;RHO GTPases Activate Formins;S33 mutants of beta-catenin aren.SQT.t phosphorylated;DARPP-32 events;Beta-catenin phosphorylation cascade;Spry regulation of FGF signaling;Truncations of AMER1 destabilize the destruction complex;CTLA4 inhibitory signaling;AXIN missense mutants destabilize the destruction complex;S45 mutants of beta-catenin aren.SQT.t phosphorylated;Cyclin D associated events in G1;Platelet sensitization by LDL;Resolution of Sister Chromatid Cohesion;Inhibition of replication initiation of damaged DNA by RB1E2F1;RAFMAP kinase cascade;Downstream signal transduction;VEGFR2 mediated cell proliferation;Signaling by SCF-KIT;Frs2-mediated activation;ARMS-mediated activation;Gastrin-CREB signalling pathway via PKC and MAPK;Interleukin receptor SHC signaling;FCERI mediated MAPK activation;DAP12 signaling;Signaling by PDGF;VEGFA-VEGFR2 Pathway;Signaling by ERBB2;IRS-related events triggered by IGF1R;Negative regulation of FGFR3 signaling;Negative regulation of FGFR4 signaling;Negative regulation of FGFR2 signaling;Negative regulation of FGFR1 signaling 
                                               hsa03015;hsa04071;hsa04114;hsa04151;hsa04152;hsa04261;hsa04350;hsa04390;hsa04530;hsa04728;hsa04730;hsa05142;hsa05160 
                                               PPP2CB 
                                               NA 
                                               slateblue1 
                                               16 
                                               -5.58715596330275 
                                             
                                             
                                               PDGFRB-mRNA 
                                               -3.42 
                                               0.725 
                                               -4.84 
                                               -2 
                                               0.0936 
                                               0.035 
                                               0.251 
                                               3.42e-05 
                                               0.000357 
                                               loglinear 
                                               ARMS-mediated activation, Constitutive Signaling by Aberrant PI3K in Cancer, DAP12 signaling, Downstream signal transduction, FCERI mediated MAPK activation, Frs2-mediated activation, GAB1 signalosome, Gastrin-CREB signalling pathway via PKC and MAPK, Interleukin receptor SHC signaling, IRS-related events triggered by IGF1R, PI3KAKT activation, PIP3 activates AKT signaling, RAFMAP kinase cascade, Role of LAT2NTALLAB on calcium mobilization, Signaling by ERBB2, Signaling by PDGF, Signaling by SCF-KIT, VEGFA-VEGFR2 Pathway, VEGFR2 mediated cell proliferation 
                                               SEBELIUS_2_C4636 
                                               PDGFRB 
                                               mRNA 
                                               0 
                                               
                                               
                                               PIP3 activates AKT signaling;Constitutive Signaling by Aberrant PI3K in Cancer;RAFMAP kinase cascade;Signaling by PDGF;Downstream signal transduction;GAB1 signalosome;Signaling by SCF-KIT;PI3KAKT activation;Role of LAT2NTALLAB on calcium mobilization;DAP12 signaling;Signaling by ERBB2;VEGFR2 mediated cell proliferation;Frs2-mediated activation;ARMS-mediated activation;Gastrin-CREB signalling pathway via PKC and MAPK;Interleukin receptor SHC signaling;FCERI mediated MAPK activation;VEGFA-VEGFR2 Pathway;IRS-related events triggered by IGF1R 
                                               hsa04010;hsa04014;hsa04015;hsa04020;hsa04060;hsa04072;hsa04151;hsa04510;hsa04540;hsa04810;hsa05166;hsa05200;hsa05206;hsa05214;hsa05215;hsa05218;hsa05230;hsa05231 
                                               PDGFRB 
                                               NA 
                                               slateblue1 
                                               16 
                                               -4.71724137931034 
                                             
                                             
                                               BRAF-mRNA 
                                               0.865 
                                               0.186 
                                               0.499 
                                               1.23 
                                               1.82 
                                               1.41 
                                               2.35 
                                               4.27e-05 
                                               0.000439 
                                               lm.nb 
                                               ARMS-mediated activation, CREB phosphorylation through the activation of Ras, DAP12 signaling, Downstream signal transduction, FCERI mediated MAPK activation, Frs2-mediated activation, Gastrin-CREB signalling pathway via PKC and MAPK, Interleukin receptor SHC signaling, IRS-related events triggered by IGF1R, MAP2K and MAPK activation, Negative feedback regulation of MAPK pathway, Negative regulation of FGFR1 signaling, Negative regulation of FGFR2 signaling, Negative regulation of FGFR3 signaling, Negative regulation of FGFR4 signaling, Negative regulation of MAPK pathway, RAF activation, RAFMAP kinase cascade, Signaling by ERBB2, Signaling by PDGF, Signaling by SCF-KIT, Spry regulation of FGF signaling, VEGFA-VEGFR2 Pathway, VEGFR2 mediated cell proliferation 
                                               SEBELIUS_2_C4636 
                                               BRAF 
                                               mRNA 
                                               0 
                                               
                                               
                                               Spry regulation of FGF signaling;MAP2K and MAPK activation;RAF activation;CREB phosphorylation through the activation of Ras;Negative feedback regulation of MAPK pathway;ARMS-mediated activation;Negative regulation of FGFR3 signaling;Negative regulation of FGFR4 signaling;Negative regulation of FGFR2 signaling;Negative regulation of FGFR1 signaling;RAFMAP kinase cascade;Downstream signal transduction;VEGFR2 mediated cell proliferation;Signaling by SCF-KIT;Frs2-mediated activation;Gastrin-CREB signalling pathway via PKC and MAPK;Interleukin receptor SHC signaling;FCERI mediated MAPK activation;DAP12 signaling;Signaling by PDGF;VEGFA-VEGFR2 Pathway;Signaling by ERBB2;IRS-related events triggered by IGF1R;Negative regulation of MAPK pathway 
                                               hsa04010;hsa04012;hsa04015;hsa04024;hsa04062;hsa04068;hsa04150;hsa04270;hsa04510;hsa04650;hsa04720;hsa04722;hsa04726;hsa04730;hsa04810;hsa04910;hsa04914;hsa05034;hsa05160;hsa05200;hsa05205;hsa05210;hsa05211;hsa05212;hsa05213;hsa05214;hsa05215;hsa05216;hsa05218;hsa05219;hsa05220;hsa05221;hsa05223 
                                               BRAF 
                                               NA 
                                               slateblue1 
                                               16 
                                               4.6505376344086 
                                             
                                             
                                               EGFR-mRNA 
                                               -1.26 
                                               0.28 
                                               -1.81 
                                               -0.711 
                                               0.418 
                                               0.286 
                                               0.611 
                                               6.51e-05 
                                               0.000656 
                                               lm.nb 
                                               ARMS-mediated activation, Constitutive Signaling by Aberrant PI3K in Cancer, Constitutive Signaling by EGFRvIII, DAP12 signaling, Downstream signal transduction, FCERI mediated MAPK activation, Frs2-mediated activation, GAB1 signalosome, Gastrin-CREB signalling pathway via PKC and MAPK, Interleukin receptor SHC signaling, IRS-related events triggered by IGF1R, PI3KAKT activation, Role of LAT2NTALLAB on calcium mobilization, Signaling by ERBB2, Signaling by PDGF, Signaling by SCF-KIT, VEGFA-VEGFR2 Pathway, VEGFR2 mediated cell proliferation 
                                               SEBELIUS_2_C4636 
                                               EGFR 
                                               mRNA 
                                               0 
                                               
                                               
                                               Constitutive Signaling by Aberrant PI3K in Cancer;Inhibition of Signaling by Overexpressed EGFR;PLCG1 events in ERBB2 signaling;EGFR interacts with phospholipase C-gamma;EGFR Transactivation by Gastrin;Constitutive Signaling by Ligand-Responsive EGFR Cancer Variants;Constitutive Signaling by EGFRvIII;GAB1 signalosome;Downstream signal transduction;Signaling by SCF-KIT;PI3KAKT activation;Role of LAT2NTALLAB on calcium mobilization;DAP12 signaling;Signaling by PDGF;Signaling by ERBB2;VEGFR2 mediated cell proliferation;Frs2-mediated activation;ARMS-mediated activation;Gastrin-CREB signalling pathway via PKC and MAPK;Interleukin receptor SHC signaling;FCERI mediated MAPK activation;VEGFA-VEGFR2 Pathway;IRS-related events triggered by IGF1R 
                                               hsa04010;hsa04012;hsa04014;hsa04015;hsa04020;hsa04060;hsa04066;hsa04068;hsa04072;hsa04144;hsa04151;hsa04320;hsa04510;hsa04520;hsa04540;hsa04810;hsa04912;hsa04915;hsa04921;hsa05120;hsa05160;hsa05200;hsa05205;hsa05206;hsa05212;hsa05213;hsa05214;hsa05215;hsa05218;hsa05219;hsa05223;hsa05230;hsa05231 
                                               EGFR 
                                               NA 
                                               slateblue1 
                                               16 
                                               -4.5 
                                             
                                             
                                               FGFR1-mRNA 
                                               -1.04 
                                               0.267 
                                               -1.57 
                                               -0.519 
                                               0.486 
                                               0.338 
                                               0.698 
                                               0.000383 
                                               0.00367 
                                               lm.nb 
                                               ARMS-mediated activation, Constitutive Signaling by Aberrant PI3K in Cancer, DAP12 signaling, Downstream signal transduction, FCERI mediated MAPK activation, Frs2-mediated activation, GAB1 signalosome, Gastrin-CREB signalling pathway via PKC and MAPK, Interleukin receptor SHC signaling, IRS-related events triggered by IGF1R, PI3K Cascade, PI3KAKT activation, RAFMAP kinase cascade, Role of LAT2NTALLAB on calcium mobilization, SHC-mediated cascade.CL.FGFR1, Signaling by ERBB2, Signaling by FGFR1 in disease, Signaling by PDGF, Signaling by SCF-KIT, VEGFA-VEGFR2 Pathway, VEGFR2 mediated cell proliferation 
                                               SEBELIUS_2_C4636 
                                               FGFR1 
                                               mRNA 
                                               0 
                                               
                                               
                                               RAFMAP kinase cascade;Constitutive Signaling by Aberrant PI3K in Cancer;SHC-mediated cascade.CL.FGFR1;Phospholipase C-mediated cascade.CL. FGFR1;Signaling by FGFR1 amplification mutants;PI3K Cascade;FGFR1b ligand binding and activation;Signaling by FGFR1 in disease;Downstream signal transduction;GAB1 signalosome;Signaling by SCF-KIT;PI3KAKT activation;Role of LAT2NTALLAB on calcium mobilization;DAP12 signaling;Signaling by PDGF;Signaling by ERBB2;VEGFR2 mediated cell proliferation;Frs2-mediated activation;ARMS-mediated activation;Gastrin-CREB signalling pathway via PKC and MAPK;Interleukin receptor SHC signaling;FCERI mediated MAPK activation;VEGFA-VEGFR2 Pathway;IRS-related events triggered by IGF1R 
                                               hsa04010;hsa04014;hsa04015;hsa04151;hsa04520;hsa04550;hsa04810;hsa05200;hsa05205;hsa05215;hsa05218;hsa05230 
                                               FGFR1 
                                               NA 
                                               slateblue1 
                                               16 
                                               -3.89513108614232 
                                             
                                             
                                               IRS1-mRNA 
                                               -0.896 
                                               0.266 
                                               -1.42 
                                               -0.374 
                                               0.538 
                                               0.374 
                                               0.772 
                                               0.0018 
                                               0.0163 
                                               lm.nb 
                                               ARMS-mediated activation, Constitutive Signaling by Aberrant PI3K in Cancer, DAP12 signaling, Downstream signal transduction, FCERI mediated MAPK activation, Frs2-mediated activation, GAB1 signalosome, Gastrin-CREB signalling pathway via PKC and MAPK, Interleukin receptor SHC signaling, IRS-related events triggered by IGF1R, PI3K Cascade, PI3KAKT activation, RAFMAP kinase cascade, Role of LAT2NTALLAB on calcium mobilization, Signaling by ERBB2, Signaling by PDGF, Signaling by SCF-KIT, VEGFA-VEGFR2 Pathway, VEGFR2 mediated cell proliferation 
                                               SEBELIUS_2_C4636 
                                               IRS1 
                                               mRNA 
                                               0 
                                               
                                               
                                               Constitutive Signaling by Aberrant PI3K in Cancer;RAFMAP kinase cascade;PI3K Cascade;Growth hormone receptor signaling;IRS-related events triggered by IGF1R;PI3KAKT activation;Downstream signal transduction;GAB1 signalosome;Signaling by SCF-KIT;Role of LAT2NTALLAB on calcium mobilization;DAP12 signaling;Signaling by PDGF;Signaling by ERBB2;VEGFR2 mediated cell proliferation;Frs2-mediated activation;ARMS-mediated activation;Gastrin-CREB signalling pathway via PKC and MAPK;Interleukin receptor SHC signaling;FCERI mediated MAPK activation;VEGFA-VEGFR2 Pathway 
                                               hsa04022;hsa04068;hsa04150;hsa04151;hsa04152;hsa04211;hsa04213;hsa04722;hsa04910;hsa04920;hsa04923;hsa04930;hsa04931;hsa04932;hsa04960;hsa05206 
                                               IRS1 
                                               NA 
                                               slateblue2 
                                               16 
                                               -3.36842105263158 
                                             
                                             
                                               KRAS-mRNA 
                                               -0.345 
                                               0.104 
                                               -0.549 
                                               -0.14 
                                               0.788 
                                               0.684 
                                               0.907 
                                               0.00211 
                                               0.0189 
                                               lm.nb 
                                               ARMS-mediated activation, CD209 (DC-SIGN) signaling, Cell surface interactions at the vascular wall, Constitutive Signaling by EGFRvIII, DAP12 signaling, Downstream signal transduction, FCERI mediated MAPK activation, Frs2-mediated activation, Gastrin-CREB signalling pathway via PKC and MAPK, Interleukin receptor SHC signaling, IRS-related events triggered by IGF1R, MAP2K and MAPK activation, RAFMAP kinase cascade, SHC-mediated cascade.CL.FGFR1, SHC-related events triggered by IGF1R, Signaling by ERBB2, Signaling by FGFR1 in disease, Signaling by FGFR2 in disease, Signaling by FGFR3 in disease, Signaling by FGFR4 in disease, Signaling by PDGF, Signaling by SCF-KIT, Tie2 Signaling, VEGFA-VEGFR2 Pathway, VEGFR2 mediated cell proliferation 
                                               SEBELIUS_2_C4636 
                                               KRAS 
                                               mRNA 
                                               0 
                                               
                                               
                                               Signaling by FGFR2 in disease;Regulation of RAS by GAPs;SHC-mediated cascade.CL.FGFR3;Signaling by FGFR3 in disease;MAP2K and MAPK activation;SHC-related events triggered by IGF1R;Signaling by FGFR4 in disease;SHC-mediated cascade.CL.FGFR1;Tie2 Signaling;Signaling by FGFR1 in disease;Activation of RAS in B cells;SHC-mediated cascade.CL.FGFR4;Constitutive Signaling by EGFRvIII;CD209 (DC-SIGN) signaling;FCERI mediated MAPK activation;RAFMAP kinase cascade;Downstream signal transduction;VEGFR2 mediated cell proliferation;Signaling by SCF-KIT;Frs2-mediated activation;ARMS-mediated activation;Gastrin-CREB signalling pathway via PKC and MAPK;Interleukin receptor SHC signaling;DAP12 signaling;Signaling by PDGF;VEGFA-VEGFR2 Pathway;Signaling by ERBB2;IRS-related events triggered by IGF1R;Cell surface interactions at the vascular wall 
                                               hsa04010;hsa04012;hsa04014;hsa04015;hsa04062;hsa04068;hsa04071;hsa04072;hsa04151;hsa04211;hsa04213;hsa04320;hsa04360;hsa04370;hsa04530;hsa04540;hsa04550;hsa04650;hsa04660;hsa04662;hsa04664;hsa04720;hsa04722;hsa04725;hsa04726;hsa04730;hsa04810;hsa04910;hsa04912;hsa04914;hsa04915;hsa04916;hsa04917;hsa04919;hsa04921;hsa04933;hsa04960;hsa05034;hsa05160;hsa05161;hsa05166;hsa05200;hsa05203;hsa05205;hsa05206;hsa05210;hsa05211;hsa05212;hsa05213;hsa05214;hsa05215;hsa05216;hsa05218;hsa05219;hsa05220;hsa05221;hsa05223;hsa05230;hsa05231 
                                               KRAS 
                                               NA 
                                               slateblue2 
                                               16 
                                               -3.31730769230769 
                                             
                                             
                                               FN1-mRNA 
                                               1.2 
                                               0.364 
                                               0.489 
                                               1.92 
                                               2.3 
                                               1.4 
                                               3.78 
                                               0.00214 
                                               0.0191 
                                               lm.nb 
                                               ARMS-mediated activation, DAP12 signaling, Downstream signal transduction, ECM proteoglycans, Elastic fibre formation, FCERI mediated MAPK activation, Frs2-mediated activation, Gastrin-CREB signalling pathway via PKC and MAPK, Integrin alphaIIb beta3 signaling, Integrin cell surface interactions, Interleukin receptor SHC signaling, IRS-related events triggered by IGF1R, MAP2K and MAPK activation, Molecules associated with elastic fibres, Non-integrin membrane-ECM interactions, p130Cas linkage to MAPK signaling for integrins, Platelet degranulation , RAFMAP kinase cascade, Signaling by ERBB2, Signaling by PDGF, Signaling by SCF-KIT, Syndecan interactions, VEGFA-VEGFR2 Pathway, VEGFR2 mediated cell proliferation 
                                               SEBELIUS_2_C4636 
                                               FN1 
                                               mRNA 
                                               0 
                                               
                                               
                                               MAP2K and MAPK activation;p130Cas linkage to MAPK signaling for integrins;Platelet degranulation ;Molecules associated with elastic fibres;Syndecan interactions;ECM proteoglycans;Fibronectin matrix formation;Integrin cell surface interactions;RAFMAP kinase cascade;Downstream signal transduction;VEGFR2 mediated cell proliferation;Signaling by SCF-KIT;Frs2-mediated activation;ARMS-mediated activation;Gastrin-CREB signalling pathway via PKC and MAPK;Interleukin receptor SHC signaling;FCERI mediated MAPK activation;DAP12 signaling;Signaling by PDGF;VEGFA-VEGFR2 Pathway;Signaling by ERBB2;IRS-related events triggered by IGF1R;Integrin alphaIIb beta3 signaling;Non-integrin membrane-ECM interactions;Elastic fibre formation 
                                               hsa04151;hsa04510;hsa04512;hsa04810;hsa04933;hsa05100;hsa05146;hsa05200;hsa05205;hsa05222 
                                               FN1 
                                               NA 
                                               slateblue2 
                                               16 
                                               3.2967032967033 
                                             
                                             
                                               PPP2CA-mRNA 
                                               0.388 
                                               0.122 
                                               0.15 
                                               0.627 
                                               1.31 
                                               1.11 
                                               1.54 
                                               0.00287 
                                               0.0255 
                                               lm.nb 
                                               ARMS-mediated activation, AXIN missense mutants destabilize the destruction complex, Beta-catenin phosphorylation cascade, CTLA4 inhibitory signaling, Cyclin AB1 associated events during G2M transition, Cyclin D associated events in G1, DAP12 signaling, DARPP-32 events, Disassembly of the destruction complex and recruitment of AXIN to the membrane, Downstream signal transduction, ERKs are inactivated, FCERI mediated MAPK activation, Frs2-mediated activation, Gastrin-CREB signalling pathway via PKC and MAPK, Interleukin receptor SHC signaling, IRS-related events triggered by IGF1R, MASTL Facilitates Mitotic Progression, Negative regulation of FGFR1 signaling, Negative regulation of FGFR2 signaling, Negative regulation of FGFR3 signaling, Negative regulation of FGFR4 signaling, Negative regulation of MAPK pathway, Nonsense Mediated Decay (NMD) enhanced by the Exon Junction Complex (EJC), RAF activation, RAFMAP kinase cascade, Resolution of Sister Chromatid Cohesion, RHO GTPases Activate Formins, S33 mutants of beta-catenin aren.SQT.t phosphorylated, S37 mutants of beta-catenin aren.SQT.t phosphorylated, S45 mutants of beta-catenin aren.SQT.t phosphorylated, Signaling by ERBB2, Signaling by PDGF, Signaling by SCF-KIT, Spry regulation of FGF signaling, T41 mutants of beta-catenin aren.SQT.t phosphorylated, Truncations of AMER1 destabilize the destruction complex, VEGFA-VEGFR2 Pathway, VEGFR2 mediated cell proliferation 
                                               SEBELIUS_2_C4636 
                                               PPP2CA 
                                               mRNA 
                                               0 
                                               
                                               
                                               MASTL Facilitates Mitotic Progression;T41 mutants of beta-catenin aren.SQT.t phosphorylated;ERKs are inactivated;Glycolysis;Negative regulation of MAPK pathway;Disassembly of the destruction complex and recruitment of AXIN to the membrane;S37 mutants of beta-catenin aren.SQT.t phosphorylated;Cyclin AB1 associated events during G2M transition;RAF activation;RHO GTPases Activate Formins;S33 mutants of beta-catenin aren.SQT.t phosphorylated;DARPP-32 events;Beta-catenin phosphorylation cascade;Initiation of Nuclear Envelope Reformation;Spry regulation of FGF signaling;Truncations of AMER1 destabilize the destruction complex;CTLA4 inhibitory signaling;AXIN missense mutants destabilize the destruction complex;S45 mutants of beta-catenin aren.SQT.t phosphorylated;Nonsense Mediated Decay (NMD) enhanced by the Exon Junction Complex (EJC);Cyclin D associated events in G1;Platelet sensitization by LDL;Resolution of Sister Chromatid Cohesion;Inhibition of replication initiation of damaged DNA by RB1E2F1;RAFMAP kinase cascade;Downstream signal transduction;VEGFR2 mediated cell proliferation;Signaling by SCF-KIT;Frs2-mediated activation;ARMS-mediated activation;Gastrin-CREB signalling pathway via PKC and MAPK;Interleukin receptor SHC signaling;FCERI mediated MAPK activation;DAP12 signaling;Signaling by PDGF;VEGFA-VEGFR2 Pathway;Signaling by ERBB2;IRS-related events triggered by IGF1R;Negative regulation of FGFR3 signaling;Negative regulation of FGFR4 signaling;Negative regulation of FGFR2 signaling;Negative regulation of FGFR1 signaling 
                                               hsa03015;hsa04071;hsa04114;hsa04151;hsa04152;hsa04261;hsa04350;hsa04390;hsa04530;hsa04728;hsa04730;hsa05142;hsa05160 
                                               PPP2CA 
                                               NA 
                                               slateblue2 
                                               16 
                                               3.18032786885246 
                                             
                                           
                                           DE Results - ARMS-mediated activation genes - Line: HEPG2 vs.Fibro  Table displaying each sample's global significance scores and directed global significance scores as defined in the heatmaps above. The global significance score is calculated as the square root of the mean squared t-statistic for the genes in a gene set, with t-statistics coming from the linear regression underlying our differential expression analysis. The directed global significance score is calculated as the square root of the mean signed squared t-statistic for the genes in a gene set, with t-statistics coming from the linear regression underlying our differential expression analysis.   
                                      
                                       
                                  
                                
                                
                                  
                                       
                                      
                                           Volcano Plot: Line: HT29 vs.Fibro   More Plot Information                Volcano Plot: Line: HT29 vs.Fibro  Volcano plot displaying each gene's -log10(p-value) and log2 fold change for the selected covariate. Highly statistically significant genes fall at the top of the plot, and highly differentially expressed genes fall to either side. Genes within the selected gene set are highlighted in orange. Horizontal lines indicate various False Discovery Rate (FDR) thresholds.   
                                           DE Results - ARMS-mediated activation genes - Line: HT29 vs.Fibro   More Plot Information      Download CSV Data     
                                           
                                             
                                             
                                           
                                           
                                             
                                               Probe Label 
                                               Log2 fold change 
                                               std error (log2) 
                                               Lower confidence limit (log2) 
                                               Upper confidence limit (log2) 
                                               Linear fold change 
                                               Lower confidence limit (linear) 
                                               Upper confidence limit (linear) 
                                               P-value 
                                               BY.p.value 
                                               method 
                                               Gene.sets 
                                               Codeset.Name 
                                               Probe.Label 
                                               Analyte.Type 
                                               Is.Control 
                                               Control.Type 
                                               Related.Probes 
                                               Probe.Annotation 
                                               KEGG.Pathways 
                                               Official.Gene.Name 
                                               Control.Conc 
                                               volcanocol 
                                               volcanopch 
                                               tstats.all 
                                             
                                           
                                           
                                             
                                               PDGFRA-mRNA 
                                               -7.96 
                                               0.273 
                                               -8.5 
                                               -7.42 
                                               0.00402 
                                               0.00277 
                                               0.00582 
                                               4.02e-27 
                                               6.33e-25 
                                               lm.nb 
                                               ARMS-mediated activation, Constitutive Signaling by Aberrant PI3K in Cancer, DAP12 signaling, Downstream signal transduction, FCERI mediated MAPK activation, Frs2-mediated activation, GAB1 signalosome, Gastrin-CREB signalling pathway via PKC and MAPK, Interleukin receptor SHC signaling, IRS-related events triggered by IGF1R, PI3KAKT activation, PIP3 activates AKT signaling, RAFMAP kinase cascade, Role of LAT2NTALLAB on calcium mobilization, Signaling by ERBB2, Signaling by PDGF, Signaling by SCF-KIT, VEGFA-VEGFR2 Pathway, VEGFR2 mediated cell proliferation 
                                               SEBELIUS_2_C4636 
                                               PDGFRA 
                                               mRNA 
                                               0 
                                               
                                               
                                               PIP3 activates AKT signaling;Constitutive Signaling by Aberrant PI3K in Cancer;RAFMAP kinase cascade;Signaling by PDGF;Downstream signal transduction;GAB1 signalosome;Signaling by SCF-KIT;PI3KAKT activation;Role of LAT2NTALLAB on calcium mobilization;DAP12 signaling;Signaling by ERBB2;VEGFR2 mediated cell proliferation;Frs2-mediated activation;ARMS-mediated activation;Gastrin-CREB signalling pathway via PKC and MAPK;Interleukin receptor SHC signaling;FCERI mediated MAPK activation;VEGFA-VEGFR2 Pathway;IRS-related events triggered by IGF1R 
                                               hsa04010;hsa04014;hsa04015;hsa04020;hsa04060;hsa04072;hsa04144;hsa04151;hsa04510;hsa04540;hsa04810;hsa05166;hsa05200;hsa05206;hsa05214;hsa05215;hsa05218;hsa05230;hsa05231 
                                               PDGFRA 
                                               NA 
                                               slateblue1 
                                               16 
                                               -29.1575091575092 
                                             
                                             
                                               FGFR1-mRNA 
                                               -9.61 
                                               0.332 
                                               -10.3 
                                               -8.96 
                                               0.00128 
                                               0.000816 
                                               0.00201 
                                               5.21e-27 
                                               7.39e-25 
                                               lm.nb 
                                               ARMS-mediated activation, Constitutive Signaling by Aberrant PI3K in Cancer, DAP12 signaling, Downstream signal transduction, FCERI mediated MAPK activation, Frs2-mediated activation, GAB1 signalosome, Gastrin-CREB signalling pathway via PKC and MAPK, Interleukin receptor SHC signaling, IRS-related events triggered by IGF1R, PI3K Cascade, PI3KAKT activation, RAFMAP kinase cascade, Role of LAT2NTALLAB on calcium mobilization, SHC-mediated cascade.CL.FGFR1, Signaling by ERBB2, Signaling by FGFR1 in disease, Signaling by PDGF, Signaling by SCF-KIT, VEGFA-VEGFR2 Pathway, VEGFR2 mediated cell proliferation 
                                               SEBELIUS_2_C4636 
                                               FGFR1 
                                               mRNA 
                                               0 
                                               
                                               
                                               RAFMAP kinase cascade;Constitutive Signaling by Aberrant PI3K in Cancer;SHC-mediated cascade.CL.FGFR1;Phospholipase C-mediated cascade.CL. FGFR1;Signaling by FGFR1 amplification mutants;PI3K Cascade;FGFR1b ligand binding and activation;Signaling by FGFR1 in disease;Downstream signal transduction;GAB1 signalosome;Signaling by SCF-KIT;PI3KAKT activation;Role of LAT2NTALLAB on calcium mobilization;DAP12 signaling;Signaling by PDGF;Signaling by ERBB2;VEGFR2 mediated cell proliferation;Frs2-mediated activation;ARMS-mediated activation;Gastrin-CREB signalling pathway via PKC and MAPK;Interleukin receptor SHC signaling;FCERI mediated MAPK activation;VEGFA-VEGFR2 Pathway;IRS-related events triggered by IGF1R 
                                               hsa04010;hsa04014;hsa04015;hsa04151;hsa04520;hsa04550;hsa04810;hsa05200;hsa05205;hsa05215;hsa05218;hsa05230 
                                               FGFR1 
                                               NA 
                                               slateblue1 
                                               16 
                                               -28.9457831325301 
                                             
                                             
                                               FN1-mRNA 
                                               -9.78 
                                               0.348 
                                               -10.5 
                                               -9.1 
                                               0.00114 
                                               0.000708 
                                               0.00182 
                                               1.46e-26 
                                               1.89e-24 
                                               lm.nb 
                                               ARMS-mediated activation, DAP12 signaling, Downstream signal transduction, ECM proteoglycans, Elastic fibre formation, FCERI mediated MAPK activation, Frs2-mediated activation, Gastrin-CREB signalling pathway via PKC and MAPK, Integrin alphaIIb beta3 signaling, Integrin cell surface interactions, Interleukin receptor SHC signaling, IRS-related events triggered by IGF1R, MAP2K and MAPK activation, Molecules associated with elastic fibres, Non-integrin membrane-ECM interactions, p130Cas linkage to MAPK signaling for integrins, Platelet degranulation , RAFMAP kinase cascade, Signaling by ERBB2, Signaling by PDGF, Signaling by SCF-KIT, Syndecan interactions, VEGFA-VEGFR2 Pathway, VEGFR2 mediated cell proliferation 
                                               SEBELIUS_2_C4636 
                                               FN1 
                                               mRNA 
                                               0 
                                               
                                               
                                               MAP2K and MAPK activation;p130Cas linkage to MAPK signaling for integrins;Platelet degranulation ;Molecules associated with elastic fibres;Syndecan interactions;ECM proteoglycans;Fibronectin matrix formation;Integrin cell surface interactions;RAFMAP kinase cascade;Downstream signal transduction;VEGFR2 mediated cell proliferation;Signaling by SCF-KIT;Frs2-mediated activation;ARMS-mediated activation;Gastrin-CREB signalling pathway via PKC and MAPK;Interleukin receptor SHC signaling;FCERI mediated MAPK activation;DAP12 signaling;Signaling by PDGF;VEGFA-VEGFR2 Pathway;Signaling by ERBB2;IRS-related events triggered by IGF1R;Integrin alphaIIb beta3 signaling;Non-integrin membrane-ECM interactions;Elastic fibre formation 
                                               hsa04151;hsa04510;hsa04512;hsa04810;hsa04933;hsa05100;hsa05146;hsa05200;hsa05205;hsa05222 
                                               FN1 
                                               NA 
                                               slateblue1 
                                               16 
                                               -28.1034482758621 
                                             
                                             
                                               PTK2-mRNA 
                                               2.36 
                                               0.112 
                                               2.14 
                                               2.58 
                                               5.14 
                                               4.41 
                                               5.98 
                                               2.87e-22 
                                               1.7e-20 
                                               lm.nb 
                                               Apoptotic cleavage of cellular proteins, ARMS-mediated activation, DAP12 signaling, DCC mediated attractive signaling, Downstream signal transduction, EPHA-mediated growth cone collapse, EPHB-mediated forward signaling, FCERI mediated MAPK activation, Frs2-mediated activation, Gastrin-CREB signalling pathway via PKC and MAPK, Integrin alphaIIb beta3 signaling, Interleukin receptor SHC signaling, IRS-related events triggered by IGF1R, p130Cas linkage to MAPK signaling for integrins, RAFMAP kinase cascade, Regulation of actin dynamics for phagocytic cup formation, RHO GTPases Activate WASPs and WAVEs, Signaling by ERBB2, Signaling by PDGF, Signaling by SCF-KIT, VEGFA-VEGFR2 Pathway, VEGFR2 mediated cell proliferation 
                                               SEBELIUS_2_C4636 
                                               PTK2 
                                               mRNA 
                                               0 
                                               
                                               
                                               Apoptotic cleavage of cellular proteins;RAFMAP kinase cascade;RHO GTPases Activate WASPs and WAVEs;DCC mediated attractive signaling;p130Cas linkage to MAPK signaling for integrins;EPHB-mediated forward signaling;EPHA-mediated growth cone collapse;Regulation of actin dynamics for phagocytic cup formation;Downstream signal transduction;VEGFR2 mediated cell proliferation;Signaling by SCF-KIT;Frs2-mediated activation;ARMS-mediated activation;Gastrin-CREB signalling pathway via PKC and MAPK;Interleukin receptor SHC signaling;FCERI mediated MAPK activation;DAP12 signaling;Signaling by PDGF;VEGFA-VEGFR2 Pathway;Signaling by ERBB2;IRS-related events triggered by IGF1R;Integrin alphaIIb beta3 signaling 
                                               hsa04012;hsa04062;hsa04151;hsa04360;hsa04370;hsa04510;hsa04670;hsa04810;hsa05100;hsa05146;hsa05200;hsa05202;hsa05205;hsa05222 
                                               PTK2 
                                               NA 
                                               slateblue1 
                                               16 
                                               21.0714285714286 
                                             
                                             
                                               PPP2CB-mRNA 
                                               -2.03 
                                               0.103 
                                               -2.23 
                                               -1.83 
                                               0.245 
                                               0.213 
                                               0.281 
                                               3.08e-21 
                                               1.56e-19 
                                               lm.nb 
                                               ARMS-mediated activation, AXIN missense mutants destabilize the destruction complex, Beta-catenin phosphorylation cascade, CTLA4 inhibitory signaling, Cyclin AB1 associated events during G2M transition, Cyclin D associated events in G1, DAP12 signaling, DARPP-32 events, Disassembly of the destruction complex and recruitment of AXIN to the membrane, Downstream signal transduction, ERKs are inactivated, FCERI mediated MAPK activation, Frs2-mediated activation, Gastrin-CREB signalling pathway via PKC and MAPK, Interleukin receptor SHC signaling, IRS-related events triggered by IGF1R, MASTL Facilitates Mitotic Progression, Misspliced GSK3beta mutants stabilize beta-catenin, Negative regulation of FGFR1 signaling, Negative regulation of FGFR2 signaling, Negative regulation of FGFR3 signaling, Negative regulation of FGFR4 signaling, Negative regulation of MAPK pathway, RAF activation, RAFMAP kinase cascade, Resolution of Sister Chromatid Cohesion, RHO GTPases Activate Formins, S33 mutants of beta-catenin aren.SQT.t phosphorylated, S37 mutants of beta-catenin aren.SQT.t phosphorylated, S45 mutants of beta-catenin aren.SQT.t phosphorylated, Signaling by ERBB2, Signaling by PDGF, Signaling by SCF-KIT, Spry regulation of FGF signaling, T41 mutants of beta-catenin aren.SQT.t phosphorylated, Truncations of AMER1 destabilize the destruction complex, VEGFA-VEGFR2 Pathway, VEGFR2 mediated cell proliferation 
                                               SEBELIUS_2_C4636 
                                               PPP2CB 
                                               mRNA 
                                               0 
                                               
                                               
                                               MASTL Facilitates Mitotic Progression;Misspliced GSK3beta mutants stabilize beta-catenin;T41 mutants of beta-catenin aren.SQT.t phosphorylated;ERKs are inactivated;Glycolysis;Negative regulation of MAPK pathway;Disassembly of the destruction complex and recruitment of AXIN to the membrane;S37 mutants of beta-catenin aren.SQT.t phosphorylated;Cyclin AB1 associated events during G2M transition;RAF activation;RHO GTPases Activate Formins;S33 mutants of beta-catenin aren.SQT.t phosphorylated;DARPP-32 events;Beta-catenin phosphorylation cascade;Spry regulation of FGF signaling;Truncations of AMER1 destabilize the destruction complex;CTLA4 inhibitory signaling;AXIN missense mutants destabilize the destruction complex;S45 mutants of beta-catenin aren.SQT.t phosphorylated;Cyclin D associated events in G1;Platelet sensitization by LDL;Resolution of Sister Chromatid Cohesion;Inhibition of replication initiation of damaged DNA by RB1E2F1;RAFMAP kinase cascade;Downstream signal transduction;VEGFR2 mediated cell proliferation;Signaling by SCF-KIT;Frs2-mediated activation;ARMS-mediated activation;Gastrin-CREB signalling pathway via PKC and MAPK;Interleukin receptor SHC signaling;FCERI mediated MAPK activation;DAP12 signaling;Signaling by PDGF;VEGFA-VEGFR2 Pathway;Signaling by ERBB2;IRS-related events triggered by IGF1R;Negative regulation of FGFR3 signaling;Negative regulation of FGFR4 signaling;Negative regulation of FGFR2 signaling;Negative regulation of FGFR1 signaling 
                                               hsa03015;hsa04071;hsa04114;hsa04151;hsa04152;hsa04261;hsa04350;hsa04390;hsa04530;hsa04728;hsa04730;hsa05142;hsa05160 
                                               PPP2CB 
                                               NA 
                                               slateblue1 
                                               16 
                                               -19.7087378640777 
                                             
                                             
                                               SRC-mRNA 
                                               2.21 
                                               0.159 
                                               1.9 
                                               2.52 
                                               4.62 
                                               3.73 
                                               5.73 
                                               2.85e-16 
                                               6.85e-15 
                                               lm.nb 
                                               ARMS-mediated activation, CD28 co-stimulation, Cell surface interactions at the vascular wall, CTLA4 inhibitory signaling, DAP12 signaling, Downstream signal transduction, FCERI mediated MAPK activation, Frs2-mediated activation, GAB1 signalosome, Gastrin-CREB signalling pathway via PKC and MAPK, Integrin alphaIIb beta3 signaling, Interleukin receptor SHC signaling, IRS-related events triggered by IGF1R, MAP2K and MAPK activation, Negative regulation of FGFR1 signaling, Negative regulation of FGFR2 signaling, Negative regulation of FGFR3 signaling, Negative regulation of FGFR4 signaling, p130Cas linkage to MAPK signaling for integrins, RAFMAP kinase cascade, Signaling by ERBB2, Signaling by PDGF, Signaling by SCF-KIT, Spry regulation of FGF signaling, VEGFA-VEGFR2 Pathway, VEGFR2 mediated cell proliferation 
                                               SEBELIUS_2_C4636 
                                               SRC 
                                               mRNA 
                                               0 
                                               
                                               
                                               FCGR activation;Spry regulation of FGF signaling;GAB1 signalosome;MAP2K and MAPK activation;CLEC7A (Dectin-1) signaling;CD28 co-stimulation;p130Cas linkage to MAPK signaling for integrins;CTLA4 inhibitory signaling;GRB2.CL.SOS provides linkage to MAPK signaling for Integrins ;Signal regulatory protein (SIRP) family interactions;PECAM1 interactions;Thrombin signalling through proteinase activated receptors (PARs);GP1b-IX-V activation signalling;c-src mediated regulation of Cx43 function and closure of gap junctions;Signaling by ERBB2;VEGFA-VEGFR2 Pathway;Signaling by SCF-KIT;VEGFR2 mediated cell proliferation;RAFMAP kinase cascade;Downstream signal transduction;Frs2-mediated activation;ARMS-mediated activation;Gastrin-CREB signalling pathway via PKC and MAPK;Interleukin receptor SHC signaling;FCERI mediated MAPK activation;DAP12 signaling;Signaling by PDGF;IRS-related events triggered by IGF1R;Negative regulation of FGFR3 signaling;Negative regulation of FGFR4 signaling;Negative regulation of FGFR2 signaling;Negative regulation of FGFR1 signaling;Integrin alphaIIb beta3 signaling;Cell surface interactions at the vascular wall 
                                               hsa04012;hsa04015;hsa04062;hsa04144;hsa04370;hsa04510;hsa04520;hsa04530;hsa04540;hsa04611;hsa04727;hsa04750;hsa04810;hsa04912;hsa04915;hsa04917;hsa04919;hsa04921;hsa05100;hsa05120;hsa05131;hsa05152;hsa05161;hsa05203;hsa05205;hsa05219 
                                               SRC 
                                               NA 
                                               slateblue1 
                                               16 
                                               13.8993710691824 
                                             
                                             
                                               VCL-mRNA 
                                               -3.07 
                                               0.225 
                                               -3.51 
                                               -2.62 
                                               0.119 
                                               0.0879 
                                               0.162 
                                               5.63e-16 
                                               1.33e-14 
                                               lm.nb 
                                               ARMS-mediated activation, DAP12 signaling, Downstream signal transduction, FCERI mediated MAPK activation, Frs2-mediated activation, Gastrin-CREB signalling pathway via PKC and MAPK, Interleukin receptor SHC signaling, IRS-related events triggered by IGF1R, MAP2K and MAPK activation, Platelet degranulation , RAFMAP kinase cascade, Signaling by ERBB2, Signaling by PDGF, Signaling by SCF-KIT, Smooth Muscle Contraction, VEGFA-VEGFR2 Pathway, VEGFR2 mediated cell proliferation 
                                               SEBELIUS_2_C4636 
                                               VCL 
                                               mRNA 
                                               0 
                                               
                                               
                                               Smooth Muscle Contraction;MAP2K and MAPK activation;Platelet degranulation ;RAFMAP kinase cascade;Downstream signal transduction;VEGFR2 mediated cell proliferation;Signaling by SCF-KIT;Frs2-mediated activation;ARMS-mediated activation;Gastrin-CREB signalling pathway via PKC and MAPK;Interleukin receptor SHC signaling;FCERI mediated MAPK activation;DAP12 signaling;Signaling by PDGF;VEGFA-VEGFR2 Pathway;Signaling by ERBB2;IRS-related events triggered by IGF1R 
                                               hsa04510;hsa04520;hsa04670;hsa04810;hsa05100;hsa05131;hsa05146 
                                               VCL 
                                               NA 
                                               slateblue1 
                                               16 
                                               -13.6444444444444 
                                             
                                             
                                               PDGFRB-mRNA 
                                               -8.37 
                                               0.683 
                                               -9.71 
                                               -7.03 
                                               0.00302 
                                               0.00119 
                                               0.00764 
                                               1.37e-14 
                                               2.77e-13 
                                               loglinear 
                                               ARMS-mediated activation, Constitutive Signaling by Aberrant PI3K in Cancer, DAP12 signaling, Downstream signal transduction, FCERI mediated MAPK activation, Frs2-mediated activation, GAB1 signalosome, Gastrin-CREB signalling pathway via PKC and MAPK, Interleukin receptor SHC signaling, IRS-related events triggered by IGF1R, PI3KAKT activation, PIP3 activates AKT signaling, RAFMAP kinase cascade, Role of LAT2NTALLAB on calcium mobilization, Signaling by ERBB2, Signaling by PDGF, Signaling by SCF-KIT, VEGFA-VEGFR2 Pathway, VEGFR2 mediated cell proliferation 
                                               SEBELIUS_2_C4636 
                                               PDGFRB 
                                               mRNA 
                                               0 
                                               
                                               
                                               PIP3 activates AKT signaling;Constitutive Signaling by Aberrant PI3K in Cancer;RAFMAP kinase cascade;Signaling by PDGF;Downstream signal transduction;GAB1 signalosome;Signaling by SCF-KIT;PI3KAKT activation;Role of LAT2NTALLAB on calcium mobilization;DAP12 signaling;Signaling by ERBB2;VEGFR2 mediated cell proliferation;Frs2-mediated activation;ARMS-mediated activation;Gastrin-CREB signalling pathway via PKC and MAPK;Interleukin receptor SHC signaling;FCERI mediated MAPK activation;VEGFA-VEGFR2 Pathway;IRS-related events triggered by IGF1R 
                                               hsa04010;hsa04014;hsa04015;hsa04020;hsa04060;hsa04072;hsa04151;hsa04510;hsa04540;hsa04810;hsa05166;hsa05200;hsa05206;hsa05214;hsa05215;hsa05218;hsa05230;hsa05231 
                                               PDGFRB 
                                               NA 
                                               slateblue1 
                                               16 
                                               -12.2547584187408 
                                             
                                             
                                               IQGAP1-mRNA 
                                               1.3 
                                               0.107 
                                               1.09 
                                               1.51 
                                               2.46 
                                               2.12 
                                               2.84 
                                               1.95e-14 
                                               3.85e-13 
                                               lm.nb 
                                               ARMS-mediated activation, DAP12 signaling, Downstream signal transduction, FCERI mediated MAPK activation, Frs2-mediated activation, Gastrin-CREB signalling pathway via PKC and MAPK, Interleukin receptor SHC signaling, IRS-related events triggered by IGF1R, MAP2K and MAPK activation, Nephrin interactions, RAFMAP kinase cascade, RHO GTPases activate IQGAPs, Signaling by ERBB2, Signaling by PDGF, Signaling by SCF-KIT, VEGFA-VEGFR2 Pathway, VEGFR2 mediated cell proliferation 
                                               SEBELIUS_2_C4636 
                                               IQGAP1 
                                               mRNA 
                                               0 
                                               
                                               
                                               MAP2K and MAPK activation;Nephrin interactions;RHO GTPases activate IQGAPs;RAFMAP kinase cascade;Downstream signal transduction;VEGFR2 mediated cell proliferation;Signaling by SCF-KIT;Frs2-mediated activation;ARMS-mediated activation;Gastrin-CREB signalling pathway via PKC and MAPK;Interleukin receptor SHC signaling;FCERI mediated MAPK activation;DAP12 signaling;Signaling by PDGF;VEGFA-VEGFR2 Pathway;Signaling by ERBB2;IRS-related events triggered by IGF1R 
                                               hsa04520;hsa04810;hsa05205 
                                               IQGAP1 
                                               NA 
                                               slateblue1 
                                               16 
                                               12.1495327102804 
                                             
                                             
                                               KRAS-mRNA 
                                               0.994 
                                               0.0971 
                                               0.803 
                                               1.18 
                                               1.99 
                                               1.75 
                                               2.27 
                                               2.41e-12 
                                               3.76e-11 
                                               lm.nb 
                                               ARMS-mediated activation, CD209 (DC-SIGN) signaling, Cell surface interactions at the vascular wall, Constitutive Signaling by EGFRvIII, DAP12 signaling, Downstream signal transduction, FCERI mediated MAPK activation, Frs2-mediated activation, Gastrin-CREB signalling pathway via PKC and MAPK, Interleukin receptor SHC signaling, IRS-related events triggered by IGF1R, MAP2K and MAPK activation, RAFMAP kinase cascade, SHC-mediated cascade.CL.FGFR1, SHC-related events triggered by IGF1R, Signaling by ERBB2, Signaling by FGFR1 in disease, Signaling by FGFR2 in disease, Signaling by FGFR3 in disease, Signaling by FGFR4 in disease, Signaling by PDGF, Signaling by SCF-KIT, Tie2 Signaling, VEGFA-VEGFR2 Pathway, VEGFR2 mediated cell proliferation 
                                               SEBELIUS_2_C4636 
                                               KRAS 
                                               mRNA 
                                               0 
                                               
                                               
                                               Signaling by FGFR2 in disease;Regulation of RAS by GAPs;SHC-mediated cascade.CL.FGFR3;Signaling by FGFR3 in disease;MAP2K and MAPK activation;SHC-related events triggered by IGF1R;Signaling by FGFR4 in disease;SHC-mediated cascade.CL.FGFR1;Tie2 Signaling;Signaling by FGFR1 in disease;Activation of RAS in B cells;SHC-mediated cascade.CL.FGFR4;Constitutive Signaling by EGFRvIII;CD209 (DC-SIGN) signaling;FCERI mediated MAPK activation;RAFMAP kinase cascade;Downstream signal transduction;VEGFR2 mediated cell proliferation;Signaling by SCF-KIT;Frs2-mediated activation;ARMS-mediated activation;Gastrin-CREB signalling pathway via PKC and MAPK;Interleukin receptor SHC signaling;DAP12 signaling;Signaling by PDGF;VEGFA-VEGFR2 Pathway;Signaling by ERBB2;IRS-related events triggered by IGF1R;Cell surface interactions at the vascular wall 
                                               hsa04010;hsa04012;hsa04014;hsa04015;hsa04062;hsa04068;hsa04071;hsa04072;hsa04151;hsa04211;hsa04213;hsa04320;hsa04360;hsa04370;hsa04530;hsa04540;hsa04550;hsa04650;hsa04660;hsa04662;hsa04664;hsa04720;hsa04722;hsa04725;hsa04726;hsa04730;hsa04810;hsa04910;hsa04912;hsa04914;hsa04915;hsa04916;hsa04917;hsa04919;hsa04921;hsa04933;hsa04960;hsa05034;hsa05160;hsa05161;hsa05166;hsa05200;hsa05203;hsa05205;hsa05206;hsa05210;hsa05211;hsa05212;hsa05213;hsa05214;hsa05215;hsa05216;hsa05218;hsa05219;hsa05220;hsa05221;hsa05223;hsa05230;hsa05231 
                                               KRAS 
                                               NA 
                                               slateblue1 
                                               16 
                                               10.2368692070031 
                                             
                                             
                                               MAP2K1-mRNA 
                                               1.39 
                                               0.139 
                                               1.12 
                                               1.66 
                                               2.62 
                                               2.17 
                                               3.17 
                                               4.65e-12 
                                               6.88e-11 
                                               lm.nb 
                                               ARMS-mediated activation, DAP12 signaling, Downstream signal transduction, FCERI mediated MAPK activation, Frs2-mediated activation, Gastrin-CREB signalling pathway via PKC and MAPK, Interleukin receptor SHC signaling, IRS-related events triggered by IGF1R, Negative feedback regulation of MAPK pathway, Negative regulation of MAPK pathway, RAF activation, RAFMAP kinase cascade, Signaling by ERBB2, Signaling by PDGF, Signaling by SCF-KIT, VEGFA-VEGFR2 Pathway, VEGFR2 mediated cell proliferation 
                                               SEBELIUS_2_C4636 
                                               MAP2K1 
                                               mRNA 
                                               0 
                                               
                                               
                                               RAF activation;Uptake and function of anthrax toxins;Negative feedback regulation of MAPK pathway;Interleukin-1 signaling;RAFMAP kinase cascade;Downstream signal transduction;VEGFR2 mediated cell proliferation;Signaling by SCF-KIT;Frs2-mediated activation;ARMS-mediated activation;Gastrin-CREB signalling pathway via PKC and MAPK;Interleukin receptor SHC signaling;FCERI mediated MAPK activation;DAP12 signaling;Signaling by PDGF;VEGFA-VEGFR2 Pathway;Signaling by ERBB2;IRS-related events triggered by IGF1R;Negative regulation of MAPK pathway 
                                               hsa04010;hsa04012;hsa04014;hsa04015;hsa04022;hsa04024;hsa04062;hsa04066;hsa04068;hsa04071;hsa04072;hsa04114;hsa04151;hsa04270;hsa04320;hsa04370;hsa04380;hsa04510;hsa04540;hsa04550;hsa04620;hsa04650;hsa04660;hsa04662;hsa04664;hsa04666;hsa04668;hsa04720;hsa04722;hsa04725;hsa04726;hsa04730;hsa04810;hsa04910;hsa04912;hsa04914;hsa04915;hsa04916;hsa04917;hsa04919;hsa04921;hsa05020;hsa05034;hsa05161;hsa05164;hsa05200;hsa05205;hsa05206;hsa05210;hsa05211;hsa05212;hsa05213;hsa05214;hsa05215;hsa05216;hsa05218;hsa05219;hsa05220;hsa05221;hsa05223;hsa05230;hsa05231 
                                               MAP2K1 
                                               NA 
                                               slateblue1 
                                               16 
                                               10 
                                             
                                             
                                               PPP2R1B-mRNA 
                                               0.978 
                                               0.107 
                                               0.769 
                                               1.19 
                                               1.97 
                                               1.7 
                                               2.28 
                                               5.09e-11 
                                               7.3e-10 
                                               lm.nb 
                                               ARMS-mediated activation, AXIN missense mutants destabilize the destruction complex, Beta-catenin phosphorylation cascade, CTLA4 inhibitory signaling, Cyclin AB1 associated events during G2M transition, Cyclin D associated events in G1, DAP12 signaling, DARPP-32 events, Disassembly of the destruction complex and recruitment of AXIN to the membrane, Downstream signal transduction, ERKs are inactivated, FCERI mediated MAPK activation, Frs2-mediated activation, Gastrin-CREB signalling pathway via PKC and MAPK, Interleukin receptor SHC signaling, IRS-related events triggered by IGF1R, MASTL Facilitates Mitotic Progression, Misspliced GSK3beta mutants stabilize beta-catenin, Negative regulation of MAPK pathway, RAF activation, RAFMAP kinase cascade, Resolution of Sister Chromatid Cohesion, RHO GTPases Activate Formins, S33 mutants of beta-catenin aren.SQT.t phosphorylated, S37 mutants of beta-catenin aren.SQT.t phosphorylated, S45 mutants of beta-catenin aren.SQT.t phosphorylated, Signaling by ERBB2, Signaling by PDGF, Signaling by SCF-KIT, T41 mutants of beta-catenin aren.SQT.t phosphorylated, Truncations of AMER1 destabilize the destruction complex, VEGFA-VEGFR2 Pathway, VEGFR2 mediated cell proliferation 
                                               SEBELIUS_2_C4636 
                                               PPP2R1B 
                                               mRNA 
                                               0 
                                               
                                               
                                               MASTL Facilitates Mitotic Progression;Misspliced GSK3beta mutants stabilize beta-catenin;T41 mutants of beta-catenin aren.SQT.t phosphorylated;ERKs are inactivated;Glycolysis;Negative regulation of MAPK pathway;Disassembly of the destruction complex and recruitment of AXIN to the membrane;S37 mutants of beta-catenin aren.SQT.t phosphorylated;Cyclin AB1 associated events during G2M transition;RAF activation;RHO GTPases Activate Formins;S33 mutants of beta-catenin aren.SQT.t phosphorylated;DARPP-32 events;Beta-catenin phosphorylation cascade;Truncations of AMER1 destabilize the destruction complex;CTLA4 inhibitory signaling;AXIN missense mutants destabilize the destruction complex;S45 mutants of beta-catenin aren.SQT.t phosphorylated;Cyclin D associated events in G1;Platelet sensitization by LDL;Resolution of Sister Chromatid Cohesion;Inhibition of replication initiation of damaged DNA by RB1E2F1;RAFMAP kinase cascade;Downstream signal transduction;VEGFR2 mediated cell proliferation;Signaling by SCF-KIT;Frs2-mediated activation;ARMS-mediated activation;Gastrin-CREB signalling pathway via PKC and MAPK;Interleukin receptor SHC signaling;FCERI mediated MAPK activation;DAP12 signaling;Signaling by PDGF;VEGFA-VEGFR2 Pathway;Signaling by ERBB2;IRS-related events triggered by IGF1R 
                                               hsa03015;hsa04071;hsa04114;hsa04151;hsa04152;hsa04261;hsa04350;hsa04390;hsa04530;hsa04728;hsa04730;hsa05142;hsa05160 
                                               PPP2R1B 
                                               NA 
                                               slateblue1 
                                               16 
                                               9.14018691588785 
                                             
                                             
                                               NRAS-mRNA 
                                               0.713 
                                               0.0909 
                                               0.535 
                                               0.892 
                                               1.64 
                                               1.45 
                                               1.86 
                                               2.17e-09 
                                               2.73e-08 
                                               lm.nb 
                                               ARMS-mediated activation, CD209 (DC-SIGN) signaling, Cell surface interactions at the vascular wall, Constitutive Signaling by EGFRvIII, DAP12 signaling, Downstream signal transduction, FCERI mediated MAPK activation, Frs2-mediated activation, Gastrin-CREB signalling pathway via PKC and MAPK, Interleukin receptor SHC signaling, IRS-related events triggered by IGF1R, MAP2K and MAPK activation, RAFMAP kinase cascade, SHC-mediated cascade.CL.FGFR1, SHC-related events triggered by IGF1R, Signaling by ERBB2, Signaling by FGFR1 in disease, Signaling by FGFR2 in disease, Signaling by FGFR3 in disease, Signaling by FGFR4 in disease, Signaling by PDGF, Signaling by SCF-KIT, Tie2 Signaling, VEGFA-VEGFR2 Pathway, VEGFR2 mediated cell proliferation 
                                               SEBELIUS_2_C4636 
                                               NRAS 
                                               mRNA 
                                               0 
                                               
                                               
                                               Signaling by FGFR2 in disease;Regulation of RAS by GAPs;SHC-mediated cascade.CL.FGFR3;Signaling by FGFR3 in disease;MAP2K and MAPK activation;SHC-related events triggered by IGF1R;Signaling by FGFR4 in disease;SHC-mediated cascade.CL.FGFR1;Tie2 Signaling;Signaling by FGFR1 in disease;Activation of RAS in B cells;SHC-mediated cascade.CL.FGFR4;Constitutive Signaling by EGFRvIII;CD209 (DC-SIGN) signaling;FCERI mediated MAPK activation;RAFMAP kinase cascade;Downstream signal transduction;VEGFR2 mediated cell proliferation;Signaling by SCF-KIT;Frs2-mediated activation;ARMS-mediated activation;Gastrin-CREB signalling pathway via PKC and MAPK;Interleukin receptor SHC signaling;DAP12 signaling;Signaling by PDGF;VEGFA-VEGFR2 Pathway;Signaling by ERBB2;IRS-related events triggered by IGF1R;Cell surface interactions at the vascular wall 
                                               hsa04010;hsa04012;hsa04014;hsa04015;hsa04062;hsa04068;hsa04071;hsa04072;hsa04151;hsa04211;hsa04213;hsa04360;hsa04370;hsa04530;hsa04540;hsa04550;hsa04650;hsa04660;hsa04662;hsa04664;hsa04720;hsa04722;hsa04725;hsa04726;hsa04730;hsa04810;hsa04910;hsa04912;hsa04915;hsa04916;hsa04917;hsa04919;hsa04921;hsa04933;hsa05034;hsa05160;hsa05161;hsa05166;hsa05200;hsa05203;hsa05205;hsa05206;hsa05211;hsa05213;hsa05214;hsa05215;hsa05216;hsa05218;hsa05219;hsa05220;hsa05221;hsa05223;hsa05230;hsa05231 
                                               NRAS 
                                               NA 
                                               slateblue1 
                                               16 
                                               7.84378437843784 
                                             
                                             
                                               RAF1-mRNA 
                                               -0.587 
                                               0.0861 
                                               -0.756 
                                               -0.418 
                                               0.666 
                                               0.592 
                                               0.748 
                                               4.99e-08 
                                               5.81e-07 
                                               loglinear 
                                               ARMS-mediated activation, CD209 (DC-SIGN) signaling, CREB phosphorylation through the activation of Ras, DAP12 signaling, Downstream signal transduction, FCERI mediated MAPK activation, Frs2-mediated activation, Gastrin-CREB signalling pathway via PKC and MAPK, Interleukin receptor SHC signaling, IRS-related events triggered by IGF1R, Negative feedback regulation of MAPK pathway, Negative regulation of MAPK pathway, RAF activation, RAFMAP kinase cascade, Signaling by ERBB2, Signaling by PDGF, Signaling by SCF-KIT, VEGFA-VEGFR2 Pathway, VEGFR2 mediated cell proliferation 
                                               SEBELIUS_2_C4636 
                                               RAF1 
                                               mRNA 
                                               0 
                                               
                                               
                                               RAF activation;Stimuli-sensing channels;CREB phosphorylation through the activation of Ras;Negative feedback regulation of MAPK pathway;CD209 (DC-SIGN) signaling;RAFMAP kinase cascade;Downstream signal transduction;VEGFR2 mediated cell proliferation;Signaling by SCF-KIT;Frs2-mediated activation;ARMS-mediated activation;Gastrin-CREB signalling pathway via PKC and MAPK;Interleukin receptor SHC signaling;FCERI mediated MAPK activation;DAP12 signaling;Signaling by PDGF;VEGFA-VEGFR2 Pathway;Signaling by ERBB2;IRS-related events triggered by IGF1R;Negative regulation of MAPK pathway 
                                               hsa04010;hsa04012;hsa04014;hsa04015;hsa04022;hsa04024;hsa04062;hsa04068;hsa04071;hsa04072;hsa04151;hsa04270;hsa04370;hsa04510;hsa04540;hsa04550;hsa04630;hsa04650;hsa04660;hsa04662;hsa04664;hsa04666;hsa04720;hsa04722;hsa04726;hsa04730;hsa04810;hsa04910;hsa04912;hsa04914;hsa04915;hsa04916;hsa04917;hsa04919;hsa04921;hsa05034;hsa05152;hsa05160;hsa05161;hsa05164;hsa05200;hsa05205;hsa05206;hsa05210;hsa05211;hsa05212;hsa05213;hsa05214;hsa05215;hsa05218;hsa05219;hsa05220;hsa05221;hsa05223;hsa05230;hsa05231 
                                               RAF1 
                                               NA 
                                               slateblue1 
                                               16 
                                               -6.81765389082462 
                                             
                                             
                                               ACTG1-mRNA 
                                               -0.902 
                                               0.146 
                                               -1.19 
                                               -0.617 
                                               0.535 
                                               0.439 
                                               0.652 
                                               3.36e-07 
                                               3.64e-06 
                                               lm.nb 
                                               ARMS-mediated activation, Cell-extracellular matrix interactions, DAP12 signaling, Downstream signal transduction, EPH-ephrin mediated repulsion of cells, EPHB-mediated forward signaling, FCERI mediated MAPK activation, Frs2-mediated activation, Gastrin-CREB signalling pathway via PKC and MAPK, Interleukin receptor SHC signaling, IRS-related events triggered by IGF1R, RAFMAP kinase cascade, Regulation of actin dynamics for phagocytic cup formation, RHO GTPases Activate Formins, RHO GTPases activate IQGAPs, RHO GTPases Activate WASPs and WAVEs, Signaling by ERBB2, Signaling by PDGF, Signaling by SCF-KIT, VEGFA-VEGFR2 Pathway, VEGFR2 mediated cell proliferation 
                                               SEBELIUS_2_C4636 
                                               ACTG1 
                                               mRNA 
                                               0 
                                               
                                               
                                               RHO GTPases Activate WASPs and WAVEs;Translocation of GLUT4 to the plasma membrane;Recycling pathway of L1;Interaction between L1 and Ankyrins;EPHB-mediated forward signaling;EPH-ephrin mediated repulsion of cells;RHO GTPases Activate Formins;RHO GTPases activate IQGAPs;Cell-extracellular matrix interactions;Regulation of actin dynamics for phagocytic cup formation;RAFMAP kinase cascade;Downstream signal transduction;VEGFR2 mediated cell proliferation;Signaling by SCF-KIT;Frs2-mediated activation;ARMS-mediated activation;Gastrin-CREB signalling pathway via PKC and MAPK;Interleukin receptor SHC signaling;FCERI mediated MAPK activation;DAP12 signaling;Signaling by PDGF;VEGFA-VEGFR2 Pathway;Signaling by ERBB2;IRS-related events triggered by IGF1R 
                                               hsa04015;hsa04145;hsa04390;hsa04510;hsa04520;hsa04530;hsa04611;hsa04670;hsa04810;hsa04919;hsa04921;hsa05100;hsa05110;hsa05130;hsa05131;hsa05132;hsa05164;hsa05205;hsa05410;hsa05412;hsa05414;hsa05416 
                                               ACTG1 
                                               NA 
                                               slateblue1 
                                               16 
                                               -6.17808219178082 
                                             
                                             
                                               CRK-mRNA 
                                               -0.616 
                                               0.1 
                                               -0.812 
                                               -0.42 
                                               0.652 
                                               0.569 
                                               0.748 
                                               3.9e-07 
                                               4.19e-06 
                                               lm.nb 
                                               ARMS-mediated activation, Downstream signal transduction, Integrin alphaIIb beta3 signaling, p130Cas linkage to MAPK signaling for integrins, Regulation of actin dynamics for phagocytic cup formation, Regulation of signaling by CBL, Signaling by PDGF, VEGFA-VEGFR2 Pathway 
                                               SEBELIUS_2_C4636 
                                               CRK 
                                               mRNA 
                                               0 
                                               
                                               
                                               ARMS-mediated activation;Downstream signal transduction;Regulation of signaling by CBL;p130Cas linkage to MAPK signaling for integrins;VEGFA-VEGFR2 Pathway;Regulation of actin dynamics for phagocytic cup formation;Signaling by PDGF;Integrin alphaIIb beta3 signaling 
                                               hsa04010;hsa04012;hsa04015;hsa04062;hsa04510;hsa04666;hsa04722;hsa04810;hsa04910;hsa05100;hsa05131;hsa05200;hsa05206;hsa05211;hsa05220 
                                               CRK 
                                               NA 
                                               slateblue1 
                                               16 
                                               -6.16 
                                             
                                             
                                               ERBB2-mRNA 
                                               1.48 
                                               0.258 
                                               0.973 
                                               1.98 
                                               2.79 
                                               1.96 
                                               3.95 
                                               1.43e-06 
                                               1.49e-05 
                                               lm.nb 
                                               ARMS-mediated activation, DAP12 signaling, Downstream signal transduction, FCERI mediated MAPK activation, Frs2-mediated activation, GAB1 signalosome, Gastrin-CREB signalling pathway via PKC and MAPK, Interleukin receptor SHC signaling, IRS-related events triggered by IGF1R, PI3KAKT activation, RAFMAP kinase cascade, Role of LAT2NTALLAB on calcium mobilization, Sema4D induced cell migration and growth-cone collapse, Signaling by ERBB2, Signaling by PDGF, Signaling by SCF-KIT, VEGFA-VEGFR2 Pathway, VEGFR2 mediated cell proliferation 
                                               SEBELIUS_2_C4636 
                                               ERBB2 
                                               mRNA 
                                               0 
                                               
                                               
                                               RAFMAP kinase cascade;GRB7 events in ERBB2 signaling;PLCG1 events in ERBB2 signaling;Sema4D induced cell migration and growth-cone collapse;Signaling by ERBB2;Downstream signal transduction;GAB1 signalosome;Signaling by SCF-KIT;PI3KAKT activation;Role of LAT2NTALLAB on calcium mobilization;DAP12 signaling;Signaling by PDGF;VEGFR2 mediated cell proliferation;Frs2-mediated activation;ARMS-mediated activation;Gastrin-CREB signalling pathway via PKC and MAPK;Interleukin receptor SHC signaling;FCERI mediated MAPK activation;VEGFA-VEGFR2 Pathway;IRS-related events triggered by IGF1R 
                                               hsa04012;hsa04020;hsa04066;hsa04510;hsa04520;hsa05200;hsa05205;hsa05206;hsa05212;hsa05213;hsa05215;hsa05219;hsa05223;hsa05230 
                                               ERBB2 
                                               NA 
                                               slateblue1 
                                               16 
                                               5.73643410852713 
                                             
                                             
                                               SPTAN1-mRNA 
                                               -0.527 
                                               0.0962 
                                               -0.716 
                                               -0.339 
                                               0.694 
                                               0.609 
                                               0.791 
                                               3.18e-06 
                                               3.24e-05 
                                               lm.nb 
                                               Apoptotic cleavage of cellular proteins, ARMS-mediated activation, DAP12 signaling, Downstream signal transduction, FCERI mediated MAPK activation, Frs2-mediated activation, Gastrin-CREB signalling pathway via PKC and MAPK, Interleukin receptor SHC signaling, IRS-related events triggered by IGF1R, Nephrin interactions, Signaling by ERBB2, Signaling by PDGF, Signaling by SCF-KIT, VEGFA-VEGFR2 Pathway, VEGFR2 mediated cell proliferation 
                                               SEBELIUS_2_C4636 
                                               SPTAN1 
                                               mRNA 
                                               0 
                                               
                                               
                                               Nephrin interactions;Caspase-mediated cleavage of cytoskeletal proteins;Downstream signal transduction;VEGFR2 mediated cell proliferation;Signaling by SCF-KIT;Frs2-mediated activation;ARMS-mediated activation;Gastrin-CREB signalling pathway via PKC and MAPK;Interleukin receptor SHC signaling;FCERI mediated MAPK activation;DAP12 signaling;Signaling by PDGF;VEGFA-VEGFR2 Pathway;Signaling by ERBB2;IRS-related events triggered by IGF1R;Apoptotic cleavage of cellular proteins 
                                               hsa04530 
                                               SPTAN1 
                                               NA 
                                               slateblue1 
                                               16 
                                               -5.47817047817048 
                                             
                                             
                                               PDGFA-mRNA 
                                               1.43 
                                               0.267 
                                               0.903 
                                               1.95 
                                               2.69 
                                               1.87 
                                               3.86 
                                               4.83e-06 
                                               4.9e-05 
                                               lm.nb 
                                               ARMS-mediated activation, Constitutive Signaling by Aberrant PI3K in Cancer, DAP12 signaling, Downstream signal transduction, FCERI mediated MAPK activation, Frs2-mediated activation, GAB1 signalosome, Gastrin-CREB signalling pathway via PKC and MAPK, Interleukin receptor SHC signaling, IRS-related events triggered by IGF1R, Non-integrin membrane-ECM interactions, PI3KAKT activation, PIP3 activates AKT signaling, Platelet degranulation , RAFMAP kinase cascade, Role of LAT2NTALLAB on calcium mobilization, Signaling by ERBB2, Signaling by PDGF, Signaling by SCF-KIT, VEGFA-VEGFR2 Pathway, VEGFR2 mediated cell proliferation 
                                               SEBELIUS_2_C4636 
                                               PDGFA 
                                               mRNA 
                                               0 
                                               
                                               
                                               Non-integrin membrane-ECM interactions;PIP3 activates AKT signaling;Constitutive Signaling by Aberrant PI3K in Cancer;RAFMAP kinase cascade;Platelet degranulation ;Signaling by PDGF;Downstream signal transduction;GAB1 signalosome;Signaling by SCF-KIT;PI3KAKT activation;Role of LAT2NTALLAB on calcium mobilization;DAP12 signaling;Signaling by ERBB2;VEGFR2 mediated cell proliferation;Frs2-mediated activation;ARMS-mediated activation;Gastrin-CREB signalling pathway via PKC and MAPK;Interleukin receptor SHC signaling;FCERI mediated MAPK activation;VEGFA-VEGFR2 Pathway;IRS-related events triggered by IGF1R 
                                               hsa04010;hsa04014;hsa04015;hsa04060;hsa04072;hsa04151;hsa04510;hsa04540;hsa04810;hsa05166;hsa05200;hsa05202;hsa05206;hsa05214;hsa05215;hsa05218;hsa05231 
                                               PDGFA 
                                               NA 
                                               slateblue1 
                                               16 
                                               5.35580524344569 
                                             
                                             
                                               ACTB-mRNA 
                                               -1.17 
                                               0.22 
                                               -1.6 
                                               -0.737 
                                               0.445 
                                               0.33 
                                               0.6 
                                               5.41e-06 
                                               5.44e-05 
                                               lm.nb 
                                               ARMS-mediated activation, Cell-extracellular matrix interactions, DAP12 signaling, Downstream signal transduction, EPH-ephrin mediated repulsion of cells, EPHB-mediated forward signaling, FCERI mediated MAPK activation, Frs2-mediated activation, Gastrin-CREB signalling pathway via PKC and MAPK, Interleukin receptor SHC signaling, IRS-related events triggered by IGF1R, RAFMAP kinase cascade, Regulation of actin dynamics for phagocytic cup formation, RHO GTPases Activate Formins, RHO GTPases activate IQGAPs, Signaling by ERBB2, Signaling by PDGF, Signaling by SCF-KIT, VEGFA-VEGFR2 Pathway, VEGFR2 mediated cell proliferation 
                                               SEBELIUS_2_C4636 
                                               ACTB 
                                               mRNA 
                                               0 
                                               
                                               
                                               EPHB-mediated forward signaling;Interaction between L1 and Ankyrins;DNA Damage Recognition in GG-NER;EPH-ephrin mediated repulsion of cells;Prefoldin mediated transfer of substrate  to CCTTriC;RHO GTPases Activate Formins;RHO GTPases activate IQGAPs;Cell-extracellular matrix interactions;Regulation of actin dynamics for phagocytic cup formation;Folding of actin by CCTTriC;B-WICH complex positively regulates rRNA expression;RAFMAP kinase cascade;Downstream signal transduction;VEGFR2 mediated cell proliferation;Signaling by SCF-KIT;Frs2-mediated activation;ARMS-mediated activation;Gastrin-CREB signalling pathway via PKC and MAPK;Interleukin receptor SHC signaling;FCERI mediated MAPK activation;DAP12 signaling;Signaling by PDGF;VEGFA-VEGFR2 Pathway;Signaling by ERBB2;IRS-related events triggered by IGF1R 
                                               hsa04015;hsa04145;hsa04390;hsa04510;hsa04520;hsa04530;hsa04611;hsa04670;hsa04810;hsa04919;hsa04921;hsa04971;hsa05100;hsa05110;hsa05130;hsa05131;hsa05132;hsa05164;hsa05205;hsa05410;hsa05412;hsa05414;hsa05416 
                                               ACTB 
                                               NA 
                                               slateblue1 
                                               16 
                                               -5.31818181818182 
                                             
                                           
                                           DE Results - ARMS-mediated activation genes - Line: HT29 vs.Fibro  Table displaying each sample's global significance scores and directed global significance scores as defined in the heatmaps above. The global significance score is calculated as the square root of the mean squared t-statistic for the genes in a gene set, with t-statistics coming from the linear regression underlying our differential expression analysis. The directed global significance score is calculated as the square root of the mean signed squared t-statistic for the genes in a gene set, with t-statistics coming from the linear regression underlying our differential expression analysis.   
                                      
                                       
                                  
                                
                              
                            
                           
                            
                          
                           
                      
                    
                    
                      
                           
                          
                               Day    Line     
                              
                                  Day  
                              
                               
                              
                                  Line: differential expression in Caco2 vs. baseline of Fibro  
                                  Line: differential expression in HEPG2 vs. baseline of Fibro  
                                  Line: differential expression in HT29 vs. baseline of Fibro  
                              
                              
                            
                              
                                
                                  
                                       
                                      
                                           Volcano Plot: Day   More Plot Information                Volcano Plot: Day  Volcano plot displaying each gene's -log10(p-value) and log2 fold change for the selected covariate. Highly statistically significant genes fall at the top of the plot, and highly differentially expressed genes fall to either side. Genes within the selected gene set are highlighted in orange. Horizontal lines indicate various False Discovery Rate (FDR) thresholds.   
                                           DE Results - AXIN missense mutants destabilize the destruction complex genes - Day   More Plot Information      Download CSV Data     
                                           
                                             
                                             
                                           
                                           
                                             
                                               Probe Label 
                                               Log2 fold change 
                                               std error (log2) 
                                               Lower confidence limit (log2) 
                                               Upper confidence limit (log2) 
                                               Linear fold change 
                                               Lower confidence limit (linear) 
                                               Upper confidence limit (linear) 
                                               P-value 
                                               BY.p.value 
                                               method 
                                               Gene.sets 
                                               Codeset.Name 
                                               Probe.Label 
                                               Analyte.Type 
                                               Is.Control 
                                               Control.Type 
                                               Related.Probes 
                                               Probe.Annotation 
                                               KEGG.Pathways 
                                               Official.Gene.Name 
                                               Control.Conc 
                                               volcanocol 
                                               volcanopch 
                                               tstats.all 
                                             
                                           
                                           
                                             
                                               PPP2R1A-mRNA 
                                               0.0412 
                                               0.0143 
                                               0.013 
                                               0.0693 
                                               1.03 
                                               1.01 
                                               1.05 
                                               0.00677 
                                               0.099 
                                               lm.nb 
                                               ARMS-mediated activation, AXIN missense mutants destabilize the destruction complex, Beta-catenin phosphorylation cascade, CTLA4 inhibitory signaling, Cyclin AB1 associated events during G2M transition, Cyclin D associated events in G1, DAP12 signaling, DARPP-32 events, Disassembly of the destruction complex and recruitment of AXIN to the membrane, Downstream signal transduction, FCERI mediated MAPK activation, Frs2-mediated activation, Gastrin-CREB signalling pathway via PKC and MAPK, Interleukin receptor SHC signaling, IRS-related events triggered by IGF1R, MASTL Facilitates Mitotic Progression, Negative regulation of FGFR1 signaling, Negative regulation of FGFR2 signaling, Negative regulation of FGFR3 signaling, Negative regulation of FGFR4 signaling, Nonsense Mediated Decay (NMD) enhanced by the Exon Junction Complex (EJC), RAF activation, RAFMAP kinase cascade, Resolution of Sister Chromatid Cohesion, RHO GTPases Activate Formins, S33 mutants of beta-catenin aren.SQT.t phosphorylated, S37 mutants of beta-catenin aren.SQT.t phosphorylated, S45 mutants of beta-catenin aren.SQT.t phosphorylated, Signaling by ERBB2, Signaling by PDGF, Signaling by SCF-KIT, Spry regulation of FGF signaling, Truncations of AMER1 destabilize the destruction complex, VEGFA-VEGFR2 Pathway, VEGFR2 mediated cell proliferation 
                                               SEBELIUS_2_C4636 
                                               PPP2R1A 
                                               mRNA 
                                               0 
                                               
                                               
                                               MASTL Facilitates Mitotic Progression;Regulation of PLK1 Activity at G2M Transition;Disassembly of the destruction complex and recruitment of AXIN to the membrane;S37 mutants of beta-catenin aren.SQT.t phosphorylated;Cyclin AB1 associated events during G2M transition;RAF activation;RHO GTPases Activate Formins;S33 mutants of beta-catenin aren.SQT.t phosphorylated;Loss of Nlp from mitotic centrosomes;DARPP-32 events;Beta-catenin phosphorylation cascade;Initiation of Nuclear Envelope Reformation;Recruitment of mitotic centrosome proteins and complexes;Spry regulation of FGF signaling;Truncations of AMER1 destabilize the destruction complex;CTLA4 inhibitory signaling;AXIN missense mutants destabilize the destruction complex;S45 mutants of beta-catenin aren.SQT.t phosphorylated;Anchoring of the basal body to the plasma membrane;Nonsense Mediated Decay (NMD) enhanced by the Exon Junction Complex (EJC);Cyclin D associated events in G1;Platelet sensitization by LDL;Resolution of Sister Chromatid Cohesion;Inhibition of replication initiation of damaged DNA by RB1E2F1;RAFMAP kinase cascade;Downstream signal transduction;VEGFR2 mediated cell proliferation;Signaling by SCF-KIT;Frs2-mediated activation;ARMS-mediated activation;Gastrin-CREB signalling pathway via PKC and MAPK;Interleukin receptor SHC signaling;FCERI mediated MAPK activation;DAP12 signaling;Signaling by PDGF;VEGFA-VEGFR2 Pathway;Signaling by ERBB2;IRS-related events triggered by IGF1R;Negative regulation of FGFR3 signaling;Negative regulation of FGFR4 signaling;Negative regulation of FGFR2 signaling;Negative regulation of FGFR1 signaling 
                                               hsa03015;hsa04071;hsa04114;hsa04151;hsa04152;hsa04261;hsa04350;hsa04390;hsa04530;hsa04728;hsa04730;hsa05142;hsa05160 
                                               PPP2R1A 
                                               NA 
                                               slateblue4 
                                               16 
                                               2.88111888111888 
                                             
                                             
                                               AXIN1-mRNA 
                                               0.091 
                                               0.0371 
                                               0.0182 
                                               0.164 
                                               1.07 
                                               1.01 
                                               1.12 
                                               0.0191 
                                               0.24 
                                               lm.nb 
                                               AXIN missense mutants destabilize the destruction complex, Beta-catenin phosphorylation cascade, Disassembly of the destruction complex and recruitment of AXIN to the membrane, Misspliced GSK3beta mutants stabilize beta-catenin, S33 mutants of beta-catenin aren.SQT.t phosphorylated, S37 mutants of beta-catenin aren.SQT.t phosphorylated, S45 mutants of beta-catenin aren.SQT.t phosphorylated, T41 mutants of beta-catenin aren.SQT.t phosphorylated, Truncations of AMER1 destabilize the destruction complex 
                                               SEBELIUS_2_C4636 
                                               AXIN1 
                                               mRNA 
                                               0 
                                               
                                               
                                               APC truncation mutants have impaired AXIN binding;Misspliced GSK3beta mutants stabilize beta-catenin;T41 mutants of beta-catenin aren.SQT.t phosphorylated;AXIN missense mutants destabilize the destruction complex;S45 mutants of beta-catenin aren.SQT.t phosphorylated;Disassembly of the destruction complex and recruitment of AXIN to the membrane;S37 mutants of beta-catenin aren.SQT.t phosphorylated;S33 mutants of beta-catenin aren.SQT.t phosphorylated;Beta-catenin phosphorylation cascade;Truncations of AMER1 destabilize the destruction complex 
                                               hsa04310;hsa04390;hsa04550;hsa05200;hsa05210;hsa05213;hsa05217 
                                               AXIN1 
                                               NA 
                                               slateblue4 
                                               16 
                                               2.45283018867925 
                                             
                                             
                                               PPP2CB-mRNA 
                                               -0.0296 
                                               0.0132 
                                               -0.0554 
                                               -0.0037 
                                               0.98 
                                               0.962 
                                               0.997 
                                               0.0311 
                                               0.348 
                                               lm.nb 
                                               ARMS-mediated activation, AXIN missense mutants destabilize the destruction complex, Beta-catenin phosphorylation cascade, CTLA4 inhibitory signaling, Cyclin AB1 associated events during G2M transition, Cyclin D associated events in G1, DAP12 signaling, DARPP-32 events, Disassembly of the destruction complex and recruitment of AXIN to the membrane, Downstream signal transduction, ERKs are inactivated, FCERI mediated MAPK activation, Frs2-mediated activation, Gastrin-CREB signalling pathway via PKC and MAPK, Interleukin receptor SHC signaling, IRS-related events triggered by IGF1R, MASTL Facilitates Mitotic Progression, Misspliced GSK3beta mutants stabilize beta-catenin, Negative regulation of FGFR1 signaling, Negative regulation of FGFR2 signaling, Negative regulation of FGFR3 signaling, Negative regulation of FGFR4 signaling, Negative regulation of MAPK pathway, RAF activation, RAFMAP kinase cascade, Resolution of Sister Chromatid Cohesion, RHO GTPases Activate Formins, S33 mutants of beta-catenin aren.SQT.t phosphorylated, S37 mutants of beta-catenin aren.SQT.t phosphorylated, S45 mutants of beta-catenin aren.SQT.t phosphorylated, Signaling by ERBB2, Signaling by PDGF, Signaling by SCF-KIT, Spry regulation of FGF signaling, T41 mutants of beta-catenin aren.SQT.t phosphorylated, Truncations of AMER1 destabilize the destruction complex, VEGFA-VEGFR2 Pathway, VEGFR2 mediated cell proliferation 
                                               SEBELIUS_2_C4636 
                                               PPP2CB 
                                               mRNA 
                                               0 
                                               
                                               
                                               MASTL Facilitates Mitotic Progression;Misspliced GSK3beta mutants stabilize beta-catenin;T41 mutants of beta-catenin aren.SQT.t phosphorylated;ERKs are inactivated;Glycolysis;Negative regulation of MAPK pathway;Disassembly of the destruction complex and recruitment of AXIN to the membrane;S37 mutants of beta-catenin aren.SQT.t phosphorylated;Cyclin AB1 associated events during G2M transition;RAF activation;RHO GTPases Activate Formins;S33 mutants of beta-catenin aren.SQT.t phosphorylated;DARPP-32 events;Beta-catenin phosphorylation cascade;Spry regulation of FGF signaling;Truncations of AMER1 destabilize the destruction complex;CTLA4 inhibitory signaling;AXIN missense mutants destabilize the destruction complex;S45 mutants of beta-catenin aren.SQT.t phosphorylated;Cyclin D associated events in G1;Platelet sensitization by LDL;Resolution of Sister Chromatid Cohesion;Inhibition of replication initiation of damaged DNA by RB1E2F1;RAFMAP kinase cascade;Downstream signal transduction;VEGFR2 mediated cell proliferation;Signaling by SCF-KIT;Frs2-mediated activation;ARMS-mediated activation;Gastrin-CREB signalling pathway via PKC and MAPK;Interleukin receptor SHC signaling;FCERI mediated MAPK activation;DAP12 signaling;Signaling by PDGF;VEGFA-VEGFR2 Pathway;Signaling by ERBB2;IRS-related events triggered by IGF1R;Negative regulation of FGFR3 signaling;Negative regulation of FGFR4 signaling;Negative regulation of FGFR2 signaling;Negative regulation of FGFR1 signaling 
                                               hsa03015;hsa04071;hsa04114;hsa04151;hsa04152;hsa04261;hsa04350;hsa04390;hsa04530;hsa04728;hsa04730;hsa05142;hsa05160 
                                               PPP2CB 
                                               NA 
                                               slateblue4 
                                               16 
                                               -2.24242424242424 
                                             
                                             
                                               APC-mRNA 
                                               0.0285 
                                               0.0157 
                                               -0.0023 
                                               0.0592 
                                               1.02 
                                               0.998 
                                               1.04 
                                               0.0779 
                                               0.762 
                                               lm.nb 
                                               Apoptotic cleavage of cellular proteins, AXIN missense mutants destabilize the destruction complex, Beta-catenin phosphorylation cascade, Deactivation of the beta-catenin transactivating complex, Disassembly of the destruction complex and recruitment of AXIN to the membrane, Misspliced GSK3beta mutants stabilize beta-catenin, S33 mutants of beta-catenin aren.SQT.t phosphorylated, S37 mutants of beta-catenin aren.SQT.t phosphorylated, S45 mutants of beta-catenin aren.SQT.t phosphorylated, T41 mutants of beta-catenin aren.SQT.t phosphorylated, Truncations of AMER1 destabilize the destruction complex 
                                               SEBELIUS_2_C4636 
                                               APC 
                                               mRNA 
                                               0 
                                               
                                               
                                               Apoptotic cleavage of cellular proteins;Misspliced GSK3beta mutants stabilize beta-catenin;APC truncation mutants have impaired AXIN binding;T41 mutants of beta-catenin aren.SQT.t phosphorylated;AXIN missense mutants destabilize the destruction complex;S45 mutants of beta-catenin aren.SQT.t phosphorylated;Deactivation of the beta-catenin transactivating complex;Disassembly of the destruction complex and recruitment of AXIN to the membrane;S37 mutants of beta-catenin aren.SQT.t phosphorylated;S33 mutants of beta-catenin aren.SQT.t phosphorylated;Beta-catenin phosphorylation cascade;Truncations of AMER1 destabilize the destruction complex 
                                               hsa04310;hsa04390;hsa04550;hsa04810;hsa05166;hsa05200;hsa05206;hsa05210;hsa05213;hsa05217 
                                               APC 
                                               NA 
                                               azure4 
                                               1 
                                               1.81528662420382 
                                             
                                             
                                               PPP2CA-mRNA 
                                               -0.026 
                                               0.0146 
                                               -0.0547 
                                               0.00266 
                                               0.982 
                                               0.963 
                                               1 
                                               0.0836 
                                               0.802 
                                               lm.nb 
                                               ARMS-mediated activation, AXIN missense mutants destabilize the destruction complex, Beta-catenin phosphorylation cascade, CTLA4 inhibitory signaling, Cyclin AB1 associated events during G2M transition, Cyclin D associated events in G1, DAP12 signaling, DARPP-32 events, Disassembly of the destruction complex and recruitment of AXIN to the membrane, Downstream signal transduction, ERKs are inactivated, FCERI mediated MAPK activation, Frs2-mediated activation, Gastrin-CREB signalling pathway via PKC and MAPK, Interleukin receptor SHC signaling, IRS-related events triggered by IGF1R, MASTL Facilitates Mitotic Progression, Negative regulation of FGFR1 signaling, Negative regulation of FGFR2 signaling, Negative regulation of FGFR3 signaling, Negative regulation of FGFR4 signaling, Negative regulation of MAPK pathway, Nonsense Mediated Decay (NMD) enhanced by the Exon Junction Complex (EJC), RAF activation, RAFMAP kinase cascade, Resolution of Sister Chromatid Cohesion, RHO GTPases Activate Formins, S33 mutants of beta-catenin aren.SQT.t phosphorylated, S37 mutants of beta-catenin aren.SQT.t phosphorylated, S45 mutants of beta-catenin aren.SQT.t phosphorylated, Signaling by ERBB2, Signaling by PDGF, Signaling by SCF-KIT, Spry regulation of FGF signaling, T41 mutants of beta-catenin aren.SQT.t phosphorylated, Truncations of AMER1 destabilize the destruction complex, VEGFA-VEGFR2 Pathway, VEGFR2 mediated cell proliferation 
                                               SEBELIUS_2_C4636 
                                               PPP2CA 
                                               mRNA 
                                               0 
                                               
                                               
                                               MASTL Facilitates Mitotic Progression;T41 mutants of beta-catenin aren.SQT.t phosphorylated;ERKs are inactivated;Glycolysis;Negative regulation of MAPK pathway;Disassembly of the destruction complex and recruitment of AXIN to the membrane;S37 mutants of beta-catenin aren.SQT.t phosphorylated;Cyclin AB1 associated events during G2M transition;RAF activation;RHO GTPases Activate Formins;S33 mutants of beta-catenin aren.SQT.t phosphorylated;DARPP-32 events;Beta-catenin phosphorylation cascade;Initiation of Nuclear Envelope Reformation;Spry regulation of FGF signaling;Truncations of AMER1 destabilize the destruction complex;CTLA4 inhibitory signaling;AXIN missense mutants destabilize the destruction complex;S45 mutants of beta-catenin aren.SQT.t phosphorylated;Nonsense Mediated Decay (NMD) enhanced by the Exon Junction Complex (EJC);Cyclin D associated events in G1;Platelet sensitization by LDL;Resolution of Sister Chromatid Cohesion;Inhibition of replication initiation of damaged DNA by RB1E2F1;RAFMAP kinase cascade;Downstream signal transduction;VEGFR2 mediated cell proliferation;Signaling by SCF-KIT;Frs2-mediated activation;ARMS-mediated activation;Gastrin-CREB signalling pathway via PKC and MAPK;Interleukin receptor SHC signaling;FCERI mediated MAPK activation;DAP12 signaling;Signaling by PDGF;VEGFA-VEGFR2 Pathway;Signaling by ERBB2;IRS-related events triggered by IGF1R;Negative regulation of FGFR3 signaling;Negative regulation of FGFR4 signaling;Negative regulation of FGFR2 signaling;Negative regulation of FGFR1 signaling 
                                               hsa03015;hsa04071;hsa04114;hsa04151;hsa04152;hsa04261;hsa04350;hsa04390;hsa04530;hsa04728;hsa04730;hsa05142;hsa05160 
                                               PPP2CA 
                                               NA 
                                               azure4 
                                               1 
                                               -1.78082191780822 
                                             
                                             
                                               GSK3B-mRNA 
                                               -0.0158 
                                               0.0129 
                                               -0.041 
                                               0.0095 
                                               0.989 
                                               0.972 
                                               1.01 
                                               0.229 
                                               1 
                                               lm.nb 
                                               AXIN missense mutants destabilize the destruction complex, DAP12 signaling, Disassembly of the destruction complex and recruitment of AXIN to the membrane, Downstream signal transduction, GAB1 signalosome, Misspliced GSK3beta mutants stabilize beta-catenin, PI3KAKT activation, PIP3 activates AKT signaling, Regulation of HSF1-mediated heat shock response, Role of LAT2NTALLAB on calcium mobilization, S33 mutants of beta-catenin aren.SQT.t phosphorylated, S37 mutants of beta-catenin aren.SQT.t phosphorylated, S45 mutants of beta-catenin aren.SQT.t phosphorylated, Signaling by ERBB2, Signaling by PDGF, Signaling by SCF-KIT, T41 mutants of beta-catenin aren.SQT.t phosphorylated, Truncations of AMER1 destabilize the destruction complex 
                                               SEBELIUS_2_C4636 
                                               GSK3B 
                                               mRNA 
                                               0 
                                               
                                               
                                               Misspliced GSK3beta mutants stabilize beta-catenin;Constitutive Signaling by AKT1 E17K in Cancer;APC truncation mutants have impaired AXIN binding;T41 mutants of beta-catenin aren.SQT.t phosphorylated;AXIN missense mutants destabilize the destruction complex;S45 mutants of beta-catenin aren.SQT.t phosphorylated;Regulation of HSF1-mediated heat shock response;AKT phosphorylates targets in the cytosol;Disassembly of the destruction complex and recruitment of AXIN to the membrane;S37 mutants of beta-catenin aren.SQT.t phosphorylated;S33 mutants of beta-catenin aren.SQT.t phosphorylated;Truncations of AMER1 destabilize the destruction complex;Degradation of GLI2 by the proteasome;PIP3 activates AKT signaling;Downstream signal transduction;GAB1 signalosome;Signaling by SCF-KIT;PI3KAKT activation;Role of LAT2NTALLAB on calcium mobilization;DAP12 signaling;Signaling by PDGF;Signaling by ERBB2 
                                               hsa04012;hsa04062;hsa04110;hsa04151;hsa04310;hsa04360;hsa04390;hsa04510;hsa04550;hsa04660;hsa04662;hsa04722;hsa04728;hsa04910;hsa04916;hsa04917;hsa04919;hsa04931;hsa04932;hsa05010;hsa05160;hsa05162;hsa05164;hsa05166;hsa05169;hsa05200;hsa05210;hsa05213;hsa05215;hsa05217 
                                               GSK3B 
                                               NA 
                                               azure4 
                                               1 
                                               -1.22480620155039 
                                             
                                             
                                               PPP2R1B-mRNA 
                                               0.0114 
                                               0.0135 
                                               -0.015 
                                               0.0379 
                                               1.01 
                                               0.99 
                                               1.03 
                                               0.402 
                                               1 
                                               lm.nb 
                                               ARMS-mediated activation, AXIN missense mutants destabilize the destruction complex, Beta-catenin phosphorylation cascade, CTLA4 inhibitory signaling, Cyclin AB1 associated events during G2M transition, Cyclin D associated events in G1, DAP12 signaling, DARPP-32 events, Disassembly of the destruction complex and recruitment of AXIN to the membrane, Downstream signal transduction, ERKs are inactivated, FCERI mediated MAPK activation, Frs2-mediated activation, Gastrin-CREB signalling pathway via PKC and MAPK, Interleukin receptor SHC signaling, IRS-related events triggered by IGF1R, MASTL Facilitates Mitotic Progression, Misspliced GSK3beta mutants stabilize beta-catenin, Negative regulation of MAPK pathway, RAF activation, RAFMAP kinase cascade, Resolution of Sister Chromatid Cohesion, RHO GTPases Activate Formins, S33 mutants of beta-catenin aren.SQT.t phosphorylated, S37 mutants of beta-catenin aren.SQT.t phosphorylated, S45 mutants of beta-catenin aren.SQT.t phosphorylated, Signaling by ERBB2, Signaling by PDGF, Signaling by SCF-KIT, T41 mutants of beta-catenin aren.SQT.t phosphorylated, Truncations of AMER1 destabilize the destruction complex, VEGFA-VEGFR2 Pathway, VEGFR2 mediated cell proliferation 
                                               SEBELIUS_2_C4636 
                                               PPP2R1B 
                                               mRNA 
                                               0 
                                               
                                               
                                               MASTL Facilitates Mitotic Progression;Misspliced GSK3beta mutants stabilize beta-catenin;T41 mutants of beta-catenin aren.SQT.t phosphorylated;ERKs are inactivated;Glycolysis;Negative regulation of MAPK pathway;Disassembly of the destruction complex and recruitment of AXIN to the membrane;S37 mutants of beta-catenin aren.SQT.t phosphorylated;Cyclin AB1 associated events during G2M transition;RAF activation;RHO GTPases Activate Formins;S33 mutants of beta-catenin aren.SQT.t phosphorylated;DARPP-32 events;Beta-catenin phosphorylation cascade;Truncations of AMER1 destabilize the destruction complex;CTLA4 inhibitory signaling;AXIN missense mutants destabilize the destruction complex;S45 mutants of beta-catenin aren.SQT.t phosphorylated;Cyclin D associated events in G1;Platelet sensitization by LDL;Resolution of Sister Chromatid Cohesion;Inhibition of replication initiation of damaged DNA by RB1E2F1;RAFMAP kinase cascade;Downstream signal transduction;VEGFR2 mediated cell proliferation;Signaling by SCF-KIT;Frs2-mediated activation;ARMS-mediated activation;Gastrin-CREB signalling pathway via PKC and MAPK;Interleukin receptor SHC signaling;FCERI mediated MAPK activation;DAP12 signaling;Signaling by PDGF;VEGFA-VEGFR2 Pathway;Signaling by ERBB2;IRS-related events triggered by IGF1R 
                                               hsa03015;hsa04071;hsa04114;hsa04151;hsa04152;hsa04261;hsa04350;hsa04390;hsa04530;hsa04728;hsa04730;hsa05142;hsa05160 
                                               PPP2R1B 
                                               NA 
                                               azure4 
                                               1 
                                               0.844444444444444 
                                             
                                             
                                               CSNK1A1-mRNA 
                                               0.0029 
                                               0.0109 
                                               -0.0186 
                                               0.0244 
                                               1 
                                               0.987 
                                               1.02 
                                               0.792 
                                               1 
                                               loglinear 
                                               AXIN missense mutants destabilize the destruction complex, Beta-catenin phosphorylation cascade, Disassembly of the destruction complex and recruitment of AXIN to the membrane, Misspliced GSK3beta mutants stabilize beta-catenin, S33 mutants of beta-catenin aren.SQT.t phosphorylated, S37 mutants of beta-catenin aren.SQT.t phosphorylated, S45 mutants of beta-catenin aren.SQT.t phosphorylated, T41 mutants of beta-catenin aren.SQT.t phosphorylated 
                                               SEBELIUS_2_C4636 
                                               CSNK1A1 
                                               mRNA 
                                               0 
                                               
                                               
                                               Misspliced GSK3beta mutants stabilize beta-catenin;APC truncation mutants have impaired AXIN binding;T41 mutants of beta-catenin aren.SQT.t phosphorylated;AXIN missense mutants destabilize the destruction complex;S45 mutants of beta-catenin aren.SQT.t phosphorylated;Disassembly of the destruction complex and recruitment of AXIN to the membrane;S37 mutants of beta-catenin aren.SQT.t phosphorylated;S33 mutants of beta-catenin aren.SQT.t phosphorylated;Beta-catenin phosphorylation cascade;Degradation of GLI2 by the proteasome 
                                               hsa04310 
                                               CSNK1A1 
                                               NA 
                                               azure4 
                                               1 
                                               0.26605504587156 
                                             
                                           
                                           DE Results - AXIN missense mutants destabilize the destruction complex genes - Day  Table displaying each sample's global significance scores and directed global significance scores as defined in the heatmaps above. The global significance score is calculated as the square root of the mean squared t-statistic for the genes in a gene set, with t-statistics coming from the linear regression underlying our differential expression analysis. The directed global significance score is calculated as the square root of the mean signed squared t-statistic for the genes in a gene set, with t-statistics coming from the linear regression underlying our differential expression analysis.   
                                      
                                       
                                  
                                
                              
                            
                            
                              
                                
                                  
                                       
                                      
                                           Volcano Plot: Line: Caco2 vs.Fibro   More Plot Information                Volcano Plot: Line: Caco2 vs.Fibro  Volcano plot displaying each gene's -log10(p-value) and log2 fold change for the selected covariate. Highly statistically significant genes fall at the top of the plot, and highly differentially expressed genes fall to either side. Genes within the selected gene set are highlighted in orange. Horizontal lines indicate various False Discovery Rate (FDR) thresholds.   
                                           DE Results - AXIN missense mutants destabilize the destruction complex genes - Line: Caco2 vs.Fibro   More Plot Information      Download CSV Data     
                                           
                                             
                                             
                                           
                                           
                                             
                                               Probe Label 
                                               Log2 fold change 
                                               std error (log2) 
                                               Lower confidence limit (log2) 
                                               Upper confidence limit (log2) 
                                               Linear fold change 
                                               Lower confidence limit (linear) 
                                               Upper confidence limit (linear) 
                                               P-value 
                                               BY.p.value 
                                               method 
                                               Gene.sets 
                                               Codeset.Name 
                                               Probe.Label 
                                               Analyte.Type 
                                               Is.Control 
                                               Control.Type 
                                               Related.Probes 
                                               Probe.Annotation 
                                               KEGG.Pathways 
                                               Official.Gene.Name 
                                               Control.Conc 
                                               volcanocol 
                                               volcanopch 
                                               tstats.all 
                                             
                                           
                                           
                                             
                                               PPP2R1B-mRNA 
                                               1.57 
                                               0.104 
                                               1.37 
                                               1.78 
                                               2.98 
                                               2.59 
                                               3.43 
                                               1.73e-17 
                                               6.65e-16 
                                               lm.nb 
                                               ARMS-mediated activation, AXIN missense mutants destabilize the destruction complex, Beta-catenin phosphorylation cascade, CTLA4 inhibitory signaling, Cyclin AB1 associated events during G2M transition, Cyclin D associated events in G1, DAP12 signaling, DARPP-32 events, Disassembly of the destruction complex and recruitment of AXIN to the membrane, Downstream signal transduction, ERKs are inactivated, FCERI mediated MAPK activation, Frs2-mediated activation, Gastrin-CREB signalling pathway via PKC and MAPK, Interleukin receptor SHC signaling, IRS-related events triggered by IGF1R, MASTL Facilitates Mitotic Progression, Misspliced GSK3beta mutants stabilize beta-catenin, Negative regulation of MAPK pathway, RAF activation, RAFMAP kinase cascade, Resolution of Sister Chromatid Cohesion, RHO GTPases Activate Formins, S33 mutants of beta-catenin aren.SQT.t phosphorylated, S37 mutants of beta-catenin aren.SQT.t phosphorylated, S45 mutants of beta-catenin aren.SQT.t phosphorylated, Signaling by ERBB2, Signaling by PDGF, Signaling by SCF-KIT, T41 mutants of beta-catenin aren.SQT.t phosphorylated, Truncations of AMER1 destabilize the destruction complex, VEGFA-VEGFR2 Pathway, VEGFR2 mediated cell proliferation 
                                               SEBELIUS_2_C4636 
                                               PPP2R1B 
                                               mRNA 
                                               0 
                                               
                                               
                                               MASTL Facilitates Mitotic Progression;Misspliced GSK3beta mutants stabilize beta-catenin;T41 mutants of beta-catenin aren.SQT.t phosphorylated;ERKs are inactivated;Glycolysis;Negative regulation of MAPK pathway;Disassembly of the destruction complex and recruitment of AXIN to the membrane;S37 mutants of beta-catenin aren.SQT.t phosphorylated;Cyclin AB1 associated events during G2M transition;RAF activation;RHO GTPases Activate Formins;S33 mutants of beta-catenin aren.SQT.t phosphorylated;DARPP-32 events;Beta-catenin phosphorylation cascade;Truncations of AMER1 destabilize the destruction complex;CTLA4 inhibitory signaling;AXIN missense mutants destabilize the destruction complex;S45 mutants of beta-catenin aren.SQT.t phosphorylated;Cyclin D associated events in G1;Platelet sensitization by LDL;Resolution of Sister Chromatid Cohesion;Inhibition of replication initiation of damaged DNA by RB1E2F1;RAFMAP kinase cascade;Downstream signal transduction;VEGFR2 mediated cell proliferation;Signaling by SCF-KIT;Frs2-mediated activation;ARMS-mediated activation;Gastrin-CREB signalling pathway via PKC and MAPK;Interleukin receptor SHC signaling;FCERI mediated MAPK activation;DAP12 signaling;Signaling by PDGF;VEGFA-VEGFR2 Pathway;Signaling by ERBB2;IRS-related events triggered by IGF1R 
                                               hsa03015;hsa04071;hsa04114;hsa04151;hsa04152;hsa04261;hsa04350;hsa04390;hsa04530;hsa04728;hsa04730;hsa05142;hsa05160 
                                               PPP2R1B 
                                               NA 
                                               slateblue1 
                                               16 
                                               15.0961538461538 
                                             
                                             
                                               PPP2CB-mRNA 
                                               -0.814 
                                               0.099 
                                               -1.01 
                                               -0.62 
                                               0.569 
                                               0.497 
                                               0.651 
                                               7.15e-10 
                                               1.11e-08 
                                               lm.nb 
                                               ARMS-mediated activation, AXIN missense mutants destabilize the destruction complex, Beta-catenin phosphorylation cascade, CTLA4 inhibitory signaling, Cyclin AB1 associated events during G2M transition, Cyclin D associated events in G1, DAP12 signaling, DARPP-32 events, Disassembly of the destruction complex and recruitment of AXIN to the membrane, Downstream signal transduction, ERKs are inactivated, FCERI mediated MAPK activation, Frs2-mediated activation, Gastrin-CREB signalling pathway via PKC and MAPK, Interleukin receptor SHC signaling, IRS-related events triggered by IGF1R, MASTL Facilitates Mitotic Progression, Misspliced GSK3beta mutants stabilize beta-catenin, Negative regulation of FGFR1 signaling, Negative regulation of FGFR2 signaling, Negative regulation of FGFR3 signaling, Negative regulation of FGFR4 signaling, Negative regulation of MAPK pathway, RAF activation, RAFMAP kinase cascade, Resolution of Sister Chromatid Cohesion, RHO GTPases Activate Formins, S33 mutants of beta-catenin aren.SQT.t phosphorylated, S37 mutants of beta-catenin aren.SQT.t phosphorylated, S45 mutants of beta-catenin aren.SQT.t phosphorylated, Signaling by ERBB2, Signaling by PDGF, Signaling by SCF-KIT, Spry regulation of FGF signaling, T41 mutants of beta-catenin aren.SQT.t phosphorylated, Truncations of AMER1 destabilize the destruction complex, VEGFA-VEGFR2 Pathway, VEGFR2 mediated cell proliferation 
                                               SEBELIUS_2_C4636 
                                               PPP2CB 
                                               mRNA 
                                               0 
                                               
                                               
                                               MASTL Facilitates Mitotic Progression;Misspliced GSK3beta mutants stabilize beta-catenin;T41 mutants of beta-catenin aren.SQT.t phosphorylated;ERKs are inactivated;Glycolysis;Negative regulation of MAPK pathway;Disassembly of the destruction complex and recruitment of AXIN to the membrane;S37 mutants of beta-catenin aren.SQT.t phosphorylated;Cyclin AB1 associated events during G2M transition;RAF activation;RHO GTPases Activate Formins;S33 mutants of beta-catenin aren.SQT.t phosphorylated;DARPP-32 events;Beta-catenin phosphorylation cascade;Spry regulation of FGF signaling;Truncations of AMER1 destabilize the destruction complex;CTLA4 inhibitory signaling;AXIN missense mutants destabilize the destruction complex;S45 mutants of beta-catenin aren.SQT.t phosphorylated;Cyclin D associated events in G1;Platelet sensitization by LDL;Resolution of Sister Chromatid Cohesion;Inhibition of replication initiation of damaged DNA by RB1E2F1;RAFMAP kinase cascade;Downstream signal transduction;VEGFR2 mediated cell proliferation;Signaling by SCF-KIT;Frs2-mediated activation;ARMS-mediated activation;Gastrin-CREB signalling pathway via PKC and MAPK;Interleukin receptor SHC signaling;FCERI mediated MAPK activation;DAP12 signaling;Signaling by PDGF;VEGFA-VEGFR2 Pathway;Signaling by ERBB2;IRS-related events triggered by IGF1R;Negative regulation of FGFR3 signaling;Negative regulation of FGFR4 signaling;Negative regulation of FGFR2 signaling;Negative regulation of FGFR1 signaling 
                                               hsa03015;hsa04071;hsa04114;hsa04151;hsa04152;hsa04261;hsa04350;hsa04390;hsa04530;hsa04728;hsa04730;hsa05142;hsa05160 
                                               PPP2CB 
                                               NA 
                                               slateblue1 
                                               16 
                                               -8.22222222222222 
                                             
                                             
                                               APC-mRNA 
                                               -0.862 
                                               0.115 
                                               -1.09 
                                               -0.637 
                                               0.55 
                                               0.471 
                                               0.643 
                                               6.14e-09 
                                               8.54e-08 
                                               lm.nb 
                                               Apoptotic cleavage of cellular proteins, AXIN missense mutants destabilize the destruction complex, Beta-catenin phosphorylation cascade, Deactivation of the beta-catenin transactivating complex, Disassembly of the destruction complex and recruitment of AXIN to the membrane, Misspliced GSK3beta mutants stabilize beta-catenin, S33 mutants of beta-catenin aren.SQT.t phosphorylated, S37 mutants of beta-catenin aren.SQT.t phosphorylated, S45 mutants of beta-catenin aren.SQT.t phosphorylated, T41 mutants of beta-catenin aren.SQT.t phosphorylated, Truncations of AMER1 destabilize the destruction complex 
                                               SEBELIUS_2_C4636 
                                               APC 
                                               mRNA 
                                               0 
                                               
                                               
                                               Apoptotic cleavage of cellular proteins;Misspliced GSK3beta mutants stabilize beta-catenin;APC truncation mutants have impaired AXIN binding;T41 mutants of beta-catenin aren.SQT.t phosphorylated;AXIN missense mutants destabilize the destruction complex;S45 mutants of beta-catenin aren.SQT.t phosphorylated;Deactivation of the beta-catenin transactivating complex;Disassembly of the destruction complex and recruitment of AXIN to the membrane;S37 mutants of beta-catenin aren.SQT.t phosphorylated;S33 mutants of beta-catenin aren.SQT.t phosphorylated;Beta-catenin phosphorylation cascade;Truncations of AMER1 destabilize the destruction complex 
                                               hsa04310;hsa04390;hsa04550;hsa04810;hsa05166;hsa05200;hsa05206;hsa05210;hsa05213;hsa05217 
                                               APC 
                                               NA 
                                               slateblue1 
                                               16 
                                               -7.49565217391304 
                                             
                                             
                                               CSNK1A1-mRNA 
                                               -0.579 
                                               0.0828 
                                               -0.741 
                                               -0.417 
                                               0.669 
                                               0.598 
                                               0.749 
                                               2.88e-08 
                                               3.73e-07 
                                               loglinear 
                                               AXIN missense mutants destabilize the destruction complex, Beta-catenin phosphorylation cascade, Disassembly of the destruction complex and recruitment of AXIN to the membrane, Misspliced GSK3beta mutants stabilize beta-catenin, S33 mutants of beta-catenin aren.SQT.t phosphorylated, S37 mutants of beta-catenin aren.SQT.t phosphorylated, S45 mutants of beta-catenin aren.SQT.t phosphorylated, T41 mutants of beta-catenin aren.SQT.t phosphorylated 
                                               SEBELIUS_2_C4636 
                                               CSNK1A1 
                                               mRNA 
                                               0 
                                               
                                               
                                               Misspliced GSK3beta mutants stabilize beta-catenin;APC truncation mutants have impaired AXIN binding;T41 mutants of beta-catenin aren.SQT.t phosphorylated;AXIN missense mutants destabilize the destruction complex;S45 mutants of beta-catenin aren.SQT.t phosphorylated;Disassembly of the destruction complex and recruitment of AXIN to the membrane;S37 mutants of beta-catenin aren.SQT.t phosphorylated;S33 mutants of beta-catenin aren.SQT.t phosphorylated;Beta-catenin phosphorylation cascade;Degradation of GLI2 by the proteasome 
                                               hsa04310 
                                               CSNK1A1 
                                               NA 
                                               slateblue1 
                                               16 
                                               -6.99275362318841 
                                             
                                             
                                               AXIN1-mRNA 
                                               0.571 
                                               0.285 
                                               0.012 
                                               1.13 
                                               1.49 
                                               1.01 
                                               2.19 
                                               0.0526 
                                               0.397 
                                               lm.nb 
                                               AXIN missense mutants destabilize the destruction complex, Beta-catenin phosphorylation cascade, Disassembly of the destruction complex and recruitment of AXIN to the membrane, Misspliced GSK3beta mutants stabilize beta-catenin, S33 mutants of beta-catenin aren.SQT.t phosphorylated, S37 mutants of beta-catenin aren.SQT.t phosphorylated, S45 mutants of beta-catenin aren.SQT.t phosphorylated, T41 mutants of beta-catenin aren.SQT.t phosphorylated, Truncations of AMER1 destabilize the destruction complex 
                                               SEBELIUS_2_C4636 
                                               AXIN1 
                                               mRNA 
                                               0 
                                               
                                               
                                               APC truncation mutants have impaired AXIN binding;Misspliced GSK3beta mutants stabilize beta-catenin;T41 mutants of beta-catenin aren.SQT.t phosphorylated;AXIN missense mutants destabilize the destruction complex;S45 mutants of beta-catenin aren.SQT.t phosphorylated;Disassembly of the destruction complex and recruitment of AXIN to the membrane;S37 mutants of beta-catenin aren.SQT.t phosphorylated;S33 mutants of beta-catenin aren.SQT.t phosphorylated;Beta-catenin phosphorylation cascade;Truncations of AMER1 destabilize the destruction complex 
                                               hsa04310;hsa04390;hsa04550;hsa05200;hsa05210;hsa05213;hsa05217 
                                               AXIN1 
                                               NA 
                                               slateblue4 
                                               16 
                                               2.00350877192982 
                                             
                                             
                                               PPP2CA-mRNA 
                                               0.168 
                                               0.111 
                                               -0.0492 
                                               0.385 
                                               1.12 
                                               0.966 
                                               1.31 
                                               0.138 
                                               0.975 
                                               lm.nb 
                                               ARMS-mediated activation, AXIN missense mutants destabilize the destruction complex, Beta-catenin phosphorylation cascade, CTLA4 inhibitory signaling, Cyclin AB1 associated events during G2M transition, Cyclin D associated events in G1, DAP12 signaling, DARPP-32 events, Disassembly of the destruction complex and recruitment of AXIN to the membrane, Downstream signal transduction, ERKs are inactivated, FCERI mediated MAPK activation, Frs2-mediated activation, Gastrin-CREB signalling pathway via PKC and MAPK, Interleukin receptor SHC signaling, IRS-related events triggered by IGF1R, MASTL Facilitates Mitotic Progression, Negative regulation of FGFR1 signaling, Negative regulation of FGFR2 signaling, Negative regulation of FGFR3 signaling, Negative regulation of FGFR4 signaling, Negative regulation of MAPK pathway, Nonsense Mediated Decay (NMD) enhanced by the Exon Junction Complex (EJC), RAF activation, RAFMAP kinase cascade, Resolution of Sister Chromatid Cohesion, RHO GTPases Activate Formins, S33 mutants of beta-catenin aren.SQT.t phosphorylated, S37 mutants of beta-catenin aren.SQT.t phosphorylated, S45 mutants of beta-catenin aren.SQT.t phosphorylated, Signaling by ERBB2, Signaling by PDGF, Signaling by SCF-KIT, Spry regulation of FGF signaling, T41 mutants of beta-catenin aren.SQT.t phosphorylated, Truncations of AMER1 destabilize the destruction complex, VEGFA-VEGFR2 Pathway, VEGFR2 mediated cell proliferation 
                                               SEBELIUS_2_C4636 
                                               PPP2CA 
                                               mRNA 
                                               0 
                                               
                                               
                                               MASTL Facilitates Mitotic Progression;T41 mutants of beta-catenin aren.SQT.t phosphorylated;ERKs are inactivated;Glycolysis;Negative regulation of MAPK pathway;Disassembly of the destruction complex and recruitment of AXIN to the membrane;S37 mutants of beta-catenin aren.SQT.t phosphorylated;Cyclin AB1 associated events during G2M transition;RAF activation;RHO GTPases Activate Formins;S33 mutants of beta-catenin aren.SQT.t phosphorylated;DARPP-32 events;Beta-catenin phosphorylation cascade;Initiation of Nuclear Envelope Reformation;Spry regulation of FGF signaling;Truncations of AMER1 destabilize the destruction complex;CTLA4 inhibitory signaling;AXIN missense mutants destabilize the destruction complex;S45 mutants of beta-catenin aren.SQT.t phosphorylated;Nonsense Mediated Decay (NMD) enhanced by the Exon Junction Complex (EJC);Cyclin D associated events in G1;Platelet sensitization by LDL;Resolution of Sister Chromatid Cohesion;Inhibition of replication initiation of damaged DNA by RB1E2F1;RAFMAP kinase cascade;Downstream signal transduction;VEGFR2 mediated cell proliferation;Signaling by SCF-KIT;Frs2-mediated activation;ARMS-mediated activation;Gastrin-CREB signalling pathway via PKC and MAPK;Interleukin receptor SHC signaling;FCERI mediated MAPK activation;DAP12 signaling;Signaling by PDGF;VEGFA-VEGFR2 Pathway;Signaling by ERBB2;IRS-related events triggered by IGF1R;Negative regulation of FGFR3 signaling;Negative regulation of FGFR4 signaling;Negative regulation of FGFR2 signaling;Negative regulation of FGFR1 signaling 
                                               hsa03015;hsa04071;hsa04114;hsa04151;hsa04152;hsa04261;hsa04350;hsa04390;hsa04530;hsa04728;hsa04730;hsa05142;hsa05160 
                                               PPP2CA 
                                               NA 
                                               azure4 
                                               1 
                                               1.51351351351351 
                                             
                                             
                                               PPP2R1A-mRNA 
                                               0.0488 
                                               0.108 
                                               -0.164 
                                               0.261 
                                               1.03 
                                               0.893 
                                               1.2 
                                               0.656 
                                               1 
                                               lm.nb 
                                               ARMS-mediated activation, AXIN missense mutants destabilize the destruction complex, Beta-catenin phosphorylation cascade, CTLA4 inhibitory signaling, Cyclin AB1 associated events during G2M transition, Cyclin D associated events in G1, DAP12 signaling, DARPP-32 events, Disassembly of the destruction complex and recruitment of AXIN to the membrane, Downstream signal transduction, FCERI mediated MAPK activation, Frs2-mediated activation, Gastrin-CREB signalling pathway via PKC and MAPK, Interleukin receptor SHC signaling, IRS-related events triggered by IGF1R, MASTL Facilitates Mitotic Progression, Negative regulation of FGFR1 signaling, Negative regulation of FGFR2 signaling, Negative regulation of FGFR3 signaling, Negative regulation of FGFR4 signaling, Nonsense Mediated Decay (NMD) enhanced by the Exon Junction Complex (EJC), RAF activation, RAFMAP kinase cascade, Resolution of Sister Chromatid Cohesion, RHO GTPases Activate Formins, S33 mutants of beta-catenin aren.SQT.t phosphorylated, S37 mutants of beta-catenin aren.SQT.t phosphorylated, S45 mutants of beta-catenin aren.SQT.t phosphorylated, Signaling by ERBB2, Signaling by PDGF, Signaling by SCF-KIT, Spry regulation of FGF signaling, Truncations of AMER1 destabilize the destruction complex, VEGFA-VEGFR2 Pathway, VEGFR2 mediated cell proliferation 
                                               SEBELIUS_2_C4636 
                                               PPP2R1A 
                                               mRNA 
                                               0 
                                               
                                               
                                               MASTL Facilitates Mitotic Progression;Regulation of PLK1 Activity at G2M Transition;Disassembly of the destruction complex and recruitment of AXIN to the membrane;S37 mutants of beta-catenin aren.SQT.t phosphorylated;Cyclin AB1 associated events during G2M transition;RAF activation;RHO GTPases Activate Formins;S33 mutants of beta-catenin aren.SQT.t phosphorylated;Loss of Nlp from mitotic centrosomes;DARPP-32 events;Beta-catenin phosphorylation cascade;Initiation of Nuclear Envelope Reformation;Recruitment of mitotic centrosome proteins and complexes;Spry regulation of FGF signaling;Truncations of AMER1 destabilize the destruction complex;CTLA4 inhibitory signaling;AXIN missense mutants destabilize the destruction complex;S45 mutants of beta-catenin aren.SQT.t phosphorylated;Anchoring of the basal body to the plasma membrane;Nonsense Mediated Decay (NMD) enhanced by the Exon Junction Complex (EJC);Cyclin D associated events in G1;Platelet sensitization by LDL;Resolution of Sister Chromatid Cohesion;Inhibition of replication initiation of damaged DNA by RB1E2F1;RAFMAP kinase cascade;Downstream signal transduction;VEGFR2 mediated cell proliferation;Signaling by SCF-KIT;Frs2-mediated activation;ARMS-mediated activation;Gastrin-CREB signalling pathway via PKC and MAPK;Interleukin receptor SHC signaling;FCERI mediated MAPK activation;DAP12 signaling;Signaling by PDGF;VEGFA-VEGFR2 Pathway;Signaling by ERBB2;IRS-related events triggered by IGF1R;Negative regulation of FGFR3 signaling;Negative regulation of FGFR4 signaling;Negative regulation of FGFR2 signaling;Negative regulation of FGFR1 signaling 
                                               hsa03015;hsa04071;hsa04114;hsa04151;hsa04152;hsa04261;hsa04350;hsa04390;hsa04530;hsa04728;hsa04730;hsa05142;hsa05160 
                                               PPP2R1A 
                                               NA 
                                               azure4 
                                               1 
                                               0.451851851851852 
                                             
                                             
                                               GSK3B-mRNA 
                                               0.00483 
                                               0.0978 
                                               -0.187 
                                               0.197 
                                               1 
                                               0.878 
                                               1.15 
                                               0.961 
                                               1 
                                               lm.nb 
                                               AXIN missense mutants destabilize the destruction complex, DAP12 signaling, Disassembly of the destruction complex and recruitment of AXIN to the membrane, Downstream signal transduction, GAB1 signalosome, Misspliced GSK3beta mutants stabilize beta-catenin, PI3KAKT activation, PIP3 activates AKT signaling, Regulation of HSF1-mediated heat shock response, Role of LAT2NTALLAB on calcium mobilization, S33 mutants of beta-catenin aren.SQT.t phosphorylated, S37 mutants of beta-catenin aren.SQT.t phosphorylated, S45 mutants of beta-catenin aren.SQT.t phosphorylated, Signaling by ERBB2, Signaling by PDGF, Signaling by SCF-KIT, T41 mutants of beta-catenin aren.SQT.t phosphorylated, Truncations of AMER1 destabilize the destruction complex 
                                               SEBELIUS_2_C4636 
                                               GSK3B 
                                               mRNA 
                                               0 
                                               
                                               
                                               Misspliced GSK3beta mutants stabilize beta-catenin;Constitutive Signaling by AKT1 E17K in Cancer;APC truncation mutants have impaired AXIN binding;T41 mutants of beta-catenin aren.SQT.t phosphorylated;AXIN missense mutants destabilize the destruction complex;S45 mutants of beta-catenin aren.SQT.t phosphorylated;Regulation of HSF1-mediated heat shock response;AKT phosphorylates targets in the cytosol;Disassembly of the destruction complex and recruitment of AXIN to the membrane;S37 mutants of beta-catenin aren.SQT.t phosphorylated;S33 mutants of beta-catenin aren.SQT.t phosphorylated;Truncations of AMER1 destabilize the destruction complex;Degradation of GLI2 by the proteasome;PIP3 activates AKT signaling;Downstream signal transduction;GAB1 signalosome;Signaling by SCF-KIT;PI3KAKT activation;Role of LAT2NTALLAB on calcium mobilization;DAP12 signaling;Signaling by PDGF;Signaling by ERBB2 
                                               hsa04012;hsa04062;hsa04110;hsa04151;hsa04310;hsa04360;hsa04390;hsa04510;hsa04550;hsa04660;hsa04662;hsa04722;hsa04728;hsa04910;hsa04916;hsa04917;hsa04919;hsa04931;hsa04932;hsa05010;hsa05160;hsa05162;hsa05164;hsa05166;hsa05169;hsa05200;hsa05210;hsa05213;hsa05215;hsa05217 
                                               GSK3B 
                                               NA 
                                               azure4 
                                               1 
                                               0.0493865030674847 
                                             
                                           
                                           DE Results - AXIN missense mutants destabilize the destruction complex genes - Line: Caco2 vs.Fibro  Table displaying each sample's global significance scores and directed global significance scores as defined in the heatmaps above. The global significance score is calculated as the square root of the mean squared t-statistic for the genes in a gene set, with t-statistics coming from the linear regression underlying our differential expression analysis. The directed global significance score is calculated as the square root of the mean signed squared t-statistic for the genes in a gene set, with t-statistics coming from the linear regression underlying our differential expression analysis.   
                                      
                                       
                                  
                                
                                
                                  
                                       
                                      
                                           Volcano Plot: Line: HEPG2 vs.Fibro   More Plot Information                Volcano Plot: Line: HEPG2 vs.Fibro  Volcano plot displaying each gene's -log10(p-value) and log2 fold change for the selected covariate. Highly statistically significant genes fall at the top of the plot, and highly differentially expressed genes fall to either side. Genes within the selected gene set are highlighted in orange. Horizontal lines indicate various False Discovery Rate (FDR) thresholds.   
                                           DE Results - AXIN missense mutants destabilize the destruction complex genes - Line: HEPG2 vs.Fibro   More Plot Information      Download CSV Data     
                                           
                                             
                                             
                                           
                                           
                                             
                                               Probe Label 
                                               Log2 fold change 
                                               std error (log2) 
                                               Lower confidence limit (log2) 
                                               Upper confidence limit (log2) 
                                               Linear fold change 
                                               Lower confidence limit (linear) 
                                               Upper confidence limit (linear) 
                                               P-value 
                                               BY.p.value 
                                               method 
                                               Gene.sets 
                                               Codeset.Name 
                                               Probe.Label 
                                               Analyte.Type 
                                               Is.Control 
                                               Control.Type 
                                               Related.Probes 
                                               Probe.Annotation 
                                               KEGG.Pathways 
                                               Official.Gene.Name 
                                               Control.Conc 
                                               volcanocol 
                                               volcanopch 
                                               tstats.all 
                                             
                                           
                                           
                                             
                                               PPP2R1B-mRNA 
                                               1.04 
                                               0.114 
                                               0.818 
                                               1.26 
                                               2.06 
                                               1.76 
                                               2.4 
                                               4.71e-11 
                                               9.7e-10 
                                               lm.nb 
                                               ARMS-mediated activation, AXIN missense mutants destabilize the destruction complex, Beta-catenin phosphorylation cascade, CTLA4 inhibitory signaling, Cyclin AB1 associated events during G2M transition, Cyclin D associated events in G1, DAP12 signaling, DARPP-32 events, Disassembly of the destruction complex and recruitment of AXIN to the membrane, Downstream signal transduction, ERKs are inactivated, FCERI mediated MAPK activation, Frs2-mediated activation, Gastrin-CREB signalling pathway via PKC and MAPK, Interleukin receptor SHC signaling, IRS-related events triggered by IGF1R, MASTL Facilitates Mitotic Progression, Misspliced GSK3beta mutants stabilize beta-catenin, Negative regulation of MAPK pathway, RAF activation, RAFMAP kinase cascade, Resolution of Sister Chromatid Cohesion, RHO GTPases Activate Formins, S33 mutants of beta-catenin aren.SQT.t phosphorylated, S37 mutants of beta-catenin aren.SQT.t phosphorylated, S45 mutants of beta-catenin aren.SQT.t phosphorylated, Signaling by ERBB2, Signaling by PDGF, Signaling by SCF-KIT, T41 mutants of beta-catenin aren.SQT.t phosphorylated, Truncations of AMER1 destabilize the destruction complex, VEGFA-VEGFR2 Pathway, VEGFR2 mediated cell proliferation 
                                               SEBELIUS_2_C4636 
                                               PPP2R1B 
                                               mRNA 
                                               0 
                                               
                                               
                                               MASTL Facilitates Mitotic Progression;Misspliced GSK3beta mutants stabilize beta-catenin;T41 mutants of beta-catenin aren.SQT.t phosphorylated;ERKs are inactivated;Glycolysis;Negative regulation of MAPK pathway;Disassembly of the destruction complex and recruitment of AXIN to the membrane;S37 mutants of beta-catenin aren.SQT.t phosphorylated;Cyclin AB1 associated events during G2M transition;RAF activation;RHO GTPases Activate Formins;S33 mutants of beta-catenin aren.SQT.t phosphorylated;DARPP-32 events;Beta-catenin phosphorylation cascade;Truncations of AMER1 destabilize the destruction complex;CTLA4 inhibitory signaling;AXIN missense mutants destabilize the destruction complex;S45 mutants of beta-catenin aren.SQT.t phosphorylated;Cyclin D associated events in G1;Platelet sensitization by LDL;Resolution of Sister Chromatid Cohesion;Inhibition of replication initiation of damaged DNA by RB1E2F1;RAFMAP kinase cascade;Downstream signal transduction;VEGFR2 mediated cell proliferation;Signaling by SCF-KIT;Frs2-mediated activation;ARMS-mediated activation;Gastrin-CREB signalling pathway via PKC and MAPK;Interleukin receptor SHC signaling;FCERI mediated MAPK activation;DAP12 signaling;Signaling by PDGF;VEGFA-VEGFR2 Pathway;Signaling by ERBB2;IRS-related events triggered by IGF1R 
                                               hsa03015;hsa04071;hsa04114;hsa04151;hsa04152;hsa04261;hsa04350;hsa04390;hsa04530;hsa04728;hsa04730;hsa05142;hsa05160 
                                               PPP2R1B 
                                               NA 
                                               slateblue1 
                                               16 
                                               9.12280701754386 
                                             
                                             
                                               AXIN1-mRNA 
                                               1.78 
                                               0.31 
                                               1.17 
                                               2.39 
                                               3.43 
                                               2.25 
                                               5.23 
                                               1.43e-06 
                                               1.75e-05 
                                               lm.nb 
                                               AXIN missense mutants destabilize the destruction complex, Beta-catenin phosphorylation cascade, Disassembly of the destruction complex and recruitment of AXIN to the membrane, Misspliced GSK3beta mutants stabilize beta-catenin, S33 mutants of beta-catenin aren.SQT.t phosphorylated, S37 mutants of beta-catenin aren.SQT.t phosphorylated, S45 mutants of beta-catenin aren.SQT.t phosphorylated, T41 mutants of beta-catenin aren.SQT.t phosphorylated, Truncations of AMER1 destabilize the destruction complex 
                                               SEBELIUS_2_C4636 
                                               AXIN1 
                                               mRNA 
                                               0 
                                               
                                               
                                               APC truncation mutants have impaired AXIN binding;Misspliced GSK3beta mutants stabilize beta-catenin;T41 mutants of beta-catenin aren.SQT.t phosphorylated;AXIN missense mutants destabilize the destruction complex;S45 mutants of beta-catenin aren.SQT.t phosphorylated;Disassembly of the destruction complex and recruitment of AXIN to the membrane;S37 mutants of beta-catenin aren.SQT.t phosphorylated;S33 mutants of beta-catenin aren.SQT.t phosphorylated;Beta-catenin phosphorylation cascade;Truncations of AMER1 destabilize the destruction complex 
                                               hsa04310;hsa04390;hsa04550;hsa05200;hsa05210;hsa05213;hsa05217 
                                               AXIN1 
                                               NA 
                                               slateblue1 
                                               16 
                                               5.74193548387097 
                                             
                                             
                                               PPP2CB-mRNA 
                                               -0.609 
                                               0.109 
                                               -0.822 
                                               -0.397 
                                               0.655 
                                               0.565 
                                               0.76 
                                               2.12e-06 
                                               2.55e-05 
                                               lm.nb 
                                               ARMS-mediated activation, AXIN missense mutants destabilize the destruction complex, Beta-catenin phosphorylation cascade, CTLA4 inhibitory signaling, Cyclin AB1 associated events during G2M transition, Cyclin D associated events in G1, DAP12 signaling, DARPP-32 events, Disassembly of the destruction complex and recruitment of AXIN to the membrane, Downstream signal transduction, ERKs are inactivated, FCERI mediated MAPK activation, Frs2-mediated activation, Gastrin-CREB signalling pathway via PKC and MAPK, Interleukin receptor SHC signaling, IRS-related events triggered by IGF1R, MASTL Facilitates Mitotic Progression, Misspliced GSK3beta mutants stabilize beta-catenin, Negative regulation of FGFR1 signaling, Negative regulation of FGFR2 signaling, Negative regulation of FGFR3 signaling, Negative regulation of FGFR4 signaling, Negative regulation of MAPK pathway, RAF activation, RAFMAP kinase cascade, Resolution of Sister Chromatid Cohesion, RHO GTPases Activate Formins, S33 mutants of beta-catenin aren.SQT.t phosphorylated, S37 mutants of beta-catenin aren.SQT.t phosphorylated, S45 mutants of beta-catenin aren.SQT.t phosphorylated, Signaling by ERBB2, Signaling by PDGF, Signaling by SCF-KIT, Spry regulation of FGF signaling, T41 mutants of beta-catenin aren.SQT.t phosphorylated, Truncations of AMER1 destabilize the destruction complex, VEGFA-VEGFR2 Pathway, VEGFR2 mediated cell proliferation 
                                               SEBELIUS_2_C4636 
                                               PPP2CB 
                                               mRNA 
                                               0 
                                               
                                               
                                               MASTL Facilitates Mitotic Progression;Misspliced GSK3beta mutants stabilize beta-catenin;T41 mutants of beta-catenin aren.SQT.t phosphorylated;ERKs are inactivated;Glycolysis;Negative regulation of MAPK pathway;Disassembly of the destruction complex and recruitment of AXIN to the membrane;S37 mutants of beta-catenin aren.SQT.t phosphorylated;Cyclin AB1 associated events during G2M transition;RAF activation;RHO GTPases Activate Formins;S33 mutants of beta-catenin aren.SQT.t phosphorylated;DARPP-32 events;Beta-catenin phosphorylation cascade;Spry regulation of FGF signaling;Truncations of AMER1 destabilize the destruction complex;CTLA4 inhibitory signaling;AXIN missense mutants destabilize the destruction complex;S45 mutants of beta-catenin aren.SQT.t phosphorylated;Cyclin D associated events in G1;Platelet sensitization by LDL;Resolution of Sister Chromatid Cohesion;Inhibition of replication initiation of damaged DNA by RB1E2F1;RAFMAP kinase cascade;Downstream signal transduction;VEGFR2 mediated cell proliferation;Signaling by SCF-KIT;Frs2-mediated activation;ARMS-mediated activation;Gastrin-CREB signalling pathway via PKC and MAPK;Interleukin receptor SHC signaling;FCERI mediated MAPK activation;DAP12 signaling;Signaling by PDGF;VEGFA-VEGFR2 Pathway;Signaling by ERBB2;IRS-related events triggered by IGF1R;Negative regulation of FGFR3 signaling;Negative regulation of FGFR4 signaling;Negative regulation of FGFR2 signaling;Negative regulation of FGFR1 signaling 
                                               hsa03015;hsa04071;hsa04114;hsa04151;hsa04152;hsa04261;hsa04350;hsa04390;hsa04530;hsa04728;hsa04730;hsa05142;hsa05160 
                                               PPP2CB 
                                               NA 
                                               slateblue1 
                                               16 
                                               -5.58715596330275 
                                             
                                             
                                               PPP2CA-mRNA 
                                               0.388 
                                               0.122 
                                               0.15 
                                               0.627 
                                               1.31 
                                               1.11 
                                               1.54 
                                               0.00287 
                                               0.0255 
                                               lm.nb 
                                               ARMS-mediated activation, AXIN missense mutants destabilize the destruction complex, Beta-catenin phosphorylation cascade, CTLA4 inhibitory signaling, Cyclin AB1 associated events during G2M transition, Cyclin D associated events in G1, DAP12 signaling, DARPP-32 events, Disassembly of the destruction complex and recruitment of AXIN to the membrane, Downstream signal transduction, ERKs are inactivated, FCERI mediated MAPK activation, Frs2-mediated activation, Gastrin-CREB signalling pathway via PKC and MAPK, Interleukin receptor SHC signaling, IRS-related events triggered by IGF1R, MASTL Facilitates Mitotic Progression, Negative regulation of FGFR1 signaling, Negative regulation of FGFR2 signaling, Negative regulation of FGFR3 signaling, Negative regulation of FGFR4 signaling, Negative regulation of MAPK pathway, Nonsense Mediated Decay (NMD) enhanced by the Exon Junction Complex (EJC), RAF activation, RAFMAP kinase cascade, Resolution of Sister Chromatid Cohesion, RHO GTPases Activate Formins, S33 mutants of beta-catenin aren.SQT.t phosphorylated, S37 mutants of beta-catenin aren.SQT.t phosphorylated, S45 mutants of beta-catenin aren.SQT.t phosphorylated, Signaling by ERBB2, Signaling by PDGF, Signaling by SCF-KIT, Spry regulation of FGF signaling, T41 mutants of beta-catenin aren.SQT.t phosphorylated, Truncations of AMER1 destabilize the destruction complex, VEGFA-VEGFR2 Pathway, VEGFR2 mediated cell proliferation 
                                               SEBELIUS_2_C4636 
                                               PPP2CA 
                                               mRNA 
                                               0 
                                               
                                               
                                               MASTL Facilitates Mitotic Progression;T41 mutants of beta-catenin aren.SQT.t phosphorylated;ERKs are inactivated;Glycolysis;Negative regulation of MAPK pathway;Disassembly of the destruction complex and recruitment of AXIN to the membrane;S37 mutants of beta-catenin aren.SQT.t phosphorylated;Cyclin AB1 associated events during G2M transition;RAF activation;RHO GTPases Activate Formins;S33 mutants of beta-catenin aren.SQT.t phosphorylated;DARPP-32 events;Beta-catenin phosphorylation cascade;Initiation of Nuclear Envelope Reformation;Spry regulation of FGF signaling;Truncations of AMER1 destabilize the destruction complex;CTLA4 inhibitory signaling;AXIN missense mutants destabilize the destruction complex;S45 mutants of beta-catenin aren.SQT.t phosphorylated;Nonsense Mediated Decay (NMD) enhanced by the Exon Junction Complex (EJC);Cyclin D associated events in G1;Platelet sensitization by LDL;Resolution of Sister Chromatid Cohesion;Inhibition of replication initiation of damaged DNA by RB1E2F1;RAFMAP kinase cascade;Downstream signal transduction;VEGFR2 mediated cell proliferation;Signaling by SCF-KIT;Frs2-mediated activation;ARMS-mediated activation;Gastrin-CREB signalling pathway via PKC and MAPK;Interleukin receptor SHC signaling;FCERI mediated MAPK activation;DAP12 signaling;Signaling by PDGF;VEGFA-VEGFR2 Pathway;Signaling by ERBB2;IRS-related events triggered by IGF1R;Negative regulation of FGFR3 signaling;Negative regulation of FGFR4 signaling;Negative regulation of FGFR2 signaling;Negative regulation of FGFR1 signaling 
                                               hsa03015;hsa04071;hsa04114;hsa04151;hsa04152;hsa04261;hsa04350;hsa04390;hsa04530;hsa04728;hsa04730;hsa05142;hsa05160 
                                               PPP2CA 
                                               NA 
                                               slateblue2 
                                               16 
                                               3.18032786885246 
                                             
                                             
                                               PPP2R1A-mRNA 
                                               -0.264 
                                               0.119 
                                               -0.498 
                                               -0.031 
                                               0.833 
                                               0.708 
                                               0.979 
                                               0.0326 
                                               0.253 
                                               lm.nb 
                                               ARMS-mediated activation, AXIN missense mutants destabilize the destruction complex, Beta-catenin phosphorylation cascade, CTLA4 inhibitory signaling, Cyclin AB1 associated events during G2M transition, Cyclin D associated events in G1, DAP12 signaling, DARPP-32 events, Disassembly of the destruction complex and recruitment of AXIN to the membrane, Downstream signal transduction, FCERI mediated MAPK activation, Frs2-mediated activation, Gastrin-CREB signalling pathway via PKC and MAPK, Interleukin receptor SHC signaling, IRS-related events triggered by IGF1R, MASTL Facilitates Mitotic Progression, Negative regulation of FGFR1 signaling, Negative regulation of FGFR2 signaling, Negative regulation of FGFR3 signaling, Negative regulation of FGFR4 signaling, Nonsense Mediated Decay (NMD) enhanced by the Exon Junction Complex (EJC), RAF activation, RAFMAP kinase cascade, Resolution of Sister Chromatid Cohesion, RHO GTPases Activate Formins, S33 mutants of beta-catenin aren.SQT.t phosphorylated, S37 mutants of beta-catenin aren.SQT.t phosphorylated, S45 mutants of beta-catenin aren.SQT.t phosphorylated, Signaling by ERBB2, Signaling by PDGF, Signaling by SCF-KIT, Spry regulation of FGF signaling, Truncations of AMER1 destabilize the destruction complex, VEGFA-VEGFR2 Pathway, VEGFR2 mediated cell proliferation 
                                               SEBELIUS_2_C4636 
                                               PPP2R1A 
                                               mRNA 
                                               0 
                                               
                                               
                                               MASTL Facilitates Mitotic Progression;Regulation of PLK1 Activity at G2M Transition;Disassembly of the destruction complex and recruitment of AXIN to the membrane;S37 mutants of beta-catenin aren.SQT.t phosphorylated;Cyclin AB1 associated events during G2M transition;RAF activation;RHO GTPases Activate Formins;S33 mutants of beta-catenin aren.SQT.t phosphorylated;Loss of Nlp from mitotic centrosomes;DARPP-32 events;Beta-catenin phosphorylation cascade;Initiation of Nuclear Envelope Reformation;Recruitment of mitotic centrosome proteins and complexes;Spry regulation of FGF signaling;Truncations of AMER1 destabilize the destruction complex;CTLA4 inhibitory signaling;AXIN missense mutants destabilize the destruction complex;S45 mutants of beta-catenin aren.SQT.t phosphorylated;Anchoring of the basal body to the plasma membrane;Nonsense Mediated Decay (NMD) enhanced by the Exon Junction Complex (EJC);Cyclin D associated events in G1;Platelet sensitization by LDL;Resolution of Sister Chromatid Cohesion;Inhibition of replication initiation of damaged DNA by RB1E2F1;RAFMAP kinase cascade;Downstream signal transduction;VEGFR2 mediated cell proliferation;Signaling by SCF-KIT;Frs2-mediated activation;ARMS-mediated activation;Gastrin-CREB signalling pathway via PKC and MAPK;Interleukin receptor SHC signaling;FCERI mediated MAPK activation;DAP12 signaling;Signaling by PDGF;VEGFA-VEGFR2 Pathway;Signaling by ERBB2;IRS-related events triggered by IGF1R;Negative regulation of FGFR3 signaling;Negative regulation of FGFR4 signaling;Negative regulation of FGFR2 signaling;Negative regulation of FGFR1 signaling 
                                               hsa03015;hsa04071;hsa04114;hsa04151;hsa04152;hsa04261;hsa04350;hsa04390;hsa04530;hsa04728;hsa04730;hsa05142;hsa05160 
                                               PPP2R1A 
                                               NA 
                                               slateblue4 
                                               16 
                                               -2.21848739495798 
                                             
                                             
                                               CSNK1A1-mRNA 
                                               -0.15 
                                               0.0908 
                                               -0.328 
                                               0.0285 
                                               0.902 
                                               0.797 
                                               1.02 
                                               0.108 
                                               0.792 
                                               loglinear 
                                               AXIN missense mutants destabilize the destruction complex, Beta-catenin phosphorylation cascade, Disassembly of the destruction complex and recruitment of AXIN to the membrane, Misspliced GSK3beta mutants stabilize beta-catenin, S33 mutants of beta-catenin aren.SQT.t phosphorylated, S37 mutants of beta-catenin aren.SQT.t phosphorylated, S45 mutants of beta-catenin aren.SQT.t phosphorylated, T41 mutants of beta-catenin aren.SQT.t phosphorylated 
                                               SEBELIUS_2_C4636 
                                               CSNK1A1 
                                               mRNA 
                                               0 
                                               
                                               
                                               Misspliced GSK3beta mutants stabilize beta-catenin;APC truncation mutants have impaired AXIN binding;T41 mutants of beta-catenin aren.SQT.t phosphorylated;AXIN missense mutants destabilize the destruction complex;S45 mutants of beta-catenin aren.SQT.t phosphorylated;Disassembly of the destruction complex and recruitment of AXIN to the membrane;S37 mutants of beta-catenin aren.SQT.t phosphorylated;S33 mutants of beta-catenin aren.SQT.t phosphorylated;Beta-catenin phosphorylation cascade;Degradation of GLI2 by the proteasome 
                                               hsa04310 
                                               CSNK1A1 
                                               NA 
                                               azure4 
                                               1 
                                               -1.65198237885463 
                                             
                                             
                                               GSK3B-mRNA 
                                               0.143 
                                               0.107 
                                               -0.0671 
                                               0.353 
                                               1.1 
                                               0.955 
                                               1.28 
                                               0.19 
                                               1 
                                               lm.nb 
                                               AXIN missense mutants destabilize the destruction complex, DAP12 signaling, Disassembly of the destruction complex and recruitment of AXIN to the membrane, Downstream signal transduction, GAB1 signalosome, Misspliced GSK3beta mutants stabilize beta-catenin, PI3KAKT activation, PIP3 activates AKT signaling, Regulation of HSF1-mediated heat shock response, Role of LAT2NTALLAB on calcium mobilization, S33 mutants of beta-catenin aren.SQT.t phosphorylated, S37 mutants of beta-catenin aren.SQT.t phosphorylated, S45 mutants of beta-catenin aren.SQT.t phosphorylated, Signaling by ERBB2, Signaling by PDGF, Signaling by SCF-KIT, T41 mutants of beta-catenin aren.SQT.t phosphorylated, Truncations of AMER1 destabilize the destruction complex 
                                               SEBELIUS_2_C4636 
                                               GSK3B 
                                               mRNA 
                                               0 
                                               
                                               
                                               Misspliced GSK3beta mutants stabilize beta-catenin;Constitutive Signaling by AKT1 E17K in Cancer;APC truncation mutants have impaired AXIN binding;T41 mutants of beta-catenin aren.SQT.t phosphorylated;AXIN missense mutants destabilize the destruction complex;S45 mutants of beta-catenin aren.SQT.t phosphorylated;Regulation of HSF1-mediated heat shock response;AKT phosphorylates targets in the cytosol;Disassembly of the destruction complex and recruitment of AXIN to the membrane;S37 mutants of beta-catenin aren.SQT.t phosphorylated;S33 mutants of beta-catenin aren.SQT.t phosphorylated;Truncations of AMER1 destabilize the destruction complex;Degradation of GLI2 by the proteasome;PIP3 activates AKT signaling;Downstream signal transduction;GAB1 signalosome;Signaling by SCF-KIT;PI3KAKT activation;Role of LAT2NTALLAB on calcium mobilization;DAP12 signaling;Signaling by PDGF;Signaling by ERBB2 
                                               hsa04012;hsa04062;hsa04110;hsa04151;hsa04310;hsa04360;hsa04390;hsa04510;hsa04550;hsa04660;hsa04662;hsa04722;hsa04728;hsa04910;hsa04916;hsa04917;hsa04919;hsa04931;hsa04932;hsa05010;hsa05160;hsa05162;hsa05164;hsa05166;hsa05169;hsa05200;hsa05210;hsa05213;hsa05215;hsa05217 
                                               GSK3B 
                                               NA 
                                               azure4 
                                               1 
                                               1.33644859813084 
                                             
                                             
                                               APC-mRNA 
                                               -0.0616 
                                               0.125 
                                               -0.306 
                                               0.183 
                                               0.958 
                                               0.809 
                                               1.14 
                                               0.624 
                                               1 
                                               lm.nb 
                                               Apoptotic cleavage of cellular proteins, AXIN missense mutants destabilize the destruction complex, Beta-catenin phosphorylation cascade, Deactivation of the beta-catenin transactivating complex, Disassembly of the destruction complex and recruitment of AXIN to the membrane, Misspliced GSK3beta mutants stabilize beta-catenin, S33 mutants of beta-catenin aren.SQT.t phosphorylated, S37 mutants of beta-catenin aren.SQT.t phosphorylated, S45 mutants of beta-catenin aren.SQT.t phosphorylated, T41 mutants of beta-catenin aren.SQT.t phosphorylated, Truncations of AMER1 destabilize the destruction complex 
                                               SEBELIUS_2_C4636 
                                               APC 
                                               mRNA 
                                               0 
                                               
                                               
                                               Apoptotic cleavage of cellular proteins;Misspliced GSK3beta mutants stabilize beta-catenin;APC truncation mutants have impaired AXIN binding;T41 mutants of beta-catenin aren.SQT.t phosphorylated;AXIN missense mutants destabilize the destruction complex;S45 mutants of beta-catenin aren.SQT.t phosphorylated;Deactivation of the beta-catenin transactivating complex;Disassembly of the destruction complex and recruitment of AXIN to the membrane;S37 mutants of beta-catenin aren.SQT.t phosphorylated;S33 mutants of beta-catenin aren.SQT.t phosphorylated;Beta-catenin phosphorylation cascade;Truncations of AMER1 destabilize the destruction complex 
                                               hsa04310;hsa04390;hsa04550;hsa04810;hsa05166;hsa05200;hsa05206;hsa05210;hsa05213;hsa05217 
                                               APC 
                                               NA 
                                               azure4 
                                               1 
                                               -0.4928 
                                             
                                           
                                           DE Results - AXIN missense mutants destabilize the destruction complex genes - Line: HEPG2 vs.Fibro  Table displaying each sample's global significance scores and directed global significance scores as defined in the heatmaps above. The global significance score is calculated as the square root of the mean squared t-statistic for the genes in a gene set, with t-statistics coming from the linear regression underlying our differential expression analysis. The directed global significance score is calculated as the square root of the mean signed squared t-statistic for the genes in a gene set, with t-statistics coming from the linear regression underlying our differential expression analysis.   
                                      
                                       
                                  
                                
                                
                                  
                                       
                                      
                                           Volcano Plot: Line: HT29 vs.Fibro   More Plot Information                Volcano Plot: Line: HT29 vs.Fibro  Volcano plot displaying each gene's -log10(p-value) and log2 fold change for the selected covariate. Highly statistically significant genes fall at the top of the plot, and highly differentially expressed genes fall to either side. Genes within the selected gene set are highlighted in orange. Horizontal lines indicate various False Discovery Rate (FDR) thresholds.   
                                           DE Results - AXIN missense mutants destabilize the destruction complex genes - Line: HT29 vs.Fibro   More Plot Information      Download CSV Data     
                                           
                                             
                                             
                                           
                                           
                                             
                                               Probe Label 
                                               Log2 fold change 
                                               std error (log2) 
                                               Lower confidence limit (log2) 
                                               Upper confidence limit (log2) 
                                               Linear fold change 
                                               Lower confidence limit (linear) 
                                               Upper confidence limit (linear) 
                                               P-value 
                                               BY.p.value 
                                               method 
                                               Gene.sets 
                                               Codeset.Name 
                                               Probe.Label 
                                               Analyte.Type 
                                               Is.Control 
                                               Control.Type 
                                               Related.Probes 
                                               Probe.Annotation 
                                               KEGG.Pathways 
                                               Official.Gene.Name 
                                               Control.Conc 
                                               volcanocol 
                                               volcanopch 
                                               tstats.all 
                                             
                                           
                                           
                                             
                                               PPP2CB-mRNA 
                                               -2.03 
                                               0.103 
                                               -2.23 
                                               -1.83 
                                               0.245 
                                               0.213 
                                               0.281 
                                               3.08e-21 
                                               1.56e-19 
                                               lm.nb 
                                               ARMS-mediated activation, AXIN missense mutants destabilize the destruction complex, Beta-catenin phosphorylation cascade, CTLA4 inhibitory signaling, Cyclin AB1 associated events during G2M transition, Cyclin D associated events in G1, DAP12 signaling, DARPP-32 events, Disassembly of the destruction complex and recruitment of AXIN to the membrane, Downstream signal transduction, ERKs are inactivated, FCERI mediated MAPK activation, Frs2-mediated activation, Gastrin-CREB signalling pathway via PKC and MAPK, Interleukin receptor SHC signaling, IRS-related events triggered by IGF1R, MASTL Facilitates Mitotic Progression, Misspliced GSK3beta mutants stabilize beta-catenin, Negative regulation of FGFR1 signaling, Negative regulation of FGFR2 signaling, Negative regulation of FGFR3 signaling, Negative regulation of FGFR4 signaling, Negative regulation of MAPK pathway, RAF activation, RAFMAP kinase cascade, Resolution of Sister Chromatid Cohesion, RHO GTPases Activate Formins, S33 mutants of beta-catenin aren.SQT.t phosphorylated, S37 mutants of beta-catenin aren.SQT.t phosphorylated, S45 mutants of beta-catenin aren.SQT.t phosphorylated, Signaling by ERBB2, Signaling by PDGF, Signaling by SCF-KIT, Spry regulation of FGF signaling, T41 mutants of beta-catenin aren.SQT.t phosphorylated, Truncations of AMER1 destabilize the destruction complex, VEGFA-VEGFR2 Pathway, VEGFR2 mediated cell proliferation 
                                               SEBELIUS_2_C4636 
                                               PPP2CB 
                                               mRNA 
                                               0 
                                               
                                               
                                               MASTL Facilitates Mitotic Progression;Misspliced GSK3beta mutants stabilize beta-catenin;T41 mutants of beta-catenin aren.SQT.t phosphorylated;ERKs are inactivated;Glycolysis;Negative regulation of MAPK pathway;Disassembly of the destruction complex and recruitment of AXIN to the membrane;S37 mutants of beta-catenin aren.SQT.t phosphorylated;Cyclin AB1 associated events during G2M transition;RAF activation;RHO GTPases Activate Formins;S33 mutants of beta-catenin aren.SQT.t phosphorylated;DARPP-32 events;Beta-catenin phosphorylation cascade;Spry regulation of FGF signaling;Truncations of AMER1 destabilize the destruction complex;CTLA4 inhibitory signaling;AXIN missense mutants destabilize the destruction complex;S45 mutants of beta-catenin aren.SQT.t phosphorylated;Cyclin D associated events in G1;Platelet sensitization by LDL;Resolution of Sister Chromatid Cohesion;Inhibition of replication initiation of damaged DNA by RB1E2F1;RAFMAP kinase cascade;Downstream signal transduction;VEGFR2 mediated cell proliferation;Signaling by SCF-KIT;Frs2-mediated activation;ARMS-mediated activation;Gastrin-CREB signalling pathway via PKC and MAPK;Interleukin receptor SHC signaling;FCERI mediated MAPK activation;DAP12 signaling;Signaling by PDGF;VEGFA-VEGFR2 Pathway;Signaling by ERBB2;IRS-related events triggered by IGF1R;Negative regulation of FGFR3 signaling;Negative regulation of FGFR4 signaling;Negative regulation of FGFR2 signaling;Negative regulation of FGFR1 signaling 
                                               hsa03015;hsa04071;hsa04114;hsa04151;hsa04152;hsa04261;hsa04350;hsa04390;hsa04530;hsa04728;hsa04730;hsa05142;hsa05160 
                                               PPP2CB 
                                               NA 
                                               slateblue1 
                                               16 
                                               -19.7087378640777 
                                             
                                             
                                               PPP2R1B-mRNA 
                                               0.978 
                                               0.107 
                                               0.769 
                                               1.19 
                                               1.97 
                                               1.7 
                                               2.28 
                                               5.09e-11 
                                               7.3e-10 
                                               lm.nb 
                                               ARMS-mediated activation, AXIN missense mutants destabilize the destruction complex, Beta-catenin phosphorylation cascade, CTLA4 inhibitory signaling, Cyclin AB1 associated events during G2M transition, Cyclin D associated events in G1, DAP12 signaling, DARPP-32 events, Disassembly of the destruction complex and recruitment of AXIN to the membrane, Downstream signal transduction, ERKs are inactivated, FCERI mediated MAPK activation, Frs2-mediated activation, Gastrin-CREB signalling pathway via PKC and MAPK, Interleukin receptor SHC signaling, IRS-related events triggered by IGF1R, MASTL Facilitates Mitotic Progression, Misspliced GSK3beta mutants stabilize beta-catenin, Negative regulation of MAPK pathway, RAF activation, RAFMAP kinase cascade, Resolution of Sister Chromatid Cohesion, RHO GTPases Activate Formins, S33 mutants of beta-catenin aren.SQT.t phosphorylated, S37 mutants of beta-catenin aren.SQT.t phosphorylated, S45 mutants of beta-catenin aren.SQT.t phosphorylated, Signaling by ERBB2, Signaling by PDGF, Signaling by SCF-KIT, T41 mutants of beta-catenin aren.SQT.t phosphorylated, Truncations of AMER1 destabilize the destruction complex, VEGFA-VEGFR2 Pathway, VEGFR2 mediated cell proliferation 
                                               SEBELIUS_2_C4636 
                                               PPP2R1B 
                                               mRNA 
                                               0 
                                               
                                               
                                               MASTL Facilitates Mitotic Progression;Misspliced GSK3beta mutants stabilize beta-catenin;T41 mutants of beta-catenin aren.SQT.t phosphorylated;ERKs are inactivated;Glycolysis;Negative regulation of MAPK pathway;Disassembly of the destruction complex and recruitment of AXIN to the membrane;S37 mutants of beta-catenin aren.SQT.t phosphorylated;Cyclin AB1 associated events during G2M transition;RAF activation;RHO GTPases Activate Formins;S33 mutants of beta-catenin aren.SQT.t phosphorylated;DARPP-32 events;Beta-catenin phosphorylation cascade;Truncations of AMER1 destabilize the destruction complex;CTLA4 inhibitory signaling;AXIN missense mutants destabilize the destruction complex;S45 mutants of beta-catenin aren.SQT.t phosphorylated;Cyclin D associated events in G1;Platelet sensitization by LDL;Resolution of Sister Chromatid Cohesion;Inhibition of replication initiation of damaged DNA by RB1E2F1;RAFMAP kinase cascade;Downstream signal transduction;VEGFR2 mediated cell proliferation;Signaling by SCF-KIT;Frs2-mediated activation;ARMS-mediated activation;Gastrin-CREB signalling pathway via PKC and MAPK;Interleukin receptor SHC signaling;FCERI mediated MAPK activation;DAP12 signaling;Signaling by PDGF;VEGFA-VEGFR2 Pathway;Signaling by ERBB2;IRS-related events triggered by IGF1R 
                                               hsa03015;hsa04071;hsa04114;hsa04151;hsa04152;hsa04261;hsa04350;hsa04390;hsa04530;hsa04728;hsa04730;hsa05142;hsa05160 
                                               PPP2R1B 
                                               NA 
                                               slateblue1 
                                               16 
                                               9.14018691588785 
                                             
                                             
                                               GSK3B-mRNA 
                                               0.74 
                                               0.101 
                                               0.543 
                                               0.938 
                                               1.67 
                                               1.46 
                                               1.92 
                                               9.76e-09 
                                               1.17e-07 
                                               lm.nb 
                                               AXIN missense mutants destabilize the destruction complex, DAP12 signaling, Disassembly of the destruction complex and recruitment of AXIN to the membrane, Downstream signal transduction, GAB1 signalosome, Misspliced GSK3beta mutants stabilize beta-catenin, PI3KAKT activation, PIP3 activates AKT signaling, Regulation of HSF1-mediated heat shock response, Role of LAT2NTALLAB on calcium mobilization, S33 mutants of beta-catenin aren.SQT.t phosphorylated, S37 mutants of beta-catenin aren.SQT.t phosphorylated, S45 mutants of beta-catenin aren.SQT.t phosphorylated, Signaling by ERBB2, Signaling by PDGF, Signaling by SCF-KIT, T41 mutants of beta-catenin aren.SQT.t phosphorylated, Truncations of AMER1 destabilize the destruction complex 
                                               SEBELIUS_2_C4636 
                                               GSK3B 
                                               mRNA 
                                               0 
                                               
                                               
                                               Misspliced GSK3beta mutants stabilize beta-catenin;Constitutive Signaling by AKT1 E17K in Cancer;APC truncation mutants have impaired AXIN binding;T41 mutants of beta-catenin aren.SQT.t phosphorylated;AXIN missense mutants destabilize the destruction complex;S45 mutants of beta-catenin aren.SQT.t phosphorylated;Regulation of HSF1-mediated heat shock response;AKT phosphorylates targets in the cytosol;Disassembly of the destruction complex and recruitment of AXIN to the membrane;S37 mutants of beta-catenin aren.SQT.t phosphorylated;S33 mutants of beta-catenin aren.SQT.t phosphorylated;Truncations of AMER1 destabilize the destruction complex;Degradation of GLI2 by the proteasome;PIP3 activates AKT signaling;Downstream signal transduction;GAB1 signalosome;Signaling by SCF-KIT;PI3KAKT activation;Role of LAT2NTALLAB on calcium mobilization;DAP12 signaling;Signaling by PDGF;Signaling by ERBB2 
                                               hsa04012;hsa04062;hsa04110;hsa04151;hsa04310;hsa04360;hsa04390;hsa04510;hsa04550;hsa04660;hsa04662;hsa04722;hsa04728;hsa04910;hsa04916;hsa04917;hsa04919;hsa04931;hsa04932;hsa05010;hsa05160;hsa05162;hsa05164;hsa05166;hsa05169;hsa05200;hsa05210;hsa05213;hsa05215;hsa05217 
                                               GSK3B 
                                               NA 
                                               slateblue1 
                                               16 
                                               7.32673267326733 
                                             
                                             
                                               AXIN1-mRNA 
                                               1.54 
                                               0.293 
                                               0.963 
                                               2.11 
                                               2.9 
                                               1.95 
                                               4.32 
                                               6.64e-06 
                                               6.63e-05 
                                               lm.nb 
                                               AXIN missense mutants destabilize the destruction complex, Beta-catenin phosphorylation cascade, Disassembly of the destruction complex and recruitment of AXIN to the membrane, Misspliced GSK3beta mutants stabilize beta-catenin, S33 mutants of beta-catenin aren.SQT.t phosphorylated, S37 mutants of beta-catenin aren.SQT.t phosphorylated, S45 mutants of beta-catenin aren.SQT.t phosphorylated, T41 mutants of beta-catenin aren.SQT.t phosphorylated, Truncations of AMER1 destabilize the destruction complex 
                                               SEBELIUS_2_C4636 
                                               AXIN1 
                                               mRNA 
                                               0 
                                               
                                               
                                               APC truncation mutants have impaired AXIN binding;Misspliced GSK3beta mutants stabilize beta-catenin;T41 mutants of beta-catenin aren.SQT.t phosphorylated;AXIN missense mutants destabilize the destruction complex;S45 mutants of beta-catenin aren.SQT.t phosphorylated;Disassembly of the destruction complex and recruitment of AXIN to the membrane;S37 mutants of beta-catenin aren.SQT.t phosphorylated;S33 mutants of beta-catenin aren.SQT.t phosphorylated;Beta-catenin phosphorylation cascade;Truncations of AMER1 destabilize the destruction complex 
                                               hsa04310;hsa04390;hsa04550;hsa05200;hsa05210;hsa05213;hsa05217 
                                               AXIN1 
                                               NA 
                                               slateblue1 
                                               16 
                                               5.25597269624573 
                                             
                                             
                                               PPP2CA-mRNA 
                                               0.495 
                                               0.115 
                                               0.271 
                                               0.719 
                                               1.41 
                                               1.21 
                                               1.65 
                                               0.000112 
                                               0.00104 
                                               lm.nb 
                                               ARMS-mediated activation, AXIN missense mutants destabilize the destruction complex, Beta-catenin phosphorylation cascade, CTLA4 inhibitory signaling, Cyclin AB1 associated events during G2M transition, Cyclin D associated events in G1, DAP12 signaling, DARPP-32 events, Disassembly of the destruction complex and recruitment of AXIN to the membrane, Downstream signal transduction, ERKs are inactivated, FCERI mediated MAPK activation, Frs2-mediated activation, Gastrin-CREB signalling pathway via PKC and MAPK, Interleukin receptor SHC signaling, IRS-related events triggered by IGF1R, MASTL Facilitates Mitotic Progression, Negative regulation of FGFR1 signaling, Negative regulation of FGFR2 signaling, Negative regulation of FGFR3 signaling, Negative regulation of FGFR4 signaling, Negative regulation of MAPK pathway, Nonsense Mediated Decay (NMD) enhanced by the Exon Junction Complex (EJC), RAF activation, RAFMAP kinase cascade, Resolution of Sister Chromatid Cohesion, RHO GTPases Activate Formins, S33 mutants of beta-catenin aren.SQT.t phosphorylated, S37 mutants of beta-catenin aren.SQT.t phosphorylated, S45 mutants of beta-catenin aren.SQT.t phosphorylated, Signaling by ERBB2, Signaling by PDGF, Signaling by SCF-KIT, Spry regulation of FGF signaling, T41 mutants of beta-catenin aren.SQT.t phosphorylated, Truncations of AMER1 destabilize the destruction complex, VEGFA-VEGFR2 Pathway, VEGFR2 mediated cell proliferation 
                                               SEBELIUS_2_C4636 
                                               PPP2CA 
                                               mRNA 
                                               0 
                                               
                                               
                                               MASTL Facilitates Mitotic Progression;T41 mutants of beta-catenin aren.SQT.t phosphorylated;ERKs are inactivated;Glycolysis;Negative regulation of MAPK pathway;Disassembly of the destruction complex and recruitment of AXIN to the membrane;S37 mutants of beta-catenin aren.SQT.t phosphorylated;Cyclin AB1 associated events during G2M transition;RAF activation;RHO GTPases Activate Formins;S33 mutants of beta-catenin aren.SQT.t phosphorylated;DARPP-32 events;Beta-catenin phosphorylation cascade;Initiation of Nuclear Envelope Reformation;Spry regulation of FGF signaling;Truncations of AMER1 destabilize the destruction complex;CTLA4 inhibitory signaling;AXIN missense mutants destabilize the destruction complex;S45 mutants of beta-catenin aren.SQT.t phosphorylated;Nonsense Mediated Decay (NMD) enhanced by the Exon Junction Complex (EJC);Cyclin D associated events in G1;Platelet sensitization by LDL;Resolution of Sister Chromatid Cohesion;Inhibition of replication initiation of damaged DNA by RB1E2F1;RAFMAP kinase cascade;Downstream signal transduction;VEGFR2 mediated cell proliferation;Signaling by SCF-KIT;Frs2-mediated activation;ARMS-mediated activation;Gastrin-CREB signalling pathway via PKC and MAPK;Interleukin receptor SHC signaling;FCERI mediated MAPK activation;DAP12 signaling;Signaling by PDGF;VEGFA-VEGFR2 Pathway;Signaling by ERBB2;IRS-related events triggered by IGF1R;Negative regulation of FGFR3 signaling;Negative regulation of FGFR4 signaling;Negative regulation of FGFR2 signaling;Negative regulation of FGFR1 signaling 
                                               hsa03015;hsa04071;hsa04114;hsa04151;hsa04152;hsa04261;hsa04350;hsa04390;hsa04530;hsa04728;hsa04730;hsa05142;hsa05160 
                                               PPP2CA 
                                               NA 
                                               slateblue1 
                                               16 
                                               4.30434782608696 
                                             
                                             
                                               PPP2R1A-mRNA 
                                               0.449 
                                               0.112 
                                               0.229 
                                               0.669 
                                               1.37 
                                               1.17 
                                               1.59 
                                               0.000287 
                                               0.00259 
                                               lm.nb 
                                               ARMS-mediated activation, AXIN missense mutants destabilize the destruction complex, Beta-catenin phosphorylation cascade, CTLA4 inhibitory signaling, Cyclin AB1 associated events during G2M transition, Cyclin D associated events in G1, DAP12 signaling, DARPP-32 events, Disassembly of the destruction complex and recruitment of AXIN to the membrane, Downstream signal transduction, FCERI mediated MAPK activation, Frs2-mediated activation, Gastrin-CREB signalling pathway via PKC and MAPK, Interleukin receptor SHC signaling, IRS-related events triggered by IGF1R, MASTL Facilitates Mitotic Progression, Negative regulation of FGFR1 signaling, Negative regulation of FGFR2 signaling, Negative regulation of FGFR3 signaling, Negative regulation of FGFR4 signaling, Nonsense Mediated Decay (NMD) enhanced by the Exon Junction Complex (EJC), RAF activation, RAFMAP kinase cascade, Resolution of Sister Chromatid Cohesion, RHO GTPases Activate Formins, S33 mutants of beta-catenin aren.SQT.t phosphorylated, S37 mutants of beta-catenin aren.SQT.t phosphorylated, S45 mutants of beta-catenin aren.SQT.t phosphorylated, Signaling by ERBB2, Signaling by PDGF, Signaling by SCF-KIT, Spry regulation of FGF signaling, Truncations of AMER1 destabilize the destruction complex, VEGFA-VEGFR2 Pathway, VEGFR2 mediated cell proliferation 
                                               SEBELIUS_2_C4636 
                                               PPP2R1A 
                                               mRNA 
                                               0 
                                               
                                               
                                               MASTL Facilitates Mitotic Progression;Regulation of PLK1 Activity at G2M Transition;Disassembly of the destruction complex and recruitment of AXIN to the membrane;S37 mutants of beta-catenin aren.SQT.t phosphorylated;Cyclin AB1 associated events during G2M transition;RAF activation;RHO GTPases Activate Formins;S33 mutants of beta-catenin aren.SQT.t phosphorylated;Loss of Nlp from mitotic centrosomes;DARPP-32 events;Beta-catenin phosphorylation cascade;Initiation of Nuclear Envelope Reformation;Recruitment of mitotic centrosome proteins and complexes;Spry regulation of FGF signaling;Truncations of AMER1 destabilize the destruction complex;CTLA4 inhibitory signaling;AXIN missense mutants destabilize the destruction complex;S45 mutants of beta-catenin aren.SQT.t phosphorylated;Anchoring of the basal body to the plasma membrane;Nonsense Mediated Decay (NMD) enhanced by the Exon Junction Complex (EJC);Cyclin D associated events in G1;Platelet sensitization by LDL;Resolution of Sister Chromatid Cohesion;Inhibition of replication initiation of damaged DNA by RB1E2F1;RAFMAP kinase cascade;Downstream signal transduction;VEGFR2 mediated cell proliferation;Signaling by SCF-KIT;Frs2-mediated activation;ARMS-mediated activation;Gastrin-CREB signalling pathway via PKC and MAPK;Interleukin receptor SHC signaling;FCERI mediated MAPK activation;DAP12 signaling;Signaling by PDGF;VEGFA-VEGFR2 Pathway;Signaling by ERBB2;IRS-related events triggered by IGF1R;Negative regulation of FGFR3 signaling;Negative regulation of FGFR4 signaling;Negative regulation of FGFR2 signaling;Negative regulation of FGFR1 signaling 
                                               hsa03015;hsa04071;hsa04114;hsa04151;hsa04152;hsa04261;hsa04350;hsa04390;hsa04530;hsa04728;hsa04730;hsa05142;hsa05160 
                                               PPP2R1A 
                                               NA 
                                               slateblue1 
                                               16 
                                               4.00892857142857 
                                             
                                             
                                               APC-mRNA 
                                               0.329 
                                               0.117 
                                               0.1 
                                               0.558 
                                               1.26 
                                               1.07 
                                               1.47 
                                               0.00766 
                                               0.0598 
                                               lm.nb 
                                               Apoptotic cleavage of cellular proteins, AXIN missense mutants destabilize the destruction complex, Beta-catenin phosphorylation cascade, Deactivation of the beta-catenin transactivating complex, Disassembly of the destruction complex and recruitment of AXIN to the membrane, Misspliced GSK3beta mutants stabilize beta-catenin, S33 mutants of beta-catenin aren.SQT.t phosphorylated, S37 mutants of beta-catenin aren.SQT.t phosphorylated, S45 mutants of beta-catenin aren.SQT.t phosphorylated, T41 mutants of beta-catenin aren.SQT.t phosphorylated, Truncations of AMER1 destabilize the destruction complex 
                                               SEBELIUS_2_C4636 
                                               APC 
                                               mRNA 
                                               0 
                                               
                                               
                                               Apoptotic cleavage of cellular proteins;Misspliced GSK3beta mutants stabilize beta-catenin;APC truncation mutants have impaired AXIN binding;T41 mutants of beta-catenin aren.SQT.t phosphorylated;AXIN missense mutants destabilize the destruction complex;S45 mutants of beta-catenin aren.SQT.t phosphorylated;Deactivation of the beta-catenin transactivating complex;Disassembly of the destruction complex and recruitment of AXIN to the membrane;S37 mutants of beta-catenin aren.SQT.t phosphorylated;S33 mutants of beta-catenin aren.SQT.t phosphorylated;Beta-catenin phosphorylation cascade;Truncations of AMER1 destabilize the destruction complex 
                                               hsa04310;hsa04390;hsa04550;hsa04810;hsa05166;hsa05200;hsa05206;hsa05210;hsa05213;hsa05217 
                                               APC 
                                               NA 
                                               slateblue3 
                                               16 
                                               2.81196581196581 
                                             
                                             
                                               CSNK1A1-mRNA 
                                               0.171 
                                               0.0856 
                                               0.00324 
                                               0.339 
                                               1.13 
                                               1 
                                               1.26 
                                               0.0531 
                                               0.389 
                                               loglinear 
                                               AXIN missense mutants destabilize the destruction complex, Beta-catenin phosphorylation cascade, Disassembly of the destruction complex and recruitment of AXIN to the membrane, Misspliced GSK3beta mutants stabilize beta-catenin, S33 mutants of beta-catenin aren.SQT.t phosphorylated, S37 mutants of beta-catenin aren.SQT.t phosphorylated, S45 mutants of beta-catenin aren.SQT.t phosphorylated, T41 mutants of beta-catenin aren.SQT.t phosphorylated 
                                               SEBELIUS_2_C4636 
                                               CSNK1A1 
                                               mRNA 
                                               0 
                                               
                                               
                                               Misspliced GSK3beta mutants stabilize beta-catenin;APC truncation mutants have impaired AXIN binding;T41 mutants of beta-catenin aren.SQT.t phosphorylated;AXIN missense mutants destabilize the destruction complex;S45 mutants of beta-catenin aren.SQT.t phosphorylated;Disassembly of the destruction complex and recruitment of AXIN to the membrane;S37 mutants of beta-catenin aren.SQT.t phosphorylated;S33 mutants of beta-catenin aren.SQT.t phosphorylated;Beta-catenin phosphorylation cascade;Degradation of GLI2 by the proteasome 
                                               hsa04310 
                                               CSNK1A1 
                                               NA 
                                               slateblue4 
                                               16 
                                               1.99766355140187 
                                             
                                           
                                           DE Results - AXIN missense mutants destabilize the destruction complex genes - Line: HT29 vs.Fibro  Table displaying each sample's global significance scores and directed global significance scores as defined in the heatmaps above. The global significance score is calculated as the square root of the mean squared t-statistic for the genes in a gene set, with t-statistics coming from the linear regression underlying our differential expression analysis. The directed global significance score is calculated as the square root of the mean signed squared t-statistic for the genes in a gene set, with t-statistics coming from the linear regression underlying our differential expression analysis.   
                                      
                                       
                                  
                                
                              
                            
                           
                            
                          
                           
                      
                    
                    
                      
                           
                          
                               Day    Line     
                              
                                  Day  
                              
                               
                              
                                  Line: differential expression in Caco2 vs. baseline of Fibro  
                                  Line: differential expression in HEPG2 vs. baseline of Fibro  
                                  Line: differential expression in HT29 vs. baseline of Fibro  
                              
                              
                            
                              
                                
                                  
                                       
                                      
                                           Volcano Plot: Day   More Plot Information                Volcano Plot: Day  Volcano plot displaying each gene's -log10(p-value) and log2 fold change for the selected covariate. Highly statistically significant genes fall at the top of the plot, and highly differentially expressed genes fall to either side. Genes within the selected gene set are highlighted in orange. Horizontal lines indicate various False Discovery Rate (FDR) thresholds.   
                                           DE Results - Beta-catenin phosphorylation cascade genes - Day   More Plot Information      Download CSV Data     
                                           
                                             
                                             
                                           
                                           
                                             
                                               Probe Label 
                                               Log2 fold change 
                                               std error (log2) 
                                               Lower confidence limit (log2) 
                                               Upper confidence limit (log2) 
                                               Linear fold change 
                                               Lower confidence limit (linear) 
                                               Upper confidence limit (linear) 
                                               P-value 
                                               BY.p.value 
                                               method 
                                               Gene.sets 
                                               Codeset.Name 
                                               Probe.Label 
                                               Analyte.Type 
                                               Is.Control 
                                               Control.Type 
                                               Related.Probes 
                                               Probe.Annotation 
                                               KEGG.Pathways 
                                               Official.Gene.Name 
                                               Control.Conc 
                                               volcanocol 
                                               volcanopch 
                                               tstats.all 
                                             
                                           
                                           
                                             
                                               PPP2R1A-mRNA 
                                               0.0412 
                                               0.0143 
                                               0.013 
                                               0.0693 
                                               1.03 
                                               1.01 
                                               1.05 
                                               0.00677 
                                               0.099 
                                               lm.nb 
                                               ARMS-mediated activation, AXIN missense mutants destabilize the destruction complex, Beta-catenin phosphorylation cascade, CTLA4 inhibitory signaling, Cyclin AB1 associated events during G2M transition, Cyclin D associated events in G1, DAP12 signaling, DARPP-32 events, Disassembly of the destruction complex and recruitment of AXIN to the membrane, Downstream signal transduction, FCERI mediated MAPK activation, Frs2-mediated activation, Gastrin-CREB signalling pathway via PKC and MAPK, Interleukin receptor SHC signaling, IRS-related events triggered by IGF1R, MASTL Facilitates Mitotic Progression, Negative regulation of FGFR1 signaling, Negative regulation of FGFR2 signaling, Negative regulation of FGFR3 signaling, Negative regulation of FGFR4 signaling, Nonsense Mediated Decay (NMD) enhanced by the Exon Junction Complex (EJC), RAF activation, RAFMAP kinase cascade, Resolution of Sister Chromatid Cohesion, RHO GTPases Activate Formins, S33 mutants of beta-catenin aren.SQT.t phosphorylated, S37 mutants of beta-catenin aren.SQT.t phosphorylated, S45 mutants of beta-catenin aren.SQT.t phosphorylated, Signaling by ERBB2, Signaling by PDGF, Signaling by SCF-KIT, Spry regulation of FGF signaling, Truncations of AMER1 destabilize the destruction complex, VEGFA-VEGFR2 Pathway, VEGFR2 mediated cell proliferation 
                                               SEBELIUS_2_C4636 
                                               PPP2R1A 
                                               mRNA 
                                               0 
                                               
                                               
                                               MASTL Facilitates Mitotic Progression;Regulation of PLK1 Activity at G2M Transition;Disassembly of the destruction complex and recruitment of AXIN to the membrane;S37 mutants of beta-catenin aren.SQT.t phosphorylated;Cyclin AB1 associated events during G2M transition;RAF activation;RHO GTPases Activate Formins;S33 mutants of beta-catenin aren.SQT.t phosphorylated;Loss of Nlp from mitotic centrosomes;DARPP-32 events;Beta-catenin phosphorylation cascade;Initiation of Nuclear Envelope Reformation;Recruitment of mitotic centrosome proteins and complexes;Spry regulation of FGF signaling;Truncations of AMER1 destabilize the destruction complex;CTLA4 inhibitory signaling;AXIN missense mutants destabilize the destruction complex;S45 mutants of beta-catenin aren.SQT.t phosphorylated;Anchoring of the basal body to the plasma membrane;Nonsense Mediated Decay (NMD) enhanced by the Exon Junction Complex (EJC);Cyclin D associated events in G1;Platelet sensitization by LDL;Resolution of Sister Chromatid Cohesion;Inhibition of replication initiation of damaged DNA by RB1E2F1;RAFMAP kinase cascade;Downstream signal transduction;VEGFR2 mediated cell proliferation;Signaling by SCF-KIT;Frs2-mediated activation;ARMS-mediated activation;Gastrin-CREB signalling pathway via PKC and MAPK;Interleukin receptor SHC signaling;FCERI mediated MAPK activation;DAP12 signaling;Signaling by PDGF;VEGFA-VEGFR2 Pathway;Signaling by ERBB2;IRS-related events triggered by IGF1R;Negative regulation of FGFR3 signaling;Negative regulation of FGFR4 signaling;Negative regulation of FGFR2 signaling;Negative regulation of FGFR1 signaling 
                                               hsa03015;hsa04071;hsa04114;hsa04151;hsa04152;hsa04261;hsa04350;hsa04390;hsa04530;hsa04728;hsa04730;hsa05142;hsa05160 
                                               PPP2R1A 
                                               NA 
                                               slateblue4 
                                               16 
                                               2.88111888111888 
                                             
                                             
                                               AXIN1-mRNA 
                                               0.091 
                                               0.0371 
                                               0.0182 
                                               0.164 
                                               1.07 
                                               1.01 
                                               1.12 
                                               0.0191 
                                               0.24 
                                               lm.nb 
                                               AXIN missense mutants destabilize the destruction complex, Beta-catenin phosphorylation cascade, Disassembly of the destruction complex and recruitment of AXIN to the membrane, Misspliced GSK3beta mutants stabilize beta-catenin, S33 mutants of beta-catenin aren.SQT.t phosphorylated, S37 mutants of beta-catenin aren.SQT.t phosphorylated, S45 mutants of beta-catenin aren.SQT.t phosphorylated, T41 mutants of beta-catenin aren.SQT.t phosphorylated, Truncations of AMER1 destabilize the destruction complex 
                                               SEBELIUS_2_C4636 
                                               AXIN1 
                                               mRNA 
                                               0 
                                               
                                               
                                               APC truncation mutants have impaired AXIN binding;Misspliced GSK3beta mutants stabilize beta-catenin;T41 mutants of beta-catenin aren.SQT.t phosphorylated;AXIN missense mutants destabilize the destruction complex;S45 mutants of beta-catenin aren.SQT.t phosphorylated;Disassembly of the destruction complex and recruitment of AXIN to the membrane;S37 mutants of beta-catenin aren.SQT.t phosphorylated;S33 mutants of beta-catenin aren.SQT.t phosphorylated;Beta-catenin phosphorylation cascade;Truncations of AMER1 destabilize the destruction complex 
                                               hsa04310;hsa04390;hsa04550;hsa05200;hsa05210;hsa05213;hsa05217 
                                               AXIN1 
                                               NA 
                                               slateblue4 
                                               16 
                                               2.45283018867925 
                                             
                                             
                                               PPP2CB-mRNA 
                                               -0.0296 
                                               0.0132 
                                               -0.0554 
                                               -0.0037 
                                               0.98 
                                               0.962 
                                               0.997 
                                               0.0311 
                                               0.348 
                                               lm.nb 
                                               ARMS-mediated activation, AXIN missense mutants destabilize the destruction complex, Beta-catenin phosphorylation cascade, CTLA4 inhibitory signaling, Cyclin AB1 associated events during G2M transition, Cyclin D associated events in G1, DAP12 signaling, DARPP-32 events, Disassembly of the destruction complex and recruitment of AXIN to the membrane, Downstream signal transduction, ERKs are inactivated, FCERI mediated MAPK activation, Frs2-mediated activation, Gastrin-CREB signalling pathway via PKC and MAPK, Interleukin receptor SHC signaling, IRS-related events triggered by IGF1R, MASTL Facilitates Mitotic Progression, Misspliced GSK3beta mutants stabilize beta-catenin, Negative regulation of FGFR1 signaling, Negative regulation of FGFR2 signaling, Negative regulation of FGFR3 signaling, Negative regulation of FGFR4 signaling, Negative regulation of MAPK pathway, RAF activation, RAFMAP kinase cascade, Resolution of Sister Chromatid Cohesion, RHO GTPases Activate Formins, S33 mutants of beta-catenin aren.SQT.t phosphorylated, S37 mutants of beta-catenin aren.SQT.t phosphorylated, S45 mutants of beta-catenin aren.SQT.t phosphorylated, Signaling by ERBB2, Signaling by PDGF, Signaling by SCF-KIT, Spry regulation of FGF signaling, T41 mutants of beta-catenin aren.SQT.t phosphorylated, Truncations of AMER1 destabilize the destruction complex, VEGFA-VEGFR2 Pathway, VEGFR2 mediated cell proliferation 
                                               SEBELIUS_2_C4636 
                                               PPP2CB 
                                               mRNA 
                                               0 
                                               
                                               
                                               MASTL Facilitates Mitotic Progression;Misspliced GSK3beta mutants stabilize beta-catenin;T41 mutants of beta-catenin aren.SQT.t phosphorylated;ERKs are inactivated;Glycolysis;Negative regulation of MAPK pathway;Disassembly of the destruction complex and recruitment of AXIN to the membrane;S37 mutants of beta-catenin aren.SQT.t phosphorylated;Cyclin AB1 associated events during G2M transition;RAF activation;RHO GTPases Activate Formins;S33 mutants of beta-catenin aren.SQT.t phosphorylated;DARPP-32 events;Beta-catenin phosphorylation cascade;Spry regulation of FGF signaling;Truncations of AMER1 destabilize the destruction complex;CTLA4 inhibitory signaling;AXIN missense mutants destabilize the destruction complex;S45 mutants of beta-catenin aren.SQT.t phosphorylated;Cyclin D associated events in G1;Platelet sensitization by LDL;Resolution of Sister Chromatid Cohesion;Inhibition of replication initiation of damaged DNA by RB1E2F1;RAFMAP kinase cascade;Downstream signal transduction;VEGFR2 mediated cell proliferation;Signaling by SCF-KIT;Frs2-mediated activation;ARMS-mediated activation;Gastrin-CREB signalling pathway via PKC and MAPK;Interleukin receptor SHC signaling;FCERI mediated MAPK activation;DAP12 signaling;Signaling by PDGF;VEGFA-VEGFR2 Pathway;Signaling by ERBB2;IRS-related events triggered by IGF1R;Negative regulation of FGFR3 signaling;Negative regulation of FGFR4 signaling;Negative regulation of FGFR2 signaling;Negative regulation of FGFR1 signaling 
                                               hsa03015;hsa04071;hsa04114;hsa04151;hsa04152;hsa04261;hsa04350;hsa04390;hsa04530;hsa04728;hsa04730;hsa05142;hsa05160 
                                               PPP2CB 
                                               NA 
                                               slateblue4 
                                               16 
                                               -2.24242424242424 
                                             
                                             
                                               APC-mRNA 
                                               0.0285 
                                               0.0157 
                                               -0.0023 
                                               0.0592 
                                               1.02 
                                               0.998 
                                               1.04 
                                               0.0779 
                                               0.762 
                                               lm.nb 
                                               Apoptotic cleavage of cellular proteins, AXIN missense mutants destabilize the destruction complex, Beta-catenin phosphorylation cascade, Deactivation of the beta-catenin transactivating complex, Disassembly of the destruction complex and recruitment of AXIN to the membrane, Misspliced GSK3beta mutants stabilize beta-catenin, S33 mutants of beta-catenin aren.SQT.t phosphorylated, S37 mutants of beta-catenin aren.SQT.t phosphorylated, S45 mutants of beta-catenin aren.SQT.t phosphorylated, T41 mutants of beta-catenin aren.SQT.t phosphorylated, Truncations of AMER1 destabilize the destruction complex 
                                               SEBELIUS_2_C4636 
                                               APC 
                                               mRNA 
                                               0 
                                               
                                               
                                               Apoptotic cleavage of cellular proteins;Misspliced GSK3beta mutants stabilize beta-catenin;APC truncation mutants have impaired AXIN binding;T41 mutants of beta-catenin aren.SQT.t phosphorylated;AXIN missense mutants destabilize the destruction complex;S45 mutants of beta-catenin aren.SQT.t phosphorylated;Deactivation of the beta-catenin transactivating complex;Disassembly of the destruction complex and recruitment of AXIN to the membrane;S37 mutants of beta-catenin aren.SQT.t phosphorylated;S33 mutants of beta-catenin aren.SQT.t phosphorylated;Beta-catenin phosphorylation cascade;Truncations of AMER1 destabilize the destruction complex 
                                               hsa04310;hsa04390;hsa04550;hsa04810;hsa05166;hsa05200;hsa05206;hsa05210;hsa05213;hsa05217 
                                               APC 
                                               NA 
                                               azure4 
                                               1 
                                               1.81528662420382 
                                             
                                             
                                               PPP2CA-mRNA 
                                               -0.026 
                                               0.0146 
                                               -0.0547 
                                               0.00266 
                                               0.982 
                                               0.963 
                                               1 
                                               0.0836 
                                               0.802 
                                               lm.nb 
                                               ARMS-mediated activation, AXIN missense mutants destabilize the destruction complex, Beta-catenin phosphorylation cascade, CTLA4 inhibitory signaling, Cyclin AB1 associated events during G2M transition, Cyclin D associated events in G1, DAP12 signaling, DARPP-32 events, Disassembly of the destruction complex and recruitment of AXIN to the membrane, Downstream signal transduction, ERKs are inactivated, FCERI mediated MAPK activation, Frs2-mediated activation, Gastrin-CREB signalling pathway via PKC and MAPK, Interleukin receptor SHC signaling, IRS-related events triggered by IGF1R, MASTL Facilitates Mitotic Progression, Negative regulation of FGFR1 signaling, Negative regulation of FGFR2 signaling, Negative regulation of FGFR3 signaling, Negative regulation of FGFR4 signaling, Negative regulation of MAPK pathway, Nonsense Mediated Decay (NMD) enhanced by the Exon Junction Complex (EJC), RAF activation, RAFMAP kinase cascade, Resolution of Sister Chromatid Cohesion, RHO GTPases Activate Formins, S33 mutants of beta-catenin aren.SQT.t phosphorylated, S37 mutants of beta-catenin aren.SQT.t phosphorylated, S45 mutants of beta-catenin aren.SQT.t phosphorylated, Signaling by ERBB2, Signaling by PDGF, Signaling by SCF-KIT, Spry regulation of FGF signaling, T41 mutants of beta-catenin aren.SQT.t phosphorylated, Truncations of AMER1 destabilize the destruction complex, VEGFA-VEGFR2 Pathway, VEGFR2 mediated cell proliferation 
                                               SEBELIUS_2_C4636 
                                               PPP2CA 
                                               mRNA 
                                               0 
                                               
                                               
                                               MASTL Facilitates Mitotic Progression;T41 mutants of beta-catenin aren.SQT.t phosphorylated;ERKs are inactivated;Glycolysis;Negative regulation of MAPK pathway;Disassembly of the destruction complex and recruitment of AXIN to the membrane;S37 mutants of beta-catenin aren.SQT.t phosphorylated;Cyclin AB1 associated events during G2M transition;RAF activation;RHO GTPases Activate Formins;S33 mutants of beta-catenin aren.SQT.t phosphorylated;DARPP-32 events;Beta-catenin phosphorylation cascade;Initiation of Nuclear Envelope Reformation;Spry regulation of FGF signaling;Truncations of AMER1 destabilize the destruction complex;CTLA4 inhibitory signaling;AXIN missense mutants destabilize the destruction complex;S45 mutants of beta-catenin aren.SQT.t phosphorylated;Nonsense Mediated Decay (NMD) enhanced by the Exon Junction Complex (EJC);Cyclin D associated events in G1;Platelet sensitization by LDL;Resolution of Sister Chromatid Cohesion;Inhibition of replication initiation of damaged DNA by RB1E2F1;RAFMAP kinase cascade;Downstream signal transduction;VEGFR2 mediated cell proliferation;Signaling by SCF-KIT;Frs2-mediated activation;ARMS-mediated activation;Gastrin-CREB signalling pathway via PKC and MAPK;Interleukin receptor SHC signaling;FCERI mediated MAPK activation;DAP12 signaling;Signaling by PDGF;VEGFA-VEGFR2 Pathway;Signaling by ERBB2;IRS-related events triggered by IGF1R;Negative regulation of FGFR3 signaling;Negative regulation of FGFR4 signaling;Negative regulation of FGFR2 signaling;Negative regulation of FGFR1 signaling 
                                               hsa03015;hsa04071;hsa04114;hsa04151;hsa04152;hsa04261;hsa04350;hsa04390;hsa04530;hsa04728;hsa04730;hsa05142;hsa05160 
                                               PPP2CA 
                                               NA 
                                               azure4 
                                               1 
                                               -1.78082191780822 
                                             
                                             
                                               PPP2R1B-mRNA 
                                               0.0114 
                                               0.0135 
                                               -0.015 
                                               0.0379 
                                               1.01 
                                               0.99 
                                               1.03 
                                               0.402 
                                               1 
                                               lm.nb 
                                               ARMS-mediated activation, AXIN missense mutants destabilize the destruction complex, Beta-catenin phosphorylation cascade, CTLA4 inhibitory signaling, Cyclin AB1 associated events during G2M transition, Cyclin D associated events in G1, DAP12 signaling, DARPP-32 events, Disassembly of the destruction complex and recruitment of AXIN to the membrane, Downstream signal transduction, ERKs are inactivated, FCERI mediated MAPK activation, Frs2-mediated activation, Gastrin-CREB signalling pathway via PKC and MAPK, Interleukin receptor SHC signaling, IRS-related events triggered by IGF1R, MASTL Facilitates Mitotic Progression, Misspliced GSK3beta mutants stabilize beta-catenin, Negative regulation of MAPK pathway, RAF activation, RAFMAP kinase cascade, Resolution of Sister Chromatid Cohesion, RHO GTPases Activate Formins, S33 mutants of beta-catenin aren.SQT.t phosphorylated, S37 mutants of beta-catenin aren.SQT.t phosphorylated, S45 mutants of beta-catenin aren.SQT.t phosphorylated, Signaling by ERBB2, Signaling by PDGF, Signaling by SCF-KIT, T41 mutants of beta-catenin aren.SQT.t phosphorylated, Truncations of AMER1 destabilize the destruction complex, VEGFA-VEGFR2 Pathway, VEGFR2 mediated cell proliferation 
                                               SEBELIUS_2_C4636 
                                               PPP2R1B 
                                               mRNA 
                                               0 
                                               
                                               
                                               MASTL Facilitates Mitotic Progression;Misspliced GSK3beta mutants stabilize beta-catenin;T41 mutants of beta-catenin aren.SQT.t phosphorylated;ERKs are inactivated;Glycolysis;Negative regulation of MAPK pathway;Disassembly of the destruction complex and recruitment of AXIN to the membrane;S37 mutants of beta-catenin aren.SQT.t phosphorylated;Cyclin AB1 associated events during G2M transition;RAF activation;RHO GTPases Activate Formins;S33 mutants of beta-catenin aren.SQT.t phosphorylated;DARPP-32 events;Beta-catenin phosphorylation cascade;Truncations of AMER1 destabilize the destruction complex;CTLA4 inhibitory signaling;AXIN missense mutants destabilize the destruction complex;S45 mutants of beta-catenin aren.SQT.t phosphorylated;Cyclin D associated events in G1;Platelet sensitization by LDL;Resolution of Sister Chromatid Cohesion;Inhibition of replication initiation of damaged DNA by RB1E2F1;RAFMAP kinase cascade;Downstream signal transduction;VEGFR2 mediated cell proliferation;Signaling by SCF-KIT;Frs2-mediated activation;ARMS-mediated activation;Gastrin-CREB signalling pathway via PKC and MAPK;Interleukin receptor SHC signaling;FCERI mediated MAPK activation;DAP12 signaling;Signaling by PDGF;VEGFA-VEGFR2 Pathway;Signaling by ERBB2;IRS-related events triggered by IGF1R 
                                               hsa03015;hsa04071;hsa04114;hsa04151;hsa04152;hsa04261;hsa04350;hsa04390;hsa04530;hsa04728;hsa04730;hsa05142;hsa05160 
                                               PPP2R1B 
                                               NA 
                                               azure4 
                                               1 
                                               0.844444444444444 
                                             
                                             
                                               CSNK1A1-mRNA 
                                               0.0029 
                                               0.0109 
                                               -0.0186 
                                               0.0244 
                                               1 
                                               0.987 
                                               1.02 
                                               0.792 
                                               1 
                                               loglinear 
                                               AXIN missense mutants destabilize the destruction complex, Beta-catenin phosphorylation cascade, Disassembly of the destruction complex and recruitment of AXIN to the membrane, Misspliced GSK3beta mutants stabilize beta-catenin, S33 mutants of beta-catenin aren.SQT.t phosphorylated, S37 mutants of beta-catenin aren.SQT.t phosphorylated, S45 mutants of beta-catenin aren.SQT.t phosphorylated, T41 mutants of beta-catenin aren.SQT.t phosphorylated 
                                               SEBELIUS_2_C4636 
                                               CSNK1A1 
                                               mRNA 
                                               0 
                                               
                                               
                                               Misspliced GSK3beta mutants stabilize beta-catenin;APC truncation mutants have impaired AXIN binding;T41 mutants of beta-catenin aren.SQT.t phosphorylated;AXIN missense mutants destabilize the destruction complex;S45 mutants of beta-catenin aren.SQT.t phosphorylated;Disassembly of the destruction complex and recruitment of AXIN to the membrane;S37 mutants of beta-catenin aren.SQT.t phosphorylated;S33 mutants of beta-catenin aren.SQT.t phosphorylated;Beta-catenin phosphorylation cascade;Degradation of GLI2 by the proteasome 
                                               hsa04310 
                                               CSNK1A1 
                                               NA 
                                               azure4 
                                               1 
                                               0.26605504587156 
                                             
                                             
                                               CTNNB1-mRNA 
                                               -0.000519 
                                               0.0215 
                                               -0.0427 
                                               0.0416 
                                               1 
                                               0.971 
                                               1.03 
                                               0.981 
                                               1 
                                               lm.nb 
                                               Apoptotic cleavage of cellular proteins, Beta-catenin phosphorylation cascade, Deactivation of the beta-catenin transactivating complex, Disassembly of the destruction complex and recruitment of AXIN to the membrane, Formation of the beta-catenin.CL.TCF transactivating complex, Misspliced GSK3beta mutants stabilize beta-catenin, Repression of WNT target genes, RHO GTPases activate IQGAPs, S37 mutants of beta-catenin aren.SQT.t phosphorylated, S45 mutants of beta-catenin aren.SQT.t phosphorylated, T41 mutants of beta-catenin aren.SQT.t phosphorylated, VEGFA-VEGFR2 Pathway, VEGFR2 mediated vascular permeability 
                                               SEBELIUS_2_C4636 
                                               CTNNB1 
                                               mRNA 
                                               0 
                                               
                                               
                                               Disassembly of the destruction complex and recruitment of AXIN to the membrane;S37 mutants of beta-catenin aren.SQT.t phosphorylated;T41 mutants of beta-catenin aren.SQT.t phosphorylated;S45 mutants of beta-catenin aren.SQT.t phosphorylated;Binding of TCFLEF.CL.CTNNB1 to target gene promoters;RHO GTPases activate IQGAPs;Deactivation of the beta-catenin transactivating complex;Repression of WNT target genes;Apoptotic cleavage of cell adhesion  proteins;Adherens junctions interactions;Misspliced GSK3beta mutants stabilize beta-catenin;VEGFR2 mediated vascular permeability;Beta-catenin phosphorylation cascade;Synthesis# secretion# and inactivation of Glucagon-like Peptide-1 (GLP-1);Formation of the beta-catenin.CL.TCF transactivating complex;Apoptotic cleavage of cellular proteins;VEGFA-VEGFR2 Pathway 
                                               hsa04015;hsa04310;hsa04390;hsa04510;hsa04520;hsa04530;hsa04550;hsa04670;hsa04916;hsa04919;hsa05100;hsa05130;hsa05166;hsa05200;hsa05205;hsa05210;hsa05213;hsa05215;hsa05216;hsa05217;hsa05412 
                                               CTNNB1 
                                               NA 
                                               azure4 
                                               1 
                                               -0.0241395348837209 
                                             
                                           
                                           DE Results - Beta-catenin phosphorylation cascade genes - Day  Table displaying each sample's global significance scores and directed global significance scores as defined in the heatmaps above. The global significance score is calculated as the square root of the mean squared t-statistic for the genes in a gene set, with t-statistics coming from the linear regression underlying our differential expression analysis. The directed global significance score is calculated as the square root of the mean signed squared t-statistic for the genes in a gene set, with t-statistics coming from the linear regression underlying our differential expression analysis.   
                                      
                                       
                                  
                                
                              
                            
                            
                              
                                
                                  
                                       
                                      
                                           Volcano Plot: Line: Caco2 vs.Fibro   More Plot Information                Volcano Plot: Line: Caco2 vs.Fibro  Volcano plot displaying each gene's -log10(p-value) and log2 fold change for the selected covariate. Highly statistically significant genes fall at the top of the plot, and highly differentially expressed genes fall to either side. Genes within the selected gene set are highlighted in orange. Horizontal lines indicate various False Discovery Rate (FDR) thresholds.   
                                           DE Results - Beta-catenin phosphorylation cascade genes - Line: Caco2 vs.Fibro   More Plot Information      Download CSV Data     
                                           
                                             
                                             
                                           
                                           
                                             
                                               Probe Label 
                                               Log2 fold change 
                                               std error (log2) 
                                               Lower confidence limit (log2) 
                                               Upper confidence limit (log2) 
                                               Linear fold change 
                                               Lower confidence limit (linear) 
                                               Upper confidence limit (linear) 
                                               P-value 
                                               BY.p.value 
                                               method 
                                               Gene.sets 
                                               Codeset.Name 
                                               Probe.Label 
                                               Analyte.Type 
                                               Is.Control 
                                               Control.Type 
                                               Related.Probes 
                                               Probe.Annotation 
                                               KEGG.Pathways 
                                               Official.Gene.Name 
                                               Control.Conc 
                                               volcanocol 
                                               volcanopch 
                                               tstats.all 
                                             
                                           
                                           
                                             
                                               PPP2R1B-mRNA 
                                               1.57 
                                               0.104 
                                               1.37 
                                               1.78 
                                               2.98 
                                               2.59 
                                               3.43 
                                               1.73e-17 
                                               6.65e-16 
                                               lm.nb 
                                               ARMS-mediated activation, AXIN missense mutants destabilize the destruction complex, Beta-catenin phosphorylation cascade, CTLA4 inhibitory signaling, Cyclin AB1 associated events during G2M transition, Cyclin D associated events in G1, DAP12 signaling, DARPP-32 events, Disassembly of the destruction complex and recruitment of AXIN to the membrane, Downstream signal transduction, ERKs are inactivated, FCERI mediated MAPK activation, Frs2-mediated activation, Gastrin-CREB signalling pathway via PKC and MAPK, Interleukin receptor SHC signaling, IRS-related events triggered by IGF1R, MASTL Facilitates Mitotic Progression, Misspliced GSK3beta mutants stabilize beta-catenin, Negative regulation of MAPK pathway, RAF activation, RAFMAP kinase cascade, Resolution of Sister Chromatid Cohesion, RHO GTPases Activate Formins, S33 mutants of beta-catenin aren.SQT.t phosphorylated, S37 mutants of beta-catenin aren.SQT.t phosphorylated, S45 mutants of beta-catenin aren.SQT.t phosphorylated, Signaling by ERBB2, Signaling by PDGF, Signaling by SCF-KIT, T41 mutants of beta-catenin aren.SQT.t phosphorylated, Truncations of AMER1 destabilize the destruction complex, VEGFA-VEGFR2 Pathway, VEGFR2 mediated cell proliferation 
                                               SEBELIUS_2_C4636 
                                               PPP2R1B 
                                               mRNA 
                                               0 
                                               
                                               
                                               MASTL Facilitates Mitotic Progression;Misspliced GSK3beta mutants stabilize beta-catenin;T41 mutants of beta-catenin aren.SQT.t phosphorylated;ERKs are inactivated;Glycolysis;Negative regulation of MAPK pathway;Disassembly of the destruction complex and recruitment of AXIN to the membrane;S37 mutants of beta-catenin aren.SQT.t phosphorylated;Cyclin AB1 associated events during G2M transition;RAF activation;RHO GTPases Activate Formins;S33 mutants of beta-catenin aren.SQT.t phosphorylated;DARPP-32 events;Beta-catenin phosphorylation cascade;Truncations of AMER1 destabilize the destruction complex;CTLA4 inhibitory signaling;AXIN missense mutants destabilize the destruction complex;S45 mutants of beta-catenin aren.SQT.t phosphorylated;Cyclin D associated events in G1;Platelet sensitization by LDL;Resolution of Sister Chromatid Cohesion;Inhibition of replication initiation of damaged DNA by RB1E2F1;RAFMAP kinase cascade;Downstream signal transduction;VEGFR2 mediated cell proliferation;Signaling by SCF-KIT;Frs2-mediated activation;ARMS-mediated activation;Gastrin-CREB signalling pathway via PKC and MAPK;Interleukin receptor SHC signaling;FCERI mediated MAPK activation;DAP12 signaling;Signaling by PDGF;VEGFA-VEGFR2 Pathway;Signaling by ERBB2;IRS-related events triggered by IGF1R 
                                               hsa03015;hsa04071;hsa04114;hsa04151;hsa04152;hsa04261;hsa04350;hsa04390;hsa04530;hsa04728;hsa04730;hsa05142;hsa05160 
                                               PPP2R1B 
                                               NA 
                                               slateblue1 
                                               16 
                                               15.0961538461538 
                                             
                                             
                                               PPP2CB-mRNA 
                                               -0.814 
                                               0.099 
                                               -1.01 
                                               -0.62 
                                               0.569 
                                               0.497 
                                               0.651 
                                               7.15e-10 
                                               1.11e-08 
                                               lm.nb 
                                               ARMS-mediated activation, AXIN missense mutants destabilize the destruction complex, Beta-catenin phosphorylation cascade, CTLA4 inhibitory signaling, Cyclin AB1 associated events during G2M transition, Cyclin D associated events in G1, DAP12 signaling, DARPP-32 events, Disassembly of the destruction complex and recruitment of AXIN to the membrane, Downstream signal transduction, ERKs are inactivated, FCERI mediated MAPK activation, Frs2-mediated activation, Gastrin-CREB signalling pathway via PKC and MAPK, Interleukin receptor SHC signaling, IRS-related events triggered by IGF1R, MASTL Facilitates Mitotic Progression, Misspliced GSK3beta mutants stabilize beta-catenin, Negative regulation of FGFR1 signaling, Negative regulation of FGFR2 signaling, Negative regulation of FGFR3 signaling, Negative regulation of FGFR4 signaling, Negative regulation of MAPK pathway, RAF activation, RAFMAP kinase cascade, Resolution of Sister Chromatid Cohesion, RHO GTPases Activate Formins, S33 mutants of beta-catenin aren.SQT.t phosphorylated, S37 mutants of beta-catenin aren.SQT.t phosphorylated, S45 mutants of beta-catenin aren.SQT.t phosphorylated, Signaling by ERBB2, Signaling by PDGF, Signaling by SCF-KIT, Spry regulation of FGF signaling, T41 mutants of beta-catenin aren.SQT.t phosphorylated, Truncations of AMER1 destabilize the destruction complex, VEGFA-VEGFR2 Pathway, VEGFR2 mediated cell proliferation 
                                               SEBELIUS_2_C4636 
                                               PPP2CB 
                                               mRNA 
                                               0 
                                               
                                               
                                               MASTL Facilitates Mitotic Progression;Misspliced GSK3beta mutants stabilize beta-catenin;T41 mutants of beta-catenin aren.SQT.t phosphorylated;ERKs are inactivated;Glycolysis;Negative regulation of MAPK pathway;Disassembly of the destruction complex and recruitment of AXIN to the membrane;S37 mutants of beta-catenin aren.SQT.t phosphorylated;Cyclin AB1 associated events during G2M transition;RAF activation;RHO GTPases Activate Formins;S33 mutants of beta-catenin aren.SQT.t phosphorylated;DARPP-32 events;Beta-catenin phosphorylation cascade;Spry regulation of FGF signaling;Truncations of AMER1 destabilize the destruction complex;CTLA4 inhibitory signaling;AXIN missense mutants destabilize the destruction complex;S45 mutants of beta-catenin aren.SQT.t phosphorylated;Cyclin D associated events in G1;Platelet sensitization by LDL;Resolution of Sister Chromatid Cohesion;Inhibition of replication initiation of damaged DNA by RB1E2F1;RAFMAP kinase cascade;Downstream signal transduction;VEGFR2 mediated cell proliferation;Signaling by SCF-KIT;Frs2-mediated activation;ARMS-mediated activation;Gastrin-CREB signalling pathway via PKC and MAPK;Interleukin receptor SHC signaling;FCERI mediated MAPK activation;DAP12 signaling;Signaling by PDGF;VEGFA-VEGFR2 Pathway;Signaling by ERBB2;IRS-related events triggered by IGF1R;Negative regulation of FGFR3 signaling;Negative regulation of FGFR4 signaling;Negative regulation of FGFR2 signaling;Negative regulation of FGFR1 signaling 
                                               hsa03015;hsa04071;hsa04114;hsa04151;hsa04152;hsa04261;hsa04350;hsa04390;hsa04530;hsa04728;hsa04730;hsa05142;hsa05160 
                                               PPP2CB 
                                               NA 
                                               slateblue1 
                                               16 
                                               -8.22222222222222 
                                             
                                             
                                               APC-mRNA 
                                               -0.862 
                                               0.115 
                                               -1.09 
                                               -0.637 
                                               0.55 
                                               0.471 
                                               0.643 
                                               6.14e-09 
                                               8.54e-08 
                                               lm.nb 
                                               Apoptotic cleavage of cellular proteins, AXIN missense mutants destabilize the destruction complex, Beta-catenin phosphorylation cascade, Deactivation of the beta-catenin transactivating complex, Disassembly of the destruction complex and recruitment of AXIN to the membrane, Misspliced GSK3beta mutants stabilize beta-catenin, S33 mutants of beta-catenin aren.SQT.t phosphorylated, S37 mutants of beta-catenin aren.SQT.t phosphorylated, S45 mutants of beta-catenin aren.SQT.t phosphorylated, T41 mutants of beta-catenin aren.SQT.t phosphorylated, Truncations of AMER1 destabilize the destruction complex 
                                               SEBELIUS_2_C4636 
                                               APC 
                                               mRNA 
                                               0 
                                               
                                               
                                               Apoptotic cleavage of cellular proteins;Misspliced GSK3beta mutants stabilize beta-catenin;APC truncation mutants have impaired AXIN binding;T41 mutants of beta-catenin aren.SQT.t phosphorylated;AXIN missense mutants destabilize the destruction complex;S45 mutants of beta-catenin aren.SQT.t phosphorylated;Deactivation of the beta-catenin transactivating complex;Disassembly of the destruction complex and recruitment of AXIN to the membrane;S37 mutants of beta-catenin aren.SQT.t phosphorylated;S33 mutants of beta-catenin aren.SQT.t phosphorylated;Beta-catenin phosphorylation cascade;Truncations of AMER1 destabilize the destruction complex 
                                               hsa04310;hsa04390;hsa04550;hsa04810;hsa05166;hsa05200;hsa05206;hsa05210;hsa05213;hsa05217 
                                               APC 
                                               NA 
                                               slateblue1 
                                               16 
                                               -7.49565217391304 
                                             
                                             
                                               CSNK1A1-mRNA 
                                               -0.579 
                                               0.0828 
                                               -0.741 
                                               -0.417 
                                               0.669 
                                               0.598 
                                               0.749 
                                               2.88e-08 
                                               3.73e-07 
                                               loglinear 
                                               AXIN missense mutants destabilize the destruction complex, Beta-catenin phosphorylation cascade, Disassembly of the destruction complex and recruitment of AXIN to the membrane, Misspliced GSK3beta mutants stabilize beta-catenin, S33 mutants of beta-catenin aren.SQT.t phosphorylated, S37 mutants of beta-catenin aren.SQT.t phosphorylated, S45 mutants of beta-catenin aren.SQT.t phosphorylated, T41 mutants of beta-catenin aren.SQT.t phosphorylated 
                                               SEBELIUS_2_C4636 
                                               CSNK1A1 
                                               mRNA 
                                               0 
                                               
                                               
                                               Misspliced GSK3beta mutants stabilize beta-catenin;APC truncation mutants have impaired AXIN binding;T41 mutants of beta-catenin aren.SQT.t phosphorylated;AXIN missense mutants destabilize the destruction complex;S45 mutants of beta-catenin aren.SQT.t phosphorylated;Disassembly of the destruction complex and recruitment of AXIN to the membrane;S37 mutants of beta-catenin aren.SQT.t phosphorylated;S33 mutants of beta-catenin aren.SQT.t phosphorylated;Beta-catenin phosphorylation cascade;Degradation of GLI2 by the proteasome 
                                               hsa04310 
                                               CSNK1A1 
                                               NA 
                                               slateblue1 
                                               16 
                                               -6.99275362318841 
                                             
                                             
                                               AXIN1-mRNA 
                                               0.571 
                                               0.285 
                                               0.012 
                                               1.13 
                                               1.49 
                                               1.01 
                                               2.19 
                                               0.0526 
                                               0.397 
                                               lm.nb 
                                               AXIN missense mutants destabilize the destruction complex, Beta-catenin phosphorylation cascade, Disassembly of the destruction complex and recruitment of AXIN to the membrane, Misspliced GSK3beta mutants stabilize beta-catenin, S33 mutants of beta-catenin aren.SQT.t phosphorylated, S37 mutants of beta-catenin aren.SQT.t phosphorylated, S45 mutants of beta-catenin aren.SQT.t phosphorylated, T41 mutants of beta-catenin aren.SQT.t phosphorylated, Truncations of AMER1 destabilize the destruction complex 
                                               SEBELIUS_2_C4636 
                                               AXIN1 
                                               mRNA 
                                               0 
                                               
                                               
                                               APC truncation mutants have impaired AXIN binding;Misspliced GSK3beta mutants stabilize beta-catenin;T41 mutants of beta-catenin aren.SQT.t phosphorylated;AXIN missense mutants destabilize the destruction complex;S45 mutants of beta-catenin aren.SQT.t phosphorylated;Disassembly of the destruction complex and recruitment of AXIN to the membrane;S37 mutants of beta-catenin aren.SQT.t phosphorylated;S33 mutants of beta-catenin aren.SQT.t phosphorylated;Beta-catenin phosphorylation cascade;Truncations of AMER1 destabilize the destruction complex 
                                               hsa04310;hsa04390;hsa04550;hsa05200;hsa05210;hsa05213;hsa05217 
                                               AXIN1 
                                               NA 
                                               slateblue4 
                                               16 
                                               2.00350877192982 
                                             
                                             
                                               PPP2CA-mRNA 
                                               0.168 
                                               0.111 
                                               -0.0492 
                                               0.385 
                                               1.12 
                                               0.966 
                                               1.31 
                                               0.138 
                                               0.975 
                                               lm.nb 
                                               ARMS-mediated activation, AXIN missense mutants destabilize the destruction complex, Beta-catenin phosphorylation cascade, CTLA4 inhibitory signaling, Cyclin AB1 associated events during G2M transition, Cyclin D associated events in G1, DAP12 signaling, DARPP-32 events, Disassembly of the destruction complex and recruitment of AXIN to the membrane, Downstream signal transduction, ERKs are inactivated, FCERI mediated MAPK activation, Frs2-mediated activation, Gastrin-CREB signalling pathway via PKC and MAPK, Interleukin receptor SHC signaling, IRS-related events triggered by IGF1R, MASTL Facilitates Mitotic Progression, Negative regulation of FGFR1 signaling, Negative regulation of FGFR2 signaling, Negative regulation of FGFR3 signaling, Negative regulation of FGFR4 signaling, Negative regulation of MAPK pathway, Nonsense Mediated Decay (NMD) enhanced by the Exon Junction Complex (EJC), RAF activation, RAFMAP kinase cascade, Resolution of Sister Chromatid Cohesion, RHO GTPases Activate Formins, S33 mutants of beta-catenin aren.SQT.t phosphorylated, S37 mutants of beta-catenin aren.SQT.t phosphorylated, S45 mutants of beta-catenin aren.SQT.t phosphorylated, Signaling by ERBB2, Signaling by PDGF, Signaling by SCF-KIT, Spry regulation of FGF signaling, T41 mutants of beta-catenin aren.SQT.t phosphorylated, Truncations of AMER1 destabilize the destruction complex, VEGFA-VEGFR2 Pathway, VEGFR2 mediated cell proliferation 
                                               SEBELIUS_2_C4636 
                                               PPP2CA 
                                               mRNA 
                                               0 
                                               
                                               
                                               MASTL Facilitates Mitotic Progression;T41 mutants of beta-catenin aren.SQT.t phosphorylated;ERKs are inactivated;Glycolysis;Negative regulation of MAPK pathway;Disassembly of the destruction complex and recruitment of AXIN to the membrane;S37 mutants of beta-catenin aren.SQT.t phosphorylated;Cyclin AB1 associated events during G2M transition;RAF activation;RHO GTPases Activate Formins;S33 mutants of beta-catenin aren.SQT.t phosphorylated;DARPP-32 events;Beta-catenin phosphorylation cascade;Initiation of Nuclear Envelope Reformation;Spry regulation of FGF signaling;Truncations of AMER1 destabilize the destruction complex;CTLA4 inhibitory signaling;AXIN missense mutants destabilize the destruction complex;S45 mutants of beta-catenin aren.SQT.t phosphorylated;Nonsense Mediated Decay (NMD) enhanced by the Exon Junction Complex (EJC);Cyclin D associated events in G1;Platelet sensitization by LDL;Resolution of Sister Chromatid Cohesion;Inhibition of replication initiation of damaged DNA by RB1E2F1;RAFMAP kinase cascade;Downstream signal transduction;VEGFR2 mediated cell proliferation;Signaling by SCF-KIT;Frs2-mediated activation;ARMS-mediated activation;Gastrin-CREB signalling pathway via PKC and MAPK;Interleukin receptor SHC signaling;FCERI mediated MAPK activation;DAP12 signaling;Signaling by PDGF;VEGFA-VEGFR2 Pathway;Signaling by ERBB2;IRS-related events triggered by IGF1R;Negative regulation of FGFR3 signaling;Negative regulation of FGFR4 signaling;Negative regulation of FGFR2 signaling;Negative regulation of FGFR1 signaling 
                                               hsa03015;hsa04071;hsa04114;hsa04151;hsa04152;hsa04261;hsa04350;hsa04390;hsa04530;hsa04728;hsa04730;hsa05142;hsa05160 
                                               PPP2CA 
                                               NA 
                                               azure4 
                                               1 
                                               1.51351351351351 
                                             
                                             
                                               PPP2R1A-mRNA 
                                               0.0488 
                                               0.108 
                                               -0.164 
                                               0.261 
                                               1.03 
                                               0.893 
                                               1.2 
                                               0.656 
                                               1 
                                               lm.nb 
                                               ARMS-mediated activation, AXIN missense mutants destabilize the destruction complex, Beta-catenin phosphorylation cascade, CTLA4 inhibitory signaling, Cyclin AB1 associated events during G2M transition, Cyclin D associated events in G1, DAP12 signaling, DARPP-32 events, Disassembly of the destruction complex and recruitment of AXIN to the membrane, Downstream signal transduction, FCERI mediated MAPK activation, Frs2-mediated activation, Gastrin-CREB signalling pathway via PKC and MAPK, Interleukin receptor SHC signaling, IRS-related events triggered by IGF1R, MASTL Facilitates Mitotic Progression, Negative regulation of FGFR1 signaling, Negative regulation of FGFR2 signaling, Negative regulation of FGFR3 signaling, Negative regulation of FGFR4 signaling, Nonsense Mediated Decay (NMD) enhanced by the Exon Junction Complex (EJC), RAF activation, RAFMAP kinase cascade, Resolution of Sister Chromatid Cohesion, RHO GTPases Activate Formins, S33 mutants of beta-catenin aren.SQT.t phosphorylated, S37 mutants of beta-catenin aren.SQT.t phosphorylated, S45 mutants of beta-catenin aren.SQT.t phosphorylated, Signaling by ERBB2, Signaling by PDGF, Signaling by SCF-KIT, Spry regulation of FGF signaling, Truncations of AMER1 destabilize the destruction complex, VEGFA-VEGFR2 Pathway, VEGFR2 mediated cell proliferation 
                                               SEBELIUS_2_C4636 
                                               PPP2R1A 
                                               mRNA 
                                               0 
                                               
                                               
                                               MASTL Facilitates Mitotic Progression;Regulation of PLK1 Activity at G2M Transition;Disassembly of the destruction complex and recruitment of AXIN to the membrane;S37 mutants of beta-catenin aren.SQT.t phosphorylated;Cyclin AB1 associated events during G2M transition;RAF activation;RHO GTPases Activate Formins;S33 mutants of beta-catenin aren.SQT.t phosphorylated;Loss of Nlp from mitotic centrosomes;DARPP-32 events;Beta-catenin phosphorylation cascade;Initiation of Nuclear Envelope Reformation;Recruitment of mitotic centrosome proteins and complexes;Spry regulation of FGF signaling;Truncations of AMER1 destabilize the destruction complex;CTLA4 inhibitory signaling;AXIN missense mutants destabilize the destruction complex;S45 mutants of beta-catenin aren.SQT.t phosphorylated;Anchoring of the basal body to the plasma membrane;Nonsense Mediated Decay (NMD) enhanced by the Exon Junction Complex (EJC);Cyclin D associated events in G1;Platelet sensitization by LDL;Resolution of Sister Chromatid Cohesion;Inhibition of replication initiation of damaged DNA by RB1E2F1;RAFMAP kinase cascade;Downstream signal transduction;VEGFR2 mediated cell proliferation;Signaling by SCF-KIT;Frs2-mediated activation;ARMS-mediated activation;Gastrin-CREB signalling pathway via PKC and MAPK;Interleukin receptor SHC signaling;FCERI mediated MAPK activation;DAP12 signaling;Signaling by PDGF;VEGFA-VEGFR2 Pathway;Signaling by ERBB2;IRS-related events triggered by IGF1R;Negative regulation of FGFR3 signaling;Negative regulation of FGFR4 signaling;Negative regulation of FGFR2 signaling;Negative regulation of FGFR1 signaling 
                                               hsa03015;hsa04071;hsa04114;hsa04151;hsa04152;hsa04261;hsa04350;hsa04390;hsa04530;hsa04728;hsa04730;hsa05142;hsa05160 
                                               PPP2R1A 
                                               NA 
                                               azure4 
                                               1 
                                               0.451851851851852 
                                             
                                             
                                               CTNNB1-mRNA 
                                               0.0018 
                                               0.163 
                                               -0.317 
                                               0.32 
                                               1 
                                               0.803 
                                               1.25 
                                               0.991 
                                               1 
                                               lm.nb 
                                               Apoptotic cleavage of cellular proteins, Beta-catenin phosphorylation cascade, Deactivation of the beta-catenin transactivating complex, Disassembly of the destruction complex and recruitment of AXIN to the membrane, Formation of the beta-catenin.CL.TCF transactivating complex, Misspliced GSK3beta mutants stabilize beta-catenin, Repression of WNT target genes, RHO GTPases activate IQGAPs, S37 mutants of beta-catenin aren.SQT.t phosphorylated, S45 mutants of beta-catenin aren.SQT.t phosphorylated, T41 mutants of beta-catenin aren.SQT.t phosphorylated, VEGFA-VEGFR2 Pathway, VEGFR2 mediated vascular permeability 
                                               SEBELIUS_2_C4636 
                                               CTNNB1 
                                               mRNA 
                                               0 
                                               
                                               
                                               Disassembly of the destruction complex and recruitment of AXIN to the membrane;S37 mutants of beta-catenin aren.SQT.t phosphorylated;T41 mutants of beta-catenin aren.SQT.t phosphorylated;S45 mutants of beta-catenin aren.SQT.t phosphorylated;Binding of TCFLEF.CL.CTNNB1 to target gene promoters;RHO GTPases activate IQGAPs;Deactivation of the beta-catenin transactivating complex;Repression of WNT target genes;Apoptotic cleavage of cell adhesion  proteins;Adherens junctions interactions;Misspliced GSK3beta mutants stabilize beta-catenin;VEGFR2 mediated vascular permeability;Beta-catenin phosphorylation cascade;Synthesis# secretion# and inactivation of Glucagon-like Peptide-1 (GLP-1);Formation of the beta-catenin.CL.TCF transactivating complex;Apoptotic cleavage of cellular proteins;VEGFA-VEGFR2 Pathway 
                                               hsa04015;hsa04310;hsa04390;hsa04510;hsa04520;hsa04530;hsa04550;hsa04670;hsa04916;hsa04919;hsa05100;hsa05130;hsa05166;hsa05200;hsa05205;hsa05210;hsa05213;hsa05215;hsa05216;hsa05217;hsa05412 
                                               CTNNB1 
                                               NA 
                                               azure4 
                                               1 
                                               0.0110429447852761 
                                             
                                           
                                           DE Results - Beta-catenin phosphorylation cascade genes - Line: Caco2 vs.Fibro  Table displaying each sample's global significance scores and directed global significance scores as defined in the heatmaps above. The global significance score is calculated as the square root of the mean squared t-statistic for the genes in a gene set, with t-statistics coming from the linear regression underlying our differential expression analysis. The directed global significance score is calculated as the square root of the mean signed squared t-statistic for the genes in a gene set, with t-statistics coming from the linear regression underlying our differential expression analysis.   
                                      
                                       
                                  
                                
                                
                                  
                                       
                                      
                                           Volcano Plot: Line: HEPG2 vs.Fibro   More Plot Information                Volcano Plot: Line: HEPG2 vs.Fibro  Volcano plot displaying each gene's -log10(p-value) and log2 fold change for the selected covariate. Highly statistically significant genes fall at the top of the plot, and highly differentially expressed genes fall to either side. Genes within the selected gene set are highlighted in orange. Horizontal lines indicate various False Discovery Rate (FDR) thresholds.   
                                           DE Results - Beta-catenin phosphorylation cascade genes - Line: HEPG2 vs.Fibro   More Plot Information      Download CSV Data     
                                           
                                             
                                             
                                           
                                           
                                             
                                               Probe Label 
                                               Log2 fold change 
                                               std error (log2) 
                                               Lower confidence limit (log2) 
                                               Upper confidence limit (log2) 
                                               Linear fold change 
                                               Lower confidence limit (linear) 
                                               Upper confidence limit (linear) 
                                               P-value 
                                               BY.p.value 
                                               method 
                                               Gene.sets 
                                               Codeset.Name 
                                               Probe.Label 
                                               Analyte.Type 
                                               Is.Control 
                                               Control.Type 
                                               Related.Probes 
                                               Probe.Annotation 
                                               KEGG.Pathways 
                                               Official.Gene.Name 
                                               Control.Conc 
                                               volcanocol 
                                               volcanopch 
                                               tstats.all 
                                             
                                           
                                           
                                             
                                               CTNNB1-mRNA 
                                               1.77 
                                               0.178 
                                               1.42 
                                               2.12 
                                               3.4 
                                               2.67 
                                               4.33 
                                               6.08e-12 
                                               1.41e-10 
                                               lm.nb 
                                               Apoptotic cleavage of cellular proteins, Beta-catenin phosphorylation cascade, Deactivation of the beta-catenin transactivating complex, Disassembly of the destruction complex and recruitment of AXIN to the membrane, Formation of the beta-catenin.CL.TCF transactivating complex, Misspliced GSK3beta mutants stabilize beta-catenin, Repression of WNT target genes, RHO GTPases activate IQGAPs, S37 mutants of beta-catenin aren.SQT.t phosphorylated, S45 mutants of beta-catenin aren.SQT.t phosphorylated, T41 mutants of beta-catenin aren.SQT.t phosphorylated, VEGFA-VEGFR2 Pathway, VEGFR2 mediated vascular permeability 
                                               SEBELIUS_2_C4636 
                                               CTNNB1 
                                               mRNA 
                                               0 
                                               
                                               
                                               Disassembly of the destruction complex and recruitment of AXIN to the membrane;S37 mutants of beta-catenin aren.SQT.t phosphorylated;T41 mutants of beta-catenin aren.SQT.t phosphorylated;S45 mutants of beta-catenin aren.SQT.t phosphorylated;Binding of TCFLEF.CL.CTNNB1 to target gene promoters;RHO GTPases activate IQGAPs;Deactivation of the beta-catenin transactivating complex;Repression of WNT target genes;Apoptotic cleavage of cell adhesion  proteins;Adherens junctions interactions;Misspliced GSK3beta mutants stabilize beta-catenin;VEGFR2 mediated vascular permeability;Beta-catenin phosphorylation cascade;Synthesis# secretion# and inactivation of Glucagon-like Peptide-1 (GLP-1);Formation of the beta-catenin.CL.TCF transactivating complex;Apoptotic cleavage of cellular proteins;VEGFA-VEGFR2 Pathway 
                                               hsa04015;hsa04310;hsa04390;hsa04510;hsa04520;hsa04530;hsa04550;hsa04670;hsa04916;hsa04919;hsa05100;hsa05130;hsa05166;hsa05200;hsa05205;hsa05210;hsa05213;hsa05215;hsa05216;hsa05217;hsa05412 
                                               CTNNB1 
                                               NA 
                                               slateblue1 
                                               16 
                                               9.9438202247191 
                                             
                                             
                                               PPP2R1B-mRNA 
                                               1.04 
                                               0.114 
                                               0.818 
                                               1.26 
                                               2.06 
                                               1.76 
                                               2.4 
                                               4.71e-11 
                                               9.7e-10 
                                               lm.nb 
                                               ARMS-mediated activation, AXIN missense mutants destabilize the destruction complex, Beta-catenin phosphorylation cascade, CTLA4 inhibitory signaling, Cyclin AB1 associated events during G2M transition, Cyclin D associated events in G1, DAP12 signaling, DARPP-32 events, Disassembly of the destruction complex and recruitment of AXIN to the membrane, Downstream signal transduction, ERKs are inactivated, FCERI mediated MAPK activation, Frs2-mediated activation, Gastrin-CREB signalling pathway via PKC and MAPK, Interleukin receptor SHC signaling, IRS-related events triggered by IGF1R, MASTL Facilitates Mitotic Progression, Misspliced GSK3beta mutants stabilize beta-catenin, Negative regulation of MAPK pathway, RAF activation, RAFMAP kinase cascade, Resolution of Sister Chromatid Cohesion, RHO GTPases Activate Formins, S33 mutants of beta-catenin aren.SQT.t phosphorylated, S37 mutants of beta-catenin aren.SQT.t phosphorylated, S45 mutants of beta-catenin aren.SQT.t phosphorylated, Signaling by ERBB2, Signaling by PDGF, Signaling by SCF-KIT, T41 mutants of beta-catenin aren.SQT.t phosphorylated, Truncations of AMER1 destabilize the destruction complex, VEGFA-VEGFR2 Pathway, VEGFR2 mediated cell proliferation 
                                               SEBELIUS_2_C4636 
                                               PPP2R1B 
                                               mRNA 
                                               0 
                                               
                                               
                                               MASTL Facilitates Mitotic Progression;Misspliced GSK3beta mutants stabilize beta-catenin;T41 mutants of beta-catenin aren.SQT.t phosphorylated;ERKs are inactivated;Glycolysis;Negative regulation of MAPK pathway;Disassembly of the destruction complex and recruitment of AXIN to the membrane;S37 mutants of beta-catenin aren.SQT.t phosphorylated;Cyclin AB1 associated events during G2M transition;RAF activation;RHO GTPases Activate Formins;S33 mutants of beta-catenin aren.SQT.t phosphorylated;DARPP-32 events;Beta-catenin phosphorylation cascade;Truncations of AMER1 destabilize the destruction complex;CTLA4 inhibitory signaling;AXIN missense mutants destabilize the destruction complex;S45 mutants of beta-catenin aren.SQT.t phosphorylated;Cyclin D associated events in G1;Platelet sensitization by LDL;Resolution of Sister Chromatid Cohesion;Inhibition of replication initiation of damaged DNA by RB1E2F1;RAFMAP kinase cascade;Downstream signal transduction;VEGFR2 mediated cell proliferation;Signaling by SCF-KIT;Frs2-mediated activation;ARMS-mediated activation;Gastrin-CREB signalling pathway via PKC and MAPK;Interleukin receptor SHC signaling;FCERI mediated MAPK activation;DAP12 signaling;Signaling by PDGF;VEGFA-VEGFR2 Pathway;Signaling by ERBB2;IRS-related events triggered by IGF1R 
                                               hsa03015;hsa04071;hsa04114;hsa04151;hsa04152;hsa04261;hsa04350;hsa04390;hsa04530;hsa04728;hsa04730;hsa05142;hsa05160 
                                               PPP2R1B 
                                               NA 
                                               slateblue1 
                                               16 
                                               9.12280701754386 
                                             
                                             
                                               AXIN1-mRNA 
                                               1.78 
                                               0.31 
                                               1.17 
                                               2.39 
                                               3.43 
                                               2.25 
                                               5.23 
                                               1.43e-06 
                                               1.75e-05 
                                               lm.nb 
                                               AXIN missense mutants destabilize the destruction complex, Beta-catenin phosphorylation cascade, Disassembly of the destruction complex and recruitment of AXIN to the membrane, Misspliced GSK3beta mutants stabilize beta-catenin, S33 mutants of beta-catenin aren.SQT.t phosphorylated, S37 mutants of beta-catenin aren.SQT.t phosphorylated, S45 mutants of beta-catenin aren.SQT.t phosphorylated, T41 mutants of beta-catenin aren.SQT.t phosphorylated, Truncations of AMER1 destabilize the destruction complex 
                                               SEBELIUS_2_C4636 
                                               AXIN1 
                                               mRNA 
                                               0 
                                               
                                               
                                               APC truncation mutants have impaired AXIN binding;Misspliced GSK3beta mutants stabilize beta-catenin;T41 mutants of beta-catenin aren.SQT.t phosphorylated;AXIN missense mutants destabilize the destruction complex;S45 mutants of beta-catenin aren.SQT.t phosphorylated;Disassembly of the destruction complex and recruitment of AXIN to the membrane;S37 mutants of beta-catenin aren.SQT.t phosphorylated;S33 mutants of beta-catenin aren.SQT.t phosphorylated;Beta-catenin phosphorylation cascade;Truncations of AMER1 destabilize the destruction complex 
                                               hsa04310;hsa04390;hsa04550;hsa05200;hsa05210;hsa05213;hsa05217 
                                               AXIN1 
                                               NA 
                                               slateblue1 
                                               16 
                                               5.74193548387097 
                                             
                                             
                                               PPP2CB-mRNA 
                                               -0.609 
                                               0.109 
                                               -0.822 
                                               -0.397 
                                               0.655 
                                               0.565 
                                               0.76 
                                               2.12e-06 
                                               2.55e-05 
                                               lm.nb 
                                               ARMS-mediated activation, AXIN missense mutants destabilize the destruction complex, Beta-catenin phosphorylation cascade, CTLA4 inhibitory signaling, Cyclin AB1 associated events during G2M transition, Cyclin D associated events in G1, DAP12 signaling, DARPP-32 events, Disassembly of the destruction complex and recruitment of AXIN to the membrane, Downstream signal transduction, ERKs are inactivated, FCERI mediated MAPK activation, Frs2-mediated activation, Gastrin-CREB signalling pathway via PKC and MAPK, Interleukin receptor SHC signaling, IRS-related events triggered by IGF1R, MASTL Facilitates Mitotic Progression, Misspliced GSK3beta mutants stabilize beta-catenin, Negative regulation of FGFR1 signaling, Negative regulation of FGFR2 signaling, Negative regulation of FGFR3 signaling, Negative regulation of FGFR4 signaling, Negative regulation of MAPK pathway, RAF activation, RAFMAP kinase cascade, Resolution of Sister Chromatid Cohesion, RHO GTPases Activate Formins, S33 mutants of beta-catenin aren.SQT.t phosphorylated, S37 mutants of beta-catenin aren.SQT.t phosphorylated, S45 mutants of beta-catenin aren.SQT.t phosphorylated, Signaling by ERBB2, Signaling by PDGF, Signaling by SCF-KIT, Spry regulation of FGF signaling, T41 mutants of beta-catenin aren.SQT.t phosphorylated, Truncations of AMER1 destabilize the destruction complex, VEGFA-VEGFR2 Pathway, VEGFR2 mediated cell proliferation 
                                               SEBELIUS_2_C4636 
                                               PPP2CB 
                                               mRNA 
                                               0 
                                               
                                               
                                               MASTL Facilitates Mitotic Progression;Misspliced GSK3beta mutants stabilize beta-catenin;T41 mutants of beta-catenin aren.SQT.t phosphorylated;ERKs are inactivated;Glycolysis;Negative regulation of MAPK pathway;Disassembly of the destruction complex and recruitment of AXIN to the membrane;S37 mutants of beta-catenin aren.SQT.t phosphorylated;Cyclin AB1 associated events during G2M transition;RAF activation;RHO GTPases Activate Formins;S33 mutants of beta-catenin aren.SQT.t phosphorylated;DARPP-32 events;Beta-catenin phosphorylation cascade;Spry regulation of FGF signaling;Truncations of AMER1 destabilize the destruction complex;CTLA4 inhibitory signaling;AXIN missense mutants destabilize the destruction complex;S45 mutants of beta-catenin aren.SQT.t phosphorylated;Cyclin D associated events in G1;Platelet sensitization by LDL;Resolution of Sister Chromatid Cohesion;Inhibition of replication initiation of damaged DNA by RB1E2F1;RAFMAP kinase cascade;Downstream signal transduction;VEGFR2 mediated cell proliferation;Signaling by SCF-KIT;Frs2-mediated activation;ARMS-mediated activation;Gastrin-CREB signalling pathway via PKC and MAPK;Interleukin receptor SHC signaling;FCERI mediated MAPK activation;DAP12 signaling;Signaling by PDGF;VEGFA-VEGFR2 Pathway;Signaling by ERBB2;IRS-related events triggered by IGF1R;Negative regulation of FGFR3 signaling;Negative regulation of FGFR4 signaling;Negative regulation of FGFR2 signaling;Negative regulation of FGFR1 signaling 
                                               hsa03015;hsa04071;hsa04114;hsa04151;hsa04152;hsa04261;hsa04350;hsa04390;hsa04530;hsa04728;hsa04730;hsa05142;hsa05160 
                                               PPP2CB 
                                               NA 
                                               slateblue1 
                                               16 
                                               -5.58715596330275 
                                             
                                             
                                               PPP2CA-mRNA 
                                               0.388 
                                               0.122 
                                               0.15 
                                               0.627 
                                               1.31 
                                               1.11 
                                               1.54 
                                               0.00287 
                                               0.0255 
                                               lm.nb 
                                               ARMS-mediated activation, AXIN missense mutants destabilize the destruction complex, Beta-catenin phosphorylation cascade, CTLA4 inhibitory signaling, Cyclin AB1 associated events during G2M transition, Cyclin D associated events in G1, DAP12 signaling, DARPP-32 events, Disassembly of the destruction complex and recruitment of AXIN to the membrane, Downstream signal transduction, ERKs are inactivated, FCERI mediated MAPK activation, Frs2-mediated activation, Gastrin-CREB signalling pathway via PKC and MAPK, Interleukin receptor SHC signaling, IRS-related events triggered by IGF1R, MASTL Facilitates Mitotic Progression, Negative regulation of FGFR1 signaling, Negative regulation of FGFR2 signaling, Negative regulation of FGFR3 signaling, Negative regulation of FGFR4 signaling, Negative regulation of MAPK pathway, Nonsense Mediated Decay (NMD) enhanced by the Exon Junction Complex (EJC), RAF activation, RAFMAP kinase cascade, Resolution of Sister Chromatid Cohesion, RHO GTPases Activate Formins, S33 mutants of beta-catenin aren.SQT.t phosphorylated, S37 mutants of beta-catenin aren.SQT.t phosphorylated, S45 mutants of beta-catenin aren.SQT.t phosphorylated, Signaling by ERBB2, Signaling by PDGF, Signaling by SCF-KIT, Spry regulation of FGF signaling, T41 mutants of beta-catenin aren.SQT.t phosphorylated, Truncations of AMER1 destabilize the destruction complex, VEGFA-VEGFR2 Pathway, VEGFR2 mediated cell proliferation 
                                               SEBELIUS_2_C4636 
                                               PPP2CA 
                                               mRNA 
                                               0 
                                               
                                               
                                               MASTL Facilitates Mitotic Progression;T41 mutants of beta-catenin aren.SQT.t phosphorylated;ERKs are inactivated;Glycolysis;Negative regulation of MAPK pathway;Disassembly of the destruction complex and recruitment of AXIN to the membrane;S37 mutants of beta-catenin aren.SQT.t phosphorylated;Cyclin AB1 associated events during G2M transition;RAF activation;RHO GTPases Activate Formins;S33 mutants of beta-catenin aren.SQT.t phosphorylated;DARPP-32 events;Beta-catenin phosphorylation cascade;Initiation of Nuclear Envelope Reformation;Spry regulation of FGF signaling;Truncations of AMER1 destabilize the destruction complex;CTLA4 inhibitory signaling;AXIN missense mutants destabilize the destruction complex;S45 mutants of beta-catenin aren.SQT.t phosphorylated;Nonsense Mediated Decay (NMD) enhanced by the Exon Junction Complex (EJC);Cyclin D associated events in G1;Platelet sensitization by LDL;Resolution of Sister Chromatid Cohesion;Inhibition of replication initiation of damaged DNA by RB1E2F1;RAFMAP kinase cascade;Downstream signal transduction;VEGFR2 mediated cell proliferation;Signaling by SCF-KIT;Frs2-mediated activation;ARMS-mediated activation;Gastrin-CREB signalling pathway via PKC and MAPK;Interleukin receptor SHC signaling;FCERI mediated MAPK activation;DAP12 signaling;Signaling by PDGF;VEGFA-VEGFR2 Pathway;Signaling by ERBB2;IRS-related events triggered by IGF1R;Negative regulation of FGFR3 signaling;Negative regulation of FGFR4 signaling;Negative regulation of FGFR2 signaling;Negative regulation of FGFR1 signaling 
                                               hsa03015;hsa04071;hsa04114;hsa04151;hsa04152;hsa04261;hsa04350;hsa04390;hsa04530;hsa04728;hsa04730;hsa05142;hsa05160 
                                               PPP2CA 
                                               NA 
                                               slateblue2 
                                               16 
                                               3.18032786885246 
                                             
                                             
                                               PPP2R1A-mRNA 
                                               -0.264 
                                               0.119 
                                               -0.498 
                                               -0.031 
                                               0.833 
                                               0.708 
                                               0.979 
                                               0.0326 
                                               0.253 
                                               lm.nb 
                                               ARMS-mediated activation, AXIN missense mutants destabilize the destruction complex, Beta-catenin phosphorylation cascade, CTLA4 inhibitory signaling, Cyclin AB1 associated events during G2M transition, Cyclin D associated events in G1, DAP12 signaling, DARPP-32 events, Disassembly of the destruction complex and recruitment of AXIN to the membrane, Downstream signal transduction, FCERI mediated MAPK activation, Frs2-mediated activation, Gastrin-CREB signalling pathway via PKC and MAPK, Interleukin receptor SHC signaling, IRS-related events triggered by IGF1R, MASTL Facilitates Mitotic Progression, Negative regulation of FGFR1 signaling, Negative regulation of FGFR2 signaling, Negative regulation of FGFR3 signaling, Negative regulation of FGFR4 signaling, Nonsense Mediated Decay (NMD) enhanced by the Exon Junction Complex (EJC), RAF activation, RAFMAP kinase cascade, Resolution of Sister Chromatid Cohesion, RHO GTPases Activate Formins, S33 mutants of beta-catenin aren.SQT.t phosphorylated, S37 mutants of beta-catenin aren.SQT.t phosphorylated, S45 mutants of beta-catenin aren.SQT.t phosphorylated, Signaling by ERBB2, Signaling by PDGF, Signaling by SCF-KIT, Spry regulation of FGF signaling, Truncations of AMER1 destabilize the destruction complex, VEGFA-VEGFR2 Pathway, VEGFR2 mediated cell proliferation 
                                               SEBELIUS_2_C4636 
                                               PPP2R1A 
                                               mRNA 
                                               0 
                                               
                                               
                                               MASTL Facilitates Mitotic Progression;Regulation of PLK1 Activity at G2M Transition;Disassembly of the destruction complex and recruitment of AXIN to the membrane;S37 mutants of beta-catenin aren.SQT.t phosphorylated;Cyclin AB1 associated events during G2M transition;RAF activation;RHO GTPases Activate Formins;S33 mutants of beta-catenin aren.SQT.t phosphorylated;Loss of Nlp from mitotic centrosomes;DARPP-32 events;Beta-catenin phosphorylation cascade;Initiation of Nuclear Envelope Reformation;Recruitment of mitotic centrosome proteins and complexes;Spry regulation of FGF signaling;Truncations of AMER1 destabilize the destruction complex;CTLA4 inhibitory signaling;AXIN missense mutants destabilize the destruction complex;S45 mutants of beta-catenin aren.SQT.t phosphorylated;Anchoring of the basal body to the plasma membrane;Nonsense Mediated Decay (NMD) enhanced by the Exon Junction Complex (EJC);Cyclin D associated events in G1;Platelet sensitization by LDL;Resolution of Sister Chromatid Cohesion;Inhibition of replication initiation of damaged DNA by RB1E2F1;RAFMAP kinase cascade;Downstream signal transduction;VEGFR2 mediated cell proliferation;Signaling by SCF-KIT;Frs2-mediated activation;ARMS-mediated activation;Gastrin-CREB signalling pathway via PKC and MAPK;Interleukin receptor SHC signaling;FCERI mediated MAPK activation;DAP12 signaling;Signaling by PDGF;VEGFA-VEGFR2 Pathway;Signaling by ERBB2;IRS-related events triggered by IGF1R;Negative regulation of FGFR3 signaling;Negative regulation of FGFR4 signaling;Negative regulation of FGFR2 signaling;Negative regulation of FGFR1 signaling 
                                               hsa03015;hsa04071;hsa04114;hsa04151;hsa04152;hsa04261;hsa04350;hsa04390;hsa04530;hsa04728;hsa04730;hsa05142;hsa05160 
                                               PPP2R1A 
                                               NA 
                                               slateblue4 
                                               16 
                                               -2.21848739495798 
                                             
                                             
                                               CSNK1A1-mRNA 
                                               -0.15 
                                               0.0908 
                                               -0.328 
                                               0.0285 
                                               0.902 
                                               0.797 
                                               1.02 
                                               0.108 
                                               0.792 
                                               loglinear 
                                               AXIN missense mutants destabilize the destruction complex, Beta-catenin phosphorylation cascade, Disassembly of the destruction complex and recruitment of AXIN to the membrane, Misspliced GSK3beta mutants stabilize beta-catenin, S33 mutants of beta-catenin aren.SQT.t phosphorylated, S37 mutants of beta-catenin aren.SQT.t phosphorylated, S45 mutants of beta-catenin aren.SQT.t phosphorylated, T41 mutants of beta-catenin aren.SQT.t phosphorylated 
                                               SEBELIUS_2_C4636 
                                               CSNK1A1 
                                               mRNA 
                                               0 
                                               
                                               
                                               Misspliced GSK3beta mutants stabilize beta-catenin;APC truncation mutants have impaired AXIN binding;T41 mutants of beta-catenin aren.SQT.t phosphorylated;AXIN missense mutants destabilize the destruction complex;S45 mutants of beta-catenin aren.SQT.t phosphorylated;Disassembly of the destruction complex and recruitment of AXIN to the membrane;S37 mutants of beta-catenin aren.SQT.t phosphorylated;S33 mutants of beta-catenin aren.SQT.t phosphorylated;Beta-catenin phosphorylation cascade;Degradation of GLI2 by the proteasome 
                                               hsa04310 
                                               CSNK1A1 
                                               NA 
                                               azure4 
                                               1 
                                               -1.65198237885463 
                                             
                                             
                                               APC-mRNA 
                                               -0.0616 
                                               0.125 
                                               -0.306 
                                               0.183 
                                               0.958 
                                               0.809 
                                               1.14 
                                               0.624 
                                               1 
                                               lm.nb 
                                               Apoptotic cleavage of cellular proteins, AXIN missense mutants destabilize the destruction complex, Beta-catenin phosphorylation cascade, Deactivation of the beta-catenin transactivating complex, Disassembly of the destruction complex and recruitment of AXIN to the membrane, Misspliced GSK3beta mutants stabilize beta-catenin, S33 mutants of beta-catenin aren.SQT.t phosphorylated, S37 mutants of beta-catenin aren.SQT.t phosphorylated, S45 mutants of beta-catenin aren.SQT.t phosphorylated, T41 mutants of beta-catenin aren.SQT.t phosphorylated, Truncations of AMER1 destabilize the destruction complex 
                                               SEBELIUS_2_C4636 
                                               APC 
                                               mRNA 
                                               0 
                                               
                                               
                                               Apoptotic cleavage of cellular proteins;Misspliced GSK3beta mutants stabilize beta-catenin;APC truncation mutants have impaired AXIN binding;T41 mutants of beta-catenin aren.SQT.t phosphorylated;AXIN missense mutants destabilize the destruction complex;S45 mutants of beta-catenin aren.SQT.t phosphorylated;Deactivation of the beta-catenin transactivating complex;Disassembly of the destruction complex and recruitment of AXIN to the membrane;S37 mutants of beta-catenin aren.SQT.t phosphorylated;S33 mutants of beta-catenin aren.SQT.t phosphorylated;Beta-catenin phosphorylation cascade;Truncations of AMER1 destabilize the destruction complex 
                                               hsa04310;hsa04390;hsa04550;hsa04810;hsa05166;hsa05200;hsa05206;hsa05210;hsa05213;hsa05217 
                                               APC 
                                               NA 
                                               azure4 
                                               1 
                                               -0.4928 
                                             
                                           
                                           DE Results - Beta-catenin phosphorylation cascade genes - Line: HEPG2 vs.Fibro  Table displaying each sample's global significance scores and directed global significance scores as defined in the heatmaps above. The global significance score is calculated as the square root of the mean squared t-statistic for the genes in a gene set, with t-statistics coming from the linear regression underlying our differential expression analysis. The directed global significance score is calculated as the square root of the mean signed squared t-statistic for the genes in a gene set, with t-statistics coming from the linear regression underlying our differential expression analysis.   
                                      
                                       
                                  
                                
                                
                                  
                                       
                                      
                                           Volcano Plot: Line: HT29 vs.Fibro   More Plot Information                Volcano Plot: Line: HT29 vs.Fibro  Volcano plot displaying each gene's -log10(p-value) and log2 fold change for the selected covariate. Highly statistically significant genes fall at the top of the plot, and highly differentially expressed genes fall to either side. Genes within the selected gene set are highlighted in orange. Horizontal lines indicate various False Discovery Rate (FDR) thresholds.   
                                           DE Results - Beta-catenin phosphorylation cascade genes - Line: HT29 vs.Fibro   More Plot Information      Download CSV Data     
                                           
                                             
                                             
                                           
                                           
                                             
                                               Probe Label 
                                               Log2 fold change 
                                               std error (log2) 
                                               Lower confidence limit (log2) 
                                               Upper confidence limit (log2) 
                                               Linear fold change 
                                               Lower confidence limit (linear) 
                                               Upper confidence limit (linear) 
                                               P-value 
                                               BY.p.value 
                                               method 
                                               Gene.sets 
                                               Codeset.Name 
                                               Probe.Label 
                                               Analyte.Type 
                                               Is.Control 
                                               Control.Type 
                                               Related.Probes 
                                               Probe.Annotation 
                                               KEGG.Pathways 
                                               Official.Gene.Name 
                                               Control.Conc 
                                               volcanocol 
                                               volcanopch 
                                               tstats.all 
                                             
                                           
                                           
                                             
                                               PPP2CB-mRNA 
                                               -2.03 
                                               0.103 
                                               -2.23 
                                               -1.83 
                                               0.245 
                                               0.213 
                                               0.281 
                                               3.08e-21 
                                               1.56e-19 
                                               lm.nb 
                                               ARMS-mediated activation, AXIN missense mutants destabilize the destruction complex, Beta-catenin phosphorylation cascade, CTLA4 inhibitory signaling, Cyclin AB1 associated events during G2M transition, Cyclin D associated events in G1, DAP12 signaling, DARPP-32 events, Disassembly of the destruction complex and recruitment of AXIN to the membrane, Downstream signal transduction, ERKs are inactivated, FCERI mediated MAPK activation, Frs2-mediated activation, Gastrin-CREB signalling pathway via PKC and MAPK, Interleukin receptor SHC signaling, IRS-related events triggered by IGF1R, MASTL Facilitates Mitotic Progression, Misspliced GSK3beta mutants stabilize beta-catenin, Negative regulation of FGFR1 signaling, Negative regulation of FGFR2 signaling, Negative regulation of FGFR3 signaling, Negative regulation of FGFR4 signaling, Negative regulation of MAPK pathway, RAF activation, RAFMAP kinase cascade, Resolution of Sister Chromatid Cohesion, RHO GTPases Activate Formins, S33 mutants of beta-catenin aren.SQT.t phosphorylated, S37 mutants of beta-catenin aren.SQT.t phosphorylated, S45 mutants of beta-catenin aren.SQT.t phosphorylated, Signaling by ERBB2, Signaling by PDGF, Signaling by SCF-KIT, Spry regulation of FGF signaling, T41 mutants of beta-catenin aren.SQT.t phosphorylated, Truncations of AMER1 destabilize the destruction complex, VEGFA-VEGFR2 Pathway, VEGFR2 mediated cell proliferation 
                                               SEBELIUS_2_C4636 
                                               PPP2CB 
                                               mRNA 
                                               0 
                                               
                                               
                                               MASTL Facilitates Mitotic Progression;Misspliced GSK3beta mutants stabilize beta-catenin;T41 mutants of beta-catenin aren.SQT.t phosphorylated;ERKs are inactivated;Glycolysis;Negative regulation of MAPK pathway;Disassembly of the destruction complex and recruitment of AXIN to the membrane;S37 mutants of beta-catenin aren.SQT.t phosphorylated;Cyclin AB1 associated events during G2M transition;RAF activation;RHO GTPases Activate Formins;S33 mutants of beta-catenin aren.SQT.t phosphorylated;DARPP-32 events;Beta-catenin phosphorylation cascade;Spry regulation of FGF signaling;Truncations of AMER1 destabilize the destruction complex;CTLA4 inhibitory signaling;AXIN missense mutants destabilize the destruction complex;S45 mutants of beta-catenin aren.SQT.t phosphorylated;Cyclin D associated events in G1;Platelet sensitization by LDL;Resolution of Sister Chromatid Cohesion;Inhibition of replication initiation of damaged DNA by RB1E2F1;RAFMAP kinase cascade;Downstream signal transduction;VEGFR2 mediated cell proliferation;Signaling by SCF-KIT;Frs2-mediated activation;ARMS-mediated activation;Gastrin-CREB signalling pathway via PKC and MAPK;Interleukin receptor SHC signaling;FCERI mediated MAPK activation;DAP12 signaling;Signaling by PDGF;VEGFA-VEGFR2 Pathway;Signaling by ERBB2;IRS-related events triggered by IGF1R;Negative regulation of FGFR3 signaling;Negative regulation of FGFR4 signaling;Negative regulation of FGFR2 signaling;Negative regulation of FGFR1 signaling 
                                               hsa03015;hsa04071;hsa04114;hsa04151;hsa04152;hsa04261;hsa04350;hsa04390;hsa04530;hsa04728;hsa04730;hsa05142;hsa05160 
                                               PPP2CB 
                                               NA 
                                               slateblue1 
                                               16 
                                               -19.7087378640777 
                                             
                                             
                                               PPP2R1B-mRNA 
                                               0.978 
                                               0.107 
                                               0.769 
                                               1.19 
                                               1.97 
                                               1.7 
                                               2.28 
                                               5.09e-11 
                                               7.3e-10 
                                               lm.nb 
                                               ARMS-mediated activation, AXIN missense mutants destabilize the destruction complex, Beta-catenin phosphorylation cascade, CTLA4 inhibitory signaling, Cyclin AB1 associated events during G2M transition, Cyclin D associated events in G1, DAP12 signaling, DARPP-32 events, Disassembly of the destruction complex and recruitment of AXIN to the membrane, Downstream signal transduction, ERKs are inactivated, FCERI mediated MAPK activation, Frs2-mediated activation, Gastrin-CREB signalling pathway via PKC and MAPK, Interleukin receptor SHC signaling, IRS-related events triggered by IGF1R, MASTL Facilitates Mitotic Progression, Misspliced GSK3beta mutants stabilize beta-catenin, Negative regulation of MAPK pathway, RAF activation, RAFMAP kinase cascade, Resolution of Sister Chromatid Cohesion, RHO GTPases Activate Formins, S33 mutants of beta-catenin aren.SQT.t phosphorylated, S37 mutants of beta-catenin aren.SQT.t phosphorylated, S45 mutants of beta-catenin aren.SQT.t phosphorylated, Signaling by ERBB2, Signaling by PDGF, Signaling by SCF-KIT, T41 mutants of beta-catenin aren.SQT.t phosphorylated, Truncations of AMER1 destabilize the destruction complex, VEGFA-VEGFR2 Pathway, VEGFR2 mediated cell proliferation 
                                               SEBELIUS_2_C4636 
                                               PPP2R1B 
                                               mRNA 
                                               0 
                                               
                                               
                                               MASTL Facilitates Mitotic Progression;Misspliced GSK3beta mutants stabilize beta-catenin;T41 mutants of beta-catenin aren.SQT.t phosphorylated;ERKs are inactivated;Glycolysis;Negative regulation of MAPK pathway;Disassembly of the destruction complex and recruitment of AXIN to the membrane;S37 mutants of beta-catenin aren.SQT.t phosphorylated;Cyclin AB1 associated events during G2M transition;RAF activation;RHO GTPases Activate Formins;S33 mutants of beta-catenin aren.SQT.t phosphorylated;DARPP-32 events;Beta-catenin phosphorylation cascade;Truncations of AMER1 destabilize the destruction complex;CTLA4 inhibitory signaling;AXIN missense mutants destabilize the destruction complex;S45 mutants of beta-catenin aren.SQT.t phosphorylated;Cyclin D associated events in G1;Platelet sensitization by LDL;Resolution of Sister Chromatid Cohesion;Inhibition of replication initiation of damaged DNA by RB1E2F1;RAFMAP kinase cascade;Downstream signal transduction;VEGFR2 mediated cell proliferation;Signaling by SCF-KIT;Frs2-mediated activation;ARMS-mediated activation;Gastrin-CREB signalling pathway via PKC and MAPK;Interleukin receptor SHC signaling;FCERI mediated MAPK activation;DAP12 signaling;Signaling by PDGF;VEGFA-VEGFR2 Pathway;Signaling by ERBB2;IRS-related events triggered by IGF1R 
                                               hsa03015;hsa04071;hsa04114;hsa04151;hsa04152;hsa04261;hsa04350;hsa04390;hsa04530;hsa04728;hsa04730;hsa05142;hsa05160 
                                               PPP2R1B 
                                               NA 
                                               slateblue1 
                                               16 
                                               9.14018691588785 
                                             
                                             
                                               AXIN1-mRNA 
                                               1.54 
                                               0.293 
                                               0.963 
                                               2.11 
                                               2.9 
                                               1.95 
                                               4.32 
                                               6.64e-06 
                                               6.63e-05 
                                               lm.nb 
                                               AXIN missense mutants destabilize the destruction complex, Beta-catenin phosphorylation cascade, Disassembly of the destruction complex and recruitment of AXIN to the membrane, Misspliced GSK3beta mutants stabilize beta-catenin, S33 mutants of beta-catenin aren.SQT.t phosphorylated, S37 mutants of beta-catenin aren.SQT.t phosphorylated, S45 mutants of beta-catenin aren.SQT.t phosphorylated, T41 mutants of beta-catenin aren.SQT.t phosphorylated, Truncations of AMER1 destabilize the destruction complex 
                                               SEBELIUS_2_C4636 
                                               AXIN1 
                                               mRNA 
                                               0 
                                               
                                               
                                               APC truncation mutants have impaired AXIN binding;Misspliced GSK3beta mutants stabilize beta-catenin;T41 mutants of beta-catenin aren.SQT.t phosphorylated;AXIN missense mutants destabilize the destruction complex;S45 mutants of beta-catenin aren.SQT.t phosphorylated;Disassembly of the destruction complex and recruitment of AXIN to the membrane;S37 mutants of beta-catenin aren.SQT.t phosphorylated;S33 mutants of beta-catenin aren.SQT.t phosphorylated;Beta-catenin phosphorylation cascade;Truncations of AMER1 destabilize the destruction complex 
                                               hsa04310;hsa04390;hsa04550;hsa05200;hsa05210;hsa05213;hsa05217 
                                               AXIN1 
                                               NA 
                                               slateblue1 
                                               16 
                                               5.25597269624573 
                                             
                                             
                                               PPP2CA-mRNA 
                                               0.495 
                                               0.115 
                                               0.271 
                                               0.719 
                                               1.41 
                                               1.21 
                                               1.65 
                                               0.000112 
                                               0.00104 
                                               lm.nb 
                                               ARMS-mediated activation, AXIN missense mutants destabilize the destruction complex, Beta-catenin phosphorylation cascade, CTLA4 inhibitory signaling, Cyclin AB1 associated events during G2M transition, Cyclin D associated events in G1, DAP12 signaling, DARPP-32 events, Disassembly of the destruction complex and recruitment of AXIN to the membrane, Downstream signal transduction, ERKs are inactivated, FCERI mediated MAPK activation, Frs2-mediated activation, Gastrin-CREB signalling pathway via PKC and MAPK, Interleukin receptor SHC signaling, IRS-related events triggered by IGF1R, MASTL Facilitates Mitotic Progression, Negative regulation of FGFR1 signaling, Negative regulation of FGFR2 signaling, Negative regulation of FGFR3 signaling, Negative regulation of FGFR4 signaling, Negative regulation of MAPK pathway, Nonsense Mediated Decay (NMD) enhanced by the Exon Junction Complex (EJC), RAF activation, RAFMAP kinase cascade, Resolution of Sister Chromatid Cohesion, RHO GTPases Activate Formins, S33 mutants of beta-catenin aren.SQT.t phosphorylated, S37 mutants of beta-catenin aren.SQT.t phosphorylated, S45 mutants of beta-catenin aren.SQT.t phosphorylated, Signaling by ERBB2, Signaling by PDGF, Signaling by SCF-KIT, Spry regulation of FGF signaling, T41 mutants of beta-catenin aren.SQT.t phosphorylated, Truncations of AMER1 destabilize the destruction complex, VEGFA-VEGFR2 Pathway, VEGFR2 mediated cell proliferation 
                                               SEBELIUS_2_C4636 
                                               PPP2CA 
                                               mRNA 
                                               0 
                                               
                                               
                                               MASTL Facilitates Mitotic Progression;T41 mutants of beta-catenin aren.SQT.t phosphorylated;ERKs are inactivated;Glycolysis;Negative regulation of MAPK pathway;Disassembly of the destruction complex and recruitment of AXIN to the membrane;S37 mutants of beta-catenin aren.SQT.t phosphorylated;Cyclin AB1 associated events during G2M transition;RAF activation;RHO GTPases Activate Formins;S33 mutants of beta-catenin aren.SQT.t phosphorylated;DARPP-32 events;Beta-catenin phosphorylation cascade;Initiation of Nuclear Envelope Reformation;Spry regulation of FGF signaling;Truncations of AMER1 destabilize the destruction complex;CTLA4 inhibitory signaling;AXIN missense mutants destabilize the destruction complex;S45 mutants of beta-catenin aren.SQT.t phosphorylated;Nonsense Mediated Decay (NMD) enhanced by the Exon Junction Complex (EJC);Cyclin D associated events in G1;Platelet sensitization by LDL;Resolution of Sister Chromatid Cohesion;Inhibition of replication initiation of damaged DNA by RB1E2F1;RAFMAP kinase cascade;Downstream signal transduction;VEGFR2 mediated cell proliferation;Signaling by SCF-KIT;Frs2-mediated activation;ARMS-mediated activation;Gastrin-CREB signalling pathway via PKC and MAPK;Interleukin receptor SHC signaling;FCERI mediated MAPK activation;DAP12 signaling;Signaling by PDGF;VEGFA-VEGFR2 Pathway;Signaling by ERBB2;IRS-related events triggered by IGF1R;Negative regulation of FGFR3 signaling;Negative regulation of FGFR4 signaling;Negative regulation of FGFR2 signaling;Negative regulation of FGFR1 signaling 
                                               hsa03015;hsa04071;hsa04114;hsa04151;hsa04152;hsa04261;hsa04350;hsa04390;hsa04530;hsa04728;hsa04730;hsa05142;hsa05160 
                                               PPP2CA 
                                               NA 
                                               slateblue1 
                                               16 
                                               4.30434782608696 
                                             
                                             
                                               PPP2R1A-mRNA 
                                               0.449 
                                               0.112 
                                               0.229 
                                               0.669 
                                               1.37 
                                               1.17 
                                               1.59 
                                               0.000287 
                                               0.00259 
                                               lm.nb 
                                               ARMS-mediated activation, AXIN missense mutants destabilize the destruction complex, Beta-catenin phosphorylation cascade, CTLA4 inhibitory signaling, Cyclin AB1 associated events during G2M transition, Cyclin D associated events in G1, DAP12 signaling, DARPP-32 events, Disassembly of the destruction complex and recruitment of AXIN to the membrane, Downstream signal transduction, FCERI mediated MAPK activation, Frs2-mediated activation, Gastrin-CREB signalling pathway via PKC and MAPK, Interleukin receptor SHC signaling, IRS-related events triggered by IGF1R, MASTL Facilitates Mitotic Progression, Negative regulation of FGFR1 signaling, Negative regulation of FGFR2 signaling, Negative regulation of FGFR3 signaling, Negative regulation of FGFR4 signaling, Nonsense Mediated Decay (NMD) enhanced by the Exon Junction Complex (EJC), RAF activation, RAFMAP kinase cascade, Resolution of Sister Chromatid Cohesion, RHO GTPases Activate Formins, S33 mutants of beta-catenin aren.SQT.t phosphorylated, S37 mutants of beta-catenin aren.SQT.t phosphorylated, S45 mutants of beta-catenin aren.SQT.t phosphorylated, Signaling by ERBB2, Signaling by PDGF, Signaling by SCF-KIT, Spry regulation of FGF signaling, Truncations of AMER1 destabilize the destruction complex, VEGFA-VEGFR2 Pathway, VEGFR2 mediated cell proliferation 
                                               SEBELIUS_2_C4636 
                                               PPP2R1A 
                                               mRNA 
                                               0 
                                               
                                               
                                               MASTL Facilitates Mitotic Progression;Regulation of PLK1 Activity at G2M Transition;Disassembly of the destruction complex and recruitment of AXIN to the membrane;S37 mutants of beta-catenin aren.SQT.t phosphorylated;Cyclin AB1 associated events during G2M transition;RAF activation;RHO GTPases Activate Formins;S33 mutants of beta-catenin aren.SQT.t phosphorylated;Loss of Nlp from mitotic centrosomes;DARPP-32 events;Beta-catenin phosphorylation cascade;Initiation of Nuclear Envelope Reformation;Recruitment of mitotic centrosome proteins and complexes;Spry regulation of FGF signaling;Truncations of AMER1 destabilize the destruction complex;CTLA4 inhibitory signaling;AXIN missense mutants destabilize the destruction complex;S45 mutants of beta-catenin aren.SQT.t phosphorylated;Anchoring of the basal body to the plasma membrane;Nonsense Mediated Decay (NMD) enhanced by the Exon Junction Complex (EJC);Cyclin D associated events in G1;Platelet sensitization by LDL;Resolution of Sister Chromatid Cohesion;Inhibition of replication initiation of damaged DNA by RB1E2F1;RAFMAP kinase cascade;Downstream signal transduction;VEGFR2 mediated cell proliferation;Signaling by SCF-KIT;Frs2-mediated activation;ARMS-mediated activation;Gastrin-CREB signalling pathway via PKC and MAPK;Interleukin receptor SHC signaling;FCERI mediated MAPK activation;DAP12 signaling;Signaling by PDGF;VEGFA-VEGFR2 Pathway;Signaling by ERBB2;IRS-related events triggered by IGF1R;Negative regulation of FGFR3 signaling;Negative regulation of FGFR4 signaling;Negative regulation of FGFR2 signaling;Negative regulation of FGFR1 signaling 
                                               hsa03015;hsa04071;hsa04114;hsa04151;hsa04152;hsa04261;hsa04350;hsa04390;hsa04530;hsa04728;hsa04730;hsa05142;hsa05160 
                                               PPP2R1A 
                                               NA 
                                               slateblue1 
                                               16 
                                               4.00892857142857 
                                             
                                             
                                               APC-mRNA 
                                               0.329 
                                               0.117 
                                               0.1 
                                               0.558 
                                               1.26 
                                               1.07 
                                               1.47 
                                               0.00766 
                                               0.0598 
                                               lm.nb 
                                               Apoptotic cleavage of cellular proteins, AXIN missense mutants destabilize the destruction complex, Beta-catenin phosphorylation cascade, Deactivation of the beta-catenin transactivating complex, Disassembly of the destruction complex and recruitment of AXIN to the membrane, Misspliced GSK3beta mutants stabilize beta-catenin, S33 mutants of beta-catenin aren.SQT.t phosphorylated, S37 mutants of beta-catenin aren.SQT.t phosphorylated, S45 mutants of beta-catenin aren.SQT.t phosphorylated, T41 mutants of beta-catenin aren.SQT.t phosphorylated, Truncations of AMER1 destabilize the destruction complex 
                                               SEBELIUS_2_C4636 
                                               APC 
                                               mRNA 
                                               0 
                                               
                                               
                                               Apoptotic cleavage of cellular proteins;Misspliced GSK3beta mutants stabilize beta-catenin;APC truncation mutants have impaired AXIN binding;T41 mutants of beta-catenin aren.SQT.t phosphorylated;AXIN missense mutants destabilize the destruction complex;S45 mutants of beta-catenin aren.SQT.t phosphorylated;Deactivation of the beta-catenin transactivating complex;Disassembly of the destruction complex and recruitment of AXIN to the membrane;S37 mutants of beta-catenin aren.SQT.t phosphorylated;S33 mutants of beta-catenin aren.SQT.t phosphorylated;Beta-catenin phosphorylation cascade;Truncations of AMER1 destabilize the destruction complex 
                                               hsa04310;hsa04390;hsa04550;hsa04810;hsa05166;hsa05200;hsa05206;hsa05210;hsa05213;hsa05217 
                                               APC 
                                               NA 
                                               slateblue3 
                                               16 
                                               2.81196581196581 
                                             
                                             
                                               CSNK1A1-mRNA 
                                               0.171 
                                               0.0856 
                                               0.00324 
                                               0.339 
                                               1.13 
                                               1 
                                               1.26 
                                               0.0531 
                                               0.389 
                                               loglinear 
                                               AXIN missense mutants destabilize the destruction complex, Beta-catenin phosphorylation cascade, Disassembly of the destruction complex and recruitment of AXIN to the membrane, Misspliced GSK3beta mutants stabilize beta-catenin, S33 mutants of beta-catenin aren.SQT.t phosphorylated, S37 mutants of beta-catenin aren.SQT.t phosphorylated, S45 mutants of beta-catenin aren.SQT.t phosphorylated, T41 mutants of beta-catenin aren.SQT.t phosphorylated 
                                               SEBELIUS_2_C4636 
                                               CSNK1A1 
                                               mRNA 
                                               0 
                                               
                                               
                                               Misspliced GSK3beta mutants stabilize beta-catenin;APC truncation mutants have impaired AXIN binding;T41 mutants of beta-catenin aren.SQT.t phosphorylated;AXIN missense mutants destabilize the destruction complex;S45 mutants of beta-catenin aren.SQT.t phosphorylated;Disassembly of the destruction complex and recruitment of AXIN to the membrane;S37 mutants of beta-catenin aren.SQT.t phosphorylated;S33 mutants of beta-catenin aren.SQT.t phosphorylated;Beta-catenin phosphorylation cascade;Degradation of GLI2 by the proteasome 
                                               hsa04310 
                                               CSNK1A1 
                                               NA 
                                               slateblue4 
                                               16 
                                               1.99766355140187 
                                             
                                             
                                               CTNNB1-mRNA 
                                               0.244 
                                               0.168 
                                               -0.0856 
                                               0.573 
                                               1.18 
                                               0.942 
                                               1.49 
                                               0.155 
                                               1 
                                               lm.nb 
                                               Apoptotic cleavage of cellular proteins, Beta-catenin phosphorylation cascade, Deactivation of the beta-catenin transactivating complex, Disassembly of the destruction complex and recruitment of AXIN to the membrane, Formation of the beta-catenin.CL.TCF transactivating complex, Misspliced GSK3beta mutants stabilize beta-catenin, Repression of WNT target genes, RHO GTPases activate IQGAPs, S37 mutants of beta-catenin aren.SQT.t phosphorylated, S45 mutants of beta-catenin aren.SQT.t phosphorylated, T41 mutants of beta-catenin aren.SQT.t phosphorylated, VEGFA-VEGFR2 Pathway, VEGFR2 mediated vascular permeability 
                                               SEBELIUS_2_C4636 
                                               CTNNB1 
                                               mRNA 
                                               0 
                                               
                                               
                                               Disassembly of the destruction complex and recruitment of AXIN to the membrane;S37 mutants of beta-catenin aren.SQT.t phosphorylated;T41 mutants of beta-catenin aren.SQT.t phosphorylated;S45 mutants of beta-catenin aren.SQT.t phosphorylated;Binding of TCFLEF.CL.CTNNB1 to target gene promoters;RHO GTPases activate IQGAPs;Deactivation of the beta-catenin transactivating complex;Repression of WNT target genes;Apoptotic cleavage of cell adhesion  proteins;Adherens junctions interactions;Misspliced GSK3beta mutants stabilize beta-catenin;VEGFR2 mediated vascular permeability;Beta-catenin phosphorylation cascade;Synthesis# secretion# and inactivation of Glucagon-like Peptide-1 (GLP-1);Formation of the beta-catenin.CL.TCF transactivating complex;Apoptotic cleavage of cellular proteins;VEGFA-VEGFR2 Pathway 
                                               hsa04015;hsa04310;hsa04390;hsa04510;hsa04520;hsa04530;hsa04550;hsa04670;hsa04916;hsa04919;hsa05100;hsa05130;hsa05166;hsa05200;hsa05205;hsa05210;hsa05213;hsa05215;hsa05216;hsa05217;hsa05412 
                                               CTNNB1 
                                               NA 
                                               azure4 
                                               1 
                                               1.45238095238095 
                                             
                                           
                                           DE Results - Beta-catenin phosphorylation cascade genes - Line: HT29 vs.Fibro  Table displaying each sample's global significance scores and directed global significance scores as defined in the heatmaps above. The global significance score is calculated as the square root of the mean squared t-statistic for the genes in a gene set, with t-statistics coming from the linear regression underlying our differential expression analysis. The directed global significance score is calculated as the square root of the mean signed squared t-statistic for the genes in a gene set, with t-statistics coming from the linear regression underlying our differential expression analysis.   
                                      
                                       
                                  
                                
                              
                            
                           
                            
                          
                           
                      
                    
                    
                      
                           
                          
                               Day    Line     
                              
                                  Day  
                              
                               
                              
                                  Line: differential expression in Caco2 vs. baseline of Fibro  
                                  Line: differential expression in HEPG2 vs. baseline of Fibro  
                                  Line: differential expression in HT29 vs. baseline of Fibro  
                              
                              
                            
                              
                                
                                  
                                       
                                      
                                           Volcano Plot: Day   More Plot Information                Volcano Plot: Day  Volcano plot displaying each gene's -log10(p-value) and log2 fold change for the selected covariate. Highly statistically significant genes fall at the top of the plot, and highly differentially expressed genes fall to either side. Genes within the selected gene set are highlighted in orange. Horizontal lines indicate various False Discovery Rate (FDR) thresholds.   
                                           DE Results - CD209 (DC-SIGN) signaling genes - Day   More Plot Information      Download CSV Data     
                                           
                                             
                                             
                                           
                                           
                                             
                                               Probe Label 
                                               Log2 fold change 
                                               std error (log2) 
                                               Lower confidence limit (log2) 
                                               Upper confidence limit (log2) 
                                               Linear fold change 
                                               Lower confidence limit (linear) 
                                               Upper confidence limit (linear) 
                                               P-value 
                                               BY.p.value 
                                               method 
                                               Gene.sets 
                                               Codeset.Name 
                                               Probe.Label 
                                               Analyte.Type 
                                               Is.Control 
                                               Control.Type 
                                               Related.Probes 
                                               Probe.Annotation 
                                               KEGG.Pathways 
                                               Official.Gene.Name 
                                               Control.Conc 
                                               volcanocol 
                                               volcanopch 
                                               tstats.all 
                                             
                                           
                                           
                                             
                                               PAK1-mRNA 
                                               0.128 
                                               0.0211 
                                               0.0869 
                                               0.169 
                                               1.09 
                                               1.06 
                                               1.12 
                                               4.84e-07 
                                               6.25e-05 
                                               lm.nb 
                                               Activation of Rac, CD209 (DC-SIGN) signaling, CD28 co-stimulation, CD28 dependent Vav1 pathway, EPHB-mediated forward signaling, Ephrin signaling, FCERI mediated MAPK activation, MAPK6MAPK4 signaling, Regulation of actin dynamics for phagocytic cup formation, RHO GTPases activate PAKs, Sema3A PAK dependent Axon repulsion, VEGFA-VEGFR2 Pathway, VEGFR2 mediated vascular permeability 
                                               SEBELIUS_2_C4636 
                                               PAK1 
                                               mRNA 
                                               0 
                                               
                                               
                                               EPHB-mediated forward signaling;Ephrin signaling;MAPK6MAPK4 signaling;DSCAM interactions;Activation of Rac;VEGFR2 mediated vascular permeability;RHO GTPases activate PAKs;Sema3A PAK dependent Axon repulsion;FCERI mediated MAPK activation;Regulation of actin dynamics for phagocytic cup formation;CD209 (DC-SIGN) signaling;CD28 dependent Vav1 pathway;VEGFA-VEGFR2 Pathway;CD28 co-stimulation 
                                               hsa04010;hsa04012;hsa04014;hsa04024;hsa04062;hsa04360;hsa04510;hsa04650;hsa04660;hsa04666;hsa04810;hsa05120;hsa05205;hsa05211 
                                               PAK1 
                                               NA 
                                               slateblue1 
                                               16 
                                               6.06635071090047 
                                             
                                             
                                               PAK2-mRNA 
                                               -0.0398 
                                               0.00994 
                                               -0.0593 
                                               -0.0203 
                                               0.973 
                                               0.96 
                                               0.986 
                                               0.000289 
                                               0.00836 
                                               loglinear 
                                               Activation of Rac, CD209 (DC-SIGN) signaling, CD28 co-stimulation, CD28 dependent Vav1 pathway, Ephrin signaling, RHO GTPases activate PAKs, Sema3A PAK dependent Axon repulsion, VEGFA-VEGFR2 Pathway, VEGFR2 mediated vascular permeability 
                                               SEBELIUS_2_C4636 
                                               PAK2 
                                               mRNA 
                                               0 
                                               
                                               
                                               Nef and signal transduction;CD28 dependent Vav1 pathway;Stimulation of the cell death response by PAK-2p34;Ephrin signaling;Activation of Rac;RHO GTPases activate PAKs;VEGFR2 mediated vascular permeability;Sema3A PAK dependent Axon repulsion;CD209 (DC-SIGN) signaling;VEGFA-VEGFR2 Pathway;CD28 co-stimulation 
                                               hsa04010;hsa04012;hsa04014;hsa04360;hsa04510;hsa04660;hsa04810;hsa05211 
                                               PAK2 
                                               NA 
                                               slateblue1 
                                               16 
                                               -4.00402414486922 
                                             
                                             
                                               RAF1-mRNA 
                                               0.0287 
                                               0.011 
                                               0.00708 
                                               0.0503 
                                               1.02 
                                               1 
                                               1.04 
                                               0.0132 
                                               0.177 
                                               loglinear 
                                               ARMS-mediated activation, CD209 (DC-SIGN) signaling, CREB phosphorylation through the activation of Ras, DAP12 signaling, Downstream signal transduction, FCERI mediated MAPK activation, Frs2-mediated activation, Gastrin-CREB signalling pathway via PKC and MAPK, Interleukin receptor SHC signaling, IRS-related events triggered by IGF1R, Negative feedback regulation of MAPK pathway, Negative regulation of MAPK pathway, RAF activation, RAFMAP kinase cascade, Signaling by ERBB2, Signaling by PDGF, Signaling by SCF-KIT, VEGFA-VEGFR2 Pathway, VEGFR2 mediated cell proliferation 
                                               SEBELIUS_2_C4636 
                                               RAF1 
                                               mRNA 
                                               0 
                                               
                                               
                                               RAF activation;Stimuli-sensing channels;CREB phosphorylation through the activation of Ras;Negative feedback regulation of MAPK pathway;CD209 (DC-SIGN) signaling;RAFMAP kinase cascade;Downstream signal transduction;VEGFR2 mediated cell proliferation;Signaling by SCF-KIT;Frs2-mediated activation;ARMS-mediated activation;Gastrin-CREB signalling pathway via PKC and MAPK;Interleukin receptor SHC signaling;FCERI mediated MAPK activation;DAP12 signaling;Signaling by PDGF;VEGFA-VEGFR2 Pathway;Signaling by ERBB2;IRS-related events triggered by IGF1R;Negative regulation of MAPK pathway 
                                               hsa04010;hsa04012;hsa04014;hsa04015;hsa04022;hsa04024;hsa04062;hsa04068;hsa04071;hsa04072;hsa04151;hsa04270;hsa04370;hsa04510;hsa04540;hsa04550;hsa04630;hsa04650;hsa04660;hsa04662;hsa04664;hsa04666;hsa04720;hsa04722;hsa04726;hsa04730;hsa04810;hsa04910;hsa04912;hsa04914;hsa04915;hsa04916;hsa04917;hsa04919;hsa04921;hsa05034;hsa05152;hsa05160;hsa05161;hsa05164;hsa05200;hsa05205;hsa05206;hsa05210;hsa05211;hsa05212;hsa05213;hsa05214;hsa05215;hsa05218;hsa05219;hsa05220;hsa05221;hsa05223;hsa05230;hsa05231 
                                               RAF1 
                                               NA 
                                               slateblue4 
                                               16 
                                               2.60909090909091 
                                             
                                             
                                               NRAS-mRNA 
                                               -0.03 
                                               0.0116 
                                               -0.0527 
                                               -0.00738 
                                               0.979 
                                               0.964 
                                               0.995 
                                               0.0134 
                                               0.177 
                                               lm.nb 
                                               ARMS-mediated activation, CD209 (DC-SIGN) signaling, Cell surface interactions at the vascular wall, Constitutive Signaling by EGFRvIII, DAP12 signaling, Downstream signal transduction, FCERI mediated MAPK activation, Frs2-mediated activation, Gastrin-CREB signalling pathway via PKC and MAPK, Interleukin receptor SHC signaling, IRS-related events triggered by IGF1R, MAP2K and MAPK activation, RAFMAP kinase cascade, SHC-mediated cascade.CL.FGFR1, SHC-related events triggered by IGF1R, Signaling by ERBB2, Signaling by FGFR1 in disease, Signaling by FGFR2 in disease, Signaling by FGFR3 in disease, Signaling by FGFR4 in disease, Signaling by PDGF, Signaling by SCF-KIT, Tie2 Signaling, VEGFA-VEGFR2 Pathway, VEGFR2 mediated cell proliferation 
                                               SEBELIUS_2_C4636 
                                               NRAS 
                                               mRNA 
                                               0 
                                               
                                               
                                               Signaling by FGFR2 in disease;Regulation of RAS by GAPs;SHC-mediated cascade.CL.FGFR3;Signaling by FGFR3 in disease;MAP2K and MAPK activation;SHC-related events triggered by IGF1R;Signaling by FGFR4 in disease;SHC-mediated cascade.CL.FGFR1;Tie2 Signaling;Signaling by FGFR1 in disease;Activation of RAS in B cells;SHC-mediated cascade.CL.FGFR4;Constitutive Signaling by EGFRvIII;CD209 (DC-SIGN) signaling;FCERI mediated MAPK activation;RAFMAP kinase cascade;Downstream signal transduction;VEGFR2 mediated cell proliferation;Signaling by SCF-KIT;Frs2-mediated activation;ARMS-mediated activation;Gastrin-CREB signalling pathway via PKC and MAPK;Interleukin receptor SHC signaling;DAP12 signaling;Signaling by PDGF;VEGFA-VEGFR2 Pathway;Signaling by ERBB2;IRS-related events triggered by IGF1R;Cell surface interactions at the vascular wall 
                                               hsa04010;hsa04012;hsa04014;hsa04015;hsa04062;hsa04068;hsa04071;hsa04072;hsa04151;hsa04211;hsa04213;hsa04360;hsa04370;hsa04530;hsa04540;hsa04550;hsa04650;hsa04660;hsa04662;hsa04664;hsa04720;hsa04722;hsa04725;hsa04726;hsa04730;hsa04810;hsa04910;hsa04912;hsa04915;hsa04916;hsa04917;hsa04919;hsa04921;hsa04933;hsa05034;hsa05160;hsa05161;hsa05166;hsa05200;hsa05203;hsa05205;hsa05206;hsa05211;hsa05213;hsa05214;hsa05215;hsa05216;hsa05218;hsa05219;hsa05220;hsa05221;hsa05223;hsa05230;hsa05231 
                                               NRAS 
                                               NA 
                                               slateblue4 
                                               16 
                                               -2.58620689655172 
                                             
                                             
                                               HRAS-mRNA 
                                               -0.0795 
                                               0.0324 
                                               -0.143 
                                               -0.016 
                                               0.946 
                                               0.906 
                                               0.989 
                                               0.019 
                                               0.24 
                                               lm.nb 
                                               ARMS-mediated activation, CD209 (DC-SIGN) signaling, Cell surface interactions at the vascular wall, Constitutive Signaling by EGFRvIII, CREB phosphorylation through the activation of Ras, DAP12 signaling, Downstream signal transduction, FCERI mediated MAPK activation, Frs2-mediated activation, Gastrin-CREB signalling pathway via PKC and MAPK, Interleukin receptor SHC signaling, IRS-related events triggered by IGF1R, MAP2K and MAPK activation, RAFMAP kinase cascade, SHC-mediated cascade.CL.FGFR1, SHC-related events triggered by IGF1R, Signaling by ERBB2, Signaling by FGFR1 in disease, Signaling by FGFR2 in disease, Signaling by FGFR3 in disease, Signaling by FGFR4 in disease, Signaling by PDGF, Signaling by SCF-KIT, Tie2 Signaling, VEGFA-VEGFR2 Pathway, VEGFR2 mediated cell proliferation 
                                               SEBELIUS_2_C4636 
                                               HRAS 
                                               mRNA 
                                               0 
                                               
                                               
                                               Signaling by FGFR2 in disease;Regulation of RAS by GAPs;SHC-mediated cascade.CL.FGFR3;Signaling by FGFR3 in disease;MAP2K and MAPK activation;SHC-related events triggered by IGF1R;Signaling by FGFR4 in disease;SHC-mediated cascade.CL.FGFR1;Tie2 Signaling;Signaling by FGFR1 in disease;Activation of RAS in B cells;SHC-mediated cascade.CL.FGFR4;Constitutive Signaling by EGFRvIII;CD209 (DC-SIGN) signaling;Ras activation uopn Ca2.PLS. infux through NMDA receptor;FCERI mediated MAPK activation;RAFMAP kinase cascade;Downstream signal transduction;VEGFR2 mediated cell proliferation;Signaling by SCF-KIT;Frs2-mediated activation;ARMS-mediated activation;Gastrin-CREB signalling pathway via PKC and MAPK;Interleukin receptor SHC signaling;DAP12 signaling;Signaling by PDGF;VEGFA-VEGFR2 Pathway;Signaling by ERBB2;IRS-related events triggered by IGF1R;Cell surface interactions at the vascular wall;CREB phosphorylation through the activation of Ras 
                                               hsa04010;hsa04012;hsa04014;hsa04015;hsa04062;hsa04068;hsa04071;hsa04072;hsa04144;hsa04151;hsa04211;hsa04213;hsa04360;hsa04370;hsa04510;hsa04530;hsa04540;hsa04550;hsa04630;hsa04650;hsa04660;hsa04662;hsa04664;hsa04720;hsa04722;hsa04725;hsa04726;hsa04730;hsa04810;hsa04910;hsa04912;hsa04915;hsa04916;hsa04917;hsa04919;hsa04921;hsa04933;hsa05034;hsa05160;hsa05161;hsa05166;hsa05200;hsa05203;hsa05205;hsa05206;hsa05211;hsa05213;hsa05214;hsa05215;hsa05216;hsa05218;hsa05219;hsa05220;hsa05221;hsa05223;hsa05230;hsa05231 
                                               HRAS 
                                               NA 
                                               slateblue4 
                                               16 
                                               -2.4537037037037 
                                             
                                             
                                               KRAS-mRNA 
                                               -0.0261 
                                               0.0121 
                                               -0.0497 
                                               -0.00237 
                                               0.982 
                                               0.966 
                                               0.998 
                                               0.0376 
                                               0.408 
                                               lm.nb 
                                               ARMS-mediated activation, CD209 (DC-SIGN) signaling, Cell surface interactions at the vascular wall, Constitutive Signaling by EGFRvIII, DAP12 signaling, Downstream signal transduction, FCERI mediated MAPK activation, Frs2-mediated activation, Gastrin-CREB signalling pathway via PKC and MAPK, Interleukin receptor SHC signaling, IRS-related events triggered by IGF1R, MAP2K and MAPK activation, RAFMAP kinase cascade, SHC-mediated cascade.CL.FGFR1, SHC-related events triggered by IGF1R, Signaling by ERBB2, Signaling by FGFR1 in disease, Signaling by FGFR2 in disease, Signaling by FGFR3 in disease, Signaling by FGFR4 in disease, Signaling by PDGF, Signaling by SCF-KIT, Tie2 Signaling, VEGFA-VEGFR2 Pathway, VEGFR2 mediated cell proliferation 
                                               SEBELIUS_2_C4636 
                                               KRAS 
                                               mRNA 
                                               0 
                                               
                                               
                                               Signaling by FGFR2 in disease;Regulation of RAS by GAPs;SHC-mediated cascade.CL.FGFR3;Signaling by FGFR3 in disease;MAP2K and MAPK activation;SHC-related events triggered by IGF1R;Signaling by FGFR4 in disease;SHC-mediated cascade.CL.FGFR1;Tie2 Signaling;Signaling by FGFR1 in disease;Activation of RAS in B cells;SHC-mediated cascade.CL.FGFR4;Constitutive Signaling by EGFRvIII;CD209 (DC-SIGN) signaling;FCERI mediated MAPK activation;RAFMAP kinase cascade;Downstream signal transduction;VEGFR2 mediated cell proliferation;Signaling by SCF-KIT;Frs2-mediated activation;ARMS-mediated activation;Gastrin-CREB signalling pathway via PKC and MAPK;Interleukin receptor SHC signaling;DAP12 signaling;Signaling by PDGF;VEGFA-VEGFR2 Pathway;Signaling by ERBB2;IRS-related events triggered by IGF1R;Cell surface interactions at the vascular wall 
                                               hsa04010;hsa04012;hsa04014;hsa04015;hsa04062;hsa04068;hsa04071;hsa04072;hsa04151;hsa04211;hsa04213;hsa04320;hsa04360;hsa04370;hsa04530;hsa04540;hsa04550;hsa04650;hsa04660;hsa04662;hsa04664;hsa04720;hsa04722;hsa04725;hsa04726;hsa04730;hsa04810;hsa04910;hsa04912;hsa04914;hsa04915;hsa04916;hsa04917;hsa04919;hsa04921;hsa04933;hsa04960;hsa05034;hsa05160;hsa05161;hsa05166;hsa05200;hsa05203;hsa05205;hsa05206;hsa05210;hsa05211;hsa05212;hsa05213;hsa05214;hsa05215;hsa05216;hsa05218;hsa05219;hsa05220;hsa05221;hsa05223;hsa05230;hsa05231 
                                               KRAS 
                                               NA 
                                               slateblue4 
                                               16 
                                               -2.15702479338843 
                                             
                                             
                                               EP300-mRNA 
                                               0.00939 
                                               0.0182 
                                               -0.0262 
                                               0.045 
                                               1.01 
                                               0.982 
                                               1.03 
                                               0.608 
                                               1 
                                               lm.nb 
                                               CD209 (DC-SIGN) signaling, Circadian Clock 
                                               SEBELIUS_2_C4636 
                                               EP300 
                                               mRNA 
                                               0 
                                               
                                               
                                               LRR FLII-interacting protein 1 (LRRFIP1) activates type I IFN production;TRAF6 mediated IRF7 activation;Constitutive Signaling by NOTCH1 PEST Domain Mutants;Polo-like kinase mediated events;B-WICH complex positively regulates rRNA expression;CD209 (DC-SIGN) signaling;Circadian Clock 
                                               hsa04024;hsa04066;hsa04068;hsa04110;hsa04310;hsa04330;hsa04350;hsa04520;hsa04630;hsa04720;hsa04916;hsa04919;hsa04922;hsa05016;hsa05152;hsa05161;hsa05164;hsa05166;hsa05168;hsa05169;hsa05200;hsa05203;hsa05206;hsa05211;hsa05215 
                                               EP300 
                                               NA 
                                               azure4 
                                               1 
                                               0.515934065934066 
                                             
                                             
                                               FYN-mRNA 
                                               0.0103 
                                               0.0454 
                                               -0.0786 
                                               0.0993 
                                               1.01 
                                               0.947 
                                               1.07 
                                               0.821 
                                               1 
                                               lm.nb 
                                               ARMS-mediated activation, CD209 (DC-SIGN) signaling, CD28 co-stimulation, CD28 dependent PI3KAkt signaling, CD28 dependent Vav1 pathway, Cell surface interactions at the vascular wall, Constitutive Signaling by Aberrant PI3K in Cancer, CTLA4 inhibitory signaling, DAP12 signaling, Downstream signal transduction, EPH-ephrin mediated repulsion of cells, FCERI mediated MAPK activation, Frs2-mediated activation, GAB1 signalosome, Gastrin-CREB signalling pathway via PKC and MAPK, GPVI-mediated activation cascade, Interleukin receptor SHC signaling, IRS-related events triggered by IGF1R, Nephrin interactions, PI3KAKT activation, RAFMAP kinase cascade, Regulation of signaling by CBL, Role of LAT2NTALLAB on calcium mobilization, Sema3A PAK dependent Axon repulsion, Signaling by ERBB2, Signaling by PDGF, Signaling by SCF-KIT, VEGFA-VEGFR2 Pathway, VEGFR2 mediated cell proliferation 
                                               SEBELIUS_2_C4636 
                                               FYN 
                                               mRNA 
                                               0 
                                               
                                               
                                               EPH-ephrin mediated repulsion of cells;Regulation of signaling by CBL;FCGR activation;Antigen activates B Cell Receptor (BCR) leading to generation of second messengers;Dectin-2 family;RAFMAP kinase cascade;Constitutive Signaling by Aberrant PI3K in Cancer;CTLA4 inhibitory signaling;CD28 dependent PI3KAkt signaling;PECAM1 interactions;CD28 dependent Vav1 pathway;GPVI-mediated activation cascade;Sema3A PAK dependent Axon repulsion;Nephrin interactions;SEMA3A-Plexin repulsion signaling by inhibiting Integrin adhesion;CRMPs in Sema3A signaling;CD209 (DC-SIGN) signaling;Downstream signal transduction;GAB1 signalosome;Signaling by SCF-KIT;PI3KAKT activation;Role of LAT2NTALLAB on calcium mobilization;DAP12 signaling;Signaling by PDGF;Signaling by ERBB2;VEGFA-VEGFR2 Pathway;VEGFR2 mediated cell proliferation;Frs2-mediated activation;ARMS-mediated activation;Gastrin-CREB signalling pathway via PKC and MAPK;Interleukin receptor SHC signaling;FCERI mediated MAPK activation;IRS-related events triggered by IGF1R;CD28 co-stimulation;Cell surface interactions at the vascular wall 
                                               hsa04071;hsa04072;hsa04360;hsa04380;hsa04510;hsa04520;hsa04611;hsa04650;hsa04660;hsa04664;hsa04725;hsa05020;hsa05130;hsa05162;hsa05416 
                                               FYN 
                                               NA 
                                               azure4 
                                               1 
                                               0.226872246696035 
                                             
                                             
                                               CREBBP-mRNA 
                                               0.000773 
                                               0.0168 
                                               -0.0321 
                                               0.0336 
                                               1 
                                               0.978 
                                               1.02 
                                               0.963 
                                               1 
                                               lm.nb 
                                               CD209 (DC-SIGN) signaling, Circadian Clock 
                                               SEBELIUS_2_C4636 
                                               CREBBP 
                                               mRNA 
                                               0 
                                               
                                               
                                               YAP1- and WWTR1 (TAZ)-stimulated gene expression;Activation of gene expression by SREBF (SREBP);Transcriptional activation of mitochondrial biogenesis;Constitutive Signaling by NOTCH1 HD.PLS.PEST Domain Mutants;Constitutive Signaling by NOTCH1 PEST Domain Mutants;CD209 (DC-SIGN) signaling;Circadian Clock 
                                               hsa04024;hsa04066;hsa04068;hsa04110;hsa04310;hsa04330;hsa04350;hsa04520;hsa04630;hsa04720;hsa04916;hsa04919;hsa04922;hsa05016;hsa05152;hsa05161;hsa05164;hsa05166;hsa05168;hsa05169;hsa05200;hsa05203;hsa05206;hsa05211;hsa05215 
                                               CREBBP 
                                               NA 
                                               azure4 
                                               1 
                                               0.0460119047619048 
                                             
                                           
                                           DE Results - CD209 (DC-SIGN) signaling genes - Day  Table displaying each sample's global significance scores and directed global significance scores as defined in the heatmaps above. The global significance score is calculated as the square root of the mean squared t-statistic for the genes in a gene set, with t-statistics coming from the linear regression underlying our differential expression analysis. The directed global significance score is calculated as the square root of the mean signed squared t-statistic for the genes in a gene set, with t-statistics coming from the linear regression underlying our differential expression analysis.   
                                      
                                       
                                  
                                
                              
                            
                            
                              
                                
                                  
                                       
                                      
                                           Volcano Plot: Line: Caco2 vs.Fibro   More Plot Information                Volcano Plot: Line: Caco2 vs.Fibro  Volcano plot displaying each gene's -log10(p-value) and log2 fold change for the selected covariate. Highly statistically significant genes fall at the top of the plot, and highly differentially expressed genes fall to either side. Genes within the selected gene set are highlighted in orange. Horizontal lines indicate various False Discovery Rate (FDR) thresholds.   
                                           DE Results - CD209 (DC-SIGN) signaling genes - Line: Caco2 vs.Fibro   More Plot Information      Download CSV Data     
                                           
                                             
                                             
                                           
                                           
                                             
                                               Probe Label 
                                               Log2 fold change 
                                               std error (log2) 
                                               Lower confidence limit (log2) 
                                               Upper confidence limit (log2) 
                                               Linear fold change 
                                               Lower confidence limit (linear) 
                                               Upper confidence limit (linear) 
                                               P-value 
                                               BY.p.value 
                                               method 
                                               Gene.sets 
                                               Codeset.Name 
                                               Probe.Label 
                                               Analyte.Type 
                                               Is.Control 
                                               Control.Type 
                                               Related.Probes 
                                               Probe.Annotation 
                                               KEGG.Pathways 
                                               Official.Gene.Name 
                                               Control.Conc 
                                               volcanocol 
                                               volcanopch 
                                               tstats.all 
                                             
                                           
                                           
                                             
                                               KRAS-mRNA 
                                               2.3 
                                               0.0937 
                                               2.11 
                                               2.48 
                                               4.91 
                                               4.32 
                                               5.58 
                                               1.8e-24 
                                               4.27e-22 
                                               lm.nb 
                                               ARMS-mediated activation, CD209 (DC-SIGN) signaling, Cell surface interactions at the vascular wall, Constitutive Signaling by EGFRvIII, DAP12 signaling, Downstream signal transduction, FCERI mediated MAPK activation, Frs2-mediated activation, Gastrin-CREB signalling pathway via PKC and MAPK, Interleukin receptor SHC signaling, IRS-related events triggered by IGF1R, MAP2K and MAPK activation, RAFMAP kinase cascade, SHC-mediated cascade.CL.FGFR1, SHC-related events triggered by IGF1R, Signaling by ERBB2, Signaling by FGFR1 in disease, Signaling by FGFR2 in disease, Signaling by FGFR3 in disease, Signaling by FGFR4 in disease, Signaling by PDGF, Signaling by SCF-KIT, Tie2 Signaling, VEGFA-VEGFR2 Pathway, VEGFR2 mediated cell proliferation 
                                               SEBELIUS_2_C4636 
                                               KRAS 
                                               mRNA 
                                               0 
                                               
                                               
                                               Signaling by FGFR2 in disease;Regulation of RAS by GAPs;SHC-mediated cascade.CL.FGFR3;Signaling by FGFR3 in disease;MAP2K and MAPK activation;SHC-related events triggered by IGF1R;Signaling by FGFR4 in disease;SHC-mediated cascade.CL.FGFR1;Tie2 Signaling;Signaling by FGFR1 in disease;Activation of RAS in B cells;SHC-mediated cascade.CL.FGFR4;Constitutive Signaling by EGFRvIII;CD209 (DC-SIGN) signaling;FCERI mediated MAPK activation;RAFMAP kinase cascade;Downstream signal transduction;VEGFR2 mediated cell proliferation;Signaling by SCF-KIT;Frs2-mediated activation;ARMS-mediated activation;Gastrin-CREB signalling pathway via PKC and MAPK;Interleukin receptor SHC signaling;DAP12 signaling;Signaling by PDGF;VEGFA-VEGFR2 Pathway;Signaling by ERBB2;IRS-related events triggered by IGF1R;Cell surface interactions at the vascular wall 
                                               hsa04010;hsa04012;hsa04014;hsa04015;hsa04062;hsa04068;hsa04071;hsa04072;hsa04151;hsa04211;hsa04213;hsa04320;hsa04360;hsa04370;hsa04530;hsa04540;hsa04550;hsa04650;hsa04660;hsa04662;hsa04664;hsa04720;hsa04722;hsa04725;hsa04726;hsa04730;hsa04810;hsa04910;hsa04912;hsa04914;hsa04915;hsa04916;hsa04917;hsa04919;hsa04921;hsa04933;hsa04960;hsa05034;hsa05160;hsa05161;hsa05166;hsa05200;hsa05203;hsa05205;hsa05206;hsa05210;hsa05211;hsa05212;hsa05213;hsa05214;hsa05215;hsa05216;hsa05218;hsa05219;hsa05220;hsa05221;hsa05223;hsa05230;hsa05231 
                                               KRAS 
                                               NA 
                                               slateblue1 
                                               16 
                                               24.5464247598719 
                                             
                                             
                                               PAK1-mRNA 
                                               1.92 
                                               0.159 
                                               1.61 
                                               2.23 
                                               3.79 
                                               3.05 
                                               4.7 
                                               2.23e-14 
                                               5.02e-13 
                                               lm.nb 
                                               Activation of Rac, CD209 (DC-SIGN) signaling, CD28 co-stimulation, CD28 dependent Vav1 pathway, EPHB-mediated forward signaling, Ephrin signaling, FCERI mediated MAPK activation, MAPK6MAPK4 signaling, Regulation of actin dynamics for phagocytic cup formation, RHO GTPases activate PAKs, Sema3A PAK dependent Axon repulsion, VEGFA-VEGFR2 Pathway, VEGFR2 mediated vascular permeability 
                                               SEBELIUS_2_C4636 
                                               PAK1 
                                               mRNA 
                                               0 
                                               
                                               
                                               EPHB-mediated forward signaling;Ephrin signaling;MAPK6MAPK4 signaling;DSCAM interactions;Activation of Rac;VEGFR2 mediated vascular permeability;RHO GTPases activate PAKs;Sema3A PAK dependent Axon repulsion;FCERI mediated MAPK activation;Regulation of actin dynamics for phagocytic cup formation;CD209 (DC-SIGN) signaling;CD28 dependent Vav1 pathway;VEGFA-VEGFR2 Pathway;CD28 co-stimulation 
                                               hsa04010;hsa04012;hsa04014;hsa04024;hsa04062;hsa04360;hsa04510;hsa04650;hsa04660;hsa04666;hsa04810;hsa05120;hsa05205;hsa05211 
                                               PAK1 
                                               NA 
                                               slateblue1 
                                               16 
                                               12.0754716981132 
                                             
                                             
                                               FYN-mRNA 
                                               -2.64 
                                               0.335 
                                               -3.3 
                                               -1.99 
                                               0.16 
                                               0.102 
                                               0.253 
                                               1.96e-09 
                                               2.84e-08 
                                               lm.nb 
                                               ARMS-mediated activation, CD209 (DC-SIGN) signaling, CD28 co-stimulation, CD28 dependent PI3KAkt signaling, CD28 dependent Vav1 pathway, Cell surface interactions at the vascular wall, Constitutive Signaling by Aberrant PI3K in Cancer, CTLA4 inhibitory signaling, DAP12 signaling, Downstream signal transduction, EPH-ephrin mediated repulsion of cells, FCERI mediated MAPK activation, Frs2-mediated activation, GAB1 signalosome, Gastrin-CREB signalling pathway via PKC and MAPK, GPVI-mediated activation cascade, Interleukin receptor SHC signaling, IRS-related events triggered by IGF1R, Nephrin interactions, PI3KAKT activation, RAFMAP kinase cascade, Regulation of signaling by CBL, Role of LAT2NTALLAB on calcium mobilization, Sema3A PAK dependent Axon repulsion, Signaling by ERBB2, Signaling by PDGF, Signaling by SCF-KIT, VEGFA-VEGFR2 Pathway, VEGFR2 mediated cell proliferation 
                                               SEBELIUS_2_C4636 
                                               FYN 
                                               mRNA 
                                               0 
                                               
                                               
                                               EPH-ephrin mediated repulsion of cells;Regulation of signaling by CBL;FCGR activation;Antigen activates B Cell Receptor (BCR) leading to generation of second messengers;Dectin-2 family;RAFMAP kinase cascade;Constitutive Signaling by Aberrant PI3K in Cancer;CTLA4 inhibitory signaling;CD28 dependent PI3KAkt signaling;PECAM1 interactions;CD28 dependent Vav1 pathway;GPVI-mediated activation cascade;Sema3A PAK dependent Axon repulsion;Nephrin interactions;SEMA3A-Plexin repulsion signaling by inhibiting Integrin adhesion;CRMPs in Sema3A signaling;CD209 (DC-SIGN) signaling;Downstream signal transduction;GAB1 signalosome;Signaling by SCF-KIT;PI3KAKT activation;Role of LAT2NTALLAB on calcium mobilization;DAP12 signaling;Signaling by PDGF;Signaling by ERBB2;VEGFA-VEGFR2 Pathway;VEGFR2 mediated cell proliferation;Frs2-mediated activation;ARMS-mediated activation;Gastrin-CREB signalling pathway via PKC and MAPK;Interleukin receptor SHC signaling;FCERI mediated MAPK activation;IRS-related events triggered by IGF1R;CD28 co-stimulation;Cell surface interactions at the vascular wall 
                                               hsa04071;hsa04072;hsa04360;hsa04380;hsa04510;hsa04520;hsa04611;hsa04650;hsa04660;hsa04664;hsa04725;hsa05020;hsa05130;hsa05162;hsa05416 
                                               FYN 
                                               NA 
                                               slateblue1 
                                               16 
                                               -7.88059701492537 
                                             
                                             
                                               EP300-mRNA 
                                               0.915 
                                               0.138 
                                               0.644 
                                               1.19 
                                               1.89 
                                               1.56 
                                               2.28 
                                               9.46e-08 
                                               1.18e-06 
                                               lm.nb 
                                               CD209 (DC-SIGN) signaling, Circadian Clock 
                                               SEBELIUS_2_C4636 
                                               EP300 
                                               mRNA 
                                               0 
                                               
                                               
                                               LRR FLII-interacting protein 1 (LRRFIP1) activates type I IFN production;TRAF6 mediated IRF7 activation;Constitutive Signaling by NOTCH1 PEST Domain Mutants;Polo-like kinase mediated events;B-WICH complex positively regulates rRNA expression;CD209 (DC-SIGN) signaling;Circadian Clock 
                                               hsa04024;hsa04066;hsa04068;hsa04110;hsa04310;hsa04330;hsa04350;hsa04520;hsa04630;hsa04720;hsa04916;hsa04919;hsa04922;hsa05016;hsa05152;hsa05161;hsa05164;hsa05166;hsa05168;hsa05169;hsa05200;hsa05203;hsa05206;hsa05211;hsa05215 
                                               EP300 
                                               NA 
                                               slateblue1 
                                               16 
                                               6.6304347826087 
                                             
                                             
                                               HRAS-mRNA 
                                               -0.868 
                                               0.244 
                                               -1.35 
                                               -0.391 
                                               0.548 
                                               0.393 
                                               0.763 
                                               0.00102 
                                               0.00903 
                                               lm.nb 
                                               ARMS-mediated activation, CD209 (DC-SIGN) signaling, Cell surface interactions at the vascular wall, Constitutive Signaling by EGFRvIII, CREB phosphorylation through the activation of Ras, DAP12 signaling, Downstream signal transduction, FCERI mediated MAPK activation, Frs2-mediated activation, Gastrin-CREB signalling pathway via PKC and MAPK, Interleukin receptor SHC signaling, IRS-related events triggered by IGF1R, MAP2K and MAPK activation, RAFMAP kinase cascade, SHC-mediated cascade.CL.FGFR1, SHC-related events triggered by IGF1R, Signaling by ERBB2, Signaling by FGFR1 in disease, Signaling by FGFR2 in disease, Signaling by FGFR3 in disease, Signaling by FGFR4 in disease, Signaling by PDGF, Signaling by SCF-KIT, Tie2 Signaling, VEGFA-VEGFR2 Pathway, VEGFR2 mediated cell proliferation 
                                               SEBELIUS_2_C4636 
                                               HRAS 
                                               mRNA 
                                               0 
                                               
                                               
                                               Signaling by FGFR2 in disease;Regulation of RAS by GAPs;SHC-mediated cascade.CL.FGFR3;Signaling by FGFR3 in disease;MAP2K and MAPK activation;SHC-related events triggered by IGF1R;Signaling by FGFR4 in disease;SHC-mediated cascade.CL.FGFR1;Tie2 Signaling;Signaling by FGFR1 in disease;Activation of RAS in B cells;SHC-mediated cascade.CL.FGFR4;Constitutive Signaling by EGFRvIII;CD209 (DC-SIGN) signaling;Ras activation uopn Ca2.PLS. infux through NMDA receptor;FCERI mediated MAPK activation;RAFMAP kinase cascade;Downstream signal transduction;VEGFR2 mediated cell proliferation;Signaling by SCF-KIT;Frs2-mediated activation;ARMS-mediated activation;Gastrin-CREB signalling pathway via PKC and MAPK;Interleukin receptor SHC signaling;DAP12 signaling;Signaling by PDGF;VEGFA-VEGFR2 Pathway;Signaling by ERBB2;IRS-related events triggered by IGF1R;Cell surface interactions at the vascular wall;CREB phosphorylation through the activation of Ras 
                                               hsa04010;hsa04012;hsa04014;hsa04015;hsa04062;hsa04068;hsa04071;hsa04072;hsa04144;hsa04151;hsa04211;hsa04213;hsa04360;hsa04370;hsa04510;hsa04530;hsa04540;hsa04550;hsa04630;hsa04650;hsa04660;hsa04662;hsa04664;hsa04720;hsa04722;hsa04725;hsa04726;hsa04730;hsa04810;hsa04910;hsa04912;hsa04915;hsa04916;hsa04917;hsa04919;hsa04921;hsa04933;hsa05034;hsa05160;hsa05161;hsa05166;hsa05200;hsa05203;hsa05205;hsa05206;hsa05211;hsa05213;hsa05214;hsa05215;hsa05216;hsa05218;hsa05219;hsa05220;hsa05221;hsa05223;hsa05230;hsa05231 
                                               HRAS 
                                               NA 
                                               slateblue1 
                                               16 
                                               -3.55737704918033 
                                             
                                             
                                               RAF1-mRNA 
                                               0.292 
                                               0.0833 
                                               0.129 
                                               0.455 
                                               1.22 
                                               1.09 
                                               1.37 
                                               0.0012 
                                               0.0103 
                                               loglinear 
                                               ARMS-mediated activation, CD209 (DC-SIGN) signaling, CREB phosphorylation through the activation of Ras, DAP12 signaling, Downstream signal transduction, FCERI mediated MAPK activation, Frs2-mediated activation, Gastrin-CREB signalling pathway via PKC and MAPK, Interleukin receptor SHC signaling, IRS-related events triggered by IGF1R, Negative feedback regulation of MAPK pathway, Negative regulation of MAPK pathway, RAF activation, RAFMAP kinase cascade, Signaling by ERBB2, Signaling by PDGF, Signaling by SCF-KIT, VEGFA-VEGFR2 Pathway, VEGFR2 mediated cell proliferation 
                                               SEBELIUS_2_C4636 
                                               RAF1 
                                               mRNA 
                                               0 
                                               
                                               
                                               RAF activation;Stimuli-sensing channels;CREB phosphorylation through the activation of Ras;Negative feedback regulation of MAPK pathway;CD209 (DC-SIGN) signaling;RAFMAP kinase cascade;Downstream signal transduction;VEGFR2 mediated cell proliferation;Signaling by SCF-KIT;Frs2-mediated activation;ARMS-mediated activation;Gastrin-CREB signalling pathway via PKC and MAPK;Interleukin receptor SHC signaling;FCERI mediated MAPK activation;DAP12 signaling;Signaling by PDGF;VEGFA-VEGFR2 Pathway;Signaling by ERBB2;IRS-related events triggered by IGF1R;Negative regulation of MAPK pathway 
                                               hsa04010;hsa04012;hsa04014;hsa04015;hsa04022;hsa04024;hsa04062;hsa04068;hsa04071;hsa04072;hsa04151;hsa04270;hsa04370;hsa04510;hsa04540;hsa04550;hsa04630;hsa04650;hsa04660;hsa04662;hsa04664;hsa04666;hsa04720;hsa04722;hsa04726;hsa04730;hsa04810;hsa04910;hsa04912;hsa04914;hsa04915;hsa04916;hsa04917;hsa04919;hsa04921;hsa05034;hsa05152;hsa05160;hsa05161;hsa05164;hsa05200;hsa05205;hsa05206;hsa05210;hsa05211;hsa05212;hsa05213;hsa05214;hsa05215;hsa05218;hsa05219;hsa05220;hsa05221;hsa05223;hsa05230;hsa05231 
                                               RAF1 
                                               NA 
                                               slateblue2 
                                               16 
                                               3.50540216086435 
                                             
                                             
                                               NRAS-mRNA 
                                               0.29 
                                               0.0881 
                                               0.118 
                                               0.463 
                                               1.22 
                                               1.09 
                                               1.38 
                                               0.00217 
                                               0.0183 
                                               lm.nb 
                                               ARMS-mediated activation, CD209 (DC-SIGN) signaling, Cell surface interactions at the vascular wall, Constitutive Signaling by EGFRvIII, DAP12 signaling, Downstream signal transduction, FCERI mediated MAPK activation, Frs2-mediated activation, Gastrin-CREB signalling pathway via PKC and MAPK, Interleukin receptor SHC signaling, IRS-related events triggered by IGF1R, MAP2K and MAPK activation, RAFMAP kinase cascade, SHC-mediated cascade.CL.FGFR1, SHC-related events triggered by IGF1R, Signaling by ERBB2, Signaling by FGFR1 in disease, Signaling by FGFR2 in disease, Signaling by FGFR3 in disease, Signaling by FGFR4 in disease, Signaling by PDGF, Signaling by SCF-KIT, Tie2 Signaling, VEGFA-VEGFR2 Pathway, VEGFR2 mediated cell proliferation 
                                               SEBELIUS_2_C4636 
                                               NRAS 
                                               mRNA 
                                               0 
                                               
                                               
                                               Signaling by FGFR2 in disease;Regulation of RAS by GAPs;SHC-mediated cascade.CL.FGFR3;Signaling by FGFR3 in disease;MAP2K and MAPK activation;SHC-related events triggered by IGF1R;Signaling by FGFR4 in disease;SHC-mediated cascade.CL.FGFR1;Tie2 Signaling;Signaling by FGFR1 in disease;Activation of RAS in B cells;SHC-mediated cascade.CL.FGFR4;Constitutive Signaling by EGFRvIII;CD209 (DC-SIGN) signaling;FCERI mediated MAPK activation;RAFMAP kinase cascade;Downstream signal transduction;VEGFR2 mediated cell proliferation;Signaling by SCF-KIT;Frs2-mediated activation;ARMS-mediated activation;Gastrin-CREB signalling pathway via PKC and MAPK;Interleukin receptor SHC signaling;DAP12 signaling;Signaling by PDGF;VEGFA-VEGFR2 Pathway;Signaling by ERBB2;IRS-related events triggered by IGF1R;Cell surface interactions at the vascular wall 
                                               hsa04010;hsa04012;hsa04014;hsa04015;hsa04062;hsa04068;hsa04071;hsa04072;hsa04151;hsa04211;hsa04213;hsa04360;hsa04370;hsa04530;hsa04540;hsa04550;hsa04650;hsa04660;hsa04662;hsa04664;hsa04720;hsa04722;hsa04725;hsa04726;hsa04730;hsa04810;hsa04910;hsa04912;hsa04915;hsa04916;hsa04917;hsa04919;hsa04921;hsa04933;hsa05034;hsa05160;hsa05161;hsa05166;hsa05200;hsa05203;hsa05205;hsa05206;hsa05211;hsa05213;hsa05214;hsa05215;hsa05216;hsa05218;hsa05219;hsa05220;hsa05221;hsa05223;hsa05230;hsa05231 
                                               NRAS 
                                               NA 
                                               slateblue2 
                                               16 
                                               3.29171396140749 
                                             
                                             
                                               PAK2-mRNA 
                                               -0.138 
                                               0.0752 
                                               -0.285 
                                               0.00968 
                                               0.909 
                                               0.821 
                                               1.01 
                                               0.0751 
                                               0.552 
                                               loglinear 
                                               Activation of Rac, CD209 (DC-SIGN) signaling, CD28 co-stimulation, CD28 dependent Vav1 pathway, Ephrin signaling, RHO GTPases activate PAKs, Sema3A PAK dependent Axon repulsion, VEGFA-VEGFR2 Pathway, VEGFR2 mediated vascular permeability 
                                               SEBELIUS_2_C4636 
                                               PAK2 
                                               mRNA 
                                               0 
                                               
                                               
                                               Nef and signal transduction;CD28 dependent Vav1 pathway;Stimulation of the cell death response by PAK-2p34;Ephrin signaling;Activation of Rac;RHO GTPases activate PAKs;VEGFR2 mediated vascular permeability;Sema3A PAK dependent Axon repulsion;CD209 (DC-SIGN) signaling;VEGFA-VEGFR2 Pathway;CD28 co-stimulation 
                                               hsa04010;hsa04012;hsa04014;hsa04360;hsa04510;hsa04660;hsa04810;hsa05211 
                                               PAK2 
                                               NA 
                                               azure4 
                                               1 
                                               -1.83510638297872 
                                             
                                             
                                               CREBBP-mRNA 
                                               0.132 
                                               0.127 
                                               -0.117 
                                               0.381 
                                               1.1 
                                               0.922 
                                               1.3 
                                               0.306 
                                               1 
                                               lm.nb 
                                               CD209 (DC-SIGN) signaling, Circadian Clock 
                                               SEBELIUS_2_C4636 
                                               CREBBP 
                                               mRNA 
                                               0 
                                               
                                               
                                               YAP1- and WWTR1 (TAZ)-stimulated gene expression;Activation of gene expression by SREBF (SREBP);Transcriptional activation of mitochondrial biogenesis;Constitutive Signaling by NOTCH1 HD.PLS.PEST Domain Mutants;Constitutive Signaling by NOTCH1 PEST Domain Mutants;CD209 (DC-SIGN) signaling;Circadian Clock 
                                               hsa04024;hsa04066;hsa04068;hsa04110;hsa04310;hsa04330;hsa04350;hsa04520;hsa04630;hsa04720;hsa04916;hsa04919;hsa04922;hsa05016;hsa05152;hsa05161;hsa05164;hsa05166;hsa05168;hsa05169;hsa05200;hsa05203;hsa05206;hsa05211;hsa05215 
                                               CREBBP 
                                               NA 
                                               azure4 
                                               1 
                                               1.03937007874016 
                                             
                                           
                                           DE Results - CD209 (DC-SIGN) signaling genes - Line: Caco2 vs.Fibro  Table displaying each sample's global significance scores and directed global significance scores as defined in the heatmaps above. The global significance score is calculated as the square root of the mean squared t-statistic for the genes in a gene set, with t-statistics coming from the linear regression underlying our differential expression analysis. The directed global significance score is calculated as the square root of the mean signed squared t-statistic for the genes in a gene set, with t-statistics coming from the linear regression underlying our differential expression analysis.   
                                      
                                       
                                  
                                
                                
                                  
                                       
                                      
                                           Volcano Plot: Line: HEPG2 vs.Fibro   More Plot Information                Volcano Plot: Line: HEPG2 vs.Fibro  Volcano plot displaying each gene's -log10(p-value) and log2 fold change for the selected covariate. Highly statistically significant genes fall at the top of the plot, and highly differentially expressed genes fall to either side. Genes within the selected gene set are highlighted in orange. Horizontal lines indicate various False Discovery Rate (FDR) thresholds.   
                                           DE Results - CD209 (DC-SIGN) signaling genes - Line: HEPG2 vs.Fibro   More Plot Information      Download CSV Data     
                                           
                                             
                                             
                                           
                                           
                                             
                                               Probe Label 
                                               Log2 fold change 
                                               std error (log2) 
                                               Lower confidence limit (log2) 
                                               Upper confidence limit (log2) 
                                               Linear fold change 
                                               Lower confidence limit (linear) 
                                               Upper confidence limit (linear) 
                                               P-value 
                                               BY.p.value 
                                               method 
                                               Gene.sets 
                                               Codeset.Name 
                                               Probe.Label 
                                               Analyte.Type 
                                               Is.Control 
                                               Control.Type 
                                               Related.Probes 
                                               Probe.Annotation 
                                               KEGG.Pathways 
                                               Official.Gene.Name 
                                               Control.Conc 
                                               volcanocol 
                                               volcanopch 
                                               tstats.all 
                                             
                                           
                                           
                                             
                                               PAK1-mRNA 
                                               -2.44 
                                               0.179 
                                               -2.79 
                                               -2.09 
                                               0.184 
                                               0.145 
                                               0.235 
                                               5.34e-16 
                                               1.72e-14 
                                               lm.nb 
                                               Activation of Rac, CD209 (DC-SIGN) signaling, CD28 co-stimulation, CD28 dependent Vav1 pathway, EPHB-mediated forward signaling, Ephrin signaling, FCERI mediated MAPK activation, MAPK6MAPK4 signaling, Regulation of actin dynamics for phagocytic cup formation, RHO GTPases activate PAKs, Sema3A PAK dependent Axon repulsion, VEGFA-VEGFR2 Pathway, VEGFR2 mediated vascular permeability 
                                               SEBELIUS_2_C4636 
                                               PAK1 
                                               mRNA 
                                               0 
                                               
                                               
                                               EPHB-mediated forward signaling;Ephrin signaling;MAPK6MAPK4 signaling;DSCAM interactions;Activation of Rac;VEGFR2 mediated vascular permeability;RHO GTPases activate PAKs;Sema3A PAK dependent Axon repulsion;FCERI mediated MAPK activation;Regulation of actin dynamics for phagocytic cup formation;CD209 (DC-SIGN) signaling;CD28 dependent Vav1 pathway;VEGFA-VEGFR2 Pathway;CD28 co-stimulation 
                                               hsa04010;hsa04012;hsa04014;hsa04024;hsa04062;hsa04360;hsa04510;hsa04650;hsa04660;hsa04666;hsa04810;hsa05120;hsa05205;hsa05211 
                                               PAK1 
                                               NA 
                                               slateblue1 
                                               16 
                                               -13.6312849162011 
                                             
                                             
                                               NRAS-mRNA 
                                               1.09 
                                               0.0962 
                                               0.904 
                                               1.28 
                                               2.13 
                                               1.87 
                                               2.43 
                                               1.28e-13 
                                               3.29e-12 
                                               lm.nb 
                                               ARMS-mediated activation, CD209 (DC-SIGN) signaling, Cell surface interactions at the vascular wall, Constitutive Signaling by EGFRvIII, DAP12 signaling, Downstream signal transduction, FCERI mediated MAPK activation, Frs2-mediated activation, Gastrin-CREB signalling pathway via PKC and MAPK, Interleukin receptor SHC signaling, IRS-related events triggered by IGF1R, MAP2K and MAPK activation, RAFMAP kinase cascade, SHC-mediated cascade.CL.FGFR1, SHC-related events triggered by IGF1R, Signaling by ERBB2, Signaling by FGFR1 in disease, Signaling by FGFR2 in disease, Signaling by FGFR3 in disease, Signaling by FGFR4 in disease, Signaling by PDGF, Signaling by SCF-KIT, Tie2 Signaling, VEGFA-VEGFR2 Pathway, VEGFR2 mediated cell proliferation 
                                               SEBELIUS_2_C4636 
                                               NRAS 
                                               mRNA 
                                               0 
                                               
                                               
                                               Signaling by FGFR2 in disease;Regulation of RAS by GAPs;SHC-mediated cascade.CL.FGFR3;Signaling by FGFR3 in disease;MAP2K and MAPK activation;SHC-related events triggered by IGF1R;Signaling by FGFR4 in disease;SHC-mediated cascade.CL.FGFR1;Tie2 Signaling;Signaling by FGFR1 in disease;Activation of RAS in B cells;SHC-mediated cascade.CL.FGFR4;Constitutive Signaling by EGFRvIII;CD209 (DC-SIGN) signaling;FCERI mediated MAPK activation;RAFMAP kinase cascade;Downstream signal transduction;VEGFR2 mediated cell proliferation;Signaling by SCF-KIT;Frs2-mediated activation;ARMS-mediated activation;Gastrin-CREB signalling pathway via PKC and MAPK;Interleukin receptor SHC signaling;DAP12 signaling;Signaling by PDGF;VEGFA-VEGFR2 Pathway;Signaling by ERBB2;IRS-related events triggered by IGF1R;Cell surface interactions at the vascular wall 
                                               hsa04010;hsa04012;hsa04014;hsa04015;hsa04062;hsa04068;hsa04071;hsa04072;hsa04151;hsa04211;hsa04213;hsa04360;hsa04370;hsa04530;hsa04540;hsa04550;hsa04650;hsa04660;hsa04662;hsa04664;hsa04720;hsa04722;hsa04725;hsa04726;hsa04730;hsa04810;hsa04910;hsa04912;hsa04915;hsa04916;hsa04917;hsa04919;hsa04921;hsa04933;hsa05034;hsa05160;hsa05161;hsa05166;hsa05200;hsa05203;hsa05205;hsa05206;hsa05211;hsa05213;hsa05214;hsa05215;hsa05216;hsa05218;hsa05219;hsa05220;hsa05221;hsa05223;hsa05230;hsa05231 
                                               NRAS 
                                               NA 
                                               slateblue1 
                                               16 
                                               11.3305613305613 
                                             
                                             
                                               EP300-mRNA 
                                               1.32 
                                               0.152 
                                               1.02 
                                               1.62 
                                               2.5 
                                               2.03 
                                               3.07 
                                               1.75e-10 
                                               3.26e-09 
                                               lm.nb 
                                               CD209 (DC-SIGN) signaling, Circadian Clock 
                                               SEBELIUS_2_C4636 
                                               EP300 
                                               mRNA 
                                               0 
                                               
                                               
                                               LRR FLII-interacting protein 1 (LRRFIP1) activates type I IFN production;TRAF6 mediated IRF7 activation;Constitutive Signaling by NOTCH1 PEST Domain Mutants;Polo-like kinase mediated events;B-WICH complex positively regulates rRNA expression;CD209 (DC-SIGN) signaling;Circadian Clock 
                                               hsa04024;hsa04066;hsa04068;hsa04110;hsa04310;hsa04330;hsa04350;hsa04520;hsa04630;hsa04720;hsa04916;hsa04919;hsa04922;hsa05016;hsa05152;hsa05161;hsa05164;hsa05166;hsa05168;hsa05169;hsa05200;hsa05203;hsa05206;hsa05211;hsa05215 
                                               EP300 
                                               NA 
                                               slateblue1 
                                               16 
                                               8.68421052631579 
                                             
                                             
                                               CREBBP-mRNA 
                                               0.78 
                                               0.139 
                                               0.507 
                                               1.05 
                                               1.72 
                                               1.42 
                                               2.07 
                                               2.16e-06 
                                               2.57e-05 
                                               lm.nb 
                                               CD209 (DC-SIGN) signaling, Circadian Clock 
                                               SEBELIUS_2_C4636 
                                               CREBBP 
                                               mRNA 
                                               0 
                                               
                                               
                                               YAP1- and WWTR1 (TAZ)-stimulated gene expression;Activation of gene expression by SREBF (SREBP);Transcriptional activation of mitochondrial biogenesis;Constitutive Signaling by NOTCH1 HD.PLS.PEST Domain Mutants;Constitutive Signaling by NOTCH1 PEST Domain Mutants;CD209 (DC-SIGN) signaling;Circadian Clock 
                                               hsa04024;hsa04066;hsa04068;hsa04110;hsa04310;hsa04330;hsa04350;hsa04520;hsa04630;hsa04720;hsa04916;hsa04919;hsa04922;hsa05016;hsa05152;hsa05161;hsa05164;hsa05166;hsa05168;hsa05169;hsa05200;hsa05203;hsa05206;hsa05211;hsa05215 
                                               CREBBP 
                                               NA 
                                               slateblue1 
                                               16 
                                               5.61151079136691 
                                             
                                             
                                               PAK2-mRNA 
                                               0.446 
                                               0.0825 
                                               0.284 
                                               0.608 
                                               1.36 
                                               1.22 
                                               1.52 
                                               4.03e-06 
                                               4.65e-05 
                                               loglinear 
                                               Activation of Rac, CD209 (DC-SIGN) signaling, CD28 co-stimulation, CD28 dependent Vav1 pathway, Ephrin signaling, RHO GTPases activate PAKs, Sema3A PAK dependent Axon repulsion, VEGFA-VEGFR2 Pathway, VEGFR2 mediated vascular permeability 
                                               SEBELIUS_2_C4636 
                                               PAK2 
                                               mRNA 
                                               0 
                                               
                                               
                                               Nef and signal transduction;CD28 dependent Vav1 pathway;Stimulation of the cell death response by PAK-2p34;Ephrin signaling;Activation of Rac;RHO GTPases activate PAKs;VEGFR2 mediated vascular permeability;Sema3A PAK dependent Axon repulsion;CD209 (DC-SIGN) signaling;VEGFA-VEGFR2 Pathway;CD28 co-stimulation 
                                               hsa04010;hsa04012;hsa04014;hsa04360;hsa04510;hsa04660;hsa04810;hsa05211 
                                               PAK2 
                                               NA 
                                               slateblue1 
                                               16 
                                               5.40606060606061 
                                             
                                             
                                               KRAS-mRNA 
                                               -0.345 
                                               0.104 
                                               -0.549 
                                               -0.14 
                                               0.788 
                                               0.684 
                                               0.907 
                                               0.00211 
                                               0.0189 
                                               lm.nb 
                                               ARMS-mediated activation, CD209 (DC-SIGN) signaling, Cell surface interactions at the vascular wall, Constitutive Signaling by EGFRvIII, DAP12 signaling, Downstream signal transduction, FCERI mediated MAPK activation, Frs2-mediated activation, Gastrin-CREB signalling pathway via PKC and MAPK, Interleukin receptor SHC signaling, IRS-related events triggered by IGF1R, MAP2K and MAPK activation, RAFMAP kinase cascade, SHC-mediated cascade.CL.FGFR1, SHC-related events triggered by IGF1R, Signaling by ERBB2, Signaling by FGFR1 in disease, Signaling by FGFR2 in disease, Signaling by FGFR3 in disease, Signaling by FGFR4 in disease, Signaling by PDGF, Signaling by SCF-KIT, Tie2 Signaling, VEGFA-VEGFR2 Pathway, VEGFR2 mediated cell proliferation 
                                               SEBELIUS_2_C4636 
                                               KRAS 
                                               mRNA 
                                               0 
                                               
                                               
                                               Signaling by FGFR2 in disease;Regulation of RAS by GAPs;SHC-mediated cascade.CL.FGFR3;Signaling by FGFR3 in disease;MAP2K and MAPK activation;SHC-related events triggered by IGF1R;Signaling by FGFR4 in disease;SHC-mediated cascade.CL.FGFR1;Tie2 Signaling;Signaling by FGFR1 in disease;Activation of RAS in B cells;SHC-mediated cascade.CL.FGFR4;Constitutive Signaling by EGFRvIII;CD209 (DC-SIGN) signaling;FCERI mediated MAPK activation;RAFMAP kinase cascade;Downstream signal transduction;VEGFR2 mediated cell proliferation;Signaling by SCF-KIT;Frs2-mediated activation;ARMS-mediated activation;Gastrin-CREB signalling pathway via PKC and MAPK;Interleukin receptor SHC signaling;DAP12 signaling;Signaling by PDGF;VEGFA-VEGFR2 Pathway;Signaling by ERBB2;IRS-related events triggered by IGF1R;Cell surface interactions at the vascular wall 
                                               hsa04010;hsa04012;hsa04014;hsa04015;hsa04062;hsa04068;hsa04071;hsa04072;hsa04151;hsa04211;hsa04213;hsa04320;hsa04360;hsa04370;hsa04530;hsa04540;hsa04550;hsa04650;hsa04660;hsa04662;hsa04664;hsa04720;hsa04722;hsa04725;hsa04726;hsa04730;hsa04810;hsa04910;hsa04912;hsa04914;hsa04915;hsa04916;hsa04917;hsa04919;hsa04921;hsa04933;hsa04960;hsa05034;hsa05160;hsa05161;hsa05166;hsa05200;hsa05203;hsa05205;hsa05206;hsa05210;hsa05211;hsa05212;hsa05213;hsa05214;hsa05215;hsa05216;hsa05218;hsa05219;hsa05220;hsa05221;hsa05223;hsa05230;hsa05231 
                                               KRAS 
                                               NA 
                                               slateblue2 
                                               16 
                                               -3.31730769230769 
                                             
                                             
                                               HRAS-mRNA 
                                               -0.461 
                                               0.267 
                                               -0.985 
                                               0.0621 
                                               0.726 
                                               0.505 
                                               1.04 
                                               0.0924 
                                               0.68 
                                               lm.nb 
                                               ARMS-mediated activation, CD209 (DC-SIGN) signaling, Cell surface interactions at the vascular wall, Constitutive Signaling by EGFRvIII, CREB phosphorylation through the activation of Ras, DAP12 signaling, Downstream signal transduction, FCERI mediated MAPK activation, Frs2-mediated activation, Gastrin-CREB signalling pathway via PKC and MAPK, Interleukin receptor SHC signaling, IRS-related events triggered by IGF1R, MAP2K and MAPK activation, RAFMAP kinase cascade, SHC-mediated cascade.CL.FGFR1, SHC-related events triggered by IGF1R, Signaling by ERBB2, Signaling by FGFR1 in disease, Signaling by FGFR2 in disease, Signaling by FGFR3 in disease, Signaling by FGFR4 in disease, Signaling by PDGF, Signaling by SCF-KIT, Tie2 Signaling, VEGFA-VEGFR2 Pathway, VEGFR2 mediated cell proliferation 
                                               SEBELIUS_2_C4636 
                                               HRAS 
                                               mRNA 
                                               0 
                                               
                                               
                                               Signaling by FGFR2 in disease;Regulation of RAS by GAPs;SHC-mediated cascade.CL.FGFR3;Signaling by FGFR3 in disease;MAP2K and MAPK activation;SHC-related events triggered by IGF1R;Signaling by FGFR4 in disease;SHC-mediated cascade.CL.FGFR1;Tie2 Signaling;Signaling by FGFR1 in disease;Activation of RAS in B cells;SHC-mediated cascade.CL.FGFR4;Constitutive Signaling by EGFRvIII;CD209 (DC-SIGN) signaling;Ras activation uopn Ca2.PLS. infux through NMDA receptor;FCERI mediated MAPK activation;RAFMAP kinase cascade;Downstream signal transduction;VEGFR2 mediated cell proliferation;Signaling by SCF-KIT;Frs2-mediated activation;ARMS-mediated activation;Gastrin-CREB signalling pathway via PKC and MAPK;Interleukin receptor SHC signaling;DAP12 signaling;Signaling by PDGF;VEGFA-VEGFR2 Pathway;Signaling by ERBB2;IRS-related events triggered by IGF1R;Cell surface interactions at the vascular wall;CREB phosphorylation through the activation of Ras 
                                               hsa04010;hsa04012;hsa04014;hsa04015;hsa04062;hsa04068;hsa04071;hsa04072;hsa04144;hsa04151;hsa04211;hsa04213;hsa04360;hsa04370;hsa04510;hsa04530;hsa04540;hsa04550;hsa04630;hsa04650;hsa04660;hsa04662;hsa04664;hsa04720;hsa04722;hsa04725;hsa04726;hsa04730;hsa04810;hsa04910;hsa04912;hsa04915;hsa04916;hsa04917;hsa04919;hsa04921;hsa04933;hsa05034;hsa05160;hsa05161;hsa05166;hsa05200;hsa05203;hsa05205;hsa05206;hsa05211;hsa05213;hsa05214;hsa05215;hsa05216;hsa05218;hsa05219;hsa05220;hsa05221;hsa05223;hsa05230;hsa05231 
                                               HRAS 
                                               NA 
                                               azure4 
                                               1 
                                               -1.72659176029963 
                                             
                                             
                                               FYN-mRNA 
                                               0.596 
                                               0.365 
                                               -0.119 
                                               1.31 
                                               1.51 
                                               0.921 
                                               2.48 
                                               0.111 
                                               0.804 
                                               lm.nb 
                                               ARMS-mediated activation, CD209 (DC-SIGN) signaling, CD28 co-stimulation, CD28 dependent PI3KAkt signaling, CD28 dependent Vav1 pathway, Cell surface interactions at the vascular wall, Constitutive Signaling by Aberrant PI3K in Cancer, CTLA4 inhibitory signaling, DAP12 signaling, Downstream signal transduction, EPH-ephrin mediated repulsion of cells, FCERI mediated MAPK activation, Frs2-mediated activation, GAB1 signalosome, Gastrin-CREB signalling pathway via PKC and MAPK, GPVI-mediated activation cascade, Interleukin receptor SHC signaling, IRS-related events triggered by IGF1R, Nephrin interactions, PI3KAKT activation, RAFMAP kinase cascade, Regulation of signaling by CBL, Role of LAT2NTALLAB on calcium mobilization, Sema3A PAK dependent Axon repulsion, Signaling by ERBB2, Signaling by PDGF, Signaling by SCF-KIT, VEGFA-VEGFR2 Pathway, VEGFR2 mediated cell proliferation 
                                               SEBELIUS_2_C4636 
                                               FYN 
                                               mRNA 
                                               0 
                                               
                                               
                                               EPH-ephrin mediated repulsion of cells;Regulation of signaling by CBL;FCGR activation;Antigen activates B Cell Receptor (BCR) leading to generation of second messengers;Dectin-2 family;RAFMAP kinase cascade;Constitutive Signaling by Aberrant PI3K in Cancer;CTLA4 inhibitory signaling;CD28 dependent PI3KAkt signaling;PECAM1 interactions;CD28 dependent Vav1 pathway;GPVI-mediated activation cascade;Sema3A PAK dependent Axon repulsion;Nephrin interactions;SEMA3A-Plexin repulsion signaling by inhibiting Integrin adhesion;CRMPs in Sema3A signaling;CD209 (DC-SIGN) signaling;Downstream signal transduction;GAB1 signalosome;Signaling by SCF-KIT;PI3KAKT activation;Role of LAT2NTALLAB on calcium mobilization;DAP12 signaling;Signaling by PDGF;Signaling by ERBB2;VEGFA-VEGFR2 Pathway;VEGFR2 mediated cell proliferation;Frs2-mediated activation;ARMS-mediated activation;Gastrin-CREB signalling pathway via PKC and MAPK;Interleukin receptor SHC signaling;FCERI mediated MAPK activation;IRS-related events triggered by IGF1R;CD28 co-stimulation;Cell surface interactions at the vascular wall 
                                               hsa04071;hsa04072;hsa04360;hsa04380;hsa04510;hsa04520;hsa04611;hsa04650;hsa04660;hsa04664;hsa04725;hsa05020;hsa05130;hsa05162;hsa05416 
                                               FYN 
                                               NA 
                                               azure4 
                                               1 
                                               1.63287671232877 
                                             
                                             
                                               RAF1-mRNA 
                                               0.103 
                                               0.0914 
                                               -0.0765 
                                               0.282 
                                               1.07 
                                               0.948 
                                               1.22 
                                               0.269 
                                               1 
                                               loglinear 
                                               ARMS-mediated activation, CD209 (DC-SIGN) signaling, CREB phosphorylation through the activation of Ras, DAP12 signaling, Downstream signal transduction, FCERI mediated MAPK activation, Frs2-mediated activation, Gastrin-CREB signalling pathway via PKC and MAPK, Interleukin receptor SHC signaling, IRS-related events triggered by IGF1R, Negative feedback regulation of MAPK pathway, Negative regulation of MAPK pathway, RAF activation, RAFMAP kinase cascade, Signaling by ERBB2, Signaling by PDGF, Signaling by SCF-KIT, VEGFA-VEGFR2 Pathway, VEGFR2 mediated cell proliferation 
                                               SEBELIUS_2_C4636 
                                               RAF1 
                                               mRNA 
                                               0 
                                               
                                               
                                               RAF activation;Stimuli-sensing channels;CREB phosphorylation through the activation of Ras;Negative feedback regulation of MAPK pathway;CD209 (DC-SIGN) signaling;RAFMAP kinase cascade;Downstream signal transduction;VEGFR2 mediated cell proliferation;Signaling by SCF-KIT;Frs2-mediated activation;ARMS-mediated activation;Gastrin-CREB signalling pathway via PKC and MAPK;Interleukin receptor SHC signaling;FCERI mediated MAPK activation;DAP12 signaling;Signaling by PDGF;VEGFA-VEGFR2 Pathway;Signaling by ERBB2;IRS-related events triggered by IGF1R;Negative regulation of MAPK pathway 
                                               hsa04010;hsa04012;hsa04014;hsa04015;hsa04022;hsa04024;hsa04062;hsa04068;hsa04071;hsa04072;hsa04151;hsa04270;hsa04370;hsa04510;hsa04540;hsa04550;hsa04630;hsa04650;hsa04660;hsa04662;hsa04664;hsa04666;hsa04720;hsa04722;hsa04726;hsa04730;hsa04810;hsa04910;hsa04912;hsa04914;hsa04915;hsa04916;hsa04917;hsa04919;hsa04921;hsa05034;hsa05152;hsa05160;hsa05161;hsa05164;hsa05200;hsa05205;hsa05206;hsa05210;hsa05211;hsa05212;hsa05213;hsa05214;hsa05215;hsa05218;hsa05219;hsa05220;hsa05221;hsa05223;hsa05230;hsa05231 
                                               RAF1 
                                               NA 
                                               azure4 
                                               1 
                                               1.12691466083151 
                                             
                                           
                                           DE Results - CD209 (DC-SIGN) signaling genes - Line: HEPG2 vs.Fibro  Table displaying each sample's global significance scores and directed global significance scores as defined in the heatmaps above. The global significance score is calculated as the square root of the mean squared t-statistic for the genes in a gene set, with t-statistics coming from the linear regression underlying our differential expression analysis. The directed global significance score is calculated as the square root of the mean signed squared t-statistic for the genes in a gene set, with t-statistics coming from the linear regression underlying our differential expression analysis.   
                                      
                                       
                                  
                                
                                
                                  
                                       
                                      
                                           Volcano Plot: Line: HT29 vs.Fibro   More Plot Information                Volcano Plot: Line: HT29 vs.Fibro  Volcano plot displaying each gene's -log10(p-value) and log2 fold change for the selected covariate. Highly statistically significant genes fall at the top of the plot, and highly differentially expressed genes fall to either side. Genes within the selected gene set are highlighted in orange. Horizontal lines indicate various False Discovery Rate (FDR) thresholds.   
                                           DE Results - CD209 (DC-SIGN) signaling genes - Line: HT29 vs.Fibro   More Plot Information      Download CSV Data     
                                           
                                             
                                             
                                           
                                           
                                             
                                               Probe Label 
                                               Log2 fold change 
                                               std error (log2) 
                                               Lower confidence limit (log2) 
                                               Upper confidence limit (log2) 
                                               Linear fold change 
                                               Lower confidence limit (linear) 
                                               Upper confidence limit (linear) 
                                               P-value 
                                               BY.p.value 
                                               method 
                                               Gene.sets 
                                               Codeset.Name 
                                               Probe.Label 
                                               Analyte.Type 
                                               Is.Control 
                                               Control.Type 
                                               Related.Probes 
                                               Probe.Annotation 
                                               KEGG.Pathways 
                                               Official.Gene.Name 
                                               Control.Conc 
                                               volcanocol 
                                               volcanopch 
                                               tstats.all 
                                             
                                           
                                           
                                             
                                               PAK1-mRNA 
                                               2.31 
                                               0.165 
                                               1.99 
                                               2.63 
                                               4.96 
                                               3.96 
                                               6.2 
                                               2.18e-16 
                                               5.51e-15 
                                               lm.nb 
                                               Activation of Rac, CD209 (DC-SIGN) signaling, CD28 co-stimulation, CD28 dependent Vav1 pathway, EPHB-mediated forward signaling, Ephrin signaling, FCERI mediated MAPK activation, MAPK6MAPK4 signaling, Regulation of actin dynamics for phagocytic cup formation, RHO GTPases activate PAKs, Sema3A PAK dependent Axon repulsion, VEGFA-VEGFR2 Pathway, VEGFR2 mediated vascular permeability 
                                               SEBELIUS_2_C4636 
                                               PAK1 
                                               mRNA 
                                               0 
                                               
                                               
                                               EPHB-mediated forward signaling;Ephrin signaling;MAPK6MAPK4 signaling;DSCAM interactions;Activation of Rac;VEGFR2 mediated vascular permeability;RHO GTPases activate PAKs;Sema3A PAK dependent Axon repulsion;FCERI mediated MAPK activation;Regulation of actin dynamics for phagocytic cup formation;CD209 (DC-SIGN) signaling;CD28 dependent Vav1 pathway;VEGFA-VEGFR2 Pathway;CD28 co-stimulation 
                                               hsa04010;hsa04012;hsa04014;hsa04024;hsa04062;hsa04360;hsa04510;hsa04650;hsa04660;hsa04666;hsa04810;hsa05120;hsa05205;hsa05211 
                                               PAK1 
                                               NA 
                                               slateblue1 
                                               16 
                                               14 
                                             
                                             
                                               KRAS-mRNA 
                                               0.994 
                                               0.0971 
                                               0.803 
                                               1.18 
                                               1.99 
                                               1.75 
                                               2.27 
                                               2.41e-12 
                                               3.76e-11 
                                               lm.nb 
                                               ARMS-mediated activation, CD209 (DC-SIGN) signaling, Cell surface interactions at the vascular wall, Constitutive Signaling by EGFRvIII, DAP12 signaling, Downstream signal transduction, FCERI mediated MAPK activation, Frs2-mediated activation, Gastrin-CREB signalling pathway via PKC and MAPK, Interleukin receptor SHC signaling, IRS-related events triggered by IGF1R, MAP2K and MAPK activation, RAFMAP kinase cascade, SHC-mediated cascade.CL.FGFR1, SHC-related events triggered by IGF1R, Signaling by ERBB2, Signaling by FGFR1 in disease, Signaling by FGFR2 in disease, Signaling by FGFR3 in disease, Signaling by FGFR4 in disease, Signaling by PDGF, Signaling by SCF-KIT, Tie2 Signaling, VEGFA-VEGFR2 Pathway, VEGFR2 mediated cell proliferation 
                                               SEBELIUS_2_C4636 
                                               KRAS 
                                               mRNA 
                                               0 
                                               
                                               
                                               Signaling by FGFR2 in disease;Regulation of RAS by GAPs;SHC-mediated cascade.CL.FGFR3;Signaling by FGFR3 in disease;MAP2K and MAPK activation;SHC-related events triggered by IGF1R;Signaling by FGFR4 in disease;SHC-mediated cascade.CL.FGFR1;Tie2 Signaling;Signaling by FGFR1 in disease;Activation of RAS in B cells;SHC-mediated cascade.CL.FGFR4;Constitutive Signaling by EGFRvIII;CD209 (DC-SIGN) signaling;FCERI mediated MAPK activation;RAFMAP kinase cascade;Downstream signal transduction;VEGFR2 mediated cell proliferation;Signaling by SCF-KIT;Frs2-mediated activation;ARMS-mediated activation;Gastrin-CREB signalling pathway via PKC and MAPK;Interleukin receptor SHC signaling;DAP12 signaling;Signaling by PDGF;VEGFA-VEGFR2 Pathway;Signaling by ERBB2;IRS-related events triggered by IGF1R;Cell surface interactions at the vascular wall 
                                               hsa04010;hsa04012;hsa04014;hsa04015;hsa04062;hsa04068;hsa04071;hsa04072;hsa04151;hsa04211;hsa04213;hsa04320;hsa04360;hsa04370;hsa04530;hsa04540;hsa04550;hsa04650;hsa04660;hsa04662;hsa04664;hsa04720;hsa04722;hsa04725;hsa04726;hsa04730;hsa04810;hsa04910;hsa04912;hsa04914;hsa04915;hsa04916;hsa04917;hsa04919;hsa04921;hsa04933;hsa04960;hsa05034;hsa05160;hsa05161;hsa05166;hsa05200;hsa05203;hsa05205;hsa05206;hsa05210;hsa05211;hsa05212;hsa05213;hsa05214;hsa05215;hsa05216;hsa05218;hsa05219;hsa05220;hsa05221;hsa05223;hsa05230;hsa05231 
                                               KRAS 
                                               NA 
                                               slateblue1 
                                               16 
                                               10.2368692070031 
                                             
                                             
                                               NRAS-mRNA 
                                               0.713 
                                               0.0909 
                                               0.535 
                                               0.892 
                                               1.64 
                                               1.45 
                                               1.86 
                                               2.17e-09 
                                               2.73e-08 
                                               lm.nb 
                                               ARMS-mediated activation, CD209 (DC-SIGN) signaling, Cell surface interactions at the vascular wall, Constitutive Signaling by EGFRvIII, DAP12 signaling, Downstream signal transduction, FCERI mediated MAPK activation, Frs2-mediated activation, Gastrin-CREB signalling pathway via PKC and MAPK, Interleukin receptor SHC signaling, IRS-related events triggered by IGF1R, MAP2K and MAPK activation, RAFMAP kinase cascade, SHC-mediated cascade.CL.FGFR1, SHC-related events triggered by IGF1R, Signaling by ERBB2, Signaling by FGFR1 in disease, Signaling by FGFR2 in disease, Signaling by FGFR3 in disease, Signaling by FGFR4 in disease, Signaling by PDGF, Signaling by SCF-KIT, Tie2 Signaling, VEGFA-VEGFR2 Pathway, VEGFR2 mediated cell proliferation 
                                               SEBELIUS_2_C4636 
                                               NRAS 
                                               mRNA 
                                               0 
                                               
                                               
                                               Signaling by FGFR2 in disease;Regulation of RAS by GAPs;SHC-mediated cascade.CL.FGFR3;Signaling by FGFR3 in disease;MAP2K and MAPK activation;SHC-related events triggered by IGF1R;Signaling by FGFR4 in disease;SHC-mediated cascade.CL.FGFR1;Tie2 Signaling;Signaling by FGFR1 in disease;Activation of RAS in B cells;SHC-mediated cascade.CL.FGFR4;Constitutive Signaling by EGFRvIII;CD209 (DC-SIGN) signaling;FCERI mediated MAPK activation;RAFMAP kinase cascade;Downstream signal transduction;VEGFR2 mediated cell proliferation;Signaling by SCF-KIT;Frs2-mediated activation;ARMS-mediated activation;Gastrin-CREB signalling pathway via PKC and MAPK;Interleukin receptor SHC signaling;DAP12 signaling;Signaling by PDGF;VEGFA-VEGFR2 Pathway;Signaling by ERBB2;IRS-related events triggered by IGF1R;Cell surface interactions at the vascular wall 
                                               hsa04010;hsa04012;hsa04014;hsa04015;hsa04062;hsa04068;hsa04071;hsa04072;hsa04151;hsa04211;hsa04213;hsa04360;hsa04370;hsa04530;hsa04540;hsa04550;hsa04650;hsa04660;hsa04662;hsa04664;hsa04720;hsa04722;hsa04725;hsa04726;hsa04730;hsa04810;hsa04910;hsa04912;hsa04915;hsa04916;hsa04917;hsa04919;hsa04921;hsa04933;hsa05034;hsa05160;hsa05161;hsa05166;hsa05200;hsa05203;hsa05205;hsa05206;hsa05211;hsa05213;hsa05214;hsa05215;hsa05216;hsa05218;hsa05219;hsa05220;hsa05221;hsa05223;hsa05230;hsa05231 
                                               NRAS 
                                               NA 
                                               slateblue1 
                                               16 
                                               7.84378437843784 
                                             
                                             
                                               RAF1-mRNA 
                                               -0.587 
                                               0.0861 
                                               -0.756 
                                               -0.418 
                                               0.666 
                                               0.592 
                                               0.748 
                                               4.99e-08 
                                               5.81e-07 
                                               loglinear 
                                               ARMS-mediated activation, CD209 (DC-SIGN) signaling, CREB phosphorylation through the activation of Ras, DAP12 signaling, Downstream signal transduction, FCERI mediated MAPK activation, Frs2-mediated activation, Gastrin-CREB signalling pathway via PKC and MAPK, Interleukin receptor SHC signaling, IRS-related events triggered by IGF1R, Negative feedback regulation of MAPK pathway, Negative regulation of MAPK pathway, RAF activation, RAFMAP kinase cascade, Signaling by ERBB2, Signaling by PDGF, Signaling by SCF-KIT, VEGFA-VEGFR2 Pathway, VEGFR2 mediated cell proliferation 
                                               SEBELIUS_2_C4636 
                                               RAF1 
                                               mRNA 
                                               0 
                                               
                                               
                                               RAF activation;Stimuli-sensing channels;CREB phosphorylation through the activation of Ras;Negative feedback regulation of MAPK pathway;CD209 (DC-SIGN) signaling;RAFMAP kinase cascade;Downstream signal transduction;VEGFR2 mediated cell proliferation;Signaling by SCF-KIT;Frs2-mediated activation;ARMS-mediated activation;Gastrin-CREB signalling pathway via PKC and MAPK;Interleukin receptor SHC signaling;FCERI mediated MAPK activation;DAP12 signaling;Signaling by PDGF;VEGFA-VEGFR2 Pathway;Signaling by ERBB2;IRS-related events triggered by IGF1R;Negative regulation of MAPK pathway 
                                               hsa04010;hsa04012;hsa04014;hsa04015;hsa04022;hsa04024;hsa04062;hsa04068;hsa04071;hsa04072;hsa04151;hsa04270;hsa04370;hsa04510;hsa04540;hsa04550;hsa04630;hsa04650;hsa04660;hsa04662;hsa04664;hsa04666;hsa04720;hsa04722;hsa04726;hsa04730;hsa04810;hsa04910;hsa04912;hsa04914;hsa04915;hsa04916;hsa04917;hsa04919;hsa04921;hsa05034;hsa05152;hsa05160;hsa05161;hsa05164;hsa05200;hsa05205;hsa05206;hsa05210;hsa05211;hsa05212;hsa05213;hsa05214;hsa05215;hsa05218;hsa05219;hsa05220;hsa05221;hsa05223;hsa05230;hsa05231 
                                               RAF1 
                                               NA 
                                               slateblue1 
                                               16 
                                               -6.81765389082462 
                                             
                                             
                                               HRAS-mRNA 
                                               1.24 
                                               0.25 
                                               0.75 
                                               1.73 
                                               2.36 
                                               1.68 
                                               3.32 
                                               1.59e-05 
                                               0.000154 
                                               lm.nb 
                                               ARMS-mediated activation, CD209 (DC-SIGN) signaling, Cell surface interactions at the vascular wall, Constitutive Signaling by EGFRvIII, CREB phosphorylation through the activation of Ras, DAP12 signaling, Downstream signal transduction, FCERI mediated MAPK activation, Frs2-mediated activation, Gastrin-CREB signalling pathway via PKC and MAPK, Interleukin receptor SHC signaling, IRS-related events triggered by IGF1R, MAP2K and MAPK activation, RAFMAP kinase cascade, SHC-mediated cascade.CL.FGFR1, SHC-related events triggered by IGF1R, Signaling by ERBB2, Signaling by FGFR1 in disease, Signaling by FGFR2 in disease, Signaling by FGFR3 in disease, Signaling by FGFR4 in disease, Signaling by PDGF, Signaling by SCF-KIT, Tie2 Signaling, VEGFA-VEGFR2 Pathway, VEGFR2 mediated cell proliferation 
                                               SEBELIUS_2_C4636 
                                               HRAS 
                                               mRNA 
                                               0 
                                               
                                               
                                               Signaling by FGFR2 in disease;Regulation of RAS by GAPs;SHC-mediated cascade.CL.FGFR3;Signaling by FGFR3 in disease;MAP2K and MAPK activation;SHC-related events triggered by IGF1R;Signaling by FGFR4 in disease;SHC-mediated cascade.CL.FGFR1;Tie2 Signaling;Signaling by FGFR1 in disease;Activation of RAS in B cells;SHC-mediated cascade.CL.FGFR4;Constitutive Signaling by EGFRvIII;CD209 (DC-SIGN) signaling;Ras activation uopn Ca2.PLS. infux through NMDA receptor;FCERI mediated MAPK activation;RAFMAP kinase cascade;Downstream signal transduction;VEGFR2 mediated cell proliferation;Signaling by SCF-KIT;Frs2-mediated activation;ARMS-mediated activation;Gastrin-CREB signalling pathway via PKC and MAPK;Interleukin receptor SHC signaling;DAP12 signaling;Signaling by PDGF;VEGFA-VEGFR2 Pathway;Signaling by ERBB2;IRS-related events triggered by IGF1R;Cell surface interactions at the vascular wall;CREB phosphorylation through the activation of Ras 
                                               hsa04010;hsa04012;hsa04014;hsa04015;hsa04062;hsa04068;hsa04071;hsa04072;hsa04144;hsa04151;hsa04211;hsa04213;hsa04360;hsa04370;hsa04510;hsa04530;hsa04540;hsa04550;hsa04630;hsa04650;hsa04660;hsa04662;hsa04664;hsa04720;hsa04722;hsa04725;hsa04726;hsa04730;hsa04810;hsa04910;hsa04912;hsa04915;hsa04916;hsa04917;hsa04919;hsa04921;hsa04933;hsa05034;hsa05160;hsa05161;hsa05166;hsa05200;hsa05203;hsa05205;hsa05206;hsa05211;hsa05213;hsa05214;hsa05215;hsa05216;hsa05218;hsa05219;hsa05220;hsa05221;hsa05223;hsa05230;hsa05231 
                                               HRAS 
                                               NA 
                                               slateblue1 
                                               16 
                                               4.96 
                                             
                                             
                                               FYN-mRNA 
                                               -1.21 
                                               0.345 
                                               -1.89 
                                               -0.534 
                                               0.432 
                                               0.271 
                                               0.69 
                                               0.00119 
                                               0.01 
                                               lm.nb 
                                               ARMS-mediated activation, CD209 (DC-SIGN) signaling, CD28 co-stimulation, CD28 dependent PI3KAkt signaling, CD28 dependent Vav1 pathway, Cell surface interactions at the vascular wall, Constitutive Signaling by Aberrant PI3K in Cancer, CTLA4 inhibitory signaling, DAP12 signaling, Downstream signal transduction, EPH-ephrin mediated repulsion of cells, FCERI mediated MAPK activation, Frs2-mediated activation, GAB1 signalosome, Gastrin-CREB signalling pathway via PKC and MAPK, GPVI-mediated activation cascade, Interleukin receptor SHC signaling, IRS-related events triggered by IGF1R, Nephrin interactions, PI3KAKT activation, RAFMAP kinase cascade, Regulation of signaling by CBL, Role of LAT2NTALLAB on calcium mobilization, Sema3A PAK dependent Axon repulsion, Signaling by ERBB2, Signaling by PDGF, Signaling by SCF-KIT, VEGFA-VEGFR2 Pathway, VEGFR2 mediated cell proliferation 
                                               SEBELIUS_2_C4636 
                                               FYN 
                                               mRNA 
                                               0 
                                               
                                               
                                               EPH-ephrin mediated repulsion of cells;Regulation of signaling by CBL;FCGR activation;Antigen activates B Cell Receptor (BCR) leading to generation of second messengers;Dectin-2 family;RAFMAP kinase cascade;Constitutive Signaling by Aberrant PI3K in Cancer;CTLA4 inhibitory signaling;CD28 dependent PI3KAkt signaling;PECAM1 interactions;CD28 dependent Vav1 pathway;GPVI-mediated activation cascade;Sema3A PAK dependent Axon repulsion;Nephrin interactions;SEMA3A-Plexin repulsion signaling by inhibiting Integrin adhesion;CRMPs in Sema3A signaling;CD209 (DC-SIGN) signaling;Downstream signal transduction;GAB1 signalosome;Signaling by SCF-KIT;PI3KAKT activation;Role of LAT2NTALLAB on calcium mobilization;DAP12 signaling;Signaling by PDGF;Signaling by ERBB2;VEGFA-VEGFR2 Pathway;VEGFR2 mediated cell proliferation;Frs2-mediated activation;ARMS-mediated activation;Gastrin-CREB signalling pathway via PKC and MAPK;Interleukin receptor SHC signaling;FCERI mediated MAPK activation;IRS-related events triggered by IGF1R;CD28 co-stimulation;Cell surface interactions at the vascular wall 
                                               hsa04071;hsa04072;hsa04360;hsa04380;hsa04510;hsa04520;hsa04611;hsa04650;hsa04660;hsa04664;hsa04725;hsa05020;hsa05130;hsa05162;hsa05416 
                                               FYN 
                                               NA 
                                               slateblue2 
                                               16 
                                               -3.50724637681159 
                                             
                                             
                                               EP300-mRNA 
                                               0.447 
                                               0.143 
                                               0.166 
                                               0.728 
                                               1.36 
                                               1.12 
                                               1.66 
                                               0.00348 
                                               0.0287 
                                               lm.nb 
                                               CD209 (DC-SIGN) signaling, Circadian Clock 
                                               SEBELIUS_2_C4636 
                                               EP300 
                                               mRNA 
                                               0 
                                               
                                               
                                               LRR FLII-interacting protein 1 (LRRFIP1) activates type I IFN production;TRAF6 mediated IRF7 activation;Constitutive Signaling by NOTCH1 PEST Domain Mutants;Polo-like kinase mediated events;B-WICH complex positively regulates rRNA expression;CD209 (DC-SIGN) signaling;Circadian Clock 
                                               hsa04024;hsa04066;hsa04068;hsa04110;hsa04310;hsa04330;hsa04350;hsa04520;hsa04630;hsa04720;hsa04916;hsa04919;hsa04922;hsa05016;hsa05152;hsa05161;hsa05164;hsa05166;hsa05168;hsa05169;hsa05200;hsa05203;hsa05206;hsa05211;hsa05215 
                                               EP300 
                                               NA 
                                               slateblue2 
                                               16 
                                               3.12587412587413 
                                             
                                             
                                               PAK2-mRNA 
                                               0.221 
                                               0.0777 
                                               0.0684 
                                               0.373 
                                               1.17 
                                               1.05 
                                               1.3 
                                               0.00728 
                                               0.0571 
                                               loglinear 
                                               Activation of Rac, CD209 (DC-SIGN) signaling, CD28 co-stimulation, CD28 dependent Vav1 pathway, Ephrin signaling, RHO GTPases activate PAKs, Sema3A PAK dependent Axon repulsion, VEGFA-VEGFR2 Pathway, VEGFR2 mediated vascular permeability 
                                               SEBELIUS_2_C4636 
                                               PAK2 
                                               mRNA 
                                               0 
                                               
                                               
                                               Nef and signal transduction;CD28 dependent Vav1 pathway;Stimulation of the cell death response by PAK-2p34;Ephrin signaling;Activation of Rac;RHO GTPases activate PAKs;VEGFR2 mediated vascular permeability;Sema3A PAK dependent Axon repulsion;CD209 (DC-SIGN) signaling;VEGFA-VEGFR2 Pathway;CD28 co-stimulation 
                                               hsa04010;hsa04012;hsa04014;hsa04360;hsa04510;hsa04660;hsa04810;hsa05211 
                                               PAK2 
                                               NA 
                                               slateblue3 
                                               16 
                                               2.84427284427284 
                                             
                                             
                                               CREBBP-mRNA 
                                               0.166 
                                               0.131 
                                               -0.0915 
                                               0.423 
                                               1.12 
                                               0.939 
                                               1.34 
                                               0.214 
                                               1 
                                               lm.nb 
                                               CD209 (DC-SIGN) signaling, Circadian Clock 
                                               SEBELIUS_2_C4636 
                                               CREBBP 
                                               mRNA 
                                               0 
                                               
                                               
                                               YAP1- and WWTR1 (TAZ)-stimulated gene expression;Activation of gene expression by SREBF (SREBP);Transcriptional activation of mitochondrial biogenesis;Constitutive Signaling by NOTCH1 HD.PLS.PEST Domain Mutants;Constitutive Signaling by NOTCH1 PEST Domain Mutants;CD209 (DC-SIGN) signaling;Circadian Clock 
                                               hsa04024;hsa04066;hsa04068;hsa04110;hsa04310;hsa04330;hsa04350;hsa04520;hsa04630;hsa04720;hsa04916;hsa04919;hsa04922;hsa05016;hsa05152;hsa05161;hsa05164;hsa05166;hsa05168;hsa05169;hsa05200;hsa05203;hsa05206;hsa05211;hsa05215 
                                               CREBBP 
                                               NA 
                                               azure4 
                                               1 
                                               1.26717557251908 
                                             
                                           
                                           DE Results - CD209 (DC-SIGN) signaling genes - Line: HT29 vs.Fibro  Table displaying each sample's global significance scores and directed global significance scores as defined in the heatmaps above. The global significance score is calculated as the square root of the mean squared t-statistic for the genes in a gene set, with t-statistics coming from the linear regression underlying our differential expression analysis. The directed global significance score is calculated as the square root of the mean signed squared t-statistic for the genes in a gene set, with t-statistics coming from the linear regression underlying our differential expression analysis.   
                                      
                                       
                                  
                                
                              
                            
                           
                            
                          
                           
                      
                    
                    
                      
                           
                          
                               Day    Line     
                              
                                  Day  
                              
                               
                              
                                  Line: differential expression in Caco2 vs. baseline of Fibro  
                                  Line: differential expression in HEPG2 vs. baseline of Fibro  
                                  Line: differential expression in HT29 vs. baseline of Fibro  
                              
                              
                            
                              
                                
                                  
                                       
                                      
                                           Volcano Plot: Day   More Plot Information                Volcano Plot: Day  Volcano plot displaying each gene's -log10(p-value) and log2 fold change for the selected covariate. Highly statistically significant genes fall at the top of the plot, and highly differentially expressed genes fall to either side. Genes within the selected gene set are highlighted in orange. Horizontal lines indicate various False Discovery Rate (FDR) thresholds.   
                                           DE Results - CD28 co-stimulation genes - Day   More Plot Information      Download CSV Data     
                                           
                                             
                                             
                                           
                                           
                                             
                                               Probe Label 
                                               Log2 fold change 
                                               std error (log2) 
                                               Lower confidence limit (log2) 
                                               Upper confidence limit (log2) 
                                               Linear fold change 
                                               Lower confidence limit (linear) 
                                               Upper confidence limit (linear) 
                                               P-value 
                                               BY.p.value 
                                               method 
                                               Gene.sets 
                                               Codeset.Name 
                                               Probe.Label 
                                               Analyte.Type 
                                               Is.Control 
                                               Control.Type 
                                               Related.Probes 
                                               Probe.Annotation 
                                               KEGG.Pathways 
                                               Official.Gene.Name 
                                               Control.Conc 
                                               volcanocol 
                                               volcanopch 
                                               tstats.all 
                                             
                                           
                                           
                                             
                                               PAK1-mRNA 
                                               0.128 
                                               0.0211 
                                               0.0869 
                                               0.169 
                                               1.09 
                                               1.06 
                                               1.12 
                                               4.84e-07 
                                               6.25e-05 
                                               lm.nb 
                                               Activation of Rac, CD209 (DC-SIGN) signaling, CD28 co-stimulation, CD28 dependent Vav1 pathway, EPHB-mediated forward signaling, Ephrin signaling, FCERI mediated MAPK activation, MAPK6MAPK4 signaling, Regulation of actin dynamics for phagocytic cup formation, RHO GTPases activate PAKs, Sema3A PAK dependent Axon repulsion, VEGFA-VEGFR2 Pathway, VEGFR2 mediated vascular permeability 
                                               SEBELIUS_2_C4636 
                                               PAK1 
                                               mRNA 
                                               0 
                                               
                                               
                                               EPHB-mediated forward signaling;Ephrin signaling;MAPK6MAPK4 signaling;DSCAM interactions;Activation of Rac;VEGFR2 mediated vascular permeability;RHO GTPases activate PAKs;Sema3A PAK dependent Axon repulsion;FCERI mediated MAPK activation;Regulation of actin dynamics for phagocytic cup formation;CD209 (DC-SIGN) signaling;CD28 dependent Vav1 pathway;VEGFA-VEGFR2 Pathway;CD28 co-stimulation 
                                               hsa04010;hsa04012;hsa04014;hsa04024;hsa04062;hsa04360;hsa04510;hsa04650;hsa04660;hsa04666;hsa04810;hsa05120;hsa05205;hsa05211 
                                               PAK1 
                                               NA 
                                               slateblue1 
                                               16 
                                               6.06635071090047 
                                             
                                             
                                               PIK3R1-mRNA 
                                               0.0895 
                                               0.021 
                                               0.0484 
                                               0.131 
                                               1.06 
                                               1.03 
                                               1.09 
                                               0.000131 
                                               0.00452 
                                               lm.nb 
                                               CD28 co-stimulation, CD28 dependent PI3KAkt signaling, Cell surface interactions at the vascular wall, Constitutive Signaling by Aberrant PI3K in Cancer, Constitutive Signaling by EGFRvIII, DAP12 signaling, Downstream signal transduction, Downstream TCR signaling, G alpha (q) signalling events, G beta.CL.gamma signalling through PI3Kgamma, GAB1 signalosome, Gastrin-CREB signalling pathway via PKC and MAPK, GPVI-mediated activation cascade, Interleukin receptor SHC signaling, IRS-related events triggered by IGF1R, Nephrin interactions, PI3KAKT activation, PIP3 activates AKT signaling, Regulation of signaling by CBL, Role of LAT2NTALLAB on calcium mobilization, Signaling by ERBB2, Signaling by FGFR1 in disease, Signaling by FGFR2 in disease, Signaling by FGFR3 in disease, Signaling by FGFR4 in disease, Signaling by PDGF, Signaling by SCF-KIT, Synthesis of PIPs at the plasma membrane, Tie2 Signaling, VEGFA-VEGFR2 Pathway 
                                               SEBELIUS_2_C4636 
                                               PIK3R1 
                                               mRNA 
                                               0 
                                               
                                               
                                               G alpha (q) signalling events;Downstream TCR signaling;Signaling by FGFR2 in disease;PIP3 activates AKT signaling;VEGFA-VEGFR2 Pathway;Synthesis of PIPs at the plasma membrane;Constitutive Signaling by Ligand-Responsive EGFR Cancer Variants;Interleukin receptor SHC signaling;G beta.CL.gamma signalling through PI3Kgamma;Regulation of signaling by CBL;Role of phospholipids in phagocytosis;Signaling by FGFR3 in disease;Constitutive Signaling by Aberrant PI3K in Cancer;Signaling by FGFR4 in disease;CD28 dependent PI3KAkt signaling;Tie2 Signaling;Signaling by FGFR1 fusion mutants;GP1b-IX-V activation signalling;GPVI-mediated activation cascade;Nephrin interactions;Constitutive Signaling by EGFRvIII;Gastrin-CREB signalling pathway via PKC and MAPK;Downstream signal transduction;GAB1 signalosome;Signaling by SCF-KIT;PI3KAKT activation;Role of LAT2NTALLAB on calcium mobilization;DAP12 signaling;Signaling by PDGF;Signaling by ERBB2;IRS-related events triggered by IGF1R;CD28 co-stimulation;Cell surface interactions at the vascular wall;Signaling by FGFR1 in disease 
                                               hsa04012;hsa04014;hsa04015;hsa04022;hsa04024;hsa04062;hsa04066;hsa04068;hsa04070;hsa04071;hsa04072;hsa04150;hsa04151;hsa04152;hsa04210;hsa04211;hsa04213;hsa04261;hsa04370;hsa04380;hsa04510;hsa04550;hsa04611;hsa04620;hsa04630;hsa04650;hsa04660;hsa04662;hsa04664;hsa04666;hsa04668;hsa04670;hsa04722;hsa04725;hsa04750;hsa04810;hsa04910;hsa04914;hsa04915;hsa04917;hsa04919;hsa04921;hsa04923;hsa04930;hsa04931;hsa04932;hsa04933;hsa04960;hsa04973;hsa05100;hsa05142;hsa05145;hsa05146;hsa05160;hsa05161;hsa05162;hsa05164;hsa05166;hsa05169;hsa05200;hsa05203;hsa05205;hsa05210;hsa05211;hsa05212;hsa05213;hsa05214;hsa05215;hsa05218;hsa05220;hsa05221;hsa05222;hsa05223;hsa05230;hsa05231 
                                               PIK3R1 
                                               NA 
                                               slateblue1 
                                               16 
                                               4.26190476190476 
                                             
                                             
                                               PAK2-mRNA 
                                               -0.0398 
                                               0.00994 
                                               -0.0593 
                                               -0.0203 
                                               0.973 
                                               0.96 
                                               0.986 
                                               0.000289 
                                               0.00836 
                                               loglinear 
                                               Activation of Rac, CD209 (DC-SIGN) signaling, CD28 co-stimulation, CD28 dependent Vav1 pathway, Ephrin signaling, RHO GTPases activate PAKs, Sema3A PAK dependent Axon repulsion, VEGFA-VEGFR2 Pathway, VEGFR2 mediated vascular permeability 
                                               SEBELIUS_2_C4636 
                                               PAK2 
                                               mRNA 
                                               0 
                                               
                                               
                                               Nef and signal transduction;CD28 dependent Vav1 pathway;Stimulation of the cell death response by PAK-2p34;Ephrin signaling;Activation of Rac;RHO GTPases activate PAKs;VEGFR2 mediated vascular permeability;Sema3A PAK dependent Axon repulsion;CD209 (DC-SIGN) signaling;VEGFA-VEGFR2 Pathway;CD28 co-stimulation 
                                               hsa04010;hsa04012;hsa04014;hsa04360;hsa04510;hsa04660;hsa04810;hsa05211 
                                               PAK2 
                                               NA 
                                               slateblue1 
                                               16 
                                               -4.00402414486922 
                                             
                                             
                                               AKT3-mRNA 
                                               0.0985 
                                               0.0254 
                                               0.0488 
                                               0.148 
                                               1.07 
                                               1.03 
                                               1.11 
                                               0.000409 
                                               0.0111 
                                               lm.nb 
                                               Activation of BAD and translocation to mitochondria , CD28 co-stimulation, CD28 dependent PI3KAkt signaling, DAP12 signaling, Downstream signal transduction, G beta.CL.gamma signalling through PI3Kgamma, GAB1 signalosome, GPVI-mediated activation cascade, PI3KAKT activation, PIP3 activates AKT signaling, Role of LAT2NTALLAB on calcium mobilization, Signaling by ERBB2, Signaling by PDGF, Signaling by SCF-KIT, TP53 Regulates Metabolic Genes, VEGFA-VEGFR2 Pathway, VEGFR2 mediated vascular permeability 
                                               SEBELIUS_2_C4636 
                                               AKT3 
                                               mRNA 
                                               0 
                                               
                                               
                                               TP53 Regulates Metabolic Genes;CD28 dependent PI3KAkt signaling;AKT phosphorylates targets in the cytosol;Negative regulation of the PI3KAKT network;Activation of BAD and translocation to mitochondria ;G beta.CL.gamma signalling through PI3Kgamma;AKT phosphorylates targets in the nucleus;VEGFR2 mediated vascular permeability;GPVI-mediated activation cascade;Signaling by ERBB2;PIP3 activates AKT signaling;Downstream signal transduction;GAB1 signalosome;Signaling by SCF-KIT;PI3KAKT activation;Role of LAT2NTALLAB on calcium mobilization;DAP12 signaling;Signaling by PDGF;CD28 co-stimulation;VEGFA-VEGFR2 Pathway 
                                               hsa04010;hsa04012;hsa04014;hsa04015;hsa04022;hsa04024;hsa04062;hsa04066;hsa04068;hsa04071;hsa04072;hsa04150;hsa04151;hsa04152;hsa04210;hsa04211;hsa04213;hsa04261;hsa04370;hsa04380;hsa04510;hsa04530;hsa04550;hsa04611;hsa04620;hsa04630;hsa04660;hsa04662;hsa04664;hsa04666;hsa04668;hsa04722;hsa04725;hsa04728;hsa04910;hsa04914;hsa04915;hsa04917;hsa04919;hsa04920;hsa04922;hsa04923;hsa04931;hsa04932;hsa04933;hsa04973;hsa05142;hsa05145;hsa05152;hsa05160;hsa05161;hsa05162;hsa05164;hsa05166;hsa05169;hsa05200;hsa05205;hsa05210;hsa05211;hsa05212;hsa05213;hsa05214;hsa05215;hsa05218;hsa05220;hsa05221;hsa05222;hsa05223;hsa05230;hsa05231 
                                               AKT3 
                                               NA 
                                               slateblue2 
                                               16 
                                               3.87795275590551 
                                             
                                             
                                               SRC-mRNA 
                                               0.0696 
                                               0.02 
                                               0.0304 
                                               0.109 
                                               1.05 
                                               1.02 
                                               1.08 
                                               0.00132 
                                               0.0268 
                                               lm.nb 
                                               ARMS-mediated activation, CD28 co-stimulation, Cell surface interactions at the vascular wall, CTLA4 inhibitory signaling, DAP12 signaling, Downstream signal transduction, FCERI mediated MAPK activation, Frs2-mediated activation, GAB1 signalosome, Gastrin-CREB signalling pathway via PKC and MAPK, Integrin alphaIIb beta3 signaling, Interleukin receptor SHC signaling, IRS-related events triggered by IGF1R, MAP2K and MAPK activation, Negative regulation of FGFR1 signaling, Negative regulation of FGFR2 signaling, Negative regulation of FGFR3 signaling, Negative regulation of FGFR4 signaling, p130Cas linkage to MAPK signaling for integrins, RAFMAP kinase cascade, Signaling by ERBB2, Signaling by PDGF, Signaling by SCF-KIT, Spry regulation of FGF signaling, VEGFA-VEGFR2 Pathway, VEGFR2 mediated cell proliferation 
                                               SEBELIUS_2_C4636 
                                               SRC 
                                               mRNA 
                                               0 
                                               
                                               
                                               FCGR activation;Spry regulation of FGF signaling;GAB1 signalosome;MAP2K and MAPK activation;CLEC7A (Dectin-1) signaling;CD28 co-stimulation;p130Cas linkage to MAPK signaling for integrins;CTLA4 inhibitory signaling;GRB2.CL.SOS provides linkage to MAPK signaling for Integrins ;Signal regulatory protein (SIRP) family interactions;PECAM1 interactions;Thrombin signalling through proteinase activated receptors (PARs);GP1b-IX-V activation signalling;c-src mediated regulation of Cx43 function and closure of gap junctions;Signaling by ERBB2;VEGFA-VEGFR2 Pathway;Signaling by SCF-KIT;VEGFR2 mediated cell proliferation;RAFMAP kinase cascade;Downstream signal transduction;Frs2-mediated activation;ARMS-mediated activation;Gastrin-CREB signalling pathway via PKC and MAPK;Interleukin receptor SHC signaling;FCERI mediated MAPK activation;DAP12 signaling;Signaling by PDGF;IRS-related events triggered by IGF1R;Negative regulation of FGFR3 signaling;Negative regulation of FGFR4 signaling;Negative regulation of FGFR2 signaling;Negative regulation of FGFR1 signaling;Integrin alphaIIb beta3 signaling;Cell surface interactions at the vascular wall 
                                               hsa04012;hsa04015;hsa04062;hsa04144;hsa04370;hsa04510;hsa04520;hsa04530;hsa04540;hsa04611;hsa04727;hsa04750;hsa04810;hsa04912;hsa04915;hsa04917;hsa04919;hsa04921;hsa05100;hsa05120;hsa05131;hsa05152;hsa05161;hsa05203;hsa05205;hsa05219 
                                               SRC 
                                               NA 
                                               slateblue2 
                                               16 
                                               3.48 
                                             
                                             
                                               PIK3R2-mRNA 
                                               0.0724 
                                               0.0251 
                                               0.0232 
                                               0.122 
                                               1.05 
                                               1.02 
                                               1.09 
                                               0.00651 
                                               0.0963 
                                               lm.nb 
                                               CD28 co-stimulation, CD28 dependent PI3KAkt signaling, Cell surface interactions at the vascular wall, Constitutive Signaling by Aberrant PI3K in Cancer, DAP12 signaling, Downstream signal transduction, Downstream TCR signaling, G alpha (1213) signalling events, G alpha (q) signalling events, G beta.CL.gamma signalling through PI3Kgamma, GAB1 signalosome, Gastrin-CREB signalling pathway via PKC and MAPK, GPVI-mediated activation cascade, Interleukin receptor SHC signaling, IRS-related events triggered by IGF1R, Nephrin interactions, PI3K Cascade, PI3KAKT activation, PIP3 activates AKT signaling, Regulation of signaling by CBL, Rho GTPase cycle, Role of LAT2NTALLAB on calcium mobilization, Signaling by ERBB2, Signaling by PDGF, Signaling by SCF-KIT, Synthesis of PIPs at the plasma membrane, Tie2 Signaling, VEGFA-VEGFR2 Pathway 
                                               SEBELIUS_2_C4636 
                                               PIK3R2 
                                               mRNA 
                                               0 
                                               
                                               
                                               G alpha (q) signalling events;Downstream TCR signaling;PIP3 activates AKT signaling;VEGFA-VEGFR2 Pathway;Synthesis of PIPs at the plasma membrane;Rho GTPase cycle;Interleukin receptor SHC signaling;G beta.CL.gamma signalling through PI3Kgamma;Regulation of signaling by CBL;Role of phospholipids in phagocytosis;G alpha (1213) signalling events;PI3K Cascade;Constitutive Signaling by Aberrant PI3K in Cancer;CD28 dependent PI3KAkt signaling;Tie2 Signaling;GPVI-mediated activation cascade;Nephrin interactions;Gastrin-CREB signalling pathway via PKC and MAPK;Downstream signal transduction;GAB1 signalosome;Signaling by SCF-KIT;PI3KAKT activation;Role of LAT2NTALLAB on calcium mobilization;DAP12 signaling;Signaling by PDGF;Signaling by ERBB2;IRS-related events triggered by IGF1R;CD28 co-stimulation;Cell surface interactions at the vascular wall 
                                               hsa04012;hsa04014;hsa04015;hsa04022;hsa04024;hsa04062;hsa04066;hsa04068;hsa04070;hsa04071;hsa04072;hsa04150;hsa04151;hsa04152;hsa04210;hsa04211;hsa04213;hsa04261;hsa04370;hsa04380;hsa04510;hsa04550;hsa04611;hsa04620;hsa04630;hsa04650;hsa04660;hsa04662;hsa04664;hsa04666;hsa04668;hsa04670;hsa04722;hsa04725;hsa04750;hsa04810;hsa04910;hsa04914;hsa04915;hsa04917;hsa04919;hsa04921;hsa04923;hsa04930;hsa04931;hsa04932;hsa04933;hsa04960;hsa04973;hsa05100;hsa05142;hsa05145;hsa05146;hsa05160;hsa05161;hsa05162;hsa05164;hsa05166;hsa05169;hsa05200;hsa05203;hsa05205;hsa05206;hsa05210;hsa05211;hsa05212;hsa05213;hsa05214;hsa05215;hsa05218;hsa05220;hsa05221;hsa05222;hsa05223;hsa05230;hsa05231 
                                               PIK3R2 
                                               NA 
                                               slateblue3 
                                               16 
                                               2.88446215139442 
                                             
                                             
                                               PIK3R3-mRNA 
                                               0.109 
                                               0.0563 
                                               -0.00124 
                                               0.22 
                                               1.08 
                                               0.999 
                                               1.16 
                                               0.0603 
                                               0.62 
                                               lm.nb 
                                               CD28 co-stimulation, CD28 dependent PI3KAkt signaling, Constitutive Signaling by Aberrant PI3K in Cancer, DAP12 signaling, Downstream signal transduction, G alpha (1213) signalling events, G alpha (q) signalling events, G beta.CL.gamma signalling through PI3Kgamma, GAB1 signalosome, Gastrin-CREB signalling pathway via PKC and MAPK, GPVI-mediated activation cascade, Interleukin receptor SHC signaling, PI3KAKT activation, PIP3 activates AKT signaling, Regulation of signaling by CBL, Role of LAT2NTALLAB on calcium mobilization, Signaling by ERBB2, Signaling by PDGF, Signaling by SCF-KIT, Synthesis of PIPs at the plasma membrane 
                                               SEBELIUS_2_C4636 
                                               PIK3R3 
                                               mRNA 
                                               0 
                                               
                                               
                                               G alpha (q) signalling events;PIP3 activates AKT signaling;Constitutive Signaling by Aberrant PI3K in Cancer;CD28 dependent PI3KAkt signaling;Synthesis of PIPs at the plasma membrane;Interleukin receptor SHC signaling;G beta.CL.gamma signalling through PI3Kgamma;Regulation of signaling by CBL;GPVI-mediated activation cascade;G alpha (1213) signalling events;Gastrin-CREB signalling pathway via PKC and MAPK;Downstream signal transduction;GAB1 signalosome;Signaling by SCF-KIT;PI3KAKT activation;Role of LAT2NTALLAB on calcium mobilization;DAP12 signaling;Signaling by PDGF;Signaling by ERBB2;CD28 co-stimulation 
                                               hsa04012;hsa04014;hsa04015;hsa04022;hsa04024;hsa04062;hsa04066;hsa04068;hsa04070;hsa04071;hsa04072;hsa04150;hsa04151;hsa04152;hsa04210;hsa04211;hsa04213;hsa04261;hsa04370;hsa04380;hsa04510;hsa04550;hsa04611;hsa04620;hsa04630;hsa04650;hsa04660;hsa04662;hsa04664;hsa04666;hsa04668;hsa04670;hsa04722;hsa04725;hsa04750;hsa04810;hsa04910;hsa04914;hsa04915;hsa04917;hsa04919;hsa04921;hsa04923;hsa04930;hsa04931;hsa04932;hsa04933;hsa04960;hsa04973;hsa05100;hsa05142;hsa05145;hsa05146;hsa05160;hsa05161;hsa05162;hsa05164;hsa05166;hsa05169;hsa05200;hsa05203;hsa05205;hsa05210;hsa05211;hsa05212;hsa05213;hsa05214;hsa05215;hsa05218;hsa05220;hsa05221;hsa05222;hsa05223;hsa05230;hsa05231 
                                               PIK3R3 
                                               NA 
                                               azure4 
                                               1 
                                               1.9360568383659 
                                             
                                             
                                               PIK3CA-mRNA 
                                               0.0374 
                                               0.0195 
                                               -0.000784 
                                               0.0756 
                                               1.03 
                                               0.999 
                                               1.05 
                                               0.0626 
                                               0.639 
                                               lm.nb 
                                               CD28 co-stimulation, CD28 dependent PI3KAkt signaling, Cell surface interactions at the vascular wall, Constitutive Signaling by Aberrant PI3K in Cancer, Constitutive Signaling by EGFRvIII, DAP12 signaling, Downstream signal transduction, Downstream TCR signaling, G alpha (1213) signalling events, G alpha (q) signalling events, G beta.CL.gamma signalling through PI3Kgamma, GAB1 signalosome, Gastrin-CREB signalling pathway via PKC and MAPK, GPVI-mediated activation cascade, Interleukin receptor SHC signaling, IRS-related events triggered by IGF1R, Nephrin interactions, PI3K Cascade, PI3KAKT activation, PIP3 activates AKT signaling, Regulation of signaling by CBL, Role of LAT2NTALLAB on calcium mobilization, Signaling by ERBB2, Signaling by FGFR1 in disease, Signaling by FGFR2 in disease, Signaling by FGFR3 in disease, Signaling by FGFR4 in disease, Signaling by PDGF, Signaling by SCF-KIT, Synthesis of PIPs at the plasma membrane, Tie2 Signaling, VEGFA-VEGFR2 Pathway 
                                               SEBELIUS_2_C4636 
                                               PIK3CA 
                                               mRNA 
                                               0 
                                               
                                               
                                               G alpha (q) signalling events;Downstream TCR signaling;Signaling by FGFR2 in disease;PIP3 activates AKT signaling;VEGFA-VEGFR2 Pathway;Synthesis of PIPs at the plasma membrane;Constitutive Signaling by Ligand-Responsive EGFR Cancer Variants;Interleukin receptor SHC signaling;G beta.CL.gamma signalling through PI3Kgamma;Regulation of signaling by CBL;Role of phospholipids in phagocytosis;G alpha (1213) signalling events;PI3K Cascade;Signaling by FGFR3 in disease;Constitutive Signaling by Aberrant PI3K in Cancer;Signaling by FGFR4 in disease;CD28 dependent PI3KAkt signaling;Tie2 Signaling;GPVI-mediated activation cascade;Nephrin interactions;Constitutive Signaling by EGFRvIII;Gastrin-CREB signalling pathway via PKC and MAPK;Downstream signal transduction;GAB1 signalosome;Signaling by SCF-KIT;PI3KAKT activation;Role of LAT2NTALLAB on calcium mobilization;DAP12 signaling;Signaling by PDGF;Signaling by ERBB2;IRS-related events triggered by IGF1R;CD28 co-stimulation;Cell surface interactions at the vascular wall;Signaling by FGFR1 in disease 
                                               hsa00562;hsa04012;hsa04014;hsa04015;hsa04022;hsa04024;hsa04062;hsa04066;hsa04068;hsa04070;hsa04071;hsa04072;hsa04150;hsa04151;hsa04152;hsa04210;hsa04211;hsa04213;hsa04261;hsa04370;hsa04380;hsa04510;hsa04550;hsa04611;hsa04620;hsa04630;hsa04650;hsa04660;hsa04662;hsa04664;hsa04666;hsa04668;hsa04670;hsa04722;hsa04725;hsa04750;hsa04810;hsa04910;hsa04914;hsa04915;hsa04917;hsa04919;hsa04921;hsa04923;hsa04930;hsa04931;hsa04932;hsa04933;hsa04960;hsa04973;hsa05100;hsa05142;hsa05145;hsa05146;hsa05160;hsa05161;hsa05162;hsa05164;hsa05166;hsa05169;hsa05200;hsa05203;hsa05205;hsa05206;hsa05210;hsa05211;hsa05212;hsa05213;hsa05214;hsa05215;hsa05218;hsa05220;hsa05221;hsa05222;hsa05223;hsa05230;hsa05231 
                                               PIK3CA 
                                               NA 
                                               azure4 
                                               1 
                                               1.91794871794872 
                                             
                                             
                                               CDC42-mRNA 
                                               -0.0218 
                                               0.0128 
                                               -0.0469 
                                               0.00331 
                                               0.985 
                                               0.968 
                                               1 
                                               0.0972 
                                               0.92 
                                               lm.nb 
                                               CD28 co-stimulation, CD28 dependent Vav1 pathway, DCC mediated attractive signaling, EPHB-mediated forward signaling, G alpha (1213) signalling events, GPVI-mediated activation cascade, Regulation of actin dynamics for phagocytic cup formation, Rho GTPase cycle, RHO GTPases Activate Formins, RHO GTPases activate IQGAPs, RHO GTPases activate PAKs, Sema4D induced cell migration and growth-cone collapse, VEGFA-VEGFR2 Pathway 
                                               SEBELIUS_2_C4636 
                                               CDC42 
                                               mRNA 
                                               0 
                                               
                                               
                                               DCC mediated attractive signaling;Factors involved in megakaryocyte development and platelet production;VEGFA-VEGFR2 Pathway;RHO GTPases activate KTN1;CD28 dependent Vav1 pathway;Inactivation of Cdc42 and Rac;Rho GTPase cycle;EPHB-mediated forward signaling;RHO GTPases Activate Formins;RHO GTPases activate PAKs;GPVI-mediated activation cascade;G alpha (1213) signalling events;RHO GTPases activate IQGAPs;Sema4D induced cell migration and growth-cone collapse;Regulation of actin dynamics for phagocytic cup formation;CD28 co-stimulation 
                                               hsa04010;hsa04014;hsa04015;hsa04062;hsa04144;hsa04360;hsa04370;hsa04510;hsa04520;hsa04530;hsa04660;hsa04666;hsa04670;hsa04722;hsa04810;hsa04912;hsa04932;hsa04933;hsa05100;hsa05120;hsa05130;hsa05131;hsa05132;hsa05200;hsa05203;hsa05205;hsa05211;hsa05212 
                                               CDC42 
                                               NA 
                                               azure4 
                                               1 
                                               -1.703125 
                                             
                                             
                                               YES1-mRNA 
                                               0.0174 
                                               0.0117 
                                               -0.00543 
                                               0.0403 
                                               1.01 
                                               0.996 
                                               1.03 
                                               0.143 
                                               1 
                                               lm.nb 
                                               CD28 co-stimulation, Cell surface interactions at the vascular wall, CTLA4 inhibitory signaling, EPH-ephrin mediated repulsion of cells, EPHA-mediated growth cone collapse, EPHB-mediated forward signaling, Regulation of signaling by CBL, Signaling by ERBB2, Signaling by SCF-KIT 
                                               SEBELIUS_2_C4636 
                                               YES1 
                                               mRNA 
                                               0 
                                               
                                               
                                               Signaling by ERBB2;EPHB-mediated forward signaling;EPHA-mediated growth cone collapse;Regulation of KIT signaling;EPH-ephrin mediated repulsion of cells;Regulation of signaling by CBL;CD28 co-stimulation;CTLA4 inhibitory signaling;Signaling by SCF-KIT;Cell surface interactions at the vascular wall 
                                               hsa04520;hsa04530 
                                               YES1 
                                               NA 
                                               azure4 
                                               1 
                                               1.48717948717949 
                                             
                                             
                                               AKT1-mRNA 
                                               0.0219 
                                               0.031 
                                               -0.0388 
                                               0.0827 
                                               1.02 
                                               0.973 
                                               1.06 
                                               0.484 
                                               1 
                                               lm.nb 
                                               Activation of BAD and translocation to mitochondria , CD28 co-stimulation, DAP12 signaling, Deactivation of the beta-catenin transactivating complex, Downstream signal transduction, G beta.CL.gamma signalling through PI3Kgamma, GAB1 signalosome, GPVI-mediated activation cascade, IRS-related events triggered by IGF1R, PI3K Cascade, PI3KAKT activation, PIP3 activates AKT signaling, Role of LAT2NTALLAB on calcium mobilization, Signaling by ERBB2, Signaling by PDGF, Signaling by SCF-KIT, TP53 Regulates Metabolic Genes, VEGFA-VEGFR2 Pathway, VEGFR2 mediated vascular permeability 
                                               SEBELIUS_2_C4636 
                                               AKT1 
                                               mRNA 
                                               0 
                                               
                                               
                                               TP53 Regulates Metabolic Genes;Deactivation of the beta-catenin transactivating complex;Activation of BAD and translocation to mitochondria ;eNOS activation;mTOR signalling;G beta.CL.gamma signalling through PI3Kgamma;KSRP (KHSRP) binds and destabilizes mRNA;AKT phosphorylates targets in the nucleus;VEGFR2 mediated vascular permeability;GPVI-mediated activation cascade;Tetrahydrobiopterin (BH4) synthesis# recycling# salvage and regulation;Butyrate Response Factor 1 (BRF1) binds and destabilizes mRNA;Signaling by ERBB2;PIP3 activates AKT signaling;Downstream signal transduction;GAB1 signalosome;Signaling by SCF-KIT;PI3KAKT activation;Role of LAT2NTALLAB on calcium mobilization;DAP12 signaling;Signaling by PDGF;CD28 co-stimulation;PI3K Cascade;IRS-related events triggered by IGF1R;VEGFA-VEGFR2 Pathway 
                                               hsa04010;hsa04012;hsa04014;hsa04015;hsa04022;hsa04024;hsa04062;hsa04066;hsa04068;hsa04071;hsa04072;hsa04150;hsa04151;hsa04152;hsa04210;hsa04211;hsa04213;hsa04261;hsa04370;hsa04380;hsa04510;hsa04530;hsa04550;hsa04611;hsa04620;hsa04630;hsa04660;hsa04662;hsa04664;hsa04666;hsa04668;hsa04722;hsa04725;hsa04728;hsa04910;hsa04914;hsa04915;hsa04917;hsa04919;hsa04920;hsa04922;hsa04923;hsa04931;hsa04932;hsa04933;hsa04973;hsa05142;hsa05145;hsa05152;hsa05160;hsa05161;hsa05162;hsa05164;hsa05166;hsa05169;hsa05200;hsa05205;hsa05210;hsa05211;hsa05212;hsa05213;hsa05214;hsa05215;hsa05218;hsa05220;hsa05221;hsa05222;hsa05223;hsa05230;hsa05231 
                                               AKT1 
                                               NA 
                                               azure4 
                                               1 
                                               0.706451612903226 
                                             
                                             
                                               AKT2-mRNA 
                                               0.00957 
                                               0.0157 
                                               -0.0212 
                                               0.0403 
                                               1.01 
                                               0.985 
                                               1.03 
                                               0.546 
                                               1 
                                               lm.nb 
                                               Activation of BAD and translocation to mitochondria , CD28 co-stimulation, DAP12 signaling, Deactivation of the beta-catenin transactivating complex, Downstream signal transduction, G beta.CL.gamma signalling through PI3Kgamma, GAB1 signalosome, GPVI-mediated activation cascade, IRS-related events triggered by IGF1R, PI3K Cascade, PI3KAKT activation, PIP3 activates AKT signaling, Role of LAT2NTALLAB on calcium mobilization, Signaling by ERBB2, Signaling by PDGF, Signaling by SCF-KIT, TP53 Regulates Metabolic Genes, VEGFA-VEGFR2 Pathway, VEGFR2 mediated vascular permeability 
                                               SEBELIUS_2_C4636 
                                               AKT2 
                                               mRNA 
                                               0 
                                               
                                               
                                               TP53 Regulates Metabolic Genes;PDE3B signalling;AKT phosphorylates targets in the cytosol;Negative regulation of the PI3KAKT network;Deactivation of the beta-catenin transactivating complex;Activation of BAD and translocation to mitochondria ;G beta.CL.gamma signalling through PI3Kgamma;AKT phosphorylates targets in the nucleus;VEGFR2 mediated vascular permeability;GPVI-mediated activation cascade;Signaling by ERBB2;PI3K Cascade;IRS-related events triggered by IGF1R;mTOR signalling;PIP3 activates AKT signaling;Downstream signal transduction;GAB1 signalosome;Signaling by SCF-KIT;PI3KAKT activation;Role of LAT2NTALLAB on calcium mobilization;DAP12 signaling;Signaling by PDGF;CD28 co-stimulation;VEGFA-VEGFR2 Pathway 
                                               hsa04010;hsa04012;hsa04014;hsa04015;hsa04022;hsa04024;hsa04062;hsa04066;hsa04068;hsa04071;hsa04072;hsa04150;hsa04151;hsa04152;hsa04210;hsa04211;hsa04213;hsa04261;hsa04370;hsa04380;hsa04510;hsa04530;hsa04550;hsa04611;hsa04620;hsa04630;hsa04660;hsa04662;hsa04664;hsa04666;hsa04668;hsa04722;hsa04725;hsa04728;hsa04910;hsa04914;hsa04915;hsa04917;hsa04919;hsa04920;hsa04922;hsa04923;hsa04931;hsa04932;hsa04933;hsa04973;hsa05142;hsa05145;hsa05152;hsa05160;hsa05161;hsa05162;hsa05164;hsa05166;hsa05169;hsa05200;hsa05205;hsa05210;hsa05211;hsa05212;hsa05213;hsa05214;hsa05215;hsa05218;hsa05220;hsa05221;hsa05222;hsa05223;hsa05230;hsa05231 
                                               AKT2 
                                               NA 
                                               azure4 
                                               1 
                                               0.609554140127389 
                                             
                                             
                                               FYN-mRNA 
                                               0.0103 
                                               0.0454 
                                               -0.0786 
                                               0.0993 
                                               1.01 
                                               0.947 
                                               1.07 
                                               0.821 
                                               1 
                                               lm.nb 
                                               ARMS-mediated activation, CD209 (DC-SIGN) signaling, CD28 co-stimulation, CD28 dependent PI3KAkt signaling, CD28 dependent Vav1 pathway, Cell surface interactions at the vascular wall, Constitutive Signaling by Aberrant PI3K in Cancer, CTLA4 inhibitory signaling, DAP12 signaling, Downstream signal transduction, EPH-ephrin mediated repulsion of cells, FCERI mediated MAPK activation, Frs2-mediated activation, GAB1 signalosome, Gastrin-CREB signalling pathway via PKC and MAPK, GPVI-mediated activation cascade, Interleukin receptor SHC signaling, IRS-related events triggered by IGF1R, Nephrin interactions, PI3KAKT activation, RAFMAP kinase cascade, Regulation of signaling by CBL, Role of LAT2NTALLAB on calcium mobilization, Sema3A PAK dependent Axon repulsion, Signaling by ERBB2, Signaling by PDGF, Signaling by SCF-KIT, VEGFA-VEGFR2 Pathway, VEGFR2 mediated cell proliferation 
                                               SEBELIUS_2_C4636 
                                               FYN 
                                               mRNA 
                                               0 
                                               
                                               
                                               EPH-ephrin mediated repulsion of cells;Regulation of signaling by CBL;FCGR activation;Antigen activates B Cell Receptor (BCR) leading to generation of second messengers;Dectin-2 family;RAFMAP kinase cascade;Constitutive Signaling by Aberrant PI3K in Cancer;CTLA4 inhibitory signaling;CD28 dependent PI3KAkt signaling;PECAM1 interactions;CD28 dependent Vav1 pathway;GPVI-mediated activation cascade;Sema3A PAK dependent Axon repulsion;Nephrin interactions;SEMA3A-Plexin repulsion signaling by inhibiting Integrin adhesion;CRMPs in Sema3A signaling;CD209 (DC-SIGN) signaling;Downstream signal transduction;GAB1 signalosome;Signaling by SCF-KIT;PI3KAKT activation;Role of LAT2NTALLAB on calcium mobilization;DAP12 signaling;Signaling by PDGF;Signaling by ERBB2;VEGFA-VEGFR2 Pathway;VEGFR2 mediated cell proliferation;Frs2-mediated activation;ARMS-mediated activation;Gastrin-CREB signalling pathway via PKC and MAPK;Interleukin receptor SHC signaling;FCERI mediated MAPK activation;IRS-related events triggered by IGF1R;CD28 co-stimulation;Cell surface interactions at the vascular wall 
                                               hsa04071;hsa04072;hsa04360;hsa04380;hsa04510;hsa04520;hsa04611;hsa04650;hsa04660;hsa04664;hsa04725;hsa05020;hsa05130;hsa05162;hsa05416 
                                               FYN 
                                               NA 
                                               azure4 
                                               1 
                                               0.226872246696035 
                                             
                                             
                                               RAC1-mRNA 
                                               -0.000703 
                                               0.0123 
                                               -0.0248 
                                               0.0234 
                                               1 
                                               0.983 
                                               1.02 
                                               0.955 
                                               1 
                                               lm.nb 
                                               Activation of Rac, CD28 co-stimulation, CD28 dependent Vav1 pathway, DAP12 signaling, EPH-ephrin mediated repulsion of cells, Ephrin signaling, G alpha (1213) signalling events, GPVI-mediated activation cascade, MAPK6MAPK4 signaling, NRAGE signals death through JNK, PCPCE pathway, Regulation of actin dynamics for phagocytic cup formation, RHO GTPases activate CIT, RHO GTPases Activate Formins, RHO GTPases activate IQGAPs, RHO GTPases activate PAKs, RHO GTPases activate PKNs, RHO GTPases Activate WASPs and WAVEs, Sema3A PAK dependent Axon repulsion, Sema4D induced cell migration and growth-cone collapse, Signal transduction by L1, Signaling by SCF-KIT, VEGFA-VEGFR2 Pathway, VEGFR2 mediated vascular permeability 
                                               SEBELIUS_2_C4636 
                                               RAC1 
                                               mRNA 
                                               0 
                                               
                                               
                                               MAPK6MAPK4 signaling;Signaling by SCF-KIT;Inactivation of Cdc42 and Rac;DAP12 signaling;Ephrin signaling;EPH-ephrin mediated repulsion of cells;Activation of Rac;RHO GTPases Activate Formins;VEGFR2 mediated vascular permeability;RHO GTPases activate PAKs;Sema4D mediated inhibition of cell attachment and migration;G alpha (1213) signalling events;Regulation of actin dynamics for phagocytic cup formation;Sema4D induced cell migration and growth-cone collapse;Signal transduction by L1;RHO GTPases activate CIT;RHO GTPases Activate WASPs and WAVEs;Factors involved in megakaryocyte development and platelet production;RHO GTPases activate KTN1;CD28 dependent Vav1 pathway;DSCAM interactions;NRAGE signals death through JNK;GPVI-mediated activation cascade;Sema3A PAK dependent Axon repulsion;RHO GTPases activate PKNs;SEMA3A-Plexin repulsion signaling by inhibiting Integrin adhesion;RHO GTPases activate IQGAPs;PCPCE pathway;VEGFA-VEGFR2 Pathway;CD28 co-stimulation 
                                               hsa04010;hsa04014;hsa04015;hsa04024;hsa04062;hsa04071;hsa04145;hsa04151;hsa04310;hsa04360;hsa04370;hsa04380;hsa04510;hsa04520;hsa04620;hsa04650;hsa04662;hsa04664;hsa04666;hsa04670;hsa04722;hsa04810;hsa04932;hsa04933;hsa04972;hsa05100;hsa05120;hsa05131;hsa05132;hsa05200;hsa05203;hsa05205;hsa05210;hsa05211;hsa05212;hsa05231;hsa05416 
                                               RAC1 
                                               NA 
                                               azure4 
                                               1 
                                               -0.0571544715447154 
                                             
                                           
                                           DE Results - CD28 co-stimulation genes - Day  Table displaying each sample's global significance scores and directed global significance scores as defined in the heatmaps above. The global significance score is calculated as the square root of the mean squared t-statistic for the genes in a gene set, with t-statistics coming from the linear regression underlying our differential expression analysis. The directed global significance score is calculated as the square root of the mean signed squared t-statistic for the genes in a gene set, with t-statistics coming from the linear regression underlying our differential expression analysis.   
                                      
                                       
                                  
                                
                              
                            
                            
                              
                                
                                  
                                       
                                      
                                           Volcano Plot: Line: Caco2 vs.Fibro   More Plot Information                Volcano Plot: Line: Caco2 vs.Fibro  Volcano plot displaying each gene's -log10(p-value) and log2 fold change for the selected covariate. Highly statistically significant genes fall at the top of the plot, and highly differentially expressed genes fall to either side. Genes within the selected gene set are highlighted in orange. Horizontal lines indicate various False Discovery Rate (FDR) thresholds.   
                                           DE Results - CD28 co-stimulation genes - Line: Caco2 vs.Fibro   More Plot Information      Download CSV Data     
                                           
                                             
                                             
                                           
                                           
                                             
                                               Probe Label 
                                               Log2 fold change 
                                               std error (log2) 
                                               Lower confidence limit (log2) 
                                               Upper confidence limit (log2) 
                                               Linear fold change 
                                               Lower confidence limit (linear) 
                                               Upper confidence limit (linear) 
                                               P-value 
                                               BY.p.value 
                                               method 
                                               Gene.sets 
                                               Codeset.Name 
                                               Probe.Label 
                                               Analyte.Type 
                                               Is.Control 
                                               Control.Type 
                                               Related.Probes 
                                               Probe.Annotation 
                                               KEGG.Pathways 
                                               Official.Gene.Name 
                                               Control.Conc 
                                               volcanocol 
                                               volcanopch 
                                               tstats.all 
                                             
                                           
                                           
                                             
                                               AKT3-mRNA 
                                               -2.88 
                                               0.162 
                                               -3.2 
                                               -2.57 
                                               0.135 
                                               0.109 
                                               0.169 
                                               9.07e-20 
                                               6.13e-18 
                                               lm.nb 
                                               Activation of BAD and translocation to mitochondria , CD28 co-stimulation, CD28 dependent PI3KAkt signaling, DAP12 signaling, Downstream signal transduction, G beta.CL.gamma signalling through PI3Kgamma, GAB1 signalosome, GPVI-mediated activation cascade, PI3KAKT activation, PIP3 activates AKT signaling, Role of LAT2NTALLAB on calcium mobilization, Signaling by ERBB2, Signaling by PDGF, Signaling by SCF-KIT, TP53 Regulates Metabolic Genes, VEGFA-VEGFR2 Pathway, VEGFR2 mediated vascular permeability 
                                               SEBELIUS_2_C4636 
                                               AKT3 
                                               mRNA 
                                               0 
                                               
                                               
                                               TP53 Regulates Metabolic Genes;CD28 dependent PI3KAkt signaling;AKT phosphorylates targets in the cytosol;Negative regulation of the PI3KAKT network;Activation of BAD and translocation to mitochondria ;G beta.CL.gamma signalling through PI3Kgamma;AKT phosphorylates targets in the nucleus;VEGFR2 mediated vascular permeability;GPVI-mediated activation cascade;Signaling by ERBB2;PIP3 activates AKT signaling;Downstream signal transduction;GAB1 signalosome;Signaling by SCF-KIT;PI3KAKT activation;Role of LAT2NTALLAB on calcium mobilization;DAP12 signaling;Signaling by PDGF;CD28 co-stimulation;VEGFA-VEGFR2 Pathway 
                                               hsa04010;hsa04012;hsa04014;hsa04015;hsa04022;hsa04024;hsa04062;hsa04066;hsa04068;hsa04071;hsa04072;hsa04150;hsa04151;hsa04152;hsa04210;hsa04211;hsa04213;hsa04261;hsa04370;hsa04380;hsa04510;hsa04530;hsa04550;hsa04611;hsa04620;hsa04630;hsa04660;hsa04662;hsa04664;hsa04666;hsa04668;hsa04722;hsa04725;hsa04728;hsa04910;hsa04914;hsa04915;hsa04917;hsa04919;hsa04920;hsa04922;hsa04923;hsa04931;hsa04932;hsa04933;hsa04973;hsa05142;hsa05145;hsa05152;hsa05160;hsa05161;hsa05162;hsa05164;hsa05166;hsa05169;hsa05200;hsa05205;hsa05210;hsa05211;hsa05212;hsa05213;hsa05214;hsa05215;hsa05218;hsa05220;hsa05221;hsa05222;hsa05223;hsa05230;hsa05231 
                                               AKT3 
                                               NA 
                                               slateblue1 
                                               16 
                                               -17.7777777777778 
                                             
                                             
                                               YES1-mRNA 
                                               1.2 
                                               0.0896 
                                               1.02 
                                               1.38 
                                               2.3 
                                               2.03 
                                               2.59 
                                               9.42e-16 
                                               2.57e-14 
                                               lm.nb 
                                               CD28 co-stimulation, Cell surface interactions at the vascular wall, CTLA4 inhibitory signaling, EPH-ephrin mediated repulsion of cells, EPHA-mediated growth cone collapse, EPHB-mediated forward signaling, Regulation of signaling by CBL, Signaling by ERBB2, Signaling by SCF-KIT 
                                               SEBELIUS_2_C4636 
                                               YES1 
                                               mRNA 
                                               0 
                                               
                                               
                                               Signaling by ERBB2;EPHB-mediated forward signaling;EPHA-mediated growth cone collapse;Regulation of KIT signaling;EPH-ephrin mediated repulsion of cells;Regulation of signaling by CBL;CD28 co-stimulation;CTLA4 inhibitory signaling;Signaling by SCF-KIT;Cell surface interactions at the vascular wall 
                                               hsa04520;hsa04530 
                                               YES1 
                                               NA 
                                               slateblue1 
                                               16 
                                               13.3928571428571 
                                             
                                             
                                               PIK3R1-mRNA 
                                               1.99 
                                               0.164 
                                               1.67 
                                               2.31 
                                               3.97 
                                               3.18 
                                               4.96 
                                               1.85e-14 
                                               4.23e-13 
                                               lm.nb 
                                               CD28 co-stimulation, CD28 dependent PI3KAkt signaling, Cell surface interactions at the vascular wall, Constitutive Signaling by Aberrant PI3K in Cancer, Constitutive Signaling by EGFRvIII, DAP12 signaling, Downstream signal transduction, Downstream TCR signaling, G alpha (q) signalling events, G beta.CL.gamma signalling through PI3Kgamma, GAB1 signalosome, Gastrin-CREB signalling pathway via PKC and MAPK, GPVI-mediated activation cascade, Interleukin receptor SHC signaling, IRS-related events triggered by IGF1R, Nephrin interactions, PI3KAKT activation, PIP3 activates AKT signaling, Regulation of signaling by CBL, Role of LAT2NTALLAB on calcium mobilization, Signaling by ERBB2, Signaling by FGFR1 in disease, Signaling by FGFR2 in disease, Signaling by FGFR3 in disease, Signaling by FGFR4 in disease, Signaling by PDGF, Signaling by SCF-KIT, Synthesis of PIPs at the plasma membrane, Tie2 Signaling, VEGFA-VEGFR2 Pathway 
                                               SEBELIUS_2_C4636 
                                               PIK3R1 
                                               mRNA 
                                               0 
                                               
                                               
                                               G alpha (q) signalling events;Downstream TCR signaling;Signaling by FGFR2 in disease;PIP3 activates AKT signaling;VEGFA-VEGFR2 Pathway;Synthesis of PIPs at the plasma membrane;Constitutive Signaling by Ligand-Responsive EGFR Cancer Variants;Interleukin receptor SHC signaling;G beta.CL.gamma signalling through PI3Kgamma;Regulation of signaling by CBL;Role of phospholipids in phagocytosis;Signaling by FGFR3 in disease;Constitutive Signaling by Aberrant PI3K in Cancer;Signaling by FGFR4 in disease;CD28 dependent PI3KAkt signaling;Tie2 Signaling;Signaling by FGFR1 fusion mutants;GP1b-IX-V activation signalling;GPVI-mediated activation cascade;Nephrin interactions;Constitutive Signaling by EGFRvIII;Gastrin-CREB signalling pathway via PKC and MAPK;Downstream signal transduction;GAB1 signalosome;Signaling by SCF-KIT;PI3KAKT activation;Role of LAT2NTALLAB on calcium mobilization;DAP12 signaling;Signaling by PDGF;Signaling by ERBB2;IRS-related events triggered by IGF1R;CD28 co-stimulation;Cell surface interactions at the vascular wall;Signaling by FGFR1 in disease 
                                               hsa04012;hsa04014;hsa04015;hsa04022;hsa04024;hsa04062;hsa04066;hsa04068;hsa04070;hsa04071;hsa04072;hsa04150;hsa04151;hsa04152;hsa04210;hsa04211;hsa04213;hsa04261;hsa04370;hsa04380;hsa04510;hsa04550;hsa04611;hsa04620;hsa04630;hsa04650;hsa04660;hsa04662;hsa04664;hsa04666;hsa04668;hsa04670;hsa04722;hsa04725;hsa04750;hsa04810;hsa04910;hsa04914;hsa04915;hsa04917;hsa04919;hsa04921;hsa04923;hsa04930;hsa04931;hsa04932;hsa04933;hsa04960;hsa04973;hsa05100;hsa05142;hsa05145;hsa05146;hsa05160;hsa05161;hsa05162;hsa05164;hsa05166;hsa05169;hsa05200;hsa05203;hsa05205;hsa05210;hsa05211;hsa05212;hsa05213;hsa05214;hsa05215;hsa05218;hsa05220;hsa05221;hsa05222;hsa05223;hsa05230;hsa05231 
                                               PIK3R1 
                                               NA 
                                               slateblue1 
                                               16 
                                               12.1341463414634 
                                             
                                             
                                               PAK1-mRNA 
                                               1.92 
                                               0.159 
                                               1.61 
                                               2.23 
                                               3.79 
                                               3.05 
                                               4.7 
                                               2.23e-14 
                                               5.02e-13 
                                               lm.nb 
                                               Activation of Rac, CD209 (DC-SIGN) signaling, CD28 co-stimulation, CD28 dependent Vav1 pathway, EPHB-mediated forward signaling, Ephrin signaling, FCERI mediated MAPK activation, MAPK6MAPK4 signaling, Regulation of actin dynamics for phagocytic cup formation, RHO GTPases activate PAKs, Sema3A PAK dependent Axon repulsion, VEGFA-VEGFR2 Pathway, VEGFR2 mediated vascular permeability 
                                               SEBELIUS_2_C4636 
                                               PAK1 
                                               mRNA 
                                               0 
                                               
                                               
                                               EPHB-mediated forward signaling;Ephrin signaling;MAPK6MAPK4 signaling;DSCAM interactions;Activation of Rac;VEGFR2 mediated vascular permeability;RHO GTPases activate PAKs;Sema3A PAK dependent Axon repulsion;FCERI mediated MAPK activation;Regulation of actin dynamics for phagocytic cup formation;CD209 (DC-SIGN) signaling;CD28 dependent Vav1 pathway;VEGFA-VEGFR2 Pathway;CD28 co-stimulation 
                                               hsa04010;hsa04012;hsa04014;hsa04024;hsa04062;hsa04360;hsa04510;hsa04650;hsa04660;hsa04666;hsa04810;hsa05120;hsa05205;hsa05211 
                                               PAK1 
                                               NA 
                                               slateblue1 
                                               16 
                                               12.0754716981132 
                                             
                                             
                                               SRC-mRNA 
                                               1.77 
                                               0.154 
                                               1.47 
                                               2.07 
                                               3.41 
                                               2.76 
                                               4.2 
                                               8.97e-14 
                                               1.9e-12 
                                               lm.nb 
                                               ARMS-mediated activation, CD28 co-stimulation, Cell surface interactions at the vascular wall, CTLA4 inhibitory signaling, DAP12 signaling, Downstream signal transduction, FCERI mediated MAPK activation, Frs2-mediated activation, GAB1 signalosome, Gastrin-CREB signalling pathway via PKC and MAPK, Integrin alphaIIb beta3 signaling, Interleukin receptor SHC signaling, IRS-related events triggered by IGF1R, MAP2K and MAPK activation, Negative regulation of FGFR1 signaling, Negative regulation of FGFR2 signaling, Negative regulation of FGFR3 signaling, Negative regulation of FGFR4 signaling, p130Cas linkage to MAPK signaling for integrins, RAFMAP kinase cascade, Signaling by ERBB2, Signaling by PDGF, Signaling by SCF-KIT, Spry regulation of FGF signaling, VEGFA-VEGFR2 Pathway, VEGFR2 mediated cell proliferation 
                                               SEBELIUS_2_C4636 
                                               SRC 
                                               mRNA 
                                               0 
                                               
                                               
                                               FCGR activation;Spry regulation of FGF signaling;GAB1 signalosome;MAP2K and MAPK activation;CLEC7A (Dectin-1) signaling;CD28 co-stimulation;p130Cas linkage to MAPK signaling for integrins;CTLA4 inhibitory signaling;GRB2.CL.SOS provides linkage to MAPK signaling for Integrins ;Signal regulatory protein (SIRP) family interactions;PECAM1 interactions;Thrombin signalling through proteinase activated receptors (PARs);GP1b-IX-V activation signalling;c-src mediated regulation of Cx43 function and closure of gap junctions;Signaling by ERBB2;VEGFA-VEGFR2 Pathway;Signaling by SCF-KIT;VEGFR2 mediated cell proliferation;RAFMAP kinase cascade;Downstream signal transduction;Frs2-mediated activation;ARMS-mediated activation;Gastrin-CREB signalling pathway via PKC and MAPK;Interleukin receptor SHC signaling;FCERI mediated MAPK activation;DAP12 signaling;Signaling by PDGF;IRS-related events triggered by IGF1R;Negative regulation of FGFR3 signaling;Negative regulation of FGFR4 signaling;Negative regulation of FGFR2 signaling;Negative regulation of FGFR1 signaling;Integrin alphaIIb beta3 signaling;Cell surface interactions at the vascular wall 
                                               hsa04012;hsa04015;hsa04062;hsa04144;hsa04370;hsa04510;hsa04520;hsa04530;hsa04540;hsa04611;hsa04727;hsa04750;hsa04810;hsa04912;hsa04915;hsa04917;hsa04919;hsa04921;hsa05100;hsa05120;hsa05131;hsa05152;hsa05161;hsa05203;hsa05205;hsa05219 
                                               SRC 
                                               NA 
                                               slateblue1 
                                               16 
                                               11.4935064935065 
                                             
                                             
                                               PIK3CA-mRNA 
                                               -1.22 
                                               0.145 
                                               -1.51 
                                               -0.94 
                                               0.428 
                                               0.352 
                                               0.521 
                                               3.77e-10 
                                               6.08e-09 
                                               lm.nb 
                                               CD28 co-stimulation, CD28 dependent PI3KAkt signaling, Cell surface interactions at the vascular wall, Constitutive Signaling by Aberrant PI3K in Cancer, Constitutive Signaling by EGFRvIII, DAP12 signaling, Downstream signal transduction, Downstream TCR signaling, G alpha (1213) signalling events, G alpha (q) signalling events, G beta.CL.gamma signalling through PI3Kgamma, GAB1 signalosome, Gastrin-CREB signalling pathway via PKC and MAPK, GPVI-mediated activation cascade, Interleukin receptor SHC signaling, IRS-related events triggered by IGF1R, Nephrin interactions, PI3K Cascade, PI3KAKT activation, PIP3 activates AKT signaling, Regulation of signaling by CBL, Role of LAT2NTALLAB on calcium mobilization, Signaling by ERBB2, Signaling by FGFR1 in disease, Signaling by FGFR2 in disease, Signaling by FGFR3 in disease, Signaling by FGFR4 in disease, Signaling by PDGF, Signaling by SCF-KIT, Synthesis of PIPs at the plasma membrane, Tie2 Signaling, VEGFA-VEGFR2 Pathway 
                                               SEBELIUS_2_C4636 
                                               PIK3CA 
                                               mRNA 
                                               0 
                                               
                                               
                                               G alpha (q) signalling events;Downstream TCR signaling;Signaling by FGFR2 in disease;PIP3 activates AKT signaling;VEGFA-VEGFR2 Pathway;Synthesis of PIPs at the plasma membrane;Constitutive Signaling by Ligand-Responsive EGFR Cancer Variants;Interleukin receptor SHC signaling;G beta.CL.gamma signalling through PI3Kgamma;Regulation of signaling by CBL;Role of phospholipids in phagocytosis;G alpha (1213) signalling events;PI3K Cascade;Signaling by FGFR3 in disease;Constitutive Signaling by Aberrant PI3K in Cancer;Signaling by FGFR4 in disease;CD28 dependent PI3KAkt signaling;Tie2 Signaling;GPVI-mediated activation cascade;Nephrin interactions;Constitutive Signaling by EGFRvIII;Gastrin-CREB signalling pathway via PKC and MAPK;Downstream signal transduction;GAB1 signalosome;Signaling by SCF-KIT;PI3KAKT activation;Role of LAT2NTALLAB on calcium mobilization;DAP12 signaling;Signaling by PDGF;Signaling by ERBB2;IRS-related events triggered by IGF1R;CD28 co-stimulation;Cell surface interactions at the vascular wall;Signaling by FGFR1 in disease 
                                               hsa00562;hsa04012;hsa04014;hsa04015;hsa04022;hsa04024;hsa04062;hsa04066;hsa04068;hsa04070;hsa04071;hsa04072;hsa04150;hsa04151;hsa04152;hsa04210;hsa04211;hsa04213;hsa04261;hsa04370;hsa04380;hsa04510;hsa04550;hsa04611;hsa04620;hsa04630;hsa04650;hsa04660;hsa04662;hsa04664;hsa04666;hsa04668;hsa04670;hsa04722;hsa04725;hsa04750;hsa04810;hsa04910;hsa04914;hsa04915;hsa04917;hsa04919;hsa04921;hsa04923;hsa04930;hsa04931;hsa04932;hsa04933;hsa04960;hsa04973;hsa05100;hsa05142;hsa05145;hsa05146;hsa05160;hsa05161;hsa05162;hsa05164;hsa05166;hsa05169;hsa05200;hsa05203;hsa05205;hsa05206;hsa05210;hsa05211;hsa05212;hsa05213;hsa05214;hsa05215;hsa05218;hsa05220;hsa05221;hsa05222;hsa05223;hsa05230;hsa05231 
                                               PIK3CA 
                                               NA 
                                               slateblue1 
                                               16 
                                               -8.41379310344828 
                                             
                                             
                                               FYN-mRNA 
                                               -2.64 
                                               0.335 
                                               -3.3 
                                               -1.99 
                                               0.16 
                                               0.102 
                                               0.253 
                                               1.96e-09 
                                               2.84e-08 
                                               lm.nb 
                                               ARMS-mediated activation, CD209 (DC-SIGN) signaling, CD28 co-stimulation, CD28 dependent PI3KAkt signaling, CD28 dependent Vav1 pathway, Cell surface interactions at the vascular wall, Constitutive Signaling by Aberrant PI3K in Cancer, CTLA4 inhibitory signaling, DAP12 signaling, Downstream signal transduction, EPH-ephrin mediated repulsion of cells, FCERI mediated MAPK activation, Frs2-mediated activation, GAB1 signalosome, Gastrin-CREB signalling pathway via PKC and MAPK, GPVI-mediated activation cascade, Interleukin receptor SHC signaling, IRS-related events triggered by IGF1R, Nephrin interactions, PI3KAKT activation, RAFMAP kinase cascade, Regulation of signaling by CBL, Role of LAT2NTALLAB on calcium mobilization, Sema3A PAK dependent Axon repulsion, Signaling by ERBB2, Signaling by PDGF, Signaling by SCF-KIT, VEGFA-VEGFR2 Pathway, VEGFR2 mediated cell proliferation 
                                               SEBELIUS_2_C4636 
                                               FYN 
                                               mRNA 
                                               0 
                                               
                                               
                                               EPH-ephrin mediated repulsion of cells;Regulation of signaling by CBL;FCGR activation;Antigen activates B Cell Receptor (BCR) leading to generation of second messengers;Dectin-2 family;RAFMAP kinase cascade;Constitutive Signaling by Aberrant PI3K in Cancer;CTLA4 inhibitory signaling;CD28 dependent PI3KAkt signaling;PECAM1 interactions;CD28 dependent Vav1 pathway;GPVI-mediated activation cascade;Sema3A PAK dependent Axon repulsion;Nephrin interactions;SEMA3A-Plexin repulsion signaling by inhibiting Integrin adhesion;CRMPs in Sema3A signaling;CD209 (DC-SIGN) signaling;Downstream signal transduction;GAB1 signalosome;Signaling by SCF-KIT;PI3KAKT activation;Role of LAT2NTALLAB on calcium mobilization;DAP12 signaling;Signaling by PDGF;Signaling by ERBB2;VEGFA-VEGFR2 Pathway;VEGFR2 mediated cell proliferation;Frs2-mediated activation;ARMS-mediated activation;Gastrin-CREB signalling pathway via PKC and MAPK;Interleukin receptor SHC signaling;FCERI mediated MAPK activation;IRS-related events triggered by IGF1R;CD28 co-stimulation;Cell surface interactions at the vascular wall 
                                               hsa04071;hsa04072;hsa04360;hsa04380;hsa04510;hsa04520;hsa04611;hsa04650;hsa04660;hsa04664;hsa04725;hsa05020;hsa05130;hsa05162;hsa05416 
                                               FYN 
                                               NA 
                                               slateblue1 
                                               16 
                                               -7.88059701492537 
                                             
                                             
                                               PIK3R3-mRNA 
                                               -2.73 
                                               0.417 
                                               -3.55 
                                               -1.91 
                                               0.151 
                                               0.0856 
                                               0.266 
                                               1.17e-07 
                                               1.44e-06 
                                               lm.nb 
                                               CD28 co-stimulation, CD28 dependent PI3KAkt signaling, Constitutive Signaling by Aberrant PI3K in Cancer, DAP12 signaling, Downstream signal transduction, G alpha (1213) signalling events, G alpha (q) signalling events, G beta.CL.gamma signalling through PI3Kgamma, GAB1 signalosome, Gastrin-CREB signalling pathway via PKC and MAPK, GPVI-mediated activation cascade, Interleukin receptor SHC signaling, PI3KAKT activation, PIP3 activates AKT signaling, Regulation of signaling by CBL, Role of LAT2NTALLAB on calcium mobilization, Signaling by ERBB2, Signaling by PDGF, Signaling by SCF-KIT, Synthesis of PIPs at the plasma membrane 
                                               SEBELIUS_2_C4636 
                                               PIK3R3 
                                               mRNA 
                                               0 
                                               
                                               
                                               G alpha (q) signalling events;PIP3 activates AKT signaling;Constitutive Signaling by Aberrant PI3K in Cancer;CD28 dependent PI3KAkt signaling;Synthesis of PIPs at the plasma membrane;Interleukin receptor SHC signaling;G beta.CL.gamma signalling through PI3Kgamma;Regulation of signaling by CBL;GPVI-mediated activation cascade;G alpha (1213) signalling events;Gastrin-CREB signalling pathway via PKC and MAPK;Downstream signal transduction;GAB1 signalosome;Signaling by SCF-KIT;PI3KAKT activation;Role of LAT2NTALLAB on calcium mobilization;DAP12 signaling;Signaling by PDGF;Signaling by ERBB2;CD28 co-stimulation 
                                               hsa04012;hsa04014;hsa04015;hsa04022;hsa04024;hsa04062;hsa04066;hsa04068;hsa04070;hsa04071;hsa04072;hsa04150;hsa04151;hsa04152;hsa04210;hsa04211;hsa04213;hsa04261;hsa04370;hsa04380;hsa04510;hsa04550;hsa04611;hsa04620;hsa04630;hsa04650;hsa04660;hsa04662;hsa04664;hsa04666;hsa04668;hsa04670;hsa04722;hsa04725;hsa04750;hsa04810;hsa04910;hsa04914;hsa04915;hsa04917;hsa04919;hsa04921;hsa04923;hsa04930;hsa04931;hsa04932;hsa04933;hsa04960;hsa04973;hsa05100;hsa05142;hsa05145;hsa05146;hsa05160;hsa05161;hsa05162;hsa05164;hsa05166;hsa05169;hsa05200;hsa05203;hsa05205;hsa05210;hsa05211;hsa05212;hsa05213;hsa05214;hsa05215;hsa05218;hsa05220;hsa05221;hsa05222;hsa05223;hsa05230;hsa05231 
                                               PIK3R3 
                                               NA 
                                               slateblue1 
                                               16 
                                               -6.54676258992806 
                                             
                                             
                                               AKT2-mRNA 
                                               0.479 
                                               0.12 
                                               0.245 
                                               0.713 
                                               1.39 
                                               1.18 
                                               1.64 
                                               0.000285 
                                               0.00263 
                                               lm.nb 
                                               Activation of BAD and translocation to mitochondria , CD28 co-stimulation, DAP12 signaling, Deactivation of the beta-catenin transactivating complex, Downstream signal transduction, G beta.CL.gamma signalling through PI3Kgamma, GAB1 signalosome, GPVI-mediated activation cascade, IRS-related events triggered by IGF1R, PI3K Cascade, PI3KAKT activation, PIP3 activates AKT signaling, Role of LAT2NTALLAB on calcium mobilization, Signaling by ERBB2, Signaling by PDGF, Signaling by SCF-KIT, TP53 Regulates Metabolic Genes, VEGFA-VEGFR2 Pathway, VEGFR2 mediated vascular permeability 
                                               SEBELIUS_2_C4636 
                                               AKT2 
                                               mRNA 
                                               0 
                                               
                                               
                                               TP53 Regulates Metabolic Genes;PDE3B signalling;AKT phosphorylates targets in the cytosol;Negative regulation of the PI3KAKT network;Deactivation of the beta-catenin transactivating complex;Activation of BAD and translocation to mitochondria ;G beta.CL.gamma signalling through PI3Kgamma;AKT phosphorylates targets in the nucleus;VEGFR2 mediated vascular permeability;GPVI-mediated activation cascade;Signaling by ERBB2;PI3K Cascade;IRS-related events triggered by IGF1R;mTOR signalling;PIP3 activates AKT signaling;Downstream signal transduction;GAB1 signalosome;Signaling by SCF-KIT;PI3KAKT activation;Role of LAT2NTALLAB on calcium mobilization;DAP12 signaling;Signaling by PDGF;CD28 co-stimulation;VEGFA-VEGFR2 Pathway 
                                               hsa04010;hsa04012;hsa04014;hsa04015;hsa04022;hsa04024;hsa04062;hsa04066;hsa04068;hsa04071;hsa04072;hsa04150;hsa04151;hsa04152;hsa04210;hsa04211;hsa04213;hsa04261;hsa04370;hsa04380;hsa04510;hsa04530;hsa04550;hsa04611;hsa04620;hsa04630;hsa04660;hsa04662;hsa04664;hsa04666;hsa04668;hsa04722;hsa04725;hsa04728;hsa04910;hsa04914;hsa04915;hsa04917;hsa04919;hsa04920;hsa04922;hsa04923;hsa04931;hsa04932;hsa04933;hsa04973;hsa05142;hsa05145;hsa05152;hsa05160;hsa05161;hsa05162;hsa05164;hsa05166;hsa05169;hsa05200;hsa05205;hsa05210;hsa05211;hsa05212;hsa05213;hsa05214;hsa05215;hsa05218;hsa05220;hsa05221;hsa05222;hsa05223;hsa05230;hsa05231 
                                               AKT2 
                                               NA 
                                               slateblue1 
                                               16 
                                               3.99166666666667 
                                             
                                             
                                               AKT1-mRNA 
                                               -0.919 
                                               0.234 
                                               -1.38 
                                               -0.461 
                                               0.529 
                                               0.385 
                                               0.726 
                                               0.000354 
                                               0.00322 
                                               lm.nb 
                                               Activation of BAD and translocation to mitochondria , CD28 co-stimulation, DAP12 signaling, Deactivation of the beta-catenin transactivating complex, Downstream signal transduction, G beta.CL.gamma signalling through PI3Kgamma, GAB1 signalosome, GPVI-mediated activation cascade, IRS-related events triggered by IGF1R, PI3K Cascade, PI3KAKT activation, PIP3 activates AKT signaling, Role of LAT2NTALLAB on calcium mobilization, Signaling by ERBB2, Signaling by PDGF, Signaling by SCF-KIT, TP53 Regulates Metabolic Genes, VEGFA-VEGFR2 Pathway, VEGFR2 mediated vascular permeability 
                                               SEBELIUS_2_C4636 
                                               AKT1 
                                               mRNA 
                                               0 
                                               
                                               
                                               TP53 Regulates Metabolic Genes;Deactivation of the beta-catenin transactivating complex;Activation of BAD and translocation to mitochondria ;eNOS activation;mTOR signalling;G beta.CL.gamma signalling through PI3Kgamma;KSRP (KHSRP) binds and destabilizes mRNA;AKT phosphorylates targets in the nucleus;VEGFR2 mediated vascular permeability;GPVI-mediated activation cascade;Tetrahydrobiopterin (BH4) synthesis# recycling# salvage and regulation;Butyrate Response Factor 1 (BRF1) binds and destabilizes mRNA;Signaling by ERBB2;PIP3 activates AKT signaling;Downstream signal transduction;GAB1 signalosome;Signaling by SCF-KIT;PI3KAKT activation;Role of LAT2NTALLAB on calcium mobilization;DAP12 signaling;Signaling by PDGF;CD28 co-stimulation;PI3K Cascade;IRS-related events triggered by IGF1R;VEGFA-VEGFR2 Pathway 
                                               hsa04010;hsa04012;hsa04014;hsa04015;hsa04022;hsa04024;hsa04062;hsa04066;hsa04068;hsa04071;hsa04072;hsa04150;hsa04151;hsa04152;hsa04210;hsa04211;hsa04213;hsa04261;hsa04370;hsa04380;hsa04510;hsa04530;hsa04550;hsa04611;hsa04620;hsa04630;hsa04660;hsa04662;hsa04664;hsa04666;hsa04668;hsa04722;hsa04725;hsa04728;hsa04910;hsa04914;hsa04915;hsa04917;hsa04919;hsa04920;hsa04922;hsa04923;hsa04931;hsa04932;hsa04933;hsa04973;hsa05142;hsa05145;hsa05152;hsa05160;hsa05161;hsa05162;hsa05164;hsa05166;hsa05169;hsa05200;hsa05205;hsa05210;hsa05211;hsa05212;hsa05213;hsa05214;hsa05215;hsa05218;hsa05220;hsa05221;hsa05222;hsa05223;hsa05230;hsa05231 
                                               AKT1 
                                               NA 
                                               slateblue1 
                                               16 
                                               -3.92735042735043 
                                             
                                             
                                               CDC42-mRNA 
                                               -0.361 
                                               0.0968 
                                               -0.551 
                                               -0.171 
                                               0.779 
                                               0.683 
                                               0.888 
                                               0.00064 
                                               0.00571 
                                               lm.nb 
                                               CD28 co-stimulation, CD28 dependent Vav1 pathway, DCC mediated attractive signaling, EPHB-mediated forward signaling, G alpha (1213) signalling events, GPVI-mediated activation cascade, Regulation of actin dynamics for phagocytic cup formation, Rho GTPase cycle, RHO GTPases Activate Formins, RHO GTPases activate IQGAPs, RHO GTPases activate PAKs, Sema4D induced cell migration and growth-cone collapse, VEGFA-VEGFR2 Pathway 
                                               SEBELIUS_2_C4636 
                                               CDC42 
                                               mRNA 
                                               0 
                                               
                                               
                                               DCC mediated attractive signaling;Factors involved in megakaryocyte development and platelet production;VEGFA-VEGFR2 Pathway;RHO GTPases activate KTN1;CD28 dependent Vav1 pathway;Inactivation of Cdc42 and Rac;Rho GTPase cycle;EPHB-mediated forward signaling;RHO GTPases Activate Formins;RHO GTPases activate PAKs;GPVI-mediated activation cascade;G alpha (1213) signalling events;RHO GTPases activate IQGAPs;Sema4D induced cell migration and growth-cone collapse;Regulation of actin dynamics for phagocytic cup formation;CD28 co-stimulation 
                                               hsa04010;hsa04014;hsa04015;hsa04062;hsa04144;hsa04360;hsa04370;hsa04510;hsa04520;hsa04530;hsa04660;hsa04666;hsa04670;hsa04722;hsa04810;hsa04912;hsa04932;hsa04933;hsa05100;hsa05120;hsa05130;hsa05131;hsa05132;hsa05200;hsa05203;hsa05205;hsa05211;hsa05212 
                                               CDC42 
                                               NA 
                                               slateblue1 
                                               16 
                                               -3.72933884297521 
                                             
                                             
                                               RAC1-mRNA 
                                               -0.326 
                                               0.093 
                                               -0.508 
                                               -0.144 
                                               0.798 
                                               0.703 
                                               0.905 
                                               0.00121 
                                               0.0104 
                                               lm.nb 
                                               Activation of Rac, CD28 co-stimulation, CD28 dependent Vav1 pathway, DAP12 signaling, EPH-ephrin mediated repulsion of cells, Ephrin signaling, G alpha (1213) signalling events, GPVI-mediated activation cascade, MAPK6MAPK4 signaling, NRAGE signals death through JNK, PCPCE pathway, Regulation of actin dynamics for phagocytic cup formation, RHO GTPases activate CIT, RHO GTPases Activate Formins, RHO GTPases activate IQGAPs, RHO GTPases activate PAKs, RHO GTPases activate PKNs, RHO GTPases Activate WASPs and WAVEs, Sema3A PAK dependent Axon repulsion, Sema4D induced cell migration and growth-cone collapse, Signal transduction by L1, Signaling by SCF-KIT, VEGFA-VEGFR2 Pathway, VEGFR2 mediated vascular permeability 
                                               SEBELIUS_2_C4636 
                                               RAC1 
                                               mRNA 
                                               0 
                                               
                                               
                                               MAPK6MAPK4 signaling;Signaling by SCF-KIT;Inactivation of Cdc42 and Rac;DAP12 signaling;Ephrin signaling;EPH-ephrin mediated repulsion of cells;Activation of Rac;RHO GTPases Activate Formins;VEGFR2 mediated vascular permeability;RHO GTPases activate PAKs;Sema4D mediated inhibition of cell attachment and migration;G alpha (1213) signalling events;Regulation of actin dynamics for phagocytic cup formation;Sema4D induced cell migration and growth-cone collapse;Signal transduction by L1;RHO GTPases activate CIT;RHO GTPases Activate WASPs and WAVEs;Factors involved in megakaryocyte development and platelet production;RHO GTPases activate KTN1;CD28 dependent Vav1 pathway;DSCAM interactions;NRAGE signals death through JNK;GPVI-mediated activation cascade;Sema3A PAK dependent Axon repulsion;RHO GTPases activate PKNs;SEMA3A-Plexin repulsion signaling by inhibiting Integrin adhesion;RHO GTPases activate IQGAPs;PCPCE pathway;VEGFA-VEGFR2 Pathway;CD28 co-stimulation 
                                               hsa04010;hsa04014;hsa04015;hsa04024;hsa04062;hsa04071;hsa04145;hsa04151;hsa04310;hsa04360;hsa04370;hsa04380;hsa04510;hsa04520;hsa04620;hsa04650;hsa04662;hsa04664;hsa04666;hsa04670;hsa04722;hsa04810;hsa04932;hsa04933;hsa04972;hsa05100;hsa05120;hsa05131;hsa05132;hsa05200;hsa05203;hsa05205;hsa05210;hsa05211;hsa05212;hsa05231;hsa05416 
                                               RAC1 
                                               NA 
                                               slateblue2 
                                               16 
                                               -3.50537634408602 
                                             
                                             
                                               PAK2-mRNA 
                                               -0.138 
                                               0.0752 
                                               -0.285 
                                               0.00968 
                                               0.909 
                                               0.821 
                                               1.01 
                                               0.0751 
                                               0.552 
                                               loglinear 
                                               Activation of Rac, CD209 (DC-SIGN) signaling, CD28 co-stimulation, CD28 dependent Vav1 pathway, Ephrin signaling, RHO GTPases activate PAKs, Sema3A PAK dependent Axon repulsion, VEGFA-VEGFR2 Pathway, VEGFR2 mediated vascular permeability 
                                               SEBELIUS_2_C4636 
                                               PAK2 
                                               mRNA 
                                               0 
                                               
                                               
                                               Nef and signal transduction;CD28 dependent Vav1 pathway;Stimulation of the cell death response by PAK-2p34;Ephrin signaling;Activation of Rac;RHO GTPases activate PAKs;VEGFR2 mediated vascular permeability;Sema3A PAK dependent Axon repulsion;CD209 (DC-SIGN) signaling;VEGFA-VEGFR2 Pathway;CD28 co-stimulation 
                                               hsa04010;hsa04012;hsa04014;hsa04360;hsa04510;hsa04660;hsa04810;hsa05211 
                                               PAK2 
                                               NA 
                                               azure4 
                                               1 
                                               -1.83510638297872 
                                             
                                             
                                               PIK3R2-mRNA 
                                               -0.152 
                                               0.189 
                                               -0.523 
                                               0.218 
                                               0.9 
                                               0.696 
                                               1.16 
                                               0.425 
                                               1 
                                               lm.nb 
                                               CD28 co-stimulation, CD28 dependent PI3KAkt signaling, Cell surface interactions at the vascular wall, Constitutive Signaling by Aberrant PI3K in Cancer, DAP12 signaling, Downstream signal transduction, Downstream TCR signaling, G alpha (1213) signalling events, G alpha (q) signalling events, G beta.CL.gamma signalling through PI3Kgamma, GAB1 signalosome, Gastrin-CREB signalling pathway via PKC and MAPK, GPVI-mediated activation cascade, Interleukin receptor SHC signaling, IRS-related events triggered by IGF1R, Nephrin interactions, PI3K Cascade, PI3KAKT activation, PIP3 activates AKT signaling, Regulation of signaling by CBL, Rho GTPase cycle, Role of LAT2NTALLAB on calcium mobilization, Signaling by ERBB2, Signaling by PDGF, Signaling by SCF-KIT, Synthesis of PIPs at the plasma membrane, Tie2 Signaling, VEGFA-VEGFR2 Pathway 
                                               SEBELIUS_2_C4636 
                                               PIK3R2 
                                               mRNA 
                                               0 
                                               
                                               
                                               G alpha (q) signalling events;Downstream TCR signaling;PIP3 activates AKT signaling;VEGFA-VEGFR2 Pathway;Synthesis of PIPs at the plasma membrane;Rho GTPase cycle;Interleukin receptor SHC signaling;G beta.CL.gamma signalling through PI3Kgamma;Regulation of signaling by CBL;Role of phospholipids in phagocytosis;G alpha (1213) signalling events;PI3K Cascade;Constitutive Signaling by Aberrant PI3K in Cancer;CD28 dependent PI3KAkt signaling;Tie2 Signaling;GPVI-mediated activation cascade;Nephrin interactions;Gastrin-CREB signalling pathway via PKC and MAPK;Downstream signal transduction;GAB1 signalosome;Signaling by SCF-KIT;PI3KAKT activation;Role of LAT2NTALLAB on calcium mobilization;DAP12 signaling;Signaling by PDGF;Signaling by ERBB2;IRS-related events triggered by IGF1R;CD28 co-stimulation;Cell surface interactions at the vascular wall 
                                               hsa04012;hsa04014;hsa04015;hsa04022;hsa04024;hsa04062;hsa04066;hsa04068;hsa04070;hsa04071;hsa04072;hsa04150;hsa04151;hsa04152;hsa04210;hsa04211;hsa04213;hsa04261;hsa04370;hsa04380;hsa04510;hsa04550;hsa04611;hsa04620;hsa04630;hsa04650;hsa04660;hsa04662;hsa04664;hsa04666;hsa04668;hsa04670;hsa04722;hsa04725;hsa04750;hsa04810;hsa04910;hsa04914;hsa04915;hsa04917;hsa04919;hsa04921;hsa04923;hsa04930;hsa04931;hsa04932;hsa04933;hsa04960;hsa04973;hsa05100;hsa05142;hsa05145;hsa05146;hsa05160;hsa05161;hsa05162;hsa05164;hsa05166;hsa05169;hsa05200;hsa05203;hsa05205;hsa05206;hsa05210;hsa05211;hsa05212;hsa05213;hsa05214;hsa05215;hsa05218;hsa05220;hsa05221;hsa05222;hsa05223;hsa05230;hsa05231 
                                               PIK3R2 
                                               NA 
                                               azure4 
                                               1 
                                               -0.804232804232804 
                                             
                                           
                                           DE Results - CD28 co-stimulation genes - Line: Caco2 vs.Fibro  Table displaying each sample's global significance scores and directed global significance scores as defined in the heatmaps above. The global significance score is calculated as the square root of the mean squared t-statistic for the genes in a gene set, with t-statistics coming from the linear regression underlying our differential expression analysis. The directed global significance score is calculated as the square root of the mean signed squared t-statistic for the genes in a gene set, with t-statistics coming from the linear regression underlying our differential expression analysis.   
                                      
                                       
                                  
                                
                                
                                  
                                       
                                      
                                           Volcano Plot: Line: HEPG2 vs.Fibro   More Plot Information                Volcano Plot: Line: HEPG2 vs.Fibro  Volcano plot displaying each gene's -log10(p-value) and log2 fold change for the selected covariate. Highly statistically significant genes fall at the top of the plot, and highly differentially expressed genes fall to either side. Genes within the selected gene set are highlighted in orange. Horizontal lines indicate various False Discovery Rate (FDR) thresholds.   
                                           DE Results - CD28 co-stimulation genes - Line: HEPG2 vs.Fibro   More Plot Information      Download CSV Data     
                                           
                                             
                                             
                                           
                                           
                                             
                                               Probe Label 
                                               Log2 fold change 
                                               std error (log2) 
                                               Lower confidence limit (log2) 
                                               Upper confidence limit (log2) 
                                               Linear fold change 
                                               Lower confidence limit (linear) 
                                               Upper confidence limit (linear) 
                                               P-value 
                                               BY.p.value 
                                               method 
                                               Gene.sets 
                                               Codeset.Name 
                                               Probe.Label 
                                               Analyte.Type 
                                               Is.Control 
                                               Control.Type 
                                               Related.Probes 
                                               Probe.Annotation 
                                               KEGG.Pathways 
                                               Official.Gene.Name 
                                               Control.Conc 
                                               volcanocol 
                                               volcanopch 
                                               tstats.all 
                                             
                                           
                                           
                                             
                                               AKT3-mRNA 
                                               -7.38 
                                               0.252 
                                               -7.87 
                                               -6.89 
                                               0.006 
                                               0.00426 
                                               0.00845 
                                               3.23e-27 
                                               1.15e-24 
                                               lm.nb 
                                               Activation of BAD and translocation to mitochondria , CD28 co-stimulation, CD28 dependent PI3KAkt signaling, DAP12 signaling, Downstream signal transduction, G beta.CL.gamma signalling through PI3Kgamma, GAB1 signalosome, GPVI-mediated activation cascade, PI3KAKT activation, PIP3 activates AKT signaling, Role of LAT2NTALLAB on calcium mobilization, Signaling by ERBB2, Signaling by PDGF, Signaling by SCF-KIT, TP53 Regulates Metabolic Genes, VEGFA-VEGFR2 Pathway, VEGFR2 mediated vascular permeability 
                                               SEBELIUS_2_C4636 
                                               AKT3 
                                               mRNA 
                                               0 
                                               
                                               
                                               TP53 Regulates Metabolic Genes;CD28 dependent PI3KAkt signaling;AKT phosphorylates targets in the cytosol;Negative regulation of the PI3KAKT network;Activation of BAD and translocation to mitochondria ;G beta.CL.gamma signalling through PI3Kgamma;AKT phosphorylates targets in the nucleus;VEGFR2 mediated vascular permeability;GPVI-mediated activation cascade;Signaling by ERBB2;PIP3 activates AKT signaling;Downstream signal transduction;GAB1 signalosome;Signaling by SCF-KIT;PI3KAKT activation;Role of LAT2NTALLAB on calcium mobilization;DAP12 signaling;Signaling by PDGF;CD28 co-stimulation;VEGFA-VEGFR2 Pathway 
                                               hsa04010;hsa04012;hsa04014;hsa04015;hsa04022;hsa04024;hsa04062;hsa04066;hsa04068;hsa04071;hsa04072;hsa04150;hsa04151;hsa04152;hsa04210;hsa04211;hsa04213;hsa04261;hsa04370;hsa04380;hsa04510;hsa04530;hsa04550;hsa04611;hsa04620;hsa04630;hsa04660;hsa04662;hsa04664;hsa04666;hsa04668;hsa04722;hsa04725;hsa04728;hsa04910;hsa04914;hsa04915;hsa04917;hsa04919;hsa04920;hsa04922;hsa04923;hsa04931;hsa04932;hsa04933;hsa04973;hsa05142;hsa05145;hsa05152;hsa05160;hsa05161;hsa05162;hsa05164;hsa05166;hsa05169;hsa05200;hsa05205;hsa05210;hsa05211;hsa05212;hsa05213;hsa05214;hsa05215;hsa05218;hsa05220;hsa05221;hsa05222;hsa05223;hsa05230;hsa05231 
                                               AKT3 
                                               NA 
                                               slateblue1 
                                               16 
                                               -29.2857142857143 
                                             
                                             
                                               SRC-mRNA 
                                               2.38 
                                               0.168 
                                               2.05 
                                               2.71 
                                               5.2 
                                               4.14 
                                               6.53 
                                               1.71e-16 
                                               6.08e-15 
                                               lm.nb 
                                               ARMS-mediated activation, CD28 co-stimulation, Cell surface interactions at the vascular wall, CTLA4 inhibitory signaling, DAP12 signaling, Downstream signal transduction, FCERI mediated MAPK activation, Frs2-mediated activation, GAB1 signalosome, Gastrin-CREB signalling pathway via PKC and MAPK, Integrin alphaIIb beta3 signaling, Interleukin receptor SHC signaling, IRS-related events triggered by IGF1R, MAP2K and MAPK activation, Negative regulation of FGFR1 signaling, Negative regulation of FGFR2 signaling, Negative regulation of FGFR3 signaling, Negative regulation of FGFR4 signaling, p130Cas linkage to MAPK signaling for integrins, RAFMAP kinase cascade, Signaling by ERBB2, Signaling by PDGF, Signaling by SCF-KIT, Spry regulation of FGF signaling, VEGFA-VEGFR2 Pathway, VEGFR2 mediated cell proliferation 
                                               SEBELIUS_2_C4636 
                                               SRC 
                                               mRNA 
                                               0 
                                               
                                               
                                               FCGR activation;Spry regulation of FGF signaling;GAB1 signalosome;MAP2K and MAPK activation;CLEC7A (Dectin-1) signaling;CD28 co-stimulation;p130Cas linkage to MAPK signaling for integrins;CTLA4 inhibitory signaling;GRB2.CL.SOS provides linkage to MAPK signaling for Integrins ;Signal regulatory protein (SIRP) family interactions;PECAM1 interactions;Thrombin signalling through proteinase activated receptors (PARs);GP1b-IX-V activation signalling;c-src mediated regulation of Cx43 function and closure of gap junctions;Signaling by ERBB2;VEGFA-VEGFR2 Pathway;Signaling by SCF-KIT;VEGFR2 mediated cell proliferation;RAFMAP kinase cascade;Downstream signal transduction;Frs2-mediated activation;ARMS-mediated activation;Gastrin-CREB signalling pathway via PKC and MAPK;Interleukin receptor SHC signaling;FCERI mediated MAPK activation;DAP12 signaling;Signaling by PDGF;IRS-related events triggered by IGF1R;Negative regulation of FGFR3 signaling;Negative regulation of FGFR4 signaling;Negative regulation of FGFR2 signaling;Negative regulation of FGFR1 signaling;Integrin alphaIIb beta3 signaling;Cell surface interactions at the vascular wall 
                                               hsa04012;hsa04015;hsa04062;hsa04144;hsa04370;hsa04510;hsa04520;hsa04530;hsa04540;hsa04611;hsa04727;hsa04750;hsa04810;hsa04912;hsa04915;hsa04917;hsa04919;hsa04921;hsa05100;hsa05120;hsa05131;hsa05152;hsa05161;hsa05203;hsa05205;hsa05219 
                                               SRC 
                                               NA 
                                               slateblue1 
                                               16 
                                               14.1666666666667 
                                             
                                             
                                               PAK1-mRNA 
                                               -2.44 
                                               0.179 
                                               -2.79 
                                               -2.09 
                                               0.184 
                                               0.145 
                                               0.235 
                                               5.34e-16 
                                               1.72e-14 
                                               lm.nb 
                                               Activation of Rac, CD209 (DC-SIGN) signaling, CD28 co-stimulation, CD28 dependent Vav1 pathway, EPHB-mediated forward signaling, Ephrin signaling, FCERI mediated MAPK activation, MAPK6MAPK4 signaling, Regulation of actin dynamics for phagocytic cup formation, RHO GTPases activate PAKs, Sema3A PAK dependent Axon repulsion, VEGFA-VEGFR2 Pathway, VEGFR2 mediated vascular permeability 
                                               SEBELIUS_2_C4636 
                                               PAK1 
                                               mRNA 
                                               0 
                                               
                                               
                                               EPHB-mediated forward signaling;Ephrin signaling;MAPK6MAPK4 signaling;DSCAM interactions;Activation of Rac;VEGFR2 mediated vascular permeability;RHO GTPases activate PAKs;Sema3A PAK dependent Axon repulsion;FCERI mediated MAPK activation;Regulation of actin dynamics for phagocytic cup formation;CD209 (DC-SIGN) signaling;CD28 dependent Vav1 pathway;VEGFA-VEGFR2 Pathway;CD28 co-stimulation 
                                               hsa04010;hsa04012;hsa04014;hsa04024;hsa04062;hsa04360;hsa04510;hsa04650;hsa04660;hsa04666;hsa04810;hsa05120;hsa05205;hsa05211 
                                               PAK1 
                                               NA 
                                               slateblue1 
                                               16 
                                               -13.6312849162011 
                                             
                                             
                                               YES1-mRNA 
                                               1.14 
                                               0.0981 
                                               0.945 
                                               1.33 
                                               2.2 
                                               1.93 
                                               2.51 
                                               7.08e-14 
                                               1.9e-12 
                                               lm.nb 
                                               CD28 co-stimulation, Cell surface interactions at the vascular wall, CTLA4 inhibitory signaling, EPH-ephrin mediated repulsion of cells, EPHA-mediated growth cone collapse, EPHB-mediated forward signaling, Regulation of signaling by CBL, Signaling by ERBB2, Signaling by SCF-KIT 
                                               SEBELIUS_2_C4636 
                                               YES1 
                                               mRNA 
                                               0 
                                               
                                               
                                               Signaling by ERBB2;EPHB-mediated forward signaling;EPHA-mediated growth cone collapse;Regulation of KIT signaling;EPH-ephrin mediated repulsion of cells;Regulation of signaling by CBL;CD28 co-stimulation;CTLA4 inhibitory signaling;Signaling by SCF-KIT;Cell surface interactions at the vascular wall 
                                               hsa04520;hsa04530 
                                               YES1 
                                               NA 
                                               slateblue1 
                                               16 
                                               11.6207951070336 
                                             
                                             
                                               PIK3R1-mRNA 
                                               1.63 
                                               0.179 
                                               1.28 
                                               1.98 
                                               3.1 
                                               2.43 
                                               3.95 
                                               5.59e-11 
                                               1.13e-09 
                                               lm.nb 
                                               CD28 co-stimulation, CD28 dependent PI3KAkt signaling, Cell surface interactions at the vascular wall, Constitutive Signaling by Aberrant PI3K in Cancer, Constitutive Signaling by EGFRvIII, DAP12 signaling, Downstream signal transduction, Downstream TCR signaling, G alpha (q) signalling events, G beta.CL.gamma signalling through PI3Kgamma, GAB1 signalosome, Gastrin-CREB signalling pathway via PKC and MAPK, GPVI-mediated activation cascade, Interleukin receptor SHC signaling, IRS-related events triggered by IGF1R, Nephrin interactions, PI3KAKT activation, PIP3 activates AKT signaling, Regulation of signaling by CBL, Role of LAT2NTALLAB on calcium mobilization, Signaling by ERBB2, Signaling by FGFR1 in disease, Signaling by FGFR2 in disease, Signaling by FGFR3 in disease, Signaling by FGFR4 in disease, Signaling by PDGF, Signaling by SCF-KIT, Synthesis of PIPs at the plasma membrane, Tie2 Signaling, VEGFA-VEGFR2 Pathway 
                                               SEBELIUS_2_C4636 
                                               PIK3R1 
                                               mRNA 
                                               0 
                                               
                                               
                                               G alpha (q) signalling events;Downstream TCR signaling;Signaling by FGFR2 in disease;PIP3 activates AKT signaling;VEGFA-VEGFR2 Pathway;Synthesis of PIPs at the plasma membrane;Constitutive Signaling by Ligand-Responsive EGFR Cancer Variants;Interleukin receptor SHC signaling;G beta.CL.gamma signalling through PI3Kgamma;Regulation of signaling by CBL;Role of phospholipids in phagocytosis;Signaling by FGFR3 in disease;Constitutive Signaling by Aberrant PI3K in Cancer;Signaling by FGFR4 in disease;CD28 dependent PI3KAkt signaling;Tie2 Signaling;Signaling by FGFR1 fusion mutants;GP1b-IX-V activation signalling;GPVI-mediated activation cascade;Nephrin interactions;Constitutive Signaling by EGFRvIII;Gastrin-CREB signalling pathway via PKC and MAPK;Downstream signal transduction;GAB1 signalosome;Signaling by SCF-KIT;PI3KAKT activation;Role of LAT2NTALLAB on calcium mobilization;DAP12 signaling;Signaling by PDGF;Signaling by ERBB2;IRS-related events triggered by IGF1R;CD28 co-stimulation;Cell surface interactions at the vascular wall;Signaling by FGFR1 in disease 
                                               hsa04012;hsa04014;hsa04015;hsa04022;hsa04024;hsa04062;hsa04066;hsa04068;hsa04070;hsa04071;hsa04072;hsa04150;hsa04151;hsa04152;hsa04210;hsa04211;hsa04213;hsa04261;hsa04370;hsa04380;hsa04510;hsa04550;hsa04611;hsa04620;hsa04630;hsa04650;hsa04660;hsa04662;hsa04664;hsa04666;hsa04668;hsa04670;hsa04722;hsa04725;hsa04750;hsa04810;hsa04910;hsa04914;hsa04915;hsa04917;hsa04919;hsa04921;hsa04923;hsa04930;hsa04931;hsa04932;hsa04933;hsa04960;hsa04973;hsa05100;hsa05142;hsa05145;hsa05146;hsa05160;hsa05161;hsa05162;hsa05164;hsa05166;hsa05169;hsa05200;hsa05203;hsa05205;hsa05210;hsa05211;hsa05212;hsa05213;hsa05214;hsa05215;hsa05218;hsa05220;hsa05221;hsa05222;hsa05223;hsa05230;hsa05231 
                                               PIK3R1 
                                               NA 
                                               slateblue1 
                                               16 
                                               9.10614525139665 
                                             
                                             
                                               AKT2-mRNA 
                                               0.873 
                                               0.131 
                                               0.617 
                                               1.13 
                                               1.83 
                                               1.53 
                                               2.19 
                                               7.75e-08 
                                               1.06e-06 
                                               lm.nb 
                                               Activation of BAD and translocation to mitochondria , CD28 co-stimulation, DAP12 signaling, Deactivation of the beta-catenin transactivating complex, Downstream signal transduction, G beta.CL.gamma signalling through PI3Kgamma, GAB1 signalosome, GPVI-mediated activation cascade, IRS-related events triggered by IGF1R, PI3K Cascade, PI3KAKT activation, PIP3 activates AKT signaling, Role of LAT2NTALLAB on calcium mobilization, Signaling by ERBB2, Signaling by PDGF, Signaling by SCF-KIT, TP53 Regulates Metabolic Genes, VEGFA-VEGFR2 Pathway, VEGFR2 mediated vascular permeability 
                                               SEBELIUS_2_C4636 
                                               AKT2 
                                               mRNA 
                                               0 
                                               
                                               
                                               TP53 Regulates Metabolic Genes;PDE3B signalling;AKT phosphorylates targets in the cytosol;Negative regulation of the PI3KAKT network;Deactivation of the beta-catenin transactivating complex;Activation of BAD and translocation to mitochondria ;G beta.CL.gamma signalling through PI3Kgamma;AKT phosphorylates targets in the nucleus;VEGFR2 mediated vascular permeability;GPVI-mediated activation cascade;Signaling by ERBB2;PI3K Cascade;IRS-related events triggered by IGF1R;mTOR signalling;PIP3 activates AKT signaling;Downstream signal transduction;GAB1 signalosome;Signaling by SCF-KIT;PI3KAKT activation;Role of LAT2NTALLAB on calcium mobilization;DAP12 signaling;Signaling by PDGF;CD28 co-stimulation;VEGFA-VEGFR2 Pathway 
                                               hsa04010;hsa04012;hsa04014;hsa04015;hsa04022;hsa04024;hsa04062;hsa04066;hsa04068;hsa04071;hsa04072;hsa04150;hsa04151;hsa04152;hsa04210;hsa04211;hsa04213;hsa04261;hsa04370;hsa04380;hsa04510;hsa04530;hsa04550;hsa04611;hsa04620;hsa04630;hsa04660;hsa04662;hsa04664;hsa04666;hsa04668;hsa04722;hsa04725;hsa04728;hsa04910;hsa04914;hsa04915;hsa04917;hsa04919;hsa04920;hsa04922;hsa04923;hsa04931;hsa04932;hsa04933;hsa04973;hsa05142;hsa05145;hsa05152;hsa05160;hsa05161;hsa05162;hsa05164;hsa05166;hsa05169;hsa05200;hsa05205;hsa05210;hsa05211;hsa05212;hsa05213;hsa05214;hsa05215;hsa05218;hsa05220;hsa05221;hsa05222;hsa05223;hsa05230;hsa05231 
                                               AKT2 
                                               NA 
                                               slateblue1 
                                               16 
                                               6.66412213740458 
                                             
                                             
                                               PAK2-mRNA 
                                               0.446 
                                               0.0825 
                                               0.284 
                                               0.608 
                                               1.36 
                                               1.22 
                                               1.52 
                                               4.03e-06 
                                               4.65e-05 
                                               loglinear 
                                               Activation of Rac, CD209 (DC-SIGN) signaling, CD28 co-stimulation, CD28 dependent Vav1 pathway, Ephrin signaling, RHO GTPases activate PAKs, Sema3A PAK dependent Axon repulsion, VEGFA-VEGFR2 Pathway, VEGFR2 mediated vascular permeability 
                                               SEBELIUS_2_C4636 
                                               PAK2 
                                               mRNA 
                                               0 
                                               
                                               
                                               Nef and signal transduction;CD28 dependent Vav1 pathway;Stimulation of the cell death response by PAK-2p34;Ephrin signaling;Activation of Rac;RHO GTPases activate PAKs;VEGFR2 mediated vascular permeability;Sema3A PAK dependent Axon repulsion;CD209 (DC-SIGN) signaling;VEGFA-VEGFR2 Pathway;CD28 co-stimulation 
                                               hsa04010;hsa04012;hsa04014;hsa04360;hsa04510;hsa04660;hsa04810;hsa05211 
                                               PAK2 
                                               NA 
                                               slateblue1 
                                               16 
                                               5.40606060606061 
                                             
                                             
                                               PIK3R3-mRNA 
                                               -1.77 
                                               0.457 
                                               -2.67 
                                               -0.874 
                                               0.293 
                                               0.157 
                                               0.546 
                                               0.000425 
                                               0.00402 
                                               lm.nb 
                                               CD28 co-stimulation, CD28 dependent PI3KAkt signaling, Constitutive Signaling by Aberrant PI3K in Cancer, DAP12 signaling, Downstream signal transduction, G alpha (1213) signalling events, G alpha (q) signalling events, G beta.CL.gamma signalling through PI3Kgamma, GAB1 signalosome, Gastrin-CREB signalling pathway via PKC and MAPK, GPVI-mediated activation cascade, Interleukin receptor SHC signaling, PI3KAKT activation, PIP3 activates AKT signaling, Regulation of signaling by CBL, Role of LAT2NTALLAB on calcium mobilization, Signaling by ERBB2, Signaling by PDGF, Signaling by SCF-KIT, Synthesis of PIPs at the plasma membrane 
                                               SEBELIUS_2_C4636 
                                               PIK3R3 
                                               mRNA 
                                               0 
                                               
                                               
                                               G alpha (q) signalling events;PIP3 activates AKT signaling;Constitutive Signaling by Aberrant PI3K in Cancer;CD28 dependent PI3KAkt signaling;Synthesis of PIPs at the plasma membrane;Interleukin receptor SHC signaling;G beta.CL.gamma signalling through PI3Kgamma;Regulation of signaling by CBL;GPVI-mediated activation cascade;G alpha (1213) signalling events;Gastrin-CREB signalling pathway via PKC and MAPK;Downstream signal transduction;GAB1 signalosome;Signaling by SCF-KIT;PI3KAKT activation;Role of LAT2NTALLAB on calcium mobilization;DAP12 signaling;Signaling by PDGF;Signaling by ERBB2;CD28 co-stimulation 
                                               hsa04012;hsa04014;hsa04015;hsa04022;hsa04024;hsa04062;hsa04066;hsa04068;hsa04070;hsa04071;hsa04072;hsa04150;hsa04151;hsa04152;hsa04210;hsa04211;hsa04213;hsa04261;hsa04370;hsa04380;hsa04510;hsa04550;hsa04611;hsa04620;hsa04630;hsa04650;hsa04660;hsa04662;hsa04664;hsa04666;hsa04668;hsa04670;hsa04722;hsa04725;hsa04750;hsa04810;hsa04910;hsa04914;hsa04915;hsa04917;hsa04919;hsa04921;hsa04923;hsa04930;hsa04931;hsa04932;hsa04933;hsa04960;hsa04973;hsa05100;hsa05142;hsa05145;hsa05146;hsa05160;hsa05161;hsa05162;hsa05164;hsa05166;hsa05169;hsa05200;hsa05203;hsa05205;hsa05210;hsa05211;hsa05212;hsa05213;hsa05214;hsa05215;hsa05218;hsa05220;hsa05221;hsa05222;hsa05223;hsa05230;hsa05231 
                                               PIK3R3 
                                               NA 
                                               slateblue1 
                                               16 
                                               -3.87308533916849 
                                             
                                             
                                               PIK3R2-mRNA 
                                               0.61 
                                               0.207 
                                               0.204 
                                               1.02 
                                               1.53 
                                               1.15 
                                               2.02 
                                               0.00552 
                                               0.0469 
                                               lm.nb 
                                               CD28 co-stimulation, CD28 dependent PI3KAkt signaling, Cell surface interactions at the vascular wall, Constitutive Signaling by Aberrant PI3K in Cancer, DAP12 signaling, Downstream signal transduction, Downstream TCR signaling, G alpha (1213) signalling events, G alpha (q) signalling events, G beta.CL.gamma signalling through PI3Kgamma, GAB1 signalosome, Gastrin-CREB signalling pathway via PKC and MAPK, GPVI-mediated activation cascade, Interleukin receptor SHC signaling, IRS-related events triggered by IGF1R, Nephrin interactions, PI3K Cascade, PI3KAKT activation, PIP3 activates AKT signaling, Regulation of signaling by CBL, Rho GTPase cycle, Role of LAT2NTALLAB on calcium mobilization, Signaling by ERBB2, Signaling by PDGF, Signaling by SCF-KIT, Synthesis of PIPs at the plasma membrane, Tie2 Signaling, VEGFA-VEGFR2 Pathway 
                                               SEBELIUS_2_C4636 
                                               PIK3R2 
                                               mRNA 
                                               0 
                                               
                                               
                                               G alpha (q) signalling events;Downstream TCR signaling;PIP3 activates AKT signaling;VEGFA-VEGFR2 Pathway;Synthesis of PIPs at the plasma membrane;Rho GTPase cycle;Interleukin receptor SHC signaling;G beta.CL.gamma signalling through PI3Kgamma;Regulation of signaling by CBL;Role of phospholipids in phagocytosis;G alpha (1213) signalling events;PI3K Cascade;Constitutive Signaling by Aberrant PI3K in Cancer;CD28 dependent PI3KAkt signaling;Tie2 Signaling;GPVI-mediated activation cascade;Nephrin interactions;Gastrin-CREB signalling pathway via PKC and MAPK;Downstream signal transduction;GAB1 signalosome;Signaling by SCF-KIT;PI3KAKT activation;Role of LAT2NTALLAB on calcium mobilization;DAP12 signaling;Signaling by PDGF;Signaling by ERBB2;IRS-related events triggered by IGF1R;CD28 co-stimulation;Cell surface interactions at the vascular wall 
                                               hsa04012;hsa04014;hsa04015;hsa04022;hsa04024;hsa04062;hsa04066;hsa04068;hsa04070;hsa04071;hsa04072;hsa04150;hsa04151;hsa04152;hsa04210;hsa04211;hsa04213;hsa04261;hsa04370;hsa04380;hsa04510;hsa04550;hsa04611;hsa04620;hsa04630;hsa04650;hsa04660;hsa04662;hsa04664;hsa04666;hsa04668;hsa04670;hsa04722;hsa04725;hsa04750;hsa04810;hsa04910;hsa04914;hsa04915;hsa04917;hsa04919;hsa04921;hsa04923;hsa04930;hsa04931;hsa04932;hsa04933;hsa04960;hsa04973;hsa05100;hsa05142;hsa05145;hsa05146;hsa05160;hsa05161;hsa05162;hsa05164;hsa05166;hsa05169;hsa05200;hsa05203;hsa05205;hsa05206;hsa05210;hsa05211;hsa05212;hsa05213;hsa05214;hsa05215;hsa05218;hsa05220;hsa05221;hsa05222;hsa05223;hsa05230;hsa05231 
                                               PIK3R2 
                                               NA 
                                               slateblue3 
                                               16 
                                               2.94685990338164 
                                             
                                             
                                               PIK3CA-mRNA 
                                               -0.437 
                                               0.159 
                                               -0.748 
                                               -0.126 
                                               0.739 
                                               0.595 
                                               0.916 
                                               0.00908 
                                               0.0753 
                                               lm.nb 
                                               CD28 co-stimulation, CD28 dependent PI3KAkt signaling, Cell surface interactions at the vascular wall, Constitutive Signaling by Aberrant PI3K in Cancer, Constitutive Signaling by EGFRvIII, DAP12 signaling, Downstream signal transduction, Downstream TCR signaling, G alpha (1213) signalling events, G alpha (q) signalling events, G beta.CL.gamma signalling through PI3Kgamma, GAB1 signalosome, Gastrin-CREB signalling pathway via PKC and MAPK, GPVI-mediated activation cascade, Interleukin receptor SHC signaling, IRS-related events triggered by IGF1R, Nephrin interactions, PI3K Cascade, PI3KAKT activation, PIP3 activates AKT signaling, Regulation of signaling by CBL, Role of LAT2NTALLAB on calcium mobilization, Signaling by ERBB2, Signaling by FGFR1 in disease, Signaling by FGFR2 in disease, Signaling by FGFR3 in disease, Signaling by FGFR4 in disease, Signaling by PDGF, Signaling by SCF-KIT, Synthesis of PIPs at the plasma membrane, Tie2 Signaling, VEGFA-VEGFR2 Pathway 
                                               SEBELIUS_2_C4636 
                                               PIK3CA 
                                               mRNA 
                                               0 
                                               
                                               
                                               G alpha (q) signalling events;Downstream TCR signaling;Signaling by FGFR2 in disease;PIP3 activates AKT signaling;VEGFA-VEGFR2 Pathway;Synthesis of PIPs at the plasma membrane;Constitutive Signaling by Ligand-Responsive EGFR Cancer Variants;Interleukin receptor SHC signaling;G beta.CL.gamma signalling through PI3Kgamma;Regulation of signaling by CBL;Role of phospholipids in phagocytosis;G alpha (1213) signalling events;PI3K Cascade;Signaling by FGFR3 in disease;Constitutive Signaling by Aberrant PI3K in Cancer;Signaling by FGFR4 in disease;CD28 dependent PI3KAkt signaling;Tie2 Signaling;GPVI-mediated activation cascade;Nephrin interactions;Constitutive Signaling by EGFRvIII;Gastrin-CREB signalling pathway via PKC and MAPK;Downstream signal transduction;GAB1 signalosome;Signaling by SCF-KIT;PI3KAKT activation;Role of LAT2NTALLAB on calcium mobilization;DAP12 signaling;Signaling by PDGF;Signaling by ERBB2;IRS-related events triggered by IGF1R;CD28 co-stimulation;Cell surface interactions at the vascular wall;Signaling by FGFR1 in disease 
                                               hsa00562;hsa04012;hsa04014;hsa04015;hsa04022;hsa04024;hsa04062;hsa04066;hsa04068;hsa04070;hsa04071;hsa04072;hsa04150;hsa04151;hsa04152;hsa04210;hsa04211;hsa04213;hsa04261;hsa04370;hsa04380;hsa04510;hsa04550;hsa04611;hsa04620;hsa04630;hsa04650;hsa04660;hsa04662;hsa04664;hsa04666;hsa04668;hsa04670;hsa04722;hsa04725;hsa04750;hsa04810;hsa04910;hsa04914;hsa04915;hsa04917;hsa04919;hsa04921;hsa04923;hsa04930;hsa04931;hsa04932;hsa04933;hsa04960;hsa04973;hsa05100;hsa05142;hsa05145;hsa05146;hsa05160;hsa05161;hsa05162;hsa05164;hsa05166;hsa05169;hsa05200;hsa05203;hsa05205;hsa05206;hsa05210;hsa05211;hsa05212;hsa05213;hsa05214;hsa05215;hsa05218;hsa05220;hsa05221;hsa05222;hsa05223;hsa05230;hsa05231 
                                               PIK3CA 
                                               NA 
                                               slateblue3 
                                               16 
                                               -2.74842767295597 
                                             
                                             
                                               FYN-mRNA 
                                               0.596 
                                               0.365 
                                               -0.119 
                                               1.31 
                                               1.51 
                                               0.921 
                                               2.48 
                                               0.111 
                                               0.804 
                                               lm.nb 
                                               ARMS-mediated activation, CD209 (DC-SIGN) signaling, CD28 co-stimulation, CD28 dependent PI3KAkt signaling, CD28 dependent Vav1 pathway, Cell surface interactions at the vascular wall, Constitutive Signaling by Aberrant PI3K in Cancer, CTLA4 inhibitory signaling, DAP12 signaling, Downstream signal transduction, EPH-ephrin mediated repulsion of cells, FCERI mediated MAPK activation, Frs2-mediated activation, GAB1 signalosome, Gastrin-CREB signalling pathway via PKC and MAPK, GPVI-mediated activation cascade, Interleukin receptor SHC signaling, IRS-related events triggered by IGF1R, Nephrin interactions, PI3KAKT activation, RAFMAP kinase cascade, Regulation of signaling by CBL, Role of LAT2NTALLAB on calcium mobilization, Sema3A PAK dependent Axon repulsion, Signaling by ERBB2, Signaling by PDGF, Signaling by SCF-KIT, VEGFA-VEGFR2 Pathway, VEGFR2 mediated cell proliferation 
                                               SEBELIUS_2_C4636 
                                               FYN 
                                               mRNA 
                                               0 
                                               
                                               
                                               EPH-ephrin mediated repulsion of cells;Regulation of signaling by CBL;FCGR activation;Antigen activates B Cell Receptor (BCR) leading to generation of second messengers;Dectin-2 family;RAFMAP kinase cascade;Constitutive Signaling by Aberrant PI3K in Cancer;CTLA4 inhibitory signaling;CD28 dependent PI3KAkt signaling;PECAM1 interactions;CD28 dependent Vav1 pathway;GPVI-mediated activation cascade;Sema3A PAK dependent Axon repulsion;Nephrin interactions;SEMA3A-Plexin repulsion signaling by inhibiting Integrin adhesion;CRMPs in Sema3A signaling;CD209 (DC-SIGN) signaling;Downstream signal transduction;GAB1 signalosome;Signaling by SCF-KIT;PI3KAKT activation;Role of LAT2NTALLAB on calcium mobilization;DAP12 signaling;Signaling by PDGF;Signaling by ERBB2;VEGFA-VEGFR2 Pathway;VEGFR2 mediated cell proliferation;Frs2-mediated activation;ARMS-mediated activation;Gastrin-CREB signalling pathway via PKC and MAPK;Interleukin receptor SHC signaling;FCERI mediated MAPK activation;IRS-related events triggered by IGF1R;CD28 co-stimulation;Cell surface interactions at the vascular wall 
                                               hsa04071;hsa04072;hsa04360;hsa04380;hsa04510;hsa04520;hsa04611;hsa04650;hsa04660;hsa04664;hsa04725;hsa05020;hsa05130;hsa05162;hsa05416 
                                               FYN 
                                               NA 
                                               azure4 
                                               1 
                                               1.63287671232877 
                                             
                                             
                                               CDC42-mRNA 
                                               -0.164 
                                               0.106 
                                               -0.372 
                                               0.0443 
                                               0.893 
                                               0.773 
                                               1.03 
                                               0.131 
                                               0.941 
                                               lm.nb 
                                               CD28 co-stimulation, CD28 dependent Vav1 pathway, DCC mediated attractive signaling, EPHB-mediated forward signaling, G alpha (1213) signalling events, GPVI-mediated activation cascade, Regulation of actin dynamics for phagocytic cup formation, Rho GTPase cycle, RHO GTPases Activate Formins, RHO GTPases activate IQGAPs, RHO GTPases activate PAKs, Sema4D induced cell migration and growth-cone collapse, VEGFA-VEGFR2 Pathway 
                                               SEBELIUS_2_C4636 
                                               CDC42 
                                               mRNA 
                                               0 
                                               
                                               
                                               DCC mediated attractive signaling;Factors involved in megakaryocyte development and platelet production;VEGFA-VEGFR2 Pathway;RHO GTPases activate KTN1;CD28 dependent Vav1 pathway;Inactivation of Cdc42 and Rac;Rho GTPase cycle;EPHB-mediated forward signaling;RHO GTPases Activate Formins;RHO GTPases activate PAKs;GPVI-mediated activation cascade;G alpha (1213) signalling events;RHO GTPases activate IQGAPs;Sema4D induced cell migration and growth-cone collapse;Regulation of actin dynamics for phagocytic cup formation;CD28 co-stimulation 
                                               hsa04010;hsa04014;hsa04015;hsa04062;hsa04144;hsa04360;hsa04370;hsa04510;hsa04520;hsa04530;hsa04660;hsa04666;hsa04670;hsa04722;hsa04810;hsa04912;hsa04932;hsa04933;hsa05100;hsa05120;hsa05130;hsa05131;hsa05132;hsa05200;hsa05203;hsa05205;hsa05211;hsa05212 
                                               CDC42 
                                               NA 
                                               azure4 
                                               1 
                                               -1.54716981132075 
                                             
                                             
                                               RAC1-mRNA 
                                               -0.0792 
                                               0.102 
                                               -0.279 
                                               0.121 
                                               0.947 
                                               0.824 
                                               1.09 
                                               0.442 
                                               1 
                                               lm.nb 
                                               Activation of Rac, CD28 co-stimulation, CD28 dependent Vav1 pathway, DAP12 signaling, EPH-ephrin mediated repulsion of cells, Ephrin signaling, G alpha (1213) signalling events, GPVI-mediated activation cascade, MAPK6MAPK4 signaling, NRAGE signals death through JNK, PCPCE pathway, Regulation of actin dynamics for phagocytic cup formation, RHO GTPases activate CIT, RHO GTPases Activate Formins, RHO GTPases activate IQGAPs, RHO GTPases activate PAKs, RHO GTPases activate PKNs, RHO GTPases Activate WASPs and WAVEs, Sema3A PAK dependent Axon repulsion, Sema4D induced cell migration and growth-cone collapse, Signal transduction by L1, Signaling by SCF-KIT, VEGFA-VEGFR2 Pathway, VEGFR2 mediated vascular permeability 
                                               SEBELIUS_2_C4636 
                                               RAC1 
                                               mRNA 
                                               0 
                                               
                                               
                                               MAPK6MAPK4 signaling;Signaling by SCF-KIT;Inactivation of Cdc42 and Rac;DAP12 signaling;Ephrin signaling;EPH-ephrin mediated repulsion of cells;Activation of Rac;RHO GTPases Activate Formins;VEGFR2 mediated vascular permeability;RHO GTPases activate PAKs;Sema4D mediated inhibition of cell attachment and migration;G alpha (1213) signalling events;Regulation of actin dynamics for phagocytic cup formation;Sema4D induced cell migration and growth-cone collapse;Signal transduction by L1;RHO GTPases activate CIT;RHO GTPases Activate WASPs and WAVEs;Factors involved in megakaryocyte development and platelet production;RHO GTPases activate KTN1;CD28 dependent Vav1 pathway;DSCAM interactions;NRAGE signals death through JNK;GPVI-mediated activation cascade;Sema3A PAK dependent Axon repulsion;RHO GTPases activate PKNs;SEMA3A-Plexin repulsion signaling by inhibiting Integrin adhesion;RHO GTPases activate IQGAPs;PCPCE pathway;VEGFA-VEGFR2 Pathway;CD28 co-stimulation 
                                               hsa04010;hsa04014;hsa04015;hsa04024;hsa04062;hsa04071;hsa04145;hsa04151;hsa04310;hsa04360;hsa04370;hsa04380;hsa04510;hsa04520;hsa04620;hsa04650;hsa04662;hsa04664;hsa04666;hsa04670;hsa04722;hsa04810;hsa04932;hsa04933;hsa04972;hsa05100;hsa05120;hsa05131;hsa05132;hsa05200;hsa05203;hsa05205;hsa05210;hsa05211;hsa05212;hsa05231;hsa05416 
                                               RAC1 
                                               NA 
                                               azure4 
                                               1 
                                               -0.776470588235294 
                                             
                                             
                                               AKT1-mRNA 
                                               0.141 
                                               0.256 
                                               -0.361 
                                               0.643 
                                               1.1 
                                               0.779 
                                               1.56 
                                               0.585 
                                               1 
                                               lm.nb 
                                               Activation of BAD and translocation to mitochondria , CD28 co-stimulation, DAP12 signaling, Deactivation of the beta-catenin transactivating complex, Downstream signal transduction, G beta.CL.gamma signalling through PI3Kgamma, GAB1 signalosome, GPVI-mediated activation cascade, IRS-related events triggered by IGF1R, PI3K Cascade, PI3KAKT activation, PIP3 activates AKT signaling, Role of LAT2NTALLAB on calcium mobilization, Signaling by ERBB2, Signaling by PDGF, Signaling by SCF-KIT, TP53 Regulates Metabolic Genes, VEGFA-VEGFR2 Pathway, VEGFR2 mediated vascular permeability 
                                               SEBELIUS_2_C4636 
                                               AKT1 
                                               mRNA 
                                               0 
                                               
                                               
                                               TP53 Regulates Metabolic Genes;Deactivation of the beta-catenin transactivating complex;Activation of BAD and translocation to mitochondria ;eNOS activation;mTOR signalling;G beta.CL.gamma signalling through PI3Kgamma;KSRP (KHSRP) binds and destabilizes mRNA;AKT phosphorylates targets in the nucleus;VEGFR2 mediated vascular permeability;GPVI-mediated activation cascade;Tetrahydrobiopterin (BH4) synthesis# recycling# salvage and regulation;Butyrate Response Factor 1 (BRF1) binds and destabilizes mRNA;Signaling by ERBB2;PIP3 activates AKT signaling;Downstream signal transduction;GAB1 signalosome;Signaling by SCF-KIT;PI3KAKT activation;Role of LAT2NTALLAB on calcium mobilization;DAP12 signaling;Signaling by PDGF;CD28 co-stimulation;PI3K Cascade;IRS-related events triggered by IGF1R;VEGFA-VEGFR2 Pathway 
                                               hsa04010;hsa04012;hsa04014;hsa04015;hsa04022;hsa04024;hsa04062;hsa04066;hsa04068;hsa04071;hsa04072;hsa04150;hsa04151;hsa04152;hsa04210;hsa04211;hsa04213;hsa04261;hsa04370;hsa04380;hsa04510;hsa04530;hsa04550;hsa04611;hsa04620;hsa04630;hsa04660;hsa04662;hsa04664;hsa04666;hsa04668;hsa04722;hsa04725;hsa04728;hsa04910;hsa04914;hsa04915;hsa04917;hsa04919;hsa04920;hsa04922;hsa04923;hsa04931;hsa04932;hsa04933;hsa04973;hsa05142;hsa05145;hsa05152;hsa05160;hsa05161;hsa05162;hsa05164;hsa05166;hsa05169;hsa05200;hsa05205;hsa05210;hsa05211;hsa05212;hsa05213;hsa05214;hsa05215;hsa05218;hsa05220;hsa05221;hsa05222;hsa05223;hsa05230;hsa05231 
                                               AKT1 
                                               NA 
                                               azure4 
                                               1 
                                               0.55078125 
                                             
                                           
                                           DE Results - CD28 co-stimulation genes - Line: HEPG2 vs.Fibro  Table displaying each sample's global significance scores and directed global significance scores as defined in the heatmaps above. The global significance score is calculated as the square root of the mean squared t-statistic for the genes in a gene set, with t-statistics coming from the linear regression underlying our differential expression analysis. The directed global significance score is calculated as the square root of the mean signed squared t-statistic for the genes in a gene set, with t-statistics coming from the linear regression underlying our differential expression analysis.   
                                      
                                       
                                  
                                
                                
                                  
                                       
                                      
                                           Volcano Plot: Line: HT29 vs.Fibro   More Plot Information                Volcano Plot: Line: HT29 vs.Fibro  Volcano plot displaying each gene's -log10(p-value) and log2 fold change for the selected covariate. Highly statistically significant genes fall at the top of the plot, and highly differentially expressed genes fall to either side. Genes within the selected gene set are highlighted in orange. Horizontal lines indicate various False Discovery Rate (FDR) thresholds.   
                                           DE Results - CD28 co-stimulation genes - Line: HT29 vs.Fibro   More Plot Information      Download CSV Data     
                                           
                                             
                                             
                                           
                                           
                                             
                                               Probe Label 
                                               Log2 fold change 
                                               std error (log2) 
                                               Lower confidence limit (log2) 
                                               Upper confidence limit (log2) 
                                               Linear fold change 
                                               Lower confidence limit (linear) 
                                               Upper confidence limit (linear) 
                                               P-value 
                                               BY.p.value 
                                               method 
                                               Gene.sets 
                                               Codeset.Name 
                                               Probe.Label 
                                               Analyte.Type 
                                               Is.Control 
                                               Control.Type 
                                               Related.Probes 
                                               Probe.Annotation 
                                               KEGG.Pathways 
                                               Official.Gene.Name 
                                               Control.Conc 
                                               volcanocol 
                                               volcanopch 
                                               tstats.all 
                                             
                                           
                                           
                                             
                                               AKT3-mRNA 
                                               -8.38 
                                               0.265 
                                               -8.9 
                                               -7.86 
                                               0.003 
                                               0.00209 
                                               0.0043 
                                               2.3e-28 
                                               4.09e-26 
                                               lm.nb 
                                               Activation of BAD and translocation to mitochondria , CD28 co-stimulation, CD28 dependent PI3KAkt signaling, DAP12 signaling, Downstream signal transduction, G beta.CL.gamma signalling through PI3Kgamma, GAB1 signalosome, GPVI-mediated activation cascade, PI3KAKT activation, PIP3 activates AKT signaling, Role of LAT2NTALLAB on calcium mobilization, Signaling by ERBB2, Signaling by PDGF, Signaling by SCF-KIT, TP53 Regulates Metabolic Genes, VEGFA-VEGFR2 Pathway, VEGFR2 mediated vascular permeability 
                                               SEBELIUS_2_C4636 
                                               AKT3 
                                               mRNA 
                                               0 
                                               
                                               
                                               TP53 Regulates Metabolic Genes;CD28 dependent PI3KAkt signaling;AKT phosphorylates targets in the cytosol;Negative regulation of the PI3KAKT network;Activation of BAD and translocation to mitochondria ;G beta.CL.gamma signalling through PI3Kgamma;AKT phosphorylates targets in the nucleus;VEGFR2 mediated vascular permeability;GPVI-mediated activation cascade;Signaling by ERBB2;PIP3 activates AKT signaling;Downstream signal transduction;GAB1 signalosome;Signaling by SCF-KIT;PI3KAKT activation;Role of LAT2NTALLAB on calcium mobilization;DAP12 signaling;Signaling by PDGF;CD28 co-stimulation;VEGFA-VEGFR2 Pathway 
                                               hsa04010;hsa04012;hsa04014;hsa04015;hsa04022;hsa04024;hsa04062;hsa04066;hsa04068;hsa04071;hsa04072;hsa04150;hsa04151;hsa04152;hsa04210;hsa04211;hsa04213;hsa04261;hsa04370;hsa04380;hsa04510;hsa04530;hsa04550;hsa04611;hsa04620;hsa04630;hsa04660;hsa04662;hsa04664;hsa04666;hsa04668;hsa04722;hsa04725;hsa04728;hsa04910;hsa04914;hsa04915;hsa04917;hsa04919;hsa04920;hsa04922;hsa04923;hsa04931;hsa04932;hsa04933;hsa04973;hsa05142;hsa05145;hsa05152;hsa05160;hsa05161;hsa05162;hsa05164;hsa05166;hsa05169;hsa05200;hsa05205;hsa05210;hsa05211;hsa05212;hsa05213;hsa05214;hsa05215;hsa05218;hsa05220;hsa05221;hsa05222;hsa05223;hsa05230;hsa05231 
                                               AKT3 
                                               NA 
                                               slateblue1 
                                               16 
                                               -31.622641509434 
                                             
                                             
                                               YES1-mRNA 
                                               1.56 
                                               0.0925 
                                               1.38 
                                               1.75 
                                               2.96 
                                               2.61 
                                               3.35 
                                               5.35e-19 
                                               2.11e-17 
                                               lm.nb 
                                               CD28 co-stimulation, Cell surface interactions at the vascular wall, CTLA4 inhibitory signaling, EPH-ephrin mediated repulsion of cells, EPHA-mediated growth cone collapse, EPHB-mediated forward signaling, Regulation of signaling by CBL, Signaling by ERBB2, Signaling by SCF-KIT 
                                               SEBELIUS_2_C4636 
                                               YES1 
                                               mRNA 
                                               0 
                                               
                                               
                                               Signaling by ERBB2;EPHB-mediated forward signaling;EPHA-mediated growth cone collapse;Regulation of KIT signaling;EPH-ephrin mediated repulsion of cells;Regulation of signaling by CBL;CD28 co-stimulation;CTLA4 inhibitory signaling;Signaling by SCF-KIT;Cell surface interactions at the vascular wall 
                                               hsa04520;hsa04530 
                                               YES1 
                                               NA 
                                               slateblue1 
                                               16 
                                               16.8648648648649 
                                             
                                             
                                               PIK3R1-mRNA 
                                               2.62 
                                               0.169 
                                               2.29 
                                               2.95 
                                               6.15 
                                               4.89 
                                               7.74 
                                               8.93e-18 
                                               2.88e-16 
                                               lm.nb 
                                               CD28 co-stimulation, CD28 dependent PI3KAkt signaling, Cell surface interactions at the vascular wall, Constitutive Signaling by Aberrant PI3K in Cancer, Constitutive Signaling by EGFRvIII, DAP12 signaling, Downstream signal transduction, Downstream TCR signaling, G alpha (q) signalling events, G beta.CL.gamma signalling through PI3Kgamma, GAB1 signalosome, Gastrin-CREB signalling pathway via PKC and MAPK, GPVI-mediated activation cascade, Interleukin receptor SHC signaling, IRS-related events triggered by IGF1R, Nephrin interactions, PI3KAKT activation, PIP3 activates AKT signaling, Regulation of signaling by CBL, Role of LAT2NTALLAB on calcium mobilization, Signaling by ERBB2, Signaling by FGFR1 in disease, Signaling by FGFR2 in disease, Signaling by FGFR3 in disease, Signaling by FGFR4 in disease, Signaling by PDGF, Signaling by SCF-KIT, Synthesis of PIPs at the plasma membrane, Tie2 Signaling, VEGFA-VEGFR2 Pathway 
                                               SEBELIUS_2_C4636 
                                               PIK3R1 
                                               mRNA 
                                               0 
                                               
                                               
                                               G alpha (q) signalling events;Downstream TCR signaling;Signaling by FGFR2 in disease;PIP3 activates AKT signaling;VEGFA-VEGFR2 Pathway;Synthesis of PIPs at the plasma membrane;Constitutive Signaling by Ligand-Responsive EGFR Cancer Variants;Interleukin receptor SHC signaling;G beta.CL.gamma signalling through PI3Kgamma;Regulation of signaling by CBL;Role of phospholipids in phagocytosis;Signaling by FGFR3 in disease;Constitutive Signaling by Aberrant PI3K in Cancer;Signaling by FGFR4 in disease;CD28 dependent PI3KAkt signaling;Tie2 Signaling;Signaling by FGFR1 fusion mutants;GP1b-IX-V activation signalling;GPVI-mediated activation cascade;Nephrin interactions;Constitutive Signaling by EGFRvIII;Gastrin-CREB signalling pathway via PKC and MAPK;Downstream signal transduction;GAB1 signalosome;Signaling by SCF-KIT;PI3KAKT activation;Role of LAT2NTALLAB on calcium mobilization;DAP12 signaling;Signaling by PDGF;Signaling by ERBB2;IRS-related events triggered by IGF1R;CD28 co-stimulation;Cell surface interactions at the vascular wall;Signaling by FGFR1 in disease 
                                               hsa04012;hsa04014;hsa04015;hsa04022;hsa04024;hsa04062;hsa04066;hsa04068;hsa04070;hsa04071;hsa04072;hsa04150;hsa04151;hsa04152;hsa04210;hsa04211;hsa04213;hsa04261;hsa04370;hsa04380;hsa04510;hsa04550;hsa04611;hsa04620;hsa04630;hsa04650;hsa04660;hsa04662;hsa04664;hsa04666;hsa04668;hsa04670;hsa04722;hsa04725;hsa04750;hsa04810;hsa04910;hsa04914;hsa04915;hsa04917;hsa04919;hsa04921;hsa04923;hsa04930;hsa04931;hsa04932;hsa04933;hsa04960;hsa04973;hsa05100;hsa05142;hsa05145;hsa05146;hsa05160;hsa05161;hsa05162;hsa05164;hsa05166;hsa05169;hsa05200;hsa05203;hsa05205;hsa05210;hsa05211;hsa05212;hsa05213;hsa05214;hsa05215;hsa05218;hsa05220;hsa05221;hsa05222;hsa05223;hsa05230;hsa05231 
                                               PIK3R1 
                                               NA 
                                               slateblue1 
                                               16 
                                               15.5029585798817 
                                             
                                             
                                               PAK1-mRNA 
                                               2.31 
                                               0.165 
                                               1.99 
                                               2.63 
                                               4.96 
                                               3.96 
                                               6.2 
                                               2.18e-16 
                                               5.51e-15 
                                               lm.nb 
                                               Activation of Rac, CD209 (DC-SIGN) signaling, CD28 co-stimulation, CD28 dependent Vav1 pathway, EPHB-mediated forward signaling, Ephrin signaling, FCERI mediated MAPK activation, MAPK6MAPK4 signaling, Regulation of actin dynamics for phagocytic cup formation, RHO GTPases activate PAKs, Sema3A PAK dependent Axon repulsion, VEGFA-VEGFR2 Pathway, VEGFR2 mediated vascular permeability 
                                               SEBELIUS_2_C4636 
                                               PAK1 
                                               mRNA 
                                               0 
                                               
                                               
                                               EPHB-mediated forward signaling;Ephrin signaling;MAPK6MAPK4 signaling;DSCAM interactions;Activation of Rac;VEGFR2 mediated vascular permeability;RHO GTPases activate PAKs;Sema3A PAK dependent Axon repulsion;FCERI mediated MAPK activation;Regulation of actin dynamics for phagocytic cup formation;CD209 (DC-SIGN) signaling;CD28 dependent Vav1 pathway;VEGFA-VEGFR2 Pathway;CD28 co-stimulation 
                                               hsa04010;hsa04012;hsa04014;hsa04024;hsa04062;hsa04360;hsa04510;hsa04650;hsa04660;hsa04666;hsa04810;hsa05120;hsa05205;hsa05211 
                                               PAK1 
                                               NA 
                                               slateblue1 
                                               16 
                                               14 
                                             
                                             
                                               SRC-mRNA 
                                               2.21 
                                               0.159 
                                               1.9 
                                               2.52 
                                               4.62 
                                               3.73 
                                               5.73 
                                               2.85e-16 
                                               6.85e-15 
                                               lm.nb 
                                               ARMS-mediated activation, CD28 co-stimulation, Cell surface interactions at the vascular wall, CTLA4 inhibitory signaling, DAP12 signaling, Downstream signal transduction, FCERI mediated MAPK activation, Frs2-mediated activation, GAB1 signalosome, Gastrin-CREB signalling pathway via PKC and MAPK, Integrin alphaIIb beta3 signaling, Interleukin receptor SHC signaling, IRS-related events triggered by IGF1R, MAP2K and MAPK activation, Negative regulation of FGFR1 signaling, Negative regulation of FGFR2 signaling, Negative regulation of FGFR3 signaling, Negative regulation of FGFR4 signaling, p130Cas linkage to MAPK signaling for integrins, RAFMAP kinase cascade, Signaling by ERBB2, Signaling by PDGF, Signaling by SCF-KIT, Spry regulation of FGF signaling, VEGFA-VEGFR2 Pathway, VEGFR2 mediated cell proliferation 
                                               SEBELIUS_2_C4636 
                                               SRC 
                                               mRNA 
                                               0 
                                               
                                               
                                               FCGR activation;Spry regulation of FGF signaling;GAB1 signalosome;MAP2K and MAPK activation;CLEC7A (Dectin-1) signaling;CD28 co-stimulation;p130Cas linkage to MAPK signaling for integrins;CTLA4 inhibitory signaling;GRB2.CL.SOS provides linkage to MAPK signaling for Integrins ;Signal regulatory protein (SIRP) family interactions;PECAM1 interactions;Thrombin signalling through proteinase activated receptors (PARs);GP1b-IX-V activation signalling;c-src mediated regulation of Cx43 function and closure of gap junctions;Signaling by ERBB2;VEGFA-VEGFR2 Pathway;Signaling by SCF-KIT;VEGFR2 mediated cell proliferation;RAFMAP kinase cascade;Downstream signal transduction;Frs2-mediated activation;ARMS-mediated activation;Gastrin-CREB signalling pathway via PKC and MAPK;Interleukin receptor SHC signaling;FCERI mediated MAPK activation;DAP12 signaling;Signaling by PDGF;IRS-related events triggered by IGF1R;Negative regulation of FGFR3 signaling;Negative regulation of FGFR4 signaling;Negative regulation of FGFR2 signaling;Negative regulation of FGFR1 signaling;Integrin alphaIIb beta3 signaling;Cell surface interactions at the vascular wall 
                                               hsa04012;hsa04015;hsa04062;hsa04144;hsa04370;hsa04510;hsa04520;hsa04530;hsa04540;hsa04611;hsa04727;hsa04750;hsa04810;hsa04912;hsa04915;hsa04917;hsa04919;hsa04921;hsa05100;hsa05120;hsa05131;hsa05152;hsa05161;hsa05203;hsa05205;hsa05219 
                                               SRC 
                                               NA 
                                               slateblue1 
                                               16 
                                               13.8993710691824 
                                             
                                             
                                               RAC1-mRNA 
                                               1.11 
                                               0.0959 
                                               0.925 
                                               1.3 
                                               2.16 
                                               1.9 
                                               2.46 
                                               6.93e-14 
                                               1.24e-12 
                                               lm.nb 
                                               Activation of Rac, CD28 co-stimulation, CD28 dependent Vav1 pathway, DAP12 signaling, EPH-ephrin mediated repulsion of cells, Ephrin signaling, G alpha (1213) signalling events, GPVI-mediated activation cascade, MAPK6MAPK4 signaling, NRAGE signals death through JNK, PCPCE pathway, Regulation of actin dynamics for phagocytic cup formation, RHO GTPases activate CIT, RHO GTPases Activate Formins, RHO GTPases activate IQGAPs, RHO GTPases activate PAKs, RHO GTPases activate PKNs, RHO GTPases Activate WASPs and WAVEs, Sema3A PAK dependent Axon repulsion, Sema4D induced cell migration and growth-cone collapse, Signal transduction by L1, Signaling by SCF-KIT, VEGFA-VEGFR2 Pathway, VEGFR2 mediated vascular permeability 
                                               SEBELIUS_2_C4636 
                                               RAC1 
                                               mRNA 
                                               0 
                                               
                                               
                                               MAPK6MAPK4 signaling;Signaling by SCF-KIT;Inactivation of Cdc42 and Rac;DAP12 signaling;Ephrin signaling;EPH-ephrin mediated repulsion of cells;Activation of Rac;RHO GTPases Activate Formins;VEGFR2 mediated vascular permeability;RHO GTPases activate PAKs;Sema4D mediated inhibition of cell attachment and migration;G alpha (1213) signalling events;Regulation of actin dynamics for phagocytic cup formation;Sema4D induced cell migration and growth-cone collapse;Signal transduction by L1;RHO GTPases activate CIT;RHO GTPases Activate WASPs and WAVEs;Factors involved in megakaryocyte development and platelet production;RHO GTPases activate KTN1;CD28 dependent Vav1 pathway;DSCAM interactions;NRAGE signals death through JNK;GPVI-mediated activation cascade;Sema3A PAK dependent Axon repulsion;RHO GTPases activate PKNs;SEMA3A-Plexin repulsion signaling by inhibiting Integrin adhesion;RHO GTPases activate IQGAPs;PCPCE pathway;VEGFA-VEGFR2 Pathway;CD28 co-stimulation 
                                               hsa04010;hsa04014;hsa04015;hsa04024;hsa04062;hsa04071;hsa04145;hsa04151;hsa04310;hsa04360;hsa04370;hsa04380;hsa04510;hsa04520;hsa04620;hsa04650;hsa04662;hsa04664;hsa04666;hsa04670;hsa04722;hsa04810;hsa04932;hsa04933;hsa04972;hsa05100;hsa05120;hsa05131;hsa05132;hsa05200;hsa05203;hsa05205;hsa05210;hsa05211;hsa05212;hsa05231;hsa05416 
                                               RAC1 
                                               NA 
                                               slateblue1 
                                               16 
                                               11.5745568300313 
                                             
                                             
                                               AKT2-mRNA 
                                               1.35 
                                               0.123 
                                               1.11 
                                               1.59 
                                               2.55 
                                               2.15 
                                               3.01 
                                               3.72e-13 
                                               6.37e-12 
                                               lm.nb 
                                               Activation of BAD and translocation to mitochondria , CD28 co-stimulation, DAP12 signaling, Deactivation of the beta-catenin transactivating complex, Downstream signal transduction, G beta.CL.gamma signalling through PI3Kgamma, GAB1 signalosome, GPVI-mediated activation cascade, IRS-related events triggered by IGF1R, PI3K Cascade, PI3KAKT activation, PIP3 activates AKT signaling, Role of LAT2NTALLAB on calcium mobilization, Signaling by ERBB2, Signaling by PDGF, Signaling by SCF-KIT, TP53 Regulates Metabolic Genes, VEGFA-VEGFR2 Pathway, VEGFR2 mediated vascular permeability 
                                               SEBELIUS_2_C4636 
                                               AKT2 
                                               mRNA 
                                               0 
                                               
                                               
                                               TP53 Regulates Metabolic Genes;PDE3B signalling;AKT phosphorylates targets in the cytosol;Negative regulation of the PI3KAKT network;Deactivation of the beta-catenin transactivating complex;Activation of BAD and translocation to mitochondria ;G beta.CL.gamma signalling through PI3Kgamma;AKT phosphorylates targets in the nucleus;VEGFR2 mediated vascular permeability;GPVI-mediated activation cascade;Signaling by ERBB2;PI3K Cascade;IRS-related events triggered by IGF1R;mTOR signalling;PIP3 activates AKT signaling;Downstream signal transduction;GAB1 signalosome;Signaling by SCF-KIT;PI3KAKT activation;Role of LAT2NTALLAB on calcium mobilization;DAP12 signaling;Signaling by PDGF;CD28 co-stimulation;VEGFA-VEGFR2 Pathway 
                                               hsa04010;hsa04012;hsa04014;hsa04015;hsa04022;hsa04024;hsa04062;hsa04066;hsa04068;hsa04071;hsa04072;hsa04150;hsa04151;hsa04152;hsa04210;hsa04211;hsa04213;hsa04261;hsa04370;hsa04380;hsa04510;hsa04530;hsa04550;hsa04611;hsa04620;hsa04630;hsa04660;hsa04662;hsa04664;hsa04666;hsa04668;hsa04722;hsa04725;hsa04728;hsa04910;hsa04914;hsa04915;hsa04917;hsa04919;hsa04920;hsa04922;hsa04923;hsa04931;hsa04932;hsa04933;hsa04973;hsa05142;hsa05145;hsa05152;hsa05160;hsa05161;hsa05162;hsa05164;hsa05166;hsa05169;hsa05200;hsa05205;hsa05210;hsa05211;hsa05212;hsa05213;hsa05214;hsa05215;hsa05218;hsa05220;hsa05221;hsa05222;hsa05223;hsa05230;hsa05231 
                                               AKT2 
                                               NA 
                                               slateblue1 
                                               16 
                                               10.9756097560976 
                                             
                                             
                                               CDC42-mRNA 
                                               0.458 
                                               0.1 
                                               0.262 
                                               0.654 
                                               1.37 
                                               1.2 
                                               1.57 
                                               5.18e-05 
                                               0.000494 
                                               lm.nb 
                                               CD28 co-stimulation, CD28 dependent Vav1 pathway, DCC mediated attractive signaling, EPHB-mediated forward signaling, G alpha (1213) signalling events, GPVI-mediated activation cascade, Regulation of actin dynamics for phagocytic cup formation, Rho GTPase cycle, RHO GTPases Activate Formins, RHO GTPases activate IQGAPs, RHO GTPases activate PAKs, Sema4D induced cell migration and growth-cone collapse, VEGFA-VEGFR2 Pathway 
                                               SEBELIUS_2_C4636 
                                               CDC42 
                                               mRNA 
                                               0 
                                               
                                               
                                               DCC mediated attractive signaling;Factors involved in megakaryocyte development and platelet production;VEGFA-VEGFR2 Pathway;RHO GTPases activate KTN1;CD28 dependent Vav1 pathway;Inactivation of Cdc42 and Rac;Rho GTPase cycle;EPHB-mediated forward signaling;RHO GTPases Activate Formins;RHO GTPases activate PAKs;GPVI-mediated activation cascade;G alpha (1213) signalling events;RHO GTPases activate IQGAPs;Sema4D induced cell migration and growth-cone collapse;Regulation of actin dynamics for phagocytic cup formation;CD28 co-stimulation 
                                               hsa04010;hsa04014;hsa04015;hsa04062;hsa04144;hsa04360;hsa04370;hsa04510;hsa04520;hsa04530;hsa04660;hsa04666;hsa04670;hsa04722;hsa04810;hsa04912;hsa04932;hsa04933;hsa05100;hsa05120;hsa05130;hsa05131;hsa05132;hsa05200;hsa05203;hsa05205;hsa05211;hsa05212 
                                               CDC42 
                                               NA 
                                               slateblue1 
                                               16 
                                               4.58 
                                             
                                             
                                               FYN-mRNA 
                                               -1.21 
                                               0.345 
                                               -1.89 
                                               -0.534 
                                               0.432 
                                               0.271 
                                               0.69 
                                               0.00119 
                                               0.01 
                                               lm.nb 
                                               ARMS-mediated activation, CD209 (DC-SIGN) signaling, CD28 co-stimulation, CD28 dependent PI3KAkt signaling, CD28 dependent Vav1 pathway, Cell surface interactions at the vascular wall, Constitutive Signaling by Aberrant PI3K in Cancer, CTLA4 inhibitory signaling, DAP12 signaling, Downstream signal transduction, EPH-ephrin mediated repulsion of cells, FCERI mediated MAPK activation, Frs2-mediated activation, GAB1 signalosome, Gastrin-CREB signalling pathway via PKC and MAPK, GPVI-mediated activation cascade, Interleukin receptor SHC signaling, IRS-related events triggered by IGF1R, Nephrin interactions, PI3KAKT activation, RAFMAP kinase cascade, Regulation of signaling by CBL, Role of LAT2NTALLAB on calcium mobilization, Sema3A PAK dependent Axon repulsion, Signaling by ERBB2, Signaling by PDGF, Signaling by SCF-KIT, VEGFA-VEGFR2 Pathway, VEGFR2 mediated cell proliferation 
                                               SEBELIUS_2_C4636 
                                               FYN 
                                               mRNA 
                                               0 
                                               
                                               
                                               EPH-ephrin mediated repulsion of cells;Regulation of signaling by CBL;FCGR activation;Antigen activates B Cell Receptor (BCR) leading to generation of second messengers;Dectin-2 family;RAFMAP kinase cascade;Constitutive Signaling by Aberrant PI3K in Cancer;CTLA4 inhibitory signaling;CD28 dependent PI3KAkt signaling;PECAM1 interactions;CD28 dependent Vav1 pathway;GPVI-mediated activation cascade;Sema3A PAK dependent Axon repulsion;Nephrin interactions;SEMA3A-Plexin repulsion signaling by inhibiting Integrin adhesion;CRMPs in Sema3A signaling;CD209 (DC-SIGN) signaling;Downstream signal transduction;GAB1 signalosome;Signaling by SCF-KIT;PI3KAKT activation;Role of LAT2NTALLAB on calcium mobilization;DAP12 signaling;Signaling by PDGF;Signaling by ERBB2;VEGFA-VEGFR2 Pathway;VEGFR2 mediated cell proliferation;Frs2-mediated activation;ARMS-mediated activation;Gastrin-CREB signalling pathway via PKC and MAPK;Interleukin receptor SHC signaling;FCERI mediated MAPK activation;IRS-related events triggered by IGF1R;CD28 co-stimulation;Cell surface interactions at the vascular wall 
                                               hsa04071;hsa04072;hsa04360;hsa04380;hsa04510;hsa04520;hsa04611;hsa04650;hsa04660;hsa04664;hsa04725;hsa05020;hsa05130;hsa05162;hsa05416 
                                               FYN 
                                               NA 
                                               slateblue2 
                                               16 
                                               -3.50724637681159 
                                             
                                             
                                               PIK3R2-mRNA 
                                               -0.659 
                                               0.196 
                                               -1.04 
                                               -0.275 
                                               0.633 
                                               0.485 
                                               0.826 
                                               0.00179 
                                               0.015 
                                               lm.nb 
                                               CD28 co-stimulation, CD28 dependent PI3KAkt signaling, Cell surface interactions at the vascular wall, Constitutive Signaling by Aberrant PI3K in Cancer, DAP12 signaling, Downstream signal transduction, Downstream TCR signaling, G alpha (1213) signalling events, G alpha (q) signalling events, G beta.CL.gamma signalling through PI3Kgamma, GAB1 signalosome, Gastrin-CREB signalling pathway via PKC and MAPK, GPVI-mediated activation cascade, Interleukin receptor SHC signaling, IRS-related events triggered by IGF1R, Nephrin interactions, PI3K Cascade, PI3KAKT activation, PIP3 activates AKT signaling, Regulation of signaling by CBL, Rho GTPase cycle, Role of LAT2NTALLAB on calcium mobilization, Signaling by ERBB2, Signaling by PDGF, Signaling by SCF-KIT, Synthesis of PIPs at the plasma membrane, Tie2 Signaling, VEGFA-VEGFR2 Pathway 
                                               SEBELIUS_2_C4636 
                                               PIK3R2 
                                               mRNA 
                                               0 
                                               
                                               
                                               G alpha (q) signalling events;Downstream TCR signaling;PIP3 activates AKT signaling;VEGFA-VEGFR2 Pathway;Synthesis of PIPs at the plasma membrane;Rho GTPase cycle;Interleukin receptor SHC signaling;G beta.CL.gamma signalling through PI3Kgamma;Regulation of signaling by CBL;Role of phospholipids in phagocytosis;G alpha (1213) signalling events;PI3K Cascade;Constitutive Signaling by Aberrant PI3K in Cancer;CD28 dependent PI3KAkt signaling;Tie2 Signaling;GPVI-mediated activation cascade;Nephrin interactions;Gastrin-CREB signalling pathway via PKC and MAPK;Downstream signal transduction;GAB1 signalosome;Signaling by SCF-KIT;PI3KAKT activation;Role of LAT2NTALLAB on calcium mobilization;DAP12 signaling;Signaling by PDGF;Signaling by ERBB2;IRS-related events triggered by IGF1R;CD28 co-stimulation;Cell surface interactions at the vascular wall 
                                               hsa04012;hsa04014;hsa04015;hsa04022;hsa04024;hsa04062;hsa04066;hsa04068;hsa04070;hsa04071;hsa04072;hsa04150;hsa04151;hsa04152;hsa04210;hsa04211;hsa04213;hsa04261;hsa04370;hsa04380;hsa04510;hsa04550;hsa04611;hsa04620;hsa04630;hsa04650;hsa04660;hsa04662;hsa04664;hsa04666;hsa04668;hsa04670;hsa04722;hsa04725;hsa04750;hsa04810;hsa04910;hsa04914;hsa04915;hsa04917;hsa04919;hsa04921;hsa04923;hsa04930;hsa04931;hsa04932;hsa04933;hsa04960;hsa04973;hsa05100;hsa05142;hsa05145;hsa05146;hsa05160;hsa05161;hsa05162;hsa05164;hsa05166;hsa05169;hsa05200;hsa05203;hsa05205;hsa05206;hsa05210;hsa05211;hsa05212;hsa05213;hsa05214;hsa05215;hsa05218;hsa05220;hsa05221;hsa05222;hsa05223;hsa05230;hsa05231 
                                               PIK3R2 
                                               NA 
                                               slateblue2 
                                               16 
                                               -3.36224489795918 
                                             
                                             
                                               PIK3CA-mRNA 
                                               -0.45 
                                               0.149 
                                               -0.743 
                                               -0.157 
                                               0.732 
                                               0.598 
                                               0.897 
                                               0.00468 
                                               0.0377 
                                               lm.nb 
                                               CD28 co-stimulation, CD28 dependent PI3KAkt signaling, Cell surface interactions at the vascular wall, Constitutive Signaling by Aberrant PI3K in Cancer, Constitutive Signaling by EGFRvIII, DAP12 signaling, Downstream signal transduction, Downstream TCR signaling, G alpha (1213) signalling events, G alpha (q) signalling events, G beta.CL.gamma signalling through PI3Kgamma, GAB1 signalosome, Gastrin-CREB signalling pathway via PKC and MAPK, GPVI-mediated activation cascade, Interleukin receptor SHC signaling, IRS-related events triggered by IGF1R, Nephrin interactions, PI3K Cascade, PI3KAKT activation, PIP3 activates AKT signaling, Regulation of signaling by CBL, Role of LAT2NTALLAB on calcium mobilization, Signaling by ERBB2, Signaling by FGFR1 in disease, Signaling by FGFR2 in disease, Signaling by FGFR3 in disease, Signaling by FGFR4 in disease, Signaling by PDGF, Signaling by SCF-KIT, Synthesis of PIPs at the plasma membrane, Tie2 Signaling, VEGFA-VEGFR2 Pathway 
                                               SEBELIUS_2_C4636 
                                               PIK3CA 
                                               mRNA 
                                               0 
                                               
                                               
                                               G alpha (q) signalling events;Downstream TCR signaling;Signaling by FGFR2 in disease;PIP3 activates AKT signaling;VEGFA-VEGFR2 Pathway;Synthesis of PIPs at the plasma membrane;Constitutive Signaling by Ligand-Responsive EGFR Cancer Variants;Interleukin receptor SHC signaling;G beta.CL.gamma signalling through PI3Kgamma;Regulation of signaling by CBL;Role of phospholipids in phagocytosis;G alpha (1213) signalling events;PI3K Cascade;Signaling by FGFR3 in disease;Constitutive Signaling by Aberrant PI3K in Cancer;Signaling by FGFR4 in disease;CD28 dependent PI3KAkt signaling;Tie2 Signaling;GPVI-mediated activation cascade;Nephrin interactions;Constitutive Signaling by EGFRvIII;Gastrin-CREB signalling pathway via PKC and MAPK;Downstream signal transduction;GAB1 signalosome;Signaling by SCF-KIT;PI3KAKT activation;Role of LAT2NTALLAB on calcium mobilization;DAP12 signaling;Signaling by PDGF;Signaling by ERBB2;IRS-related events triggered by IGF1R;CD28 co-stimulation;Cell surface interactions at the vascular wall;Signaling by FGFR1 in disease 
                                               hsa00562;hsa04012;hsa04014;hsa04015;hsa04022;hsa04024;hsa04062;hsa04066;hsa04068;hsa04070;hsa04071;hsa04072;hsa04150;hsa04151;hsa04152;hsa04210;hsa04211;hsa04213;hsa04261;hsa04370;hsa04380;hsa04510;hsa04550;hsa04611;hsa04620;hsa04630;hsa04650;hsa04660;hsa04662;hsa04664;hsa04666;hsa04668;hsa04670;hsa04722;hsa04725;hsa04750;hsa04810;hsa04910;hsa04914;hsa04915;hsa04917;hsa04919;hsa04921;hsa04923;hsa04930;hsa04931;hsa04932;hsa04933;hsa04960;hsa04973;hsa05100;hsa05142;hsa05145;hsa05146;hsa05160;hsa05161;hsa05162;hsa05164;hsa05166;hsa05169;hsa05200;hsa05203;hsa05205;hsa05206;hsa05210;hsa05211;hsa05212;hsa05213;hsa05214;hsa05215;hsa05218;hsa05220;hsa05221;hsa05222;hsa05223;hsa05230;hsa05231 
                                               PIK3CA 
                                               NA 
                                               slateblue2 
                                               16 
                                               -3.02013422818792 
                                             
                                             
                                               PAK2-mRNA 
                                               0.221 
                                               0.0777 
                                               0.0684 
                                               0.373 
                                               1.17 
                                               1.05 
                                               1.3 
                                               0.00728 
                                               0.0571 
                                               loglinear 
                                               Activation of Rac, CD209 (DC-SIGN) signaling, CD28 co-stimulation, CD28 dependent Vav1 pathway, Ephrin signaling, RHO GTPases activate PAKs, Sema3A PAK dependent Axon repulsion, VEGFA-VEGFR2 Pathway, VEGFR2 mediated vascular permeability 
                                               SEBELIUS_2_C4636 
                                               PAK2 
                                               mRNA 
                                               0 
                                               
                                               
                                               Nef and signal transduction;CD28 dependent Vav1 pathway;Stimulation of the cell death response by PAK-2p34;Ephrin signaling;Activation of Rac;RHO GTPases activate PAKs;VEGFR2 mediated vascular permeability;Sema3A PAK dependent Axon repulsion;CD209 (DC-SIGN) signaling;VEGFA-VEGFR2 Pathway;CD28 co-stimulation 
                                               hsa04010;hsa04012;hsa04014;hsa04360;hsa04510;hsa04660;hsa04810;hsa05211 
                                               PAK2 
                                               NA 
                                               slateblue3 
                                               16 
                                               2.84427284427284 
                                             
                                             
                                               PIK3R3-mRNA 
                                               0.688 
                                               0.429 
                                               -0.152 
                                               1.53 
                                               1.61 
                                               0.9 
                                               2.88 
                                               0.117 
                                               0.829 
                                               lm.nb 
                                               CD28 co-stimulation, CD28 dependent PI3KAkt signaling, Constitutive Signaling by Aberrant PI3K in Cancer, DAP12 signaling, Downstream signal transduction, G alpha (1213) signalling events, G alpha (q) signalling events, G beta.CL.gamma signalling through PI3Kgamma, GAB1 signalosome, Gastrin-CREB signalling pathway via PKC and MAPK, GPVI-mediated activation cascade, Interleukin receptor SHC signaling, PI3KAKT activation, PIP3 activates AKT signaling, Regulation of signaling by CBL, Role of LAT2NTALLAB on calcium mobilization, Signaling by ERBB2, Signaling by PDGF, Signaling by SCF-KIT, Synthesis of PIPs at the plasma membrane 
                                               SEBELIUS_2_C4636 
                                               PIK3R3 
                                               mRNA 
                                               0 
                                               
                                               
                                               G alpha (q) signalling events;PIP3 activates AKT signaling;Constitutive Signaling by Aberrant PI3K in Cancer;CD28 dependent PI3KAkt signaling;Synthesis of PIPs at the plasma membrane;Interleukin receptor SHC signaling;G beta.CL.gamma signalling through PI3Kgamma;Regulation of signaling by CBL;GPVI-mediated activation cascade;G alpha (1213) signalling events;Gastrin-CREB signalling pathway via PKC and MAPK;Downstream signal transduction;GAB1 signalosome;Signaling by SCF-KIT;PI3KAKT activation;Role of LAT2NTALLAB on calcium mobilization;DAP12 signaling;Signaling by PDGF;Signaling by ERBB2;CD28 co-stimulation 
                                               hsa04012;hsa04014;hsa04015;hsa04022;hsa04024;hsa04062;hsa04066;hsa04068;hsa04070;hsa04071;hsa04072;hsa04150;hsa04151;hsa04152;hsa04210;hsa04211;hsa04213;hsa04261;hsa04370;hsa04380;hsa04510;hsa04550;hsa04611;hsa04620;hsa04630;hsa04650;hsa04660;hsa04662;hsa04664;hsa04666;hsa04668;hsa04670;hsa04722;hsa04725;hsa04750;hsa04810;hsa04910;hsa04914;hsa04915;hsa04917;hsa04919;hsa04921;hsa04923;hsa04930;hsa04931;hsa04932;hsa04933;hsa04960;hsa04973;hsa05100;hsa05142;hsa05145;hsa05146;hsa05160;hsa05161;hsa05162;hsa05164;hsa05166;hsa05169;hsa05200;hsa05203;hsa05205;hsa05210;hsa05211;hsa05212;hsa05213;hsa05214;hsa05215;hsa05218;hsa05220;hsa05221;hsa05222;hsa05223;hsa05230;hsa05231 
                                               PIK3R3 
                                               NA 
                                               azure4 
                                               1 
                                               1.6037296037296 
                                             
                                             
                                               AKT1-mRNA 
                                               0.00517 
                                               0.241 
                                               -0.468 
                                               0.478 
                                               1 
                                               0.723 
                                               1.39 
                                               0.983 
                                               1 
                                               lm.nb 
                                               Activation of BAD and translocation to mitochondria , CD28 co-stimulation, DAP12 signaling, Deactivation of the beta-catenin transactivating complex, Downstream signal transduction, G beta.CL.gamma signalling through PI3Kgamma, GAB1 signalosome, GPVI-mediated activation cascade, IRS-related events triggered by IGF1R, PI3K Cascade, PI3KAKT activation, PIP3 activates AKT signaling, Role of LAT2NTALLAB on calcium mobilization, Signaling by ERBB2, Signaling by PDGF, Signaling by SCF-KIT, TP53 Regulates Metabolic Genes, VEGFA-VEGFR2 Pathway, VEGFR2 mediated vascular permeability 
                                               SEBELIUS_2_C4636 
                                               AKT1 
                                               mRNA 
                                               0 
                                               
                                               
                                               TP53 Regulates Metabolic Genes;Deactivation of the beta-catenin transactivating complex;Activation of BAD and translocation to mitochondria ;eNOS activation;mTOR signalling;G beta.CL.gamma signalling through PI3Kgamma;KSRP (KHSRP) binds and destabilizes mRNA;AKT phosphorylates targets in the nucleus;VEGFR2 mediated vascular permeability;GPVI-mediated activation cascade;Tetrahydrobiopterin (BH4) synthesis# recycling# salvage and regulation;Butyrate Response Factor 1 (BRF1) binds and destabilizes mRNA;Signaling by ERBB2;PIP3 activates AKT signaling;Downstream signal transduction;GAB1 signalosome;Signaling by SCF-KIT;PI3KAKT activation;Role of LAT2NTALLAB on calcium mobilization;DAP12 signaling;Signaling by PDGF;CD28 co-stimulation;PI3K Cascade;IRS-related events triggered by IGF1R;VEGFA-VEGFR2 Pathway 
                                               hsa04010;hsa04012;hsa04014;hsa04015;hsa04022;hsa04024;hsa04062;hsa04066;hsa04068;hsa04071;hsa04072;hsa04150;hsa04151;hsa04152;hsa04210;hsa04211;hsa04213;hsa04261;hsa04370;hsa04380;hsa04510;hsa04530;hsa04550;hsa04611;hsa04620;hsa04630;hsa04660;hsa04662;hsa04664;hsa04666;hsa04668;hsa04722;hsa04725;hsa04728;hsa04910;hsa04914;hsa04915;hsa04917;hsa04919;hsa04920;hsa04922;hsa04923;hsa04931;hsa04932;hsa04933;hsa04973;hsa05142;hsa05145;hsa05152;hsa05160;hsa05161;hsa05162;hsa05164;hsa05166;hsa05169;hsa05200;hsa05205;hsa05210;hsa05211;hsa05212;hsa05213;hsa05214;hsa05215;hsa05218;hsa05220;hsa05221;hsa05222;hsa05223;hsa05230;hsa05231 
                                               AKT1 
                                               NA 
                                               azure4 
                                               1 
                                               0.0214522821576764 
                                             
                                           
                                           DE Results - CD28 co-stimulation genes - Line: HT29 vs.Fibro  Table displaying each sample's global significance scores and directed global significance scores as defined in the heatmaps above. The global significance score is calculated as the square root of the mean squared t-statistic for the genes in a gene set, with t-statistics coming from the linear regression underlying our differential expression analysis. The directed global significance score is calculated as the square root of the mean signed squared t-statistic for the genes in a gene set, with t-statistics coming from the linear regression underlying our differential expression analysis.   
                                      
                                       
                                  
                                
                              
                            
                           
                            
                          
                           
                      
                    
                    
                      
                           
                          
                               Day    Line     
                              
                                  Day  
                              
                               
                              
                                  Line: differential expression in Caco2 vs. baseline of Fibro  
                                  Line: differential expression in HEPG2 vs. baseline of Fibro  
                                  Line: differential expression in HT29 vs. baseline of Fibro  
                              
                              
                            
                              
                                
                                  
                                       
                                      
                                           Volcano Plot: Day   More Plot Information                Volcano Plot: Day  Volcano plot displaying each gene's -log10(p-value) and log2 fold change for the selected covariate. Highly statistically significant genes fall at the top of the plot, and highly differentially expressed genes fall to either side. Genes within the selected gene set are highlighted in orange. Horizontal lines indicate various False Discovery Rate (FDR) thresholds.   
                                           DE Results - CD28 dependent PI3KAkt signaling genes - Day   More Plot Information      Download CSV Data     
                                           
                                             
                                             
                                           
                                           
                                             
                                               Probe Label 
                                               Log2 fold change 
                                               std error (log2) 
                                               Lower confidence limit (log2) 
                                               Upper confidence limit (log2) 
                                               Linear fold change 
                                               Lower confidence limit (linear) 
                                               Upper confidence limit (linear) 
                                               P-value 
                                               BY.p.value 
                                               method 
                                               Gene.sets 
                                               Codeset.Name 
                                               Probe.Label 
                                               Analyte.Type 
                                               Is.Control 
                                               Control.Type 
                                               Related.Probes 
                                               Probe.Annotation 
                                               KEGG.Pathways 
                                               Official.Gene.Name 
                                               Control.Conc 
                                               volcanocol 
                                               volcanopch 
                                               tstats.all 
                                             
                                           
                                           
                                             
                                               PIK3R1-mRNA 
                                               0.0895 
                                               0.021 
                                               0.0484 
                                               0.131 
                                               1.06 
                                               1.03 
                                               1.09 
                                               0.000131 
                                               0.00452 
                                               lm.nb 
                                               CD28 co-stimulation, CD28 dependent PI3KAkt signaling, Cell surface interactions at the vascular wall, Constitutive Signaling by Aberrant PI3K in Cancer, Constitutive Signaling by EGFRvIII, DAP12 signaling, Downstream signal transduction, Downstream TCR signaling, G alpha (q) signalling events, G beta.CL.gamma signalling through PI3Kgamma, GAB1 signalosome, Gastrin-CREB signalling pathway via PKC and MAPK, GPVI-mediated activation cascade, Interleukin receptor SHC signaling, IRS-related events triggered by IGF1R, Nephrin interactions, PI3KAKT activation, PIP3 activates AKT signaling, Regulation of signaling by CBL, Role of LAT2NTALLAB on calcium mobilization, Signaling by ERBB2, Signaling by FGFR1 in disease, Signaling by FGFR2 in disease, Signaling by FGFR3 in disease, Signaling by FGFR4 in disease, Signaling by PDGF, Signaling by SCF-KIT, Synthesis of PIPs at the plasma membrane, Tie2 Signaling, VEGFA-VEGFR2 Pathway 
                                               SEBELIUS_2_C4636 
                                               PIK3R1 
                                               mRNA 
                                               0 
                                               
                                               
                                               G alpha (q) signalling events;Downstream TCR signaling;Signaling by FGFR2 in disease;PIP3 activates AKT signaling;VEGFA-VEGFR2 Pathway;Synthesis of PIPs at the plasma membrane;Constitutive Signaling by Ligand-Responsive EGFR Cancer Variants;Interleukin receptor SHC signaling;G beta.CL.gamma signalling through PI3Kgamma;Regulation of signaling by CBL;Role of phospholipids in phagocytosis;Signaling by FGFR3 in disease;Constitutive Signaling by Aberrant PI3K in Cancer;Signaling by FGFR4 in disease;CD28 dependent PI3KAkt signaling;Tie2 Signaling;Signaling by FGFR1 fusion mutants;GP1b-IX-V activation signalling;GPVI-mediated activation cascade;Nephrin interactions;Constitutive Signaling by EGFRvIII;Gastrin-CREB signalling pathway via PKC and MAPK;Downstream signal transduction;GAB1 signalosome;Signaling by SCF-KIT;PI3KAKT activation;Role of LAT2NTALLAB on calcium mobilization;DAP12 signaling;Signaling by PDGF;Signaling by ERBB2;IRS-related events triggered by IGF1R;CD28 co-stimulation;Cell surface interactions at the vascular wall;Signaling by FGFR1 in disease 
                                               hsa04012;hsa04014;hsa04015;hsa04022;hsa04024;hsa04062;hsa04066;hsa04068;hsa04070;hsa04071;hsa04072;hsa04150;hsa04151;hsa04152;hsa04210;hsa04211;hsa04213;hsa04261;hsa04370;hsa04380;hsa04510;hsa04550;hsa04611;hsa04620;hsa04630;hsa04650;hsa04660;hsa04662;hsa04664;hsa04666;hsa04668;hsa04670;hsa04722;hsa04725;hsa04750;hsa04810;hsa04910;hsa04914;hsa04915;hsa04917;hsa04919;hsa04921;hsa04923;hsa04930;hsa04931;hsa04932;hsa04933;hsa04960;hsa04973;hsa05100;hsa05142;hsa05145;hsa05146;hsa05160;hsa05161;hsa05162;hsa05164;hsa05166;hsa05169;hsa05200;hsa05203;hsa05205;hsa05210;hsa05211;hsa05212;hsa05213;hsa05214;hsa05215;hsa05218;hsa05220;hsa05221;hsa05222;hsa05223;hsa05230;hsa05231 
                                               PIK3R1 
                                               NA 
                                               slateblue1 
                                               16 
                                               4.26190476190476 
                                             
                                             
                                               AKT3-mRNA 
                                               0.0985 
                                               0.0254 
                                               0.0488 
                                               0.148 
                                               1.07 
                                               1.03 
                                               1.11 
                                               0.000409 
                                               0.0111 
                                               lm.nb 
                                               Activation of BAD and translocation to mitochondria , CD28 co-stimulation, CD28 dependent PI3KAkt signaling, DAP12 signaling, Downstream signal transduction, G beta.CL.gamma signalling through PI3Kgamma, GAB1 signalosome, GPVI-mediated activation cascade, PI3KAKT activation, PIP3 activates AKT signaling, Role of LAT2NTALLAB on calcium mobilization, Signaling by ERBB2, Signaling by PDGF, Signaling by SCF-KIT, TP53 Regulates Metabolic Genes, VEGFA-VEGFR2 Pathway, VEGFR2 mediated vascular permeability 
                                               SEBELIUS_2_C4636 
                                               AKT3 
                                               mRNA 
                                               0 
                                               
                                               
                                               TP53 Regulates Metabolic Genes;CD28 dependent PI3KAkt signaling;AKT phosphorylates targets in the cytosol;Negative regulation of the PI3KAKT network;Activation of BAD and translocation to mitochondria ;G beta.CL.gamma signalling through PI3Kgamma;AKT phosphorylates targets in the nucleus;VEGFR2 mediated vascular permeability;GPVI-mediated activation cascade;Signaling by ERBB2;PIP3 activates AKT signaling;Downstream signal transduction;GAB1 signalosome;Signaling by SCF-KIT;PI3KAKT activation;Role of LAT2NTALLAB on calcium mobilization;DAP12 signaling;Signaling by PDGF;CD28 co-stimulation;VEGFA-VEGFR2 Pathway 
                                               hsa04010;hsa04012;hsa04014;hsa04015;hsa04022;hsa04024;hsa04062;hsa04066;hsa04068;hsa04071;hsa04072;hsa04150;hsa04151;hsa04152;hsa04210;hsa04211;hsa04213;hsa04261;hsa04370;hsa04380;hsa04510;hsa04530;hsa04550;hsa04611;hsa04620;hsa04630;hsa04660;hsa04662;hsa04664;hsa04666;hsa04668;hsa04722;hsa04725;hsa04728;hsa04910;hsa04914;hsa04915;hsa04917;hsa04919;hsa04920;hsa04922;hsa04923;hsa04931;hsa04932;hsa04933;hsa04973;hsa05142;hsa05145;hsa05152;hsa05160;hsa05161;hsa05162;hsa05164;hsa05166;hsa05169;hsa05200;hsa05205;hsa05210;hsa05211;hsa05212;hsa05213;hsa05214;hsa05215;hsa05218;hsa05220;hsa05221;hsa05222;hsa05223;hsa05230;hsa05231 
                                               AKT3 
                                               NA 
                                               slateblue2 
                                               16 
                                               3.87795275590551 
                                             
                                             
                                               PIK3R2-mRNA 
                                               0.0724 
                                               0.0251 
                                               0.0232 
                                               0.122 
                                               1.05 
                                               1.02 
                                               1.09 
                                               0.00651 
                                               0.0963 
                                               lm.nb 
                                               CD28 co-stimulation, CD28 dependent PI3KAkt signaling, Cell surface interactions at the vascular wall, Constitutive Signaling by Aberrant PI3K in Cancer, DAP12 signaling, Downstream signal transduction, Downstream TCR signaling, G alpha (1213) signalling events, G alpha (q) signalling events, G beta.CL.gamma signalling through PI3Kgamma, GAB1 signalosome, Gastrin-CREB signalling pathway via PKC and MAPK, GPVI-mediated activation cascade, Interleukin receptor SHC signaling, IRS-related events triggered by IGF1R, Nephrin interactions, PI3K Cascade, PI3KAKT activation, PIP3 activates AKT signaling, Regulation of signaling by CBL, Rho GTPase cycle, Role of LAT2NTALLAB on calcium mobilization, Signaling by ERBB2, Signaling by PDGF, Signaling by SCF-KIT, Synthesis of PIPs at the plasma membrane, Tie2 Signaling, VEGFA-VEGFR2 Pathway 
                                               SEBELIUS_2_C4636 
                                               PIK3R2 
                                               mRNA 
                                               0 
                                               
                                               
                                               G alpha (q) signalling events;Downstream TCR signaling;PIP3 activates AKT signaling;VEGFA-VEGFR2 Pathway;Synthesis of PIPs at the plasma membrane;Rho GTPase cycle;Interleukin receptor SHC signaling;G beta.CL.gamma signalling through PI3Kgamma;Regulation of signaling by CBL;Role of phospholipids in phagocytosis;G alpha (1213) signalling events;PI3K Cascade;Constitutive Signaling by Aberrant PI3K in Cancer;CD28 dependent PI3KAkt signaling;Tie2 Signaling;GPVI-mediated activation cascade;Nephrin interactions;Gastrin-CREB signalling pathway via PKC and MAPK;Downstream signal transduction;GAB1 signalosome;Signaling by SCF-KIT;PI3KAKT activation;Role of LAT2NTALLAB on calcium mobilization;DAP12 signaling;Signaling by PDGF;Signaling by ERBB2;IRS-related events triggered by IGF1R;CD28 co-stimulation;Cell surface interactions at the vascular wall 
                                               hsa04012;hsa04014;hsa04015;hsa04022;hsa04024;hsa04062;hsa04066;hsa04068;hsa04070;hsa04071;hsa04072;hsa04150;hsa04151;hsa04152;hsa04210;hsa04211;hsa04213;hsa04261;hsa04370;hsa04380;hsa04510;hsa04550;hsa04611;hsa04620;hsa04630;hsa04650;hsa04660;hsa04662;hsa04664;hsa04666;hsa04668;hsa04670;hsa04722;hsa04725;hsa04750;hsa04810;hsa04910;hsa04914;hsa04915;hsa04917;hsa04919;hsa04921;hsa04923;hsa04930;hsa04931;hsa04932;hsa04933;hsa04960;hsa04973;hsa05100;hsa05142;hsa05145;hsa05146;hsa05160;hsa05161;hsa05162;hsa05164;hsa05166;hsa05169;hsa05200;hsa05203;hsa05205;hsa05206;hsa05210;hsa05211;hsa05212;hsa05213;hsa05214;hsa05215;hsa05218;hsa05220;hsa05221;hsa05222;hsa05223;hsa05230;hsa05231 
                                               PIK3R2 
                                               NA 
                                               slateblue3 
                                               16 
                                               2.88446215139442 
                                             
                                             
                                               PIK3R3-mRNA 
                                               0.109 
                                               0.0563 
                                               -0.00124 
                                               0.22 
                                               1.08 
                                               0.999 
                                               1.16 
                                               0.0603 
                                               0.62 
                                               lm.nb 
                                               CD28 co-stimulation, CD28 dependent PI3KAkt signaling, Constitutive Signaling by Aberrant PI3K in Cancer, DAP12 signaling, Downstream signal transduction, G alpha (1213) signalling events, G alpha (q) signalling events, G beta.CL.gamma signalling through PI3Kgamma, GAB1 signalosome, Gastrin-CREB signalling pathway via PKC and MAPK, GPVI-mediated activation cascade, Interleukin receptor SHC signaling, PI3KAKT activation, PIP3 activates AKT signaling, Regulation of signaling by CBL, Role of LAT2NTALLAB on calcium mobilization, Signaling by ERBB2, Signaling by PDGF, Signaling by SCF-KIT, Synthesis of PIPs at the plasma membrane 
                                               SEBELIUS_2_C4636 
                                               PIK3R3 
                                               mRNA 
                                               0 
                                               
                                               
                                               G alpha (q) signalling events;PIP3 activates AKT signaling;Constitutive Signaling by Aberrant PI3K in Cancer;CD28 dependent PI3KAkt signaling;Synthesis of PIPs at the plasma membrane;Interleukin receptor SHC signaling;G beta.CL.gamma signalling through PI3Kgamma;Regulation of signaling by CBL;GPVI-mediated activation cascade;G alpha (1213) signalling events;Gastrin-CREB signalling pathway via PKC and MAPK;Downstream signal transduction;GAB1 signalosome;Signaling by SCF-KIT;PI3KAKT activation;Role of LAT2NTALLAB on calcium mobilization;DAP12 signaling;Signaling by PDGF;Signaling by ERBB2;CD28 co-stimulation 
                                               hsa04012;hsa04014;hsa04015;hsa04022;hsa04024;hsa04062;hsa04066;hsa04068;hsa04070;hsa04071;hsa04072;hsa04150;hsa04151;hsa04152;hsa04210;hsa04211;hsa04213;hsa04261;hsa04370;hsa04380;hsa04510;hsa04550;hsa04611;hsa04620;hsa04630;hsa04650;hsa04660;hsa04662;hsa04664;hsa04666;hsa04668;hsa04670;hsa04722;hsa04725;hsa04750;hsa04810;hsa04910;hsa04914;hsa04915;hsa04917;hsa04919;hsa04921;hsa04923;hsa04930;hsa04931;hsa04932;hsa04933;hsa04960;hsa04973;hsa05100;hsa05142;hsa05145;hsa05146;hsa05160;hsa05161;hsa05162;hsa05164;hsa05166;hsa05169;hsa05200;hsa05203;hsa05205;hsa05210;hsa05211;hsa05212;hsa05213;hsa05214;hsa05215;hsa05218;hsa05220;hsa05221;hsa05222;hsa05223;hsa05230;hsa05231 
                                               PIK3R3 
                                               NA 
                                               azure4 
                                               1 
                                               1.9360568383659 
                                             
                                             
                                               PIK3CA-mRNA 
                                               0.0374 
                                               0.0195 
                                               -0.000784 
                                               0.0756 
                                               1.03 
                                               0.999 
                                               1.05 
                                               0.0626 
                                               0.639 
                                               lm.nb 
                                               CD28 co-stimulation, CD28 dependent PI3KAkt signaling, Cell surface interactions at the vascular wall, Constitutive Signaling by Aberrant PI3K in Cancer, Constitutive Signaling by EGFRvIII, DAP12 signaling, Downstream signal transduction, Downstream TCR signaling, G alpha (1213) signalling events, G alpha (q) signalling events, G beta.CL.gamma signalling through PI3Kgamma, GAB1 signalosome, Gastrin-CREB signalling pathway via PKC and MAPK, GPVI-mediated activation cascade, Interleukin receptor SHC signaling, IRS-related events triggered by IGF1R, Nephrin interactions, PI3K Cascade, PI3KAKT activation, PIP3 activates AKT signaling, Regulation of signaling by CBL, Role of LAT2NTALLAB on calcium mobilization, Signaling by ERBB2, Signaling by FGFR1 in disease, Signaling by FGFR2 in disease, Signaling by FGFR3 in disease, Signaling by FGFR4 in disease, Signaling by PDGF, Signaling by SCF-KIT, Synthesis of PIPs at the plasma membrane, Tie2 Signaling, VEGFA-VEGFR2 Pathway 
                                               SEBELIUS_2_C4636 
                                               PIK3CA 
                                               mRNA 
                                               0 
                                               
                                               
                                               G alpha (q) signalling events;Downstream TCR signaling;Signaling by FGFR2 in disease;PIP3 activates AKT signaling;VEGFA-VEGFR2 Pathway;Synthesis of PIPs at the plasma membrane;Constitutive Signaling by Ligand-Responsive EGFR Cancer Variants;Interleukin receptor SHC signaling;G beta.CL.gamma signalling through PI3Kgamma;Regulation of signaling by CBL;Role of phospholipids in phagocytosis;G alpha (1213) signalling events;PI3K Cascade;Signaling by FGFR3 in disease;Constitutive Signaling by Aberrant PI3K in Cancer;Signaling by FGFR4 in disease;CD28 dependent PI3KAkt signaling;Tie2 Signaling;GPVI-mediated activation cascade;Nephrin interactions;Constitutive Signaling by EGFRvIII;Gastrin-CREB signalling pathway via PKC and MAPK;Downstream signal transduction;GAB1 signalosome;Signaling by SCF-KIT;PI3KAKT activation;Role of LAT2NTALLAB on calcium mobilization;DAP12 signaling;Signaling by PDGF;Signaling by ERBB2;IRS-related events triggered by IGF1R;CD28 co-stimulation;Cell surface interactions at the vascular wall;Signaling by FGFR1 in disease 
                                               hsa00562;hsa04012;hsa04014;hsa04015;hsa04022;hsa04024;hsa04062;hsa04066;hsa04068;hsa04070;hsa04071;hsa04072;hsa04150;hsa04151;hsa04152;hsa04210;hsa04211;hsa04213;hsa04261;hsa04370;hsa04380;hsa04510;hsa04550;hsa04611;hsa04620;hsa04630;hsa04650;hsa04660;hsa04662;hsa04664;hsa04666;hsa04668;hsa04670;hsa04722;hsa04725;hsa04750;hsa04810;hsa04910;hsa04914;hsa04915;hsa04917;hsa04919;hsa04921;hsa04923;hsa04930;hsa04931;hsa04932;hsa04933;hsa04960;hsa04973;hsa05100;hsa05142;hsa05145;hsa05146;hsa05160;hsa05161;hsa05162;hsa05164;hsa05166;hsa05169;hsa05200;hsa05203;hsa05205;hsa05206;hsa05210;hsa05211;hsa05212;hsa05213;hsa05214;hsa05215;hsa05218;hsa05220;hsa05221;hsa05222;hsa05223;hsa05230;hsa05231 
                                               PIK3CA 
                                               NA 
                                               azure4 
                                               1 
                                               1.91794871794872 
                                             
                                             
                                               FYN-mRNA 
                                               0.0103 
                                               0.0454 
                                               -0.0786 
                                               0.0993 
                                               1.01 
                                               0.947 
                                               1.07 
                                               0.821 
                                               1 
                                               lm.nb 
                                               ARMS-mediated activation, CD209 (DC-SIGN) signaling, CD28 co-stimulation, CD28 dependent PI3KAkt signaling, CD28 dependent Vav1 pathway, Cell surface interactions at the vascular wall, Constitutive Signaling by Aberrant PI3K in Cancer, CTLA4 inhibitory signaling, DAP12 signaling, Downstream signal transduction, EPH-ephrin mediated repulsion of cells, FCERI mediated MAPK activation, Frs2-mediated activation, GAB1 signalosome, Gastrin-CREB signalling pathway via PKC and MAPK, GPVI-mediated activation cascade, Interleukin receptor SHC signaling, IRS-related events triggered by IGF1R, Nephrin interactions, PI3KAKT activation, RAFMAP kinase cascade, Regulation of signaling by CBL, Role of LAT2NTALLAB on calcium mobilization, Sema3A PAK dependent Axon repulsion, Signaling by ERBB2, Signaling by PDGF, Signaling by SCF-KIT, VEGFA-VEGFR2 Pathway, VEGFR2 mediated cell proliferation 
                                               SEBELIUS_2_C4636 
                                               FYN 
                                               mRNA 
                                               0 
                                               
                                               
                                               EPH-ephrin mediated repulsion of cells;Regulation of signaling by CBL;FCGR activation;Antigen activates B Cell Receptor (BCR) leading to generation of second messengers;Dectin-2 family;RAFMAP kinase cascade;Constitutive Signaling by Aberrant PI3K in Cancer;CTLA4 inhibitory signaling;CD28 dependent PI3KAkt signaling;PECAM1 interactions;CD28 dependent Vav1 pathway;GPVI-mediated activation cascade;Sema3A PAK dependent Axon repulsion;Nephrin interactions;SEMA3A-Plexin repulsion signaling by inhibiting Integrin adhesion;CRMPs in Sema3A signaling;CD209 (DC-SIGN) signaling;Downstream signal transduction;GAB1 signalosome;Signaling by SCF-KIT;PI3KAKT activation;Role of LAT2NTALLAB on calcium mobilization;DAP12 signaling;Signaling by PDGF;Signaling by ERBB2;VEGFA-VEGFR2 Pathway;VEGFR2 mediated cell proliferation;Frs2-mediated activation;ARMS-mediated activation;Gastrin-CREB signalling pathway via PKC and MAPK;Interleukin receptor SHC signaling;FCERI mediated MAPK activation;IRS-related events triggered by IGF1R;CD28 co-stimulation;Cell surface interactions at the vascular wall 
                                               hsa04071;hsa04072;hsa04360;hsa04380;hsa04510;hsa04520;hsa04611;hsa04650;hsa04660;hsa04664;hsa04725;hsa05020;hsa05130;hsa05162;hsa05416 
                                               FYN 
                                               NA 
                                               azure4 
                                               1 
                                               0.226872246696035 
                                             
                                           
                                           DE Results - CD28 dependent PI3KAkt signaling genes - Day  Table displaying each sample's global significance scores and directed global significance scores as defined in the heatmaps above. The global significance score is calculated as the square root of the mean squared t-statistic for the genes in a gene set, with t-statistics coming from the linear regression underlying our differential expression analysis. The directed global significance score is calculated as the square root of the mean signed squared t-statistic for the genes in a gene set, with t-statistics coming from the linear regression underlying our differential expression analysis.   
                                      
                                       
                                  
                                
                              
                            
                            
                              
                                
                                  
                                       
                                      
                                           Volcano Plot: Line: Caco2 vs.Fibro   More Plot Information                Volcano Plot: Line: Caco2 vs.Fibro  Volcano plot displaying each gene's -log10(p-value) and log2 fold change for the selected covariate. Highly statistically significant genes fall at the top of the plot, and highly differentially expressed genes fall to either side. Genes within the selected gene set are highlighted in orange. Horizontal lines indicate various False Discovery Rate (FDR) thresholds.   
                                           DE Results - CD28 dependent PI3KAkt signaling genes - Line: Caco2 vs.Fibro   More Plot Information      Download CSV Data     
                                           
                                             
                                             
                                           
                                           
                                             
                                               Probe Label 
                                               Log2 fold change 
                                               std error (log2) 
                                               Lower confidence limit (log2) 
                                               Upper confidence limit (log2) 
                                               Linear fold change 
                                               Lower confidence limit (linear) 
                                               Upper confidence limit (linear) 
                                               P-value 
                                               BY.p.value 
                                               method 
                                               Gene.sets 
                                               Codeset.Name 
                                               Probe.Label 
                                               Analyte.Type 
                                               Is.Control 
                                               Control.Type 
                                               Related.Probes 
                                               Probe.Annotation 
                                               KEGG.Pathways 
                                               Official.Gene.Name 
                                               Control.Conc 
                                               volcanocol 
                                               volcanopch 
                                               tstats.all 
                                             
                                           
                                           
                                             
                                               AKT3-mRNA 
                                               -2.88 
                                               0.162 
                                               -3.2 
                                               -2.57 
                                               0.135 
                                               0.109 
                                               0.169 
                                               9.07e-20 
                                               6.13e-18 
                                               lm.nb 
                                               Activation of BAD and translocation to mitochondria , CD28 co-stimulation, CD28 dependent PI3KAkt signaling, DAP12 signaling, Downstream signal transduction, G beta.CL.gamma signalling through PI3Kgamma, GAB1 signalosome, GPVI-mediated activation cascade, PI3KAKT activation, PIP3 activates AKT signaling, Role of LAT2NTALLAB on calcium mobilization, Signaling by ERBB2, Signaling by PDGF, Signaling by SCF-KIT, TP53 Regulates Metabolic Genes, VEGFA-VEGFR2 Pathway, VEGFR2 mediated vascular permeability 
                                               SEBELIUS_2_C4636 
                                               AKT3 
                                               mRNA 
                                               0 
                                               
                                               
                                               TP53 Regulates Metabolic Genes;CD28 dependent PI3KAkt signaling;AKT phosphorylates targets in the cytosol;Negative regulation of the PI3KAKT network;Activation of BAD and translocation to mitochondria ;G beta.CL.gamma signalling through PI3Kgamma;AKT phosphorylates targets in the nucleus;VEGFR2 mediated vascular permeability;GPVI-mediated activation cascade;Signaling by ERBB2;PIP3 activates AKT signaling;Downstream signal transduction;GAB1 signalosome;Signaling by SCF-KIT;PI3KAKT activation;Role of LAT2NTALLAB on calcium mobilization;DAP12 signaling;Signaling by PDGF;CD28 co-stimulation;VEGFA-VEGFR2 Pathway 
                                               hsa04010;hsa04012;hsa04014;hsa04015;hsa04022;hsa04024;hsa04062;hsa04066;hsa04068;hsa04071;hsa04072;hsa04150;hsa04151;hsa04152;hsa04210;hsa04211;hsa04213;hsa04261;hsa04370;hsa04380;hsa04510;hsa04530;hsa04550;hsa04611;hsa04620;hsa04630;hsa04660;hsa04662;hsa04664;hsa04666;hsa04668;hsa04722;hsa04725;hsa04728;hsa04910;hsa04914;hsa04915;hsa04917;hsa04919;hsa04920;hsa04922;hsa04923;hsa04931;hsa04932;hsa04933;hsa04973;hsa05142;hsa05145;hsa05152;hsa05160;hsa05161;hsa05162;hsa05164;hsa05166;hsa05169;hsa05200;hsa05205;hsa05210;hsa05211;hsa05212;hsa05213;hsa05214;hsa05215;hsa05218;hsa05220;hsa05221;hsa05222;hsa05223;hsa05230;hsa05231 
                                               AKT3 
                                               NA 
                                               slateblue1 
                                               16 
                                               -17.7777777777778 
                                             
                                             
                                               PIK3R1-mRNA 
                                               1.99 
                                               0.164 
                                               1.67 
                                               2.31 
                                               3.97 
                                               3.18 
                                               4.96 
                                               1.85e-14 
                                               4.23e-13 
                                               lm.nb 
                                               CD28 co-stimulation, CD28 dependent PI3KAkt signaling, Cell surface interactions at the vascular wall, Constitutive Signaling by Aberrant PI3K in Cancer, Constitutive Signaling by EGFRvIII, DAP12 signaling, Downstream signal transduction, Downstream TCR signaling, G alpha (q) signalling events, G beta.CL.gamma signalling through PI3Kgamma, GAB1 signalosome, Gastrin-CREB signalling pathway via PKC and MAPK, GPVI-mediated activation cascade, Interleukin receptor SHC signaling, IRS-related events triggered by IGF1R, Nephrin interactions, PI3KAKT activation, PIP3 activates AKT signaling, Regulation of signaling by CBL, Role of LAT2NTALLAB on calcium mobilization, Signaling by ERBB2, Signaling by FGFR1 in disease, Signaling by FGFR2 in disease, Signaling by FGFR3 in disease, Signaling by FGFR4 in disease, Signaling by PDGF, Signaling by SCF-KIT, Synthesis of PIPs at the plasma membrane, Tie2 Signaling, VEGFA-VEGFR2 Pathway 
                                               SEBELIUS_2_C4636 
                                               PIK3R1 
                                               mRNA 
                                               0 
                                               
                                               
                                               G alpha (q) signalling events;Downstream TCR signaling;Signaling by FGFR2 in disease;PIP3 activates AKT signaling;VEGFA-VEGFR2 Pathway;Synthesis of PIPs at the plasma membrane;Constitutive Signaling by Ligand-Responsive EGFR Cancer Variants;Interleukin receptor SHC signaling;G beta.CL.gamma signalling through PI3Kgamma;Regulation of signaling by CBL;Role of phospholipids in phagocytosis;Signaling by FGFR3 in disease;Constitutive Signaling by Aberrant PI3K in Cancer;Signaling by FGFR4 in disease;CD28 dependent PI3KAkt signaling;Tie2 Signaling;Signaling by FGFR1 fusion mutants;GP1b-IX-V activation signalling;GPVI-mediated activation cascade;Nephrin interactions;Constitutive Signaling by EGFRvIII;Gastrin-CREB signalling pathway via PKC and MAPK;Downstream signal transduction;GAB1 signalosome;Signaling by SCF-KIT;PI3KAKT activation;Role of LAT2NTALLAB on calcium mobilization;DAP12 signaling;Signaling by PDGF;Signaling by ERBB2;IRS-related events triggered by IGF1R;CD28 co-stimulation;Cell surface interactions at the vascular wall;Signaling by FGFR1 in disease 
                                               hsa04012;hsa04014;hsa04015;hsa04022;hsa04024;hsa04062;hsa04066;hsa04068;hsa04070;hsa04071;hsa04072;hsa04150;hsa04151;hsa04152;hsa04210;hsa04211;hsa04213;hsa04261;hsa04370;hsa04380;hsa04510;hsa04550;hsa04611;hsa04620;hsa04630;hsa04650;hsa04660;hsa04662;hsa04664;hsa04666;hsa04668;hsa04670;hsa04722;hsa04725;hsa04750;hsa04810;hsa04910;hsa04914;hsa04915;hsa04917;hsa04919;hsa04921;hsa04923;hsa04930;hsa04931;hsa04932;hsa04933;hsa04960;hsa04973;hsa05100;hsa05142;hsa05145;hsa05146;hsa05160;hsa05161;hsa05162;hsa05164;hsa05166;hsa05169;hsa05200;hsa05203;hsa05205;hsa05210;hsa05211;hsa05212;hsa05213;hsa05214;hsa05215;hsa05218;hsa05220;hsa05221;hsa05222;hsa05223;hsa05230;hsa05231 
                                               PIK3R1 
                                               NA 
                                               slateblue1 
                                               16 
                                               12.1341463414634 
                                             
                                             
                                               PIK3CA-mRNA 
                                               -1.22 
                                               0.145 
                                               -1.51 
                                               -0.94 
                                               0.428 
                                               0.352 
                                               0.521 
                                               3.77e-10 
                                               6.08e-09 
                                               lm.nb 
                                               CD28 co-stimulation, CD28 dependent PI3KAkt signaling, Cell surface interactions at the vascular wall, Constitutive Signaling by Aberrant PI3K in Cancer, Constitutive Signaling by EGFRvIII, DAP12 signaling, Downstream signal transduction, Downstream TCR signaling, G alpha (1213) signalling events, G alpha (q) signalling events, G beta.CL.gamma signalling through PI3Kgamma, GAB1 signalosome, Gastrin-CREB signalling pathway via PKC and MAPK, GPVI-mediated activation cascade, Interleukin receptor SHC signaling, IRS-related events triggered by IGF1R, Nephrin interactions, PI3K Cascade, PI3KAKT activation, PIP3 activates AKT signaling, Regulation of signaling by CBL, Role of LAT2NTALLAB on calcium mobilization, Signaling by ERBB2, Signaling by FGFR1 in disease, Signaling by FGFR2 in disease, Signaling by FGFR3 in disease, Signaling by FGFR4 in disease, Signaling by PDGF, Signaling by SCF-KIT, Synthesis of PIPs at the plasma membrane, Tie2 Signaling, VEGFA-VEGFR2 Pathway 
                                               SEBELIUS_2_C4636 
                                               PIK3CA 
                                               mRNA 
                                               0 
                                               
                                               
                                               G alpha (q) signalling events;Downstream TCR signaling;Signaling by FGFR2 in disease;PIP3 activates AKT signaling;VEGFA-VEGFR2 Pathway;Synthesis of PIPs at the plasma membrane;Constitutive Signaling by Ligand-Responsive EGFR Cancer Variants;Interleukin receptor SHC signaling;G beta.CL.gamma signalling through PI3Kgamma;Regulation of signaling by CBL;Role of phospholipids in phagocytosis;G alpha (1213) signalling events;PI3K Cascade;Signaling by FGFR3 in disease;Constitutive Signaling by Aberrant PI3K in Cancer;Signaling by FGFR4 in disease;CD28 dependent PI3KAkt signaling;Tie2 Signaling;GPVI-mediated activation cascade;Nephrin interactions;Constitutive Signaling by EGFRvIII;Gastrin-CREB signalling pathway via PKC and MAPK;Downstream signal transduction;GAB1 signalosome;Signaling by SCF-KIT;PI3KAKT activation;Role of LAT2NTALLAB on calcium mobilization;DAP12 signaling;Signaling by PDGF;Signaling by ERBB2;IRS-related events triggered by IGF1R;CD28 co-stimulation;Cell surface interactions at the vascular wall;Signaling by FGFR1 in disease 
                                               hsa00562;hsa04012;hsa04014;hsa04015;hsa04022;hsa04024;hsa04062;hsa04066;hsa04068;hsa04070;hsa04071;hsa04072;hsa04150;hsa04151;hsa04152;hsa04210;hsa04211;hsa04213;hsa04261;hsa04370;hsa04380;hsa04510;hsa04550;hsa04611;hsa04620;hsa04630;hsa04650;hsa04660;hsa04662;hsa04664;hsa04666;hsa04668;hsa04670;hsa04722;hsa04725;hsa04750;hsa04810;hsa04910;hsa04914;hsa04915;hsa04917;hsa04919;hsa04921;hsa04923;hsa04930;hsa04931;hsa04932;hsa04933;hsa04960;hsa04973;hsa05100;hsa05142;hsa05145;hsa05146;hsa05160;hsa05161;hsa05162;hsa05164;hsa05166;hsa05169;hsa05200;hsa05203;hsa05205;hsa05206;hsa05210;hsa05211;hsa05212;hsa05213;hsa05214;hsa05215;hsa05218;hsa05220;hsa05221;hsa05222;hsa05223;hsa05230;hsa05231 
                                               PIK3CA 
                                               NA 
                                               slateblue1 
                                               16 
                                               -8.41379310344828 
                                             
                                             
                                               FYN-mRNA 
                                               -2.64 
                                               0.335 
                                               -3.3 
                                               -1.99 
                                               0.16 
                                               0.102 
                                               0.253 
                                               1.96e-09 
                                               2.84e-08 
                                               lm.nb 
                                               ARMS-mediated activation, CD209 (DC-SIGN) signaling, CD28 co-stimulation, CD28 dependent PI3KAkt signaling, CD28 dependent Vav1 pathway, Cell surface interactions at the vascular wall, Constitutive Signaling by Aberrant PI3K in Cancer, CTLA4 inhibitory signaling, DAP12 signaling, Downstream signal transduction, EPH-ephrin mediated repulsion of cells, FCERI mediated MAPK activation, Frs2-mediated activation, GAB1 signalosome, Gastrin-CREB signalling pathway via PKC and MAPK, GPVI-mediated activation cascade, Interleukin receptor SHC signaling, IRS-related events triggered by IGF1R, Nephrin interactions, PI3KAKT activation, RAFMAP kinase cascade, Regulation of signaling by CBL, Role of LAT2NTALLAB on calcium mobilization, Sema3A PAK dependent Axon repulsion, Signaling by ERBB2, Signaling by PDGF, Signaling by SCF-KIT, VEGFA-VEGFR2 Pathway, VEGFR2 mediated cell proliferation 
                                               SEBELIUS_2_C4636 
                                               FYN 
                                               mRNA 
                                               0 
                                               
                                               
                                               EPH-ephrin mediated repulsion of cells;Regulation of signaling by CBL;FCGR activation;Antigen activates B Cell Receptor (BCR) leading to generation of second messengers;Dectin-2 family;RAFMAP kinase cascade;Constitutive Signaling by Aberrant PI3K in Cancer;CTLA4 inhibitory signaling;CD28 dependent PI3KAkt signaling;PECAM1 interactions;CD28 dependent Vav1 pathway;GPVI-mediated activation cascade;Sema3A PAK dependent Axon repulsion;Nephrin interactions;SEMA3A-Plexin repulsion signaling by inhibiting Integrin adhesion;CRMPs in Sema3A signaling;CD209 (DC-SIGN) signaling;Downstream signal transduction;GAB1 signalosome;Signaling by SCF-KIT;PI3KAKT activation;Role of LAT2NTALLAB on calcium mobilization;DAP12 signaling;Signaling by PDGF;Signaling by ERBB2;VEGFA-VEGFR2 Pathway;VEGFR2 mediated cell proliferation;Frs2-mediated activation;ARMS-mediated activation;Gastrin-CREB signalling pathway via PKC and MAPK;Interleukin receptor SHC signaling;FCERI mediated MAPK activation;IRS-related events triggered by IGF1R;CD28 co-stimulation;Cell surface interactions at the vascular wall 
                                               hsa04071;hsa04072;hsa04360;hsa04380;hsa04510;hsa04520;hsa04611;hsa04650;hsa04660;hsa04664;hsa04725;hsa05020;hsa05130;hsa05162;hsa05416 
                                               FYN 
                                               NA 
                                               slateblue1 
                                               16 
                                               -7.88059701492537 
                                             
                                             
                                               PIK3R3-mRNA 
                                               -2.73 
                                               0.417 
                                               -3.55 
                                               -1.91 
                                               0.151 
                                               0.0856 
                                               0.266 
                                               1.17e-07 
                                               1.44e-06 
                                               lm.nb 
                                               CD28 co-stimulation, CD28 dependent PI3KAkt signaling, Constitutive Signaling by Aberrant PI3K in Cancer, DAP12 signaling, Downstream signal transduction, G alpha (1213) signalling events, G alpha (q) signalling events, G beta.CL.gamma signalling through PI3Kgamma, GAB1 signalosome, Gastrin-CREB signalling pathway via PKC and MAPK, GPVI-mediated activation cascade, Interleukin receptor SHC signaling, PI3KAKT activation, PIP3 activates AKT signaling, Regulation of signaling by CBL, Role of LAT2NTALLAB on calcium mobilization, Signaling by ERBB2, Signaling by PDGF, Signaling by SCF-KIT, Synthesis of PIPs at the plasma membrane 
                                               SEBELIUS_2_C4636 
                                               PIK3R3 
                                               mRNA 
                                               0 
                                               
                                               
                                               G alpha (q) signalling events;PIP3 activates AKT signaling;Constitutive Signaling by Aberrant PI3K in Cancer;CD28 dependent PI3KAkt signaling;Synthesis of PIPs at the plasma membrane;Interleukin receptor SHC signaling;G beta.CL.gamma signalling through PI3Kgamma;Regulation of signaling by CBL;GPVI-mediated activation cascade;G alpha (1213) signalling events;Gastrin-CREB signalling pathway via PKC and MAPK;Downstream signal transduction;GAB1 signalosome;Signaling by SCF-KIT;PI3KAKT activation;Role of LAT2NTALLAB on calcium mobilization;DAP12 signaling;Signaling by PDGF;Signaling by ERBB2;CD28 co-stimulation 
                                               hsa04012;hsa04014;hsa04015;hsa04022;hsa04024;hsa04062;hsa04066;hsa04068;hsa04070;hsa04071;hsa04072;hsa04150;hsa04151;hsa04152;hsa04210;hsa04211;hsa04213;hsa04261;hsa04370;hsa04380;hsa04510;hsa04550;hsa04611;hsa04620;hsa04630;hsa04650;hsa04660;hsa04662;hsa04664;hsa04666;hsa04668;hsa04670;hsa04722;hsa04725;hsa04750;hsa04810;hsa04910;hsa04914;hsa04915;hsa04917;hsa04919;hsa04921;hsa04923;hsa04930;hsa04931;hsa04932;hsa04933;hsa04960;hsa04973;hsa05100;hsa05142;hsa05145;hsa05146;hsa05160;hsa05161;hsa05162;hsa05164;hsa05166;hsa05169;hsa05200;hsa05203;hsa05205;hsa05210;hsa05211;hsa05212;hsa05213;hsa05214;hsa05215;hsa05218;hsa05220;hsa05221;hsa05222;hsa05223;hsa05230;hsa05231 
                                               PIK3R3 
                                               NA 
                                               slateblue1 
                                               16 
                                               -6.54676258992806 
                                             
                                             
                                               PIK3R2-mRNA 
                                               -0.152 
                                               0.189 
                                               -0.523 
                                               0.218 
                                               0.9 
                                               0.696 
                                               1.16 
                                               0.425 
                                               1 
                                               lm.nb 
                                               CD28 co-stimulation, CD28 dependent PI3KAkt signaling, Cell surface interactions at the vascular wall, Constitutive Signaling by Aberrant PI3K in Cancer, DAP12 signaling, Downstream signal transduction, Downstream TCR signaling, G alpha (1213) signalling events, G alpha (q) signalling events, G beta.CL.gamma signalling through PI3Kgamma, GAB1 signalosome, Gastrin-CREB signalling pathway via PKC and MAPK, GPVI-mediated activation cascade, Interleukin receptor SHC signaling, IRS-related events triggered by IGF1R, Nephrin interactions, PI3K Cascade, PI3KAKT activation, PIP3 activates AKT signaling, Regulation of signaling by CBL, Rho GTPase cycle, Role of LAT2NTALLAB on calcium mobilization, Signaling by ERBB2, Signaling by PDGF, Signaling by SCF-KIT, Synthesis of PIPs at the plasma membrane, Tie2 Signaling, VEGFA-VEGFR2 Pathway 
                                               SEBELIUS_2_C4636 
                                               PIK3R2 
                                               mRNA 
                                               0 
                                               
                                               
                                               G alpha (q) signalling events;Downstream TCR signaling;PIP3 activates AKT signaling;VEGFA-VEGFR2 Pathway;Synthesis of PIPs at the plasma membrane;Rho GTPase cycle;Interleukin receptor SHC signaling;G beta.CL.gamma signalling through PI3Kgamma;Regulation of signaling by CBL;Role of phospholipids in phagocytosis;G alpha (1213) signalling events;PI3K Cascade;Constitutive Signaling by Aberrant PI3K in Cancer;CD28 dependent PI3KAkt signaling;Tie2 Signaling;GPVI-mediated activation cascade;Nephrin interactions;Gastrin-CREB signalling pathway via PKC and MAPK;Downstream signal transduction;GAB1 signalosome;Signaling by SCF-KIT;PI3KAKT activation;Role of LAT2NTALLAB on calcium mobilization;DAP12 signaling;Signaling by PDGF;Signaling by ERBB2;IRS-related events triggered by IGF1R;CD28 co-stimulation;Cell surface interactions at the vascular wall 
                                               hsa04012;hsa04014;hsa04015;hsa04022;hsa04024;hsa04062;hsa04066;hsa04068;hsa04070;hsa04071;hsa04072;hsa04150;hsa04151;hsa04152;hsa04210;hsa04211;hsa04213;hsa04261;hsa04370;hsa04380;hsa04510;hsa04550;hsa04611;hsa04620;hsa04630;hsa04650;hsa04660;hsa04662;hsa04664;hsa04666;hsa04668;hsa04670;hsa04722;hsa04725;hsa04750;hsa04810;hsa04910;hsa04914;hsa04915;hsa04917;hsa04919;hsa04921;hsa04923;hsa04930;hsa04931;hsa04932;hsa04933;hsa04960;hsa04973;hsa05100;hsa05142;hsa05145;hsa05146;hsa05160;hsa05161;hsa05162;hsa05164;hsa05166;hsa05169;hsa05200;hsa05203;hsa05205;hsa05206;hsa05210;hsa05211;hsa05212;hsa05213;hsa05214;hsa05215;hsa05218;hsa05220;hsa05221;hsa05222;hsa05223;hsa05230;hsa05231 
                                               PIK3R2 
                                               NA 
                                               azure4 
                                               1 
                                               -0.804232804232804 
                                             
                                           
                                           DE Results - CD28 dependent PI3KAkt signaling genes - Line: Caco2 vs.Fibro  Table displaying each sample's global significance scores and directed global significance scores as defined in the heatmaps above. The global significance score is calculated as the square root of the mean squared t-statistic for the genes in a gene set, with t-statistics coming from the linear regression underlying our differential expression analysis. The directed global significance score is calculated as the square root of the mean signed squared t-statistic for the genes in a gene set, with t-statistics coming from the linear regression underlying our differential expression analysis.   
                                      
                                       
                                  
                                
                                
                                  
                                       
                                      
                                           Volcano Plot: Line: HEPG2 vs.Fibro   More Plot Information                Volcano Plot: Line: HEPG2 vs.Fibro  Volcano plot displaying each gene's -log10(p-value) and log2 fold change for the selected covariate. Highly statistically significant genes fall at the top of the plot, and highly differentially expressed genes fall to either side. Genes within the selected gene set are highlighted in orange. Horizontal lines indicate various False Discovery Rate (FDR) thresholds.   
                                           DE Results - CD28 dependent PI3KAkt signaling genes - Line: HEPG2 vs.Fibro   More Plot Information      Download CSV Data     
                                           
                                             
                                             
                                           
                                           
                                             
                                               Probe Label 
                                               Log2 fold change 
                                               std error (log2) 
                                               Lower confidence limit (log2) 
                                               Upper confidence limit (log2) 
                                               Linear fold change 
                                               Lower confidence limit (linear) 
                                               Upper confidence limit (linear) 
                                               P-value 
                                               BY.p.value 
                                               method 
                                               Gene.sets 
                                               Codeset.Name 
                                               Probe.Label 
                                               Analyte.Type 
                                               Is.Control 
                                               Control.Type 
                                               Related.Probes 
                                               Probe.Annotation 
                                               KEGG.Pathways 
                                               Official.Gene.Name 
                                               Control.Conc 
                                               volcanocol 
                                               volcanopch 
                                               tstats.all 
                                             
                                           
                                           
                                             
                                               AKT3-mRNA 
                                               -7.38 
                                               0.252 
                                               -7.87 
                                               -6.89 
                                               0.006 
                                               0.00426 
                                               0.00845 
                                               3.23e-27 
                                               1.15e-24 
                                               lm.nb 
                                               Activation of BAD and translocation to mitochondria , CD28 co-stimulation, CD28 dependent PI3KAkt signaling, DAP12 signaling, Downstream signal transduction, G beta.CL.gamma signalling through PI3Kgamma, GAB1 signalosome, GPVI-mediated activation cascade, PI3KAKT activation, PIP3 activates AKT signaling, Role of LAT2NTALLAB on calcium mobilization, Signaling by ERBB2, Signaling by PDGF, Signaling by SCF-KIT, TP53 Regulates Metabolic Genes, VEGFA-VEGFR2 Pathway, VEGFR2 mediated vascular permeability 
                                               SEBELIUS_2_C4636 
                                               AKT3 
                                               mRNA 
                                               0 
                                               
                                               
                                               TP53 Regulates Metabolic Genes;CD28 dependent PI3KAkt signaling;AKT phosphorylates targets in the cytosol;Negative regulation of the PI3KAKT network;Activation of BAD and translocation to mitochondria ;G beta.CL.gamma signalling through PI3Kgamma;AKT phosphorylates targets in the nucleus;VEGFR2 mediated vascular permeability;GPVI-mediated activation cascade;Signaling by ERBB2;PIP3 activates AKT signaling;Downstream signal transduction;GAB1 signalosome;Signaling by SCF-KIT;PI3KAKT activation;Role of LAT2NTALLAB on calcium mobilization;DAP12 signaling;Signaling by PDGF;CD28 co-stimulation;VEGFA-VEGFR2 Pathway 
                                               hsa04010;hsa04012;hsa04014;hsa04015;hsa04022;hsa04024;hsa04062;hsa04066;hsa04068;hsa04071;hsa04072;hsa04150;hsa04151;hsa04152;hsa04210;hsa04211;hsa04213;hsa04261;hsa04370;hsa04380;hsa04510;hsa04530;hsa04550;hsa04611;hsa04620;hsa04630;hsa04660;hsa04662;hsa04664;hsa04666;hsa04668;hsa04722;hsa04725;hsa04728;hsa04910;hsa04914;hsa04915;hsa04917;hsa04919;hsa04920;hsa04922;hsa04923;hsa04931;hsa04932;hsa04933;hsa04973;hsa05142;hsa05145;hsa05152;hsa05160;hsa05161;hsa05162;hsa05164;hsa05166;hsa05169;hsa05200;hsa05205;hsa05210;hsa05211;hsa05212;hsa05213;hsa05214;hsa05215;hsa05218;hsa05220;hsa05221;hsa05222;hsa05223;hsa05230;hsa05231 
                                               AKT3 
                                               NA 
                                               slateblue1 
                                               16 
                                               -29.2857142857143 
                                             
                                             
                                               PIK3R1-mRNA 
                                               1.63 
                                               0.179 
                                               1.28 
                                               1.98 
                                               3.1 
                                               2.43 
                                               3.95 
                                               5.59e-11 
                                               1.13e-09 
                                               lm.nb 
                                               CD28 co-stimulation, CD28 dependent PI3KAkt signaling, Cell surface interactions at the vascular wall, Constitutive Signaling by Aberrant PI3K in Cancer, Constitutive Signaling by EGFRvIII, DAP12 signaling, Downstream signal transduction, Downstream TCR signaling, G alpha (q) signalling events, G beta.CL.gamma signalling through PI3Kgamma, GAB1 signalosome, Gastrin-CREB signalling pathway via PKC and MAPK, GPVI-mediated activation cascade, Interleukin receptor SHC signaling, IRS-related events triggered by IGF1R, Nephrin interactions, PI3KAKT activation, PIP3 activates AKT signaling, Regulation of signaling by CBL, Role of LAT2NTALLAB on calcium mobilization, Signaling by ERBB2, Signaling by FGFR1 in disease, Signaling by FGFR2 in disease, Signaling by FGFR3 in disease, Signaling by FGFR4 in disease, Signaling by PDGF, Signaling by SCF-KIT, Synthesis of PIPs at the plasma membrane, Tie2 Signaling, VEGFA-VEGFR2 Pathway 
                                               SEBELIUS_2_C4636 
                                               PIK3R1 
                                               mRNA 
                                               0 
                                               
                                               
                                               G alpha (q) signalling events;Downstream TCR signaling;Signaling by FGFR2 in disease;PIP3 activates AKT signaling;VEGFA-VEGFR2 Pathway;Synthesis of PIPs at the plasma membrane;Constitutive Signaling by Ligand-Responsive EGFR Cancer Variants;Interleukin receptor SHC signaling;G beta.CL.gamma signalling through PI3Kgamma;Regulation of signaling by CBL;Role of phospholipids in phagocytosis;Signaling by FGFR3 in disease;Constitutive Signaling by Aberrant PI3K in Cancer;Signaling by FGFR4 in disease;CD28 dependent PI3KAkt signaling;Tie2 Signaling;Signaling by FGFR1 fusion mutants;GP1b-IX-V activation signalling;GPVI-mediated activation cascade;Nephrin interactions;Constitutive Signaling by EGFRvIII;Gastrin-CREB signalling pathway via PKC and MAPK;Downstream signal transduction;GAB1 signalosome;Signaling by SCF-KIT;PI3KAKT activation;Role of LAT2NTALLAB on calcium mobilization;DAP12 signaling;Signaling by PDGF;Signaling by ERBB2;IRS-related events triggered by IGF1R;CD28 co-stimulation;Cell surface interactions at the vascular wall;Signaling by FGFR1 in disease 
                                               hsa04012;hsa04014;hsa04015;hsa04022;hsa04024;hsa04062;hsa04066;hsa04068;hsa04070;hsa04071;hsa04072;hsa04150;hsa04151;hsa04152;hsa04210;hsa04211;hsa04213;hsa04261;hsa04370;hsa04380;hsa04510;hsa04550;hsa04611;hsa04620;hsa04630;hsa04650;hsa04660;hsa04662;hsa04664;hsa04666;hsa04668;hsa04670;hsa04722;hsa04725;hsa04750;hsa04810;hsa04910;hsa04914;hsa04915;hsa04917;hsa04919;hsa04921;hsa04923;hsa04930;hsa04931;hsa04932;hsa04933;hsa04960;hsa04973;hsa05100;hsa05142;hsa05145;hsa05146;hsa05160;hsa05161;hsa05162;hsa05164;hsa05166;hsa05169;hsa05200;hsa05203;hsa05205;hsa05210;hsa05211;hsa05212;hsa05213;hsa05214;hsa05215;hsa05218;hsa05220;hsa05221;hsa05222;hsa05223;hsa05230;hsa05231 
                                               PIK3R1 
                                               NA 
                                               slateblue1 
                                               16 
                                               9.10614525139665 
                                             
                                             
                                               PIK3R3-mRNA 
                                               -1.77 
                                               0.457 
                                               -2.67 
                                               -0.874 
                                               0.293 
                                               0.157 
                                               0.546 
                                               0.000425 
                                               0.00402 
                                               lm.nb 
                                               CD28 co-stimulation, CD28 dependent PI3KAkt signaling, Constitutive Signaling by Aberrant PI3K in Cancer, DAP12 signaling, Downstream signal transduction, G alpha (1213) signalling events, G alpha (q) signalling events, G beta.CL.gamma signalling through PI3Kgamma, GAB1 signalosome, Gastrin-CREB signalling pathway via PKC and MAPK, GPVI-mediated activation cascade, Interleukin receptor SHC signaling, PI3KAKT activation, PIP3 activates AKT signaling, Regulation of signaling by CBL, Role of LAT2NTALLAB on calcium mobilization, Signaling by ERBB2, Signaling by PDGF, Signaling by SCF-KIT, Synthesis of PIPs at the plasma membrane 
                                               SEBELIUS_2_C4636 
                                               PIK3R3 
                                               mRNA 
                                               0 
                                               
                                               
                                               G alpha (q) signalling events;PIP3 activates AKT signaling;Constitutive Signaling by Aberrant PI3K in Cancer;CD28 dependent PI3KAkt signaling;Synthesis of PIPs at the plasma membrane;Interleukin receptor SHC signaling;G beta.CL.gamma signalling through PI3Kgamma;Regulation of signaling by CBL;GPVI-mediated activation cascade;G alpha (1213) signalling events;Gastrin-CREB signalling pathway via PKC and MAPK;Downstream signal transduction;GAB1 signalosome;Signaling by SCF-KIT;PI3KAKT activation;Role of LAT2NTALLAB on calcium mobilization;DAP12 signaling;Signaling by PDGF;Signaling by ERBB2;CD28 co-stimulation 
                                               hsa04012;hsa04014;hsa04015;hsa04022;hsa04024;hsa04062;hsa04066;hsa04068;hsa04070;hsa04071;hsa04072;hsa04150;hsa04151;hsa04152;hsa04210;hsa04211;hsa04213;hsa04261;hsa04370;hsa04380;hsa04510;hsa04550;hsa04611;hsa04620;hsa04630;hsa04650;hsa04660;hsa04662;hsa04664;hsa04666;hsa04668;hsa04670;hsa04722;hsa04725;hsa04750;hsa04810;hsa04910;hsa04914;hsa04915;hsa04917;hsa04919;hsa04921;hsa04923;hsa04930;hsa04931;hsa04932;hsa04933;hsa04960;hsa04973;hsa05100;hsa05142;hsa05145;hsa05146;hsa05160;hsa05161;hsa05162;hsa05164;hsa05166;hsa05169;hsa05200;hsa05203;hsa05205;hsa05210;hsa05211;hsa05212;hsa05213;hsa05214;hsa05215;hsa05218;hsa05220;hsa05221;hsa05222;hsa05223;hsa05230;hsa05231 
                                               PIK3R3 
                                               NA 
                                               slateblue1 
                                               16 
                                               -3.87308533916849 
                                             
                                             
                                               PIK3R2-mRNA 
                                               0.61 
                                               0.207 
                                               0.204 
                                               1.02 
                                               1.53 
                                               1.15 
                                               2.02 
                                               0.00552 
                                               0.0469 
                                               lm.nb 
                                               CD28 co-stimulation, CD28 dependent PI3KAkt signaling, Cell surface interactions at the vascular wall, Constitutive Signaling by Aberrant PI3K in Cancer, DAP12 signaling, Downstream signal transduction, Downstream TCR signaling, G alpha (1213) signalling events, G alpha (q) signalling events, G beta.CL.gamma signalling through PI3Kgamma, GAB1 signalosome, Gastrin-CREB signalling pathway via PKC and MAPK, GPVI-mediated activation cascade, Interleukin receptor SHC signaling, IRS-related events triggered by IGF1R, Nephrin interactions, PI3K Cascade, PI3KAKT activation, PIP3 activates AKT signaling, Regulation of signaling by CBL, Rho GTPase cycle, Role of LAT2NTALLAB on calcium mobilization, Signaling by ERBB2, Signaling by PDGF, Signaling by SCF-KIT, Synthesis of PIPs at the plasma membrane, Tie2 Signaling, VEGFA-VEGFR2 Pathway 
                                               SEBELIUS_2_C4636 
                                               PIK3R2 
                                               mRNA 
                                               0 
                                               
                                               
                                               G alpha (q) signalling events;Downstream TCR signaling;PIP3 activates AKT signaling;VEGFA-VEGFR2 Pathway;Synthesis of PIPs at the plasma membrane;Rho GTPase cycle;Interleukin receptor SHC signaling;G beta.CL.gamma signalling through PI3Kgamma;Regulation of signaling by CBL;Role of phospholipids in phagocytosis;G alpha (1213) signalling events;PI3K Cascade;Constitutive Signaling by Aberrant PI3K in Cancer;CD28 dependent PI3KAkt signaling;Tie2 Signaling;GPVI-mediated activation cascade;Nephrin interactions;Gastrin-CREB signalling pathway via PKC and MAPK;Downstream signal transduction;GAB1 signalosome;Signaling by SCF-KIT;PI3KAKT activation;Role of LAT2NTALLAB on calcium mobilization;DAP12 signaling;Signaling by PDGF;Signaling by ERBB2;IRS-related events triggered by IGF1R;CD28 co-stimulation;Cell surface interactions at the vascular wall 
                                               hsa04012;hsa04014;hsa04015;hsa04022;hsa04024;hsa04062;hsa04066;hsa04068;hsa04070;hsa04071;hsa04072;hsa04150;hsa04151;hsa04152;hsa04210;hsa04211;hsa04213;hsa04261;hsa04370;hsa04380;hsa04510;hsa04550;hsa04611;hsa04620;hsa04630;hsa04650;hsa04660;hsa04662;hsa04664;hsa04666;hsa04668;hsa04670;hsa04722;hsa04725;hsa04750;hsa04810;hsa04910;hsa04914;hsa04915;hsa04917;hsa04919;hsa04921;hsa04923;hsa04930;hsa04931;hsa04932;hsa04933;hsa04960;hsa04973;hsa05100;hsa05142;hsa05145;hsa05146;hsa05160;hsa05161;hsa05162;hsa05164;hsa05166;hsa05169;hsa05200;hsa05203;hsa05205;hsa05206;hsa05210;hsa05211;hsa05212;hsa05213;hsa05214;hsa05215;hsa05218;hsa05220;hsa05221;hsa05222;hsa05223;hsa05230;hsa05231 
                                               PIK3R2 
                                               NA 
                                               slateblue3 
                                               16 
                                               2.94685990338164 
                                             
                                             
                                               PIK3CA-mRNA 
                                               -0.437 
                                               0.159 
                                               -0.748 
                                               -0.126 
                                               0.739 
                                               0.595 
                                               0.916 
                                               0.00908 
                                               0.0753 
                                               lm.nb 
                                               CD28 co-stimulation, CD28 dependent PI3KAkt signaling, Cell surface interactions at the vascular wall, Constitutive Signaling by Aberrant PI3K in Cancer, Constitutive Signaling by EGFRvIII, DAP12 signaling, Downstream signal transduction, Downstream TCR signaling, G alpha (1213) signalling events, G alpha (q) signalling events, G beta.CL.gamma signalling through PI3Kgamma, GAB1 signalosome, Gastrin-CREB signalling pathway via PKC and MAPK, GPVI-mediated activation cascade, Interleukin receptor SHC signaling, IRS-related events triggered by IGF1R, Nephrin interactions, PI3K Cascade, PI3KAKT activation, PIP3 activates AKT signaling, Regulation of signaling by CBL, Role of LAT2NTALLAB on calcium mobilization, Signaling by ERBB2, Signaling by FGFR1 in disease, Signaling by FGFR2 in disease, Signaling by FGFR3 in disease, Signaling by FGFR4 in disease, Signaling by PDGF, Signaling by SCF-KIT, Synthesis of PIPs at the plasma membrane, Tie2 Signaling, VEGFA-VEGFR2 Pathway 
                                               SEBELIUS_2_C4636 
                                               PIK3CA 
                                               mRNA 
                                               0 
                                               
                                               
                                               G alpha (q) signalling events;Downstream TCR signaling;Signaling by FGFR2 in disease;PIP3 activates AKT signaling;VEGFA-VEGFR2 Pathway;Synthesis of PIPs at the plasma membrane;Constitutive Signaling by Ligand-Responsive EGFR Cancer Variants;Interleukin receptor SHC signaling;G beta.CL.gamma signalling through PI3Kgamma;Regulation of signaling by CBL;Role of phospholipids in phagocytosis;G alpha (1213) signalling events;PI3K Cascade;Signaling by FGFR3 in disease;Constitutive Signaling by Aberrant PI3K in Cancer;Signaling by FGFR4 in disease;CD28 dependent PI3KAkt signaling;Tie2 Signaling;GPVI-mediated activation cascade;Nephrin interactions;Constitutive Signaling by EGFRvIII;Gastrin-CREB signalling pathway via PKC and MAPK;Downstream signal transduction;GAB1 signalosome;Signaling by SCF-KIT;PI3KAKT activation;Role of LAT2NTALLAB on calcium mobilization;DAP12 signaling;Signaling by PDGF;Signaling by ERBB2;IRS-related events triggered by IGF1R;CD28 co-stimulation;Cell surface interactions at the vascular wall;Signaling by FGFR1 in disease 
                                               hsa00562;hsa04012;hsa04014;hsa04015;hsa04022;hsa04024;hsa04062;hsa04066;hsa04068;hsa04070;hsa04071;hsa04072;hsa04150;hsa04151;hsa04152;hsa04210;hsa04211;hsa04213;hsa04261;hsa04370;hsa04380;hsa04510;hsa04550;hsa04611;hsa04620;hsa04630;hsa04650;hsa04660;hsa04662;hsa04664;hsa04666;hsa04668;hsa04670;hsa04722;hsa04725;hsa04750;hsa04810;hsa04910;hsa04914;hsa04915;hsa04917;hsa04919;hsa04921;hsa04923;hsa04930;hsa04931;hsa04932;hsa04933;hsa04960;hsa04973;hsa05100;hsa05142;hsa05145;hsa05146;hsa05160;hsa05161;hsa05162;hsa05164;hsa05166;hsa05169;hsa05200;hsa05203;hsa05205;hsa05206;hsa05210;hsa05211;hsa05212;hsa05213;hsa05214;hsa05215;hsa05218;hsa05220;hsa05221;hsa05222;hsa05223;hsa05230;hsa05231 
                                               PIK3CA 
                                               NA 
                                               slateblue3 
                                               16 
                                               -2.74842767295597 
                                             
                                             
                                               FYN-mRNA 
                                               0.596 
                                               0.365 
                                               -0.119 
                                               1.31 
                                               1.51 
                                               0.921 
                                               2.48 
                                               0.111 
                                               0.804 
                                               lm.nb 
                                               ARMS-mediated activation, CD209 (DC-SIGN) signaling, CD28 co-stimulation, CD28 dependent PI3KAkt signaling, CD28 dependent Vav1 pathway, Cell surface interactions at the vascular wall, Constitutive Signaling by Aberrant PI3K in Cancer, CTLA4 inhibitory signaling, DAP12 signaling, Downstream signal transduction, EPH-ephrin mediated repulsion of cells, FCERI mediated MAPK activation, Frs2-mediated activation, GAB1 signalosome, Gastrin-CREB signalling pathway via PKC and MAPK, GPVI-mediated activation cascade, Interleukin receptor SHC signaling, IRS-related events triggered by IGF1R, Nephrin interactions, PI3KAKT activation, RAFMAP kinase cascade, Regulation of signaling by CBL, Role of LAT2NTALLAB on calcium mobilization, Sema3A PAK dependent Axon repulsion, Signaling by ERBB2, Signaling by PDGF, Signaling by SCF-KIT, VEGFA-VEGFR2 Pathway, VEGFR2 mediated cell proliferation 
                                               SEBELIUS_2_C4636 
                                               FYN 
                                               mRNA 
                                               0 
                                               
                                               
                                               EPH-ephrin mediated repulsion of cells;Regulation of signaling by CBL;FCGR activation;Antigen activates B Cell Receptor (BCR) leading to generation of second messengers;Dectin-2 family;RAFMAP kinase cascade;Constitutive Signaling by Aberrant PI3K in Cancer;CTLA4 inhibitory signaling;CD28 dependent PI3KAkt signaling;PECAM1 interactions;CD28 dependent Vav1 pathway;GPVI-mediated activation cascade;Sema3A PAK dependent Axon repulsion;Nephrin interactions;SEMA3A-Plexin repulsion signaling by inhibiting Integrin adhesion;CRMPs in Sema3A signaling;CD209 (DC-SIGN) signaling;Downstream signal transduction;GAB1 signalosome;Signaling by SCF-KIT;PI3KAKT activation;Role of LAT2NTALLAB on calcium mobilization;DAP12 signaling;Signaling by PDGF;Signaling by ERBB2;VEGFA-VEGFR2 Pathway;VEGFR2 mediated cell proliferation;Frs2-mediated activation;ARMS-mediated activation;Gastrin-CREB signalling pathway via PKC and MAPK;Interleukin receptor SHC signaling;FCERI mediated MAPK activation;IRS-related events triggered by IGF1R;CD28 co-stimulation;Cell surface interactions at the vascular wall 
                                               hsa04071;hsa04072;hsa04360;hsa04380;hsa04510;hsa04520;hsa04611;hsa04650;hsa04660;hsa04664;hsa04725;hsa05020;hsa05130;hsa05162;hsa05416 
                                               FYN 
                                               NA 
                                               azure4 
                                               1 
                                               1.63287671232877 
                                             
                                           
                                           DE Results - CD28 dependent PI3KAkt signaling genes - Line: HEPG2 vs.Fibro  Table displaying each sample's global significance scores and directed global significance scores as defined in the heatmaps above. The global significance score is calculated as the square root of the mean squared t-statistic for the genes in a gene set, with t-statistics coming from the linear regression underlying our differential expression analysis. The directed global significance score is calculated as the square root of the mean signed squared t-statistic for the genes in a gene set, with t-statistics coming from the linear regression underlying our differential expression analysis.   
                                      
                                       
                                  
                                
                                
                                  
                                       
                                      
                                           Volcano Plot: Line: HT29 vs.Fibro   More Plot Information                Volcano Plot: Line: HT29 vs.Fibro  Volcano plot displaying each gene's -log10(p-value) and log2 fold change for the selected covariate. Highly statistically significant genes fall at the top of the plot, and highly differentially expressed genes fall to either side. Genes within the selected gene set are highlighted in orange. Horizontal lines indicate various False Discovery Rate (FDR) thresholds.   
                                           DE Results - CD28 dependent PI3KAkt signaling genes - Line: HT29 vs.Fibro   More Plot Information      Download CSV Data     
                                           
                                             
                                             
                                           
                                           
                                             
                                               Probe Label 
                                               Log2 fold change 
                                               std error (log2) 
                                               Lower confidence limit (log2) 
                                               Upper confidence limit (log2) 
                                               Linear fold change 
                                               Lower confidence limit (linear) 
                                               Upper confidence limit (linear) 
                                               P-value 
                                               BY.p.value 
                                               method 
                                               Gene.sets 
                                               Codeset.Name 
                                               Probe.Label 
                                               Analyte.Type 
                                               Is.Control 
                                               Control.Type 
                                               Related.Probes 
                                               Probe.Annotation 
                                               KEGG.Pathways 
                                               Official.Gene.Name 
                                               Control.Conc 
                                               volcanocol 
                                               volcanopch 
                                               tstats.all 
                                             
                                           
                                           
                                             
                                               AKT3-mRNA 
                                               -8.38 
                                               0.265 
                                               -8.9 
                                               -7.86 
                                               0.003 
                                               0.00209 
                                               0.0043 
                                               2.3e-28 
                                               4.09e-26 
                                               lm.nb 
                                               Activation of BAD and translocation to mitochondria , CD28 co-stimulation, CD28 dependent PI3KAkt signaling, DAP12 signaling, Downstream signal transduction, G beta.CL.gamma signalling through PI3Kgamma, GAB1 signalosome, GPVI-mediated activation cascade, PI3KAKT activation, PIP3 activates AKT signaling, Role of LAT2NTALLAB on calcium mobilization, Signaling by ERBB2, Signaling by PDGF, Signaling by SCF-KIT, TP53 Regulates Metabolic Genes, VEGFA-VEGFR2 Pathway, VEGFR2 mediated vascular permeability 
                                               SEBELIUS_2_C4636 
                                               AKT3 
                                               mRNA 
                                               0 
                                               
                                               
                                               TP53 Regulates Metabolic Genes;CD28 dependent PI3KAkt signaling;AKT phosphorylates targets in the cytosol;Negative regulation of the PI3KAKT network;Activation of BAD and translocation to mitochondria ;G beta.CL.gamma signalling through PI3Kgamma;AKT phosphorylates targets in the nucleus;VEGFR2 mediated vascular permeability;GPVI-mediated activation cascade;Signaling by ERBB2;PIP3 activates AKT signaling;Downstream signal transduction;GAB1 signalosome;Signaling by SCF-KIT;PI3KAKT activation;Role of LAT2NTALLAB on calcium mobilization;DAP12 signaling;Signaling by PDGF;CD28 co-stimulation;VEGFA-VEGFR2 Pathway 
                                               hsa04010;hsa04012;hsa04014;hsa04015;hsa04022;hsa04024;hsa04062;hsa04066;hsa04068;hsa04071;hsa04072;hsa04150;hsa04151;hsa04152;hsa04210;hsa04211;hsa04213;hsa04261;hsa04370;hsa04380;hsa04510;hsa04530;hsa04550;hsa04611;hsa04620;hsa04630;hsa04660;hsa04662;hsa04664;hsa04666;hsa04668;hsa04722;hsa04725;hsa04728;hsa04910;hsa04914;hsa04915;hsa04917;hsa04919;hsa04920;hsa04922;hsa04923;hsa04931;hsa04932;hsa04933;hsa04973;hsa05142;hsa05145;hsa05152;hsa05160;hsa05161;hsa05162;hsa05164;hsa05166;hsa05169;hsa05200;hsa05205;hsa05210;hsa05211;hsa05212;hsa05213;hsa05214;hsa05215;hsa05218;hsa05220;hsa05221;hsa05222;hsa05223;hsa05230;hsa05231 
                                               AKT3 
                                               NA 
                                               slateblue1 
                                               16 
                                               -31.622641509434 
                                             
                                             
                                               PIK3R1-mRNA 
                                               2.62 
                                               0.169 
                                               2.29 
                                               2.95 
                                               6.15 
                                               4.89 
                                               7.74 
                                               8.93e-18 
                                               2.88e-16 
                                               lm.nb 
                                               CD28 co-stimulation, CD28 dependent PI3KAkt signaling, Cell surface interactions at the vascular wall, Constitutive Signaling by Aberrant PI3K in Cancer, Constitutive Signaling by EGFRvIII, DAP12 signaling, Downstream signal transduction, Downstream TCR signaling, G alpha (q) signalling events, G beta.CL.gamma signalling through PI3Kgamma, GAB1 signalosome, Gastrin-CREB signalling pathway via PKC and MAPK, GPVI-mediated activation cascade, Interleukin receptor SHC signaling, IRS-related events triggered by IGF1R, Nephrin interactions, PI3KAKT activation, PIP3 activates AKT signaling, Regulation of signaling by CBL, Role of LAT2NTALLAB on calcium mobilization, Signaling by ERBB2, Signaling by FGFR1 in disease, Signaling by FGFR2 in disease, Signaling by FGFR3 in disease, Signaling by FGFR4 in disease, Signaling by PDGF, Signaling by SCF-KIT, Synthesis of PIPs at the plasma membrane, Tie2 Signaling, VEGFA-VEGFR2 Pathway 
                                               SEBELIUS_2_C4636 
                                               PIK3R1 
                                               mRNA 
                                               0 
                                               
                                               
                                               G alpha (q) signalling events;Downstream TCR signaling;Signaling by FGFR2 in disease;PIP3 activates AKT signaling;VEGFA-VEGFR2 Pathway;Synthesis of PIPs at the plasma membrane;Constitutive Signaling by Ligand-Responsive EGFR Cancer Variants;Interleukin receptor SHC signaling;G beta.CL.gamma signalling through PI3Kgamma;Regulation of signaling by CBL;Role of phospholipids in phagocytosis;Signaling by FGFR3 in disease;Constitutive Signaling by Aberrant PI3K in Cancer;Signaling by FGFR4 in disease;CD28 dependent PI3KAkt signaling;Tie2 Signaling;Signaling by FGFR1 fusion mutants;GP1b-IX-V activation signalling;GPVI-mediated activation cascade;Nephrin interactions;Constitutive Signaling by EGFRvIII;Gastrin-CREB signalling pathway via PKC and MAPK;Downstream signal transduction;GAB1 signalosome;Signaling by SCF-KIT;PI3KAKT activation;Role of LAT2NTALLAB on calcium mobilization;DAP12 signaling;Signaling by PDGF;Signaling by ERBB2;IRS-related events triggered by IGF1R;CD28 co-stimulation;Cell surface interactions at the vascular wall;Signaling by FGFR1 in disease 
                                               hsa04012;hsa04014;hsa04015;hsa04022;hsa04024;hsa04062;hsa04066;hsa04068;hsa04070;hsa04071;hsa04072;hsa04150;hsa04151;hsa04152;hsa04210;hsa04211;hsa04213;hsa04261;hsa04370;hsa04380;hsa04510;hsa04550;hsa04611;hsa04620;hsa04630;hsa04650;hsa04660;hsa04662;hsa04664;hsa04666;hsa04668;hsa04670;hsa04722;hsa04725;hsa04750;hsa04810;hsa04910;hsa04914;hsa04915;hsa04917;hsa04919;hsa04921;hsa04923;hsa04930;hsa04931;hsa04932;hsa04933;hsa04960;hsa04973;hsa05100;hsa05142;hsa05145;hsa05146;hsa05160;hsa05161;hsa05162;hsa05164;hsa05166;hsa05169;hsa05200;hsa05203;hsa05205;hsa05210;hsa05211;hsa05212;hsa05213;hsa05214;hsa05215;hsa05218;hsa05220;hsa05221;hsa05222;hsa05223;hsa05230;hsa05231 
                                               PIK3R1 
                                               NA 
                                               slateblue1 
                                               16 
                                               15.5029585798817 
                                             
                                             
                                               FYN-mRNA 
                                               -1.21 
                                               0.345 
                                               -1.89 
                                               -0.534 
                                               0.432 
                                               0.271 
                                               0.69 
                                               0.00119 
                                               0.01 
                                               lm.nb 
                                               ARMS-mediated activation, CD209 (DC-SIGN) signaling, CD28 co-stimulation, CD28 dependent PI3KAkt signaling, CD28 dependent Vav1 pathway, Cell surface interactions at the vascular wall, Constitutive Signaling by Aberrant PI3K in Cancer, CTLA4 inhibitory signaling, DAP12 signaling, Downstream signal transduction, EPH-ephrin mediated repulsion of cells, FCERI mediated MAPK activation, Frs2-mediated activation, GAB1 signalosome, Gastrin-CREB signalling pathway via PKC and MAPK, GPVI-mediated activation cascade, Interleukin receptor SHC signaling, IRS-related events triggered by IGF1R, Nephrin interactions, PI3KAKT activation, RAFMAP kinase cascade, Regulation of signaling by CBL, Role of LAT2NTALLAB on calcium mobilization, Sema3A PAK dependent Axon repulsion, Signaling by ERBB2, Signaling by PDGF, Signaling by SCF-KIT, VEGFA-VEGFR2 Pathway, VEGFR2 mediated cell proliferation 
                                               SEBELIUS_2_C4636 
                                               FYN 
                                               mRNA 
                                               0 
                                               
                                               
                                               EPH-ephrin mediated repulsion of cells;Regulation of signaling by CBL;FCGR activation;Antigen activates B Cell Receptor (BCR) leading to generation of second messengers;Dectin-2 family;RAFMAP kinase cascade;Constitutive Signaling by Aberrant PI3K in Cancer;CTLA4 inhibitory signaling;CD28 dependent PI3KAkt signaling;PECAM1 interactions;CD28 dependent Vav1 pathway;GPVI-mediated activation cascade;Sema3A PAK dependent Axon repulsion;Nephrin interactions;SEMA3A-Plexin repulsion signaling by inhibiting Integrin adhesion;CRMPs in Sema3A signaling;CD209 (DC-SIGN) signaling;Downstream signal transduction;GAB1 signalosome;Signaling by SCF-KIT;PI3KAKT activation;Role of LAT2NTALLAB on calcium mobilization;DAP12 signaling;Signaling by PDGF;Signaling by ERBB2;VEGFA-VEGFR2 Pathway;VEGFR2 mediated cell proliferation;Frs2-mediated activation;ARMS-mediated activation;Gastrin-CREB signalling pathway via PKC and MAPK;Interleukin receptor SHC signaling;FCERI mediated MAPK activation;IRS-related events triggered by IGF1R;CD28 co-stimulation;Cell surface interactions at the vascular wall 
                                               hsa04071;hsa04072;hsa04360;hsa04380;hsa04510;hsa04520;hsa04611;hsa04650;hsa04660;hsa04664;hsa04725;hsa05020;hsa05130;hsa05162;hsa05416 
                                               FYN 
                                               NA 
                                               slateblue2 
                                               16 
                                               -3.50724637681159 
                                             
                                             
                                               PIK3R2-mRNA 
                                               -0.659 
                                               0.196 
                                               -1.04 
                                               -0.275 
                                               0.633 
                                               0.485 
                                               0.826 
                                               0.00179 
                                               0.015 
                                               lm.nb 
                                               CD28 co-stimulation, CD28 dependent PI3KAkt signaling, Cell surface interactions at the vascular wall, Constitutive Signaling by Aberrant PI3K in Cancer, DAP12 signaling, Downstream signal transduction, Downstream TCR signaling, G alpha (1213) signalling events, G alpha (q) signalling events, G beta.CL.gamma signalling through PI3Kgamma, GAB1 signalosome, Gastrin-CREB signalling pathway via PKC and MAPK, GPVI-mediated activation cascade, Interleukin receptor SHC signaling, IRS-related events triggered by IGF1R, Nephrin interactions, PI3K Cascade, PI3KAKT activation, PIP3 activates AKT signaling, Regulation of signaling by CBL, Rho GTPase cycle, Role of LAT2NTALLAB on calcium mobilization, Signaling by ERBB2, Signaling by PDGF, Signaling by SCF-KIT, Synthesis of PIPs at the plasma membrane, Tie2 Signaling, VEGFA-VEGFR2 Pathway 
                                               SEBELIUS_2_C4636 
                                               PIK3R2 
                                               mRNA 
                                               0 
                                               
                                               
                                               G alpha (q) signalling events;Downstream TCR signaling;PIP3 activates AKT signaling;VEGFA-VEGFR2 Pathway;Synthesis of PIPs at the plasma membrane;Rho GTPase cycle;Interleukin receptor SHC signaling;G beta.CL.gamma signalling through PI3Kgamma;Regulation of signaling by CBL;Role of phospholipids in phagocytosis;G alpha (1213) signalling events;PI3K Cascade;Constitutive Signaling by Aberrant PI3K in Cancer;CD28 dependent PI3KAkt signaling;Tie2 Signaling;GPVI-mediated activation cascade;Nephrin interactions;Gastrin-CREB signalling pathway via PKC and MAPK;Downstream signal transduction;GAB1 signalosome;Signaling by SCF-KIT;PI3KAKT activation;Role of LAT2NTALLAB on calcium mobilization;DAP12 signaling;Signaling by PDGF;Signaling by ERBB2;IRS-related events triggered by IGF1R;CD28 co-stimulation;Cell surface interactions at the vascular wall 
                                               hsa04012;hsa04014;hsa04015;hsa04022;hsa04024;hsa04062;hsa04066;hsa04068;hsa04070;hsa04071;hsa04072;hsa04150;hsa04151;hsa04152;hsa04210;hsa04211;hsa04213;hsa04261;hsa04370;hsa04380;hsa04510;hsa04550;hsa04611;hsa04620;hsa04630;hsa04650;hsa04660;hsa04662;hsa04664;hsa04666;hsa04668;hsa04670;hsa04722;hsa04725;hsa04750;hsa04810;hsa04910;hsa04914;hsa04915;hsa04917;hsa04919;hsa04921;hsa04923;hsa04930;hsa04931;hsa04932;hsa04933;hsa04960;hsa04973;hsa05100;hsa05142;hsa05145;hsa05146;hsa05160;hsa05161;hsa05162;hsa05164;hsa05166;hsa05169;hsa05200;hsa05203;hsa05205;hsa05206;hsa05210;hsa05211;hsa05212;hsa05213;hsa05214;hsa05215;hsa05218;hsa05220;hsa05221;hsa05222;hsa05223;hsa05230;hsa05231 
                                               PIK3R2 
                                               NA 
                                               slateblue2 
                                               16 
                                               -3.36224489795918 
                                             
                                             
                                               PIK3CA-mRNA 
                                               -0.45 
                                               0.149 
                                               -0.743 
                                               -0.157 
                                               0.732 
                                               0.598 
                                               0.897 
                                               0.00468 
                                               0.0377 
                                               lm.nb 
                                               CD28 co-stimulation, CD28 dependent PI3KAkt signaling, Cell surface interactions at the vascular wall, Constitutive Signaling by Aberrant PI3K in Cancer, Constitutive Signaling by EGFRvIII, DAP12 signaling, Downstream signal transduction, Downstream TCR signaling, G alpha (1213) signalling events, G alpha (q) signalling events, G beta.CL.gamma signalling through PI3Kgamma, GAB1 signalosome, Gastrin-CREB signalling pathway via PKC and MAPK, GPVI-mediated activation cascade, Interleukin receptor SHC signaling, IRS-related events triggered by IGF1R, Nephrin interactions, PI3K Cascade, PI3KAKT activation, PIP3 activates AKT signaling, Regulation of signaling by CBL, Role of LAT2NTALLAB on calcium mobilization, Signaling by ERBB2, Signaling by FGFR1 in disease, Signaling by FGFR2 in disease, Signaling by FGFR3 in disease, Signaling by FGFR4 in disease, Signaling by PDGF, Signaling by SCF-KIT, Synthesis of PIPs at the plasma membrane, Tie2 Signaling, VEGFA-VEGFR2 Pathway 
                                               SEBELIUS_2_C4636 
                                               PIK3CA 
                                               mRNA 
                                               0 
                                               
                                               
                                               G alpha (q) signalling events;Downstream TCR signaling;Signaling by FGFR2 in disease;PIP3 activates AKT signaling;VEGFA-VEGFR2 Pathway;Synthesis of PIPs at the plasma membrane;Constitutive Signaling by Ligand-Responsive EGFR Cancer Variants;Interleukin receptor SHC signaling;G beta.CL.gamma signalling through PI3Kgamma;Regulation of signaling by CBL;Role of phospholipids in phagocytosis;G alpha (1213) signalling events;PI3K Cascade;Signaling by FGFR3 in disease;Constitutive Signaling by Aberrant PI3K in Cancer;Signaling by FGFR4 in disease;CD28 dependent PI3KAkt signaling;Tie2 Signaling;GPVI-mediated activation cascade;Nephrin interactions;Constitutive Signaling by EGFRvIII;Gastrin-CREB signalling pathway via PKC and MAPK;Downstream signal transduction;GAB1 signalosome;Signaling by SCF-KIT;PI3KAKT activation;Role of LAT2NTALLAB on calcium mobilization;DAP12 signaling;Signaling by PDGF;Signaling by ERBB2;IRS-related events triggered by IGF1R;CD28 co-stimulation;Cell surface interactions at the vascular wall;Signaling by FGFR1 in disease 
                                               hsa00562;hsa04012;hsa04014;hsa04015;hsa04022;hsa04024;hsa04062;hsa04066;hsa04068;hsa04070;hsa04071;hsa04072;hsa04150;hsa04151;hsa04152;hsa04210;hsa04211;hsa04213;hsa04261;hsa04370;hsa04380;hsa04510;hsa04550;hsa04611;hsa04620;hsa04630;hsa04650;hsa04660;hsa04662;hsa04664;hsa04666;hsa04668;hsa04670;hsa04722;hsa04725;hsa04750;hsa04810;hsa04910;hsa04914;hsa04915;hsa04917;hsa04919;hsa04921;hsa04923;hsa04930;hsa04931;hsa04932;hsa04933;hsa04960;hsa04973;hsa05100;hsa05142;hsa05145;hsa05146;hsa05160;hsa05161;hsa05162;hsa05164;hsa05166;hsa05169;hsa05200;hsa05203;hsa05205;hsa05206;hsa05210;hsa05211;hsa05212;hsa05213;hsa05214;hsa05215;hsa05218;hsa05220;hsa05221;hsa05222;hsa05223;hsa05230;hsa05231 
                                               PIK3CA 
                                               NA 
                                               slateblue2 
                                               16 
                                               -3.02013422818792 
                                             
                                             
                                               PIK3R3-mRNA 
                                               0.688 
                                               0.429 
                                               -0.152 
                                               1.53 
                                               1.61 
                                               0.9 
                                               2.88 
                                               0.117 
                                               0.829 
                                               lm.nb 
                                               CD28 co-stimulation, CD28 dependent PI3KAkt signaling, Constitutive Signaling by Aberrant PI3K in Cancer, DAP12 signaling, Downstream signal transduction, G alpha (1213) signalling events, G alpha (q) signalling events, G beta.CL.gamma signalling through PI3Kgamma, GAB1 signalosome, Gastrin-CREB signalling pathway via PKC and MAPK, GPVI-mediated activation cascade, Interleukin receptor SHC signaling, PI3KAKT activation, PIP3 activates AKT signaling, Regulation of signaling by CBL, Role of LAT2NTALLAB on calcium mobilization, Signaling by ERBB2, Signaling by PDGF, Signaling by SCF-KIT, Synthesis of PIPs at the plasma membrane 
                                               SEBELIUS_2_C4636 
                                               PIK3R3 
                                               mRNA 
                                               0 
                                               
                                               
                                               G alpha (q) signalling events;PIP3 activates AKT signaling;Constitutive Signaling by Aberrant PI3K in Cancer;CD28 dependent PI3KAkt signaling;Synthesis of PIPs at the plasma membrane;Interleukin receptor SHC signaling;G beta.CL.gamma signalling through PI3Kgamma;Regulation of signaling by CBL;GPVI-mediated activation cascade;G alpha (1213) signalling events;Gastrin-CREB signalling pathway via PKC and MAPK;Downstream signal transduction;GAB1 signalosome;Signaling by SCF-KIT;PI3KAKT activation;Role of LAT2NTALLAB on calcium mobilization;DAP12 signaling;Signaling by PDGF;Signaling by ERBB2;CD28 co-stimulation 
                                               hsa04012;hsa04014;hsa04015;hsa04022;hsa04024;hsa04062;hsa04066;hsa04068;hsa04070;hsa04071;hsa04072;hsa04150;hsa04151;hsa04152;hsa04210;hsa04211;hsa04213;hsa04261;hsa04370;hsa04380;hsa04510;hsa04550;hsa04611;hsa04620;hsa04630;hsa04650;hsa04660;hsa04662;hsa04664;hsa04666;hsa04668;hsa04670;hsa04722;hsa04725;hsa04750;hsa04810;hsa04910;hsa04914;hsa04915;hsa04917;hsa04919;hsa04921;hsa04923;hsa04930;hsa04931;hsa04932;hsa04933;hsa04960;hsa04973;hsa05100;hsa05142;hsa05145;hsa05146;hsa05160;hsa05161;hsa05162;hsa05164;hsa05166;hsa05169;hsa05200;hsa05203;hsa05205;hsa05210;hsa05211;hsa05212;hsa05213;hsa05214;hsa05215;hsa05218;hsa05220;hsa05221;hsa05222;hsa05223;hsa05230;hsa05231 
                                               PIK3R3 
                                               NA 
                                               azure4 
                                               1 
                                               1.6037296037296 
                                             
                                           
                                           DE Results - CD28 dependent PI3KAkt signaling genes - Line: HT29 vs.Fibro  Table displaying each sample's global significance scores and directed global significance scores as defined in the heatmaps above. The global significance score is calculated as the square root of the mean squared t-statistic for the genes in a gene set, with t-statistics coming from the linear regression underlying our differential expression analysis. The directed global significance score is calculated as the square root of the mean signed squared t-statistic for the genes in a gene set, with t-statistics coming from the linear regression underlying our differential expression analysis.   
                                      
                                       
                                  
                                
                              
                            
                           
                            
                          
                           
                      
                    
                    
                      
                           
                          
                               Day    Line     
                              
                                  Day  
                              
                               
                              
                                  Line: differential expression in Caco2 vs. baseline of Fibro  
                                  Line: differential expression in HEPG2 vs. baseline of Fibro  
                                  Line: differential expression in HT29 vs. baseline of Fibro  
                              
                              
                            
                              
                                
                                  
                                       
                                      
                                           Volcano Plot: Day   More Plot Information                Volcano Plot: Day  Volcano plot displaying each gene's -log10(p-value) and log2 fold change for the selected covariate. Highly statistically significant genes fall at the top of the plot, and highly differentially expressed genes fall to either side. Genes within the selected gene set are highlighted in orange. Horizontal lines indicate various False Discovery Rate (FDR) thresholds.   
                                           DE Results - CD28 dependent Vav1 pathway genes - Day   More Plot Information      Download CSV Data     
                                           
                                             
                                             
                                           
                                           
                                             
                                               Probe Label 
                                               Log2 fold change 
                                               std error (log2) 
                                               Lower confidence limit (log2) 
                                               Upper confidence limit (log2) 
                                               Linear fold change 
                                               Lower confidence limit (linear) 
                                               Upper confidence limit (linear) 
                                               P-value 
                                               BY.p.value 
                                               method 
                                               Gene.sets 
                                               Codeset.Name 
                                               Probe.Label 
                                               Analyte.Type 
                                               Is.Control 
                                               Control.Type 
                                               Related.Probes 
                                               Probe.Annotation 
                                               KEGG.Pathways 
                                               Official.Gene.Name 
                                               Control.Conc 
                                               volcanocol 
                                               volcanopch 
                                               tstats.all 
                                             
                                           
                                           
                                             
                                               PAK1-mRNA 
                                               0.128 
                                               0.0211 
                                               0.0869 
                                               0.169 
                                               1.09 
                                               1.06 
                                               1.12 
                                               4.84e-07 
                                               6.25e-05 
                                               lm.nb 
                                               Activation of Rac, CD209 (DC-SIGN) signaling, CD28 co-stimulation, CD28 dependent Vav1 pathway, EPHB-mediated forward signaling, Ephrin signaling, FCERI mediated MAPK activation, MAPK6MAPK4 signaling, Regulation of actin dynamics for phagocytic cup formation, RHO GTPases activate PAKs, Sema3A PAK dependent Axon repulsion, VEGFA-VEGFR2 Pathway, VEGFR2 mediated vascular permeability 
                                               SEBELIUS_2_C4636 
                                               PAK1 
                                               mRNA 
                                               0 
                                               
                                               
                                               EPHB-mediated forward signaling;Ephrin signaling;MAPK6MAPK4 signaling;DSCAM interactions;Activation of Rac;VEGFR2 mediated vascular permeability;RHO GTPases activate PAKs;Sema3A PAK dependent Axon repulsion;FCERI mediated MAPK activation;Regulation of actin dynamics for phagocytic cup formation;CD209 (DC-SIGN) signaling;CD28 dependent Vav1 pathway;VEGFA-VEGFR2 Pathway;CD28 co-stimulation 
                                               hsa04010;hsa04012;hsa04014;hsa04024;hsa04062;hsa04360;hsa04510;hsa04650;hsa04660;hsa04666;hsa04810;hsa05120;hsa05205;hsa05211 
                                               PAK1 
                                               NA 
                                               slateblue1 
                                               16 
                                               6.06635071090047 
                                             
                                             
                                               PAK2-mRNA 
                                               -0.0398 
                                               0.00994 
                                               -0.0593 
                                               -0.0203 
                                               0.973 
                                               0.96 
                                               0.986 
                                               0.000289 
                                               0.00836 
                                               loglinear 
                                               Activation of Rac, CD209 (DC-SIGN) signaling, CD28 co-stimulation, CD28 dependent Vav1 pathway, Ephrin signaling, RHO GTPases activate PAKs, Sema3A PAK dependent Axon repulsion, VEGFA-VEGFR2 Pathway, VEGFR2 mediated vascular permeability 
                                               SEBELIUS_2_C4636 
                                               PAK2 
                                               mRNA 
                                               0 
                                               
                                               
                                               Nef and signal transduction;CD28 dependent Vav1 pathway;Stimulation of the cell death response by PAK-2p34;Ephrin signaling;Activation of Rac;RHO GTPases activate PAKs;VEGFR2 mediated vascular permeability;Sema3A PAK dependent Axon repulsion;CD209 (DC-SIGN) signaling;VEGFA-VEGFR2 Pathway;CD28 co-stimulation 
                                               hsa04010;hsa04012;hsa04014;hsa04360;hsa04510;hsa04660;hsa04810;hsa05211 
                                               PAK2 
                                               NA 
                                               slateblue1 
                                               16 
                                               -4.00402414486922 
                                             
                                             
                                               CDC42-mRNA 
                                               -0.0218 
                                               0.0128 
                                               -0.0469 
                                               0.00331 
                                               0.985 
                                               0.968 
                                               1 
                                               0.0972 
                                               0.92 
                                               lm.nb 
                                               CD28 co-stimulation, CD28 dependent Vav1 pathway, DCC mediated attractive signaling, EPHB-mediated forward signaling, G alpha (1213) signalling events, GPVI-mediated activation cascade, Regulation of actin dynamics for phagocytic cup formation, Rho GTPase cycle, RHO GTPases Activate Formins, RHO GTPases activate IQGAPs, RHO GTPases activate PAKs, Sema4D induced cell migration and growth-cone collapse, VEGFA-VEGFR2 Pathway 
                                               SEBELIUS_2_C4636 
                                               CDC42 
                                               mRNA 
                                               0 
                                               
                                               
                                               DCC mediated attractive signaling;Factors involved in megakaryocyte development and platelet production;VEGFA-VEGFR2 Pathway;RHO GTPases activate KTN1;CD28 dependent Vav1 pathway;Inactivation of Cdc42 and Rac;Rho GTPase cycle;EPHB-mediated forward signaling;RHO GTPases Activate Formins;RHO GTPases activate PAKs;GPVI-mediated activation cascade;G alpha (1213) signalling events;RHO GTPases activate IQGAPs;Sema4D induced cell migration and growth-cone collapse;Regulation of actin dynamics for phagocytic cup formation;CD28 co-stimulation 
                                               hsa04010;hsa04014;hsa04015;hsa04062;hsa04144;hsa04360;hsa04370;hsa04510;hsa04520;hsa04530;hsa04660;hsa04666;hsa04670;hsa04722;hsa04810;hsa04912;hsa04932;hsa04933;hsa05100;hsa05120;hsa05130;hsa05131;hsa05132;hsa05200;hsa05203;hsa05205;hsa05211;hsa05212 
                                               CDC42 
                                               NA 
                                               azure4 
                                               1 
                                               -1.703125 
                                             
                                             
                                               FYN-mRNA 
                                               0.0103 
                                               0.0454 
                                               -0.0786 
                                               0.0993 
                                               1.01 
                                               0.947 
                                               1.07 
                                               0.821 
                                               1 
                                               lm.nb 
                                               ARMS-mediated activation, CD209 (DC-SIGN) signaling, CD28 co-stimulation, CD28 dependent PI3KAkt signaling, CD28 dependent Vav1 pathway, Cell surface interactions at the vascular wall, Constitutive Signaling by Aberrant PI3K in Cancer, CTLA4 inhibitory signaling, DAP12 signaling, Downstream signal transduction, EPH-ephrin mediated repulsion of cells, FCERI mediated MAPK activation, Frs2-mediated activation, GAB1 signalosome, Gastrin-CREB signalling pathway via PKC and MAPK, GPVI-mediated activation cascade, Interleukin receptor SHC signaling, IRS-related events triggered by IGF1R, Nephrin interactions, PI3KAKT activation, RAFMAP kinase cascade, Regulation of signaling by CBL, Role of LAT2NTALLAB on calcium mobilization, Sema3A PAK dependent Axon repulsion, Signaling by ERBB2, Signaling by PDGF, Signaling by SCF-KIT, VEGFA-VEGFR2 Pathway, VEGFR2 mediated cell proliferation 
                                               SEBELIUS_2_C4636 
                                               FYN 
                                               mRNA 
                                               0 
                                               
                                               
                                               EPH-ephrin mediated repulsion of cells;Regulation of signaling by CBL;FCGR activation;Antigen activates B Cell Receptor (BCR) leading to generation of second messengers;Dectin-2 family;RAFMAP kinase cascade;Constitutive Signaling by Aberrant PI3K in Cancer;CTLA4 inhibitory signaling;CD28 dependent PI3KAkt signaling;PECAM1 interactions;CD28 dependent Vav1 pathway;GPVI-mediated activation cascade;Sema3A PAK dependent Axon repulsion;Nephrin interactions;SEMA3A-Plexin repulsion signaling by inhibiting Integrin adhesion;CRMPs in Sema3A signaling;CD209 (DC-SIGN) signaling;Downstream signal transduction;GAB1 signalosome;Signaling by SCF-KIT;PI3KAKT activation;Role of LAT2NTALLAB on calcium mobilization;DAP12 signaling;Signaling by PDGF;Signaling by ERBB2;VEGFA-VEGFR2 Pathway;VEGFR2 mediated cell proliferation;Frs2-mediated activation;ARMS-mediated activation;Gastrin-CREB signalling pathway via PKC and MAPK;Interleukin receptor SHC signaling;FCERI mediated MAPK activation;IRS-related events triggered by IGF1R;CD28 co-stimulation;Cell surface interactions at the vascular wall 
                                               hsa04071;hsa04072;hsa04360;hsa04380;hsa04510;hsa04520;hsa04611;hsa04650;hsa04660;hsa04664;hsa04725;hsa05020;hsa05130;hsa05162;hsa05416 
                                               FYN 
                                               NA 
                                               azure4 
                                               1 
                                               0.226872246696035 
                                             
                                             
                                               RAC1-mRNA 
                                               -0.000703 
                                               0.0123 
                                               -0.0248 
                                               0.0234 
                                               1 
                                               0.983 
                                               1.02 
                                               0.955 
                                               1 
                                               lm.nb 
                                               Activation of Rac, CD28 co-stimulation, CD28 dependent Vav1 pathway, DAP12 signaling, EPH-ephrin mediated repulsion of cells, Ephrin signaling, G alpha (1213) signalling events, GPVI-mediated activation cascade, MAPK6MAPK4 signaling, NRAGE signals death through JNK, PCPCE pathway, Regulation of actin dynamics for phagocytic cup formation, RHO GTPases activate CIT, RHO GTPases Activate Formins, RHO GTPases activate IQGAPs, RHO GTPases activate PAKs, RHO GTPases activate PKNs, RHO GTPases Activate WASPs and WAVEs, Sema3A PAK dependent Axon repulsion, Sema4D induced cell migration and growth-cone collapse, Signal transduction by L1, Signaling by SCF-KIT, VEGFA-VEGFR2 Pathway, VEGFR2 mediated vascular permeability 
                                               SEBELIUS_2_C4636 
                                               RAC1 
                                               mRNA 
                                               0 
                                               
                                               
                                               MAPK6MAPK4 signaling;Signaling by SCF-KIT;Inactivation of Cdc42 and Rac;DAP12 signaling;Ephrin signaling;EPH-ephrin mediated repulsion of cells;Activation of Rac;RHO GTPases Activate Formins;VEGFR2 mediated vascular permeability;RHO GTPases activate PAKs;Sema4D mediated inhibition of cell attachment and migration;G alpha (1213) signalling events;Regulation of actin dynamics for phagocytic cup formation;Sema4D induced cell migration and growth-cone collapse;Signal transduction by L1;RHO GTPases activate CIT;RHO GTPases Activate WASPs and WAVEs;Factors involved in megakaryocyte development and platelet production;RHO GTPases activate KTN1;CD28 dependent Vav1 pathway;DSCAM interactions;NRAGE signals death through JNK;GPVI-mediated activation cascade;Sema3A PAK dependent Axon repulsion;RHO GTPases activate PKNs;SEMA3A-Plexin repulsion signaling by inhibiting Integrin adhesion;RHO GTPases activate IQGAPs;PCPCE pathway;VEGFA-VEGFR2 Pathway;CD28 co-stimulation 
                                               hsa04010;hsa04014;hsa04015;hsa04024;hsa04062;hsa04071;hsa04145;hsa04151;hsa04310;hsa04360;hsa04370;hsa04380;hsa04510;hsa04520;hsa04620;hsa04650;hsa04662;hsa04664;hsa04666;hsa04670;hsa04722;hsa04810;hsa04932;hsa04933;hsa04972;hsa05100;hsa05120;hsa05131;hsa05132;hsa05200;hsa05203;hsa05205;hsa05210;hsa05211;hsa05212;hsa05231;hsa05416 
                                               RAC1 
                                               NA 
                                               azure4 
                                               1 
                                               -0.0571544715447154 
                                             
                                           
                                           DE Results - CD28 dependent Vav1 pathway genes - Day  Table displaying each sample's global significance scores and directed global significance scores as defined in the heatmaps above. The global significance score is calculated as the square root of the mean squared t-statistic for the genes in a gene set, with t-statistics coming from the linear regression underlying our differential expression analysis. The directed global significance score is calculated as the square root of the mean signed squared t-statistic for the genes in a gene set, with t-statistics coming from the linear regression underlying our differential expression analysis.   
                                      
                                       
                                  
                                
                              
                            
                            
                              
                                
                                  
                                       
                                      
                                           Volcano Plot: Line: Caco2 vs.Fibro   More Plot Information                Volcano Plot: Line: Caco2 vs.Fibro  Volcano plot displaying each gene's -log10(p-value) and log2 fold change for the selected covariate. Highly statistically significant genes fall at the top of the plot, and highly differentially expressed genes fall to either side. Genes within the selected gene set are highlighted in orange. Horizontal lines indicate various False Discovery Rate (FDR) thresholds.   
                                           DE Results - CD28 dependent Vav1 pathway genes - Line: Caco2 vs.Fibro   More Plot Information      Download CSV Data     
                                           
                                             
                                             
                                           
                                           
                                             
                                               Probe Label 
                                               Log2 fold change 
                                               std error (log2) 
                                               Lower confidence limit (log2) 
                                               Upper confidence limit (log2) 
                                               Linear fold change 
                                               Lower confidence limit (linear) 
                                               Upper confidence limit (linear) 
                                               P-value 
                                               BY.p.value 
                                               method 
                                               Gene.sets 
                                               Codeset.Name 
                                               Probe.Label 
                                               Analyte.Type 
                                               Is.Control 
                                               Control.Type 
                                               Related.Probes 
                                               Probe.Annotation 
                                               KEGG.Pathways 
                                               Official.Gene.Name 
                                               Control.Conc 
                                               volcanocol 
                                               volcanopch 
                                               tstats.all 
                                             
                                           
                                           
                                             
                                               PAK1-mRNA 
                                               1.92 
                                               0.159 
                                               1.61 
                                               2.23 
                                               3.79 
                                               3.05 
                                               4.7 
                                               2.23e-14 
                                               5.02e-13 
                                               lm.nb 
                                               Activation of Rac, CD209 (DC-SIGN) signaling, CD28 co-stimulation, CD28 dependent Vav1 pathway, EPHB-mediated forward signaling, Ephrin signaling, FCERI mediated MAPK activation, MAPK6MAPK4 signaling, Regulation of actin dynamics for phagocytic cup formation, RHO GTPases activate PAKs, Sema3A PAK dependent Axon repulsion, VEGFA-VEGFR2 Pathway, VEGFR2 mediated vascular permeability 
                                               SEBELIUS_2_C4636 
                                               PAK1 
                                               mRNA 
                                               0 
                                               
                                               
                                               EPHB-mediated forward signaling;Ephrin signaling;MAPK6MAPK4 signaling;DSCAM interactions;Activation of Rac;VEGFR2 mediated vascular permeability;RHO GTPases activate PAKs;Sema3A PAK dependent Axon repulsion;FCERI mediated MAPK activation;Regulation of actin dynamics for phagocytic cup formation;CD209 (DC-SIGN) signaling;CD28 dependent Vav1 pathway;VEGFA-VEGFR2 Pathway;CD28 co-stimulation 
                                               hsa04010;hsa04012;hsa04014;hsa04024;hsa04062;hsa04360;hsa04510;hsa04650;hsa04660;hsa04666;hsa04810;hsa05120;hsa05205;hsa05211 
                                               PAK1 
                                               NA 
                                               slateblue1 
                                               16 
                                               12.0754716981132 
                                             
                                             
                                               FYN-mRNA 
                                               -2.64 
                                               0.335 
                                               -3.3 
                                               -1.99 
                                               0.16 
                                               0.102 
                                               0.253 
                                               1.96e-09 
                                               2.84e-08 
                                               lm.nb 
                                               ARMS-mediated activation, CD209 (DC-SIGN) signaling, CD28 co-stimulation, CD28 dependent PI3KAkt signaling, CD28 dependent Vav1 pathway, Cell surface interactions at the vascular wall, Constitutive Signaling by Aberrant PI3K in Cancer, CTLA4 inhibitory signaling, DAP12 signaling, Downstream signal transduction, EPH-ephrin mediated repulsion of cells, FCERI mediated MAPK activation, Frs2-mediated activation, GAB1 signalosome, Gastrin-CREB signalling pathway via PKC and MAPK, GPVI-mediated activation cascade, Interleukin receptor SHC signaling, IRS-related events triggered by IGF1R, Nephrin interactions, PI3KAKT activation, RAFMAP kinase cascade, Regulation of signaling by CBL, Role of LAT2NTALLAB on calcium mobilization, Sema3A PAK dependent Axon repulsion, Signaling by ERBB2, Signaling by PDGF, Signaling by SCF-KIT, VEGFA-VEGFR2 Pathway, VEGFR2 mediated cell proliferation 
                                               SEBELIUS_2_C4636 
                                               FYN 
                                               mRNA 
                                               0 
                                               
                                               
                                               EPH-ephrin mediated repulsion of cells;Regulation of signaling by CBL;FCGR activation;Antigen activates B Cell Receptor (BCR) leading to generation of second messengers;Dectin-2 family;RAFMAP kinase cascade;Constitutive Signaling by Aberrant PI3K in Cancer;CTLA4 inhibitory signaling;CD28 dependent PI3KAkt signaling;PECAM1 interactions;CD28 dependent Vav1 pathway;GPVI-mediated activation cascade;Sema3A PAK dependent Axon repulsion;Nephrin interactions;SEMA3A-Plexin repulsion signaling by inhibiting Integrin adhesion;CRMPs in Sema3A signaling;CD209 (DC-SIGN) signaling;Downstream signal transduction;GAB1 signalosome;Signaling by SCF-KIT;PI3KAKT activation;Role of LAT2NTALLAB on calcium mobilization;DAP12 signaling;Signaling by PDGF;Signaling by ERBB2;VEGFA-VEGFR2 Pathway;VEGFR2 mediated cell proliferation;Frs2-mediated activation;ARMS-mediated activation;Gastrin-CREB signalling pathway via PKC and MAPK;Interleukin receptor SHC signaling;FCERI mediated MAPK activation;IRS-related events triggered by IGF1R;CD28 co-stimulation;Cell surface interactions at the vascular wall 
                                               hsa04071;hsa04072;hsa04360;hsa04380;hsa04510;hsa04520;hsa04611;hsa04650;hsa04660;hsa04664;hsa04725;hsa05020;hsa05130;hsa05162;hsa05416 
                                               FYN 
                                               NA 
                                               slateblue1 
                                               16 
                                               -7.88059701492537 
                                             
                                             
                                               CDC42-mRNA 
                                               -0.361 
                                               0.0968 
                                               -0.551 
                                               -0.171 
                                               0.779 
                                               0.683 
                                               0.888 
                                               0.00064 
                                               0.00571 
                                               lm.nb 
                                               CD28 co-stimulation, CD28 dependent Vav1 pathway, DCC mediated attractive signaling, EPHB-mediated forward signaling, G alpha (1213) signalling events, GPVI-mediated activation cascade, Regulation of actin dynamics for phagocytic cup formation, Rho GTPase cycle, RHO GTPases Activate Formins, RHO GTPases activate IQGAPs, RHO GTPases activate PAKs, Sema4D induced cell migration and growth-cone collapse, VEGFA-VEGFR2 Pathway 
                                               SEBELIUS_2_C4636 
                                               CDC42 
                                               mRNA 
                                               0 
                                               
                                               
                                               DCC mediated attractive signaling;Factors involved in megakaryocyte development and platelet production;VEGFA-VEGFR2 Pathway;RHO GTPases activate KTN1;CD28 dependent Vav1 pathway;Inactivation of Cdc42 and Rac;Rho GTPase cycle;EPHB-mediated forward signaling;RHO GTPases Activate Formins;RHO GTPases activate PAKs;GPVI-mediated activation cascade;G alpha (1213) signalling events;RHO GTPases activate IQGAPs;Sema4D induced cell migration and growth-cone collapse;Regulation of actin dynamics for phagocytic cup formation;CD28 co-stimulation 
                                               hsa04010;hsa04014;hsa04015;hsa04062;hsa04144;hsa04360;hsa04370;hsa04510;hsa04520;hsa04530;hsa04660;hsa04666;hsa04670;hsa04722;hsa04810;hsa04912;hsa04932;hsa04933;hsa05100;hsa05120;hsa05130;hsa05131;hsa05132;hsa05200;hsa05203;hsa05205;hsa05211;hsa05212 
                                               CDC42 
                                               NA 
                                               slateblue1 
                                               16 
                                               -3.72933884297521 
                                             
                                             
                                               RAC1-mRNA 
                                               -0.326 
                                               0.093 
                                               -0.508 
                                               -0.144 
                                               0.798 
                                               0.703 
                                               0.905 
                                               0.00121 
                                               0.0104 
                                               lm.nb 
                                               Activation of Rac, CD28 co-stimulation, CD28 dependent Vav1 pathway, DAP12 signaling, EPH-ephrin mediated repulsion of cells, Ephrin signaling, G alpha (1213) signalling events, GPVI-mediated activation cascade, MAPK6MAPK4 signaling, NRAGE signals death through JNK, PCPCE pathway, Regulation of actin dynamics for phagocytic cup formation, RHO GTPases activate CIT, RHO GTPases Activate Formins, RHO GTPases activate IQGAPs, RHO GTPases activate PAKs, RHO GTPases activate PKNs, RHO GTPases Activate WASPs and WAVEs, Sema3A PAK dependent Axon repulsion, Sema4D induced cell migration and growth-cone collapse, Signal transduction by L1, Signaling by SCF-KIT, VEGFA-VEGFR2 Pathway, VEGFR2 mediated vascular permeability 
                                               SEBELIUS_2_C4636 
                                               RAC1 
                                               mRNA 
                                               0 
                                               
                                               
                                               MAPK6MAPK4 signaling;Signaling by SCF-KIT;Inactivation of Cdc42 and Rac;DAP12 signaling;Ephrin signaling;EPH-ephrin mediated repulsion of cells;Activation of Rac;RHO GTPases Activate Formins;VEGFR2 mediated vascular permeability;RHO GTPases activate PAKs;Sema4D mediated inhibition of cell attachment and migration;G alpha (1213) signalling events;Regulation of actin dynamics for phagocytic cup formation;Sema4D induced cell migration and growth-cone collapse;Signal transduction by L1;RHO GTPases activate CIT;RHO GTPases Activate WASPs and WAVEs;Factors involved in megakaryocyte development and platelet production;RHO GTPases activate KTN1;CD28 dependent Vav1 pathway;DSCAM interactions;NRAGE signals death through JNK;GPVI-mediated activation cascade;Sema3A PAK dependent Axon repulsion;RHO GTPases activate PKNs;SEMA3A-Plexin repulsion signaling by inhibiting Integrin adhesion;RHO GTPases activate IQGAPs;PCPCE pathway;VEGFA-VEGFR2 Pathway;CD28 co-stimulation 
                                               hsa04010;hsa04014;hsa04015;hsa04024;hsa04062;hsa04071;hsa04145;hsa04151;hsa04310;hsa04360;hsa04370;hsa04380;hsa04510;hsa04520;hsa04620;hsa04650;hsa04662;hsa04664;hsa04666;hsa04670;hsa04722;hsa04810;hsa04932;hsa04933;hsa04972;hsa05100;hsa05120;hsa05131;hsa05132;hsa05200;hsa05203;hsa05205;hsa05210;hsa05211;hsa05212;hsa05231;hsa05416 
                                               RAC1 
                                               NA 
                                               slateblue2 
                                               16 
                                               -3.50537634408602 
                                             
                                             
                                               PAK2-mRNA 
                                               -0.138 
                                               0.0752 
                                               -0.285 
                                               0.00968 
                                               0.909 
                                               0.821 
                                               1.01 
                                               0.0751 
                                               0.552 
                                               loglinear 
                                               Activation of Rac, CD209 (DC-SIGN) signaling, CD28 co-stimulation, CD28 dependent Vav1 pathway, Ephrin signaling, RHO GTPases activate PAKs, Sema3A PAK dependent Axon repulsion, VEGFA-VEGFR2 Pathway, VEGFR2 mediated vascular permeability 
                                               SEBELIUS_2_C4636 
                                               PAK2 
                                               mRNA 
                                               0 
                                               
                                               
                                               Nef and signal transduction;CD28 dependent Vav1 pathway;Stimulation of the cell death response by PAK-2p34;Ephrin signaling;Activation of Rac;RHO GTPases activate PAKs;VEGFR2 mediated vascular permeability;Sema3A PAK dependent Axon repulsion;CD209 (DC-SIGN) signaling;VEGFA-VEGFR2 Pathway;CD28 co-stimulation 
                                               hsa04010;hsa04012;hsa04014;hsa04360;hsa04510;hsa04660;hsa04810;hsa05211 
                                               PAK2 
                                               NA 
                                               azure4 
                                               1 
                                               -1.83510638297872 
                                             
                                           
                                           DE Results - CD28 dependent Vav1 pathway genes - Line: Caco2 vs.Fibro  Table displaying each sample's global significance scores and directed global significance scores as defined in the heatmaps above. The global significance score is calculated as the square root of the mean squared t-statistic for the genes in a gene set, with t-statistics coming from the linear regression underlying our differential expression analysis. The directed global significance score is calculated as the square root of the mean signed squared t-statistic for the genes in a gene set, with t-statistics coming from the linear regression underlying our differential expression analysis.   
                                      
                                       
                                  
                                
                                
                                  
                                       
                                      
                                           Volcano Plot: Line: HEPG2 vs.Fibro   More Plot Information                Volcano Plot: Line: HEPG2 vs.Fibro  Volcano plot displaying each gene's -log10(p-value) and log2 fold change for the selected covariate. Highly statistically significant genes fall at the top of the plot, and highly differentially expressed genes fall to either side. Genes within the selected gene set are highlighted in orange. Horizontal lines indicate various False Discovery Rate (FDR) thresholds.   
                                           DE Results - CD28 dependent Vav1 pathway genes - Line: HEPG2 vs.Fibro   More Plot Information      Download CSV Data     
                                           
                                             
                                             
                                           
                                           
                                             
                                               Probe Label 
                                               Log2 fold change 
                                               std error (log2) 
                                               Lower confidence limit (log2) 
                                               Upper confidence limit (log2) 
                                               Linear fold change 
                                               Lower confidence limit (linear) 
                                               Upper confidence limit (linear) 
                                               P-value 
                                               BY.p.value 
                                               method 
                                               Gene.sets 
                                               Codeset.Name 
                                               Probe.Label 
                                               Analyte.Type 
                                               Is.Control 
                                               Control.Type 
                                               Related.Probes 
                                               Probe.Annotation 
                                               KEGG.Pathways 
                                               Official.Gene.Name 
                                               Control.Conc 
                                               volcanocol 
                                               volcanopch 
                                               tstats.all 
                                             
                                           
                                           
                                             
                                               PAK1-mRNA 
                                               -2.44 
                                               0.179 
                                               -2.79 
                                               -2.09 
                                               0.184 
                                               0.145 
                                               0.235 
                                               5.34e-16 
                                               1.72e-14 
                                               lm.nb 
                                               Activation of Rac, CD209 (DC-SIGN) signaling, CD28 co-stimulation, CD28 dependent Vav1 pathway, EPHB-mediated forward signaling, Ephrin signaling, FCERI mediated MAPK activation, MAPK6MAPK4 signaling, Regulation of actin dynamics for phagocytic cup formation, RHO GTPases activate PAKs, Sema3A PAK dependent Axon repulsion, VEGFA-VEGFR2 Pathway, VEGFR2 mediated vascular permeability 
                                               SEBELIUS_2_C4636 
                                               PAK1 
                                               mRNA 
                                               0 
                                               
                                               
                                               EPHB-mediated forward signaling;Ephrin signaling;MAPK6MAPK4 signaling;DSCAM interactions;Activation of Rac;VEGFR2 mediated vascular permeability;RHO GTPases activate PAKs;Sema3A PAK dependent Axon repulsion;FCERI mediated MAPK activation;Regulation of actin dynamics for phagocytic cup formation;CD209 (DC-SIGN) signaling;CD28 dependent Vav1 pathway;VEGFA-VEGFR2 Pathway;CD28 co-stimulation 
                                               hsa04010;hsa04012;hsa04014;hsa04024;hsa04062;hsa04360;hsa04510;hsa04650;hsa04660;hsa04666;hsa04810;hsa05120;hsa05205;hsa05211 
                                               PAK1 
                                               NA 
                                               slateblue1 
                                               16 
                                               -13.6312849162011 
                                             
                                             
                                               PAK2-mRNA 
                                               0.446 
                                               0.0825 
                                               0.284 
                                               0.608 
                                               1.36 
                                               1.22 
                                               1.52 
                                               4.03e-06 
                                               4.65e-05 
                                               loglinear 
                                               Activation of Rac, CD209 (DC-SIGN) signaling, CD28 co-stimulation, CD28 dependent Vav1 pathway, Ephrin signaling, RHO GTPases activate PAKs, Sema3A PAK dependent Axon repulsion, VEGFA-VEGFR2 Pathway, VEGFR2 mediated vascular permeability 
                                               SEBELIUS_2_C4636 
                                               PAK2 
                                               mRNA 
                                               0 
                                               
                                               
                                               Nef and signal transduction;CD28 dependent Vav1 pathway;Stimulation of the cell death response by PAK-2p34;Ephrin signaling;Activation of Rac;RHO GTPases activate PAKs;VEGFR2 mediated vascular permeability;Sema3A PAK dependent Axon repulsion;CD209 (DC-SIGN) signaling;VEGFA-VEGFR2 Pathway;CD28 co-stimulation 
                                               hsa04010;hsa04012;hsa04014;hsa04360;hsa04510;hsa04660;hsa04810;hsa05211 
                                               PAK2 
                                               NA 
                                               slateblue1 
                                               16 
                                               5.40606060606061 
                                             
                                             
                                               FYN-mRNA 
                                               0.596 
                                               0.365 
                                               -0.119 
                                               1.31 
                                               1.51 
                                               0.921 
                                               2.48 
                                               0.111 
                                               0.804 
                                               lm.nb 
                                               ARMS-mediated activation, CD209 (DC-SIGN) signaling, CD28 co-stimulation, CD28 dependent PI3KAkt signaling, CD28 dependent Vav1 pathway, Cell surface interactions at the vascular wall, Constitutive Signaling by Aberrant PI3K in Cancer, CTLA4 inhibitory signaling, DAP12 signaling, Downstream signal transduction, EPH-ephrin mediated repulsion of cells, FCERI mediated MAPK activation, Frs2-mediated activation, GAB1 signalosome, Gastrin-CREB signalling pathway via PKC and MAPK, GPVI-mediated activation cascade, Interleukin receptor SHC signaling, IRS-related events triggered by IGF1R, Nephrin interactions, PI3KAKT activation, RAFMAP kinase cascade, Regulation of signaling by CBL, Role of LAT2NTALLAB on calcium mobilization, Sema3A PAK dependent Axon repulsion, Signaling by ERBB2, Signaling by PDGF, Signaling by SCF-KIT, VEGFA-VEGFR2 Pathway, VEGFR2 mediated cell proliferation 
                                               SEBELIUS_2_C4636 
                                               FYN 
                                               mRNA 
                                               0 
                                               
                                               
                                               EPH-ephrin mediated repulsion of cells;Regulation of signaling by CBL;FCGR activation;Antigen activates B Cell Receptor (BCR) leading to generation of second messengers;Dectin-2 family;RAFMAP kinase cascade;Constitutive Signaling by Aberrant PI3K in Cancer;CTLA4 inhibitory signaling;CD28 dependent PI3KAkt signaling;PECAM1 interactions;CD28 dependent Vav1 pathway;GPVI-mediated activation cascade;Sema3A PAK dependent Axon repulsion;Nephrin interactions;SEMA3A-Plexin repulsion signaling by inhibiting Integrin adhesion;CRMPs in Sema3A signaling;CD209 (DC-SIGN) signaling;Downstream signal transduction;GAB1 signalosome;Signaling by SCF-KIT;PI3KAKT activation;Role of LAT2NTALLAB on calcium mobilization;DAP12 signaling;Signaling by PDGF;Signaling by ERBB2;VEGFA-VEGFR2 Pathway;VEGFR2 mediated cell proliferation;Frs2-mediated activation;ARMS-mediated activation;Gastrin-CREB signalling pathway via PKC and MAPK;Interleukin receptor SHC signaling;FCERI mediated MAPK activation;IRS-related events triggered by IGF1R;CD28 co-stimulation;Cell surface interactions at the vascular wall 
                                               hsa04071;hsa04072;hsa04360;hsa04380;hsa04510;hsa04520;hsa04611;hsa04650;hsa04660;hsa04664;hsa04725;hsa05020;hsa05130;hsa05162;hsa05416 
                                               FYN 
                                               NA 
                                               azure4 
                                               1 
                                               1.63287671232877 
                                             
                                             
                                               CDC42-mRNA 
                                               -0.164 
                                               0.106 
                                               -0.372 
                                               0.0443 
                                               0.893 
                                               0.773 
                                               1.03 
                                               0.131 
                                               0.941 
                                               lm.nb 
                                               CD28 co-stimulation, CD28 dependent Vav1 pathway, DCC mediated attractive signaling, EPHB-mediated forward signaling, G alpha (1213) signalling events, GPVI-mediated activation cascade, Regulation of actin dynamics for phagocytic cup formation, Rho GTPase cycle, RHO GTPases Activate Formins, RHO GTPases activate IQGAPs, RHO GTPases activate PAKs, Sema4D induced cell migration and growth-cone collapse, VEGFA-VEGFR2 Pathway 
                                               SEBELIUS_2_C4636 
                                               CDC42 
                                               mRNA 
                                               0 
                                               
                                               
                                               DCC mediated attractive signaling;Factors involved in megakaryocyte development and platelet production;VEGFA-VEGFR2 Pathway;RHO GTPases activate KTN1;CD28 dependent Vav1 pathway;Inactivation of Cdc42 and Rac;Rho GTPase cycle;EPHB-mediated forward signaling;RHO GTPases Activate Formins;RHO GTPases activate PAKs;GPVI-mediated activation cascade;G alpha (1213) signalling events;RHO GTPases activate IQGAPs;Sema4D induced cell migration and growth-cone collapse;Regulation of actin dynamics for phagocytic cup formation;CD28 co-stimulation 
                                               hsa04010;hsa04014;hsa04015;hsa04062;hsa04144;hsa04360;hsa04370;hsa04510;hsa04520;hsa04530;hsa04660;hsa04666;hsa04670;hsa04722;hsa04810;hsa04912;hsa04932;hsa04933;hsa05100;hsa05120;hsa05130;hsa05131;hsa05132;hsa05200;hsa05203;hsa05205;hsa05211;hsa05212 
                                               CDC42 
                                               NA 
                                               azure4 
                                               1 
                                               -1.54716981132075 
                                             
                                             
                                               RAC1-mRNA 
                                               -0.0792 
                                               0.102 
                                               -0.279 
                                               0.121 
                                               0.947 
                                               0.824 
                                               1.09 
                                               0.442 
                                               1 
                                               lm.nb 
                                               Activation of Rac, CD28 co-stimulation, CD28 dependent Vav1 pathway, DAP12 signaling, EPH-ephrin mediated repulsion of cells, Ephrin signaling, G alpha (1213) signalling events, GPVI-mediated activation cascade, MAPK6MAPK4 signaling, NRAGE signals death through JNK, PCPCE pathway, Regulation of actin dynamics for phagocytic cup formation, RHO GTPases activate CIT, RHO GTPases Activate Formins, RHO GTPases activate IQGAPs, RHO GTPases activate PAKs, RHO GTPases activate PKNs, RHO GTPases Activate WASPs and WAVEs, Sema3A PAK dependent Axon repulsion, Sema4D induced cell migration and growth-cone collapse, Signal transduction by L1, Signaling by SCF-KIT, VEGFA-VEGFR2 Pathway, VEGFR2 mediated vascular permeability 
                                               SEBELIUS_2_C4636 
                                               RAC1 
                                               mRNA 
                                               0 
                                               
                                               
                                               MAPK6MAPK4 signaling;Signaling by SCF-KIT;Inactivation of Cdc42 and Rac;DAP12 signaling;Ephrin signaling;EPH-ephrin mediated repulsion of cells;Activation of Rac;RHO GTPases Activate Formins;VEGFR2 mediated vascular permeability;RHO GTPases activate PAKs;Sema4D mediated inhibition of cell attachment and migration;G alpha (1213) signalling events;Regulation of actin dynamics for phagocytic cup formation;Sema4D induced cell migration and growth-cone collapse;Signal transduction by L1;RHO GTPases activate CIT;RHO GTPases Activate WASPs and WAVEs;Factors involved in megakaryocyte development and platelet production;RHO GTPases activate KTN1;CD28 dependent Vav1 pathway;DSCAM interactions;NRAGE signals death through JNK;GPVI-mediated activation cascade;Sema3A PAK dependent Axon repulsion;RHO GTPases activate PKNs;SEMA3A-Plexin repulsion signaling by inhibiting Integrin adhesion;RHO GTPases activate IQGAPs;PCPCE pathway;VEGFA-VEGFR2 Pathway;CD28 co-stimulation 
                                               hsa04010;hsa04014;hsa04015;hsa04024;hsa04062;hsa04071;hsa04145;hsa04151;hsa04310;hsa04360;hsa04370;hsa04380;hsa04510;hsa04520;hsa04620;hsa04650;hsa04662;hsa04664;hsa04666;hsa04670;hsa04722;hsa04810;hsa04932;hsa04933;hsa04972;hsa05100;hsa05120;hsa05131;hsa05132;hsa05200;hsa05203;hsa05205;hsa05210;hsa05211;hsa05212;hsa05231;hsa05416 
                                               RAC1 
                                               NA 
                                               azure4 
                                               1 
                                               -0.776470588235294 
                                             
                                           
                                           DE Results - CD28 dependent Vav1 pathway genes - Line: HEPG2 vs.Fibro  Table displaying each sample's global significance scores and directed global significance scores as defined in the heatmaps above. The global significance score is calculated as the square root of the mean squared t-statistic for the genes in a gene set, with t-statistics coming from the linear regression underlying our differential expression analysis. The directed global significance score is calculated as the square root of the mean signed squared t-statistic for the genes in a gene set, with t-statistics coming from the linear regression underlying our differential expression analysis.   
                                      
                                       
                                  
                                
                                
                                  
                                       
                                      
                                           Volcano Plot: Line: HT29 vs.Fibro   More Plot Information                Volcano Plot: Line: HT29 vs.Fibro  Volcano plot displaying each gene's -log10(p-value) and log2 fold change for the selected covariate. Highly statistically significant genes fall at the top of the plot, and highly differentially expressed genes fall to either side. Genes within the selected gene set are highlighted in orange. Horizontal lines indicate various False Discovery Rate (FDR) thresholds.   
                                           DE Results - CD28 dependent Vav1 pathway genes - Line: HT29 vs.Fibro   More Plot Information      Download CSV Data     
                                           
                                             
                                             
                                           
                                           
                                             
                                               Probe Label 
                                               Log2 fold change 
                                               std error (log2) 
                                               Lower confidence limit (log2) 
                                               Upper confidence limit (log2) 
                                               Linear fold change 
                                               Lower confidence limit (linear) 
                                               Upper confidence limit (linear) 
                                               P-value 
                                               BY.p.value 
                                               method 
                                               Gene.sets 
                                               Codeset.Name 
                                               Probe.Label 
                                               Analyte.Type 
                                               Is.Control 
                                               Control.Type 
                                               Related.Probes 
                                               Probe.Annotation 
                                               KEGG.Pathways 
                                               Official.Gene.Name 
                                               Control.Conc 
                                               volcanocol 
                                               volcanopch 
                                               tstats.all 
                                             
                                           
                                           
                                             
                                               PAK1-mRNA 
                                               2.31 
                                               0.165 
                                               1.99 
                                               2.63 
                                               4.96 
                                               3.96 
                                               6.2 
                                               2.18e-16 
                                               5.51e-15 
                                               lm.nb 
                                               Activation of Rac, CD209 (DC-SIGN) signaling, CD28 co-stimulation, CD28 dependent Vav1 pathway, EPHB-mediated forward signaling, Ephrin signaling, FCERI mediated MAPK activation, MAPK6MAPK4 signaling, Regulation of actin dynamics for phagocytic cup formation, RHO GTPases activate PAKs, Sema3A PAK dependent Axon repulsion, VEGFA-VEGFR2 Pathway, VEGFR2 mediated vascular permeability 
                                               SEBELIUS_2_C4636 
                                               PAK1 
                                               mRNA 
                                               0 
                                               
                                               
                                               EPHB-mediated forward signaling;Ephrin signaling;MAPK6MAPK4 signaling;DSCAM interactions;Activation of Rac;VEGFR2 mediated vascular permeability;RHO GTPases activate PAKs;Sema3A PAK dependent Axon repulsion;FCERI mediated MAPK activation;Regulation of actin dynamics for phagocytic cup formation;CD209 (DC-SIGN) signaling;CD28 dependent Vav1 pathway;VEGFA-VEGFR2 Pathway;CD28 co-stimulation 
                                               hsa04010;hsa04012;hsa04014;hsa04024;hsa04062;hsa04360;hsa04510;hsa04650;hsa04660;hsa04666;hsa04810;hsa05120;hsa05205;hsa05211 
                                               PAK1 
                                               NA 
                                               slateblue1 
                                               16 
                                               14 
                                             
                                             
                                               RAC1-mRNA 
                                               1.11 
                                               0.0959 
                                               0.925 
                                               1.3 
                                               2.16 
                                               1.9 
                                               2.46 
                                               6.93e-14 
                                               1.24e-12 
                                               lm.nb 
                                               Activation of Rac, CD28 co-stimulation, CD28 dependent Vav1 pathway, DAP12 signaling, EPH-ephrin mediated repulsion of cells, Ephrin signaling, G alpha (1213) signalling events, GPVI-mediated activation cascade, MAPK6MAPK4 signaling, NRAGE signals death through JNK, PCPCE pathway, Regulation of actin dynamics for phagocytic cup formation, RHO GTPases activate CIT, RHO GTPases Activate Formins, RHO GTPases activate IQGAPs, RHO GTPases activate PAKs, RHO GTPases activate PKNs, RHO GTPases Activate WASPs and WAVEs, Sema3A PAK dependent Axon repulsion, Sema4D induced cell migration and growth-cone collapse, Signal transduction by L1, Signaling by SCF-KIT, VEGFA-VEGFR2 Pathway, VEGFR2 mediated vascular permeability 
                                               SEBELIUS_2_C4636 
                                               RAC1 
                                               mRNA 
                                               0 
                                               
                                               
                                               MAPK6MAPK4 signaling;Signaling by SCF-KIT;Inactivation of Cdc42 and Rac;DAP12 signaling;Ephrin signaling;EPH-ephrin mediated repulsion of cells;Activation of Rac;RHO GTPases Activate Formins;VEGFR2 mediated vascular permeability;RHO GTPases activate PAKs;Sema4D mediated inhibition of cell attachment and migration;G alpha (1213) signalling events;Regulation of actin dynamics for phagocytic cup formation;Sema4D induced cell migration and growth-cone collapse;Signal transduction by L1;RHO GTPases activate CIT;RHO GTPases Activate WASPs and WAVEs;Factors involved in megakaryocyte development and platelet production;RHO GTPases activate KTN1;CD28 dependent Vav1 pathway;DSCAM interactions;NRAGE signals death through JNK;GPVI-mediated activation cascade;Sema3A PAK dependent Axon repulsion;RHO GTPases activate PKNs;SEMA3A-Plexin repulsion signaling by inhibiting Integrin adhesion;RHO GTPases activate IQGAPs;PCPCE pathway;VEGFA-VEGFR2 Pathway;CD28 co-stimulation 
                                               hsa04010;hsa04014;hsa04015;hsa04024;hsa04062;hsa04071;hsa04145;hsa04151;hsa04310;hsa04360;hsa04370;hsa04380;hsa04510;hsa04520;hsa04620;hsa04650;hsa04662;hsa04664;hsa04666;hsa04670;hsa04722;hsa04810;hsa04932;hsa04933;hsa04972;hsa05100;hsa05120;hsa05131;hsa05132;hsa05200;hsa05203;hsa05205;hsa05210;hsa05211;hsa05212;hsa05231;hsa05416 
                                               RAC1 
                                               NA 
                                               slateblue1 
                                               16 
                                               11.5745568300313 
                                             
                                             
                                               CDC42-mRNA 
                                               0.458 
                                               0.1 
                                               0.262 
                                               0.654 
                                               1.37 
                                               1.2 
                                               1.57 
                                               5.18e-05 
                                               0.000494 
                                               lm.nb 
                                               CD28 co-stimulation, CD28 dependent Vav1 pathway, DCC mediated attractive signaling, EPHB-mediated forward signaling, G alpha (1213) signalling events, GPVI-mediated activation cascade, Regulation of actin dynamics for phagocytic cup formation, Rho GTPase cycle, RHO GTPases Activate Formins, RHO GTPases activate IQGAPs, RHO GTPases activate PAKs, Sema4D induced cell migration and growth-cone collapse, VEGFA-VEGFR2 Pathway 
                                               SEBELIUS_2_C4636 
                                               CDC42 
                                               mRNA 
                                               0 
                                               
                                               
                                               DCC mediated attractive signaling;Factors involved in megakaryocyte development and platelet production;VEGFA-VEGFR2 Pathway;RHO GTPases activate KTN1;CD28 dependent Vav1 pathway;Inactivation of Cdc42 and Rac;Rho GTPase cycle;EPHB-mediated forward signaling;RHO GTPases Activate Formins;RHO GTPases activate PAKs;GPVI-mediated activation cascade;G alpha (1213) signalling events;RHO GTPases activate IQGAPs;Sema4D induced cell migration and growth-cone collapse;Regulation of actin dynamics for phagocytic cup formation;CD28 co-stimulation 
                                               hsa04010;hsa04014;hsa04015;hsa04062;hsa04144;hsa04360;hsa04370;hsa04510;hsa04520;hsa04530;hsa04660;hsa04666;hsa04670;hsa04722;hsa04810;hsa04912;hsa04932;hsa04933;hsa05100;hsa05120;hsa05130;hsa05131;hsa05132;hsa05200;hsa05203;hsa05205;hsa05211;hsa05212 
                                               CDC42 
                                               NA 
                                               slateblue1 
                                               16 
                                               4.58 
                                             
                                             
                                               FYN-mRNA 
                                               -1.21 
                                               0.345 
                                               -1.89 
                                               -0.534 
                                               0.432 
                                               0.271 
                                               0.69 
                                               0.00119 
                                               0.01 
                                               lm.nb 
                                               ARMS-mediated activation, CD209 (DC-SIGN) signaling, CD28 co-stimulation, CD28 dependent PI3KAkt signaling, CD28 dependent Vav1 pathway, Cell surface interactions at the vascular wall, Constitutive Signaling by Aberrant PI3K in Cancer, CTLA4 inhibitory signaling, DAP12 signaling, Downstream signal transduction, EPH-ephrin mediated repulsion of cells, FCERI mediated MAPK activation, Frs2-mediated activation, GAB1 signalosome, Gastrin-CREB signalling pathway via PKC and MAPK, GPVI-mediated activation cascade, Interleukin receptor SHC signaling, IRS-related events triggered by IGF1R, Nephrin interactions, PI3KAKT activation, RAFMAP kinase cascade, Regulation of signaling by CBL, Role of LAT2NTALLAB on calcium mobilization, Sema3A PAK dependent Axon repulsion, Signaling by ERBB2, Signaling by PDGF, Signaling by SCF-KIT, VEGFA-VEGFR2 Pathway, VEGFR2 mediated cell proliferation 
                                               SEBELIUS_2_C4636 
                                               FYN 
                                               mRNA 
                                               0 
                                               
                                               
                                               EPH-ephrin mediated repulsion of cells;Regulation of signaling by CBL;FCGR activation;Antigen activates B Cell Receptor (BCR) leading to generation of second messengers;Dectin-2 family;RAFMAP kinase cascade;Constitutive Signaling by Aberrant PI3K in Cancer;CTLA4 inhibitory signaling;CD28 dependent PI3KAkt signaling;PECAM1 interactions;CD28 dependent Vav1 pathway;GPVI-mediated activation cascade;Sema3A PAK dependent Axon repulsion;Nephrin interactions;SEMA3A-Plexin repulsion signaling by inhibiting Integrin adhesion;CRMPs in Sema3A signaling;CD209 (DC-SIGN) signaling;Downstream signal transduction;GAB1 signalosome;Signaling by SCF-KIT;PI3KAKT activation;Role of LAT2NTALLAB on calcium mobilization;DAP12 signaling;Signaling by PDGF;Signaling by ERBB2;VEGFA-VEGFR2 Pathway;VEGFR2 mediated cell proliferation;Frs2-mediated activation;ARMS-mediated activation;Gastrin-CREB signalling pathway via PKC and MAPK;Interleukin receptor SHC signaling;FCERI mediated MAPK activation;IRS-related events triggered by IGF1R;CD28 co-stimulation;Cell surface interactions at the vascular wall 
                                               hsa04071;hsa04072;hsa04360;hsa04380;hsa04510;hsa04520;hsa04611;hsa04650;hsa04660;hsa04664;hsa04725;hsa05020;hsa05130;hsa05162;hsa05416 
                                               FYN 
                                               NA 
                                               slateblue2 
                                               16 
                                               -3.50724637681159 
                                             
                                             
                                               PAK2-mRNA 
                                               0.221 
                                               0.0777 
                                               0.0684 
                                               0.373 
                                               1.17 
                                               1.05 
                                               1.3 
                                               0.00728 
                                               0.0571 
                                               loglinear 
                                               Activation of Rac, CD209 (DC-SIGN) signaling, CD28 co-stimulation, CD28 dependent Vav1 pathway, Ephrin signaling, RHO GTPases activate PAKs, Sema3A PAK dependent Axon repulsion, VEGFA-VEGFR2 Pathway, VEGFR2 mediated vascular permeability 
                                               SEBELIUS_2_C4636 
                                               PAK2 
                                               mRNA 
                                               0 
                                               
                                               
                                               Nef and signal transduction;CD28 dependent Vav1 pathway;Stimulation of the cell death response by PAK-2p34;Ephrin signaling;Activation of Rac;RHO GTPases activate PAKs;VEGFR2 mediated vascular permeability;Sema3A PAK dependent Axon repulsion;CD209 (DC-SIGN) signaling;VEGFA-VEGFR2 Pathway;CD28 co-stimulation 
                                               hsa04010;hsa04012;hsa04014;hsa04360;hsa04510;hsa04660;hsa04810;hsa05211 
                                               PAK2 
                                               NA 
                                               slateblue3 
                                               16 
                                               2.84427284427284 
                                             
                                           
                                           DE Results - CD28 dependent Vav1 pathway genes - Line: HT29 vs.Fibro  Table displaying each sample's global significance scores and directed global significance scores as defined in the heatmaps above. The global significance score is calculated as the square root of the mean squared t-statistic for the genes in a gene set, with t-statistics coming from the linear regression underlying our differential expression analysis. The directed global significance score is calculated as the square root of the mean signed squared t-statistic for the genes in a gene set, with t-statistics coming from the linear regression underlying our differential expression analysis.   
                                      
                                       
                                  
                                
                              
                            
                           
                            
                          
                           
                      
                    
                    
                      
                           
                          
                               Day    Line     
                              
                                  Day  
                              
                               
                              
                                  Line: differential expression in Caco2 vs. baseline of Fibro  
                                  Line: differential expression in HEPG2 vs. baseline of Fibro  
                                  Line: differential expression in HT29 vs. baseline of Fibro  
                              
                              
                            
                              
                                
                                  
                                       
                                      
                                           Volcano Plot: Day   More Plot Information                Volcano Plot: Day  Volcano plot displaying each gene's -log10(p-value) and log2 fold change for the selected covariate. Highly statistically significant genes fall at the top of the plot, and highly differentially expressed genes fall to either side. Genes within the selected gene set are highlighted in orange. Horizontal lines indicate various False Discovery Rate (FDR) thresholds.   
                                           DE Results - Cell-extracellular matrix interactions genes - Day   More Plot Information      Download CSV Data     
                                           
                                             
                                             
                                           
                                           
                                             
                                               Probe Label 
                                               Log2 fold change 
                                               std error (log2) 
                                               Lower confidence limit (log2) 
                                               Upper confidence limit (log2) 
                                               Linear fold change 
                                               Lower confidence limit (linear) 
                                               Upper confidence limit (linear) 
                                               P-value 
                                               BY.p.value 
                                               method 
                                               Gene.sets 
                                               Codeset.Name 
                                               Probe.Label 
                                               Analyte.Type 
                                               Is.Control 
                                               Control.Type 
                                               Related.Probes 
                                               Probe.Annotation 
                                               KEGG.Pathways 
                                               Official.Gene.Name 
                                               Control.Conc 
                                               volcanocol 
                                               volcanopch 
                                               tstats.all 
                                             
                                           
                                           
                                             
                                               FLNA-mRNA 
                                               -0.107 
                                               0.0254 
                                               -0.156 
                                               -0.0569 
                                               0.929 
                                               0.897 
                                               0.961 
                                               0.00016 
                                               0.00515 
                                               lm.nb 
                                               Cell-extracellular matrix interactions, Platelet degranulation , RHO GTPases activate PAKs 
                                               SEBELIUS_2_C4636 
                                               FLNA 
                                               mRNA 
                                               0 
                                               
                                               
                                               RHO GTPases activate PAKs;Platelet degranulation ;Cell-extracellular matrix interactions 
                                               hsa04010;hsa04510;hsa05132;hsa05205 
                                               FLNA 
                                               NA 
                                               slateblue1 
                                               16 
                                               -4.21259842519685 
                                             
                                             
                                               ITGB1-mRNA 
                                               -0.0466 
                                               0.0124 
                                               -0.071 
                                               -0.0222 
                                               0.968 
                                               0.952 
                                               0.985 
                                               0.000611 
                                               0.0149 
                                               lm.nb 
                                               Cell-extracellular matrix interactions, Cell surface interactions at the vascular wall, ECM proteoglycans, Elastic fibre formation, Integrin cell surface interactions, Laminin interactions, Non-integrin membrane-ECM interactions, RHO GTPases Activate Formins 
                                               SEBELIUS_2_C4636 
                                               ITGB1 
                                               mRNA 
                                               0 
                                               
                                               
                                               Laminin interactions;RHO GTPases Activate Formins;ECM proteoglycans;Fibronectin matrix formation;Integrin cell surface interactions;Immunoregulatory interactions between a Lymphoid and a non-Lymphoid cell;Localization of the PINCH-ILK-PARVIN complex to focal adhesions;Elastic fibre formation;Cell surface interactions at the vascular wall;Non-integrin membrane-ECM interactions;Cell-extracellular matrix interactions 
                                               hsa04015;hsa04145;hsa04151;hsa04360;hsa04510;hsa04512;hsa04514;hsa04611;hsa04670;hsa04810;hsa05100;hsa05130;hsa05131;hsa05133;hsa05140;hsa05145;hsa05200;hsa05205;hsa05222;hsa05410;hsa05412;hsa05414 
                                               ITGB1 
                                               NA 
                                               slateblue2 
                                               16 
                                               -3.75806451612903 
                                             
                                             
                                               ACTB-mRNA 
                                               -0.0865 
                                               0.0282 
                                               -0.142 
                                               -0.0313 
                                               0.942 
                                               0.906 
                                               0.979 
                                               0.00397 
                                               0.0647 
                                               lm.nb 
                                               ARMS-mediated activation, Cell-extracellular matrix interactions, DAP12 signaling, Downstream signal transduction, EPH-ephrin mediated repulsion of cells, EPHB-mediated forward signaling, FCERI mediated MAPK activation, Frs2-mediated activation, Gastrin-CREB signalling pathway via PKC and MAPK, Interleukin receptor SHC signaling, IRS-related events triggered by IGF1R, RAFMAP kinase cascade, Regulation of actin dynamics for phagocytic cup formation, RHO GTPases Activate Formins, RHO GTPases activate IQGAPs, Signaling by ERBB2, Signaling by PDGF, Signaling by SCF-KIT, VEGFA-VEGFR2 Pathway, VEGFR2 mediated cell proliferation 
                                               SEBELIUS_2_C4636 
                                               ACTB 
                                               mRNA 
                                               0 
                                               
                                               
                                               EPHB-mediated forward signaling;Interaction between L1 and Ankyrins;DNA Damage Recognition in GG-NER;EPH-ephrin mediated repulsion of cells;Prefoldin mediated transfer of substrate  to CCTTriC;RHO GTPases Activate Formins;RHO GTPases activate IQGAPs;Cell-extracellular matrix interactions;Regulation of actin dynamics for phagocytic cup formation;Folding of actin by CCTTriC;B-WICH complex positively regulates rRNA expression;RAFMAP kinase cascade;Downstream signal transduction;VEGFR2 mediated cell proliferation;Signaling by SCF-KIT;Frs2-mediated activation;ARMS-mediated activation;Gastrin-CREB signalling pathway via PKC and MAPK;Interleukin receptor SHC signaling;FCERI mediated MAPK activation;DAP12 signaling;Signaling by PDGF;VEGFA-VEGFR2 Pathway;Signaling by ERBB2;IRS-related events triggered by IGF1R 
                                               hsa04015;hsa04145;hsa04390;hsa04510;hsa04520;hsa04530;hsa04611;hsa04670;hsa04810;hsa04919;hsa04921;hsa04971;hsa05100;hsa05110;hsa05130;hsa05131;hsa05132;hsa05164;hsa05205;hsa05410;hsa05412;hsa05414;hsa05416 
                                               ACTB 
                                               NA 
                                               slateblue3 
                                               16 
                                               -3.06737588652482 
                                             
                                             
                                               ACTN1-mRNA 
                                               -0.0584 
                                               0.0246 
                                               -0.107 
                                               -0.0101 
                                               0.96 
                                               0.929 
                                               0.993 
                                               0.023 
                                               0.272 
                                               lm.nb 
                                               Cell-extracellular matrix interactions, Nephrin interactions, Non-integrin membrane-ECM interactions, Platelet degranulation , Syndecan interactions 
                                               SEBELIUS_2_C4636 
                                               ACTN1 
                                               mRNA 
                                               0 
                                               
                                               
                                               Syndecan interactions;Nephrin interactions;Platelet degranulation ;Non-integrin membrane-ECM interactions;Cell-extracellular matrix interactions 
                                               hsa04510;hsa04520;hsa04530;hsa04670;hsa04810;hsa05146;hsa05203;hsa05412 
                                               ACTN1 
                                               NA 
                                               slateblue4 
                                               16 
                                               -2.3739837398374 
                                             
                                             
                                               PXN-mRNA 
                                               0.0257 
                                               0.0143 
                                               -0.00222 
                                               0.0537 
                                               1.02 
                                               0.998 
                                               1.04 
                                               0.0794 
                                               0.771 
                                               lm.nb 
                                               Cell-extracellular matrix interactions, GAB1 signalosome, Smooth Muscle Contraction, VEGFA-VEGFR2 Pathway 
                                               SEBELIUS_2_C4636 
                                               PXN 
                                               mRNA 
                                               0 
                                               
                                               
                                               Smooth Muscle Contraction;GAB1 signalosome;VEGFA-VEGFR2 Pathway;Cell-extracellular matrix interactions 
                                               hsa04062;hsa04370;hsa04510;hsa04670;hsa04810;hsa05100;hsa05203;hsa05205 
                                               PXN 
                                               NA 
                                               azure4 
                                               1 
                                               1.7972027972028 
                                             
                                             
                                               ACTG1-mRNA 
                                               -0.0332 
                                               0.0186 
                                               -0.0696 
                                               0.00335 
                                               0.977 
                                               0.953 
                                               1 
                                               0.0832 
                                               0.802 
                                               lm.nb 
                                               ARMS-mediated activation, Cell-extracellular matrix interactions, DAP12 signaling, Downstream signal transduction, EPH-ephrin mediated repulsion of cells, EPHB-mediated forward signaling, FCERI mediated MAPK activation, Frs2-mediated activation, Gastrin-CREB signalling pathway via PKC and MAPK, Interleukin receptor SHC signaling, IRS-related events triggered by IGF1R, RAFMAP kinase cascade, Regulation of actin dynamics for phagocytic cup formation, RHO GTPases Activate Formins, RHO GTPases activate IQGAPs, RHO GTPases Activate WASPs and WAVEs, Signaling by ERBB2, Signaling by PDGF, Signaling by SCF-KIT, VEGFA-VEGFR2 Pathway, VEGFR2 mediated cell proliferation 
                                               SEBELIUS_2_C4636 
                                               ACTG1 
                                               mRNA 
                                               0 
                                               
                                               
                                               RHO GTPases Activate WASPs and WAVEs;Translocation of GLUT4 to the plasma membrane;Recycling pathway of L1;Interaction between L1 and Ankyrins;EPHB-mediated forward signaling;EPH-ephrin mediated repulsion of cells;RHO GTPases Activate Formins;RHO GTPases activate IQGAPs;Cell-extracellular matrix interactions;Regulation of actin dynamics for phagocytic cup formation;RAFMAP kinase cascade;Downstream signal transduction;VEGFR2 mediated cell proliferation;Signaling by SCF-KIT;Frs2-mediated activation;ARMS-mediated activation;Gastrin-CREB signalling pathway via PKC and MAPK;Interleukin receptor SHC signaling;FCERI mediated MAPK activation;DAP12 signaling;Signaling by PDGF;VEGFA-VEGFR2 Pathway;Signaling by ERBB2;IRS-related events triggered by IGF1R 
                                               hsa04015;hsa04145;hsa04390;hsa04510;hsa04520;hsa04530;hsa04611;hsa04670;hsa04810;hsa04919;hsa04921;hsa05100;hsa05110;hsa05130;hsa05131;hsa05132;hsa05164;hsa05205;hsa05410;hsa05412;hsa05414;hsa05416 
                                               ACTG1 
                                               NA 
                                               azure4 
                                               1 
                                               -1.78494623655914 
                                             
                                           
                                           DE Results - Cell-extracellular matrix interactions genes - Day  Table displaying each sample's global significance scores and directed global significance scores as defined in the heatmaps above. The global significance score is calculated as the square root of the mean squared t-statistic for the genes in a gene set, with t-statistics coming from the linear regression underlying our differential expression analysis. The directed global significance score is calculated as the square root of the mean signed squared t-statistic for the genes in a gene set, with t-statistics coming from the linear regression underlying our differential expression analysis.   
                                      
                                       
                                  
                                
                              
                            
                            
                              
                                
                                  
                                       
                                      
                                           Volcano Plot: Line: Caco2 vs.Fibro   More Plot Information                Volcano Plot: Line: Caco2 vs.Fibro  Volcano plot displaying each gene's -log10(p-value) and log2 fold change for the selected covariate. Highly statistically significant genes fall at the top of the plot, and highly differentially expressed genes fall to either side. Genes within the selected gene set are highlighted in orange. Horizontal lines indicate various False Discovery Rate (FDR) thresholds.   
                                           DE Results - Cell-extracellular matrix interactions genes - Line: Caco2 vs.Fibro   More Plot Information      Download CSV Data     
                                           
                                             
                                             
                                           
                                           
                                             
                                               Probe Label 
                                               Log2 fold change 
                                               std error (log2) 
                                               Lower confidence limit (log2) 
                                               Upper confidence limit (log2) 
                                               Linear fold change 
                                               Lower confidence limit (linear) 
                                               Upper confidence limit (linear) 
                                               P-value 
                                               BY.p.value 
                                               method 
                                               Gene.sets 
                                               Codeset.Name 
                                               Probe.Label 
                                               Analyte.Type 
                                               Is.Control 
                                               Control.Type 
                                               Related.Probes 
                                               Probe.Annotation 
                                               KEGG.Pathways 
                                               Official.Gene.Name 
                                               Control.Conc 
                                               volcanocol 
                                               volcanopch 
                                               tstats.all
