## Supplementary material for "Comparative pathway enrichment analysis in gastrointestinal cell lines Caco-2, HT-29, HEPG2, and colon fibroblasts using a custom expression panel for tight-junction and cytoskeletal regulatory genes": Sup. Results 1: panel2_2.js

```
document.write('      	  <li>Volcano Plot - Day<br><img src="results//DE//volcano plotDay.png"></img></li>');
document.write('      	  <li>Volcano Plot - LineCaco2<br><img src="results//DE//volcano plotLineCaco2.png"></img></li>');
document.write('      	  <li>Volcano Plot - LineHT29<br><img src="results//DE//volcano plotLineHT29.png"></img></li>');
document.write('      	  <li>Volcano Plot - LineHEPG2<br><img src="results//DE//volcano plotLineHEPG2.png"></img></li>');
```
