## Supplementary material for "Comparative pathway enrichment analysis in gastrointestinal cell lines Caco-2, HT-29, HEPG2, and colon fibroblasts using a custom expression panel for tight-junction and cytoskeletal regulatory genes": Sup. Results 1: advanced.analysis.js

```
	heatmapData_UQh1YWZvRyUX_kmCGIztRmQ = "Dimname,20161013_sebelius2-101316_01_10.RCC,20161013_sebelius2-101316_01_10.RCC,20161013_sebelius2-101316_01_09.RCC,20161013_sebelius2-101316_01_10.RCC,20170307_20170307-caco_01_05.RCC,20170307_20170307-caco_01_06.RCC,20170307_20170307-caco_01_03.RCC,20170307_20170307-caco_01_04.RCC,20170307_20170307-caco_01_01.RCC,20170307_20170307-caco_01_02.RCC,20180926_Polarization01_FibD4B_06.RCC,20180926_Polarization01_FibD4A_05.RCC,20180928_Polarization-02_FibD7A_05.RCC,20180928_Polarization-02_FibD7A_05.RCC,20180926_Polarization01_Caco2D4A_01.RCC,20180926_Polarization01_Caco2D4A_01.RCC,20180926_Polarization01_Caco2D4A_01.RCC,20180926_Polarization01_Caco2D4A_01.RCC,20180928_Polarization-02_Caco2D7A_01.RCC,20180928_Polarization-02_Caco2D7A_01.RCC,20180928_Polarization-02_Caco2D7A_01.RCC,20180928_Polarization-02_Caco2D7A_01.RCC,20181004_Pol03_HepG2D4A_01.RCC,20181004_Pol03_HepG2D4B_02.RCC,20181004_Pol03_HepG2D4C_03.RCC,20181004_Pol03_HepG2D4D_04.RCC,20181004_Pol03_HepG2D7A_05.RCC,20181004_Pol03_HepG2D7B_06.RCC,20181004_Pol03_HepG2D7C_07.RCC,20181004_Pol03_HepG2D7D_08.RCC,20180926_Polarization01_HT29D4A_07.RCC,20180926_Polarization01_HT29D4A_07.RCC,20180926_Polarization01_HT29D4A_07.RCC,20180926_Polarization01_HT29D4A_07.RCC,20180926_Polarization01_HT29D4A_07.RCC,20180926_Polarization01_HT29D4A_07.RCC,20180928_Polarization-02_HT29D7A_07.RCC,20180928_Polarization-02_HT29D7A_07.RCC,20180928_Polarization-02_HT29D7A_07.RCC,20180928_Polarization-02_HT29D7A_07.RCC,20180928_Polarization-02_HT29D7A_07.RCC,20180928_Polarization-02_HT29D7A_07.RCC\nACTB - mRNA,4134,3131,550,666,854,887,2915,1920,7892,5346,82078,94908,95873,89698,46819,43028,35762,54864,38077,36843,39973,36828,51540,44899,24673,22289,35511,32730,28566,27136,23248,23553,41311,37104,38785,38739,34299,33935,36465,43933,38419,38656\nCFL1 - mRNA,2612,1356,196,526,355,456,906,947,2042,1494,34355,40050,40808,40579,27447,28217,25184,27379,17229,20445,19454,21185,31296,31652,15454,14738,21906,21879,14489,14526,25961,27465,61136,54252,58616,56979,41071,44003,45940,46347,46521,47053\nMYL12B - mRNA,2588,1983,194,446,669,763,1044,960,2494,1887,25001,29372,29457,29334,21722,22115,20052,20039,15256,17201,16279,19061,18125,18563,9901,10174,12708,12809,9454,9364,40728,43046,60800,54779,57314,55616,52956,53608,53665,53837,52223,54912\nPOS_A(128) - mRNA,10837,6212,11130,1696,3732,3638,10421,6113,8301,7349,31405,32980,26132,23476,23664,18900,15126,24013,36747,17960,19243,18759,26107,22680,20544,19333,21440,22530,24096,19316,13445,16986,18127,17597,15978,17990,23003,18276,18059,24725,17997,19368\nCCND1 - mRNA,1391,1302,141,201,240,267,620,630,1535,1068,25892,30975,35849,36605,16326,15898,13697,16219,10041,12715,11491,12063,21357,22900,19635,18345,19931,18787,19414,18962,7112,7603,6299,5165,6054,5664,3712,4073,3659,3949,3971,3683\nITGB1 - mRNA,1593,1269,136,213,390,452,633,580,1513,1050,39157,47249,52042,51157,15608,15336,14296,17011,11424,12544,11896,13221,18111,17640,17964,17868,13753,13819,11713,10923,14653,14928,12763,11630,12176,11833,14982,14268,13024,13613,13164,13656\nCTNNB1 - mRNA,1727,1101,96,150,392,498,882,766,1129,843,10651,12093,14492,14244,22915,23583,22492,24177,12520,16101,15576,17004,53095,52187,35457,35348,40104,40393,40837,40381,11775,12752,14094,12311,13633,13443,13597,13521,13150,13462,13877,13959\nPPP1CA - mRNA,3819,2050,206,469,729,787,1577,1506,2413,1862,8004,9645,9977,10304,34768,35209,31532,33173,20353,24029,24027,25998,19114,19317,13004,12958,15452,15231,12112,11889,19895,20795,33281,28833,31990,30776,20041,20783,25312,26280,25525,25756\nMYL9 - mRNA,1540,609,190,132,197,235,529,597,2035,1558,60241,72020,71373,71851,9461,9368,8088,9070,6248,7087,6420,7802,648,441,121,144,278,253,109,126,31,24,45,39,35,26,42,39,30,23,28,23\nROCK2 - mRNA,532,596,65,78,156,181,234,197,308,259,4498,5690,6290,6162,4485,4412,3960,4024,3431,4149,3882,3966,14571,15230,22340,22066,17682,15623,22619,20407,8086,8486,8065,6640,7460,7197,7154,7464,8264,9212,9018,8492\nTUBB - mRNA,809,447,57,114,120,151,517,334,649,513,4159,5146,6236,6429,8936,8048,7172,7693,7976,8018,8143,7487,13002,13173,4580,4248,13206,13181,7807,7395,1893,2139,9245,8110,8559,8211,5427,4780,8373,9202,8200,8579\nPPP2CA - mRNA,1141,843,87,147,308,386,516,496,890,686,6102,6802,8147,8083,12534,12582,11407,12655,8248,9886,9324,10499,13606,13304,8138,8126,8922,9161,6874,6985,7791,8565,10876,9849,10309,10264,7608,7684,8647,8772,8729,8997\nNCKAP1 - mRNA,905,711,85,124,246,262,375,307,566,488,7487,9371,9586,9853,8693,8547,7673,8616,5354,6189,6355,6625,10795,11175,8134,8123,8363,8087,8114,7691,9888,10179,9092,7996,8681,8414,9426,9378,8887,9354,8793,9135\nPPP1CB - mRNA,927,824,46,139,245,320,357,327,456,318,5584,6676,7387,7186,7247,6832,6549,6927,6339,6978,6372,7178,10293,10778,9622,9199,9485,9064,9830,9055,8803,9300,8711,8008,8163,7891,10295,10177,9382,9784,9362,9525\nALDH18A1 - mRNA,955,896,34,177,224,274,409,364,440,336,2394,2804,3272,3044,7088,6988,6364,6958,6314,6962,6639,7351,12137,11630,10734,9782,12960,12386,15011,15064,4546,5349,6267,5393,5807,6014,9312,9786,6189,6340,6319,6630\nCSNK2B - mRNA,1069,1023,47,156,284,342,519,454,675,475,6743,8304,7994,8350,11376,11471,10318,11072,7853,8997,8844,9241,19545,19989,15151,15321,14843,14958,14887,13649,7021,7563,10414,9562,9890,9937,7350,7597,7399,7856,7611,7720\nCTNNA1 - mRNA,2201,1939,105,315,570,662,987,832,1328,1069,12798,15486,16210,15912,17516,17249,15387,16779,13097,14937,14320,15237,12641,13100,11493,11382,11682,11314,12895,12305,13924,14751,18387,16242,17580,17391,17085,17801,16617,17350,16811,17392\nCDC42 - mRNA,2046,1334,131,259,489,532,790,717,1406,1101,13126,15358,16629,16825,16506,16983,15265,16543,11134,14179,13414,14681,18068,18326,11069,11259,13348,13510,11425,11344,13709,14821,23205,20201,21608,21176,17756,17652,18214,18332,18636,18784\nPPP2R1A - mRNA,1905,1394,70,269,438,444,761,636,773,584,9244,11109,11572,11406,16907,16810,15064,16340,11833,13948,13543,14108,10058,10161,7488,7481,8451,8404,8848,8160,11023,11986,14709,12949,14135,13581,12796,13400,12073,12530,12385,12499\nCSNK1A1 - mRNA,1532,1242,81,202,357,415,582,504,776,564,10729,13099,14609,13805,12349,12016,10976,10856,9201,10722,10136,10641,12061,12749,13593,13471,9980,9684,10174,9594,15251,15294,12814,10923,11923,11276,12807,13240,12343,13321,12533,12903\nACTN4 - mRNA,861,878,42,189,241,257,387,360,750,607,10595,12712,12814,12233,9067,9160,8223,9430,7635,8368,7967,8277,11290,10826,9671,9240,11585,11437,12755,11432,12265,12310,18185,16749,17570,17256,14448,13969,14659,16067,15652,15448\nRAB11A - mRNA,1044,995,58,218,316,396,384,390,569,432,5222,5964,6358,6506,8378,8877,8291,7838,6961,8634,7845,8753,7377,7632,4522,4533,5863,6034,5545,5256,14533,16135,21363,18910,20551,19934,19773,20481,17391,16899,17298,17733\nRAC1 - mRNA,692,640,65,105,214,231,346,278,629,486,5539,6297,7864,7698,7386,7013,6098,8027,6042,6894,6263,6922,7347,6899,6417,6316,5967,5857,5695,5374,9256,9648,13958,12331,13375,13011,13129,13905,13065,14911,13891,14292\nIQGAP1 - mRNA,754,481,42,82,147,147,230,227,440,355,4414,5093,5768,5956,5432,5629,4892,5332,3582,4349,4079,4572,3132,3281,1967,2085,2775,2690,1864,1782,9201,9605,11491,9703,10927,10554,12642,12900,12396,12232,12488,12716\nPPP2CB - mRNA,634,607,38,89,190,165,289,225,633,442,6206,7686,8100,8045,5997,5685,5092,6034,4346,4670,4763,4841,5972,6032,4552,4590,4613,4465,4279,4081,1472,1479,1790,1540,1663,1596,1757,1777,1524,1690,1620,1610\nEPB41L2 - mRNA,407,449,43,43,103,122,175,163,276,182,9359,10996,10484,10915,3638,3670,3353,3541,2865,3567,3275,3227,6863,7086,4184,3766,6277,6057,5527,5054,167,185,321,283,300,293,268,289,259,250,239,257\nMYH10 - mRNA,309,229,18,33,61,73,152,133,211,171,6627,8933,8699,8334,3128,3055,2765,2903,2200,2606,2655,2582,5812,6154,3371,3187,5991,5687,5690,5320,365,446,690,556,694,634,611,617,692,640,667,624\nHSPB1 - mRNA,177,48,12,17,9,18,114,79,114,82,6753,8180,7981,7394,2321,2094,1883,2354,1958,1613,1634,1723,827,732,393,341,683,615,230,193,680,790,554,467,450,547,959,891,692,890,729,715\nPDGFRB - mRNA,73,95,3,11,13,27,38,30,24,22,8360,10401,13236,13166,762,766,654,634,730,825,895,805,785,893,1091,1041,941,939,920,826,48,28,39,38,39,41,34,49,46,42,43,37\nITGA5 - mRNA,96,87,8,23,15,15,22,25,57,40,8949,10376,11273,11464,628,612,515,676,698,724,723,697,63,34,34,27,32,20,11,8,11,16,16,14,16,16,13,13,11,17,14,16\nGSTM3 - mRNA,16,16,1,1,3,7,6,5,10,24,8283,9706,9187,9298,176,162,175,181,126,161,121,156,5923,6139,3803,3722,6117,6168,4511,4549,14,23,22,17,23,14,15,25,20,21,39,22\nADAM9 - mRNA,333,301,33,46,70,100,117,126,283,220,7726,8440,8877,9377,2503,2673,2410,2500,1687,2017,1820,2241,5084,5312,4318,4186,3614,3772,2643,2598,2490,2466,2253,1992,2221,2060,1958,2058,1921,1843,1958,1969\nACTN1 - mRNA,219,133,14,35,34,33,101,97,267,187,9146,11134,10580,10766,2503,2427,2096,2401,1597,1663,1702,1710,3967,4104,2723,2595,3340,3349,3132,2956,1346,1401,1563,1411,1490,1509,1401,1521,1273,1436,1378,1342\nFGFR1 - mRNA,107,62,8,8,18,29,34,40,99,68,4528,5303,6765,6614,1236,1189,1151,1451,1013,1141,1026,1141,3653,3405,2245,2213,2864,2890,2363,2179,13,18,18,24,20,12,17,18,17,11,23,24\nTGFB1 - mRNA,47,34,3,6,10,12,40,29,28,15,2765,3386,3894,4031,441,440,380,441,427,475,471,365,2108,1939,2614,2542,1918,1772,1947,1851,313,346,235,205,216,220,454,449,328,396,312,349\nITGAV - mRNA,112,74,20,16,35,40,53,59,103,83,3104,4029,4376,4377,922,963,823,898,624,724,706,717,3392,3541,2494,2508,2472,2378,1953,1899,1264,1347,1097,954,1047,1039,1291,1311,1189,1271,1152,1216\nPDGFRA - mRNA,292,211,13,7,60,80,138,121,233,143,4107,4560,5483,5203,1625,1554,1280,1836,1374,1405,1339,1484,40,38,31,37,32,36,35,28,25,30,29,38,28,26,34,42,21,21,34,30\nJAM3 - mRNA,11,11,4,2,3,1,1,5,11,1,3234,3721,3859,3853,84,69,75,85,68,82,72,63,384,408,204,206,259,251,164,155,25,17,31,15,16,19,30,20,24,17,25,18\nAKT3 - mRNA,75,68,5,15,18,14,25,26,20,20,2168,2753,3079,2935,423,456,385,363,391,442,409,445,30,27,33,25,27,27,27,21,22,13,14,23,16,21,10,21,32,13,22,16\nZEB1 - mRNA,7,4,1,1,2,1,2,1,1,1,2141,2485,3040,2958,35,42,27,26,36,50,39,33,593,556,659,616,584,537,675,622,31,32,20,27,43,31,46,45,23,34,35,34\nRAB8B - mRNA,185,109,7,21,44,54,79,74,78,61,1737,2055,2363,2378,1177,1167,1044,994,1010,1217,1143,1243,1145,1174,885,854,1049,1005,1066,977,291,326,246,224,253,282,635,617,458,397,429,436\nLIMK1 - mRNA,111,76,10,19,37,23,65,58,100,80,2148,2340,3073,2883,1255,1155,989,1086,971,1053,991,1106,1145,1218,872,778,934,939,585,546,855,983,972,808,831,829,1165,1281,955,1244,1098,1012\nBHLHE40 - mRNA,107,246,2,30,16,14,23,20,22,11,379,497,469,505,263,268,235,172,809,1123,851,929,622,771,4072,3528,4029,2999,1770,1464,204,294,134,108,122,150,416,571,390,376,429,396\nIL1R1 b - mRNA,99,29,5,7,9,14,34,35,33,24,700,948,893,916,866,916,820,1009,335,494,510,559,1212,1160,2160,2191,1666,1677,1682,1662,50,50,22,21,24,30,33,30,29,25,38,20\nSMAD4 - mRNA,188,166,10,27,41,53,94,62,77,71,951,1160,1168,1183,1110,972,835,980,1097,953,948,1047,1559,1672,2247,2031,1862,1595,2325,1935,274,318,235,244,215,217,312,339,297,377,332,283\nIGF1R - mRNA,90,119,10,9,33,29,40,42,69,49,435,521,622,600,656,662,542,624,645,796,729,715,1012,1041,2161,2035,1600,1470,2453,2183,930,964,809,711,724,779,1304,1469,1098,1159,1191,1223\nLIMK2 - mRNA,134,118,5,13,30,41,72,48,49,52,451,533,624,624,932,904,810,1031,713,835,766,849,1822,1736,1779,1664,1854,1786,1641,1546,419,510,736,641,640,696,500,512,611,638,595,674\nPIK3R2 - mRNA,133,106,5,18,45,44,84,68,44,51,850,1008,1107,1082,1238,1248,1052,1314,1095,1117,1054,1082,1340,1189,1581,1471,1498,1465,1577,1387,573,619,435,456,468,485,623,678,562,653,572,593\nDVL2 - mRNA,121,114,13,18,32,28,56,47,48,45,448,535,646,680,1145,1062,998,1063,943,1044,1006,1076,1467,1474,1252,1243,1459,1471,1547,1370,580,584,636,548,626,602,689,790,799,819,820,830\nCCND3 - mRNA,32,17,7,3,6,10,15,10,27,21,1683,1897,1613,1783,435,467,338,564,226,263,267,259,538,429,115,130,262,264,78,62,820,780,1049,962,1023,1063,1051,981,839,982,931,960\nRRAS - mRNA,13,14,1,1,3,3,4,3,7,9,888,1050,1179,1086,49,59,59,76,79,56,80,55,79,78,53,50,74,84,34,51,692,733,664,616,617,615,1225,1241,902,884,877,937\nCA2 - mRNA,82,158,2,25,36,57,27,34,17,17,13,23,20,15,560,594,503,549,562,645,684,630,34,44,13,31,21,22,17,20,498,594,1466,1283,1382,1419,1036,996,1169,1213,1144,1289\nCXCR4 - mRNA,38,63,3,6,12,12,19,17,9,9,20,15,16,29,711,695,642,558,547,722,666,636,113,110,101,78,96,80,36,40,859,919,1461,1300,1412,1425,1103,1020,1295,1312,1319,1333\nPIK3R3 - mRNA,22,16,2,4,3,7,2,6,7,7,551,630,825,735,178,177,152,184,156,160,146,177,394,401,131,118,210,224,56,56,682,721,1100,932,1093,1028,891,815,920,1039,955,981\nHRAS - mRNA,48,31,4,4,4,17,19,16,40,32,396,417,480,503,405,451,350,366,275,303,305,317,494,490,279,299,307,307,262,226,771,762,1369,1214,1309,1239,629,621,800,889,865,777\nMYH14 - mRNA,29,37,2,7,13,11,12,9,21,8,15,26,28,27,264,275,202,275,296,296,291,334,92,93,92,104,95,86,111,97,1440,1582,793,669,751,770,2144,2242,1310,1475,1468,1384\nMAGI3 - mRNA,129,130,14,14,34,38,50,49,65,57,174,190,261,291,974,1032,912,959,837,909,921,1025,489,494,662,688,632,624,714,653,985,1105,1343,1174,1323,1269,1591,1769,1553,1758,1675,1636\nIRS1 - mRNA,60,100,4,11,34,32,37,22,36,22,754,971,1122,1171,557,546,503,503,562,679,638,648,291,266,774,766,361,341,549,520,1501,1477,1205,982,1147,1074,1556,1636,1561,1607,1533,1473\nEXOC3 - mRNA,82,89,7,14,30,21,34,29,27,28,426,544,668,678,649,617,573,570,470,543,521,557,504,527,527,551,513,484,540,486,981,1132,1213,1070,1209,1165,1155,1268,1095,1154,1162,1203\nTCF7 - mRNA,66,22,6,4,6,7,17,19,46,36,60,74,96,100,763,736,652,792,289,384,353,418,727,715,846,804,662,592,583,492,767,875,735,637,697,681,736,732,890,875,898,826\nBAIAP2 - mRNA,73,69,11,10,25,17,50,41,78,54,293,380,407,365,799,672,607,782,575,539,583,625,734,658,886,884,768,639,743,693,878,948,788,747,763,711,911,908,755,885,824,774\nAPC - mRNA,58,48,5,6,16,17,24,25,24,13,519,554,634,668,466,456,448,516,371,405,396,396,632,637,542,519,520,522,486,497,630,639,710,629,643,682,628,654,602,619,646,666\nDVL1 - mRNA,30,58,3,13,24,21,18,18,30,19,266,310,466,464,367,392,378,368,304,332,328,334,814,926,879,918,828,843,918,899,741,748,727,663,688,710,820,942,690,737,753,766\nBCAR1 - mRNA,62,40,11,10,16,9,32,18,40,31,454,549,592,612,580,555,442,678,367,449,425,470,844,869,1084,1027,753,791,783,678,600,632,607,589,569,597,724,724,542,635,581,572\nSOS2 - mRNA,84,72,4,17,29,24,40,21,35,23,589,750,1000,971,603,572,501,590,477,631,565,592,799,732,1075,1013,825,826,860,826,811,920,343,317,340,350,885,909,610,668,684,704\nWASF1 - mRNA,60,78,4,7,20,24,23,25,25,33,525,616,739,763,619,628,520,567,506,602,606,593,1291,1172,1456,1476,1233,1223,1276,1198,573,716,703,612,646,659,678,674,731,814,752,830\nTARBP2 - mRNA,110,93,3,15,23,25,51,45,37,55,364,448,586,565,1170,1144,986,1106,732,911,861,918,1205,1188,846,851,1035,973,923,797,640,636,909,821,918,824,818,834,768,863,818,816\nABI2 - mRNA,123,119,6,17,36,29,58,56,46,37,591,731,796,773,979,1077,861,1069,932,1029,936,1031,1169,1148,1280,1307,1220,1142,1136,1066,946,1013,930,867,862,882,939,1007,1036,1092,1116,1038\nPRKCZ - mRNA,139,164,5,26,46,59,53,47,54,46,148,155,186,200,902,959,808,923,825,995,831,999,1270,1213,1653,1612,1274,1287,1498,1376,669,838,895,813,899,864,1094,1173,1096,1098,1123,1121\nPIK3CB - mRNA,117,152,11,20,24,43,64,47,89,65,254,283,332,361,1030,1006,833,954,764,849,811,902,982,1055,984,1009,1147,1068,1030,954,1164,1253,1152,1018,1036,1006,1256,1306,1233,1282,1187,1211\nLLGL2 - mRNA,82,36,9,10,16,18,43,24,81,50,23,28,28,19,1099,1146,920,1490,599,638,669,645,645,591,644,629,580,626,547,498,505,555,498,477,453,487,534,547,495,563,553,551\nDIAPH3 - mRNA,153,134,39,19,24,38,63,53,191,123,299,310,500,533,1212,1224,1096,1375,1037,1116,1026,1172,698,697,180,168,910,827,393,392,332,367,949,880,984,904,576,573,1225,1304,1259,1285\nTJP2 - mRNA,36,61,5,14,10,19,16,16,17,14,244,279,318,325,354,331,346,332,284,347,286,328,2028,2107,939,953,1450,1404,1272,1180,1463,1479,1637,1328,1573,1419,1293,1488,1471,1456,1530,1445\nPIK3CA - mRNA,93,121,16,19,46,48,52,43,44,26,1348,1497,1832,1900,777,734,656,675,609,766,673,754,1127,1136,1219,1257,1115,1009,1164,1137,864,984,998,933,951,985,1134,1281,1052,1083,1145,1149\nJUN - mRNA,28,58,10,13,6,6,22,14,87,44,811,915,1189,1262,205,191,169,194,301,276,272,260,686,620,1228,1125,1035,784,289,266,1079,969,756,713,714,777,1066,1164,866,1044,860,890\nOCLN - mRNA,252,416,18,45,102,100,140,112,72,64,58,71,95,113,1838,1809,1616,1653,1919,2401,2235,2217,475,443,787,690,530,444,604,541,534,524,377,275,307,342,550,635,424,497,486,518\nCDX2 - mRNA,354,422,12,77,124,92,92,98,78,60,23,15,18,26,1631,1551,1180,1397,1567,1842,1668,1702,12,10,20,15,16,17,23,18,156,207,161,132,135,148,471,535,193,210,228,228\nRAC3 - mRNA,222,176,9,39,43,44,106,60,126,84,92,135,133,134,2026,1983,1735,2121,1661,1554,1687,1621,184,153,157,153,171,197,127,134,264,313,396,319,344,365,409,396,424,412,399,401\nMAPK7 - mRNA,126,143,7,22,39,39,65,52,84,71,765,971,1212,1154,1100,1096,931,1123,1095,1245,1212,1225,781,748,774,750,915,829,671,601,488,490,475,403,437,426,528,598,521,517,560,535\nTGFB2 - mRNA,88,76,15,9,42,34,52,36,55,48,559,639,613,585,882,893,821,780,1046,1351,1128,1295,29,34,53,39,28,31,40,35,700,735,544,431,480,450,611,681,530,547,540,516\nNLK - mRNA,171,168,8,29,48,55,72,67,81,60,292,342,354,372,1473,1516,1430,1362,950,1258,1062,1309,197,256,267,291,226,233,205,246,354,393,333,318,311,320,362,355,347,306,327,357\nCLDN23 - mRNA,259,180,5,36,46,53,70,68,123,97,739,864,937,936,1351,1397,1220,1296,992,1239,1114,1197,44,52,36,33,40,40,49,37,260,273,181,145,194,165,320,309,212,237,234,253\nCYFIP2 - mRNA,3,18,1,4,3,3,1,2,1,3,41,67,59,52,25,33,36,27,86,81,72,62,327,337,750,742,506,427,897,793,445,532,136,141,134,147,891,942,475,532,507,550\nMYL12A b - mRNA,79,72,2,12,33,35,27,29,46,45,327,471,436,434,485,556,474,436,449,506,512,540,601,626,559,529,505,492,580,576,266,292,463,439,432,442,494,535,441,400,415,443\nPOS_D(2) - mRNA,229,146,242,38,74,115,196,122,115,134,671,642,524,495,462,425,291,441,640,349,385,404,488,417,401,396,441,442,483,393,264,328,330,359,297,350,480,350,372,509,374,398\nMAGI1 - mRNA,51,57,8,6,12,10,30,16,30,28,120,155,162,167,314,272,224,277,326,272,336,286,236,213,339,316,275,225,344,344,76,100,82,76,73,76,135,115,83,104,94,98\nCLDN15 - mRNA,31,29,2,4,4,8,8,9,9,11,41,53,70,70,173,163,150,169,196,222,209,192,188,195,313,248,215,242,216,197,187,200,146,150,145,166,356,366,180,234,206,180\nHCLS1 - mRNA,19,31,2,7,12,10,9,6,8,11,94,92,121,114,184,182,170,164,167,172,195,172,157,146,137,147,128,124,151,128,211,221,164,169,146,151,158,189,176,155,155,137\nARAF - mRNA,25,20,2,5,3,6,14,13,16,11,173,193,235,216,248,284,224,353,149,179,199,201,147,161,153,180,131,157,180,161,218,205,273,222,216,229,241,258,227,238,224,237\nBAD - mRNA,49,22,3,9,14,17,24,23,33,18,319,398,410,360,359,329,291,503,333,302,258,312,292,265,131,136,206,209,132,121,314,337,338,308,328,315,417,419,391,441,407,391\nMYL5 - mRNA,29,44,6,15,13,19,16,17,8,4,293,390,372,371,229,209,212,198,228,263,264,239,244,265,220,274,291,279,325,289,362,406,280,251,265,265,519,539,322,373,342,376\nMYL12A c - mRNA,23,17,8,5,4,6,12,8,11,11,183,218,328,288,123,147,108,140,162,204,189,215,122,98,192,175,167,110,136,128,399,343,247,226,238,218,389,466,343,363,376,362\nAXIN1 - mRNA,43,32,2,10,8,12,15,9,21,14,121,145,202,159,267,264,223,303,253,289,235,273,348,300,451,433,482,441,683,563,458,484,326,299,313,326,360,330,337,404,349,349\nHSPA1A - mRNA,38,16,4,3,8,6,22,18,22,11,147,168,178,184,441,392,348,578,446,405,344,415,314,266,393,303,435,376,188,153,356,396,320,292,324,331,293,335,299,336,292,299\nIL1R2 - mRNA,58,48,5,9,14,14,24,18,38,31,25,37,33,38,405,399,374,379,417,467,436,399,269,285,379,404,354,349,376,382,204,239,311,259,268,287,214,193,278,244,255,262\nFYN - mRNA,20,28,2,3,3,4,7,6,17,6,592,646,726,796,184,185,136,184,163,198,180,187,1510,1442,941,948,873,854,794,698,431,497,253,201,252,224,185,215,201,252,267,217\nVAV3 b - mRNA,24,39,3,13,4,1,9,11,8,4,29,24,32,34,61,69,69,40,63,72,62,62,858,948,1057,1109,1037,966,1352,1338,16,18,29,25,29,28,26,19,13,16,22,22\nPIK3CD - mRNA,10,5,3,6,2,5,4,4,2,3,218,267,288,281,36,43,38,33,68,59,64,73,777,794,942,967,827,826,1047,957,23,28,32,13,19,25,29,34,23,23,24,32\nMRAS - mRNA,21,8,3,3,6,6,11,8,9,10,1138,1399,1684,1520,158,142,147,130,154,173,176,182,408,432,204,201,259,275,130,139,41,45,71,52,62,49,55,54,50,52,54,64\nITGA4 - mRNA,5,2,1,2,1,1,5,1,1,3,638,657,917,874,13,23,34,29,26,28,32,41,28,14,25,18,20,22,18,10,21,16,23,27,24,20,17,17,22,15,14,19\nMAPK10 - mRNA,5,1,2,3,3,3,6,5,3,5,748,986,1246,1224,50,62,56,54,40,56,53,74,44,47,27,31,44,23,31,21,116,141,117,106,107,98,160,177,122,103,131,131\nPDGFD - mRNA,9,8,3,5,2,5,4,5,5,1,1043,1285,1344,1326,78,75,75,69,80,80,81,102,89,97,68,104,81,83,65,89,63,71,68,58,60,58,63,55,55,53,34,71\nBCL2 - mRNA,26,11,5,1,8,7,12,20,29,19,118,156,212,235,228,208,187,234,168,193,207,214,127,104,81,80,136,101,111,118,38,49,50,60,55,57,46,41,60,67,52,69\nEPB41L3 - mRNA,32,25,5,1,1,4,7,9,7,15,28,22,32,31,124,124,124,108,91,76,68,96,27,16,22,20,17,13,7,9,13,16,31,19,20,17,12,20,17,8,12,19\nALPI - mRNA,54,36,2,9,8,7,30,10,8,10,35,14,21,24,212,218,157,180,176,186,164,180,28,22,28,25,24,21,32,24,30,20,26,23,27,21,31,31,25,22,25,18\nHPSE - mRNA,7,4,3,2,1,1,4,3,6,6,111,139,98,119,62,68,65,78,72,60,59,77,36,30,21,31,27,28,17,11,85,94,125,119,101,120,120,106,164,199,177,184\nPIK3CG - mRNA,5,2,3,2,2,2,2,2,4,3,20,25,21,30,20,27,23,23,24,36,21,32,22,21,23,14,23,19,14,19,153,163,152,127,152,115,235,226,235,216,222,225\nRAC2 - mRNA,1,2,1,2,1,1,1,3,4,2,65,101,120,105,20,13,27,27,9,17,19,25,24,17,12,22,13,32,12,17,161,146,118,141,140,122,218,229,144,136,155,158\nCLDN8 - mRNA,1,5,2,1,3,1,2,3,6,2,25,18,26,32,29,28,32,25,32,37,33,38,40,42,39,35,45,49,52,45,32,32,59,47,50,35,49,42,42,41,38,46\nPOS_F(0.125) - mRNA,18,13,26,4,6,12,13,12,14,15,67,54,49,50,54,44,28,52,62,45,44,33,43,39,32,42,42,51,45,35,31,33,27,32,30,37,30,27,34,52,35,40\nCA1 - mRNA,3,2,2,2,1,3,4,1,3,1,24,16,18,18,40,45,47,51,35,36,59,59,39,30,26,25,28,35,17,26,35,32,41,25,29,30,23,14,29,20,14,28\nACTN3 - mRNA,5,4,1,1,2,1,1,3,2,1,14,15,18,18,31,34,23,45,33,48,40,51,19,17,33,25,25,17,21,18,17,23,15,13,11,7,11,13,14,12,7,14\nAPC2 - mRNA,5,8,3,1,2,2,3,4,4,3,11,21,20,31,69,92,54,71,67,61,85,71,22,31,22,23,26,18,25,23,15,21,20,20,24,18,15,22,19,15,16,18\nWAS - mRNA,11,14,1,1,3,1,1,5,3,3,20,15,21,25,61,55,41,52,65,67,52,67,23,21,31,27,25,17,32,24,38,48,30,40,31,29,49,52,50,41,42,43\nMYH11 - mRNA,3,4,1,1,1,1,1,1,1,1,51,82,129,110,18,18,11,17,17,20,12,25,18,15,17,9,21,12,10,11,10,4,12,15,13,10,11,13,6,8,10,18\nNEG_C(0) - mRNA,2,2,2,1,1,1,3,1,3,1,3,8,9,5,4,4,7,9,2,12,9,5,11,13,6,7,15,15,5,10,5,8,9,8,7,8,9,7,8,8,12,11\nNEG_H(0) - mRNA,1,1,1,1,1,1,1,1,1,1,7,5,9,7,3,8,8,2,9,1,6,5,10,6,4,5,8,9,9,3,2,4,8,6,4,2,4,8,7,5,6,2\nNEG_G(0) - mRNA,1,4,1,1,1,1,1,2,1,1,6,6,7,9,9,7,7,5,8,9,5,11,10,9,9,7,7,9,6,12,5,5,5,8,5,5,8,7,9,7,5,5\nNEG_B(0) - mRNA,1,2,4,1,1,1,3,1,1,1,10,13,15,13,6,8,6,8,4,3,2,5,9,7,9,9,9,5,7,7,10,7,13,8,11,16,11,13,13,7,8,13\nPIK3R5 - mRNA,3,3,1,1,1,1,1,1,2,1,12,16,9,9,12,8,12,12,11,11,5,13,11,11,14,6,13,8,10,10,21,8,12,11,13,3,7,9,12,7,6,6\nNEG_D(0) - mRNA,1,1,1,1,1,1,1,2,1,1,17,12,9,6,7,7,7,6,18,6,9,8,11,4,8,5,7,8,5,8,11,8,2,5,18,10,5,9,8,4,2,6\nNEG_E(0) - mRNA,4,4,5,2,1,6,2,4,4,5,17,15,19,23,29,21,25,25,32,26,17,19,17,12,16,28,19,15,11,23,27,13,24,26,22,33,15,23,23,23,17,20\nNEG_A(0) - mRNA,6,5,5,1,1,1,6,3,3,1,16,19,20,13,24,23,17,18,21,25,19,23,29,25,14,17,24,32,24,19,27,21,29,24,40,26,12,22,26,23,18,28\nNEG_F(0) - mRNA,7,3,5,1,4,1,2,2,1,3,18,25,25,30,18,18,15,7,18,17,15,16,22,18,14,11,7,8,14,9,7,12,13,13,12,13,21,10,13,22,15,8\nVAV3 - mRNA,8,2,1,2,1,2,2,1,2,1,13,20,20,26,24,23,24,13,14,13,19,17,12,17,10,8,13,4,17,17,16,14,29,32,24,19,21,23,17,10,17,23\nPRKCB - mRNA,2,2,1,1,1,1,1,1,2,1,20,13,19,22,16,9,6,16,12,17,9,22,14,19,13,14,8,12,16,13,26,18,21,22,27,16,16,12,13,16,18,16\nJAM2 - mRNA,2,3,1,1,1,1,3,1,1,1,169,199,252,198,14,24,6,15,9,12,8,17,10,12,18,15,16,11,6,4,12,13,14,7,11,10,3,10,13,17,9,15\nMYL12A - mRNA,9,10,2,1,3,1,2,1,8,5,91,89,120,104,56,71,60,71,57,91,65,76,69,75,63,52,65,56,62,74,65,66,78,55,75,63,77,88,66,55,76,58\nPOS_E(0.5) - mRNA,45,17,30,8,14,7,20,24,26,22,119,115,96,86,78,72,70,81,130,59,71,70,75,91,67,66,90,79,90,72,57,64,71,71,62,60,83,60,55,88,51,77\nCCND2 - mRNA,27,16,6,5,3,5,16,8,31,45,333,454,648,670,539,548,483,601,271,298,286,324,394,341,269,266,254,250,129,124,25,19,23,21,29,16,10,19,23,13,18,30\nTCF7L1 - mRNA,54,83,5,15,12,24,43,17,29,35,439,608,572,631,643,666,561,642,565,644,718,630,84,68,125,133,140,96,155,154,30,36,25,33,24,21,60,54,46,34,45,30\nITGA7 - mRNA,123,85,2,22,28,30,66,75,29,22,49,56,79,76,1045,1001,915,985,782,854,926,878,41,39,38,35,39,22,29,27,20,25,24,26,21,28,29,29,34,33,26,21\nPDGFA - mRNA,73,113,6,16,29,22,35,33,51,45,76,102,94,75,703,657,582,651,701,695,692,677,128,85,145,155,123,94,104,78,356,306,115,104,136,97,205,235,145,186,188,162\nLEF1 - mRNA,39,20,4,6,7,7,25,15,39,22,102,112,107,120,792,775,711,921,348,460,423,497,34,30,28,32,24,25,9,20,22,37,22,12,23,14,63,94,31,16,26,25\nCGN - mRNA,485,516,27,59,101,124,227,182,244,173,8,14,22,12,3977,3923,3304,3994,2873,3114,2911,3016,64,56,32,29,32,33,42,26,1355,1330,851,682,761,768,1294,1387,1019,1113,1013,1060\nITGA3 - mRNA,29,76,2,13,17,22,11,13,10,4,1557,1819,1906,2065,188,205,146,172,197,210,178,174,155,133,36,31,76,78,15,28,3473,3477,2000,1789,1928,1908,2786,2598,2271,2557,2348,2281\nPOS_C(8) - mRNA,943,602,900,158,349,331,853,551,689,623,2359,2647,2058,1752,1802,1585,1104,1655,2774,1420,1509,1441,1835,1588,1441,1511,1568,1616,1746,1531,1001,1299,1400,1264,1229,1349,1752,1424,1301,1888,1347,1413\nPXN - mRNA,206,198,12,33,56,52,89,90,146,119,1682,1834,2369,2310,2133,2054,1706,2277,1689,1841,1922,1985,1018,973,971,859,907,835,899,863,1738,1800,1563,1358,1511,1446,1786,1868,1660,1826,1787,1740\nMPP5 - mRNA,381,374,35,57,103,160,127,132,179,155,1333,1643,1915,1908,2441,2420,2214,2259,1850,2341,2083,2434,1657,1696,1130,1102,1442,1290,1157,1174,1190,1227,1401,1119,1254,1221,1494,1635,1297,1337,1325,1276\nPDCD4 - mRNA,307,450,7,65,117,139,95,119,23,19,1155,1405,1410,1434,1571,1680,1545,1318,1793,2349,1992,2227,783,859,1269,1386,1283,1197,1714,1708,1121,1274,962,841,940,955,1861,2006,1555,1475,1592,1568\nWASL - mRNA,276,382,19,68,113,109,130,126,116,90,920,1074,1212,1304,2320,2330,1877,2016,1867,2220,1928,2201,1244,1367,1800,1875,1336,1259,1549,1439,1917,2162,1539,1277,1461,1374,1853,2080,1672,1750,1771,1842\nSYMPK - mRNA,232,245,9,30,66,67,87,100,80,61,617,732,863,922,1743,1791,1474,1626,1400,1650,1591,1703,1356,1337,1550,1500,1316,1209,1344,1297,1628,1706,1725,1533,1742,1678,1794,2039,1736,1829,1816,1703\nASH1L - mRNA,204,236,21,34,56,74,105,85,91,70,829,1021,1207,1253,1794,1854,1559,1719,1432,1681,1640,1804,1014,1066,1829,1851,1387,1197,1709,1637,1911,1997,1659,1515,1624,1657,1983,2207,1904,1933,1936,1949\nERBB2 - mRNA,144,160,4,44,65,47,73,73,39,40,524,612,743,754,1099,1059,899,1203,1364,1375,1235,1295,1426,1396,1847,1798,1917,1784,2676,2396,1515,1642,1012,914,1019,1013,2515,2479,1493,1684,1673,1615\nPRKCD - mRNA,200,238,5,32,57,79,83,81,79,70,481,559,588,633,1676,1645,1396,1761,1620,1661,1560,1694,1211,1098,1825,1679,1545,1529,1872,1708,1209,1283,1158,1034,1114,1115,2006,2059,1417,1562,1492,1493\nGUSB - mRNA,250,311,8,72,88,81,128,113,96,59,379,454,495,462,2508,2435,2181,2281,2801,2989,2880,2864,1223,1242,1268,1180,1441,1417,1567,1296,1214,1425,1505,1261,1426,1354,1698,1672,1790,1912,1820,1919\nCSNK2A2 - mRNA,300,406,19,31,106,115,134,94,183,120,593,651,754,778,2152,2163,1906,2250,2070,2299,2174,2239,1264,1183,2306,2417,1354,1254,1993,1669,929,1075,595,514,569,540,1127,1186,943,972,924,981\nBRAF - mRNA,323,514,28,73,146,129,153,128,101,90,579,796,890,884,2343,2372,2144,2258,2227,2691,2457,2595,969,977,1682,1613,1161,1042,1542,1380,1168,1252,942,810,857,920,1245,1396,1083,1138,1140,1136\nTJP3 - mRNA,268,214,7,40,74,85,105,83,75,59,17,11,16,9,1812,1864,1563,1926,1705,1930,1718,1831,301,281,439,450,310,310,466,437,1928,2044,1368,1313,1327,1347,2634,2741,1573,1553,1454,1619\nCRB3 - mRNA,293,283,16,55,96,95,132,105,204,146,17,31,21,37,2337,2395,2039,2432,1903,2138,2070,2176,2044,1934,1180,1153,1488,1529,1309,1298,2248,2329,2130,1878,2007,1919,2403,2365,2243,2369,2293,2515\nPAK4 - mRNA,255,207,20,28,69,75,101,111,125,111,326,339,409,407,2254,2288,1957,2151,1770,1974,1864,2082,1228,1232,1092,975,1340,1396,1439,1190,1818,2012,2467,2247,2470,2346,2638,2740,2398,2706,2480,2423\nPIK3R1 - mRNA,298,276,11,51,100,115,108,108,71,65,316,459,427,491,1847,1797,1827,1648,1397,1862,1717,1830,1065,1068,1218,1202,1187,1142,1314,1306,1969,2205,2331,1919,2165,2153,2659,2852,1922,1933,1941,1904\nCTTN - mRNA,293,231,15,43,60,75,102,119,174,144,1617,1967,2028,2079,2357,2231,2094,2336,1363,1651,1680,1807,1247,1232,1126,1167,1005,1028,920,924,2078,2201,2645,2345,2487,2566,2671,2870,2313,2238,2434,2472\nGSK3B - mRNA,227,279,34,41,57,72,116,107,140,108,1335,1463,1731,1831,1947,2061,1726,1848,1437,1794,1616,1790,1833,1898,1874,1777,1701,1625,1557,1329,2555,2686,2132,1881,1954,1893,2370,2669,2351,2441,2434,2492\nAGO2 - mRNA,80,95,8,13,26,21,49,36,79,64,563,685,749,733,974,971,810,1094,782,865,905,861,766,769,1191,1153,796,785,1018,951,2280,2459,2771,2435,2754,2615,1966,2010,2267,2594,2502,2414\nNUMA1 - mRNA,97,109,4,15,28,18,82,67,56,42,603,731,859,872,1470,1523,1338,1653,984,1305,1352,1211,888,809,746,752,1066,990,1292,1211,1689,1779,1582,1410,1474,1427,2466,2592,1987,2113,2085,2327\nDGCR8 - mRNA,171,184,20,36,52,53,101,70,121,80,659,758,839,808,1711,1599,1411,1559,1211,1545,1493,1459,1410,1365,1309,1174,1047,1073,942,909,1775,1971,2332,2096,2203,2217,1770,1818,2205,2322,2157,2234\nEGFR - mRNA,228,447,7,49,107,111,90,94,100,74,1532,1858,2442,2386,1094,1060,929,934,1629,1745,1505,1550,580,600,1036,858,625,616,943,953,2500,2757,2289,1994,2150,2139,2041,2091,1990,2285,2236,2083\nPDGFC - mRNA,78,115,1,6,55,30,26,25,12,15,1549,1828,1552,1657,647,664,606,536,483,583,570,579,960,975,1011,1049,979,897,1221,1140,1765,1818,2238,1883,2151,1982,2061,2039,1632,1611,1557,1631\nIL1R1 - mRNA,373,137,21,21,62,59,118,128,101,91,2060,2453,2667,2588,2534,2755,2474,2502,1154,1492,1513,1771,3350,3559,6447,6386,4870,4798,4986,4939,133,145,100,80,73,69,93,91,104,84,94,80\nEPB41 - mRNA,146,171,6,26,53,72,65,90,90,72,153,210,226,258,2053,2102,1844,2165,1472,1841,1795,1883,3120,3194,2846,2893,2659,2490,2612,2345,2279,2330,2375,2140,2312,2285,2715,2890,2232,2253,2240,2293\nEPB41L1 - mRNA,403,300,79,44,45,53,142,114,606,370,635,822,871,822,1759,1744,1324,2246,1652,1697,1628,1642,3126,2859,1619,1534,2562,2625,2046,1786,2071,2034,1760,1492,1732,1679,1854,1829,1593,1903,1715,1805\nCDK4 - mRNA,213,163,13,39,70,63,74,82,146,89,677,866,919,1016,2215,2299,2193,2136,1383,1724,1661,1768,2690,2583,1663,1779,1691,1704,1363,1288,1182,1344,2284,2005,2163,2081,1245,1262,1575,1668,1616,1634\nXPO5 - mRNA,218,166,16,42,49,70,81,97,141,104,543,539,747,641,2167,2185,1992,2107,1282,1770,1468,1790,2592,2591,2377,2404,2136,1987,2161,2001,1086,1151,1619,1399,1635,1536,1098,1273,1361,1355,1324,1391\nPPP2R2D - mRNA,202,222,7,40,49,67,83,64,86,74,761,900,969,1082,1886,1913,1620,1869,1513,1686,1684,1702,2032,2045,1908,1825,1937,1922,2017,1947,1319,1464,1600,1475,1505,1437,1422,1561,1410,1483,1402,1419\nPOLR2A - mRNA,114,132,9,25,36,34,81,54,90,67,1754,2108,2469,2490,1567,1539,1365,1955,1262,1622,1498,1527,3680,3325,2741,2439,2735,2708,2307,2047,1250,1353,1328,1222,1288,1307,1417,1485,1400,1551,1532,1557\nAKT1 - mRNA,107,92,4,16,28,35,70,53,103,76,1396,1711,1838,1804,1413,1387,1194,1705,1022,1070,1121,1107,2230,2091,1594,1525,1952,1953,1514,1453,1298,1446,1390,1382,1348,1382,1522,1419,1593,1662,1611,1664\nGNAI3 - mRNA,296,290,22,47,85,94,98,94,167,128,1910,2187,2486,2525,2146,2142,1932,1824,1643,1930,1805,1910,3051,3153,2181,2097,2567,2464,2406,2398,1616,1685,3183,2801,3101,2940,2073,2144,2454,2504,2561,2466\nDVL3 - mRNA,285,308,17,37,69,75,135,95,168,103,1652,2033,2146,2142,2430,2395,1943,2209,1897,1940,2052,2136,1842,1884,2521,2558,2065,1995,2378,2147,1882,1959,1607,1499,1570,1584,1866,1944,1833,1931,1776,1907\nPPP1R12A - mRNA,298,363,26,60,89,118,124,117,140,95,1961,2286,2880,3108,2179,2299,2037,2162,1704,2102,1846,2142,2058,2035,1779,1835,1921,1812,1803,1759,1953,2057,2256,1965,2237,2054,2178,2428,2164,2210,2273,2152\nMAPK8 - mRNA,558,363,32,38,110,117,218,183,196,152,1059,1200,1458,1497,3973,4071,3804,4130,2337,2809,2836,3152,1791,1884,2273,2226,1702,1677,1707,1762,1899,1991,2077,1846,2040,1974,1809,1733,1959,1963,1823,2033\nCRK - mRNA,310,302,29,39,76,106,189,146,251,206,2134,2572,2787,2718,3718,3652,3254,3674,2690,3028,3134,3247,2454,2320,2478,2533,2072,2166,2014,1887,1622,1779,1566,1299,1531,1485,1359,1319,1448,1511,1378,1369\nRAF1 - mRNA,367,347,30,54,83,102,145,146,156,118,1609,1781,2057,2016,3162,3139,2740,3265,2345,2745,2481,2907,1880,1912,1959,1809,1948,1828,2066,1865,929,987,1267,1103,1140,1136,973,997,1075,1157,1173,1114\nVAV2 - mRNA,341,464,25,69,131,138,161,154,134,122,645,789,871,870,2703,2696,2257,2531,2470,2967,2671,2808,2107,2061,3126,3026,2437,2200,3320,3072,1271,1476,1320,1088,1283,1266,1418,1506,1680,1836,1826,1739\nTBP - mRNA,279,289,21,57,81,85,129,138,140,118,772,910,1165,1206,2421,2434,2285,2304,1788,2202,1961,2320,3091,3256,2829,2902,3018,2855,3148,3035,1438,1524,1605,1451,1571,1502,1686,1891,1732,1692,1735,1770\nDICER1 - mRNA,315,326,21,44,84,102,155,129,158,142,1144,1431,1706,1670,3143,3099,2708,2901,2251,2541,2526,2645,3898,4177,3974,3828,3654,3584,3824,3595,993,1009,1199,1050,1148,1140,1027,1084,921,964,1001,907\nDNMT3A - mRNA,364,448,13,71,122,168,157,185,100,98,388,448,497,521,2958,3044,2645,2655,2235,2700,2708,2781,3662,3736,3705,3547,3159,3204,3547,3245,847,958,683,615,643,700,1156,1248,907,920,957,948\nPPP1CC - mRNA,578,585,24,85,210,253,191,207,233,180,803,919,1019,1100,4138,4158,3637,3926,3099,3333,3282,3627,3872,3971,4118,3916,3269,3366,3952,3668,1624,1864,3034,2746,2897,2914,2712,2586,2603,2621,2691,2738\nCREBBP - mRNA,342,373,30,46,107,103,162,160,173,142,1586,1986,2495,2372,3041,3061,2612,3003,2407,2674,2658,2648,2523,2595,4471,4280,2913,2636,4104,3694,2631,2657,1878,1672,1811,1764,1888,1956,1950,2147,2116,2129\nSOS1 - mRNA,723,515,31,72,142,149,226,208,127,96,1154,1325,1712,1757,3607,3523,3160,3430,3001,3604,3624,3609,2817,3097,4506,4489,3533,3311,4887,4475,1529,1505,1301,1152,1231,1252,1722,1864,1508,1619,1612,1551\nEP300 - mRNA,457,417,25,49,100,92,204,180,226,140,1148,1399,1610,1641,3845,3973,3363,4054,2948,3473,3484,3258,2595,2541,4390,4285,3102,2747,3922,3546,1985,1909,1630,1508,1602,1559,1704,1820,1589,1823,1678,1732\nSMAD2 - mRNA,201,200,8,17,65,64,91,68,93,79,1880,2241,2495,2582,1494,1436,1239,1305,1143,1333,1300,1397,3717,3868,4032,3872,3527,3232,3540,3428,2566,2723,2918,2457,2860,2653,2766,2803,2618,2995,2763,2749\nPRKCA - mRNA,214,187,14,29,43,41,109,79,96,64,2384,2902,3573,3568,2173,2182,1926,2224,1392,1735,1727,1711,3208,3340,3707,3562,3380,3092,3423,3210,1957,2092,1291,1008,1130,1073,2938,3686,2098,2385,2333,2229\nMAPK3 - mRNA,344,315,9,58,74,110,115,122,114,92,2972,3369,3638,3577,2681,2588,2263,2413,2197,2460,2428,2480,3080,3124,2185,2111,3259,3398,3201,3049,2166,2407,1474,1377,1445,1377,3167,3186,1936,2043,2108,2110\nRRAS2 - mRNA,395,480,43,58,170,198,173,181,236,209,2375,3059,3588,3440,3113,3076,2622,2802,2290,2818,2580,2827,2683,2865,2837,2723,2108,2045,2341,2294,3068,3275,2962,2565,2739,2743,3046,3299,2529,2552,2692,2633\nMAP3K7 - mRNA,426,445,36,68,137,126,184,171,192,154,2562,2980,3311,3582,3393,3626,3217,3142,2644,3311,3091,3390,4020,4050,3043,2998,3391,3407,3363,3174,2219,2419,2939,2576,2838,2770,2533,2731,2475,2511,2505,2420\nROCK1 - mRNA,492,406,41,49,113,144,220,161,294,205,2224,2788,2927,2913,3403,3445,3118,3240,2441,2691,2690,2824,2803,2819,2660,2600,2756,2533,2805,2594,1957,1976,2170,1856,2132,1911,2372,2378,2225,2318,2211,2207\nBRK1 - mRNA,440,314,11,61,110,109,166,134,168,98,2568,3072,3346,3116,3222,2980,2830,3278,3108,2961,2599,3158,3222,3293,3183,2893,3089,3045,2970,2690,1450,1624,2120,2027,2026,2046,1936,1943,1752,2146,2016,1853\nTP53 - mRNA,27,56,5,13,13,13,32,11,15,13,1860,2204,2628,2704,278,286,280,335,280,306,330,348,3617,3540,3058,3049,3390,3207,2854,2678,3628,3694,4767,4189,4668,4669,3809,3901,5319,5680,5469,5437\nCCNG2 - mRNA,331,519,5,70,113,79,135,115,50,34,1435,1799,1954,1806,1821,1782,1509,1479,3127,3360,3126,3040,663,700,1448,1312,1793,1581,1648,1495,2000,2141,1829,1532,1746,1725,3634,3634,4196,4476,4295,4263\nYBX3 - mRNA,302,221,30,41,83,81,165,153,349,219,1293,1527,1633,1598,3801,3610,3258,3452,2745,2890,2766,3044,2098,1975,2724,2897,1778,1721,2046,1879,3002,3302,3541,3135,3427,3371,3484,3458,3419,3771,3529,3607\nDROSHA - mRNA,329,229,23,46,75,84,152,155,196,153,1438,1639,1908,1998,3486,3417,3026,3271,2267,2670,2787,2787,2722,2855,1795,1785,2514,2518,2201,2033,2345,2367,3701,3155,3650,3466,2406,2534,3156,3338,3205,3276\nPTEN - mRNA,364,455,22,64,156,162,135,144,129,134,1884,2231,2875,2902,2637,2646,2421,2410,2200,2504,2326,2569,2094,2088,3263,3231,2208,2050,2413,2335,3166,3615,3069,2706,2913,2926,4250,4509,3655,3943,3857,3762\nSPRY2 - mRNA,307,243,28,37,89,90,131,145,300,208,1301,1361,2399,2533,3495,3410,3091,3152,2231,2684,2538,2780,1479,1476,1621,1599,2037,1747,2003,1840,3648,4020,2640,2277,2577,2344,4261,4444,3727,3737,3722,3735\nSMAD3 - mRNA,322,453,19,69,86,91,152,115,164,138,4820,5835,7516,7593,2028,2038,1774,1912,1824,2347,2166,2169,2099,2141,4520,4340,3056,2837,4943,4592,5420,5290,3761,3340,3597,3644,4736,5351,3926,4214,4008,4113\nTRIO - mRNA,350,361,37,49,93,110,131,153,161,125,4230,4937,6057,6250,2060,2147,1799,1970,1628,1961,1851,1961,1818,1924,2522,2521,2281,1939,2484,2361,2211,2438,1801,1490,1739,1615,2261,2485,2062,2004,2126,2070\nGNAI1 - mRNA,8,18,1,2,4,9,7,8,8,8,3955,4666,5055,5268,327,296,355,315,229,286,236,304,960,985,1130,1116,587,577,572,623,2330,2449,3192,2744,2971,2912,3002,3118,2368,2402,2401,2514\nG6PD - mRNA,110,42,16,7,7,15,51,50,159,120,4123,4792,4724,4756,1456,1285,1202,1612,812,766,866,916,3058,2914,989,856,2027,2021,1194,1035,2189,2067,4270,3848,4147,4095,2207,2131,3310,3773,3699,3618\nRAB13 - mRNA,1329,1039,60,168,392,513,569,558,406,341,9770,11579,12800,13113,8925,9116,8123,8237,7567,8337,7788,8608,4594,4520,3525,3559,3641,3564,3300,3070,5011,5528,5171,4565,4826,4770,6042,6182,5362,5472,5387,5378\nCD9 - mRNA,1094,824,61,216,267,323,346,314,479,379,6981,7837,9105,9168,8703,8419,7943,8269,7735,8836,7821,8629,684,637,998,969,677,697,755,653,5777,5855,6308,5754,5825,6073,6239,6089,5727,5798,5745,5754\nPSEN1 - mRNA,658,672,41,106,182,240,263,229,300,249,2555,3118,3312,3334,4809,4972,4375,5009,4022,5059,4835,4934,2969,2854,3686,3651,3038,2952,3522,3488,2628,2832,2237,2020,2130,2043,2764,2909,2456,2484,2545,2628\nPPP2R2A - mRNA,769,878,42,116,228,245,363,289,445,345,1926,2253,2756,2547,6143,5957,5178,5259,5680,6493,6263,6425,3203,3223,2744,2807,2949,2832,2677,2428,1761,1900,2101,1781,1906,1926,2193,2363,1975,2119,2060,1989\nDOCK1 - mRNA,1122,800,64,120,261,293,415,382,378,380,2249,2855,3097,3154,7759,8021,7615,7493,5477,7004,6689,6853,2338,2390,2653,2655,2495,2315,2682,2645,2914,3084,3420,2973,3245,3169,2736,3130,2990,3118,2984,3014\nPPP2R1B - mRNA,889,720,68,131,231,239,384,367,478,360,1545,1730,2057,2108,7118,7215,6302,7323,5079,6434,6045,6443,4709,4624,2625,2401,3886,3703,3846,3555,2289,2506,3995,3486,3854,3787,2750,3033,3001,3157,3123,3093\nMAPK9 - mRNA,785,717,58,104,222,260,296,304,423,363,2340,2765,3209,3294,6860,7111,6206,6746,4896,5809,5718,6405,4647,4997,3260,3163,4231,4064,3818,3589,3437,3505,3399,3023,3434,3279,4347,4665,4377,4454,4432,4475\nCASK - mRNA,1533,1499,101,196,492,545,581,570,847,663,2679,3157,3452,3541,10318,10558,9549,9764,7327,9409,8810,9958,3131,3268,3795,3957,3211,2992,3640,3513,2243,2376,1908,1699,1770,1812,2404,2583,2301,2225,2350,2430\nKRAS - mRNA,1340,888,103,127,275,271,409,448,650,525,1148,1383,1549,1459,9253,9645,8722,9277,6326,7732,7311,7730,1484,1467,979,1000,1146,1028,767,734,1835,2045,2592,2124,2445,2442,2676,2928,2418,2485,2460,2511\nPOS_B(32) - mRNA,3563,2122,3429,541,1284,1212,2515,1891,1999,1944,8264,8671,7032,6234,6249,5074,3999,5781,10181,4893,4924,5069,6469,5970,5311,5086,5708,5891,6369,4909,3362,4324,4763,4530,4256,4581,6067,4884,4789,6795,4822,5172\nCLTC - mRNA,693,523,51,102,155,187,199,234,364,301,3970,4470,5017,5120,5710,5822,5147,5849,4015,5096,4522,5214,8160,8384,4761,4692,6542,6271,5145,4943,3639,3807,5201,4472,4966,4678,4496,4804,4432,4470,4385,4462\nRHOA - mRNA,616,352,31,63,124,144,288,219,390,305,5208,6005,7163,6531,5619,5322,4780,5760,5307,4991,4429,5271,5866,5936,4358,4006,4912,5145,3935,3501,2971,3301,4723,4501,4509,4548,4423,4548,4514,5042,4718,4866\nSPTAN1 - mRNA,690,593,56,109,132,166,307,294,642,454,5074,5729,6367,6316,6190,6100,5447,6385,4423,5171,4770,5194,5773,5658,4928,4827,5429,5515,3997,3686,3861,3936,3718,3357,3598,3447,3380,3645,3393,3635,3641,3641\nCSNK2A1 - mRNA,432,504,28,73,143,163,264,201,327,249,1697,2015,2475,2520,5361,5444,4863,5401,4564,5515,5347,5590,8625,8323,7196,7324,7292,7118,7534,7330,3865,3868,5178,4541,4869,4749,4827,4762,4742,4638,4691,4952\nMAPK1 - mRNA,806,854,44,149,266,280,318,333,421,328,4108,4827,5572,5522,6403,6515,5795,6135,5096,6308,5815,6561,7872,7981,7861,7874,7476,7028,7704,7505,2793,2841,3803,3327,3726,3561,3638,3995,4088,4212,4400,4332\nAXIN2 - mRNA,498,203,47,26,52,73,267,182,413,266,127,169,187,172,6212,5985,5299,7177,2751,2931,2756,3374,5287,4987,7927,7376,6647,6170,8032,7395,1971,2346,1497,1431,1446,1461,2236,2535,1469,1749,1618,1652\nPARD3 - mRNA,806,894,48,94,237,291,298,257,405,302,1954,2353,2744,2754,5238,5102,4770,4938,4378,5512,4918,5208,4715,4910,7771,7636,5046,4846,6352,6113,2808,3142,2669,2256,2621,2494,3679,3970,3444,3488,3416,3553\nMLLT4 - mRNA,635,643,49,73,177,170,287,250,323,237,438,502,656,692,4220,4141,3522,3966,3720,4072,3812,4152,4915,4973,8765,8477,6252,5480,9070,8361,4011,4509,3477,3063,3227,3302,4678,5185,4090,4709,4489,4491\nMYC - mRNA,341,216,14,49,117,108,184,154,130,118,537,589,937,1003,3078,2918,2705,2839,2069,2312,2589,2460,3887,3836,6111,6337,3873,3594,6162,5762,2567,2827,3728,3344,3612,3528,4168,4444,3778,3708,3821,3887\nSRC - mRNA,552,579,20,95,187,164,242,196,192,140,823,963,1050,1026,3980,4039,3364,3812,3350,3723,3379,3801,3867,3695,5913,5757,4041,4041,4992,4562,4033,4530,3116,2947,3065,2968,4684,4881,3748,4084,3919,3979\nABCF1 - mRNA,386,279,27,31,71,70,188,152,282,198,1822,2247,2360,2424,4720,4715,4156,4979,2898,3310,3471,3456,7029,6982,5176,5088,5488,5394,4794,4610,2271,2575,3738,3393,3591,3603,2476,2607,2894,3075,2943,3025\nNRAS - mRNA,403,329,30,47,101,138,164,157,267,224,1896,2026,2239,2202,3633,3701,3277,3946,2515,2884,2938,3148,6110,5757,3556,3472,4466,4367,3726,3594,2962,3144,3609,3262,3544,3407,2690,2893,2624,2708,2766,2707\nPAK2 - mRNA,476,427,39,44,96,131,212,154,342,238,2543,2940,3517,3458,4025,4057,3396,4184,2908,3327,3367,3632,4540,4730,4307,4183,3833,3455,4062,3970,3169,3277,3580,3102,3459,3274,2925,2992,3076,3195,3180,3102\nHPRT1 - mRNA,861,687,54,111,221,248,338,265,410,266,1192,1453,1683,1700,5401,5397,4853,5002,4519,5735,5253,5239,3831,4028,3360,3112,4129,3884,4904,4911,1578,1881,4940,4364,4829,4649,3948,4061,5054,5229,5149,5313\nEXOC4 - mRNA,498,545,40,100,175,183,223,209,192,136,1574,1931,2379,2333,3522,3706,3226,3543,3288,4041,3809,3962,2409,2303,2871,2678,2847,2570,3334,3121,3849,4040,4088,3549,3890,3817,4920,5424,4836,5078,5076,5313\nPRKCI - mRNA,429,492,37,68,135,157,184,157,276,193,1893,2261,2634,2705,3486,3315,2937,3198,2714,3359,3112,3311,2502,2474,1860,1745,2223,2181,1929,1899,3883,4010,4314,3585,4067,3882,4363,4513,4349,4715,4624,4528\nTCF7L2 - mRNA,447,596,25,87,158,143,158,149,131,116,960,1162,1314,1346,3322,3349,2956,3217,3136,3825,3541,3593,1988,2099,3219,3016,2506,2357,4534,4077,3833,4106,3408,2999,3260,3161,3670,4038,3548,3581,3482,3632\nCTBP1 - mRNA,334,275,13,41,91,104,201,172,184,161,1758,2123,2476,2330,3652,3464,3185,3635,2763,2927,2913,3091,4329,4143,3809,3776,3754,3643,3602,3541,4235,4458,4265,3898,3947,4086,4149,4112,3879,4435,4121,4160\nAKT2 - mRNA,402,319,20,42,96,122,208,163,211,148,1586,1839,1967,2013,4127,4124,3536,4118,2679,2960,3074,3029,3182,3009,3343,3149,3077,3089,3644,3357,3518,3884,4728,4351,4409,4440,4055,4025,3733,4308,3836,3912\nITGA6 - mRNA,296,219,23,25,40,56,139,102,185,119,569,580,1099,1076,2893,2686,2596,2597,1635,1853,1826,1966,8961,9216,10199,10006,7177,6855,7067,6766,10735,11365,10496,9123,9954,9528,7841,7813,9013,9847,9091,9131\nITGB5 - mRNA,433,234,39,47,72,79,198,163,281,194,6633,7693,9371,9435,3920,3649,3352,3989,2455,2639,2571,3037,7474,7361,6327,6071,6268,6252,4743,4527,7630,7770,7914,6752,7479,7222,7884,8059,8922,10272,9564,9580\nPTK2 - mRNA,689,743,38,100,215,219,220,202,180,146,1751,2068,2268,2431,4548,4613,4246,4350,3757,4595,4245,4553,2998,3168,3174,3095,3152,3109,3685,3421,8962,9412,9062,8090,8574,8394,10687,11188,9320,9365,9455,9745\nPRKAA1 - mRNA,429,549,32,91,166,186,188,166,162,126,3651,4166,4612,4683,3512,3719,3161,3148,2797,3452,3037,3239,2993,3200,4041,3849,2961,2655,3446,3197,8406,8943,7040,5885,6653,6333,8666,9235,7365,7982,7656,7786\nWASF2 - mRNA,709,827,45,109,211,270,286,287,306,248,4008,4648,5608,5656,5062,5011,4364,4572,4411,5507,4964,5557,4447,4635,4380,4416,4893,4501,5606,5166,7392,7652,6969,5570,6635,6355,8924,9790,8407,9031,8957,8921\nVAPA - mRNA,579,609,33,101,180,241,174,201,334,240,2813,3115,3725,3882,3886,3977,3609,3760,3021,3626,3266,3815,6208,6236,4547,4525,4635,4567,4412,4249,4838,5346,6501,5785,6138,5906,6756,6964,6255,6383,6516,6770\nMAP2K1 - mRNA,524,414,47,74,130,152,208,162,436,312,2244,2450,3197,2990,3568,3571,3131,3694,3144,3592,3314,3744,3180,3076,2658,2508,3601,3542,4053,3812,4034,4365,6090,5216,5569,5566,7205,7360,7043,7257,7230,7631\nHSPA1B - mRNA,411,241,23,42,38,68,205,165,293,210,945,1245,1440,1437,4742,4460,3920,5222,3349,3613,3671,3765,6130,5872,5406,4894,6467,5942,3035,3024,6161,6461,6381,5738,5943,6192,5888,5760,4870,4965,5109,4937\nYES1 - mRNA,858,980,71,125,289,325,302,350,457,336,1893,2084,2464,2650,5722,5748,4887,5556,4310,5314,4832,5511,4707,5022,4639,4529,4794,4444,4967,4835,5025,5455,6092,5185,5864,5584,5898,6427,5878,6087,6190,6209\nCRKL - mRNA,493,529,24,95,125,140,194,184,276,198,1656,1900,2306,2437,4748,5217,4631,5514,3213,4859,4656,4615,5080,4644,5239,5054,5981,5596,6078,5993,4286,5138,4968,4270,4764,4730,5879,6336,5555,5017,5417,5698\nDIAPH1 - mRNA,893,605,91,117,154,186,362,330,731,542,1424,1533,1876,1986,7679,7814,6770,7981,4913,5766,5851,6144,6594,6332,4268,4283,5521,5369,4446,4314,4735,4837,8073,6902,7757,7561,6139,6338,7294,7834,7459,7609\nTJP1 - mRNA,762,616,96,110,175,210,288,297,663,565,4483,5276,6011,6167,6865,7183,6330,6741,4667,5691,5324,6137,3841,3790,4028,4186,3029,3091,2752,2729,7910,8325,7503,6467,7318,7187,6265,6753,7139,7195,7149,7477\nF11R - mRNA,1450,1181,61,234,347,386,518,496,610,450,67,79,103,96,10619,10975,9659,9969,8198,9637,9252,9655,2884,2929,2549,2467,3054,2946,3334,3024,5028,5778,5795,5056,5525,5279,6437,6986,6142,6047,6086,6375\nCLDN4 - mRNA,1081,1026,50,225,249,274,461,375,811,508,25,37,53,54,8360,8017,7170,8888,7862,7965,7713,7969,3178,2826,1492,1511,1854,2036,1318,1294,9124,9546,9012,8713,8376,8675,11633,11471,8573,9468,8709,8900\nPAK1 - mRNA,955,1595,41,247,440,552,407,408,320,241,1447,1668,2079,2187,6402,6640,5922,6129,6183,7547,6687,7350,297,300,369,351,293,286,335,298,6801,7072,7008,6262,6727,6697,9467,10195,7915,7959,8053,8431\nITGB4 - mRNA,103,79,3,17,28,30,31,37,48,25,33,35,26,24,582,628,564,556,578,644,658,684,140,113,153,140,152,136,138,136,6912,6933,4567,3953,4462,4244,6445,6548,6112,6602,6423,6214\nCLDN3 - mRNA,214,185,8,32,67,47,98,77,57,40,8,13,15,21,1335,1216,1036,1190,1854,1991,1734,1926,47,47,37,58,62,57,106,86,3223,3770,3403,3324,3197,3139,6227,6234,4161,4706,4514,4402\nINADL - mRNA,327,344,51,52,99,121,163,125,154,109,180,211,274,228,2411,2523,2245,2656,2157,2571,2567,2490,1110,1087,1926,1880,1379,1252,2222,2011,4704,4769,3652,3080,3400,3329,4575,4976,4524,4817,4582,4716\nBAIAP2L1 - mRNA,461,417,28,61,117,117,125,125,204,152,360,392,429,435,2655,2810,2404,2440,2238,2442,2355,2449,1728,1787,1629,1671,1311,1267,1574,1440,3549,3824,3222,2906,3118,3117,3792,4423,3586,3874,3826,3917\nCTBP2 - mRNA,595,401,30,66,146,171,247,231,321,272,2093,2455,2932,3049,5521,5675,5149,5361,3840,4756,4230,5073,62,83,103,94,111,79,88,77,5938,6384,6131,5181,5797,5589,4418,4748,4986,5340,5193,5275\nCLDN7 - mRNA,432,384,36,91,87,86,168,165,344,242,19,22,18,25,3899,3653,3293,3933,3422,3544,3200,3503,20,32,18,16,29,17,29,26,8041,8813,5204,4987,4908,4926,7191,7345,6257,7154,6510,6644\nMET - mRNA,605,687,37,97,182,186,270,226,182,146,414,383,596,589,4108,4016,3542,3669,3637,4143,3934,3987,2190,2213,3850,3832,2093,1873,2805,2399,7390,7693,7172,6062,6645,6437,6483,6522,6956,7782,7211,7285\nITGA1 - mRNA,53,32,3,6,8,15,23,25,15,28,17529,21897,24302,23643,688,666,587,580,380,436,409,482,2527,2556,6393,6322,3043,3024,2890,2598,3429,3667,2623,2227,2491,2589,5285,5402,4116,4105,4211,4447\nVCL - mRNA,288,186,45,24,49,50,118,103,393,308,13359,15659,17406,16585,2617,2679,2315,2812,2002,2250,2093,2524,2145,2222,933,859,1774,1519,1065,1068,1188,1237,1982,1734,1842,1803,1533,1603,1759,1893,1795,1807\nGNAI2 - mRNA,767,589,44,111,200,254,270,291,360,246,16582,19690,21118,21394,5697,5865,5363,5485,4134,5045,4875,5253,7674,7626,3202,3173,6060,6002,3684,3567,8439,8852,10930,9321,10507,9936,9950,10607,10078,9792,10237,10211\nMYH9 - mRNA,963,734,110,121,175,213,501,446,881,682,15179,18439,18834,19361,8923,8345,7287,8587,6154,6749,6561,7079,7384,7524,6731,6501,7381,7198,8113,7681,3974,4144,5645,4927,5493,5091,7348,8017,6047,6453,6134,6198\nCD44 - mRNA,52,33,1,11,10,17,47,27,47,30,36834,41486,39098,38312,1164,1118,1114,1248,754,722,698,752,216,94,109,124,103,83,119,110,9665,12330,7797,7284,7258,7588,12294,12014,9113,10209,9184,9815\nFLNA - mRNA,614,406,100,99,120,136,309,277,1006,726,28064,35796,35518,35297,5701,5589,4937,5824,4801,5798,5255,5367,8451,8314,5231,4910,6677,6825,4111,4139,3593,3120,4314,3870,4334,4076,2331,2534,3259,3579,3359,3379\nMYLK - mRNA,46,32,6,11,6,13,30,24,22,27,26587,34565,30162,29931,581,670,564,665,522,596,575,620,1835,1883,1439,1424,1678,1745,1526,1559,697,783,486,408,505,473,1237,1327,814,795,756,843\nFN1 - mRNA,22662,10613,996,1179,2722,2742,9836,7979,8660,6356,76714,90575,103654,101410,125229,118874,111531,114470,101743,114807,113529,109422,148729,146786,251520,244081,194640,188503,232159,214283,73,76,91,88,73,71,65,74,64,74,449,59\nHSPA8 - mRNA,20754,8967,981,2187,3301,4276,8533,8100,12042,9744,60377,70272,83045,83246,179040,182664,171296,172184,116083,136367,135470,146788,153113,151766,82175,85134,85066,89978,44585,42964,117511,119092,157482,142914,148310,148315,82807,79745,110979,110114,112089,114613\nRPL19 - mRNA,18432,20804,574,4160,8022,9337,8061,8271,7566,6202,55887,67497,67889,70715,153993,157279,144429,142025,130613,145545,145715,151046,188285,183747,174708,184328,144251,144296,187069,181284,115121,129505,144086,131123,135402,136877,128279,130264,121015,121579,119686,125866\nRPLP0 - mRNA,17074,19906,500,3268,7196,8048,8938,6974,7480,5914,63650,76476,85578,82686,146659,141130,128011,137712,146270,158453,150259,153407,141056,136311,133726,130012,115694,113802,147675,140200,161669,176900,150692,138921,139419,142375,151283,145586,134590,139726,135537,135913\nACTG1 - mRNA,5905,6634,589,996,1901,2034,2834,2701,7010,4894,89917,104200,116639,113700,45148,41769,36423,40547,48554,53443,47376,51580,88600,90870,110609,101465,74096,62580,76418,73817,37125,40529,53368,45175,50630,48967,45308,49627,52772,60777,58558,56126\nLDHA - mRNA,14135,16082,221,2298,4615,5042,5524,4616,2856,2205,27823,31188,37188,37080,58507,55615,52718,38269,102252,114929,107513,96316,97706,104595,94668,89040,166381,153131,183217,172912,24246,35690,70680,61206,65417,65171,205083,201734,139204,139884,139290,138170\nLYZ - mRNA,1387,291,23,29,83,136,430,337,214,266,35,38,35,32,7852,7752,8221,7512,3361,4199,4054,4181,266557,259137,199741,189922,283016,287536,230397,225972,182732,182309,199196,174931,188592,186283,227893,232935,205907,213753,213972,215607\nGAPDH - mRNA,51077,44906,1145,6982,14027,16460,16927,17266,14518,10997,119167,134053,147922,148375,261564,257226,232573,223465,315195,358583,332800,335929,184297,192914,224924,221458,225441,215350,228075,219785,127163,156478,174190,154451,163945,163129,249712,253234,182367,187549,189159,187537\n";
	colAnnotation_UQh1YWZvRyUX_kmCGIztRmQ = "File.Name,Description,Batch.ID,Sample.Name,Cartridge.Id,Lane.Number,Analyte.Types,System.Type,Gene.RLF,Scanner.Id,Stage.Position,Scanned.Date,Gender,QC.Flag,FOV.Count,FOV.Counted,Binding.Density,Is.Reference,Imaging.QC.Flag,Imaging.QC,Binding.Density.QC.Flag,Binding.Density.QC,Positive.Control.QC.Flag,Positive.Control.QC,Limit.of.Detection.QC.Flag.of.mRNA,Limit.of.Detection.QC.of.mRNA,Line,Day,\n20161013_sebelius2-101316_01_09.RCC,Caco2 Control D1,JR2,01,sebelius2-101316,9,mRNA,NONE,SEBELIUS_2_C4636,1507C0300,3,2016-10-13 00:00:00.0,Unknown,false,280,278,0.25,false,false,0.992857,false,0.25,false,0.986733,false,7,Caco2,1,\n20161013_sebelius2-101316_01_10.RCC,Caco2 Control D3,JR2,01,sebelius2-101316,10,mRNA,NONE,SEBELIUS_2_C4636,1507C0300,3,2016-10-13 00:00:00.0,Unknown,false,280,280,0.46,false,false,1,false,0.46,false,0.995246,false,7.824747,Caco2,3,\n20161013_sebelius2-101316_01_11.RCC,Caco2 Control D7,JR2,01,sebelius2-101316,11,mRNA,NONE,SEBELIUS_2_C4636,1507C0300,3,2016-10-13 00:00:00.0,Unknown,false,280,277,0.43,false,false,0.989286,false,0.43,false,0.980532,false,6.75,Caco2,7,\n20161013_sebelius2-101316_01_12.RCC,Caco2 Control D10,JR2,01,sebelius2-101316,12,mRNA,NONE,SEBELIUS_2_C4636,1507C0300,3,2016-10-13 00:00:00.0,Unknown,false,280,280,0.25,false,false,1,false,0.25,false,0.995783,false,5.125,Caco2,10,\n20170307_20170307-caco_01_01.RCC,Caco2 Control D1,JR2,01,20170307-caco,1,mRNA,NONE,SEBELIUS_2_C4636,1507C0300,1,2017-03-07 00:00:00.0,Unknown,false,280,280,0.32,false,false,1,false,0.32,false,0.994436,false,5.875,Caco2,1,\n20170307_20170307-caco_01_02.RCC,Caco2 Control D1,JR2,01,20170307-caco,2,mRNA,NONE,SEBELIUS_2_C4636,1507C0300,1,2017-03-07 00:00:00.0,Unknown,false,280,280,0.31,false,false,1,false,0.31,false,0.992261,false,5.75,Caco2,1,\n20170307_20170307-caco_01_03.RCC,Caco2 Control D4,JR2,01,20170307-caco,3,mRNA,NONE,SEBELIUS_2_C4636,1507C0300,1,2017-03-07 00:00:00.0,Unknown,false,280,280,0.33,false,false,1,false,0.33,false,0.980453,false,6.375,Caco2,4,\n20170307_20170307-caco_01_04.RCC,Caco2 Control D4,JR2,01,20170307-caco,4,mRNA,NONE,SEBELIUS_2_C4636,1507C0300,1,2017-03-07 00:00:00.0,Unknown,false,280,280,0.3,false,false,1,false,0.3,false,0.994702,false,6,Caco2,4,\n20170307_20170307-caco_01_05.RCC,Caco2 Control D7,JR2,01,20170307-caco,5,mRNA,NONE,SEBELIUS_2_C4636,1507C0300,1,2017-03-07 00:00:00.0,Unknown,false,280,280,0.36,false,false,1,false,0.36,false,0.992391,false,5.375,Caco2,7,\n20170307_20170307-caco_01_06.RCC,Caco2 Control D7,JR2,01,20170307-caco,6,mRNA,NONE,SEBELIUS_2_C4636,1507C0300,1,2017-03-07 00:00:00.0,Unknown,false,280,280,0.28,false,false,1,false,0.28,false,0.955249,false,5.625,Caco2,7,\n20180926_Polarization01_Caco2D4A_01.RCC,Caco2 Control D5,ST,Caco2D4A,Polarization01,1,mRNA,NONE,SEBELIUS_2_C4636,1507C0300,5,2018-09-26 00:00:00.0,Unknown,false,555,524,1.8,false,false,0.944144,false,1.8,false,0.995004,false,32.241182,Caco2,5,\n20180926_Polarization01_Caco2D4B_02.RCC,Caco2 Control D5,ST,Caco2D4B,Polarization01,2,mRNA,NONE,SEBELIUS_2_C4636,1507C0300,5,2018-09-26 00:00:00.0,Unknown,true,555,522,2.27,false,false,0.940541,true,2.27,false,0.993594,false,26.813122,Caco2,5,\n20180926_Polarization01_Caco2D4C_03.RCC,Caco2 Control D5,ST,Caco2D4C,Polarization01,3,mRNA,NONE,SEBELIUS_2_C4636,1507C0300,5,2018-09-26 00:00:00.0,Unknown,true,555,513,2.78,false,false,0.924324,true,2.78,false,0.999676,false,25.190455,Caco2,5,\n20180926_Polarization01_Caco2D4D_04.RCC,Caco2 Control D5,ST,Caco2D4D,Polarization01,4,mRNA,NONE,SEBELIUS_2_C4636,1507C0300,5,2018-09-26 00:00:00.0,Unknown,true,555,506,2.38,false,false,0.911712,true,2.38,false,0.997102,false,25.269016,Caco2,5,\n20180926_Polarization01_FibD4A_05.RCC,Fibro Control D5,ST,FibD4A,Polarization01,5,mRNA,NONE,SEBELIUS_2_C4636,1507C0300,5,2018-09-26 00:00:00.0,Unknown,false,555,495,1.49,false,false,0.891892,false,1.49,false,0.99523,false,26.458078,Fibro,5,\n20180926_Polarization01_FibD4B_06.RCC,Fibro Control D5,ST,FibD4B,Polarization01,6,mRNA,NONE,SEBELIUS_2_C4636,1507C0300,5,2018-09-26 00:00:00.0,Unknown,false,555,491,1.15,false,false,0.884685,false,1.15,false,0.995856,false,23.642375,Fibro,5,\n20180926_Polarization01_HT29D4A_07.RCC,HT29 Control D5,ST,HT29D4A,Polarization01,7,mRNA,NONE,SEBELIUS_2_C4636,1507C0300,5,2018-09-26 00:00:00.0,Unknown,false,555,494,2.25,false,false,0.89009,false,2.25,false,0.99817,false,31.435711,HT29,5,\n20180926_Polarization01_HT29D4B_08.RCC,HT29 Control D5,ST,HT29D4B,Polarization01,8,mRNA,NONE,SEBELIUS_2_C4636,1507C0300,5,2018-09-26 00:00:00.0,Unknown,true,555,487,2.45,false,false,0.877477,true,2.45,false,0.997999,false,38.69426,HT29,5,\n20180926_Polarization01_HT29D4C_09.RCC,HT29 Control D5,ST,HT29D4C,Polarization01,9,mRNA,NONE,SEBELIUS_2_C4636,1507C0300,5,2018-09-26 00:00:00.0,Unknown,true,555,481,2.67,false,false,0.866667,true,2.67,false,0.997853,false,28.699056,HT29,5,\n20180926_Polarization01_HT29D4D_10.RCC,HT29 Control D5,ST,HT29D4D,Polarization01,10,mRNA,NONE,SEBELIUS_2_C4636,1507C0300,5,2018-09-26 00:00:00.0,Unknown,true,555,479,2.47,false,false,0.863063,true,2.47,false,0.995337,false,35.329784,HT29,5,\n20180926_Polarization01_HT29D4E_11.RCC,HT29 Control D4,ST,HT29D4E,Polarization01,11,mRNA,NONE,SEBELIUS_2_C4636,1507C0300,5,2018-09-26 00:00:00.0,Unknown,true,555,453,2.78,false,false,0.816216,true,2.78,false,0.998514,false,31.425946,HT29,4,\n20180926_Polarization01_HT29D4F_12.RCC,HT29 Control D4,ST,HT29D4F,Polarization01,12,mRNA,NONE,SEBELIUS_2_C4636,1507C0300,5,2018-09-26 00:00:00.0,Unknown,true,555,452,2.33,false,false,0.814414,true,2.33,false,0.997126,false,20.743505,HT29,4,\n20180928_Polarization-02_Caco2D7A_01.RCC,Caco2 Control D8,ST,Caco2D7A,Polarization-02,1,mRNA,NONE,SEBELIUS_2_C4636,1507C0300,2,2018-09-27 00:00:00.0,Unknown,true,555,534,2.53,false,false,0.962162,true,2.53,false,0.995977,false,22.474098,Caco2,8,\n20180928_Polarization-02_Caco2D7B_02.RCC,Caco2 Control D8,ST,Caco2D7B,Polarization-02,2,mRNA,NONE,SEBELIUS_2_C4636,1507C0300,2,2018-09-27 00:00:00.0,Unknown,false,555,533,1.92,false,false,0.96036,false,1.92,false,0.994538,false,31.436929,Caco2,8,\n20180928_Polarization-02_Caco2D7C_03.RCC,Caco2 Control D8,ST,Caco2D7C,Polarization-02,3,mRNA,NONE,SEBELIUS_2_C4636,1507C0300,2,2018-09-27 00:00:00.0,Unknown,false,555,535,1.34,false,false,0.963964,false,1.34,false,0.997677,false,34.142351,Caco2,8,\n20180928_Polarization-02_Caco2D7D_04.RCC,Caco2 Control D8,ST,Caco2D7D,Polarization-02,4,mRNA,NONE,SEBELIUS_2_C4636,1507C0300,2,2018-09-27 00:00:00.0,Unknown,false,555,533,2.05,false,false,0.96036,false,2.05,false,0.995298,false,25.601672,Caco2,8,\n20180928_Polarization-02_FibD7A_05.RCC,Fibro Control D8,ST,FibD7A,Polarization-02,5,mRNA,NONE,SEBELIUS_2_C4636,1507C0300,2,2018-09-27 00:00:00.0,Unknown,false,555,539,1.58,false,false,0.971171,false,1.58,false,0.996031,false,27.367248,Fibro,8,\n20180928_Polarization-02_FibD7B_06.RCC,Fibro Control D8,ST,FibD7B,Polarization-02,6,mRNA,NONE,SEBELIUS_2_C4636,1507C0300,2,2018-09-27 00:00:00.0,Unknown,false,555,532,1.47,false,false,0.958559,false,1.47,false,0.995629,false,31.034423,Fibro,8,\n20180928_Polarization-02_HT29D7A_07.RCC,HT29 Control D8,ST,HT29D7A,Polarization-02,7,mRNA,NONE,SEBELIUS_2_C4636,1507C0300,2,2018-09-27 00:00:00.0,Unknown,false,555,527,1.64,false,false,0.94955,false,1.64,false,0.995438,false,29.61076,HT29,8,\n20180928_Polarization-02_HT29D7B_08.RCC,HT29 Control D8,ST,HT29D7B,Polarization-02,8,mRNA,NONE,SEBELIUS_2_C4636,1507C0300,2,2018-09-27 00:00:00.0,Unknown,false,555,527,1.61,false,false,0.94955,false,1.61,false,0.990571,false,22.324298,HT29,8,\n20180928_Polarization-02_HT29D7C_09.RCC,HT29 Control D8,ST,HT29D7C,Polarization-02,9,mRNA,NONE,SEBELIUS_2_C4636,1507C0300,2,2018-09-27 00:00:00.0,Unknown,false,555,529,1.89,false,false,0.953153,false,1.89,false,0.997566,false,28.893882,HT29,8,\n20180928_Polarization-02_HT29D7D_10.RCC,HT29 Control D8,ST,HT29D7D,Polarization-02,10,mRNA,NONE,SEBELIUS_2_C4636,1507C0300,2,2018-09-27 00:00:00.0,Unknown,false,555,531,2.16,false,false,0.956757,false,2.16,false,0.992868,false,27.913127,HT29,8,\n20180928_Polarization-02_HT29D7E_11.RCC,HT29 Control D7,ST,HT29D7E,Polarization-02,11,mRNA,NONE,SEBELIUS_2_C4636,1507C0300,2,2018-09-27 00:00:00.0,Unknown,false,555,527,1.96,false,false,0.94955,false,1.96,false,0.995349,false,21.680057,HT29,7,\n20180928_Polarization-02_HT29D7F_12.RCC,HT29 Control D7,ST,HT29D7F,Polarization-02,12,mRNA,NONE,SEBELIUS_2_C4636,1507C0300,2,2018-09-27 00:00:00.0,Unknown,false,555,528,1.85,false,false,0.951351,false,1.85,false,0.994969,false,25.465345,HT29,7,\n20181004_Pol03_HepG2D4A_01.RCC,HEPG2 Control D5,ST,HepG2D4A,Pol03,1,mRNA,NONE,SEBELIUS_2_C4636,1507C0300,1,2018-10-04 00:00:00.0,Unknown,false,555,544,1.97,false,false,0.98018,false,1.97,false,0.994053,false,29.354049,HEPG2,5,\n20181004_Pol03_HepG2D4B_02.RCC,HEPG2 Control D5,ST,HepG2D4B,Pol03,2,mRNA,NONE,SEBELIUS_2_C4636,1507C0300,1,2018-10-04 00:00:00.0,Unknown,false,555,541,2.07,false,false,0.974775,false,2.07,false,0.999207,false,25.678388,HEPG2,5,\n20181004_Pol03_HepG2D4C_03.RCC,HEPG2 Control D4,ST,HepG2D4C,Pol03,3,mRNA,NONE,SEBELIUS_2_C4636,1507C0300,1,2018-10-04 00:00:00.0,Unknown,false,555,538,1.74,false,false,0.969369,false,1.74,false,0.99564,false,18.485281,HEPG2,4,\n20181004_Pol03_HepG2D4D_04.RCC,HEPG2 Control D4,ST,HepG2D4D,Pol03,4,mRNA,NONE,SEBELIUS_2_C4636,1507C0300,1,2018-10-04 00:00:00.0,Unknown,false,555,538,2.12,false,false,0.969369,false,2.12,false,0.994346,false,26.852592,HEPG2,4,\n20181004_Pol03_HepG2D7A_05.RCC,HEPG2 Control D8,ST,HepG2D7A,Pol03,5,mRNA,NONE,SEBELIUS_2_C4636,1507C0300,1,2018-10-04 00:00:00.0,Unknown,false,555,540,1.96,false,false,0.972973,false,1.96,false,0.998224,false,25.136644,HEPG2,8,\n20181004_Pol03_HepG2D7B_06.RCC,HEPG2 Control D8,ST,HepG2D7B,Pol03,6,mRNA,NONE,SEBELIUS_2_C4636,1507C0300,1,2018-10-04 00:00:00.0,Unknown,false,555,534,2,false,false,0.962162,false,2,false,0.996591,false,29.761011,HEPG2,8,\n20181004_Pol03_HepG2D7C_07.RCC,HEPG2 Control D7,ST,HepG2D7C,Pol03,7,mRNA,NONE,SEBELIUS_2_C4636,1507C0300,1,2018-10-04 00:00:00.0,Unknown,false,555,532,2.07,false,false,0.958559,false,2.07,false,0.99712,false,22.973013,HEPG2,7,\n20181004_Pol03_HepG2D7D_08.RCC,HEPG2 Control D7,ST,HepG2D7D,Pol03,8,mRNA,NONE,SEBELIUS_2_C4636,1507C0300,1,2018-10-04 00:00:00.0,Unknown,false,555,531,1.96,false,false,0.956757,false,1.96,false,0.995664,false,24.508926,HEPG2,7,\n";
	rowAnnotation_UQh1YWZvRyUX_kmCGIztRmQ = "Dimname,ProbeID,Codeset.Name,Probe.Label,Analyte.Type,Is.Control,Control.Type,Related.Probes,Probe.Annotation,KEGG.Pathways,Official.Gene.Name,Control.Conc,probeLabel.Type,below.threshold\nFN1 - mRNA,NM_212482.1:1776,SEBELIUS_2_C4636,FN1,mRNA,0,,,MAP2K and MAPK activation;p130Cas linkage to MAPK signaling for integrins;Platelet degranulation ;Molecules associated with elastic fibres;Syndecan interactions;ECM proteoglycans;Fibronectin matrix formation;Integrin cell surface interactions;RAFMAP kinase cascade;Downstream signal transduction;VEGFR2 mediated cell proliferation;Signaling by SCF-KIT;Frs2-mediated activation;ARMS-mediated activation;Gastrin-CREB signalling pathway via PKC and MAPK;Interleukin receptor SHC signaling;FCERI mediated MAPK activation;DAP12 signaling;Signaling by PDGF;VEGFA-VEGFR2 Pathway;Signaling by ERBB2;IRS-related events triggered by IGF1R;Integrin alphaIIb beta3 signaling;Non-integrin membrane-ECM interactions;Elastic fibre formation,hsa04151;hsa04510;hsa04512;hsa04810;hsa04933;hsa05100;hsa05146;hsa05200;hsa05205;hsa05222,FN1,NA,FN1 - mRNA,False\nLYZ - mRNA,NM_000239.2:305,SEBELIUS_2_C4636,LYZ,mRNA,0,,,,,LYZ,NA,LYZ - mRNA,False\nLDHA - mRNA,NM_001165414.1:1690,SEBELIUS_2_C4636,LDHA,mRNA,0,,,,hsa01100;hsa04066;hsa04922;hsa05230,LDHA,NA,LDHA - mRNA,False\nRPL19 - mRNA,NM_000981.3:315,SEBELIUS_2_C4636,RPL19,mRNA,0,,,Nonsense Mediated Decay (NMD) enhanced by the Exon Junction Complex (EJC),,RPL19,NA,RPL19 - mRNA,False\nRPLP0 - mRNA,NM_001002.3:250,SEBELIUS_2_C4636,RPLP0,mRNA,0,,,Nonsense Mediated Decay (NMD) enhanced by the Exon Junction Complex (EJC),,RPLP0,NA,RPLP0 - mRNA,False\nCCND1 - mRNA,NM_053056.2:2725,SEBELIUS_2_C4636,CCND1,mRNA,0,,,Cyclin D associated events in G1;Pre-NOTCH Transcription and Translation,hsa04068;hsa04110;hsa04115;hsa04151;hsa04152;hsa04310;hsa04390;hsa04510;hsa04630;hsa04917;hsa04919;hsa04921;hsa04933;hsa05161;hsa05162;hsa05166;hsa05200;hsa05203;hsa05205;hsa05206;hsa05210;hsa05212;hsa05213;hsa05214;hsa05215;hsa05216;hsa05218;hsa05219;hsa05220;hsa05221;hsa05222;hsa05223;hsa05416,CCND1,NA,CCND1 - mRNA,False\nCTNNB1 - mRNA,NM_001098210.1:1815,SEBELIUS_2_C4636,CTNNB1,mRNA,0,,,Disassembly of the destruction complex and recruitment of AXIN to the membrane;S37 mutants of beta-catenin aren.SQT.t phosphorylated;T41 mutants of beta-catenin aren.SQT.t phosphorylated;S45 mutants of beta-catenin aren.SQT.t phosphorylated;Binding of TCFLEF.CL.CTNNB1 to target gene promoters;RHO GTPases activate IQGAPs;Deactivation of the beta-catenin transactivating complex;Repression of WNT target genes;Apoptotic cleavage of cell adhesion  proteins;Adherens junctions interactions;Misspliced GSK3beta mutants stabilize beta-catenin;VEGFR2 mediated vascular permeability;Beta-catenin phosphorylation cascade;Synthesis# secretion# and inactivation of Glucagon-like Peptide-1 (GLP-1);Formation of the beta-catenin.CL.TCF transactivating complex;Apoptotic cleavage of cellular proteins;VEGFA-VEGFR2 Pathway,hsa04015;hsa04310;hsa04390;hsa04510;hsa04520;hsa04530;hsa04550;hsa04670;hsa04916;hsa04919;hsa05100;hsa05130;hsa05166;hsa05200;hsa05205;hsa05210;hsa05213;hsa05215;hsa05216;hsa05217;hsa05412,CTNNB1,NA,CTNNB1 - mRNA,False\nITGB1 - mRNA,NM_033666.2:2000,SEBELIUS_2_C4636,ITGB1,mRNA,0,,,Laminin interactions;RHO GTPases Activate Formins;ECM proteoglycans;Fibronectin matrix formation;Integrin cell surface interactions;Immunoregulatory interactions between a Lymphoid and a non-Lymphoid cell;Localization of the PINCH-ILK-PARVIN complex to focal adhesions;Elastic fibre formation;Cell surface interactions at the vascular wall;Non-integrin membrane-ECM interactions;Cell-extracellular matrix interactions,hsa04015;hsa04145;hsa04151;hsa04360;hsa04510;hsa04512;hsa04514;hsa04611;hsa04670;hsa04810;hsa05100;hsa05130;hsa05131;hsa05133;hsa05140;hsa05145;hsa05200;hsa05205;hsa05222;hsa05410;hsa05412;hsa05414,ITGB1,NA,ITGB1 - mRNA,False\nCDC42 - mRNA,NM_001039802.1:390,SEBELIUS_2_C4636,CDC42,mRNA,0,,,DCC mediated attractive signaling;Factors involved in megakaryocyte development and platelet production;VEGFA-VEGFR2 Pathway;RHO GTPases activate KTN1;CD28 dependent Vav1 pathway;Inactivation of Cdc42 and Rac;Rho GTPase cycle;EPHB-mediated forward signaling;RHO GTPases Activate Formins;RHO GTPases activate PAKs;GPVI-mediated activation cascade;G alpha (1213) signalling events;RHO GTPases activate IQGAPs;Sema4D induced cell migration and growth-cone collapse;Regulation of actin dynamics for phagocytic cup formation;CD28 co-stimulation,hsa04010;hsa04014;hsa04015;hsa04062;hsa04144;hsa04360;hsa04370;hsa04510;hsa04520;hsa04530;hsa04660;hsa04666;hsa04670;hsa04722;hsa04810;hsa04912;hsa04932;hsa04933;hsa05100;hsa05120;hsa05130;hsa05131;hsa05132;hsa05200;hsa05203;hsa05205;hsa05211;hsa05212,CDC42,NA,CDC42 - mRNA,False\nCSNK1A1 - mRNA,NM_001892.4:2053,SEBELIUS_2_C4636,CSNK1A1,mRNA,0,,,Misspliced GSK3beta mutants stabilize beta-catenin;APC truncation mutants have impaired AXIN binding;T41 mutants of beta-catenin aren.SQT.t phosphorylated;AXIN missense mutants destabilize the destruction complex;S45 mutants of beta-catenin aren.SQT.t phosphorylated;Disassembly of the destruction complex and recruitment of AXIN to the membrane;S37 mutants of beta-catenin aren.SQT.t phosphorylated;S33 mutants of beta-catenin aren.SQT.t phosphorylated;Beta-catenin phosphorylation cascade;Degradation of GLI2 by the proteasome,hsa04310,CSNK1A1,NA,CSNK1A1 - mRNA,False\nPPP2R1A - mRNA,NM_014225.3:1440,SEBELIUS_2_C4636,PPP2R1A,mRNA,0,,,MASTL Facilitates Mitotic Progression;Regulation of PLK1 Activity at G2M Transition;Disassembly of the destruction complex and recruitment of AXIN to the membrane;S37 mutants of beta-catenin aren.SQT.t phosphorylated;Cyclin AB1 associated events during G2M transition;RAF activation;RHO GTPases Activate Formins;S33 mutants of beta-catenin aren.SQT.t phosphorylated;Loss of Nlp from mitotic centrosomes;DARPP-32 events;Beta-catenin phosphorylation cascade;Initiation of Nuclear Envelope Reformation;Recruitment of mitotic centrosome proteins and complexes;Spry regulation of FGF signaling;Truncations of AMER1 destabilize the destruction complex;CTLA4 inhibitory signaling;AXIN missense mutants destabilize the destruction complex;S45 mutants of beta-catenin aren.SQT.t phosphorylated;Anchoring of the basal body to the plasma membrane;Nonsense Mediated Decay (NMD) enhanced by the Exon Junction Complex (EJC);Cyclin D associated events in G1;Platelet sensitization by LDL;Resolution of Sister Chromatid Cohesion;Inhibition of replication initiation of damaged DNA by RB1E2F1;RAFMAP kinase cascade;Downstream signal transduction;VEGFR2 mediated cell proliferation;Signaling by SCF-KIT;Frs2-mediated activation;ARMS-mediated activation;Gastrin-CREB signalling pathway via PKC and MAPK;Interleukin receptor SHC signaling;FCERI mediated MAPK activation;DAP12 signaling;Signaling by PDGF;VEGFA-VEGFR2 Pathway;Signaling by ERBB2;IRS-related events triggered by IGF1R;Negative regulation of FGFR3 signaling;Negative regulation of FGFR4 signaling;Negative regulation of FGFR2 signaling;Negative regulation of FGFR1 signaling,hsa03015;hsa04071;hsa04114;hsa04151;hsa04152;hsa04261;hsa04350;hsa04390;hsa04530;hsa04728;hsa04730;hsa05142;hsa05160,PPP2R1A,NA,PPP2R1A - mRNA,False\nACTB - mRNA,NM_001101.2:1010,SEBELIUS_2_C4636,ACTB,mRNA,0,,,EPHB-mediated forward signaling;Interaction between L1 and Ankyrins;DNA Damage Recognition in GG-NER;EPH-ephrin mediated repulsion of cells;Prefoldin mediated transfer of substrate  to CCTTriC;RHO GTPases Activate Formins;RHO GTPases activate IQGAPs;Cell-extracellular matrix interactions;Regulation of actin dynamics for phagocytic cup formation;Folding of actin by CCTTriC;B-WICH complex positively regulates rRNA expression;RAFMAP kinase cascade;Downstream signal transduction;VEGFR2 mediated cell proliferation;Signaling by SCF-KIT;Frs2-mediated activation;ARMS-mediated activation;Gastrin-CREB signalling pathway via PKC and MAPK;Interleukin receptor SHC signaling;FCERI mediated MAPK activation;DAP12 signaling;Signaling by PDGF;VEGFA-VEGFR2 Pathway;Signaling by ERBB2;IRS-related events triggered by IGF1R,hsa04015;hsa04145;hsa04390;hsa04510;hsa04520;hsa04530;hsa04611;hsa04670;hsa04810;hsa04919;hsa04921;hsa04971;hsa05100;hsa05110;hsa05130;hsa05131;hsa05132;hsa05164;hsa05205;hsa05410;hsa05412;hsa05414;hsa05416,ACTB,NA,ACTB - mRNA,False\nACTG1 - mRNA,NM_001614.2:1615,SEBELIUS_2_C4636,ACTG1,mRNA,0,,,RHO GTPases Activate WASPs and WAVEs;Translocation of GLUT4 to the plasma membrane;Recycling pathway of L1;Interaction between L1 and Ankyrins;EPHB-mediated forward signaling;EPH-ephrin mediated repulsion of cells;RHO GTPases Activate Formins;RHO GTPases activate IQGAPs;Cell-extracellular matrix interactions;Regulation of actin dynamics for phagocytic cup formation;RAFMAP kinase cascade;Downstream signal transduction;VEGFR2 mediated cell proliferation;Signaling by SCF-KIT;Frs2-mediated activation;ARMS-mediated activation;Gastrin-CREB signalling pathway via PKC and MAPK;Interleukin receptor SHC signaling;FCERI mediated MAPK activation;DAP12 signaling;Signaling by PDGF;VEGFA-VEGFR2 Pathway;Signaling by ERBB2;IRS-related events triggered by IGF1R,hsa04015;hsa04145;hsa04390;hsa04510;hsa04520;hsa04530;hsa04611;hsa04670;hsa04810;hsa04919;hsa04921;hsa05100;hsa05110;hsa05130;hsa05131;hsa05132;hsa05164;hsa05205;hsa05410;hsa05412;hsa05414;hsa05416,ACTG1,NA,ACTG1 - mRNA,False\nPPP1CA - mRNA,NM_002708.3:505,SEBELIUS_2_C4636,PPP1CA,mRNA,0,,,DARPP-32 events;Downregulation of TGF-beta receptor signaling;Circadian Clock;TGF-beta receptor signaling activates SMADs,hsa03015;hsa04022;hsa04024;hsa04114;hsa04261;hsa04270;hsa04390;hsa04510;hsa04611;hsa04720;hsa04728;hsa04750;hsa04810;hsa04910;hsa04921;hsa04931;hsa05031;hsa05034;hsa05168;hsa05205,PPP1CA,NA,PPP1CA - mRNA,False\nCFL1 - mRNA,NM_005507.2:225,SEBELIUS_2_C4636,CFL1,mRNA,0,,,EPHB-mediated forward signaling;Sema3A PAK dependent Axon repulsion;Platelet degranulation ;Regulation of actin dynamics for phagocytic cup formation,hsa04360;hsa04666;hsa04810;hsa05133,CFL1,NA,CFL1 - mRNA,False\nMYL12B - mRNA,NM_001144945.1:749,SEBELIUS_2_C4636,MYL12B,mRNA,0,,,EPHA-mediated growth cone collapse;RHO GTPases Activate ROCKs;Smooth Muscle Contraction;RHO GTPases activate CIT;RHO GTPases activate PAKs;RHO GTPases activate PKNs;Sema4D induced cell migration and growth-cone collapse,hsa04510;hsa04530;hsa04611;hsa04670;hsa04810,MYL12B,NA,MYL12B - mRNA,False\nEPB41L3 - mRNA,NM_012307.2:3758,SEBELIUS_2_C4636,EPB41L3,mRNA,0,,,,hsa04530,EPB41L3,NA,EPB41L3 - mRNA,True\nALPI - mRNA,NM_001631.3:1270,SEBELIUS_2_C4636,ALPI,mRNA,0,,,,hsa01100,ALPI,NA,ALPI - mRNA,False\nCLDN8 - mRNA,NM_199328.2:805,SEBELIUS_2_C4636,CLDN8,mRNA,0,,,Tight junction interactions,hsa04514;hsa04530;hsa04670;hsa05160,CLDN8,NA,CLDN8 - mRNA,True\nWAS - mRNA,NM_000377.2:90,SEBELIUS_2_C4636,WAS,mRNA,0,,,RHO GTPases Activate WASPs and WAVEs;Regulation of actin dynamics for phagocytic cup formation,hsa04062;hsa04144;hsa04520;hsa04666;hsa04810;hsa05100;hsa05130;hsa05131;hsa05132;hsa05231,WAS,NA,WAS - mRNA,True\nCA1 - mRNA,NM_001738.2:895,SEBELIUS_2_C4636,CA1,mRNA,0,,,,,CA1,NA,CA1 - mRNA,True\nAPC2 - mRNA,NM_005883.2:8120,SEBELIUS_2_C4636,APC2,mRNA,0,,,,hsa04310;hsa04390;hsa04550;hsa04810;hsa05166;hsa05200;hsa05206;hsa05210;hsa05213;hsa05217,APC2,NA,APC2 - mRNA,True\nACTN3 - mRNA,NM_001104.1:296,SEBELIUS_2_C4636,ACTN3,mRNA,0,,,Nephrin interactions,hsa04510;hsa04520;hsa04530;hsa04670;hsa04810;hsa05146;hsa05203;hsa05412,ACTN3,NA,ACTN3 - mRNA,True\nPIK3R5 - mRNA,NM_001142633.1:3335,SEBELIUS_2_C4636,PIK3R5,mRNA,0,,,Synthesis of PIPs at the plasma membrane;G beta.CL.gamma signalling through PI3Kgamma;GPVI-mediated activation cascade,hsa04012;hsa04014;hsa04015;hsa04022;hsa04024;hsa04062;hsa04066;hsa04068;hsa04070;hsa04071;hsa04072;hsa04150;hsa04151;hsa04152;hsa04210;hsa04211;hsa04213;hsa04261;hsa04370;hsa04380;hsa04510;hsa04550;hsa04611;hsa04620;hsa04630;hsa04650;hsa04660;hsa04662;hsa04664;hsa04666;hsa04668;hsa04670;hsa04722;hsa04725;hsa04750;hsa04810;hsa04910;hsa04914;hsa04915;hsa04917;hsa04919;hsa04921;hsa04923;hsa04930;hsa04931;hsa04932;hsa04933;hsa04960;hsa04973;hsa05100;hsa05142;hsa05145;hsa05146;hsa05160;hsa05161;hsa05162;hsa05164;hsa05166;hsa05169;hsa05200;hsa05203;hsa05205;hsa05210;hsa05211;hsa05212;hsa05213;hsa05214;hsa05215;hsa05218;hsa05220;hsa05221;hsa05222;hsa05223;hsa05230;hsa05231,PIK3R5,NA,PIK3R5 - mRNA,True\nVAV3 - mRNA,NM_006113.4:1632,SEBELIUS_2_C4636,VAV3,mRNA,0,,NM_001079874.1:2680,Rho GTPase cycle;DAP12 signaling;EPH-ephrin mediated repulsion of cells;VEGFR2 mediated vascular permeability;NRAGE signals death through JNK;GPVI-mediated activation cascade;G alpha (1213) signalling events;FCERI mediated MAPK activation;Regulation of actin dynamics for phagocytic cup formation;VEGFA-VEGFR2 Pathway,hsa04024;hsa04062;hsa04510;hsa04650;hsa04660;hsa04662;hsa04664;hsa04666;hsa04670;hsa04810,VAV3,NA,VAV3 - mRNA,True\nPRKCB - mRNA,NM_212535.1:1750,SEBELIUS_2_C4636,PRKCB,mRNA,0,,,Disinhibition of SNARE formation;Trafficking of AMPA receptors;VEGFA-VEGFR2 Pathway;PCPCE pathway,hsa04010;hsa04012;hsa04014;hsa04015;hsa04020;hsa04062;hsa04064;hsa04066;hsa04070;hsa04071;hsa04150;hsa04270;hsa04310;hsa04370;hsa04510;hsa04530;hsa04540;hsa04650;hsa04662;hsa04666;hsa04670;hsa04713;hsa04720;hsa04723;hsa04724;hsa04725;hsa04726;hsa04727;hsa04728;hsa04730;hsa04750;hsa04912;hsa04916;hsa04919;hsa04921;hsa04931;hsa04933;hsa04960;hsa04971;hsa04972;hsa04973;hsa05031;hsa05032;hsa05140;hsa05146;hsa05161;hsa05164;hsa05200;hsa05205;hsa05206;hsa05214;hsa05223;hsa05231,PRKCB,NA,PRKCB - mRNA,True\nITGA4 - mRNA,NM_000885.4:975,SEBELIUS_2_C4636,ITGA4,mRNA,0,,,Integrin cell surface interactions;Cell surface interactions at the vascular wall,hsa04151;hsa04510;hsa04512;hsa04514;hsa04640;hsa04670;hsa04810;hsa05140;hsa05410;hsa05412;hsa05414,ITGA4,NA,ITGA4 - mRNA,True\nJAM2 - mRNA,NM_001270407.1:840,SEBELIUS_2_C4636,JAM2,mRNA,0,,,Integrin cell surface interactions;Cell surface interactions at the vascular wall,hsa04514;hsa04530;hsa04670;hsa05120,JAM2,NA,JAM2 - mRNA,True\nMYH11 - mRNA,NM_001040113.1:920,SEBELIUS_2_C4636,MYH11,mRNA,0,,,EPHA-mediated growth cone collapse;RHO GTPases Activate ROCKs;Smooth Muscle Contraction;RHO GTPases activate CIT;RHO GTPases activate PAKs;RHO GTPases activate PKNs;Sema4D induced cell migration and growth-cone collapse,hsa04270;hsa04530,MYH11,NA,MYH11 - mRNA,True\nVAV3 b - mRNA,NM_001079874.1:2680,SEBELIUS_2_C4636,VAV3 b,mRNA,0,,NM_006113.4:1632,Rho GTPase cycle;DAP12 signaling;EPH-ephrin mediated repulsion of cells;VEGFR2 mediated vascular permeability;NRAGE signals death through JNK;GPVI-mediated activation cascade;G alpha (1213) signalling events;FCERI mediated MAPK activation;Regulation of actin dynamics for phagocytic cup formation;VEGFA-VEGFR2 Pathway,hsa04024;hsa04062;hsa04510;hsa04650;hsa04660;hsa04662;hsa04664;hsa04666;hsa04670;hsa04810,VAV3,NA,VAV3 b - mRNA,False\nPIK3CD - mRNA,NM_005026.3:2978,SEBELIUS_2_C4636,PIK3CD,mRNA,0,,,Synthesis of PIPs at the plasma membrane;Interleukin receptor SHC signaling;G beta.CL.gamma signalling through PI3Kgamma;PIP3 activates AKT signaling;Constitutive Signaling by Aberrant PI3K in Cancer;Regulation of signaling by CBL;GPVI-mediated activation cascade;Downstream signal transduction;GAB1 signalosome;Signaling by SCF-KIT;PI3KAKT activation;Role of LAT2NTALLAB on calcium mobilization;DAP12 signaling;Signaling by PDGF;Signaling by ERBB2,hsa00562;hsa04012;hsa04014;hsa04015;hsa04022;hsa04024;hsa04062;hsa04066;hsa04068;hsa04070;hsa04071;hsa04072;hsa04150;hsa04151;hsa04152;hsa04210;hsa04211;hsa04213;hsa04261;hsa04370;hsa04380;hsa04510;hsa04550;hsa04611;hsa04620;hsa04630;hsa04650;hsa04660;hsa04662;hsa04664;hsa04666;hsa04668;hsa04670;hsa04722;hsa04725;hsa04750;hsa04810;hsa04910;hsa04914;hsa04915;hsa04917;hsa04919;hsa04921;hsa04923;hsa04930;hsa04931;hsa04932;hsa04933;hsa04960;hsa04973;hsa05100;hsa05142;hsa05145;hsa05146;hsa05160;hsa05161;hsa05162;hsa05164;hsa05166;hsa05169;hsa05200;hsa05203;hsa05205;hsa05210;hsa05211;hsa05212;hsa05213;hsa05214;hsa05215;hsa05218;hsa05220;hsa05221;hsa05222;hsa05223;hsa05230;hsa05231,PIK3CD,NA,PIK3CD - mRNA,False\nMRAS - mRNA,NM_012219.3:4210,SEBELIUS_2_C4636,MRAS,mRNA,0,,,,hsa04010;hsa04014;hsa04015;hsa04072;hsa04530;hsa04810;hsa05166;hsa05205,MRAS,NA,MRAS - mRNA,False\nPDGFD - mRNA,NM_025208.4:1120,SEBELIUS_2_C4636,PDGFD,mRNA,0,,,Signaling by PDGF,hsa04014;hsa04015;hsa04072;hsa04151;hsa04510;hsa04540;hsa04810;hsa05215;hsa05218;hsa05231,PDGFD,NA,PDGFD - mRNA,False\nJAM3 - mRNA,NM_032801.3:890,SEBELIUS_2_C4636,JAM3,mRNA,0,,,Integrin cell surface interactions;Cell surface interactions at the vascular wall,hsa04514;hsa04530;hsa04670;hsa05120,JAM3,NA,JAM3 - mRNA,False\nZEB1 - mRNA,NM_001128128.1:1450,SEBELIUS_2_C4636,ZEB1,mRNA,0,,,,hsa05202;hsa05206,ZEB1,NA,ZEB1 - mRNA,True\nCYFIP2 - mRNA,NM_001037332.2:4990,SEBELIUS_2_C4636,CYFIP2,mRNA,0,,,RHO GTPases Activate WASPs and WAVEs;VEGFA-VEGFR2 Pathway;Regulation of actin dynamics for phagocytic cup formation,hsa04810,CYFIP2,NA,CYFIP2 - mRNA,False\nPIK3CG - mRNA,NM_002649.2:2125,SEBELIUS_2_C4636,PIK3CG,mRNA,0,,,Synthesis of PIPs at the plasma membrane;G beta.CL.gamma signalling through PI3Kgamma;GPVI-mediated activation cascade,hsa00562;hsa04012;hsa04014;hsa04015;hsa04022;hsa04024;hsa04062;hsa04066;hsa04068;hsa04070;hsa04071;hsa04072;hsa04150;hsa04151;hsa04152;hsa04210;hsa04211;hsa04213;hsa04261;hsa04370;hsa04380;hsa04510;hsa04550;hsa04611;hsa04620;hsa04630;hsa04650;hsa04660;hsa04662;hsa04664;hsa04666;hsa04668;hsa04670;hsa04722;hsa04725;hsa04750;hsa04810;hsa04910;hsa04914;hsa04915;hsa04917;hsa04919;hsa04921;hsa04923;hsa04930;hsa04931;hsa04932;hsa04933;hsa04960;hsa04973;hsa05100;hsa05142;hsa05145;hsa05146;hsa05160;hsa05161;hsa05162;hsa05164;hsa05166;hsa05169;hsa05200;hsa05203;hsa05205;hsa05210;hsa05211;hsa05212;hsa05213;hsa05214;hsa05215;hsa05218;hsa05220;hsa05221;hsa05222;hsa05223;hsa05230;hsa05231,PIK3CG,NA,PIK3CG - mRNA,True\nRAC2 - mRNA,NM_002872.4:341,SEBELIUS_2_C4636,RAC2,mRNA,0,,,Rho GTPase cycle;GPVI-mediated activation cascade;G alpha (1213) signalling events;PCPCE pathway;Sema4D induced cell migration and growth-cone collapse,hsa04010;hsa04014;hsa04015;hsa04024;hsa04062;hsa04071;hsa04310;hsa04360;hsa04370;hsa04510;hsa04520;hsa04650;hsa04662;hsa04664;hsa04666;hsa04670;hsa04810;hsa05200;hsa05210;hsa05212;hsa05231;hsa05416,RAC2,NA,RAC2 - mRNA,True\nMAPK10 - mRNA,NM_138981.2:450,SEBELIUS_2_C4636,MAPK10,mRNA,0,,,Activation of the AP-1 family of transcription factors;Oxidative Stress Induced Senescence;FCERI mediated MAPK activation,hsa04010;hsa04012;hsa04014;hsa04024;hsa04068;hsa04071;hsa04141;hsa04310;hsa04380;hsa04510;hsa04620;hsa04621;hsa04664;hsa04668;hsa04722;hsa04723;hsa04728;hsa04750;hsa04910;hsa04912;hsa04914;hsa04917;hsa04920;hsa04930;hsa04931;hsa04932;hsa04933;hsa05120;hsa05131;hsa05132;hsa05133;hsa05142;hsa05145;hsa05152;hsa05160;hsa05161;hsa05164;hsa05168;hsa05169;hsa05200;hsa05210;hsa05212;hsa05231,MAPK10,NA,MAPK10 - mRNA,False\nHPSE - mRNA,NM_006665.3:1625,SEBELIUS_2_C4636,HPSE,mRNA,0,,,,hsa01100;hsa05205,HPSE,NA,HPSE - mRNA,False\nMYL12A - mRNA,NM_006471.2:78,SEBELIUS_2_C4636,MYL12A,mRNA,0,,NM_001303049.1:9;NM_001303048.1:27,Smooth Muscle Contraction;Ephrin signaling,hsa04510;hsa04530;hsa04611;hsa04670;hsa04810,MYL12A,NA,MYL12A - mRNA,False\nMYL9 - mRNA,NM_181526.1:133,SEBELIUS_2_C4636,MYL9,mRNA,0,,,EPHA-mediated growth cone collapse;RHO GTPases Activate ROCKs;Smooth Muscle Contraction;RHO GTPases activate CIT;RHO GTPases activate PAKs;RHO GTPases activate PKNs;Sema4D induced cell migration and growth-cone collapse,hsa04022;hsa04024;hsa04270;hsa04510;hsa04530;hsa04670;hsa04810;hsa04921,MYL9,NA,MYL9 - mRNA,False\nGSTM3 - mRNA,NM_000849.4:602,SEBELIUS_2_C4636,GSTM3,mRNA,0,,,,,GSTM3,NA,GSTM3 - mRNA,False\nIL1R1 b - mRNA,NM_001320984.1:2649,SEBELIUS_2_C4636,IL1R1 b,mRNA,0,,,,,,NA,IL1R1 b - mRNA,False\nPDGFRB - mRNA,NM_002609.3:840,SEBELIUS_2_C4636,PDGFRB,mRNA,0,,,PIP3 activates AKT signaling;Constitutive Signaling by Aberrant PI3K in Cancer;RAFMAP kinase cascade;Signaling by PDGF;Downstream signal transduction;GAB1 signalosome;Signaling by SCF-KIT;PI3KAKT activation;Role of LAT2NTALLAB on calcium mobilization;DAP12 signaling;Signaling by ERBB2;VEGFR2 mediated cell proliferation;Frs2-mediated activation;ARMS-mediated activation;Gastrin-CREB signalling pathway via PKC and MAPK;Interleukin receptor SHC signaling;FCERI mediated MAPK activation;VEGFA-VEGFR2 Pathway;IRS-related events triggered by IGF1R,hsa04010;hsa04014;hsa04015;hsa04020;hsa04060;hsa04072;hsa04151;hsa04510;hsa04540;hsa04810;hsa05166;hsa05200;hsa05206;hsa05214;hsa05215;hsa05218;hsa05230;hsa05231,PDGFRB,NA,PDGFRB - mRNA,False\nFGFR1 - mRNA,NM_015850.2:1335,SEBELIUS_2_C4636,FGFR1,mRNA,0,,,RAFMAP kinase cascade;Constitutive Signaling by Aberrant PI3K in Cancer;SHC-mediated cascade.CL.FGFR1;Phospholipase C-mediated cascade.CL. FGFR1;Signaling by FGFR1 amplification mutants;PI3K Cascade;FGFR1b ligand binding and activation;Signaling by FGFR1 in disease;Downstream signal transduction;GAB1 signalosome;Signaling by SCF-KIT;PI3KAKT activation;Role of LAT2NTALLAB on calcium mobilization;DAP12 signaling;Signaling by PDGF;Signaling by ERBB2;VEGFR2 mediated cell proliferation;Frs2-mediated activation;ARMS-mediated activation;Gastrin-CREB signalling pathway via PKC and MAPK;Interleukin receptor SHC signaling;FCERI mediated MAPK activation;VEGFA-VEGFR2 Pathway;IRS-related events triggered by IGF1R,hsa04010;hsa04014;hsa04015;hsa04151;hsa04520;hsa04550;hsa04810;hsa05200;hsa05205;hsa05215;hsa05218;hsa05230,FGFR1,NA,FGFR1 - mRNA,False\nPDGFRA - mRNA,NM_006206.3:1925,SEBELIUS_2_C4636,PDGFRA,mRNA,0,,,PIP3 activates AKT signaling;Constitutive Signaling by Aberrant PI3K in Cancer;RAFMAP kinase cascade;Signaling by PDGF;Downstream signal transduction;GAB1 signalosome;Signaling by SCF-KIT;PI3KAKT activation;Role of LAT2NTALLAB on calcium mobilization;DAP12 signaling;Signaling by ERBB2;VEGFR2 mediated cell proliferation;Frs2-mediated activation;ARMS-mediated activation;Gastrin-CREB signalling pathway via PKC and MAPK;Interleukin receptor SHC signaling;FCERI mediated MAPK activation;VEGFA-VEGFR2 Pathway;IRS-related events triggered by IGF1R,hsa04010;hsa04014;hsa04015;hsa04020;hsa04060;hsa04072;hsa04144;hsa04151;hsa04510;hsa04540;hsa04810;hsa05166;hsa05200;hsa05206;hsa05214;hsa05215;hsa05218;hsa05230;hsa05231,PDGFRA,NA,PDGFRA - mRNA,False\nAKT3 - mRNA,NM_005465.4:287,SEBELIUS_2_C4636,AKT3,mRNA,0,,,TP53 Regulates Metabolic Genes;CD28 dependent PI3KAkt signaling;AKT phosphorylates targets in the cytosol;Negative regulation of the PI3KAKT network;Activation of BAD and translocation to mitochondria ;G beta.CL.gamma signalling through PI3Kgamma;AKT phosphorylates targets in the nucleus;VEGFR2 mediated vascular permeability;GPVI-mediated activation cascade;Signaling by ERBB2;PIP3 activates AKT signaling;Downstream signal transduction;GAB1 signalosome;Signaling by SCF-KIT;PI3KAKT activation;Role of LAT2NTALLAB on calcium mobilization;DAP12 signaling;Signaling by PDGF;CD28 co-stimulation;VEGFA-VEGFR2 Pathway,hsa04010;hsa04012;hsa04014;hsa04015;hsa04022;hsa04024;hsa04062;hsa04066;hsa04068;hsa04071;hsa04072;hsa04150;hsa04151;hsa04152;hsa04210;hsa04211;hsa04213;hsa04261;hsa04370;hsa04380;hsa04510;hsa04530;hsa04550;hsa04611;hsa04620;hsa04630;hsa04660;hsa04662;hsa04664;hsa04666;hsa04668;hsa04722;hsa04725;hsa04728;hsa04910;hsa04914;hsa04915;hsa04917;hsa04919;hsa04920;hsa04922;hsa04923;hsa04931;hsa04932;hsa04933;hsa04973;hsa05142;hsa05145;hsa05152;hsa05160;hsa05161;hsa05162;hsa05164;hsa05166;hsa05169;hsa05200;hsa05205;hsa05210;hsa05211;hsa05212;hsa05213;hsa05214;hsa05215;hsa05218;hsa05220;hsa05221;hsa05222;hsa05223;hsa05230;hsa05231,AKT3,NA,AKT3 - mRNA,False\nITGA5 - mRNA,NM_002205.2:925,SEBELIUS_2_C4636,ITGA5,mRNA,0,,,Signal transduction by L1;Elastic fibre formation;Integrin cell surface interactions;Cell surface interactions at the vascular wall,hsa04145;hsa04151;hsa04510;hsa04512;hsa04640;hsa04810;hsa05100;hsa05131;hsa05133;hsa05205;hsa05206;hsa05410;hsa05412;hsa05414,ITGA5,NA,ITGA5 - mRNA,False\nITGA7 - mRNA,NM_002206.1:1170,SEBELIUS_2_C4636,ITGA7,mRNA,0,,,Laminin interactions;ECM proteoglycans;Integrin cell surface interactions,hsa04151;hsa04510;hsa04512;hsa04810;hsa05410;hsa05412;hsa05414,ITGA7,NA,ITGA7 - mRNA,False\nLEF1 - mRNA,NM_016269.3:1165,SEBELIUS_2_C4636,LEF1,mRNA,0,,,Ca2.PLS. pathway;Binding of TCFLEF.CL.CTNNB1 to target gene promoters;Deactivation of the beta-catenin transactivating complex;Repression of WNT target genes;Formation of the beta-catenin.CL.TCF transactivating complex,hsa04310;hsa04390;hsa04520;hsa04916;hsa05200;hsa05210;hsa05213;hsa05215;hsa05216;hsa05217;hsa05221;hsa05412,LEF1,NA,LEF1 - mRNA,False\nBCL2 - mRNA,NM_000657.2:5,SEBELIUS_2_C4636,BCL2,mRNA,0,,,Activation of BAD and translocation to mitochondria ,hsa04064;hsa04066;hsa04071;hsa04141;hsa04151;hsa04210;hsa04261;hsa04510;hsa04630;hsa04722;hsa04725;hsa04933;hsa05145;hsa05152;hsa05161;hsa05169;hsa05200;hsa05206;hsa05210;hsa05215;hsa05222,BCL2,NA,BCL2 - mRNA,False\nMAGI1 - mRNA,NM_004742.2:325,SEBELIUS_2_C4636,MAGI1,mRNA,0,,,,hsa04015;hsa04530,MAGI1,NA,MAGI1 - mRNA,False\nCCND2 - mRNA,NM_001759.3:503,SEBELIUS_2_C4636,CCND2,mRNA,0,,,Cyclin D associated events in G1,hsa04068;hsa04110;hsa04115;hsa04151;hsa04310;hsa04390;hsa04510;hsa04630;hsa04917;hsa05162;hsa05166;hsa05202;hsa05203;hsa05206,CCND2,NA,CCND2 - mRNA,False\nTCF7L1 - mRNA,NM_031283.1:2215,SEBELIUS_2_C4636,TCF7L1,mRNA,0,,,Ca2.PLS. pathway;Binding of TCFLEF.CL.CTNNB1 to target gene promoters;Deactivation of the beta-catenin transactivating complex;Repression of WNT target genes;Formation of the beta-catenin.CL.TCF transactivating complex,hsa04310;hsa04390;hsa04520;hsa04916;hsa05200;hsa05210;hsa05213;hsa05215;hsa05216;hsa05217;hsa05221;hsa05412,TCF7L1,NA,TCF7L1 - mRNA,False\nTARBP2 - mRNA,NM_134323.1:1060,SEBELIUS_2_C4636,TARBP2,mRNA,0,,,MicroRNA (miRNA) biogenesis,,TARBP2,NA,TARBP2 - mRNA,False\nABI2 - mRNA,NM_005759.5:1160,SEBELIUS_2_C4636,ABI2,mRNA,0,,,RHO GTPases Activate WASPs and WAVEs;VEGFA-VEGFR2 Pathway;Regulation of actin dynamics for phagocytic cup formation,hsa04810,ABI2,NA,ABI2 - mRNA,False\nPIK3R2 - mRNA,NM_005027.2:3100,SEBELIUS_2_C4636,PIK3R2,mRNA,0,,,G alpha (q) signalling events;Downstream TCR signaling;PIP3 activates AKT signaling;VEGFA-VEGFR2 Pathway;Synthesis of PIPs at the plasma membrane;Rho GTPase cycle;Interleukin receptor SHC signaling;G beta.CL.gamma signalling through PI3Kgamma;Regulation of signaling by CBL;Role of phospholipids in phagocytosis;G alpha (1213) signalling events;PI3K Cascade;Constitutive Signaling by Aberrant PI3K in Cancer;CD28 dependent PI3KAkt signaling;Tie2 Signaling;GPVI-mediated activation cascade;Nephrin interactions;Gastrin-CREB signalling pathway via PKC and MAPK;Downstream signal transduction;GAB1 signalosome;Signaling by SCF-KIT;PI3KAKT activation;Role of LAT2NTALLAB on calcium mobilization;DAP12 signaling;Signaling by PDGF;Signaling by ERBB2;IRS-related events triggered by IGF1R;CD28 co-stimulation;Cell surface interactions at the vascular wall,hsa04012;hsa04014;hsa04015;hsa04022;hsa04024;hsa04062;hsa04066;hsa04068;hsa04070;hsa04071;hsa04072;hsa04150;hsa04151;hsa04152;hsa04210;hsa04211;hsa04213;hsa04261;hsa04370;hsa04380;hsa04510;hsa04550;hsa04611;hsa04620;hsa04630;hsa04650;hsa04660;hsa04662;hsa04664;hsa04666;hsa04668;hsa04670;hsa04722;hsa04725;hsa04750;hsa04810;hsa04910;hsa04914;hsa04915;hsa04917;hsa04919;hsa04921;hsa04923;hsa04930;hsa04931;hsa04932;hsa04933;hsa04960;hsa04973;hsa05100;hsa05142;hsa05145;hsa05146;hsa05160;hsa05161;hsa05162;hsa05164;hsa05166;hsa05169;hsa05200;hsa05203;hsa05205;hsa05206;hsa05210;hsa05211;hsa05212;hsa05213;hsa05214;hsa05215;hsa05218;hsa05220;hsa05221;hsa05222;hsa05223;hsa05230;hsa05231,PIK3R2,NA,PIK3R2 - mRNA,False\nLIMK2 - mRNA,NM_001031801.1:1590,SEBELIUS_2_C4636,LIMK2,mRNA,0,,,EPHB-mediated forward signaling;RHO GTPases Activate ROCKs;Sema4D induced cell migration and growth-cone collapse,hsa04360;hsa04666;hsa04810,LIMK2,NA,LIMK2 - mRNA,False\nDVL2 - mRNA,NM_004422.2:1025,SEBELIUS_2_C4636,DVL2,mRNA,0,,,Disassembly of the destruction complex and recruitment of AXIN to the membrane;WNT mediated activation of DVL;Negative regulation of TCF-dependent signaling by DVL-interacting proteins;PCPCE pathway,hsa04310;hsa04330;hsa04390;hsa04550;hsa04916;hsa05166;hsa05200;hsa05217,DVL2,NA,DVL2 - mRNA,False\nERBB2 - mRNA,NM_001005862.1:1255,SEBELIUS_2_C4636,ERBB2,mRNA,0,,,RAFMAP kinase cascade;GRB7 events in ERBB2 signaling;PLCG1 events in ERBB2 signaling;Sema4D induced cell migration and growth-cone collapse;Signaling by ERBB2;Downstream signal transduction;GAB1 signalosome;Signaling by SCF-KIT;PI3KAKT activation;Role of LAT2NTALLAB on calcium mobilization;DAP12 signaling;Signaling by PDGF;VEGFR2 mediated cell proliferation;Frs2-mediated activation;ARMS-mediated activation;Gastrin-CREB signalling pathway via PKC and MAPK;Interleukin receptor SHC signaling;FCERI mediated MAPK activation;VEGFA-VEGFR2 Pathway;IRS-related events triggered by IGF1R,hsa04012;hsa04020;hsa04066;hsa04510;hsa04520;hsa05200;hsa05205;hsa05206;hsa05212;hsa05213;hsa05215;hsa05219;hsa05223;hsa05230,ERBB2,NA,ERBB2 - mRNA,False\nPRKCD - mRNA,NM_006254.3:2165,SEBELIUS_2_C4636,PRKCD,mRNA,0,,,HuR (ELAVL1) binds and stabilizes mRNA;Downstream signal transduction;EGFR interacts with phospholipase C-gamma;Phospholipase C-mediated cascade.CL. FGFR1;PLCG1 events in ERBB2 signaling;DAP12 signaling;PLC beta mediated events;Signaling by PDGF;Signaling by ERBB2;VEGFA-VEGFR2 Pathway;G alpha (q) signalling events;Gastrin-CREB signalling pathway via PKC and MAPK,hsa04062;hsa04270;hsa04530;hsa04666;hsa04722;hsa04750;hsa04912;hsa04915;hsa04930;hsa04931;hsa04933,PRKCD,NA,PRKCD - mRNA,False\nIGF1R - mRNA,NM_000875.3:10810,SEBELIUS_2_C4636,IGF1R,mRNA,0,,,IRS-related events triggered by IGF1R;SHC-related events triggered by IGF1R,hsa04014;hsa04015;hsa04066;hsa04068;hsa04114;hsa04144;hsa04151;hsa04152;hsa04211;hsa04213;hsa04510;hsa04520;hsa04550;hsa04730;hsa04914;hsa05200;hsa05202;hsa05205;hsa05214;hsa05215;hsa05218,IGF1R,NA,IGF1R - mRNA,False\nPRKCZ - mRNA,NM_002744.4:771,SEBELIUS_2_C4636,PRKCZ,mRNA,0,,,TGF-beta receptor signaling in EMT (epithelial to mesenchymal transition);VEGFR2 mediated cell proliferation;GPVI-mediated activation cascade;VEGFA-VEGFR2 Pathway,hsa04015;hsa04062;hsa04071;hsa04144;hsa04390;hsa04530;hsa04611;hsa04910;hsa04930;hsa04931;hsa04933,PRKCZ,NA,PRKCZ - mRNA,False\nPDGFC - mRNA,NM_016205.2:2596,SEBELIUS_2_C4636,PDGFC,mRNA,0,,,Signaling by PDGF,hsa04014;hsa04015;hsa04060;hsa04072;hsa04151;hsa04510;hsa04540;hsa04810;hsa05215;hsa05218;hsa05231,PDGFC,NA,PDGFC - mRNA,False\nIRS1 - mRNA,NM_005544.2:6224,SEBELIUS_2_C4636,IRS1,mRNA,0,,,Constitutive Signaling by Aberrant PI3K in Cancer;RAFMAP kinase cascade;PI3K Cascade;Growth hormone receptor signaling;IRS-related events triggered by IGF1R;PI3KAKT activation;Downstream signal transduction;GAB1 signalosome;Signaling by SCF-KIT;Role of LAT2NTALLAB on calcium mobilization;DAP12 signaling;Signaling by PDGF;Signaling by ERBB2;VEGFR2 mediated cell proliferation;Frs2-mediated activation;ARMS-mediated activation;Gastrin-CREB signalling pathway via PKC and MAPK;Interleukin receptor SHC signaling;FCERI mediated MAPK activation;VEGFA-VEGFR2 Pathway,hsa04022;hsa04068;hsa04150;hsa04151;hsa04152;hsa04211;hsa04213;hsa04722;hsa04910;hsa04920;hsa04923;hsa04930;hsa04931;hsa04932;hsa04960;hsa05206,IRS1,NA,IRS1 - mRNA,False\nEXOC3 - mRNA,NM_007277.4:1820,SEBELIUS_2_C4636,EXOC3,mRNA,0,,,Translocation of GLUT4 to the plasma membrane,hsa04530,EXOC3,NA,EXOC3 - mRNA,False\nNUMA1 - mRNA,NM_006185.2:575,SEBELIUS_2_C4636,NUMA1,mRNA,0,,,,,NUMA1,NA,NUMA1 - mRNA,False\nAGO2 - mRNA,NM_012154.2:625,SEBELIUS_2_C4636,AGO2,mRNA,0,,,Pre-NOTCH Transcription and Translation;PIP3 activates AKT signaling;Post-transcriptional silencing by small RNAs;MicroRNA (miRNA) biogenesis;Small interfering RNA (siRNA) biogenesis;Downstream signal transduction;GAB1 signalosome;Signaling by SCF-KIT;PI3KAKT activation;Role of LAT2NTALLAB on calcium mobilization;DAP12 signaling;Signaling by PDGF;Signaling by ERBB2,,AGO2,NA,AGO2 - mRNA,False\nBHLHE40 - mRNA,NM_003670.1:560,SEBELIUS_2_C4636,BHLHE40,mRNA,0,,,Circadian Clock,,BHLHE40,NA,BHLHE40 - mRNA,False\nTGFB1 - mRNA,NM_000660.3:1260,SEBELIUS_2_C4636,TGFB1,mRNA,0,,,TGF-beta receptor signaling in EMT (epithelial to mesenchymal transition);TGFBR1 LBD Mutants in Cancer;Influenza Virus Induced Apoptosis;Molecules associated with elastic fibres;Syndecan interactions;ECM proteoglycans;Transcriptional regulation of white adipocyte differentiation;TGF-beta receptor signaling activates SMADs;Elastic fibre formation;Non-integrin membrane-ECM interactions,hsa04010;hsa04060;hsa04068;hsa04110;hsa04144;hsa04350;hsa04380;hsa04390;hsa04932;hsa04933;hsa05140;hsa05142;hsa05145;hsa05146;hsa05152;hsa05161;hsa05166;hsa05200;hsa05205;hsa05210;hsa05211;hsa05212;hsa05220;hsa05321;hsa05410;hsa05414,TGFB1,NA,TGFB1 - mRNA,False\nRAB8B - mRNA,NM_016530.2:820,SEBELIUS_2_C4636,RAB8B,mRNA,0,,,,,RAB8B,NA,RAB8B - mRNA,False\nSMAD4 - mRNA,NM_005359.3:1370,SEBELIUS_2_C4636,SMAD4,mRNA,0,,,Signaling by NODAL,hsa04068;hsa04110;hsa04310;hsa04350;hsa04390;hsa04520;hsa04550;hsa04933;hsa05161;hsa05166;hsa05200;hsa05210;hsa05212;hsa05220,SMAD4,NA,SMAD4 - mRNA,False\nHSPB1 - mRNA,NM_001540.3:374,SEBELIUS_2_C4636,HSPB1,mRNA,0,,,MAPK6MAPK4 signaling;VEGFA-VEGFR2 Pathway,hsa04010;hsa04370;hsa05146;hsa05169,HSPB1,NA,HSPB1 - mRNA,False\nMYLK - mRNA,NM_053032.2:710,SEBELIUS_2_C4636,MYLK,mRNA,0,,,Smooth Muscle Contraction;RHO GTPases activate PAKs,hsa04020;hsa04022;hsa04270;hsa04510;hsa04611;hsa04810;hsa04921;hsa04971,MYLK,NA,MYLK - mRNA,False\nIL1R1 - mRNA,NM_000877.3:582,SEBELIUS_2_C4636,IL1R1,mRNA,0,,,,hsa04010;hsa04060;hsa04064;hsa04210;hsa04380;hsa04640;hsa04750;hsa05146;hsa05166,IL1R1,NA,IL1R1 - mRNA,False\nEPB41L2 - mRNA,NM_001199389.1:1880,SEBELIUS_2_C4636,EPB41L2,mRNA,0,,,,hsa04530,EPB41L2,NA,EPB41L2 - mRNA,False\nMYH10 - mRNA,NM_005964.1:7092,SEBELIUS_2_C4636,MYH10,mRNA,0,,,EPHA-mediated growth cone collapse;RHO GTPases Activate ROCKs;RHO GTPases activate CIT;RHO GTPases activate PAKs;RHO GTPases activate PKNs;Sema4D induced cell migration and growth-cone collapse,hsa04530;hsa04810;hsa05132,MYH10,NA,MYH10 - mRNA,False\nCD44 - mRNA,NM_001001392.1:429,SEBELIUS_2_C4636,CD44,mRNA,0,,,Hyaluronan uptake and degradation;Cell surface interactions at the vascular wall,hsa04512;hsa04640;hsa05131;hsa05169;hsa05205;hsa05206,CD44,NA,CD44 - mRNA,False\nGNAI1 - mRNA,NM_002069.4:670,SEBELIUS_2_C4636,GNAI1,mRNA,0,,,ADP signalling through P2Y purinoceptor 12;Adrenaline#noradrenaline inhibits insulin secretion,hsa04015;hsa04022;hsa04024;hsa04062;hsa04071;hsa04261;hsa04360;hsa04530;hsa04540;hsa04611;hsa04670;hsa04713;hsa04723;hsa04724;hsa04725;hsa04726;hsa04727;hsa04728;hsa04730;hsa04914;hsa04915;hsa04916;hsa04921;hsa04923;hsa04971;hsa05032;hsa05034;hsa05133;hsa05142;hsa05145;hsa05200,GNAI1,NA,GNAI1 - mRNA,False\nITGA1 - mRNA,NM_181501.1:1875,SEBELIUS_2_C4636,ITGA1,mRNA,0,,,Laminin interactions;Other semaphorin interactions;Integrin cell surface interactions,hsa04151;hsa04510;hsa04512;hsa04640;hsa04810;hsa05410;hsa05412;hsa05414,ITGA1,NA,ITGA1 - mRNA,False\nTP53 - mRNA,NM_000546.2:1330,SEBELIUS_2_C4636,TP53,mRNA,0,,,Activation of PUMA and translocation to mitochondria;Recruitment and ATM-mediated phosphorylation of repair and signaling proteins at DNA double strand breaks;Factors involved in megakaryocyte development and platelet production;Oncogene Induced Senescence,hsa04010;hsa04071;hsa04110;hsa04115;hsa04151;hsa04210;hsa04211;hsa04310;hsa04722;hsa04919;hsa05016;hsa05160;hsa05161;hsa05162;hsa05166;hsa05168;hsa05169;hsa05200;hsa05202;hsa05203;hsa05205;hsa05206;hsa05210;hsa05212;hsa05213;hsa05214;hsa05215;hsa05216;hsa05217;hsa05218;hsa05219;hsa05220;hsa05222;hsa05223;hsa05230,TP53,NA,TP53 - mRNA,False\nCTBP2 - mRNA,NM_001329.2:1884,SEBELIUS_2_C4636,CTBP2,mRNA,0,,,Repression of WNT target genes,hsa04310;hsa04330;hsa05200;hsa05220,CTBP2,NA,CTBP2 - mRNA,False\nKRAS - mRNA,NM_004985.3:327,SEBELIUS_2_C4636,KRAS,mRNA,0,,,Signaling by FGFR2 in disease;Regulation of RAS by GAPs;SHC-mediated cascade.CL.FGFR3;Signaling by FGFR3 in disease;MAP2K and MAPK activation;SHC-related events triggered by IGF1R;Signaling by FGFR4 in disease;SHC-mediated cascade.CL.FGFR1;Tie2 Signaling;Signaling by FGFR1 in disease;Activation of RAS in B cells;SHC-mediated cascade.CL.FGFR4;Constitutive Signaling by EGFRvIII;CD209 (DC-SIGN) signaling;FCERI mediated MAPK activation;RAFMAP kinase cascade;Downstream signal transduction;VEGFR2 mediated cell proliferation;Signaling by SCF-KIT;Frs2-mediated activation;ARMS-mediated activation;Gastrin-CREB signalling pathway via PKC and MAPK;Interleukin receptor SHC signaling;DAP12 signaling;Signaling by PDGF;VEGFA-VEGFR2 Pathway;Signaling by ERBB2;IRS-related events triggered by IGF1R;Cell surface interactions at the vascular wall,hsa04010;hsa04012;hsa04014;hsa04015;hsa04062;hsa04068;hsa04071;hsa04072;hsa04151;hsa04211;hsa04213;hsa04320;hsa04360;hsa04370;hsa04530;hsa04540;hsa04550;hsa04650;hsa04660;hsa04662;hsa04664;hsa04720;hsa04722;hsa04725;hsa04726;hsa04730;hsa04810;hsa04910;hsa04912;hsa04914;hsa04915;hsa04916;hsa04917;hsa04919;hsa04921;hsa04933;hsa04960;hsa05034;hsa05160;hsa05161;hsa05166;hsa05200;hsa05203;hsa05205;hsa05206;hsa05210;hsa05211;hsa05212;hsa05213;hsa05214;hsa05215;hsa05216;hsa05218;hsa05219;hsa05220;hsa05221;hsa05223;hsa05230;hsa05231,KRAS,NA,KRAS - mRNA,False\nPAK1 - mRNA,NM_002576.3:975,SEBELIUS_2_C4636,PAK1,mRNA,0,,,EPHB-mediated forward signaling;Ephrin signaling;MAPK6MAPK4 signaling;DSCAM interactions;Activation of Rac;VEGFR2 mediated vascular permeability;RHO GTPases activate PAKs;Sema3A PAK dependent Axon repulsion;FCERI mediated MAPK activation;Regulation of actin dynamics for phagocytic cup formation;CD209 (DC-SIGN) signaling;CD28 dependent Vav1 pathway;VEGFA-VEGFR2 Pathway;CD28 co-stimulation,hsa04010;hsa04012;hsa04014;hsa04024;hsa04062;hsa04360;hsa04510;hsa04650;hsa04660;hsa04666;hsa04810;hsa05120;hsa05205;hsa05211,PAK1,NA,PAK1 - mRNA,False\nCD9 - mRNA,NM_001769.2:405,SEBELIUS_2_C4636,CD9,mRNA,0,,,Platelet degranulation ,hsa04640,CD9,NA,CD9 - mRNA,False\nDICER1 - mRNA,NM_177438.2:5060,SEBELIUS_2_C4636,DICER1,mRNA,0,,,MicroRNA (miRNA) biogenesis,hsa05206,DICER1,NA,DICER1 - mRNA,False\nRAF1 - mRNA,NM_002880.3:1444,SEBELIUS_2_C4636,RAF1,mRNA,0,,,RAF activation;Stimuli-sensing channels;CREB phosphorylation through the activation of Ras;Negative feedback regulation of MAPK pathway;CD209 (DC-SIGN) signaling;RAFMAP kinase cascade;Downstream signal transduction;VEGFR2 mediated cell proliferation;Signaling by SCF-KIT;Frs2-mediated activation;ARMS-mediated activation;Gastrin-CREB signalling pathway via PKC and MAPK;Interleukin receptor SHC signaling;FCERI mediated MAPK activation;DAP12 signaling;Signaling by PDGF;VEGFA-VEGFR2 Pathway;Signaling by ERBB2;IRS-related events triggered by IGF1R;Negative regulation of MAPK pathway,hsa04010;hsa04012;hsa04014;hsa04015;hsa04022;hsa04024;hsa04062;hsa04068;hsa04071;hsa04072;hsa04151;hsa04270;hsa04370;hsa04510;hsa04540;hsa04550;hsa04630;hsa04650;hsa04660;hsa04662;hsa04664;hsa04666;hsa04720;hsa04722;hsa04726;hsa04730;hsa04810;hsa04910;hsa04912;hsa04914;hsa04915;hsa04916;hsa04917;hsa04919;hsa04921;hsa05034;hsa05152;hsa05160;hsa05161;hsa05164;hsa05200;hsa05205;hsa05206;hsa05210;hsa05211;hsa05212;hsa05213;hsa05214;hsa05215;hsa05218;hsa05219;hsa05220;hsa05221;hsa05223;hsa05230;hsa05231,RAF1,NA,RAF1 - mRNA,False\nSOS1 - mRNA,NM_005633.2:1635,SEBELIUS_2_C4636,SOS1,mRNA,0,,,Signaling by FGFR2 in disease;Antigen activates B Cell Receptor (BCR) leading to generation of second messengers;SHC-mediated cascade.CL.FGFR3;Signaling by FGFR3 in disease;RAFMAP kinase cascade;SHC-related events triggered by IGF1R;Signaling by FGFR4 in disease;SHC-mediated cascade.CL.FGFR1;GRB2.CL.SOS provides linkage to MAPK signaling for Integrins ;Tie2 Signaling;Role of LAT2NTALLAB on calcium mobilization;Signaling by FGFR1 in disease;Signal attenuation;NRAGE signals death through JNK;SHC-mediated cascade.CL.FGFR4;FCERI mediated Ca.PLS.2 mobilization;Constitutive Signaling by EGFRvIII;FCERI mediated MAPK activation;Signaling by ERBB2;Signaling by SCF-KIT;DAP12 signaling;Gastrin-CREB signalling pathway via PKC and MAPK;Interleukin receptor SHC signaling;IRS-related events triggered by IGF1R;Downstream signal transduction;VEGFR2 mediated cell proliferation;Frs2-mediated activation;ARMS-mediated activation;Signaling by PDGF;VEGFA-VEGFR2 Pathway;Integrin alphaIIb beta3 signaling;Cell surface interactions at the vascular wall,hsa04010;hsa04012;hsa04014;hsa04062;hsa04068;hsa04072;hsa04151;hsa04320;hsa04510;hsa04540;hsa04630;hsa04650;hsa04660;hsa04662;hsa04664;hsa04722;hsa04810;hsa04910;hsa04912;hsa04915;hsa04917;hsa05034;hsa05160;hsa05200;hsa05205;hsa05206;hsa05211;hsa05213;hsa05214;hsa05215;hsa05220;hsa05221;hsa05223;hsa05231,SOS1,NA,SOS1 - mRNA,False\nEP300 - mRNA,NM_001429.2:715,SEBELIUS_2_C4636,EP300,mRNA,0,,,LRR FLII-interacting protein 1 (LRRFIP1) activates type I IFN production;TRAF6 mediated IRF7 activation;Constitutive Signaling by NOTCH1 PEST Domain Mutants;Polo-like kinase mediated events;B-WICH complex positively regulates rRNA expression;CD209 (DC-SIGN) signaling;Circadian Clock,hsa04024;hsa04066;hsa04068;hsa04110;hsa04310;hsa04330;hsa04350;hsa04520;hsa04630;hsa04720;hsa04916;hsa04919;hsa04922;hsa05016;hsa05152;hsa05161;hsa05164;hsa05166;hsa05168;hsa05169;hsa05200;hsa05203;hsa05206;hsa05211;hsa05215,EP300,NA,EP300 - mRNA,False\nVAV2 - mRNA,NM_001134398.1:1615,SEBELIUS_2_C4636,VAV2,mRNA,0,,,Rho GTPase cycle;DAP12 signaling;Signal transduction by L1;EPH-ephrin mediated repulsion of cells;VEGFR2 mediated vascular permeability;NRAGE signals death through JNK;GPVI-mediated activation cascade;G alpha (1213) signalling events;FCERI mediated MAPK activation;Regulation of actin dynamics for phagocytic cup formation;VEGFA-VEGFR2 Pathway,hsa04015;hsa04024;hsa04062;hsa04510;hsa04650;hsa04660;hsa04662;hsa04664;hsa04666;hsa04670;hsa04810;hsa05205,VAV2,NA,VAV2 - mRNA,False\nTBP - mRNA,NM_001172085.1:587,SEBELIUS_2_C4636,TBP,mRNA,0,,,,hsa05016;hsa05166;hsa05168;hsa05169;hsa05203,TBP,NA,TBP - mRNA,False\nDNMT3A - mRNA,NM_022552.4:2056,SEBELIUS_2_C4636,DNMT3A,mRNA,0,,,,hsa01100;hsa05206,DNMT3A,NA,DNMT3A - mRNA,False\nCSNK2A2 - mRNA,NM_001896.2:1350,SEBELIUS_2_C4636,CSNK2A2,mRNA,0,,,WNT mediated activation of DVL;Signal transduction by L1,hsa04064;hsa04310;hsa04520;hsa04530;hsa05162;hsa05168;hsa05169,CSNK2A2,NA,CSNK2A2 - mRNA,False\nBRAF - mRNA,NM_004333.3:565,SEBELIUS_2_C4636,BRAF,mRNA,0,,,Spry regulation of FGF signaling;MAP2K and MAPK activation;RAF activation;CREB phosphorylation through the activation of Ras;Negative feedback regulation of MAPK pathway;ARMS-mediated activation;Negative regulation of FGFR3 signaling;Negative regulation of FGFR4 signaling;Negative regulation of FGFR2 signaling;Negative regulation of FGFR1 signaling;RAFMAP kinase cascade;Downstream signal transduction;VEGFR2 mediated cell proliferation;Signaling by SCF-KIT;Frs2-mediated activation;Gastrin-CREB signalling pathway via PKC and MAPK;Interleukin receptor SHC signaling;FCERI mediated MAPK activation;DAP12 signaling;Signaling by PDGF;VEGFA-VEGFR2 Pathway;Signaling by ERBB2;IRS-related events triggered by IGF1R;Negative regulation of MAPK pathway,hsa04010;hsa04012;hsa04015;hsa04024;hsa04062;hsa04068;hsa04150;hsa04270;hsa04510;hsa04650;hsa04720;hsa04722;hsa04726;hsa04730;hsa04810;hsa04910;hsa04914;hsa05034;hsa05160;hsa05200;hsa05205;hsa05210;hsa05211;hsa05212;hsa05213;hsa05214;hsa05215;hsa05216;hsa05218;hsa05219;hsa05220;hsa05221;hsa05223,BRAF,NA,BRAF - mRNA,False\nEPB41L1 - mRNA,NM_001258330.1:900,SEBELIUS_2_C4636,EPB41L1,mRNA,0,,,,hsa04530,EPB41L1,NA,EPB41L1 - mRNA,False\nDVL3 - mRNA,NM_004423.3:1012,SEBELIUS_2_C4636,DVL3,mRNA,0,,,Disassembly of the destruction complex and recruitment of AXIN to the membrane;WNT mediated activation of DVL;RHO GTPases Activate Formins;PCPCE pathway,hsa04310;hsa04330;hsa04390;hsa04550;hsa04916;hsa05166;hsa05200;hsa05217,DVL3,NA,DVL3 - mRNA,False\nGSK3B - mRNA,NM_002093.2:925,SEBELIUS_2_C4636,GSK3B,mRNA,0,,,Misspliced GSK3beta mutants stabilize beta-catenin;Constitutive Signaling by AKT1 E17K in Cancer;APC truncation mutants have impaired AXIN binding;T41 mutants of beta-catenin aren.SQT.t phosphorylated;AXIN missense mutants destabilize the destruction complex;S45 mutants of beta-catenin aren.SQT.t phosphorylated;Regulation of HSF1-mediated heat shock response;AKT phosphorylates targets in the cytosol;Disassembly of the destruction complex and recruitment of AXIN to the membrane;S37 mutants of beta-catenin aren.SQT.t phosphorylated;S33 mutants of beta-catenin aren.SQT.t phosphorylated;Truncations of AMER1 destabilize the destruction complex;Degradation of GLI2 by the proteasome;PIP3 activates AKT signaling;Downstream signal transduction;GAB1 signalosome;Signaling by SCF-KIT;PI3KAKT activation;Role of LAT2NTALLAB on calcium mobilization;DAP12 signaling;Signaling by PDGF;Signaling by ERBB2,hsa04012;hsa04062;hsa04110;hsa04151;hsa04310;hsa04360;hsa04390;hsa04510;hsa04550;hsa04660;hsa04662;hsa04722;hsa04728;hsa04910;hsa04916;hsa04917;hsa04919;hsa04931;hsa04932;hsa05010;hsa05160;hsa05162;hsa05164;hsa05166;hsa05169;hsa05200;hsa05210;hsa05213;hsa05215;hsa05217,GSK3B,NA,GSK3B - mRNA,False\nGNAI3 - mRNA,NM_006496.1:1150,SEBELIUS_2_C4636,GNAI3,mRNA,0,,,G alpha (i) signalling events;ADP signalling through P2Y purinoceptor 12,hsa04015;hsa04022;hsa04024;hsa04062;hsa04071;hsa04261;hsa04360;hsa04530;hsa04540;hsa04611;hsa04670;hsa04713;hsa04723;hsa04724;hsa04725;hsa04726;hsa04727;hsa04728;hsa04730;hsa04914;hsa04915;hsa04916;hsa04921;hsa04923;hsa04971;hsa05032;hsa05034;hsa05133;hsa05142;hsa05145;hsa05200,GNAI3,NA,GNAI3 - mRNA,False\nDROSHA - mRNA,NM_013235.4:1860,SEBELIUS_2_C4636,DROSHA,mRNA,0,,,MicroRNA (miRNA) biogenesis,hsa05205,DROSHA,NA,DROSHA - mRNA,False\nEPB41 - mRNA,NM_001166005.1:1330,SEBELIUS_2_C4636,EPB41,mRNA,0,,,,hsa04530,EPB41,NA,EPB41 - mRNA,False\nGUSB - mRNA,NM_000181.1:1350,SEBELIUS_2_C4636,GUSB,mRNA,0,,,,hsa01100,GUSB,NA,GUSB - mRNA,False\nPAK4 - mRNA,NM_005884.3:2130,SEBELIUS_2_C4636,PAK4,mRNA,0,,,Activation of Rac,hsa04012;hsa04014;hsa04360;hsa04510;hsa04660;hsa04810;hsa05206;hsa05211,PAK4,NA,PAK4 - mRNA,False\nPIK3R1 - mRNA,NM_181504.2:1105,SEBELIUS_2_C4636,PIK3R1,mRNA,0,,,G alpha (q) signalling events;Downstream TCR signaling;Signaling by FGFR2 in disease;PIP3 activates AKT signaling;VEGFA-VEGFR2 Pathway;Synthesis of PIPs at the plasma membrane;Constitutive Signaling by Ligand-Responsive EGFR Cancer Variants;Interleukin receptor SHC signaling;G beta.CL.gamma signalling through PI3Kgamma;Regulation of signaling by CBL;Role of phospholipids in phagocytosis;Signaling by FGFR3 in disease;Constitutive Signaling by Aberrant PI3K in Cancer;Signaling by FGFR4 in disease;CD28 dependent PI3KAkt signaling;Tie2 Signaling;Signaling by FGFR1 fusion mutants;GP1b-IX-V activation signalling;GPVI-mediated activation cascade;Nephrin interactions;Constitutive Signaling by EGFRvIII;Gastrin-CREB signalling pathway via PKC and MAPK;Downstream signal transduction;GAB1 signalosome;Signaling by SCF-KIT;PI3KAKT activation;Role of LAT2NTALLAB on calcium mobilization;DAP12 signaling;Signaling by PDGF;Signaling by ERBB2;IRS-related events triggered by IGF1R;CD28 co-stimulation;Cell surface interactions at the vascular wall;Signaling by FGFR1 in disease,hsa04012;hsa04014;hsa04015;hsa04022;hsa04024;hsa04062;hsa04066;hsa04068;hsa04070;hsa04071;hsa04072;hsa04150;hsa04151;hsa04152;hsa04210;hsa04211;hsa04213;hsa04261;hsa04370;hsa04380;hsa04510;hsa04550;hsa04611;hsa04620;hsa04630;hsa04650;hsa04660;hsa04662;hsa04664;hsa04666;hsa04668;hsa04670;hsa04722;hsa04725;hsa04750;hsa04810;hsa04910;hsa04914;hsa04915;hsa04917;hsa04919;hsa04921;hsa04923;hsa04930;hsa04931;hsa04932;hsa04933;hsa04960;hsa04973;hsa05100;hsa05142;hsa05145;hsa05146;hsa05160;hsa05161;hsa05162;hsa05164;hsa05166;hsa05169;hsa05200;hsa05203;hsa05205;hsa05210;hsa05211;hsa05212;hsa05213;hsa05214;hsa05215;hsa05218;hsa05220;hsa05221;hsa05222;hsa05223;hsa05230;hsa05231,PIK3R1,NA,PIK3R1 - mRNA,False\nPPP2R2D - mRNA,NM_001291310.1:2320,SEBELIUS_2_C4636,PPP2R2D,mRNA,0,,,MASTL Facilitates Mitotic Progression,hsa03015;hsa04071;hsa04151;hsa04152;hsa04261;hsa04390;hsa04530;hsa04728;hsa05142;hsa05160,PPP2R2D,NA,PPP2R2D - mRNA,False\nXPO5 - mRNA,NM_020750.2:375,SEBELIUS_2_C4636,XPO5,mRNA,0,,,MicroRNA (miRNA) biogenesis,,XPO5,NA,XPO5 - mRNA,False\nCDK4 - mRNA,NM_000075.2:1055,SEBELIUS_2_C4636,CDK4,mRNA,0,,,Cyclin D associated events in G1;Oncogene Induced Senescence;Ubiquitin-dependent degradation of Cyclin D1;RMTs methylate histone arginines,hsa04110;hsa04115;hsa04151;hsa04530;hsa04660;hsa04933;hsa05161;hsa05162;hsa05166;hsa05200;hsa05203;hsa05212;hsa05214;hsa05218;hsa05219;hsa05220;hsa05222;hsa05223,CDK4,NA,CDK4 - mRNA,False\nDGCR8 - mRNA,NM_022720.5:1655,SEBELIUS_2_C4636,DGCR8,mRNA,0,,,MicroRNA (miRNA) biogenesis,,DGCR8,NA,DGCR8 - mRNA,False\nSYMPK - mRNA,NM_004819.2:460,SEBELIUS_2_C4636,SYMPK,mRNA,0,,,,hsa03015;hsa04530,SYMPK,NA,SYMPK - mRNA,False\nASH1L - mRNA,NM_018489.2:5005,SEBELIUS_2_C4636,ASH1L,mRNA,0,,,,hsa04530,ASH1L,NA,ASH1L - mRNA,False\nPIK3CA - mRNA,NM_006218.2:2445,SEBELIUS_2_C4636,PIK3CA,mRNA,0,,,G alpha (q) signalling events;Downstream TCR signaling;Signaling by FGFR2 in disease;PIP3 activates AKT signaling;VEGFA-VEGFR2 Pathway;Synthesis of PIPs at the plasma membrane;Constitutive Signaling by Ligand-Responsive EGFR Cancer Variants;Interleukin receptor SHC signaling;G beta.CL.gamma signalling through PI3Kgamma;Regulation of signaling by CBL;Role of phospholipids in phagocytosis;G alpha (1213) signalling events;PI3K Cascade;Signaling by FGFR3 in disease;Constitutive Signaling by Aberrant PI3K in Cancer;Signaling by FGFR4 in disease;CD28 dependent PI3KAkt signaling;Tie2 Signaling;GPVI-mediated activation cascade;Nephrin interactions;Constitutive Signaling by EGFRvIII;Gastrin-CREB signalling pathway via PKC and MAPK;Downstream signal transduction;GAB1 signalosome;Signaling by SCF-KIT;PI3KAKT activation;Role of LAT2NTALLAB on calcium mobilization;DAP12 signaling;Signaling by PDGF;Signaling by ERBB2;IRS-related events triggered by IGF1R;CD28 co-stimulation;Cell surface interactions at the vascular wall;Signaling by FGFR1 in disease,hsa00562;hsa04012;hsa04014;hsa04015;hsa04022;hsa04024;hsa04062;hsa04066;hsa04068;hsa04070;hsa04071;hsa04072;hsa04150;hsa04151;hsa04152;hsa04210;hsa04211;hsa04213;hsa04261;hsa04370;hsa04380;hsa04510;hsa04550;hsa04611;hsa04620;hsa04630;hsa04650;hsa04660;hsa04662;hsa04664;hsa04666;hsa04668;hsa04670;hsa04722;hsa04725;hsa04750;hsa04810;hsa04910;hsa04914;hsa04915;hsa04917;hsa04919;hsa04921;hsa04923;hsa04930;hsa04931;hsa04932;hsa04933;hsa04960;hsa04973;hsa05100;hsa05142;hsa05145;hsa05146;hsa05160;hsa05161;hsa05162;hsa05164;hsa05166;hsa05169;hsa05200;hsa05203;hsa05205;hsa05206;hsa05210;hsa05211;hsa05212;hsa05213;hsa05214;hsa05215;hsa05218;hsa05220;hsa05221;hsa05222;hsa05223;hsa05230;hsa05231,PIK3CA,NA,PIK3CA - mRNA,False\nLIMK1 - mRNA,NM_002314.3:3133,SEBELIUS_2_C4636,LIMK1,mRNA,0,,,EPHB-mediated forward signaling;RHO GTPases Activate ROCKs;RHO GTPases activate PAKs;Sema3A PAK dependent Axon repulsion;Regulation of actin dynamics for phagocytic cup formation;Sema4D induced cell migration and growth-cone collapse,hsa04360;hsa04666;hsa04810,LIMK1,NA,LIMK1 - mRNA,False\nITGAV - mRNA,NM_002210.2:2615,SEBELIUS_2_C4636,ITGAV,mRNA,0,,,Molecules associated with elastic fibres;Syndecan interactions;Signal transduction by L1;Laminin interactions;ECM proteoglycans;Integrin cell surface interactions;VEGFA-VEGFR2 Pathway;Elastic fibre formation;Non-integrin membrane-ECM interactions;Cell surface interactions at the vascular wall,hsa04145;hsa04151;hsa04510;hsa04512;hsa04514;hsa04810;hsa04919;hsa05200;hsa05205;hsa05222;hsa05410;hsa05412;hsa05414,ITGAV,NA,ITGAV - mRNA,False\nPOLR2A - mRNA,NM_000937.2:3775,SEBELIUS_2_C4636,POLR2A,mRNA,0,,,RNA Polymerase II Transcription Initiation,hsa01100;hsa05016;hsa05168;hsa05169,POLR2A,NA,POLR2A - mRNA,False\nAKT1 - mRNA,NM_001014432.1:1275,SEBELIUS_2_C4636,AKT1,mRNA,0,,,TP53 Regulates Metabolic Genes;Deactivation of the beta-catenin transactivating complex;Activation of BAD and translocation to mitochondria ;eNOS activation;mTOR signalling;G beta.CL.gamma signalling through PI3Kgamma;KSRP (KHSRP) binds and destabilizes mRNA;AKT phosphorylates targets in the nucleus;VEGFR2 mediated vascular permeability;GPVI-mediated activation cascade;Tetrahydrobiopterin (BH4) synthesis# recycling# salvage and regulation;Butyrate Response Factor 1 (BRF1) binds and destabilizes mRNA;Signaling by ERBB2;PIP3 activates AKT signaling;Downstream signal transduction;GAB1 signalosome;Signaling by SCF-KIT;PI3KAKT activation;Role of LAT2NTALLAB on calcium mobilization;DAP12 signaling;Signaling by PDGF;CD28 co-stimulation;PI3K Cascade;IRS-related events triggered by IGF1R;VEGFA-VEGFR2 Pathway,hsa04010;hsa04012;hsa04014;hsa04015;hsa04022;hsa04024;hsa04062;hsa04066;hsa04068;hsa04071;hsa04072;hsa04150;hsa04151;hsa04152;hsa04210;hsa04211;hsa04213;hsa04261;hsa04370;hsa04380;hsa04510;hsa04530;hsa04550;hsa04611;hsa04620;hsa04630;hsa04660;hsa04662;hsa04664;hsa04666;hsa04668;hsa04722;hsa04725;hsa04728;hsa04910;hsa04914;hsa04915;hsa04917;hsa04919;hsa04920;hsa04922;hsa04923;hsa04931;hsa04932;hsa04933;hsa04973;hsa05142;hsa05145;hsa05152;hsa05160;hsa05161;hsa05162;hsa05164;hsa05166;hsa05169;hsa05200;hsa05205;hsa05210;hsa05211;hsa05212;hsa05213;hsa05214;hsa05215;hsa05218;hsa05220;hsa05221;hsa05222;hsa05223;hsa05230;hsa05231,AKT1,NA,AKT1 - mRNA,False\nEGFR - mRNA,NM_201282.1:360,SEBELIUS_2_C4636,EGFR,mRNA,0,,,Constitutive Signaling by Aberrant PI3K in Cancer;Inhibition of Signaling by Overexpressed EGFR;PLCG1 events in ERBB2 signaling;EGFR interacts with phospholipase C-gamma;EGFR Transactivation by Gastrin;Constitutive Signaling by Ligand-Responsive EGFR Cancer Variants;Constitutive Signaling by EGFRvIII;GAB1 signalosome;Downstream signal transduction;Signaling by SCF-KIT;PI3KAKT activation;Role of LAT2NTALLAB on calcium mobilization;DAP12 signaling;Signaling by PDGF;Signaling by ERBB2;VEGFR2 mediated cell proliferation;Frs2-mediated activation;ARMS-mediated activation;Gastrin-CREB signalling pathway via PKC and MAPK;Interleukin receptor SHC signaling;FCERI mediated MAPK activation;VEGFA-VEGFR2 Pathway;IRS-related events triggered by IGF1R,hsa04010;hsa04012;hsa04014;hsa04015;hsa04020;hsa04060;hsa04066;hsa04068;hsa04072;hsa04144;hsa04151;hsa04320;hsa04510;hsa04520;hsa04540;hsa04810;hsa04912;hsa04915;hsa04921;hsa05120;hsa05160;hsa05200;hsa05205;hsa05206;hsa05212;hsa05213;hsa05214;hsa05215;hsa05218;hsa05219;hsa05223;hsa05230;hsa05231,EGFR,NA,EGFR - mRNA,False\nPXN - mRNA,NM_001080855.1:2495,SEBELIUS_2_C4636,PXN,mRNA,0,,,Smooth Muscle Contraction;GAB1 signalosome;VEGFA-VEGFR2 Pathway;Cell-extracellular matrix interactions,hsa04062;hsa04370;hsa04510;hsa04670;hsa04810;hsa05100;hsa05203;hsa05205,PXN,NA,PXN - mRNA,False\nCTTN - mRNA,NM_001184740.1:240,SEBELIUS_2_C4636,CTTN,mRNA,0,,,RHO GTPases activate PAKs,hsa04530;hsa05100;hsa05130;hsa05131;hsa05205,CTTN,NA,CTTN - mRNA,False\nPDCD4 - mRNA,NM_014456.3:1115,SEBELIUS_2_C4636,PDCD4,mRNA,0,,,,hsa05205;hsa05206,PDCD4,NA,PDCD4 - mRNA,False\nCCNG2 - mRNA,NM_004354.2:336,SEBELIUS_2_C4636,CCNG2,mRNA,0,,,,hsa04068;hsa04115,CCNG2,NA,CCNG2 - mRNA,False\nYBX3 - mRNA,NM_003651.3:660,SEBELIUS_2_C4636,YBX3,mRNA,0,,,,hsa04530,YBX3,NA,YBX3 - mRNA,False\nSPRY2 - mRNA,NM_005842.2:1481,SEBELIUS_2_C4636,SPRY2,mRNA,0,,,Spry regulation of FGF signaling;Negative regulation of FGFR3 signaling;Negative regulation of FGFR4 signaling;Negative regulation of FGFR2 signaling;Negative regulation of FGFR1 signaling,hsa05206,SPRY2,NA,SPRY2 - mRNA,False\nEXOC4 - mRNA,NM_001037126.1:560,SEBELIUS_2_C4636,EXOC4,mRNA,0,,,,hsa04530,EXOC4,NA,EXOC4 - mRNA,False\nPRKCI - mRNA,NM_002740.4:530,SEBELIUS_2_C4636,PRKCI,mRNA,0,,,Tight junction interactions,hsa04015;hsa04144;hsa04390;hsa04530;hsa04611;hsa04910,PRKCI,NA,PRKCI - mRNA,False\nSMAD2 - mRNA,NM_005901.5:1678,SEBELIUS_2_C4636,SMAD2,mRNA,0,,,Transcriptional regulation of pluripotent stem cells;TGF-beta receptor signaling activates SMADs,hsa04068;hsa04110;hsa04144;hsa04350;hsa04390;hsa04520;hsa04550;hsa04933;hsa05142;hsa05166;hsa05200;hsa05205;hsa05210;hsa05212;hsa05321,SMAD2,NA,SMAD2 - mRNA,False\nPRKCA - mRNA,NM_002737.2:680,SEBELIUS_2_C4636,PRKCA,mRNA,0,,,WNT5A-dependent internalization of FZD4;Disinhibition of SNARE formation;Trafficking of AMPA receptors;Downstream signal transduction;EGFR interacts with phospholipase C-gamma;Phospholipase C-mediated cascade.CL. FGFR1;PLCG1 events in ERBB2 signaling;DAP12 signaling;PLC beta mediated events;Signaling by PDGF;Signaling by ERBB2;Signaling by SCF-KIT;Non-integrin membrane-ECM interactions;Gastrin-CREB signalling pathway via PKC and MAPK;VEGFA-VEGFR2 Pathway;PCPCE pathway,hsa04010;hsa04012;hsa04014;hsa04015;hsa04020;hsa04066;hsa04070;hsa04071;hsa04072;hsa04150;hsa04151;hsa04261;hsa04270;hsa04310;hsa04370;hsa04510;hsa04530;hsa04540;hsa04650;hsa04664;hsa04666;hsa04670;hsa04713;hsa04720;hsa04723;hsa04724;hsa04725;hsa04726;hsa04727;hsa04728;hsa04730;hsa04750;hsa04912;hsa04916;hsa04919;hsa04921;hsa04933;hsa04960;hsa04971;hsa04972;hsa05031;hsa05032;hsa05110;hsa05130;hsa05146;hsa05161;hsa05164;hsa05200;hsa05205;hsa05206;hsa05214;hsa05223;hsa05231,PRKCA,NA,PRKCA - mRNA,False\nMAPK3 - mRNA,NM_001040056.1:580,SEBELIUS_2_C4636,MAPK3,mRNA,0,,,ERKs are inactivated;ISG15 antiviral mechanism;RHO GTPases Activate WASPs and WAVEs;Senescence-Associated Secretory Phenotype (SASP);Regulation of HSF1-mediated heat shock response;Thrombin signalling through proteinase activated receptors (PARs);Negative regulation of FGFR4 signaling;Oxidative Stress Induced Senescence;Signal attenuation;RNA Polymerase I Promoter Opening;Negative regulation of FGFR3 signaling;Negative feedback regulation of MAPK pathway;Growth hormone receptor signaling;Golgi Cisternae Pericentriolar Stack Reorganization;FCERI mediated MAPK activation;Negative regulation of MAPK pathway;RAFMAP kinase cascade;Downstream signal transduction;VEGFR2 mediated cell proliferation;Signaling by SCF-KIT;Frs2-mediated activation;ARMS-mediated activation;Gastrin-CREB signalling pathway via PKC and MAPK;Interleukin receptor SHC signaling;DAP12 signaling;Signaling by PDGF;VEGFA-VEGFR2 Pathway;Signaling by ERBB2;IRS-related events triggered by IGF1R,hsa04010;hsa04012;hsa04014;hsa04015;hsa04022;hsa04024;hsa04062;hsa04066;hsa04068;hsa04071;hsa04072;hsa04114;hsa04150;hsa04151;hsa04261;hsa04270;hsa04320;hsa04350;hsa04360;hsa04370;hsa04380;hsa04510;hsa04520;hsa04540;hsa04550;hsa04611;hsa04620;hsa04621;hsa04650;hsa04660;hsa04662;hsa04664;hsa04666;hsa04668;hsa04713;hsa04720;hsa04722;hsa04723;hsa04724;hsa04725;hsa04726;hsa04730;hsa04810;hsa04910;hsa04912;hsa04914;hsa04915;hsa04916;hsa04917;hsa04919;hsa04921;hsa04930;hsa04933;hsa04960;hsa05010;hsa05020;hsa05034;hsa05131;hsa05132;hsa05133;hsa05140;hsa05142;hsa05145;hsa05152;hsa05160;hsa05161;hsa05164;hsa05200;hsa05203;hsa05205;hsa05210;hsa05211;hsa05212;hsa05213;hsa05214;hsa05215;hsa05216;hsa05218;hsa05219;hsa05220;hsa05221;hsa05223;hsa05230;hsa05231,MAPK3,NA,MAPK3 - mRNA,False\nSMAD3 - mRNA,NM_005902.3:4220,SEBELIUS_2_C4636,SMAD3,mRNA,0,,,Signaling by NODAL;TGF-beta receptor signaling activates SMADs,hsa04068;hsa04110;hsa04144;hsa04310;hsa04350;hsa04390;hsa04520;hsa04550;hsa04933;hsa05142;hsa05161;hsa05166;hsa05200;hsa05210;hsa05212;hsa05220;hsa05321,SMAD3,NA,SMAD3 - mRNA,False\nADAM9 - mRNA,NM_001005845.1:695,SEBELIUS_2_C4636,ADAM9,mRNA,0,,,,,ADAM9,NA,ADAM9 - mRNA,False\nTRIO - mRNA,NM_007118.2:1735,SEBELIUS_2_C4636,TRIO,mRNA,0,,,G alpha (q) signalling events;Rho GTPase cycle;NRAGE signals death through JNK;DCC mediated attractive signaling;G alpha (1213) signalling events;Gastrin-CREB signalling pathway via PKC and MAPK,,TRIO,NA,TRIO - mRNA,False\nG6PD - mRNA,NM_000402.2:1155,SEBELIUS_2_C4636,G6PD,mRNA,0,,,TP53 Regulates Metabolic Genes,hsa01100;hsa05230,G6PD,NA,G6PD - mRNA,False\nVCL - mRNA,NM_014000.2:4780,SEBELIUS_2_C4636,VCL,mRNA,0,,,Smooth Muscle Contraction;MAP2K and MAPK activation;Platelet degranulation ;RAFMAP kinase cascade;Downstream signal transduction;VEGFR2 mediated cell proliferation;Signaling by SCF-KIT;Frs2-mediated activation;ARMS-mediated activation;Gastrin-CREB signalling pathway via PKC and MAPK;Interleukin receptor SHC signaling;FCERI mediated MAPK activation;DAP12 signaling;Signaling by PDGF;VEGFA-VEGFR2 Pathway;Signaling by ERBB2;IRS-related events triggered by IGF1R,hsa04510;hsa04520;hsa04670;hsa04810;hsa05100;hsa05131;hsa05146,VCL,NA,VCL - mRNA,False\nACTN1 - mRNA,NM_001102.3:2760,SEBELIUS_2_C4636,ACTN1,mRNA,0,,,Syndecan interactions;Nephrin interactions;Platelet degranulation ;Non-integrin membrane-ECM interactions;Cell-extracellular matrix interactions,hsa04510;hsa04520;hsa04530;hsa04670;hsa04810;hsa05146;hsa05203;hsa05412,ACTN1,NA,ACTN1 - mRNA,False\nPPP2CB - mRNA,NM_001009552.1:1075,SEBELIUS_2_C4636,PPP2CB,mRNA,0,,,MASTL Facilitates Mitotic Progression;Misspliced GSK3beta mutants stabilize beta-catenin;T41 mutants of beta-catenin aren.SQT.t phosphorylated;ERKs are inactivated;Glycolysis;Negative regulation of MAPK pathway;Disassembly of the destruction complex and recruitment of AXIN to the membrane;S37 mutants of beta-catenin aren.SQT.t phosphorylated;Cyclin AB1 associated events during G2M transition;RAF activation;RHO GTPases Activate Formins;S33 mutants of beta-catenin aren.SQT.t phosphorylated;DARPP-32 events;Beta-catenin phosphorylation cascade;Spry regulation of FGF signaling;Truncations of AMER1 destabilize the destruction complex;CTLA4 inhibitory signaling;AXIN missense mutants destabilize the destruction complex;S45 mutants of beta-catenin aren.SQT.t phosphorylated;Cyclin D associated events in G1;Platelet sensitization by LDL;Resolution of Sister Chromatid Cohesion;Inhibition of replication initiation of damaged DNA by RB1E2F1;RAFMAP kinase cascade;Downstream signal transduction;VEGFR2 mediated cell proliferation;Signaling by SCF-KIT;Frs2-mediated activation;ARMS-mediated activation;Gastrin-CREB signalling pathway via PKC and MAPK;Interleukin receptor SHC signaling;FCERI mediated MAPK activation;DAP12 signaling;Signaling by PDGF;VEGFA-VEGFR2 Pathway;Signaling by ERBB2;IRS-related events triggered by IGF1R;Negative regulation of FGFR3 signaling;Negative regulation of FGFR4 signaling;Negative regulation of FGFR2 signaling;Negative regulation of FGFR1 signaling,hsa03015;hsa04071;hsa04114;hsa04151;hsa04152;hsa04261;hsa04350;hsa04390;hsa04530;hsa04728;hsa04730;hsa05142;hsa05160,PPP2CB,NA,PPP2CB - mRNA,False\nCASK - mRNA,NM_001126054.2:2190,SEBELIUS_2_C4636,CASK,mRNA,0,,,Syndecan interactions;Nephrin interactions;Non-integrin membrane-ECM interactions,hsa04530,CASK,NA,CASK - mRNA,False\nPPP2R2A - mRNA,NM_001177591.1:430,SEBELIUS_2_C4636,PPP2R2A,mRNA,0,,,Nonsense Mediated Decay (NMD) enhanced by the Exon Junction Complex (EJC);Cyclin D associated events in G1;Cyclin AB1 associated events during G2M transition,hsa03015;hsa04071;hsa04151;hsa04152;hsa04261;hsa04390;hsa04530;hsa04728;hsa05142;hsa05160,PPP2R2A,NA,PPP2R2A - mRNA,False\nPPP2R1B - mRNA,NM_002716.4:1445,SEBELIUS_2_C4636,PPP2R1B,mRNA,0,,,MASTL Facilitates Mitotic Progression;Misspliced GSK3beta mutants stabilize beta-catenin;T41 mutants of beta-catenin aren.SQT.t phosphorylated;ERKs are inactivated;Glycolysis;Negative regulation of MAPK pathway;Disassembly of the destruction complex and recruitment of AXIN to the membrane;S37 mutants of beta-catenin aren.SQT.t phosphorylated;Cyclin AB1 associated events during G2M transition;RAF activation;RHO GTPases Activate Formins;S33 mutants of beta-catenin aren.SQT.t phosphorylated;DARPP-32 events;Beta-catenin phosphorylation cascade;Truncations of AMER1 destabilize the destruction complex;CTLA4 inhibitory signaling;AXIN missense mutants destabilize the destruction complex;S45 mutants of beta-catenin aren.SQT.t phosphorylated;Cyclin D associated events in G1;Platelet sensitization by LDL;Resolution of Sister Chromatid Cohesion;Inhibition of replication initiation of damaged DNA by RB1E2F1;RAFMAP kinase cascade;Downstream signal transduction;VEGFR2 mediated cell proliferation;Signaling by SCF-KIT;Frs2-mediated activation;ARMS-mediated activation;Gastrin-CREB signalling pathway via PKC and MAPK;Interleukin receptor SHC signaling;FCERI mediated MAPK activation;DAP12 signaling;Signaling by PDGF;VEGFA-VEGFR2 Pathway;Signaling by ERBB2;IRS-related events triggered by IGF1R,hsa03015;hsa04071;hsa04114;hsa04151;hsa04152;hsa04261;hsa04350;hsa04390;hsa04530;hsa04728;hsa04730;hsa05142;hsa05160,PPP2R1B,NA,PPP2R1B - mRNA,False\nDOCK1 - mRNA,NM_001380.3:840,SEBELIUS_2_C4636,DOCK1,mRNA,0,,,DCC mediated attractive signaling;Factors involved in megakaryocyte development and platelet production;VEGFA-VEGFR2 Pathway;Regulation of actin dynamics for phagocytic cup formation,hsa04510;hsa04810;hsa05100;hsa05131,DOCK1,NA,DOCK1 - mRNA,False\nFLNA - mRNA,NM_001456.3:7335,SEBELIUS_2_C4636,FLNA,mRNA,0,,,RHO GTPases activate PAKs;Platelet degranulation ;Cell-extracellular matrix interactions,hsa04010;hsa04510;hsa05132;hsa05205,FLNA,NA,FLNA - mRNA,False\nRAB13 - mRNA,NM_002870.3:332,SEBELIUS_2_C4636,RAB13,mRNA,0,,,Translocation of GLUT4 to the plasma membrane,hsa04530,RAB13,NA,RAB13 - mRNA,False\nMYH9 - mRNA,NM_002473.3:3153,SEBELIUS_2_C4636,MYH9,mRNA,0,,,EPHA-mediated growth cone collapse;RHO GTPases Activate ROCKs;RHO GTPases activate CIT;RHO GTPases activate PAKs;RHO GTPases activate PKNs;Regulation of actin dynamics for phagocytic cup formation;Sema4D induced cell migration and growth-cone collapse,hsa04530;hsa04810;hsa05132,MYH9,NA,MYH9 - mRNA,False\nVAPA - mRNA,NM_194434.2:763,SEBELIUS_2_C4636,VAPA,mRNA,0,,,,hsa04530,VAPA,NA,VAPA - mRNA,False\nMAP2K1 - mRNA,NM_002755.2:970,SEBELIUS_2_C4636,MAP2K1,mRNA,0,,,RAF activation;Uptake and function of anthrax toxins;Negative feedback regulation of MAPK pathway;Interleukin-1 signaling;RAFMAP kinase cascade;Downstream signal transduction;VEGFR2 mediated cell proliferation;Signaling by SCF-KIT;Frs2-mediated activation;ARMS-mediated activation;Gastrin-CREB signalling pathway via PKC and MAPK;Interleukin receptor SHC signaling;FCERI mediated MAPK activation;DAP12 signaling;Signaling by PDGF;VEGFA-VEGFR2 Pathway;Signaling by ERBB2;IRS-related events triggered by IGF1R;Negative regulation of MAPK pathway,hsa04010;hsa04012;hsa04014;hsa04015;hsa04022;hsa04024;hsa04062;hsa04066;hsa04068;hsa04071;hsa04072;hsa04114;hsa04151;hsa04270;hsa04320;hsa04370;hsa04380;hsa04510;hsa04540;hsa04550;hsa04620;hsa04650;hsa04660;hsa04662;hsa04664;hsa04666;hsa04668;hsa04720;hsa04722;hsa04725;hsa04726;hsa04730;hsa04810;hsa04910;hsa04912;hsa04914;hsa04915;hsa04916;hsa04917;hsa04919;hsa04921;hsa05020;hsa05034;hsa05161;hsa05164;hsa05200;hsa05205;hsa05206;hsa05210;hsa05211;hsa05212;hsa05213;hsa05214;hsa05215;hsa05216;hsa05218;hsa05219;hsa05220;hsa05221;hsa05223;hsa05230;hsa05231,MAP2K1,NA,MAP2K1 - mRNA,False\nPTK2 - mRNA,NM_153831.2:965,SEBELIUS_2_C4636,PTK2,mRNA,0,,,Apoptotic cleavage of cellular proteins;RAFMAP kinase cascade;RHO GTPases Activate WASPs and WAVEs;DCC mediated attractive signaling;p130Cas linkage to MAPK signaling for integrins;EPHB-mediated forward signaling;EPHA-mediated growth cone collapse;Regulation of actin dynamics for phagocytic cup formation;Downstream signal transduction;VEGFR2 mediated cell proliferation;Signaling by SCF-KIT;Frs2-mediated activation;ARMS-mediated activation;Gastrin-CREB signalling pathway via PKC and MAPK;Interleukin receptor SHC signaling;FCERI mediated MAPK activation;DAP12 signaling;Signaling by PDGF;VEGFA-VEGFR2 Pathway;Signaling by ERBB2;IRS-related events triggered by IGF1R;Integrin alphaIIb beta3 signaling,hsa04012;hsa04062;hsa04151;hsa04360;hsa04370;hsa04510;hsa04670;hsa04810;hsa05100;hsa05146;hsa05200;hsa05202;hsa05205;hsa05222,PTK2,NA,PTK2 - mRNA,False\nPRKAA1 - mRNA,NM_006251.5:1455,SEBELIUS_2_C4636,PRKAA1,mRNA,0,,,TP53 Regulates Metabolic Genes;PI3K Cascade;IRS-related events triggered by IGF1R,hsa04068;hsa04150;hsa04151;hsa04152;hsa04211;hsa04213;hsa04910;hsa04920;hsa04921;hsa04922;hsa04931;hsa04932;hsa05410,PRKAA1,NA,PRKAA1 - mRNA,False\nIQGAP1 - mRNA,NM_003870.3:820,SEBELIUS_2_C4636,IQGAP1,mRNA,0,,,MAP2K and MAPK activation;Nephrin interactions;RHO GTPases activate IQGAPs;RAFMAP kinase cascade;Downstream signal transduction;VEGFR2 mediated cell proliferation;Signaling by SCF-KIT;Frs2-mediated activation;ARMS-mediated activation;Gastrin-CREB signalling pathway via PKC and MAPK;Interleukin receptor SHC signaling;FCERI mediated MAPK activation;DAP12 signaling;Signaling by PDGF;VEGFA-VEGFR2 Pathway;Signaling by ERBB2;IRS-related events triggered by IGF1R,hsa04520;hsa04810;hsa05205,IQGAP1,NA,IQGAP1 - mRNA,False\nWASF2 - mRNA,NM_006990.2:485,SEBELIUS_2_C4636,WASF2,mRNA,0,,,RHO GTPases Activate WASPs and WAVEs;VEGFA-VEGFR2 Pathway;Regulation of actin dynamics for phagocytic cup formation,hsa04520;hsa04666;hsa04810;hsa05100;hsa05131;hsa05132;hsa05231,WASF2,NA,WASF2 - mRNA,False\nTJP1 - mRNA,NM_003257.3:6277,SEBELIUS_2_C4636,TJP1,mRNA,0,,,Apoptotic cleavage of cellular proteins,hsa04520;hsa04530;hsa04540;hsa05110;hsa05120;hsa05132,TJP1,NA,TJP1 - mRNA,False\nITGB5 - mRNA,NM_002213.3:2560,SEBELIUS_2_C4636,ITGB5,mRNA,0,,,Molecules associated with elastic fibres;Syndecan interactions;Smooth Muscle Contraction;ECM proteoglycans;Integrin cell surface interactions;Elastic fibre formation;Non-integrin membrane-ECM interactions,hsa04145;hsa04151;hsa04510;hsa04512;hsa04810;hsa05205;hsa05410;hsa05412;hsa05414,ITGB5,NA,ITGB5 - mRNA,False\nGNAI2 - mRNA,NM_002070.2:1000,SEBELIUS_2_C4636,GNAI2,mRNA,0,,,ADP signalling through P2Y purinoceptor 12;Adrenaline#noradrenaline inhibits insulin secretion,hsa04015;hsa04022;hsa04024;hsa04062;hsa04071;hsa04261;hsa04360;hsa04530;hsa04540;hsa04611;hsa04670;hsa04713;hsa04723;hsa04724;hsa04725;hsa04726;hsa04727;hsa04728;hsa04730;hsa04914;hsa04915;hsa04916;hsa04921;hsa04923;hsa04971;hsa05032;hsa05034;hsa05133;hsa05142;hsa05145;hsa05200,GNAI2,NA,GNAI2 - mRNA,False\nACTN4 - mRNA,NM_004924.4:1768,SEBELIUS_2_C4636,ACTN4,mRNA,0,,,Nephrin interactions;Platelet degranulation ,hsa04510;hsa04520;hsa04530;hsa04670;hsa04810;hsa05146;hsa05203;hsa05412,ACTN4,NA,ACTN4 - mRNA,False\nRAC1 - mRNA,NM_198829.1:1250,SEBELIUS_2_C4636,RAC1,mRNA,0,,,MAPK6MAPK4 signaling;Signaling by SCF-KIT;Inactivation of Cdc42 and Rac;DAP12 signaling;Ephrin signaling;EPH-ephrin mediated repulsion of cells;Activation of Rac;RHO GTPases Activate Formins;VEGFR2 mediated vascular permeability;RHO GTPases activate PAKs;Sema4D mediated inhibition of cell attachment and migration;G alpha (1213) signalling events;Regulation of actin dynamics for phagocytic cup formation;Sema4D induced cell migration and growth-cone collapse;Signal transduction by L1;RHO GTPases activate CIT;RHO GTPases Activate WASPs and WAVEs;Factors involved in megakaryocyte development and platelet production;RHO GTPases activate KTN1;CD28 dependent Vav1 pathway;DSCAM interactions;NRAGE signals death through JNK;GPVI-mediated activation cascade;Sema3A PAK dependent Axon repulsion;RHO GTPases activate PKNs;SEMA3A-Plexin repulsion signaling by inhibiting Integrin adhesion;RHO GTPases activate IQGAPs;PCPCE pathway;VEGFA-VEGFR2 Pathway;CD28 co-stimulation,hsa04010;hsa04014;hsa04015;hsa04024;hsa04062;hsa04071;hsa04145;hsa04151;hsa04310;hsa04360;hsa04370;hsa04380;hsa04510;hsa04520;hsa04620;hsa04650;hsa04662;hsa04664;hsa04666;hsa04670;hsa04722;hsa04810;hsa04932;hsa04933;hsa04972;hsa05100;hsa05120;hsa05131;hsa05132;hsa05200;hsa05203;hsa05205;hsa05210;hsa05211;hsa05212;hsa05231;hsa05416,RAC1,NA,RAC1 - mRNA,False\nRAB11A - mRNA,NM_004663.3:710,SEBELIUS_2_C4636,RAB11A,mRNA,0,,,Anchoring of the basal body to the plasma membrane,hsa04144;hsa04972,RAB11A,NA,RAB11A - mRNA,False\nROCK2 - mRNA,NM_004850.3:3140,SEBELIUS_2_C4636,ROCK2,mRNA,0,,,EPHB-mediated forward signaling;EPHA-mediated growth cone collapse;RHO GTPases Activate ROCKs;G alpha (1213) signalling events;VEGFA-VEGFR2 Pathway;Sema4D induced cell migration and growth-cone collapse,hsa04022;hsa04024;hsa04062;hsa04071;hsa04270;hsa04310;hsa04360;hsa04510;hsa04611;hsa04670;hsa04810;hsa04921;hsa05130;hsa05131;hsa05132;hsa05200;hsa05205,ROCK2,NA,ROCK2 - mRNA,False\nCSNK2B - mRNA,NM_001320.5:375,SEBELIUS_2_C4636,CSNK2B,mRNA,0,,,WNT mediated activation of DVL;Signal transduction by L1,hsa04064;hsa04310;hsa04520;hsa04530;hsa05162;hsa05168;hsa05169,CSNK2B,NA,CSNK2B - mRNA,False\nNCKAP1 - mRNA,NM_013436.3:2353,SEBELIUS_2_C4636,NCKAP1,mRNA,0,,,RHO GTPases Activate WASPs and WAVEs;VEGFA-VEGFR2 Pathway;Regulation of actin dynamics for phagocytic cup formation,hsa04810,NCKAP1,NA,NCKAP1 - mRNA,False\nPPP1CB - mRNA,NM_002709.2:393,SEBELIUS_2_C4636,PPP1CB,mRNA,0,,,RHO GTPases Activate ROCKs;Downregulation of TGF-beta receptor signaling;RHO GTPases activate PKNs;Circadian Clock;Regulation of PLK1 Activity at G2M Transition;TGF-beta receptor signaling activates SMADs,hsa03015;hsa04022;hsa04024;hsa04114;hsa04261;hsa04270;hsa04390;hsa04510;hsa04611;hsa04720;hsa04728;hsa04750;hsa04810;hsa04910;hsa04921;hsa04931;hsa05031;hsa05034;hsa05168;hsa05205,PPP1CB,NA,PPP1CB - mRNA,False\nTUBB - mRNA,NM_178014.2:1955,SEBELIUS_2_C4636,TUBB,mRNA,0,,,,hsa04145;hsa04540;hsa05130,TUBB,NA,TUBB - mRNA,False\nALDH18A1 - mRNA,NM_001017423.1:1980,SEBELIUS_2_C4636,ALDH18A1,mRNA,0,,,,hsa01100,ALDH18A1,NA,ALDH18A1 - mRNA,False\nCLDN4 - mRNA,NM_001305.3:1242,SEBELIUS_2_C4636,CLDN4,mRNA,0,,,Tight junction interactions,hsa04514;hsa04530;hsa04670;hsa05160,CLDN4,NA,CLDN4 - mRNA,False\nF11R - mRNA,NM_144503.1:3650,SEBELIUS_2_C4636,F11R,mRNA,0,,,TGF-beta receptor signaling in EMT (epithelial to mesenchymal transition);Tight junction interactions;Integrin cell surface interactions;Cell surface interactions at the vascular wall,hsa04514;hsa04530;hsa04670;hsa05120,F11R,NA,F11R - mRNA,False\nITGA6 - mRNA,NM_000210.1:3065,SEBELIUS_2_C4636,ITGA6,mRNA,0,,,Syndecan interactions;Assembly of collagen fibrils and other multimeric structures;Basigin interactions;Non-integrin membrane-ECM interactions;Cell surface interactions at the vascular wall,hsa04151;hsa04510;hsa04512;hsa04514;hsa04640;hsa04810;hsa05145;hsa05200;hsa05222;hsa05410;hsa05412;hsa05414,ITGA6,NA,ITGA6 - mRNA,False\nHPRT1 - mRNA,NM_000194.1:240,SEBELIUS_2_C4636,HPRT1,mRNA,0,,,,hsa01100,HPRT1,NA,HPRT1 - mRNA,False\nDIAPH1 - mRNA,NM_005219.4:1398,SEBELIUS_2_C4636,DIAPH1,mRNA,0,,,RHO GTPases Activate Formins,hsa04510;hsa04810;hsa04933;hsa05131,DIAPH1,NA,DIAPH1 - mRNA,False\nYES1 - mRNA,NM_005433.3:265,SEBELIUS_2_C4636,YES1,mRNA,0,,,Signaling by ERBB2;EPHB-mediated forward signaling;EPHA-mediated growth cone collapse;Regulation of KIT signaling;EPH-ephrin mediated repulsion of cells;Regulation of signaling by CBL;CD28 co-stimulation;CTLA4 inhibitory signaling;Signaling by SCF-KIT;Cell surface interactions at the vascular wall,hsa04520;hsa04530,YES1,NA,YES1 - mRNA,False\nCSNK2A1 - mRNA,NM_177559.2:1930,SEBELIUS_2_C4636,CSNK2A1,mRNA,0,,,WNT mediated activation of DVL;Signal transduction by L1,hsa04064;hsa04310;hsa04520;hsa04530;hsa05162;hsa05168;hsa05169,CSNK2A1,NA,CSNK2A1 - mRNA,False\nCRKL - mRNA,NM_005207.3:1372,SEBELIUS_2_C4636,CRKL,mRNA,0,,,Downstream signal transduction;Regulation of signaling by CBL;Frs2-mediated activation;Signaling by PDGF,hsa04010;hsa04012;hsa04015;hsa04062;hsa04510;hsa04666;hsa04722;hsa04810;hsa04910;hsa05100;hsa05131;hsa05200;hsa05206;hsa05211;hsa05220,CRKL,NA,CRKL - mRNA,False\nRHOA - mRNA,NM_001664.2:1230,SEBELIUS_2_C4636,RHOA,mRNA,0,,,RHO GTPases activate KTN1;EPHB-mediated forward signaling;Rho GTPase cycle;EPHA-mediated growth cone collapse;RHO GTPases Activate ROCKs;Axonal growth inhibition (RHOA activation);PI3KAKT activation;G beta.CL.gamma signalling through PI3Kgamma;RHO GTPases Activate Formins;RHO GTPases activate PKNs;GPVI-mediated activation cascade;G alpha (1213) signalling events;Sema4D mediated inhibition of cell attachment and migration;PCPCE pathway;Sema4D induced cell migration and growth-cone collapse,hsa04014;hsa04015;hsa04022;hsa04024;hsa04062;hsa04071;hsa04072;hsa04144;hsa04270;hsa04310;hsa04350;hsa04360;hsa04510;hsa04520;hsa04530;hsa04611;hsa04660;hsa04670;hsa04722;hsa04810;hsa04921;hsa04972;hsa05100;hsa05130;hsa05133;hsa05152;hsa05200;hsa05203;hsa05205;hsa05206;hsa05210,RHOA,NA,RHOA - mRNA,False\nCLTC - mRNA,NM_004859.2:290,SEBELIUS_2_C4636,CLTC,mRNA,0,,,MHC class II antigen presentation;PCPCE pathway,hsa04144;hsa05016;hsa05100,CLTC,NA,CLTC - mRNA,False\nHSPA1B - mRNA,NM_005346.3:2095,SEBELIUS_2_C4636,HSPA1B,mRNA,0,,,Regulation of HSF1-mediated heat shock response;AUF1 (hnRNP D0) binds and destabilizes mRNA,hsa04010;hsa04141;hsa04144;hsa04213;hsa04915;hsa05145;hsa05162;hsa05164;hsa05169,HSPA1B,NA,HSPA1B - mRNA,False\nCTBP1 - mRNA,NM_001328.2:425,SEBELIUS_2_C4636,CTBP1,mRNA,0,,,Deactivation of the beta-catenin transactivating complex;Repression of WNT target genes,hsa04310;hsa04330;hsa05200;hsa05220,CTBP1,NA,CTBP1 - mRNA,False\nAKT2 - mRNA,NM_001626.4:699,SEBELIUS_2_C4636,AKT2,mRNA,0,,,TP53 Regulates Metabolic Genes;PDE3B signalling;AKT phosphorylates targets in the cytosol;Negative regulation of the PI3KAKT network;Deactivation of the beta-catenin transactivating complex;Activation of BAD and translocation to mitochondria ;G beta.CL.gamma signalling through PI3Kgamma;AKT phosphorylates targets in the nucleus;VEGFR2 mediated vascular permeability;GPVI-mediated activation cascade;Signaling by ERBB2;PI3K Cascade;IRS-related events triggered by IGF1R;mTOR signalling;PIP3 activates AKT signaling;Downstream signal transduction;GAB1 signalosome;Signaling by SCF-KIT;PI3KAKT activation;Role of LAT2NTALLAB on calcium mobilization;DAP12 signaling;Signaling by PDGF;CD28 co-stimulation;VEGFA-VEGFR2 Pathway,hsa04010;hsa04012;hsa04014;hsa04015;hsa04022;hsa04024;hsa04062;hsa04066;hsa04068;hsa04071;hsa04072;hsa04150;hsa04151;hsa04152;hsa04210;hsa04211;hsa04213;hsa04261;hsa04370;hsa04380;hsa04510;hsa04530;hsa04550;hsa04611;hsa04620;hsa04630;hsa04660;hsa04662;hsa04664;hsa04666;hsa04668;hsa04722;hsa04725;hsa04728;hsa04910;hsa04914;hsa04915;hsa04917;hsa04919;hsa04920;hsa04922;hsa04923;hsa04931;hsa04932;hsa04933;hsa04973;hsa05142;hsa05145;hsa05152;hsa05160;hsa05161;hsa05162;hsa05164;hsa05166;hsa05169;hsa05200;hsa05205;hsa05210;hsa05211;hsa05212;hsa05213;hsa05214;hsa05215;hsa05218;hsa05220;hsa05221;hsa05222;hsa05223;hsa05230;hsa05231,AKT2,NA,AKT2 - mRNA,False\nABCF1 - mRNA,NM_001090.2:850,SEBELIUS_2_C4636,ABCF1,mRNA,0,,,,,ABCF1,NA,ABCF1 - mRNA,False\nNRAS - mRNA,NM_002524.3:877,SEBELIUS_2_C4636,NRAS,mRNA,0,,,Signaling by FGFR2 in disease;Regulation of RAS by GAPs;SHC-mediated cascade.CL.FGFR3;Signaling by FGFR3 in disease;MAP2K and MAPK activation;SHC-related events triggered by IGF1R;Signaling by FGFR4 in disease;SHC-mediated cascade.CL.FGFR1;Tie2 Signaling;Signaling by FGFR1 in disease;Activation of RAS in B cells;SHC-mediated cascade.CL.FGFR4;Constitutive Signaling by EGFRvIII;CD209 (DC-SIGN) signaling;FCERI mediated MAPK activation;RAFMAP kinase cascade;Downstream signal transduction;VEGFR2 mediated cell proliferation;Signaling by SCF-KIT;Frs2-mediated activation;ARMS-mediated activation;Gastrin-CREB signalling pathway via PKC and MAPK;Interleukin receptor SHC signaling;DAP12 signaling;Signaling by PDGF;VEGFA-VEGFR2 Pathway;Signaling by ERBB2;IRS-related events triggered by IGF1R;Cell surface interactions at the vascular wall,hsa04010;hsa04012;hsa04014;hsa04015;hsa04062;hsa04068;hsa04071;hsa04072;hsa04151;hsa04211;hsa04213;hsa04360;hsa04370;hsa04530;hsa04540;hsa04550;hsa04650;hsa04660;hsa04662;hsa04664;hsa04720;hsa04722;hsa04725;hsa04726;hsa04730;hsa04810;hsa04910;hsa04912;hsa04915;hsa04916;hsa04917;hsa04919;hsa04921;hsa04933;hsa05034;hsa05160;hsa05161;hsa05166;hsa05200;hsa05203;hsa05205;hsa05206;hsa05211;hsa05213;hsa05214;hsa05215;hsa05216;hsa05218;hsa05219;hsa05220;hsa05221;hsa05223;hsa05230;hsa05231,NRAS,NA,NRAS - mRNA,False\nPAK2 - mRNA,NM_002577.3:3785,SEBELIUS_2_C4636,PAK2,mRNA,0,,,Nef and signal transduction;CD28 dependent Vav1 pathway;Stimulation of the cell death response by PAK-2p34;Ephrin signaling;Activation of Rac;RHO GTPases activate PAKs;VEGFR2 mediated vascular permeability;Sema3A PAK dependent Axon repulsion;CD209 (DC-SIGN) signaling;VEGFA-VEGFR2 Pathway;CD28 co-stimulation,hsa04010;hsa04012;hsa04014;hsa04360;hsa04510;hsa04660;hsa04810;hsa05211,PAK2,NA,PAK2 - mRNA,False\nMLLT4 - mRNA,NM_005936.2:1560,SEBELIUS_2_C4636,MLLT4,mRNA,0,,,Adherens junctions interactions,hsa04014;hsa04015;hsa04024;hsa04520;hsa04530;hsa04670,MLLT4,NA,MLLT4 - mRNA,False\nMYC - mRNA,NM_002467.3:1610,SEBELIUS_2_C4636,MYC,mRNA,0,,,Constitutive Signaling by NOTCH1 PEST Domain Mutants;SMAD2SMAD3.CL.SMAD4 heterotrimer regulates transcription;Formation of the beta-catenin.CL.TCF transactivating complex,hsa04010;hsa04012;hsa04110;hsa04151;hsa04310;hsa04350;hsa04390;hsa04550;hsa04630;hsa04919;hsa05161;hsa05166;hsa05169;hsa05200;hsa05202;hsa05205;hsa05206;hsa05210;hsa05213;hsa05216;hsa05219;hsa05220;hsa05221;hsa05222;hsa05230,MYC,NA,MYC - mRNA,False\nPPP1CC - mRNA,NM_002710.2:267,SEBELIUS_2_C4636,PPP1CC,mRNA,0,,,Separation of Sister Chromatids;RHO GTPases Activate Formins;Resolution of Sister Chromatid Cohesion;Downregulation of TGF-beta receptor signaling;Circadian Clock;TGF-beta receptor signaling activates SMADs,hsa03015;hsa04022;hsa04024;hsa04114;hsa04261;hsa04270;hsa04390;hsa04510;hsa04611;hsa04720;hsa04728;hsa04750;hsa04810;hsa04910;hsa04921;hsa04931;hsa05031;hsa05034;hsa05168;hsa05205,PPP1CC,NA,PPP1CC - mRNA,False\nSRC - mRNA,NM_005417.3:1410,SEBELIUS_2_C4636,SRC,mRNA,0,,,FCGR activation;Spry regulation of FGF signaling;GAB1 signalosome;MAP2K and MAPK activation;CLEC7A (Dectin-1) signaling;CD28 co-stimulation;p130Cas linkage to MAPK signaling for integrins;CTLA4 inhibitory signaling;GRB2.CL.SOS provides linkage to MAPK signaling for Integrins ;Signal regulatory protein (SIRP) family interactions;PECAM1 interactions;Thrombin signalling through proteinase activated receptors (PARs);GP1b-IX-V activation signalling;c-src mediated regulation of Cx43 function and closure of gap junctions;Signaling by ERBB2;VEGFA-VEGFR2 Pathway;Signaling by SCF-KIT;VEGFR2 mediated cell proliferation;RAFMAP kinase cascade;Downstream signal transduction;Frs2-mediated activation;ARMS-mediated activation;Gastrin-CREB signalling pathway via PKC and MAPK;Interleukin receptor SHC signaling;FCERI mediated MAPK activation;DAP12 signaling;Signaling by PDGF;IRS-related events triggered by IGF1R;Negative regulation of FGFR3 signaling;Negative regulation of FGFR4 signaling;Negative regulation of FGFR2 signaling;Negative regulation of FGFR1 signaling;Integrin alphaIIb beta3 signaling;Cell surface interactions at the vascular wall,hsa04012;hsa04015;hsa04062;hsa04144;hsa04370;hsa04510;hsa04520;hsa04530;hsa04540;hsa04611;hsa04727;hsa04750;hsa04810;hsa04912;hsa04915;hsa04917;hsa04919;hsa04921;hsa05100;hsa05120;hsa05131;hsa05152;hsa05161;hsa05203;hsa05205;hsa05219,SRC,NA,SRC - mRNA,False\nTCF7L2 - mRNA,NM_001146274.1:1393,SEBELIUS_2_C4636,TCF7L2,mRNA,0,,,Ca2.PLS. pathway;Deactivation of the beta-catenin transactivating complex;Repression of WNT target genes;Synthesis# secretion# and inactivation of Glucagon-like Peptide-1 (GLP-1);Formation of the beta-catenin.CL.TCF transactivating complex,hsa04310;hsa04390;hsa04520;hsa04916;hsa05200;hsa05210;hsa05213;hsa05215;hsa05216;hsa05217;hsa05221;hsa05412,TCF7L2,NA,TCF7L2 - mRNA,False\nAXIN2 - mRNA,NM_004655.3:1035,SEBELIUS_2_C4636,AXIN2,mRNA,0,,,Binding of TCFLEF.CL.CTNNB1 to target gene promoters;Formation of the beta-catenin.CL.TCF transactivating complex,hsa04310;hsa04390;hsa04550;hsa05200;hsa05210;hsa05213;hsa05217,AXIN2,NA,AXIN2 - mRNA,False\nMET - mRNA,NM_001127500.1:1925,SEBELIUS_2_C4636,MET,mRNA,0,,,,hsa04014;hsa04015;hsa04060;hsa04144;hsa04151;hsa04360;hsa04510;hsa04520;hsa05100;hsa05120;hsa05200;hsa05202;hsa05205;hsa05206;hsa05211;hsa05218;hsa05230,MET,NA,MET - mRNA,False\nINADL - mRNA,XM_005270347.1:343,SEBELIUS_2_C4636,INADL,mRNA,0,,,,hsa04390;hsa04530,INADL,NA,INADL - mRNA,False\nBAIAP2L1 - mRNA,NM_018842.3:259,SEBELIUS_2_C4636,BAIAP2L1,mRNA,0,,,,,BAIAP2L1,NA,BAIAP2L1 - mRNA,False\nITGA3 - mRNA,NM_002204.2:4550,SEBELIUS_2_C4636,ITGA3,mRNA,0,,,Laminin interactions;Integrin cell surface interactions;Cell surface interactions at the vascular wall,hsa04151;hsa04510;hsa04512;hsa04640;hsa04810;hsa05200;hsa05222;hsa05410;hsa05412;hsa05414,ITGA3,NA,ITGA3 - mRNA,False\nPIK3R3 - mRNA,NM_003629.3:1800,SEBELIUS_2_C4636,PIK3R3,mRNA,0,,,G alpha (q) signalling events;PIP3 activates AKT signaling;Constitutive Signaling by Aberrant PI3K in Cancer;CD28 dependent PI3KAkt signaling;Synthesis of PIPs at the plasma membrane;Interleukin receptor SHC signaling;G beta.CL.gamma signalling through PI3Kgamma;Regulation of signaling by CBL;GPVI-mediated activation cascade;G alpha (1213) signalling events;Gastrin-CREB signalling pathway via PKC and MAPK;Downstream signal transduction;GAB1 signalosome;Signaling by SCF-KIT;PI3KAKT activation;Role of LAT2NTALLAB on calcium mobilization;DAP12 signaling;Signaling by PDGF;Signaling by ERBB2;CD28 co-stimulation,hsa04012;hsa04014;hsa04015;hsa04022;hsa04024;hsa04062;hsa04066;hsa04068;hsa04070;hsa04071;hsa04072;hsa04150;hsa04151;hsa04152;hsa04210;hsa04211;hsa04213;hsa04261;hsa04370;hsa04380;hsa04510;hsa04550;hsa04611;hsa04620;hsa04630;hsa04650;hsa04660;hsa04662;hsa04664;hsa04666;hsa04668;hsa04670;hsa04722;hsa04725;hsa04750;hsa04810;hsa04910;hsa04914;hsa04915;hsa04917;hsa04919;hsa04921;hsa04923;hsa04930;hsa04931;hsa04932;hsa04933;hsa04960;hsa04973;hsa05100;hsa05142;hsa05145;hsa05146;hsa05160;hsa05161;hsa05162;hsa05164;hsa05166;hsa05169;hsa05200;hsa05203;hsa05205;hsa05210;hsa05211;hsa05212;hsa05213;hsa05214;hsa05215;hsa05218;hsa05220;hsa05221;hsa05222;hsa05223;hsa05230;hsa05231,PIK3R3,NA,PIK3R3 - mRNA,False\nRRAS - mRNA,NM_006270.3:330,SEBELIUS_2_C4636,RRAS,mRNA,0,,,CREB phosphorylation through the activation of Ras,hsa04010;hsa04014;hsa04015;hsa04024;hsa04072;hsa04530;hsa04810;hsa05166;hsa05205,RRAS,NA,RRAS - mRNA,False\nPDGFA - mRNA,NM_002607.5:2460,SEBELIUS_2_C4636,PDGFA,mRNA,0,,,Non-integrin membrane-ECM interactions;PIP3 activates AKT signaling;Constitutive Signaling by Aberrant PI3K in Cancer;RAFMAP kinase cascade;Platelet degranulation ;Signaling by PDGF;Downstream signal transduction;GAB1 signalosome;Signaling by SCF-KIT;PI3KAKT activation;Role of LAT2NTALLAB on calcium mobilization;DAP12 signaling;Signaling by ERBB2;VEGFR2 mediated cell proliferation;Frs2-mediated activation;ARMS-mediated activation;Gastrin-CREB signalling pathway via PKC and MAPK;Interleukin receptor SHC signaling;FCERI mediated MAPK activation;VEGFA-VEGFR2 Pathway;IRS-related events triggered by IGF1R,hsa04010;hsa04014;hsa04015;hsa04060;hsa04072;hsa04151;hsa04510;hsa04540;hsa04810;hsa05166;hsa05200;hsa05202;hsa05206;hsa05214;hsa05215;hsa05218;hsa05231,PDGFA,NA,PDGFA - mRNA,False\nIL1R2 - mRNA,NM_001261419.1:186,SEBELIUS_2_C4636,IL1R2,mRNA,0,,,,hsa04010;hsa04060;hsa04640;hsa05146;hsa05166;hsa05202,IL1R2,NA,IL1R2 - mRNA,False\nHSPA1A - mRNA,NM_005345.5:98,SEBELIUS_2_C4636,HSPA1A,mRNA,0,,,Regulation of HSF1-mediated heat shock response;AUF1 (hnRNP D0) binds and destabilizes mRNA,hsa04010;hsa04141;hsa04144;hsa04213;hsa04915;hsa05020;hsa05145;hsa05162;hsa05164;hsa05169,HSPA1A,NA,HSPA1A - mRNA,False\nAXIN1 - mRNA,NM_181050.1:2742,SEBELIUS_2_C4636,AXIN1,mRNA,0,,,APC truncation mutants have impaired AXIN binding;Misspliced GSK3beta mutants stabilize beta-catenin;T41 mutants of beta-catenin aren.SQT.t phosphorylated;AXIN missense mutants destabilize the destruction complex;S45 mutants of beta-catenin aren.SQT.t phosphorylated;Disassembly of the destruction complex and recruitment of AXIN to the membrane;S37 mutants of beta-catenin aren.SQT.t phosphorylated;S33 mutants of beta-catenin aren.SQT.t phosphorylated;Beta-catenin phosphorylation cascade;Truncations of AMER1 destabilize the destruction complex,hsa04310;hsa04390;hsa04550;hsa05200;hsa05210;hsa05213;hsa05217,AXIN1,NA,AXIN1 - mRNA,False\nMYL5 - mRNA,NM_002477.1:398,SEBELIUS_2_C4636,MYL5,mRNA,0,,,Smooth Muscle Contraction,hsa04510;hsa04530;hsa04670;hsa04810,MYL5,NA,MYL5 - mRNA,False\nBAD - mRNA,NM_004322.3:963,SEBELIUS_2_C4636,BAD,mRNA,0,,,Activation of BAD and translocation to mitochondria ;NRAGE signals death through JNK;Constitutive Signaling by AKT1 E17K in Cancer;AKT phosphorylates targets in the cytosol;PIP3 activates AKT signaling;Downstream signal transduction;GAB1 signalosome;Signaling by SCF-KIT;PI3KAKT activation;Role of LAT2NTALLAB on calcium mobilization;DAP12 signaling;Signaling by PDGF;Signaling by ERBB2,hsa04012;hsa04014;hsa04022;hsa04024;hsa04151;hsa04210;hsa04370;hsa04510;hsa04722;hsa04910;hsa04919;hsa05010;hsa05145;hsa05152;hsa05160;hsa05161;hsa05200;hsa05203;hsa05210;hsa05212;hsa05213;hsa05215;hsa05218;hsa05220;hsa05221;hsa05223,BAD,NA,BAD - mRNA,False\nMYL12A c - mRNA,NM_001303049.1:9,SEBELIUS_2_C4636,MYL12A c,mRNA,0,,NM_006471.2:78;NM_001303048.1:27,EPHA-mediated growth cone collapse;RHO GTPases Activate ROCKs;Smooth Muscle Contraction;RHO GTPases activate CIT;RHO GTPases activate PAKs;RHO GTPases activate PKNs;Sema4D induced cell migration and growth-cone collapse;Ephrin signaling,hsa04510;hsa04530;hsa04611;hsa04670;hsa04810,MYL12A,NA,MYL12A c - mRNA,False\nCLDN15 - mRNA,NM_014343.1:1129,SEBELIUS_2_C4636,CLDN15,mRNA,0,,,Tight junction interactions,hsa04514;hsa04530;hsa04670;hsa05160,CLDN15,NA,CLDN15 - mRNA,False\nHCLS1 - mRNA,NM_005335.4:1846,SEBELIUS_2_C4636,HCLS1,mRNA,0,,,,hsa04530;hsa05100;hsa05130;hsa05131;hsa05205,HCLS1,NA,HCLS1 - mRNA,False\nCCND3 - mRNA,NM_001760.2:1215,SEBELIUS_2_C4636,CCND3,mRNA,0,,,Cyclin D associated events in G1;MAPK6MAPK4 signaling;Transcriptional regulation of white adipocyte differentiation,hsa04110;hsa04115;hsa04151;hsa04310;hsa04390;hsa04510;hsa04630;hsa05162;hsa05166;hsa05203,CCND3,NA,CCND3 - mRNA,False\nHRAS - mRNA,NM_005343.2:396,SEBELIUS_2_C4636,HRAS,mRNA,0,,,Signaling by FGFR2 in disease;Regulation of RAS by GAPs;SHC-mediated cascade.CL.FGFR3;Signaling by FGFR3 in disease;MAP2K and MAPK activation;SHC-related events triggered by IGF1R;Signaling by FGFR4 in disease;SHC-mediated cascade.CL.FGFR1;Tie2 Signaling;Signaling by FGFR1 in disease;Activation of RAS in B cells;SHC-mediated cascade.CL.FGFR4;Constitutive Signaling by EGFRvIII;CD209 (DC-SIGN) signaling;Ras activation uopn Ca2.PLS. infux through NMDA receptor;FCERI mediated MAPK activation;RAFMAP kinase cascade;Downstream signal transduction;VEGFR2 mediated cell proliferation;Signaling by SCF-KIT;Frs2-mediated activation;ARMS-mediated activation;Gastrin-CREB signalling pathway via PKC and MAPK;Interleukin receptor SHC signaling;DAP12 signaling;Signaling by PDGF;VEGFA-VEGFR2 Pathway;Signaling by ERBB2;IRS-related events triggered by IGF1R;Cell surface interactions at the vascular wall;CREB phosphorylation through the activation of Ras,hsa04010;hsa04012;hsa04014;hsa04015;hsa04062;hsa04068;hsa04071;hsa04072;hsa04144;hsa04151;hsa04211;hsa04213;hsa04360;hsa04370;hsa04510;hsa04530;hsa04540;hsa04550;hsa04630;hsa04650;hsa04660;hsa04662;hsa04664;hsa04720;hsa04722;hsa04725;hsa04726;hsa04730;hsa04810;hsa04910;hsa04912;hsa04915;hsa04916;hsa04917;hsa04919;hsa04921;hsa04933;hsa05034;hsa05160;hsa05161;hsa05166;hsa05200;hsa05203;hsa05205;hsa05206;hsa05211;hsa05213;hsa05214;hsa05215;hsa05216;hsa05218;hsa05219;hsa05220;hsa05221;hsa05223;hsa05230;hsa05231,HRAS,NA,HRAS - mRNA,False\nFYN - mRNA,NM_002037.3:765,SEBELIUS_2_C4636,FYN,mRNA,0,,,EPH-ephrin mediated repulsion of cells;Regulation of signaling by CBL;FCGR activation;Antigen activates B Cell Receptor (BCR) leading to generation of second messengers;Dectin-2 family;RAFMAP kinase cascade;Constitutive Signaling by Aberrant PI3K in Cancer;CTLA4 inhibitory signaling;CD28 dependent PI3KAkt signaling;PECAM1 interactions;CD28 dependent Vav1 pathway;GPVI-mediated activation cascade;Sema3A PAK dependent Axon repulsion;Nephrin interactions;SEMA3A-Plexin repulsion signaling by inhibiting Integrin adhesion;CRMPs in Sema3A signaling;CD209 (DC-SIGN) signaling;Downstream signal transduction;GAB1 signalosome;Signaling by SCF-KIT;PI3KAKT activation;Role of LAT2NTALLAB on calcium mobilization;DAP12 signaling;Signaling by PDGF;Signaling by ERBB2;VEGFA-VEGFR2 Pathway;VEGFR2 mediated cell proliferation;Frs2-mediated activation;ARMS-mediated activation;Gastrin-CREB signalling pathway via PKC and MAPK;Interleukin receptor SHC signaling;FCERI mediated MAPK activation;IRS-related events triggered by IGF1R;CD28 co-stimulation;Cell surface interactions at the vascular wall,hsa04071;hsa04072;hsa04360;hsa04380;hsa04510;hsa04520;hsa04611;hsa04650;hsa04660;hsa04664;hsa04725;hsa05020;hsa05130;hsa05162;hsa05416,FYN,NA,FYN - mRNA,False\nTJP2 - mRNA,NM_004817.2:2785,SEBELIUS_2_C4636,TJP2,mRNA,0,,,Apoptotic cleavage of cellular proteins,hsa04530;hsa05110,TJP2,NA,TJP2 - mRNA,False\nJUN - mRNA,NM_002228.3:140,SEBELIUS_2_C4636,JUN,mRNA,0,,,Activation of the AP-1 family of transcription factors;MAPK6MAPK4 signaling;Pre-NOTCH Transcription and Translation;Oxidative Stress Induced Senescence;FCERI mediated MAPK activation;Senescence-Associated Secretory Phenotype (SASP),hsa04010;hsa04012;hsa04024;hsa04310;hsa04380;hsa04510;hsa04620;hsa04660;hsa04662;hsa04668;hsa04722;hsa04912;hsa04915;hsa04921;hsa04932;hsa04933;hsa05031;hsa05120;hsa05132;hsa05133;hsa05140;hsa05142;hsa05161;hsa05164;hsa05166;hsa05168;hsa05169;hsa05200;hsa05203;hsa05210;hsa05211;hsa05231;hsa05321,JUN,NA,JUN - mRNA,False\nSOS2 - mRNA,NM_006939.2:3845,SEBELIUS_2_C4636,SOS2,mRNA,0,,,Rho GTPase cycle;Activation of Rac;NRAGE signals death through JNK;G alpha (1213) signalling events,hsa04010;hsa04012;hsa04014;hsa04062;hsa04068;hsa04072;hsa04151;hsa04320;hsa04510;hsa04540;hsa04630;hsa04650;hsa04660;hsa04662;hsa04664;hsa04722;hsa04810;hsa04910;hsa04912;hsa04915;hsa04917;hsa05034;hsa05160;hsa05200;hsa05205;hsa05206;hsa05211;hsa05213;hsa05214;hsa05215;hsa05220;hsa05221;hsa05223;hsa05231,SOS2,NA,SOS2 - mRNA,False\nWASF1 - mRNA,NM_001024934.1:1425,SEBELIUS_2_C4636,WASF1,mRNA,0,,,RHO GTPases Activate WASPs and WAVEs;VEGFA-VEGFR2 Pathway;Regulation of actin dynamics for phagocytic cup formation,hsa04520;hsa04666;hsa04810;hsa05100;hsa05131;hsa05132;hsa05231,WASF1,NA,WASF1 - mRNA,False\nDVL1 - mRNA,NM_004421.2:789,SEBELIUS_2_C4636,DVL1,mRNA,0,,,Disassembly of the destruction complex and recruitment of AXIN to the membrane;WNT mediated activation of DVL;RHO GTPases Activate Formins;PCPCE pathway,hsa04310;hsa04330;hsa04390;hsa04550;hsa04916;hsa05166;hsa05200;hsa05217,DVL1,NA,DVL1 - mRNA,False\nBCAR1 - mRNA,NM_001170717.1:1341,SEBELIUS_2_C4636,BCAR1,mRNA,0,,,Downstream signal transduction;p130Cas linkage to MAPK signaling for integrins;VEGFA-VEGFR2 Pathway;Signaling by PDGF;Integrin alphaIIb beta3 signaling,hsa04015;hsa04062;hsa04510;hsa04670;hsa04810;hsa05100,BCAR1,NA,BCAR1 - mRNA,False\nAPC - mRNA,NM_000038.3:6850,SEBELIUS_2_C4636,APC,mRNA,0,,,Apoptotic cleavage of cellular proteins;Misspliced GSK3beta mutants stabilize beta-catenin;APC truncation mutants have impaired AXIN binding;T41 mutants of beta-catenin aren.SQT.t phosphorylated;AXIN missense mutants destabilize the destruction complex;S45 mutants of beta-catenin aren.SQT.t phosphorylated;Deactivation of the beta-catenin transactivating complex;Disassembly of the destruction complex and recruitment of AXIN to the membrane;S37 mutants of beta-catenin aren.SQT.t phosphorylated;S33 mutants of beta-catenin aren.SQT.t phosphorylated;Beta-catenin phosphorylation cascade;Truncations of AMER1 destabilize the destruction complex,hsa04310;hsa04390;hsa04550;hsa04810;hsa05166;hsa05200;hsa05206;hsa05210;hsa05213;hsa05217,APC,NA,APC - mRNA,False\nCRB3 - mRNA,NM_139161.3:300,SEBELIUS_2_C4636,CRB3,mRNA,0,,,Tight junction interactions,hsa04530,CRB3,NA,CRB3 - mRNA,False\nTJP3 - mRNA,NM_014428.1:1925,SEBELIUS_2_C4636,TJP3,mRNA,0,,,,hsa04530,TJP3,NA,TJP3 - mRNA,False\nOCLN - mRNA,NM_002538.3:5130,SEBELIUS_2_C4636,OCLN,mRNA,0,,,Apoptotic cleavage of cellular proteins,hsa04514;hsa04530;hsa04670;hsa05130;hsa05160,OCLN,NA,OCLN - mRNA,False\nNLK - mRNA,NM_016231.4:618,SEBELIUS_2_C4636,NLK,mRNA,0,,,Ca2.PLS. pathway,hsa04010;hsa04068;hsa04310;hsa04520,NLK,NA,NLK - mRNA,False\nRAC3 - mRNA,NM_005052.2:702,SEBELIUS_2_C4636,RAC3,mRNA,0,,,Rho GTPase cycle;PCPCE pathway,hsa04010;hsa04014;hsa04015;hsa04024;hsa04071;hsa04310;hsa04360;hsa04370;hsa04510;hsa04520;hsa04650;hsa04662;hsa04664;hsa04810;hsa05200;hsa05210;hsa05212;hsa05231;hsa05416,RAC3,NA,RAC3 - mRNA,False\nLLGL2 - mRNA,NM_001015002.1:1065,SEBELIUS_2_C4636,LLGL2,mRNA,0,,,,hsa04390;hsa04530,LLGL2,NA,LLGL2 - mRNA,False\nTCF7 - mRNA,NM_201633.2:519,SEBELIUS_2_C4636,TCF7,mRNA,0,,,,,,NA,TCF7 - mRNA,False\nDIAPH3 - mRNA,NM_001042517.1:1832,SEBELIUS_2_C4636,DIAPH3,mRNA,0,,,RHO GTPases Activate Formins,hsa04810,DIAPH3,NA,DIAPH3 - mRNA,False\nBAIAP2 - mRNA,NM_017450.2:653,SEBELIUS_2_C4636,BAIAP2,mRNA,0,,,RHO GTPases Activate WASPs and WAVEs;VEGFA-VEGFR2 Pathway;Regulation of actin dynamics for phagocytic cup formation,hsa04520;hsa04810,BAIAP2,NA,BAIAP2 - mRNA,False\nPIK3CB - mRNA,NM_006219.1:2945,SEBELIUS_2_C4636,PIK3CB,mRNA,0,,,Downstream TCR signaling;PIP3 activates AKT signaling;Constitutive Signaling by Aberrant PI3K in Cancer;Tie2 Signaling;VEGFA-VEGFR2 Pathway;Synthesis of PIPs at the plasma membrane;Interleukin receptor SHC signaling;G beta.CL.gamma signalling through PI3Kgamma;Regulation of signaling by CBL;GPVI-mediated activation cascade;Role of phospholipids in phagocytosis;Nephrin interactions;PI3K Cascade;Downstream signal transduction;GAB1 signalosome;Signaling by SCF-KIT;PI3KAKT activation;Role of LAT2NTALLAB on calcium mobilization;DAP12 signaling;Signaling by PDGF;Signaling by ERBB2;Cell surface interactions at the vascular wall;IRS-related events triggered by IGF1R,hsa00562;hsa04012;hsa04014;hsa04015;hsa04022;hsa04024;hsa04062;hsa04066;hsa04068;hsa04070;hsa04071;hsa04072;hsa04150;hsa04151;hsa04152;hsa04210;hsa04211;hsa04213;hsa04261;hsa04370;hsa04380;hsa04510;hsa04550;hsa04611;hsa04620;hsa04630;hsa04650;hsa04660;hsa04662;hsa04664;hsa04666;hsa04668;hsa04670;hsa04722;hsa04725;hsa04750;hsa04810;hsa04910;hsa04914;hsa04915;hsa04917;hsa04919;hsa04921;hsa04923;hsa04930;hsa04931;hsa04932;hsa04933;hsa04960;hsa04973;hsa05100;hsa05142;hsa05145;hsa05146;hsa05160;hsa05161;hsa05162;hsa05164;hsa05166;hsa05169;hsa05200;hsa05203;hsa05205;hsa05210;hsa05211;hsa05212;hsa05213;hsa05214;hsa05215;hsa05218;hsa05220;hsa05221;hsa05222;hsa05223;hsa05230;hsa05231,PIK3CB,NA,PIK3CB - mRNA,False\nMAGI3 - mRNA,NM_152900.2:6685,SEBELIUS_2_C4636,MAGI3,mRNA,0,,,,hsa04015;hsa04530,MAGI3,NA,MAGI3 - mRNA,False\nCDX2 - mRNA,NM_001265.2:1812,SEBELIUS_2_C4636,CDX2,mRNA,0,,,,,CDX2,NA,CDX2 - mRNA,False\nTGFB2 - mRNA,NM_003238.2:1125,SEBELIUS_2_C4636,TGFB2,mRNA,0,,,Molecules associated with elastic fibres;ECM proteoglycans;Platelet degranulation ;Elastic fibre formation,hsa04010;hsa04060;hsa04068;hsa04110;hsa04144;hsa04350;hsa04380;hsa04390;hsa04933;hsa05140;hsa05142;hsa05145;hsa05146;hsa05152;hsa05161;hsa05166;hsa05200;hsa05205;hsa05206;hsa05210;hsa05211;hsa05212;hsa05220;hsa05321;hsa05410;hsa05414,TGFB2,NA,TGFB2 - mRNA,False\nCLDN23 - mRNA,NM_194284.2:1768,SEBELIUS_2_C4636,CLDN23,mRNA,0,,,Tight junction interactions,hsa04514;hsa04530;hsa04670;hsa05160,CLDN23,NA,CLDN23 - mRNA,False\nCLDN7 - mRNA,NM_001307.3:175,SEBELIUS_2_C4636,CLDN7,mRNA,0,,,Tight junction interactions,hsa04514;hsa04530;hsa04670;hsa05160,CLDN7,NA,CLDN7 - mRNA,False\nCGN - mRNA,NM_020770.2:1112,SEBELIUS_2_C4636,CGN,mRNA,0,,,TGF-beta receptor signaling in EMT (epithelial to mesenchymal transition),hsa04530,CGN,NA,CGN - mRNA,False\nCLDN3 - mRNA,NM_001306.3:606,SEBELIUS_2_C4636,CLDN3,mRNA,0,,,Tight junction interactions,hsa04514;hsa04530;hsa04670;hsa05160,CLDN3,NA,CLDN3 - mRNA,False\nITGB4 - mRNA,NM_001005731.1:4151,SEBELIUS_2_C4636,ITGB4,mRNA,0,,,Syndecan interactions;Assembly of collagen fibrils and other multimeric structures;Non-integrin membrane-ECM interactions,hsa04151;hsa04510;hsa04512;hsa04810;hsa05410;hsa05412;hsa05414,ITGB4,NA,ITGB4 - mRNA,False\nCA2 - mRNA,NM_000067.2:575,SEBELIUS_2_C4636,CA2,mRNA,0,,,,hsa04971;hsa04972,CA2,NA,CA2 - mRNA,False\nMYH14 - mRNA,NM_001077186.1:6304,SEBELIUS_2_C4636,MYH14,mRNA,0,,,EPHA-mediated growth cone collapse;RHO GTPases Activate ROCKs;RHO GTPases activate CIT;RHO GTPases activate PAKs;RHO GTPases activate PKNs;Sema4D induced cell migration and growth-cone collapse,hsa04530;hsa04810;hsa05132,MYH14,NA,MYH14 - mRNA,False\nCXCR4 - mRNA,NM_003467.2:1335,SEBELIUS_2_C4636,CXCR4,mRNA,0,,,,hsa04060;hsa04062;hsa04144;hsa04360;hsa04670;hsa05200,CXCR4,NA,CXCR4 - mRNA,False\n";
	heatmapData_UyySR3RU_9hQ_hWdumMiueg = "Dimname,20180928_Polarization-02_FibD7A_05.RCC,20180928_Polarization-02_FibD7B_06.RCC,20180926_Polarization01_FibD4A_05.RCC,20180926_Polarization01_FibD4B_06.RCC,20180926_Polarization01_HT29D4D_10.RCC,20180926_Polarization01_HT29D4C_09.RCC,20180926_Polarization01_HT29D4A_07.RCC,20180926_Polarization01_HT29D4B_08.RCC,20180926_Polarization01_HT29D4E_11.RCC,20180926_Polarization01_HT29D4F_12.RCC,20180928_Polarization-02_HT29D7E_11.RCC,20180928_Polarization-02_HT29D7F_12.RCC,20180928_Polarization-02_HT29D7B_08.RCC,20180928_Polarization-02_HT29D7A_07.RCC,20180928_Polarization-02_HT29D7C_09.RCC,20180928_Polarization-02_HT29D7D_10.RCC,20181004_Pol03_HepG2D4A_01.RCC,20181004_Pol03_HepG2D4B_02.RCC,20181004_Pol03_HepG2D7A_05.RCC,20181004_Pol03_HepG2D7B_06.RCC,20181004_Pol03_HepG2D7C_07.RCC,20181004_Pol03_HepG2D7D_08.RCC,20181004_Pol03_HepG2D4C_03.RCC,20181004_Pol03_HepG2D4D_04.RCC,20161013_sebelius2-101316_01_12.RCC,20161013_sebelius2-101316_01_11.RCC,20170307_20170307-caco_01_05.RCC,20170307_20170307-caco_01_06.RCC,20161013_sebelius2-101316_01_09.RCC,20170307_20170307-caco_01_01.RCC,20170307_20170307-caco_01_02.RCC,20180928_Polarization-02_Caco2D7C_03.RCC,20180928_Polarization-02_Caco2D7A_01.RCC,20180928_Polarization-02_Caco2D7B_02.RCC,20180928_Polarization-02_Caco2D7D_04.RCC,20180926_Polarization01_Caco2D4D_04.RCC,20180926_Polarization01_Caco2D4C_03.RCC,20180926_Polarization01_Caco2D4A_01.RCC,20180926_Polarization01_Caco2D4B_02.RCC,20170307_20170307-caco_01_03.RCC,20161013_sebelius2-101316_01_10.RCC,20170307_20170307-caco_01_04.RCC\nFN1 - mRNA,15.5161289134903,15.4728451717661,15.4897197624664,15.503573346043,5.38168264061771,5.77442531273256,5.63499095957823,5.36491397069513,5.44108271650874,5.41673174364114,5.18955844065191,5.30673500297172,7.98005456923237,5.3680020298439,5.0541144370371,5.19941341587416,16.0624363525319,16.0380430825591,16.6380353626414,16.6182016510502,16.9501479240332,16.8965211191785,17.0103617706689,16.9811028239372,14.7040750782607,15.1347951689678,14.974018342194,14.7010703333264,15.3362922019809,15.5036087750865,15.4109707232919,15.686117512564,15.6664929698428,15.6111810407228,15.4955879909289,15.4536085424866,15.5358590683858,15.5501262467899,15.4539986234173,16.1369112942441,16.1667820230365,15.9824820184315\nLYZ - mRNA,3.98399566498464,3.84300477438578,4.270821995682,4.40565409557399,16.7390720169321,16.7314201443739,16.7310254711515,16.699998364995,16.7306279326908,16.6448304889125,16.9651877132787,16.9268495419367,16.8765454293672,16.8641338068597,16.8895158801902,16.8510467668747,16.9041944419945,16.8580437282162,17.1781107538013,17.2273567623323,16.939156645323,16.973147740469,16.6778191471186,16.6191498608464,9.35870807040847,9.94612560865168,9.93860642203693,10.3675203187975,9.89985222596986,10.1649244518735,10.8323497494446,10.7662223261052,10.8589220003247,10.8381600305795,10.7856770140841,11.5239798656102,11.7738720097539,11.554761535668,11.5152777995966,11.6212481242375,12.1365465261988,11.4170945686844\nLDHA - mRNA,14.037262188347,14.0213583978987,13.9515979014318,14.040361444262,15.2238780328574,15.21637916032,15.2362107388185,15.1724673908286,13.8167172945333,14.2920365111339,16.8130390091283,16.7193763662689,16.2572150956664,16.2524200944301,16.2475562683626,16.286254557656,15.4562694371603,15.5491425951181,16.4117178305221,16.3183805401743,16.6085880855204,16.5870417145966,15.6006373964739,15.526268325966,15.666890157927,15.7344094803331,15.7356819232231,15.5798378804057,13.1641928293043,13.9032377295015,13.8836302543883,15.6933170350989,15.5879432160079,15.6127133109071,15.3115325438582,13.8728871002136,14.4547818499071,14.4522387022957,14.358111370475,15.3045528789061,15.4857787165937,15.1929172961481\nRPL19 - mRNA,14.9056028622793,14.9527334245608,15.0654302427468,15.0465961917534,16.2944560405851,16.3155549362853,16.2637669578384,16.2219789351262,16.0640496612992,16.1514525039821,16.1361161144552,16.0883605448989,16.038377815079,16.05008319628,16.1130004846772,16.0842397732189,16.4026684432026,16.3620494032116,16.2058084574406,16.2326454748035,16.6386052170082,16.6552553193184,16.4846343197295,16.5760180656798,16.5230948883094,16.1058235883396,16.5333132415045,16.4688008452962,14.5411971965885,15.3087725265218,15.3755851245561,16.0464864696754,16.0265814983383,15.9534357106725,15.9606731396001,15.7647838953728,15.9087747570095,15.8484223250211,15.857891472599,15.8497983813824,15.8687194553175,16.0343358402339\nRPLP0 - mRNA,15.239664990451,15.1783602337505,15.2456139283717,15.2342439723051,16.3512718374832,16.3988988694054,16.3284396223866,16.264157071116,16.5539417008965,16.6013787526627,16.3740809799411,16.2487937399695,16.2178101529783,16.2507896388618,16.2237953177652,16.2376251123962,15.986018407218,15.9312306968395,15.8875411841118,15.8901400794185,16.2974603257174,16.284490069229,16.098968640125,16.0723876171311,16.1749193434266,16.042165993929,16.3765464619415,16.2544720673527,14.3420745545749,15.2922800202474,15.3069858420638,16.2098218760707,16.0708835073159,16.0760254067903,15.9830494753573,15.7202932714462,15.7346820924543,15.7780231600919,15.7015901127959,15.998791604043,15.7583079178089,15.7882604362793\nCCND1 - mRNA,13.9843579034155,14.0027578784698,13.9417111441537,13.9365898671646,11.699541071196,11.6495463278477,11.7481034152649,11.7387623716426,12.0472977664585,12.0611573059317,11.025171622751,11.0891577363462,11.1247695800157,11.1058203166753,11.018137069575,11.0366471162132,13.2625312852831,13.3577482110944,13.3503131388365,13.2914207807409,13.3702040159331,13.3981867642245,13.3311881510767,13.2472016378726,12.1517787664598,12.107763998909,11.4704575862985,11.3407534092347,12.5158416223116,13.0074804057299,12.837763245631,12.3451628352048,12.3620083598538,12.4365737993219,12.3143458075615,12.6343957688788,12.5103461510327,12.6108008201727,12.5514826067521,12.1491796799223,12.1407011584174,12.3196978058899\nCTNNB1 - mRNA,12.677681738493,12.6410714947488,12.5847790508107,12.6550723465932,12.9465030329867,12.9026540277047,12.9099898181129,12.9099048596618,12.7746975970162,12.807240144188,12.89819138199,12.8201958727793,12.929891026559,12.8751538055503,12.9403793601537,12.8821885949412,14.5763982858389,14.5460910829521,14.3590451815979,14.3957913715934,14.4429835045735,14.4887522631805,14.1838310287276,14.1934433225299,11.7295457657768,11.8658490193078,12.1782768348052,12.2400594096152,11.961252770634,12.5642872362454,12.4964561348855,12.6634944406171,12.8008287955781,12.7771926312387,12.8096312296792,13.2103461365661,13.225898163055,13.0999217434636,13.1203845793609,12.6576901202056,12.4528468214041,12.6016903694275\nITGB1 - mRNA,14.5221014341997,14.4856488166687,14.5508906575701,14.5333536957729,12.7624637846893,12.8205571704558,12.7668761797884,12.7468420553187,13.0901666040123,13.0345375010493,13.0381332342728,12.8977771378096,12.8537932456156,12.8912460832868,12.908718730703,12.8682984001404,13.0246884851808,12.9812511736677,12.8150453964959,12.8483392923106,12.6412172532746,12.6024447716093,13.2028691803025,13.2091928930399,12.2354366955067,12.0707266195845,12.1708973044396,12.1002364399624,12.4637531111632,12.9866537376891,12.8132409264969,12.5313277628263,12.4119805589102,12.4170398018597,12.4465883680386,12.7031788505374,12.5720977505557,12.5459151408109,12.4995595658497,12.1791169640764,12.3363250054005,12.2003988775175\nCDC42 - mRNA,12.8761264410478,12.8813236587708,12.9295972146992,12.9565107963922,13.6020780006084,13.6171327854006,13.6293443964371,13.5743673363241,12.9940938750598,13.0241594019501,13.2832096209486,13.2048256693188,13.3552875495997,13.3206252345893,13.3686880807853,13.3521735846496,13.0212590970844,13.0362925372256,12.7719226339979,12.8157137468514,12.6053007841527,12.6570050626832,12.5042854445499,12.5428934293448,12.5175353629675,12.142793216928,12.4972576456422,12.3353399130484,12.4097132714503,12.8808383445449,12.8816660675078,12.494231866663,12.5852435135726,12.5937981247434,12.5977073066524,12.6629317821153,12.6667138006083,12.6266200032392,12.6467286532243,12.4987641177125,12.6973848836205,12.5063190962045\nCSNK1A1 - mRNA,12.6892824552571,12.5959080418612,12.7000635399545,12.6655990736697,12.6929032836834,12.7300752202609,12.7726295945438,12.7165490758704,13.1478743797056,13.0694823889409,12.8118355755923,12.7898971390373,12.7829271051632,12.8599634325466,12.8268903043411,12.7908188835174,12.4381718069294,12.5127847028665,12.3524107630402,12.3353610566373,12.4379936181826,12.4152799696451,12.8006244497575,12.8016716844198,12.1589385580327,12.0396997239639,12.0433472547191,11.9770250037815,11.7161402727975,12.0233703076595,11.9166186663422,12.2191224507853,12.1809924359986,12.1906163775071,12.1333908016963,12.0552035233752,12.190838597687,12.2080236774194,12.1475940416958,12.0579306175563,12.2799950358001,11.9977697110025\nPPP2R1A - mRNA,12.3530632559693,12.3205100917555,12.4623358171312,12.4506726230967,12.961237613082,12.9755467622325,12.9716078663864,12.9620736745336,12.6794874558338,12.717866888768,12.8105959047391,12.8072270175753,12.7657891826249,12.7716474584242,12.7809964418919,12.7589100089928,12.1761657367257,12.1854430054381,12.1124930119117,12.1308341393114,12.2365299095546,12.1817166823025,11.9404128931064,11.9531177692888,12.5721894378375,12.2062651116118,12.3383540502912,12.0744733438973,11.5055732868578,12.0177820693972,11.9668918728234,12.5820757623735,12.599051361695,12.5701006065212,12.5402705464662,12.6451188809894,12.6475911161284,12.6612501465299,12.6319570483051,12.4448079181516,12.5943697362559,12.3333727427869\nACTB - mRNA,15.4035496045575,15.2957929716076,15.5571365620362,15.6010790353072,14.4734346136728,14.4942807987413,14.4614346823275,14.4183005930099,13.7560771452616,13.6924274406879,14.2330695218779,14.1477680300168,14.3990152819297,14.5815660621422,14.4098761270003,14.3536381895114,14.5335148153939,14.3290839260882,14.1835650300043,14.0922996745705,13.9274056535974,13.9152837000069,13.6606936972465,13.5281466245628,13.8801054423522,13.3736580588821,13.3016592503105,13.0728477718482,14.4795780783249,15.3696327061959,15.1613114361865,14.2681796948176,14.1605298584427,13.9714328772494,13.9245601172276,14.392570706892,13.8949136329707,14.1307235782233,13.9879131071467,14.3823354427359,13.7121171273006,13.9273803831111\nACTG1 - mRNA,15.6864033784163,15.6378775030527,15.6918902342916,15.7326773603151,14.8114577116223,14.7782306747346,14.8308836453471,14.8027942756123,14.4313615630916,14.4754711889601,14.6346688155311,14.6961192618569,15.0070615944496,15.0497865078326,14.9478530118836,14.8868984565253,15.3151289769766,15.3462046916267,15.2446947045193,15.0273877307667,15.3470191190689,15.3590287311343,15.8251574270867,15.714724988743,14.460729007349,14.4569135625314,14.4561098073386,14.2701614414047,14.5784140937433,15.1986561943704,15.0338657000228,14.6188500317141,14.4056604020353,14.5080433324495,14.4105688387166,13.9563060734716,13.9213359666259,14.0782915372849,13.945069927794,14.341679317566,14.2265157981597,14.4197677120115\nPPP1CA - mRNA,12.1391030072055,12.1739216525004,12.2584600051749,12.2428765305348,14.1414536354096,14.1304270270211,14.1496074819683,14.1404227838181,13.5313764362098,13.5127532947313,13.4578575046804,13.440397937387,13.8091062751438,13.8402263194246,13.8240923223134,13.826947300347,13.102452001896,13.1122716685821,12.9830926250475,12.9886967380892,12.6895437293057,12.7247029981328,12.7367160142133,12.7456577962044,13.3741711877963,12.7626584601362,13.0733419950169,12.9002773030547,13.0627907970961,13.6600696656881,13.6397046715022,13.3645013394785,13.4261605318907,13.3548189472656,13.422157708033,13.6667203856466,13.713302890713,13.7013895373832,13.6985815763359,13.4960222194385,13.5977736584977,13.5769958421745\nCFL1 - mRNA,14.1712770334366,14.1514504667179,14.3124091044218,14.3446035412177,15.0300782027788,15.0423823971354,15.026930861737,15.0140963121545,13.9153164932138,13.9141109016034,14.493023081772,14.5225959028599,14.6750774228796,14.6587370012733,14.6934704763027,14.6868766521394,13.8138005535991,13.8246972751089,13.4866251169716,13.5112328721094,12.9480646454533,13.0137131101637,12.9857408561384,12.9313554920575,13.5396460645732,12.1663917288797,12.0352422050821,12.1129474917119,12.9910001140281,13.4192246163502,13.3220317467327,13.1240988459043,13.121570829323,13.1217904001582,13.1268001942912,13.3897806515747,13.3889936570217,13.3602779267761,13.3792019340902,12.6964225146943,13.0497336354216,12.907710402971\nMYL12B - mRNA,13.7010355365267,13.6832909651151,13.8650483668733,13.8860692597133,14.9951478879274,15.056329003284,15.0189800225978,14.981689378227,14.5649915078925,14.5623957928215,14.8596971603206,14.8074423291257,14.8418806510634,14.874859063562,14.9163052251748,14.9111072730402,13.0258032733239,13.0548304986348,12.7010360375291,12.7388439209076,12.3321033886786,12.3802724645557,12.3434067027897,12.3967017325993,13.3016269752012,12.7147192279722,12.9494293913315,12.8555967243817,12.9762031121,13.7077032153021,13.6589460215439,12.9486366271099,12.8645160856581,12.8725347774017,12.9743808751414,12.9395214134722,13.0602323502965,13.022786399625,13.0276625659676,12.9009612712179,13.0364163558677,12.9273803831111\nEPB41L3 - mRNA,3.85471264803967,3.79720108477265,3.48232610087571,4.08372600068663,3.31939836236336,3.56292120753885,4.08139262976641,3.49701750670247,2.95169787576981,3.16880423019755,2.75215312834461,3.41920973223013,2.75444543521238,2.15854866421495,3.41939890111884,3.2868762571245,3.63499740081035,2.87468823322278,3.15504950406594,2.79441341077947,1.93274916137754,2.35728824688597,3.52947979216026,3.40603044862402,4.5007270752809,6.40510645551812,3.56356699069001,5.28005747754717,7.69821836480021,5.23081238752999,6.68395790955195,5.55934213899586,4.96125446810813,5.05025747118804,5.34100718676115,5.40388602011985,5.72297014190106,5.57011338858875,5.54912125456841,5.68041019670564,6.69879445387518,6.19041478894489\nALPI - mRNA,3.24703007081843,3.42796727510693,2.83024940429602,4.40565409557399,3.62425294389178,3.83855565015227,3.82763603752062,3.92997691397858,4.15814875323724,3.49073232508491,4.12138693801033,4.05147794772964,3.81333912426595,3.61798028285224,3.34139638911757,3.84326960564888,3.68746482070448,3.33411985186008,3.65254916353675,3.48629111541713,4.12539423931994,3.77232574616481,3.87740309558057,3.72795854351138,7.67065207672321,6.9311752671857,6.56356699069001,6.08741239960477,6.37629026991285,5.42345746547238,6.09899540883079,6.51097911743446,6.23134363147587,6.34148876885249,6.24789778236967,6.14085161428606,6.06339458040581,6.34383753276508,6.36310926895846,7.77994587025655,7.45368195603865,6.34241788238994\nCLDN8 - mRNA,3.55515236618076,3.84300477438578,3.19281948368073,3.92022726840375,4.36121853805799,4.8695825457729,5.00983936874137,4.81894560158983,4.25125815762872,4.16880423019755,4.78190047173867,4.48959906012153,4.41741044793481,4.51610066883303,4.69503334373227,4.59173083865292,4.20203799353424,4.26700565600154,4.55943975914527,4.70868353675358,4.82583395746103,4.67921634177333,4.35545039238522,4.21338537068163,4.5007270752809,4.08317836063075,5.14852949141116,3.28005747754717,6.37629026991285,5.00841996619354,3.77706731394343,4.05154749879716,3.91818574621624,4.01178332337341,4.00397219948358,3.29285470773111,3.76877383151418,3.47389807332945,3.40227986623914,3.87305527464803,1.69879445387518,4.60545228822373\nWAS - mRNA,3.24703007081843,3.4868609641605,2.92978507784693,3.59829917351639,4.08991651624059,4.63692178898262,4.03408691498805,4.12928572220198,4.4991856710723,4.75376673091871,4.78190047173867,4.79772135548386,4.56180035726999,4.51610066883303,4.59773614237735,4.84326960564888,3.40367185470389,3.26700565600154,3.71144285259032,3.18143653388871,4.12539423931994,3.77232574616481,4.02424448390984,3.83898985590013,4.5007270752809,5.568605187801,5.14852949141116,3.28005747754717,5.37629026991285,4.00841996619354,4.36202981466459,5.07391531182562,4.57423134499888,4.86841914820223,4.82213387649777,4.34943823609748,4.12632583613227,4.54665441576476,4.3762846577062,2.87305527464803,5.15822607251248,5.34241788238994\nCA1 - mRNA,3.02463764948198,3.01292977582809,3.02289448223842,3.86133357935018,4.13882611672154,3.95884988386999,4.48474832399761,4.03307040694268,4.38054117457369,4.16880423019755,3.69075258368047,2.90463655940037,2.97683785654883,3.48047675910231,3.97882630973286,4.05739441100173,4.16551211750913,3.7815788288313,3.8749415848732,4.22325670958334,3.21285708057028,3.88780296358475,3.77048789166406,3.72795854351138,5.5007270752809,2.76125026574339,3.56356699069001,4.86501997826832,6.37629026991285,4.00841996619354,2.77706731394343,4.18083051574213,4.75643467621963,3.97225495918677,4.63868773540184,4.32142385992788,4.32336268319182,3.93784517308924,4.08677804051121,4.87305527464803,3.28375695459633,3.02048978750257\nAPC2 - mRNA,3.17664074292704,3.79720108477265,3.41521190501718,2.73580269726632,3.40186052255534,3.63692178898262,3.44912441426689,3.76005191253626,3.15814875323724,3.56112165297631,3.07408122323198,3.55671325598006,3.16948293449122,3.06543925982347,3.34139638911757,3.44734092931775,3.33954151728418,3.82888454360966,3.76802638095669,3.26389869408069,3.76925042909466,3.71092520150067,3.52947979216026,3.60766430979367,4.5007270752809,4.76125026574339,4.56356699069001,4.28005747754717,6.961252770634,4.42345746547238,4.36202981466459,5.11763668925493,5.28318256299549,4.73306729530734,4.90579180554468,4.79874563746107,4.52366133367765,4.72444153498005,5.11848690023855,4.45801777536919,4.02072254876254,5.02048978750257\nACTN3 - mRNA,3.02463764948198,3.01292977582809,2.92978507784693,3.08372600068663,2.03929044317063,3.01543341223635,3.03408691498805,2.63452103045241,3.33872099887906,3.69236618625456,2.62662224626075,2.79772135548386,1.97683785654883,2.7435111649361,2.97882630973286,3.00676833793176,3.12803741209046,2.96215107447312,3.71144285259032,3.18143653388871,3.5177116620987,3.35728824688597,4.11444229288142,3.72795854351138,4.5007270752809,3.76125026574339,4.56356699069001,3.28005747754717,5.37629026991285,3.42345746547238,2.77706731394343,4.09594161815561,4.19571972174515,4.38729245846561,4.42847002801149,4.14085161428606,3.2923357875712,3.57011338858875,3.68238778543187,2.87305527464803,4.02072254876254,4.60545228822373\nPIK3R5 - mRNA,2.02463764948198,2.01292977582809,3.02289448223842,2.86133357935018,0.816898021834179,2.77442531273256,2.71215882010069,2.8755291299562,3.64357558040748,2.16880423019755,1.97454554968106,2.26720663878508,1.75444543521238,1.96590358627255,1.75643388839641,2.78437591659532,2.33954151728418,2.33411985186008,2.76802638095669,2.09397369263837,2.4473223342073,2.50929134033102,2.87740309558057,1.66906485445782,4.5007270752809,3.34621276646455,3.56356699069001,3.28005747754717,5.37629026991285,3.42345746547238,2.77706731394343,2.51097911743446,1.19571972174515,2.26176157638175,2.45648440418109,2.23396101867754,2.35373633223534,2.20087957892303,1.59492494418154,2.87305527464803,3.28375695459633,3.02048978750257\nVAV3 - mRNA,3.17664074292704,3.54344449252687,3.34482257712578,2.97681079677012,3.47986303455661,4.31499369409526,3.9851773145071,3.76005191253626,3.25125815762872,2.97615915225516,3.55950805040222,3.62084359339978,3.25694577574156,2.48047675910231,3.69503334373227,3.2868762571245,2.46507239936803,2.96215107447312,2.76802638095669,1.09397369263837,3.21285708057028,3.274826086694,2.39197626841033,2.08410235373666,5.5007270752809,2.76125026574339,3.56356699069001,4.28005747754717,5.37629026991285,3.42345746547238,2.77706731394343,2.85890242085477,3.12171914030137,2.50276967588555,2.84350752729033,2.34943823609748,3.35373633223534,3.20087957892303,3.11848690023855,3.87305527464803,4.69879445387518,3.02048978750257\nPRKCB - mRNA,3.10264016148326,3.30243639302307,2.72333420037951,3.59829917351639,3.23193552111302,3.77442531273256,3.51951374215829,3.92997691397858,3.95169787576981,3.33872923163986,3.16719062762346,2.68224413806392,3.33940793593354,3.15854866421495,3.17147138767526,2.89985313401525,2.68746482070448,3.12261574666637,2.0675866628156,2.67893619335953,3.12539423931994,2.88780296358475,2.77048789166406,2.89145727579426,4.5007270752809,2.76125026574339,3.56356699069001,3.28005747754717,5.37629026991285,3.42345746547238,2.77706731394343,2.63650999951832,2.0437166283001,2.8897927989948,3.21547630467729,2.64899851795638,1.35373633223534,2.61591707820188,1.76484994562385,2.87305527464803,2.69879445387518,3.02048978750257\nITGA4 - mRNA,8.69549057163473,8.61449424388637,8.38264404256075,8.5937836923939,3.55386361600039,4.06988119625873,3.65075827543654,3.76005191253626,3.64357558040748,3.16880423019755,3.2546534688738,3.18474447859311,2.97683785654883,3.06543925982347,3.41939890111884,3.65884503451146,3.68746482070448,2.68204315528038,3.38951475770296,3.55340531127567,3.29531924076225,2.50929134033102,3.71390436329769,3.25402735517897,5.5007270752809,2.76125026574339,3.56356699069001,3.28005747754717,5.37629026991285,2.42345746547238,4.36202981466459,3.75198721693825,3.87379162685779,3.60968487980206,4.11359669065808,3.50697951308396,3.85623667276452,2.31635679634297,3.11848690023855,5.19498336953539,4.02072254876254,3.02048978750257\nJAM2 - mRNA,6.83199257153959,6.47236139446539,6.65951910278206,6.67725051491121,2.55386361600039,2.12234861615286,2.93455124143713,2.63452103045241,2.83622065834988,2.86924394833864,0.752153128344613,2.41920973223013,2.33940793593354,3.24601150546529,3.07836198328377,2.89985313401525,2.20203799353424,2.45965073394394,3.0675866628156,2.55340531127567,1.7103567400411,1.18736324544366,3.23997317496528,2.99099294934518,4.5007270752809,3.34621276646455,3.56356699069001,3.28005747754717,5.37629026991285,2.42345746547238,2.77706731394343,2.22147250023947,1.87379162685779,2.38729245846561,2.84350752729033,2.5558891135649,1.35373633223534,2.42327200025948,3.17988744490269,4.45801777536919,2.69879445387518,3.02048978750257\nMYH11 - mRNA,5.86593990346293,5.62436448791044,5.3804464868565,4.94879642060052,2.55386361600039,3.22188428970378,2.71215882010069,2.8755291299562,2.57318625251608,1.16880423019755,2.62662224626075,2.79772135548386,2.49141102937859,2.15854866421495,3.34139638911757,1.78437591659532,3.05003490008919,2.7815788288313,3.45990408559436,2.67893619335953,2.4473223342073,2.64679486408095,3.15751101477331,2.25402735517897,4.5007270752809,3.76125026574339,3.56356699069001,3.28005747754717,5.37629026991285,2.42345746547238,2.77706731394343,3.1390103400475,2.45875412757895,3.12425805263182,3.39990087581472,2.73646135920672,2.22820545015148,2.78584207964419,2.76484994562385,2.87305527464803,3.28375695459633,3.02048978750257\nVAV3 b - mRNA,3.85471264803967,3.93046761563611,3.60785698295957,4.1343520737566,4.03929044317063,3.95884988386999,3.9851773145071,4.03307040694268,3.25125815762872,3.33872923163986,3.86763034576455,3.34520915078635,3.62891455312852,3.15854866421495,3.63090300631255,2.89985313401525,8.62494373614642,8.76343148212104,9.08578684162882,9.00985307147415,9.52627367560212,9.57322564608512,9.11580783490565,9.19914600389837,8.20116679342199,7.04665248460564,5.56356699069001,3.28005747754717,6.961252770634,5.42345746547238,4.77706731394343,5.02882742229708,4.82798793724466,4.97225495918677,4.71024099642687,3.97092661284375,4.87729828829235,4.54665441576476,4.7034494009597,6.04298027609034,6.28375695459633,6.47992140613987\nPIK3CD - mRNA,7.02463764948198,6.9774310946067,7.08359041392597,7.04455540340595,3.87579171088775,3.01543341223635,4.12719631937953,3.42301692525869,3.77482011368573,3.97615915225516,4.02517162275103,4.18474447859311,3.75444543521238,3.68211062027196,4.17147138767526,3.72297537193117,8.48188068705459,8.50768343036574,8.75933018198687,8.78397166405782,9.15743996625075,9.08973835992968,8.94963142313575,9.0014744332135,7.08568957600206,4.08317836063075,4.56356699069001,5.60198557243453,6.961252770634,3.42345746547238,4.36202981466459,5.1390103400475,4.87379162685779,4.6849730071063,4.94586924492001,3.69339263731484,4.01670134495777,3.78584207964419,4.02118969888363,4.87305527464803,5.02072254876254,5.02048978750257\nMRAS - mRNA,9.57238907110607,9.41286038271672,9.47307472940317,9.42865592093561,4.84664536522823,5.01543341223635,5.27694343888421,5.12928572220198,4.6088101622468,4.66065732652723,4.94855034114812,4.85216913950624,4.92437043665469,4.85898838235604,5.17147138767526,4.84326960564888,7.55253524061837,7.62957573538625,7.08439495050215,7.1972615010504,6.14776205234839,6.30630431816716,6.74247351549446,6.73515404491559,6.08568957600206,4.76125026574339,6.14852949141116,5.86501997826832,6.961252770634,5.59338246691469,6.09899540883079,6.31833403949206,6.33322324549509,6.23695818538118,6.26383932623869,5.67136633098484,5.96844617635055,5.91969782637898,5.74467206368622,6.33248689328533,6.09111187665394,6.02048978750257\nPDGFD - mRNA,9.24703007081843,9.21586983449836,9.35044712631966,9.30289452114879,5.08991651624059,5.17297468922283,5.21465916062987,5.08198000742363,5.22853808112864,5.31855134970223,5.14447055112337,4.87864135086743,4.25694577574156,4.88646911877815,5.32121850717994,4.98077312939882,5.35584332961328,5.47460107540991,5.40743666570022,5.4690131239853,5.14776205234839,5.66309667641005,5.15751101477331,5.78454207187775,6.82265517016826,4.76125026574339,4.56356699069001,5.60198557243453,6.961252770634,4.74538556035974,2.77706731394343,5.37347559368452,5.21364162974241,5.12425805263182,5.42847002801149,4.75752297473455,4.99759252201006,4.90131929706413,4.82374363467742,4.87305527464803,4.86871945531749,5.34241788238994\nJAM3 - mRNA,10.7687239765809,10.7547712450453,10.8843691573642,10.9354750421027,3.47986303455661,3.22188428970378,4.08139262976641,3.17508941181511,3.89511434740344,3.25626707144789,4.07408122323198,3.41920973223013,3.81333912426595,3.24601150546529,3.34139638911757,3.78437591659532,7.46507239936803,7.54711357519428,7.08439495050215,7.06551724658915,6.48294624393802,6.46348765071789,6.74247351549446,6.77060288091988,5.5007270752809,5.22068188438069,5.14852949141116,3.28005747754717,7.37629026991285,5.88288908410968,2.77706731394343,5.1390103400475,5.0437166283001,5.15988196236254,4.73332460953991,5.05838945409409,4.99759252201006,5.00823450098064,4.7034494009597,2.87305527464803,5.15822607251248,5.34241788238994\nZEB1 - mRNA,10.4245682563706,10.3734111114848,10.3019246186881,10.3404401590145,4.1861318314999,4.06988119625873,3.44912441426689,4.60135416651721,4.20545446801559,4.16880423019755,4.69075258368047,4.58913473367244,4.29876595143619,4.24601150546529,4.25893422892559,3.72297537193117,8.09199819319288,7.99362930594629,8.25741122169562,8.16275197062379,8.52413793125813,8.46813401557426,8.43418282853102,8.35088889443156,4.5007270752809,3.76125026574339,4.56356699069001,3.28005747754717,5.37629026991285,2.42345746547238,2.77706731394343,4.22147250023947,4.15919384572004,4.44618614751918,3.80043880539845,3.34943823609748,3.52366133367765,3.74520009514684,3.9872423669603,3.87305527464803,4.50614937593278,3.02048978750257\nCYFIP2 - mRNA,4.73735569740151,4.54344449252687,5.08898367269619,4.63392308324711,6.43160786594939,6.45454504649405,6.21465916062987,6.24117860227288,8.04891968348248,8.22408666569874,8.96647224914538,8.97686488695555,8.15532487149457,8.21383109971614,8.27475919608728,8.09119711909247,7.23325672414496,7.27129301440464,8.05058023750991,7.83206595225886,8.9343584142392,8.81854030114764,8.62079495890621,8.61937773035746,6.5007270752809,5.9311752671857,5.14852949141116,4.86501997826832,5.37629026991285,2.42345746547238,4.36202981466459,5.47781225349926,5.0437166283001,5.14217996062908,4.71024099642687,3.40388602011985,3.9386988329565,3.2597732679766,3.63931906353999,2.87305527464803,3.28375695459633,4.02048978750257\nPIK3CG - mRNA,3.24703007081843,3.74989536999429,3.66675067201314,3.59829917351639,6.0774255720574,6.30367838086743,6.37512383282312,6.42301692525869,6.50864600032137,6.51753238442863,7.04370757418846,6.91746059975796,6.96389880084133,6.91343616637842,6.98525257889229,7.07593036243916,3.33954151728418,3.26700565600154,3.59114861887261,3.34190120608196,2.93274916137754,3.43529075888724,3.59361012957998,2.89145727579426,5.5007270752809,2.76125026574339,4.56356699069001,4.28005747754717,6.961252770634,4.42345746547238,4.36202981466459,3.63650999951832,3.26610904963655,3.97225495918677,3.75604468603999,3.1725604740134,3.2923357875712,2.93784517308924,3.349812446345,3.87305527464803,4.02072254876254,4.02048978750257\nRAC2 - mRNA,5.76160324364819,5.5572502920519,5.68110596499021,5.29873889165748,6.16267285867591,6.45454504649405,6.00983936874137,6.30437242876007,6.58217503574334,6.35862878907757,6.93537495240038,6.93648542543971,6.44560733976546,6.24601150546529,6.47525213585236,6.36933841731647,3.46507239936803,2.96215107447312,2.76802638095669,4.09397369263837,2.7103567400411,3.274826086694,2.65501067424412,3.54353397237396,5.5007270752809,2.76125026574339,3.56356699069001,3.28005747754717,5.37629026991285,4.42345746547238,3.77706731394343,2.22147250023947,3.12171914030137,2.8897927989948,3.39990087581472,3.40388602011985,3.52366133367765,2.93784517308924,2.29536466232263,2.87305527464803,1.69879445387518,4.60545228822373\nMAPK10 - mRNA,9.13780100106367,9.10039261707843,8.96833831861633,8.82326553851666,5.84664536522823,6.04291414865846,5.99756103896294,5.91655639821626,6.10923915275629,6.30835558259635,6.48911872251082,6.56488718742577,6.20290593602867,5.84504919139817,6.20489438921271,6.13015075343705,4.33954151728418,4.42927708490042,4.5270182814529,3.61753564869539,4.07959054970681,3.57968066822242,3.82493567568643,4.03829866412353,6.08568957600206,1.76125026574339,5.14852949141116,4.86501997826832,6.37629026991285,4.00841996619354,5.09899540883079,4.37347559368452,4.60171208142099,4.60968487980206,4.96549805166894,4.40388602011985,4.57612875357179,4.2597732679766,4.54912125456841,5.45801777536919,4.02072254876254,5.34241788238994\nHPSE - mRNA,5.46942249215488,5.73782253769372,6.14183555496192,6.07078694497913,6.13882611672154,6.2098114574032,6.09298060404162,5.8333008945669,5.66064909376642,5.72339308187519,6.07408122323198,5.82520209190597,6.63708848457422,6.7951732847586,6.69503334373227,6.55696542049224,4.05003490008919,3.7815788288313,3.82247416497907,3.90132861469598,3.21285708057028,2.64679486408095,3.46236559630173,4.03829866412353,5.5007270752809,3.76125026574339,3.56356699069001,3.28005747754717,6.961252770634,5.00841996619354,5.36202981466459,5.22147250023947,4.75643467621963,4.70922055335298,5.02283122673489,4.93440073681863,4.79114164454264,4.57011338858875,4.68238778543187,4.87305527464803,4.50614937593278,4.60545228822373\nMYL12A - mRNA,5.76160324364819,5.54344449252687,5.49862791320481,5.78416571882772,5.20921544461294,5.09635340761992,5.41259853824178,5.40390810231099,5.27362597065717,5.213198349556,5.43397716831836,5.55671325598006,5.41741044793481,4.93990837773961,5.02945238280283,5.24380753523261,4.98863435542505,5.10350692371866,5.08995447584405,4.90132861469598,5.07959054970681,5.39681661107261,5.04732809702288,4.78454207187775,4.5007270752809,5.08317836063075,5.14852949141116,3.28005747754717,6.37629026991285,5.42345746547238,5.09899540883079,4.8844375129619,4.89615943988624,5.31012459794315,5.00397219948358,4.79874563746107,4.6756644271227,4.42327200025948,4.74467206368622,3.87305527464803,4.86871945531749,3.02048978750257\nMYL9 - mRNA,14.9778034417177,14.975725389741,15.1590044613161,15.1548291771104,3.93237523925411,4.60039591295751,4.61904941570921,4.30437242876007,4.20545446801559,3.75376673091871,4.55950805040222,4.38268385620502,3.97683785654883,3.68211062027196,3.69503334373227,4.10630401148268,8.2199599015315,7.6593230787803,7.18652773553911,7.07696726733268,5.89357856409687,6.16464316894357,5.98891141079756,6.25402735517897,11.5451211946394,11.0115486836497,11.1856188101464,11.1565744241122,12.9461458782438,13.414270544626,13.3825468320051,11.6607262369391,11.5221492088675,11.5932892914433,11.6856729690646,11.7958853533723,11.7503411134172,11.8236940431299,11.7884503046828,11.9201791867621,12.2875090894574,12.2420769087674\nGSTM3 - mRNA,12.0200907613858,12.0257094849047,12.2675556270026,12.2923087519383,3.03929044317063,3.4024565353456,3.58662793801683,3.69865136787212,3.05861307968632,3.69236618625456,3.07408122323198,3.74113782711749,4.45488515335347,3.55086608699371,3.63090300631255,3.52134151076152,11.4122222694419,11.4584761877251,11.6461952235534,11.6845607425534,11.2646258100393,11.3386969647353,10.9629703971536,10.9459646937958,4.5007270752809,5.76125026574339,5.14852949141116,6.08741239960477,5.37629026991285,5.74538556035974,7.36202981466459,6.02882742229708,5.79265486413238,6.13324683585907,6.04144690490224,6.14884440503959,6.21998494334651,6.07534869683917,5.93477494706616,5.45801777536919,5.69879445387518,5.34241788238994\nIL1R1 b - mRNA,8.6572290131609,8.68220856248272,8.91163773113667,8.72758219046136,4.13882611672154,3.70731111687402,3.58662793801683,3.76005191253626,4.89511434740344,4.81266041997228,4.21158474698191,4.00417223295129,4.41741044793481,3.80240485398967,3.49339948256262,4.05739441100173,9.12328388211983,9.05459732323772,9.76975934818115,9.80564066620272,9.84135622957508,9.886067912214,10.1468637705738,10.1814761227269,7.3080819973385,6.61923126087096,6.73349199213232,7.08741239960477,7.69821836480021,7.46785158483083,7.36202981466459,7.4395647841423,7.86814506371665,7.75069718932913,7.88274915888318,8.62770897706274,8.44825393101963,8.3741402929286,8.43412873227848,7.96051811589837,8.32815107395479,8.14977280444754\nPDGFRB - mRNA,12.5468922243522,12.5275342568043,12.3673291041624,12.3056583055973,4.58948752573111,4.56292120753885,4.41259853824178,4.46049163067736,4.83622065834988,3.97615915225516,4.2546534688738,4.71199148145798,4.59574768919332,4.55086608699371,4.38092475330421,4.72297537193117,8.49665874242587,8.677204598344,8.94563757554413,8.96895504029215,8.97088429026432,8.8773612168631,9.16148355984657,9.10785670703608,7.9601586939182,8.33110587407434,7.2640067088311,8.03494497971063,6.961252770634,7.00841996619354,7.23649893258073,8.56330015256454,8.67953549900941,8.49058026687763,8.40888965904197,7.95733754809579,8.12192065701227,8.1895642656952,8.17612552610649,8.12098278809162,7.8886190127552,7.92738038311109\nFGFR1 - mRNA,11.5785868669036,11.5343121045769,11.3954875153792,11.4210293214609,2.81689802183418,3.89995619481642,3.29712132082184,3.49701750670247,2.95169787576981,3.33872923163986,3.2546534688738,3.26720663878508,3.69304489054824,2.61798028285224,3.75643388839641,3.2868762571245,10.7149759370089,10.6081273161222,10.5514024400799,10.5908274700264,10.3317981532938,10.2768137265596,10.2025479031515,10.1958900895377,7.5007270752809,7.71544657613027,7.73349199213232,8.13803847267474,8.37629026991285,9.05281408555199,8.86453015519377,9.0359659575983,8.87660664246484,8.95841303403314,8.91212776232867,9.15183032202313,8.93744594964641,8.88738010610625,8.81045794392719,7.96051811589837,8.44026144027633,8.34241788238994\nPDGFRA - mRNA,11.2754624063365,11.1881327663625,11.1777125912905,11.2802403106081,3.93237523925411,4.56292120753885,3.9851773145071,3.98244433387271,3.89511434740344,4.07569482580607,4.2546534688738,4.48959906012153,4.25694577574156,3.55086608699371,4.07836198328377,3.59173083865292,4.20203799353424,4.12261574666637,4.0675866628156,4.26389869408069,4.25467725626491,3.99471816750126,4.02424448390984,4.29355571936561,7.3080819973385,9.48234945445058,9.47045758629853,9.60198557243453,9.07672998805394,10.2876436101267,9.93693865072182,9.47571378761526,9.2607318721821,9.25868437285274,9.2913200626608,9.49134886137019,9.09070192640154,9.28214108100506,9.19669573258925,9.9815797314262,9.8886190127552,9.93935302477717\nAKT3 - mRNA,10.4429588000845,10.3621495623832,10.449683376834,10.3585201199829,3.62425294389178,3.83855565015227,2.93455124143713,3.17508941181511,3.71068977626602,2.86924394833864,2.48911872251082,3.48959906012153,3.62891455312852,2.85898838235604,3.17147138767526,4.19941341587416,3.7870004942554,3.62957573538625,3.82247416497907,3.84886119480184,3.88028174148341,3.57968066822242,4.11444229288142,3.72795854351138,8.40761767088942,7.84871310699373,7.73349199213232,7.08741239960477,7.69821836480021,6.74538556035974,7.09899540883079,7.66257229610451,7.54974865979954,7.59023251713589,7.55370621189375,7.15282425595213,7.35748846709645,7.34043093132183,7.42781495834628,7.51691146442276,7.92761314437106,7.72092950564367\nITGA5 - mRNA,12.3152965279414,12.3278276686479,12.3638572464901,12.4038818416854,3.23193552111302,3.12234861615286,3.12719631937953,3.17508941181511,2.71068977626602,3.16880423019755,2.86763034576455,2.79772135548386,2.97683785654883,3.24601150546529,3.17147138767526,2.65884503451146,4.8573898221468,3.96215107447312,4.0675866628156,3.41590178752574,2.58482585795724,2.18736324544366,4.15751101477331,3.83898985590013,9.02428903133791,8.20419376159212,7.47045758629853,7.18694807315569,8.37629026991285,8.25634747963712,8.09899540883079,8.49863072500681,8.37164346380891,8.30217584482766,8.20105953190842,8.04987795423857,7.77720245358476,7.9105378270935,7.85231278687419,7.33248689328533,8.28375695459633,7.6643459772773\nITGA7 - mRNA,5.15849339621678,5.09093228782936,4.83024940429602,4.89108092274423,4.03929044317063,4.01543341223635,3.71215882010069,3.56740683459387,3.57318625251608,3.81266041997228,4.02517162275103,3.95526263247034,3.86992265263232,4.2029427835734,3.56378881045402,4.2868762571245,4.23766190326496,4.16009045208503,4.35298888167785,3.55340531127567,3.98337523444751,3.94225074760712,4.31797568696655,4.21338537068163,8.9601586939182,8.17064120188109,8.37092191274761,8.18694807315569,6.37629026991285,7.28143846059995,7.23649893258073,8.66257229610451,8.72866001011803,8.54042221736495,8.53412181557535,8.59297843230012,8.60640176468559,8.64520430517012,8.56215120301753,8.91744939400649,8.64130895921442,9.24930847799845\nLEF1 - mRNA,5.59617963444082,5.74989536999429,5.83024940429602,5.94879642060052,3.03929044317063,2.89995619481642,3.58662793801683,3.69865136787212,3.71068977626602,4.3782575958265,5.14447055112337,5.65187048902041,3.86992265263232,3.15854866421495,3.81532757744998,4.15360972626103,3.96757273989722,3.7815788288313,3.65254916353675,3.7378298824131,2.29531924076225,3.50929134033102,3.87740309558057,4.08410235373666,7.08568957600206,6.08317836063075,6.37092191274761,6.08741239960477,7.37629026991285,7.70885968433463,7.23649893258073,7.49449099464589,7.59830547997774,7.64782000868883,7.71314672760228,8.49605586404772,8.2424795811336,8.24527369828149,8.19297744434314,7.51691146442276,6.98419667273743,6.92738038311109\nBCL2 - mRNA,6.58263310260287,6.71952172095077,6.30829670110066,6.15901412799087,5.06482553527776,5.22188428970378,4.77105250915425,4.95644912533977,4.4991856710723,4.78351407431276,4.69075258368047,4.45483364196085,4.86992265263232,5.22463785467272,5.27999584445342,5.10630401148268,5.86879458541904,5.57512795136387,6.15504950406594,5.75218517539017,5.91981010567005,6.0700062948055,5.40989817640759,5.40603044862402,4.5007270752809,5.22068188438069,6.56356699069001,6.08741239960477,7.69821836480021,7.28143846059995,7.02499482738702,6.44386492157592,6.56727858435711,6.39478699501254,6.49751167244114,6.51936323753979,6.31566829140182,6.44880709236662,6.29536466232263,6.45801777536919,6.39923417201627,7.34241788238994\nMAGI1 - mRNA,6.1945626509243,6.22670906685983,6.29901888751265,6.18326167423754,5.47986303455661,5.56292120753885,5.48474832399761,5.36491397069513,5.4991856710723,5.81266041997228,6.24400622467429,5.94277168828714,5.72407178616886,5.85898838235604,5.78618123179046,5.57445284722108,6.76275294800872,6.60939785344862,7.17087447122762,6.90775488385541,7.55165899402204,7.61362800014575,7.47518963665931,7.38788310191376,7.08568957600206,7.59414027990813,7.14852949141116,6.60198557243453,8.37629026991285,7.3303480610809,7.58442223600103,7.40027565302824,7.26610904963655,6.8897927989948,6.91591602281838,6.76274068400557,6.57612875357179,6.9105378270935,6.68238778543187,7.77994587025655,7.37121979584667,7.02048978750257\nCCND2 - mRNA,8.1945626509243,8.23102205973091,7.84944296952933,7.65574944570029,3.23193552111302,3.70731111687402,3.65075827543654,4.03307040694268,3.89511434740344,3.41673174364114,2.48911872251082,3.34520915078635,3.33940793593354,2.85898838235604,4.07836198328377,3.72297537193117,7.50216171810325,7.28831616224695,7.05627134958776,7.05975797730046,6.13662149474319,6.14155955583053,7.14151053607959,7.13938478923785,6.82265517016826,5.76125026574339,5.14852949141116,5.60198557243453,7.961252770634,7.37765377585926,8.2689204102731,7.13369654015103,7.03366296363618,7.02149847820662,7.09589468892462,7.88021969866757,7.68465321034996,7.69005854095438,7.69295702714206,6.87305527464803,6.45368195603865,6.02048978750257\nTCF7L1 - mRNA,8.01458398481806,8.14450096936833,8.270821995682,8.05444820816438,3.62425294389178,4.35938781345371,3.77105250915425,3.76005191253626,4.15814875323724,4.33872923163986,5.07408122323198,4.85216913950624,4.6613360308209,4.24601150546529,4.07836198328377,4.72297537193117,5.27242732142564,4.96215107447312,6.19686967976056,5.67893619335953,6.40151864459418,6.45414978613856,6.03583245818505,6.13938478923785,8.40761767088942,8.13628969709032,7.14852949141116,7.86501997826832,7.69821836480021,7.28143846059995,7.9063503308884,8.19365455609971,8.36163166068084,8.13324683585907,8.05525270442727,7.97542800507869,7.90063079212298,7.94459200552982,7.9743033112528,8.29932002935013,7.45368195603865,7.10795262875291\nTARBP2 - mRNA,8.04946950246192,7.98511183168833,7.83024940429602,7.78416571882772,8.91843604829624,8.99623210587307,8.95533280357364,9.01743975522892,8.57318625251608,8.48168718548191,8.84314766056521,8.80118521078743,8.84543996743297,8.9117654133939,8.84389672964675,8.78437591659532,9.1149273297642,9.08900735402355,9.08300171520229,9.02026968741949,8.97558107696571,8.82579915943413,8.79456202664292,8.81711767542262,8.40761767088942,8.30040907685142,8.08712894674702,7.92391366732189,6.961252770634,7.63291083110133,8.55842702746809,8.5672473370812,8.62366105425463,8.63363720154651,8.5983950294538,8.76013418819109,8.7142176678921,8.80820989267264,8.75479628095992,8.54548061661953,8.48015416739984,8.51234288383225\nABI2 - mRNA,8.49133726858332,8.43732937831062,8.53662207819085,8.48338539880656,9.01657036667054,9.07488187731709,8.98828322537492,8.92663347090421,9.13695453096811,9.15322268899869,9.04217197527723,9.07312960534955,9.29360424632041,9.2513058051348,9.19106211603314,9.21622170356071,9.07116911317853,9.03959515989847,9.32025209526585,9.25132062800122,9.27514135882462,9.24535496820283,9.39197626841033,9.43614577953209,8.58818991653124,8.65606802905134,8.73349199213232,8.13803847267474,7.961252770634,7.94701942152939,7.98652067957238,8.91573364345144,8.74415634644119,8.80935722463843,8.7658733034081,8.71104465567687,8.51864325891103,8.55108212780557,8.66772747872574,8.7310362697756,8.64130895921442,8.82784470956018\nPIK3R2 - mRNA,8.96715215482122,8.92248955821259,9.00017440573833,9.00769010965409,8.15377645818751,8.14788370826,7.89206791011562,8.04545413139851,8.41364948638562,8.44259982941182,8.45027898064746,8.50242310047911,8.32935427126961,8.50948784576138,8.38335968222126,8.33383973609509,9.26812718399201,9.09022123296844,9.61640857127435,9.61065864194798,9.74836118411045,9.62511531776726,9.69666982588134,9.60668388499149,8.67065207672321,8.48917072030659,9.05542008701968,8.73948909618446,7.69821836480021,7.88288908410968,8.44949265591493,9.1482626532857,8.915450778495,8.92774342822777,8.83552946986681,9.00874807827871,8.80769282080649,8.88971267741614,8.88032716304378,9.26537269742679,8.75407688937637,9.10795262875291\nLIMK2 - mRNA,8.14011486690192,8.12840699324802,8.08088620499759,8.09335470188441,8.67487901696175,8.639174240714,8.65075827543654,8.49701750670247,7.96206459132807,8.16315766705641,8.13297491228554,8.09728163734277,8.38622879268653,8.47596127797982,8.56807616885711,8.45444198569289,9.71141714244893,9.63623946566726,9.92401219144113,9.8964900577596,9.80575376283365,9.7816878493685,9.86689896879013,9.78454207187775,8.20116679342199,8.64389331510523,8.47045758629853,8.63760948216525,7.69821836480021,8.03816730958759,8.47750703208452,8.52930576524105,8.45499220878275,8.50796234510587,8.48566542959303,8.65882713532449,8.43055192928617,8.48010322285616,8.41510390659672,9.04298027609034,8.76488364433295,8.60545228822373\nDVL2 - mRNA,8.1901030027336,8.25239571052348,8.08628956352693,8.08372600068663,8.46555519787272,8.41302577705579,8.44007927466389,8.46510825874773,8.43116724764365,8.35862878907757,8.59555080032775,8.72299048040723,8.84896303399667,8.83626830585596,8.86843891390954,8.84146510880214,9.39876305432027,9.40020904231785,9.57835083073349,9.6165552238932,9.72065172076898,9.60732342329154,9.36006702045558,9.36371293478914,8.67065207672321,8.59414027990813,8.56356699069001,8.08741239960477,9.07672998805394,8.00841996619354,8.2689204102731,8.93266145947226,8.84820621666332,8.83023595431434,8.82750704859662,8.70292439948749,8.73166983685144,8.77704896118618,8.64749299498569,8.68041019670564,8.61765769114977,8.57507863918021\nERBB2 - mRNA,8.39193104857827,8.40142548765444,8.28028232493107,8.30979408016648,9.21635397991416,9.15104404915333,9.11018989407384,9.16802774798092,9.81636023598903,9.85004264197536,10.4635333123163,10.3728241934295,9.87770466452958,9.87622508728134,9.82878983725654,9.7434118660087,9.35786816509077,9.32177145943243,9.97222128448375,9.89487359255868,10.5112566399614,10.4137754382324,9.92101632450541,9.89627965925111,9.9601586939182,9.08317836063075,9.58593480371846,8.8346463292248,7.37629026991285,7.70885968433463,8.09899540883079,9.46517542782133,9.14408695332983,9.22754586104384,9.09478106861391,8.88141944513246,8.58095113702863,8.71789274915111,8.64341181817387,9.06287983352805,8.86871945531749,9.21031434638259\nPRKCD - mRNA,8.05438499287604,8.14906646381412,8.14959895508161,8.18626416239907,9.35476351592069,9.32901416441019,9.30461585736877,9.29662292915514,9.49085668687784,9.49410968528723,10.1372965182356,10.105009751975,9.71251475474646,9.76772740235693,9.7154698378098,9.66803745879218,9.12209304834121,8.97535057222798,9.66097778560733,9.67234638399918,9.99575895890334,9.92545550506415,9.90372892217771,9.79748886867369,9.5007270752809,9.65606802905134,9.39645700485475,9.58383822572427,7.69821836480021,8.72723821364949,8.9063503308884,9.71332559656915,9.4811219406074,9.50016631570683,9.48226284537219,9.43117771178759,9.21585705772384,9.32672351190123,9.27879681280414,9.24809470599495,9.3426506436499,9.3603397903872\nIGF1R - mRNA,8.13548341817028,8.07182346488166,8.04803404451692,8.04124266936512,8.83741503917469,8.78869944371468,8.78719221180951,8.67493529889831,9.11234506362411,9.08169356642751,9.51591878185453,9.61790031789905,9.38744063235534,9.33721351522142,9.42768007620264,9.30007575487936,8.86310347334119,8.8984425865222,9.71144285259032,9.6155741323621,10.3857257578775,10.2794596604344,10.1475315303825,10.0749154328903,7.67065207672321,8.65606802905134,8.60796111004846,8.13803847267474,8.69821836480021,8.53198192225055,8.39177715805864,8.38470284910778,8.38356663118473,8.43895457828811,8.23784411770575,7.93440073681863,7.85092287286805,7.97346908281996,7.96561235098875,8.19498336953539,8.19064755020485,8.41280721028133\nPRKCZ - mRNA,6.3938714591477,6.4868609641605,6.29901888751265,6.48582444425798,8.98682302327649,8.98210523617029,8.93294019153115,8.98726671732956,8.63712055827018,8.8796106638969,9.26258765041601,9.29326893537128,9.30262514689183,9.25921100322015,9.30204195048068,9.29744549883469,9.19072268030641,9.11905206834329,9.3827362250719,9.42377003085907,9.67421614777869,9.61362800014575,9.76091918281528,9.73873838226463,9.20116679342199,9.11880227036148,9.08712894674702,9.16270052690901,7.69821836480021,8.17834496763585,8.30062927000044,8.73979780793034,8.57249629362813,8.76088267317547,8.72038555383241,8.49918535560216,8.42698531426598,8.43290070145726,8.50031194919967,8.60097572921123,8.81773552659869,8.57507863918021\nPDGFC - mRNA,9.4546254902268,9.53736266160723,9.85894483729649,9.87349250741859,10.1846767682995,10.1938109787095,10.2551906403559,10.2458812212382,10.0367106257873,9.99694071439166,10.1763194171627,10.0909276973594,9.77403616357026,9.81228944292152,9.84301245438714,9.87183875784566,8.7870004942554,8.80394664185975,9.00275171241929,8.90293786755763,9.37924172430734,9.34218135449576,9.05161545542598,9.11890131631393,7.08568957600206,8.60674031668777,9.34492670421467,8.18694807315569,5.37629026991285,6.00841996619354,6.68395790955195,7.96742687763293,8.02860973590989,7.98968203094495,7.93346422402923,7.71508770841416,8.01194781498713,7.95353898019438,7.96996437552846,7.57349499278912,7.98419667273743,7.6643459772773\nIRS1 - mRNA,8.98656960864847,9.03653013488699,8.94622196765761,8.83479179189769,9.30071379909844,9.25457290840995,9.36201375049686,9.33873908783093,9.80296641924941,9.69725834096234,9.77081697260965,9.77323867028452,9.75162491615,9.80870287789004,9.6960131027111,9.80766823785207,7.06498524155516,6.92997066872397,7.56344168970277,7.50760162166255,8.22605657832514,8.20973105847211,8.66623792966738,8.66530293566162,7.9601586939182,8.40510645551812,8.65102983194035,8.28005747754717,7.37629026991285,7.59338246691469,7.23649893258073,8.18597381901809,8.19120424062266,8.20959772198919,8.09589468892462,7.62341310776191,7.74318842131971,7.73745059554191,7.68768208510139,8.08250864027698,7.6056850494837,7.47992140613987\nEXOC3 - mRNA,8.23841694051372,8.24814623752212,8.11035732348875,8.01108069885486,9.41804976065467,9.37838877538377,9.37156015450004,9.41468794106423,9.18936748384796,9.31346247302943,9.34086776392688,9.40562066748218,9.35187728789575,9.33097617286043,9.40389231485133,9.2961285703627,7.85738982214679,7.91634738485999,8.07040167842265,8.01283692991297,8.20220983637077,8.11217574904944,8.11170732516018,8.19001086230782,8.3080819973385,8.23698369670979,8.47045758629853,7.67237490032593,8.18364519197045,7.17834496763585,7.58442223600103,7.92806444536216,7.8989311891363,7.88713834554882,7.87757820338003,7.80381662700849,7.93116516027109,7.95799174620102,7.86405162333095,7.96051811589837,8.05634645849326,7.87847078263015\nNUMA1 - mRNA,8.60122696917814,8.6111890991627,8.53662207819085,8.51238527052911,9.71070514058879,9.77647314138142,9.75473020385232,9.70061022091018,9.9732117704301,9.96565502546471,10.4351477120263,10.4371316402274,10.1953146028233,10.2036257161499,10.3557268830806,10.1557895731238,8.67452576499698,8.53468412565276,9.12557838557477,9.04525840760535,9.46078459401386,9.42934639513799,8.6130799937782,8.6386912054143,8.40761767088942,8.52943459052032,8.37092191274761,7.44998247898948,7.37629026991285,8.23081238752999,8.16938473672219,8.9940620041364,9.27467106313997,9.1521640492017,8.99802783573432,9.3398695272487,9.15463623215564,9.1375175179256,9.16762517066883,9.23060727926612,8.29870729606231,9.08657897796035\nAGO2 - mRNA,8.40353455649842,8.36067416251959,8.4428546600863,8.41336219070926,10.5845307522473,10.564699751065,10.5633873148609,10.6024022559501,10.4060762666808,10.4326602497936,10.108238233964,10.0702614234091,10.458349008657,10.4995114284666,10.4086813484303,10.3459820916149,8.46131048057184,8.46152802118461,8.70421128335925,8.71052253641736,9.11691608539564,9.08066477630422,9.28800587138708,9.25527915138843,8.20116679342199,8.33110587407434,8.2640067088311,7.67237490032593,8.37629026991285,8.72723821364949,8.77706731394343,8.66257229610451,8.69556560882836,8.55888628026854,8.50591411343684,8.74439554074894,8.43055192928617,8.54369504028422,8.51825242960073,8.48776511876324,8.02072254876254,8.19041478894489\nBHLHE40 - mRNA,7.72815676055505,7.82314435202493,7.9799965238007,7.84242511680012,6.4607542116089,6.06988119625873,6.1932855098373,6.10582674937799,6.92368349960021,7.36847657503391,7.86763034576455,8.25462857270561,7.91431677199077,7.71313751589259,7.80082800775487,7.80674372962377,8.16088066877748,8.46527528313782,11.0437927529642,10.6442394995071,9.9149278842903,9.7030630837277,11.0615700195987,10.8687371992942,9.40761767088942,9.70376477108263,7.56356699069001,7.08741239960477,6.37629026991285,6.88288908410968,6.23649893258073,8.71154339122714,8.60680694854375,8.93547217014506,8.61557947242265,6.07526327265848,6.64529077807918,6.65483606749418,6.66101413463931,7.39661723070504,8.44026144027633,7.34241788238994\nTGFB1 - mRNA,10.78174981676,10.8199268422369,10.7482607401341,10.7094348437511,7.01329523463768,6.99447379360071,7.00371326594453,6.92997691397858,7.54127700456134,7.60343245783428,7.99373911491437,7.90785327208391,7.45488515335347,7.78790528429456,7.61855461388491,7.55696542049224,9.92176905028409,9.79578532132957,9.97297366783373,9.88513658119339,10.0524314080402,10.0414524253142,10.4220915993184,10.3958506687416,7.08568957600206,6.84871310699373,6.88549508557737,6.86501997826832,6.961252770634,7.23081238752999,6.68395790955195,7.78963975841765,7.75337487647057,7.69411366096277,7.26779733980737,7.43363336351391,7.33862943984513,7.4005519237594,7.3762846577062,8.19498336953539,7.25338330555282,7.87847078263015\nRAB8B - mRNA,10.0611165620135,10.0585377741314,10.0278171608075,10.0387531173394,7.37148687351182,7.12234861615286,7.06971082471877,7.15808298650942,7.436133500537,7.51753238442863,8.47780340928299,8.36640831649219,7.91431677199077,7.7915438613579,7.93965571245218,8.0386172039711,9.04124178163119,9.07190492633283,9.10238562539287,9.06695347870467,9.18338596207912,9.11957799741204,8.85958181849333,8.82219461335715,8.89304449805966,8.52943459052032,9.02299860932731,9.03494497971063,8.18364519197045,8.70885968433463,8.70780465150632,9.03168707643632,9.03240131507227,9.05144341045819,9.03565526709248,8.60610055951867,8.79667982808407,8.81681568324032,8.78351378988888,9.17683602282514,9.23017591439149,9.22994315313152\nSMAD4 - mRNA,9.04453720691969,9.05123913272556,9.20280357225335,9.16967260948959,6.9934867535575,7.24573103165815,7.00371326594453,6.92328226140457,7.34929024058925,7.48168718548191,7.45259284648571,7.50242310047911,7.54452236583815,7.71696937748361,7.31612963050714,7.41373253667493,9.48651511134466,9.58204736530366,9.93022402037439,9.7333144012906,10.3084092402027,10.1054810964754,10.2038325827029,10.0720768780328,9.25561457744437,9.13628969709032,8.92111899530809,9.00797793211036,8.69821836480021,8.69024400616728,8.92681443344811,9.15089530920035,8.76253487575605,8.6986623616544,8.7880904129708,8.58563645695896,8.4744062188756,8.73226103943935,8.51973744778732,9.42764412632567,9.25338330555282,8.97468609788945\nHSPB1 - mRNA,11.8170664566749,11.695144102573,12.0207796100675,11.9976839208043,8.32733254390558,8.18227243380492,8.24093848542872,7.98887060303215,8.66064909376642,8.79451307326202,9.07257763264159,8.89656325886469,8.67925793881816,8.95621019006871,8.65327081934101,8.63404164351089,8.57185341781815,8.39038807150682,8.48332843110569,8.35841629286498,6.97088429026431,6.77982028271174,7.68843367578157,7.49773028276083,8.58818991653124,7.34621276646455,6.73349199213232,7.44998247898948,8.961252770634,9.25634747963712,9.13461931856151,9.98671254840086,9.54798389500347,9.4578606809001,9.50675167226494,9.84989712299483,9.64759111612841,9.79644788554636,9.62697067111234,9.70594528881277,9.16640000395818,9.32427053567968\nMYLK - mRNA,13.7351571224562,13.7123576342728,14.099918787433,13.9748044550319,8.11763189445242,7.98741903606676,8.05200882298531,8.15522898945427,8.69627300349714,8.78167272748859,9.43982041259983,9.47123429271296,8.7317253587123,8.79335971438667,8.89086020861734,8.86829840014041,9.72167424637745,9.75350551783701,9.78011366325542,9.86298501373539,9.70093348615447,9.79376845814144,9.56089905026326,9.55983578470306,7.9601586939182,6.76125026574339,6.14852949141116,6.98049719568826,7.961252770634,6.88288908410968,7.5319548161069,8.07945349536705,8.04120977268953,8.02149847820662,8.03216909131423,8.02620904834494,7.90832518391298,7.79831143160641,7.98294222952667,7.77994587025655,7.22235640993219,7.60545228822373\nIL1R1 - mRNA,10.2357147575906,10.1806266763783,10.283226000796,10.2847997006996,5.34045997789119,5.63692178898262,5.77105250915425,5.36491397069513,6.30654059312991,6.34871332021249,5.70634943873149,5.60507627754146,5.72407178616886,5.55086608699371,5.49339948256262,5.89985313401525,10.5900552788794,10.6719444507326,11.3172927197853,11.3221911349318,11.4090614068114,11.4573664988221,11.7244604416719,11.7247991913068,8.89304449805966,8.85928234870392,9.51776330107688,9.16270052690901,9.76860769269161,9.08166894822418,9.28486195414213,9.22397500744264,9.43698789907453,9.34536177799969,9.54639318279191,9.9378645921222,10.0414036164906,9.92311788734269,10.0227615476401,9.75569832400987,10.2418262741304,10.0204897875026\nEPB41L2 - mRNA,12.2106142863681,12.2570292842639,12.4475856737514,12.46850975078,7.42669237553527,7.45965193692714,7.45362580650183,7.40390810231099,6.63496245010277,6.70018569071386,7.23327981808123,7.27220731984345,7.07034974247197,7.12433294887703,7.17709593686913,7.21622170356071,11.6247335379905,11.6654439833411,11.6834461563987,11.658361388691,11.5576751365341,11.4905731943884,11.1007153097699,10.9629196383509,9.926991829983,10.5718219004845,10.2500675178732,10.2107948151101,10.8025550246149,10.5319819222505,10.2848619541421,10.5358669224414,10.55107081817,10.6028254582113,10.4120225485269,10.4389396463473,10.480010599308,10.4448469058534,10.4364892919121,10.3242663864804,10.3676794381414,10.3692179417337\nMYH10 - mRNA,11.9413464971159,11.8677981590883,12.1478235288409,11.9705112827055,8.54027455125243,8.43393476681877,8.55764887104506,8.61388126439337,7.7630108113961,7.96970413011786,8.42221919744219,8.36640831649219,8.55102588567581,8.48047675910231,8.4568736065375,8.63404164351089,11.3849288862797,11.4619969615563,11.6161677809807,11.5674257815906,11.5996071765139,11.5645737758322,10.7890090855072,10.7220856575145,9.54512119463935,9.60045405384034,9.49430432825289,9.46988203642718,9.54621527135516,10.1445566541796,10.1949198288293,10.1548353072092,10.2482877725493,10.1499513263126,10.0903179713472,10.152327374583,10.2018375966363,10.2269418755092,10.1718816088876,10.1209827880916,9.97025748177955,10.0757722230038\nCD44 - mRNA,14.1095198380916,14.0685134946218,14.3632314238333,14.4451215338525,12.1214394845245,12.1455089010721,12.0558997385729,12.0004457532944,12.4898121747273,12.7586894094878,12.7528573973559,12.6497105858692,12.3343898611669,12.4761026009011,12.4322439400288,12.3531237680259,6.63499740081035,5.42927708490042,5.75408718999882,5.4690131239853,6.02021200262788,5.96872295896832,5.83823249829989,6.03829866412353,7.9601586939182,6.80564438510185,6.88549508557737,7.36752031879751,5.37629026991285,7.97804631715002,7.68395790955195,8.60996821206582,8.32087485306744,8.29818498463163,8.31063353771763,8.93440073681863,8.89030734885421,8.80079242111016,8.72162941702473,8.42764412632567,7.39923417201627,7.77537728966604\nGNAI1 - mRNA,11.15820802483,11.2060444046423,11.2108650742226,11.2258330579892,10.7397301613117,10.7370584602681,10.7674412556019,10.7118223016505,10.4373723971704,10.4267812887615,10.7188988892441,10.7036868500406,10.3989026227216,10.3885690999206,10.4672403220958,10.4088667815031,8.7870004942554,8.81866814756652,8.26480335592565,8.26640120128386,8.28526557609833,8.47045159846766,9.21215523082552,9.20822366556585,5.5007270752809,5.9311752671857,5.56356699069001,6.44998247898948,5.37629026991285,5.42345746547238,5.77706731394343,6.8907512868941,6.75643467621963,6.96220129452285,7.00397219948358,6.94820653634366,7.24044904590623,6.96906390369996,6.80437830981048,5.68041019670564,4.69879445387518,6.02048978750257\nITGA1 - mRNA,13.4235000768666,13.3721302609617,13.4413400886192,13.3738271532952,10.5701147710131,10.435879536883,10.4841984116445,10.4575987180559,10.9948303465644,11.0091887809092,11.5348783838935,11.4965595618517,11.2094300946603,11.1617151708801,11.2900880770268,11.2064406827681,10.1833198475917,10.1943603541698,10.6388652813302,10.6562161168594,10.622248016708,10.5305489608915,11.7123255520268,11.7102676736412,7.08568957600206,6.76125026574339,6.56356699069001,7.18694807315569,6.961252770634,6.3303480610809,7.58442223600103,7.62140310712811,7.54974865979954,7.57051428252138,7.66893402226995,7.82890760797132,7.96599052462424,8.04218183290397,7.9743033112528,7.39661723070504,7.42671490843838,7.6643459772773\nTP53 - mRNA,10.214462208362,10.243884210668,10.1288029908096,10.1374579846244,11.4208333943552,11.3473838629618,11.3460622294492,11.3636782575225,11.0762168981572,11.01977238118,11.0623872001868,11.0269099203673,11.5865442812515,11.630223878607,11.5800665004734,11.576352737494,10.7006877813027,10.6642218781931,10.7946562208393,10.74098233108,10.6041638587957,10.574303490768,10.6484208648838,10.6582227887464,8.20116679342199,7.568605187801,7.2640067088311,6.98049719568826,7.69821836480021,6.3303480610809,6.47750703208452,7.18083051574213,7.24011384110361,7.05971780043711,7.19898818188872,7.03701580330152,6.89805684845915,6.73485815092538,6.75479628095992,7.87305527464803,6.45368195603865,6.47992140613987\nCTBP2 - mRNA,10.3723820361735,10.4171252093955,10.2844017914405,10.3077276748847,11.6803099807758,11.6540085623453,11.7091030086883,11.6761801820896,11.7870195356154,11.8090491664199,11.2763683309787,11.3103858569847,11.511835444579,11.5411726907899,11.5364267661572,11.4830805833657,4.83430620903375,5.24972766456971,5.8620025291657,5.39775444081548,5.58482585795724,5.45414978613856,5.75654870070619,5.6386912054143,10.5451211946394,10.4087086921983,10.75339154957,10.6979099924331,10.2831808655214,10.7498869525947,10.8645301551938,10.9584380944057,10.9202335748651,11.0178629574901,11.0646681311711,11.037284937719,11.0988503863116,11.0466309639938,11.0653296212472,10.8214225062327,10.9155403120705,10.8722388289186\nKRAS - mRNA,9.45183407682924,9.35376894230367,9.45647992338891,9.44127800530471,10.4857830061004,10.3675617448994,10.4670463222642,10.4307081616547,10.0928225053593,10.1666893580267,10.5530530282649,10.6129814756268,10.4339255347178,10.4375788006646,10.4655177009468,10.4390119451233,9.41538527526768,9.39334138889617,9.2299779915725,9.09959824183225,8.70847700682287,8.70699949828687,9.00521322312666,9.04988663839875,11.4894117620531,11.5556661320935,11.666854799102,11.362206518901,12.0627907970961,11.7677533733882,11.8132409264969,11.6786253393146,11.7096446633023,11.7189558799557,11.6722973848522,11.828441144086,11.8592171065958,11.7916225537278,11.830490467118,11.5490123075898,12.0868117392203,11.8278447095602\nPAK1 - mRNA,9.87638669089804,9.93774227943387,9.72679805568308,9.77522028516075,11.9412347738252,11.9274014872339,11.9019833789807,11.8908369546465,11.9827893340907,11.956706789589,12.3758822344085,12.4128557922467,12.144793552548,12.1169201251648,12.2129594315567,12.1497870508987,7.09442901944765,7.10350692371866,7.26234351723785,7.25384502941676,7.51341152466508,7.40653176590582,7.59752517958336,7.53942957404122,12.4490943068656,12.4005909743956,12.3449267042147,12.3885819343253,10.7338422745309,10.7453855603597,10.6899566501734,11.6456387890576,11.5809350288387,11.6840175166145,11.5995732206511,11.2304345074108,11.3006426066917,11.2602240397475,11.2918924704158,11.5419402589143,11.5981513767983,11.6929151294741\nCD9 - mRNA,12.0071559499039,12.0053961031427,11.9589802639181,12.0455890747555,11.8001291749631,11.8053431998345,11.75016326417,11.6831317462441,11.7473629369688,11.6842576855861,11.774289721717,11.6692712349473,11.6575741120247,11.6598882822763,11.6618208934145,11.6829773039993,8.29796241353278,8.18983779547908,8.47059868639059,8.5389885385068,8.68572707353238,8.53830242699009,9.03294417886023,9.00445520915174,12.2556145774444,11.4477507929266,11.6242629223776,11.6154478322411,11.3070276074757,11.3273393112086,11.3431213521145,11.9687330751336,11.8069289951145,11.9115076610997,11.8310223483411,11.6624956722481,11.724242119474,11.703214159121,11.634358110506,11.3076835022848,11.7941914766677,11.3151105363942\nDICER1 - mRNA,9.59111457935796,9.54863716174719,9.50570243896508,9.43624241540742,9.38675363016513,9.35116730664875,9.35481226279375,9.3399963384908,9.20690806515709,9.14751468930391,9.17141109394165,9.17943067869664,9.13670919332722,9.07143800044491,8.99643012820378,9.0464707619655,10.8086282739048,10.9029396613856,10.9028475814431,10.901328614696,11.0262610473007,10.9991393007703,11.0264243307726,10.9864774682227,9.9601586939182,10.1099784199745,9.95588441346877,9.95248281951866,9.76860769269161,9.72723821364949,9.92681443344811,10.1878978402513,10.1764305506453,10.1135106177978,10.1250966930414,10.1513330981544,10.1717858550892,10.2338436350006,10.1925119837678,10.1491796799223,9.99800247226246,10.0317170429258\nRAF1 - mRNA,9.86103872656461,9.82028469788569,9.82136628334003,9.92831968935247,9.3816826406177,9.42221076968664,9.43439712852034,9.32990752086721,9.11079294401137,9.11571050465395,9.09348662240457,9.05873133174096,9.36547023251973,9.33472181332244,9.29300490501529,9.26953436035098,9.75662684521188,9.77555504120353,9.99536462489794,9.93002404769644,10.138018778185,10.0523231605863,10.0059498557906,9.9050790463579,10.2556145774444,10.2000421183217,9.93860642203693,9.95248281951866,10.2831808655214,9.70885968433463,9.65971036330527,10.2469197061999,10.1504976467502,10.224920391637,10.2613600421762,10.3218657943902,10.1887340093621,10.2425387305602,10.2110142577637,10.052964364663,10.2184307067184,10.2103143463826\nSOS1 - mRNA,9.59617963444082,9.62190325039415,9.39467112657634,9.44879858727451,9.52195436804564,9.48491869553757,9.47260156609733,9.44070445829957,9.82963084898952,9.72435200184462,9.9170600550203,9.96146778199705,9.82411896301919,9.81943593451751,9.77045435871135,9.75783412914282,10.3400537470218,10.4713439008974,10.8542646967063,10.7870249880354,11.3801276285666,11.3150352124826,11.2076797717584,11.2162807346522,10.6706520767232,10.769678887814,10.7133141101947,10.4992259980093,10.3304865802997,9.41214215224455,9.36202981466459,10.6027751023954,10.697158866904,10.617713253558,10.5734281139372,10.3929913790166,10.3944826745786,10.4325007859894,10.3775137036765,10.6932342370632,11.1966462908263,10.7209295056437\nEP300 - mRNA,9.50755762104165,9.52336429789949,9.47307472940317,9.44127800530471,9.83834073381081,9.87341440736392,9.79785256849797,9.82074784422382,10.206181449659,10.0674056176015,9.90190024784929,9.92700437242883,9.88200993493105,9.99064751025408,9.92969460240198,9.83331682522268,10.2216287218921,10.1858688932761,10.6665696338517,10.5176148877142,11.0627680595121,10.9793400663423,11.1700533979457,11.1491818427365,10.1154369193961,10.4651538391881,10.2074231804647,9.80361943360418,10.0201464596876,10.2436364278876,9.9063503308884,10.5770683078922,10.6403205354567,10.5642966531265,10.4258155745655,10.6341288914233,10.4843068950408,10.5246849610511,10.5509380222696,10.5454806166195,10.5348448089332,10.5123428838322\nVAV2 - mRNA,8.62124155663854,8.60787636512187,8.64677597225187,8.60952642893964,9.53799721054136,9.4024565353456,9.49351853362535,9.50039486690479,9.56300647263368,9.69628123625795,9.63683244486297,9.6537876920147,10.0039539844754,10.0008990076288,9.93551360498184,9.91365893354028,9.92108449746418,9.88381702276204,10.3184771985389,10.1972615010504,10.8223617655542,10.7723257461648,10.6801502365226,10.6472986259534,10.6092515320591,10.619231260871,10.5969899922275,10.3885819343253,10.0201464596876,9.48954665593015,9.70780465150632,10.3218428252692,10.2569558874929,10.3371191692247,10.2113719063446,9.95449031006707,9.90896453470602,10.0162628747366,9.99152972536339,10.2039721527626,10.1124223828994,10.2872763281975\nTBP - mRNA,9.04082688758132,9.07901896628586,8.85261721732447,8.86882811589711,9.7846046186273,9.81782549816201,9.7755539013892,9.7925568771093,9.74110611806802,9.74245141769087,9.88657944856554,9.98221528529253,9.93020288195685,9.8830625173349,9.96100503264562,9.95763663060088,10.4739678375051,10.543573217489,10.6269637533423,10.5732487228886,10.7456140648274,10.7548440465867,10.5361246349192,10.5869341578034,10.3336170894456,9.93617594824407,9.90341699357463,9.68944841368487,9.76860769269161,9.55274048241735,9.65971036330527,9.85567851998048,9.81116544704994,9.90692871130883,9.93595377605493,9.8189235193987,9.92675228145962,9.85730444220081,9.84403839689526,9.88428253007129,9.82291576570437,10.1290142442807\nDNMT3A - mRNA,7.81181468960196,7.86814433666428,7.83024940429602,7.87628392081615,8.68314663294535,8.57943629432186,8.54293808766962,8.50376433914306,8.97747631696092,9.07268607593373,9.34211631012414,9.38268385620502,9.07185804897725,9.00403871515932,9.06021463657351,9.02437215640268,10.7185259744442,10.7419669729324,10.6928388845625,10.7396321250471,10.9177778542551,10.8513660083302,10.9253060007162,10.8764859686718,10.6504741947856,10.568605187801,10.4943043282529,10.6723749003259,9.07672998805394,9.0673136552471,9.39177715805864,10.1776066148678,10.2768036504328,10.2010736496826,10.1974327153867,10.0234946636479,10.1378258385156,10.1463234153009,10.1666775876851,10.1676760235397,10.2065890940739,10.5518712480189\nCSNK2A2 - mRNA,8.41313336130834,8.44663111937197,8.36940821540405,8.48825937317503,8.30875111816385,8.32061824328914,8.34394217757484,8.32737425412169,9.11079294401137,9.23892517467437,9.30546242779568,9.30916993188877,9.02123197590728,9.08336116782073,9.10958071389449,9.08052737654926,9.18389064682398,9.08292259156257,9.47059868639059,9.38629532544041,10.086120234159,9.89213148480625,10.2412249711747,10.3231041115022,9.45492338566777,10.4265861829286,10.2914874452532,10.1255475284915,9.62421778335643,9.93915730375642,9.68395790955195,10.0669625511838,9.9599278518851,9.96912070846264,9.88468349889259,9.78470780406078,9.66510623542412,9.6873794407585,9.67374289414351,9.9391444651058,9.92761314437106,9.57507863918021\nBRAF - mRNA,8.65237417389343,8.63090733377721,8.65951910278207,8.45379061661826,9.0774255720574,8.97677179186725,9.00677956899231,8.91824080592761,9.44108271650874,9.45882307713017,9.4491206545789,9.54436486355242,9.32430104354333,9.31083350652153,9.32121850717994,9.28023094348249,8.80046275406196,8.80690298519117,9.24873891968116,9.11911325491688,9.71598128923497,9.61781579710919,9.78601016377811,9.7396330768923,10.6905516341609,10.7668748149373,10.75339154957,10.2912847329704,10.1836451919705,9.08166894822418,9.2689204102731,10.172433341585,10.13647376922,10.1962566333849,10.0975635092852,9.78982828872939,9.83486302197196,9.81005831706501,9.80681323872754,10.1304431173407,10.0341848085691,10.0204897875026\nEPB41L1 - mRNA,8.62124155663854,8.52599935806746,8.7058890659201,8.58698386028856,9.94532203605005,9.8580255143505,9.90855603290419,9.93331262654183,10.2673699958492,10.1589081940551,10.023616156249,9.93412099709148,9.9134757955514,10.052608510489,9.98925450944979,9.83694396739947,10.4902119616465,10.3559831378539,10.3906414231572,10.4520754000792,10.1239846690653,9.98987961056488,9.73093544382553,9.66718512123959,9.9601586939182,9.99006895623927,9.05542008701968,9.00797793211036,11.68007101809,11.6666314489453,11.3084487744597,9.74154547021661,9.54267661112404,9.53110080728712,9.4372830978178,9.78214073035699,9.1394612383214,9.39645684567384,9.36310926895846,10.0228023941527,10.3534304824031,9.85337980166732\nDVL3 - mRNA,9.92214700879619,9.90774753913603,10.0122889820831,9.96636905004847,9.86129214119263,9.86477836204312,9.77735053305462,9.7916382555941,10.1293090703573,10.1047059124651,10.0329238984752,10.0220941409485,9.96389880084133,10.0736811131656,10.068560515988,10.039404486414,9.72716724473822,9.75427148283556,10.0795127291224,10.0561467237481,10.3409272390656,10.2554697213024,10.3698285763815,10.4049029026183,9.71018044090985,10.0280368064383,9.67209144746818,9.50887616804305,9.46375311116319,9.81577488825114,9.46356784112665,9.94105146220864,9.87660664246484,9.72417089481895,9.81674061772755,9.75817622131166,9.69284401709953,9.86265767669502,9.82073488480508,9.94987087169886,9.85361256292728,9.59034539583352\nGSK3B - mRNA,9.61210265740631,9.68142085018311,9.5376085363769,9.65899510520394,10.1183942168167,10.1922778276185,10.1851880421387,10.1073041637835,10.5703657334537,10.560047819625,10.3778619714091,10.4793652274847,10.418596387205,10.4118052439961,10.4545597406993,10.3984722395272,9.72010096918677,9.76495251024389,9.79975408847898,9.76019769544155,9.72994746839898,9.56348863471983,9.94195341118215,9.87933031976534,9.85827907989898,9.88537157757258,9.39645700485475,9.44998247898948,10.4637531111632,9.55274048241735,9.5319548161069,9.5403918452545,9.53200310960958,9.61129413266372,9.56178855819161,9.50074755937244,9.52199058069314,9.54295424692217,9.60405373372079,9.7310362697756,9.52534294116609,9.76195677390372\nGNAI3 - mRNA,10.1343232290922,10.1450724469123,10.1176319872865,10.1757280015521,10.7535359608368,10.7667199621698,10.7633677610155,10.7736072224277,9.90946964038051,9.88733710626677,10.1846950287329,10.1633708278005,10.4919744720887,10.4485675111476,10.4394284720781,10.4603329495371,10.4551763632255,10.4971976879061,10.393454243391,10.3607602333333,10.357815166496,10.4149791888579,10.1608222281637,10.1182134998333,10.0553159269585,9.94115935575833,9.97295792682771,9.8346463292248,9.83572188855014,9.80716175794643,9.77706731394343,9.73366426374723,9.69157474863232,9.7167150898999,9.65540160896311,9.48188853212113,9.68465321034996,9.6833514389584,9.65966770893179,9.48776511876324,9.90824781950413,9.57507863918021\nDROSHA - mRNA,9.75255810404518,9.80734564217819,9.70149462133787,9.76621903906832,10.9909914602732,10.9384179839652,10.9808957396798,11.0087701604698,10.4466303650315,10.3776482209322,10.3996115547995,10.4044824464836,10.8155915759973,10.8633169035775,10.8491910293163,10.8232949058876,10.2905612501509,10.353963263473,10.3633555972361,10.3920362603574,10.2293376692115,10.1767577452884,9.87981630223338,9.88581071265312,10.0242890313379,9.60045405384034,9.79238568118589,9.67237490032593,9.89985222596986,10.0381673095876,10.0344551566361,10.1981161745379,10.3182889139616,10.1849539843194,10.2005419731438,10.3245145620392,10.3319701037309,10.3832739323276,10.3334394766108,10.1209827880916,10.0607382276104,10.2966141927768\nEPB41 - mRNA,6.67489161045486,6.85423202980903,6.73713999990454,6.53375892132168,10.3899139710585,10.3783887753838,10.3409081174852,10.3500150943158,10.405443366894,10.3549184697392,10.5739271103152,10.5941354147308,10.2987659514362,10.2961802624504,10.334492027256,10.3235347277033,10.4874402123965,10.515836830634,10.4442547246733,10.3759037195938,10.4763334208664,10.3827354528464,10.5447681200494,10.5824529616663,9.20116679342199,9.17910278062929,9.29148744525321,9.44998247898948,7.961252770634,8.91531056180206,8.94699231538574,9.57510945485417,9.6835597555682,9.64860386909436,9.63486197065422,9.72914982757047,9.61739677194352,9.61943499031048,9.63247189814371,8.89542308767649,8.8886190127552,9.51234288383225\nGUSB - mRNA,7.80599736300664,7.69475381580183,7.84944296952933,7.84242511680012,9.63494754468802,9.61534625442348,9.6827440910266,9.652847678259,9.4968108638844,9.64555043413702,9.8968113711765,9.80464076942365,9.99920566957728,10.0594154721957,10.0776103838706,10.0051572880258,9.13631858717427,9.15313769144326,9.56044128299035,9.56259773555639,9.73918370379252,9.52721324832828,9.37838720366237,9.28867349798586,10.6706520767232,10.042021035874,10.0229986093273,9.61990748043179,8.37629026991285,9.00841996619354,8.65971036330527,10.5032737668717,10.3656447231875,10.3477771394226,10.2398604633043,9.80444924926852,9.85954788615493,9.90823871100392,9.84463100115124,9.87305527464803,9.66457873853727,9.84066874991776\nPAK4 - mRNA,7.53066968098142,7.51188975865202,7.42803594537476,7.6250992328601,10.4279228191415,10.4487781032752,10.3957383196797,10.4453847382871,10.0793946418228,10.1432188200031,10.532419476875,10.5172418151907,10.4456073397655,10.5604947881915,10.4140500771266,10.4270293592884,9.14220474401706,9.14147477391768,9.45560394816073,9.54105691884803,9.61624511606024,9.40410910363896,9.16280531444282,9.01336076237363,9.3080819973385,9.45473722324272,9.67209144746818,9.50887616804305,9.69821836480021,9.38924175013447,9.57148318029354,9.84108114376752,9.73797777151207,9.74923623220085,9.77979903933941,9.71979032737945,9.70320187214099,9.7541888783741,9.75479628095992,9.53126675739983,9.69314789073404,9.81490565385268\nPIK3R1 - mRNA,7.59280490766016,7.78258398870047,7.86524482565222,7.58015182680613,10.3040681252803,10.2211326902906,10.3139296085084,10.2552407214292,10.1945055535303,10.2753671706424,10.5438586894811,10.5750399037867,10.0920673372049,10.0751745864262,10.0662891509832,10.1078060366479,8.93674761376008,8.93538416491034,9.2806908824575,9.25132062800122,9.48514379964227,9.53830242699009,9.3203465914293,9.31532353444785,10.1731524172524,9.86977472252156,10.2074231804647,10.1255475284915,8.83572188855014,8.57320458497706,8.79943512697188,9.49966380420662,9.61946595085992,9.66496731530325,9.5936726192114,9.3354990451396,9.60403475014167,9.46688522918432,9.40629963769148,9.6279427768115,9.91796297433734,9.77537728966604\nPPP2R2D - mRNA,8.77506550345475,8.92248955821259,8.83667567345545,8.84812372213257,9.72077986757036,9.84149293323183,9.77105250915426,9.73063718346217,9.61648700688026,9.68450406848159,9.64089637724287,9.70553645932068,9.62275356850185,9.69285154666958,9.64213026173581,9.66089286316032,9.86879458541904,9.87257336105189,9.98719490141885,10.0023663134121,10.1033896079329,10.114400414164,9.96789362952848,9.9177831023914,9.82265517016826,9.5556661320935,9.17827683480522,9.34614666800494,8.18364519197045,8.84972222017448,8.98652067957238,9.6147437710139,9.59146804992418,9.52171877868654,9.48906000772596,9.51704937170154,9.43055192928617,9.49703103887697,9.49654610252765,9.24809470599495,9.35700593662697,9.02048978750257\nXPO5 - mRNA,8.39967708082891,8.16718532100452,8.09703594499092,8.36117946643339,9.81689802183418,9.76517394126001,9.7880835896821,9.85016433220055,9.33606654543308,9.33747634832978,9.26785296662866,9.41129834124412,9.54017034129844,9.56262580045618,9.61337809221749,9.60986476737815,10.2199599015315,10.2139815334029,10.1282825945032,10.050349849888,10.2028775961794,10.1538686973494,10.2849743614654,10.3153235344478,9.89304449805966,9.13628969709032,9.17827683480522,9.40934049449213,9.37629026991285,9.56300881787118,9.47750703208452,9.3757280454159,9.393427879701,9.59186360271482,9.56178855819161,9.68997311677369,9.72877576358226,9.69740051629555,9.6883425085695,9.21290527753266,9.46697877865211,9.6204026296897\nCDK4 - mRNA,8.69863369932871,8.83168946115794,8.78111769696514,8.67938310220402,10.2549967708484,10.2843802154375,10.2845432547424,10.2539073617771,9.45827247780625,9.56112165297631,9.4491206545789,9.39877783232532,9.82769441724302,9.86245223765961,9.84566365582094,9.8205495291488,10.2735003560909,10.2095201613408,9.79124760722558,9.82868331286421,9.53796408612515,9.51828012355827,9.76962062681024,9.88095314900382,9.78612929414315,9.10997841997447,9.69285000763497,9.25733740104708,9.07672998805394,9.6132820243524,9.25280074490983,9.48513293994766,9.57162798482017,9.55387401683356,9.54394724543143,9.70969444964394,9.86746392818778,9.72900806164426,9.76171569489972,9.08250864027698,9.43350407410102,9.37804179212066\nDGCR8 - mRNA,8.5672396484795,8.50121625713757,8.58894852040951,8.64050573363708,10.3463285759803,10.3484166956327,10.31454839258,10.280343191516,10.0448614669081,10.113516291241,9.95672427259382,9.92541812153688,10.2442933956517,10.3397009210805,10.2968848581586,10.305976356319,9.34158934593303,9.28937346902999,9.09963238974641,9.1614080533949,9.00497748893272,9.01549972963776,9.42429755546821,9.28131904684671,9.67065207672321,9.2848122218004,9.2640067088311,9.00797793211036,9.69821836480021,9.34232070274698,9.09899540883079,9.29353064849149,9.41779007699233,9.39572108053619,9.26680885395789,9.25540373065417,9.23127610411145,9.35654112269129,9.23787916766187,9.53126675739983,9.11664696876108,9.14977280444754\nSYMPK - mRNA,8.60792939721863,8.69162771481511,8.53859432052246,8.54549775777844,9.94446252155285,9.89713567575404,9.87957696593242,9.94161832041397,9.92014314189497,9.90520616151584,9.97615480254272,10.0909276973594,9.99603142178214,9.99538802396366,9.9053341073538,9.96096464831864,9.28525136178322,9.25947198331613,9.42953043655084,9.3335722218875,9.5177116620987,9.52832600969536,9.66810067368457,9.6348491391199,9.40761767088942,9.69788820474596,9.60796111004846,9.34614666800494,8.54621527135516,8.74538556035974,8.70780465150632,9.50275861062949,9.50950974718884,9.49058026687763,9.48990740571854,9.31611006003141,9.29429464060925,9.38327393232756,9.4014745661675,9.31599877049676,9.55677544900275,9.6643459772773\nASH1L - mRNA,9.09192260879469,9.13417547370764,9.01866163311622,8.97159937012478,9.92629340833447,9.88009577245557,9.82329449033539,9.84042532900028,10.1513702206062,10.1324228476415,10.1206595898523,10.2051525516224,10.0883461717658,10.0751745864262,10.0999897629331,10.0942311791821,8.86595183565022,8.93267995598196,9.5053387351392,9.31918112958187,9.86433092099608,9.86420185191781,9.90688753327168,9.93819153360723,9.58818991653124,9.64389331510523,9.37092191274761,9.48951084317612,9.76860769269161,8.93125210567108,8.9063503308884,9.53536327606142,9.55327172636323,9.51743396698062,9.57302830929537,9.39635234743445,9.37517904421197,9.42488125312114,9.45135047280707,9.58730079231415,9.37121979584667,9.42988072364028\nPIK3CA - mRNA,9.69391643613662,9.73478847760409,9.57075298829683,9.67297585981088,9.17591543545676,9.18072696494702,9.09009232471679,9.06839094267567,9.00614565979219,9.11131873553679,9.31439555256568,9.42033639768441,9.33061481747553,9.23936619182327,9.33763447032137,9.23833240516646,9.0183816988191,9.02443535272746,9.19041465762326,9.07268415174473,9.31026958222822,9.3383797843359,9.32153058414318,9.37987128815717,8.74865458872448,8.68011350301799,9.08712894674702,8.86501997826832,9.37629026991285,7.88288908410968,7.47750703208452,8.30184591670349,8.26825432146811,8.38353053966941,8.31446539930866,8.04774220989458,8.12632583613227,8.21768786660959,8.11456119702475,8.57349499278912,8.23795326498321,8.44675454220467\nLIMK1 - mRNA,10.4401446996326,10.3363598958807,10.2151872967092,10.3451493566144,8.92716381260877,8.97320517684706,9.05200882298531,8.87379407858545,8.99103876740198,9.10985183653813,9.3533048671651,9.42033639768441,9.27014527349642,9.43931943434555,9.15446496236957,9.09877033879727,9.04124178163119,9.12498665112911,8.93486540252526,8.96895504029215,8.31768705379071,8.28012038636351,8.83823249829989,8.68772869872285,8.74865458872448,8.00917777918698,8.77302035631896,7.80361943360418,8.69821836480021,9.0673136552471,9.09899540883079,8.97487498421635,8.82653287404428,8.84261967877017,8.8671803562747,8.73380690576075,8.71860054227329,8.90938872704001,8.76860208048496,8.89542308767649,8.49321032022528,8.87847078263015\nITGAV - mRNA,10.9501096708322,10.9387314430248,10.999100572387,10.8762839208162,9.25291546001723,9.21283915010077,9.22654412978272,9.20713513874592,9.55503890580582,9.56433836565985,9.50146391293065,9.45373360756452,9.33940793593354,9.47029697921991,9.41939890111884,9.41494641561982,10.6080303532101,10.6646293616137,10.33904969072,10.309506692384,10.0568704732067,10.0783874355932,10.3542939233526,10.3764239865387,8.5007270752809,7.97070363137234,8.69285000763497,8.60198557243453,9.69821836480021,9.1099579926556,9.15210674529035,8.33694971765941,8.33731600012897,8.30217584482766,8.2418739947419,8.45957015269753,8.45352245193581,8.46454001863122,8.50631693202499,8.60097572921123,8.50614937593278,8.90313283686442\nPOLR2A - mRNA,10.1244237691825,10.1249348013412,10.0645536338756,10.0528041110738,9.58397894690846,9.57002226391399,9.50223575072646,9.50600628992973,9.53897053717817,9.57075035417409,9.63581467054147,9.6335288530309,9.75068351641618,9.75753163525104,9.77602461675429,9.65062452770649,10.7255999495913,10.5738268584987,10.4849117804955,10.4969857162134,10.2971965280029,10.1866586324671,10.4905347863487,10.3361763965328,9.14458326505562,8.80564438510185,8.73349199213232,8.36752031879751,8.54621527135516,8.91531056180206,8.8431565044012,9.35304369377971,9.42261353531654,9.46588806196173,9.33252903283685,9.5819514101511,9.1834590673214,9.22970654267446,9.18270246050974,9.21290527753266,8.53168446803992,8.77537728966604\nAKT1 - mRNA,9.69863369932871,9.65998839764116,9.76351852672783,9.72345430483868,9.66447742150128,9.74753559448352,9.5680654869904,9.57169419299697,9.59333282562786,9.66665606714867,9.738943271127,9.56794051140332,9.82322371319779,9.85725333098529,9.87191110581635,9.83694396739947,10.0029378934545,9.90466557981236,9.99832400037849,10.0254499265252,9.68954372930567,9.6921822330765,9.70848408751344,9.65869588107427,8.5007270752809,8.2848122218004,8.37092191274761,8.40934049449213,7.37629026991285,9.1099579926556,9.02499482738702,9.04872697973478,9.00436218966322,8.86572503903297,8.86848419282155,9.38455454186792,8.99036095277899,9.08046282853582,9.03267701650514,9.002338291593,8.44026144027633,8.74841024206577\nEGFR - mRNA,10.1085601330271,10.063383102081,9.88242926862107,9.85757166055398,10.2946562882781,10.2764433884935,10.2876980669352,10.2452103562919,10.5389705371782,10.5976877822077,10.1622510946562,10.1272589839323,10.2961874073344,10.3165271141604,10.1959185117159,10.1579661313052,8.06001898866181,8.10350692371866,8.35529904236505,8.36076023333328,9.00650819999504,9.0836956493536,9.08685646120952,8.8289361912362,10.1154369193961,10.5653812869267,10.3050339770912,10.0744733438973,8.18364519197045,9.0673136552471,8.98652067957238,9.72131838732268,9.42933939850485,9.57134127884147,9.35409718620159,8.51627725766605,8.62830861789684,8.71131410099443,8.6447734936321,9.36490837097771,9.53168446803992,9.57507863918021\nPXN - mRNA,10.0647751312799,10.0166819106892,9.86367240583347,9.99233306888417,9.72978735806414,9.72226145833999,9.73729838237919,9.73637735697994,10.0144705244431,9.98258542141459,9.96970699274468,9.96456037705243,9.97280685353272,9.99301971419917,9.93634297841135,9.89638094210845,8.87163174472257,8.80098422800389,8.89254540334412,8.79960607999979,8.93757154483439,8.94057999462261,8.99337565894216,8.83061667487512,9.54512119463935,9.390606885823,9.37092191274761,8.98049719568826,8.961252770634,9.6132820243524,9.67188507725137,9.77350111159855,9.78218424763154,9.64860386909436,9.71096797807031,9.80191709409301,9.50517576283247,9.67458532854245,9.59914541049973,9.34878870561443,9.3852949810584,9.51234288383225\nCTTN - mRNA,9.84055458504301,9.86467881724415,9.96467572451695,9.93547504210269,10.5572409762027,10.510365901498,10.4962483263809,10.455280204032,10.2722380965329,10.2727476600891,10.5503548882586,10.5841166589058,10.418596387205,10.2865429851913,10.4429344155796,10.3749629665104,9.16435564847655,9.14147477391768,9.04056644888189,9.09959824183225,8.97088429026432,9.03911228685971,9.2070392856032,9.27269119944401,9.926991829983,9.61299930715945,9.47045758629853,9.50887616804305,9.28318086552137,9.86640096132111,9.94699231538574,9.46411734560237,9.58803714452391,9.49145436265771,9.57542547690459,9.8388230768364,9.80081955844499,9.81865312145204,9.71839974242568,9.54548061661953,9.89355130829743,9.91530755081052\nPDCD4 - mRNA,9.31619209532583,9.32883408308768,9.4792488973467,9.45004821493245,9.13129244403613,9.0309556843504,9.03708940314957,9.05160635838011,9.38182872043414,9.48395379245385,10.0290529676826,10.0673875279549,9.80610755503487,9.68504790335151,9.78618123179046,9.80211228089212,8.49297839593792,8.62120255436124,9.39289211790528,9.31918112958187,9.86854563343244,9.92545550506415,9.37952452736407,9.52081389587387,10.5230948883094,10.5750314569604,10.4339317102734,10.3989985502707,8.18364519197045,6.94701942152939,7.02499482738702,9.85970727166554,9.83379355892587,10.0001609557567,9.87693052882778,9.01313317296444,9.36216495430592,9.23338454349607,9.30917046184766,9.44291088297898,9.96088929924536,9.91530755081052\nCCNG2 - mRNA,9.78692740000806,9.66158695186663,9.8358739534899,9.76320610019208,9.98431616766592,9.89619427602022,9.96403567912825,9.94492725544455,10.2170424422908,10.2328733105831,10.9945333318576,10.9246243415769,11.2379253505171,11.2865429851913,11.2291247276392,11.2342123784514,8.25297495875947,8.32589934505511,9.87574643568397,9.72059534499674,9.81189476650316,9.73329301452912,9.56989406060617,9.44165435835474,10.6300100922259,10.7808409941013,10.3837459531052,9.58383822572427,7.69821836480021,8.0673136552471,7.86453015519377,10.6621110027222,10.4838936898575,10.5165754754106,10.3259002943709,9.17940485505545,9.32815092204087,9.44643228517871,9.39420656570346,9.94987087169886,10.0694818606824,9.86597983844695\nYBX3 - mRNA,9.52802172360137,9.48505646731375,9.59937882903527,9.61287763843928,10.9508964330972,10.9292434217847,10.9171374477704,10.9178198883343,10.8029664192494,10.8579286351108,10.9337195362223,10.853003790985,10.9545266492709,11.0392800993566,10.9880550954628,10.9387721233076,9.91490886122415,9.82232517117461,9.86362627164537,9.84300507467858,10.1239846690653,10.0631125968637,10.481559161535,10.5844463230071,9.85827907989898,9.54915282513482,9.93860642203693,9.61990748043179,10.2831808655214,10.870540691682,10.5518543735446,10.4741379326897,10.3073770680083,10.2991837351325,10.3277973295435,10.4022152671353,10.4385447200397,10.5080803880638,10.4127080659561,10.2393774888938,9.93719919320026,10.2778776301952\nSPRY2 - mRNA,10.0829300903331,10.1496361360983,9.43334583374241,9.62177632534682,10.4266923755353,10.4679122702319,10.4935185336253,10.5065662336747,11.0841481717935,11.1417840162638,11.2241669638017,11.2149247387319,11.0313452745504,11.0262135125957,11.0383639153519,11.063212519603,9.41051623574595,9.40216523928318,10.0598169277815,9.86463758555513,10.0933409451326,10.032853296388,9.73271654904051,9.72705657721699,9.71018044090985,9.68606276934917,10.0393004216564,9.77191057387684,10.1836451919705,10.6522761559683,10.4775070320845,10.1750222970413,10.1832679806989,10.1924989139446,10.1969138536509,10.2710503374128,10.3626317703724,10.3869938184647,10.3304809680931,9.90647827618548,9.96088929924536,10.2003988775175\nEXOC4 - mRNA,10.0708522044648,10.03097536637,9.93802693118907,9.89659090413651,11.1301589923286,11.1081905531562,11.1243758003172,11.1006438516887,11.1615261134396,11.1489438078367,11.4316332278501,11.5024231004791,11.4789592883323,11.4685933438625,11.5467823851483,11.4390119451233,10.1143285768853,10.0439869290156,10.5428134405579,10.4215263367196,10.8284326283064,10.7951558873629,10.5573857887301,10.471042599061,11.1445832650556,10.8513626854077,11.0147781025223,10.7957573158312,10.6982183648002,10.0084199661935,9.86453015519377,10.7345420824788,10.7689881994211,10.7828265939279,10.7080578509295,10.4397542680747,10.4243045546698,10.3980962720331,10.4505721102088,10.6739551745683,10.6587963859433,10.7278489195835\nPRKCI - mRNA,10.2177522782962,10.2444176531,10.1656397589899,10.1628297743327,11.1545199238267,11.1227510976845,11.20200678054,11.1648386872772,11.1742141495372,11.1381907515398,11.258295386,11.2371527005553,11.3444086169919,11.3615907197774,11.3161296305071,11.2858814093382,10.1689759728127,10.1473180181992,10.1858788958426,10.1847477472791,10.039031667369,10.0783874355932,9.93113507951836,9.85311367483367,10.5881899165312,10.7037647710826,10.6403825877408,10.5746782264388,10.5857436355418,10.5319819222505,10.3695243512115,10.457752504216,10.477417971844,10.516146037979,10.449095981437,10.2919527414367,10.288901381839,10.3832739323276,10.2897180991815,10.396617230705,10.4436282913747,10.3151105363942\nSMAD2 - mRNA,10.1395367482643,10.177278059693,10.1528214157488,10.152888025194,10.6053444813417,10.5776758364574,10.6379604872974,10.6568888434809,10.5765636127184,10.5797854972413,10.600776068774,10.5500376663753,10.6015027813037,10.7068889518911,10.5961625791882,10.5536627978194,10.7400328715086,10.7920603126996,10.8518125232419,10.7521851753902,10.9149278842903,10.9305146395562,11.0473280970229,11.0029655910113,8.58818991653124,9.40510645551812,9.58593480371846,9.28005747754717,8.37629026991285,8.96261627658041,9.08084806212053,9.21015718701164,9.21808753477361,9.18279102283343,9.20416099144946,8.99883260941363,9.04373430365479,9.16088151099111,9.08276497800459,9.38084991484673,9.34984614505411,9.10795262875291\nPRKCA - mRNA,10.6576328466249,10.6439046743061,10.5257262863052,10.4955395990912,9.29936988186954,9.29227361759518,9.46146960468672,9.31719646911766,10.1856861982555,10.1994713664445,10.6878093081797,10.9451219929735,10.3574535264754,10.3783222151078,10.293652288935,10.2342123784514,10.5275683251018,10.5803206205842,10.7903941939851,10.6882982965632,10.8664398163298,10.8357208274533,10.9260845733421,10.8825741548383,9.35870807040847,9.30814472563103,8.98983174539211,8.63760948216525,9.18364519197045,9.00841996619354,8.77706731394343,9.49449099464589,9.62784399438671,9.56304990521008,9.49666873052941,9.76793959067989,9.68016581935764,9.70138953738316,9.68636033050514,9.64123959942496,9.44026144027633,9.32427053567968\nMAPK3 - mRNA,10.6836424756912,10.647539177381,10.7409991953602,10.8135894791676,9.65924836524799,9.74230653823022,9.6527171284746,9.67194318920315,10.332075685237,10.401824656672,10.7960917796516,10.7348121888681,10.2111420861284,10.1550221529482,10.2144986712698,10.1182766531488,10.4688245342291,10.4838669713648,10.7378002995542,10.8244438298226,10.7697012008655,10.7614836804534,10.1634657379109,10.1278132170838,10.3587080704085,10.0604582841307,9.77302035631896,10.0614171910718,8.54621527135516,9.25634747963712,9.30062927000044,10.1528666532204,10.1193443331135,10.0667725579711,10.0321690913142,9.88561071817213,9.91279470078108,10.0044725821844,9.93254684617404,9.71854532559241,10.1250592085773,9.95122712506546\nSMAD3 - mRNA,11.7304619994597,11.7334590672176,11.5334114228331,11.5111885097464,11.0632427649151,11.0206260814567,11.0040969104198,10.9876678559505,11.65533529387,11.5378562371989,11.3766439932524,11.4828744509441,11.1381497276864,11.1995232630323,11.1774467441756,11.1382578733403,9.91559634993903,9.93875731360829,10.6450154908513,10.5641241279127,11.3965654304597,11.3522701721193,11.2121552308255,11.1675816810685,10.6092515320591,10.5846175057896,9.98983174539211,9.78785211774586,9.62421778335643,9.78100947009047,9.8855917707216,9.8844375129619,9.95460915446612,9.99893208426763,9.83885902371227,9.54986532593713,9.56156412581525,9.60175901520522,9.58786328034735,10.1209827880916,10.0297113319898,9.86597983844695\nADAM9 - mRNA,11.9705691299548,12.0379154916611,12.065921765833,12.191877040861,10.2403641431836,10.2749956261633,10.264827917615,10.2920830893621,10.5331881845842,10.4367613146004,10.1023556772272,10.1043089042367,10.1046479840949,10.0063890198457,10.1147187835768,10.1070552195397,11.1918582136518,11.2497276645697,10.8869674536802,10.9750876533135,10.4933549489615,10.5305489608915,11.1461957015455,11.1154589499924,10.0242890313379,9.99486994250309,9.69285000763497,9.92391366732189,10.4206843892713,10.5681157083043,10.5584270274681,9.77179175708473,9.70351436194385,9.78032532635742,9.88597161955039,9.93671089750583,10.0035912626315,9.90535965389021,9.97916906642461,9.74341999423144,10.0781728209464,9.99776971100249\nTRIO - mRNA,11.4191003440923,11.4526452488226,11.2923134119795,11.3228130266363,9.88925397069431,9.85609030944478,9.94177878528581,9.93913162912169,10.3617414674449,10.4202866408178,10.309935904375,10.3763117737924,10.2234088160223,10.1272154574102,10.1868864400619,10.2092420332423,9.70824638284099,9.78458131699282,10.2230373941277,10.0150707807452,10.4038436975404,10.3925455716723,10.3704007338512,10.3838827565952,10.1154369193961,10.2571052926306,10.102725801798,10.0614171910718,10.5857436355418,9.754374343587,9.74285159860552,9.72043248306341,9.72788080672836,9.73970377793661,9.69341850623214,9.59297843230012,9.58175330276567,9.62434570027246,9.66303142004031,9.90647827618548,10.1500055657075,10.2778776301952\nG6PD - mRNA,11.0605058974368,11.0585377741314,11.2493066750272,11.2858498245171,11.2315832576414,11.2248867778653,11.1872166738874,11.1929417437211,10.3473143968097,10.182126903623,10.2750615419031,10.1545965151255,11.0224025196152,11.0400450490251,10.9924480802965,10.8920289175687,10.4584825900077,10.3834733952873,10.0527170362825,10.0748272990181,9.34698136058475,9.20277829783034,9.01987488428208,8.82556934013781,7.3080819973385,7.15356768852215,6.37092191274761,7.18694807315569,9.37629026991285,9.73634042075674,9.68395790955195,8.71688341598234,8.63201484158451,8.38353053966941,8.59524847413694,9.30363454648435,8.99999501222537,9.12371171840057,8.92247758826278,8.54548061661953,8.48015416739984,8.6643459772773\nVCL - mRNA,12.9420097289588,12.8605961660612,12.9575989453551,12.9818954757931,10.0481192025454,10.0748818773171,10.079937566566,10.0221467579064,9.46557727842949,9.44143401517392,9.74933260928223,9.74384034749731,9.97925106320164,10.0450073599187,9.99085217853986,9.97995318334612,9.94687183103379,9.99233133461187,9.86037695711666,9.66287984714046,9.18203195443314,9.24805917713121,8.93578144437473,8.83061667487512,9.08568957600206,9.30040907685142,9.17827683480522,8.92391366732189,10.8681433662425,11.041842967731,11.0438538546383,10.0187737576332,9.9051482231135,9.93803924384886,10.0575408810225,10.1063793970289,9.94557030966178,9.96961528724991,9.98240381002391,9.75569832400987,9.86871945531749,9.70699031468579\nACTN1 - mRNA,12.2237646550411,12.2371994844052,12.4655788501042,12.435296282436,9.79131261163971,9.77749596669253,9.73729838237919,9.71618602716463,9.64572085223904,9.62104547062837,9.61943186805427,9.66808607506727,9.59784310719552,9.646388698038,9.56164034387544,9.51343011977552,10.8339425807922,10.8775032488298,10.773219050177,10.8034883533451,10.738262736611,10.7167937995898,10.4810294405668,10.4256211769819,9.63001009222587,8.81653270124458,8.65102983194035,8.32445159690562,9.18364519197045,10.4841533971599,10.3239617738311,9.69269609620839,9.60680694854375,9.50190241103173,9.49582529581325,9.8784182061868,9.80219683305163,9.90535965389021,9.83988333733267,9.53126675739983,9.47358151347635,9.6204026296897\nPPP2CB - mRNA,11.838418840699,11.8168814799966,11.9309117433012,11.8758190601577,9.87218045733537,9.90370832967752,9.93294019153115,9.87466186510238,9.77482011368573,9.69921056729662,9.94608910363183,9.89250960337145,9.83126103226321,9.88135619538449,9.82431636067724,9.77306060336748,11.4241083487814,11.4331089464914,11.2390762398813,11.218418152647,11.1884522029434,11.1820702407018,11.2223330158295,11.2483807920378,10.9764605062473,11.0068029719991,11.133422599021,10.646379691793,10.6242177833564,11.7295191549007,11.5649698733349,11.1370199579784,11.0914464602218,10.9915367923415,10.9971340649008,11.2078974991031,11.0827905354155,11.1659423356677,11.0678695779611,11.0479809571487,11.0071334840146,10.8342709787196\nCASK - mRNA,10.6079293972186,10.6329459027767,10.6472330275514,10.6638499444714,10.0553027611593,10.0454638312795,10.025041775385,9.96462305678547,10.382472062715,10.3831233509983,10.3984118083346,10.4321135654608,10.3679279759436,10.2781382849561,10.4182119861684,10.3674586840982,10.4925176917855,10.5488805013685,10.7163936125414,10.640868152526,10.955116974406,10.965851106666,10.9599323438258,11.0342937037094,12.1154369193961,12.3110349336913,12.5060814960292,12.3701698972115,12.0345017526646,12.1496756248046,12.149932374056,11.8905543991366,11.9787179306656,12.0021556421187,12.037684986106,11.9022550977375,11.9899077738713,11.9487928093546,11.9609738948522,12.0554496280526,12.280936435534,12.1753078965547\nPPP2R2A - mRNA,10.283072820744,10.15758801866,10.1605260804738,10.1877630664725,10.1433275089565,10.1134654951969,10.1640567660526,10.071421815725,10.0334373514599,10.0605879334159,10.2658807242971,10.3036855513166,10.1779115565618,10.2077165365871,10.129298948509,10.147050353826,10.5253179814216,10.5288767062823,10.5935967698773,10.5615792427214,10.5117956629607,10.4329159516993,10.4921129396958,10.5389157022492,11.3587080704085,11.5393273952788,11.3964570048547,11.2166954165497,10.7686076926916,11.2211189913261,11.207519865609,11.5232227131892,11.4864297904626,11.4669994517716,11.4055254250086,11.0095712994371,11.1069530814143,11.2006447461658,11.1352951879251,11.3768810126438,11.285634241837,11.1954154700033\nPPP2R1B - mRNA,9.86103872656461,9.88466392602299,9.7794508047625,9.86976220142076,11.1187752269974,11.0823505482209,11.0911761071949,11.0872302681108,10.4117599051844,10.4599749295194,10.5924065309228,10.6638114199669,10.7781997885118,10.7828872095279,10.7662625057645,10.7506410194724,11.0813149059464,11.0496139157235,10.9916568484009,10.94845252681,11.0345372866598,10.9829970899504,10.4281498809638,10.3135220419671,11.5341500768183,11.2531033620731,11.4153160321061,11.1809242855279,11.4637531111632,11.3243242734531,11.2689204102731,11.361876257414,11.4353182509943,11.4538301783972,11.4095615654614,11.4872175984587,11.3903678705317,11.4131732957117,11.4117086235601,11.4580177753692,11.4948340627049,11.5401260403458\nDOCK1 - mRNA,10.4513683157143,10.4659717191763,10.5021695124886,10.4114390244068,10.8617474654953,10.8526974420029,10.8669769291528,10.8390921747016,10.7600433196932,10.7593912801126,10.5850431425094,10.7092285791627,10.7125147547465,10.7649538769127,10.7289350892731,10.7453431849596,10.0711691131785,10.0974831360909,10.3524107630402,10.270770170786,10.5144877612244,10.556415252445,10.4434571337517,10.4585984994282,11.4076176708894,11.4051064555181,11.5914729872599,11.4748143319694,11.3762902699128,10.9856998896935,11.3469229222744,11.4707176638257,11.5813664567848,11.576293326178,11.4985646577013,11.5203262540904,11.6634021528888,11.5375720889869,11.564491341472,11.5700228008823,11.830651414484,11.5979186155383\nFLNA - mRNA,13.9709753741612,13.9502627235777,14.1504052452948,14.0528041110738,11.2248738572788,11.233111545127,11.20200678054,11.2565728499088,11.0622313776309,10.7761345439472,10.3539239167523,10.4044824464836,10.8832990147257,10.9638894926313,10.8938520228391,10.8696270526127,11.925016247743,11.8960152658472,11.7725709877086,11.830587023333,11.1306678958839,11.2024297786739,11.422919228201,11.3456096629387,11.1300836953605,10.4265861829286,10.4704575862985,10.3675203187975,12.0201464596876,12.3978720552779,12.2808930519392,11.2806667194558,11.2332666757073,11.3036695758059,11.1459448588131,11.1567931581551,11.0381927612553,11.0929163643349,11.0432994038443,11.1445183025524,10.9608892992454,11.1342319535518\nRAB13 - mRNA,12.4985688378144,12.5217149378926,12.5221175246714,12.5305139254848,11.4517090720059,11.4713928389668,11.4634239088629,11.4117016120309,11.5421409800583,11.6013461305858,11.7280010963514,11.691139576201,11.5647492843303,11.5764011791008,11.5643254275481,11.5879689198567,11.0456450400248,11.0167952905253,10.8977056882601,10.8932553141603,10.8136445484531,10.7713861857012,10.8534557156965,10.8813585712464,11.8930444980597,11.7822302046476,12.1782768348052,12.2828724931542,11.2831808655214,11.0887933826576,11.1906952429676,11.9370532285894,11.8008287955781,11.8276425776662,11.8275070485966,11.6569017913384,11.7565707605567,11.7395535320057,11.7491101534468,12.0253401169546,12.0749198431514,12.1446110993318\nMYH9 - mRNA,13.0557644623429,13.0838706239952,13.1933672781755,13.1661602066023,11.5456688713768,11.5814874504074,11.5899541850399,11.5984679883336,11.2076343148784,11.1856125178841,12.0103265387348,12.0661283962391,11.7520953857582,11.8143029743592,11.7690584272615,11.7614172795257,11.7302967362927,11.751972366746,11.9171872376531,11.9073540795631,12.111414012384,12.0944416797401,11.7866533148583,11.7505482925515,11.4195903125555,11.2808865185866,11.0147781025223,11.014767097773,12.1576499834375,12.2064556743928,12.1906952429676,11.6388562271307,11.553491632627,11.5227879961378,11.5453745461324,11.7169369943924,11.5998831068712,11.7392302027706,11.6216212784315,11.8417220678432,11.6101864417186,11.8213896874229\nVAPA - mRNA,10.7177373582766,10.7655891770994,10.6279109301498,10.7342649159437,11.759901162247,11.8130949380901,11.7936422581944,11.7586423422816,11.4914532116086,11.5530483524406,11.8891442404248,11.8629821249939,11.8392538230167,11.7985675966707,11.8964115061376,11.8102075849273,11.480022740834,11.4810934459206,11.2459402863285,11.2510047670167,11.2326113149113,11.2402708737569,11.2207474622675,11.2278044305946,11.1589385580327,11.0115486836497,11.0554200870197,11.1929468137771,10.4206843892713,10.8071617579464,10.6839579095519,10.6123579675251,10.5471007024195,10.6264931674886,10.6535120277619,10.5255154645214,10.5861572594114,10.5399872637872,10.5523897909867,10.3159987704968,10.8762139918644,10.6715414786815\nMAP2K1 - mRNA,10.4972156768202,10.3889345434712,10.2814605161283,10.4082280392378,11.6743607144446,11.6637218483263,11.6994228321732,11.6182919891121,11.2292535262417,11.2605701687144,11.9819733426856,11.9427716882871,11.9892628663297,11.983706218916,12.0691277989264,11.9813877832635,10.5149209488186,10.4615280211846,10.8817685470059,10.8843221893903,11.1101686993668,11.0836956493536,10.4461735627991,10.3764239865387,10.7101804409098,10.4547372232427,10.5859348037185,10.5279849909908,10.9308791215905,11.1916417902493,11.0624695328057,10.6699330010558,10.5681495140792,10.6129015924856,10.6264094056234,10.4999666689388,10.3811816246529,10.4168169781222,10.3970373627676,10.5734949927891,10.7322174554126,10.3603397903872\nPTK2 - mRNA,10.0019175729819,10.0903389524145,10.0369149525533,10.0503344470626,12.2670782689989,12.2969176814126,12.2728100948929,12.2408421136321,12.3808631687812,12.3690898354218,12.5507599310877,12.546946175819,12.3763446773283,12.3516119433454,12.4219178578205,12.3855276554158,10.4298945665947,10.5040448533024,10.689638482272,10.6962085939794,10.9728431433024,10.9275656342116,10.7021345863581,10.6798260478383,11.1445832650556,11.298468666282,11.3117598402795,11.0548445371483,10.6242177833564,9.91531056180206,9.96689187282345,10.9269128994389,10.9253404652982,10.9681791039209,10.9086464300592,10.7357982035798,10.8206624874196,10.7669336170941,10.7664145212472,10.6544149881727,11.1271546265795,10.6787012702544\nPRKAA1 - mRNA,11.0258894456914,11.0362220970848,11.0473416062791,11.1104470320639,11.8606088851813,11.8378203940211,11.9085560329042,11.8748787301972,12.2884618987807,12.2953473909099,12.2483411488953,12.2701778832126,12.0718580489772,12.1210832277262,12.0981380170225,12.0458832257974,10.427486462145,10.5185444229975,10.5994554379932,10.4684698383299,10.8761012255449,10.8298662742242,11.0505448097064,10.9943703095475,11.0085217154796,10.8619126047486,10.9386064220369,10.8192162886552,10.3762902699128,9.76330746835701,9.75434723744335,10.5012120372734,10.4422228200252,10.5555467069234,10.4173774388352,10.2692183434639,10.3949391514187,10.3939942077372,10.4556239767935,10.4276441263257,10.4436282913747,10.3955292188495\nIQGAP1 - mRNA,11.3485680972805,11.3831328129679,11.3371944841359,11.3842419929086,12.5974377893061,12.5592088514148,12.6154130523755,12.5906991558506,12.4188331096169,12.3983741287504,12.7931277271619,12.7523650825407,12.7777377564691,12.7369213555758,12.8058286888132,12.7970004554604,10.4929783959379,10.5546081117412,10.5058587189404,10.4873641500824,9.98958038397422,9.98664486696558,10.0118294158015,10.1099340220687,10.858279079899,10.6711433495134,10.7632393355264,10.4797298223835,10.7686076926916,11.204817178997,11.2487425283355,10.8580971207831,10.8677914186503,10.8887979512085,10.9146543742545,11.0294595830454,11.0249825200416,11.0231848424466,11.0535878770494,10.7185453255924,11.2572151671438,10.8470382747935\nWASF2 - mRNA,11.3079832820503,11.3085711791952,11.2052888356429,11.2450378718242,11.8656119310053,11.7584553063227,11.8939322589095,11.8709701620727,12.1030071990448,12.0704253885437,12.2906654258676,12.3543747818338,12.2982828253366,12.29921869463,12.2944611105298,12.2367887736374,10.9987265879984,11.0530418567357,11.3240902302486,11.2300035420239,11.5781502683524,11.5221951735617,11.1667633280115,11.1926268105148,11.2689114000578,11.4529937849147,11.2846661793972,11.356873074598,10.8681433662425,10.680845308165,10.7312636243303,11.1584375438894,11.1510790269881,11.2293808508953,11.1961352109905,10.8076082061709,10.8602092178378,10.9214088703126,10.8858077666111,11.0329266114264,11.1684362711147,11.1853967141783\nTJP1 - mRNA,11.4081019527613,11.4333579053638,11.38812333149,11.4066198626797,12.0431094918343,11.9738745891431,12.0004481622817,12.0123231121542,12.2007201369889,12.1920387870435,11.7802894218327,11.8185944795181,11.9730086714027,11.9713276358426,12.0397152053799,12.0009197025106,10.7873761471779,10.7626703662812,10.632212525611,10.6878316314966,10.551658994022,10.6015199247178,11.0458961415298,11.1154589499924,11.2820867888056,11.0280368064383,11.0147781025223,10.9943029952133,11.961252770634,11.796322525585,11.919174371246,11.2398272507613,11.2520864827697,11.2767964218771,11.3393625541775,11.3677454278574,11.3967636158299,11.3609610832363,11.4052957443653,11.0429802760903,11.2724416413685,11.2348089083033\nITGB5 - mRNA,12.0486999419589,12.0468115768736,11.9322250752453,11.9718168891664,12.0501182260617,12.0360928828024,12.0773876693522,12.0437190800278,12.1487254993506,12.0925031134926,12.1119026886669,12.0736667552314,12.3928813492029,12.4849781513373,12.3972813282988,12.3225648486477,11.7477747470276,11.7203742891998,11.6813761272882,11.704075755638,11.3369783923995,11.3317028366915,11.6973540540378,11.6518208115874,10.0553159269585,9.6316149853268,9.73349199213232,9.58383822572427,10.6616924887751,10.5578837856933,10.3769801561306,10.3130548079992,10.2019055216914,10.1681055930707,10.3244758792075,10.6108099260435,10.4795802652135,10.5525550172044,10.428210379766,10.5024118947276,10.4570176686019,10.3692179417337\nGNAI2 - mRNA,13.2208982369678,13.2279233999096,13.2880699730273,13.293701482869,12.5103849793335,12.5012627209383,12.5432020999388,12.5341525954934,12.2941144956646,12.2805919659986,12.4476714379418,12.4700106903809,12.49098830289,12.4159365069076,12.4893079286798,12.4983351568439,11.7858729482497,11.7713990489489,11.6326887411759,11.6452012962367,10.9724515854113,10.9878587459105,10.7148057660392,10.7157341596543,11.295142941631,10.9633740895739,11.2074231804647,11.2687421643193,10.8357218885501,10.9153105618021,10.7195818192827,11.0648701722226,11.1249781303821,11.1029685117382,11.1149705551947,11.0702744232469,11.1575983695378,11.0919037690725,11.1128403370974,10.9498708716989,11.2818772213781,11.2053651304109\nACTN4 - mRNA,12.5001459232039,12.4214954052554,12.6567978915869,12.6474670458886,13.3067459822735,13.3467810355967,13.2776576247225,13.2759159827108,12.8335177710737,12.7563473715694,12.9857728051043,12.8672227630026,13.103542329392,13.1303616206204,13.0866038366259,13.0389124852358,12.3428677643072,12.2769009059374,12.5675570865678,12.5753947455702,12.7641695178098,12.6681534465395,12.3094975330422,12.2577794900401,12.062969499502,11.5393273952788,11.47645632692,11.285682026741,10.7686076926916,11.9742042508556,12.0226200201991,11.9499599404814,11.8336124864385,11.8329970939914,11.7709369281144,11.8520405735188,11.7742229450611,11.7623266477897,11.7560568271658,11.4692450307924,11.448663881272,11.5123428838322\nRAC1 - mRNA,11.7957602543803,11.7532727301966,11.6433434342517,11.7117809017923,12.8993797496973,12.904995875538,12.8960009358353,12.8823406828783,12.4274313067362,12.4048184220976,12.8476600414553,12.8605977615768,12.931345775355,13.0226380583719,12.9743915862605,12.8728329200815,11.723049457932,11.6268597783852,11.6103767243705,11.6099198717479,11.6008743659154,11.5791438514568,11.7177314414092,11.7088978095969,11.214972592947,11.0831783606308,11.3050339770912,11.1318065189632,11.3986580829413,11.7203736723517,11.7018798175492,11.6123579675251,11.4864297904626,11.5534555411117,11.5130179122054,11.619643700223,11.3428942665239,11.4664946254075,11.3707409568546,11.3076835022848,11.1334226815119,11.1394308602261\nRAB11A - mRNA,11.4890699491776,11.5105598814269,11.5649590245219,11.6267578209419,13.5148791342281,13.5218554369324,13.5100229574344,13.5020103877017,13.0783030817439,13.1467101875719,13.438434783489,13.4192801746042,13.2478005567073,13.2031989211757,13.2856203936473,13.2854666868698,11.7289284189641,11.7725336892283,11.5850100042121,11.6528726737851,11.562365984369,11.547112805766,11.2127934502745,11.2303527996226,12.2689114000578,11.7198029811744,11.8673477388671,11.9094140976268,11.2342712650404,11.575742307779,11.5319548161069,11.8166263584138,11.8113493587132,11.8781433352059,11.851606541171,11.5852683755586,11.7861042357542,11.6483072470684,11.7107814260966,11.4580177753692,11.7267004504451,11.6278201012522\nROCK2 - mRNA,11.4735569498063,11.4321877414251,11.4971074194324,11.4114390244068,12.0451154641276,12.0119612203295,12.1046551374225,12.0400493269577,12.2324686459004,12.2196731934883,11.9717250306187,11.9630149081945,12.3080747291287,12.3278473600078,12.2233600435806,12.2120379547392,12.7109221703918,12.7693165545974,13.1775805086608,13.0253575851777,13.5906417671657,13.5041397346915,13.5173897388936,13.5136398629861,10.7861292941431,10.9804187862056,10.8489692095523,10.7799033646304,11.3986580829413,10.6902440061673,10.79387560163,10.7959609093548,10.7963760295715,10.8208778996161,10.7095136482688,10.6234131077619,10.7200585464812,10.7468093480085,10.7021420197729,10.7434199942314,10.7540768893764,10.642541606959\nCSNK2B - mRNA,11.8194145054601,11.8705652566346,12.0424852105963,11.9955459609571,12.5105301708036,12.538090384443,12.4734330107698,12.4468442174616,12.0287189702985,12.0535471337772,12.0107191622345,11.9884957432486,12.0633532392763,12.0981278785296,12.0858565198307,12.0525279993144,13.1346218628235,13.1616069122121,12.9250817547892,12.962603360851,12.9871596729112,12.9238708803222,12.9571735710234,12.9873251981702,11.7861292941431,11.7598406954887,11.7133141101947,11.6979099924331,10.9308791215905,11.8222011574106,11.6688510171617,11.9905756805604,11.984274936674,11.9375582653344,11.9298779497221,12.0836267455931,12.1016495626669,12.0896228278213,12.0806284894661,11.8926460030059,11.7608405915957,11.8470382747935\nNCKAP1 - mRNA,12.0814258726586,12.1093521162217,12.2168817761576,12.1465431177392,12.2705116238818,12.2800564508399,12.2775782881974,12.2587349386795,12.5227211855331,12.4821124456297,12.3696205936388,12.2923462010636,12.2716226879019,12.3499163776339,12.3286604011901,12.3168941887071,12.2781855215567,12.3226754441808,12.0973915104463,12.0753625887556,12.1115918265262,12.0963187227331,12.0597974489851,12.0718992827792,11.4549233856678,11.2349560153628,11.5060814960292,11.3134804790846,11.7856812060505,11.5681157083043,11.7078046515063,11.4379489224379,11.5074680367501,11.3978205643521,11.4497494252653,11.7218010525006,11.6743488710768,11.7015555074772,11.6561273512265,11.4238020600313,11.5205684358457,11.2825846328728\nPPP1CB - mRNA,11.705485510353,11.6539779943879,11.727662721601,11.7234543048387,12.1779279455098,12.2822199529313,12.2158189499873,12.1699731537859,12.3550377100803,12.3518192310803,12.4968468371431,12.4103063599383,12.3620839842032,12.4147573527423,12.3889747649433,12.3950932013701,12.2094858099257,12.2704901046125,12.2790187211144,12.2399058384589,12.3883699405479,12.3318621712812,12.3021692576052,12.2513636763669,11.6196681480044,11.4477507929266,11.5002049296926,11.6019855724345,10.8998522259699,11.2563474796371,11.0899502692278,11.6815870514533,11.5113221783831,11.5709278399388,11.5654108938561,11.4070134764517,11.4458327472261,11.4390852571866,11.333017203802,11.3528355386771,11.5552199825007,11.3736366130007\nTUBB - mRNA,11.4611178607375,11.4933834097835,11.3521302239722,11.2983920482754,12.2352777411991,12.3004798931999,12.3016539236711,12.238315944118,10.1377168533324,10.2315249973626,11.5731298209659,11.3200765402109,12.170891128884,12.3262804053182,12.2380651639875,12.2309423256614,12.5465558374617,12.5599845530705,12.7564925934283,12.7801458993102,12.0559467936561,12.0396976774301,11.2311800565073,11.1366704045408,11.3336170894456,10.5653812869267,10.4704575862985,10.5184622168722,11.2091802840776,11.7655321334715,11.7798823295505,12.0129971931954,11.8651363138443,11.7713566598952,11.6262167251502,11.5583291109632,11.5769336043826,11.7413305486852,11.5693395339871,11.887075744963,11.3587903463052,11.4041940799766\nALDH18A1 - mRNA,10.5306696809814,10.4147574178893,10.476165116249,10.5015785155725,11.786044673687,11.7118660126058,11.7407555971678,11.6786667305178,11.4016401264466,11.5538577185437,12.3520659705317,12.3537852047758,11.7949634849623,11.7888157892417,11.8662645426752,11.7949040224818,12.447234141307,12.3802517095834,12.7293647605876,12.6903964233559,12.9991267082187,13.0661805300579,12.459948346296,12.3400161030744,11.9683326253639,11.568605187801,11.3709219127476,11.3780895605077,10.4637531111632,11.204817178997,11.1693847367222,11.6758860441101,11.57054186338,11.5676160564681,11.5997694923391,11.4134554815763,11.4044919518452,11.4070799667569,11.3655888370983,11.5490123075898,11.5981513767983,11.5282844277013\nCLDN4 - mRNA,4.58263310260287,4.59789227654924,4.23234784786737,3.92022726840375,12.314583563466,12.4039475219132,12.264827917615,12.2071351387459,12.4067088889409,12.3894848512498,12.6731262043276,12.5829851826274,12.257774291839,12.3673926549496,12.2910610084164,12.2649978439357,10.5140133079954,10.3392339835567,9.92401219144113,10.0854955387141,9.48952889432799,9.52498514743616,9.6130799937782,9.64539029890149,12.3145082664979,11.7640652813504,11.5235689227581,11.3780895605077,11.0201464596876,12.0870155696897,11.7657520007156,11.9922281472213,11.7868680224126,11.7617886041571,11.7162276678719,11.7666416193455,11.5765312351035,11.6452043051701,11.5637717030778,11.7216782150774,11.7769452616098,11.5712365728858\nF11R - mRNA,5.54121317522289,5.42796727510693,5.32667523041552,5.3424602690868,11.5979844717837,11.6187744422724,11.6277892652734,11.6068481609813,11.5470270920492,11.6651587187622,11.8193633811332,11.8675325647376,11.7407615530058,11.7205525278665,11.8096810143088,11.78390621285,10.3739653478877,10.3908807111021,10.6440710096124,10.6185154076742,10.8284326283064,10.7496056696647,10.3857638315452,10.3526443540368,12.3710917948643,11.9670435151405,12.0023588432683,11.8725145148152,11.3070276074757,11.6761228979226,11.5908485051605,12.0526037734319,12.049341177494,12.0366983475008,11.993105229878,11.9322315963696,12.0064319501939,11.9902773705065,12.0168582634916,11.8898635623346,12.2006316387775,11.9746860978894\nITGA6 - mRNA,8.9566883189889,8.9144671369424,8.20280357225335,8.42865592093561,12.4498932189771,12.4702862958472,12.4847483239976,12.4561500834921,12.6412927283749,12.6411142939028,12.1040125721158,12.028942536195,12.319706217169,12.424017207014,12.3280285408752,12.3372050912294,12.0095539216788,12.0446132346651,11.8767518681875,11.836914643305,11.9122764329846,11.9114507072376,12.3861882576451,12.3726800907294,9.14458326505562,9.53603732534457,8.88549508557737,9.08741239960477,9.89985222596986,9.95483892598869,9.67188507725137,9.72662241918261,9.70826267684201,9.65797712377172,9.69709229238057,9.99162881663479,10.1108484995133,10.1142676861315,9.98616853360898,9.99199634737154,9.90824781950413,9.69291512947407\nHPRT1 - mRNA,9.57153210936962,9.57432380541084,9.52771346987126,9.49553959909119,11.414640231632,11.4064290804189,11.3974916458516,11.4125981599878,9.87513964764218,10.0460883637207,11.1140969020799,11.0849009492671,11.4995594892887,11.5108680190618,11.5467823851483,11.5026233648189,10.7836152090708,10.8505362012296,11.0791633659918,11.0173011780576,11.3851375030107,11.449164352639,10.7842936911891,10.6877286987229,11.295142941631,11.1854165545615,11.3514695500814,11.234253787934,11.1311777720763,11.1029375649778,10.8323497494446,11.1933353404682,11.2327174960125,11.2879077286476,11.1111204327091,10.9372878601374,11.0134349762784,11.0149279111539,10.9928669161542,11.2739347109302,11.448663881272,11.0703383369531\nDIAPH1 - mRNA,9.72815676055505,9.79865468191415,9.60503646389719,9.75210450959542,12.1162968602787,12.0677924525308,12.1060854957411,12.0963724981225,11.4604068679845,11.4087010526434,11.7509785821258,11.727093581725,12.0342494459147,12.0940820771657,12.0649625347602,12.0319079002103,11.5670480703173,11.50313377336,11.4983005486075,11.4844113821989,11.2436864723469,11.2621737066041,11.1293926343474,11.1485083156277,11.3710917948643,11.0020415979053,10.8303535313849,10.8192162886552,11.8840849101115,11.9371850614248,11.8592163552973,11.3139360225482,11.3882591296771,11.2956850792394,11.3410071867611,11.6113523265917,11.4937139499765,11.5226198106627,11.5267704843652,11.3729011617312,11.5013108189964,11.3868120017484\nYES1 - mRNA,10.1214991887346,10.2147814187237,10.0480340445169,10.1628297743327,11.6790187473227,11.6551219677958,11.6998965458668,11.6927587999489,11.5461660385824,11.5821677114085,11.6932007346852,11.7472113943794,11.7652066285929,11.7300643146928,11.7716166040339,11.7205230595255,11.0807020363361,11.1687345464943,11.2946008564647,11.2116167940275,11.4035532707572,11.4266634198079,11.2496463043728,11.2290791777868,11.466511359943,11.697888204746,11.7384926731907,11.624353385463,11.5260373894175,11.2595078205305,11.1693847367222,11.1250196527736,11.1121963661829,11.177912470235,11.1841430978725,11.0888294019378,11.0235072207609,11.0982208605251,11.0837692906389,11.1114600139731,11.4436282913747,11.4717008993349\nCSNK2A1 - mRNA,10.127925457894,10.1422127927731,9.99945860565374,10.0051419281717,11.4453435619714,11.4637880155196,11.4653755692796,11.4244991970842,11.1675108564409,11.0861763096744,11.4041017390681,11.3146335426131,11.3651627199872,11.337835768861,11.4452669868895,11.4106933633562,11.9544186400871,11.897576155026,11.8996855088547,11.8912299101482,12.0045945572673,12.0269607284648,11.8830276447744,11.922518429534,10.6905516341609,10.7385301892433,10.7234383274684,10.6287856317782,10.1836451919705,10.7766042909705,10.7370692460115,11.2076305750858,11.2583055882564,11.2314751282232,11.2046772537705,11.0480093509084,11.0164047077529,11.0042034979645,11.0053762956855,10.9174493940065,10.4536819560386,10.6715414786815\nCRKL - mRNA,10.0258894456914,10.0938953101091,9.91467818545673,9.96985803612835,11.4395599893398,11.3750140486031,11.4056457776,11.3930471096792,11.3166742923053,11.4957954050984,11.6885456901817,11.7266382574224,11.5727613099623,11.4511578869792,11.6477112939991,11.6389846121506,11.1907226803064,11.0558404900883,11.6137576650829,11.5441539398031,11.6947751988422,11.7364259040141,11.425123920192,11.3873123026814,11.0705826836118,10.8083741778574,10.5293512753521,10.4093404944921,9.961252770634,10.5319819222505,10.406423934023,10.7012527642686,11.0586669697661,11.0487736748617,10.9281596185731,11.0778820699666,10.9458818732179,10.8290212978438,10.9439296622093,10.4729681168352,10.6442382902531,10.5440517435596\nRHOA - mRNA,11.6610608747519,11.5160929672951,11.5748430128314,11.6228848117947,11.3829520600053,11.4510235434808,11.3326841395301,11.3136812064526,10.7879910474641,10.8574916534201,11.2780001545221,11.248298176235,11.3734426377307,11.4583290670735,11.4199919926137,11.339604119066,11.3982712544038,11.4099636098436,11.3296815081858,11.4229290544197,11.0675421597148,10.9609145918722,11.1594986546389,11.0520490595494,10.4780069987808,10.2206818843807,10.5177633010769,10.4499824789895,10.3304865802997,11.030787779222,11.0297327463937,11.4252283322088,10.9865569087431,11.0874431462459,11.1199056627695,11.1408516142861,10.9915687343823,11.0720147631661,10.9726777392038,11.0429802760903,10.9655809945701,10.7952768471037\nCLTC - mRNA,11.147321870804,11.1649328692731,11.1489535983091,11.2312943706593,11.4236116674655,11.4416981669385,11.4717696419757,11.4529579579919,11.0805844934704,11.0632430847598,11.3016169478444,11.3273020730484,11.2678440618233,11.2846077802856,11.2949461859194,11.3131555819233,11.8744633355057,11.9081112347602,11.7431027068984,11.7084534968837,11.4543496011013,11.4585344365834,11.2870970870793,11.2800896517652,11.1731524172524,10.7919174019903,10.8396913959642,10.8269519374348,11.0487156118843,10.9312521056711,11.0106869907031,11.0227317712018,11.0165369036093,11.1174795200008,11.1042195535756,11.162972791697,11.0982898987207,11.0951921084521,11.1022240683043,10.5096798951917,11.1355059960124,10.890854507086\nHSPA1B - mRNA,9.34656574436935,9.33184912084311,9.30482450919386,9.16054159773746,11.8281252772574,11.8013259468639,11.7667631380614,11.7120650774258,11.8402069779434,11.8263459899009,11.6907525836805,11.5891347336724,11.4883081548909,11.4361266666307,11.4408903174164,11.4491194728439,11.461781257225,11.394324486066,11.7264675578635,11.6307065824738,10.692875040463,10.7496056696647,11.4703939700577,11.340900739816,9.89304449805966,9.67413960197335,8.81149450413359,9.36752031879751,9.89985222596986,10.6182143198946,10.4913128316096,10.7610621595039,10.7157490259573,10.6213114971769,10.6344788355993,10.9993852602692,10.7054117705168,10.8271970256839,10.7177529389892,10.5525353741535,10.3817890375569,10.3868120017484\nCTBP1 - mRNA,10.1285082472539,10.0291190139274,10.0747831381438,10.0560904337724,11.2284090098463,11.2435120693531,11.1855263455852,11.1216302157082,11.2994044118483,11.2909851314573,11.1857385694951,11.1029061865366,11.1782616827717,11.2732670534034,11.1938392007037,11.1208824764056,10.9599279838592,10.891148338202,10.9417995976465,10.9248849723812,10.9399767052262,10.9773043738346,10.9652447460863,10.9667454030985,9.85827907989898,9.86453807415541,10.0713616308887,9.98049719568826,9.07672998805394,9.94701942152939,10.107984192058,10.4835673456097,10.3820816129981,10.3175369881532,10.3499026248982,10.4767381667633,10.4058514886578,10.4503881281861,10.3531481589083,10.524106965827,10.0824987463492,10.4467545422047\nAKT2 - mRNA,9.7964938903182,9.81813623130711,9.86760024665035,9.90754813433301,11.3482794823505,11.4021249956358,11.3342106395571,11.2813251734309,11.0317979251007,11.0921317156167,11.1526768267281,11.0720547052321,11.0748699395094,11.2313511987592,11.1051620426275,11.0655332091772,10.5158280189779,10.4297566245561,10.6548953911491,10.686897848266,10.956701483122,10.9003200671412,10.7769758131404,10.7047803958721,9.89304449805966,10.0786628795083,10.1485294914112,10.2107948151101,9.69821836480021,10.1445566541796,9.98652067957238,10.4390263646395,10.4596930765486,10.3337114182608,10.3206705488354,10.6567266325886,10.5566763909056,10.6267948012089,10.6047535615496,10.5734949927891,10.3498461450541,10.3692179417337\nABCF1 - mRNA,10.0592837922889,10.0861787578587,10.1566788914183,10.1076783224311,11.0469184575398,11.0433394088062,10.9952471731247,10.9852593494798,10.4003701483356,10.4991609471555,10.4409862268377,10.4454565048783,10.6925547614316,10.7449193593289,10.7341908147246,10.6982626224059,11.6592136382312,11.6441128747755,11.4896514289884,11.491113498874,11.3524084329691,11.3579142807604,11.4076700755155,11.396985309021,9.45492338566777,9.88537157757258,9.71331411019469,9.40934049449213,10.1311777720763,10.5630088178712,10.406423934023,10.5523893783541,10.6349272766864,10.494945459439,10.5109321882035,10.9306388180224,10.7897537704184,10.8204882224511,10.797966999744,10.4276441263257,10.2912514911433,10.2684173009462\nNRAS - mRNA,9.98335146089227,9.94760352795014,10.0073129410396,10.1651143275273,10.9662217532094,10.9865347608071,10.9445797472768,10.9662523003701,10.7836140829176,10.7871897324562,10.5605810850674,10.5956322452724,10.6030683756417,10.5615606877899,10.5739505598902,10.5569654204922,11.457066563353,11.3657897301393,11.1923541986381,11.1864005084186,10.9888061982616,10.9987379389536,10.8660877823527,10.8456535861811,10.0553159269585,10.1231940394786,10.2217784734418,10.3885819343253,10.2831808655214,10.4841533971599,10.584422236001,10.3478901834901,10.3944103074141,10.2961854069853,10.3762645115475,10.5951737586799,10.4469337890569,10.4428627286174,10.4486243644818,10.2306072792661,10.3534304824031,10.3151105363942\nPAK2 - mRNA,10.634842267665,10.5987269280281,10.5444949219621,10.5886868253191,10.9087741275872,10.9139766651314,10.9329401915311,10.931228710188,10.881070102011,10.8469641877403,10.6814115369816,10.6441760972304,10.8042939846629,10.8001488800493,10.7704543587113,10.786253203836,11.0285864808252,11.0823127014495,10.9718449456161,10.848443687914,11.1133687636161,11.142286537474,11.1425157817648,11.1144246363807,9.9601586939182,10.4993425253639,10.1485294914112,10.3134804790846,10.6616924887751,10.8413099803583,10.6718850772514,10.5573590527168,10.5910396326854,10.5023361086273,10.5825931733309,10.6796656542033,10.4983945750672,10.5906901460912,10.5811225313878,10.6009757292112,10.5936122171831,10.2872763281975\nMLLT4 - mRNA,8.21226465265776,8.2776330020225,7.99443803618919,8.0511581382302,10.9210599260263,10.8957233456942,10.8908236711072,10.8310672743021,11.2210044084432,11.3073960248351,11.3588667739759,11.4374099110434,11.3016613154068,11.3597536715145,11.3042923944548,11.1972985437033,11.1430855998748,11.1545889490255,11.6776887258153,11.5139338704863,12.2722810747358,12.2168230332686,12.1675865491699,12.1334404254886,10.6905516341609,11.0899251930713,11.031172540773,10.6894484136849,10.9910001140281,10.7588478201663,10.6658105628417,10.9126344047926,10.7701240307677,10.7938518038202,10.7756354143979,10.6024674801852,10.5509530253454,10.6589443617964,10.6106884315514,11.0379622013237,11.0094072355347,10.9862740721647\nMYC - mRNA,8.72661788569886,8.81311066499796,8.22501830606888,8.34514935661444,11.0165703666705,11.0223528261761,10.9913824640338,10.993671589296,10.5771257382041,10.6338603980287,11.192330189902,11.2149247387319,11.0692174768487,11.0149741928405,11.0959127800145,11.0828204022625,10.8045512909861,10.7800752382409,10.9868224488439,10.9053483861483,11.7145772063593,11.6797171906035,11.64724093921,11.7136866539305,10.1154369193961,9.51613776790686,10.4339317102734,10.0349449797106,9.18364519197045,9.44582527850084,9.65971036330527,10.0662654286572,10.2119708767579,9.97725564024513,10.0204872862666,10.1201656516808,10.1701867102284,10.2036945945301,10.1056891120994,10.396617230705,10.1124223828994,10.2872763281975\nPPP1CC - mRNA,8.84765098420549,8.9462925827978,8.86681553352745,8.92562725614634,10.7407206831775,10.7381096042424,10.6942016896266,10.6754333810855,9.9165940748139,10.0329903748518,10.5723320907598,10.433788197153,10.5634096101316,10.5144503025434,10.5903781189332,10.5453730122783,10.7989731359215,10.8299748787472,10.7422203245829,10.8107931539683,11.0737614709046,11.0281411690387,11.0777762881552,11.0192674033404,10.9101180114186,10.9535430802142,11.2778125083561,11.2630510522415,9.961252770634,10.2876436101267,10.2689204102731,10.6491345383834,10.5541511503715,10.5049355598547,10.5806057160103,10.5878429754226,10.5973070419698,10.6306350080619,10.6165989870399,10.4504841026838,10.8737201363759,10.7139767450019\nSRC - mRNA,8.89088626059316,8.84581978999283,8.93428647008187,8.96111969905065,10.7672108977338,10.8400250392193,10.7326758374412,10.7567607703933,11.2288958480346,11.3140995651311,11.3607159881247,11.3502426737365,11.1057527920934,11.1543158150927,11.129661571369,11.0713186535333,10.7971089477297,10.7260468822563,11.0480832989991,11.0744703288218,11.4107964581822,11.3428139767558,11.5997227352876,11.5752038506532,11.0705826836118,10.9386698037326,11.1104614505776,10.6376094821652,9.69821836480021,10.0084199661935,9.9063503308884,10.7614928790297,10.5961722620216,10.664579858596,10.648207995902,10.5453309218663,10.4847358217693,10.5744697936329,10.5747073748846,10.7919185119226,10.8073189106533,10.6351996316178\nTCF7L2 - mRNA,9.214462208362,9.23746746899609,9.20528883564294,9.18326167423754,10.8581008410175,10.865259500964,10.8619059396054,10.8457456609335,11.1555164404292,11.1723221423062,11.008754975354,11.0767068326742,10.9351834221423,10.9646954675355,10.9980198749662,10.9922037101752,9.83721194020917,9.91017468451493,10.3587573621375,10.2967097358885,11.2719629150607,11.1806554869322,10.7224450350006,10.6425230670053,10.9436705711296,10.9804187862056,10.8673477388671,10.4399288143256,10.0201464596876,9.45688046700983,9.635048309071,10.6662573429124,10.6637327552487,10.7035739902118,10.5670179060422,10.3004987386091,10.2982043856603,10.3137534361643,10.3044396048883,10.1768360228251,10.5029254750585,10.2396583079647\nAXIN2 - mRNA,6.40160710792731,6.26926952908787,6.4237739185206,6.26505576540119,9.74467598391652,9.79780165082193,9.67505482543795,9.67294124876623,10.1959702186722,10.3647915282261,10.2938951004666,10.4050516692335,9.8294788269212,9.9308632381366,9.8614693590947,9.72003209643044,11.2483435100278,11.1586447208879,11.7660743013733,11.6850284666752,12.0969377932707,12.0396976774301,12.0226074345798,11.932725294166,9.20116679342199,9.42658618292857,9.2640067088311,9.46988203642718,10.9308791215905,11.1134554368918,10.8323497494446,10.4772879231134,10.3021517995621,10.319507210434,10.4762889443276,11.4581637233283,11.1402782435575,11.2167591925892,11.1420604760124,10.9337512063356,10.6587963859433,10.5282844277013\nMET - mRNA,8.07388116850183,8.04512859821624,7.60409506416337,7.96985803612835,11.8841082746227,11.8805718308796,11.9353560922479,11.8731428959786,12.1026168066622,12.0781348232044,11.8296364852041,11.7683803630912,11.9854665611439,12.0844739291878,12.0021846446271,11.9634556331807,9.97682505313541,9.98647596902379,10.0989432590713,9.96510887688142,10.5791792948161,10.4155806877371,10.9806909039926,10.9879841994728,11.100639917468,11.1854165545615,11.0713616308887,10.8192162886552,10.5857436355418,9.93125210567108,9.96689187282345,10.8800807092528,10.8155728691363,10.8187900627237,10.7171325760373,10.4901697072049,10.5591223282661,10.6201375445201,10.5664684981323,10.9498708716989,10.9395857860371,10.8406687499178\nINADL - mRNA,6.9527447310002,6.67589478855052,6.7439936709456,6.7682241749587,10.9328086762533,10.9037083296775,10.9616673693638,10.9064084428402,11.4509305024651,11.3882752975919,11.3267466556822,11.3780524074734,11.3312446777959,11.3924678728019,11.374819390655,11.342796629864,8.99645385988435,8.96082445825009,9.49699340432958,9.38399253957099,10.243037340709,10.161060611749,9.98144016136643,9.96061930030166,10.201166793422,10.1875150204455,10.1929236107696,10.1989207148218,11.0487156118843,9.69024400616728,9.54525163872036,10.1263579599576,10.1996592074332,10.1304438525781,10.0379747129954,10.0240379493033,9.90127356114269,9.8513330138044,9.89584943515784,10.2217834288791,10.0519412793733,9.98627407216466\nBAIAP2L1 - mRNA,7.59954648553922,7.60787636512187,7.63760432635362,7.7682241749587,10.8378779607384,10.8198126817281,10.7809370980861,10.7814946245129,11.0444550166897,11.0696710381783,11.0559338765217,11.2080911642414,11.0711040928373,11.0781569028182,11.107004800626,11.0075731887425,9.63499740081035,9.67801215226427,9.42403863303735,9.40117450177918,9.74561406482743,9.67921634177333,9.73981906204848,9.79059837179785,10.4314644128438,10.4651538391881,10.4339317102734,10.1504221971306,10.1836451919705,10.0958828074439,10.024994827387,10.1795418197736,10.0753029713579,10.0561774427319,10.014021744604,9.90166395040663,9.99999501222537,9.99041322389339,10.0512793592898,9.83883955931012,10.5474173943045,9.98627407216466\nITGA3 - mRNA,9.75104505194961,9.85493084069258,9.8518243098899,9.88092430770806,10.1297809771185,10.1199313661472,10.0929806040416,10.0879787480451,11.0132248530108,10.9324315819252,10.6111701702247,10.4404673527906,10.3666996276013,10.4787851094566,10.3269221189874,10.348525406581,6.15623430392112,5.92997066872397,5.31551417625918,5.37937591150062,3.03228483492846,3.99471816750126,4.23997317496528,4.03829866412353,8.20116679342199,8.00917777918698,7.65102983194035,7.73948909618446,6.37629026991285,5.74538556035974,4.77706731394343,6.67359931825354,6.34952505782419,6.51657547541058,6.19898818188872,6.07526327265848,5.9585983903942,6.17050592987951,6.27440504368698,6.33248689328533,6.55677544900275,6.72092950564367\nPIK3R3 - mRNA,8.54296295717285,8.3646052141095,8.32210250062569,8.38227958720018,9.2375600703069,9.17917983874954,9.23048412779155,9.26916709748701,8.66488608665289,8.66265967943837,8.96647224914538,8.76793788646121,9.06883985741434,9.17952860311916,9.10958071389449,9.04490346681854,7.50216171810325,7.5221466596777,6.78183218048172,6.90132861469598,4.93274916137754,4.99471816750126,6.10347117506042,5.9667454030985,6.5007270752809,5.76125026574339,5.14852949141116,6.08741239960477,6.37629026991285,5.23081238752999,5.58442223600103,6.33694971765941,6.06361618573781,6.12425805263182,6.22365023612299,6.1725604740134,6.01670134495777,6.09165050916827,6.06253049426453,3.87305527464803,6.15822607251248,5.60545228822373\nRRAS - mRNA,9.05806065101944,8.92781316219014,9.0590680947919,9.07078694497913,8.49637812133962,8.58178023479016,8.50223575072646,8.44421609096453,8.68588638526544,8.68647361833136,9.42575666151339,9.37456903747312,8.94591596693596,8.94645122360638,9.04337662533444,9.01639703912954,5.18389064682398,5.16009045208503,5.27704002844455,5.48629111541713,4.21285708057028,4.85978858741515,4.79796862808617,4.72795854351138,4.5007270752809,5.568605187801,5.14852949141116,4.86501997826832,5.37629026991285,5.23081238752999,5.94699231538574,5.35532824697426,5.19571972174515,4.60968487980206,4.53740439956465,4.89692603139997,4.65141688087602,4.23062692231708,4.47756799354338,4.87305527464803,5.39923417201627,4.60545228822373\nPDGFA - mRNA,5.40930149971731,5.07182346488166,5.69531982420991,5.52429859207261,5.83184836330015,6.01543341223635,5.97268637032391,6.26255225306545,7.72699158859512,7.4261920728902,6.8466707271289,6.97379858390777,6.72407178616886,6.69770747532298,6.51132139055988,6.37932250588909,5.88010989864688,5.28407916936048,6.01010116815484,5.64856254431601,5.82583395746103,5.4727654643059,6.2499572635379,6.3602267590109,8.5007270752809,8.58142922815858,8.42154798581758,7.73948909618446,7.961252770634,8.09588280744388,8.2689204102731,8.50481813280778,8.30841985449451,8.24319912535533,8.15905670961499,7.99551225112202,7.95364917442247,8.07329795727441,7.95467450450387,8.002338291593,7.8886190127552,8.06488390686103\nIL1R2 - mRNA,3.89910676739813,4.09093228782936,4.23234784786737,3.92022726840375,7.39684244778871,7.33180198178182,7.40796708951013,7.24117860227288,6.92368349960021,7.0696710381783,6.9086576140246,6.68973867461085,7.16383637135008,7.08928600177783,7.20489438921271,7.31835448859767,6.9515722612035,7.02950634227489,7.5351922128986,7.54105691884803,7.67998309099758,7.76479207347941,7.63610221169406,7.74231383648845,7.67065207672321,7.34621276646455,7.37092191274761,7.08741239960477,7.69821836480021,7.67138497891597,7.73126362433031,7.75545107224182,7.64197595163472,7.66960869745412,7.39628962226234,7.21505255612748,7.31566829140182,7.27769517597386,7.23516988040388,7.45801777536919,7.55677544900275,7.19041478894489\nHSPA1A - mRNA,6.33044607900607,6.36656673044279,6.41521190501718,6.47604342346539,7.60262292792024,7.50481825297528,7.44912441426689,7.51493941469973,7.72699158859512,7.79816085027716,7.36194748204571,7.4852989226879,7.35930749337124,7.55086608699371,7.39547306187336,7.42341509007226,7.1747306475385,6.92997066872397,7.83245825355169,7.64856254431601,6.67998309099758,6.44475108813631,7.68843367578157,7.32727633720961,6.08568957600206,5.76125026574339,6.56356699069001,5.86501997826832,7.37629026991285,6.88288908410968,6.23649893258073,7.85244739871747,7.30005638155989,7.46410805551644,7.45301221227428,7.82392420045706,7.21171732736291,7.4005519237594,7.20963478829674,7.33248689328533,6.94672196731876,7.19041478894489\nAXIN1 - mRNA,6.51292413079147,6.15588772967013,6.20280357225335,6.19523431590362,7.5806636753441,7.53899536829337,7.47592447361061,7.46510825874773,8.09046194572566,8.08766746747215,7.65904372395313,7.46360385158858,7.61656616070088,7.81676014696674,7.61855461388491,7.59601819705602,7.32305339449561,7.10350692371866,7.98047599904556,7.87860853819589,8.54113600761003,8.32435435752389,7.88703179677835,7.84232556846338,7.82265517016826,6.76125026574339,6.56356699069001,6.86501997826832,6.37629026991285,6.81577488825114,6.58442223600103,7.03454107349147,6.75030857342279,6.97725564024513,6.84880182695985,6.89217250142934,6.56967373143449,6.67661300988943,6.63931906353999,6.77994587025655,7.12505920857728,6.19041478894489\nMYL5 - mRNA,7.3938714591477,7.37828015100658,7.63022479598803,7.47112793305127,7.28178407056359,7.28653724804603,7.2564793363245,7.22493796126567,7.75110404471192,7.83414014738273,8.18678135598134,8.17142310009527,7.58733544937712,7.70158048447018,7.72606023935289,7.53033029398878,6.81084723620977,6.92453678267334,7.25246200572388,7.21809500446756,7.46969014723576,7.36228892794433,6.85140788704763,7.18213443669719,8.40761767088942,7.22068188438069,7.2640067088311,7.52798499099075,7.961252770634,5.42345746547238,4.77706731394343,6.8844375129619,6.91818574621624,6.84124894703676,6.65691149402074,6.27835513803599,6.49669428607738,6.45512086629882,6.30228407626242,6.87305527464803,6.55677544900275,7.10795262875291\nBAD - mRNA,7.53419274754512,7.33485787071545,7.65951910278207,7.5937836923939,7.5311435395003,7.58178023479016,7.52807575566171,7.53264141643319,7.54587890652035,7.56540901137941,7.87109420106812,7.80808807104212,7.83836791875747,7.94318350977247,7.78249618498261,7.81043821318151,7.0699344575269,6.92453678267334,6.75408718999882,6.80133282471926,6.16978835867839,6.10622648271825,6.10347117506042,6.171565194987,7.67065207672321,6.22068188438069,7.37092191274761,7.36752031879751,6.961252770634,7.46785158483083,6.94699231538574,7.43092586586842,6.88501888228104,7.04073469706954,7.04144690490224,7.62341310776191,6.95364917442247,7.10375711202493,6.95686871791678,7.45801777536919,7.31350429799039,7.54405174355959\nMYL12A c - mRNA,7.21226465265776,7.01292977582809,6.79107880701534,6.79207091691307,7.00011984588995,7.13517265651045,7.07556355096421,7.06990717512305,7.89150309385106,7.59086899637036,7.77081697260965,7.96146778199705,7.72407178616886,7.6623744022107,7.67131727475846,7.62147818204697,5.81084723620977,5.48939807733799,6.45129095528965,5.87533340616303,6.21285708057028,6.18736324544366,6.65501067424412,6.53531346556899,6.82265517016826,5.84871310699373,5.56356699069001,5.86501997826832,8.37629026991285,5.88288908410968,6.23649893258073,6.39139750168179,6.43603405107886,6.47475529971595,6.50423753562945,5.77828153490135,5.52366133367765,5.55843158354112,5.7945972890179,6.45801777536919,6.22235640993219,6.02048978750257\nCLDN15 - mRNA,4.98399566498464,4.97228779133074,4.75081493680161,4.63392308324711,6.60697495245995,6.54381238459114,6.31702087825955,6.35499850183004,6.79815261751636,6.81266041997228,7.64292405858986,7.61298147562681,6.85598346167444,7.02891338379835,6.66332448400493,6.69126651220383,6.43469875032452,6.48201854697239,6.81577951240506,7.01283692991297,6.88028174148341,6.80941506490003,7.36006702045558,7.03829866412353,6.5007270752809,6.61923126087096,5.56356699069001,6.28005747754717,6.37629026991285,5.59338246691469,6.23649893258073,6.66625734291237,6.58115075893867,6.59674582409456,6.34100718676115,6.04987795423857,5.99759252201006,6.0505453058386,5.94365309841261,5.87305527464803,6.65299076426205,6.19041478894489\nHCLS1 - mRNA,5.77357588531427,5.67589478855052,5.54645643829543,5.83095993030666,6.4703402604381,6.71587313037745,6.48474832399761,6.36491397069513,6.9723573463359,6.95670678958898,6.47097137580056,6.65952406156384,6.44560733976546,6.43467306948919,6.26950347063578,6.65884503451146,6.17473064753851,6.0645127921028,6.0675866628156,6.04817000302525,6.36379897864502,6.18736324544366,6.16808025648349,6.28377469857302,7.3080819973385,6.71544657613027,7.14852949141116,6.60198557243453,6.37629026991285,5.42345746547238,6.23649893258073,6.43525179127121,6.4811219406074,6.22859471244655,6.18230944074209,6.00655052257447,6.17816476765188,6.13947903425889,6.10271958438023,6.04298027609034,5.94672196731876,5.60545228822373\nCCND3 - mRNA,9.51024337119532,9.64309576201077,9.91239844564989,9.99319053995898,9.28586140264413,9.2248867778653,9.1619952819568,9.17367984156044,8.93073825713416,8.77613454394716,9.20473758158563,9.03539096356201,9.03212029205002,9.09812787852964,9.07836198328377,8.91194041631398,7.9515722612035,7.61952207072233,7.10100966435305,7.13836781199683,5.41079645818219,5.14155955583053,5.91553822446734,6.10647016676511,6.08568957600206,5.84871310699373,6.14852949141116,6.60198557243453,8.18364519197045,7.17834496763585,7.16938473672219,6.87172646121235,6.93448755854534,6.84124894703676,6.77285297372655,7.78854987035518,7.16965326779637,7.38078866893797,7.4622036838912,6.77994587025655,6.69879445387518,6.34241788238994\nHRAS - mRNA,7.76160324364819,7.8174193641913,7.72679805568308,7.90572769870864,9.50689599325363,9.56054640035094,9.54610305063743,9.52933879376035,8.84184520754375,8.74245141769087,8.46410683450275,8.37573109556325,8.92603925701531,8.95458827304472,8.77324217608297,8.84326960564889,7.82847713023156,7.81132617222535,7.32968150818578,7.35606853800855,7.15881724085739,7.00754220785884,7.19416948535215,7.30810402793476,6.5007270752809,6.71544657613027,5.56356699069001,7.36752031879751,7.37629026991285,7.74538556035974,7.77706731394343,7.15483530720918,7.12645705930804,7.04550394121741,7.0643837161794,7.16469835624043,7.21998494334651,7.27769517597386,7.41190856743692,7.12098278809162,7.28375695459633,7.02048978750257\nFYN - mRNA,8.35853838603542,8.47962939492943,8.35828483693234,8.48582444425798,7.03929044317063,6.96604538527419,7.11018989407384,7.15236933531502,8.00280221671782,8.12590627175984,6.69857208813977,6.84547448693223,7.23017886617878,7.13582858771486,6.93302262011974,6.85046510705309,9.44044273285932,9.3685436824636,8.83742450644504,8.83206595225886,8.7583894364629,8.63444647165331,8.9480990862515,8.97284560263492,6.08568957600206,6.568605187801,5.14852949141116,5.28005747754717,6.37629026991285,6.51092030672272,5.36202981466459,6.40027565302824,6.36564472318746,6.43168657782407,6.30293914592763,6.1725604740134,5.85623667276452,6.13947903425889,6.12630640469785,5.68041019670564,6.02072254876254,5.60545228822373\nTJP2 - mRNA,7.16759560332403,7.18730068230159,7.1470157940676,7.20710841619191,9.70259439517358,9.69003312544219,9.80403492585368,9.7943923672308,9.76597221176721,9.69921056729662,9.50369718743371,9.6364404484508,9.74879887207124,9.66634330441364,9.6683251650633,9.72199494712899,9.86595183565022,9.91566283204009,9.56942384771789,9.54930091294293,9.43827719460429,9.39193438969286,8.94502952117674,8.9804347576466,8.3080819973385,7.69198760330628,6.88549508557737,7.52798499099075,7.69821836480021,6.51092030672272,6.58442223600103,7.20129461830184,7.03366296363618,7.24112181032272,7.11359669065808,7.02403794930331,7.20340205915091,7.08352262828487,6.96561235098875,6.87305527464803,6.86871945531749,7.02048978750257\nJUN - mRNA,9.07024564778533,9.14450096936833,8.86052241540982,8.9399291828463,8.83370630952073,8.79275196053915,8.6894387436006,8.65486967584421,9.32673730711674,9.08915708561263,9.22518235038263,9.28215698025105,8.91767578408068,9.18645466078483,8.96913291352902,8.95763663060088,8.30217466481969,8.15081263849702,9.08300171520229,8.70868353675358,7.30031992182062,7.24264568094484,9.33214301889315,9.21981163984106,8.20116679342199,7.61923126087096,6.14852949141116,5.86501997826832,8.69821836480021,8.86640096132111,8.23649893258073,7.28516717555686,6.96125446810813,6.91085441452262,6.77841249906845,6.24891136014351,6.16965326779637,6.29539717770732,6.17235377221728,7.33248689328533,6.50614937593278,6.82784470956018\nSOS2 - mRNA,8.82049693270176,8.76633225980497,8.57364126762166,8.47849490245949,7.68314663294535,7.62333272423467,7.54926108555234,7.58448034795281,8.91481626184599,9.01429428114193,8.95672427259382,8.92541812153688,8.58733544937712,8.542252956689,8.63090300631255,8.45207884832441,8.52216159157486,8.39038807150682,8.75583697194878,8.78397166405782,8.8735870889094,8.8773612168631,9.14016911799979,9.0685208125378,8.58818991653124,7.9311752671857,8.42154798581758,7.86501997826832,7.37629026991285,7.55274048241735,7.30062927000044,7.94939295480267,8.01589868416034,8.10382615272701,7.96549805166894,7.85356966220559,7.73744062470939,7.85193127010196,7.75479628095992,8.19498336953539,8.09111187665394,7.41280721028133\nWASF1 - mRNA,8.38414320218582,8.41854402122031,8.28968102293332,8.31254469118251,8.59607017612107,8.57238153678791,8.58457719845206,8.51047976650903,8.41364948638562,8.65262000746181,8.5723320907598,8.49388641852463,8.72407178616886,8.8274336484812,8.86843891390954,8.7131410118266,9.21438318395407,9.06944508764503,9.33554374721844,9.35018238116576,9.44280685308481,9.41377543823244,9.57784281372166,9.61157935979706,7.3080819973385,8.04665248460564,7.88549508557737,7.86501997826832,7.37629026991285,7.06731365524711,7.82146143330188,8.03454107349147,8.11696561033074,8.03594963450416,7.967932980586,7.79620344289861,7.79114164454264,7.88971267741614,7.88954569307316,7.39661723070504,7.6056850494837,7.6643459772773\nDVL1 - mRNA,7.71889879269395,7.70098576951335,7.29901888751265,7.33165351413022,8.70361073550507,8.68785875420785,8.63300787329912,8.60135416651721,8.78458788993455,8.71569869008519,8.8466707271289,8.97686488695555,8.72598898916315,8.68406947331002,8.75267196960021,8.62986596753969,8.54899488291313,8.72955661648302,8.76107362031492,8.81336251358046,8.96774458273375,8.9995405509581,8.84976752866637,8.92645269715047,8.20116679342199,7.61923126087096,8.14852949141116,7.67237490032593,6.961252770634,7.3303480610809,7.02499482738702,7.29947501224075,7.23134363147587,7.17736938909138,7.13974897851405,7.1725604740134,7.33101625573526,7.13555333104509,7.20963478829674,7.04298027609034,6.6056850494837,7.19041478894489\nBCAR1 - mRNA,8.06416601366862,8.10039261707843,8.12355682124361,8.10291956591994,8.45352264237783,8.51711751792572,8.37274902563521,8.32737425412169,8.4800768481246,8.47258497837465,8.66703651470666,8.59712752442597,8.35187728789575,8.46916144587448,8.33134272445364,8.28156245722803,8.60120908735406,8.63790060003718,8.62409271748753,8.72150757711116,8.73826273661098,8.59250470858,9.15219721487684,9.08832282005485,7.82265517016826,7.08317836063075,7.56356699069001,6.44998247898948,8.83572188855014,7.74538556035974,7.73126362433031,7.57118375164037,7.60511065788285,7.6129015924856,7.63256163260499,8.05413998109273,7.55667639090561,7.79582616821681,7.711268905419,7.87305527464803,7.65299076426205,7.19041478894489\nAPC - mRNA,8.16305167817908,8.22670906685983,8.1366366482876,8.29596180698691,8.6455634501372,8.61190990097455,8.59887153377158,8.50376433914306,8.550466176016,8.48847635114455,8.46181137651508,8.45042846284085,8.50487328918515,8.43234426342921,8.55084975474652,8.43303309263386,8.18389064682398,8.18983779547908,8.08995447584405,8.12187968920826,8.05020674292572,8.14446528700594,8.15219721487684,8.10369308209454,7.08568957600206,7.34621276646455,7.56356699069001,7.36752031879751,7.69821836480021,7.00841996619354,6.47750703208452,7.58682287541796,7.5031482469374,7.46410805551644,7.3854013061196,7.66022577337964,7.57612875357179,7.48010322285616,7.42781495834628,7.45801777536919,7.55677544900275,7.6643459772773\nCRB3 - mRNA,3.24703007081843,4.05245814001473,3.97709079262529,3.36383391987937,10.1380745173084,10.189975041749,10.1838340344927,10.1459143131777,10.3856844778496,10.3542991544084,10.3978115607533,10.3049061055695,10.332503574072,10.3686111474552,10.4678140723681,10.3306273209604,9.8772893795845,9.79206031269962,9.60674547392363,9.67234638399918,9.47964362126518,9.52943791344279,9.27461931777217,9.25527915138843,10.2820867888056,9.90590850857527,10.1485294914112,9.84991308587811,9.37629026991285,10.0958828074439,9.96689187282345,9.94560734507118,9.88920667924448,9.86437609546495,9.84350752729033,9.89692603139997,9.76241989153078,9.8063590969847,9.82073488480508,9.91744939400649,9.89355130829743,9.7347353051687\nTJP3 - mRNA,2.85471264803967,2.01292977582809,2.48232610087571,3.36383391987937,9.62746965657533,9.67364489498815,9.54504883426543,9.5490420671853,10.1641474938587,10.1659837111352,10.53023025788,10.5177682501685,9.67529448841082,9.75939077860225,9.83235865797782,9.81871637128985,7.11372957540658,7.00911455344371,7.34371106808984,7.37009809791261,7.98958038397422,7.95885271494426,7.84812530305832,7.8978835449537,9.82265517016826,9.50271725214454,9.77302035631896,9.68944841368487,8.18364519197045,8.65227615596826,8.65971036330527,9.7871035227087,9.62030594799625,9.7167150898999,9.59446076183733,9.56039050579984,9.37887589451385,9.43928431824811,9.45911108883582,9.58730079231415,9.76488364433295,9.3955292188495\nOCLN - mRNA,5.42456825637062,5.66318373680096,5.1726416017431,5.1343520737566,7.64978803599892,7.41828150250728,7.68561703264819,7.43718425718529,8.31195408931627,8.202227231735,8.27047843603548,8.4078944190023,8.094295438097,8.11565070577723,8.18827967536181,7.92733387043736,7.77189360186519,7.6658511217778,8.11743521226616,7.88838955898848,8.36379897864502,8.26684802927047,8.69026799903045,8.51455490540219,9.99258017161057,10.4616899838845,10.2359923326615,9.92391366732189,9.54621527135516,8.59338246691469,8.77706731394343,9.95768649499254,9.99985074292847,10.0317496459749,9.8704377409073,9.3398695272487,9.42698531426598,9.45983812949091,9.41590163680278,10.002338291593,9.67607437737509,9.82784470956018\nNLK - mRNA,7.32231819812267,7.38216358549381,7.44074699712431,7.46619563750904,7.55386361600038,7.62787664937962,7.50657468645079,7.45586018194571,7.71886370771172,7.78718973245616,7.66703651470666,7.56895685173481,7.52262975998931,7.4159365069076,7.65125165170435,7.63820526845242,6.50216171810326,6.87468823322278,6.88776562523079,6.95815983729265,6.80487433882539,7.1298777507829,7.13074410521052,7.26897769664494,9.35870807040847,9.15356768852215,9.14852949141116,9.06141719107183,8.37629026991285,8.76330746835701,8.68395790955195,8.94333120201547,8.92635967766194,9.09924616462374,9.11029406798523,9.06050950596846,9.25057326317994,9.14045879323773,9.16097898235263,9.04298027609034,9.11664696876108,9.08657897796035\nRAC3 - mRNA,5.90999508354086,5.90909396484355,6.09971007928925,5.79993303468604,7.7436881748804,7.63240630786013,7.75655293945914,7.60135416651721,7.29565227698717,7.45882307713017,7.84314766056521,7.72663825742238,7.80972787071357,7.84504919139817,7.81892981413018,7.92733387043736,6.40367185470389,6.13207607591543,6.4854391777015,6.71602551209475,6.11407892609211,6.25345243590143,6.36466892241459,6.34149019642931,9.78612929414315,9.22068188438069,8.98983174539211,8.73948909618446,8.54621527135516,9.4007373889723,9.16938473672219,9.74938385675954,9.59403588514536,9.40410074615217,9.41871306155753,9.69952742348694,9.52949377897981,9.60033553700301,9.54839390641038,9.60097572921123,9.49321032022528,8.92738038311109\nLLGL2 - mRNA,3.66206757009728,3.09093228782936,3.83024940429602,3.79993303468604,8.15971348319536,8.21283915010077,8.08719825144761,7.99845665186134,8.23139773526788,8.28514819143502,8.22788655931101,8.19267866013532,8.28061860472593,8.29553977629518,8.27737989624641,8.15069813084113,8.21326524895749,8.08170255340031,8.24749575283053,8.38399253957099,8.2207912621125,8.14736517751174,8.40096505163758,8.38101856061595,7.82265517016826,6.9311752671857,7.56356699069001,7.44998247898948,8.54621527135516,8.76330746835701,8.42092350371815,8.27795969158595,8.25965402749925,8.11974257150933,8.08920003635061,9.19009513330591,8.61426388245856,8.71789274915111,8.75731627293844,8.29932002935013,8.05634645849326,7.60545228822373\nTCF7 - mRNA,5.43967514876083,5.4868609641605,5.23234784786737,5.18326167423754,8.64344650912509,8.63014325635156,8.64879675910326,8.62010425768353,8.83434092513165,8.94194343691724,8.69075258368047,8.61298147562681,8.98005456923237,8.93168787093464,8.8614693590947,8.99707494172792,8.38592145256647,8.35648766488853,8.43827406962282,8.30342705826732,8.31274631252044,8.1298777507829,8.79456202664292,8.73515404491559,6.5007270752809,6.22068188438069,6.14852949141116,6.08741239960477,7.961252770634,7.94701942152939,7.94699231538574,7.22647318129784,7.33731600012897,7.38729245846561,7.46340381812088,8.27835513803599,8.11750198574526,8.19145632503641,8.11848690023855,6.96051811589837,7.74318857323363,7.26841730094616\nDIAPH3 - mRNA,7.82049693270176,7.90099649714495,7.29901888751265,7.50037275282713,9.05211448352821,9.09635340761992,9.01746059640064,9.11760391715435,7.62629758897564,7.68844048304076,8.33711562906577,8.25967296609967,9.46754550221024,9.50727681844602,9.4990240317565,9.45797944976409,8.32719312485653,8.3197030790912,8.89730939790166,8.78571721180965,7.74377974157855,7.80207308955886,6.56190126985264,6.47641977651542,8.74865458872448,8.82733945620116,8.14852949141116,8.52798499099075,10.6616924887751,10.0008862935081,9.71958181928267,9.06974767761039,8.87660664246484,8.92645126957364,8.95080154046224,9.07421442125577,8.86680591447471,8.85909106167483,8.85231278687419,8.85033519814795,8.95618229656783,8.74841024206577\nBAIAP2 - mRNA,7.52359763230592,7.35475742815315,7.59275009056936,7.47112793305127,8.70564127073244,8.8599581268845,8.74924813883591,8.75062865864964,9.02933528716408,9.05754747909581,8.99849787142551,8.92383012463368,8.85598346167444,8.94808230918531,8.76766114381967,8.7597462500866,8.39974615149009,8.23663200695802,8.65254916353675,8.41364581358537,8.66261263985854,8.62407478758087,8.86121106207798,8.87200491312809,7.82265517016826,7.86977472252156,8.20742318046473,7.36752031879751,8.83572188855014,8.70885968433463,8.5319548161069,8.2189656446289,8.06114370005829,7.87647142049696,8.04375706558944,8.26002331526374,8.01432653776986,8.25796877112985,7.9872423669603,8.51691146442275,7.8886190127552,8.37804179212066\nPIK3CB - mRNA,7.2297520793866,7.33885980127295,7.1675527250703,7.26505576540119,9.20635011091855,9.30651554017096,9.29712132082184,9.19189769950166,9.436133500537,9.45997492951941,9.46181137651508,9.4482208188892,9.38258715413313,9.48272921083369,9.41345453736959,9.467370500277,8.81968911296157,8.91771551681733,9.23123633883142,9.15466962432593,9.13382286139052,9.08520870144917,9.01256267886221,9.06281281284302,8.82265517016826,9.00917777918698,8.14852949141116,8.70632223224926,8.83572188855014,8.89919089643878,8.79943512697188,8.62897632683291,8.53734973107506,8.53195070129749,8.57302830929537,8.5468439739619,8.47094651687973,8.62434570027246,8.56933953398706,8.87305527464803,8.56915917345858,8.57507863918021\nMAGI3 - mRNA,6.88261864460956,7.02788011729406,6.59275009056936,6.71931457447775,9.54141187495413,9.51221038720531,9.51843990880697,9.54468675809379,9.19523807197246,9.27863488447634,9.8029087479545,9.88599997003323,9.87942831472372,9.93826801935835,9.84742842061701,9.80025553026146,7.81380055359911,7.82305546480746,8.3713674109927,8.37937591150062,8.60517450334904,8.53830242699009,8.44073558033018,8.51036710843876,8.3080819973385,8.78361807877185,8.65102983194035,8.52798499099075,9.18364519197045,8.44582527850084,8.60995732810817,8.7606313113475,8.72084897294913,8.63046644193856,8.7574528804328,8.55438552297452,8.60166384567892,8.54369504028422,8.60615219960479,8.51691146442275,8.71002170929843,8.63519963161778\nCDX2 - mRNA,3.02463764948198,3.54344449252687,2.92978507784693,3.79993303468604,6.44138888674197,6.35938781345371,6.45811319749415,6.25190500886594,6.53666037649097,6.86229118769688,8.04677387723624,8.16067671863128,7.00237294865597,6.87279418188107,7.00436140184,6.79187045314224,2.46507239936803,2.19661632811014,3.0675866628156,3.18143653388871,3.64895619537695,3.35728824688597,3.39197626841033,2.99099294934518,10.7675136159758,10.4823494544506,10.5177633010769,9.80361943360418,8.961252770634,8.70885968433463,8.68395790955195,9.66533696326974,9.57769520030245,9.64938730383579,9.48906000772596,9.09711482336585,8.97334497576339,9.28745814491376,9.19390791521763,9.39661723070504,10.1664000039582,9.63519963161778\nTGFB2 - mRNA,8.11445591173045,8.03529758885654,8.34256660318541,8.40307555147222,8.04571671233006,8.06653775318436,8.21465916062987,8.08198000742363,8.70246926946105,8.69040466992128,8.42221919744219,8.50879262535484,8.24629853154206,8.2539456870075,8.18269864309851,8.24926196532472,3.73809089377445,3.96215107447312,3.8749415848732,4.04817000302525,4.4473223342073,4.31664626238862,4.79796862808617,4.36950457259891,7.67065207672321,8.00917777918698,8.95588441346877,8.36752031879751,9.28318086552137,8.20481717899704,8.36202981466459,9.0822146350441,9.01334297925658,9.20214191707014,9.09478106861391,8.25632883170599,8.45001224329199,8.4005519237594,8.39744130930276,8.57349499278912,8.15822607251248,8.19041478894489\nCLDN23 - mRNA,8.72661788569886,8.71336949396918,8.77778198440188,8.80580163277518,6.59825773535884,6.4949027841102,6.62704220646274,6.77500225400224,7.27362597065717,7.2615613711174,7.48911872251082,7.36874466524714,7.03984765407463,7.04729191311321,7.15446496236957,6.92733387043736,4.33954151728418,4.57512795136387,4.38951475770296,4.41590178752574,4.74010408343515,4.39681661107261,4.23997317496528,4.12849647309511,9.67065207672321,9.25310336207307,9.08712894674702,9.00797793211036,7.69821836480021,9.36597197081162,9.37698015613056,9.00574380918404,8.99532514419782,9.07729042988506,8.98125212298349,8.98884852084101,9.02143926396443,9.01572903752756,9.043041249591,9.002338291593,9.71560274156173,9.10795262875291\nCLDN7 - mRNA,3.02463764948198,3.4868609641605,3.48232610087571,3.52429859207261,11.498136433612,11.5989501817604,11.4726015660973,11.436008945478,12.2244173722185,12.2742217199742,11.9791673219937,11.9398284127864,11.8379247625442,11.9630830672102,11.8693077456376,11.8106688045735,3.20203799353424,3.87468823322278,3.92556765794317,3.18143653388871,3.98337523444751,3.88780296358475,3.23997317496528,3.08410235373666,11.0085217154796,10.3462127664645,10.0065104865387,9.70632223224926,10.5462152713552,10.8497222201745,10.695930551218,10.7921715432866,10.5176478166325,10.5934928462995,10.5304199588421,10.5904129888993,10.4539606281095,10.5448055175995,10.4297909825436,10.2653726974268,10.4536819560386,10.3868120017484\nCGN - mRNA,3.31414426667697,2.42796727510693,2.83024940429602,2.27637107862903,8.81689802183418,8.72862162311943,8.86021164106549,8.74684205531865,9.65533529386995,9.5460147605861,9.50481252961596,9.53503370966637,9.15390139329236,9.27878654155691,9.22131993712582,9.19235175203997,4.88010989864688,4.68204315528038,4.0675866628156,4.13836781199683,4.5177116620987,3.88780296358475,4.07004817352297,3.94208334886423,10.3833701246427,10.7724775211666,10.2217784734418,10.234253787934,10.1311777720763,10.3541948030353,10.2116955415802,10.5398897763297,10.3810907509806,10.4068831868235,10.3144653993087,10.6126171354004,10.4587718029336,10.5733819250071,10.5326665642586,10.6996037619389,10.6206353909497,10.5282844277013\nCLDN3 - mRNA,2.76160324364819,3.23532219716454,2.72333420037951,2.27637107862903,10.8480248346951,11.0136983608656,10.8597877553445,10.8175924405956,10.9054466306883,11.0491530383536,11.7715121915007,11.7032240769681,11.3096736376831,11.3588342694392,11.2754148175668,11.2221279889189,4.43469875032452,4.42927708490042,5.02178297320247,4.92686370680311,5.85331469388314,5.61362800014575,4.27950153915192,4.94208334886423,9.5007270752809,9.2926317262597,9.62965618114778,8.8346463292248,8.37629026991285,8.25634747963712,8.09899540883079,9.90797302742269,9.63367981007962,9.76160746346496,9.66743667388345,8.86574437615169,8.78558211920074,8.99854110477679,8.84285245762512,9.48776511876324,9.44026144027633,9.28727632819747\nITGB4 - mRNA,3.55515236618076,3.42796727510693,4.15217749918338,4.32076519798748,11.2831444620278,11.2637259339149,11.2842273937579,11.2985642100592,12.0061456597922,11.9280682753863,11.8211552708725,11.7741202438169,11.8185145155979,11.8472360874375,11.7727779137971,11.7768422439099,6.00939291559185,5.69486719563797,6.31551417625918,6.18143653388871,6.23391869609811,6.274826086694,6.32743601621562,6.21338537068163,8.58818991653124,8.0650310139205,8.37092191274761,8.18694807315569,6.961252770634,8.00841996619354,7.42092350371815,8.22647318129784,8.23573540059303,8.13324683585907,8.17389720092589,7.76793959067989,7.90832518391298,7.80079242111016,7.88954569307316,7.82725158503491,8.3852949810584,8.22994315313152\nCA2 - mRNA,3.17664074292704,2.74989536999429,3.54645643829543,2.97681079677012,9.70259439517358,9.64029914918494,9.64486570751334,9.60763131220337,8.2112600896968,8.38312335099832,9.18399891531001,9.05728356941085,9.32935427126961,9.40291249933546,9.50350793603691,9.39047263040582,3.96757273989722,4.33411985186008,3.45990408559436,3.55340531127567,3.21285708057028,3.50929134033102,2.77048789166406,4.03829866412353,9.14458326505562,9.0650310139205,8.73349199213232,9.11294749171191,6.37629026991285,6.51092030672272,6.86453015519377,8.18597381901809,8.29164414174369,8.13548530805507,8.05525270442727,7.74966085696158,7.74318842131971,7.74520009514684,7.8092440649823,7.6279427768115,8.05634645849326,8.10795262875291\nMYH14 - mRNA,3.66206757009728,3.59789227654924,3.72333420037951,3.18326167423754,8.82065015669529,8.70085609473672,8.75837337508351,8.72775850932938,9.74311125395839,9.79633811467034,10.2332798180812,10.2278522001482,9.68911918733444,9.68504790335151,9.60609961531198,9.55476451229897,5.40367185470389,5.41384704433081,5.63744227114655,5.52023844734047,5.91981010567005,5.78727608763078,5.59361012957998,5.78454207187775,7.3080819973385,6.97070363137234,7.2640067088311,6.73948909618446,6.37629026991285,6.81577488825114,5.77706731394343,7.26100086442611,7.05866696976607,7.01178332337341,7.13974897851405,6.75228632636841,6.42698531426598,6.66031119756033,6.69821275259356,6.45801777536919,6.55677544900275,6.19041478894489\nCXCR4 - mRNA,2.85471264803967,3.70098576951335,2.92978507784693,3.59829917351639,9.70868172505249,9.65928960201108,9.63993678218303,9.63861378508629,8.99777247876718,9.01272528148659,9.27440770321484,9.09163507420163,9.53471178374277,9.51610066883303,9.55193245276423,9.53814979844808,5.70028886106207,5.65604794674744,5.65254916353675,5.41590178752574,4.29531924076225,4.50929134033102,5.72825965627476,5.36950457259891,7.08568957600206,7.73853018924331,7.14852949141116,6.86501997826832,6.961252770634,5.59338246691469,5.94699231538574,8.14694452158972,8.25316999392905,8.29818498463163,8.06892764132435,7.77311982978557,8.09520331863649,8.08962282782129,8.03579411179241,7.12098278809162,6.94672196731876,7.10795262875291\n";
	colAnnotation_UyySR3RU_9hQ_hWdumMiueg = "File.Name,Description,Batch.ID,Sample.Name,Cartridge.Id,Lane.Number,Analyte.Types,System.Type,Gene.RLF,Scanner.Id,Stage.Position,Scanned.Date,Gender,QC.Flag,FOV.Count,FOV.Counted,Binding.Density,Is.Reference,Imaging.QC.Flag,Imaging.QC,Binding.Density.QC.Flag,Binding.Density.QC,Positive.Control.QC.Flag,Positive.Control.QC,Limit.of.Detection.QC.Flag.of.mRNA,Limit.of.Detection.QC.of.mRNA,Line,Day,\n20161013_sebelius2-101316_01_09.RCC,Caco2 Control D1,JR2,01,sebelius2-101316,9,mRNA,NONE,SEBELIUS_2_C4636,1507C0300,3,2016-10-13 00:00:00.0,Unknown,false,280,278,0.25,false,false,0.992857,false,0.25,false,0.986733,false,7,Caco2,1,\n20161013_sebelius2-101316_01_10.RCC,Caco2 Control D3,JR2,01,sebelius2-101316,10,mRNA,NONE,SEBELIUS_2_C4636,1507C0300,3,2016-10-13 00:00:00.0,Unknown,false,280,280,0.46,false,false,1,false,0.46,false,0.995246,false,7.824747,Caco2,3,\n20161013_sebelius2-101316_01_11.RCC,Caco2 Control D7,JR2,01,sebelius2-101316,11,mRNA,NONE,SEBELIUS_2_C4636,1507C0300,3,2016-10-13 00:00:00.0,Unknown,false,280,277,0.43,false,false,0.989286,false,0.43,false,0.980532,false,6.75,Caco2,7,\n20161013_sebelius2-101316_01_12.RCC,Caco2 Control D10,JR2,01,sebelius2-101316,12,mRNA,NONE,SEBELIUS_2_C4636,1507C0300,3,2016-10-13 00:00:00.0,Unknown,false,280,280,0.25,false,false,1,false,0.25,false,0.995783,false,5.125,Caco2,10,\n20170307_20170307-caco_01_01.RCC,Caco2 Control D1,JR2,01,20170307-caco,1,mRNA,NONE,SEBELIUS_2_C4636,1507C0300,1,2017-03-07 00:00:00.0,Unknown,false,280,280,0.32,false,false,1,false,0.32,false,0.994436,false,5.875,Caco2,1,\n20170307_20170307-caco_01_02.RCC,Caco2 Control D1,JR2,01,20170307-caco,2,mRNA,NONE,SEBELIUS_2_C4636,1507C0300,1,2017-03-07 00:00:00.0,Unknown,false,280,280,0.31,false,false,1,false,0.31,false,0.992261,false,5.75,Caco2,1,\n20170307_20170307-caco_01_03.RCC,Caco2 Control D4,JR2,01,20170307-caco,3,mRNA,NONE,SEBELIUS_2_C4636,1507C0300,1,2017-03-07 00:00:00.0,Unknown,false,280,280,0.33,false,false,1,false,0.33,false,0.980453,false,6.375,Caco2,4,\n20170307_20170307-caco_01_04.RCC,Caco2 Control D4,JR2,01,20170307-caco,4,mRNA,NONE,SEBELIUS_2_C4636,1507C0300,1,2017-03-07 00:00:00.0,Unknown,false,280,280,0.3,false,false,1,false,0.3,false,0.994702,false,6,Caco2,4,\n20170307_20170307-caco_01_05.RCC,Caco2 Control D7,JR2,01,20170307-caco,5,mRNA,NONE,SEBELIUS_2_C4636,1507C0300,1,2017-03-07 00:00:00.0,Unknown,false,280,280,0.36,false,false,1,false,0.36,false,0.992391,false,5.375,Caco2,7,\n20170307_20170307-caco_01_06.RCC,Caco2 Control D7,JR2,01,20170307-caco,6,mRNA,NONE,SEBELIUS_2_C4636,1507C0300,1,2017-03-07 00:00:00.0,Unknown,false,280,280,0.28,false,false,1,false,0.28,false,0.955249,false,5.625,Caco2,7,\n20180926_Polarization01_Caco2D4A_01.RCC,Caco2 Control D5,ST,Caco2D4A,Polarization01,1,mRNA,NONE,SEBELIUS_2_C4636,1507C0300,5,2018-09-26 00:00:00.0,Unknown,false,555,524,1.8,false,false,0.944144,false,1.8,false,0.995004,false,32.241182,Caco2,5,\n20180926_Polarization01_Caco2D4B_02.RCC,Caco2 Control D5,ST,Caco2D4B,Polarization01,2,mRNA,NONE,SEBELIUS_2_C4636,1507C0300,5,2018-09-26 00:00:00.0,Unknown,true,555,522,2.27,false,false,0.940541,true,2.27,false,0.993594,false,26.813122,Caco2,5,\n20180926_Polarization01_Caco2D4C_03.RCC,Caco2 Control D5,ST,Caco2D4C,Polarization01,3,mRNA,NONE,SEBELIUS_2_C4636,1507C0300,5,2018-09-26 00:00:00.0,Unknown,true,555,513,2.78,false,false,0.924324,true,2.78,false,0.999676,false,25.190455,Caco2,5,\n20180926_Polarization01_Caco2D4D_04.RCC,Caco2 Control D5,ST,Caco2D4D,Polarization01,4,mRNA,NONE,SEBELIUS_2_C4636,1507C0300,5,2018-09-26 00:00:00.0,Unknown,true,555,506,2.38,false,false,0.911712,true,2.38,false,0.997102,false,25.269016,Caco2,5,\n20180926_Polarization01_FibD4A_05.RCC,Fibro Control D5,ST,FibD4A,Polarization01,5,mRNA,NONE,SEBELIUS_2_C4636,1507C0300,5,2018-09-26 00:00:00.0,Unknown,false,555,495,1.49,false,false,0.891892,false,1.49,false,0.99523,false,26.458078,Fibro,5,\n20180926_Polarization01_FibD4B_06.RCC,Fibro Control D5,ST,FibD4B,Polarization01,6,mRNA,NONE,SEBELIUS_2_C4636,1507C0300,5,2018-09-26 00:00:00.0,Unknown,false,555,491,1.15,false,false,0.884685,false,1.15,false,0.995856,false,23.642375,Fibro,5,\n20180926_Polarization01_HT29D4A_07.RCC,HT29 Control D5,ST,HT29D4A,Polarization01,7,mRNA,NONE,SEBELIUS_2_C4636,1507C0300,5,2018-09-26 00:00:00.0,Unknown,false,555,494,2.25,false,false,0.89009,false,2.25,false,0.99817,false,31.435711,HT29,5,\n20180926_Polarization01_HT29D4B_08.RCC,HT29 Control D5,ST,HT29D4B,Polarization01,8,mRNA,NONE,SEBELIUS_2_C4636,1507C0300,5,2018-09-26 00:00:00.0,Unknown,true,555,487,2.45,false,false,0.877477,true,2.45,false,0.997999,false,38.69426,HT29,5,\n20180926_Polarization01_HT29D4C_09.RCC,HT29 Control D5,ST,HT29D4C,Polarization01,9,mRNA,NONE,SEBELIUS_2_C4636,1507C0300,5,2018-09-26 00:00:00.0,Unknown,true,555,481,2.67,false,false,0.866667,true,2.67,false,0.997853,false,28.699056,HT29,5,\n20180926_Polarization01_HT29D4D_10.RCC,HT29 Control D5,ST,HT29D4D,Polarization01,10,mRNA,NONE,SEBELIUS_2_C4636,1507C0300,5,2018-09-26 00:00:00.0,Unknown,true,555,479,2.47,false,false,0.863063,true,2.47,false,0.995337,false,35.329784,HT29,5,\n20180926_Polarization01_HT29D4E_11.RCC,HT29 Control D4,ST,HT29D4E,Polarization01,11,mRNA,NONE,SEBELIUS_2_C4636,1507C0300,5,2018-09-26 00:00:00.0,Unknown,true,555,453,2.78,false,false,0.816216,true,2.78,false,0.998514,false,31.425946,HT29,4,\n20180926_Polarization01_HT29D4F_12.RCC,HT29 Control D4,ST,HT29D4F,Polarization01,12,mRNA,NONE,SEBELIUS_2_C4636,1507C0300,5,2018-09-26 00:00:00.0,Unknown,true,555,452,2.33,false,false,0.814414,true,2.33,false,0.997126,false,20.743505,HT29,4,\n20180928_Polarization-02_Caco2D7A_01.RCC,Caco2 Control D8,ST,Caco2D7A,Polarization-02,1,mRNA,NONE,SEBELIUS_2_C4636,1507C0300,2,2018-09-27 00:00:00.0,Unknown,true,555,534,2.53,false,false,0.962162,true,2.53,false,0.995977,false,22.474098,Caco2,8,\n20180928_Polarization-02_Caco2D7B_02.RCC,Caco2 Control D8,ST,Caco2D7B,Polarization-02,2,mRNA,NONE,SEBELIUS_2_C4636,1507C0300,2,2018-09-27 00:00:00.0,Unknown,false,555,533,1.92,false,false,0.96036,false,1.92,false,0.994538,false,31.436929,Caco2,8,\n20180928_Polarization-02_Caco2D7C_03.RCC,Caco2 Control D8,ST,Caco2D7C,Polarization-02,3,mRNA,NONE,SEBELIUS_2_C4636,1507C0300,2,2018-09-27 00:00:00.0,Unknown,false,555,535,1.34,false,false,0.963964,false,1.34,false,0.997677,false,34.142351,Caco2,8,\n20180928_Polarization-02_Caco2D7D_04.RCC,Caco2 Control D8,ST,Caco2D7D,Polarization-02,4,mRNA,NONE,SEBELIUS_2_C4636,1507C0300,2,2018-09-27 00:00:00.0,Unknown,false,555,533,2.05,false,false,0.96036,false,2.05,false,0.995298,false,25.601672,Caco2,8,\n20180928_Polarization-02_FibD7A_05.RCC,Fibro Control D8,ST,FibD7A,Polarization-02,5,mRNA,NONE,SEBELIUS_2_C4636,1507C0300,2,2018-09-27 00:00:00.0,Unknown,false,555,539,1.58,false,false,0.971171,false,1.58,false,0.996031,false,27.367248,Fibro,8,\n20180928_Polarization-02_FibD7B_06.RCC,Fibro Control D8,ST,FibD7B,Polarization-02,6,mRNA,NONE,SEBELIUS_2_C4636,1507C0300,2,2018-09-27 00:00:00.0,Unknown,false,555,532,1.47,false,false,0.958559,false,1.47,false,0.995629,false,31.034423,Fibro,8,\n20180928_Polarization-02_HT29D7A_07.RCC,HT29 Control D8,ST,HT29D7A,Polarization-02,7,mRNA,NONE,SEBELIUS_2_C4636,1507C0300,2,2018-09-27 00:00:00.0,Unknown,false,555,527,1.64,false,false,0.94955,false,1.64,false,0.995438,false,29.61076,HT29,8,\n20180928_Polarization-02_HT29D7B_08.RCC,HT29 Control D8,ST,HT29D7B,Polarization-02,8,mRNA,NONE,SEBELIUS_2_C4636,1507C0300,2,2018-09-27 00:00:00.0,Unknown,false,555,527,1.61,false,false,0.94955,false,1.61,false,0.990571,false,22.324298,HT29,8,\n20180928_Polarization-02_HT29D7C_09.RCC,HT29 Control D8,ST,HT29D7C,Polarization-02,9,mRNA,NONE,SEBELIUS_2_C4636,1507C0300,2,2018-09-27 00:00:00.0,Unknown,false,555,529,1.89,false,false,0.953153,false,1.89,false,0.997566,false,28.893882,HT29,8,\n20180928_Polarization-02_HT29D7D_10.RCC,HT29 Control D8,ST,HT29D7D,Polarization-02,10,mRNA,NONE,SEBELIUS_2_C4636,1507C0300,2,2018-09-27 00:00:00.0,Unknown,false,555,531,2.16,false,false,0.956757,false,2.16,false,0.992868,false,27.913127,HT29,8,\n20180928_Polarization-02_HT29D7E_11.RCC,HT29 Control D7,ST,HT29D7E,Polarization-02,11,mRNA,NONE,SEBELIUS_2_C4636,1507C0300,2,2018-09-27 00:00:00.0,Unknown,false,555,527,1.96,false,false,0.94955,false,1.96,false,0.995349,false,21.680057,HT29,7,\n20180928_Polarization-02_HT29D7F_12.RCC,HT29 Control D7,ST,HT29D7F,Polarization-02,12,mRNA,NONE,SEBELIUS_2_C4636,1507C0300,2,2018-09-27 00:00:00.0,Unknown,false,555,528,1.85,false,false,0.951351,false,1.85,false,0.994969,false,25.465345,HT29,7,\n20181004_Pol03_HepG2D4A_01.RCC,HEPG2 Control D5,ST,HepG2D4A,Pol03,1,mRNA,NONE,SEBELIUS_2_C4636,1507C0300,1,2018-10-04 00:00:00.0,Unknown,false,555,544,1.97,false,false,0.98018,false,1.97,false,0.994053,false,29.354049,HEPG2,5,\n20181004_Pol03_HepG2D4B_02.RCC,HEPG2 Control D5,ST,HepG2D4B,Pol03,2,mRNA,NONE,SEBELIUS_2_C4636,1507C0300,1,2018-10-04 00:00:00.0,Unknown,false,555,541,2.07,false,false,0.974775,false,2.07,false,0.999207,false,25.678388,HEPG2,5,\n20181004_Pol03_HepG2D4C_03.RCC,HEPG2 Control D4,ST,HepG2D4C,Pol03,3,mRNA,NONE,SEBELIUS_2_C4636,1507C0300,1,2018-10-04 00:00:00.0,Unknown,false,555,538,1.74,false,false,0.969369,false,1.74,false,0.99564,false,18.485281,HEPG2,4,\n20181004_Pol03_HepG2D4D_04.RCC,HEPG2 Control D4,ST,HepG2D4D,Pol03,4,mRNA,NONE,SEBELIUS_2_C4636,1507C0300,1,2018-10-04 00:00:00.0,Unknown,false,555,538,2.12,false,false,0.969369,false,2.12,false,0.994346,false,26.852592,HEPG2,4,\n20181004_Pol03_HepG2D7A_05.RCC,HEPG2 Control D8,ST,HepG2D7A,Pol03,5,mRNA,NONE,SEBELIUS_2_C4636,1507C0300,1,2018-10-04 00:00:00.0,Unknown,false,555,540,1.96,false,false,0.972973,false,1.96,false,0.998224,false,25.136644,HEPG2,8,\n20181004_Pol03_HepG2D7B_06.RCC,HEPG2 Control D8,ST,HepG2D7B,Pol03,6,mRNA,NONE,SEBELIUS_2_C4636,1507C0300,1,2018-10-04 00:00:00.0,Unknown,false,555,534,2,false,false,0.962162,false,2,false,0.996591,false,29.761011,HEPG2,8,\n20181004_Pol03_HepG2D7C_07.RCC,HEPG2 Control D7,ST,HepG2D7C,Pol03,7,mRNA,NONE,SEBELIUS_2_C4636,1507C0300,1,2018-10-04 00:00:00.0,Unknown,false,555,532,2.07,false,false,0.958559,false,2.07,false,0.99712,false,22.973013,HEPG2,7,\n20181004_Pol03_HepG2D7D_08.RCC,HEPG2 Control D7,ST,HepG2D7D,Pol03,8,mRNA,NONE,SEBELIUS_2_C4636,1507C0300,1,2018-10-04 00:00:00.0,Unknown,false,555,531,1.96,false,false,0.956757,false,1.96,false,0.995664,false,24.508926,HEPG2,7,\n";
```
