## Supplementary material for "Comparative pathway enrichment analysis in gastrointestinal cell lines Caco-2, HT-29, HEPG2, and colon fibroblasts using a custom expression panel for tight-junction and cytoskeletal regulatory genes": Sup. Results 1: d3.min.js

```
// https://d3js.org Version 4.2.2. Copyright 2016 Mike Bostock.
!function(t,n){"object"==typeof exports&&"undefined"!=typeof module?n(exports):"function"==typeof define&&define.amd?define(["exports"],n):n(t.d3=t.d3||{})}(this,function(t){"use strict";function n(t,n){return t<n?-1:t>n?1:t>=n?0:NaN}function e(t){return 1===t.length&&(t=r(t)),{left:function(n,e,r,i){for(null==r&&(r=0),null==i&&(i=n.length);r<i;){var o=r+i>>>1;t(n[o],e)<0?r=o+1:i=o}return r},right:function(n,e,r,i){for(null==r&&(r=0),null==i&&(i=n.length);r<i;){var o=r+i>>>1;t(n[o],e)>0?i=o:r=o+1}return r}}}function r(t){return function(e,r){return n(t(e),r)}}function i(t,n){return n<t?-1:n>t?1:n>=t?0:NaN}function o(t){return null===t?NaN:+t}function u(t,n){var e,r,i=t.length,u=0,a=0,c=-1,s=0;if(null==n)for(;++c<i;)isNaN(e=o(t[c]))||(r=e-u,u+=r/++s,a+=r*(e-u));else for(;++c<i;)isNaN(e=o(n(t[c],c,t)))||(r=e-u,u+=r/++s,a+=r*(e-u));if(s>1)return a/(s-1)}function a(t,n){var e=u(t,n);return e?Math.sqrt(e):e}function c(t,n){var e,r,i,o=-1,u=t.length;if(null==n){for(;++o<u;)if(null!=(r=t[o])&&r>=r){e=i=r;break}for(;++o<u;)null!=(r=t[o])&&(e>r&&(e=r),i<r&&(i=r))}else{for(;++o<u;)if(null!=(r=n(t[o],o,t))&&r>=r){e=i=r;break}for(;++o<u;)null!=(r=n(t[o],o,t))&&(e>r&&(e=r),i<r&&(i=r))}return[e,i]}function s(t){return function(){return t}}function f(t){return t}function l(t,n,e){t=+t,n=+n,e=(i=arguments.length)<2?(n=t,t=0,1):i<3?1:+e;for(var r=-1,i=0|Math.max(0,Math.ceil((n-t)/e)),o=new Array(i);++r<i;)o[r]=t+r*e;return o}function h(t,n,e){var r=p(t,n,e);return l(Math.ceil(t/r)*r,Math.floor(n/r)*r+r/2,r)}function p(t,n,e){var r=Math.abs(n-t)/Math.max(0,e),i=Math.pow(10,Math.floor(Math.log(r)/Math.LN10)),o=r/i;return o>=Rd?i*=10:o>=Ud?i*=5:o>=Dd&&(i*=2),n<t?-i:i}function d(t){return Math.ceil(Math.log(t.length)/Math.LN2)+1}function v(){function t(t){var i,o,u=t.length,a=new Array(u);for(i=0;i<u;++i)a[i]=n(t[i],i,t);var c=e(a),s=c[0],f=c[1],l=r(a,s,f);Array.isArray(l)||(l=h(s,f,l));for(var p=l.length;l[0]<=s;)l.shift(),--p;for(;l[p-1]>=f;)l.pop(),--p;var d,v=new Array(p+1);for(i=0;i<=p;++i)d=v[i]=[],d.x0=i>0?l[i-1]:s,d.x1=i<p?l[i]:f;for(i=0;i<u;++i)o=a[i],s<=o&&o<=f&&v[Cd(l,o,0,p)].push(t[i]);return v}var n=f,e=c,r=d;return t.value=function(e){return arguments.length?(n="function"==typeof e?e:s(e),t):n},t.domain=function(n){return arguments.length?(e="function"==typeof n?n:s([n[0],n[1]]),t):e},t.thresholds=function(n){return arguments.length?(r="function"==typeof n?n:s(Array.isArray(n)?qd.call(n):n),t):r},t}function _(t,n,e){if(null==e&&(e=o),r=t.length){if((n=+n)<=0||r<2)return+e(t[0],0,t);if(n>=1)return+e(t[r-1],r-1,t);var r,i=(r-1)*n,u=Math.floor(i),a=+e(t[u],u,t),c=+e(t[u+1],u+1,t);return a+(c-a)*(i-u)}}function y(t,e,r){return t=Ld.call(t,o).sort(n),Math.ceil((r-e)/(2*(_(t,.75)-_(t,.25))*Math.pow(t.length,-1/3)))}function g(t,n,e){return Math.ceil((e-n)/(3.5*a(t)*Math.pow(t.length,-1/3)))}function m(t,n){var e,r,i=-1,o=t.length;if(null==n){for(;++i<o;)if(null!=(r=t[i])&&r>=r){e=r;break}for(;++i<o;)null!=(r=t[i])&&r>e&&(e=r)}else{for(;++i<o;)if(null!=(r=n(t[i],i,t))&&r>=r){e=r;break}for(;++i<o;)null!=(r=n(t[i],i,t))&&r>e&&(e=r)}return e}function x(t,n){var e,r=0,i=t.length,u=-1,a=i;if(null==n)for(;++u<i;)isNaN(e=o(t[u]))?--a:r+=e;else for(;++u<i;)isNaN(e=o(n(t[u],u,t)))?--a:r+=e;if(a)return r/a}function b(t,e){var r,i=[],u=t.length,a=-1;if(null==e)for(;++a<u;)isNaN(r=o(t[a]))||i.push(r);else for(;++a<u;)isNaN(r=o(e(t[a],a,t)))||i.push(r);return _(i.sort(n),.5)}function w(t){for(var n,e,r,i=t.length,o=-1,u=0;++o<i;)u+=t[o].length;for(e=new Array(u);--i>=0;)for(r=t[i],n=r.length;--n>=0;)e[--u]=r[n];return e}function M(t,n){var e,r,i=-1,o=t.length;if(null==n){for(;++i<o;)if(null!=(r=t[i])&&r>=r){e=r;break}for(;++i<o;)null!=(r=t[i])&&e>r&&(e=r)}else{for(;++i<o;)if(null!=(r=n(t[i],i,t))&&r>=r){e=r;break}for(;++i<o;)null!=(r=n(t[i],i,t))&&e>r&&(e=r)}return e}function T(t){for(var n=0,e=t.length-1,r=t[0],i=new Array(e<0?0:e);n<e;)i[n]=[r,r=t[++n]];return i}function k(t,n){for(var e=n.length,r=new Array(e);e--;)r[e]=t[n[e]];return r}function S(t,e){if(r=t.length){var r,i,o=0,u=0,a=t[u];for(e||(e=n);++o<r;)(e(i=t[o],a)<0||0!==e(a,a))&&(a=i,u=o);return 0===e(a,a)?u:void 0}}function N(t,n,e){for(var r,i,o=(null==e?t.length:e)-(n=null==n?0:+n);o;)i=Math.random()*o--|0,r=t[o+n],t[o+n]=t[i+n],t[i+n]=r;return t}function A(t,n){var e,r=0,i=t.length,o=-1;if(null==n)for(;++o<i;)(e=+t[o])&&(r+=e);else for(;++o<i;)(e=+n(t[o],o,t))&&(r+=e);return r}function E(t){if(!(i=t.length))return[];for(var n=-1,e=M(t,C),r=new Array(e);++n<e;)for(var i,o=-1,u=r[n]=new Array(i);++o<i;)u[o]=t[o][n];return r}function C(t){return t.length}function z(){return E(arguments)}function P(){}function q(t,n){var e=new P;if(t instanceof P)t.each(function(t,n){e.set(n,t)});else if(Array.isArray(t)){var r,i=-1,o=t.length;if(null==n)for(;++i<o;)e.set(i,t[i]);else for(;++i<o;)e.set(n(r=t[i],i,t),r)}else if(t)for(var u in t)e.set(u,t[u]);return e}function L(){function t(n,i,u,a){if(i>=o.length)return null!=r?r(n):null!=e?n.sort(e):n;for(var c,s,f,l=-1,h=n.length,p=o[i++],d=q(),v=u();++l<h;)(f=d.get(c=p(s=n[l])+""))?f.push(s):d.set(c,[s]);return d.each(function(n,e){a(v,e,t(n,i,u,a))}),v}function n(t,e){if(++e>o.length)return t;var i,a=u[e-1];return null!=r&&e>=o.length?i=t.entries():(i=[],t.each(function(t,r){i.push({key:r,values:n(t,e)})})),null!=a?i.sort(function(t,n){return a(t.key,n.key)}):i}var e,r,i,o=[],u=[];return i={object:function(n){return t(n,0,R,U)},map:function(n){return t(n,0,D,O)},entries:function(e){return n(t(e,0,D,O),0)},key:function(t){return o.push(t),i},sortKeys:function(t){return u[o.length-1]=t,i},sortValues:function(t){return e=t,i},rollup:function(t){return r=t,i}}}function R(){return{}}function U(t,n,e){t[n]=e}function D(){return q()}function O(t,n,e){t.set(n,e)}function F(){}function I(t,n){var e=new F;if(t instanceof F)t.each(function(t){e.add(t)});else if(t){var r=-1,i=t.length;if(null==n)for(;++r<i;)e.add(t[r]);else for(;++r<i;)e.add(n(t[r],r,t))}return e}function Y(t){var n=[];for(var e in t)n.push(e);return n}function B(t){var n=[];for(var e in t)n.push(t[e]);return n}function j(t){var n=[];for(var e in t)n.push({key:e,value:t[e]});return n}function H(t,n){return t=null==t?0:+t,n=null==n?1:+n,1===arguments.length?(n=t,t=0):n-=t,function(){return Math.random()*n+t}}function X(t,n){var e,r;return t=null==t?0:+t,n=null==n?1:+n,function(){var i;if(null!=e)i=e,e=null;else do e=2*Math.random()-1,i=2*Math.random()-1,r=e*e+i*i;while(!r||r>1);return t+n*i*Math.sqrt(-2*Math.log(r)/r)}}function V(){var t=X.apply(this,arguments);return function(){return Math.exp(t())}}function W(t){return function(){for(var n=0,e=0;e<t;++e)n+=Math.random();return n}}function $(t){var n=W(t);return function(){return n()/t}}function Z(t){return function(){return-Math.log(1-Math.random())/t}}function G(t){return+t}function J(t){return t*t}function Q(t){return t*(2-t)}function K(t){return((t*=2)<=1?t*t:--t*(2-t)+1)/2}function tt(t){return t*t*t}function nt(t){return--t*t*t+1}function et(t){return((t*=2)<=1?t*t*t:(t-=2)*t*t+2)/2}function rt(t){return 1-Math.cos(t*Xd)}function it(t){return Math.sin(t*Xd)}function ot(t){return(1-Math.cos(Hd*t))/2}function ut(t){return Math.pow(2,10*t-10)}function at(t){return 1-Math.pow(2,-10*t)}function ct(t){return((t*=2)<=1?Math.pow(2,10*t-10):2-Math.pow(2,10-10*t))/2}function st(t){return 1-Math.sqrt(1-t*t)}function ft(t){return Math.sqrt(1- --t*t)}function lt(t){return((t*=2)<=1?1-Math.sqrt(1-t*t):Math.sqrt(1-(t-=2)*t)+1)/2}function ht(t){return 1-pt(1-t)}function pt(t){return(t=+t)<Vd?nv*t*t:t<$d?nv*(t-=Wd)*t+Zd:t<Jd?nv*(t-=Gd)*t+Qd:nv*(t-=Kd)*t+tv}function dt(t){return((t*=2)<=1?1-pt(1-t):pt(t-1)+1)/2}function vt(t){for(var n,e=-1,r=t.length,i=t[r-1],o=0;++e<r;)n=i,i=t[e],o+=n[1]*i[0]-n[0]*i[1];return o/2}function _t(t){for(var n,e,r=-1,i=t.length,o=0,u=0,a=t[i-1],c=0;++r<i;)n=a,a=t[r],c+=e=n[0]*a[1]-a[0]*n[1],o+=(n[0]+a[0])*e,u+=(n[1]+a[1])*e;return c*=3,[o/c,u/c]}function yt(t,n,e){return(n[0]-t[0])*(e[1]-t[1])-(n[1]-t[1])*(e[0]-t[0])}function gt(t,n){return t[0]-n[0]||t[1]-n[1]}function mt(t){for(var n=t.length,e=[0,1],r=2,i=2;i<n;++i){for(;r>1&&yt(t[e[r-2]],t[e[r-1]],t[i])<=0;)--r;e[r++]=i}return e.slice(0,r)}function xt(t){if((e=t.length)<3)return null;var n,e,r=new Array(e),i=new Array(e);for(n=0;n<e;++n)r[n]=[+t[n][0],+t[n][1],n];for(r.sort(gt),n=0;n<e;++n)i[n]=[r[n][0],-r[n][1]];var o=mt(r),u=mt(i),a=u[0]===o[0],c=u[u.length-1]===o[o.length-1],s=[];for(n=o.length-1;n>=0;--n)s.push(t[r[o[n]][2]]);for(n=+a;n<u.length-c;++n)s.push(t[r[u[n]][2]]);return s}function bt(t,n){for(var e,r,i=t.length,o=t[i-1],u=n[0],a=n[1],c=o[0],s=o[1],f=!1,l=0;l<i;++l)o=t[l],e=o[0],r=o[1],r>a!=s>a&&u<(c-e)*(a-r)/(s-r)+e&&(f=!f),c=e,s=r;return f}function wt(t){for(var n,e,r=-1,i=t.length,o=t[i-1],u=o[0],a=o[1],c=0;++r<i;)n=u,e=a,o=t[r],u=o[0],a=o[1],n-=u,e-=a,c+=Math.sqrt(n*n+e*e);return c}function Mt(){this._x0=this._y0=this._x1=this._y1=null,this._=[]}function Tt(){return new Mt}function kt(t){var n=+this._x.call(null,t),e=+this._y.call(null,t);return St(this.cover(n,e),n,e,t)}function St(t,n,e,r){if(isNaN(n)||isNaN(e))return t;var i,o,u,a,c,s,f,l,h,p=t._root,d={data:r},v=t._x0,_=t._y0,y=t._x1,g=t._y1;if(!p)return t._root=d,t;for(;p.length;)if((s=n>=(o=(v+y)/2))?v=o:y=o,(f=e>=(u=(_+g)/2))?_=u:g=u,i=p,!(p=p[l=f<<1|s]))return i[l]=d,t;if(a=+t._x.call(null,p.data),c=+t._y.call(null,p.data),n===a&&e===c)return d.next=p,i?i[l]=d:t._root=d,t;do i=i?i[l]=new Array(4):t._root=new Array(4),(s=n>=(o=(v+y)/2))?v=o:y=o,(f=e>=(u=(_+g)/2))?_=u:g=u;while((l=f<<1|s)===(h=(c>=u)<<1|a>=o));return i[h]=p,i[l]=d,t}function Nt(t){var n,e,r,i,o=t.length,u=new Array(o),a=new Array(o),c=1/0,s=1/0,f=-(1/0),l=-(1/0);for(e=0;e<o;++e)isNaN(r=+this._x.call(null,n=t[e]))||isNaN(i=+this._y.call(null,n))||(u[e]=r,a[e]=i,r<c&&(c=r),r>f&&(f=r),i<s&&(s=i),i>l&&(l=i));for(f<c&&(c=this._x0,f=this._x1),l<s&&(s=this._y0,l=this._y1),this.cover(c,s).cover(f,l),e=0;e<o;++e)St(this,u[e],a[e],t[e]);return this}function At(t,n){if(isNaN(t=+t)||isNaN(n=+n))return this;var e=this._x0,r=this._y0,i=this._x1,o=this._y1;if(isNaN(e))i=(e=Math.floor(t))+1,o=(r=Math.floor(n))+1;else{if(!(e>t||t>i||r>n||n>o))return this;var u,a,c=i-e,s=this._root;switch(a=(n<(r+o)/2)<<1|t<(e+i)/2){case 0:do u=new Array(4),u[a]=s,s=u;while(c*=2,i=e+c,o=r+c,t>i||n>o);break;case 1:do u=new Array(4),u[a]=s,s=u;while(c*=2,e=i-c,o=r+c,e>t||n>o);break;case 2:do u=new Array(4),u[a]=s,s=u;while(c*=2,i=e+c,r=o-c,t>i||r>n);break;case 3:do u=new Array(4),u[a]=s,s=u;while(c*=2,e=i-c,r=o-c,e>t||r>n)}this._root&&this._root.length&&(this._root=s)}return this._x0=e,this._y0=r,this._x1=i,this._y1=o,this}function Et(){var t=[];return this.visit(function(n){if(!n.length)do t.push(n.data);while(n=n.next)}),t}function Ct(t){return arguments.length?this.cover(+t[0][0],+t[0][1]).cover(+t[1][0],+t[1][1]):isNaN(this._x0)?void 0:[[this._x0,this._y0],[this._x1,this._y1]]}function zt(t,n,e,r,i){this.node=t,this.x0=n,this.y0=e,this.x1=r,this.y1=i}function Pt(t,n,e){var r,i,o,u,a,c,s,f=this._x0,l=this._y0,h=this._x1,p=this._y1,d=[],v=this._root;for(v&&d.push(new zt(v,f,l,h,p)),null==e?e=1/0:(f=t-e,l=n-e,h=t+e,p=n+e,e*=e);c=d.pop();)if(!(!(v=c.node)||(i=c.x0)>h||(o=c.y0)>p||(u=c.x1)<f||(a=c.y1)<l))if(v.length){var _=(i+u)/2,y=(o+a)/2;d.push(new zt(v[3],_,y,u,a),new zt(v[2],i,y,_,a),new zt(v[1],_,o,u,y),new zt(v[0],i,o,_,y)),(s=(n>=y)<<1|t>=_)&&(c=d[d.length-1],d[d.length-1]=d[d.length-1-s],d[d.length-1-s]=c)}else{var g=t-+this._x.call(null,v.data),m=n-+this._y.call(null,v.data),x=g*g+m*m;if(x<e){var b=Math.sqrt(e=x);f=t-b,l=n-b,h=t+b,p=n+b,r=v.data}}return r}function qt(t){if(isNaN(o=+this._x.call(null,t))||isNaN(u=+this._y.call(null,t)))return this;var n,e,r,i,o,u,a,c,s,f,l,h,p=this._root,d=this._x0,v=this._y0,_=this._x1,y=this._y1;if(!p)return this;if(p.length)for(;;){if((s=o>=(a=(d+_)/2))?d=a:_=a,(f=u>=(c=(v+y)/2))?v=c:y=c,n=p,!(p=p[l=f<<1|s]))return this;if(!p.length)break;(n[l+1&3]||n[l+2&3]||n[l+3&3])&&(e=n,h=l)}for(;p.data!==t;)if(r=p,!(p=p.next))return this;return(i=p.next)&&delete p.next,r?(i?r.next=i:delete r.next,this):n?(i?n[l]=i:delete n[l],(p=n[0]||n[1]||n[2]||n[3])&&p===(n[3]||n[2]||n[1]||n[0])&&!p.length&&(e?e[h]=p:this._root=p),this):(this._root=i,this)}function Lt(t){for(var n=0,e=t.length;n<e;++n)this.remove(t[n]);return this}function Rt(){return this._root}function Ut(){var t=0;return this.visit(function(n){if(!n.length)do++t;while(n=n.next)}),t}function Dt(t){var n,e,r,i,o,u,a=[],c=this._root;for(c&&a.push(new zt(c,this._x0,this._y0,this._x1,this._y1));n=a.pop();)if(!t(c=n.node,r=n.x0,i=n.y0,o=n.x1,u=n.y1)&&c.length){var s=(r+o)/2,f=(i+u)/2;(e=c[3])&&a.push(new zt(e,s,f,o,u)),(e=c[2])&&a.push(new zt(e,r,f,s,u)),(e=c[1])&&a.push(new zt(e,s,i,o,f)),(e=c[0])&&a.push(new zt(e,r,i,s,f))}return this}function Ot(t){var n,e=[],r=[];for(this._root&&e.push(new zt(this._root,this._x0,this._y0,this._x1,this._y1));n=e.pop();){var i=n.node;if(i.length){var o,u=n.x0,a=n.y0,c=n.x1,s=n.y1,f=(u+c)/2,l=(a+s)/2;(o=i[0])&&e.push(new zt(o,u,a,f,l)),(o=i[1])&&e.push(new zt(o,f,a,c,l)),(o=i[2])&&e.push(new zt(o,u,l,f,s)),(o=i[3])&&e.push(new zt(o,f,l,c,s))}r.push(n)}for(;n=r.pop();)t(n.node,n.x0,n.y0,n.x1,n.y1);return this}function Ft(t){return t[0]}function It(t){return arguments.length?(this._x=t,this):this._x}function Yt(t){return t[1]}function Bt(t){return arguments.length?(this._y=t,this):this._y}function jt(t,n,e){var r=new Ht(null==n?Ft:n,null==e?Yt:e,NaN,NaN,NaN,NaN);return null==t?r:r.addAll(t)}function Ht(t,n,e,r,i,o){this._x=t,this._y=n,this._x0=e,this._y0=r,this._x1=i,this._y1=o,this._root=void 0}function Xt(t){for(var n={data:t.data},e=n;t=t.next;)e=e.next={data:t.data};return n}function Vt(t){if(!(t>=1))throw new Error;this._size=t,this._call=this._error=null,this._tasks=[],this._data=[],this._waiting=this._active=this._ended=this._start=0}function Wt(t){if(!t._start)try{$t(t)}catch(n){t._tasks[t._ended+t._active-1]&&Gt(t,n)}}function $t(t){for(;t._start=t._waiting&&t._active<t._size;){var n=t._ended+t._active,e=t._tasks[n],r=e.length-1,i=e[r];e[r]=Zt(t,n),--t._waiting,++t._active,e=i.apply(null,e),t._tasks[n]&&(t._tasks[n]=e||gv)}}function Zt(t,n){return function(e,r){t._tasks[n]&&(--t._active,++t._ended,t._tasks[n]=null,null==t._error&&(null!=e?Gt(t,e):(t._data[n]=r,t._waiting?Wt(t):Jt(t))))}}function Gt(t,n){var e,r=t._tasks.length;for(t._error=n,t._data=void 0,t._waiting=NaN;--r>=0;)if((e=t._tasks[r])&&(t._tasks[r]=null,e.abort))try{e.abort()}catch(t){}t._active=NaN,Jt(t)}function Jt(t){!t._active&&t._call&&t._call(t._error,t._data)}function Qt(t){return new Vt(arguments.length?+t:1/0)}function Kt(t){return function(){return t}}function tn(t){return t.innerRadius}function nn(t){return t.outerRadius}function en(t){return t.startAngle}function rn(t){return t.endAngle}function on(t){return t&&t.padAngle}function un(t){return t>=1?bv:t<=-1?-bv:Math.asin(t)}function an(t,n,e,r,i,o,u,a){var c=e-t,s=r-n,f=u-i,l=a-o,h=(f*(n-o)-l*(t-i))/(l*c-f*s);return[t+h*c,n+h*s]}function cn(t,n,e,r,i,o,u){var a=t-e,c=n-r,s=(u?o:-o)/Math.sqrt(a*a+c*c),f=s*c,l=-s*a,h=t+f,p=n+l,d=e+f,v=r+l,_=(h+d)/2,y=(p+v)/2,g=d-h,m=v-p,x=g*g+m*m,b=i-o,w=h*v-d*p,M=(m<0?-1:1)*Math.sqrt(Math.max(0,b*b*x-w*w)),T=(w*m-g*M)/x,k=(-w*g-m*M)/x,S=(w*m+g*M)/x,N=(-w*g+m*M)/x,A=T-_,E=k-y,C=S-_,z=N-y;return A*A+E*E>C*C+z*z&&(T=S,k=N),{cx:T,cy:k,x01:-f,y01:-l,x11:T*(i/b-1),y11:k*(i/b-1)}}function sn(){function t(){var t,s,f=+n.apply(this,arguments),l=+e.apply(this,arguments),h=o.apply(this,arguments)-bv,p=u.apply(this,arguments)-bv,d=Math.abs(p-h),v=p>h;if(c||(c=t=Tt()),l<f&&(s=l,l=f,f=s),l>mv)if(d>wv-mv)c.moveTo(l*Math.cos(h),l*Math.sin(h)),c.arc(0,0,l,h,p,!v),f>mv&&(c.moveTo(f*Math.cos(p),f*Math.sin(p)),c.arc(0,0,f,p,h,v));else{var _,y,g=h,m=p,x=h,b=p,w=d,M=d,T=a.apply(this,arguments)/2,k=T>mv&&(i?+i.apply(this,arguments):Math.sqrt(f*f+l*l)),S=Math.min(Math.abs(l-f)/2,+r.apply(this,arguments)),N=S,A=S;if(k>mv){var E=un(k/f*Math.sin(T)),C=un(k/l*Math.sin(T));(w-=2*E)>mv?(E*=v?1:-1,x+=E,b-=E):(w=0,x=b=(h+p)/2),(M-=2*C)>mv?(C*=v?1:-1,g+=C,m-=C):(M=0,g=m=(h+p)/2)}var z=l*Math.cos(g),P=l*Math.sin(g),q=f*Math.cos(b),L=f*Math.sin(b);if(S>mv){var R=l*Math.cos(m),U=l*Math.sin(m),D=f*Math.cos(x),O=f*Math.sin(x);if(d<xv){var F=w>mv?an(z,P,D,O,R,U,q,L):[q,L],I=z-F[0],Y=P-F[1],B=R-F[0],j=U-F[1],H=1/Math.sin(Math.acos((I*B+Y*j)/(Math.sqrt(I*I+Y*Y)*Math.sqrt(B*B+j*j)))/2),X=Math.sqrt(F[0]*F[0]+F[1]*F[1]);N=Math.min(S,(f-X)/(H-1)),A=Math.min(S,(l-X)/(H+1))}}M>mv?A>mv?(_=cn(D,O,z,P,l,A,v),y=cn(R,U,q,L,l,A,v),c.moveTo(_.cx+_.x01,_.cy+_.y01),A<S?c.arc(_.cx,_.cy,A,Math.atan2(_.y01,_.x01),Math.atan2(y.y01,y.x01),!v):(c.arc(_.cx,_.cy,A,Math.atan2(_.y01,_.x01),Math.atan2(_.y11,_.x11),!v),c.arc(0,0,l,Math.atan2(_.cy+_.y11,_.cx+_.x11),Math.atan2(y.cy+y.y11,y.cx+y.x11),!v),c.arc(y.cx,y.cy,A,Math.atan2(y.y11,y.x11),Math.atan2(y.y01,y.x01),!v))):(c.moveTo(z,P),c.arc(0,0,l,g,m,!v)):c.moveTo(z,P),f>mv&&w>mv?N>mv?(_=cn(q,L,R,U,f,-N,v),y=cn(z,P,D,O,f,-N,v),c.lineTo(_.cx+_.x01,_.cy+_.y01),N<S?c.arc(_.cx,_.cy,N,Math.atan2(_.y01,_.x01),Math.atan2(y.y01,y.x01),!v):(c.arc(_.cx,_.cy,N,Math.atan2(_.y01,_.x01),Math.atan2(_.y11,_.x11),!v),c.arc(0,0,f,Math.atan2(_.cy+_.y11,_.cx+_.x11),Math.atan2(y.cy+y.y11,y.cx+y.x11),v),c.arc(y.cx,y.cy,N,Math.atan2(y.y11,y.x11),Math.atan2(y.y01,y.x01),!v))):c.arc(0,0,f,b,x,v):c.lineTo(q,L)}else c.moveTo(0,0);if(c.closePath(),t)return c=null,t+""||null}var n=tn,e=nn,r=Kt(0),i=null,o=en,u=rn,a=on,c=null;return t.centroid=function(){var t=(+n.apply(this,arguments)+ +e.apply(this,arguments))/2,r=(+o.apply(this,arguments)+ +u.apply(this,arguments))/2-xv/2;return[Math.cos(r)*t,Math.sin(r)*t]},t.innerRadius=function(e){return arguments.length?(n="function"==typeof e?e:Kt(+e),t):n},t.outerRadius=function(n){return arguments.length?(e="function"==typeof n?n:Kt(+n),t):e},t.cornerRadius=function(n){return arguments.length?(r="function"==typeof n?n:Kt(+n),t):r},t.padRadius=function(n){return arguments.length?(i=null==n?null:"function"==typeof n?n:Kt(+n),t):i},t.startAngle=function(n){return arguments.length?(o="function"==typeof n?n:Kt(+n),t):o},t.endAngle=function(n){return arguments.length?(u="function"==typeof n?n:Kt(+n),t):u},t.padAngle=function(n){return arguments.length?(a="function"==typeof n?n:Kt(+n),t):a},t.context=function(n){return arguments.length?(c=null==n?null:n,t):c},t}function fn(t){this._context=t}function ln(t){return new fn(t)}function hn(t){return t[0]}function pn(t){return t[1]}function dn(){function t(t){var a,c,s,f=t.length,l=!1;for(null==i&&(u=o(s=Tt())),a=0;a<=f;++a)!(a<f&&r(c=t[a],a,t))===l&&((l=!l)?u.lineStart():u.lineEnd()),l&&u.point(+n(c,a,t),+e(c,a,t));if(s)return u=null,s+""||null}var n=hn,e=pn,r=Kt(!0),i=null,o=ln,u=null;return t.x=function(e){return arguments.length?(n="function"==typeof e?e:Kt(+e),t):n},t.y=function(n){return arguments.length?(e="function"==typeof n?n:Kt(+n),t):e},t.defined=function(n){return arguments.length?(r="function"==typeof n?n:Kt(!!n),t):r},t.curve=function(n){return arguments.length?(o=n,null!=i&&(u=o(i)),t):o},t.context=function(n){return arguments.length?(null==n?i=u=null:u=o(i=n),t):i},t}function vn(){function t(t){var n,f,l,h,p,d=t.length,v=!1,_=new Array(d),y=new Array(d);for(null==a&&(s=c(p=Tt())),n=0;n<=d;++n){if(!(n<d&&u(h=t[n],n,t))===v)if(v=!v)f=n,s.areaStart(),s.lineStart();else{for(s.lineEnd(),s.lineStart(),l=n-1;l>=f;--l)s.point(_[l],y[l]);s.lineEnd(),s.areaEnd()}v&&(_[n]=+e(h,n,t),y[n]=+i(h,n,t),s.point(r?+r(h,n,t):_[n],o?+o(h,n,t):y[n]))}if(p)return s=null,p+""||null}function n(){return dn().defined(u).curve(c).context(a)}var e=hn,r=null,i=Kt(0),o=pn,u=Kt(!0),a=null,c=ln,s=null;return t.x=function(n){return arguments.length?(e="function"==typeof n?n:Kt(+n),r=null,t):e},t.x0=function(n){return arguments.length?(e="function"==typeof n?n:Kt(+n),t):e},t.x1=function(n){return arguments.length?(r=null==n?null:"function"==typeof n?n:Kt(+n),t):r},t.y=function(n){return arguments.length?(i="function"==typeof n?n:Kt(+n),o=null,t):i},t.y0=function(n){return arguments.length?(i="function"==typeof n?n:Kt(+n),t):i},t.y1=function(n){return arguments.length?(o=null==n?null:"function"==typeof n?n:Kt(+n),t):o},t.lineX0=t.lineY0=function(){return n().x(e).y(i)},t.lineY1=function(){return n().x(e).y(o)},t.lineX1=function(){return n().x(r).y(i)},t.defined=function(n){return arguments.length?(u="function"==typeof n?n:Kt(!!n),t):u},t.curve=function(n){return arguments.length?(c=n,null!=a&&(s=c(a)),t):c},t.context=function(n){return arguments.length?(null==n?a=s=null:s=c(a=n),t):a},t}function _n(t,n){return n<t?-1:n>t?1:n>=t?0:NaN}function yn(t){return t}function gn(){function t(t){var a,c,s,f,l,h=t.length,p=0,d=new Array(h),v=new Array(h),_=+i.apply(this,arguments),y=Math.min(wv,Math.max(-wv,o.apply(this,arguments)-_)),g=Math.min(Math.abs(y)/h,u.apply(this,arguments)),m=g*(y<0?-1:1);for(a=0;a<h;++a)(l=v[d[a]=a]=+n(t[a],a,t))>0&&(p+=l);for(null!=e?d.sort(function(t,n){return e(v[t],v[n])}):null!=r&&d.sort(function(n,e){return r(t[n],t[e])}),a=0,s=p?(y-h*m)/p:0;a<h;++a,_=f)c=d[a],l=v[c],f=_+(l>0?l*s:0)+m,v[c]={data:t[c],index:a,value:l,startAngle:_,endAngle:f,padAngle:g};return v}var n=yn,e=_n,r=null,i=Kt(0),o=Kt(wv),u=Kt(0);return t.value=function(e){return arguments.length?(n="function"==typeof e?e:Kt(+e),t):n},t.sortValues=function(n){return arguments.length?(e=n,r=null,t):e},t.sort=function(n){return arguments.length?(r=n,e=null,t):r},t.startAngle=function(n){return arguments.length?(i="function"==typeof n?n:Kt(+n),t):i},t.endAngle=function(n){return arguments.length?(o="function"==typeof n?n:Kt(+n),t):o},t.padAngle=function(n){return arguments.length?(u="function"==typeof n?n:Kt(+n),t):u},t}function mn(t){this._curve=t}function xn(t){function n(n){return new mn(t(n))}return n._curve=t,n}function bn(t){var n=t.curve;return t.angle=t.x,delete t.x,t.radius=t.y,delete t.y,t.curve=function(t){return arguments.length?n(xn(t)):n()._curve},t}function wn(){return bn(dn().curve(Mv))}function Mn(){var t=vn().curve(Mv),n=t.curve,e=t.lineX0,r=t.lineX1,i=t.lineY0,o=t.lineY1;return t.angle=t.x,delete t.x,t.startAngle=t.x0,delete t.x0,t.endAngle=t.x1,delete t.x1,t.radius=t.y,delete t.y,t.innerRadius=t.y0,delete t.y0,t.outerRadius=t.y1,delete t.y1,t.lineStartAngle=function(){return bn(e())},delete t.lineX0,t.lineEndAngle=function(){return bn(r())},delete t.lineX1,t.lineInnerRadius=function(){return bn(i())},delete t.lineY0,t.lineOuterRadius=function(){return bn(o())},delete t.lineY1,t.curve=function(t){return arguments.length?n(xn(t)):n()._curve},t}function Tn(){function t(){var t;if(r||(r=t=Tt()),n.apply(this,arguments).draw(r,+e.apply(this,arguments)),t)return r=null,t+""||null}var n=Kt(Tv),e=Kt(64),r=null;return t.type=function(e){return arguments.length?(n="function"==typeof e?e:Kt(e),t):n},t.size=function(n){return arguments.length?(e="function"==typeof n?n:Kt(+n),t):e},t.context=function(n){return arguments.length?(r=null==n?null:n,t):r},t}function kn(){}function Sn(t,n,e){t._context.bezierCurveTo((2*t._x0+t._x1)/3,(2*t._y0+t._y1)/3,(t._x0+2*t._x1)/3,(t._y0+2*t._y1)/3,(t._x0+4*t._x1+n)/6,(t._y0+4*t._y1+e)/6)}function Nn(t){this._context=t}function An(t){return new Nn(t)}function En(t){this._context=t}function Cn(t){return new En(t)}function zn(t){this._context=t}function Pn(t){return new zn(t)}function qn(t,n){this._basis=new Nn(t),this._beta=n}function Ln(t,n,e){t._context.bezierCurveTo(t._x1+t._k*(t._x2-t._x0),t._y1+t._k*(t._y2-t._y0),t._x2+t._k*(t._x1-n),t._y2+t._k*(t._y1-e),t._x2,t._y2)}function Rn(t,n){this._context=t,this._k=(1-n)/6}function Un(t,n){this._context=t,this._k=(1-n)/6}function Dn(t,n){this._context=t,this._k=(1-n)/6}function On(t,n,e){var r=t._x1,i=t._y1,o=t._x2,u=t._y2;if(t._l01_a>mv){var a=2*t._l01_2a+3*t._l01_a*t._l12_a+t._l12_2a,c=3*t._l01_a*(t._l01_a+t._l12_a);r=(r*a-t._x0*t._l12_2a+t._x2*t._l01_2a)/c,i=(i*a-t._y0*t._l12_2a+t._y2*t._l01_2a)/c}if(t._l23_a>mv){var s=2*t._l23_2a+3*t._l23_a*t._l12_a+t._l12_2a,f=3*t._l23_a*(t._l23_a+t._l12_a);o=(o*s+t._x1*t._l23_2a-n*t._l12_2a)/f,u=(u*s+t._y1*t._l23_2a-e*t._l12_2a)/f}t._context.bezierCurveTo(r,i,o,u,t._x2,t._y2)}function Fn(t,n){this._context=t,this._alpha=n}function In(t,n){this._context=t,this._alpha=n}function Yn(t,n){this._context=t,this._alpha=n}function Bn(t){this._context=t}function jn(t){return new Bn(t)}function Hn(t){return t<0?-1:1}function Xn(t,n,e){var r=t._x1-t._x0,i=n-t._x1,o=(t._y1-t._y0)/(r||i<0&&-0),u=(e-t._y1)/(i||r<0&&-0),a=(o*i+u*r)/(r+i);return(Hn(o)+Hn(u))*Math.min(Math.abs(o),Math.abs(u),.5*Math.abs(a))||0}function Vn(t,n){var e=t._x1-t._x0;return e?(3*(t._y1-t._y0)/e-n)/2:n}function Wn(t,n,e){var r=t._x0,i=t._y0,o=t._x1,u=t._y1,a=(o-r)/3;t._context.bezierCurveTo(r+a,i+a*n,o-a,u-a*e,o,u)}function $n(t){this._context=t}function Zn(t){this._context=new Gn(t)}function Gn(t){this._context=t}function Jn(t){return new $n(t)}function Qn(t){return new Zn(t)}function Kn(t){this._context=t}function te(t){var n,e,r=t.length-1,i=new Array(r),o=new Array(r),u=new Array(r);for(i[0]=0,o[0]=2,u[0]=t[0]+2*t[1],n=1;n<r-1;++n)i[n]=1,o[n]=4,u[n]=4*t[n]+2*t[n+1];for(i[r-1]=2,o[r-1]=7,u[r-1]=8*t[r-1]+t[r],n=1;n<r;++n)e=i[n]/o[n-1],o[n]-=e,u[n]-=e*u[n-1];for(i[r-1]=u[r-1]/o[r-1],n=r-2;n>=0;--n)i[n]=(u[n]-i[n+1])/o[n];for(o[r-1]=(t[r]+i[r-1])/2,n=0;n<r-1;++n)o[n]=2*t[n+1]-i[n+1];return[i,o]}function ne(t){return new Kn(t)}function ee(t,n){this._context=t,this._t=n}function re(t){return new ee(t,.5)}function ie(t){return new ee(t,0)}function oe(t){return new ee(t,1)}function ue(t,n){if((r=t.length)>1)for(var e,r,i=1,o=t[n[0]],u=o.length;i<r;++i){e=o,o=t[n[i]];for(var a=0;a<u;++a)o[a][1]+=o[a][0]=isNaN(e[a][1])?e[a][0]:e[a][1]}}function ae(t){for(var n=t.length,e=new Array(n);--n>=0;)e[n]=n;return e}function ce(t,n){return t[n]}function se(){function t(t){var o,u,a=n.apply(this,arguments),c=t.length,s=a.length,f=new Array(s);for(o=0;o<s;++o){for(var l,h=a[o],p=f[o]=new Array(c),d=0;d<c;++d)p[d]=l=[0,+i(t[d],h,d,t)],l.data=t[d];p.key=h}for(o=0,u=e(f);o<s;++o)f[u[o]].index=o;return r(f,u),f}var n=Kt([]),e=ae,r=ue,i=ce;return t.keys=function(e){return arguments.length?(n="function"==typeof e?e:Kt(Gv.call(e)),t):n},t.value=function(n){return arguments.length?(i="function"==typeof n?n:Kt(+n),t):i},t.order=function(n){return arguments.length?(e=null==n?ae:"function"==typeof n?n:Kt(Gv.call(n)),t):e},t.offset=function(n){return arguments.length?(r=null==n?ue:n,t):r},t}function fe(t,n){if((r=t.length)>0){for(var e,r,i,o=0,u=t[0].length;o<u;++o){for(i=e=0;e<r;++e)i+=t[e][o][1]||0;if(i)for(e=0;e<r;++e)t[e][o][1]/=i}ue(t,n)}}function le(t,n){if((e=t.length)>0){for(var e,r=0,i=t[n[0]],o=i.length;r<o;++r){for(var u=0,a=0;u<e;++u)a+=t[u][r][1]||0;i[r][1]+=i[r][0]=-a/2}ue(t,n)}}function he(t,n){if((i=t.length)>0&&(r=(e=t[n[0]]).length)>0){for(var e,r,i,o=0,u=1;u<r;++u){for(var a=0,c=0,s=0;a<i;++a){for(var f=t[n[a]],l=f[u][1]||0,h=f[u-1][1]||0,p=(l-h)/2,d=0;d<a;++d){var v=t[n[d]],_=v[u][1]||0,y=v[u-1][1]||0;p+=_-y}c+=l,s+=p*l}e[u-1][1]+=e[u-1][0]=o,c&&(o-=s/c)}e[u-1][1]+=e[u-1][0]=o,ue(t,n)}}function pe(t){var n=t.map(de);return ae(t).sort(function(t,e){return n[t]-n[e]})}function de(t){for(var n,e=0,r=-1,i=t.length;++r<i;)(n=+t[r][1])&&(e+=n);return e}function ve(t){return pe(t).reverse()}function _e(t){var n,e,r=t.length,i=t.map(de),o=ae(t).sort(function(t,n){return i[n]-i[t]}),u=0,a=0,c=[],s=[];for(n=0;n<r;++n)e=o[n],u<a?(u+=i[e],c.push(e)):(a+=i[e],s.push(e));return s.reverse().concat(c)}function ye(t){return ae(t).reverse()}function ge(t,n,e){t.prototype=n.prototype=e,e.constructor=t}function me(t,n){var e=Object.create(t.prototype);for(var r in n)e[r]=n[r];return e}function xe(){}function be(t){var n;return t=(t+"").trim().toLowerCase(),(n=Kv.exec(t))?(n=parseInt(n[1],16),new Se(n>>8&15|n>>4&240,n>>4&15|240&n,(15&n)<<4|15&n,1)):(n=t_.exec(t))?we(parseInt(n[1],16)):(n=n_.exec(t))?new Se(n[1],n[2],n[3],1):(n=e_.exec(t))?new Se(255*n[1]/100,255*n[2]/100,255*n[3]/100,1):(n=r_.exec(t))?Me(n[1],n[2],n[3],n[4]):(n=i_.exec(t))?Me(255*n[1]/100,255*n[2]/100,255*n[3]/100,n[4]):(n=o_.exec(t))?Ne(n[1],n[2]/100,n[3]/100,1):(n=u_.exec(t))?Ne(n[1],n[2]/100,n[3]/100,n[4]):a_.hasOwnProperty(t)?we(a_[t]):"transparent"===t?new Se(NaN,NaN,NaN,0):null}function we(t){return new Se(t>>16&255,t>>8&255,255&t,1)}function Me(t,n,e,r){return r<=0&&(t=n=e=NaN),new Se(t,n,e,r)}function Te(t){return t instanceof xe||(t=be(t)),t?(t=t.rgb(),new Se(t.r,t.g,t.b,t.opacity)):new Se}function ke(t,n,e,r){return 1===arguments.length?Te(t):new Se(t,n,e,null==r?1:r)}function Se(t,n,e,r){this.r=+t,this.g=+n,this.b=+e,this.opacity=+r}function Ne(t,n,e,r){return r<=0?t=n=e=NaN:e<=0||e>=1?t=n=NaN:n<=0&&(t=NaN),new Ce(t,n,e,r)}function Ae(t){if(t instanceof Ce)return new Ce(t.h,t.s,t.l,t.opacity);if(t instanceof xe||(t=be(t)),!t)return new Ce;if(t instanceof Ce)return t;t=t.rgb();var n=t.r/255,e=t.g/255,r=t.b/255,i=Math.min(n,e,r),o=Math.max(n,e,r),u=NaN,a=o-i,c=(o+i)/2;return a?(u=n===o?(e-r)/a+6*(e<r):e===o?(r-n)/a+2:(n-e)/a+4,a/=c<.5?o+i:2-o-i,u*=60):a=c>0&&c<1?0:u,new Ce(u,a,c,t.opacity)}function Ee(t,n,e,r){return 1===arguments.length?Ae(t):new Ce(t,n,e,null==r?1:r)}function Ce(t,n,e,r){this.h=+t,this.s=+n,this.l=+e,this.opacity=+r}function ze(t,n,e){return 255*(t<60?n+(e-n)*t/60:t<180?e:t<240?n+(e-n)*(240-t)/60:n)}function Pe(t){if(t instanceof Le)return new Le(t.l,t.a,t.b,t.opacity);if(t instanceof Ye){var n=t.h*c_;return new Le(t.l,Math.cos(n)*t.c,Math.sin(n)*t.c,t.opacity)}t instanceof Se||(t=Te(t));var e=Oe(t.r),r=Oe(t.g),i=Oe(t.b),o=Re((.4124564*e+.3575761*r+.1804375*i)/l_),u=Re((.2126729*e+.7151522*r+.072175*i)/h_),a=Re((.0193339*e+.119192*r+.9503041*i)/p_);return new Le(116*u-16,500*(o-u),200*(u-a),t.opacity)}function qe(t,n,e,r){return 1===arguments.length?Pe(t):new Le(t,n,e,null==r?1:r)}function Le(t,n,e,r){this.l=+t,this.a=+n,this.b=+e,this.opacity=+r}function Re(t){return t>y_?Math.pow(t,1/3):t/__+d_}function Ue(t){return t>v_?t*t*t:__*(t-d_)}function De(t){return 255*(t<=.0031308?12.92*t:1.055*Math.pow(t,1/2.4)-.055)}function Oe(t){return(t/=255)<=.04045?t/12.92:Math.pow((t+.055)/1.055,2.4)}function Fe(t){if(t instanceof Ye)return new Ye(t.h,t.c,t.l,t.opacity);t instanceof Le||(t=Pe(t));var n=Math.atan2(t.b,t.a)*s_;return new Ye(n<0?n+360:n,Math.sqrt(t.a*t.a+t.b*t.b),t.l,t.opacity)}function Ie(t,n,e,r){return 1===arguments.length?Fe(t):new Ye(t,n,e,null==r?1:r)}function Ye(t,n,e,r){this.h=+t,this.c=+n,this.l=+e,this.opacity=+r}function Be(t){if(t instanceof He)return new He(t.h,t.s,t.l,t.opacity);t instanceof Se||(t=Te(t));var n=t.r/255,e=t.g/255,r=t.b/255,i=(k_*r+M_*n-T_*e)/(k_+M_-T_),o=r-i,u=(w_*(e-i)-x_*o)/b_,a=Math.sqrt(u*u+o*o)/(w_*i*(1-i)),c=a?Math.atan2(u,o)*s_-120:NaN;return new He(c<0?c+360:c,a,i,t.opacity)}function je(t,n,e,r){return 1===arguments.length?Be(t):new He(t,n,e,null==r?1:r)}function He(t,n,e,r){this.h=+t,this.s=+n,this.l=+e,this.opacity=+r}function Xe(t,n,e,r,i){var o=t*t,u=o*t;return((1-3*t+3*o-u)*n+(4-6*o+3*u)*e+(1+3*t+3*o-3*u)*r+u*i)/6}function Ve(t){var n=t.length-1;return function(e){var r=e<=0?e=0:e>=1?(e=1,n-1):Math.floor(e*n),i=t[r],o=t[r+1],u=r>0?t[r-1]:2*i-o,a=r<n-1?t[r+2]:2*o-i;return Xe((e-r/n)*n,u,i,o,a)}}function We(t){var n=t.length;return function(e){var r=Math.floor(((e%=1)<0?++e:e)*n),i=t[(r+n-1)%n],o=t[r%n],u=t[(r+1)%n],a=t[(r+2)%n];return Xe((e-r/n)*n,i,o,u,a)}}function $e(t){return function(){return t}}function Ze(t,n){return function(e){return t+e*n}}function Ge(t,n,e){return t=Math.pow(t,e),n=Math.pow(n,e)-t,e=1/e,function(r){return Math.pow(t+r*n,e)}}function Je(t,n){var e=n-t;return e?Ze(t,e>180||e<-180?e-360*Math.round(e/360):e):$e(isNaN(t)?n:t)}function Qe(t){return 1===(t=+t)?Ke:function(n,e){return e-n?Ge(n,e,t):$e(isNaN(n)?e:n)}}function Ke(t,n){var e=n-t;return e?Ze(t,e):$e(isNaN(t)?n:t)}function tr(t){return function(n){var e,r,i=n.length,o=new Array(i),u=new Array(i),a=new Array(i);for(e=0;e<i;++e)r=ke(n[e]),o[e]=r.r||0,u[e]=r.g||0,a[e]=r.b||0;return o=t(o),u=t(u),a=t(a),r.opacity=1,function(t){return r.r=o(t),r.g=u(t),r.b=a(t),r+""}}}function nr(t,n){var e,r=n?n.length:0,i=t?Math.min(r,t.length):0,o=new Array(r),u=new Array(r);for(e=0;e<i;++e)o[e]=cr(t[e],n[e]);
for(;e<r;++e)u[e]=n[e];return function(t){for(e=0;e<i;++e)u[e]=o[e](t);return u}}function er(t,n){var e=new Date;return t=+t,n-=t,function(r){return e.setTime(t+n*r),e}}function rr(t,n){return t=+t,n-=t,function(e){return t+n*e}}function ir(t,n){var e,r={},i={};null!==t&&"object"==typeof t||(t={}),null!==n&&"object"==typeof n||(n={});for(e in n)e in t?r[e]=cr(t[e],n[e]):i[e]=n[e];return function(t){for(e in r)i[e]=r[e](t);return i}}function or(t){return function(){return t}}function ur(t){return function(n){return t(n)+""}}function ar(t,n){var e,r,i,o=q_.lastIndex=L_.lastIndex=0,u=-1,a=[],c=[];for(t+="",n+="";(e=q_.exec(t))&&(r=L_.exec(n));)(i=r.index)>o&&(i=n.slice(o,i),a[u]?a[u]+=i:a[++u]=i),(e=e[0])===(r=r[0])?a[u]?a[u]+=r:a[++u]=r:(a[++u]=null,c.push({i:u,x:rr(e,r)})),o=L_.lastIndex;return o<n.length&&(i=n.slice(o),a[u]?a[u]+=i:a[++u]=i),a.length<2?c[0]?ur(c[0].x):or(n):(n=c.length,function(t){for(var e,r=0;r<n;++r)a[(e=c[r]).i]=e.x(t);return a.join("")})}function cr(t,n){var e,r=typeof n;return null==n||"boolean"===r?$e(n):("number"===r?rr:"string"===r?(e=be(n))?(n=e,C_):ar:n instanceof be?C_:n instanceof Date?er:Array.isArray(n)?nr:isNaN(n)?ir:rr)(t,n)}function sr(t,n){return t=+t,n-=t,function(e){return Math.round(t+n*e)}}function fr(t,n,e,r,i,o){var u,a,c;return(u=Math.sqrt(t*t+n*n))&&(t/=u,n/=u),(c=t*e+n*r)&&(e-=t*c,r-=n*c),(a=Math.sqrt(e*e+r*r))&&(e/=a,r/=a,c/=a),t*r<n*e&&(t=-t,n=-n,c=-c,u=-u),{translateX:i,translateY:o,rotate:Math.atan2(n,t)*R_,skewX:Math.atan(c)*R_,scaleX:u,scaleY:a}}function lr(t){return"none"===t?U_:(S_||(S_=document.createElement("DIV"),N_=document.documentElement,A_=document.defaultView),S_.style.transform=t,t=A_.getComputedStyle(N_.appendChild(S_),null).getPropertyValue("transform"),N_.removeChild(S_),t=t.slice(7,-1).split(","),fr(+t[0],+t[1],+t[2],+t[3],+t[4],+t[5]))}function hr(t){return null==t?U_:(E_||(E_=document.createElementNS("http://www.w3.org/2000/svg","g")),E_.setAttribute("transform",t),(t=E_.transform.baseVal.consolidate())?(t=t.matrix,fr(t.a,t.b,t.c,t.d,t.e,t.f)):U_)}function pr(t,n,e,r){function i(t){return t.length?t.pop()+" ":""}function o(t,r,i,o,u,a){if(t!==i||r!==o){var c=u.push("translate(",null,n,null,e);a.push({i:c-4,x:rr(t,i)},{i:c-2,x:rr(r,o)})}else(i||o)&&u.push("translate("+i+n+o+e)}function u(t,n,e,o){t!==n?(t-n>180?n+=360:n-t>180&&(t+=360),o.push({i:e.push(i(e)+"rotate(",null,r)-2,x:rr(t,n)})):n&&e.push(i(e)+"rotate("+n+r)}function a(t,n,e,o){t!==n?o.push({i:e.push(i(e)+"skewX(",null,r)-2,x:rr(t,n)}):n&&e.push(i(e)+"skewX("+n+r)}function c(t,n,e,r,o,u){if(t!==e||n!==r){var a=o.push(i(o)+"scale(",null,",",null,")");u.push({i:a-4,x:rr(t,e)},{i:a-2,x:rr(n,r)})}else 1===e&&1===r||o.push(i(o)+"scale("+e+","+r+")")}return function(n,e){var r=[],i=[];return n=t(n),e=t(e),o(n.translateX,n.translateY,e.translateX,e.translateY,r,i),u(n.rotate,e.rotate,r,i),a(n.skewX,e.skewX,r,i),c(n.scaleX,n.scaleY,e.scaleX,e.scaleY,r,i),n=e=null,function(t){for(var n,e=-1,o=i.length;++e<o;)r[(n=i[e]).i]=n.x(t);return r.join("")}}}function dr(t){return((t=Math.exp(t))+1/t)/2}function vr(t){return((t=Math.exp(t))-1/t)/2}function _r(t){return((t=Math.exp(2*t))-1)/(t+1)}function yr(t,n){var e,r,i=t[0],o=t[1],u=t[2],a=n[0],c=n[1],s=n[2],f=a-i,l=c-o,h=f*f+l*l;if(h<B_)r=Math.log(s/u)/F_,e=function(t){return[i+t*f,o+t*l,u*Math.exp(F_*t*r)]};else{var p=Math.sqrt(h),d=(s*s-u*u+Y_*h)/(2*u*I_*p),v=(s*s-u*u-Y_*h)/(2*s*I_*p),_=Math.log(Math.sqrt(d*d+1)-d),y=Math.log(Math.sqrt(v*v+1)-v);r=(y-_)/F_,e=function(t){var n=t*r,e=dr(_),a=u/(I_*p)*(e*_r(F_*n+_)-vr(_));return[i+a*f,o+a*l,u*e/dr(F_*n+_)]}}return e.duration=1e3*r,e}function gr(t){return function(n,e){var r=t((n=Ee(n)).h,(e=Ee(e)).h),i=Ke(n.s,e.s),o=Ke(n.l,e.l),u=Ke(n.opacity,e.opacity);return function(t){return n.h=r(t),n.s=i(t),n.l=o(t),n.opacity=u(t),n+""}}}function mr(t,n){var e=Ke((t=qe(t)).l,(n=qe(n)).l),r=Ke(t.a,n.a),i=Ke(t.b,n.b),o=Ke(t.opacity,n.opacity);return function(n){return t.l=e(n),t.a=r(n),t.b=i(n),t.opacity=o(n),t+""}}function xr(t){return function(n,e){var r=t((n=Ie(n)).h,(e=Ie(e)).h),i=Ke(n.c,e.c),o=Ke(n.l,e.l),u=Ke(n.opacity,e.opacity);return function(t){return n.h=r(t),n.c=i(t),n.l=o(t),n.opacity=u(t),n+""}}}function br(t){return function n(e){function r(n,r){var i=t((n=je(n)).h,(r=je(r)).h),o=Ke(n.s,r.s),u=Ke(n.l,r.l),a=Ke(n.opacity,r.opacity);return function(t){return n.h=i(t),n.s=o(t),n.l=u(Math.pow(t,e)),n.opacity=a(t),n+""}}return e=+e,r.gamma=n,r}(1)}function wr(t,n){for(var e=new Array(n),r=0;r<n;++r)e[r]=t(r/(n-1));return e}function Mr(){for(var t,n=0,e=arguments.length,r={};n<e;++n){if(!(t=arguments[n]+"")||t in r)throw new Error("illegal type: "+t);r[t]=[]}return new Tr(r)}function Tr(t){this._=t}function kr(t,n){return t.trim().split(/^|\s+/).map(function(t){var e="",r=t.indexOf(".");if(r>=0&&(e=t.slice(r+1),t=t.slice(0,r)),t&&!n.hasOwnProperty(t))throw new Error("unknown type: "+t);return{type:t,name:e}})}function Sr(t,n){for(var e,r=0,i=t.length;r<i;++r)if((e=t[r]).name===n)return e.value}function Nr(t,n,e){for(var r=0,i=t.length;r<i;++r)if(t[r].name===n){t[r]=Z_,t=t.slice(0,r).concat(t.slice(r+1));break}return null!=e&&t.push({name:n,value:e}),t}function Ar(t){return new Function("d","return {"+t.map(function(t,n){return JSON.stringify(t)+": d["+n+"]"}).join(",")+"}")}function Er(t,n){var e=Ar(t);return function(r,i){return n(e(r),i,t)}}function Cr(t){var n=Object.create(null),e=[];return t.forEach(function(t){for(var r in t)r in n||e.push(n[r]=r)}),e}function zr(t){function n(t,n){var r,i,o=e(t,function(t,e){return r?r(t,e-1):(i=t,void(r=n?Er(t,n):Ar(t)))});return o.columns=i,o}function e(t,n){function e(){if(f>=s)return u;if(i)return i=!1,o;var n,e=f;if(34===t.charCodeAt(e)){for(var r=e;r++<s;)if(34===t.charCodeAt(r)){if(34!==t.charCodeAt(r+1))break;++r}return f=r+2,n=t.charCodeAt(r+1),13===n?(i=!0,10===t.charCodeAt(r+2)&&++f):10===n&&(i=!0),t.slice(e+1,r).replace(/""/g,'"')}for(;f<s;){var a=1;if(n=t.charCodeAt(f++),10===n)i=!0;else if(13===n)i=!0,10===t.charCodeAt(f)&&(++f,++a);else if(n!==c)continue;return t.slice(e,f-a)}return t.slice(e)}for(var r,i,o={},u={},a=[],s=t.length,f=0,l=0;(r=e())!==u;){for(var h=[];r!==o&&r!==u;)h.push(r),r=e();n&&null==(h=n(h,l++))||a.push(h)}return a}function r(n,e){return null==e&&(e=Cr(n)),[e.map(u).join(t)].concat(n.map(function(n){return e.map(function(t){return u(n[t])}).join(t)})).join("\n")}function i(t){return t.map(o).join("\n")}function o(n){return n.map(u).join(t)}function u(t){return null==t?"":a.test(t+="")?'"'+t.replace(/\"/g,'""')+'"':t}var a=new RegExp('["'+t+"\n]"),c=t.charCodeAt(0);return{parse:n,parseRows:e,format:r,formatRows:i}}function Pr(t,n){function e(t){var n,e=s.status;if(!e&&Lr(s)||e>=200&&e<300||304===e){if(o)try{n=o.call(r,s)}catch(t){return void a.call("error",r,t)}else n=s;a.call("load",r,n)}else a.call("error",r,t)}var r,i,o,u,a=Mr("beforesend","progress","load","error"),c=q(),s=new XMLHttpRequest,f=null,l=null,h=0;if("undefined"==typeof XDomainRequest||"withCredentials"in s||!/^(http(s)?:)?\/\//.test(t)||(s=new XDomainRequest),"onload"in s?s.onload=s.onerror=s.ontimeout=e:s.onreadystatechange=function(t){s.readyState>3&&e(t)},s.onprogress=function(t){a.call("progress",r,t)},r={header:function(t,n){return t=(t+"").toLowerCase(),arguments.length<2?c.get(t):(null==n?c.remove(t):c.set(t,n+""),r)},mimeType:function(t){return arguments.length?(i=null==t?null:t+"",r):i},responseType:function(t){return arguments.length?(u=t,r):u},timeout:function(t){return arguments.length?(h=+t,r):h},user:function(t){return arguments.length<1?f:(f=null==t?null:t+"",r)},password:function(t){return arguments.length<1?l:(l=null==t?null:t+"",r)},response:function(t){return o=t,r},get:function(t,n){return r.send("GET",t,n)},post:function(t,n){return r.send("POST",t,n)},send:function(n,e,o){return s.open(n,t,!0,f,l),null==i||c.has("accept")||c.set("accept",i+",*/*"),s.setRequestHeader&&c.each(function(t,n){s.setRequestHeader(n,t)}),null!=i&&s.overrideMimeType&&s.overrideMimeType(i),null!=u&&(s.responseType=u),h>0&&(s.timeout=h),null==o&&"function"==typeof e&&(o=e,e=null),null!=o&&1===o.length&&(o=qr(o)),null!=o&&r.on("error",o).on("load",function(t){o(null,t)}),a.call("beforesend",r,s),s.send(null==e?null:e),r},abort:function(){return s.abort(),r},on:function(){var t=a.on.apply(a,arguments);return t===a?r:t}},null!=n){if("function"!=typeof n)throw new Error("invalid callback: "+n);return r.get(n)}return r}function qr(t){return function(n,e){t(null==n?e:null)}}function Lr(t){var n=t.responseType;return n&&"text"!==n?t.response:t.responseText}function Rr(t,n){return function(e,r){var i=Pr(e).mimeType(t).response(n);if(null!=r){if("function"!=typeof r)throw new Error("invalid callback: "+r);return i.get(r)}return i}}function Ur(t,n){return function(e,r,i){arguments.length<3&&(i=r,r=null);var o=Pr(e).mimeType(t);return o.row=function(t){return arguments.length?o.response(Dr(n,r=t)):r},o.row(r),i?o.get(i):o}}function Dr(t,n){return function(e){return t(e.responseText,n)}}function Or(){return my||(wy(Fr),my=by.now()+xy)}function Fr(){my=0}function Ir(){this._call=this._time=this._next=null}function Yr(t,n,e){var r=new Ir;return r.restart(t,n,e),r}function Br(){Or(),++dy;for(var t,n=G_;n;)(t=my-n._time)>=0&&n._call.call(null,t),n=n._next;--dy}function jr(t){my=(gy=t||by.now())+xy,dy=vy=0;try{Br()}finally{dy=0,Xr(),my=0}}function Hr(){var t=by.now(),n=t-gy;n>yy&&(xy-=n,gy=t)}function Xr(){for(var t,n,e=G_,r=1/0;e;)e._call?(r>e._time&&(r=e._time),t=e,e=e._next):(n=e._next,e._next=null,e=t?t._next=n:G_=n);J_=t,Vr(r)}function Vr(t){if(!dy){vy&&(vy=clearTimeout(vy));var n=t-my;n>24?(t<1/0&&(vy=setTimeout(jr,n)),_y&&(_y=clearInterval(_y))):(_y||(_y=setInterval(Hr,yy)),dy=1,wy(jr))}}function Wr(t,n,e){var r=new Ir;return n=null==n?0:+n,r.restart(function(e){r.stop(),t(e+n)},n,e),r}function $r(t,n,e){var r=new Ir,i=n;return null==n?(r.restart(t,n,e),r):(n=+n,e=null==e?Or():+e,r.restart(function o(u){u+=i,r.restart(o,i+=n,e),t(u)},n,e),r)}function Zr(t,n,e,r){function i(n){return t(n=new Date(+n)),n}return i.floor=i,i.ceil=function(e){return t(e=new Date(e-1)),n(e,1),t(e),e},i.round=function(t){var n=i(t),e=i.ceil(t);return t-n<e-t?n:e},i.offset=function(t,e){return n(t=new Date(+t),null==e?1:Math.floor(e)),t},i.range=function(e,r,o){var u=[];if(e=i.ceil(e),o=null==o?1:Math.floor(o),!(e<r&&o>0))return u;do u.push(new Date(+e));while(n(e,o),t(e),e<r);return u},i.filter=function(e){return Zr(function(n){for(;t(n),!e(n);)n.setTime(n-1)},function(t,r){for(;--r>=0;)for(;n(t,1),!e(t););})},e&&(i.count=function(n,r){return My.setTime(+n),Ty.setTime(+r),t(My),t(Ty),Math.floor(e(My,Ty))},i.every=function(t){return t=Math.floor(t),isFinite(t)&&t>0?t>1?i.filter(r?function(n){return r(n)%t===0}:function(n){return i.count(0,n)%t===0}):i:null}),i}function Gr(t){return Zr(function(n){n.setDate(n.getDate()-(n.getDay()+7-t)%7),n.setHours(0,0,0,0)},function(t,n){t.setDate(t.getDate()+7*n)},function(t,n){return(n-t-(n.getTimezoneOffset()-t.getTimezoneOffset())*Ay)/zy})}function Jr(t){return Zr(function(n){n.setUTCDate(n.getUTCDate()-(n.getUTCDay()+7-t)%7),n.setUTCHours(0,0,0,0)},function(t,n){t.setUTCDate(t.getUTCDate()+7*n)},function(t,n){return(n-t)/zy})}function Qr(t,n){if((e=(t=n?t.toExponential(n-1):t.toExponential()).indexOf("e"))<0)return null;var e,r=t.slice(0,e);return[r.length>1?r[0]+r.slice(2):r,+t.slice(e+1)]}function Kr(t){return t=Qr(Math.abs(t)),t?t[1]:NaN}function ti(t,n){return function(e,r){for(var i=e.length,o=[],u=0,a=t[0],c=0;i>0&&a>0&&(c+a+1>r&&(a=Math.max(1,r-c)),o.push(e.substring(i-=a,i+a)),!((c+=a+1)>r));)a=t[u=(u+1)%t.length];return o.reverse().join(n)}}function ni(t,n){t=t.toPrecision(n);t:for(var e,r=t.length,i=1,o=-1;i<r;++i)switch(t[i]){case".":o=e=i;break;case"0":0===o&&(o=i),e=i;break;case"e":break t;default:o>0&&(o=0)}return o>0?t.slice(0,o)+t.slice(e+1):t}function ei(t,n){var e=Qr(t,n);if(!e)return t+"";var r=e[0],i=e[1],o=i-(Ng=3*Math.max(-8,Math.min(8,Math.floor(i/3))))+1,u=r.length;return o===u?r:o>u?r+new Array(o-u+1).join("0"):o>0?r.slice(0,o)+"."+r.slice(o):"0."+new Array(1-o).join("0")+Qr(t,Math.max(0,n+o-1))[0]}function ri(t,n){var e=Qr(t,n);if(!e)return t+"";var r=e[0],i=e[1];return i<0?"0."+new Array(-i).join("0")+r:r.length>i+1?r.slice(0,i+1)+"."+r.slice(i+1):r+new Array(i-r.length+2).join("0")}function ii(t){return new oi(t)}function oi(t){if(!(n=Cg.exec(t)))throw new Error("invalid format: "+t);var n,e=n[1]||" ",r=n[2]||">",i=n[3]||"-",o=n[4]||"",u=!!n[5],a=n[6]&&+n[6],c=!!n[7],s=n[8]&&+n[8].slice(1),f=n[9]||"";"n"===f?(c=!0,f="g"):Eg[f]||(f=""),(u||"0"===e&&"="===r)&&(u=!0,e="0",r="="),this.fill=e,this.align=r,this.sign=i,this.symbol=o,this.zero=u,this.width=a,this.comma=c,this.precision=s,this.type=f}function ui(t){return t}function ai(t){function n(t){function n(t){var n,i,c,g=d,m=v;if("c"===p)m=_(t)+m,t="";else{t=+t;var x=(t<0||1/t<0)&&(t*=-1,!0);if(t=_(t,h),x)for(n=-1,i=t.length,x=!1;++n<i;)if(c=t.charCodeAt(n),48<c&&c<58||"x"===p&&96<c&&c<103||"X"===p&&64<c&&c<71){x=!0;break}if(g=(x?"("===a?a:"-":"-"===a||"("===a?"":a)+g,m=m+("s"===p?Pg[8+Ng/3]:"")+(x&&"("===a?")":""),y)for(n=-1,i=t.length;++n<i;)if(c=t.charCodeAt(n),48>c||c>57){m=(46===c?o+t.slice(n+1):t.slice(n))+m,t=t.slice(0,n);break}}l&&!s&&(t=r(t,1/0));var b=g.length+t.length+m.length,w=b<f?new Array(f-b+1).join(e):"";switch(l&&s&&(t=r(w+t,w.length?f-m.length:1/0),w=""),u){case"<":return g+t+m+w;case"=":return g+w+t+m;case"^":return w.slice(0,b=w.length>>1)+g+t+m+w.slice(b)}return w+g+t+m}t=ii(t);var e=t.fill,u=t.align,a=t.sign,c=t.symbol,s=t.zero,f=t.width,l=t.comma,h=t.precision,p=t.type,d="$"===c?i[0]:"#"===c&&/[boxX]/.test(p)?"0"+p.toLowerCase():"",v="$"===c?i[1]:/[%p]/.test(p)?"%":"",_=Eg[p],y=!p||/[defgprs%]/.test(p);return h=null==h?p?6:12:/[gprs]/.test(p)?Math.max(1,Math.min(21,h)):Math.max(0,Math.min(20,h)),n.toString=function(){return t+""},n}function e(t,e){var r=n((t=ii(t),t.type="f",t)),i=3*Math.max(-8,Math.min(8,Math.floor(Kr(e)/3))),o=Math.pow(10,-i),u=Pg[8+i/3];return function(t){return r(o*t)+u}}var r=t.grouping&&t.thousands?ti(t.grouping,t.thousands):ui,i=t.currency,o=t.decimal;return{format:n,formatPrefix:e}}function ci(n){return zg=ai(n),t.format=zg.format,t.formatPrefix=zg.formatPrefix,zg}function si(t){return Math.max(0,-Kr(Math.abs(t)))}function fi(t,n){return Math.max(0,3*Math.max(-8,Math.min(8,Math.floor(Kr(n)/3)))-Kr(Math.abs(t)))}function li(t,n){return t=Math.abs(t),n=Math.abs(n)-t,Math.max(0,Kr(n)-Kr(t))+1}function hi(t){if(0<=t.y&&t.y<100){var n=new Date(-1,t.m,t.d,t.H,t.M,t.S,t.L);return n.setFullYear(t.y),n}return new Date(t.y,t.m,t.d,t.H,t.M,t.S,t.L)}function pi(t){if(0<=t.y&&t.y<100){var n=new Date(Date.UTC(-1,t.m,t.d,t.H,t.M,t.S,t.L));return n.setUTCFullYear(t.y),n}return new Date(Date.UTC(t.y,t.m,t.d,t.H,t.M,t.S,t.L))}function di(t){return{y:t,m:0,d:1,H:0,M:0,S:0,L:0}}function vi(t){function n(t,n){return function(e){var r,i,o,u=[],a=-1,c=0,s=t.length;for(e instanceof Date||(e=new Date(+e));++a<s;)37===t.charCodeAt(a)&&(u.push(t.slice(c,a)),null!=(i=Lg[r=t.charAt(++a)])?r=t.charAt(++a):i="e"===r?" ":"0",(o=n[r])&&(r=o(e,i)),u.push(r),c=a+1);return u.push(t.slice(c,a)),u.join("")}}function e(t,n){return function(e){var i=di(1900),o=r(i,t,e+="",0);if(o!=e.length)return null;if("p"in i&&(i.H=i.H%12+12*i.p),"W"in i||"U"in i){"w"in i||(i.w="W"in i?1:0);var u="Z"in i?pi(di(i.y)).getUTCDay():n(di(i.y)).getDay();i.m=0,i.d="W"in i?(i.w+6)%7+7*i.W-(u+5)%7:i.w+7*i.U-(u+6)%7}return"Z"in i?(i.H+=i.Z/100|0,i.M+=i.Z%100,pi(i)):n(i)}}function r(t,n,e,r){for(var i,o,u=0,a=n.length,c=e.length;u<a;){if(r>=c)return-1;if(i=n.charCodeAt(u++),37===i){if(i=n.charAt(u++),o=B[i in Lg?n.charAt(u++):i],!o||(r=o(t,e,r))<0)return-1}else if(i!=e.charCodeAt(r++))return-1}return r}function i(t,n,e){var r=C.exec(n.slice(e));return r?(t.p=z[r[0].toLowerCase()],e+r[0].length):-1}function o(t,n,e){var r=L.exec(n.slice(e));return r?(t.w=R[r[0].toLowerCase()],e+r[0].length):-1}function u(t,n,e){var r=P.exec(n.slice(e));return r?(t.w=q[r[0].toLowerCase()],e+r[0].length):-1}function a(t,n,e){var r=O.exec(n.slice(e));return r?(t.m=F[r[0].toLowerCase()],e+r[0].length):-1}function c(t,n,e){var r=U.exec(n.slice(e));return r?(t.m=D[r[0].toLowerCase()],e+r[0].length):-1}function s(t,n,e){return r(t,w,n,e)}function f(t,n,e){return r(t,M,n,e)}function l(t,n,e){return r(t,T,n,e)}function h(t){return N[t.getDay()]}function p(t){return S[t.getDay()]}function d(t){return E[t.getMonth()]}function v(t){return A[t.getMonth()]}function _(t){return k[+(t.getHours()>=12)]}function y(t){return N[t.getUTCDay()]}function g(t){return S[t.getUTCDay()]}function m(t){return E[t.getUTCMonth()]}function x(t){return A[t.getUTCMonth()]}function b(t){return k[+(t.getUTCHours()>=12)]}var w=t.dateTime,M=t.date,T=t.time,k=t.periods,S=t.days,N=t.shortDays,A=t.months,E=t.shortMonths,C=gi(k),z=mi(k),P=gi(S),q=mi(S),L=gi(N),R=mi(N),U=gi(A),D=mi(A),O=gi(E),F=mi(E),I={a:h,A:p,b:d,B:v,c:null,d:Li,e:Li,H:Ri,I:Ui,j:Di,L:Oi,m:Fi,M:Ii,p:_,S:Yi,U:Bi,w:ji,W:Hi,x:null,X:null,y:Xi,Y:Vi,Z:Wi,"%":co},Y={a:y,A:g,b:m,B:x,c:null,d:$i,e:$i,H:Zi,I:Gi,j:Ji,L:Qi,m:Ki,M:to,p:b,S:no,U:eo,w:ro,W:io,x:null,X:null,y:oo,Y:uo,Z:ao,"%":co},B={a:o,A:u,b:a,B:c,c:s,d:Ni,e:Ni,H:Ei,I:Ei,j:Ai,L:Pi,m:Si,M:Ci,p:i,S:zi,U:bi,w:xi,W:wi,x:f,X:l,y:Ti,Y:Mi,Z:ki,"%":qi};return I.x=n(M,I),I.X=n(T,I),I.c=n(w,I),Y.x=n(M,Y),Y.X=n(T,Y),Y.c=n(w,Y),{format:function(t){var e=n(t+="",I);return e.toString=function(){return t},e},parse:function(t){var n=e(t+="",hi);return n.toString=function(){return t},n},utcFormat:function(t){var e=n(t+="",Y);return e.toString=function(){return t},e},utcParse:function(t){var n=e(t,pi);return n.toString=function(){return t},n}}}function _i(t,n,e){var r=t<0?"-":"",i=(r?-t:t)+"",o=i.length;return r+(o<e?new Array(e-o+1).join(n)+i:i)}function yi(t){return t.replace(Dg,"\\$&")}function gi(t){return new RegExp("^(?:"+t.map(yi).join("|")+")","i")}function mi(t){for(var n={},e=-1,r=t.length;++e<r;)n[t[e].toLowerCase()]=e;return n}function xi(t,n,e){var r=Rg.exec(n.slice(e,e+1));return r?(t.w=+r[0],e+r[0].length):-1}function bi(t,n,e){var r=Rg.exec(n.slice(e));return r?(t.U=+r[0],e+r[0].length):-1}function wi(t,n,e){var r=Rg.exec(n.slice(e));return r?(t.W=+r[0],e+r[0].length):-1}function Mi(t,n,e){var r=Rg.exec(n.slice(e,e+4));return r?(t.y=+r[0],e+r[0].length):-1}function Ti(t,n,e){var r=Rg.exec(n.slice(e,e+2));return r?(t.y=+r[0]+(+r[0]>68?1900:2e3),e+r[0].length):-1}function ki(t,n,e){var r=/^(Z)|([+-]\d\d)(?:\:?(\d\d))?/.exec(n.slice(e,e+6));return r?(t.Z=r[1]?0:-(r[2]+(r[3]||"00")),e+r[0].length):-1}function Si(t,n,e){var r=Rg.exec(n.slice(e,e+2));return r?(t.m=r[0]-1,e+r[0].length):-1}function Ni(t,n,e){var r=Rg.exec(n.slice(e,e+2));return r?(t.d=+r[0],e+r[0].length):-1}function Ai(t,n,e){var r=Rg.exec(n.slice(e,e+3));return r?(t.m=0,t.d=+r[0],e+r[0].length):-1}function Ei(t,n,e){var r=Rg.exec(n.slice(e,e+2));return r?(t.H=+r[0],e+r[0].length):-1}function Ci(t,n,e){var r=Rg.exec(n.slice(e,e+2));return r?(t.M=+r[0],e+r[0].length):-1}function zi(t,n,e){var r=Rg.exec(n.slice(e,e+2));return r?(t.S=+r[0],e+r[0].length):-1}function Pi(t,n,e){var r=Rg.exec(n.slice(e,e+3));return r?(t.L=+r[0],e+r[0].length):-1}function qi(t,n,e){var r=Ug.exec(n.slice(e,e+1));return r?e+r[0].length:-1}function Li(t,n){return _i(t.getDate(),n,2)}function Ri(t,n){return _i(t.getHours(),n,2)}function Ui(t,n){return _i(t.getHours()%12||12,n,2)}function Di(t,n){return _i(1+Oy.count(eg(t),t),n,3)}function Oi(t,n){return _i(t.getMilliseconds(),n,3)}function Fi(t,n){return _i(t.getMonth()+1,n,2)}function Ii(t,n){return _i(t.getMinutes(),n,2)}function Yi(t,n){return _i(t.getSeconds(),n,2)}function Bi(t,n){return _i(Iy.count(eg(t),t),n,2)}function ji(t){return t.getDay()}function Hi(t,n){return _i(Yy.count(eg(t),t),n,2)}function Xi(t,n){return _i(t.getFullYear()%100,n,2)}function Vi(t,n){return _i(t.getFullYear()%1e4,n,4)}function Wi(t){var n=t.getTimezoneOffset();return(n>0?"-":(n*=-1,"+"))+_i(n/60|0,"0",2)+_i(n%60,"0",2)}function $i(t,n){return _i(t.getUTCDate(),n,2)}function Zi(t,n){return _i(t.getUTCHours(),n,2)}function Gi(t,n){return _i(t.getUTCHours()%12||12,n,2)}function Ji(t,n){return _i(1+cg.count(Sg(t),t),n,3)}function Qi(t,n){return _i(t.getUTCMilliseconds(),n,3)}function Ki(t,n){return _i(t.getUTCMonth()+1,n,2)}function to(t,n){return _i(t.getUTCMinutes(),n,2)}function no(t,n){return _i(t.getUTCSeconds(),n,2)}function eo(t,n){return _i(fg.count(Sg(t),t),n,2)}function ro(t){return t.getUTCDay()}function io(t,n){return _i(lg.count(Sg(t),t),n,2)}function oo(t,n){return _i(t.getUTCFullYear()%100,n,2)}function uo(t,n){return _i(t.getUTCFullYear()%1e4,n,4)}function ao(){return"+0000"}function co(){return"%"}function so(n){return qg=vi(n),t.timeFormat=qg.format,t.timeParse=qg.parse,t.utcFormat=qg.utcFormat,t.utcParse=qg.utcParse,qg}function fo(t){return t.toISOString()}function lo(t){var n=new Date(t);return isNaN(n)?null:n}function ho(t){function n(n){var o=n+"",u=e.get(o);if(!u){if(i!==Hg)return i;e.set(o,u=r.push(n))}return t[(u-1)%t.length]}var e=q(),r=[],i=Hg;return t=null==t?[]:jg.call(t),n.domain=function(t){if(!arguments.length)return r.slice();r=[],e=q();for(var i,o,u=-1,a=t.length;++u<a;)e.has(o=(i=t[u])+"")||e.set(o,r.push(i));return n},n.range=function(e){return arguments.length?(t=jg.call(e),n):t.slice()},n.unknown=function(t){return arguments.length?(i=t,n):i},n.copy=function(){return ho().domain(r).range(t).unknown(i)},n}function po(){function t(){var t=i().length,r=u[1]<u[0],h=u[r-0],p=u[1-r];n=(p-h)/Math.max(1,t-c+2*s),a&&(n=Math.floor(n)),h+=(p-h-n*(t-c))*f,e=n*(1-c),a&&(h=Math.round(h),e=Math.round(e));var d=l(t).map(function(t){return h+n*t});return o(r?d.reverse():d)}var n,e,r=ho().unknown(void 0),i=r.domain,o=r.range,u=[0,1],a=!1,c=0,s=0,f=.5;return delete r.unknown,r.domain=function(n){return arguments.length?(i(n),t()):i()},r.range=function(n){return arguments.length?(u=[+n[0],+n[1]],t()):u.slice()},r.rangeRound=function(n){return u=[+n[0],+n[1]],a=!0,t()},r.bandwidth=function(){return e},r.step=function(){return n},r.round=function(n){return arguments.length?(a=!!n,t()):a},r.padding=function(n){return arguments.length?(c=s=Math.max(0,Math.min(1,n)),t()):c},r.paddingInner=function(n){return arguments.length?(c=Math.max(0,Math.min(1,n)),t()):c},r.paddingOuter=function(n){return arguments.length?(s=Math.max(0,Math.min(1,n)),t()):s},r.align=function(n){return arguments.length?(f=Math.max(0,Math.min(1,n)),t()):f},r.copy=function(){return po().domain(i()).range(u).round(a).paddingInner(c).paddingOuter(s).align(f)},t()}function vo(t){var n=t.copy;return t.padding=t.paddingOuter,delete t.paddingInner,delete t.paddingOuter,t.copy=function(){return vo(n())},t}function _o(){return vo(po().paddingInner(1))}function yo(t){return function(){return t}}function go(t){return+t}function mo(t,n){return(n-=t=+t)?function(e){return(e-t)/n}:yo(n)}function xo(t){return function(n,e){var r=t(n=+n,e=+e);return function(t){return t<=n?0:t>=e?1:r(t)}}}function bo(t){return function(n,e){var r=t(n=+n,e=+e);return function(t){return t<=0?n:t>=1?e:r(t)}}}function wo(t,n,e,r){var i=t[0],o=t[1],u=n[0],a=n[1];return o<i?(i=e(o,i),u=r(a,u)):(i=e(i,o),u=r(u,a)),function(t){return u(i(t))}}function Mo(t,n,e,r){var i=Math.min(t.length,n.length)-1,o=new Array(i),u=new Array(i),a=-1;for(t[i]<t[0]&&(t=t.slice().reverse(),n=n.slice().reverse());++a<i;)o[a]=e(t[a],t[a+1]),u[a]=r(n[a],n[a+1]);return function(n){var e=Cd(t,n,1,i)-1;return u[e](o[e](n))}}function To(t,n){return n.domain(t.domain()).range(t.range()).interpolate(t.interpolate()).clamp(t.clamp())}function ko(t,n){function e(){return i=Math.min(a.length,c.length)>2?Mo:wo,o=u=null,r}function r(n){return(o||(o=i(a,c,f?xo(t):t,s)))(+n)}var i,o,u,a=Xg,c=Xg,s=cr,f=!1;return r.invert=function(t){return(u||(u=i(c,a,mo,f?bo(n):n)))(+t)},r.domain=function(t){return arguments.length?(a=Bg.call(t,go),e()):a.slice()},r.range=function(t){return arguments.length?(c=jg.call(t),e()):c.slice()},r.rangeRound=function(t){return c=jg.call(t),s=sr,e()},r.clamp=function(t){return arguments.length?(f=!!t,e()):f},r.interpolate=function(t){return arguments.length?(s=t,e()):s},e()}function So(n,e,r){var i,o=n[0],u=n[n.length-1],a=p(o,u,null==e?10:e);switch(r=ii(null==r?",f":r),r.type){case"s":var c=Math.max(Math.abs(o),Math.abs(u));return null!=r.precision||isNaN(i=fi(a,c))||(r.precision=i),t.formatPrefix(r,c);case"":case"e":case"g":case"p":case"r":null!=r.precision||isNaN(i=li(a,Math.max(Math.abs(o),Math.abs(u))))||(r.precision=i-("e"===r.type));break;case"f":case"%":null!=r.precision||isNaN(i=si(a))||(r.precision=i-2*("%"===r.type))}return t.format(r)}function No(t){var n=t.domain;return t.ticks=function(t){var e=n();return h(e[0],e[e.length-1],null==t?10:t)},t.tickFormat=function(t,e){return So(n(),t,e)},t.nice=function(e){var r=n(),i=r.length-1,o=null==e?10:e,u=r[0],a=r[i],c=p(u,a,o);return c&&(c=p(Math.floor(u/c)*c,Math.ceil(a/c)*c,o),r[0]=Math.floor(u/c)*c,r[i]=Math.ceil(a/c)*c,n(r)),t},t}function Ao(){var t=ko(mo,rr);return t.copy=function(){return To(t,Ao())},No(t)}function Eo(){function t(t){return+t}var n=[0,1];return t.invert=t,t.domain=t.range=function(e){return arguments.length?(n=Bg.call(e,go),t):n.slice()},t.copy=function(){return Eo().domain(n)},No(t)}function Co(t,n){t=t.slice();var e,r=0,i=t.length-1,o=t[r],u=t[i];return u<o&&(e=r,r=i,i=e,e=o,o=u,u=e),t[r]=n.floor(o),t[i]=n.ceil(u),t}function zo(t,n){return(n=Math.log(n/t))?function(e){return Math.log(e/t)/n}:yo(n)}function Po(t,n){return t<0?function(e){return-Math.pow(-n,e)*Math.pow(-t,1-e)}:function(e){return Math.pow(n,e)*Math.pow(t,1-e)}}function qo(t){return isFinite(t)?+("1e"+t):t<0?0:t}function Lo(t){return 10===t?qo:t===Math.E?Math.exp:function(n){return Math.pow(t,n)}}function Ro(t){return t===Math.E?Math.log:10===t&&Math.log10||2===t&&Math.log2||(t=Math.log(t),function(n){return Math.log(n)/t})}function Uo(t){return function(n){return-t(-n)}}function Do(){function n(){return o=Ro(i),u=Lo(i),r()[0]<0&&(o=Uo(o),u=Uo(u)),e}var e=ko(zo,Po).domain([1,10]),r=e.domain,i=10,o=Ro(10),u=Lo(10);return e.base=function(t){return arguments.length?(i=+t,n()):i},e.domain=function(t){return arguments.length?(r(t),n()):r()},e.ticks=function(t){var n,e=r(),a=e[0],c=e[e.length-1];(n=c<a)&&(p=a,a=c,c=p);var s,f,l,p=o(a),d=o(c),v=null==t?10:+t,_=[];if(!(i%1)&&d-p<v){if(p=Math.round(p)-1,d=Math.round(d)+1,a>0){for(;p<d;++p)for(f=1,s=u(p);f<i;++f)if(l=s*f,!(l<a)){if(l>c)break;_.push(l)}}else for(;p<d;++p)for(f=i-1,s=u(p);f>=1;--f)if(l=s*f,!(l<a)){if(l>c)break;_.push(l)}}else _=h(p,d,Math.min(d-p,v)).map(u);return n?_.reverse():_},e.tickFormat=function(n,r){if(null==r&&(r=10===i?".0e":","),"function"!=typeof r&&(r=t.format(r)),n===1/0)return r;null==n&&(n=10);var a=Math.max(1,i*n/e.ticks().length);return function(t){var n=t/u(Math.round(o(t)));return n*i<i-.5&&(n*=i),n<=a?r(t):""}},e.nice=function(){return r(Co(r(),{floor:function(t){return u(Math.floor(o(t)))},ceil:function(t){return u(Math.ceil(o(t)))}}))},e.copy=function(){return To(e,Do().base(i))},e}function Oo(t,n){return t<0?-Math.pow(-t,n):Math.pow(t,n)}function Fo(){function t(t,n){return(n=Oo(n,e)-(t=Oo(t,e)))?function(r){return(Oo(r,e)-t)/n}:yo(n)}function n(t,n){return n=Oo(n,e)-(t=Oo(t,e)),function(r){return Oo(t+n*r,1/e)}}var e=1,r=ko(t,n),i=r.domain;return r.exponent=function(t){return arguments.length?(e=+t,i(i())):e},r.copy=function(){return To(r,Fo().exponent(e))},No(r)}function Io(){return Fo().exponent(.5)}function Yo(){function t(){var t=0,n=Math.max(1,i.length);for(o=new Array(n-1);++t<n;)o[t-1]=_(r,t/n);return e}function e(t){if(!isNaN(t=+t))return i[Cd(o,t)]}var r=[],i=[],o=[];return e.invertExtent=function(t){var n=i.indexOf(t);return n<0?[NaN,NaN]:[n>0?o[n-1]:r[0],n<o.length?o[n]:r[r.length-1]]},e.domain=function(e){if(!arguments.length)return r.slice();r=[];for(var i,o=0,u=e.length;o<u;++o)i=e[o],null==i||isNaN(i=+i)||r.push(i);return r.sort(n),t()},e.range=function(n){return arguments.length?(i=jg.call(n),t()):i.slice()},e.quantiles=function(){return o.slice()},e.copy=function(){return Yo().domain(r).range(i)},e}function Bo(){function t(t){if(t<=t)return u[Cd(o,t,0,i)]}function n(){var n=-1;for(o=new Array(i);++n<i;)o[n]=((n+1)*r-(n-i)*e)/(i+1);return t}var e=0,r=1,i=1,o=[.5],u=[0,1];return t.domain=function(t){return arguments.length?(e=+t[0],r=+t[1],n()):[e,r]},t.range=function(t){return arguments.length?(i=(u=jg.call(t)).length-1,n()):u.slice()},t.invertExtent=function(t){var n=u.indexOf(t);return n<0?[NaN,NaN]:n<1?[e,o[0]]:n>=i?[o[i-1],r]:[o[n-1],o[n]]},t.copy=function(){return Bo().domain([e,r]).range(u)},No(t)}function jo(){function t(t){if(t<=t)return e[Cd(n,t,0,r)]}var n=[.5],e=[0,1],r=1;return t.domain=function(i){return arguments.length?(n=jg.call(i),r=Math.min(n.length,e.length-1),t):n.slice()},t.range=function(i){return arguments.length?(e=jg.call(i),r=Math.min(n.length,e.length-1),t):e.slice()},t.invertExtent=function(t){var r=e.indexOf(t);return[n[r-1],n[r]]},t.copy=function(){return jo().domain(n).range(e)},t}function Ho(t){return new Date(t)}function Xo(t){return t instanceof Date?+t:+new Date(+t)}function Vo(t,n,r,i,o,u,a,c,s){function f(e){return(a(e)<e?_:u(e)<e?y:o(e)<e?g:i(e)<e?m:n(e)<e?r(e)<e?x:b:t(e)<e?w:M)(e)}function l(n,r,i,o){if(null==n&&(n=10),"number"==typeof n){var u=Math.abs(i-r)/n,a=e(function(t){return t[2]}).right(T,u);a===T.length?(o=p(r/Qg,i/Qg,n),n=t):a?(a=T[u/T[a-1][2]<T[a][2]/u?a-1:a],o=a[1],n=a[0]):(o=p(r,i,n),n=c)}return null==o?n:n.every(o)}var h=ko(mo,rr),d=h.invert,v=h.domain,_=s(".%L"),y=s(":%S"),g=s("%I:%M"),m=s("%I %p"),x=s("%a %d"),b=s("%b %d"),w=s("%B"),M=s("%Y"),T=[[a,1,Vg],[a,5,5*Vg],[a,15,15*Vg],[a,30,30*Vg],[u,1,Wg],[u,5,5*Wg],[u,15,15*Wg],[u,30,30*Wg],[o,1,$g],[o,3,3*$g],[o,6,6*$g],[o,12,12*$g],[i,1,Zg],[i,2,2*Zg],[r,1,Gg],[n,1,Jg],[n,3,3*Jg],[t,1,Qg]];return h.invert=function(t){return new Date(d(t))},h.domain=function(t){return arguments.length?v(Bg.call(t,Xo)):v().map(Ho)},h.ticks=function(t,n){var e,r=v(),i=r[0],o=r[r.length-1],u=o<i;return u&&(e=i,i=o,o=e),e=l(t,i,o,n),e=e?e.range(i,o+1):[],u?e.reverse():e},h.tickFormat=function(t,n){return null==n?f:s(n)},h.nice=function(t,n){var e=v();return(t=l(t,e[0],e[e.length-1],n))?v(Co(e,t)):h},h.copy=function(){return To(h,Vo(t,n,r,i,o,u,a,c,s))},h}function Wo(){return Vo(eg,tg,Iy,Oy,Uy,Ly,Py,ky,t.timeFormat).domain([new Date(2e3,0,1),new Date(2e3,0,2)])}function $o(){return Vo(Sg,Tg,fg,cg,ug,ig,Py,ky,t.utcFormat).domain([Date.UTC(2e3,0,1),Date.UTC(2e3,0,2)])}function Zo(t){return t.match(/.{6}/g).map(function(t){return"#"+t})}function Go(t){(t<0||t>1)&&(t-=Math.floor(t));var n=Math.abs(t-.5);return um.h=360*t-100,um.s=1.5-1.5*n,um.l=.8-.9*n,um+""}function Jo(t){var n=t.length;return function(e){return t[Math.max(0,Math.min(n-1,Math.floor(e*n)))]}}function Qo(t){function n(n){var o=(n-e)/(r-e);return t(i?Math.max(0,Math.min(1,o)):o)}var e=0,r=1,i=!1;return n.domain=function(t){return arguments.length?(e=+t[0],r=+t[1],n):[e,r]},n.clamp=function(t){return arguments.length?(i=!!t,n):i},n.interpolator=function(e){return arguments.length?(t=e,n):t},n.copy=function(){return Qo(t).domain([e,r]).clamp(i)},No(n)}function Ko(t){var n=t+="",e=n.indexOf(":");return e>=0&&"xmlns"!==(n=t.slice(0,e))&&(t=t.slice(e+1)),hm.hasOwnProperty(n)?{space:hm[n],local:t}:t}function tu(t){return function(){var n=this.ownerDocument,e=this.namespaceURI;return e===lm&&n.documentElement.namespaceURI===lm?n.createElement(t):n.createElementNS(e,t)}}function nu(t){return function(){return this.ownerDocument.createElementNS(t.space,t.local)}}function eu(t){var n=Ko(t);return(n.local?nu:tu)(n)}function ru(){return new iu}function iu(){this._="@"+(++pm).toString(36)}function ou(t,n,e){return t=uu(t,n,e),function(n){var e=n.relatedTarget;e&&(e===this||8&e.compareDocumentPosition(this))||t.call(this,n)}}function uu(n,e,r){return function(i){var o=t.event;t.event=i;try{n.call(this,this.__data__,e,r)}finally{t.event=o}}}function au(t){return t.trim().split(/^|\s+/).map(function(t){var n="",e=t.indexOf(".");return e>=0&&(n=t.slice(e+1),t=t.slice(0,e)),
{type:t,name:n}})}function cu(t){return function(){var n=this.__on;if(n){for(var e,r=0,i=-1,o=n.length;r<o;++r)e=n[r],t.type&&e.type!==t.type||e.name!==t.name?n[++i]=e:this.removeEventListener(e.type,e.listener,e.capture);++i?n.length=i:delete this.__on}}}function su(t,n,e){var r=gm.hasOwnProperty(t.type)?ou:uu;return function(i,o,u){var a,c=this.__on,s=r(n,o,u);if(c)for(var f=0,l=c.length;f<l;++f)if((a=c[f]).type===t.type&&a.name===t.name)return this.removeEventListener(a.type,a.listener,a.capture),this.addEventListener(a.type,a.listener=s,a.capture=e),void(a.value=n);this.addEventListener(t.type,s,e),a={type:t.type,name:t.name,value:n,listener:s,capture:e},c?c.push(a):this.__on=[a]}}function fu(t,n,e){var r,i,o=au(t+""),u=o.length;{if(!(arguments.length<2)){for(a=n?su:cu,null==e&&(e=!1),r=0;r<u;++r)this.each(a(o[r],n,e));return this}var a=this.node().__on;if(a)for(var c,s=0,f=a.length;s<f;++s)for(r=0,c=a[s];r<u;++r)if((i=o[r]).type===c.type&&i.name===c.name)return c.value}}function lu(n,e,r,i){var o=t.event;n.sourceEvent=t.event,t.event=n;try{return e.apply(r,i)}finally{t.event=o}}function hu(){for(var n,e=t.event;n=e.sourceEvent;)e=n;return e}function pu(t,n){var e=t.ownerSVGElement||t;if(e.createSVGPoint){var r=e.createSVGPoint();return r.x=n.clientX,r.y=n.clientY,r=r.matrixTransform(t.getScreenCTM().inverse()),[r.x,r.y]}var i=t.getBoundingClientRect();return[n.clientX-i.left-t.clientLeft,n.clientY-i.top-t.clientTop]}function du(t){var n=hu();return n.changedTouches&&(n=n.changedTouches[0]),pu(t,n)}function vu(){}function _u(t){return null==t?vu:function(){return this.querySelector(t)}}function yu(t){"function"!=typeof t&&(t=_u(t));for(var n=this._groups,e=n.length,r=new Array(e),i=0;i<e;++i)for(var o,u,a=n[i],c=a.length,s=r[i]=new Array(c),f=0;f<c;++f)(o=a[f])&&(u=t.call(o,o.__data__,f,a))&&("__data__"in o&&(u.__data__=o.__data__),s[f]=u);return new qa(r,this._parents)}function gu(){return[]}function mu(t){return null==t?gu:function(){return this.querySelectorAll(t)}}function xu(t){"function"!=typeof t&&(t=mu(t));for(var n=this._groups,e=n.length,r=[],i=[],o=0;o<e;++o)for(var u,a=n[o],c=a.length,s=0;s<c;++s)(u=a[s])&&(r.push(t.call(u,u.__data__,s,a)),i.push(u));return new qa(r,i)}function bu(t){"function"!=typeof t&&(t=ym(t));for(var n=this._groups,e=n.length,r=new Array(e),i=0;i<e;++i)for(var o,u=n[i],a=u.length,c=r[i]=[],s=0;s<a;++s)(o=u[s])&&t.call(o,o.__data__,s,u)&&c.push(o);return new qa(r,this._parents)}function wu(t){return new Array(t.length)}function Mu(){return new qa(this._enter||this._groups.map(wu),this._parents)}function Tu(t,n){this.ownerDocument=t.ownerDocument,this.namespaceURI=t.namespaceURI,this._next=null,this._parent=t,this.__data__=n}function ku(t){return function(){return t}}function Su(t,n,e,r,i,o){for(var u,a=0,c=n.length,s=o.length;a<s;++a)(u=n[a])?(u.__data__=o[a],r[a]=u):e[a]=new Tu(t,o[a]);for(;a<c;++a)(u=n[a])&&(i[a]=u)}function Nu(t,n,e,r,i,o,u){var a,c,s,f={},l=n.length,h=o.length,p=new Array(l);for(a=0;a<l;++a)(c=n[a])&&(p[a]=s=xm+u.call(c,c.__data__,a,n),s in f?i[a]=c:f[s]=c);for(a=0;a<h;++a)s=xm+u.call(t,o[a],a,o),(c=f[s])?(r[a]=c,c.__data__=o[a],f[s]=null):e[a]=new Tu(t,o[a]);for(a=0;a<l;++a)(c=n[a])&&f[p[a]]===c&&(i[a]=c)}function Au(t,n){if(!t)return p=new Array(this.size()),s=-1,this.each(function(t){p[++s]=t}),p;var e=n?Nu:Su,r=this._parents,i=this._groups;"function"!=typeof t&&(t=ku(t));for(var o=i.length,u=new Array(o),a=new Array(o),c=new Array(o),s=0;s<o;++s){var f=r[s],l=i[s],h=l.length,p=t.call(f,f&&f.__data__,s,r),d=p.length,v=a[s]=new Array(d),_=u[s]=new Array(d),y=c[s]=new Array(h);e(f,l,v,_,y,p,n);for(var g,m,x=0,b=0;x<d;++x)if(g=v[x]){for(x>=b&&(b=x+1);!(m=_[b])&&++b<d;);g._next=m||null}}return u=new qa(u,r),u._enter=a,u._exit=c,u}function Eu(){return new qa(this._exit||this._groups.map(wu),this._parents)}function Cu(t){for(var n=this._groups,e=t._groups,r=n.length,i=e.length,o=Math.min(r,i),u=new Array(r),a=0;a<o;++a)for(var c,s=n[a],f=e[a],l=s.length,h=u[a]=new Array(l),p=0;p<l;++p)(c=s[p]||f[p])&&(h[p]=c);for(;a<r;++a)u[a]=n[a];return new qa(u,this._parents)}function zu(){for(var t=this._groups,n=-1,e=t.length;++n<e;)for(var r,i=t[n],o=i.length-1,u=i[o];--o>=0;)(r=i[o])&&(u&&u!==r.nextSibling&&u.parentNode.insertBefore(r,u),u=r);return this}function Pu(t){function n(n,e){return n&&e?t(n.__data__,e.__data__):!n-!e}t||(t=qu);for(var e=this._groups,r=e.length,i=new Array(r),o=0;o<r;++o){for(var u,a=e[o],c=a.length,s=i[o]=new Array(c),f=0;f<c;++f)(u=a[f])&&(s[f]=u);s.sort(n)}return new qa(i,this._parents).order()}function qu(t,n){return t<n?-1:t>n?1:t>=n?0:NaN}function Lu(){var t=arguments[0];return arguments[0]=this,t.apply(null,arguments),this}function Ru(){var t=new Array(this.size()),n=-1;return this.each(function(){t[++n]=this}),t}function Uu(){for(var t=this._groups,n=0,e=t.length;n<e;++n)for(var r=t[n],i=0,o=r.length;i<o;++i){var u=r[i];if(u)return u}return null}function Du(){var t=0;return this.each(function(){++t}),t}function Ou(){return!this.node()}function Fu(t){for(var n=this._groups,e=0,r=n.length;e<r;++e)for(var i,o=n[e],u=0,a=o.length;u<a;++u)(i=o[u])&&t.call(i,i.__data__,u,o);return this}function Iu(t){return function(){this.removeAttribute(t)}}function Yu(t){return function(){this.removeAttributeNS(t.space,t.local)}}function Bu(t,n){return function(){this.setAttribute(t,n)}}function ju(t,n){return function(){this.setAttributeNS(t.space,t.local,n)}}function Hu(t,n){return function(){var e=n.apply(this,arguments);null==e?this.removeAttribute(t):this.setAttribute(t,e)}}function Xu(t,n){return function(){var e=n.apply(this,arguments);null==e?this.removeAttributeNS(t.space,t.local):this.setAttributeNS(t.space,t.local,e)}}function Vu(t,n){var e=Ko(t);if(arguments.length<2){var r=this.node();return e.local?r.getAttributeNS(e.space,e.local):r.getAttribute(e)}return this.each((null==n?e.local?Yu:Iu:"function"==typeof n?e.local?Xu:Hu:e.local?ju:Bu)(e,n))}function Wu(t){return t.ownerDocument&&t.ownerDocument.defaultView||t.document&&t||t.defaultView}function $u(t){return function(){this.style.removeProperty(t)}}function Zu(t,n,e){return function(){this.style.setProperty(t,n,e)}}function Gu(t,n,e){return function(){var r=n.apply(this,arguments);null==r?this.style.removeProperty(t):this.style.setProperty(t,r,e)}}function Ju(t,n,e){var r;return arguments.length>1?this.each((null==n?$u:"function"==typeof n?Gu:Zu)(t,n,null==e?"":e)):Wu(r=this.node()).getComputedStyle(r,null).getPropertyValue(t)}function Qu(t){return function(){delete this[t]}}function Ku(t,n){return function(){this[t]=n}}function ta(t,n){return function(){var e=n.apply(this,arguments);null==e?delete this[t]:this[t]=e}}function na(t,n){return arguments.length>1?this.each((null==n?Qu:"function"==typeof n?ta:Ku)(t,n)):this.node()[t]}function ea(t){return t.trim().split(/^|\s+/)}function ra(t){return t.classList||new ia(t)}function ia(t){this._node=t,this._names=ea(t.getAttribute("class")||"")}function oa(t,n){for(var e=ra(t),r=-1,i=n.length;++r<i;)e.add(n[r])}function ua(t,n){for(var e=ra(t),r=-1,i=n.length;++r<i;)e.remove(n[r])}function aa(t){return function(){oa(this,t)}}function ca(t){return function(){ua(this,t)}}function sa(t,n){return function(){(n.apply(this,arguments)?oa:ua)(this,t)}}function fa(t,n){var e=ea(t+"");if(arguments.length<2){for(var r=ra(this.node()),i=-1,o=e.length;++i<o;)if(!r.contains(e[i]))return!1;return!0}return this.each(("function"==typeof n?sa:n?aa:ca)(e,n))}function la(){this.textContent=""}function ha(t){return function(){this.textContent=t}}function pa(t){return function(){var n=t.apply(this,arguments);this.textContent=null==n?"":n}}function da(t){return arguments.length?this.each(null==t?la:("function"==typeof t?pa:ha)(t)):this.node().textContent}function va(){this.innerHTML=""}function _a(t){return function(){this.innerHTML=t}}function ya(t){return function(){var n=t.apply(this,arguments);this.innerHTML=null==n?"":n}}function ga(t){return arguments.length?this.each(null==t?va:("function"==typeof t?ya:_a)(t)):this.node().innerHTML}function ma(){this.nextSibling&&this.parentNode.appendChild(this)}function xa(){return this.each(ma)}function ba(){this.previousSibling&&this.parentNode.insertBefore(this,this.parentNode.firstChild)}function wa(){return this.each(ba)}function Ma(t){var n="function"==typeof t?t:eu(t);return this.select(function(){return this.appendChild(n.apply(this,arguments))})}function Ta(){return null}function ka(t,n){var e="function"==typeof t?t:eu(t),r=null==n?Ta:"function"==typeof n?n:_u(n);return this.select(function(){return this.insertBefore(e.apply(this,arguments),r.apply(this,arguments)||null)})}function Sa(){var t=this.parentNode;t&&t.removeChild(this)}function Na(){return this.each(Sa)}function Aa(t){return arguments.length?this.property("__data__",t):this.node().__data__}function Ea(t,n,e){var r=Wu(t),i=r.CustomEvent;i?i=new i(n,e):(i=r.document.createEvent("Event"),e?(i.initEvent(n,e.bubbles,e.cancelable),i.detail=e.detail):i.initEvent(n,!1,!1)),t.dispatchEvent(i)}function Ca(t,n){return function(){return Ea(this,t,n)}}function za(t,n){return function(){return Ea(this,t,n.apply(this,arguments))}}function Pa(t,n){return this.each(("function"==typeof n?za:Ca)(t,n))}function qa(t,n){this._groups=t,this._parents=n}function La(){return new qa([[document.documentElement]],bm)}function Ra(t){return"string"==typeof t?new qa([[document.querySelector(t)]],[document.documentElement]):new qa([[t]],bm)}function Ua(t){return"string"==typeof t?new qa([document.querySelectorAll(t)],[document.documentElement]):new qa([null==t?[]:t],bm)}function Da(t,n,e){arguments.length<3&&(e=n,n=hu().changedTouches);for(var r,i=0,o=n?n.length:0;i<o;++i)if((r=n[i]).identifier===e)return pu(t,r);return null}function Oa(t,n){null==n&&(n=hu().touches);for(var e=0,r=n?n.length:0,i=new Array(r);e<r;++e)i[e]=pu(t,n[e]);return i}function Fa(t,n,e,r,i,o){var u=t.__transition;if(u){if(e in u)return}else t.__transition={};ja(t,e,{name:n,index:r,group:i,on:wm,tween:Mm,time:o.time,delay:o.delay,duration:o.duration,ease:o.ease,timer:null,state:Tm})}function Ia(t,n){var e=t.__transition;if(!e||!(e=e[n])||e.state>Tm)throw new Error("too late");return e}function Ya(t,n){var e=t.__transition;if(!e||!(e=e[n])||e.state>Sm)throw new Error("too late");return e}function Ba(t,n){var e=t.__transition;if(!e||!(e=e[n]))throw new Error("too late");return e}function ja(t,n,e){function r(t){e.state=km,e.delay<=t?i(t-e.delay):e.timer.restart(i,e.delay,e.time)}function i(r){var i,c,s,f;for(i in a)f=a[i],f.name===e.name&&(f.state===Nm?(f.state=Em,f.timer.stop(),f.on.call("interrupt",t,t.__data__,f.index,f.group),delete a[i]):+i<n&&(f.state=Em,f.timer.stop(),delete a[i]));if(Wr(function(){e.state===Nm&&(e.timer.restart(o,e.delay,e.time),o(r))}),e.state=Sm,e.on.call("start",t,t.__data__,e.index,e.group),e.state===Sm){for(e.state=Nm,u=new Array(s=e.tween.length),i=0,c=-1;i<s;++i)(f=e.tween[i].value.call(t,t.__data__,e.index,e.group))&&(u[++c]=f);u.length=c+1}}function o(r){for(var i=r<e.duration?e.ease.call(null,r/e.duration):(e.state=Am,1),o=-1,c=u.length;++o<c;)u[o].call(null,i);if(e.state===Am){e.state=Em,e.timer.stop(),e.on.call("end",t,t.__data__,e.index,e.group);for(o in a)if(+o!==n)return void delete a[n];delete t.__transition}}var u,a=t.__transition;a[n]=e,e.timer=Yr(r,0,e.time)}function Ha(t,n){var e,r,i,o=t.__transition,u=!0;if(o){n=null==n?null:n+"";for(i in o)(e=o[i]).name===n?(r=e.state===Nm,e.state=Em,e.timer.stop(),r&&e.on.call("interrupt",t,t.__data__,e.index,e.group),delete o[i]):u=!1;u&&delete t.__transition}}function Xa(t){return this.each(function(){Ha(this,t)})}function Va(t,n){var e,r;return function(){var i=Ya(this,t),o=i.tween;if(o!==e){r=e=o;for(var u=0,a=r.length;u<a;++u)if(r[u].name===n){r=r.slice(),r.splice(u,1);break}}i.tween=r}}function Wa(t,n,e){var r,i;if("function"!=typeof e)throw new Error;return function(){var o=Ya(this,t),u=o.tween;if(u!==r){i=(r=u).slice();for(var a={name:n,value:e},c=0,s=i.length;c<s;++c)if(i[c].name===n){i[c]=a;break}c===s&&i.push(a)}o.tween=i}}function $a(t,n){var e=this._id;if(t+="",arguments.length<2){for(var r,i=Ba(this.node(),e).tween,o=0,u=i.length;o<u;++o)if((r=i[o]).name===t)return r.value;return null}return this.each((null==n?Va:Wa)(e,t,n))}function Za(t,n,e){var r=t._id;return t.each(function(){var t=Ya(this,r);(t.value||(t.value={}))[n]=e.apply(this,arguments)}),function(t){return Ba(t,r).value[n]}}function Ga(t,n){var e;return("number"==typeof n?rr:n instanceof be?C_:(e=be(n))?(n=e,C_):ar)(t,n)}function Ja(t){return function(){this.removeAttribute(t)}}function Qa(t){return function(){this.removeAttributeNS(t.space,t.local)}}function Ka(t,n,e){var r,i;return function(){var o=this.getAttribute(t);return o===e?null:o===r?i:i=n(r=o,e)}}function tc(t,n,e){var r,i;return function(){var o=this.getAttributeNS(t.space,t.local);return o===e?null:o===r?i:i=n(r=o,e)}}function nc(t,n,e){var r,i,o;return function(){var u,a=e(this);return null==a?void this.removeAttribute(t):(u=this.getAttribute(t),u===a?null:u===r&&a===i?o:o=n(r=u,i=a))}}function ec(t,n,e){var r,i,o;return function(){var u,a=e(this);return null==a?void this.removeAttributeNS(t.space,t.local):(u=this.getAttributeNS(t.space,t.local),u===a?null:u===r&&a===i?o:o=n(r=u,i=a))}}function rc(t,n){var e=Ko(t),r="transform"===e?O_:Ga;return this.attrTween(t,"function"==typeof n?(e.local?ec:nc)(e,r,Za(this,"attr."+t,n)):null==n?(e.local?Qa:Ja)(e):(e.local?tc:Ka)(e,r,n))}function ic(t,n){function e(){var e=this,r=n.apply(e,arguments);return r&&function(n){e.setAttributeNS(t.space,t.local,r(n))}}return e._value=n,e}function oc(t,n){function e(){var e=this,r=n.apply(e,arguments);return r&&function(n){e.setAttribute(t,r(n))}}return e._value=n,e}function uc(t,n){var e="attr."+t;if(arguments.length<2)return(e=this.tween(e))&&e._value;if(null==n)return this.tween(e,null);if("function"!=typeof n)throw new Error;var r=Ko(t);return this.tween(e,(r.local?ic:oc)(r,n))}function ac(t,n){return function(){Ia(this,t).delay=+n.apply(this,arguments)}}function cc(t,n){return n=+n,function(){Ia(this,t).delay=n}}function sc(t){var n=this._id;return arguments.length?this.each(("function"==typeof t?ac:cc)(n,t)):Ba(this.node(),n).delay}function fc(t,n){return function(){Ya(this,t).duration=+n.apply(this,arguments)}}function lc(t,n){return n=+n,function(){Ya(this,t).duration=n}}function hc(t){var n=this._id;return arguments.length?this.each(("function"==typeof t?fc:lc)(n,t)):Ba(this.node(),n).duration}function pc(t,n){if("function"!=typeof n)throw new Error;return function(){Ya(this,t).ease=n}}function dc(t){var n=this._id;return arguments.length?this.each(pc(n,t)):Ba(this.node(),n).ease}function vc(t){"function"!=typeof t&&(t=ym(t));for(var n=this._groups,e=n.length,r=new Array(e),i=0;i<e;++i)for(var o,u=n[i],a=u.length,c=r[i]=[],s=0;s<a;++s)(o=u[s])&&t.call(o,o.__data__,s,u)&&c.push(o);return new Uc(r,this._parents,this._name,this._id)}function _c(t){if(t._id!==this._id)throw new Error;for(var n=this._groups,e=t._groups,r=n.length,i=e.length,o=Math.min(r,i),u=new Array(r),a=0;a<o;++a)for(var c,s=n[a],f=e[a],l=s.length,h=u[a]=new Array(l),p=0;p<l;++p)(c=s[p]||f[p])&&(h[p]=c);for(;a<r;++a)u[a]=n[a];return new Uc(u,this._parents,this._name,this._id)}function yc(t){return(t+"").trim().split(/^|\s+/).every(function(t){var n=t.indexOf(".");return n>=0&&(t=t.slice(0,n)),!t||"start"===t})}function gc(t,n,e){var r,i,o=yc(n)?Ia:Ya;return function(){var u=o(this,t),a=u.on;a!==r&&(i=(r=a).copy()).on(n,e),u.on=i}}function mc(t,n){var e=this._id;return arguments.length<2?Ba(this.node(),e).on.on(t):this.each(gc(e,t,n))}function xc(t){return function(){var n=this.parentNode;for(var e in this.__transition)if(+e!==t)return;n&&n.removeChild(this)}}function bc(){return this.on("end.remove",xc(this._id))}function wc(t){var n=this._name,e=this._id;"function"!=typeof t&&(t=_u(t));for(var r=this._groups,i=r.length,o=new Array(i),u=0;u<i;++u)for(var a,c,s=r[u],f=s.length,l=o[u]=new Array(f),h=0;h<f;++h)(a=s[h])&&(c=t.call(a,a.__data__,h,s))&&("__data__"in a&&(c.__data__=a.__data__),l[h]=c,Fa(l[h],n,e,h,l,Ba(a,e)));return new Uc(o,this._parents,n,e)}function Mc(t){var n=this._name,e=this._id;"function"!=typeof t&&(t=mu(t));for(var r=this._groups,i=r.length,o=[],u=[],a=0;a<i;++a)for(var c,s=r[a],f=s.length,l=0;l<f;++l)if(c=s[l]){for(var h,p=t.call(c,c.__data__,l,s),d=Ba(c,e),v=0,_=p.length;v<_;++v)(h=p[v])&&Fa(h,n,e,v,p,d);o.push(p),u.push(c)}return new Uc(o,u,n,e)}function Tc(){return new Cm(this._groups,this._parents)}function kc(t,n){var e,r,i;return function(){var o=Wu(this).getComputedStyle(this,null),u=o.getPropertyValue(t),a=(this.style.removeProperty(t),o.getPropertyValue(t));return u===a?null:u===e&&a===r?i:i=n(e=u,r=a)}}function Sc(t){return function(){this.style.removeProperty(t)}}function Nc(t,n,e){var r,i;return function(){var o=Wu(this).getComputedStyle(this,null).getPropertyValue(t);return o===e?null:o===r?i:i=n(r=o,e)}}function Ac(t,n,e){var r,i,o;return function(){var u=Wu(this).getComputedStyle(this,null),a=u.getPropertyValue(t),c=e(this);return null==c&&(this.style.removeProperty(t),c=u.getPropertyValue(t)),a===c?null:a===r&&c===i?o:o=n(r=a,i=c)}}function Ec(t,n,e){var r="transform"==(t+="")?D_:Ga;return null==n?this.styleTween(t,kc(t,r)).on("end.style."+t,Sc(t)):this.styleTween(t,"function"==typeof n?Ac(t,r,Za(this,"style."+t,n)):Nc(t,r,n),e)}function Cc(t,n,e){function r(){var r=this,i=n.apply(r,arguments);return i&&function(n){r.style.setProperty(t,i(n),e)}}return r._value=n,r}function zc(t,n,e){var r="style."+(t+="");if(arguments.length<2)return(r=this.tween(r))&&r._value;if(null==n)return this.tween(r,null);if("function"!=typeof n)throw new Error;return this.tween(r,Cc(t,n,null==e?"":e))}function Pc(t){return function(){this.textContent=t}}function qc(t){return function(){var n=t(this);this.textContent=null==n?"":n}}function Lc(t){return this.tween("text","function"==typeof t?qc(Za(this,"text",t)):Pc(null==t?"":t+""))}function Rc(){for(var t=this._name,n=this._id,e=Oc(),r=this._groups,i=r.length,o=0;o<i;++o)for(var u,a=r[o],c=a.length,s=0;s<c;++s)if(u=a[s]){var f=Ba(u,n);Fa(u,t,e,s,a,{time:f.time+f.delay+f.duration,delay:0,duration:f.duration,ease:f.ease})}return new Uc(r,this._parents,t,e)}function Uc(t,n,e,r){this._groups=t,this._parents=n,this._name=e,this._id=r}function Dc(t){return La().transition(t)}function Oc(){return++zm}function Fc(t,n){for(var e;!(e=t.__transition)||!(e=e[n]);)if(!(t=t.parentNode))return qm.time=Or(),qm;return e}function Ic(t){var n,e;t instanceof Uc?(n=t._id,t=t._name):(n=Oc(),(e=qm).time=Or(),t=null==t?null:t+"");for(var r=this._groups,i=r.length,o=0;o<i;++o)for(var u,a=r[o],c=a.length,s=0;s<c;++s)(u=a[s])&&Fa(u,t,n,s,a,e||Fc(u,n));return new Uc(r,this._parents,t,n)}function Yc(t,n){var e,r,i=t.__transition;if(i){n=null==n?null:n+"";for(r in i)if((e=i[r]).state>km&&e.name===n)return new Uc([[t]],Lm,n,+r)}return null}function Bc(t){return t}function jc(t,n,e){var r=t(e);return"translate("+(isFinite(r)?r:n(e))+",0)"}function Hc(t,n,e){var r=t(e);return"translate(0,"+(isFinite(r)?r:n(e))+")"}function Xc(t){var n=t.bandwidth()/2;return t.round()&&(n=Math.round(n)),function(e){return t(e)+n}}function Vc(){return!this.__axis}function Wc(t,n){function e(e){var s,f=null==i?n.ticks?n.ticks.apply(n,r):n.domain():i,l=null==o?n.tickFormat?n.tickFormat.apply(n,r):Bc:o,h=Math.max(u,0)+c,p=t===Um||t===Om?jc:Hc,d=n.range(),v=d[0]+.5,_=d[d.length-1]+.5,y=(n.bandwidth?Xc:Bc)(n.copy()),g=e.selection?e.selection():e,m=g.selectAll(".domain").data([null]),x=g.selectAll(".tick").data(f,n).order(),b=x.exit(),w=x.enter().append("g").attr("class","tick"),M=x.select("line"),T=x.select("text"),k=t===Um||t===Fm?-1:1,S=t===Fm||t===Dm?(s="x","y"):(s="y","x");m=m.merge(m.enter().insert("path",".tick").attr("class","domain").attr("stroke","#000")),x=x.merge(w),M=M.merge(w.append("line").attr("stroke","#000").attr(s+"2",k*u).attr(S+"1",.5).attr(S+"2",.5)),T=T.merge(w.append("text").attr("fill","#000").attr(s,k*h).attr(S,.5).attr("dy",t===Um?"0em":t===Om?"0.71em":"0.32em")),e!==g&&(m=m.transition(e),x=x.transition(e),M=M.transition(e),T=T.transition(e),b=b.transition(e).attr("opacity",Im).attr("transform",function(t){return p(y,this.parentNode.__axis||y,t)}),w.attr("opacity",Im).attr("transform",function(t){return p(this.parentNode.__axis||y,y,t)})),b.remove(),m.attr("d",t===Fm||t==Dm?"M"+k*a+","+v+"H0.5V"+_+"H"+k*a:"M"+v+","+k*a+"V0.5H"+_+"V"+k*a),x.attr("opacity",1).attr("transform",function(t){return p(y,y,t)}),M.attr(s+"2",k*u),T.attr(s,k*h).text(l),g.filter(Vc).attr("fill","none").attr("font-size",10).attr("font-family","sans-serif").attr("text-anchor",t===Dm?"start":t===Fm?"end":"middle"),g.each(function(){this.__axis=y})}var r=[],i=null,o=null,u=6,a=6,c=3;return e.scale=function(t){return arguments.length?(n=t,e):n},e.ticks=function(){return r=Rm.call(arguments),e},e.tickArguments=function(t){return arguments.length?(r=null==t?[]:Rm.call(t),e):r.slice()},e.tickValues=function(t){return arguments.length?(i=null==t?null:Rm.call(t),e):i&&i.slice()},e.tickFormat=function(t){return arguments.length?(o=t,e):o},e.tickSize=function(t){return arguments.length?(u=a=+t,e):u},e.tickSizeInner=function(t){return arguments.length?(u=+t,e):u},e.tickSizeOuter=function(t){return arguments.length?(a=+t,e):a},e.tickPadding=function(t){return arguments.length?(c=+t,e):c},e}function $c(t){return Wc(Um,t)}function Zc(t){return Wc(Dm,t)}function Gc(t){return Wc(Om,t)}function Jc(t){return Wc(Fm,t)}function Qc(t,n){return t.parent===n.parent?1:2}function Kc(t){return t.reduce(ts,0)/t.length}function ts(t,n){return t+n.x}function ns(t){return 1+t.reduce(es,0)}function es(t,n){return Math.max(t,n.y)}function rs(t){for(var n;n=t.children;)t=n[0];return t}function is(t){for(var n;n=t.children;)t=n[n.length-1];return t}function os(){function t(t){var o,u=0;t.eachAfter(function(t){var e=t.children;e?(t.x=Kc(e),t.y=ns(e)):(t.x=o?u+=n(t,o):0,t.y=0,o=t)});var a=rs(t),c=is(t),s=a.x-n(a,c)/2,f=c.x+n(c,a)/2;return t.eachAfter(i?function(n){n.x=(n.x-t.x)*e,n.y=(t.y-n.y)*r}:function(n){n.x=(n.x-s)/(f-s)*e,n.y=(1-(t.y?n.y/t.y:1))*r})}var n=Qc,e=1,r=1,i=!1;return t.separation=function(e){return arguments.length?(n=e,t):n},t.size=function(n){return arguments.length?(i=!1,e=+n[0],r=+n[1],t):i?null:[e,r]},t.nodeSize=function(n){return arguments.length?(i=!0,e=+n[0],r=+n[1],t):i?[e,r]:null},t}function us(t){var n,e,r,i,o=this,u=[o];do for(n=u.reverse(),u=[];o=n.pop();)if(t(o),e=o.children)for(r=0,i=e.length;r<i;++r)u.push(e[r]);while(u.length);return this}function as(t){for(var n,e,r=this,i=[r];r=i.pop();)if(t(r),n=r.children)for(e=n.length-1;e>=0;--e)i.push(n[e]);return this}function cs(t){for(var n,e,r,i=this,o=[i],u=[];i=o.pop();)if(u.push(i),n=i.children)for(e=0,r=n.length;e<r;++e)o.push(n[e]);for(;i=u.pop();)t(i);return this}function ss(t){return this.eachAfter(function(n){for(var e=+t(n.data)||0,r=n.children,i=r&&r.length;--i>=0;)e+=r[i].value;n.value=e})}function fs(t){return this.eachBefore(function(n){n.children&&n.children.sort(t)})}function ls(t){for(var n=this,e=hs(n,t),r=[n];n!==e;)n=n.parent,r.push(n);for(var i=r.length;t!==e;)r.splice(i,0,t),t=t.parent;return r}function hs(t,n){if(t===n)return t;var e=t.ancestors(),r=n.ancestors(),i=null;for(t=e.pop(),n=r.pop();t===n;)i=t,t=e.pop(),n=r.pop();return i}function ps(){for(var t=this,n=[t];t=t.parent;)n.push(t);return n}function ds(){var t=[];return this.each(function(n){t.push(n)}),t}function vs(){var t=[];return this.eachBefore(function(n){n.children||t.push(n)}),t}function _s(){var t=this,n=[];return t.each(function(e){e!==t&&n.push({source:e.parent,target:e})}),n}function ys(t,n){var e,r,i,o,u,a=new ws(t),c=+t.value&&(a.value=t.value),s=[a];for(null==n&&(n=ms);e=s.pop();)if(c&&(e.value=+e.data.value),(i=n(e.data))&&(u=i.length))for(e.children=new Array(u),o=u-1;o>=0;--o)s.push(r=e.children[o]=new ws(i[o])),r.parent=e,r.depth=e.depth+1;return a.eachBefore(bs)}function gs(){return ys(this).eachBefore(xs)}function ms(t){return t.children}function xs(t){t.data=t.data.data}function bs(t){var n=0;do t.height=n;while((t=t.parent)&&t.height<++n)}function ws(t){this.data=t,this.depth=this.height=0,this.parent=null}function Ms(t){this._=t,this.next=null}function Ts(t){for(var n,e=(t=t.slice()).length,r=null,i=r;e;){var o=new Ms(t[e-1]);i=i?i.next=o:r=o,t[n]=t[--e]}return{head:r,tail:i}}function ks(t){return Ns(Ts(t),[])}function Ss(t,n){var e=n.x-t.x,r=n.y-t.y,i=t.r-n.r;return i*i+1e-6>e*e+r*r}function Ns(t,n){var e,r,i,o=null,u=t.head;switch(n.length){case 1:e=As(n[0]);break;case 2:e=Es(n[0],n[1]);break;case 3:e=Cs(n[0],n[1],n[2])}for(;u;)i=u._,r=u.next,e&&Ss(e,i)?o=u:(o?(t.tail=o,o.next=null):t.head=t.tail=null,n.push(i),e=Ns(t,n),n.pop(),t.head?(u.next=t.head,t.head=u):(u.next=null,t.head=t.tail=u),o=t.tail,o.next=r),u=r;return t.tail=o,e}function As(t){return{x:t.x,y:t.y,r:t.r}}function Es(t,n){var e=t.x,r=t.y,i=t.r,o=n.x,u=n.y,a=n.r,c=o-e,s=u-r,f=a-i,l=Math.sqrt(c*c+s*s);return{x:(e+o+c/l*f)/2,y:(r+u+s/l*f)/2,r:(l+i+a)/2}}function Cs(t,n,e){var r=t.x,i=t.y,o=t.r,u=n.x,a=n.y,c=n.r,s=e.x,f=e.y,l=e.r,h=2*(r-u),p=2*(i-a),d=2*(c-o),v=r*r+i*i-o*o-u*u-a*a+c*c,_=2*(r-s),y=2*(i-f),g=2*(l-o),m=r*r+i*i-o*o-s*s-f*f+l*l,x=_*p-h*y,b=(p*m-y*v)/x-r,w=(y*d-p*g)/x,M=(_*v-h*m)/x-i,T=(h*g-_*d)/x,k=w*w+T*T-1,S=2*(b*w+M*T+o),N=b*b+M*M-o*o,A=(-S-Math.sqrt(S*S-4*k*N))/(2*k);return{x:b+w*A+r,y:M+T*A+i,r:A}}function zs(t,n,e){var r=t.x,i=t.y,o=n.r+e.r,u=t.r+e.r,a=n.x-r,c=n.y-i,s=a*a+c*c;if(s){var f=.5+((u*=u)-(o*=o))/(2*s),l=Math.sqrt(Math.max(0,2*o*(u+s)-(u-=s)*u-o*o))/(2*s);e.x=r+f*a+l*c,e.y=i+f*c-l*a}else e.x=r+u,e.y=i}function Ps(t,n){var e=n.x-t.x,r=n.y-t.y,i=t.r+n.r;return i*i>e*e+r*r}function qs(t,n,e){var r=t.x-n,i=t.y-e;return r*r+i*i}function Ls(t){this._=t,this.next=null,this.previous=null}function Rs(t){if(!(i=t.length))return 0;var n,e,r,i;if(n=t[0],n.x=0,n.y=0,!(i>1))return n.r;if(e=t[1],n.x=-e.r,e.x=n.r,e.y=0,!(i>2))return n.r+e.r;zs(e,n,r=t[2]);var o,u,a,c,s,f,l,h=n.r*n.r,p=e.r*e.r,d=r.r*r.r,v=h+p+d,_=h*n.x+p*e.x+d*r.x,y=h*n.y+p*e.y+d*r.y;n=new Ls(n),e=new Ls(e),r=new Ls(r),n.next=r.previous=e,e.next=n.previous=r,r.next=e.previous=n;t:for(a=3;a<i;++a){if(zs(n._,e._,r=t[a]),r=new Ls(r),(s=n.previous)===(c=e.next)){if(Ps(c._,r._)){n=e,e=c,--a;continue t}}else{f=c._.r,l=s._.r;do if(f<=l){if(Ps(c._,r._)){e=c,n.next=e,e.previous=n,--a;continue t}c=c.next,f+=c._.r}else{if(Ps(s._,r._)){n=s,n.next=e,e.previous=n,--a;continue t}s=s.previous,l+=s._.r}while(c!==s.next)}for(r.previous=n,r.next=e,n.next=e.previous=e=r,v+=d=r._.r*r._.r,_+=d*r._.x,y+=d*r._.y,h=qs(n._,o=_/v,u=y/v);(r=r.next)!==e;)(d=qs(r._,o,u))<h&&(n=r,h=d);e=n.next}for(n=[e._],r=e;(r=r.next)!==e;)n.push(r._);for(r=ks(n),a=0;a<i;++a)n=t[a],n.x-=r.x,n.y-=r.y;return r.r}function Us(t){return Rs(t),t}function Ds(t){return null==t?null:Os(t)}function Os(t){if("function"!=typeof t)throw new Error;return t}function Fs(){return 0}function Is(t){return function(){return t}}function Ys(t){return Math.sqrt(t.value)}function Bs(){function t(t){return t.x=e/2,t.y=r/2,n?t.eachBefore(js(n)).eachAfter(Hs(i,.5)).eachBefore(Xs(1)):t.eachBefore(js(Ys)).eachAfter(Hs(Fs,1)).eachAfter(Hs(i,t.r/Math.min(e,r))).eachBefore(Xs(Math.min(e,r)/(2*t.r))),t}var n=null,e=1,r=1,i=Fs;return t.radius=function(e){return arguments.length?(n=Ds(e),t):n},t.size=function(n){return arguments.length?(e=+n[0],r=+n[1],t):[e,r]},t.padding=function(n){return arguments.length?(i="function"==typeof n?n:Is(+n),t):i},t}function js(t){return function(n){n.children||(n.r=Math.max(0,+t(n)||0))}}function Hs(t,n){return function(e){if(r=e.children){var r,i,o,u=r.length,a=t(e)*n||0;if(a)for(i=0;i<u;++i)r[i].r+=a;if(o=Rs(r),a)for(i=0;i<u;++i)r[i].r-=a;e.r=o+a}}}function Xs(t){return function(n){var e=n.parent;n.r*=t,e&&(n.x=e.x+t*n.x,n.y=e.y+t*n.y)}}function Vs(t){t.x0=Math.round(t.x0),t.y0=Math.round(t.y0),t.x1=Math.round(t.x1),t.y1=Math.round(t.y1)}function Ws(t,n,e,r,i){for(var o,u=t.children,a=-1,c=u.length,s=t.value&&(r-n)/t.value;++a<c;)o=u[a],o.y0=e,o.y1=i,o.x0=n,o.x1=n+=o.value*s}function $s(){function t(t){var u=t.height+1;return t.x0=t.y0=i,t.x1=e,t.y1=r/u,t.eachBefore(n(r,u)),o&&t.eachBefore(Vs),t}function n(t,n){return function(e){e.children&&Ws(e,e.x0,t*(e.depth+1)/n,e.x1,t*(e.depth+2)/n);var r=e.x0,o=e.y0,u=e.x1-i,a=e.y1-i;u<r&&(r=u=(r+u)/2),a<o&&(o=a=(o+a)/2),e.x0=r,e.y0=o,e.x1=u,e.y1=a}}var e=1,r=1,i=0,o=!1;return t.round=function(n){return arguments.length?(o=!!n,t):o},t.size=function(n){return arguments.length?(e=+n[0],r=+n[1],t):[e,r]},t.padding=function(n){return arguments.length?(i=+n,t):i},t}function Zs(t){return t.id}function Gs(t){return t.parentId}function Js(){function t(t){var r,i,o,u,a,c,s,f=t.length,l=new Array(f),h={};for(i=0;i<f;++i)r=t[i],a=l[i]=new ws(r),null!=(c=n(r,i,t))&&(c+="")&&(s=Ym+(a.id=c),h[s]=s in h?jm:a);for(i=0;i<f;++i)if(a=l[i],c=e(t[i],i,t),null!=c&&(c+="")){if(u=h[Ym+c],!u)throw new Error("missing: "+c);if(u===jm)throw new Error("ambiguous: "+c);u.children?u.children.push(a):u.children=[a],a.parent=u}else{if(o)throw new Error("multiple roots");o=a}if(!o)throw new Error("no root");if(o.parent=Bm,o.eachBefore(function(t){t.depth=t.parent.depth+1,--f}).eachBefore(bs),o.parent=null,f>0)throw new Error("cycle");return o}var n=Zs,e=Gs;return t.id=function(e){return arguments.length?(n=Os(e),t):n},t.parentId=function(n){return arguments.length?(e=Os(n),t):e},t}function Qs(t,n){return t.parent===n.parent?1:2}function Ks(t){var n=t.children;return n?n[0]:t.t}function tf(t){var n=t.children;return n?n[n.length-1]:t.t}function nf(t,n,e){var r=e/(n.i-t.i);n.c-=r,n.s+=e,t.c+=r,n.z+=e,n.m+=e}function ef(t){for(var n,e=0,r=0,i=t.children,o=i.length;--o>=0;)n=i[o],n.z+=e,n.m+=e,e+=n.s+(r+=n.c)}function rf(t,n,e){return t.a.parent===n.parent?t.a:e}function of(t,n){this._=t,this.parent=null,this.children=null,this.A=null,this.a=this,this.z=0,this.m=0,this.c=0,this.s=0,this.t=null,this.i=n}function uf(t){for(var n,e,r,i,o,u=new of(t,0),a=[u];n=a.pop();)if(r=n._.children)for(n.children=new Array(o=r.length),i=o-1;i>=0;--i)a.push(e=n.children[i]=new of(r[i],i)),e.parent=n;return(u.parent=new of(null,0)).children=[u],u}function af(){function t(t){var r=uf(t);if(r.eachAfter(n),r.parent.m=-r.z,r.eachBefore(e),c)t.eachBefore(i);else{var s=t,f=t,l=t;t.eachBefore(function(t){t.x<s.x&&(s=t),t.x>f.x&&(f=t),t.depth>l.depth&&(l=t)});var h=s===f?1:o(s,f)/2,p=h-s.x,d=u/(f.x+h+p),v=a/(l.depth||1);t.eachBefore(function(t){t.x=(t.x+p)*d,t.y=t.depth*v})}return t}function n(t){var n=t.children,e=t.parent.children,i=t.i?e[t.i-1]:null;if(n){ef(t);var u=(n[0].z+n[n.length-1].z)/2;i?(t.z=i.z+o(t._,i._),t.m=t.z-u):t.z=u}else i&&(t.z=i.z+o(t._,i._));t.parent.A=r(t,i,t.parent.A||e[0])}function e(t){t._.x=t.z+t.parent.m,t.m+=t.parent.m}function r(t,n,e){if(n){for(var r,i=t,u=t,a=n,c=i.parent.children[0],s=i.m,f=u.m,l=a.m,h=c.m;a=tf(a),i=Ks(i),a&&i;)c=Ks(c),u=tf(u),u.a=t,r=a.z+l-i.z-s+o(a._,i._),r>0&&(nf(rf(a,t,e),t,r),s+=r,f+=r),l+=a.m,s+=i.m,h+=c.m,f+=u.m;a&&!tf(u)&&(u.t=a,u.m+=l-f),i&&!Ks(c)&&(c.t=i,c.m+=s-h,e=t)}return e}function i(t){t.x*=u,t.y=t.depth*a}var o=Qs,u=1,a=1,c=null;return t.separation=function(n){return arguments.length?(o=n,t):o},t.size=function(n){return arguments.length?(c=!1,u=+n[0],a=+n[1],t):c?null:[u,a]},t.nodeSize=function(n){return arguments.length?(c=!0,u=+n[0],a=+n[1],t):c?[u,a]:null},t}function cf(t,n,e,r,i){for(var o,u=t.children,a=-1,c=u.length,s=t.value&&(i-e)/t.value;++a<c;)o=u[a],o.x0=n,o.x1=r,o.y0=e,o.y1=e+=o.value*s}function sf(t,n,e,r,i,o){for(var u,a,c,s,f,l,h,p,d,v,_,y,g=[],m=n.children,x=0,b=m.length,w=n.value;x<b;){for(s=i-e,f=o-r,h=p=l=m[x].value,_=Math.max(f/s,s/f)/(w*t),y=l*l*_,v=Math.max(p/y,y/h),c=x+1;c<b;++c){if(l+=a=m[c].value,a<h&&(h=a),a>p&&(p=a),y=l*l*_,d=Math.max(p/y,y/h),d>v){l-=a;break}v=d}g.push(u={value:l,dice:s<f,children:m.slice(x,c)}),u.dice?Ws(u,e,r,i,w?r+=f*l/w:o):cf(u,e,r,w?e+=s*l/w:i,o),w-=l,x=c}return g}function ff(){function t(t){return t.x0=t.y0=0,t.x1=i,t.y1=o,t.eachBefore(n),u=[0],r&&t.eachBefore(Vs),t}function n(t){
var n=u[t.depth],r=t.x0+n,i=t.y0+n,o=t.x1-n,h=t.y1-n;o<r&&(r=o=(r+o)/2),h<i&&(i=h=(i+h)/2),t.x0=r,t.y0=i,t.x1=o,t.y1=h,t.children&&(n=u[t.depth+1]=a(t)/2,r+=l(t)-n,i+=c(t)-n,o-=s(t)-n,h-=f(t)-n,o<r&&(r=o=(r+o)/2),h<i&&(i=h=(i+h)/2),e(t,r,i,o,h))}var e=Xm,r=!1,i=1,o=1,u=[0],a=Fs,c=Fs,s=Fs,f=Fs,l=Fs;return t.round=function(n){return arguments.length?(r=!!n,t):r},t.size=function(n){return arguments.length?(i=+n[0],o=+n[1],t):[i,o]},t.tile=function(n){return arguments.length?(e=Os(n),t):e},t.padding=function(n){return arguments.length?t.paddingInner(n).paddingOuter(n):t.paddingInner()},t.paddingInner=function(n){return arguments.length?(a="function"==typeof n?n:Is(+n),t):a},t.paddingOuter=function(n){return arguments.length?t.paddingTop(n).paddingRight(n).paddingBottom(n).paddingLeft(n):t.paddingTop()},t.paddingTop=function(n){return arguments.length?(c="function"==typeof n?n:Is(+n),t):c},t.paddingRight=function(n){return arguments.length?(s="function"==typeof n?n:Is(+n),t):s},t.paddingBottom=function(n){return arguments.length?(f="function"==typeof n?n:Is(+n),t):f},t.paddingLeft=function(n){return arguments.length?(l="function"==typeof n?n:Is(+n),t):l},t}function lf(t,n,e,r,i){function o(t,n,e,r,i,u,a){if(t>=n-1){var s=c[t];return s.x0=r,s.y0=i,s.x1=u,s.y1=a,void 0}for(var l=f[t],h=e/2+l,p=t+1,d=n-1;p<d;){var v=p+d>>>1;f[v]<h?p=v+1:d=v}var _=f[p]-l,y=e-_;if(a-i>u-r){var g=(i*y+a*_)/e;o(t,p,_,r,i,u,g),o(p,n,y,r,g,u,a)}else{var m=(r*y+u*_)/e;o(t,p,_,r,i,m,a),o(p,n,y,m,i,u,a)}}var u,a,c=t.children,s=c.length,f=new Array(s+1);for(f[0]=a=u=0;u<s;++u)f[u+1]=a+=c[u].value;o(0,s,t.value,n,e,r,i)}function hf(t,n,e,r,i){(1&t.depth?cf:Ws)(t,n,e,r,i)}function pf(t,n){function e(){var e,i,o=r.length,u=0,a=0;for(e=0;e<o;++e)i=r[e],u+=i.x,a+=i.y;for(u=u/o-t,a=a/o-n,e=0;e<o;++e)i=r[e],i.x-=u,i.y-=a}var r;return null==t&&(t=0),null==n&&(n=0),e.initialize=function(t){r=t},e.x=function(n){return arguments.length?(t=+n,e):t},e.y=function(t){return arguments.length?(n=+t,e):n},e}function df(t){return function(){return t}}function vf(){return 1e-6*(Math.random()-.5)}function _f(t){return t.x+t.vx}function yf(t){return t.y+t.vy}function gf(t){function n(){function t(t,e,r,i,u){var a=t.data,p=t.r,d=l+p;{if(!a)return e>s+d||i<s-d||r>f+d||u<f-d;if(a.index>n){var v=s-a.x-a.vx,_=f-a.y-a.vy,y=v*v+_*_;y<d*d&&(0===v&&(v=vf(),y+=v*v),0===_&&(_=vf(),y+=_*_),y=(d-(y=Math.sqrt(y)))/y*o,c.vx+=(v*=y)*(d=(p*=p)/(h+p)),c.vy+=(_*=y)*d,a.vx-=v*(d=1-d),a.vy-=_*d)}}}for(var n,a,c,s,f,l,h,p=r.length,d=0;d<u;++d)for(a=jt(r,_f,yf).visitAfter(e),n=0;n<p;++n)c=r[n],l=i[n],h=l*l,s=c.x+c.vx,f=c.y+c.vy,a.visit(t)}function e(t){if(t.data)return t.r=i[t.data.index];for(var n=t.r=0;n<4;++n)t[n]&&t[n].r>t.r&&(t.r=t[n].r)}var r,i,o=1,u=1;return"function"!=typeof t&&(t=df(null==t?1:+t)),n.initialize=function(n){var e,o=(r=n).length;for(i=new Array(o),e=0;e<o;++e)i[e]=+t(r[e],e,r)},n.iterations=function(t){return arguments.length?(u=+t,n):u},n.strength=function(t){return arguments.length?(o=+t,n):o},n.radius=function(e){return arguments.length?(t="function"==typeof e?e:df(+e),n):t},n}function mf(t,n){return n}function xf(t){function n(t){return 1/Math.min(s[t.source.index],s[t.target.index])}function e(n){for(var e=0,r=t.length;e<d;++e)for(var i,o,c,s,l,h,p,v=0;v<r;++v)i=t[v],o=i.source,c=i.target,s=c.x+c.vx-o.x-o.vx||vf(),l=c.y+c.vy-o.y-o.vy||vf(),h=Math.sqrt(s*s+l*l),h=(h-a[v])/h*n*u[v],s*=h,l*=h,c.vx-=s*(p=f[v]),c.vy-=l*p,o.vx+=s*(p=1-p),o.vy+=l*p}function r(){if(c){var n,e,r=c.length,h=t.length,p=q(c,l);for(n=0,s=new Array(r);n<r;++n)s[n]=0;for(n=0;n<h;++n)e=t[n],e.index=n,"object"!=typeof e.source&&(e.source=p.get(e.source)),"object"!=typeof e.target&&(e.target=p.get(e.target)),++s[e.source.index],++s[e.target.index];for(n=0,f=new Array(h);n<h;++n)e=t[n],f[n]=s[e.source.index]/(s[e.source.index]+s[e.target.index]);u=new Array(h),i(),a=new Array(h),o()}}function i(){if(c)for(var n=0,e=t.length;n<e;++n)u[n]=+h(t[n],n,t)}function o(){if(c)for(var n=0,e=t.length;n<e;++n)a[n]=+p(t[n],n,t)}var u,a,c,s,f,l=mf,h=n,p=df(30),d=1;return null==t&&(t=[]),e.initialize=function(t){c=t,r()},e.links=function(n){return arguments.length?(t=n,r(),e):t},e.id=function(t){return arguments.length?(l=t,e):l},e.iterations=function(t){return arguments.length?(d=+t,e):d},e.strength=function(t){return arguments.length?(h="function"==typeof t?t:df(+t),i(),e):h},e.distance=function(t){return arguments.length?(p="function"==typeof t?t:df(+t),o(),e):p},e}function bf(t){return t.x}function wf(t){return t.y}function Mf(t){function n(){e(),p.call("tick",o),u<a&&(h.stop(),p.call("end",o))}function e(){var n,e,r=t.length;for(u+=(s-u)*c,l.each(function(t){t(u)}),n=0;n<r;++n)e=t[n],null==e.fx?e.x+=e.vx*=f:(e.x=e.fx,e.vx=0),null==e.fy?e.y+=e.vy*=f:(e.y=e.fy,e.vy=0)}function r(){for(var n,e=0,r=t.length;e<r;++e){if(n=t[e],n.index=e,isNaN(n.x)||isNaN(n.y)){var i=Wm*Math.sqrt(e),o=e*$m;n.x=i*Math.cos(o),n.y=i*Math.sin(o)}(isNaN(n.vx)||isNaN(n.vy))&&(n.vx=n.vy=0)}}function i(n){return n.initialize&&n.initialize(t),n}var o,u=1,a=.001,c=1-Math.pow(a,1/300),s=0,f=.6,l=q(),h=Yr(n),p=Mr("tick","end");return null==t&&(t=[]),r(),o={tick:e,restart:function(){return h.restart(n),o},stop:function(){return h.stop(),o},nodes:function(n){return arguments.length?(t=n,r(),l.each(i),o):t},alpha:function(t){return arguments.length?(u=+t,o):u},alphaMin:function(t){return arguments.length?(a=+t,o):a},alphaDecay:function(t){return arguments.length?(c=+t,o):+c},alphaTarget:function(t){return arguments.length?(s=+t,o):s},velocityDecay:function(t){return arguments.length?(f=1-t,o):1-f},force:function(t,n){return arguments.length>1?(null==n?l.remove(t):l.set(t,i(n)),o):l.get(t)},find:function(n,e,r){var i,o,u,a,c,s=0,f=t.length;for(null==r?r=1/0:r*=r,s=0;s<f;++s)a=t[s],i=n-a.x,o=e-a.y,u=i*i+o*o,u<r&&(c=a,r=u);return c},on:function(t,n){return arguments.length>1?(p.on(t,n),o):p.on(t)}}}function Tf(){function t(t){var n,a=i.length,c=jt(i,bf,wf).visitAfter(e);for(u=t,n=0;n<a;++n)o=i[n],c.visit(r)}function n(){if(i){var t,n=i.length;for(a=new Array(n),t=0;t<n;++t)a[t]=+c(i[t],t,i)}}function e(t){var n,e,r,i,o,u=0;if(t.length){for(r=i=o=0;o<4;++o)(n=t[o])&&(e=n.value)&&(u+=e,r+=e*n.x,i+=e*n.y);t.x=r/u,t.y=i/u}else{n=t,n.x=n.data.x,n.y=n.data.y;do u+=a[n.data.index];while(n=n.next)}t.value=u}function r(t,n,e,r){if(!t.value)return!0;var i=t.x-o.x,c=t.y-o.y,h=r-n,p=i*i+c*c;if(h*h/l<p)return p<f&&(0===i&&(i=vf(),p+=i*i),0===c&&(c=vf(),p+=c*c),p<s&&(p=Math.sqrt(s*p)),o.vx+=i*t.value*u/p,o.vy+=c*t.value*u/p),!0;if(!(t.length||p>=f)){(t.data!==o||t.next)&&(0===i&&(i=vf(),p+=i*i),0===c&&(c=vf(),p+=c*c),p<s&&(p=Math.sqrt(s*p)));do t.data!==o&&(h=a[t.data.index]*u/p,o.vx+=i*h,o.vy+=c*h);while(t=t.next)}}var i,o,u,a,c=df(-30),s=1,f=1/0,l=.81;return t.initialize=function(t){i=t,n()},t.strength=function(e){return arguments.length?(c="function"==typeof e?e:df(+e),n(),t):c},t.distanceMin=function(n){return arguments.length?(s=n*n,t):Math.sqrt(s)},t.distanceMax=function(n){return arguments.length?(f=n*n,t):Math.sqrt(f)},t.theta=function(n){return arguments.length?(l=n*n,t):Math.sqrt(l)},t}function kf(t){function n(t){for(var n,e=0,u=r.length;e<u;++e)n=r[e],n.vx+=(o[e]-n.x)*i[e]*t}function e(){if(r){var n,e=r.length;for(i=new Array(e),o=new Array(e),n=0;n<e;++n)i[n]=isNaN(o[n]=+t(r[n],n,r))?0:+u(r[n],n,r)}}var r,i,o,u=df(.1);return"function"!=typeof t&&(t=df(null==t?0:+t)),n.initialize=function(t){r=t,e()},n.strength=function(t){return arguments.length?(u="function"==typeof t?t:df(+t),e(),n):u},n.x=function(r){return arguments.length?(t="function"==typeof r?r:df(+r),e(),n):t},n}function Sf(t){function n(t){for(var n,e=0,u=r.length;e<u;++e)n=r[e],n.vy+=(o[e]-n.y)*i[e]*t}function e(){if(r){var n,e=r.length;for(i=new Array(e),o=new Array(e),n=0;n<e;++n)i[n]=isNaN(o[n]=+t(r[n],n,r))?0:+u(r[n],n,r)}}var r,i,o,u=df(.1);return"function"!=typeof t&&(t=df(null==t?0:+t)),n.initialize=function(t){r=t,e()},n.strength=function(t){return arguments.length?(u="function"==typeof t?t:df(+t),e(),n):u},n.y=function(r){return arguments.length?(t="function"==typeof r?r:df(+r),e(),n):t},n}function Nf(){t.event.stopImmediatePropagation()}function Af(){t.event.preventDefault(),t.event.stopImmediatePropagation()}function Ef(t){var n=t.document.documentElement,e=Ra(t).on("dragstart.drag",Af,!0);"onselectstart"in n?e.on("selectstart.drag",Af,!0):(n.__noselect=n.style.MozUserSelect,n.style.MozUserSelect="none")}function Cf(t,n){var e=t.document.documentElement,r=Ra(t).on("dragstart.drag",null);n&&(r.on("click.drag",Af,!0),setTimeout(function(){r.on("click.drag",null)},0)),"onselectstart"in e?r.on("selectstart.drag",null):(e.style.MozUserSelect=e.__noselect,delete e.__noselect)}function zf(t){return function(){return t}}function Pf(t,n,e,r,i,o,u,a,c,s){this.target=t,this.type=n,this.subject=e,this.identifier=r,this.active=i,this.x=o,this.y=u,this.dx=a,this.dy=c,this._=s}function qf(){return!t.event.button}function Lf(){return this.parentNode}function Rf(n){return null==n?{x:t.event.x,y:t.event.y}:n}function Uf(){function n(t){t.on("mousedown.drag",e).on("touchstart.drag",o).on("touchmove.drag",u).on("touchend.drag touchcancel.drag",a).style("-webkit-tap-highlight-color","rgba(0,0,0,0)")}function e(){if(!f&&l.apply(this,arguments)){var n=c("mouse",h.apply(this,arguments),du,this,arguments);n&&(Ra(t.event.view).on("mousemove.drag",r,!0).on("mouseup.drag",i,!0),Ef(t.event.view),Nf(),s=!1,n("start"))}}function r(){Af(),s=!0,d.mouse("drag")}function i(){Ra(t.event.view).on("mousemove.drag mouseup.drag",null),Cf(t.event.view,s),Af(),d.mouse("end")}function o(){if(l.apply(this,arguments)){var n,e,r=t.event.changedTouches,i=h.apply(this,arguments),o=r.length;for(n=0;n<o;++n)(e=c(r[n].identifier,i,Da,this,arguments))&&(Nf(),e("start"))}}function u(){var n,e,r=t.event.changedTouches,i=r.length;for(n=0;n<i;++n)(e=d[r[n].identifier])&&(Af(),e("drag"))}function a(){var n,e,r=t.event.changedTouches,i=r.length;for(f&&clearTimeout(f),f=setTimeout(function(){f=null},500),n=0;n<i;++n)(e=d[r[n].identifier])&&(Nf(),e("end"))}function c(e,r,i,o,u){var a,c,s,f=i(r,e),l=v.copy();if(lu(new Pf(n,"beforestart",a,e,_,f[0],f[1],0,0,l),function(){return null!=(t.event.subject=a=p.apply(o,u))&&(c=a.x-f[0]||0,s=a.y-f[1]||0,!0)}))return function t(h){var p,v=f;switch(h){case"start":d[e]=t,p=_++;break;case"end":delete d[e],--_;case"drag":f=i(r,e),p=_}lu(new Pf(n,h,a,e,p,f[0]+c,f[1]+s,f[0]-v[0],f[1]-v[1],l),l.apply,l,[h,o,u])}}var s,f,l=qf,h=Lf,p=Rf,d={},v=Mr("start","drag","end"),_=0;return n.filter=function(t){return arguments.length?(l="function"==typeof t?t:zf(!!t),n):l},n.container=function(t){return arguments.length?(h="function"==typeof t?t:zf(t),n):h},n.subject=function(t){return arguments.length?(p="function"==typeof t?t:zf(t),n):p},n.on=function(){var t=v.on.apply(v,arguments);return t===v?n:t},n}function Df(t){return function(){return t}}function Of(t){return t[0]}function Ff(t){return t[1]}function If(){this._=null}function Yf(t){t.U=t.C=t.L=t.R=t.P=t.N=null}function Bf(t,n){var e=n,r=n.R,i=e.U;i?i.L===e?i.L=r:i.R=r:t._=r,r.U=i,e.U=r,e.R=r.L,e.R&&(e.R.U=e),r.L=e}function jf(t,n){var e=n,r=n.L,i=e.U;i?i.L===e?i.L=r:i.R=r:t._=r,r.U=i,e.U=r,e.L=r.R,e.L&&(e.L.U=e),r.R=e}function Hf(t){for(;t.L;)t=t.L;return t}function Xf(t,n,e,r){var i=[null,null],o=Km.push(i)-1;return i.left=t,i.right=n,e&&Wf(i,t,n,e),r&&Wf(i,n,t,r),Jm[t.index].halfedges.push(o),Jm[n.index].halfedges.push(o),i}function Vf(t,n,e){var r=[n,e];return r.left=t,r}function Wf(t,n,e,r){t[0]||t[1]?t.left===e?t[1]=r:t[0]=r:(t[0]=r,t.left=n,t.right=e)}function $f(t,n,e,r,i){var o,u=t[0],a=t[1],c=u[0],s=u[1],f=a[0],l=a[1],h=0,p=1,d=f-c,v=l-s;if(o=n-c,d||!(o>0)){if(o/=d,d<0){if(o<h)return;o<p&&(p=o)}else if(d>0){if(o>p)return;o>h&&(h=o)}if(o=r-c,d||!(o<0)){if(o/=d,d<0){if(o>p)return;o>h&&(h=o)}else if(d>0){if(o<h)return;o<p&&(p=o)}if(o=e-s,v||!(o>0)){if(o/=v,v<0){if(o<h)return;o<p&&(p=o)}else if(v>0){if(o>p)return;o>h&&(h=o)}if(o=i-s,v||!(o<0)){if(o/=v,v<0){if(o>p)return;o>h&&(h=o)}else if(v>0){if(o<h)return;o<p&&(p=o)}return!(h>0||p<1)||(h>0&&(t[0]=[c+h*d,s+h*v]),p<1&&(t[1]=[c+p*d,s+p*v]),!0)}}}}}function Zf(t,n,e,r,i){var o=t[1];if(o)return!0;var u,a,c=t[0],s=t.left,f=t.right,l=s[0],h=s[1],p=f[0],d=f[1],v=(l+p)/2,_=(h+d)/2;if(d===h){if(v<n||v>=r)return;if(l>p){if(c){if(c[1]>=i)return}else c=[v,e];o=[v,i]}else{if(c){if(c[1]<e)return}else c=[v,i];o=[v,e]}}else if(u=(l-p)/(d-h),a=_-u*v,u<-1||u>1)if(l>p){if(c){if(c[1]>=i)return}else c=[(e-a)/u,e];o=[(i-a)/u,i]}else{if(c){if(c[1]<e)return}else c=[(i-a)/u,i];o=[(e-a)/u,e]}else if(h<d){if(c){if(c[0]>=r)return}else c=[n,u*n+a];o=[r,u*r+a]}else{if(c){if(c[0]<n)return}else c=[r,u*r+a];o=[n,u*n+a]}return t[0]=c,t[1]=o,!0}function Gf(t,n,e,r){for(var i,o=Km.length;o--;)Zf(i=Km[o],t,n,e,r)&&$f(i,t,n,e,r)&&(Math.abs(i[0][0]-i[1][0])>ex||Math.abs(i[0][1]-i[1][1])>ex)||delete Km[o]}function Jf(t){return Jm[t.index]={site:t,halfedges:[]}}function Qf(t,n){var e=t.site,r=n.left,i=n.right;return e===i&&(i=r,r=e),i?Math.atan2(i[1]-r[1],i[0]-r[0]):(e===r?(r=n[1],i=n[0]):(r=n[0],i=n[1]),Math.atan2(r[0]-i[0],i[1]-r[1]))}function Kf(t,n){return n[+(n.left!==t.site)]}function tl(t,n){return n[+(n.left===t.site)]}function nl(){for(var t,n,e,r,i=0,o=Jm.length;i<o;++i)if((t=Jm[i])&&(r=(n=t.halfedges).length)){var u=new Array(r),a=new Array(r);for(e=0;e<r;++e)u[e]=e,a[e]=Qf(t,Km[n[e]]);for(u.sort(function(t,n){return a[n]-a[t]}),e=0;e<r;++e)a[e]=n[u[e]];for(e=0;e<r;++e)n[e]=a[e]}}function el(t,n,e,r){var i,o,u,a,c,s,f,l,h,p,d,v,_=Jm.length,y=!0;for(i=0;i<_;++i)if(o=Jm[i]){for(u=o.site,c=o.halfedges,a=c.length;a--;)Km[c[a]]||c.splice(a,1);for(a=0,s=c.length;a<s;)p=tl(o,Km[c[a]]),d=p[0],v=p[1],f=Kf(o,Km[c[++a%s]]),l=f[0],h=f[1],(Math.abs(d-l)>ex||Math.abs(v-h)>ex)&&(c.splice(a,0,Km.push(Vf(u,p,Math.abs(d-t)<ex&&r-v>ex?[t,Math.abs(l-t)<ex?h:r]:Math.abs(v-r)<ex&&e-d>ex?[Math.abs(h-r)<ex?l:e,r]:Math.abs(d-e)<ex&&v-n>ex?[e,Math.abs(l-e)<ex?h:n]:Math.abs(v-n)<ex&&d-t>ex?[Math.abs(h-n)<ex?l:t,n]:null))-1),++s);s&&(y=!1)}if(y){var g,m,x,b=1/0;for(i=0,y=null;i<_;++i)(o=Jm[i])&&(u=o.site,g=u[0]-t,m=u[1]-n,x=g*g+m*m,x<b&&(b=x,y=o));if(y){var w=[t,n],M=[t,r],T=[e,r],k=[e,n];y.halfedges.push(Km.push(Vf(u=y.site,w,M))-1,Km.push(Vf(u,M,T))-1,Km.push(Vf(u,T,k))-1,Km.push(Vf(u,k,w))-1)}}for(i=0;i<_;++i)(o=Jm[i])&&(o.halfedges.length||delete Jm[i])}function rl(){Yf(this),this.x=this.y=this.arc=this.site=this.cy=null}function il(t){var n=t.P,e=t.N;if(n&&e){var r=n.site,i=t.site,o=e.site;if(r!==o){var u=i[0],a=i[1],c=r[0]-u,s=r[1]-a,f=o[0]-u,l=o[1]-a,h=2*(c*l-s*f);if(!(h>=-rx)){var p=c*c+s*s,d=f*f+l*l,v=(l*p-s*d)/h,_=(c*d-f*p)/h,y=tx.pop()||new rl;y.arc=t,y.site=i,y.x=v+u,y.y=(y.cy=_+a)+Math.sqrt(v*v+_*_),t.circle=y;for(var g=null,m=Qm._;m;)if(y.y<m.y||y.y===m.y&&y.x<=m.x){if(!m.L){g=m.P;break}m=m.L}else{if(!m.R){g=m;break}m=m.R}Qm.insert(g,y),g||(Zm=y)}}}}function ol(t){var n=t.circle;n&&(n.P||(Zm=n.N),Qm.remove(n),tx.push(n),Yf(n),t.circle=null)}function ul(){Yf(this),this.edge=this.site=this.circle=null}function al(t){var n=nx.pop()||new ul;return n.site=t,n}function cl(t){ol(t),Gm.remove(t),nx.push(t),Yf(t)}function sl(t){var n=t.circle,e=n.x,r=n.cy,i=[e,r],o=t.P,u=t.N,a=[t];cl(t);for(var c=o;c.circle&&Math.abs(e-c.circle.x)<ex&&Math.abs(r-c.circle.cy)<ex;)o=c.P,a.unshift(c),cl(c),c=o;a.unshift(c),ol(c);for(var s=u;s.circle&&Math.abs(e-s.circle.x)<ex&&Math.abs(r-s.circle.cy)<ex;)u=s.N,a.push(s),cl(s),s=u;a.push(s),ol(s);var f,l=a.length;for(f=1;f<l;++f)s=a[f],c=a[f-1],Wf(s.edge,c.site,s.site,i);c=a[0],s=a[l-1],s.edge=Xf(c.site,s.site,null,i),il(c),il(s)}function fl(t){for(var n,e,r,i,o=t[0],u=t[1],a=Gm._;a;)if(r=ll(a,u)-o,r>ex)a=a.L;else{if(i=o-hl(a,u),!(i>ex)){r>-ex?(n=a.P,e=a):i>-ex?(n=a,e=a.N):n=e=a;break}if(!a.R){n=a;break}a=a.R}Jf(t);var c=al(t);if(Gm.insert(n,c),n||e){if(n===e)return ol(n),e=al(n.site),Gm.insert(c,e),c.edge=e.edge=Xf(n.site,c.site),il(n),void il(e);if(!e)return void(c.edge=Xf(n.site,c.site));ol(n),ol(e);var s=n.site,f=s[0],l=s[1],h=t[0]-f,p=t[1]-l,d=e.site,v=d[0]-f,_=d[1]-l,y=2*(h*_-p*v),g=h*h+p*p,m=v*v+_*_,x=[(_*g-p*m)/y+f,(h*m-v*g)/y+l];Wf(e.edge,s,d,x),c.edge=Xf(s,t,null,x),e.edge=Xf(t,d,null,x),il(n),il(e)}}function ll(t,n){var e=t.site,r=e[0],i=e[1],o=i-n;if(!o)return r;var u=t.P;if(!u)return-(1/0);e=u.site;var a=e[0],c=e[1],s=c-n;if(!s)return a;var f=a-r,l=1/o-1/s,h=f/s;return l?(-h+Math.sqrt(h*h-2*l*(f*f/(-2*s)-c+s/2+i-o/2)))/l+r:(r+a)/2}function hl(t,n){var e=t.N;if(e)return ll(e,n);var r=t.site;return r[1]===n?r[0]:1/0}function pl(t,n,e){return(t[0]-e[0])*(n[1]-t[1])-(t[0]-n[0])*(e[1]-t[1])}function dl(t,n){return n[1]-t[1]||n[0]-t[0]}function vl(t,n){var e,r,i,o=t.sort(dl).pop();for(Km=[],Jm=new Array(t.length),Gm=new If,Qm=new If;;)if(i=Zm,o&&(!i||o[1]<i.y||o[1]===i.y&&o[0]<i.x))o[0]===e&&o[1]===r||(fl(o),e=o[0],r=o[1]),o=t.pop();else{if(!i)break;sl(i.arc)}if(nl(),n){var u=+n[0][0],a=+n[0][1],c=+n[1][0],s=+n[1][1];Gf(u,a,c,s),el(u,a,c,s)}this.edges=Km,this.cells=Jm,Gm=Qm=Km=Jm=null}function _l(){function t(t){return new vl(t.map(function(r,i){var o=[Math.round(n(r,i,t)/ex)*ex,Math.round(e(r,i,t)/ex)*ex];return o.index=i,o.data=r,o}),r)}var n=Of,e=Ff,r=null;return t.polygons=function(n){return t(n).polygons()},t.links=function(n){return t(n).links()},t.triangles=function(n){return t(n).triangles()},t.x=function(e){return arguments.length?(n="function"==typeof e?e:Df(+e),t):n},t.y=function(n){return arguments.length?(e="function"==typeof n?n:Df(+n),t):e},t.extent=function(n){return arguments.length?(r=null==n?null:[[+n[0][0],+n[0][1]],[+n[1][0],+n[1][1]]],t):r&&[[r[0][0],r[0][1]],[r[1][0],r[1][1]]]},t.size=function(n){return arguments.length?(r=null==n?null:[[0,0],[+n[0],+n[1]]],t):r&&[r[1][0]-r[0][0],r[1][1]-r[0][1]]},t}function yl(t){return function(){return t}}function gl(t,n,e){this.target=t,this.type=n,this.transform=e}function ml(t,n,e){this.k=t,this.x=n,this.y=e}function xl(t){return t.__zoom||ix}function bl(){t.event.stopImmediatePropagation()}function wl(){t.event.preventDefault(),t.event.stopImmediatePropagation()}function Ml(){return!t.event.button}function Tl(){var t,n,e=this;return e instanceof SVGElement?(e=e.ownerSVGElement||e,t=e.width.baseVal.value,n=e.height.baseVal.value):(t=e.clientWidth,n=e.clientHeight),[[0,0],[t,n]]}function kl(){return this.__zoom||ix}function Sl(){function n(t){t.on("wheel.zoom",s).on("mousedown.zoom",f).on("dblclick.zoom",l).on("touchstart.zoom",h).on("touchmove.zoom",p).on("touchend.zoom touchcancel.zoom",d).style("-webkit-tap-highlight-color","rgba(0,0,0,0)").property("__zoom",kl)}function e(t,n){return n=Math.max(m,Math.min(x,n)),n===t.k?t:new ml(n,t.x,t.y)}function r(t,n,e){var r=n[0]-e[0]*t.k,i=n[1]-e[1]*t.k;return r===t.x&&i===t.y?t:new ml(t.k,r,i)}function i(t,n){var e=Math.min(0,t.invertX(n[0][0])-b)||Math.max(0,t.invertX(n[1][0])-w),r=Math.min(0,t.invertY(n[0][1])-M)||Math.max(0,t.invertY(n[1][1])-T);return e||r?t.translate(e,r):t}function o(t){return[(+t[0][0]+ +t[1][0])/2,(+t[0][1]+ +t[1][1])/2]}function u(t,n,e){t.on("start.zoom",function(){a(this,arguments).start()}).on("interrupt.zoom end.zoom",function(){a(this,arguments).end()}).tween("zoom",function(){var t=this,r=arguments,i=a(t,r),u=g.apply(t,r),c=e||o(u),s=Math.max(u[1][0]-u[0][0],u[1][1]-u[0][1]),f=t.__zoom,l="function"==typeof n?n.apply(t,r):n,h=yr(f.invert(c).concat(s/f.k),l.invert(c).concat(s/l.k));return function(t){if(1===t)t=l;else{var n=h(t),e=s/n[2];t=new ml(e,c[0]-n[0]*e,c[1]-n[1]*e)}i.zoom(null,t)}})}function a(t,n){for(var e,r=0,i=S.length;r<i;++r)if((e=S[r]).that===t)return e;return new c(t,n)}function c(t,n){this.that=t,this.args=n,this.index=-1,this.active=0,this.extent=g.apply(t,n)}function s(){function n(){o.wheel=null,o.end()}if(y.apply(this,arguments)){var o=a(this,arguments),u=this.__zoom,c=Math.max(m,Math.min(x,u.k*Math.pow(2,-t.event.deltaY*(t.event.deltaMode?120:1)/500))),s=du(this);if(o.wheel)o.mouse[0][0]===s[0]&&o.mouse[0][1]===s[1]||(o.mouse[1]=u.invert(o.mouse[0]=s)),clearTimeout(o.wheel);else{if(u.k===c)return;o.mouse=[s,u.invert(s)],Ha(this),o.start()}wl(),o.wheel=setTimeout(n,E),o.zoom("mouse",i(r(e(u,c),o.mouse[0],o.mouse[1]),o.extent))}}function f(){function n(){wl(),o.moved=!0,o.zoom("mouse",i(r(o.that.__zoom,o.mouse[0]=du(o.that),o.mouse[1]),o.extent))}function e(){u.on("mousemove.zoom mouseup.zoom",null),Cf(t.event.view,o.moved),wl(),o.end()}if(!_&&y.apply(this,arguments)){var o=a(this,arguments),u=Ra(t.event.view).on("mousemove.zoom",n,!0).on("mouseup.zoom",e,!0),c=du(this);Ef(t.event.view),bl(),o.mouse=[c,this.__zoom.invert(c)],Ha(this),o.start()}}function l(){if(y.apply(this,arguments)){var o=this.__zoom,a=du(this),c=o.invert(a),s=o.k*(t.event.shiftKey?.5:2),f=i(r(e(o,s),a,c),g.apply(this,arguments));wl(),k>0?Ra(this).transition().duration(k).call(u,f,a):Ra(this).call(n.transform,f)}}function h(){if(y.apply(this,arguments)){var n,e,r,i=a(this,arguments),o=t.event.changedTouches,u=o.length;for(bl(),n=0;n<u;++n)e=o[n],r=Da(this,o,e.identifier),r=[r,this.__zoom.invert(r),e.identifier],i.touch0?i.touch1||(i.touch1=r):i.touch0=r;return v&&(v=clearTimeout(v),!i.touch1)?(i.end(),l.apply(this,arguments)):void(t.event.touches.length===u&&(v=setTimeout(function(){v=null},A),Ha(this),i.start()))}}function p(){var n,o,u,c,s=a(this,arguments),f=t.event.changedTouches,l=f.length;for(wl(),v&&(v=clearTimeout(v)),n=0;n<l;++n)o=f[n],u=Da(this,f,o.identifier),s.touch0&&s.touch0[2]===o.identifier?s.touch0[0]=u:s.touch1&&s.touch1[2]===o.identifier&&(s.touch1[0]=u);if(o=s.that.__zoom,s.touch1){var h=s.touch0[0],p=s.touch0[1],d=s.touch1[0],_=s.touch1[1],y=(y=d[0]-h[0])*y+(y=d[1]-h[1])*y,g=(g=_[0]-p[0])*g+(g=_[1]-p[1])*g;o=e(o,Math.sqrt(y/g)),u=[(h[0]+d[0])/2,(h[1]+d[1])/2],c=[(p[0]+_[0])/2,(p[1]+_[1])/2]}else{if(!s.touch0)return;u=s.touch0[0],c=s.touch0[1]}s.zoom("touch",i(r(o,u,c),s.extent))}function d(){var n,e,r=a(this,arguments),i=t.event.changedTouches,o=i.length;for(bl(),_&&clearTimeout(_),_=setTimeout(function(){_=null},A),n=0;n<o;++n)e=i[n],r.touch0&&r.touch0[2]===e.identifier?delete r.touch0:r.touch1&&r.touch1[2]===e.identifier&&delete r.touch1;r.touch1&&!r.touch0&&(r.touch0=r.touch1,delete r.touch1),r.touch0||r.end()}var v,_,y=Ml,g=Tl,m=0,x=1/0,b=-x,w=x,M=b,T=w,k=250,S=[],N=Mr("start","zoom","end"),A=500,E=150;return n.transform=function(t,n){var e=t.selection?t.selection():t;e.property("__zoom",kl),t!==e?u(t,n):e.interrupt().each(function(){a(this,arguments).start().zoom(null,"function"==typeof n?n.apply(this,arguments):n).end()})},n.scaleBy=function(t,e){n.scaleTo(t,function(){var t=this.__zoom.k,n="function"==typeof e?e.apply(this,arguments):e;return t*n})},n.scaleTo=function(t,u){n.transform(t,function(){var t=g.apply(this,arguments),n=this.__zoom,a=o(t),c=n.invert(a),s="function"==typeof u?u.apply(this,arguments):u;return i(r(e(n,s),a,c),t)})},n.translateBy=function(t,e,r){n.transform(t,function(){return i(this.__zoom.translate("function"==typeof e?e.apply(this,arguments):e,"function"==typeof r?r.apply(this,arguments):r),g.apply(this,arguments))})},c.prototype={start:function(){return 1===++this.active&&(this.index=S.push(this)-1,this.emit("start")),this},zoom:function(t,n){return this.mouse&&"mouse"!==t&&(this.mouse[1]=n.invert(this.mouse[0])),this.touch0&&"touch"!==t&&(this.touch0[1]=n.invert(this.touch0[0])),this.touch1&&"touch"!==t&&(this.touch1[1]=n.invert(this.touch1[0])),this.that.__zoom=n,this.emit("zoom"),this},end:function(){return 0===--this.active&&(S.splice(this.index,1),this.index=-1,this.emit("end")),this},emit:function(t){lu(new gl(n,t,this.that.__zoom),N.apply,N,[t,this.that,this.args])}},n.filter=function(t){return arguments.length?(y="function"==typeof t?t:yl(!!t),n):y},n.extent=function(t){return arguments.length?(g="function"==typeof t?t:yl([[+t[0][0],+t[0][1]],[+t[1][0],+t[1][1]]]),n):g},n.scaleExtent=function(t){return arguments.length?(m=+t[0],x=+t[1],n):[m,x]},n.translateExtent=function(t){return arguments.length?(b=+t[0][0],w=+t[1][0],M=+t[0][1],T=+t[1][1],n):[[b,M],[w,T]]},n.duration=function(t){return arguments.length?(k=+t,n):k},n.on=function(){var t=N.on.apply(N,arguments);return t===N?n:t},n}function Nl(t){return function(){return t}}function Al(t,n,e){this.target=t,this.type=n,this.selection=e}function El(){t.event.stopImmediatePropagation()}function Cl(){t.event.preventDefault(),t.event.stopImmediatePropagation()}function zl(t){return{type:t}}function Pl(){return!t.event.button}function ql(){var t=this.ownerSVGElement||this;return[[0,0],[t.width.baseVal.value,t.height.baseVal.value]]}function Ll(t){for(;!t.__brush;)if(!(t=t.parentNode))return;return t.__brush}function Rl(t){return t[0][0]===t[1][0]||t[0][1]===t[1][1]}function Ul(t){var n=t.__brush;return n?n.dim.output(n.selection):null}function Dl(){return Il(sx)}function Ol(){return Il(fx)}function Fl(){return Il(lx)}function Il(n){function e(t){var e=t.property("__brush",a).selectAll(".overlay").data([zl("overlay")]);e.enter().append("rect").attr("class","overlay").attr("pointer-events","all").attr("cursor",hx.overlay).merge(e).each(function(){var t=Ll(this).extent;Ra(this).attr("x",t[0][0]).attr("y",t[0][1]).attr("width",t[1][0]-t[0][0]).attr("height",t[1][1]-t[0][1])}),t.selectAll(".selection").data([zl("selection")]).enter().append("rect").attr("class","selection").attr("cursor",hx.selection).attr("fill","#777").attr("fill-opacity",.3).attr("stroke","#fff").attr("shape-rendering","crispEdges");var i=t.selectAll(".handle").data(n.handles,function(t){return t.type});i.exit().remove(),i.enter().append("rect").attr("class",function(t){return"handle handle--"+t.type}).attr("cursor",function(t){return hx[t.type]}),t.each(r).attr("fill","none").attr("pointer-events","all").style("-webkit-tap-highlight-color","rgba(0,0,0,0)").on("mousedown.brush touchstart.brush",u)}function r(){var t=Ra(this),n=Ll(this).selection;n?(t.selectAll(".selection").style("display",null).attr("x",n[0][0]).attr("y",n[0][1]).attr("width",n[1][0]-n[0][0]).attr("height",n[1][1]-n[0][1]),t.selectAll(".handle").style("display",null).attr("x",function(t){return"e"===t.type[t.type.length-1]?n[1][0]-h/2:n[0][0]-h/2}).attr("y",function(t){return"s"===t.type[0]?n[1][1]-h/2:n[0][1]-h/2}).attr("width",function(t){return"n"===t.type||"s"===t.type?n[1][0]-n[0][0]+h:h}).attr("height",function(t){return"e"===t.type||"w"===t.type?n[1][1]-n[0][1]+h:h})):t.selectAll(".selection,.handle").style("display","none").attr("x",null).attr("y",null).attr("width",null).attr("height",null)}function i(t,n){return t.__brush.emitter||new o(t,n)}function o(t,n){this.that=t,this.args=n,this.state=t.__brush,this.active=0}function u(){function e(){var t=du(T);!U||w||M||(Math.abs(t[0]-O[0])>Math.abs(t[1]-O[1])?M=!0:w=!0),O=t,b=!0,Cl(),o()}function o(){var t;switch(m=O[0]-D[0],x=O[1]-D[1],S){case ux:case ox:N&&(m=Math.max(P-l,Math.min(L-v,m)),h=l+m,_=v+m),A&&(x=Math.max(q-p,Math.min(R-y,x)),d=p+x,g=y+x);break;case ax:N<0?(m=Math.max(P-l,Math.min(L-l,m)),h=l+m,_=v):N>0&&(m=Math.max(P-v,Math.min(L-v,m)),h=l,_=v+m),A<0?(x=Math.max(q-p,Math.min(R-p,x)),d=p+x,g=y):A>0&&(x=Math.max(q-y,Math.min(R-y,x)),d=p,g=y+x);break;case cx:N&&(h=Math.max(P,Math.min(L,l-m*N)),_=Math.max(P,Math.min(L,v+m*N))),A&&(d=Math.max(q,Math.min(R,p-x*A)),g=Math.max(q,Math.min(R,y+x*A)))}_<h&&(N*=-1,t=l,l=v,v=t,t=h,h=_,_=t,k in px&&Y.attr("cursor",hx[k=px[k]])),g<d&&(A*=-1,t=p,p=y,y=t,t=d,d=g,g=t,k in dx&&Y.attr("cursor",hx[k=dx[k]])),z=E.selection,w&&(h=z[0][0],_=z[1][0]),M&&(d=z[0][1],g=z[1][1]),z[0][0]===h&&z[0][1]===d&&z[1][0]===_&&z[1][1]===g||(E.selection=[[h,d],[_,g]],r.call(T),F.brush())}function u(){if(El(),t.event.touches){if(t.event.touches.length)return;c&&clearTimeout(c),c=setTimeout(function(){c=null},500),I.on("touchmove.brush touchend.brush touchcancel.brush",null)}else Cf(t.event.view,b),B.on("keydown.brush keyup.brush mousemove.brush mouseup.brush",null);I.attr("pointer-events","all"),Y.attr("cursor",hx.overlay),Rl(z)&&(E.selection=null,r.call(T)),F.end()}function a(){switch(t.event.keyCode){case 16:U=N&&A;break;case 18:S===ax&&(N&&(v=_-m*N,l=h+m*N),A&&(y=g-x*A,p=d+x*A),S=cx,o());break;case 32:S!==ax&&S!==cx||(N<0?v=_-m:N>0&&(l=h-m),A<0?y=g-x:A>0&&(p=d-x),S=ux,Y.attr("cursor",hx.selection),o());break;default:return}Cl()}function s(){switch(t.event.keyCode){case 16:U&&(w=M=U=!1,o());break;case 18:S===cx&&(N<0?v=_:N>0&&(l=h),A<0?y=g:A>0&&(p=d),S=ax,o());break;case 32:S===ux&&(t.event.altKey?(N&&(v=_-m*N,l=h+m*N),A&&(y=g-x*A,p=d+x*A),S=cx):(N<0?v=_:N>0&&(l=h),A<0?y=g:A>0&&(p=d),S=ax),Y.attr("cursor",hx[k]),o());break;default:return}Cl()}if(t.event.touches){if(t.event.changedTouches.length<t.event.touches.length)return Cl()}else if(c)return;if(f.apply(this,arguments)){var l,h,p,d,v,_,y,g,m,x,b,w,M,T=this,k=t.event.target.__data__.type,S="selection"===(t.event.metaKey?k="overlay":k)?ox:t.event.altKey?cx:ax,N=n===fx?null:vx[k],A=n===sx?null:_x[k],E=Ll(T),C=E.extent,z=E.selection,P=C[0][0],q=C[0][1],L=C[1][0],R=C[1][1],U=N&&A&&t.event.shiftKey,D=du(T),O=D,F=i(T,arguments).beforestart();"overlay"===k?E.selection=z=[[l=n===fx?P:D[0],p=n===sx?q:D[1]],[v=n===fx?L:l,y=n===sx?R:p]]:(l=z[0][0],p=z[0][1],v=z[1][0],y=z[1][1]),h=l,d=p,_=v,g=y;var I=Ra(T).attr("pointer-events","none"),Y=I.selectAll(".overlay").attr("cursor",hx[k]);if(t.event.touches)I.on("touchmove.brush",e,!0).on("touchend.brush touchcancel.brush",u,!0);else{var B=Ra(t.event.view).on("keydown.brush",a,!0).on("keyup.brush",s,!0).on("mousemove.brush",e,!0).on("mouseup.brush",u,!0);Ef(t.event.view)}El(),Ha(T),r.call(T),F.start()}}function a(){var t=this.__brush||{selection:null};return t.extent=s.apply(this,arguments),t.dim=n,t}var c,s=ql,f=Pl,l=Mr(e,"start","brush","end"),h=6;return e.move=function(t,e){t.selection?t.on("start.brush",function(){i(this,arguments).beforestart().start()}).on("interrupt.brush end.brush",function(){i(this,arguments).end()}).tween("brush",function(){function t(t){u.selection=1===t&&Rl(s)?null:f(t),r.call(o),a.brush()}var o=this,u=o.__brush,a=i(o,arguments),c=u.selection,s=n.input("function"==typeof e?e.apply(this,arguments):e,u.extent),f=cr(c,s);return c&&s?t:t(1)}):t.each(function(){var t=this,o=arguments,u=t.__brush,a=n.input("function"==typeof e?e.apply(t,o):e,u.extent),c=i(t,o).beforestart();Ha(t),u.selection=null==a||Rl(a)?null:a,r.call(t),c.start().brush().end()})},o.prototype={beforestart:function(){return 1===++this.active&&(this.state.emitter=this,this.starting=!0),this},start:function(){return this.starting&&(this.starting=!1,this.emit("start")),this},brush:function(){return this.emit("brush"),this},end:function(){return 0===--this.active&&(delete this.state.emitter,this.emit("end")),this},emit:function(t){lu(new Al(e,t,n.output(this.state.selection)),l.apply,l,[t,this.that,this.args])}},e.extent=function(t){return arguments.length?(s="function"==typeof t?t:Nl([[+t[0][0],+t[0][1]],[+t[1][0],+t[1][1]]]),e):s},e.filter=function(t){return arguments.length?(f="function"==typeof t?t:Nl(!!t),e):f},e.handleSize=function(t){return arguments.length?(h=+t,e):h},e.on=function(){var t=l.on.apply(l,arguments);return t===l?e:t},e}function Yl(t){return function(n,e){return t(n.source.value+n.target.value,e.source.value+e.target.value)}}function Bl(){function t(t){var o,u,a,c,s,f,h=t.length,p=[],d=l(h),v=[],_=[],y=_.groups=new Array(h),g=new Array(h*h);for(o=0,s=-1;++s<h;){for(u=0,f=-1;++f<h;)u+=t[s][f];p.push(u),v.push(l(h)),o+=u}for(e&&d.sort(function(t,n){return e(p[t],p[n])}),r&&v.forEach(function(n,e){n.sort(function(n,i){return r(t[e][n],t[e][i])})}),o=wx(0,bx-n*h)/o,c=o?n:bx/h,u=0,s=-1;++s<h;){for(a=u,f=-1;++f<h;){var m=d[s],x=v[m][f],b=t[m][x],w=u,M=u+=b*o;g[x*h+m]={index:m,subindex:x,startAngle:w,endAngle:M,value:b}}y[m]={index:m,startAngle:a,endAngle:u,value:p[m]},u+=c}for(s=-1;++s<h;)for(f=s-1;++f<h;){var T=g[f*h+s],k=g[s*h+f];(T.value||k.value)&&_.push(T.value<k.value?{source:k,target:T}:{source:T,target:k})}return i?_.sort(i):_}var n=0,e=null,r=null,i=null;return t.padAngle=function(e){return arguments.length?(n=wx(0,e),t):n},t.sortGroups=function(n){return arguments.length?(e=n,t):e},t.sortSubgroups=function(n){return arguments.length?(r=n,t):r},t.sortChords=function(n){
return arguments.length?(null==n?i=null:(i=Yl(n))._=n,t):i&&i._},t}function jl(t){return function(){return t}}function Hl(t){return t.source}function Xl(t){return t.target}function Vl(t){return t.radius}function Wl(t){return t.startAngle}function $l(t){return t.endAngle}function Zl(){function t(){var t,a=Mx.call(arguments),c=n.apply(this,a),s=e.apply(this,a),f=+r.apply(this,(a[0]=c,a)),l=i.apply(this,a)-xx,h=o.apply(this,a)-xx,p=f*yx(l),d=f*gx(l),v=+r.apply(this,(a[0]=s,a)),_=i.apply(this,a)-xx,y=o.apply(this,a)-xx;if(u||(u=t=Tt()),u.moveTo(p,d),u.arc(0,0,f,l,h),l===_&&h===y||(u.quadraticCurveTo(0,0,v*yx(_),v*gx(_)),u.arc(0,0,v,_,y)),u.quadraticCurveTo(0,0,p,d),u.closePath(),t)return u=null,t+""||null}var n=Hl,e=Xl,r=Vl,i=Wl,o=$l,u=null;return t.radius=function(n){return arguments.length?(r="function"==typeof n?n:jl(+n),t):r},t.startAngle=function(n){return arguments.length?(i="function"==typeof n?n:jl(+n),t):i},t.endAngle=function(n){return arguments.length?(o="function"==typeof n?n:jl(+n),t):o},t.source=function(e){return arguments.length?(n=e,t):n},t.target=function(n){return arguments.length?(e=n,t):e},t.context=function(n){return arguments.length?(u=null==n?null:n,t):u},t}function Gl(){return new Jl}function Jl(){this.reset()}function Ql(t,n,e){var r=t.s=n+e,i=r-n,o=r-i;t.t=n-o+(e-i)}function Kl(t){return t>1?0:t<-1?ib:Math.acos(t)}function th(t){return t>1?ob:t<-1?-ob:Math.asin(t)}function nh(t){return(t=gb(t/2))*t}function eh(){}function rh(t,n){t&&Mb.hasOwnProperty(t.type)&&Mb[t.type](t,n)}function ih(t,n,e){var r,i=-1,o=t.length-e;for(n.lineStart();++i<o;)r=t[i],n.point(r[0],r[1],r[2]);n.lineEnd()}function oh(t,n){var e=-1,r=t.length;for(n.polygonStart();++e<r;)ih(t[e],n,1);n.polygonEnd()}function uh(t,n){t&&wb.hasOwnProperty(t.type)?wb[t.type](t,n):rh(t,n)}function ah(){Sb.point=sh}function ch(){fh(Tx,kx)}function sh(t,n){Sb.point=fh,Tx=t,kx=n,t*=sb,n*=sb,Sx=t,Nx=pb(n=n/2+ub),Ax=gb(n)}function fh(t,n){t*=sb,n*=sb,n=n/2+ub;var e=t-Sx,r=e>=0?1:-1,i=r*e,o=pb(n),u=gb(n),a=Ax*u,c=Nx*o+a*pb(i),s=a*r*gb(i);Tb.add(hb(s,c)),Sx=t,Nx=o,Ax=u}function lh(t){return kb.reset(),uh(t,Sb),2*kb}function hh(t){return[hb(t[1],t[0]),th(t[2])]}function ph(t){var n=t[0],e=t[1],r=pb(e);return[r*pb(n),r*gb(n),gb(e)]}function dh(t,n){return t[0]*n[0]+t[1]*n[1]+t[2]*n[2]}function vh(t,n){return[t[1]*n[2]-t[2]*n[1],t[2]*n[0]-t[0]*n[2],t[0]*n[1]-t[1]*n[0]]}function _h(t,n){t[0]+=n[0],t[1]+=n[1],t[2]+=n[2]}function yh(t,n){return[t[0]*n,t[1]*n,t[2]*n]}function gh(t){var n=xb(t[0]*t[0]+t[1]*t[1]+t[2]*t[2]);t[0]/=n,t[1]/=n,t[2]/=n}function mh(t,n){Dx.push(Ox=[Ex=t,zx=t]),n<Cx&&(Cx=n),n>Px&&(Px=n)}function xh(t,n){var e=ph([t*sb,n*sb]);if(Ux){var r=vh(Ux,e),i=[r[1],-r[0],0],o=vh(i,r);gh(o),o=hh(o);var u,a=t-qx,c=a>0?1:-1,s=o[0]*cb*c,f=fb(a)>180;f^(c*qx<s&&s<c*t)?(u=o[1]*cb,u>Px&&(Px=u)):(s=(s+360)%360-180,f^(c*qx<s&&s<c*t)?(u=-o[1]*cb,u<Cx&&(Cx=u)):(n<Cx&&(Cx=n),n>Px&&(Px=n))),f?t<qx?Sh(Ex,t)>Sh(Ex,zx)&&(zx=t):Sh(t,zx)>Sh(Ex,zx)&&(Ex=t):zx>=Ex?(t<Ex&&(Ex=t),t>zx&&(zx=t)):t>qx?Sh(Ex,t)>Sh(Ex,zx)&&(zx=t):Sh(t,zx)>Sh(Ex,zx)&&(Ex=t)}else mh(t,n);Ux=e,qx=t}function bh(){Ab.point=xh}function wh(){Ox[0]=Ex,Ox[1]=zx,Ab.point=mh,Ux=null}function Mh(t,n){if(Ux){var e=t-qx;Nb.add(fb(e)>180?e+(e>0?360:-360):e)}else Lx=t,Rx=n;Sb.point(t,n),xh(t,n)}function Th(){Sb.lineStart()}function kh(){Mh(Lx,Rx),Sb.lineEnd(),fb(Nb)>eb&&(Ex=-(zx=180)),Ox[0]=Ex,Ox[1]=zx,Ux=null}function Sh(t,n){return(n-=t)<0?n+360:n}function Nh(t,n){return t[0]-n[0]}function Ah(t,n){return t[0]<=t[1]?t[0]<=n&&n<=t[1]:n<t[0]||t[1]<n}function Eh(t){var n,e,r,i,o,u,a;if(Px=zx=-(Ex=Cx=1/0),Dx=[],uh(t,Ab),e=Dx.length){for(Dx.sort(Nh),n=1,r=Dx[0],o=[r];n<e;++n)i=Dx[n],Ah(r,i[0])||Ah(r,i[1])?(Sh(r[0],i[1])>Sh(r[0],r[1])&&(r[1]=i[1]),Sh(i[0],r[1])>Sh(r[0],r[1])&&(r[0]=i[0])):o.push(r=i);for(u=-(1/0),e=o.length-1,n=0,r=o[e];n<=e;r=i,++n)i=o[n],(a=Sh(r[1],i[0]))>u&&(u=a,Ex=i[0],zx=r[1])}return Dx=Ox=null,Ex===1/0||Cx===1/0?[[NaN,NaN],[NaN,NaN]]:[[Ex,Cx],[zx,Px]]}function Ch(t,n){t*=sb,n*=sb;var e=pb(n);zh(e*pb(t),e*gb(t),gb(n))}function zh(t,n,e){++Fx,Yx+=(t-Yx)/Fx,Bx+=(n-Bx)/Fx,jx+=(e-jx)/Fx}function Ph(){Eb.point=qh}function qh(t,n){t*=sb,n*=sb;var e=pb(n);Qx=e*pb(t),Kx=e*gb(t),tb=gb(n),Eb.point=Lh,zh(Qx,Kx,tb)}function Lh(t,n){t*=sb,n*=sb;var e=pb(n),r=e*pb(t),i=e*gb(t),o=gb(n),u=hb(xb((u=Kx*o-tb*i)*u+(u=tb*r-Qx*o)*u+(u=Qx*i-Kx*r)*u),Qx*r+Kx*i+tb*o);Ix+=u,Hx+=u*(Qx+(Qx=r)),Xx+=u*(Kx+(Kx=i)),Vx+=u*(tb+(tb=o)),zh(Qx,Kx,tb)}function Rh(){Eb.point=Ch}function Uh(){Eb.point=Oh}function Dh(){Fh(Gx,Jx),Eb.point=Ch}function Oh(t,n){Gx=t,Jx=n,t*=sb,n*=sb,Eb.point=Fh;var e=pb(n);Qx=e*pb(t),Kx=e*gb(t),tb=gb(n),zh(Qx,Kx,tb)}function Fh(t,n){t*=sb,n*=sb;var e=pb(n),r=e*pb(t),i=e*gb(t),o=gb(n),u=Kx*o-tb*i,a=tb*r-Qx*o,c=Qx*i-Kx*r,s=xb(u*u+a*a+c*c),f=Qx*r+Kx*i+tb*o,l=s&&-Kl(f)/s,h=hb(s,f);Wx+=l*u,$x+=l*a,Zx+=l*c,Ix+=h,Hx+=h*(Qx+(Qx=r)),Xx+=h*(Kx+(Kx=i)),Vx+=h*(tb+(tb=o)),zh(Qx,Kx,tb)}function Ih(t){Fx=Ix=Yx=Bx=jx=Hx=Xx=Vx=Wx=$x=Zx=0,uh(t,Eb);var n=Wx,e=$x,r=Zx,i=n*n+e*e+r*r;return i<rb&&(n=Hx,e=Xx,r=Vx,Ix<eb&&(n=Yx,e=Bx,r=jx),i=n*n+e*e+r*r,i<rb)?[NaN,NaN]:[hb(e,n)*cb,th(r/xb(i))*cb]}function Yh(t){return function(){return t}}function Bh(t,n){function e(e,r){return e=t(e,r),n(e[0],e[1])}return t.invert&&n.invert&&(e.invert=function(e,r){return e=n.invert(e,r),e&&t.invert(e[0],e[1])}),e}function jh(t,n){return[t>ib?t-ab:t<-ib?t+ab:t,n]}function Hh(t,n,e){return(t%=ab)?n||e?Bh(Vh(t),Wh(n,e)):Vh(t):n||e?Wh(n,e):jh}function Xh(t){return function(n,e){return n+=t,[n>ib?n-ab:n<-ib?n+ab:n,e]}}function Vh(t){var n=Xh(t);return n.invert=Xh(-t),n}function Wh(t,n){function e(t,n){var e=pb(n),a=pb(t)*e,c=gb(t)*e,s=gb(n),f=s*r+a*i;return[hb(c*o-f*u,a*r-s*i),th(f*o+c*u)]}var r=pb(t),i=gb(t),o=pb(n),u=gb(n);return e.invert=function(t,n){var e=pb(n),a=pb(t)*e,c=gb(t)*e,s=gb(n),f=s*o-c*u;return[hb(c*o+s*u,a*r+f*i),th(f*r-a*i)]},e}function $h(t){function n(n){return n=t(n[0]*sb,n[1]*sb),n[0]*=cb,n[1]*=cb,n}return t=Hh(t[0]*sb,t[1]*sb,t.length>2?t[2]*sb:0),n.invert=function(n){return n=t.invert(n[0]*sb,n[1]*sb),n[0]*=cb,n[1]*=cb,n},n}function Zh(t,n,e,r,i,o){if(e){var u=pb(n),a=gb(n),c=r*e;null==i?(i=n+r*ab,o=n-c/2):(i=Gh(u,i),o=Gh(u,o),(r>0?i<o:i>o)&&(i+=r*ab));for(var s,f=i;r>0?f>o:f<o;f-=c)s=hh([u,-a*pb(f),-a*gb(f)]),t.point(s[0],s[1])}}function Gh(t,n){n=ph(n),n[0]-=t,gh(n);var e=Kl(-n[1]);return((-n[2]<0?-e:e)+ab-eb)%ab}function Jh(){function t(t,n){e.push(t=r(t,n)),t[0]*=cb,t[1]*=cb}function n(){var t=i.apply(this,arguments),n=o.apply(this,arguments)*sb,c=u.apply(this,arguments)*sb;return e=[],r=Hh(-t[0]*sb,-t[1]*sb,0).invert,Zh(a,n,c,1),t={type:"Polygon",coordinates:[e]},e=r=null,t}var e,r,i=Yh([0,0]),o=Yh(90),u=Yh(6),a={point:t};return n.center=function(t){return arguments.length?(i="function"==typeof t?t:Yh([+t[0],+t[1]]),n):i},n.radius=function(t){return arguments.length?(o="function"==typeof t?t:Yh(+t),n):o},n.precision=function(t){return arguments.length?(u="function"==typeof t?t:Yh(+t),n):u},n}function Qh(){var t,n=[];return{point:function(n,e){t.push([n,e])},lineStart:function(){n.push(t=[])},lineEnd:eh,rejoin:function(){n.length>1&&n.push(n.pop().concat(n.shift()))},result:function(){var e=n;return n=[],t=null,e}}}function Kh(t,n,e,r,i,o){var u,a=t[0],c=t[1],s=n[0],f=n[1],l=0,h=1,p=s-a,d=f-c;if(u=e-a,p||!(u>0)){if(u/=p,p<0){if(u<l)return;u<h&&(h=u)}else if(p>0){if(u>h)return;u>l&&(l=u)}if(u=i-a,p||!(u<0)){if(u/=p,p<0){if(u>h)return;u>l&&(l=u)}else if(p>0){if(u<l)return;u<h&&(h=u)}if(u=r-c,d||!(u>0)){if(u/=d,d<0){if(u<l)return;u<h&&(h=u)}else if(d>0){if(u>h)return;u>l&&(l=u)}if(u=o-c,d||!(u<0)){if(u/=d,d<0){if(u>h)return;u>l&&(l=u)}else if(d>0){if(u<l)return;u<h&&(h=u)}return l>0&&(t[0]=a+l*p,t[1]=c+l*d),h<1&&(n[0]=a+h*p,n[1]=c+h*d),!0}}}}}function tp(t,n){return fb(t[0]-n[0])<eb&&fb(t[1]-n[1])<eb}function np(t,n,e,r){this.x=t,this.z=n,this.o=e,this.e=r,this.v=!1,this.n=this.p=null}function ep(t,n,e,r,i){var o,u,a=[],c=[];if(t.forEach(function(t){if(!((n=t.length-1)<=0)){var n,e,r=t[0],u=t[n];if(tp(r,u)){for(i.lineStart(),o=0;o<n;++o)i.point((r=t[o])[0],r[1]);return void i.lineEnd()}a.push(e=new np(r,t,null,!0)),c.push(e.o=new np(r,null,e,!1)),a.push(e=new np(u,t,null,!1)),c.push(e.o=new np(u,null,e,!0))}}),a.length){for(c.sort(n),rp(a),rp(c),o=0,u=c.length;o<u;++o)c[o].e=e=!e;for(var s,f,l=a[0];;){for(var h=l,p=!0;h.v;)if((h=h.n)===l)return;s=h.z,i.lineStart();do{if(h.v=h.o.v=!0,h.e){if(p)for(o=0,u=s.length;o<u;++o)i.point((f=s[o])[0],f[1]);else r(h.x,h.n.x,1,i);h=h.n}else{if(p)for(s=h.p.z,o=s.length-1;o>=0;--o)i.point((f=s[o])[0],f[1]);else r(h.x,h.p.x,-1,i);h=h.p}h=h.o,s=h.z,p=!p}while(!h.v);i.lineEnd()}}}function rp(t){if(n=t.length){for(var n,e,r=0,i=t[0];++r<n;)i.n=e=t[r],e.p=i,i=e;i.n=e=t[0],e.p=i}}function ip(t,n,e,r){function i(i,o){return t<=i&&i<=e&&n<=o&&o<=r}function o(i,o,a,s){var f=0,l=0;if(null==i||(f=u(i,a))!==(l=u(o,a))||c(i,o)<0^a>0){do s.point(0===f||3===f?t:e,f>1?r:n);while((f=(f+a+4)%4)!==l)}else s.point(o[0],o[1])}function u(r,i){return fb(r[0]-t)<eb?i>0?0:3:fb(r[0]-e)<eb?i>0?2:1:fb(r[1]-n)<eb?i>0?1:0:i>0?3:2}function a(t,n){return c(t.x,n.x)}function c(t,n){var e=u(t,1),r=u(n,1);return e!==r?e-r:0===e?n[1]-t[1]:1===e?t[0]-n[0]:2===e?t[1]-n[1]:n[0]-t[0]}return function(u){function c(t,n){i(t,n)&&N.point(t,n)}function s(){for(var n=0,e=0,i=_.length;e<i;++e)for(var o,u,a=_[e],c=1,s=a.length,f=a[0],l=f[0],h=f[1];c<s;++c)o=l,u=h,f=a[c],l=f[0],h=f[1],u<=r?h>r&&(l-o)*(r-u)>(h-u)*(t-o)&&++n:h<=r&&(l-o)*(r-u)<(h-u)*(t-o)&&--n;return n}function f(){N=A,v=[],_=[],S=!0}function l(){var t=s(),n=S&&t,e=(v=w(v)).length;(n||e)&&(u.polygonStart(),n&&(u.lineStart(),o(null,null,1,u),u.lineEnd()),e&&ep(v,a,t,o,u),u.polygonEnd()),N=u,v=_=y=null}function h(){E.point=d,_&&_.push(y=[]),k=!0,T=!1,b=M=NaN}function p(){v&&(d(g,m),x&&T&&A.rejoin(),v.push(A.result())),E.point=c,T&&N.lineEnd()}function d(o,u){var a=i(o,u);if(_&&y.push([o,u]),k)g=o,m=u,x=a,k=!1,a&&(N.lineStart(),N.point(o,u));else if(a&&T)N.point(o,u);else{var c=[b=Math.max(Bb,Math.min(Yb,b)),M=Math.max(Bb,Math.min(Yb,M))],s=[o=Math.max(Bb,Math.min(Yb,o)),u=Math.max(Bb,Math.min(Yb,u))];Kh(c,s,t,n,e,r)?(T||(N.lineStart(),N.point(c[0],c[1])),N.point(s[0],s[1]),a||N.lineEnd(),S=!1):a&&(N.lineStart(),N.point(o,u),S=!1)}b=o,M=u,T=a}var v,_,y,g,m,x,b,M,T,k,S,N=u,A=Qh(),E={point:c,lineStart:h,lineEnd:p,polygonStart:f,polygonEnd:l};return E}}function op(){var t,n,e,r=0,i=0,o=960,u=500;return e={stream:function(e){return t&&n===e?t:t=ip(r,i,o,u)(n=e)},extent:function(a){return arguments.length?(r=+a[0][0],i=+a[0][1],o=+a[1][0],u=+a[1][1],t=n=null,e):[[r,i],[o,u]]}}}function up(){Hb.point=cp,Hb.lineEnd=ap}function ap(){Hb.point=Hb.lineEnd=eh}function cp(t,n){t*=sb,n*=sb,Cb=t,zb=gb(n),Pb=pb(n),Hb.point=sp}function sp(t,n){t*=sb,n*=sb;var e=gb(n),r=pb(n),i=fb(t-Cb),o=pb(i),u=gb(i),a=r*u,c=Pb*e-zb*r*o,s=zb*e+Pb*r*o;jb.add(hb(xb(a*a+c*c),s)),Cb=t,zb=e,Pb=r}function fp(t){return jb.reset(),uh(t,Hb),+jb}function lp(t,n){return Xb[0]=t,Xb[1]=n,fp(Vb)}function hp(t,n,e){var r=l(t,n-eb,e).concat(n);return function(t){return r.map(function(n){return[t,n]})}}function pp(t,n,e){var r=l(t,n-eb,e).concat(n);return function(t){return r.map(function(n){return[n,t]})}}function dp(){function t(){return{type:"MultiLineString",coordinates:n()}}function n(){return l(db(o/y)*y,i,y).map(p).concat(l(db(s/g)*g,c,g).map(d)).concat(l(db(r/v)*v,e,v).filter(function(t){return fb(t%y)>eb}).map(f)).concat(l(db(a/_)*_,u,_).filter(function(t){return fb(t%g)>eb}).map(h))}var e,r,i,o,u,a,c,s,f,h,p,d,v=10,_=v,y=90,g=360,m=2.5;return t.lines=function(){return n().map(function(t){return{type:"LineString",coordinates:t}})},t.outline=function(){return{type:"Polygon",coordinates:[p(o).concat(d(c).slice(1),p(i).reverse().slice(1),d(s).reverse().slice(1))]}},t.extent=function(n){return arguments.length?t.extentMajor(n).extentMinor(n):t.extentMinor()},t.extentMajor=function(n){return arguments.length?(o=+n[0][0],i=+n[1][0],s=+n[0][1],c=+n[1][1],o>i&&(n=o,o=i,i=n),s>c&&(n=s,s=c,c=n),t.precision(m)):[[o,s],[i,c]]},t.extentMinor=function(n){return arguments.length?(r=+n[0][0],e=+n[1][0],a=+n[0][1],u=+n[1][1],r>e&&(n=r,r=e,e=n),a>u&&(n=a,a=u,u=n),t.precision(m)):[[r,a],[e,u]]},t.step=function(n){return arguments.length?t.stepMajor(n).stepMinor(n):t.stepMinor()},t.stepMajor=function(n){return arguments.length?(y=+n[0],g=+n[1],t):[y,g]},t.stepMinor=function(n){return arguments.length?(v=+n[0],_=+n[1],t):[v,_]},t.precision=function(n){return arguments.length?(m=+n,f=hp(a,u,90),h=pp(r,e,m),p=hp(s,c,90),d=pp(o,i,m),t):m},t.extentMajor([[-180,-90+eb],[180,90-eb]]).extentMinor([[-180,-80-eb],[180,80+eb]])}function vp(t,n){var e=t[0]*sb,r=t[1]*sb,i=n[0]*sb,o=n[1]*sb,u=pb(r),a=gb(r),c=pb(o),s=gb(o),f=u*pb(e),l=u*gb(e),h=c*pb(i),p=c*gb(i),d=2*th(xb(nh(o-r)+u*c*nh(i-e))),v=gb(d),_=d?function(t){var n=gb(t*=d)/v,e=gb(d-t)/v,r=e*f+n*h,i=e*l+n*p,o=e*a+n*s;return[hb(i,r)*cb,hb(o,xb(r*r+i*i))*cb]}:function(){return[e*cb,r*cb]};return _.distance=d,_}function _p(t){return t}function yp(){Zb.point=gp}function gp(t,n){Zb.point=mp,qb=Rb=t,Lb=Ub=n}function mp(t,n){$b.add(Ub*t-Rb*n),Rb=t,Ub=n}function xp(){mp(qb,Lb)}function bp(t,n){t<Gb&&(Gb=t),t>Qb&&(Qb=t),n<Jb&&(Jb=n),n>Kb&&(Kb=n)}function wp(t,n){nw+=t,ew+=n,++rw}function Mp(){fw.point=Tp}function Tp(t,n){fw.point=kp,wp(Fb=t,Ib=n)}function kp(t,n){var e=t-Fb,r=n-Ib,i=xb(e*e+r*r);iw+=i*(Fb+t)/2,ow+=i*(Ib+n)/2,uw+=i,wp(Fb=t,Ib=n)}function Sp(){fw.point=wp}function Np(){fw.point=Ep}function Ap(){Cp(Db,Ob)}function Ep(t,n){fw.point=Cp,wp(Db=Fb=t,Ob=Ib=n)}function Cp(t,n){var e=t-Fb,r=n-Ib,i=xb(e*e+r*r);iw+=i*(Fb+t)/2,ow+=i*(Ib+n)/2,uw+=i,i=Ib*t-Fb*n,aw+=i*(Fb+t),cw+=i*(Ib+n),sw+=3*i,wp(Fb=t,Ib=n)}function zp(t){function n(n,e){t.moveTo(n+u,e),t.arc(n,e,u,0,ab)}function e(n,e){t.moveTo(n,e),a.point=r}function r(n,e){t.lineTo(n,e)}function i(){a.point=n}function o(){t.closePath()}var u=4.5,a={point:n,lineStart:function(){a.point=e},lineEnd:i,polygonStart:function(){a.lineEnd=o},polygonEnd:function(){a.lineEnd=i,a.point=n},pointRadius:function(t){return u=t,a},result:eh};return a}function Pp(){function t(t,n){a.push("M",t,",",n,u)}function n(t,n){a.push("M",t,",",n),c.point=e}function e(t,n){a.push("L",t,",",n)}function r(){c.point=n}function i(){c.point=t}function o(){a.push("Z")}var u=qp(4.5),a=[],c={point:t,lineStart:r,lineEnd:i,polygonStart:function(){c.lineEnd=o},polygonEnd:function(){c.lineEnd=i,c.point=t},pointRadius:function(t){return u=qp(t),c},result:function(){if(a.length){var t=a.join("");return a=[],t}}};return c}function qp(t){return"m0,"+t+"a"+t+","+t+" 0 1,1 0,"+-2*t+"a"+t+","+t+" 0 1,1 0,"+2*t+"z"}function Lp(){function t(t){return t&&("function"==typeof o&&i.pointRadius(+o.apply(this,arguments)),uh(t,e(i))),i.result()}var n,e,r,i,o=4.5;return t.area=function(t){return uh(t,e(Zb)),Zb.result()},t.bounds=function(t){return uh(t,e(tw)),tw.result()},t.centroid=function(t){return uh(t,e(fw)),fw.result()},t.projection=function(r){return arguments.length?(e=null==(n=r)?_p:r.stream,t):n},t.context=function(n){return arguments.length?(i=null==(r=n)?new Pp:new zp(n),"function"!=typeof o&&i.pointRadius(o),t):r},t.pointRadius=function(n){return arguments.length?(o="function"==typeof n?n:(i.pointRadius(+n),+n),t):o},t.projection(null).context(null)}function Rp(t,n){var e=n[0],r=n[1],i=[gb(e),-pb(e),0],o=0,u=0;lw.reset();for(var a=0,c=t.length;a<c;++a)if(f=(s=t[a]).length)for(var s,f,l=s[f-1],h=l[0],p=l[1]/2+ub,d=gb(p),v=pb(p),_=0;_<f;++_,h=g,d=x,v=b,l=y){var y=s[_],g=y[0],m=y[1]/2+ub,x=gb(m),b=pb(m),w=g-h,M=w>=0?1:-1,T=M*w,k=T>ib,S=d*x;if(lw.add(hb(S*M*gb(T),v*b+S*pb(T))),o+=k?w+M*ab:w,k^h>=e^g>=e){var N=vh(ph(l),ph(y));gh(N);var A=vh(i,N);gh(A);var E=(k^w>=0?-1:1)*th(A[2]);(r>E||r===E&&(N[0]||N[1]))&&(u+=k^w>=0?1:-1)}}return(o<-eb||o<eb&&lw<-eb)^1&u}function Up(t,n,e,r){return function(i,o){function u(n,e){var r=i(n,e);t(n=r[0],e=r[1])&&o.point(n,e)}function a(t,n){var e=i(t,n);_.point(e[0],e[1])}function c(){b.point=a,_.lineStart()}function s(){b.point=u,_.lineEnd()}function f(t,n){v.push([t,n]);var e=i(t,n);m.point(e[0],e[1])}function l(){m.lineStart(),v=[]}function h(){f(v[0][0],v[0][1]),m.lineEnd();var t,n,e,r,i=m.clean(),u=g.result(),a=u.length;if(v.pop(),p.push(v),v=null,a)if(1&i){if(e=u[0],(n=e.length-1)>0){for(x||(o.polygonStart(),x=!0),o.lineStart(),t=0;t<n;++t)o.point((r=e[t])[0],r[1]);o.lineEnd()}}else a>1&&2&i&&u.push(u.pop().concat(u.shift())),d.push(u.filter(Dp))}var p,d,v,_=n(o),y=i.invert(r[0],r[1]),g=Qh(),m=n(g),x=!1,b={point:u,lineStart:c,lineEnd:s,polygonStart:function(){b.point=f,b.lineStart=l,b.lineEnd=h,d=[],p=[]},polygonEnd:function(){b.point=u,b.lineStart=c,b.lineEnd=s,d=w(d);var t=Rp(p,y);d.length?(x||(o.polygonStart(),x=!0),ep(d,Op,t,e,o)):t&&(x||(o.polygonStart(),x=!0),o.lineStart(),e(null,null,1,o),o.lineEnd()),x&&(o.polygonEnd(),x=!1),d=p=null},sphere:function(){o.polygonStart(),o.lineStart(),e(null,null,1,o),o.lineEnd(),o.polygonEnd()}};return b}}function Dp(t){return t.length>1}function Op(t,n){return((t=t.x)[0]<0?t[1]-ob-eb:ob-t[1])-((n=n.x)[0]<0?n[1]-ob-eb:ob-n[1])}function Fp(t){var n,e=NaN,r=NaN,i=NaN;return{lineStart:function(){t.lineStart(),n=1},point:function(o,u){var a=o>0?ib:-ib,c=fb(o-e);fb(c-ib)<eb?(t.point(e,r=(r+u)/2>0?ob:-ob),t.point(i,r),t.lineEnd(),t.lineStart(),t.point(a,r),t.point(o,r),n=0):i!==a&&c>=ib&&(fb(e-i)<eb&&(e-=i*eb),fb(o-a)<eb&&(o-=a*eb),r=Ip(e,r,o,u),t.point(i,r),t.lineEnd(),t.lineStart(),t.point(a,r),n=0),t.point(e=o,r=u),i=a},lineEnd:function(){t.lineEnd(),e=r=NaN},clean:function(){return 2-n}}}function Ip(t,n,e,r){var i,o,u=gb(t-e);return fb(u)>eb?lb((gb(n)*(o=pb(r))*gb(e)-gb(r)*(i=pb(n))*gb(t))/(i*o*u)):(n+r)/2}function Yp(t,n,e,r){var i;if(null==t)i=e*ob,r.point(-ib,i),r.point(0,i),r.point(ib,i),r.point(ib,0),r.point(ib,-i),r.point(0,-i),r.point(-ib,-i),r.point(-ib,0),r.point(-ib,i);else if(fb(t[0]-n[0])>eb){var o=t[0]<n[0]?ib:-ib;i=e*o/2,r.point(-o,i),r.point(0,i),r.point(o,i)}else r.point(n[0],n[1])}function Bp(t,n){function e(e,r,i,o){Zh(o,t,n,i,e,r)}function r(t,n){return pb(t)*pb(n)>a}function i(t){var n,e,i,a,f;return{lineStart:function(){a=i=!1,f=1},point:function(l,h){var p,d=[l,h],v=r(l,h),_=c?v?0:u(l,h):v?u(l+(l<0?ib:-ib),h):0;if(!n&&(a=i=v)&&t.lineStart(),v!==i&&(p=o(n,d),(tp(n,p)||tp(d,p))&&(d[0]+=eb,d[1]+=eb,v=r(d[0],d[1]))),v!==i)f=0,v?(t.lineStart(),p=o(d,n),t.point(p[0],p[1])):(p=o(n,d),t.point(p[0],p[1]),t.lineEnd()),n=p;else if(s&&n&&c^v){var y;_&e||!(y=o(d,n,!0))||(f=0,c?(t.lineStart(),t.point(y[0][0],y[0][1]),t.point(y[1][0],y[1][1]),t.lineEnd()):(t.point(y[1][0],y[1][1]),t.lineEnd(),t.lineStart(),t.point(y[0][0],y[0][1])))}!v||n&&tp(n,d)||t.point(d[0],d[1]),n=d,i=v,e=_},lineEnd:function(){i&&t.lineEnd(),n=null},clean:function(){return f|(a&&i)<<1}}}function o(t,n,e){var r=ph(t),i=ph(n),o=[1,0,0],u=vh(r,i),c=dh(u,u),s=u[0],f=c-s*s;if(!f)return!e&&t;var l=a*c/f,h=-a*s/f,p=vh(o,u),d=yh(o,l),v=yh(u,h);_h(d,v);var _=p,y=dh(d,_),g=dh(_,_),m=y*y-g*(dh(d,d)-1);if(!(m<0)){var x=xb(m),b=yh(_,(-y-x)/g);if(_h(b,d),b=hh(b),!e)return b;var w,M=t[0],T=n[0],k=t[1],S=n[1];T<M&&(w=M,M=T,T=w);var N=T-M,A=fb(N-ib)<eb,E=A||N<eb;if(!A&&S<k&&(w=k,k=S,S=w),E?A?k+S>0^b[1]<(fb(b[0]-M)<eb?k:S):k<=b[1]&&b[1]<=S:N>ib^(M<=b[0]&&b[0]<=T)){var C=yh(_,(-y+x)/g);return _h(C,d),[b,hh(C)]}}}function u(n,e){var r=c?t:ib-t,i=0;return n<-r?i|=1:n>r&&(i|=2),e<-r?i|=4:e>r&&(i|=8),i}var a=pb(t),c=a>0,s=fb(a)>eb;return Up(r,i,e,c?[0,-t]:[-ib,t-ib])}function jp(t){return{stream:Hp(t)}}function Hp(t){function n(){}var e=n.prototype=Object.create(Xp.prototype);for(var r in t)e[r]=t[r];return function(t){var e=new n;return e.stream=t,e}}function Xp(){}function Vp(t,n,e){var r=n[1][0]-n[0][0],i=n[1][1]-n[0][1],o=t.clipExtent&&t.clipExtent();t.scale(150).translate([0,0]),null!=o&&t.clipExtent(null),uh(e,t.stream(tw));var u=tw.result(),a=Math.min(r/(u[1][0]-u[0][0]),i/(u[1][1]-u[0][1])),c=+n[0][0]+(r-a*(u[1][0]+u[0][0]))/2,s=+n[0][1]+(i-a*(u[1][1]+u[0][1]))/2;return null!=o&&t.clipExtent(o),t.scale(150*a).translate([c,s])}function Wp(t){return function(n,e){return Vp(t,[[0,0],n],e)}}function $p(t){return function(n,e){return Vp(t,n,e)}}function Zp(t,n){return+n?Jp(t,n):Gp(t)}function Gp(t){return Hp({point:function(n,e){n=t(n,e),this.stream.point(n[0],n[1])}})}function Jp(t,n){function e(r,i,o,u,a,c,s,f,l,h,p,d,v,_){var y=s-r,g=f-i,m=y*y+g*g;if(m>4*n&&v--){var x=u+h,b=a+p,w=c+d,M=xb(x*x+b*b+w*w),T=th(w/=M),k=fb(fb(w)-1)<eb||fb(o-l)<eb?(o+l)/2:hb(b,x),S=t(k,T),N=S[0],A=S[1],E=N-r,C=A-i,z=g*E-y*C;(z*z/m>n||fb((y*E+g*C)/m-.5)>.3||u*h+a*p+c*d<dw)&&(e(r,i,o,u,a,c,N,A,k,x/=M,b/=M,w,v,_),_.point(N,A),e(N,A,k,x,b,w,s,f,l,h,p,d,v,_))}}return function(n){function r(e,r){e=t(e,r),n.point(e[0],e[1])}function i(){y=NaN,w.point=o,n.lineStart()}function o(r,i){var o=ph([r,i]),u=t(r,i);e(y,g,_,m,x,b,y=u[0],g=u[1],_=r,m=o[0],x=o[1],b=o[2],pw,n),n.point(y,g)}function u(){w.point=r,n.lineEnd()}function a(){i(),w.point=c,w.lineEnd=s}function c(t,n){o(f=t,n),l=y,h=g,p=m,d=x,v=b,w.point=o}function s(){e(y,g,_,m,x,b,l,h,f,p,d,v,pw,n),w.lineEnd=u,u()}var f,l,h,p,d,v,_,y,g,m,x,b,w={point:r,lineStart:i,lineEnd:u,polygonStart:function(){n.polygonStart(),w.lineStart=a},polygonEnd:function(){n.polygonEnd(),w.lineStart=i}};return w}}function Qp(t){return Kp(function(){return t})()}function Kp(t){function n(t){return t=f(t[0]*sb,t[1]*sb),[t[0]*_+a,c-t[1]*_]}function e(t){return t=f.invert((t[0]-a)/_,(c-t[1])/_),t&&[t[0]*cb,t[1]*cb]}function r(t,n){return t=u(t,n),[t[0]*_+a,c-t[1]*_]}function i(){f=Bh(s=Hh(b,w,M),u);var t=u(m,x);return a=y-t[0]*_,c=g+t[1]*_,o()}function o(){return d=v=null,n}var u,a,c,s,f,l,h,p,d,v,_=150,y=480,g=250,m=0,x=0,b=0,w=0,M=0,T=null,k=hw,S=null,N=_p,A=.5,E=Zp(r,A);return n.stream=function(t){return d&&v===t?d:d=vw(k(s,E(N(v=t))))},n.clipAngle=function(t){return arguments.length?(k=+t?Bp(T=t*sb,6*sb):(T=null,hw),o()):T*cb},n.clipExtent=function(t){return arguments.length?(N=null==t?(S=l=h=p=null,_p):ip(S=+t[0][0],l=+t[0][1],h=+t[1][0],p=+t[1][1]),o()):null==S?null:[[S,l],[h,p]]},n.scale=function(t){return arguments.length?(_=+t,i()):_},n.translate=function(t){return arguments.length?(y=+t[0],g=+t[1],i()):[y,g]},n.center=function(t){return arguments.length?(m=t[0]%360*sb,x=t[1]%360*sb,i()):[m*cb,x*cb]},n.rotate=function(t){return arguments.length?(b=t[0]%360*sb,w=t[1]%360*sb,M=t.length>2?t[2]%360*sb:0,i()):[b*cb,w*cb,M*cb]},n.precision=function(t){return arguments.length?(E=Zp(r,A=t*t),o()):xb(A)},n.fitExtent=$p(n),n.fitSize=Wp(n),function(){return u=t.apply(this,arguments),n.invert=u.invert&&e,i()}}function td(t){var n=0,e=ib/3,r=Kp(t),i=r(n,e);return i.parallels=function(t){return arguments.length?r(n=t[0]*sb,e=t[1]*sb):[n*cb,e*cb]},i}function nd(t,n){function e(t,n){var e=xb(o-2*i*gb(n))/i;return[e*gb(t*=i),u-e*pb(t)]}var r=gb(t),i=(r+gb(n))/2,o=1+r*(2*i-r),u=xb(o)/i;return e.invert=function(t,n){var e=u-n;return[hb(t,e)/i,th((o-(t*t+e*e)*i*i)/(2*i))]},e}function ed(){return td(nd).scale(155.424).center([0,33.6442])}function rd(){return ed().parallels([29.5,45.5]).scale(1070).translate([480,250]).rotate([96,0]).center([-.6,38.7])}function id(t){var n=t.length;return{point:function(e,r){for(var i=-1;++i<n;)t[i].point(e,r)},sphere:function(){for(var e=-1;++e<n;)t[e].sphere()},lineStart:function(){for(var e=-1;++e<n;)t[e].lineStart()},lineEnd:function(){for(var e=-1;++e<n;)t[e].lineEnd()},polygonStart:function(){for(var e=-1;++e<n;)t[e].polygonStart()},polygonEnd:function(){for(var e=-1;++e<n;)t[e].polygonEnd()}}}function od(){function t(t){var n=t[0],e=t[1];return u=null,r.point(n,e),u||(i.point(n,e),u)||(o.point(n,e),u)}var n,e,r,i,o,u,a=rd(),c=ed().rotate([154,0]).center([-2,58.5]).parallels([55,65]),s=ed().rotate([157,0]).center([-3,19.9]).parallels([8,18]),f={point:function(t,n){u=[t,n]}};return t.invert=function(t){var n=a.scale(),e=a.translate(),r=(t[0]-e[0])/n,i=(t[1]-e[1])/n;return(i>=.12&&i<.234&&r>=-.425&&r<-.214?c:i>=.166&&i<.234&&r>=-.214&&r<-.115?s:a).invert(t)},t.stream=function(t){return n&&e===t?n:n=id([a.stream(e=t),c.stream(t),s.stream(t)])},t.precision=function(n){return arguments.length?(a.precision(n),c.precision(n),s.precision(n),t):a.precision()},t.scale=function(n){return arguments.length?(a.scale(n),c.scale(.35*n),s.scale(n),t.translate(a.translate())):a.scale()},t.translate=function(n){if(!arguments.length)return a.translate();var e=a.scale(),u=+n[0],l=+n[1];return r=a.translate(n).clipExtent([[u-.455*e,l-.238*e],[u+.455*e,l+.238*e]]).stream(f),i=c.translate([u-.307*e,l+.201*e]).clipExtent([[u-.425*e+eb,l+.12*e+eb],[u-.214*e-eb,l+.234*e-eb]]).stream(f),o=s.translate([u-.205*e,l+.212*e]).clipExtent([[u-.214*e+eb,l+.166*e+eb],[u-.115*e-eb,l+.234*e-eb]]).stream(f),t},t.fitExtent=$p(t),t.fitSize=Wp(t),t.scale(1070)}function ud(t){return function(n,e){var r=pb(n),i=pb(e),o=t(r*i);return[o*i*gb(n),o*gb(e)]}}function ad(t){return function(n,e){var r=xb(n*n+e*e),i=t(r),o=gb(i),u=pb(i);return[hb(n*o,r*u),th(r&&e*o/r)]}}function cd(){return Qp(_w).scale(124.75).clipAngle(179.999)}function sd(){return Qp(yw).scale(79.4188).clipAngle(179.999)}function fd(t,n){return[t,_b(bb((ob+n)/2))]}function ld(){return hd(fd).scale(961/ab)}function hd(t){var n,e=Qp(t),r=e.scale,i=e.translate,o=e.clipExtent;return e.scale=function(t){return arguments.length?(r(t),n&&e.clipExtent(null),e):r()},e.translate=function(t){return arguments.length?(i(t),n&&e.clipExtent(null),e):i()},e.clipExtent=function(t){if(!arguments.length)return n?null:o();if(n=null==t){var u=ib*r(),a=i();t=[[a[0]-u,a[1]-u],[a[0]+u,a[1]+u]]}return o(t),e},e.clipExtent(null)}function pd(t){return bb((ob+t)/2)}function dd(t,n){function e(t,n){o>0?n<-ob+eb&&(n=-ob+eb):n>ob-eb&&(n=ob-eb);var e=o/yb(pd(n),i);return[e*gb(i*t),o-e*pb(i*t)]}var r=pb(t),i=t===n?gb(t):_b(r/pb(n))/_b(pd(n)/pd(t)),o=r*yb(pd(t),i)/i;return i?(e.invert=function(t,n){var e=o-n,r=mb(i)*xb(t*t+e*e);return[hb(t,e)/i,2*lb(yb(o/r,1/i))-ob]},e):fd}function vd(){return td(dd).scale(109.5).parallels([30,30])}function _d(t,n){return[t,n]}function yd(){return Qp(_d).scale(152.63)}function gd(t,n){function e(t,n){var e=o-n,r=i*t;return[e*gb(r),o-e*pb(r)]}var r=pb(t),i=t===n?gb(t):(r-pb(n))/(n-t),o=r/i+t;return fb(i)<eb?_d:(e.invert=function(t,n){var e=o-n;return[hb(t,e)/i,o-mb(i)*xb(t*t+e*e)]},e)}function md(){return td(gd).scale(131.154).center([0,13.9389])}function xd(t,n){var e=pb(n),r=pb(t)*e;return[e*gb(t)/r,gb(n)/r]}function bd(){return Qp(xd).scale(144.049).clipAngle(60)}function wd(t,n){return[pb(n)*gb(t),gb(n)]}function Md(){return Qp(wd).scale(249.5).clipAngle(90+eb)}function Td(t,n){var e=pb(n),r=1+pb(t)*e;return[e*gb(t)/r,gb(n)/r]}function kd(){return Qp(Td).scale(250).clipAngle(142)}function Sd(t,n){return[_b(bb((ob+n)/2)),-t]}function Nd(){var t=hd(Sd),n=t.center,e=t.rotate;return t.center=function(t){return arguments.length?n([-t[1],t[0]]):(t=n(),[t[1],-t[0]])},t.rotate=function(t){return arguments.length?e([t[0],t[1],t.length>2?t[2]+90:90]):(t=e(),[t[0],t[1],t[2]-90])},e([0,0,90]).scale(159.155)}var Ad="4.2.2",Ed=e(n),Cd=Ed.right,zd=Ed.left,Pd=Array.prototype,qd=Pd.slice,Ld=Pd.map,Rd=Math.sqrt(50),Ud=Math.sqrt(10),Dd=Math.sqrt(2),Od="$";P.prototype=q.prototype={constructor:P,has:function(t){return Od+t in this},get:function(t){return this[Od+t]},set:function(t,n){return this[Od+t]=n,this},remove:function(t){var n=Od+t;return n in this&&delete this[n]},clear:function(){for(var t in this)t[0]===Od&&delete this[t]},keys:function(){var t=[];for(var n in this)n[0]===Od&&t.push(n.slice(1));return t},values:function(){var t=[];for(var n in this)n[0]===Od&&t.push(this[n]);return t},entries:function(){var t=[];for(var n in this)n[0]===Od&&t.push({key:n.slice(1),value:this[n]});return t},size:function(){var t=0;for(var n in this)n[0]===Od&&++t;return t},empty:function(){for(var t in this)if(t[0]===Od)return!1;return!0},each:function(t){for(var n in this)n[0]===Od&&t(this[n],n.slice(1),this)}};var Fd=q.prototype;F.prototype=I.prototype={constructor:F,has:Fd.has,add:function(t){return t+="",this[Od+t]=t,this},remove:Fd.remove,clear:Fd.clear,values:Fd.keys,size:Fd.size,empty:Fd.empty,each:Fd.each};var Id=3,Yd=function t(n){function e(t){return Math.pow(t,n)}return n=+n,e.exponent=t,e}(Id),Bd=function t(n){function e(t){return 1-Math.pow(1-t,n)}return n=+n,e.exponent=t,e}(Id),jd=function t(n){function e(t){return((t*=2)<=1?Math.pow(t,n):2-Math.pow(2-t,n))/2}return n=+n,e.exponent=t,e}(Id),Hd=Math.PI,Xd=Hd/2,Vd=4/11,Wd=6/11,$d=8/11,Zd=.75,Gd=9/11,Jd=10/11,Qd=.9375,Kd=21/22,tv=63/64,nv=1/Vd/Vd,ev=1.70158,rv=function t(n){function e(t){return t*t*((n+1)*t-n)}return n=+n,e.overshoot=t,e}(ev),iv=function t(n){function e(t){return--t*t*((n+1)*t+n)+1}return n=+n,e.overshoot=t,e}(ev),ov=function t(n){function e(t){return((t*=2)<1?t*t*((n+1)*t-n):(t-=2)*t*((n+1)*t+n)+2)/2}return n=+n,e.overshoot=t,e}(ev),uv=2*Math.PI,av=1,cv=.3,sv=function t(n,e){function r(t){return n*Math.pow(2,10*--t)*Math.sin((i-t)/e)}var i=Math.asin(1/(n=Math.max(1,n)))*(e/=uv);return r.amplitude=function(n){return t(n,e*uv)},r.period=function(e){return t(n,e)},r}(av,cv),fv=function t(n,e){function r(t){return 1-n*Math.pow(2,-10*(t=+t))*Math.sin((t+i)/e)}var i=Math.asin(1/(n=Math.max(1,n)))*(e/=uv);return r.amplitude=function(n){return t(n,e*uv)},r.period=function(e){return t(n,e)},r}(av,cv),lv=function t(n,e){function r(t){return((t=2*t-1)<0?n*Math.pow(2,10*t)*Math.sin((i-t)/e):2-n*Math.pow(2,-10*t)*Math.sin((i+t)/e))/2}var i=Math.asin(1/(n=Math.max(1,n)))*(e/=uv);return r.amplitude=function(n){return t(n,e*uv)},r.period=function(e){return t(n,e)},r}(av,cv),hv=Math.PI,pv=2*hv,dv=1e-6,vv=pv-dv;Mt.prototype=Tt.prototype={constructor:Mt,moveTo:function(t,n){this._.push("M",this._x0=this._x1=+t,",",this._y0=this._y1=+n)},closePath:function(){null!==this._x1&&(this._x1=this._x0,this._y1=this._y0,this._.push("Z"))},lineTo:function(t,n){this._.push("L",this._x1=+t,",",this._y1=+n)},quadraticCurveTo:function(t,n,e,r){this._.push("Q",+t,",",+n,",",this._x1=+e,",",this._y1=+r)},bezierCurveTo:function(t,n,e,r,i,o){this._.push("C",+t,",",+n,",",+e,",",+r,",",this._x1=+i,",",this._y1=+o)},arcTo:function(t,n,e,r,i){t=+t,n=+n,e=+e,r=+r,i=+i;var o=this._x1,u=this._y1,a=e-t,c=r-n,s=o-t,f=u-n,l=s*s+f*f;if(i<0)throw new Error("negative radius: "+i);if(null===this._x1)this._.push("M",this._x1=t,",",this._y1=n);else if(l>dv)if(Math.abs(f*a-c*s)>dv&&i){var h=e-o,p=r-u,d=a*a+c*c,v=h*h+p*p,_=Math.sqrt(d),y=Math.sqrt(l),g=i*Math.tan((hv-Math.acos((d+l-v)/(2*_*y)))/2),m=g/y,x=g/_;Math.abs(m-1)>dv&&this._.push("L",t+m*s,",",n+m*f),this._.push("A",i,",",i,",0,0,",+(f*h>s*p),",",this._x1=t+x*a,",",this._y1=n+x*c)}else this._.push("L",this._x1=t,",",this._y1=n);else;},arc:function(t,n,e,r,i,o){t=+t,n=+n,e=+e;var u=e*Math.cos(r),a=e*Math.sin(r),c=t+u,s=n+a,f=1^o,l=o?r-i:i-r;if(e<0)throw new Error("negative radius: "+e);null===this._x1?this._.push("M",c,",",s):(Math.abs(this._x1-c)>dv||Math.abs(this._y1-s)>dv)&&this._.push("L",c,",",s),e&&(l>vv?this._.push("A",e,",",e,",0,1,",f,",",t-u,",",n-a,"A",e,",",e,",0,1,",f,",",this._x1=c,",",this._y1=s):(l<0&&(l=l%pv+pv),this._.push("A",e,",",e,",0,",+(l>=hv),",",f,",",this._x1=t+e*Math.cos(i),",",this._y1=n+e*Math.sin(i))))},rect:function(t,n,e,r){this._.push("M",this._x0=this._x1=+t,",",this._y0=this._y1=+n,"h",+e,"v",+r,"h",-e,"Z")},toString:function(){return this._.join("")}};var _v=jt.prototype=Ht.prototype;_v.copy=function(){var t,n,e=new Ht(this._x,this._y,this._x0,this._y0,this._x1,this._y1),r=this._root;if(!r)return e;if(!r.length)return e._root=Xt(r),e;for(t=[{source:r,target:e._root=new Array(4)}];r=t.pop();)for(var i=0;i<4;++i)(n=r.source[i])&&(n.length?t.push({source:n,target:r.target[i]=new Array(4)}):r.target[i]=Xt(n));return e},_v.add=kt,_v.addAll=Nt,_v.cover=At,_v.data=Et,_v.extent=Ct,_v.find=Pt,_v.remove=qt,_v.removeAll=Lt,_v.root=Rt,_v.size=Ut,_v.visit=Dt,_v.visitAfter=Ot,_v.x=It,_v.y=Bt;var yv=[].slice,gv={};Vt.prototype=Qt.prototype={constructor:Vt,defer:function(t){if("function"!=typeof t||this._call)throw new Error;if(null!=this._error)return this;var n=yv.call(arguments,1);return n.push(t),++this._waiting,this._tasks.push(n),Wt(this),this},abort:function(){return null==this._error&&Gt(this,new Error("abort")),
this},await:function(t){if("function"!=typeof t||this._call)throw new Error;return this._call=function(n,e){t.apply(null,[n].concat(e))},Jt(this),this},awaitAll:function(t){if("function"!=typeof t||this._call)throw new Error;return this._call=t,Jt(this),this}};var mv=1e-12,xv=Math.PI,bv=xv/2,wv=2*xv;fn.prototype={areaStart:function(){this._line=0},areaEnd:function(){this._line=NaN},lineStart:function(){this._point=0},lineEnd:function(){(this._line||0!==this._line&&1===this._point)&&this._context.closePath(),this._line=1-this._line},point:function(t,n){switch(t=+t,n=+n,this._point){case 0:this._point=1,this._line?this._context.lineTo(t,n):this._context.moveTo(t,n);break;case 1:this._point=2;default:this._context.lineTo(t,n)}}};var Mv=xn(ln);mn.prototype={areaStart:function(){this._curve.areaStart()},areaEnd:function(){this._curve.areaEnd()},lineStart:function(){this._curve.lineStart()},lineEnd:function(){this._curve.lineEnd()},point:function(t,n){this._curve.point(n*Math.sin(t),n*-Math.cos(t))}};var Tv={draw:function(t,n){var e=Math.sqrt(n/xv);t.moveTo(e,0),t.arc(0,0,e,0,wv)}},kv={draw:function(t,n){var e=Math.sqrt(n/5)/2;t.moveTo(-3*e,-e),t.lineTo(-e,-e),t.lineTo(-e,-3*e),t.lineTo(e,-3*e),t.lineTo(e,-e),t.lineTo(3*e,-e),t.lineTo(3*e,e),t.lineTo(e,e),t.lineTo(e,3*e),t.lineTo(-e,3*e),t.lineTo(-e,e),t.lineTo(-3*e,e),t.closePath()}},Sv=Math.sqrt(1/3),Nv=2*Sv,Av={draw:function(t,n){var e=Math.sqrt(n/Nv),r=e*Sv;t.moveTo(0,-e),t.lineTo(r,0),t.lineTo(0,e),t.lineTo(-r,0),t.closePath()}},Ev=.8908130915292852,Cv=Math.sin(xv/10)/Math.sin(7*xv/10),zv=Math.sin(wv/10)*Cv,Pv=-Math.cos(wv/10)*Cv,qv={draw:function(t,n){var e=Math.sqrt(n*Ev),r=zv*e,i=Pv*e;t.moveTo(0,-e),t.lineTo(r,i);for(var o=1;o<5;++o){var u=wv*o/5,a=Math.cos(u),c=Math.sin(u);t.lineTo(c*e,-a*e),t.lineTo(a*r-c*i,c*r+a*i)}t.closePath()}},Lv={draw:function(t,n){var e=Math.sqrt(n),r=-e/2;t.rect(r,r,e,e)}},Rv=Math.sqrt(3),Uv={draw:function(t,n){var e=-Math.sqrt(n/(3*Rv));t.moveTo(0,2*e),t.lineTo(-Rv*e,-e),t.lineTo(Rv*e,-e),t.closePath()}},Dv=-.5,Ov=Math.sqrt(3)/2,Fv=1/Math.sqrt(12),Iv=3*(Fv/2+1),Yv={draw:function(t,n){var e=Math.sqrt(n/Iv),r=e/2,i=e*Fv,o=r,u=e*Fv+e,a=-o,c=u;t.moveTo(r,i),t.lineTo(o,u),t.lineTo(a,c),t.lineTo(Dv*r-Ov*i,Ov*r+Dv*i),t.lineTo(Dv*o-Ov*u,Ov*o+Dv*u),t.lineTo(Dv*a-Ov*c,Ov*a+Dv*c),t.lineTo(Dv*r+Ov*i,Dv*i-Ov*r),t.lineTo(Dv*o+Ov*u,Dv*u-Ov*o),t.lineTo(Dv*a+Ov*c,Dv*c-Ov*a),t.closePath()}},Bv=[Tv,kv,Av,Lv,qv,Uv,Yv];Nn.prototype={areaStart:function(){this._line=0},areaEnd:function(){this._line=NaN},lineStart:function(){this._x0=this._x1=this._y0=this._y1=NaN,this._point=0},lineEnd:function(){switch(this._point){case 3:Sn(this,this._x1,this._y1);case 2:this._context.lineTo(this._x1,this._y1)}(this._line||0!==this._line&&1===this._point)&&this._context.closePath(),this._line=1-this._line},point:function(t,n){switch(t=+t,n=+n,this._point){case 0:this._point=1,this._line?this._context.lineTo(t,n):this._context.moveTo(t,n);break;case 1:this._point=2;break;case 2:this._point=3,this._context.lineTo((5*this._x0+this._x1)/6,(5*this._y0+this._y1)/6);default:Sn(this,t,n)}this._x0=this._x1,this._x1=t,this._y0=this._y1,this._y1=n}},En.prototype={areaStart:kn,areaEnd:kn,lineStart:function(){this._x0=this._x1=this._x2=this._x3=this._x4=this._y0=this._y1=this._y2=this._y3=this._y4=NaN,this._point=0},lineEnd:function(){switch(this._point){case 1:this._context.moveTo(this._x2,this._y2),this._context.closePath();break;case 2:this._context.moveTo((this._x2+2*this._x3)/3,(this._y2+2*this._y3)/3),this._context.lineTo((this._x3+2*this._x2)/3,(this._y3+2*this._y2)/3),this._context.closePath();break;case 3:this.point(this._x2,this._y2),this.point(this._x3,this._y3),this.point(this._x4,this._y4)}},point:function(t,n){switch(t=+t,n=+n,this._point){case 0:this._point=1,this._x2=t,this._y2=n;break;case 1:this._point=2,this._x3=t,this._y3=n;break;case 2:this._point=3,this._x4=t,this._y4=n,this._context.moveTo((this._x0+4*this._x1+t)/6,(this._y0+4*this._y1+n)/6);break;default:Sn(this,t,n)}this._x0=this._x1,this._x1=t,this._y0=this._y1,this._y1=n}},zn.prototype={areaStart:function(){this._line=0},areaEnd:function(){this._line=NaN},lineStart:function(){this._x0=this._x1=this._y0=this._y1=NaN,this._point=0},lineEnd:function(){(this._line||0!==this._line&&3===this._point)&&this._context.closePath(),this._line=1-this._line},point:function(t,n){switch(t=+t,n=+n,this._point){case 0:this._point=1;break;case 1:this._point=2;break;case 2:this._point=3;var e=(this._x0+4*this._x1+t)/6,r=(this._y0+4*this._y1+n)/6;this._line?this._context.lineTo(e,r):this._context.moveTo(e,r);break;case 3:this._point=4;default:Sn(this,t,n)}this._x0=this._x1,this._x1=t,this._y0=this._y1,this._y1=n}},qn.prototype={lineStart:function(){this._x=[],this._y=[],this._basis.lineStart()},lineEnd:function(){var t=this._x,n=this._y,e=t.length-1;if(e>0)for(var r,i=t[0],o=n[0],u=t[e]-i,a=n[e]-o,c=-1;++c<=e;)r=c/e,this._basis.point(this._beta*t[c]+(1-this._beta)*(i+r*u),this._beta*n[c]+(1-this._beta)*(o+r*a));this._x=this._y=null,this._basis.lineEnd()},point:function(t,n){this._x.push(+t),this._y.push(+n)}};var jv=function t(n){function e(t){return 1===n?new Nn(t):new qn(t,n)}return e.beta=function(n){return t(+n)},e}(.85);Rn.prototype={areaStart:function(){this._line=0},areaEnd:function(){this._line=NaN},lineStart:function(){this._x0=this._x1=this._x2=this._y0=this._y1=this._y2=NaN,this._point=0},lineEnd:function(){switch(this._point){case 2:this._context.lineTo(this._x2,this._y2);break;case 3:Ln(this,this._x1,this._y1)}(this._line||0!==this._line&&1===this._point)&&this._context.closePath(),this._line=1-this._line},point:function(t,n){switch(t=+t,n=+n,this._point){case 0:this._point=1,this._line?this._context.lineTo(t,n):this._context.moveTo(t,n);break;case 1:this._point=2,this._x1=t,this._y1=n;break;case 2:this._point=3;default:Ln(this,t,n)}this._x0=this._x1,this._x1=this._x2,this._x2=t,this._y0=this._y1,this._y1=this._y2,this._y2=n}};var Hv=function t(n){function e(t){return new Rn(t,n)}return e.tension=function(n){return t(+n)},e}(0);Un.prototype={areaStart:kn,areaEnd:kn,lineStart:function(){this._x0=this._x1=this._x2=this._x3=this._x4=this._x5=this._y0=this._y1=this._y2=this._y3=this._y4=this._y5=NaN,this._point=0},lineEnd:function(){switch(this._point){case 1:this._context.moveTo(this._x3,this._y3),this._context.closePath();break;case 2:this._context.lineTo(this._x3,this._y3),this._context.closePath();break;case 3:this.point(this._x3,this._y3),this.point(this._x4,this._y4),this.point(this._x5,this._y5)}},point:function(t,n){switch(t=+t,n=+n,this._point){case 0:this._point=1,this._x3=t,this._y3=n;break;case 1:this._point=2,this._context.moveTo(this._x4=t,this._y4=n);break;case 2:this._point=3,this._x5=t,this._y5=n;break;default:Ln(this,t,n)}this._x0=this._x1,this._x1=this._x2,this._x2=t,this._y0=this._y1,this._y1=this._y2,this._y2=n}};var Xv=function t(n){function e(t){return new Un(t,n)}return e.tension=function(n){return t(+n)},e}(0);Dn.prototype={areaStart:function(){this._line=0},areaEnd:function(){this._line=NaN},lineStart:function(){this._x0=this._x1=this._x2=this._y0=this._y1=this._y2=NaN,this._point=0},lineEnd:function(){(this._line||0!==this._line&&3===this._point)&&this._context.closePath(),this._line=1-this._line},point:function(t,n){switch(t=+t,n=+n,this._point){case 0:this._point=1;break;case 1:this._point=2;break;case 2:this._point=3,this._line?this._context.lineTo(this._x2,this._y2):this._context.moveTo(this._x2,this._y2);break;case 3:this._point=4;default:Ln(this,t,n)}this._x0=this._x1,this._x1=this._x2,this._x2=t,this._y0=this._y1,this._y1=this._y2,this._y2=n}};var Vv=function t(n){function e(t){return new Dn(t,n)}return e.tension=function(n){return t(+n)},e}(0);Fn.prototype={areaStart:function(){this._line=0},areaEnd:function(){this._line=NaN},lineStart:function(){this._x0=this._x1=this._x2=this._y0=this._y1=this._y2=NaN,this._l01_a=this._l12_a=this._l23_a=this._l01_2a=this._l12_2a=this._l23_2a=this._point=0},lineEnd:function(){switch(this._point){case 2:this._context.lineTo(this._x2,this._y2);break;case 3:this.point(this,this._x2,this._y2)}(this._line||0!==this._line&&1===this._point)&&this._context.closePath(),this._line=1-this._line},point:function(t,n){if(t=+t,n=+n,this._point){var e=this._x2-t,r=this._y2-n;this._l23_a=Math.sqrt(this._l23_2a=Math.pow(e*e+r*r,this._alpha))}switch(this._point){case 0:this._point=1,this._line?this._context.lineTo(t,n):this._context.moveTo(t,n);break;case 1:this._point=2;break;case 2:this._point=3;default:On(this,t,n)}this._l01_a=this._l12_a,this._l12_a=this._l23_a,this._l01_2a=this._l12_2a,this._l12_2a=this._l23_2a,this._x0=this._x1,this._x1=this._x2,this._x2=t,this._y0=this._y1,this._y1=this._y2,this._y2=n}};var Wv=function t(n){function e(t){return n?new Fn(t,n):new Rn(t,0)}return e.alpha=function(n){return t(+n)},e}(.5);In.prototype={areaStart:kn,areaEnd:kn,lineStart:function(){this._x0=this._x1=this._x2=this._x3=this._x4=this._x5=this._y0=this._y1=this._y2=this._y3=this._y4=this._y5=NaN,this._l01_a=this._l12_a=this._l23_a=this._l01_2a=this._l12_2a=this._l23_2a=this._point=0},lineEnd:function(){switch(this._point){case 1:this._context.moveTo(this._x3,this._y3),this._context.closePath();break;case 2:this._context.lineTo(this._x3,this._y3),this._context.closePath();break;case 3:this.point(this._x3,this._y3),this.point(this._x4,this._y4),this.point(this._x5,this._y5)}},point:function(t,n){if(t=+t,n=+n,this._point){var e=this._x2-t,r=this._y2-n;this._l23_a=Math.sqrt(this._l23_2a=Math.pow(e*e+r*r,this._alpha))}switch(this._point){case 0:this._point=1,this._x3=t,this._y3=n;break;case 1:this._point=2,this._context.moveTo(this._x4=t,this._y4=n);break;case 2:this._point=3,this._x5=t,this._y5=n;break;default:On(this,t,n)}this._l01_a=this._l12_a,this._l12_a=this._l23_a,this._l01_2a=this._l12_2a,this._l12_2a=this._l23_2a,this._x0=this._x1,this._x1=this._x2,this._x2=t,this._y0=this._y1,this._y1=this._y2,this._y2=n}};var $v=function t(n){function e(t){return n?new In(t,n):new Un(t,0)}return e.alpha=function(n){return t(+n)},e}(.5);Yn.prototype={areaStart:function(){this._line=0},areaEnd:function(){this._line=NaN},lineStart:function(){this._x0=this._x1=this._x2=this._y0=this._y1=this._y2=NaN,this._l01_a=this._l12_a=this._l23_a=this._l01_2a=this._l12_2a=this._l23_2a=this._point=0},lineEnd:function(){(this._line||0!==this._line&&3===this._point)&&this._context.closePath(),this._line=1-this._line},point:function(t,n){if(t=+t,n=+n,this._point){var e=this._x2-t,r=this._y2-n;this._l23_a=Math.sqrt(this._l23_2a=Math.pow(e*e+r*r,this._alpha))}switch(this._point){case 0:this._point=1;break;case 1:this._point=2;break;case 2:this._point=3,this._line?this._context.lineTo(this._x2,this._y2):this._context.moveTo(this._x2,this._y2);break;case 3:this._point=4;default:On(this,t,n)}this._l01_a=this._l12_a,this._l12_a=this._l23_a,this._l01_2a=this._l12_2a,this._l12_2a=this._l23_2a,this._x0=this._x1,this._x1=this._x2,this._x2=t,this._y0=this._y1,this._y1=this._y2,this._y2=n}};var Zv=function t(n){function e(t){return n?new Yn(t,n):new Dn(t,0)}return e.alpha=function(n){return t(+n)},e}(.5);Bn.prototype={areaStart:kn,areaEnd:kn,lineStart:function(){this._point=0},lineEnd:function(){this._point&&this._context.closePath()},point:function(t,n){t=+t,n=+n,this._point?this._context.lineTo(t,n):(this._point=1,this._context.moveTo(t,n))}},$n.prototype={areaStart:function(){this._line=0},areaEnd:function(){this._line=NaN},lineStart:function(){this._x0=this._x1=this._y0=this._y1=this._t0=NaN,this._point=0},lineEnd:function(){switch(this._point){case 2:this._context.lineTo(this._x1,this._y1);break;case 3:Wn(this,this._t0,Vn(this,this._t0))}(this._line||0!==this._line&&1===this._point)&&this._context.closePath(),this._line=1-this._line},point:function(t,n){var e=NaN;if(t=+t,n=+n,t!==this._x1||n!==this._y1){switch(this._point){case 0:this._point=1,this._line?this._context.lineTo(t,n):this._context.moveTo(t,n);break;case 1:this._point=2;break;case 2:this._point=3,Wn(this,Vn(this,e=Xn(this,t,n)),e);break;default:Wn(this,this._t0,e=Xn(this,t,n))}this._x0=this._x1,this._x1=t,this._y0=this._y1,this._y1=n,this._t0=e}}},(Zn.prototype=Object.create($n.prototype)).point=function(t,n){$n.prototype.point.call(this,n,t)},Gn.prototype={moveTo:function(t,n){this._context.moveTo(n,t)},closePath:function(){this._context.closePath()},lineTo:function(t,n){this._context.lineTo(n,t)},bezierCurveTo:function(t,n,e,r,i,o){this._context.bezierCurveTo(n,t,r,e,o,i)}},Kn.prototype={areaStart:function(){this._line=0},areaEnd:function(){this._line=NaN},lineStart:function(){this._x=[],this._y=[]},lineEnd:function(){var t=this._x,n=this._y,e=t.length;if(e)if(this._line?this._context.lineTo(t[0],n[0]):this._context.moveTo(t[0],n[0]),2===e)this._context.lineTo(t[1],n[1]);else for(var r=te(t),i=te(n),o=0,u=1;u<e;++o,++u)this._context.bezierCurveTo(r[0][o],i[0][o],r[1][o],i[1][o],t[u],n[u]);(this._line||0!==this._line&&1===e)&&this._context.closePath(),this._line=1-this._line,this._x=this._y=null},point:function(t,n){this._x.push(+t),this._y.push(+n)}},ee.prototype={areaStart:function(){this._line=0},areaEnd:function(){this._line=NaN},lineStart:function(){this._x=this._y=NaN,this._point=0},lineEnd:function(){0<this._t&&this._t<1&&2===this._point&&this._context.lineTo(this._x,this._y),(this._line||0!==this._line&&1===this._point)&&this._context.closePath(),this._line>=0&&(this._t=1-this._t,this._line=1-this._line)},point:function(t,n){switch(t=+t,n=+n,this._point){case 0:this._point=1,this._line?this._context.lineTo(t,n):this._context.moveTo(t,n);break;case 1:this._point=2;default:if(this._t<=0)this._context.lineTo(this._x,n),this._context.lineTo(t,n);else{var e=this._x*(1-this._t)+t*this._t;this._context.lineTo(e,this._y),this._context.lineTo(e,n)}}this._x=t,this._y=n}};var Gv=Array.prototype.slice,Jv=.7,Qv=1/Jv,Kv=/^#([0-9a-f]{3})$/,t_=/^#([0-9a-f]{6})$/,n_=/^rgb\(\s*([-+]?\d+)\s*,\s*([-+]?\d+)\s*,\s*([-+]?\d+)\s*\)$/,e_=/^rgb\(\s*([-+]?\d+(?:\.\d+)?)%\s*,\s*([-+]?\d+(?:\.\d+)?)%\s*,\s*([-+]?\d+(?:\.\d+)?)%\s*\)$/,r_=/^rgba\(\s*([-+]?\d+)\s*,\s*([-+]?\d+)\s*,\s*([-+]?\d+)\s*,\s*([-+]?\d+(?:\.\d+)?)\s*\)$/,i_=/^rgba\(\s*([-+]?\d+(?:\.\d+)?)%\s*,\s*([-+]?\d+(?:\.\d+)?)%\s*,\s*([-+]?\d+(?:\.\d+)?)%\s*,\s*([-+]?\d+(?:\.\d+)?)\s*\)$/,o_=/^hsl\(\s*([-+]?\d+(?:\.\d+)?)\s*,\s*([-+]?\d+(?:\.\d+)?)%\s*,\s*([-+]?\d+(?:\.\d+)?)%\s*\)$/,u_=/^hsla\(\s*([-+]?\d+(?:\.\d+)?)\s*,\s*([-+]?\d+(?:\.\d+)?)%\s*,\s*([-+]?\d+(?:\.\d+)?)%\s*,\s*([-+]?\d+(?:\.\d+)?)\s*\)$/,a_={aliceblue:15792383,antiquewhite:16444375,aqua:65535,aquamarine:8388564,azure:15794175,beige:16119260,bisque:16770244,black:0,blanchedalmond:16772045,blue:255,blueviolet:9055202,brown:10824234,burlywood:14596231,cadetblue:6266528,chartreuse:8388352,chocolate:13789470,coral:16744272,cornflowerblue:6591981,cornsilk:16775388,crimson:14423100,cyan:65535,darkblue:139,darkcyan:35723,darkgoldenrod:12092939,darkgray:11119017,darkgreen:25600,darkgrey:11119017,darkkhaki:12433259,darkmagenta:9109643,darkolivegreen:5597999,darkorange:16747520,darkorchid:10040012,darkred:9109504,darksalmon:15308410,darkseagreen:9419919,darkslateblue:4734347,darkslategray:3100495,darkslategrey:3100495,darkturquoise:52945,darkviolet:9699539,deeppink:16716947,deepskyblue:49151,dimgray:6908265,dimgrey:6908265,dodgerblue:2003199,firebrick:11674146,floralwhite:16775920,forestgreen:2263842,fuchsia:16711935,gainsboro:14474460,ghostwhite:16316671,gold:16766720,goldenrod:14329120,gray:8421504,green:32768,greenyellow:11403055,grey:8421504,honeydew:15794160,hotpink:16738740,indianred:13458524,indigo:4915330,ivory:16777200,khaki:15787660,lavender:15132410,lavenderblush:16773365,lawngreen:8190976,lemonchiffon:16775885,lightblue:11393254,lightcoral:15761536,lightcyan:14745599,lightgoldenrodyellow:16448210,lightgray:13882323,lightgreen:9498256,lightgrey:13882323,lightpink:16758465,lightsalmon:16752762,lightseagreen:2142890,lightskyblue:8900346,lightslategray:7833753,lightslategrey:7833753,lightsteelblue:11584734,lightyellow:16777184,lime:65280,limegreen:3329330,linen:16445670,magenta:16711935,maroon:8388608,mediumaquamarine:6737322,mediumblue:205,mediumorchid:12211667,mediumpurple:9662683,mediumseagreen:3978097,mediumslateblue:8087790,mediumspringgreen:64154,mediumturquoise:4772300,mediumvioletred:13047173,midnightblue:1644912,mintcream:16121850,mistyrose:16770273,moccasin:16770229,navajowhite:16768685,navy:128,oldlace:16643558,olive:8421376,olivedrab:7048739,orange:16753920,orangered:16729344,orchid:14315734,palegoldenrod:15657130,palegreen:10025880,paleturquoise:11529966,palevioletred:14381203,papayawhip:16773077,peachpuff:16767673,peru:13468991,pink:16761035,plum:14524637,powderblue:11591910,purple:8388736,rebeccapurple:6697881,red:16711680,rosybrown:12357519,royalblue:4286945,saddlebrown:9127187,salmon:16416882,sandybrown:16032864,seagreen:3050327,seashell:16774638,sienna:10506797,silver:12632256,skyblue:8900331,slateblue:6970061,slategray:7372944,slategrey:7372944,snow:16775930,springgreen:65407,steelblue:4620980,tan:13808780,teal:32896,thistle:14204888,tomato:16737095,turquoise:4251856,violet:15631086,wheat:16113331,white:16777215,whitesmoke:16119285,yellow:16776960,yellowgreen:10145074};ge(xe,be,{displayable:function(){return this.rgb().displayable()},toString:function(){return this.rgb()+""}}),ge(Se,ke,me(xe,{brighter:function(t){return t=null==t?Qv:Math.pow(Qv,t),new Se(this.r*t,this.g*t,this.b*t,this.opacity)},darker:function(t){return t=null==t?Jv:Math.pow(Jv,t),new Se(this.r*t,this.g*t,this.b*t,this.opacity)},rgb:function(){return this},displayable:function(){return 0<=this.r&&this.r<=255&&0<=this.g&&this.g<=255&&0<=this.b&&this.b<=255&&0<=this.opacity&&this.opacity<=1},toString:function(){var t=this.opacity;return t=isNaN(t)?1:Math.max(0,Math.min(1,t)),(1===t?"rgb(":"rgba(")+Math.max(0,Math.min(255,Math.round(this.r)||0))+", "+Math.max(0,Math.min(255,Math.round(this.g)||0))+", "+Math.max(0,Math.min(255,Math.round(this.b)||0))+(1===t?")":", "+t+")")}})),ge(Ce,Ee,me(xe,{brighter:function(t){return t=null==t?Qv:Math.pow(Qv,t),new Ce(this.h,this.s,this.l*t,this.opacity)},darker:function(t){return t=null==t?Jv:Math.pow(Jv,t),new Ce(this.h,this.s,this.l*t,this.opacity)},rgb:function(){var t=this.h%360+360*(this.h<0),n=isNaN(t)||isNaN(this.s)?0:this.s,e=this.l,r=e+(e<.5?e:1-e)*n,i=2*e-r;return new Se(ze(t>=240?t-240:t+120,i,r),ze(t,i,r),ze(t<120?t+240:t-120,i,r),this.opacity)},displayable:function(){return(0<=this.s&&this.s<=1||isNaN(this.s))&&0<=this.l&&this.l<=1&&0<=this.opacity&&this.opacity<=1}}));var c_=Math.PI/180,s_=180/Math.PI,f_=18,l_=.95047,h_=1,p_=1.08883,d_=4/29,v_=6/29,__=3*v_*v_,y_=v_*v_*v_;ge(Le,qe,me(xe,{brighter:function(t){return new Le(this.l+f_*(null==t?1:t),this.a,this.b,this.opacity)},darker:function(t){return new Le(this.l-f_*(null==t?1:t),this.a,this.b,this.opacity)},rgb:function(){var t=(this.l+16)/116,n=isNaN(this.a)?t:t+this.a/500,e=isNaN(this.b)?t:t-this.b/200;return t=h_*Ue(t),n=l_*Ue(n),e=p_*Ue(e),new Se(De(3.2404542*n-1.5371385*t-.4985314*e),De(-.969266*n+1.8760108*t+.041556*e),De(.0556434*n-.2040259*t+1.0572252*e),this.opacity)}})),ge(Ye,Ie,me(xe,{brighter:function(t){return new Ye(this.h,this.c,this.l+f_*(null==t?1:t),this.opacity)},darker:function(t){return new Ye(this.h,this.c,this.l-f_*(null==t?1:t),this.opacity)},rgb:function(){return Pe(this).rgb()}}));var g_=-.14861,m_=1.78277,x_=-.29227,b_=-.90649,w_=1.97294,M_=w_*b_,T_=w_*m_,k_=m_*x_-b_*g_;ge(He,je,me(xe,{brighter:function(t){return t=null==t?Qv:Math.pow(Qv,t),new He(this.h,this.s,this.l*t,this.opacity)},darker:function(t){return t=null==t?Jv:Math.pow(Jv,t),new He(this.h,this.s,this.l*t,this.opacity)},rgb:function(){var t=isNaN(this.h)?0:(this.h+120)*c_,n=+this.l,e=isNaN(this.s)?0:this.s*n*(1-n),r=Math.cos(t),i=Math.sin(t);return new Se(255*(n+e*(g_*r+m_*i)),255*(n+e*(x_*r+b_*i)),255*(n+e*(w_*r)),this.opacity)}}));var S_,N_,A_,E_,C_=function t(n){function e(t,n){var e=r((t=ke(t)).r,(n=ke(n)).r),i=r(t.g,n.g),o=r(t.b,n.b),u=r(t.opacity,n.opacity);return function(n){return t.r=e(n),t.g=i(n),t.b=o(n),t.opacity=u(n),t+""}}var r=Qe(n);return e.gamma=t,e}(1),z_=tr(Ve),P_=tr(We),q_=/[-+]?(?:\d+\.?\d*|\.?\d+)(?:[eE][-+]?\d+)?/g,L_=new RegExp(q_.source,"g"),R_=180/Math.PI,U_={translateX:0,translateY:0,rotate:0,skewX:0,scaleX:1,scaleY:1},D_=pr(lr,"px, ","px)","deg)"),O_=pr(hr,", ",")",")"),F_=Math.SQRT2,I_=2,Y_=4,B_=1e-12,j_=gr(Je),H_=gr(Ke),X_=xr(Je),V_=xr(Ke),W_=br(Je),$_=br(Ke),Z_={value:function(){}};Tr.prototype=Mr.prototype={constructor:Tr,on:function(t,n){var e,r=this._,i=kr(t+"",r),o=-1,u=i.length;{if(!(arguments.length<2)){if(null!=n&&"function"!=typeof n)throw new Error("invalid callback: "+n);for(;++o<u;)if(e=(t=i[o]).type)r[e]=Nr(r[e],t.name,n);else if(null==n)for(e in r)r[e]=Nr(r[e],t.name,null);return this}for(;++o<u;)if((e=(t=i[o]).type)&&(e=Sr(r[e],t.name)))return e}},copy:function(){var t={},n=this._;for(var e in n)t[e]=n[e].slice();return new Tr(t)},call:function(t,n){if((e=arguments.length-2)>0)for(var e,r,i=new Array(e),o=0;o<e;++o)i[o]=arguments[o+2];if(!this._.hasOwnProperty(t))throw new Error("unknown type: "+t);for(r=this._[t],o=0,e=r.length;o<e;++o)r[o].value.apply(n,i)},apply:function(t,n,e){if(!this._.hasOwnProperty(t))throw new Error("unknown type: "+t);for(var r=this._[t],i=0,o=r.length;i<o;++i)r[i].value.apply(n,e)}};var G_,J_,Q_=zr(","),K_=Q_.parse,ty=Q_.parseRows,ny=Q_.format,ey=Q_.formatRows,ry=zr("\t"),iy=ry.parse,oy=ry.parseRows,uy=ry.format,ay=ry.formatRows,cy=Rr("text/html",function(t){return document.createRange().createContextualFragment(t.responseText)}),sy=Rr("application/json",function(t){return JSON.parse(t.responseText)}),fy=Rr("text/plain",function(t){return t.responseText}),ly=Rr("application/xml",function(t){var n=t.responseXML;if(!n)throw new Error("parse error");return n}),hy=Ur("text/csv",K_),py=Ur("text/tab-separated-values",iy),dy=0,vy=0,_y=0,yy=1e3,gy=0,my=0,xy=0,by="object"==typeof performance&&performance.now?performance:Date,wy="function"==typeof requestAnimationFrame?by===Date?function(t){requestAnimationFrame(function(){t(by.now())})}:requestAnimationFrame:function(t){setTimeout(t,17)};Ir.prototype=Yr.prototype={constructor:Ir,restart:function(t,n,e){if("function"!=typeof t)throw new TypeError("callback is not a function");e=(null==e?Or():+e)+(null==n?0:+n),this._next||J_===this||(J_?J_._next=this:G_=this,J_=this),this._call=t,this._time=e,Vr()},stop:function(){this._call&&(this._call=null,this._time=1/0,Vr())}};var My=new Date,Ty=new Date,ky=Zr(function(){},function(t,n){t.setTime(+t+n)},function(t,n){return n-t});ky.every=function(t){return t=Math.floor(t),isFinite(t)&&t>0?t>1?Zr(function(n){n.setTime(Math.floor(n/t)*t)},function(n,e){n.setTime(+n+e*t)},function(n,e){return(e-n)/t}):ky:null};var Sy=ky.range,Ny=1e3,Ay=6e4,Ey=36e5,Cy=864e5,zy=6048e5,Py=Zr(function(t){t.setTime(Math.floor(t/Ny)*Ny)},function(t,n){t.setTime(+t+n*Ny)},function(t,n){return(n-t)/Ny},function(t){return t.getUTCSeconds()}),qy=Py.range,Ly=Zr(function(t){t.setTime(Math.floor(t/Ay)*Ay)},function(t,n){t.setTime(+t+n*Ay)},function(t,n){return(n-t)/Ay},function(t){return t.getMinutes()}),Ry=Ly.range,Uy=Zr(function(t){var n=t.getTimezoneOffset()*Ay%Ey;n<0&&(n+=Ey),t.setTime(Math.floor((+t-n)/Ey)*Ey+n)},function(t,n){t.setTime(+t+n*Ey)},function(t,n){return(n-t)/Ey},function(t){return t.getHours()}),Dy=Uy.range,Oy=Zr(function(t){t.setHours(0,0,0,0)},function(t,n){t.setDate(t.getDate()+n)},function(t,n){return(n-t-(n.getTimezoneOffset()-t.getTimezoneOffset())*Ay)/Cy},function(t){return t.getDate()-1}),Fy=Oy.range,Iy=Gr(0),Yy=Gr(1),By=Gr(2),jy=Gr(3),Hy=Gr(4),Xy=Gr(5),Vy=Gr(6),Wy=Iy.range,$y=Yy.range,Zy=By.range,Gy=jy.range,Jy=Hy.range,Qy=Xy.range,Ky=Vy.range,tg=Zr(function(t){t.setDate(1),t.setHours(0,0,0,0)},function(t,n){t.setMonth(t.getMonth()+n)},function(t,n){return n.getMonth()-t.getMonth()+12*(n.getFullYear()-t.getFullYear())},function(t){return t.getMonth()}),ng=tg.range,eg=Zr(function(t){t.setMonth(0,1),t.setHours(0,0,0,0)},function(t,n){t.setFullYear(t.getFullYear()+n)},function(t,n){return n.getFullYear()-t.getFullYear()},function(t){return t.getFullYear()});eg.every=function(t){return isFinite(t=Math.floor(t))&&t>0?Zr(function(n){n.setFullYear(Math.floor(n.getFullYear()/t)*t),n.setMonth(0,1),n.setHours(0,0,0,0)},function(n,e){n.setFullYear(n.getFullYear()+e*t)}):null};var rg=eg.range,ig=Zr(function(t){t.setUTCSeconds(0,0)},function(t,n){t.setTime(+t+n*Ay)},function(t,n){return(n-t)/Ay},function(t){return t.getUTCMinutes()}),og=ig.range,ug=Zr(function(t){t.setUTCMinutes(0,0,0)},function(t,n){t.setTime(+t+n*Ey)},function(t,n){return(n-t)/Ey},function(t){return t.getUTCHours()}),ag=ug.range,cg=Zr(function(t){t.setUTCHours(0,0,0,0)},function(t,n){t.setUTCDate(t.getUTCDate()+n)},function(t,n){return(n-t)/Cy},function(t){return t.getUTCDate()-1}),sg=cg.range,fg=Jr(0),lg=Jr(1),hg=Jr(2),pg=Jr(3),dg=Jr(4),vg=Jr(5),_g=Jr(6),yg=fg.range,gg=lg.range,mg=hg.range,xg=pg.range,bg=dg.range,wg=vg.range,Mg=_g.range,Tg=Zr(function(t){t.setUTCDate(1),t.setUTCHours(0,0,0,0)},function(t,n){t.setUTCMonth(t.getUTCMonth()+n)},function(t,n){return n.getUTCMonth()-t.getUTCMonth()+12*(n.getUTCFullYear()-t.getUTCFullYear())},function(t){return t.getUTCMonth()}),kg=Tg.range,Sg=Zr(function(t){t.setUTCMonth(0,1),t.setUTCHours(0,0,0,0)},function(t,n){t.setUTCFullYear(t.getUTCFullYear()+n)},function(t,n){return n.getUTCFullYear()-t.getUTCFullYear()},function(t){return t.getUTCFullYear()});Sg.every=function(t){return isFinite(t=Math.floor(t))&&t>0?Zr(function(n){n.setUTCFullYear(Math.floor(n.getUTCFullYear()/t)*t),n.setUTCMonth(0,1),n.setUTCHours(0,0,0,0)},function(n,e){n.setUTCFullYear(n.getUTCFullYear()+e*t)}):null};var Ng,Ag=Sg.range,Eg={"":ni,"%":function(t,n){return(100*t).toFixed(n)},b:function(t){return Math.round(t).toString(2)},c:function(t){return t+""},d:function(t){return Math.round(t).toString(10)},e:function(t,n){return t.toExponential(n)},f:function(t,n){return t.toFixed(n)},g:function(t,n){return t.toPrecision(n)},o:function(t){return Math.round(t).toString(8)},p:function(t,n){return ri(100*t,n)},r:ri,s:ei,X:function(t){return Math.round(t).toString(16).toUpperCase()},x:function(t){return Math.round(t).toString(16)}},Cg=/^(?:(.)?([<>=^]))?([+\-\( ])?([$#])?(0)?(\d+)?(,)?(\.\d+)?([a-z%])?$/i;oi.prototype.toString=function(){return this.fill+this.align+this.sign+this.symbol+(this.zero?"0":"")+(null==this.width?"":Math.max(1,0|this.width))+(this.comma?",":"")+(null==this.precision?"":"."+Math.max(0,0|this.precision))+this.type};var zg,Pg=["y","z","a","f","p","n","µ","m","","k","M","G","T","P","E","Z","Y"];ci({decimal:".",thousands:",",grouping:[3],currency:["$",""]});var qg,Lg={"-":"",_:" ",0:"0"},Rg=/^\s*\d+/,Ug=/^%/,Dg=/[\\\^\$\*\+\?\|\[\]\(\)\.\{\}]/g;so({dateTime:"%x, %X",date:"%-m/%-d/%Y",time:"%-I:%M:%S %p",periods:["AM","PM"],days:["Sunday","Monday","Tuesday","Wednesday","Thursday","Friday","Saturday"],shortDays:["Sun","Mon","Tue","Wed","Thu","Fri","Sat"],months:["January","February","March","April","May","June","July","August","September","October","November","December"],shortMonths:["Jan","Feb","Mar","Apr","May","Jun","Jul","Aug","Sep","Oct","Nov","Dec"]});var Og="%Y-%m-%dT%H:%M:%S.%LZ",Fg=Date.prototype.toISOString?fo:t.utcFormat(Og),Ig=+new Date("2000-01-01T00:00:00.000Z")?lo:t.utcParse(Og),Yg=Array.prototype,Bg=Yg.map,jg=Yg.slice,Hg={name:"implicit"},Xg=[0,1],Vg=1e3,Wg=60*Vg,$g=60*Wg,Zg=24*$g,Gg=7*Zg,Jg=30*Zg,Qg=365*Zg,Kg=Zo("1f77b4ff7f0e2ca02cd627289467bd8c564be377c27f7f7fbcbd2217becf"),tm=Zo("393b795254a36b6ecf9c9ede6379398ca252b5cf6bcedb9c8c6d31bd9e39e7ba52e7cb94843c39ad494ad6616be7969c7b4173a55194ce6dbdde9ed6"),nm=Zo("3182bd6baed69ecae1c6dbefe6550dfd8d3cfdae6bfdd0a231a35474c476a1d99bc7e9c0756bb19e9ac8bcbddcdadaeb636363969696bdbdbdd9d9d9"),em=Zo("1f77b4aec7e8ff7f0effbb782ca02c98df8ad62728ff98969467bdc5b0d58c564bc49c94e377c2f7b6d27f7f7fc7c7c7bcbd22dbdb8d17becf9edae5"),rm=$_(je(300,.5,0),je(-240,.5,1)),im=$_(je(-100,.75,.35),je(80,1.5,.8)),om=$_(je(260,.75,.35),je(80,1.5,.8)),um=je(),am=Jo(Zo("44015444025645045745055946075a46085c460a5d460b5e470d60470e6147106347116447136548146748166848176948186a481a6c481b6d481c6e481d6f481f70482071482173482374482475482576482677482878482979472a7a472c7a472d7b472e7c472f7d46307e46327e46337f463480453581453781453882443983443a83443b84433d84433e85423f854240864241864142874144874045884046883f47883f48893e49893e4a893e4c8a3d4d8a3d4e8a3c4f8a3c508b3b518b3b528b3a538b3a548c39558c39568c38588c38598c375a8c375b8d365c8d365d8d355e8d355f8d34608d34618d33628d33638d32648e32658e31668e31678e31688e30698e306a8e2f6b8e2f6c8e2e6d8e2e6e8e2e6f8e2d708e2d718e2c718e2c728e2c738e2b748e2b758e2a768e2a778e2a788e29798e297a8e297b8e287c8e287d8e277e8e277f8e27808e26818e26828e26828e25838e25848e25858e24868e24878e23888e23898e238a8d228b8d228c8d228d8d218e8d218f8d21908d21918c20928c20928c20938c1f948c1f958b1f968b1f978b1f988b1f998a1f9a8a1e9b8a1e9c891e9d891f9e891f9f881fa0881fa1881fa1871fa28720a38620a48621a58521a68522a78522a88423a98324aa8325ab8225ac8226ad8127ad8128ae8029af7f2ab07f2cb17e2db27d2eb37c2fb47c31b57b32b67a34b67935b77937b87838b9773aba763bbb753dbc743fbc7340bd7242be7144bf7046c06f48c16e4ac16d4cc26c4ec36b50c46a52c56954c56856c66758c7655ac8645cc8635ec96260ca6063cb5f65cb5e67cc5c69cd5b6ccd5a6ece5870cf5773d05675d05477d1537ad1517cd2507fd34e81d34d84d44b86d54989d5488bd6468ed64590d74393d74195d84098d83e9bd93c9dd93ba0da39a2da37a5db36a8db34aadc32addc30b0dd2fb2dd2db5de2bb8de29bade28bddf26c0df25c2df23c5e021c8e020cae11fcde11dd0e11cd2e21bd5e21ad8e219dae319dde318dfe318e2e418e5e419e7e419eae51aece51befe51cf1e51df4e61ef6e620f8e621fbe723fde725")),cm=Jo(Zo("00000401000501010601010802010902020b02020d03030f03031204041405041606051806051a07061c08071e0907200a08220b09240c09260d0a290e0b2b100b2d110c2f120d31130d34140e36150e38160f3b180f3d19103f1a10421c10441d11471e114920114b21114e22115024125325125527125829115a2a115c2c115f2d11612f116331116533106734106936106b38106c390f6e3b0f703d0f713f0f72400f74420f75440f764510774710784910784a10794c117a4e117b4f127b51127c52137c54137d56147d57157e59157e5a167e5c167f5d177f5f187f601880621980641a80651a80671b80681c816a1c816b1d816d1d816e1e81701f81721f817320817521817621817822817922827b23827c23827e24828025828125818326818426818627818827818928818b29818c29818e2a81902a81912b81932b80942c80962c80982d80992d809b2e7f9c2e7f9e2f7fa02f7fa1307ea3307ea5317ea6317da8327daa337dab337cad347cae347bb0357bb2357bb3367ab5367ab73779b83779ba3878bc3978bd3977bf3a77c03a76c23b75c43c75c53c74c73d73c83e73ca3e72cc3f71cd4071cf4070d0416fd2426fd3436ed5446dd6456cd8456cd9466bdb476adc4869de4968df4a68e04c67e24d66e34e65e44f64e55064e75263e85362e95462ea5661eb5760ec5860ed5a5fee5b5eef5d5ef05f5ef1605df2625df2645cf3655cf4675cf4695cf56b5cf66c5cf66e5cf7705cf7725cf8745cf8765cf9785df9795df97b5dfa7d5efa7f5efa815ffb835ffb8560fb8761fc8961fc8a62fc8c63fc8e64fc9065fd9266fd9467fd9668fd9869fd9a6afd9b6bfe9d6cfe9f6dfea16efea36ffea571fea772fea973feaa74feac76feae77feb078feb27afeb47bfeb67cfeb77efeb97ffebb81febd82febf84fec185fec287fec488fec68afec88cfeca8dfecc8ffecd90fecf92fed194fed395fed597fed799fed89afdda9cfddc9efddea0fde0a1fde2a3fde3a5fde5a7fde7a9fde9aafdebacfcecaefceeb0fcf0b2fcf2b4fcf4b6fcf6b8fcf7b9fcf9bbfcfbbdfcfdbf")),sm=Jo(Zo("00000401000501010601010802010a02020c02020e03021004031204031405041706041907051b08051d09061f0a07220b07240c08260d08290e092b10092d110a30120a32140b34150b37160b39180c3c190c3e1b0c411c0c431e0c451f0c48210c4a230c4c240c4f260c51280b53290b552b0b572d0b592f0a5b310a5c320a5e340a5f3609613809623909633b09643d09653e0966400a67420a68440a68450a69470b6a490b6a4a0c6b4c0c6b4d0d6c4f0d6c510e6c520e6d540f6d550f6d57106e59106e5a116e5c126e5d126e5f136e61136e62146e64156e65156e67166e69166e6a176e6c186e6d186e6f196e71196e721a6e741a6e751b6e771c6d781c6d7a1d6d7c1d6d7d1e6d7f1e6c801f6c82206c84206b85216b87216b88226a8a226a8c23698d23698f24699025689225689326679526679727669827669a28659b29649d29649f2a63a02a63a22b62a32c61a52c60a62d60a82e5fa92e5eab2f5ead305dae305cb0315bb1325ab3325ab43359b63458b73557b93556ba3655bc3754bd3853bf3952c03a51c13a50c33b4fc43c4ec63d4dc73e4cc83f4bca404acb4149cc4248ce4347cf4446d04545d24644d34743d44842d54a41d74b3fd84c3ed94d3dda4e3cdb503bdd513ade5238df5337e05536e15635e25734e35933e45a31e55c30e65d2fe75e2ee8602de9612bea632aeb6429eb6628ec6726ed6925ee6a24ef6c23ef6e21f06f20f1711ff1731df2741cf3761bf37819f47918f57b17f57d15f67e14f68013f78212f78410f8850ff8870ef8890cf98b0bf98c0af98e09fa9008fa9207fa9407fb9606fb9706fb9906fb9b06fb9d07fc9f07fca108fca309fca50afca60cfca80dfcaa0ffcac11fcae12fcb014fcb216fcb418fbb61afbb81dfbba1ffbbc21fbbe23fac026fac228fac42afac62df9c72ff9c932f9cb35f8cd37f8cf3af7d13df7d340f6d543f6d746f5d949f5db4cf4dd4ff4df53f4e156f3e35af3e55df2e661f2e865f2ea69f1ec6df1ed71f1ef75f1f179f2f27df2f482f3f586f3f68af4f88ef5f992f6fa96f8fb9af9fc9dfafda1fcffa4")),fm=Jo(Zo("0d088710078813078916078a19068c1b068d1d068e20068f2206902406912605912805922a05932c05942e05952f059631059733059735049837049938049a3a049a3c049b3e049c3f049c41049d43039e44039e46039f48039f4903a04b03a14c02a14e02a25002a25102a35302a35502a45601a45801a45901a55b01a55c01a65e01a66001a66100a76300a76400a76600a76700a86900a86a00a86c00a86e00a86f00a87100a87201a87401a87501a87701a87801a87a02a87b02a87d03a87e03a88004a88104a78305a78405a78606a68707a68808a68a09a58b0aa58d0ba58e0ca48f0da4910ea3920fa39410a29511a19613a19814a099159f9a169f9c179e9d189d9e199da01a9ca11b9ba21d9aa31e9aa51f99a62098a72197a82296aa2395ab2494ac2694ad2793ae2892b02991b12a90b22b8fb32c8eb42e8db52f8cb6308bb7318ab83289ba3388bb3488bc3587bd3786be3885bf3984c03a83c13b82c23c81c33d80c43e7fc5407ec6417dc7427cc8437bc9447aca457acb4679cc4778cc4977cd4a76ce4b75cf4c74d04d73d14e72d24f71d35171d45270d5536fd5546ed6556dd7566cd8576bd9586ada5a6ada5b69db5c68dc5d67dd5e66de5f65de6164df6263e06363e16462e26561e26660e3685fe4695ee56a5de56b5de66c5ce76e5be76f5ae87059e97158e97257ea7457eb7556eb7655ec7754ed7953ed7a52ee7b51ef7c51ef7e50f07f4ff0804ef1814df1834cf2844bf3854bf3874af48849f48948f58b47f58c46f68d45f68f44f79044f79143f79342f89441f89540f9973ff9983ef99a3efa9b3dfa9c3cfa9e3bfb9f3afba139fba238fca338fca537fca636fca835fca934fdab33fdac33fdae32fdaf31fdb130fdb22ffdb42ffdb52efeb72dfeb82cfeba2cfebb2bfebd2afebe2afec029fdc229fdc328fdc527fdc627fdc827fdca26fdcb26fccd25fcce25fcd025fcd225fbd324fbd524fbd724fad824fada24f9dc24f9dd25f8df25f8e125f7e225f7e425f6e626f6e826f5e926f5eb27f4ed27f3ee27f3f027f2f227f1f426f1f525f0f724f0f921")),lm="http://www.w3.org/1999/xhtml",hm={
svg:"http://www.w3.org/2000/svg",xhtml:lm,xlink:"http://www.w3.org/1999/xlink",xml:"http://www.w3.org/XML/1998/namespace",xmlns:"http://www.w3.org/2000/xmlns/"},pm=0;iu.prototype=ru.prototype={constructor:iu,get:function(t){for(var n=this._;!(n in t);)if(!(t=t.parentNode))return;return t[n]},set:function(t,n){return t[this._]=n},remove:function(t){return this._ in t&&delete t[this._]},toString:function(){return this._}};var dm=function(t){return function(){return this.matches(t)}};if("undefined"!=typeof document){var vm=document.documentElement;if(!vm.matches){var _m=vm.webkitMatchesSelector||vm.msMatchesSelector||vm.mozMatchesSelector||vm.oMatchesSelector;dm=function(t){return function(){return _m.call(this,t)}}}}var ym=dm,gm={};if(t.event=null,"undefined"!=typeof document){var mm=document.documentElement;"onmouseenter"in mm||(gm={mouseenter:"mouseover",mouseleave:"mouseout"})}Tu.prototype={constructor:Tu,appendChild:function(t){return this._parent.insertBefore(t,this._next)},insertBefore:function(t,n){return this._parent.insertBefore(t,n)},querySelector:function(t){return this._parent.querySelector(t)},querySelectorAll:function(t){return this._parent.querySelectorAll(t)}};var xm="$";ia.prototype={add:function(t){var n=this._names.indexOf(t);n<0&&(this._names.push(t),this._node.setAttribute("class",this._names.join(" ")))},remove:function(t){var n=this._names.indexOf(t);n>=0&&(this._names.splice(n,1),this._node.setAttribute("class",this._names.join(" ")))},contains:function(t){return this._names.indexOf(t)>=0}};var bm=[null];qa.prototype=La.prototype={constructor:qa,select:yu,selectAll:xu,filter:bu,data:Au,enter:Mu,exit:Eu,merge:Cu,order:zu,sort:Pu,call:Lu,nodes:Ru,node:Uu,size:Du,empty:Ou,each:Fu,attr:Vu,style:Ju,property:na,classed:fa,text:da,html:ga,raise:xa,lower:wa,append:Ma,insert:ka,remove:Na,datum:Aa,on:fu,dispatch:Pa};var wm=Mr("start","end","interrupt"),Mm=[],Tm=0,km=1,Sm=2,Nm=3,Am=4,Em=5,Cm=La.prototype.constructor,zm=0,Pm=La.prototype;Uc.prototype=Dc.prototype={constructor:Uc,select:wc,selectAll:Mc,filter:vc,merge:_c,selection:Tc,transition:Rc,call:Pm.call,nodes:Pm.nodes,node:Pm.node,size:Pm.size,empty:Pm.empty,each:Pm.each,on:mc,attr:rc,attrTween:uc,style:Ec,styleTween:zc,text:Lc,remove:bc,tween:$a,delay:sc,duration:hc,ease:dc};var qm={time:null,delay:0,duration:250,ease:et};La.prototype.interrupt=Xa,La.prototype.transition=Ic;var Lm=[null],Rm=Array.prototype.slice,Um=1,Dm=2,Om=3,Fm=4,Im=1e-6;ws.prototype=ys.prototype={constructor:ws,each:us,eachAfter:cs,eachBefore:as,sum:ss,sort:fs,path:ls,ancestors:ps,descendants:ds,leaves:vs,links:_s,copy:gs};var Ym="$",Bm={depth:-1},jm={};of.prototype=Object.create(ws.prototype);var Hm=(1+Math.sqrt(5))/2,Xm=function t(n){function e(t,e,r,i,o){sf(n,t,e,r,i,o)}return e.ratio=function(n){return t((n=+n)>1?n:1)},e}(Hm),Vm=function t(n){function e(t,e,r,i,o){if((u=t._squarify)&&u.ratio===n)for(var u,a,c,s,f,l=-1,h=u.length,p=t.value;++l<h;){for(a=u[l],c=a.children,s=a.value=0,f=c.length;s<f;++s)a.value+=c[s].value;a.dice?Ws(a,e,r,i,r+=(o-r)*a.value/p):cf(a,e,r,e+=(i-e)*a.value/p,o),p-=a.value}else t._squarify=u=sf(n,t,e,r,i,o),u.ratio=n}return e.ratio=function(n){return t((n=+n)>1?n:1)},e}(Hm),Wm=10,$m=Math.PI*(3-Math.sqrt(5));Pf.prototype.on=function(){var t=this._.on.apply(this._,arguments);return t===this._?this:t},If.prototype={constructor:If,insert:function(t,n){var e,r,i;if(t){if(n.P=t,n.N=t.N,t.N&&(t.N.P=n),t.N=n,t.R){for(t=t.R;t.L;)t=t.L;t.L=n}else t.R=n;e=t}else this._?(t=Hf(this._),n.P=null,n.N=t,t.P=t.L=n,e=t):(n.P=n.N=null,this._=n,e=null);for(n.L=n.R=null,n.U=e,n.C=!0,t=n;e&&e.C;)r=e.U,e===r.L?(i=r.R,i&&i.C?(e.C=i.C=!1,r.C=!0,t=r):(t===e.R&&(Bf(this,e),t=e,e=t.U),e.C=!1,r.C=!0,jf(this,r))):(i=r.L,i&&i.C?(e.C=i.C=!1,r.C=!0,t=r):(t===e.L&&(jf(this,e),t=e,e=t.U),e.C=!1,r.C=!0,Bf(this,r))),e=t.U;this._.C=!1},remove:function(t){t.N&&(t.N.P=t.P),t.P&&(t.P.N=t.N),t.N=t.P=null;var n,e,r,i=t.U,o=t.L,u=t.R;if(e=o?u?Hf(u):o:u,i?i.L===t?i.L=e:i.R=e:this._=e,o&&u?(r=e.C,e.C=t.C,e.L=o,o.U=e,e!==u?(i=e.U,e.U=t.U,t=e.R,i.L=t,e.R=u,u.U=e):(e.U=i,i=e,t=e.R)):(r=t.C,t=e),t&&(t.U=i),!r){if(t&&t.C)return void(t.C=!1);do{if(t===this._)break;if(t===i.L){if(n=i.R,n.C&&(n.C=!1,i.C=!0,Bf(this,i),n=i.R),n.L&&n.L.C||n.R&&n.R.C){n.R&&n.R.C||(n.L.C=!1,n.C=!0,jf(this,n),n=i.R),n.C=i.C,i.C=n.R.C=!1,Bf(this,i),t=this._;break}}else if(n=i.L,n.C&&(n.C=!1,i.C=!0,jf(this,i),n=i.L),n.L&&n.L.C||n.R&&n.R.C){n.L&&n.L.C||(n.R.C=!1,n.C=!0,Bf(this,n),n=i.L),n.C=i.C,i.C=n.L.C=!1,jf(this,i),t=this._;break}n.C=!0,t=i,i=i.U}while(!t.C);t&&(t.C=!1)}}};var Zm,Gm,Jm,Qm,Km,tx=[],nx=[],ex=1e-6,rx=1e-12;vl.prototype={constructor:vl,polygons:function(){var t=this.edges;return this.cells.map(function(n){var e=n.halfedges.map(function(e){return Kf(n,t[e])});return e.data=n.site.data,e})},triangles:function(){var t=[],n=this.edges;return this.cells.forEach(function(e,r){for(var i,o=e.site,u=e.halfedges,a=-1,c=u.length,s=n[u[c-1]],f=s.left===o?s.right:s.left;++a<c;)i=f,s=n[u[a]],f=s.left===o?s.right:s.left,r<i.index&&r<f.index&&pl(o,i,f)<0&&t.push([o.data,i.data,f.data])}),t},links:function(){return this.edges.filter(function(t){return t.right}).map(function(t){return{source:t.left.data,target:t.right.data}})}},ml.prototype={constructor:ml,scale:function(t){return 1===t?this:new ml(this.k*t,this.x,this.y)},translate:function(t,n){return 0===t&0===n?this:new ml(this.k,this.x+this.k*t,this.y+this.k*n)},apply:function(t){return[t[0]*this.k+this.x,t[1]*this.k+this.y]},applyX:function(t){return t*this.k+this.x},applyY:function(t){return t*this.k+this.y},invert:function(t){return[(t[0]-this.x)/this.k,(t[1]-this.y)/this.k]},invertX:function(t){return(t-this.x)/this.k},invertY:function(t){return(t-this.y)/this.k},rescaleX:function(t){return t.copy().domain(t.range().map(this.invertX,this).map(t.invert,t))},rescaleY:function(t){return t.copy().domain(t.range().map(this.invertY,this).map(t.invert,t))},toString:function(){return"translate("+this.x+","+this.y+") scale("+this.k+")"}};var ix=new ml(1,0,0);xl.prototype=ml.prototype;var ox={name:"drag"},ux={name:"space"},ax={name:"handle"},cx={name:"center"},sx={name:"x",handles:["e","w"].map(zl),input:function(t,n){return t&&[[t[0],n[0][1]],[t[1],n[1][1]]]},output:function(t){return t&&[t[0][0],t[1][0]]}},fx={name:"y",handles:["n","s"].map(zl),input:function(t,n){return t&&[[n[0][0],t[0]],[n[1][0],t[1]]]},output:function(t){return t&&[t[0][1],t[1][1]]}},lx={name:"xy",handles:["n","e","s","w","nw","ne","se","sw"].map(zl),input:function(t){return t},output:function(t){return t}},hx={overlay:"crosshair",selection:"move",n:"ns-resize",e:"ew-resize",s:"ns-resize",w:"ew-resize",nw:"nwse-resize",ne:"nesw-resize",se:"nwse-resize",sw:"nesw-resize"},px={e:"w",w:"e",nw:"ne",ne:"nw",se:"sw",sw:"se"},dx={n:"s",s:"n",nw:"sw",ne:"se",se:"ne",sw:"nw"},vx={overlay:1,selection:1,n:null,e:1,s:null,w:-1,nw:-1,ne:1,se:1,sw:-1},_x={overlay:1,selection:1,n:-1,e:null,s:1,w:null,nw:-1,ne:-1,se:1,sw:1},yx=Math.cos,gx=Math.sin,mx=Math.PI,xx=mx/2,bx=2*mx,wx=Math.max,Mx=Array.prototype.slice;Jl.prototype={constructor:Jl,reset:function(){this.s=this.t=0},add:function(t){Ql(nb,t,this.t),Ql(this,nb.s,this.s),this.s?this.t+=nb.t:this.s=nb.t},valueOf:function(){return this.s}};var Tx,kx,Sx,Nx,Ax,Ex,Cx,zx,Px,qx,Lx,Rx,Ux,Dx,Ox,Fx,Ix,Yx,Bx,jx,Hx,Xx,Vx,Wx,$x,Zx,Gx,Jx,Qx,Kx,tb,nb=new Jl,eb=1e-6,rb=1e-12,ib=Math.PI,ob=ib/2,ub=ib/4,ab=2*ib,cb=180/ib,sb=ib/180,fb=Math.abs,lb=Math.atan,hb=Math.atan2,pb=Math.cos,db=Math.ceil,vb=Math.exp,_b=Math.log,yb=Math.pow,gb=Math.sin,mb=Math.sign||function(t){return t>0?1:t<0?-1:0},xb=Math.sqrt,bb=Math.tan,wb={Feature:function(t,n){rh(t.geometry,n)},FeatureCollection:function(t,n){for(var e=t.features,r=-1,i=e.length;++r<i;)rh(e[r].geometry,n)}},Mb={Sphere:function(t,n){n.sphere()},Point:function(t,n){t=t.coordinates,n.point(t[0],t[1],t[2])},MultiPoint:function(t,n){for(var e=t.coordinates,r=-1,i=e.length;++r<i;)t=e[r],n.point(t[0],t[1],t[2])},LineString:function(t,n){ih(t.coordinates,n,0)},MultiLineString:function(t,n){for(var e=t.coordinates,r=-1,i=e.length;++r<i;)ih(e[r],n,0)},Polygon:function(t,n){oh(t.coordinates,n)},MultiPolygon:function(t,n){for(var e=t.coordinates,r=-1,i=e.length;++r<i;)oh(e[r],n)},GeometryCollection:function(t,n){for(var e=t.geometries,r=-1,i=e.length;++r<i;)rh(e[r],n)}},Tb=Gl(),kb=Gl(),Sb={point:eh,lineStart:eh,lineEnd:eh,polygonStart:function(){Tb.reset(),Sb.lineStart=ah,Sb.lineEnd=ch},polygonEnd:function(){var t=+Tb;kb.add(t<0?ab+t:t),this.lineStart=this.lineEnd=this.point=eh},sphere:function(){kb.add(ab)}},Nb=Gl(),Ab={point:mh,lineStart:bh,lineEnd:wh,polygonStart:function(){Ab.point=Mh,Ab.lineStart=Th,Ab.lineEnd=kh,Nb.reset(),Sb.polygonStart()},polygonEnd:function(){Sb.polygonEnd(),Ab.point=mh,Ab.lineStart=bh,Ab.lineEnd=wh,Tb<0?(Ex=-(zx=180),Cx=-(Px=90)):Nb>eb?Px=90:Nb<-eb&&(Cx=-90),Ox[0]=Ex,Ox[1]=zx}},Eb={sphere:eh,point:Ch,lineStart:Ph,lineEnd:Rh,polygonStart:function(){Eb.lineStart=Uh,Eb.lineEnd=Dh},polygonEnd:function(){Eb.lineStart=Ph,Eb.lineEnd=Rh}};jh.invert=jh;var Cb,zb,Pb,qb,Lb,Rb,Ub,Db,Ob,Fb,Ib,Yb=1e9,Bb=-Yb,jb=Gl(),Hb={sphere:eh,point:eh,lineStart:up,lineEnd:eh,polygonStart:eh,polygonEnd:eh},Xb=[null,null],Vb={type:"LineString",coordinates:Xb},Wb=Gl(),$b=Gl(),Zb={point:eh,lineStart:eh,lineEnd:eh,polygonStart:function(){Zb.lineStart=yp,Zb.lineEnd=xp},polygonEnd:function(){Zb.lineStart=Zb.lineEnd=Zb.point=eh,Wb.add(fb($b)),$b.reset()},result:function(){var t=Wb/2;return Wb.reset(),t}},Gb=1/0,Jb=Gb,Qb=-Gb,Kb=Qb,tw={point:bp,lineStart:eh,lineEnd:eh,polygonStart:eh,polygonEnd:eh,result:function(){var t=[[Gb,Jb],[Qb,Kb]];return Qb=Kb=-(Jb=Gb=1/0),t}},nw=0,ew=0,rw=0,iw=0,ow=0,uw=0,aw=0,cw=0,sw=0,fw={point:wp,lineStart:Mp,lineEnd:Sp,polygonStart:function(){fw.lineStart=Np,fw.lineEnd=Ap},polygonEnd:function(){fw.point=wp,fw.lineStart=Mp,fw.lineEnd=Sp},result:function(){var t=sw?[aw/sw,cw/sw]:uw?[iw/uw,ow/uw]:rw?[nw/rw,ew/rw]:[NaN,NaN];return nw=ew=rw=iw=ow=uw=aw=cw=sw=0,t}},lw=Gl(),hw=Up(function(){return!0},Fp,Yp,[-ib,-ob]);Xp.prototype={point:function(t,n){this.stream.point(t,n)},sphere:function(){this.stream.sphere()},lineStart:function(){this.stream.lineStart()},lineEnd:function(){this.stream.lineEnd()},polygonStart:function(){this.stream.polygonStart()},polygonEnd:function(){this.stream.polygonEnd()}};var pw=16,dw=pb(30*sb),vw=Hp({point:function(t,n){this.stream.point(t*sb,n*sb)}}),_w=ud(function(t){return xb(2/(1+t))});_w.invert=ad(function(t){return 2*th(t/2)});var yw=ud(function(t){return(t=Kl(t))&&t/gb(t)});yw.invert=ad(function(t){return t}),fd.invert=function(t,n){return[t,2*lb(vb(n))-ob]},_d.invert=_d,xd.invert=ad(lb),wd.invert=ad(th),Td.invert=ad(function(t){return 2*lb(t)}),Sd.invert=function(t,n){return[-n,2*lb(vb(t))-ob]},t.version=Ad,t.bisect=Cd,t.bisectRight=Cd,t.bisectLeft=zd,t.ascending=n,t.bisector=e,t.descending=i,t.deviation=a,t.extent=c,t.histogram=v,t.thresholdFreedmanDiaconis=y,t.thresholdScott=g,t.thresholdSturges=d,t.max=m,t.mean=x,t.median=b,t.merge=w,t.min=M,t.pairs=T,t.permute=k,t.quantile=_,t.range=l,t.scan=S,t.shuffle=N,t.sum=A,t.ticks=h,t.tickStep=p,t.transpose=E,t.variance=u,t.zip=z,t.entries=j,t.keys=Y,t.values=B,t.map=q,t.set=I,t.nest=L,t.randomUniform=H,t.randomNormal=X,t.randomLogNormal=V,t.randomBates=$,t.randomIrwinHall=W,t.randomExponential=Z,t.easeLinear=G,t.easeQuad=K,t.easeQuadIn=J,t.easeQuadOut=Q,t.easeQuadInOut=K,t.easeCubic=et,t.easeCubicIn=tt,t.easeCubicOut=nt,t.easeCubicInOut=et,t.easePoly=jd,t.easePolyIn=Yd,t.easePolyOut=Bd,t.easePolyInOut=jd,t.easeSin=ot,t.easeSinIn=rt,t.easeSinOut=it,t.easeSinInOut=ot,t.easeExp=ct,t.easeExpIn=ut,t.easeExpOut=at,t.easeExpInOut=ct,t.easeCircle=lt,t.easeCircleIn=st,t.easeCircleOut=ft,t.easeCircleInOut=lt,t.easeBounce=pt,t.easeBounceIn=ht,t.easeBounceOut=pt,t.easeBounceInOut=dt,t.easeBack=ov,t.easeBackIn=rv,t.easeBackOut=iv,t.easeBackInOut=ov,t.easeElastic=fv,t.easeElasticIn=sv,t.easeElasticOut=fv,t.easeElasticInOut=lv,t.polygonArea=vt,t.polygonCentroid=_t,t.polygonHull=xt,t.polygonContains=bt,t.polygonLength=wt,t.path=Tt,t.quadtree=jt,t.queue=Qt,t.arc=sn,t.area=vn,t.line=dn,t.pie=gn,t.radialArea=Mn,t.radialLine=wn,t.symbol=Tn,t.symbols=Bv,t.symbolCircle=Tv,t.symbolCross=kv,t.symbolDiamond=Av,t.symbolSquare=Lv,t.symbolStar=qv,t.symbolTriangle=Uv,t.symbolWye=Yv,t.curveBasisClosed=Cn,t.curveBasisOpen=Pn,t.curveBasis=An,t.curveBundle=jv,t.curveCardinalClosed=Xv,t.curveCardinalOpen=Vv,t.curveCardinal=Hv,t.curveCatmullRomClosed=$v,t.curveCatmullRomOpen=Zv,t.curveCatmullRom=Wv,t.curveLinearClosed=jn,t.curveLinear=ln,t.curveMonotoneX=Jn,t.curveMonotoneY=Qn,t.curveNatural=ne,t.curveStep=re,t.curveStepAfter=oe,t.curveStepBefore=ie,t.stack=se,t.stackOffsetExpand=fe,t.stackOffsetNone=ue,t.stackOffsetSilhouette=le,t.stackOffsetWiggle=he,t.stackOrderAscending=pe,t.stackOrderDescending=ve,t.stackOrderInsideOut=_e,t.stackOrderNone=ae,t.stackOrderReverse=ye,t.color=be,t.rgb=ke,t.hsl=Ee,t.lab=qe,t.hcl=Ie,t.cubehelix=je,t.interpolate=cr,t.interpolateArray=nr,t.interpolateDate=er,t.interpolateNumber=rr,t.interpolateObject=ir,t.interpolateRound=sr,t.interpolateString=ar,t.interpolateTransformCss=D_,t.interpolateTransformSvg=O_,t.interpolateZoom=yr,t.interpolateRgb=C_,t.interpolateRgbBasis=z_,t.interpolateRgbBasisClosed=P_,t.interpolateHsl=j_,t.interpolateHslLong=H_,t.interpolateLab=mr,t.interpolateHcl=X_,t.interpolateHclLong=V_,t.interpolateCubehelix=W_,t.interpolateCubehelixLong=$_,t.interpolateBasis=Ve,t.interpolateBasisClosed=We,t.quantize=wr,t.dispatch=Mr,t.dsvFormat=zr,t.csvParse=K_,t.csvParseRows=ty,t.csvFormat=ny,t.csvFormatRows=ey,t.tsvParse=iy,t.tsvParseRows=oy,t.tsvFormat=uy,t.tsvFormatRows=ay,t.request=Pr,t.html=cy,t.json=sy,t.text=fy,t.xml=ly,t.csv=hy,t.tsv=py,t.now=Or,t.timer=Yr,t.timerFlush=Br,t.timeout=Wr,t.interval=$r,t.timeInterval=Zr,t.timeMillisecond=ky,t.timeMilliseconds=Sy,t.timeSecond=Py,t.timeSeconds=qy,t.timeMinute=Ly,t.timeMinutes=Ry,t.timeHour=Uy,t.timeHours=Dy,t.timeDay=Oy,t.timeDays=Fy,t.timeWeek=Iy;t.timeWeeks=Wy;t.timeSunday=Iy,t.timeSundays=Wy,t.timeMonday=Yy,t.timeMondays=$y,t.timeTuesday=By,t.timeTuesdays=Zy,t.timeWednesday=jy,t.timeWednesdays=Gy,t.timeThursday=Hy,t.timeThursdays=Jy,t.timeFriday=Xy,t.timeFridays=Qy,t.timeSaturday=Vy,t.timeSaturdays=Ky,t.timeMonth=tg,t.timeMonths=ng,t.timeYear=eg,t.timeYears=rg,t.utcMillisecond=ky,t.utcMilliseconds=Sy,t.utcSecond=Py,t.utcSeconds=qy,t.utcMinute=ig,t.utcMinutes=og,t.utcHour=ug,t.utcHours=ag,t.utcDay=cg,t.utcDays=sg,t.utcWeek=fg,t.utcWeeks=yg,t.utcSunday=fg,t.utcSundays=yg,t.utcMonday=lg,t.utcMondays=gg,t.utcTuesday=hg,t.utcTuesdays=mg,t.utcWednesday=pg,t.utcWednesdays=xg,t.utcThursday=dg,t.utcThursdays=bg,t.utcFriday=vg,t.utcFridays=wg,t.utcSaturday=_g,t.utcSaturdays=Mg,t.utcMonth=Tg,t.utcMonths=kg,t.utcYear=Sg,t.utcYears=Ag,t.formatLocale=ai,t.formatDefaultLocale=ci,t.formatSpecifier=ii,t.precisionFixed=si,t.precisionPrefix=fi,t.precisionRound=li,t.isoFormat=Fg,t.isoParse=Ig,t.timeFormatLocale=vi,t.timeFormatDefaultLocale=so,t.scaleBand=po,t.scalePoint=_o,t.scaleIdentity=Eo,t.scaleLinear=Ao,t.scaleLog=Do,t.scaleOrdinal=ho,t.scaleImplicit=Hg,t.scalePow=Fo,t.scaleSqrt=Io,t.scaleQuantile=Yo,t.scaleQuantize=Bo,t.scaleThreshold=jo,t.scaleTime=Wo,t.scaleUtc=$o,t.schemeCategory10=Kg,t.schemeCategory20b=tm,t.schemeCategory20c=nm,t.schemeCategory20=em,t.scaleSequential=Qo,t.interpolateCubehelixDefault=rm,t.interpolateRainbow=Go,t.interpolateWarm=im,t.interpolateCool=om,t.interpolateViridis=am,t.interpolateMagma=cm,t.interpolateInferno=sm,t.interpolatePlasma=fm,t.creator=eu,t.customEvent=lu,t.local=ru,t.matcher=ym,t.mouse=du,t.namespace=Ko,t.namespaces=hm,t.select=Ra,t.selectAll=Ua,t.selection=La,t.selector=_u,t.selectorAll=mu,t.touch=Da,t.touches=Oa,t.window=Wu,t.active=Yc,t.interrupt=Ha,t.transition=Dc,t.axisTop=$c,t.axisRight=Zc,t.axisBottom=Gc,t.axisLeft=Jc,t.cluster=os,t.hierarchy=ys,t.pack=Bs,t.packSiblings=Us,t.packEnclose=ks,t.partition=$s,t.stratify=Js,t.tree=af,t.treemap=ff,t.treemapBinary=lf,t.treemapDice=Ws,t.treemapSlice=cf,t.treemapSliceDice=hf,t.treemapSquarify=Xm,t.treemapResquarify=Vm,t.forceCenter=pf,t.forceCollide=gf,t.forceLink=xf,t.forceManyBody=Tf,t.forceSimulation=Mf,t.forceX=kf,t.forceY=Sf,t.drag=Uf,t.dragDisable=Ef,t.dragEnable=Cf,t.voronoi=_l,t.zoom=Sl,t.zoomIdentity=ix,t.zoomTransform=xl,t.brush=Fl,t.brushX=Dl,t.brushY=Ol,t.brushSelection=Ul,t.chord=Bl,t.ribbon=Zl,t.geoAlbers=rd,t.geoAlbersUsa=od,t.geoArea=lh,t.geoAzimuthalEqualArea=cd,t.geoAzimuthalEqualAreaRaw=_w,t.geoAzimuthalEquidistant=sd,t.geoAzimuthalEquidistantRaw=yw,t.geoBounds=Eh,t.geoCentroid=Ih,t.geoCircle=Jh,t.geoClipExtent=op,t.geoConicConformal=vd,t.geoConicConformalRaw=dd,t.geoConicEqualArea=ed,t.geoConicEqualAreaRaw=nd,t.geoConicEquidistant=md,t.geoConicEquidistantRaw=gd,t.geoDistance=lp,t.geoEquirectangular=yd,t.geoEquirectangularRaw=_d,t.geoGnomonic=bd,t.geoGnomonicRaw=xd,t.geoGraticule=dp,t.geoInterpolate=vp,t.geoLength=fp,t.geoMercator=ld,t.geoMercatorRaw=fd,t.geoOrthographic=Md,t.geoOrthographicRaw=wd,t.geoPath=Lp,t.geoProjection=Qp,t.geoProjectionMutator=Kp,t.geoRotation=$h,t.geoStereographic=kd,t.geoStereographicRaw=Td,t.geoStream=uh,t.geoTransform=jp,t.geoTransverseMercator=Nd,t.geoTransverseMercatorRaw=Sd,Object.defineProperty(t,"__esModule",{value:!0})});
```
