## Supplementary material for "Comparative pathway enrichment analysis in gastrointestinal cell lines Caco-2, HT-29, HEPG2, and colon fibroblasts using a custom expression panel for tight-junction and cytoskeletal regulatory genes": Sup. Results 1: fastclick.js

```
/**
 * @preserve FastClick: polyfill to remove click delays on browsers with touch UIs.
 *
 * @version 1.0.2
 * @codingstandard ftlabs-jsv2
 * @copyright The Financial Times Limited [All Rights Reserved]
 * @license MIT License (see LICENSE.txt)
 */
function FastClick(a,b){"use strict";function c(a,b){return function(){return a.apply(b,arguments)}}var d;if(b=b||{},this.trackingClick=!1,this.trackingClickStart=0,this.targetElement=null,this.touchStartX=0,this.touchStartY=0,this.lastTouchIdentifier=0,this.touchBoundary=b.touchBoundary||10,this.layer=a,this.tapDelay=b.tapDelay||200,!FastClick.notNeeded(a)){for(var e=["onMouse","onClick","onTouchStart","onTouchMove","onTouchEnd","onTouchCancel"],f=this,g=0,h=e.length;h>g;g++)f[e[g]]=c(f[e[g]],f);deviceIsAndroid&&(a.addEventListener("mouseover",this.onMouse,!0),a.addEventListener("mousedown",this.onMouse,!0),a.addEventListener("mouseup",this.onMouse,!0)),a.addEventListener("click",this.onClick,!0),a.addEventListener("touchstart",this.onTouchStart,!1),a.addEventListener("touchmove",this.onTouchMove,!1),a.addEventListener("touchend",this.onTouchEnd,!1),a.addEventListener("touchcancel",this.onTouchCancel,!1),Event.prototype.stopImmediatePropagation||(a.removeEventListener=function(b,c,d){var e=Node.prototype.removeEventListener;"click"===b?e.call(a,b,c.hijacked||c,d):e.call(a,b,c,d)},a.addEventListener=function(b,c,d){var e=Node.prototype.addEventListener;"click"===b?e.call(a,b,c.hijacked||(c.hijacked=function(a){a.propagationStopped||c(a)}),d):e.call(a,b,c,d)}),"function"==typeof a.onclick&&(d=a.onclick,a.addEventListener("click",function(a){d(a)},!1),a.onclick=null)}}var deviceIsAndroid=navigator.userAgent.indexOf("Android")>0,deviceIsIOS=/iP(ad|hone|od)/.test(navigator.userAgent),deviceIsIOS4=deviceIsIOS&&/OS 4_\d(_\d)?/.test(navigator.userAgent),deviceIsIOSWithBadTarget=deviceIsIOS&&/OS ([6-9]|\d{2})_\d/.test(navigator.userAgent);FastClick.prototype.needsClick=function(a){"use strict";switch(a.nodeName.toLowerCase()){case"button":case"select":case"textarea":if(a.disabled)return!0;break;case"input":if(deviceIsIOS&&"file"===a.type||a.disabled)return!0;break;case"label":case"video":return!0}return/\bneedsclick\b/.test(a.className)},FastClick.prototype.needsFocus=function(a){"use strict";switch(a.nodeName.toLowerCase()){case"textarea":return!0;case"select":return!deviceIsAndroid;case"input":switch(a.type){case"button":case"checkbox":case"file":case"image":case"radio":case"submit":return!1}return!a.disabled&&!a.readOnly;default:return/\bneedsfocus\b/.test(a.className)}},FastClick.prototype.sendClick=function(a,b){"use strict";var c,d;document.activeElement&&document.activeElement!==a&&document.activeElement.blur(),d=b.changedTouches[0],c=document.createEvent("MouseEvents"),c.initMouseEvent(this.determineEventType(a),!0,!0,window,1,d.screenX,d.screenY,d.clientX,d.clientY,!1,!1,!1,!1,0,null),c.forwardedTouchEvent=!0,a.dispatchEvent(c)},FastClick.prototype.determineEventType=function(a){"use strict";return deviceIsAndroid&&"select"===a.tagName.toLowerCase()?"mousedown":"click"},FastClick.prototype.focus=function(a){"use strict";var b;deviceIsIOS&&a.setSelectionRange&&0!==a.type.indexOf("date")&&"time"!==a.type?(b=a.value.length,a.setSelectionRange(b,b)):a.focus()},FastClick.prototype.updateScrollParent=function(a){"use strict";var b,c;if(b=a.fastClickScrollParent,!b||!b.contains(a)){c=a;do{if(c.scrollHeight>c.offsetHeight){b=c,a.fastClickScrollParent=c;break}c=c.parentElement}while(c)}b&&(b.fastClickLastScrollTop=b.scrollTop)},FastClick.prototype.getTargetElementFromEventTarget=function(a){"use strict";return a.nodeType===Node.TEXT_NODE?a.parentNode:a},FastClick.prototype.onTouchStart=function(a){"use strict";var b,c,d;if(a.targetTouches.length>1)return!0;if(b=this.getTargetElementFromEventTarget(a.target),c=a.targetTouches[0],deviceIsIOS){if(d=window.getSelection(),d.rangeCount&&!d.isCollapsed)return!0;if(!deviceIsIOS4){if(c.identifier===this.lastTouchIdentifier)return a.preventDefault(),!1;this.lastTouchIdentifier=c.identifier,this.updateScrollParent(b)}}return this.trackingClick=!0,this.trackingClickStart=a.timeStamp,this.targetElement=b,this.touchStartX=c.pageX,this.touchStartY=c.pageY,a.timeStamp-this.lastClickTime<this.tapDelay&&a.preventDefault(),!0},FastClick.prototype.touchHasMoved=function(a){"use strict";var b=a.changedTouches[0],c=this.touchBoundary;return Math.abs(b.pageX-this.touchStartX)>c||Math.abs(b.pageY-this.touchStartY)>c?!0:!1},FastClick.prototype.onTouchMove=function(a){"use strict";return this.trackingClick?((this.targetElement!==this.getTargetElementFromEventTarget(a.target)||this.touchHasMoved(a))&&(this.trackingClick=!1,this.targetElement=null),!0):!0},FastClick.prototype.findControl=function(a){"use strict";return void 0!==a.control?a.control:a.htmlFor?document.getElementById(a.htmlFor):a.querySelector("button, input:not([type=hidden]), keygen, meter, output, progress, select, textarea")},FastClick.prototype.onTouchEnd=function(a){"use strict";var b,c,d,e,f,g=this.targetElement;if(!this.trackingClick)return!0;if(a.timeStamp-this.lastClickTime<this.tapDelay)return this.cancelNextClick=!0,!0;if(this.cancelNextClick=!1,this.lastClickTime=a.timeStamp,c=this.trackingClickStart,this.trackingClick=!1,this.trackingClickStart=0,deviceIsIOSWithBadTarget&&(f=a.changedTouches[0],g=document.elementFromPoint(f.pageX-window.pageXOffset,f.pageY-window.pageYOffset)||g,g.fastClickScrollParent=this.targetElement.fastClickScrollParent),d=g.tagName.toLowerCase(),"label"===d){if(b=this.findControl(g)){if(this.focus(g),deviceIsAndroid)return!1;g=b}}else if(this.needsFocus(g))return a.timeStamp-c>100||deviceIsIOS&&window.top!==window&&"input"===d?(this.targetElement=null,!1):(this.focus(g),this.sendClick(g,a),deviceIsIOS&&"select"===d||(this.targetElement=null,a.preventDefault()),!1);return deviceIsIOS&&!deviceIsIOS4&&(e=g.fastClickScrollParent,e&&e.fastClickLastScrollTop!==e.scrollTop)?!0:(this.needsClick(g)||(a.preventDefault(),this.sendClick(g,a)),!1)},FastClick.prototype.onTouchCancel=function(){"use strict";this.trackingClick=!1,this.targetElement=null},FastClick.prototype.onMouse=function(a){"use strict";return this.targetElement?a.forwardedTouchEvent?!0:a.cancelable&&(!this.needsClick(this.targetElement)||this.cancelNextClick)?(a.stopImmediatePropagation?a.stopImmediatePropagation():a.propagationStopped=!0,a.stopPropagation(),a.preventDefault(),!1):!0:!0},FastClick.prototype.onClick=function(a){"use strict";var b;return this.trackingClick?(this.targetElement=null,this.trackingClick=!1,!0):"submit"===a.target.type&&0===a.detail?!0:(b=this.onMouse(a),b||(this.targetElement=null),b)},FastClick.prototype.destroy=function(){"use strict";var a=this.layer;deviceIsAndroid&&(a.removeEventListener("mouseover",this.onMouse,!0),a.removeEventListener("mousedown",this.onMouse,!0),a.removeEventListener("mouseup",this.onMouse,!0)),a.removeEventListener("click",this.onClick,!0),a.removeEventListener("touchstart",this.onTouchStart,!1),a.removeEventListener("touchmove",this.onTouchMove,!1),a.removeEventListener("touchend",this.onTouchEnd,!1),a.removeEventListener("touchcancel",this.onTouchCancel,!1)},FastClick.notNeeded=function(a){"use strict";var b,c;if("undefined"==typeof window.ontouchstart)return!0;if(c=+(/Chrome\/([0-9]+)/.exec(navigator.userAgent)||[,0])[1]){if(!deviceIsAndroid)return!0;if(b=document.querySelector("meta[name=viewport]")){if(-1!==b.content.indexOf("user-scalable=no"))return!0;if(c>31&&document.documentElement.scrollWidth<=window.outerWidth)return!0}}return"none"===a.style.msTouchAction?!0:!1},FastClick.attach=function(a,b){"use strict";return new FastClick(a,b)},"function"==typeof define&&"object"==typeof define.amd&&define.amd?define(function(){"use strict";return FastClick}):"undefined"!=typeof module&&module.exports?(module.exports=FastClick.attach,module.exports.FastClick=FastClick):window.FastClick=FastClick;
```
