## Supplementary material for "Comparative pathway enrichment analysis in gastrointestinal cell lines Caco-2, HT-29, HEPG2, and colon fibroblasts using a custom expression panel for tight-junction and cytoskeletal regulatory genes": Sup. Results 1: foundation.abide.js

```
;(function ($, window, document, undefined) {
  'use strict';

  Foundation.libs.abide = {
    name : 'abide',

    version : '5.3.0',

    settings : {
      live_validate : true,
      focus_on_invalid : true,
      error_labels: true, // labels with a for="inputId" will recieve an `error` class
      timeout : 1000,
      patterns : {
        alpha: /^[a-zA-Z]+$/,
        alpha_numeric : /^[a-zA-Z0-9]+$/,
        integer: /^[-+]?\d+$/,
        number: /^[-+]?\d*(?:[\.\,]\d+)?$/,

        // amex, visa, diners
        card : /^(?:4[0-9]{12}(?:[0-9]{3})?|5[1-5][0-9]{14}|6(?:011|5[0-9][0-9])[0-9]{12}|3[47][0-9]{13}|3(?:0[0-5]|[68][0-9])[0-9]{11}|(?:2131|1800|35\d{3})\d{11})$/,
        cvv : /^([0-9]){3,4}$/,

        // http://www.whatwg.org/specs/web-apps/current-work/multipage/states-of-the-type-attribute.html#valid-e-mail-address
        email : /^[a-zA-Z0-9.!#$%&'*+\/=?^_`{|}~-]+@[a-zA-Z0-9](?:[a-zA-Z0-9-]{0,61}[a-zA-Z0-9])?(?:\.[a-zA-Z0-9](?:[a-zA-Z0-9-]{0,61}[a-zA-Z0-9])?)*$/,

        url: /^(https?|ftp|file|ssh):\/\/(((([a-zA-Z]|\d|-|\.|_|~|[\u00A0-\uD7FF\uF900-\uFDCF\uFDF0-\uFFEF])|(%[\da-f]{2})|[!\$&'\(\)\*\+,;=]|:)*@)?(((\d|[1-9]\d|1\d\d|2[0-4]\d|25[0-5])\.(\d|[1-9]\d|1\d\d|2[0-4]\d|25[0-5])\.(\d|[1-9]\d|1\d\d|2[0-4]\d|25[0-5])\.(\d|[1-9]\d|1\d\d|2[0-4]\d|25[0-5]))|((([a-zA-Z]|\d|[\u00A0-\uD7FF\uF900-\uFDCF\uFDF0-\uFFEF])|(([a-zA-Z]|\d|[\u00A0-\uD7FF\uF900-\uFDCF\uFDF0-\uFFEF])([a-zA-Z]|\d|-|\.|_|~|[\u00A0-\uD7FF\uF900-\uFDCF\uFDF0-\uFFEF])*([a-zA-Z]|\d|[\u00A0-\uD7FF\uF900-\uFDCF\uFDF0-\uFFEF])))\.)+(([a-zA-Z]|[\u00A0-\uD7FF\uF900-\uFDCF\uFDF0-\uFFEF])|(([a-zA-Z]|[\u00A0-\uD7FF\uF900-\uFDCF\uFDF0-\uFFEF])([a-zA-Z]|\d|-|\.|_|~|[\u00A0-\uD7FF\uF900-\uFDCF\uFDF0-\uFFEF])*([a-zA-Z]|[\u00A0-\uD7FF\uF900-\uFDCF\uFDF0-\uFFEF])))\.?)(:\d*)?)(\/((([a-zA-Z]|\d|-|\.|_|~|[\u00A0-\uD7FF\uF900-\uFDCF\uFDF0-\uFFEF])|(%[\da-f]{2})|[!\$&'\(\)\*\+,;=]|:|@)+(\/(([a-zA-Z]|\d|-|\.|_|~|[\u00A0-\uD7FF\uF900-\uFDCF\uFDF0-\uFFEF])|(%[\da-f]{2})|[!\$&'\(\)\*\+,;=]|:|@)*)*)?)?(\?((([a-zA-Z]|\d|-|\.|_|~|[\u00A0-\uD7FF\uF900-\uFDCF\uFDF0-\uFFEF])|(%[\da-f]{2})|[!\$&'\(\)\*\+,;=]|:|@)|[\uE000-\uF8FF]|\/|\?)*)?(\#((([a-zA-Z]|\d|-|\.|_|~|[\u00A0-\uD7FF\uF900-\uFDCF\uFDF0-\uFFEF])|(%[\da-f]{2})|[!\$&'\(\)\*\+,;=]|:|@)|\/|\?)*)?$/,
        // abc.de
        domain: /^([a-zA-Z0-9]([a-zA-Z0-9\-]{0,61}[a-zA-Z0-9])?\.)+[a-zA-Z]{2,6}$/,

        datetime: /^([0-2][0-9]{3})\-([0-1][0-9])\-([0-3][0-9])T([0-5][0-9])\:([0-5][0-9])\:([0-5][0-9])(Z|([\-\+]([0-1][0-9])\:00))$/,
        // YYYY-MM-DD
        date: /(?:19|20)[0-9]{2}-(?:(?:0[1-9]|1[0-2])-(?:0[1-9]|1[0-9]|2[0-9])|(?:(?!02)(?:0[1-9]|1[0-2])-(?:30))|(?:(?:0[13578]|1[02])-31))$/,
        // HH:MM:SS
        time : /^(0[0-9]|1[0-9]|2[0-3])(:[0-5][0-9]){2}$/,
        dateISO: /^\d{4}[\/\-]\d{1,2}[\/\-]\d{1,2}$/,
        // MM/DD/YYYY
        month_day_year : /^(0[1-9]|1[012])[- \/.](0[1-9]|[12][0-9]|3[01])[- \/.]\d{4}$/,

        // #FFF or #FFFFFF
        color: /^#?([a-fA-F0-9]{6}|[a-fA-F0-9]{3})$/
      },
      validators : {
        equalTo: function(el, required, parent) {
          var from  = document.getElementById(el.getAttribute(this.add_namespace('data-equalto'))).value,
              to    = el.value,
              valid = (from === to);

          return valid;
        }
      }
    },

    timer : null,

    init : function (scope, method, options) {
      this.bindings(method, options);
    },

    events : function (scope) {
      var self = this,
          form = self.S(scope).attr('novalidate', 'novalidate'),
          settings = form.data(this.attr_name(true) + '-init') || {};

      this.invalid_attr = this.add_namespace('data-invalid');

      form
        .off('.abide')
        .on('submit.fndtn.abide validate.fndtn.abide', function (e) {
          var is_ajax = /ajax/i.test(self.S(this).attr(self.attr_name()));
          return self.validate(self.S(this).find('input, textarea, select').get(), e, is_ajax);
        })
        .on('reset', function() {
          return self.reset($(this));
        })
        .find('input, textarea, select')
          .off('.abide')
          .on('blur.fndtn.abide change.fndtn.abide', function (e) {
            self.validate([this], e);
          })
          .on('keydown.fndtn.abide', function (e) {
            if (settings.live_validate === true) {
              clearTimeout(self.timer);
              self.timer = setTimeout(function () {
                self.validate([this], e);
              }.bind(this), settings.timeout);
            }
          });
    },

    reset : function (form) {
      form.removeAttr(this.invalid_attr);
      $(this.invalid_attr, form).removeAttr(this.invalid_attr);
      $('.error', form).not('small').removeClass('error');
    },

    validate : function (els, e, is_ajax) {
      var validations = this.parse_patterns(els),
          validation_count = validations.length,
          form = this.S(els[0]).closest('[data-' + this.attr_name(true) + ']'),
          settings = form.data(this.attr_name(true) + '-init') || {},
          submit_event = /submit/.test(e.type);

      form.trigger('validated').trigger('validated.fndtn.abide');
      // Has to count up to make sure the focus gets applied to the top error
      for (var i=0; i < validation_count; i++) {
        if (!validations[i] && (submit_event || is_ajax)) {
          if (settings.focus_on_invalid) els[i].focus();
          form.trigger('invalid').trigger('invalid.fndtn.abide');
          this.S(els[i]).closest('[data-' + this.attr_name(true) + ']').attr(this.invalid_attr, '');
          return false;
        }
      }

      if (submit_event || is_ajax) {
        form.trigger('valid').trigger('valid.fndtn.abide');
      }

      form.removeAttr(this.invalid_attr);

      if (is_ajax) return false;

      return true;
    },

    parse_patterns : function (els) {
      var i = els.length,
          el_patterns = [];

      while (i--) {
        el_patterns.push(this.pattern(els[i]));
      }

      return this.check_validation_and_apply_styles(el_patterns);
    },

    pattern : function (el) {
      var type = el.getAttribute('type'),
          required = typeof el.getAttribute('required') === 'string';

      var pattern = el.getAttribute('pattern') || '';

      if (this.settings.patterns.hasOwnProperty(pattern) && pattern.length > 0) {
        return [el, this.settings.patterns[pattern], required];
      } else if (pattern.length > 0) {
        return [el, new RegExp('^'+pattern+'$'), required];
      }

      if (this.settings.patterns.hasOwnProperty(type)) {
        return [el, this.settings.patterns[type], required];
      }

      pattern = /.*/;

      return [el, pattern, required];
    },

    check_validation_and_apply_styles : function (el_patterns) {
      var i = el_patterns.length,
          validations = [],
          form = this.S(el_patterns[0][0]).closest('[data-' + this.attr_name(true) + ']'),
          settings = form.data(this.attr_name(true) + '-init') || {};

      while (i--) {
        var el = el_patterns[i][0],
            required = el_patterns[i][2],
            value = el.value.trim(),
            direct_parent = this.S(el).parent(),
            validator = el.getAttribute(this.add_namespace('data-abide-validator')),
            is_radio = el.type === "radio",
            is_checkbox = el.type === "checkbox",
            label = this.S('label[for="' + el.getAttribute('id') + '"]'),
            valid_length = (required) ? (el.value.length > 0) : true;

        var parent, valid;

        // support old way to do equalTo validations
        if(el.getAttribute(this.add_namespace('data-equalto'))) { validator = "equalTo" }

        if (!direct_parent.is('label')) {
          parent = direct_parent;
        } else {
          parent = direct_parent.parent();
        }

        if (is_radio && required) {
          validations.push(this.valid_radio(el, required));
        } else if (is_checkbox && required) {
          validations.push(this.valid_checkbox(el, required));
        } else if (validator) {
          valid = this.settings.validators[validator].apply(this, [el, required, parent])
          validations.push(valid);

          if (valid) {
            this.S(el).removeAttr(this.invalid_attr);
            parent.removeClass('error');
          } else {
            this.S(el).attr(this.invalid_attr, '');
            parent.addClass('error');
          }

        } else {

          if (el_patterns[i][1].test(value) && valid_length ||
            !required && el.value.length < 1 || $(el).attr('disabled')) {
            this.S(el).removeAttr(this.invalid_attr);
            parent.removeClass('error');
            if (label.length > 0 && settings.error_labels) label.removeClass('error');

            validations.push(true);
            $(el).triggerHandler('valid');
          } else {
            this.S(el).attr(this.invalid_attr, '');
            parent.addClass('error');
            if (label.length > 0 && settings.error_labels) label.addClass('error');

            validations.push(false);
            $(el).triggerHandler('invalid');
          }
        }
      }

      return validations;
    },

    valid_checkbox : function(el, required) {
      var el = this.S(el),
          valid = (el.is(':checked') || !required);

      if (valid) {
        el.removeAttr(this.invalid_attr).parent().removeClass('error');
      } else {
        el.attr(this.invalid_attr, '').parent().addClass('error');
      }

      return valid;
    },

    valid_radio : function (el, required) {
      var name = el.getAttribute('name'),
          group = this.S(el).closest('[data-' + this.attr_name(true) + ']').find("[name='"+name+"']"),
          count = group.length,
          valid = false;

      // Has to count up to make sure the focus gets applied to the top error
      for (var i=0; i < count; i++) {
        if (group[i].checked) valid = true;
      }

      // Has to count up to make sure the focus gets applied to the top error
      for (var i=0; i < count; i++) {
        if (valid) {
          this.S(group[i]).removeAttr(this.invalid_attr).parent().removeClass('error');
        } else {
          this.S(group[i]).attr(this.invalid_attr, '').parent().addClass('error');
        }
      }

      return valid;
    },

    valid_equal: function(el, required, parent) {
      var from  = document.getElementById(el.getAttribute(this.add_namespace('data-equalto'))).value,
          to    = el.value,
          valid = (from === to);

      if (valid) {
        this.S(el).removeAttr(this.invalid_attr);
        parent.removeClass('error');
      } else {
        this.S(el).attr(this.invalid_attr, '');
        parent.addClass('error');
      }

      return valid;
    },

    valid_oneof: function(el, required, parent, doNotValidateOthers) {
      var el = this.S(el),
        others = this.S('[' + this.add_namespace('data-oneof') + ']'),
        valid = others.filter(':checked').length > 0;

      if (valid) {
        el.removeAttr(this.invalid_attr).parent().removeClass('error');
      } else {
        el.attr(this.invalid_attr, '').parent().addClass('error');
      }

      if (!doNotValidateOthers) {
        var _this = this;
        others.each(function() {
          _this.valid_oneof.call(_this, this, null, null, true);
        });
      }

      return valid;
    }
  };
}(jQuery, window, window.document));
```
