## Supplementary material for "Comparative pathway enrichment analysis in gastrointestinal cell lines Caco-2, HT-29, HEPG2, and colon fibroblasts using a custom expression panel for tight-junction and cytoskeletal regulatory genes": Sup. Results 1: foundation.accordion.js

```
;(function ($, window, document, undefined) {
  'use strict';

  Foundation.libs.accordion = {
    name : 'accordion',

    version : '5.3.0',

    settings : {
      active_class: 'active',
      multi_expand: false,
      toggleable: true,
      callback : function () {}
    },

    init : function (scope, method, options) {
      this.bindings(method, options);
    },

    events : function () {
      var self = this;
      var S = this.S;
      S(this.scope)
      .off('.fndtn.accordion')
      .on('click.fndtn.accordion', '[' + this.attr_name() + '] > dd > a', function (e) {
        var accordion = S(this).closest('[' + self.attr_name() + ']'),
            target = S('#' + this.href.split('#')[1]),
            siblings = S('dd > .content', accordion),
            aunts = $('dd', accordion),
            groupSelector = self.attr_name() + '=' + accordion.attr(self.attr_name()),
            settings = accordion.data(self.attr_name(true) + '-init'),
            active_content = S('dd > .content.' + settings.active_class, accordion);
        e.preventDefault();

        if (accordion.attr(self.attr_name())) {
          siblings = siblings.add('[' + groupSelector + '] dd > .content');
          aunts = aunts.add('[' + groupSelector + '] dd');
        }

        if (settings.toggleable && target.is(active_content)) {
          target.parent('dd').toggleClass(settings.active_class, false);
          target.toggleClass(settings.active_class, false);
          settings.callback(target);
          target.triggerHandler('toggled', [accordion]);
          accordion.triggerHandler('toggled', [target]);
          return;
        }

        if (!settings.multi_expand) {
          siblings.removeClass(settings.active_class);
          aunts.removeClass(settings.active_class);
        }

        target.addClass(settings.active_class).parent().addClass(settings.active_class);
        settings.callback(target);
        target.triggerHandler('toggled', [accordion]);
        accordion.triggerHandler('toggled', [target]);
      });
    },

    off : function () {},

    reflow : function () {}
  };
}(jQuery, window, window.document));
```
