## Supplementary material for "Comparative pathway enrichment analysis in gastrointestinal cell lines Caco-2, HT-29, HEPG2, and colon fibroblasts using a custom expression panel for tight-junction and cytoskeletal regulatory genes": Sup. Results 1: foundation.alert.js

```
;(function ($, window, document, undefined) {
  'use strict';

  Foundation.libs.alert = {
    name : 'alert',

    version : '5.3.0',

    settings : {
      callback: function (){}
    },

    init : function (scope, method, options) {
      this.bindings(method, options);
    },

    events : function () {
      var self = this,
          S = this.S;

      $(this.scope).off('.alert').on('click.fndtn.alert', '[' + this.attr_name() + '] a.close', function (e) {
          var alertBox = S(this).closest('[' + self.attr_name() + ']'),
              settings = alertBox.data(self.attr_name(true) + '-init') || self.settings;

        e.preventDefault();
        if (Modernizr.csstransitions) {
          alertBox.addClass("alert-close");
          alertBox.on('transitionend webkitTransitionEnd oTransitionEnd', function(e) {
            S(this).trigger('close').trigger('close.fndtn.alert').remove();
            settings.callback();
          });
        } else {
          alertBox.fadeOut(300, function () {
            S(this).trigger('close').trigger('close.fndtn.alert').remove();
            settings.callback();
          });
        }
      });
    },

    reflow : function () {}
  };
}(jQuery, window, window.document));
```
