## Supplementary material for "Comparative pathway enrichment analysis in gastrointestinal cell lines Caco-2, HT-29, HEPG2, and colon fibroblasts using a custom expression panel for tight-junction and cytoskeletal regulatory genes": Sup. Results 1: foundation.clearing.js

```
;(function ($, window, document, undefined) {
  'use strict';

  Foundation.libs.clearing = {
    name : 'clearing',

    version: '5.3.0',

    settings : {
      templates : {
        viewing : '<a href="#" class="clearing-close">&times;</a>' +
          '<div class="visible-img" style="display: none"><div class="clearing-touch-label"></div><img src="data:image/gif;base64,R0lGODlhAQABAAD/ACwAAAAAAQABAAACADs%3D" alt="" />' +
          '<p class="clearing-caption"></p><a href="#" class="clearing-main-prev"><span></span></a>' +
          '<a href="#" class="clearing-main-next"><span></span></a></div>'
      },

      // comma delimited list of selectors that, on click, will close clearing,
      // add 'div.clearing-blackout, div.visible-img' to close on background click
      close_selectors : '.clearing-close, div.clearing-blackout', 

      // Default to the entire li element.
      open_selectors : '',

      touch_label : '',

      // event initializers and locks
      init : false,
      locked : false
    },

    init : function (scope, method, options) {
      var self = this;
      Foundation.inherit(this, 'throttle image_loaded');

      this.bindings(method, options);

      if (self.S(this.scope).is('[' + this.attr_name() + ']')) {
        this.assemble(self.S('li', this.scope));
      } else {
        self.S('[' + this.attr_name() + ']', this.scope).each(function () {
          self.assemble(self.S('li', this));
        });
      }
    },

    events : function (scope) {
      var self = this,
          S = self.S,
          $scroll_container = $('.scroll-container');

      if ($scroll_container.length > 0) {
        this.scope = $scroll_container;
      }

      S(this.scope)
        .off('.clearing')
        .on('click.fndtn.clearing', 'ul[' + this.attr_name() + '] li ' + this.settings.open_selectors,
          function (e, current, target) {
            var current = current || S(this),
                target = target || current,
                next = current.next('li'),
                settings = current.closest('[' + self.attr_name() + ']').data(self.attr_name(true) + '-init'),
                image = S(e.target);

            e.preventDefault();

            if (!settings) {
              self.init();
              settings = current.closest('[' + self.attr_name() + ']').data(self.attr_name(true) + '-init');
            }

            // if clearing is open and the current image is
            // clicked, go to the next image in sequence
            if (target.hasClass('visible') &&
              current[0] === target[0] &&
              next.length > 0 && self.is_open(current)) {
              target = next;
              image = S('img', target);
            }

            // set current and target to the clicked li if not otherwise defined.
            self.open(image, current, target);
            self.update_paddles(target);
          })

        .on('click.fndtn.clearing', '.clearing-main-next',
          function (e) { self.nav(e, 'next') })
        .on('click.fndtn.clearing', '.clearing-main-prev',
          function (e) { self.nav(e, 'prev') })
        .on('click.fndtn.clearing', this.settings.close_selectors,
          function (e) { Foundation.libs.clearing.close(e, this) });

      $(document).on('keydown.fndtn.clearing',
          function (e) { self.keydown(e) });

      S(window).off('.clearing').on('resize.fndtn.clearing',
        function () { self.resize() });

      this.swipe_events(scope);
    },

    swipe_events : function (scope) {
      var self = this,
      S = self.S;

      S(this.scope)
        .on('touchstart.fndtn.clearing', '.visible-img', function(e) {
          if (!e.touches) { e = e.originalEvent; }
          var data = {
                start_page_x: e.touches[0].pageX,
                start_page_y: e.touches[0].pageY,
                start_time: (new Date()).getTime(),
                delta_x: 0,
                is_scrolling: undefined
              };

          S(this).data('swipe-transition', data);
          e.stopPropagation();
        })
        .on('touchmove.fndtn.clearing', '.visible-img', function(e) {
          if (!e.touches) { e = e.originalEvent; }
          // Ignore pinch/zoom events
          if(e.touches.length > 1 || e.scale && e.scale !== 1) return;

          var data = S(this).data('swipe-transition');

          if (typeof data === 'undefined') {
            data = {};
          }

          data.delta_x = e.touches[0].pageX - data.start_page_x;

          if (Foundation.rtl) {
            data.delta_x = -data.delta_x;
          }

          if (typeof data.is_scrolling === 'undefined') {
            data.is_scrolling = !!( data.is_scrolling || Math.abs(data.delta_x) < Math.abs(e.touches[0].pageY - data.start_page_y) );
          }

          if (!data.is_scrolling && !data.active) {
            e.preventDefault();
            var direction = (data.delta_x < 0) ? 'next' : 'prev';
            data.active = true;
            self.nav(e, direction);
          }
        })
        .on('touchend.fndtn.clearing', '.visible-img', function(e) {
          S(this).data('swipe-transition', {});
          e.stopPropagation();
        });
    },

    assemble : function ($li) {
      var $el = $li.parent();

      if ($el.parent().hasClass('carousel')) {
        return;
      }
      
      $el.after('<div id="foundationClearingHolder"></div>');

      var grid = $el.detach(),
          grid_outerHTML = '';

      if (grid[0] == null) {
        return;
      } else {
        grid_outerHTML = grid[0].outerHTML;
      }
      
      var holder = this.S('#foundationClearingHolder'),
          settings = $el.data(this.attr_name(true) + '-init'),
          data = {
            grid: '<div class="carousel">' + grid_outerHTML + '</div>',
            viewing: settings.templates.viewing
          },
          wrapper = '<div class="clearing-assembled"><div>' + data.viewing +
            data.grid + '</div></div>',
          touch_label = this.settings.touch_label;

      if (Modernizr.touch) {
        wrapper = $(wrapper).find('.clearing-touch-label').html(touch_label).end();
      }

      holder.after(wrapper).remove();
    },

    open : function ($image, current, target) {
      var self = this,
          body = $(document.body),
          root = target.closest('.clearing-assembled'),
          container = self.S('div', root).first(),
          visible_image = self.S('.visible-img', container),
          image = self.S('img', visible_image).not($image),
          label = self.S('.clearing-touch-label', container),
          error = false;

      image.error(function () {
        error = true;
      });

      function startLoad() {
        setTimeout(function () {
          this.image_loaded(image, function () {
            if (image.outerWidth() === 1 && !error) {
              startLoad.call(this);
            } else {
              cb.call(this, image);
            }
          }.bind(this));
        }.bind(this), 100);
      }

      function cb (image) {
        var $image = $(image);
        $image.css('visibility', 'visible');
        // toggle the gallery
        body.css('overflow', 'hidden');
        root.addClass('clearing-blackout');
        container.addClass('clearing-container');
        visible_image.show();
        this.fix_height(target)
          .caption(self.S('.clearing-caption', visible_image), self.S('img', target))
          .center_and_label(image, label)
          .shift(current, target, function () {
            target.closest('li').siblings().removeClass('visible');
            target.closest('li').addClass('visible');
          });
        visible_image.trigger('opened.fndtn.clearing')
      }

      if (!this.locked()) {
        visible_image.trigger('open.fndtn.clearing');
        // set the image to the selected thumbnail
        image
          .attr('src', this.load($image))
          .css('visibility', 'hidden');

        startLoad.call(this);
      }
    },

    close : function (e, el) {
      e.preventDefault();

      var root = (function (target) {
            if (/blackout/.test(target.selector)) {
              return target;
            } else {
              return target.closest('.clearing-blackout');
            }
          }($(el))),
          body = $(document.body), container, visible_image;

      if (el === e.target && root) {
        body.css('overflow', '');
        container = $('div', root).first();
        visible_image = $('.visible-img', container);
        visible_image.trigger('close.fndtn.clearing');
        this.settings.prev_index = 0;
        $('ul[' + this.attr_name() + ']', root)
          .attr('style', '').closest('.clearing-blackout')
          .removeClass('clearing-blackout');
        container.removeClass('clearing-container');
        visible_image.hide();
        visible_image.trigger('closed.fndtn.clearing');        
      }

      return false;
    },

    is_open : function (current) {
      return current.parent().prop('style').length > 0;
    },

    keydown : function (e) {
      var clearing = $('.clearing-blackout ul[' + this.attr_name() + ']'),
          NEXT_KEY = this.rtl ? 37 : 39,
          PREV_KEY = this.rtl ? 39 : 37,
          ESC_KEY = 27;

      if (e.which === NEXT_KEY) this.go(clearing, 'next');
      if (e.which === PREV_KEY) this.go(clearing, 'prev');
      if (e.which === ESC_KEY) this.S('a.clearing-close').trigger('click').trigger('click.fndtn.clearing');
    },

    nav : function (e, direction) {
      var clearing = $('ul[' + this.attr_name() + ']', '.clearing-blackout');

      e.preventDefault();
      this.go(clearing, direction);
    },

    resize : function () {
      var image = $('img', '.clearing-blackout .visible-img'),
          label = $('.clearing-touch-label', '.clearing-blackout');

      if (image.length) {
        this.center_and_label(image, label);
        image.trigger('resized.fndtn.clearing')
      }
    },

    // visual adjustments
    fix_height : function (target) {
      var lis = target.parent().children(),
          self = this;

      lis.each(function () {
        var li = self.S(this),
            image = li.find('img');

        if (li.height() > image.outerHeight()) {
          li.addClass('fix-height');
        }
      })
      .closest('ul')
      .width(lis.length * 100 + '%');

      return this;
    },

    update_paddles : function (target) {
      target = target.closest('li');
      var visible_image = target
        .closest('.carousel')
        .siblings('.visible-img');

      if (target.next().length > 0) {
        this.S('.clearing-main-next', visible_image).removeClass('disabled');
      } else {
        this.S('.clearing-main-next', visible_image).addClass('disabled');
      }

      if (target.prev().length > 0) {
        this.S('.clearing-main-prev', visible_image).removeClass('disabled');
      } else {
        this.S('.clearing-main-prev', visible_image).addClass('disabled');
      }
    },

    center_and_label : function (target, label) {
      if (!this.rtl) {
        target.css({
          marginLeft : -(target.outerWidth() / 2),
          marginTop : -(target.outerHeight() / 2)
        });

        if (label.length > 0) {
          label.css({
            marginLeft : -(label.outerWidth() / 2),
            marginTop : -(target.outerHeight() / 2)-label.outerHeight()-10
          });
        }
      } else {
        target.css({
          marginRight : -(target.outerWidth() / 2),
          marginTop : -(target.outerHeight() / 2),
          left: 'auto',
          right: '50%'
        });

        if (label.length > 0) {
          label.css({
            marginRight : -(label.outerWidth() / 2),
            marginTop : -(target.outerHeight() / 2)-label.outerHeight()-10,
            left: 'auto',
            right: '50%'
          });
        }
      }
      return this;
    },

    // image loading and preloading

    load : function ($image) {
      var href;

      if ($image[0].nodeName === "A") {
        href = $image.attr('href');
      } else {
        href = $image.parent().attr('href');
      }

      this.preload($image);

      if (href) return href;
      return $image.attr('src');
    },

    preload : function ($image) {
      this
        .img($image.closest('li').next())
        .img($image.closest('li').prev());
    },

    img : function (img) {
      if (img.length) {
        var new_img = new Image(),
            new_a = this.S('a', img);

        if (new_a.length) {
          new_img.src = new_a.attr('href');
        } else {
          new_img.src = this.S('img', img).attr('src');
        }
      }
      return this;
    },

    // image caption

    caption : function (container, $image) {
      var caption = $image.attr('data-caption');

      if (caption) {
        container
          .html(caption)
          .show();
      } else {
        container
          .text('')
          .hide();
      }
      return this;
    }, 

    // directional methods

    go : function ($ul, direction) {
      var current = this.S('.visible', $ul),
          target = current[direction]();

      if (target.length) {
        this.S('img', target)
          .trigger('click', [current, target]).trigger('click.fndtn.clearing', [current, target])
          .trigger('change.fndtn.clearing');
      }
    },

    shift : function (current, target, callback) {
      var clearing = target.parent(),
          old_index = this.settings.prev_index || target.index(),
          direction = this.direction(clearing, current, target),
          dir = this.rtl ? 'right' : 'left',
          left = parseInt(clearing.css('left'), 10),
          width = target.outerWidth(),
          skip_shift;

      var dir_obj = {};

      // we use jQuery animate instead of CSS transitions because we
      // need a callback to unlock the next animation
      // needs support for RTL **
      if (target.index() !== old_index && !/skip/.test(direction)){
        if (/left/.test(direction)) {
          this.lock();
          dir_obj[dir] = left + width;
          clearing.animate(dir_obj, 300, this.unlock());
        } else if (/right/.test(direction)) {
          this.lock();
          dir_obj[dir] = left - width;
          clearing.animate(dir_obj, 300, this.unlock());
        }
      } else if (/skip/.test(direction)) {
        // the target image is not adjacent to the current image, so
        // do we scroll right or not
        skip_shift = target.index() - this.settings.up_count;
        this.lock();

        if (skip_shift > 0) {
          dir_obj[dir] = -(skip_shift * width);
          clearing.animate(dir_obj, 300, this.unlock());
        } else {
          dir_obj[dir] = 0;
          clearing.animate(dir_obj, 300, this.unlock());
        }
      }

      callback();
    },

    direction : function ($el, current, target) {
      var lis = this.S('li', $el),
          li_width = lis.outerWidth() + (lis.outerWidth() / 4),
          up_count = Math.floor(this.S('.clearing-container').outerWidth() / li_width) - 1,
          target_index = lis.index(target),
          response;

      this.settings.up_count = up_count;

      if (this.adjacent(this.settings.prev_index, target_index)) {
        if ((target_index > up_count) && target_index > this.settings.prev_index) {
          response = 'right';
        } else if ((target_index > up_count - 1) && target_index <= this.settings.prev_index) {
          response = 'left';
        } else {
          response = false;
        }
      } else {
        response = 'skip';
      }

      this.settings.prev_index = target_index;

      return response;
    },

    adjacent : function (current_index, target_index) {
      for (var i = target_index + 1; i >= target_index - 1; i--) {
        if (i === current_index) return true;
      }
      return false;
    },

    // lock management

    lock : function () {
      this.settings.locked = true;
    },

    unlock : function () {
      this.settings.locked = false;
    },

    locked : function () {
      return this.settings.locked;
    },

    off : function () {
      this.S(this.scope).off('.fndtn.clearing');
      this.S(window).off('.fndtn.clearing');
    },

    reflow : function () {
      this.init();
    }
  };

}(jQuery, window, window.document));
```
