## Supplementary material for "Comparative pathway enrichment analysis in gastrointestinal cell lines Caco-2, HT-29, HEPG2, and colon fibroblasts using a custom expression panel for tight-junction and cytoskeletal regulatory genes": Sup. Results 1: foundation.dropdown.js

```
;(function ($, window, document, undefined) {
  'use strict';

  Foundation.libs.dropdown = {
    name : 'dropdown',

    version : '5.3.0',

    settings : {
      active_class: 'open',
      align: 'bottom',
      is_hover: false,
      opened: function(){},
      closed: function(){}
    },

    init : function (scope, method, options) {
      Foundation.inherit(this, 'throttle');

      this.bindings(method, options);
    },

    events : function (scope) {
      var self = this,
          S = self.S;

      S(this.scope)
        .off('.dropdown')
        .on('click.fndtn.dropdown', '[' + this.attr_name() + ']', function (e) {
          var settings = S(this).data(self.attr_name(true) + '-init') || self.settings;
          if (!settings.is_hover || Modernizr.touch) {
            e.preventDefault();
            self.toggle($(this));
          }
        })
        .on('mouseenter.fndtn.dropdown', '[' + this.attr_name() + '], [' + this.attr_name() + '-content]', function (e) {
          var $this = S(this),
              dropdown,
              target;

          clearTimeout(self.timeout);

          if ($this.data(self.data_attr())) {
            dropdown = S('#' + $this.data(self.data_attr()));
            target = $this;
          } else {
            dropdown = $this;
            target = S("[" + self.attr_name() + "='" + dropdown.attr('id') + "']");
          }

          var settings = target.data(self.attr_name(true) + '-init') || self.settings;
          
          if(S(e.target).data(self.data_attr()) && settings.is_hover) {
            self.closeall.call(self);
          }
          
          if (settings.is_hover) self.open.apply(self, [dropdown, target]);
        })
        .on('mouseleave.fndtn.dropdown', '[' + this.attr_name() + '], [' + this.attr_name() + '-content]', function (e) {
          var $this = S(this);
          self.timeout = setTimeout(function () {
            if ($this.data(self.data_attr())) {
              var settings = $this.data(self.data_attr(true) + '-init') || self.settings;
              if (settings.is_hover) self.close.call(self, S('#' + $this.data(self.data_attr())));
            } else {
              var target = S('[' + self.attr_name() + '="' + S(this).attr('id') + '"]'),
                  settings = target.data(self.attr_name(true) + '-init') || self.settings;
              if (settings.is_hover) self.close.call(self, $this);
            }
          }.bind(this), 150);
        })
        .on('click.fndtn.dropdown', function (e) {
          var parent = S(e.target).closest('[' + self.attr_name() + '-content]');

          if (S(e.target).data(self.data_attr()) || S(e.target).parent().data(self.data_attr())) {
            return;
          }
          if (!(S(e.target).data('revealId')) && 
            (parent.length > 0 && (S(e.target).is('[' + self.attr_name() + '-content]') || 
              $.contains(parent.first()[0], e.target)))) {
            e.stopPropagation();
            return;
          }

          self.close.call(self, S('[' + self.attr_name() + '-content]'));
        })
        .on('opened.fndtn.dropdown', '[' + self.attr_name() + '-content]', function () {
            self.settings.opened.call(this);
        })
        .on('closed.fndtn.dropdown', '[' + self.attr_name() + '-content]', function () {
            self.settings.closed.call(this);
        });

      S(window)
        .off('.dropdown')
        .on('resize.fndtn.dropdown', self.throttle(function () {
          self.resize.call(self);
        }, 50));

      this.resize();
    },

    close: function (dropdown) {
      var self = this;
      dropdown.each(function () {
        if (self.S(this).hasClass(self.settings.active_class)) {
          self.S(this)
            .css(Foundation.rtl ? 'right':'left', '-99999px')
            .removeClass(self.settings.active_class)
            .prev('[' + self.attr_name() + ']')
            .removeClass(self.settings.active_class)
            .removeData('target');

          self.S(this).trigger('closed').trigger('closed.fndtn.dropdown', [dropdown]);
        }
      });
    },

    closeall: function() {
      var self = this;
      $.each(self.S('[' + this.attr_name() + '-content]'), function() {
        self.close.call(self, self.S(this))
      });
    },

    open: function (dropdown, target) {
        this
          .css(dropdown
            .addClass(this.settings.active_class), target);
        dropdown.prev('[' + this.attr_name() + ']').addClass(this.settings.active_class);
        dropdown.data('target', target.get(0)).trigger('opened').trigger('opened.fndtn.dropdown', [dropdown, target]);
    },

    data_attr: function () {
      if (this.namespace.length > 0) {
        return this.namespace + '-' + this.name;
      }

      return this.name;
    },

    toggle : function (target) {
      var dropdown = this.S('#' + target.data(this.data_attr()));
      if (dropdown.length === 0) {
        // No dropdown found, not continuing
        return;
      }

      this.close.call(this, this.S('[' + this.attr_name() + '-content]').not(dropdown));

      if (dropdown.hasClass(this.settings.active_class)) {
        this.close.call(this, dropdown);
        if (dropdown.data('target') !== target.get(0))
          this.open.call(this, dropdown, target);
      } else {
        this.open.call(this, dropdown, target);
      }
    },

    resize : function () {
      var dropdown = this.S('[' + this.attr_name() + '-content].open'),
          target = this.S("[" + this.attr_name() + "='" + dropdown.attr('id') + "']");

      if (dropdown.length && target.length) {
        this.css(dropdown, target);
      }
    },

    css : function (dropdown, target) {
      var left_offset = Math.max((target.width() - dropdown.width()) / 2, 8);
      
      this.clear_idx();

      if (this.small()) {
        var p = this.dirs.bottom.call(dropdown, target);

        dropdown.attr('style', '').removeClass('drop-left drop-right drop-top').css({
          position : 'absolute',
          width: '95%',
          'max-width': 'none',
          top: p.top
        });

        dropdown.css(Foundation.rtl ? 'right':'left', left_offset);
      } else {
        var settings = target.data(this.attr_name(true) + '-init') || this.settings;

        this.style(dropdown, target, settings);
      }

      return dropdown;
    },

    style : function (dropdown, target, settings) {
      var css = $.extend({position: 'absolute'}, 
        this.dirs[settings.align].call(dropdown, target, settings));

      dropdown.attr('style', '').css(css);
    },

    // return CSS property object
    // `this` is the dropdown
    dirs : {
      // Calculate target offset
      _base : function (t) {
        var o_p = this.offsetParent(),
            o = o_p.offset(),
            p = t.offset();

        p.top -= o.top;
        p.left -= o.left;

        return p;
      },
      top: function (t, s) {
        var self = Foundation.libs.dropdown,
            p = self.dirs._base.call(this, t),
            pip_offset_base = 8;

        this.addClass('drop-top');

        if (t.outerWidth() < this.outerWidth() || self.small()) {
          self.adjust_pip(pip_offset_base, p);
        }

        if (Foundation.rtl) {
          return {left: p.left - this.outerWidth() + t.outerWidth(), 
            top: p.top - this.outerHeight()};
        }

        return {left: p.left, top: p.top - this.outerHeight()};
      },
      bottom: function (t, s) {
        var self = Foundation.libs.dropdown,
            p = self.dirs._base.call(this, t),
            pip_offset_base = 8;

        if (t.outerWidth() < this.outerWidth() || self.small()) {
          self.adjust_pip(pip_offset_base, p);
        }

        if (self.rtl) {
          return {left: p.left - this.outerWidth() + t.outerWidth(), top: p.top + t.outerHeight()};
        }

        return {left: p.left, top: p.top + t.outerHeight()};
      },
      left: function (t, s) {
        var p = Foundation.libs.dropdown.dirs._base.call(this, t);

        this.addClass('drop-left');

        return {left: p.left - this.outerWidth(), top: p.top};
      },
      right: function (t, s) {
        var p = Foundation.libs.dropdown.dirs._base.call(this, t);

        this.addClass('drop-right');

        return {left: p.left + t.outerWidth(), top: p.top};
      }
    },

    // Insert rule to style psuedo elements
    adjust_pip : function (pip_offset_base, p) {
      var sheet = Foundation.stylesheet;

      if (this.small()) {
        pip_offset_base += p.left - 8;
      }

      this.rule_idx = sheet.cssRules.length;

      var sel_before = '.f-dropdown.open:before',
          sel_after  = '.f-dropdown.open:after',
          css_before = 'left: ' + pip_offset_base + 'px;',
          css_after  = 'left: ' + (pip_offset_base - 1) + 'px;';

      if (sheet.insertRule) {
        sheet.insertRule([sel_before, '{', css_before, '}'].join(' '), this.rule_idx);
        sheet.insertRule([sel_after, '{', css_after, '}'].join(' '), this.rule_idx + 1);
      } else {
        sheet.addRule(sel_before, css_before, this.rule_idx);
        sheet.addRule(sel_after, css_after, this.rule_idx + 1);
      }
    },

    // Remove old dropdown rule index
    clear_idx : function () {
      var sheet = Foundation.stylesheet;

      if (this.rule_idx) {
        sheet.deleteRule(this.rule_idx);
        sheet.deleteRule(this.rule_idx);
        delete this.rule_idx;
      }
    },

    small : function () {
      return matchMedia(Foundation.media_queries.small).matches &&
        !matchMedia(Foundation.media_queries.medium).matches;
    },

    off: function () {
      this.S(this.scope).off('.fndtn.dropdown');
      this.S('html, body').off('.fndtn.dropdown');
      this.S(window).off('.fndtn.dropdown');
      this.S('[data-dropdown-content]').off('.fndtn.dropdown');
    },

    reflow : function () {}
  };
}(jQuery, window, window.document));
```
