## Supplementary material for "Comparative pathway enrichment analysis in gastrointestinal cell lines Caco-2, HT-29, HEPG2, and colon fibroblasts using a custom expression panel for tight-junction and cytoskeletal regulatory genes": Sup. Results 1: foundation.equalizer.js

```
;(function ($, window, document, undefined) {
  'use strict';

  Foundation.libs.equalizer = {
    name : 'equalizer',

    version : '5.3.0',

    settings : {
      use_tallest: true,
      before_height_change: $.noop,
      after_height_change: $.noop,
      equalize_on_stack: false
    },

    init : function (scope, method, options) {
      Foundation.inherit(this, 'image_loaded');
      this.bindings(method, options);
      this.reflow();
    },

    events : function () {
      this.S(window).off('.equalizer').on('resize.fndtn.equalizer', function(e){
        this.reflow();
      }.bind(this));
    },

    equalize: function(equalizer) {
      var isStacked = false,
          vals = equalizer.find('[' + this.attr_name() + '-watch]:visible'),
          settings = equalizer.data(this.attr_name(true)+'-init');

      if (vals.length === 0) return;
      var firstTopOffset = vals.first().offset().top;
      settings.before_height_change();
      equalizer.trigger('before-height-change').trigger('before-height-change.fndth.equalizer');
      vals.height('inherit');
      vals.each(function(){
        var el = $(this);
        if (el.offset().top !== firstTopOffset) {
          isStacked = true;
        }
      });

      if (settings.equalize_on_stack === false) {
        if (isStacked) return;
      };

      var heights = vals.map(function(){ return $(this).outerHeight(false) }).get();

      if (settings.use_tallest) {
        var max = Math.max.apply(null, heights);
        vals.css('height', max);
      } else {
        var min = Math.min.apply(null, heights);
        vals.css('height', min);
      }
      settings.after_height_change();
      equalizer.trigger('after-height-change').trigger('after-height-change.fndtn.equalizer');
    },

    reflow : function () {
      var self = this;

      this.S('[' + this.attr_name() + ']', this.scope).each(function(){
        var $eq_target = $(this);
        self.image_loaded(self.S('img', this), function(){
          self.equalize($eq_target)
        });
      });
    }
  };
})(jQuery, window, window.document);
```
