## Supplementary material for "Comparative pathway enrichment analysis in gastrointestinal cell lines Caco-2, HT-29, HEPG2, and colon fibroblasts using a custom expression panel for tight-junction and cytoskeletal regulatory genes": Sup. Results 1: foundation.interchange.js

```
;(function ($, window, document, undefined) {
  'use strict';

  Foundation.libs.interchange = {
    name : 'interchange',

    version : '5.3.0',

    cache : {},

    images_loaded : false,
    nodes_loaded : false,

    settings : {
      load_attr : 'interchange',

      named_queries : {
        'default' : 'only screen',
        small : Foundation.media_queries.small,
        medium : Foundation.media_queries.medium,
        large : Foundation.media_queries.large,
        xlarge : Foundation.media_queries.xlarge,
        xxlarge: Foundation.media_queries.xxlarge,
        landscape : 'only screen and (orientation: landscape)',
        portrait : 'only screen and (orientation: portrait)',
        retina : 'only screen and (-webkit-min-device-pixel-ratio: 2),' +
          'only screen and (min--moz-device-pixel-ratio: 2),' +
          'only screen and (-o-min-device-pixel-ratio: 2/1),' +
          'only screen and (min-device-pixel-ratio: 2),' +
          'only screen and (min-resolution: 192dpi),' +
          'only screen and (min-resolution: 2dppx)'
      },

      directives : {
        replace: function (el, path, trigger) {
          // The trigger argument, if called within the directive, fires
          // an event named after the directive on the element, passing
          // any parameters along to the event that you pass to trigger.
          //
          // ex. trigger(), trigger([a, b, c]), or trigger(a, b, c)
          //
          // This allows you to bind a callback like so:
          // $('#interchangeContainer').on('replace', function (e, a, b, c) {
          //   console.log($(this).html(), a, b, c);
          // });

          if (/IMG/.test(el[0].nodeName)) {
            var orig_path = el[0].src;

            if (new RegExp(path, 'i').test(orig_path)) return;

            el[0].src = path;

            return trigger(el[0].src);
          }
          var last_path = el.data(this.data_attr + '-last-path'),
              self = this;

          if (last_path == path) return;

          if (/\.(gif|jpg|jpeg|tiff|png)([?#].*)?/i.test(path)) {
            $(el).css('background-image', 'url('+path+')');
            el.data('interchange-last-path', path);
            return trigger(path);
          }

          return $.get(path, function (response) {
            el.html(response);
            el.data(self.data_attr + '-last-path', path);
            trigger();
          });

        }
      }
    },

    init : function (scope, method, options) {
      Foundation.inherit(this, 'throttle random_str');

      this.data_attr = this.set_data_attr();
      $.extend(true, this.settings, method, options);
      this.bindings(method, options);
      this.load('images');
      this.load('nodes');
    },

    get_media_hash : function() {
        var mediaHash='';
        for (var queryName in this.settings.named_queries ) {
            mediaHash += matchMedia(this.settings.named_queries[queryName]).matches.toString();
        }
        return mediaHash;
    },

    events : function () {
      var self = this, prevMediaHash;

      $(window)
        .off('.interchange')
        .on('resize.fndtn.interchange', self.throttle(function () {
            var currMediaHash = self.get_media_hash();
            if (currMediaHash !== prevMediaHash) {
                self.resize();
            }
            prevMediaHash = currMediaHash;
        }, 50));

      return this;
    },

    resize : function () {
      var cache = this.cache;

      if(!this.images_loaded || !this.nodes_loaded) {
        setTimeout($.proxy(this.resize, this), 50);
        return;
      }

      for (var uuid in cache) {
        if (cache.hasOwnProperty(uuid)) {
          var passed = this.results(uuid, cache[uuid]);

          if (passed) {
            this.settings.directives[passed
              .scenario[1]].call(this, passed.el, passed.scenario[0], function () {
                if (arguments[0] instanceof Array) { 
                  var args = arguments[0];
                } else { 
                  var args = Array.prototype.slice.call(arguments, 0);
                }

                passed.el.trigger(passed.scenario[1], args);
              });
          }
        }
      }

    },

    results : function (uuid, scenarios) {
      var count = scenarios.length;

      if (count > 0) {
        var el = this.S('[' + this.add_namespace('data-uuid') + '="' + uuid + '"]');

        while (count--) {
          var mq, rule = scenarios[count][2];
          if (this.settings.named_queries.hasOwnProperty(rule)) {
            mq = matchMedia(this.settings.named_queries[rule]);
          } else {
            mq = matchMedia(rule);
          }
          if (mq.matches) {
            return {el: el, scenario: scenarios[count]};
          }
        }
      }

      return false;
    },

    load : function (type, force_update) {
      if (typeof this['cached_' + type] === 'undefined' || force_update) {
        this['update_' + type]();
      }

      return this['cached_' + type];
    },

    update_images : function () {
      var images = this.S('img[' + this.data_attr + ']'),
          count = images.length,
          i = count,
          loaded_count = 0,
          data_attr = this.data_attr;

      this.cache = {};
      this.cached_images = [];
      this.images_loaded = (count === 0);

      while (i--) {
        loaded_count++;
        if (images[i]) {
          var str = images[i].getAttribute(data_attr) || '';

          if (str.length > 0) {
            this.cached_images.push(images[i]);
          }
        }

        if (loaded_count === count) {
          this.images_loaded = true;
          this.enhance('images');
        }
      }

      return this;
    },

    update_nodes : function () {
      var nodes = this.S('[' + this.data_attr + ']').not('img'),
          count = nodes.length,
          i = count,
          loaded_count = 0,
          data_attr = this.data_attr;

      this.cached_nodes = [];
      this.nodes_loaded = (count === 0);


      while (i--) {
        loaded_count++;
        var str = nodes[i].getAttribute(data_attr) || '';

        if (str.length > 0) {
          this.cached_nodes.push(nodes[i]);
        }

        if(loaded_count === count) {
          this.nodes_loaded = true;
          this.enhance('nodes');
        }
      }

      return this;
    },

    enhance : function (type) {
      var i = this['cached_' + type].length;

      while (i--) {
        this.object($(this['cached_' + type][i]));
      }

      return $(window).trigger('resize').trigger('resize.fndtn.interchange');
    },

    convert_directive : function (directive) {

      var trimmed = this.trim(directive);

      if (trimmed.length > 0) {
        return trimmed;
      }

      return 'replace';
    },

    parse_scenario : function (scenario) {
      // This logic had to be made more complex since some users were using commas in the url path
      // So we cannot simply just split on a comma
      var directive_match = scenario[0].match(/(.+),\s*(\w+)\s*$/),
      media_query         = scenario[1];

      if (directive_match) {
        var path  = directive_match[1],
        directive = directive_match[2];
      }
      else {
        var cached_split = scenario[0].split(/,\s*$/),
        path             = cached_split[0],
        directive        = '';               
      }

      return [this.trim(path), this.convert_directive(directive), this.trim(media_query)];
    },

    object : function(el) {
      var raw_arr = this.parse_data_attr(el),
          scenarios = [], 
          i = raw_arr.length;

      if (i > 0) {
        while (i--) {
          var split = raw_arr[i].split(/\((.*?)(\))$/);

          if (split.length > 1) {
            var params = this.parse_scenario(split);
            scenarios.push(params);
          }
        }
      }

      return this.store(el, scenarios);
    },

    store : function (el, scenarios) {
      var uuid = this.random_str(),
          current_uuid = el.data(this.add_namespace('uuid', true));

      if (this.cache[current_uuid]) return this.cache[current_uuid];

      el.attr(this.add_namespace('data-uuid'), uuid);

      return this.cache[uuid] = scenarios;
    },

    trim : function(str) {

      if (typeof str === 'string') {
        return $.trim(str);
      }

      return str;
    },

    set_data_attr: function (init) {
      if (init) {
        if (this.namespace.length > 0) {
          return this.namespace + '-' + this.settings.load_attr;
        }

        return this.settings.load_attr;
      }

      if (this.namespace.length > 0) {
        return 'data-' + this.namespace + '-' + this.settings.load_attr;
      }

      return 'data-' + this.settings.load_attr;
    },

    parse_data_attr : function (el) {
      var raw = el.attr(this.attr_name()).split(/\[(.*?)\]/),
          i = raw.length, 
          output = [];

      while (i--) {
        if (raw[i].replace(/[\W\d]+/, '').length > 4) {
          output.push(raw[i]);
        }
      }

      return output;
    },

    reflow : function () {
      this.load('images', true);
      this.load('nodes', true);
    }

  };

}(jQuery, window, window.document));
```
