## Supplementary material for "Comparative pathway enrichment analysis in gastrointestinal cell lines Caco-2, HT-29, HEPG2, and colon fibroblasts using a custom expression panel for tight-junction and cytoskeletal regulatory genes": Sup. Results 1: foundation.joyride.js

```
;(function ($, window, document, undefined) {
  'use strict';

  var Modernizr = Modernizr || false;

  Foundation.libs.joyride = {
    name : 'joyride',

    version : '5.3.0',

    defaults : {
      expose                   : false,     // turn on or off the expose feature
      modal                    : true,      // Whether to cover page with modal during the tour
      tip_location             : 'bottom',  // 'top' or 'bottom' in relation to parent
      nub_position             : 'auto',    // override on a per tooltip bases
      scroll_speed             : 1500,      // Page scrolling speed in milliseconds, 0 = no scroll animation
      scroll_animation         : 'linear',  // supports 'swing' and 'linear', extend with jQuery UI.
      timer                    : 0,         // 0 = no timer , all other numbers = timer in milliseconds
      start_timer_on_click     : true,      // true or false - true requires clicking the first button start the timer
      start_offset             : 0,         // the index of the tooltip you want to start on (index of the li)
      next_button              : true,      // true or false to control whether a next button is used
      tip_animation            : 'fade',    // 'pop' or 'fade' in each tip
      pause_after              : [],        // array of indexes where to pause the tour after
      exposed                  : [],        // array of expose elements
      tip_animation_fade_speed : 300,       // when tipAnimation = 'fade' this is speed in milliseconds for the transition
      cookie_monster           : false,     // true or false to control whether cookies are used
      cookie_name              : 'joyride', // Name the cookie you'll use
      cookie_domain            : false,     // Will this cookie be attached to a domain, ie. '.notableapp.com'
      cookie_expires           : 365,       // set when you would like the cookie to expire.
      tip_container            : 'body',    // Where will the tip be attached
      abort_on_close           : true,      // When true, the close event will not fire any callback
      tip_location_patterns    : {
        top: ['bottom'],
        bottom: [], // bottom should not need to be repositioned
        left: ['right', 'top', 'bottom'],
        right: ['left', 'top', 'bottom']
      },
      post_ride_callback     : function (){},    // A method to call once the tour closes (canceled or complete)
      post_step_callback     : function (){},    // A method to call after each step
      pre_step_callback      : function (){},    // A method to call before each step
      pre_ride_callback      : function (){},    // A method to call before the tour starts (passed index, tip, and cloned exposed element)
      post_expose_callback   : function (){},    // A method to call after an element has been exposed
      template : { // HTML segments for tip layout
        link    : '<a href="#close" class="joyride-close-tip">&times;</a>',
        timer   : '<div class="joyride-timer-indicator-wrap"><span class="joyride-timer-indicator"></span></div>',
        tip     : '<div class="joyride-tip-guide"><span class="joyride-nub"></span></div>',
        wrapper : '<div class="joyride-content-wrapper"></div>',
        button  : '<a href="#" class="small button joyride-next-tip"></a>',
        modal   : '<div class="joyride-modal-bg"></div>',
        expose  : '<div class="joyride-expose-wrapper"></div>',
        expose_cover: '<div class="joyride-expose-cover"></div>'
      },
      expose_add_class : '' // One or more space-separated class names to be added to exposed element
    },

    init : function (scope, method, options) {
      Foundation.inherit(this, 'throttle random_str');

      this.settings = this.settings || $.extend({}, this.defaults, (options || method));

      this.bindings(method, options)
    },

    events : function () {
      var self = this;

      $(this.scope)
        .off('.joyride')
        .on('click.fndtn.joyride', '.joyride-next-tip, .joyride-modal-bg', function (e) {
          e.preventDefault();

          if (this.settings.$li.next().length < 1) {
            this.end();
          } else if (this.settings.timer > 0) {
            clearTimeout(this.settings.automate);
            this.hide();
            this.show();
            this.startTimer();
          } else {
            this.hide();
            this.show();
          }

        }.bind(this))

        .on('click.fndtn.joyride', '.joyride-close-tip', function (e) {
          e.preventDefault();
          this.end(this.settings.abort_on_close);
        }.bind(this));

      $(window)
        .off('.joyride')
        .on('resize.fndtn.joyride', self.throttle(function () {
          if ($('[' + self.attr_name() + ']').length > 0 && self.settings.$next_tip) {
            if (self.settings.exposed.length > 0) {
              var $els = $(self.settings.exposed);

              $els.each(function () {
                var $this = $(this);
                self.un_expose($this);
                self.expose($this);
              });
            }

            if (self.is_phone()) {
              self.pos_phone();
            } else {
              self.pos_default(false);
            }
          }
        }, 100));
    },

    start : function () {
      var self = this,
          $this = $('[' + this.attr_name() + ']', this.scope),
          integer_settings = ['timer', 'scrollSpeed', 'startOffset', 'tipAnimationFadeSpeed', 'cookieExpires'],
          int_settings_count = integer_settings.length;

      if (!$this.length > 0) return;

      if (!this.settings.init) this.events();

      this.settings = $this.data(this.attr_name(true) + '-init');

      // non configureable settings
      this.settings.$content_el = $this;
      this.settings.$body = $(this.settings.tip_container);
      this.settings.body_offset = $(this.settings.tip_container).position();
      this.settings.$tip_content = this.settings.$content_el.find('> li');
      this.settings.paused = false;
      this.settings.attempts = 0;

      // can we create cookies?
      if (typeof $.cookie !== 'function') {
        this.settings.cookie_monster = false;
      }

      // generate the tips and insert into dom.
      if (!this.settings.cookie_monster || this.settings.cookie_monster && !$.cookie(this.settings.cookie_name)) {
        this.settings.$tip_content.each(function (index) {
          var $this = $(this);
          this.settings = $.extend({}, self.defaults, self.data_options($this));

          // Make sure that settings parsed from data_options are integers where necessary
          var i = int_settings_count;
          while (i--) {
            self.settings[integer_settings[i]] = parseInt(self.settings[integer_settings[i]], 10);
          }
          self.create({$li : $this, index : index});
        });

        // show first tip
        if (!this.settings.start_timer_on_click && this.settings.timer > 0) {
          this.show('init');
          this.startTimer();
        } else {
          this.show('init');
        }

      }
    },

    resume : function () {
      this.set_li();
      this.show();
    },

    tip_template : function (opts) {
      var $blank, content;

      opts.tip_class = opts.tip_class || '';

      $blank = $(this.settings.template.tip).addClass(opts.tip_class);
      content = $.trim($(opts.li).html()) +
        this.button_text(opts.button_text) +
        this.settings.template.link +
        this.timer_instance(opts.index);

      $blank.append($(this.settings.template.wrapper));
      $blank.first().attr(this.add_namespace('data-index'), opts.index);
      $('.joyride-content-wrapper', $blank).append(content);

      return $blank[0];
    },

    timer_instance : function (index) {
      var txt;

      if ((index === 0 && this.settings.start_timer_on_click && this.settings.timer > 0) || this.settings.timer === 0) {
        txt = '';
      } else {
        txt = $(this.settings.template.timer)[0].outerHTML;
      }
      return txt;
    },

    button_text : function (txt) {
      if (this.settings.next_button) {
        txt = $.trim(txt) || 'Next';
        txt = $(this.settings.template.button).append(txt)[0].outerHTML;
      } else {
        txt = '';
      }
      return txt;
    },

    create : function (opts) {
      var buttonText = opts.$li.attr(this.add_namespace('data-button')) 
        || opts.$li.attr(this.add_namespace('data-text')),
        tipClass = opts.$li.attr('class'),
        $tip_content = $(this.tip_template({
          tip_class : tipClass,
          index : opts.index,
          button_text : buttonText,
          li : opts.$li
        }));

      $(this.settings.tip_container).append($tip_content);
    },

    show : function (init) {
      var $timer = null;

      // are we paused?
      if (this.settings.$li === undefined
        || ($.inArray(this.settings.$li.index(), this.settings.pause_after) === -1)) {

        // don't go to the next li if the tour was paused
        if (this.settings.paused) {
          this.settings.paused = false;
        } else {
          this.set_li(init);
        }

        this.settings.attempts = 0;

        if (this.settings.$li.length && this.settings.$target.length > 0) {
          if (init) { //run when we first start
            this.settings.pre_ride_callback(this.settings.$li.index(), this.settings.$next_tip);
            if (this.settings.modal) {
              this.show_modal();
            }
          }

          this.settings.pre_step_callback(this.settings.$li.index(), this.settings.$next_tip);

          if (this.settings.modal && this.settings.expose) {
            this.expose();
          }

          this.settings.tip_settings = $.extend({}, this.settings, this.data_options(this.settings.$li));

          this.settings.timer = parseInt(this.settings.timer, 10);

          this.settings.tip_settings.tip_location_pattern = this.settings.tip_location_patterns[this.settings.tip_settings.tip_location];

          // scroll if not modal
          if (!/body/i.test(this.settings.$target.selector)) {
            this.scroll_to();
          }

          if (this.is_phone()) {
            this.pos_phone(true);
          } else {
            this.pos_default(true);
          }

          $timer = this.settings.$next_tip.find('.joyride-timer-indicator');

          if (/pop/i.test(this.settings.tip_animation)) {

            $timer.width(0);

            if (this.settings.timer > 0) {

              this.settings.$next_tip.show();

              setTimeout(function () {
                $timer.animate({
                  width: $timer.parent().width()
                }, this.settings.timer, 'linear');
              }.bind(this), this.settings.tip_animation_fade_speed);

            } else {
              this.settings.$next_tip.show();

            }


          } else if (/fade/i.test(this.settings.tip_animation)) {

            $timer.width(0);

            if (this.settings.timer > 0) {

              this.settings.$next_tip
                .fadeIn(this.settings.tip_animation_fade_speed)
                .show();

              setTimeout(function () {
                $timer.animate({
                  width: $timer.parent().width()
                }, this.settings.timer, 'linear');
              }.bind(this), this.settings.tip_animation_fade_speed);

            } else {
              this.settings.$next_tip.fadeIn(this.settings.tip_animation_fade_speed);
            }
          }

          this.settings.$current_tip = this.settings.$next_tip;

        // skip non-existant targets
        } else if (this.settings.$li && this.settings.$target.length < 1) {

          this.show();

        } else {

          this.end();

        }
      } else {

        this.settings.paused = true;

      }

    },

    is_phone : function () {
      return matchMedia(Foundation.media_queries.small).matches &&
        !matchMedia(Foundation.media_queries.medium).matches;
    },

    hide : function () {
      if (this.settings.modal && this.settings.expose) {
        this.un_expose();
      }

      if (!this.settings.modal) {
        $('.joyride-modal-bg').hide();
      }

      // Prevent scroll bouncing...wait to remove from layout
      this.settings.$current_tip.css('visibility', 'hidden');
      setTimeout($.proxy(function() {
        this.hide();
        this.css('visibility', 'visible');
      }, this.settings.$current_tip), 0);
      this.settings.post_step_callback(this.settings.$li.index(),
        this.settings.$current_tip);
    },

    set_li : function (init) {
      if (init) {
        this.settings.$li = this.settings.$tip_content.eq(this.settings.start_offset);
        this.set_next_tip();
        this.settings.$current_tip = this.settings.$next_tip;
      } else {
        this.settings.$li = this.settings.$li.next();
        this.set_next_tip();
      }

      this.set_target();
    },

    set_next_tip : function () {
      this.settings.$next_tip = $(".joyride-tip-guide").eq(this.settings.$li.index());
      this.settings.$next_tip.data('closed', '');
    },

    set_target : function () {
      var cl = this.settings.$li.attr(this.add_namespace('data-class')),
          id = this.settings.$li.attr(this.add_namespace('data-id')),
          $sel = function () {
            if (id) {
              return $(document.getElementById(id));
            } else if (cl) {
              return $('.' + cl).first();
            } else {
              return $('body');
            }
          };

      this.settings.$target = $sel();
    },

    scroll_to : function () {
      var window_half, tipOffset;

      window_half = $(window).height() / 2;
      tipOffset = Math.ceil(this.settings.$target.offset().top - window_half + this.settings.$next_tip.outerHeight());

      if (tipOffset != 0) {
        $('html, body').stop().animate({
          scrollTop: tipOffset
        }, this.settings.scroll_speed, 'swing');
      }
    },

    paused : function () {
      return ($.inArray((this.settings.$li.index() + 1), this.settings.pause_after) === -1);
    },

    restart : function () {
      this.hide();
      this.settings.$li = undefined;
      this.show('init');
    },

    pos_default : function (init) {
      var $nub = this.settings.$next_tip.find('.joyride-nub'),
          nub_width = Math.ceil($nub.outerWidth() / 2),
          nub_height = Math.ceil($nub.outerHeight() / 2),
          toggle = init || false;

      // tip must not be "display: none" to calculate position
      if (toggle) {
        this.settings.$next_tip.css('visibility', 'hidden');
        this.settings.$next_tip.show();
      }

      if (!/body/i.test(this.settings.$target.selector)) {
      	  var topAdjustment = this.settings.tip_settings.tipAdjustmentY ? parseInt(this.settings.tip_settings.tipAdjustmentY) : 0,
			        leftAdjustment = this.settings.tip_settings.tipAdjustmentX ? parseInt(this.settings.tip_settings.tipAdjustmentX) : 0;
          
          if (this.bottom()) {
            if (this.rtl) {
              this.settings.$next_tip.css({
                top: (this.settings.$target.offset().top + nub_height + this.settings.$target.outerHeight() + topAdjustment),
                left: this.settings.$target.offset().left + this.settings.$target.outerWidth() - this.settings.$next_tip.outerWidth() + leftAdjustment});
            } else {
              this.settings.$next_tip.css({
                top: (this.settings.$target.offset().top + nub_height + this.settings.$target.outerHeight() + topAdjustment),
                left: this.settings.$target.offset().left + leftAdjustment});
            }

            this.nub_position($nub, this.settings.tip_settings.nub_position, 'top');

          } else if (this.top()) {
            if (this.rtl) {
              this.settings.$next_tip.css({
                top: (this.settings.$target.offset().top - this.settings.$next_tip.outerHeight() - nub_height + topAdjustment),
                left: this.settings.$target.offset().left + this.settings.$target.outerWidth() - this.settings.$next_tip.outerWidth()});
            } else {
              this.settings.$next_tip.css({
                top: (this.settings.$target.offset().top - this.settings.$next_tip.outerHeight() - nub_height + topAdjustment),
                left: this.settings.$target.offset().left + leftAdjustment});
            }

            this.nub_position($nub, this.settings.tip_settings.nub_position, 'bottom');

          } else if (this.right()) {

            this.settings.$next_tip.css({
              top: this.settings.$target.offset().top + topAdjustment,
              left: (this.settings.$target.outerWidth() + this.settings.$target.offset().left + nub_width + leftAdjustment)});

            this.nub_position($nub, this.settings.tip_settings.nub_position, 'left');

          } else if (this.left()) {

            this.settings.$next_tip.css({
              top: this.settings.$target.offset().top + topAdjustment,
              left: (this.settings.$target.offset().left - this.settings.$next_tip.outerWidth() - nub_width + leftAdjustment)});

            this.nub_position($nub, this.settings.tip_settings.nub_position, 'right');

          }

          if (!this.visible(this.corners(this.settings.$next_tip)) && this.settings.attempts < this.settings.tip_settings.tip_location_pattern.length) {

            $nub.removeClass('bottom')
              .removeClass('top')
              .removeClass('right')
              .removeClass('left');

            this.settings.tip_settings.tip_location = this.settings.tip_settings.tip_location_pattern[this.settings.attempts];

            this.settings.attempts++;

            this.pos_default();

          }

      } else if (this.settings.$li.length) {

        this.pos_modal($nub);

      }

      if (toggle) {
        this.settings.$next_tip.hide();
        this.settings.$next_tip.css('visibility', 'visible');
      }

    },

    pos_phone : function (init) {
      var tip_height = this.settings.$next_tip.outerHeight(),
          tip_offset = this.settings.$next_tip.offset(),
          target_height = this.settings.$target.outerHeight(),
          $nub = $('.joyride-nub', this.settings.$next_tip),
          nub_height = Math.ceil($nub.outerHeight() / 2),
          toggle = init || false;

      $nub.removeClass('bottom')
        .removeClass('top')
        .removeClass('right')
        .removeClass('left');

      if (toggle) {
        this.settings.$next_tip.css('visibility', 'hidden');
        this.settings.$next_tip.show();
      }

      if (!/body/i.test(this.settings.$target.selector)) {

        if (this.top()) {

            this.settings.$next_tip.offset({top: this.settings.$target.offset().top - tip_height - nub_height});
            $nub.addClass('bottom');

        } else {

          this.settings.$next_tip.offset({top: this.settings.$target.offset().top + target_height + nub_height});
          $nub.addClass('top');

        }

      } else if (this.settings.$li.length) {
        this.pos_modal($nub);
      }

      if (toggle) {
        this.settings.$next_tip.hide();
        this.settings.$next_tip.css('visibility', 'visible');
      }
    },

    pos_modal : function ($nub) {
      this.center();
      $nub.hide();

      this.show_modal();
    },

    show_modal : function () {
      if (!this.settings.$next_tip.data('closed')) {
        var joyridemodalbg =  $('.joyride-modal-bg');
        if (joyridemodalbg.length < 1) {
          $('body').append(this.settings.template.modal).show();
        }

        if (/pop/i.test(this.settings.tip_animation)) {
            joyridemodalbg.show();
        } else {
            joyridemodalbg.fadeIn(this.settings.tip_animation_fade_speed);
        }
      }
    },

    expose : function () {
      var expose,
          exposeCover,
          el,
          origCSS,
          origClasses,
          randId = 'expose-' + this.random_str(6);

      if (arguments.length > 0 && arguments[0] instanceof $) {
        el = arguments[0];
      } else if(this.settings.$target && !/body/i.test(this.settings.$target.selector)){
        el = this.settings.$target;
      }  else {
        return false;
      }

      if(el.length < 1){
        if(window.console){
          console.error('element not valid', el);
        }
        return false;
      }

      expose = $(this.settings.template.expose);
      this.settings.$body.append(expose);
      expose.css({
        top: el.offset().top,
        left: el.offset().left,
        width: el.outerWidth(true),
        height: el.outerHeight(true)
      });

      exposeCover = $(this.settings.template.expose_cover);

      origCSS = {
        zIndex: el.css('z-index'),
        position: el.css('position')
      };

      origClasses = el.attr('class') == null ? '' : el.attr('class');

      el.css('z-index',parseInt(expose.css('z-index'))+1);

      if (origCSS.position == 'static') {
        el.css('position','relative');
      }

      el.data('expose-css',origCSS);
      el.data('orig-class', origClasses);
      el.attr('class', origClasses + ' ' + this.settings.expose_add_class);

      exposeCover.css({
        top: el.offset().top,
        left: el.offset().left,
        width: el.outerWidth(true),
        height: el.outerHeight(true)
      });

      if (this.settings.modal) this.show_modal();

      this.settings.$body.append(exposeCover);
      expose.addClass(randId);
      exposeCover.addClass(randId);
      el.data('expose', randId);
      this.settings.post_expose_callback(this.settings.$li.index(), this.settings.$next_tip, el);
      this.add_exposed(el);
    },

    un_expose : function () {
      var exposeId,
          el,
          expose ,
          origCSS,
          origClasses,
          clearAll = false;

      if (arguments.length > 0 && arguments[0] instanceof $) {
        el = arguments[0];
      } else if(this.settings.$target && !/body/i.test(this.settings.$target.selector)){
        el = this.settings.$target;
      }  else {
        return false;
      }

      if(el.length < 1){
        if (window.console) {
          console.error('element not valid', el);
        }
        return false;
      }

      exposeId = el.data('expose');
      expose = $('.' + exposeId);

      if (arguments.length > 1) {
        clearAll = arguments[1];
      }

      if (clearAll === true) {
        $('.joyride-expose-wrapper,.joyride-expose-cover').remove();
      } else {
        expose.remove();
      }

      origCSS = el.data('expose-css');

      if (origCSS.zIndex == 'auto') {
        el.css('z-index', '');
      } else {
        el.css('z-index', origCSS.zIndex);
      }

      if (origCSS.position != el.css('position')) {
        if(origCSS.position == 'static') {// this is default, no need to set it.
          el.css('position', '');
        } else {
          el.css('position', origCSS.position);
        }
      }

      origClasses = el.data('orig-class');
      el.attr('class', origClasses);
      el.removeData('orig-classes');

      el.removeData('expose');
      el.removeData('expose-z-index');
      this.remove_exposed(el);
    },

    add_exposed: function(el){
      this.settings.exposed = this.settings.exposed || [];
      if (el instanceof $ || typeof el === 'object') {
        this.settings.exposed.push(el[0]);
      } else if (typeof el == 'string') {
        this.settings.exposed.push(el);
      }
    },

    remove_exposed: function(el){
      var search, i;
      if (el instanceof $) {
        search = el[0]
      } else if (typeof el == 'string'){
        search = el;
      }

      this.settings.exposed = this.settings.exposed || [];
      i = this.settings.exposed.length;

      while (i--) {
        if (this.settings.exposed[i] == search) {
          this.settings.exposed.splice(i, 1);
          return;
        }
      }
    },

    center : function () {
      var $w = $(window);

      this.settings.$next_tip.css({
        top : ((($w.height() - this.settings.$next_tip.outerHeight()) / 2) + $w.scrollTop()),
        left : ((($w.width() - this.settings.$next_tip.outerWidth()) / 2) + $w.scrollLeft())
      });

      return true;
    },

    bottom : function () {
      return /bottom/i.test(this.settings.tip_settings.tip_location);
    },

    top : function () {
      return /top/i.test(this.settings.tip_settings.tip_location);
    },

    right : function () {
      return /right/i.test(this.settings.tip_settings.tip_location);
    },

    left : function () {
      return /left/i.test(this.settings.tip_settings.tip_location);
    },

    corners : function (el) {
      var w = $(window),
          window_half = w.height() / 2,
          //using this to calculate since scroll may not have finished yet.
          tipOffset = Math.ceil(this.settings.$target.offset().top - window_half + this.settings.$next_tip.outerHeight()),
          right = w.width() + w.scrollLeft(),
          offsetBottom =  w.height() + tipOffset,
          bottom = w.height() + w.scrollTop(),
          top = w.scrollTop();

      if (tipOffset < top) {
        if (tipOffset < 0) {
          top = 0;
        } else {
          top = tipOffset;
        }
      }

      if (offsetBottom > bottom) {
        bottom = offsetBottom;
      }

      return [
        el.offset().top < top,
        right < el.offset().left + el.outerWidth(),
        bottom < el.offset().top + el.outerHeight(),
        w.scrollLeft() > el.offset().left
      ];
    },

    visible : function (hidden_corners) {
      var i = hidden_corners.length;

      while (i--) {
        if (hidden_corners[i]) return false;
      }

      return true;
    },

    nub_position : function (nub, pos, def) {
      if (pos === 'auto') {
        nub.addClass(def);
      } else {
        nub.addClass(pos);
      }
    },

    startTimer : function () {
      if (this.settings.$li.length) {
        this.settings.automate = setTimeout(function () {
          this.hide();
          this.show();
          this.startTimer();
        }.bind(this), this.settings.timer);
      } else {
        clearTimeout(this.settings.automate);
      }
    },

    end : function (abort) {
      if (this.settings.cookie_monster) {
        $.cookie(this.settings.cookie_name, 'ridden', { expires: this.settings.cookie_expires, domain: this.settings.cookie_domain });
      }

      if (this.settings.timer > 0) {
        clearTimeout(this.settings.automate);
      }

      if (this.settings.modal && this.settings.expose) {
        this.un_expose();
      }

      this.settings.$next_tip.data('closed', true);

      $('.joyride-modal-bg').hide();
      this.settings.$current_tip.hide();

      if (typeof abort === 'undefined' || abort === false) {
        this.settings.post_step_callback(this.settings.$li.index(), this.settings.$current_tip);
        this.settings.post_ride_callback(this.settings.$li.index(), this.settings.$current_tip);
      }

      $('.joyride-tip-guide').remove();
    },

    off : function () {
      $(this.scope).off('.joyride');
      $(window).off('.joyride');
      $('.joyride-close-tip, .joyride-next-tip, .joyride-modal-bg').off('.joyride');
      $('.joyride-tip-guide, .joyride-modal-bg').remove();
      clearTimeout(this.settings.automate);
      this.settings = {};
    },

    reflow : function () {}
  };
}(jQuery, window, window.document));
```
