## Supplementary material for "Comparative pathway enrichment analysis in gastrointestinal cell lines Caco-2, HT-29, HEPG2, and colon fibroblasts using a custom expression panel for tight-junction and cytoskeletal regulatory genes": Sup. Results 1: foundation.js

```
/*
 * Foundation Responsive Library
 * http://foundation.zurb.com
 * Copyright 2014, ZURB
 * Free to use under the MIT license.
 * http://www.opensource.org/licenses/mit-license.php
*/

(function ($, window, document, undefined) {
  'use strict';

  var header_helpers = function (class_array) {
    var i = class_array.length;
    var head = $('head');

    while (i--) {
      if(head.has('.' + class_array[i]).length === 0) {
        head.append('<meta class="' + class_array[i] + '" />');
      }
    }
  };

  header_helpers([
    'foundation-mq-small',
    'foundation-mq-medium',
    'foundation-mq-large',
    'foundation-mq-xlarge',
    'foundation-mq-xxlarge',
    'foundation-data-attribute-namespace']);

  // Enable FastClick if present

  $(function() {
    if (typeof FastClick !== 'undefined') {
      // Don't attach to body if undefined
      if (typeof document.body !== 'undefined') {
        FastClick.attach(document.body);
      }
    }
  });

  // private Fast Selector wrapper,
  // returns jQuery object. Only use where
  // getElementById is not available.
  var S = function (selector, context) {
    if (typeof selector === 'string') {
      if (context) {
        var cont;
        if (context.jquery) {
          cont = context[0];
          if (!cont) return context;
        } else {
          cont = context;
        }
        return $(cont.querySelectorAll(selector));
      }

      return $(document.querySelectorAll(selector));
    }

    return $(selector, context);
  };

  // Namespace functions.

  var attr_name = function (init) {
    var arr = [];
    if (!init) arr.push('data');
    if (this.namespace.length > 0) arr.push(this.namespace);
    arr.push(this.name);

    return arr.join('-');
  };

  var add_namespace = function (str) {
    var parts = str.split('-'),
        i = parts.length,
        arr = [];

    while (i--) {
      if (i !== 0) {
        arr.push(parts[i]);
      } else {
        if (this.namespace.length > 0) {
          arr.push(this.namespace, parts[i]);
        } else {
          arr.push(parts[i]);
        }
      }
    }

    return arr.reverse().join('-');
  };

  // Event binding and data-options updating.

  var bindings = function (method, options) {
    var self = this,
        should_bind_events = !S(this).data(this.attr_name(true));


    if (S(this.scope).is('[' + this.attr_name() +']')) {
      S(this.scope).data(this.attr_name(true) + '-init', $.extend({}, this.settings, (options || method), this.data_options(S(this.scope))));

      if (should_bind_events) {
        this.events(this.scope);
      }

    } else {
      S('[' + this.attr_name() +']', this.scope).each(function () {
        var should_bind_events = !S(this).data(self.attr_name(true) + '-init');
        S(this).data(self.attr_name(true) + '-init', $.extend({}, self.settings, (options || method), self.data_options(S(this))));

        if (should_bind_events) {
          self.events(this);
        }
      });
    }
    // # Patch to fix #5043 to move this *after* the if/else clause in order for Backbone and similar frameworks to have improved control over event binding and data-options updating. 
    if (typeof method === 'string') {
      return this[method].call(this, options);
    }

  };

  var single_image_loaded = function (image, callback) {
    function loaded () {
      callback(image[0]);
    }

    function bindLoad () {
      this.one('load', loaded);

      if (/MSIE (\d+\.\d+);/.test(navigator.userAgent)) {
        var src = this.attr( 'src' ),
            param = src.match( /\?/ ) ? '&' : '?';

        param += 'random=' + (new Date()).getTime();
        this.attr('src', src + param);
      }
    }

    if (!image.attr('src')) {
      loaded();
      return;
    }

    if (image[0].complete || image[0].readyState === 4) {
      loaded();
    } else {
      bindLoad.call(image);
    }
  };

  /*
    https://github.com/paulirish/matchMedia.js
  */

  window.matchMedia = window.matchMedia || (function( doc ) {

    "use strict";

    var bool,
        docElem = doc.documentElement,
        refNode = docElem.firstElementChild || docElem.firstChild,
        // fakeBody required for <FF4 when executed in <head>
        fakeBody = doc.createElement( "body" ),
        div = doc.createElement( "div" );

    div.id = "mq-test-1";
    div.style.cssText = "position:absolute;top:-100em";
    fakeBody.style.background = "none";
    fakeBody.appendChild(div);

    return function (q) {

      div.innerHTML = "&shy;<style media=\"" + q + "\"> #mq-test-1 { width: 42px; }</style>";

      docElem.insertBefore( fakeBody, refNode );
      bool = div.offsetWidth === 42;
      docElem.removeChild( fakeBody );

      return {
        matches: bool,
        media: q
      };

    };

  }( document ));

  /*
   * jquery.requestAnimationFrame
   * https://github.com/gnarf37/jquery-requestAnimationFrame
   * Requires jQuery 1.8+
   *
   * Copyright (c) 2012 Corey Frang
   * Licensed under the MIT license.
   */

  (function($) {

  // requestAnimationFrame polyfill adapted from Erik Möller
  // fixes from Paul Irish and Tino Zijdel
  // http://paulirish.com/2011/requestanimationframe-for-smart-animating/
  // http://my.opera.com/emoller/blog/2011/12/20/requestanimationframe-for-smart-er-animating

  var animating,
      lastTime = 0,
      vendors = ['webkit', 'moz'],
      requestAnimationFrame = window.requestAnimationFrame,
      cancelAnimationFrame = window.cancelAnimationFrame,
      jqueryFxAvailable = 'undefined' !== typeof jQuery.fx;

  for (; lastTime < vendors.length && !requestAnimationFrame; lastTime++) {
    requestAnimationFrame = window[ vendors[lastTime] + "RequestAnimationFrame" ];
    cancelAnimationFrame = cancelAnimationFrame ||
      window[ vendors[lastTime] + "CancelAnimationFrame" ] ||
      window[ vendors[lastTime] + "CancelRequestAnimationFrame" ];
  }

  function raf() {
    if (animating) {
      requestAnimationFrame(raf);

      if (jqueryFxAvailable) {
        jQuery.fx.tick();
      }
    }
  }

  if (requestAnimationFrame) {
    // use rAF
    window.requestAnimationFrame = requestAnimationFrame;
    window.cancelAnimationFrame = cancelAnimationFrame;

    if (jqueryFxAvailable) {
      jQuery.fx.timer = function (timer) {
        if (timer() && jQuery.timers.push(timer) && !animating) {
          animating = true;
          raf();
        }
      };

      jQuery.fx.stop = function () {
        animating = false;
      };
    }
  } else {
    // polyfill
    window.requestAnimationFrame = function (callback) {
      var currTime = new Date().getTime(),
        timeToCall = Math.max(0, 16 - (currTime - lastTime)),
        id = window.setTimeout(function () {
          callback(currTime + timeToCall);
        }, timeToCall);
      lastTime = currTime + timeToCall;
      return id;
    };

    window.cancelAnimationFrame = function (id) {
      clearTimeout(id);
    };

  }

  }( jQuery ));


  function removeQuotes (string) {
    if (typeof string === 'string' || string instanceof String) {
      string = string.replace(/^['\\/"]+|(;\s?})+|['\\/"]+$/g, '');
    }

    return string;
  }

  window.Foundation = {
    name : 'Foundation',

    version : '5.3.0',

    media_queries : {
      small : S('.foundation-mq-small').css('font-family').replace(/^[\/\\'"]+|(;\s?})+|[\/\\'"]+$/g, ''),
      medium : S('.foundation-mq-medium').css('font-family').replace(/^[\/\\'"]+|(;\s?})+|[\/\\'"]+$/g, ''),
      large : S('.foundation-mq-large').css('font-family').replace(/^[\/\\'"]+|(;\s?})+|[\/\\'"]+$/g, ''),
      xlarge: S('.foundation-mq-xlarge').css('font-family').replace(/^[\/\\'"]+|(;\s?})+|[\/\\'"]+$/g, ''),
      xxlarge: S('.foundation-mq-xxlarge').css('font-family').replace(/^[\/\\'"]+|(;\s?})+|[\/\\'"]+$/g, '')
    },

    stylesheet : $('<style></style>').appendTo('head')[0].sheet,

    global: {
      namespace: undefined
    },

    init : function (scope, libraries, method, options, response) {
      var args = [scope, method, options, response],
          responses = [];

      // check RTL
      this.rtl = /rtl/i.test(S('html').attr('dir'));

      // set foundation global scope
      this.scope = scope || this.scope;

      this.set_namespace();

      if (libraries && typeof libraries === 'string' && !/reflow/i.test(libraries)) {
        if (this.libs.hasOwnProperty(libraries)) {
          responses.push(this.init_lib(libraries, args));
        }
      } else {
        for (var lib in this.libs) {
          responses.push(this.init_lib(lib, libraries));
        }
      }

      return scope;
    },

    init_lib : function (lib, args) {
      if (this.libs.hasOwnProperty(lib)) {
        this.patch(this.libs[lib]);

        if (args && args.hasOwnProperty(lib)) {
            if (typeof this.libs[lib].settings !== 'undefined') {
                $.extend(true, this.libs[lib].settings, args[lib]);
            }
            else if (typeof this.libs[lib].defaults !== 'undefined') {
                $.extend(true, this.libs[lib].defaults, args[lib]);
            }
          return this.libs[lib].init.apply(this.libs[lib], [this.scope, args[lib]]);
        }

        args = args instanceof Array ? args : new Array(args);    // PATCH: added this line
        return this.libs[lib].init.apply(this.libs[lib], args);
      }

      return function () {};
    },

    patch : function (lib) {
      lib.scope = this.scope;
      lib.namespace = this.global.namespace;
      lib.rtl = this.rtl;
      lib['data_options'] = this.utils.data_options;
      lib['attr_name'] = attr_name;
      lib['add_namespace'] = add_namespace;
      lib['bindings'] = bindings;
      lib['S'] = this.utils.S;
    },

    inherit : function (scope, methods) {
      var methods_arr = methods.split(' '),
          i = methods_arr.length;

      while (i--) {
        if (this.utils.hasOwnProperty(methods_arr[i])) {
          scope[methods_arr[i]] = this.utils[methods_arr[i]];
        }
      }
    },

    set_namespace: function () {

      // Description:
      //    Don't bother reading the namespace out of the meta tag
      //    if the namespace has been set globally in javascript
      //
      // Example:
      //    Foundation.global.namespace = 'my-namespace';
      // or make it an empty string:
      //    Foundation.global.namespace = '';
      //
      //

      // If the namespace has not been set (is undefined), try to read it out of the meta element.
      // Otherwise use the globally defined namespace, even if it's empty ('')
      var namespace = ( this.global.namespace === undefined ) ? $('.foundation-data-attribute-namespace').css('font-family') : this.global.namespace;

      // Finally, if the namsepace is either undefined or false, set it to an empty string.
      // Otherwise use the namespace value.
      this.global.namespace = ( namespace === undefined || /false/i.test(namespace) ) ? '' : namespace;
    },

    libs : {},

    // methods that can be inherited in libraries
    utils : {

      // Description:
      //    Fast Selector wrapper returns jQuery object. Only use where getElementById
      //    is not available.
      //
      // Arguments:
      //    Selector (String): CSS selector describing the element(s) to be
      //    returned as a jQuery object.
      //
      //    Scope (String): CSS selector describing the area to be searched. Default
      //    is document.
      //
      // Returns:
      //    Element (jQuery Object): jQuery object containing elements matching the
      //    selector within the scope.
      S : S,

      // Description:
      //    Executes a function a max of once every n milliseconds
      //
      // Arguments:
      //    Func (Function): Function to be throttled.
      //
      //    Delay (Integer): Function execution threshold in milliseconds.
      //
      // Returns:
      //    Lazy_function (Function): Function with throttling applied.
      throttle : function (func, delay) {
        var timer = null;

        return function () {
          var context = this, args = arguments;

          if (timer == null) {
            timer = setTimeout(function () {
              func.apply(context, args);
              timer = null;
            }, delay);
          }
        };
      },

      // Description:
      //    Executes a function when it stops being invoked for n seconds
      //    Modified version of _.debounce() http://underscorejs.org
      //
      // Arguments:
      //    Func (Function): Function to be debounced.
      //
      //    Delay (Integer): Function execution threshold in milliseconds.
      //
      //    Immediate (Bool): Whether the function should be called at the beginning
      //    of the delay instead of the end. Default is false.
      //
      // Returns:
      //    Lazy_function (Function): Function with debouncing applied.
      debounce : function (func, delay, immediate) {
        var timeout, result;
        return function () {
          var context = this, args = arguments;
          var later = function () {
            timeout = null;
            if (!immediate) result = func.apply(context, args);
          };
          var callNow = immediate && !timeout;
          clearTimeout(timeout);
          timeout = setTimeout(later, delay);
          if (callNow) result = func.apply(context, args);
          return result;
        };
      },

      // Description:
      //    Parses data-options attribute
      //
      // Arguments:
      //    El (jQuery Object): Element to be parsed.
      //
      // Returns:
      //    Options (Javascript Object): Contents of the element's data-options
      //    attribute.
      data_options : function (el, data_attr_name) {
        data_attr_name = data_attr_name || 'options';
        var opts = {}, ii, p, opts_arr,
            data_options = function (el) {
              var namespace = Foundation.global.namespace;

              if (namespace.length > 0) {
                return el.data(namespace + '-' + data_attr_name);
              }

              return el.data(data_attr_name);
            };

        var cached_options = data_options(el);

        if (typeof cached_options === 'object') {
          return cached_options;
        }

        opts_arr = (cached_options || ':').split(';');
        ii = opts_arr.length;

        function isNumber (o) {
          return ! isNaN (o-0) && o !== null && o !== "" && o !== false && o !== true;
        }

        function trim (str) {
          if (typeof str === 'string') return $.trim(str);
          return str;
        }

        while (ii--) {
          p = opts_arr[ii].split(':');
          p = [p[0], p.slice(1).join(':')];

          if (/true/i.test(p[1])) p[1] = true;
          if (/false/i.test(p[1])) p[1] = false;
          if (isNumber(p[1])) {
            if (p[1].indexOf('.') === -1) {
              p[1] = parseInt(p[1], 10);
            } else {
              p[1] = parseFloat(p[1]);
            }
          }

          if (p.length === 2 && p[0].length > 0) {
            opts[trim(p[0])] = trim(p[1]);
          }
        }

        return opts;
      },

      // Description:
      //    Adds JS-recognizable media queries
      //
      // Arguments:
      //    Media (String): Key string for the media query to be stored as in
      //    Foundation.media_queries
      //
      //    Class (String): Class name for the generated <meta> tag
      register_media : function (media, media_class) {
        if(Foundation.media_queries[media] === undefined) {
          $('head').append('<meta class="' + media_class + '"/>');
          Foundation.media_queries[media] = removeQuotes($('.' + media_class).css('font-family'));
        }
      },

      // Description:
      //    Add custom CSS within a JS-defined media query
      //
      // Arguments:
      //    Rule (String): CSS rule to be appended to the document.
      //
      //    Media (String): Optional media query string for the CSS rule to be
      //    nested under.
      add_custom_rule : function (rule, media) {
        if (media === undefined && Foundation.stylesheet) {
          Foundation.stylesheet.insertRule(rule, Foundation.stylesheet.cssRules.length);
        } else {
          var query = Foundation.media_queries[media];

          if (query !== undefined) {
            Foundation.stylesheet.insertRule('@media ' +
              Foundation.media_queries[media] + '{ ' + rule + ' }');
          }
        }
      },

      // Description:
      //    Performs a callback function when an image is fully loaded
      //
      // Arguments:
      //    Image (jQuery Object): Image(s) to check if loaded.
      //
      //    Callback (Function): Function to execute when image is fully loaded.
      image_loaded : function (images, callback) {
        var self = this,
            unloaded = images.length;

        if (unloaded === 0) {
          callback(images);
        }

        images.each(function () {
          single_image_loaded(self.S(this), function () {
            unloaded -= 1;
            if (unloaded === 0) {
              callback(images);
            }
          });
        });
      },

      // Description:
      //    Returns a random, alphanumeric string
      //
      // Arguments:
      //    Length (Integer): Length of string to be generated. Defaults to random
      //    integer.
      //
      // Returns:
      //    Rand (String): Pseudo-random, alphanumeric string.
      random_str : function () {
        if (!this.fidx) this.fidx = 0;
        this.prefix = this.prefix || [(this.name || 'F'), (+new Date).toString(36)].join('-');

        return this.prefix + (this.fidx++).toString(36);
      }
    }
  };

  $.fn.foundation = function () {
    var args = Array.prototype.slice.call(arguments, 0);

    return this.each(function () {
      Foundation.init.apply(Foundation, [this].concat(args));
      return this;
    });
  };

}(jQuery, window, window.document));
```
