## Supplementary material for "Comparative pathway enrichment analysis in gastrointestinal cell lines Caco-2, HT-29, HEPG2, and colon fibroblasts using a custom expression panel for tight-junction and cytoskeletal regulatory genes": Sup. Results 1: foundation.magellan.js

```
;(function ($, window, document, undefined) {
  'use strict';

  Foundation.libs['magellan-expedition'] = {
    name : 'magellan-expedition',

    version : '5.3.0',

    settings : {
      active_class: 'active',
      threshold: 0, // pixels from the top of the expedition for it to become fixes
      destination_threshold: 20, // pixels from the top of destination for it to be considered active
      throttle_delay: 30, // calculation throttling to increase framerate
      fixed_top: 0 // top distance in pixels assigend to the fixed element on scroll
    }, 

    init : function (scope, method, options) {
      Foundation.inherit(this, 'throttle');
      this.bindings(method, options);
    },

    events : function () {
      var self = this,
          S = self.S,
          settings = self.settings;

      // initialize expedition offset
      self.set_expedition_position();

      S(self.scope)
        .off('.magellan')
        .on('click.fndtn.magellan', '[' + self.add_namespace('data-magellan-arrival') + '] a[href^="#"]', function (e) {
          e.preventDefault();
          var expedition = $(this).closest('[' + self.attr_name() + ']'),
              settings = expedition.data('magellan-expedition-init'),
              hash = this.hash.split('#').join(''),
              target = $("a[name='"+hash+"']");
          
          if (target.length === 0) {
            target = $('#'+hash);
          }

          // Account for expedition height if fixed position
          var scroll_top = target.offset().top - settings.destination_threshold;
          scroll_top = scroll_top - expedition.outerHeight();

          $('html, body').stop().animate({
            'scrollTop': scroll_top
          }, 700, 'swing', function () {
            if(history.pushState) {
              history.pushState(null, null, '#'+hash);
            }
            else {
              location.hash = '#'+hash;
            }
          });
        })
        .on('scroll.fndtn.magellan', self.throttle(this.check_for_arrivals.bind(this), settings.throttle_delay));
      
      $(window)
        .on('resize.fndtn.magellan', self.throttle(this.set_expedition_position.bind(this), settings.throttle_delay));
    },

    check_for_arrivals : function() {
      var self = this;
      self.update_arrivals();
      self.update_expedition_positions();
    },

    set_expedition_position : function() {
      var self = this;
      $('[' + this.attr_name() + '=fixed]', self.scope).each(function(idx, el) {
        var expedition = $(this),
            settings = expedition.data('magellan-expedition-init'),
            styles = expedition.attr('styles'), // save styles
            top_offset;

        expedition.attr('style', '');
        top_offset = expedition.offset().top + settings.threshold;

        expedition.data(self.data_attr('magellan-top-offset'), top_offset);
        expedition.attr('style', styles);
      });
    },

    update_expedition_positions : function() {
      var self = this,
          window_top_offset = $(window).scrollTop();

      $('[' + this.attr_name() + '=fixed]', self.scope).each(function() {
        var expedition = $(this),
            settings = expedition.data('magellan-expedition-init'),
            top_offset = expedition.data('magellan-top-offset');

        if (window_top_offset >= top_offset) {
          // Placeholder allows height calculations to be consistent even when
          // appearing to switch between fixed/non-fixed placement
          var placeholder = expedition.prev('[' + self.add_namespace('data-magellan-expedition-clone') + ']');
          if (placeholder.length === 0) {
            placeholder = expedition.clone();
            placeholder.removeAttr(self.attr_name());
            placeholder.attr(self.add_namespace('data-magellan-expedition-clone'),'');
            expedition.before(placeholder);
          }
          expedition.css({position:'fixed', top: settings.fixed_top});
        } else {
          expedition.prev('[' + self.add_namespace('data-magellan-expedition-clone') + ']').remove();
          expedition.attr('style','').removeClass('fixed');
        }
      });
    },

    update_arrivals : function() {
      var self = this,
          window_top_offset = $(window).scrollTop();

      $('[' + this.attr_name() + ']', self.scope).each(function() {
        var expedition = $(this),
            settings = expedition.data(self.attr_name(true) + '-init'),
            offsets = self.offsets(expedition, window_top_offset),
            arrivals = expedition.find('[' + self.add_namespace('data-magellan-arrival') + ']'),
            active_item = false;
        offsets.each(function(idx, item) {
          if (item.viewport_offset >= item.top_offset) {
            var arrivals = expedition.find('[' + self.add_namespace('data-magellan-arrival') + ']');
            arrivals.not(item.arrival).removeClass(settings.active_class);
            item.arrival.addClass(settings.active_class);
            active_item = true;
            return true;
          }
        });

        if (!active_item) arrivals.removeClass(settings.active_class);
      });
    },

    offsets : function(expedition, window_offset) {
      var self = this,
          settings = expedition.data(self.attr_name(true) + '-init'),
          viewport_offset = window_offset;

      return expedition.find('[' + self.add_namespace('data-magellan-arrival') + ']').map(function(idx, el) {
        var name = $(this).data(self.data_attr('magellan-arrival')),
            dest = $('[' + self.add_namespace('data-magellan-destination') + '=' + name + ']');
        if (dest.length > 0) {
          var top_offset = dest.offset().top - settings.destination_threshold - expedition.outerHeight();
          return {
            destination : dest,
            arrival : $(this),
            top_offset : top_offset,
            viewport_offset : viewport_offset
          }
        }
      }).sort(function(a, b) {
        if (a.top_offset < b.top_offset) return -1;
        if (a.top_offset > b.top_offset) return 1;
        return 0;
      });
    },

    data_attr: function (str) {
      if (this.namespace.length > 0) {
        return this.namespace + '-' + str;
      }

      return str;
    },

    off : function () {
      this.S(this.scope).off('.magellan');
      this.S(window).off('.magellan');
    },

    reflow : function () {
      var self = this;
      // remove placeholder expeditions used for height calculation purposes
      $('[' + self.add_namespace('data-magellan-expedition-clone') + ']', self.scope).remove();
    }
  };
}(jQuery, window, window.document));
```
