## Supplementary material for "Comparative pathway enrichment analysis in gastrointestinal cell lines Caco-2, HT-29, HEPG2, and colon fibroblasts using a custom expression panel for tight-junction and cytoskeletal regulatory genes": Sup. Results 1: foundation.offcanvas.js

```
;(function ($, window, document, undefined) {
  'use strict';

  Foundation.libs.offcanvas = {
    name : 'offcanvas',

    version : '5.3.0',

    settings : {
      open_method: 'move',
      close_on_click: true
    },

    init : function (scope, method, options) {
      this.bindings(method, options);
    },

    events : function () {
      var self = this,
          S = self.S,
          move_class = '',
          right_postfix = '',
          left_postfix = '';

      if (this.settings.open_method === 'move') {
        move_class = 'move-';
        right_postfix = 'right';
        left_postfix = 'left';
      } else if (this.settings.open_method === 'overlap') {
        move_class = 'offcanvas-overlap';
      }

      S(this.scope).off('.offcanvas')
        .on('click.fndtn.offcanvas', '.left-off-canvas-toggle', function (e) {
          self.click_toggle_class(e, move_class + right_postfix);
        })
        .on('click.fndtn.offcanvas', '.left-off-canvas-menu a', function (e) {
          var settings = self.get_settings(e);
          if (settings.close_on_click) {
            self.hide.call(self, move_class + right_postfix, self.get_wrapper(e));
          }
        })
        .on('click.fndtn.offcanvas', '.right-off-canvas-toggle', function (e) {
          self.click_toggle_class(e, move_class + left_postfix);
        })
        .on('click.fndtn.offcanvas', '.right-off-canvas-menu a', function (e) {
          var settings = self.get_settings(e);
          if (settings.close_on_click) {
            self.hide.call(self, move_class + left_postfix, self.get_wrapper(e));
          }
        })
        .on('click.fndtn.offcanvas', '.exit-off-canvas', function (e) {
          self.click_remove_class(e, move_class + left_postfix);
          if (right_postfix) self.click_remove_class(e, move_class + right_postfix);
        });

    },

    toggle: function(class_name, $off_canvas) {
      $off_canvas = $off_canvas || this.get_wrapper();
      if ($off_canvas.is('.' + class_name)) {
        this.hide(class_name, $off_canvas);
      } else {
        this.show(class_name, $off_canvas);
      }
    },

    show: function(class_name, $off_canvas) {
      $off_canvas = $off_canvas || this.get_wrapper();
      $off_canvas.trigger('open').trigger('open.fndtn.offcanvas');
      $off_canvas.addClass(class_name);
    },

    hide: function(class_name, $off_canvas) {
      $off_canvas = $off_canvas || this.get_wrapper();
      $off_canvas.trigger('close').trigger('close.fndtn.offcanvas');
      $off_canvas.removeClass(class_name);
    },

    click_toggle_class: function(e, class_name) {
      e.preventDefault();
      var $off_canvas = this.get_wrapper(e);
      this.toggle(class_name, $off_canvas);
    },

    click_remove_class: function(e, class_name) {
      e.preventDefault();
      var $off_canvas = this.get_wrapper(e);
      this.hide(class_name, $off_canvas);
    },

    get_settings: function(e) {
      var offcanvas  = this.S(e.target).closest('[' + this.attr_name() + ']');
      return offcanvas.data(this.attr_name(true) + '-init') || this.settings;
    },

    get_wrapper: function(e) {
      var $off_canvas = this.S(e ? e.target : this.scope).closest('.off-canvas-wrap');

      if ($off_canvas.length === 0) {
        $off_canvas = this.S('.off-canvas-wrap');
      }
      return $off_canvas;
    },

    reflow : function () {}
  };
}(jQuery, window, window.document));
```
