## Supplementary material for "Comparative pathway enrichment analysis in gastrointestinal cell lines Caco-2, HT-29, HEPG2, and colon fibroblasts using a custom expression panel for tight-junction and cytoskeletal regulatory genes": Sup. Results 1: foundation.orbit.js

```
;(function ($, window, document, undefined) {
  'use strict';

  var noop = function() {};

  var Orbit = function(el, settings) {
    // Don't reinitialize plugin
    if (el.hasClass(settings.slides_container_class)) {
      return this;
    }

    var self = this,
        container,
        slides_container = el,
        number_container,
        bullets_container,
        timer_container,
        idx = 0,
        animate,
        timer,
        locked = false,
        adjust_height_after = false;


    self.slides = function() {
      return slides_container.children(settings.slide_selector);
    };

    self.slides().first().addClass(settings.active_slide_class);

    self.update_slide_number = function(index) {
      if (settings.slide_number) {
        number_container.find('span:first').text(parseInt(index)+1);
        number_container.find('span:last').text(self.slides().length);
      }
      if (settings.bullets) {
        bullets_container.children().removeClass(settings.bullets_active_class);
        $(bullets_container.children().get(index)).addClass(settings.bullets_active_class);
      }
    };

    self.update_active_link = function(index) {
      var link = $('a[data-orbit-link="'+self.slides().eq(index).attr('data-orbit-slide')+'"]');
      link.siblings().removeClass(settings.bullets_active_class);
      link.addClass(settings.bullets_active_class);
    };

    self.build_markup = function() {
      slides_container.wrap('<div class="'+settings.container_class+'"></div>');
      container = slides_container.parent();
      slides_container.addClass(settings.slides_container_class);
      
      if (settings.stack_on_small) {
        container.addClass(settings.stack_on_small_class);
      }

      if (settings.navigation_arrows) {
        container.append($('<a href="#"><span></span></a>').addClass(settings.prev_class));
        container.append($('<a href="#"><span></span></a>').addClass(settings.next_class));
      }

      if (settings.timer) {
        timer_container = $('<div>').addClass(settings.timer_container_class);
        timer_container.append('<span>');
        timer_container.append($('<div>').addClass(settings.timer_progress_class));
        timer_container.addClass(settings.timer_paused_class);
        container.append(timer_container);
      }

      if (settings.slide_number) {
        number_container = $('<div>').addClass(settings.slide_number_class);
        number_container.append('<span></span> ' + settings.slide_number_text + ' <span></span>');
        container.append(number_container);
      }

      if (settings.bullets) {
        bullets_container = $('<ol>').addClass(settings.bullets_container_class);
        container.append(bullets_container);
        bullets_container.wrap('<div class="orbit-bullets-container"></div>');
        self.slides().each(function(idx, el) {
          var bullet = $('<li>').attr('data-orbit-slide', idx).on('click', self.link_bullet);;
          bullets_container.append(bullet);
        });
      }

    };

    self._goto = function(next_idx, start_timer) {
      // if (locked) {return false;}
      if (next_idx === idx) {return false;}
      if (typeof timer === 'object') {timer.restart();}
      var slides = self.slides();

      var dir = 'next';
      locked = true;
      if (next_idx < idx) {dir = 'prev';}
      if (next_idx >= slides.length) {
        if (!settings.circular) return false;
        next_idx = 0;
      } else if (next_idx < 0) {
        if (!settings.circular) return false;
        next_idx = slides.length - 1;
      }
      
      var current = $(slides.get(idx));
      var next = $(slides.get(next_idx));

      current.css('zIndex', 2);
      current.removeClass(settings.active_slide_class);
      next.css('zIndex', 4).addClass(settings.active_slide_class);

      slides_container.trigger('before-slide-change.fndtn.orbit');
      settings.before_slide_change();
      self.update_active_link(next_idx);
      
      var callback = function() {
        var unlock = function() {
          idx = next_idx;
          locked = false;
          if (start_timer === true) {timer = self.create_timer(); timer.start();}
          self.update_slide_number(idx);
          slides_container.trigger('after-slide-change.fndtn.orbit',[{slide_number: idx, total_slides: slides.length}]);
          settings.after_slide_change(idx, slides.length);
        };
        if (slides_container.height() != next.height() && settings.variable_height) {
          slides_container.animate({'height': next.height()}, 250, 'linear', unlock);
        } else {
          unlock();
        }
      };

      if (slides.length === 1) {callback(); return false;}

      var start_animation = function() {
        if (dir === 'next') {animate.next(current, next, callback);}
        if (dir === 'prev') {animate.prev(current, next, callback);}        
      };

      if (next.height() > slides_container.height() && settings.variable_height) {
        slides_container.animate({'height': next.height()}, 250, 'linear', start_animation);
      } else {
        start_animation();
      }
    };
    
    self.next = function(e) {
      e.stopImmediatePropagation();
      e.preventDefault();
      self._goto(idx + 1);
    };
    
    self.prev = function(e) {
      e.stopImmediatePropagation();
      e.preventDefault();
      self._goto(idx - 1);
    };

    self.link_custom = function(e) {
      e.preventDefault();
      var link = $(this).attr('data-orbit-link');
      if ((typeof link === 'string') && (link = $.trim(link)) != "") {
        var slide = container.find('[data-orbit-slide='+link+']');
        if (slide.index() != -1) {self._goto(slide.index());}
      }
    };

    self.link_bullet = function(e) {    
      var index = $(this).attr('data-orbit-slide');
      if ((typeof index === 'string') && (index = $.trim(index)) != "") {
        if(isNaN(parseInt(index)))
        {
          var slide = container.find('[data-orbit-slide='+index+']');
          if (slide.index() != -1) {self._goto(slide.index() + 1);}
        }
        else
        {
          self._goto(parseInt(index));
        }
      }

    }

    self.timer_callback = function() {
      self._goto(idx + 1, true);
    }
    
    self.compute_dimensions = function() {
      var current = $(self.slides().get(idx));
      var h = current.height();
      if (!settings.variable_height) {
        self.slides().each(function(){
          if ($(this).height() > h) { h = $(this).height(); }
        });
      }
      slides_container.height(h);
    };

    self.create_timer = function() {
      var t = new Timer(
        container.find('.'+settings.timer_container_class), 
        settings, 
        self.timer_callback
      );
      return t;
    };

    self.stop_timer = function() {
      if (typeof timer === 'object') timer.stop();
    };

    self.toggle_timer = function() {
      var t = container.find('.'+settings.timer_container_class);
      if (t.hasClass(settings.timer_paused_class)) {
        if (typeof timer === 'undefined') {timer = self.create_timer();}
        timer.start();     
      }
      else {
        if (typeof timer === 'object') {timer.stop();}
      }
    };

    self.init = function() {
      self.build_markup();
      if (settings.timer) {
        timer = self.create_timer(); 
        Foundation.utils.image_loaded(this.slides().children('img'), timer.start);
      }
      animate = new FadeAnimation(settings, slides_container);
      if (settings.animation === 'slide') 
        animate = new SlideAnimation(settings, slides_container);        

      container.on('click', '.'+settings.next_class, self.next);
      container.on('click', '.'+settings.prev_class, self.prev);

      if (settings.next_on_click) {
        container.on('click', '.'+settings.slides_container_class+' [data-orbit-slide]', self.link_bullet);
      }

      container.on('click', self.toggle_timer);
      if (settings.swipe) {
        container.on('touchstart.fndtn.orbit', function(e) {
          if (!e.touches) {e = e.originalEvent;}
          var data = {
            start_page_x: e.touches[0].pageX,
            start_page_y: e.touches[0].pageY,
            start_time: (new Date()).getTime(),
            delta_x: 0,
            is_scrolling: undefined
          };
          container.data('swipe-transition', data);
          e.stopPropagation();
        })
        .on('touchmove.fndtn.orbit', function(e) {
          if (!e.touches) { e = e.originalEvent; }
          // Ignore pinch/zoom events
          if(e.touches.length > 1 || e.scale && e.scale !== 1) return;

          var data = container.data('swipe-transition');
          if (typeof data === 'undefined') {data = {};}

          data.delta_x = e.touches[0].pageX - data.start_page_x;

          if ( typeof data.is_scrolling === 'undefined') {
            data.is_scrolling = !!( data.is_scrolling || Math.abs(data.delta_x) < Math.abs(e.touches[0].pageY - data.start_page_y) );
          }

          if (!data.is_scrolling && !data.active) {
            e.preventDefault();
            var direction = (data.delta_x < 0) ? (idx+1) : (idx-1);
            data.active = true;
            self._goto(direction);
          }
        })
        .on('touchend.fndtn.orbit', function(e) {
          container.data('swipe-transition', {});
          e.stopPropagation();
        })
      }
      container.on('mouseenter.fndtn.orbit', function(e) {
        if (settings.timer && settings.pause_on_hover) {
          self.stop_timer();
        }
      })
      .on('mouseleave.fndtn.orbit', function(e) {
        if (settings.timer && settings.resume_on_mouseout) {
          timer.start();
        }
      });
      
      $(document).on('click', '[data-orbit-link]', self.link_custom);
      $(window).on('load resize', self.compute_dimensions);
      Foundation.utils.image_loaded(this.slides().children('img'), self.compute_dimensions);
      Foundation.utils.image_loaded(this.slides().children('img'), function() {
        container.prev('.'+settings.preloader_class).css('display', 'none');
        self.update_slide_number(0);
        self.update_active_link(0);
        slides_container.trigger('ready.fndtn.orbit');
      });
    };

    self.init();
  };

  var Timer = function(el, settings, callback) {
    var self = this,
        duration = settings.timer_speed,
        progress = el.find('.'+settings.timer_progress_class),
        start, 
        timeout,
        left = -1;

    this.update_progress = function(w) {
      var new_progress = progress.clone();
      new_progress.attr('style', '');
      new_progress.css('width', w+'%');
      progress.replaceWith(new_progress);
      progress = new_progress;
    };

    this.restart = function() {
      clearTimeout(timeout);
      el.addClass(settings.timer_paused_class);
      left = -1;
      self.update_progress(0);
    };

    this.start = function() {
      if (!el.hasClass(settings.timer_paused_class)) {return true;}
      left = (left === -1) ? duration : left;
      el.removeClass(settings.timer_paused_class);
      start = new Date().getTime();
      progress.animate({'width': '100%'}, left, 'linear');
      timeout = setTimeout(function() {
        self.restart();
        callback();
      }, left);
      el.trigger('timer-started.fndtn.orbit')
    };

    this.stop = function() {
      if (el.hasClass(settings.timer_paused_class)) {return true;}
      clearTimeout(timeout);
      el.addClass(settings.timer_paused_class);
      var end = new Date().getTime();
      left = left - (end - start);
      var w = 100 - ((left / duration) * 100);
      self.update_progress(w);
      el.trigger('timer-stopped.fndtn.orbit');
    };
  };
  
  var SlideAnimation = function(settings, container) {
    var duration = settings.animation_speed;
    var is_rtl = ($('html[dir=rtl]').length === 1);
    var margin = is_rtl ? 'marginRight' : 'marginLeft';
    var animMargin = {};
    animMargin[margin] = '0%';

    this.next = function(current, next, callback) {
      current.animate({marginLeft:'-100%'}, duration);
      next.animate(animMargin, duration, function() {
        current.css(margin, '100%');
        callback();
      });
    };

    this.prev = function(current, prev, callback) {
      current.animate({marginLeft:'100%'}, duration);
      prev.css(margin, '-100%');
      prev.animate(animMargin, duration, function() {
        current.css(margin, '100%');
        callback();
      });
    };
  };

  var FadeAnimation = function(settings, container) {
    var duration = settings.animation_speed;
    var is_rtl = ($('html[dir=rtl]').length === 1);
    var margin = is_rtl ? 'marginRight' : 'marginLeft';

    this.next = function(current, next, callback) {
      next.css({'margin':'0%', 'opacity':'0.01'});
      next.animate({'opacity':'1'}, duration, 'linear', function() {
        current.css('margin', '100%');
        callback();
      });
    };

    this.prev = function(current, prev, callback) {
      prev.css({'margin':'0%', 'opacity':'0.01'});
      prev.animate({'opacity':'1'}, duration, 'linear', function() {
        current.css('margin', '100%');
        callback();
      });
    };
  };


  Foundation.libs = Foundation.libs || {};

  Foundation.libs.orbit = {
    name: 'orbit',

    version: '5.3.0',

    settings: {
      animation: 'slide',
      timer_speed: 10000,
      pause_on_hover: true,
      resume_on_mouseout: false,
      next_on_click: true,
      animation_speed: 500,
      stack_on_small: false,
      navigation_arrows: true,
      slide_number: true,
      slide_number_text: 'of',
      container_class: 'orbit-container',
      stack_on_small_class: 'orbit-stack-on-small',
      next_class: 'orbit-next',
      prev_class: 'orbit-prev',
      timer_container_class: 'orbit-timer',
      timer_paused_class: 'paused',
      timer_progress_class: 'orbit-progress',
      slides_container_class: 'orbit-slides-container',
      preloader_class: 'preloader',
      slide_selector: '*',
      bullets_container_class: 'orbit-bullets',
      bullets_active_class: 'active',
      slide_number_class: 'orbit-slide-number',
      caption_class: 'orbit-caption',
      active_slide_class: 'active',
      orbit_transition_class: 'orbit-transitioning',
      bullets: true,
      circular: true,
      timer: true,
      variable_height: false,
      swipe: true,
      before_slide_change: noop,
      after_slide_change: noop
    },

    init : function (scope, method, options) {
      var self = this;
      this.bindings(method, options);
    },

    events : function (instance) {
      var orbit_instance = new Orbit(this.S(instance), this.S(instance).data('orbit-init'));
      this.S(instance).data(self.name + '-instance', orbit_instance);
    },

    reflow : function () {
      var self = this;

      if (self.S(self.scope).is('[data-orbit]')) {
        var $el = self.S(self.scope);
        var instance = $el.data(self.name + '-instance');
        instance.compute_dimensions();
      } else {
        self.S('[data-orbit]', self.scope).each(function(idx, el) {
          var $el = self.S(el);
          var opts = self.data_options($el);
          var instance = $el.data(self.name + '-instance');
          instance.compute_dimensions();
        });
      }
    }
  };

    
}(jQuery, window, window.document));
```
