## Supplementary material for "Comparative pathway enrichment analysis in gastrointestinal cell lines Caco-2, HT-29, HEPG2, and colon fibroblasts using a custom expression panel for tight-junction and cytoskeletal regulatory genes": Sup. Results 1: foundation.reveal.js

```
;(function ($, window, document, undefined) {
  'use strict';

  Foundation.libs.reveal = {
    name : 'reveal',

    version : '5.3.0',

    locked : false,

    settings : {
      animation: 'fadeAndPop',
      animation_speed: 250,
      close_on_background_click: true,
      close_on_esc: true,
      dismiss_modal_class: 'close-reveal-modal',
      bg_class: 'reveal-modal-bg',
      root_element: 'body',
      open: function(){},
      opened: function(){},
      close: function(){},
      closed: function(){},
      bg : $('.reveal-modal-bg'),
      css : {
        open : {
          'opacity': 0,
          'visibility': 'visible',
          'display' : 'block'
        },
        close : {
          'opacity': 1,
          'visibility': 'hidden',
          'display': 'none'
        }
      }
    },

    init : function (scope, method, options) {
      $.extend(true, this.settings, method, options);
      this.bindings(method, options);
    },

    events : function (scope) {
      var self = this,
          S = self.S;

      S(this.scope)
        .off('.reveal')
        .on('click.fndtn.reveal', '[' + this.add_namespace('data-reveal-id') + ']:not([disabled])', function (e) {
          e.preventDefault();

          if (!self.locked) {
            var element = S(this),
                ajax = element.data(self.data_attr('reveal-ajax'));

            self.locked = true;

            if (typeof ajax === 'undefined') {
              self.open.call(self, element);
            } else {
              var url = ajax === true ? element.attr('href') : ajax;

              self.open.call(self, element, {url: url});
            }
          }
        });

      S(document)
        .on('touchend.fndtn.reveal click.fndtn.reveal', this.close_targets(), function (e) {

          e.preventDefault();

          if (!self.locked) {
            var settings = S('[' + self.attr_name() + '].open').data(self.attr_name(true) + '-init'),
                bg_clicked = S(e.target)[0] === S('.' + settings.bg_class)[0];

            if (bg_clicked) {
              if (settings.close_on_background_click) {
                e.stopPropagation();
              } else {
                return;
              }
            }

            self.locked = true;
            self.close.call(self, bg_clicked ? S('[' + self.attr_name() + '].open') : S(this).closest('[' + self.attr_name() + ']'));
          }
        });

      if(S('[' + self.attr_name() + ']', this.scope).length > 0) {
        S(this.scope)
          // .off('.reveal')
          .on('open.fndtn.reveal', this.settings.open)
          .on('opened.fndtn.reveal', this.settings.opened)
          .on('opened.fndtn.reveal', this.open_video)
          .on('close.fndtn.reveal', this.settings.close)
          .on('closed.fndtn.reveal', this.settings.closed)
          .on('closed.fndtn.reveal', this.close_video);
      } else {
        S(this.scope)
          // .off('.reveal')
          .on('open.fndtn.reveal', '[' + self.attr_name() + ']', this.settings.open)
          .on('opened.fndtn.reveal', '[' + self.attr_name() + ']', this.settings.opened)
          .on('opened.fndtn.reveal', '[' + self.attr_name() + ']', this.open_video)
          .on('close.fndtn.reveal', '[' + self.attr_name() + ']', this.settings.close)
          .on('closed.fndtn.reveal', '[' + self.attr_name() + ']', this.settings.closed)
          .on('closed.fndtn.reveal', '[' + self.attr_name() + ']', this.close_video);
      }

      return true;
    },

    // PATCH #3: turning on key up capture only when a reveal window is open
    key_up_on : function (scope) {
      var self = this;

      // PATCH #1: fixing multiple keyup event trigger from single key press
      self.S('body').off('keyup.fndtn.reveal').on('keyup.fndtn.reveal', function ( event ) {
        var open_modal = self.S('[' + self.attr_name() + '].open'),
            settings = open_modal.data(self.attr_name(true) + '-init');
        // PATCH #2: making sure that the close event can be called only while unlocked,
        //           so that multiple keyup.fndtn.reveal events don't prevent clean closing of the reveal window.
        if ( settings && event.which === 27  && settings.close_on_esc && !self.locked) { // 27 is the keycode for the Escape key
          self.close.call(self, open_modal);
        }
      });

      return true;
    },

    // PATCH #3: turning on key up capture only when a reveal window is open
    key_up_off : function (scope) {
      this.S('body').off('keyup.fndtn.reveal');
      return true;
    },

    open : function (target, ajax_settings) {
      var self = this,
          modal;

      if (target) {
        if (typeof target.selector !== 'undefined') {
          // Find the named node; only use the first one found, since the rest of the code assumes there's only one node
          modal = self.S('#' + target.data(self.data_attr('reveal-id'))).first();
        } else {
          modal = self.S(this.scope);

          ajax_settings = target;
        }
      } else {
        modal = self.S(this.scope);
      }

      var settings = modal.data(self.attr_name(true) + '-init');
      settings = settings || this.settings;

      if (!modal.hasClass('open')) {
        var open_modal = self.S('[' + self.attr_name() + '].open');

        if (typeof modal.data('css-top') === 'undefined') {
          modal.data('css-top', parseInt(modal.css('top'), 10))
            .data('offset', this.cache_offset(modal));
        }

        this.key_up_on(modal);    // PATCH #3: turning on key up capture only when a reveal window is open
        modal.trigger('open').trigger('open.fndtn.reveal');

        if (open_modal.length < 1) {
          this.toggle_bg(modal, true);
        }

        if (typeof ajax_settings === 'string') {
          ajax_settings = {
            url: ajax_settings
          };
        }

        if (typeof ajax_settings === 'undefined' || !ajax_settings.url) {
          if (open_modal.length > 0) {
            this.hide(open_modal, settings.css.close);
          }

          this.show(modal, settings.css.open);
        } else {
          var old_success = typeof ajax_settings.success !== 'undefined' ? ajax_settings.success : null;

          $.extend(ajax_settings, {
            success: function (data, textStatus, jqXHR) {
              if ( $.isFunction(old_success) ) {
                old_success(data, textStatus, jqXHR);
              }

              modal.html(data);
              self.S(modal).foundation('section', 'reflow');
              self.S(modal).children().foundation();

              if (open_modal.length > 0) {
                self.hide(open_modal, settings.css.close);
              }
              self.show(modal, settings.css.open);
            }
          });

          $.ajax(ajax_settings);
        }
      }
    },

    close : function (modal) {
      var modal = modal && modal.length ? modal : this.S(this.scope),
          open_modals = this.S('[' + this.attr_name() + '].open'),
          settings = modal.data(this.attr_name(true) + '-init') || this.settings;

      if (open_modals.length > 0) {
        this.locked = true;
        this.key_up_off(modal);   // PATCH #3: turning on key up capture only when a reveal window is open
        modal.trigger('close').trigger('close.fndtn.reveal');
        this.toggle_bg(modal, false);
        this.hide(open_modals, settings.css.close, settings);
      }
    },

    close_targets : function () {
      var base = '.' + this.settings.dismiss_modal_class;

      if (this.settings.close_on_background_click) {
        return base + ', .' + this.settings.bg_class;
      }

      return base;
    },

    toggle_bg : function (modal, state) {
      if (this.S('.' + this.settings.bg_class).length === 0) {
        this.settings.bg = $('<div />', {'class': this.settings.bg_class})
          .appendTo('body').hide();
      }

      var visible = this.settings.bg.filter(':visible').length > 0;
      if ( state != visible ) {
        if ( state == undefined ? visible : !state ) {
          this.hide(this.settings.bg);
        } else {
          this.show(this.settings.bg);
        }
      }
    },

    show : function (el, css) {
      // is modal
      if (css) {
        var settings = el.data(this.attr_name(true) + '-init') || this.settings,
            root_element = settings.root_element;

        if (el.parent(root_element).length === 0) {
          var placeholder = el.wrap('<div style="display: none;" />').parent();

          el.on('closed.fndtn.reveal.wrapped', function() {
            el.detach().appendTo(placeholder);
            el.unwrap().unbind('closed.fndtn.reveal.wrapped');
          });

          el.detach().appendTo(root_element);
        }

        var animData = getAnimationData(settings.animation);
        if (!animData.animate) {
          this.locked = false;
        }
        if (animData.pop) {
          css.top = $(window).scrollTop() - el.data('offset') + 'px';
          var end_css = {
            top: $(window).scrollTop() + el.data('css-top') + 'px',
            opacity: 1
          };

          return setTimeout(function () {
            return el
              .css(css)
              .animate(end_css, settings.animation_speed, 'linear', function () {
                this.locked = false;
                el.trigger('opened').trigger('opened.fndtn.reveal');
              }.bind(this))
              .addClass('open');
          }.bind(this), settings.animation_speed / 2);
        }

        if (animData.fade) {
          css.top = $(window).scrollTop() + el.data('css-top') + 'px';
          var end_css = {opacity: 1};

          return setTimeout(function () {
            return el
              .css(css)
              .animate(end_css, settings.animation_speed, 'linear', function () {
                this.locked = false;
                el.trigger('opened').trigger('opened.fndtn.reveal');
              }.bind(this))
              .addClass('open');
          }.bind(this), settings.animation_speed / 2);
        }

        return el.css(css).show().css({opacity: 1}).addClass('open').trigger('opened').trigger('opened.fndtn.reveal');
      }

      var settings = this.settings;

      // should we animate the background?
      if (getAnimationData(settings.animation).fade) {
        return el.fadeIn(settings.animation_speed / 2);
      }

      this.locked = false;

      return el.show();
    },

    hide : function (el, css) {
      // is modal
      if (css) {
        var settings = el.data(this.attr_name(true) + '-init');
        settings = settings || this.settings;

        var animData = getAnimationData(settings.animation);
        if (!animData.animate) {
          this.locked = false;
        }
        if (animData.pop) {
          var end_css = {
            top: - $(window).scrollTop() - el.data('offset') + 'px',
            opacity: 0
          };

          return setTimeout(function () {
            return el
              .animate(end_css, settings.animation_speed, 'linear', function () {
                this.locked = false;
                el.css(css).trigger('closed').trigger('closed.fndtn.reveal');
              }.bind(this))
              .removeClass('open');
          }.bind(this), settings.animation_speed / 2);
        }

        if (animData.fade) {
          var end_css = {opacity: 0};

          return setTimeout(function () {
            return el
              .animate(end_css, settings.animation_speed, 'linear', function () {
                this.locked = false;
                el.css(css).trigger('closed').trigger('closed.fndtn.reveal');
              }.bind(this))
              .removeClass('open');
          }.bind(this), settings.animation_speed / 2);
        }

        return el.hide().css(css).removeClass('open').trigger('closed').trigger('closed.fndtn.reveal');
      }

      var settings = this.settings;

      // should we animate the background?
      if (getAnimationData(settings.animation).fade) {
        return el.fadeOut(settings.animation_speed / 2);
      }

      return el.hide();
    },

    close_video : function (e) {
      var video = $('.flex-video', e.target),
          iframe = $('iframe', video);

      if (iframe.length > 0) {
        iframe.attr('data-src', iframe[0].src);
        iframe.attr('src', 'about:blank');
        video.hide();
      }
    },

    open_video : function (e) {
      var video = $('.flex-video', e.target),
          iframe = video.find('iframe');

      if (iframe.length > 0) {
        var data_src = iframe.attr('data-src');
        if (typeof data_src === 'string') {
          iframe[0].src = iframe.attr('data-src');
        } else {
          var src = iframe[0].src;
          iframe[0].src = undefined;
          iframe[0].src = src;
        }
        video.show();
      }
    },

    data_attr: function (str) {
      if (this.namespace.length > 0) {
        return this.namespace + '-' + str;
      }

      return str;
    },

    cache_offset : function (modal) {
      var offset = modal.show().height() + parseInt(modal.css('top'), 10);

      modal.hide();

      return offset;
    },

    off : function () {
      $(this.scope).off('.fndtn.reveal');
    },

    reflow : function () {}
  };

  /*
   * getAnimationData('popAndFade') // {animate: true,  pop: true,  fade: true}
   * getAnimationData('fade')       // {animate: true,  pop: false, fade: true}
   * getAnimationData('pop')        // {animate: true,  pop: true,  fade: false}
   * getAnimationData('foo')        // {animate: false, pop: false, fade: false}
   * getAnimationData(null)         // {animate: false, pop: false, fade: false}
   */
  function getAnimationData(str) {
    var fade = /fade/i.test(str);
    var pop = /pop/i.test(str);
    return {
      animate: fade || pop,
      pop: pop,
      fade: fade
    };
  }
}(jQuery, window, window.document));
```
