## Supplementary material for "Comparative pathway enrichment analysis in gastrointestinal cell lines Caco-2, HT-29, HEPG2, and colon fibroblasts using a custom expression panel for tight-junction and cytoskeletal regulatory genes": Sup. Results 1: foundation.slider.js

```
;(function ($, window, document, undefined) {
  'use strict';

  Foundation.libs.slider = {
    name : 'slider',

    version : '5.3.0',

    settings: {
      start: 0,
      end: 100,
      step: 1,
      initial: null,
      display_selector: '',
      vertical: false,
      on_change: function(){}
    },

    cache : {},

    init : function (scope, method, options) {
      Foundation.inherit(this,'throttle');
      this.bindings(method, options);
      this.reflow();
    },

    events : function() {
      var self = this;

      $(this.scope)
        .off('.slider')
        .on('mousedown.fndtn.slider touchstart.fndtn.slider pointerdown.fndtn.slider',
        '[' + self.attr_name() + ']:not(.disabled, [disabled]) .range-slider-handle', function(e) {
          if (!self.cache.active) {
            e.preventDefault();
            self.set_active_slider($(e.target));
          }
        })
        .on('mousemove.fndtn.slider touchmove.fndtn.slider pointermove.fndtn.slider', function(e) {
          if (!!self.cache.active) {
            e.preventDefault();
            if ($.data(self.cache.active[0], 'settings').vertical) {
              self.calculate_position(self.cache.active, e.pageY || 
                                                         e.originalEvent.clientY || 
                                                         e.originalEvent.touches[0].clientY || 
                                                         e.currentPoint.y);
            } else {
              self.calculate_position(self.cache.active, e.pageX || 
                                                         e.originalEvent.clientX || 
                                                         e.originalEvent.touches[0].clientX || 
                                                         e.currentPoint.x);
            }
          }
        })
        .on('mouseup.fndtn.slider touchend.fndtn.slider pointerup.fndtn.slider', function(e) {
          self.remove_active_slider();
        })
        .on('change.fndtn.slider', function(e) {
          self.settings.on_change();
        });

      self.S(window)
        .on('resize.fndtn.slider', self.throttle(function(e) {
          self.reflow();
        }, 300));
    },

    set_active_slider : function($handle) {
      this.cache.active = $handle;
    },

    remove_active_slider : function() {
      this.cache.active = null;
    },

    calculate_position : function($handle, cursor_x) {
      var self = this,
          settings = $.data($handle[0], 'settings'),
          handle_l = $.data($handle[0], 'handle_l'),
          handle_o = $.data($handle[0], 'handle_o'),
          bar_l = $.data($handle[0], 'bar_l'),
          bar_o = $.data($handle[0], 'bar_o');

      requestAnimationFrame(function(){
        var pct;

        if (Foundation.rtl && !settings.vertical) {
          pct = self.limit_to(((bar_o+bar_l-cursor_x)/bar_l),0,1);
        } else {
          pct = self.limit_to(((cursor_x-bar_o)/bar_l),0,1);
        }

        pct = settings.vertical ? 1-pct : pct;

        var norm = self.normalized_value(pct, settings.start, settings.end, settings.step);

        self.set_ui($handle, norm);
      });
    },

    set_ui : function($handle, value) {
      var settings = $.data($handle[0], 'settings'),
          handle_l = $.data($handle[0], 'handle_l'),
          bar_l = $.data($handle[0], 'bar_l'),
          norm_pct = this.normalized_percentage(value, settings.start, settings.end),
          handle_offset = norm_pct*(bar_l-handle_l)-1,
          progress_bar_length = norm_pct*100;

      if (Foundation.rtl && !settings.vertical) {
        handle_offset = -handle_offset;
      }

      handle_offset = settings.vertical ? -handle_offset + bar_l - handle_l + 1 : handle_offset;
      this.set_translate($handle, handle_offset, settings.vertical);

      if (settings.vertical) {
        $handle.siblings('.range-slider-active-segment').css('height', progress_bar_length + '%');
      } else {
        $handle.siblings('.range-slider-active-segment').css('width', progress_bar_length + '%');
      }

      $handle.parent().attr(this.attr_name(), value).trigger('change').trigger('change.fndtn.slider');

      $handle.parent().children('input[type=hidden]').val(value);

      if (settings.input_id != '') {
        $(settings.display_selector).each(function(){
          if (this.hasOwnProperty('value')) {
            $(this).val(value);
          } else {
            $(this).text(value);
          }
        });
      }

    },

    normalized_percentage : function(val, start, end) {
      return (val - start)/(end - start);
    },

    normalized_value : function(val, start, end, step) {
      var range = end - start,
          point = val*range,
          mod = (point-(point%step)) / step,
          rem = point % step,
          round = ( rem >= step*0.5 ? step : 0);
      return (mod*step + round) + start;
    },

    set_translate : function(ele, offset, vertical) {
      if (vertical) {
        $(ele)
          .css('-webkit-transform', 'translateY('+offset+'px)')
          .css('-moz-transform', 'translateY('+offset+'px)')
          .css('-ms-transform', 'translateY('+offset+'px)')
          .css('-o-transform', 'translateY('+offset+'px)')
          .css('transform', 'translateY('+offset+'px)');
      } else {
        $(ele)
          .css('-webkit-transform', 'translateX('+offset+'px)')
          .css('-moz-transform', 'translateX('+offset+'px)')
          .css('-ms-transform', 'translateX('+offset+'px)')
          .css('-o-transform', 'translateX('+offset+'px)')
          .css('transform', 'translateX('+offset+'px)');
      }
    },

    limit_to : function(val, min, max) {
      return Math.min(Math.max(val, min), max);
    },

    initialize_settings : function(handle) {
      var settings = $.extend({}, this.settings, this.data_options($(handle).parent()));

      if (settings.vertical) {
        $.data(handle, 'bar_o', $(handle).parent().offset().top);
        $.data(handle, 'bar_l', $(handle).parent().outerHeight());
        $.data(handle, 'handle_o', $(handle).offset().top);
        $.data(handle, 'handle_l', $(handle).outerHeight());
      } else {
        $.data(handle, 'bar_o', $(handle).parent().offset().left);
        $.data(handle, 'bar_l', $(handle).parent().outerWidth());
        $.data(handle, 'handle_o', $(handle).offset().left);
        $.data(handle, 'handle_l', $(handle).outerWidth());
      }

      $.data(handle, 'bar', $(handle).parent());
      $.data(handle, 'settings', settings);
    },

    set_initial_position : function($ele) {
      var settings = $.data($ele.children('.range-slider-handle')[0], 'settings'),
          initial = (!!settings.initial ? settings.initial : Math.floor((settings.end-settings.start)*0.5/settings.step)*settings.step+settings.start),
          $handle = $ele.children('.range-slider-handle');
      this.set_ui($handle, initial);
    },

    set_value : function(value) {
      var self = this;
      $('[' + self.attr_name() + ']', this.scope).each(function(){
        $(this).attr(self.attr_name(), value);
      });
      if (!!$(this.scope).attr(self.attr_name())) {
        $(this.scope).attr(self.attr_name(), value);
      }
      self.reflow();
    },

    reflow : function() {
      var self = this;
      self.S('[' + this.attr_name() + ']').each(function() {
        var handle = $(this).children('.range-slider-handle')[0],
            val = $(this).attr(self.attr_name());
        self.initialize_settings(handle);

        if (val) {
          self.set_ui($(handle), parseFloat(val));
        } else {
          self.set_initial_position($(this));
        }
      });
    }
  };

}(jQuery, window, window.document));
```
