## Supplementary material for "Comparative pathway enrichment analysis in gastrointestinal cell lines Caco-2, HT-29, HEPG2, and colon fibroblasts using a custom expression panel for tight-junction and cytoskeletal regulatory genes": Sup. Results 1: foundation.tab.js

```
;(function ($, window, document, undefined) {
  'use strict';

  Foundation.libs.tab = {
    name : 'tab',

    version : '5.3.0',

    settings : {
      active_class: 'active',
      callback : function () {},
      deep_linking: false,
      scroll_to_content: true,
      is_hover: false
    },

    default_tab_hashes: [],

    init : function (scope, method, options) {
      var self = this,
          S = this.S;

      this.bindings(method, options);
      this.handle_location_hash_change();

      // Store the default active tabs which will be referenced when the
      // location hash is absent, as in the case of navigating the tabs and
      // returning to the first viewing via the browser Back button.
      S('[' + this.attr_name() + '] > .active > a', this.scope).each(function () {
        self.default_tab_hashes.push(this.hash);
      });
    },

    events : function () {
      var self = this,
          S = this.S;

      S(this.scope)
        .off('.tab')
        // Click event: tab title
        .on('click.fndtn.tab', '[' + this.attr_name() + '] > * > a', function (e) {
          var settings = S(this).closest('[' + self.attr_name() +']').data(self.attr_name(true) + '-init');
          if (!settings.is_hover || Modernizr.touch) {
            e.preventDefault();
            e.stopPropagation();
            self.toggle_active_tab(S(this).parent());
          }
        })
        // Hover event: tab title
        .on('mouseenter.fndtn.tab', '[' + this.attr_name() + '] > * > a', function (e) {
          var settings = S(this).closest('[' + self.attr_name() +']').data(self.attr_name(true) + '-init');
          if (settings.is_hover) self.toggle_active_tab(S(this).parent());
        });

      // Location hash change event
      S(window).on('hashchange.fndtn.tab', function (e) {
        e.preventDefault();
        self.handle_location_hash_change();
      });
    },

    handle_location_hash_change : function () {
      var self = this,
          S = this.S;

      S('[' + this.attr_name() + ']', this.scope).each(function () {
        var settings = S(this).data(self.attr_name(true) + '-init');
        if (settings.deep_linking) {
          // Match the location hash to a label
          var hash = self.scope.location.hash;
          if (hash != '') {
            // Check whether the location hash references a tab content div or
            // another element on the page (inside or outside the tab content div)
            var hash_element = S(hash);
            if (hash_element.hasClass('content') && hash_element.parent().hasClass('tab-content')) {
              // Tab content div
              self.toggle_active_tab($('[' + self.attr_name() + '] > * > a[href=' + hash + ']').parent());
            } else {
              // Not the tab content div. If inside the tab content, find the
              // containing tab and toggle it as active.
              var hash_tab_container_id = hash_element.closest('.content').attr('id');
              if (hash_tab_container_id != undefined) {
                self.toggle_active_tab($('[' + self.attr_name() + '] > * > a[href=#' + hash_tab_container_id + ']').parent(), hash);
              }
            }
          } else {
            // Reference the default tab hashes which were initialized in the init function
            for (var ind in self.default_tab_hashes) {
              self.toggle_active_tab($('[' + self.attr_name() + '] > * > a[href=' + self.default_tab_hashes[ind] + ']').parent());
            }
          }
        }
       });
     },

    toggle_active_tab: function (tab, location_hash) {
      var S = this.S,
          tabs = tab.closest('[' + this.attr_name() + ']'),
          anchor = tab.children('a').first(),
          target_hash = '#' + anchor.attr('href').split('#')[1],
          target = S(target_hash),
          siblings = tab.siblings(),
          settings = tabs.data(this.attr_name(true) + '-init');

      // allow usage of data-tab-content attribute instead of href
      if (S(this).data(this.data_attr('tab-content'))) {
        target_hash = '#' + S(this).data(this.data_attr('tab-content')).split('#')[1];
        target = S(target_hash);
      }

      if (settings.deep_linking) {
        // Get the scroll Y position prior to moving to the hash ID
        var cur_ypos = $('body,html').scrollTop();

        // Update the location hash to preserve browser history
        // Note that the hash does not need to correspond to the
        // tab content ID anchor; it can be an ID inside or outside of the tab
        // content div.
        if (location_hash != undefined) {
          window.location.hash = location_hash;
        } else {
          window.location.hash = target_hash;
        }

        if (settings.scroll_to_content) {
          // If the user is requesting the content of a tab, then scroll to the
          // top of the title area; otherwise, scroll to the element within
          // the content area as defined by the hash value.
          if (location_hash == undefined || location_hash == target_hash) {
            tab.parent()[0].scrollIntoView();
          } else {
            S(target_hash)[0].scrollIntoView();
          }
        } else {
          // Adjust the scrollbar to the Y position prior to setting the hash
          // Only do this for the tab content anchor, otherwise there will be
          // conflicts with in-tab anchor links nested in the tab-content div
          if (location_hash == undefined || location_hash == target_hash) {
            $('body,html').scrollTop(cur_ypos);
          }
        }
      }

      // WARNING: The activation and deactivation of the tab content must
      // occur after the deep linking in order to properly refresh the browser
      // window (notably in Chrome).
      tab.addClass(settings.active_class).triggerHandler('opened');
      siblings.removeClass(settings.active_class);
      target.siblings().removeClass(settings.active_class).end().addClass(settings.active_class);
      settings.callback(tab);
      target.triggerHandler('toggled', [tab]);
      tabs.triggerHandler('toggled', [target]);
    },

    data_attr: function (str) {
      if (this.namespace.length > 0) {
        return this.namespace + '-' + str;
      }

      return str;
    },

    off : function () {},

    reflow : function () {}
  };
}(jQuery, window, window.document));
```
