## Supplementary material for "Comparative pathway enrichment analysis in gastrointestinal cell lines Caco-2, HT-29, HEPG2, and colon fibroblasts using a custom expression panel for tight-junction and cytoskeletal regulatory genes": Sup. Results 1: foundation.tooltip.js

```
;(function ($, window, document, undefined) {
  'use strict';

  Foundation.libs.tooltip = {
    name : 'tooltip',

    version : '5.3.0',

    settings : {
      additional_inheritable_classes : [],
      tooltip_class : '.tooltip',
      append_to: 'body',
      touch_close_text: 'Tap To Close',
      disable_for_touch: false,
      hover_delay: 200,
      show_on : 'all',
      tip_template : function (selector, content) {
        return '<span data-selector="' + selector + '" class="'
          + Foundation.libs.tooltip.settings.tooltip_class.substring(1)
          + '">' + content + '<span class="nub"></span></span>';
      }
    },

    cache : {},

    init : function (scope, method, options) {
      Foundation.inherit(this, 'random_str');
      this.bindings(method, options);
    },

    should_show: function (target, tip) {
      var settings = $.extend({}, this.settings, this.data_options(target));

      if (settings.show_on === 'all') {
        return true;
      } else if (this.small() && settings.show_on === 'small') {
        return true;
      } else if (this.medium() && settings.show_on === 'medium') {
        return true;
      } else if (this.large() && settings.show_on === 'large') {
        return true;
      }
      return false;
    },

    medium : function () {
      return matchMedia(Foundation.media_queries['medium']).matches;
    },

    large : function () {
      return matchMedia(Foundation.media_queries['large']).matches;
    },

    events : function (instance) {
      var self = this,
          S = self.S;

      self.create(this.S(instance));

      $(this.scope)
        .off('.tooltip')
        .on('mouseenter.fndtn.tooltip mouseleave.fndtn.tooltip touchstart.fndtn.tooltip MSPointerDown.fndtn.tooltip',
          '[' + this.attr_name() + ']', function (e) {
          var $this = S(this),
              settings = $.extend({}, self.settings, self.data_options($this)),
              is_touch = false;

          if (Modernizr.touch && /touchstart|MSPointerDown/i.test(e.type) && S(e.target).is('a')) {
            return false;
          }

          if (/mouse/i.test(e.type) && self.ie_touch(e)) return false;

          if ($this.hasClass('open')) {
            if (Modernizr.touch && /touchstart|MSPointerDown/i.test(e.type)) e.preventDefault();
            self.hide($this);
          } else {
            if (settings.disable_for_touch && Modernizr.touch && /touchstart|MSPointerDown/i.test(e.type)) {
              return;
            } else if(!settings.disable_for_touch && Modernizr.touch && /touchstart|MSPointerDown/i.test(e.type)) {
              e.preventDefault();
              S(settings.tooltip_class + '.open').hide();
              is_touch = true;
            }

            if (/enter|over/i.test(e.type)) {
              this.timer = setTimeout(function () {
                var tip = self.showTip($this);
              }.bind(this), self.settings.hover_delay);
            } else if (e.type === 'mouseout' || e.type === 'mouseleave') {
              clearTimeout(this.timer);
              self.hide($this);
            } else {
              self.showTip($this);
            }
          }
        })
        .on('mouseleave.fndtn.tooltip touchstart.fndtn.tooltip MSPointerDown.fndtn.tooltip', '[' + this.attr_name() + '].open', function (e) {
          if (/mouse/i.test(e.type) && self.ie_touch(e)) return false;

          if($(this).data('tooltip-open-event-type') == 'touch' && e.type == 'mouseleave') {
            return;
          }
          else if($(this).data('tooltip-open-event-type') == 'mouse' && /MSPointerDown|touchstart/i.test(e.type)) {
            self.convert_to_touch($(this));
          } else {
            self.hide($(this));
          }
        })
        .on('DOMNodeRemoved DOMAttrModified', '[' + this.attr_name() + ']:not(a)', function (e) {
          self.hide(S(this));
        });
    },

    ie_touch : function (e) {
      // How do I distinguish between IE11 and Windows Phone 8?????
      return false;
    },

    showTip : function ($target) {
      var $tip = this.getTip($target);
      if (this.should_show($target, $tip)){
        return this.show($target);
      }
      return;
    },

    getTip : function ($target) {
      var selector = this.selector($target),
          settings = $.extend({}, this.settings, this.data_options($target)),
          tip = null;

      if (selector) {
        tip = this.S('span[data-selector="' + selector + '"]' + settings.tooltip_class);
      }

      return (typeof tip === 'object') ? tip : false;
    },

    selector : function ($target) {
      var id = $target.attr('id'),
          dataSelector = $target.attr(this.attr_name()) || $target.attr('data-selector');

      if ((id && id.length < 1 || !id) && typeof dataSelector != 'string') {
        dataSelector = this.random_str(6);
        $target.attr('data-selector', dataSelector);
      }

      return (id && id.length > 0) ? id : dataSelector;
    },

    create : function ($target) {
      var self = this,
          settings = $.extend({}, this.settings, this.data_options($target)),
          tip_template = this.settings.tip_template;

      if (typeof settings.tip_template === 'string' && window.hasOwnProperty(settings.tip_template)) {
        tip_template = window[settings.tip_template];
      }

      var $tip = $(tip_template(this.selector($target), $('<div></div>').html($target.attr('title')).html())),
          classes = this.inheritable_classes($target);

      $tip.addClass(classes).appendTo(settings.append_to);

      if (Modernizr.touch) {
        $tip.append('<span class="tap-to-close">'+settings.touch_close_text+'</span>');
        $tip.on('touchstart.fndtn.tooltip MSPointerDown.fndtn.tooltip', function(e) {
          self.hide($target);
        });
      }

      $target.removeAttr('title').attr('title','');
    },

    reposition : function (target, tip, classes) {
      var width, nub, nubHeight, nubWidth, column, objPos;

      tip.css('visibility', 'hidden').show();

      width = target.data('width');
      nub = tip.children('.nub');
      nubHeight = nub.outerHeight();
      nubWidth = nub.outerHeight();

      if (this.small()) {
        tip.css({'width' : '100%' });
      } else {
        tip.css({'width' : (width) ? width : 'auto'});
      }

      objPos = function (obj, top, right, bottom, left, width) {
        return obj.css({
          'top' : (top) ? top : 'auto',
          'bottom' : (bottom) ? bottom : 'auto',
          'left' : (left) ? left : 'auto',
          'right' : (right) ? right : 'auto'
        }).end();
      };

      objPos(tip, (target.offset().top + target.outerHeight() + 10), 'auto', 'auto', target.offset().left);

      if (this.small()) {
        objPos(tip, (target.offset().top + target.outerHeight() + 10), 'auto', 'auto', 12.5, $(this.scope).width());
        tip.addClass('tip-override');
        objPos(nub, -nubHeight, 'auto', 'auto', target.offset().left);
      } else {
        var left = target.offset().left;
        if (Foundation.rtl) {
          nub.addClass('rtl');
          left = target.offset().left + target.outerWidth() - tip.outerWidth();
        }
        objPos(tip, (target.offset().top + target.outerHeight() + 10), 'auto', 'auto', left);
        tip.removeClass('tip-override');
        if (classes && classes.indexOf('tip-top') > -1) {
          if (Foundation.rtl) nub.addClass('rtl');
          objPos(tip, (target.offset().top - tip.outerHeight()), 'auto', 'auto', left)
            .removeClass('tip-override');
        } else if (classes && classes.indexOf('tip-left') > -1) {
          objPos(tip, (target.offset().top + (target.outerHeight() / 2) - (tip.outerHeight() / 2)), 'auto', 'auto', (target.offset().left - tip.outerWidth() - nubHeight))
            .removeClass('tip-override');
          nub.removeClass('rtl');
        } else if (classes && classes.indexOf('tip-right') > -1) {
          objPos(tip, (target.offset().top + (target.outerHeight() / 2) - (tip.outerHeight() / 2)), 'auto', 'auto', (target.offset().left + target.outerWidth() + nubHeight))
            .removeClass('tip-override');
          nub.removeClass('rtl');
        }
      }

      tip.css('visibility', 'visible').hide();
    },

    small : function () {
      return matchMedia(Foundation.media_queries.small).matches &&
        !matchMedia(Foundation.media_queries.medium).matches;
    },

    inheritable_classes : function ($target) {
      var settings = $.extend({}, this.settings, this.data_options($target)),
          inheritables = ['tip-top', 'tip-left', 'tip-bottom', 'tip-right', 'radius', 'round'].concat(settings.additional_inheritable_classes),
          classes = $target.attr('class'),
          filtered = classes ? $.map(classes.split(' '), function (el, i) {
            if ($.inArray(el, inheritables) !== -1) {
              return el;
            }
          }).join(' ') : '';

      return $.trim(filtered);
    },

    convert_to_touch : function($target) {
      var self = this,
          $tip = self.getTip($target),
          settings = $.extend({}, self.settings, self.data_options($target));

      if ($tip.find('.tap-to-close').length === 0) {
        $tip.append('<span class="tap-to-close">'+settings.touch_close_text+'</span>');
        $tip.on('click.fndtn.tooltip.tapclose touchstart.fndtn.tooltip.tapclose MSPointerDown.fndtn.tooltip.tapclose', function(e) {
          self.hide($target);
        });
      }

      $target.data('tooltip-open-event-type', 'touch');
    },

    show : function ($target) {
      var $tip = this.getTip($target);

      if ($target.data('tooltip-open-event-type') == 'touch') {
        this.convert_to_touch($target);
      }

      this.reposition($target, $tip, $target.attr('class'));
      $target.addClass('open');
      $tip.fadeIn(150);
    },

    hide : function ($target) {
      var $tip = this.getTip($target);

      $tip.fadeOut(150, function() {
        $tip.find('.tap-to-close').remove();
        $tip.off('click.fndtn.tooltip.tapclose touchstart.fndtn.tooltip.tapclose MSPointerDown.fndtn.tapclose');
        $target.removeClass('open');
      });
    },

    off : function () {
      var self = this;
      this.S(this.scope).off('.fndtn.tooltip');
      this.S(this.settings.tooltip_class).each(function (i) {
        $('[' + self.attr_name() + ']').eq(i).attr('title', $(this).text());
      }).remove();
    },

    reflow : function () {}
  };
}(jQuery, window, window.document));
```
