## Supplementary material for "Comparative pathway enrichment analysis in gastrointestinal cell lines Caco-2, HT-29, HEPG2, and colon fibroblasts using a custom expression panel for tight-junction and cytoskeletal regulatory genes": Sup. Results 1: foundation.topbar.js

```
;(function ($, window, document, undefined) {
  'use strict';

  Foundation.libs.topbar = {
    name : 'topbar',

    version: '5.3.0',

    settings : {
      index : 0,
      sticky_class : 'sticky',
      custom_back_text: true,
      back_text: 'Back',
      is_hover: true,
      scrolltop : true, // jump to top when sticky nav menu toggle is clicked
      sticky_on : 'all'
    },
    
    init : function (section, method, options) {
      Foundation.inherit(this, 'add_custom_rule register_media throttle');
      var self = this;

      self.register_media('topbar', 'foundation-mq-topbar');

      this.bindings(method, options);

      self.S('[' + this.attr_name() + ']', this.scope).each(function () {
        var topbar = $(this),
            settings = topbar.data(self.attr_name(true) + '-init'),
            section = self.S('section', this);
        topbar.data('index', 0);
        var topbarContainer = topbar.parent();
        if (topbarContainer.hasClass('fixed') || self.is_sticky(topbar, topbarContainer, settings) ) {
          self.settings.sticky_class = settings.sticky_class;
          self.settings.sticky_topbar = topbar;
          topbar.data('height', topbarContainer.outerHeight());
          topbar.data('stickyoffset', topbarContainer.offset().top);
        } else {
          topbar.data('height', topbar.outerHeight());
        }

        if (!settings.assembled) {
          self.assemble(topbar);
        }

        if (settings.is_hover) {
          self.S('.has-dropdown', topbar).addClass('not-click');
        } else {
          self.S('.has-dropdown', topbar).removeClass('not-click');
        }

        // Pad body when sticky (scrolled) or fixed.
        self.add_custom_rule('.f-topbar-fixed { padding-top: ' + topbar.data('height') + 'px }');

        if (topbarContainer.hasClass('fixed')) {
          self.S('body').addClass('f-topbar-fixed');
        }
      });

    },

    is_sticky: function (topbar, topbarContainer, settings) {
      var sticky = topbarContainer.hasClass(settings.sticky_class);

      if (sticky && settings.sticky_on === 'all') {
        return true;
      } else if (sticky && this.small() && settings.sticky_on === 'small') {
        return (matchMedia(Foundation.media_queries.small).matches && !matchMedia(Foundation.media_queries.medium).matches &&
            !matchMedia(Foundation.media_queries.large).matches); 
        //return true;
      } else if (sticky && this.medium() && settings.sticky_on === 'medium') {
        return (matchMedia(Foundation.media_queries.small).matches && matchMedia(Foundation.media_queries.medium).matches &&
            !matchMedia(Foundation.media_queries.large).matches);
        //return true;
      } else if(sticky && this.large() && settings.sticky_on === 'large') {
        return (matchMedia(Foundation.media_queries.small).matches && matchMedia(Foundation.media_queries.medium).matches &&
            matchMedia(Foundation.media_queries.large).matches);
        //return true;
      }

      return false;
    },

    toggle: function (toggleEl) {
      var self = this,
          topbar;

      if (toggleEl) {
        topbar = self.S(toggleEl).closest('[' + this.attr_name() + ']');
      } else {
        topbar = self.S('[' + this.attr_name() + ']');
      }

      var settings = topbar.data(this.attr_name(true) + '-init');

      var section = self.S('section, .section', topbar);

      if (self.breakpoint()) {
        if (!self.rtl) {
          section.css({left: '0%'});
          $('>.name', section).css({left: '100%'});
        } else {
          section.css({right: '0%'});
          $('>.name', section).css({right: '100%'});
        }

        self.S('li.moved', section).removeClass('moved');
        topbar.data('index', 0);

        topbar
          .toggleClass('expanded')
          .css('height', '');
      }

      if (settings.scrolltop) {
        if (!topbar.hasClass('expanded')) {
          if (topbar.hasClass('fixed')) {
            topbar.parent().addClass('fixed');
            topbar.removeClass('fixed');
            self.S('body').addClass('f-topbar-fixed');
          }
        } else if (topbar.parent().hasClass('fixed')) {
          if (settings.scrolltop) {
            topbar.parent().removeClass('fixed');
            topbar.addClass('fixed');
            self.S('body').removeClass('f-topbar-fixed');

            window.scrollTo(0,0);
          } else {
            topbar.parent().removeClass('expanded');
          }
        }
      } else {
        if (self.is_sticky(topbar, topbar.parent(), settings)) {
          topbar.parent().addClass('fixed');
        }

        if (topbar.parent().hasClass('fixed')) {
          if (!topbar.hasClass('expanded')) {
            topbar.removeClass('fixed');
            topbar.parent().removeClass('expanded');
            self.update_sticky_positioning();
          } else {
            topbar.addClass('fixed');
            topbar.parent().addClass('expanded');
            self.S('body').addClass('f-topbar-fixed');
          }
        }
      }
    },

    timer : null,

    events : function (bar) {
      var self = this,
          S = this.S;

      S(this.scope)
        .off('.topbar')
        .on('click.fndtn.topbar', '[' + this.attr_name() + '] .toggle-topbar', function (e) {
          e.preventDefault();
          self.toggle(this);
        })
        .on('click.fndtn.topbar','.top-bar .top-bar-section li a[href^="#"],[' + this.attr_name() + '] .top-bar-section li a[href^="#"]',function (e) {
            var li = $(this).closest('li');
            if(self.breakpoint() && !li.hasClass('back') && !li.hasClass('has-dropdown'))
            {
            self.toggle();
            }
        })
        .on('click.fndtn.topbar', '[' + this.attr_name() + '] li.has-dropdown', function (e) {
          var li = S(this),
              target = S(e.target),
              topbar = li.closest('[' + self.attr_name() + ']'),
              settings = topbar.data(self.attr_name(true) + '-init');

          if(target.data('revealId')) {
            self.toggle();
            return;
          }

          if (self.breakpoint()) return;
          if (settings.is_hover && !Modernizr.touch) return;

          e.stopImmediatePropagation();

          if (li.hasClass('hover')) {
            li
              .removeClass('hover')
              .find('li')
              .removeClass('hover');

            li.parents('li.hover')
              .removeClass('hover');
          } else {
            li.addClass('hover');

            $(li).siblings().removeClass('hover');

            if (target[0].nodeName === 'A' && target.parent().hasClass('has-dropdown')) {
              e.preventDefault();
            }
          }
        })
        .on('click.fndtn.topbar', '[' + this.attr_name() + '] .has-dropdown>a', function (e) {
          if (self.breakpoint()) {

            e.preventDefault();

            var $this = S(this),
                topbar = $this.closest('[' + self.attr_name() + ']'),
                section = topbar.find('section, .section'),
                dropdownHeight = $this.next('.dropdown').outerHeight(),
                $selectedLi = $this.closest('li');

            topbar.data('index', topbar.data('index') + 1);
            $selectedLi.addClass('moved');

            if (!self.rtl) {
              section.css({left: -(100 * topbar.data('index')) + '%'});
              section.find('>.name').css({left: 100 * topbar.data('index') + '%'});
            } else {
              section.css({right: -(100 * topbar.data('index')) + '%'});
              section.find('>.name').css({right: 100 * topbar.data('index') + '%'});
            }

            topbar.css('height', $this.siblings('ul').outerHeight(true) + topbar.data('height'));
          }
        });
      
      S(window).off('.topbar').on('resize.fndtn.topbar', self.throttle(function () {
        self.resize.call(self);
      }, 50)).trigger('resize').trigger('resize.fndtn.topbar');

      S('body').off('.topbar').on('click.fndtn.topbar touchstart.fndtn.topbar', function (e) {
        var parent = S(e.target).closest('li').closest('li.hover');

        if (parent.length > 0) {
          return;
        }

        S('[' + self.attr_name() + '] li.hover').removeClass('hover');
      });

      // Go up a level on Click
      S(this.scope).on('click.fndtn.topbar', '[' + this.attr_name() + '] .has-dropdown .back', function (e) {
        e.preventDefault();

        var $this = S(this),
            topbar = $this.closest('[' + self.attr_name() + ']'),
            section = topbar.find('section, .section'),
            settings = topbar.data(self.attr_name(true) + '-init'),
            $movedLi = $this.closest('li.moved'),
            $previousLevelUl = $movedLi.parent();

        topbar.data('index', topbar.data('index') - 1);

        if (!self.rtl) {
          section.css({left: -(100 * topbar.data('index')) + '%'});
          section.find('>.name').css({left: 100 * topbar.data('index') + '%'});
        } else {
          section.css({right: -(100 * topbar.data('index')) + '%'});
          section.find('>.name').css({right: 100 * topbar.data('index') + '%'});
        }

        if (topbar.data('index') === 0) {
          topbar.css('height', '');
        } else {
          topbar.css('height', $previousLevelUl.outerHeight(true) + topbar.data('height'));
        }

        setTimeout(function () {
          $movedLi.removeClass('moved');
        }, 300);
      });
    },

    resize : function () {
      var self = this;
      self.S('[' + this.attr_name() + ']').each(function () {
        var topbar = self.S(this),
            settings = topbar.data(self.attr_name(true) + '-init');

        var stickyContainer = topbar.parent('.' + self.settings.sticky_class);
        var stickyOffset;

        if (!self.breakpoint()) {
          var doToggle = topbar.hasClass('expanded');
          topbar
            .css('height', '')
            .removeClass('expanded')
            .find('li')
            .removeClass('hover');

            if(doToggle) {
              self.toggle(topbar);
            }
        }

        if(self.is_sticky(topbar, stickyContainer, settings)) {
          if(stickyContainer.hasClass('fixed')) {
            // Remove the fixed to allow for correct calculation of the offset.
            stickyContainer.removeClass('fixed');

            stickyOffset = stickyContainer.offset().top;
            if(self.S(document.body).hasClass('f-topbar-fixed')) {
              stickyOffset -= topbar.data('height');
            }

            topbar.data('stickyoffset', stickyOffset);
            stickyContainer.addClass('fixed');
          } else {
            stickyOffset = stickyContainer.offset().top;
            topbar.data('stickyoffset', stickyOffset);
          }
        }

      });
    },

    breakpoint : function () {
      return !matchMedia(Foundation.media_queries['topbar']).matches;
    },

    small : function () {
      return matchMedia(Foundation.media_queries['small']).matches;
    },

    medium : function () {
      return matchMedia(Foundation.media_queries['medium']).matches;
    },

    large : function () {
      return matchMedia(Foundation.media_queries['large']).matches;
    },

    assemble : function (topbar) {
      var self = this,
          settings = topbar.data(this.attr_name(true) + '-init'),
          section = self.S('section', topbar);

      // Pull element out of the DOM for manipulation
      section.detach();

      self.S('.has-dropdown>a', section).each(function () {
        var $link = self.S(this),
            $dropdown = $link.siblings('.dropdown'),
            url = $link.attr('href'),
            $titleLi;

        if (!$dropdown.find('.title.back').length) {
          $titleLi = $('<li class="title back js-generated"><h5><a href="javascript:void(0)"></a></h5></li>');
  
          // Copy link to subnav
          if (settings.custom_back_text == true) {
            $('h5>a', $titleLi).html(settings.back_text);
          } else {
            $('h5>a', $titleLi).html('&laquo; ' + $link.html());
          }
          $dropdown.prepend($titleLi);
        }
      });

      // Put element back in the DOM
      section.appendTo(topbar);

      // check for sticky
      this.sticky();

      this.assembled(topbar);
    },

    assembled : function (topbar) {
      topbar.data(this.attr_name(true), $.extend({}, topbar.data(this.attr_name(true)), {assembled: true}));
    },

    height : function (ul) {
      var total = 0,
          self = this;

      $('> li', ul).each(function () { 
        total += self.S(this).outerHeight(true); 
      });

      return total;
    },

    sticky : function () {
      var self = this;

      this.S(window).on('scroll', function() {
        self.update_sticky_positioning();
      });
    },

    update_sticky_positioning: function() {
      var klass = '.' + this.settings.sticky_class,
          $window = this.S(window), 
          self = this;

      if (self.settings.sticky_topbar && self.is_sticky(this.settings.sticky_topbar,this.settings.sticky_topbar.parent(), this.settings)) {
        var distance = this.settings.sticky_topbar.data('stickyoffset');
        if (!self.S(klass).hasClass('expanded')) {
          if ($window.scrollTop() > (distance)) {
            if (!self.S(klass).hasClass('fixed')) {
              self.S(klass).addClass('fixed');
              self.S('body').addClass('f-topbar-fixed');
            }
          } else if ($window.scrollTop() <= distance) {
            if (self.S(klass).hasClass('fixed')) {
              self.S(klass).removeClass('fixed');
              self.S('body').removeClass('f-topbar-fixed');
            }
          }
        }
      }
    },

    off : function () {
      this.S(this.scope).off('.fndtn.topbar');
      this.S(window).off('.fndtn.topbar');
    },

    reflow : function () {}
  };
}(jQuery, this, this.document));
```
