## Supplementary material for "Comparative pathway enrichment analysis in gastrointestinal cell lines Caco-2, HT-29, HEPG2, and colon fibroblasts using a custom expression panel for tight-junction and cytoskeletal regulatory genes": Sup. Results 1: heatmap.js

```
//--------------------------------------------------------------------------------------------------
//                                      BEGIN HEATMAP FUNCTION
//--------------------------------------------------------------------------------------------------

function heatmap(id, datasetFile, colAnnoFile, rowAnnoFile, colClustOrder, rowClustOrder,
                 height, renderOnBrushEnd, categorical,
                 colCatScheme, colConScheme, colAnnoHeatScheme,
                 rowCatScheme, rowConScheme, rowAnnoHeatScheme,
                 bucketDividers, bucketColors,
                 animDuration, sideColorPad, annoTitlePad, axisPad,
                 fontSize, fontSizeCK, lowColor, midColor, highColor, numColors) {

  // assign parameters to defaults if not given
  height            = height || 600;
  colCatScheme      = colCatScheme || "google";
  colConScheme      = colConScheme || "rainbow";
  colAnnoHeatScheme = colAnnoHeatScheme || "plasma";
  rowCatScheme      = rowCatScheme || "ns";
  rowConScheme      = rowConScheme || "cubehelix";
  rowAnnoHeatScheme = rowAnnoHeatScheme || "magma";
  bucketDividers    = bucketDividers || [25, 50, 100, 500];
  bucketColors      = bucketColors || ["cornflowerblue", "navy", "gray", "#7F5217", "goldenrod"];
  animDuration      = animDuration || 1200;
  sideColorPad      = sideColorPad || 3;
  annoTitlePad      = annoTitlePad || 7;
  axisPad           = axisPad || 5;
  fontSize          = fontSize || 9;
  fontSizeCK        = fontSizeCK || 11;
  lowColor          = lowColor || "cornflowerblue";
  midColor          = midColor || "black";
  highColor         = highColor || "orange";
  numColors         = numColors || 256;

  // the array of colors used for the heatmap
  var heatmapColors = interpolateColors(lowColor, midColor, highColor, numColors);

  // the DOM element passed in
  var parent = document.getElementById(id),
  // the width of the SVG will be the same as the parent
      width = parent.clientWidth,
  // holds all DOM elements of the heatmap (SVG and divs for the tooltips)
      container = d3.select("#" + id).append("div").attr("class", "heatmap");

  // margin convention for D3
  var margin = {top: 3, right: 3, bottom: 3, left: 3};
  // width and height will refer to the 'inner' width/height of the widget, with margins applied
  width = width - margin.left - margin.right;
  height = height - margin.top - margin.bottom;

  // the actual SVG, whose width is the same as the parent
  var SVG = container.append("svg");
  svgSetup(width, height);
  // the pseudo-SVG, with margins applied, to which all subsequent SVG elements are appended
  var svg = SVG.append("g").attr("transform", "translate(" + margin.left + "," + margin.top + ")");
  // when the browser window resizes, the SVG also resizes to fit the parent's width
  window.addEventListener("resize", resizeSVG);

  // the "dims" will hold all elements relevant to the columns and rows of the data, separately
  var col = {}, row = {};
  // parse the files (.csv strings) and assign the data structures to col and row fields
  var dataset    = parseDataMatrix(datasetFile);
  col.stats      = dataset.stats.col;
  row.stats      = dataset.stats.row;
  col.clustOrder = colClustOrder || dataset.colnames;
  row.clustOrder = rowClustOrder || dataset.rownames;
  col.names      = colClustOrder || dataset.colnames;
  row.names      = rowClustOrder || dataset.rownames;
  col.annotated  = colAnnoFile ? true : false;
  row.annotated  = rowAnnoFile ? true : false;
  if (col.annotated) annoSetup(col, colAnnoFile);
  if (row.annotated) annoSetup(row, rowAnnoFile);
  if (col.annotated) colorsSetup(col, categorical, colCatScheme, colConScheme, colAnnoHeatScheme);
  if (row.annotated) colorsSetup(row, categorical, rowCatScheme, rowConScheme, rowAnnoHeatScheme);

  function annoSetup(dim, annoFile) {
    var annosParsed = parseAnnotations(annoFile);
    dim.annotations = annosParsed.annotations;
    dim.labelsAnnotated = annosParsed.labels;
  }

  //================================================================================================
  //                                        REFERENCES BY DIM
  //
  // Note that the row and col objects are mirror images of each other; every field that col has,
  // row also has, and vice-versa. They could be made into objects of the same new class, but its
  // much easier to just build them as we go along and think of them more as growing lists
  // containing everything that's relevant to their respective dimension.
  //
  // For example, when we zoom and pan using the brush for the columns, the only things that need
  // to get visually updated are:
  //    * column labels
  //    * x-coordinates of the heatmap cells
  //    * widths of the heatmap cells
  //    * x-coordinates of the column side colors
  //    * heights of the column side colors
  // Similarly, when doing the same thing to the rows, we need only be concerned with updating:
  //    * row labels
  //    * y-coordinates of the heatmap cells
  //    * heights of the heatmap cells
  //    * y-coordinates of the row side colors
  //    * heights of the row side colors
  // Grouping these another way, we see that there are different "types" of things that get updated:
  //    * labels (column, row)
  //    * coordinates (x, y)
  //    * lengths (width, height)
  //    * side colors (column, row)
  //    * heatmap cells
  // For each of these types, col and row should store a reference (with the same name) to the value
  // of that type that is relevant to them (note that we update the heatmap cells regardless, so we
  // can just store this as a global variable):
  //    * col.labels = column labels
  //      row.labels = row labels
  //    * col.coordinate = x
  //      row.coordiante = y
  //    * col.length = width
  //      row.length = height
  //    * col.sideColors = column side colors
  //      row.sideColors = row side colors
  // We can thus create a function which handles the event where either dim has been zoomed/panned,
  // needing only parameter, the dim, whose "labels", "coordinate", "length", and "sideColors"
  // fields will be used (along with the global reference to the heatmap cells) to determine the
  // visual updates. NOTE: the function that actually does this modifies more variables than just
  // those listed here, and additionally the actual field names may be different than they are here.
  //
  // And that's the concept behind the "dim".
  //
  //================================================================================================

  // set the current scope for each dimension (these get modified by interactivity functions)
  col.currentScope = [0, col.names.length];
  row.currentScope = [0, row.names.length];
  col.other        = row;
  row.other        = col;
  col.self         = "col";
  row.self         = "row";
  col.title        = "Sample";
  row.title        = "Probe";
  col.pos          = "x";
  row.pos          = "y";
  col.size         = "width";
  row.size         = "height";
  col.sizeHeatmap  = function() { return sizeHeatmap(row) - marginAnnoColor - marginAnnoLabel; };
  row.sizeHeatmap  = function() { return sizeHeatmap(col); };
  if (col.annotated) annotypesSetup(col);
  if (row.annotated) annotypesSetup(row);

  function annotypesSetup(dim) {
    dim.annotypes = Object.keys(dim.annotations).sort(function(a, b) {
      return a.localeCompare(b);
    });
    dim.annoBy = dim.annotypes[0];
  }

  //================================================================================================
  //                                              MARGINS
  // A margin describes a visual element's length in pixels along one axis/dimension.
  //================================================================================================

  var marginAnnoColor, marginAnnoLabel, marginAnnoTitle, marginColorKey;
  marginsSetup(width, height); // initialize

  function marginsSetup(w, h) {
    marginAnnoColor = Math.floor(w / 50);
    marginAnnoLabel = Math.min(Math.floor(w / 4), Math.floor(annoMax() + axisPad));
    marginAnnoTitle = fontSizeCK + 2 * annoTitlePad;
    col.marginTotal = h;
    row.marginTotal = w;
    col.marginLabel = col.labels ? Math.ceil(col.labels.getBox().height + 2 * axisPad) : 0;
    row.marginLabel = row.labels ? Math.ceil(row.labels.getBox().width + 2 * axisPad) : 0;
    col.marginLabelSub = col.labelsSub ? Math.ceil(col.labelsSub.getBox().height + 2 * axisPad) : 0;
    row.marginLabelSub = row.labelsSub ? Math.ceil(row.labelsSub.getBox().width + 2 * axisPad) : 0;
    col.marginBrush = Math.floor(h / 10);
    row.marginBrush = Math.floor(h / 10);
    marginColorKey = Math.floor(h / 4) - marginAnnoTitle;
    sideAndAnnoMarginsSetup(col);
    sideAndAnnoMarginsSetup(row);

    function sideAndAnnoMarginsSetup(dim) {
      dim.marginSideColor = dim.annotated ? Math.floor(h / 40) : 0;
      dim.marginAnnoTotal = dim.annotated ? Math.floor(3 * h / 8) : 0;
      dim.marginAnnoHeight = dim.annotated ? dim.marginAnnoTotal - marginAnnoTitle : 0;
    }

    function annoMax() {
      var cM1 = col.labelsAnno ? col.labelsAnno.getBox().width + 2 * axisPad : 0,
          rM1 = row.labelsAnno ? row.labelsAnno.getBox().width + 2 * axisPad : 0,
          cM2 = col.annoTitle ? col.annoTitle.getBox().width - marginAnnoColor + axisPad : 0,
          rM2 = row.annoTitle ? row.annoTitle.getBox().width - marginAnnoColor + axisPad : 0,
          ck1 = colorKey ? colorKey.labels[scalingDim].getBox().width + 2 * axisPad : 0,
          ck2 = colorKey ? colorKey.titles[scalingDim].getBox().width
                                                                    - marginAnnoColor + axisPad : 0;
      return Math.max(cM1, rM1, cM2, rM2, ck1, ck2);
    }
  }

  //================================================================================================
  //                                      TOOLTIPS/SETTINGS PANEL
  // Tooltips provide information for rows, columns, matrix data, and annotations when hovering over
  // the side colors, heatmap cells, and color key.
  //================================================================================================

  var scaleBy, scalingDim, settingsHidden = true,
      settingsPanel              = settingsPanelSetup(),
      cellTooltip                = cellTooltipSetup(),
      annoTooltip                = annoTooltipSetup();
  if (col.annotated) col.tooltip = sideTooltipSetup(col);
  if (row.annotated) row.tooltip = sideTooltipSetup(row);

  //================================================================================================
  //                                              SCALES
  // Scales are very useful for determining where visual elements should be placed relative to each
  // other. For example, to determine the sizes and positions of the cells that make up the heatmap,
  // a scale can map an array of row or column names to a continuous range of pixels.
  //================================================================================================

  // scales for determining cell color
  var mainColorScale = d3.scaleQuantize()
                        .domain([-dataset.stats.zMax[scalingDim], dataset.stats.zMax[scalingDim]])
                        .range(heatmapColors);
  var bucketizer = new Bucketizer(bucketDividers, bucketColors);
  if (col.annotated) colorScalesSetup(col);
  if (row.annotated) colorScalesSetup(row);

  function colorScalesSetup(dim) {
    dim.annoToNum = categorical ?
      d3.scaleOrdinal().domain(dim.annotations[dim.annoBy])
                       .range(d3.range(dim.annotations[dim.annoBy].length))
      : d3.scalePoint().domain(dim.annotations[dim.annoBy])
                       .range([0, 0.9]); // must be within [0, 1]
    dim.numToColor = dim.annoReg;
  }

  // for cell position/size - map row/col.names to x/y/width/height based on the margins
  col.scaleCell    = d3.scaleBand(); // col.names, col.sizeHeatmap -> x, width of cells
  row.scaleCell    = d3.scaleBand(); // row.names, row.sizeHeatmap -> y, height of cells
  col.scaleCellSub = d3.scaleBand(); // col.names, row.marginBrush -> x, width of cellsRight
  row.scaleCellSub = d3.scaleBand(); // row.names, col.marginBrush -> y, height of cellsBottom
  if (col.annotated) col.scaleAnnoColor = d3.scaleBand().domain(col.annotations[col.annoBy]);
  if (row.annotated) row.scaleAnnoColor = d3.scaleBand().domain(row.annotations[row.annoBy]);
  var scaleBucket = d3.scaleBand().domain(bucketColors),
      scaleGradient = d3.scaleBand().domain(heatmapColors);

  scalesSetup(width, height); // initialize

  function scalesSetup(width, height) {
    col.scaleCell.domain(col.names).range([0, col.sizeHeatmap()]);
    row.scaleCell.domain(row.names).range([0, row.sizeHeatmap()]);
    col.scaleCellSub.domain(col.names).range([0, row.marginBrush]);
    row.scaleCellSub.domain(row.names).range([0, col.marginBrush]);
    if (col.annotated) col.scaleAnnoColor.range([0, col.marginAnnoHeight]);
    if (row.annotated) row.scaleAnnoColor.range([0, row.marginAnnoHeight]);
    scaleBucket.range([0, marginColorKey]);
    scaleGradient.range([0, marginColorKey]);
  }

  //================================================================================================
  //                                            COLOR KEY
  // This holds all the elements that make up the color keys for the scaling options (row, col, none,
  // and bucket).
  //================================================================================================

  var colorKey = {
    cells: {},
    labels: {},
    titles: {},
    updateCells: function(attrs) {
      var names = Object.keys(this.cells);
      for (var j = 0; j < names.length; j++) this.cells[names[j]].update(attrs);
    },
    positionCells: function() {
      var names = Object.keys(this.cells);
      for (var j = 0; j < names.length; j++)
        positionElement(this.cells[names[j]].group, this.cellsAnchor);
    },
    updateNTLabels: function() {
      var names = Object.keys(this.labels);
      for (var j = 0; j < names.length; j++) this.labels[names[j]].updateNT();
    },
    positionLabels: function() {
      var names = Object.keys(this.labels);
      for (var j = 0; j < names.length; j++)
        positionElement(this.labels[names[j]].group, this.labelsAnchor);
    },
    positionTitles: function() {
      var names = Object.keys(this.titles);
      for (var j = 0; j < names.length; j++)
        positionElement(this.titles[names[j]].selection, this.titleAnchor);
    },
    addTitle: function(name, text) {
      this.titles[name] = new Title(name + "CKTitle", text);
    },
    addLabels: function(name, labels) {
      this.labels[name] = new Labels(name + "CKLabels", labels,
        function() { return marginColorKey; }, this.cells[name].height, "right", false);
    },
    change: function(type) {
      var names = Object.keys(this.titles);
      for (var j = 0; j < names.length; j++) {
        this.cells[names[j]].group.classed("hidden", names[j] !== type);
        this.labels[names[j]].group.classed("hidden", names[j] !== type);
        this.titles[names[j]].selection.classed("hidden", names[j] !== type);
      }
    }
  };

  //================================================================================================
  //                                          CELlS
  // These represent groupings of colorful rectangles (row side colors, color keys, heatmap itself).
  //
  // In combination with the axes, these make up all the SVG elements. With the exception of the
  // annotation titles and axes, every visual component can be decomposed into 2 parts:
  //    * group - a g element which gets positioned at an anchor point
  //    * cells - rect elements which live inside the group
  // When a group is tranlated to a new position, all the elements inside of it move as well, and
  // this makes it so that the coordinates (x and y) of cells are relative their group, not to the
  // SVG as a whole.
  //================================================================================================

  function Cells(type, dim, x, y, width, height, fill) {
    this.dim = dim;
    this.x = x;
    this.y = y;
    this.width = width;
    this.height = height;
    this.fill = fill;
    this.group = svg.append("g");
    this.update = function(attrs) {
      for (var j = 0; j < attrs.length; j++) this.selection.attr(attrs[j], this[attrs[j]]);
    };
    this.position = function() { positionElement(this.group, this.anchor); };
    switch(type) {
      case "heatmap":
        this.selection = this.group.selectAll("g") // first, we add the rows in (not visible)
          .data(dataset.matrix) // each "d" is an array of cells
          .enter()              // selects all the new data (i.e., all of it)
          .append("g")          // the rows have now been added
          .selectAll("rect")    // then, we add the cells in (visible)
          .data(identity, key)  // in the key function, "d" is now a cell
          .enter()              // from here on, "d" refers to an individual cell
          .append("rect");      // the cells have now been added, but still invisible
        if (!dim) this.selection.on("mouseover", function(d) { displayCellTooltip(d, this); })
                .on("mouseout", function() { cellTooltip.classed("hidden", true); })
                .on("click", function() { toggleSettingsPanel(this, width, height, cellTooltip); });
        break;
      case "sideColors":
        this.selection = this.group.selectAll("rect")
              .data(dim.labelsAnnotated, key)
              .enter()
              .append("rect")
              .on("mouseover", function(d) { displaySideTooltip(d, this, dim); })
              .on("mouseout", function() { dim.tooltip.classed("hidden", true); })
              .on("click", function() { toggleSettingsPanel(this, width, height, dim.tooltip); });
        break;
      case "annoColors":
        if (!dim.self) {
          this.selection = this.group.selectAll("rect")
                .data(dim, identity)
                .enter()
                .append("rect");
        } else {
          this.setup = function(data) {
            this.group.selectAll("rect").remove();
            this.selection = this.group.selectAll("rect")
                  .data(data, identity)
                  .enter()
                  .append("rect")
                  .on("mouseover", function(d) { displayAnnoTooltip(d, this, dim); })
                  .on("mouseout", function() { annoTooltip.classed("hidden", true); });
            this.update(["x", "y", "width", "height", "fill"]);
          };
          this.setup(dim.annotations[dim.annoBy]); // initialize
        }
        break;
    }
    this.update(["x", "y", "width", "height", "fill"]); // initialize
  }

  var cells = new Cells("heatmap", null,
    function(d) { return col.scaleCell(d.col); },
    function(d) { return row.scaleCell(d.row); },
    function() { return col.scaleCell.bandwidth(); },
    function() { return row.scaleCell.bandwidth(); },
    function(d) {
      if (scalingDim === "none") return mainColorScale(d.value);
      if (scalingDim === "bucket") return bucketizer.bucketize(d.value);
      var ref = dataset.stats[scalingDim][dotsToUnders(d[scalingDim])];
      return mainColorScale((d.value - ref.mean) / ref.stdev);
    });

  col.cellsSub = new Cells("heatmap", col,
    cells.x, // inherit x attribute from cells
    function(d) { return row.scaleCellSub(d.row); },
    cells.width, // inherit width attribute from cells
    function() { return row.scaleCellSub.bandwidth(); },
    cells.fill); // inherit fill attribute from cells
  row.cellsSub = new Cells("heatmap", row,
    function(d) { return col.scaleCellSub(d.col); },
    cells.y, // inherit y attribute from cells
    function() { return col.scaleCellSub.bandwidth(); },
    cells.height, // inherit height attribute from cells
    cells.fill); // inherit fill attribute from cells

  colorKey.cells.none = new Cells("annoColors", heatmapColors,
    function() { return 0; },
    function(d) { return scaleGradient(d); },
    function() { return marginAnnoColor; },
    function() { return scaleGradient.bandwidth(); },
    identity);
  colorKey.cells.col = new Cells("annoColors", heatmapColors, // same as none
    function() { return 0; },
    function(d) { return scaleGradient(d); },
    function() { return marginAnnoColor; },
    function() { return scaleGradient.bandwidth(); },
    identity);
  colorKey.cells.row = new Cells("annoColors", heatmapColors, // same as none
    function() { return 0; },
    function(d) { return scaleGradient(d); },
    function() { return marginAnnoColor; },
    function() { return scaleGradient.bandwidth(); },
    identity);
  colorKey.cells.bucket = new Cells("annoColors", bucketColors,
    function() { return 0; },
    function(d) { return scaleBucket(d); },
    function() { return marginAnnoColor; },
    function() { return scaleBucket.bandwidth(); },
    identity);

  if (col.annotated) sideAndAnnoColorsSetup(col);
  if (row.annotated) sideAndAnnoColorsSetup(row);

  function sideAndAnnoColorsSetup(dim) {
    dim.sideColors = new Cells("sideColors", dim,
      dim.self === "col" ? function(d) { return col.scaleCell(d.key); } : function() { return 0; },
      dim.self === "row" ? function(d) { return row.scaleCell(d.key); } : function() { return 0; },
      dim.self === "col" ? cells.width : function() { return row.marginSideColor - sideColorPad; },
      dim.self === "row" ? cells.height : function() { return col.marginSideColor - sideColorPad; },
      function(d) { return dim.numToColor(dim.annoToNum(d.annos[dim.annoBy])); });
    dim.annoColors = new Cells("annoColors", dim,
      function() { return 0; },
      function(d) { return dim.scaleAnnoColor(d); },
      function() { return marginAnnoColor; },
      function() { return dim.scaleAnnoColor.bandwidth(); },
      function(d) { return dim.numToColor(dim.annoToNum(d)); });
  }

  //================================================================================================
  //                                              LABELS
  // These make up all the tickmark-prefaced pieces of text next to cells.
  //
  // The axes provide a visualization for the labels of rows, columns, and annotations. There are 3
  // parts that go into making a visible axis:
  //    * scale - a d3.scalePoint object whose domain is the labels and range is the pixel
  //              coordinate extent in which to display them
  //    * axis component - a d3.axis object determining the axis orientation (top/bottom/left/right)
  //    * SVG element - a g element which makes the axis visible
  // When an axis is to be visually updated, first update its scale, then call its axis component on
  // its SVG element.
  //================================================================================================

  function Labels(id, names, room, offset, orientation, angled) {
    this.id = id;
    this.names = names;
    this.room = room;
    this.offset = offset;
    this.angled = angled;
    this.factor = this.angled ? 0.75 : 1;
    this.scale = d3.scalePoint();
    this.updateScale = function(newNames) {
      this.names = newNames;
      this.scale.domain(sample(this.names, Math.floor(this.factor * this.room() / fontSize)))
                .range([this.offset() / 2, this.room() - this.offset() / 2]);
    };
    this.updateScale(this.names);
    switch(orientation) {
      case "left": this.axis = d3.axisLeft(this.scale); break;
      case "top": this.axis = d3.axisTop(this.scale); break;
      case "right": this.axis = d3.axisRight(this.scale); break;
      case "bottom": this.axis = d3.axisBottom(this.scale); break;
    }
    this.group = svg.append("g").attr("class", "axis").attr("id", id).style("font-size", fontSize);
    this.update = function() {
      this.updateScale(this.names);
      if (this.angled) {
        this.group.transition().duration(animDuration).call(this.axis)
               .selectAll("text")                // to angle the other way:
               .style("text-anchor", "start")    // end
               .attr("dx", ".8em")               // -.8em
               .attr("dy", ".15em")              // .15em
               .attr("transform", "rotate(45)"); // rotate(-45)
      } else {
        this.group.transition().duration(animDuration).call(this.axis);
      }
    };
    this.updateNT = function() {
      this.updateScale(this.names);
      if (this.angled) {
        this.group.call(this.axis)
               .selectAll("text")                // to angle the other way:
               .style("text-anchor", "start")    // end
               .attr("dx", ".8em")               // -.8em
               .attr("dy", ".15em")              // .15em
               .attr("transform", "rotate(45)"); // rotate(-45)
      } else {
        this.group.call(this.axis);
      }
    };
    this.updateNT(); // for initial angling
    this.position = function() { positionElement(this.group, this.anchor); };
    this.getBox = function() { return document.getElementById(this.id).getBoundingClientRect(); };
  }

  row.labels = new Labels("rLabs", row.names, row.sizeHeatmap, cells.height, "right", false);
  col.labels = new Labels("cLabs", col.names, col.sizeHeatmap, cells.width, "bottom", true);
  row.labelsSub = new Labels("rSubs", row.names, row.sizeHeatmap, cells.height, "right", false);
  col.labelsSub = new Labels("cSubs", col.names, col.sizeHeatmap, cells.width, "bottom", true);
  if (row.annotated) row.labelsAnno = new Labels("rAnnos", row.annotations[row.annoBy],
          function() { return row.marginAnnoHeight; }, row.annoColors.height, "right", false);
  if (col.annotated) col.labelsAnno = new Labels("cAnnos", col.annotations[col.annoBy],
          function() { return col.marginAnnoHeight; }, col.annoColors.height, "right", false);
  colorKey.addLabels("bucket", bucketDividers.concat([bucketDividers[bucketDividers.length - 1]])
            .map(function(d, i) { return i < bucketDividers.length - 1 ? "< " + d : ">= " + d; }));
  colorKey.addLabels("none", [dataset.stats.totalMin,
                  (dataset.stats.totalMin + dataset.stats.totalMax ) / 2, dataset.stats.totalMax]);
  colorKey.addLabels("row", [-dataset.stats.zMax.row.toFixed(2),
                           0, dataset.stats.zMax.row.toFixed(2)]);
  colorKey.addLabels("col", [-dataset.stats.zMax.col.toFixed(2),
                           0, dataset.stats.zMax.col.toFixed(2)]);

  //================================================================================================
  //                                            TITLES
  // These represent the titles on the columns of cells at the right.
  //================================================================================================

  function Title(id, text) {
    this.id = id;
    this.text = text;
    this.selection = svg.append("text").attr("class", "annoTitle").attr("id", this.id)
                                  .style("font-size", fontSizeCK).text(this.text);
    this.setText = function(text) {
      this.text = text;
      this.selection.text(this.text);
    };
    this.position = function() {
      positionElement(this.selection, this.anchor);
    };
    this.getBox = function() { return document.getElementById(this.id).getBoundingClientRect(); };
  }

  if (col.annotated) col.annoTitle = new Title("cTitle", undersToSpaces(col.annoBy));
  if (row.annotated) row.annoTitle = new Title("rTitle", undersToSpaces(row.annoBy));
  colorKey.addTitle("bucket", "Buckets");
  colorKey.addTitle("none", "Linear Gradient");
  colorKey.addTitle("row", "Row Z-Score");
  colorKey.addTitle("col", "Column Z-Score");

  //================================================================================================
  //                                             ANCHORS
  // An anchor is a 2-element array describing the pixel coordinates (relative to the SVG, not the
  // webpage as a whole) of the upper-left corner of a visual element. Using the "transform"
  // attribute of SVG elements, we can position each group of visual elements (for example, all the
  // cells of the heatmap) by simply translating it to the right by its anchor at index 0, and down
  // by its anchor at index 1. Anchors are determined by the margins.
  //================================================================================================

  anchorsSetup(width, height);

  function anchorsSetup(w, h) { // w not yet used
    cells.anchor         = [row.marginSideColor, col.marginSideColor];
    col.labels.anchor    = [cells.anchor[0], cells.anchor[1] + row.sizeHeatmap() + axisPad];
    row.labels.anchor    = [cells.anchor[0] + col.sizeHeatmap() + axisPad, cells.anchor[1]];
    col.cellsSub.anchor  = [cells.anchor[0], col.labels.anchor[1] + col.marginLabel];
    row.cellsSub.anchor  = [row.labels.anchor[0] + row.marginLabel, cells.anchor[1]];
    col.labelsSub.anchor = [cells.anchor[0], col.cellsSub.anchor[1] + col.marginBrush + axisPad];
    row.labelsSub.anchor = [row.cellsSub.anchor[0] + row.marginBrush + axisPad, cells.anchor[1]];
    if (col.annotated) {
      col.sideColors.anchor = [cells.anchor[0], 0];
      col.annoColors.anchor = [row.labelsSub.anchor[0] + row.marginLabelSub, marginAnnoTitle];
      col.annoTitle.anchor = [col.annoColors.anchor[0], col.annoColors.anchor[1] - annoTitlePad];
      col.labelsAnno.anchor = [col.annoColors.anchor[0] + marginAnnoColor + axisPad,
                                                                          col.annoColors.anchor[1]];
    }
    if (row.annotated) {
      row.sideColors.anchor = [0, cells.anchor[1]];
      row.annoColors.anchor = [row.labelsSub.anchor[0] + row.marginLabelSub,
                               col.marginAnnoTotal + marginAnnoTitle];
      row.annoTitle.anchor = [row.annoColors.anchor[0], row.annoColors.anchor[1] - annoTitlePad];
      row.labelsAnno.anchor = [row.annoColors.anchor[0] + marginAnnoColor + axisPad,
                                                                          row.annoColors.anchor[1]];
    }
    colorKey.cellsAnchor = [row.labelsSub.anchor[0] + row.marginLabelSub,
                            col.marginAnnoTotal + row.marginAnnoTotal + marginAnnoTitle];
    colorKey.labelsAnchor = [colorKey.cellsAnchor[0] + marginAnnoColor + axisPad,
                             colorKey.cellsAnchor[1]];
    colorKey.titleAnchor = [colorKey.cellsAnchor[0],
                            colorKey.cellsAnchor[1] - annoTitlePad];
  }

  //================================================================================================
  //                                           BRUSHES
  // The brushes provide a way to zoom and pan on the main heatmap by selecting regions on brushable
  // heatmaps, and they are made up of 2 parts:
  //    * brush component - a d3.brush object (brushX of col, brushY for row) which defines
  //                        important properties of the brush, namely its extent (the maximum pixel
  //                        area that the user can brush) and its interactive behavior (what to do
  //                        when the user starts/stops brushing, or while they are brushing)
  //    * SVG element - a g element which makes the brush visible and usable. By default, it
  //                    contains 4 rect elements; an overlay which lets you create a selection, a
  //                    selection which is draggable, and 2 "handles" on either side of the
  //                    selection which allow it to be resized. The attributes of these elements
  //                    can be programmatically controlled with CSS and JS
  //================================================================================================

  function Brush(dim, upperLeft, lowerRight) {
    this.brush = dim.self === "col" ? d3.brushX() : d3.brushY();
    this.upperLeft = upperLeft;
    this.lowerRight = lowerRight;
    this.index = dim.self === "col" ? 0 : 1;
    this.inverter = d3.scaleQuantize().range(dim.names);
    this.brush.on("brush", function() { brushed(dim); })
              .on("end", function() { ended(dim); });
    this.group = svg.append("g").attr("class", "brush").call(this.brush);
    this.brushToScope = function() {
      this.group.call(this.brush.move,
                    [this.inverter.invertExtent(dim.names[dim.currentScope[0]])[0],
                     this.inverter.invertExtent(dim.names[dim.currentScope[1] - 1])[1] - 1]);
    };
    this.callBrush = function() { this.group.call(this.brush); };
    this.clearBrush = function() { this.group.call(this.brush.move, null); };
    this.extentsSetup = function() {
      this.brush.extent([this.upperLeft(), this.lowerRight()]);
      this.inverter.domain([this.upperLeft()[this.index], this.lowerRight()[this.index]]);
    };
    this.extentsSetup();
  }

  col.brusher = new Brush(col, function() { return col.cellsSub.anchor; },
                               function() { return [col.cellsSub.anchor[0] + col.sizeHeatmap(),
                                                    col.cellsSub.anchor[1] + col.marginBrush]; });
  row.brusher = new Brush(row, function() { return row.cellsSub.anchor; },
                               function() { return [row.cellsSub.anchor[0] + row.marginBrush,
                                                    row.cellsSub.anchor[1] + row.sizeHeatmap()]; });

  //================================================================================================
  //                                           INITIALIZATION
  // One final call to resizeSVG completes the initial rendering of the widget.
  //================================================================================================

  resizeSVG();

  //================================================================================================
  //                                   INTERACTIVITY FUNCTIONS
  // These functions determine all the things that can happen in the widget once it is initially
  // rendered.
  //
  // For the brushes (the tools used to zoom/pan), there are 2 functions, brushed and ended, which
  // handle all the updates to the data structures and DOM that are necessary to perform zoom/pan
  // (with the help of helper functions).
  //
  // For the dropdowns in the settings panel, there are 3 functions, annoUpdate, sortUpdate, and
  // updateColorScaling.
  //================================================================================================

  // updates the current scope of the given dimension and, if renderOnBrushEnd is false, performs
  // visual updates on the main heatmap and side colors
  function brushed(dim) {
    settingsPanel.classed("hidden", true); // hide the settings panel in case it's visible
    settingsHidden = true;
    if (!renderOnBrushEnd) {
      var inverses = d3.event.selection.map(dim.brusher.inverter); // bounds of brushed -> row/column
      dim.currentScope = [dim.names.indexOf(inverses[0]), dim.names.indexOf(inverses[1]) + 1];
      renderScope(dim, false);
    }
  }

  // resets the scope of the given dim only if there is no current selection (i.e., the user clicks
  // off of the selected area, otherwise renders the dim's current scope if renderOnBrushEnd is true
  function ended(dim) {
    if (d3.event.selection) {
      if (renderOnBrushEnd) {
        var inverses = d3.event.selection.map(dim.brusher.inverter); // pixel bounds -> row/column
        dim.currentScope = [dim.names.indexOf(inverses[0]), dim.names.indexOf(inverses[1]) + 1];
        renderScope(dim, true);
      }
    } else {
      settingsPanel.classed("hidden", true); // hide the settings panel in case it's visible
      settingsHidden = true;
      dim.currentScope = [0, dim.names.length];
      // scale updates
      dim.scaleCell.domain(dim.names);
      dim.labels.updateScale(dim.names);
      // visual updates
      dim.labels.update();
      cells.update([dim.pos, dim.size]);
      if (dim.annotated) dim.sideColors.update([dim.pos, dim.size]);
    }
  }

  // renders the currentScope for the given dim. If transition is true, the labels will update with
  // a transition, else they will update without a transition
  function renderScope(dim, transition) {
    var scopeArray = dim.names.slice(dim.currentScope[0], dim.currentScope[1]);
    var inScope = {};
    for (var j = 0; j < scopeArray.length; j++) inScope[scopeArray[j]] = true; // undefined is falsy
    // scale updates
    dim.scaleCell.domain(scopeArray);
    dim.labels.updateScale(scopeArray);
    // visual updates
    transition ? dim.labels.update() : dim.labels.updateNT();
    updateVisualScope(dim, inScope);
  }

  // repositions and resizes the cells of the main heatmap and the side colors of the given
  // dimension, showing only those that are in visible (for which vis[d[dim.self]] is true)
  function updateVisualScope(dim, vis) {
    cells.selection.attr(dim.pos, function(d) { return vis[d[dim.self]] ? cells[dim.pos](d) : 0; })
                  .attr(dim.size, function(d) { return vis[d[dim.self]] ? cells[dim.size]() : 0; });
    if (dim.annotated) dim.sideColors.selection
                .attr(dim.pos, function(d) { return vis[d.key] ? dim.sideColors[dim.pos](d) : 0; })
               .attr(dim.size, function(d) { return vis[d.key] ? dim.sideColors[dim.size]() : 0; });
  }

  // annotates the rows/columns (depending on dim) and updates the respective annotation colors by
  // the currently selected annotation option for the given dimension
  function annoUpdate(dim, newAnnotype) {
    dim.annoBy = newAnnotype;
    var values = dim.annotations[dim.annoBy];
    // scale updates
    dim.annoToNum.domain(values);
    if (categorical) dim.annoToNum.range(d3.range(values.length));
    if (values.every(function(value) { return !isNaN(value); }) && values.length > 2) {
      dim.numToColor = categorical ? function(index) { return dim.annoHeat(index / values.length); }
                                    : dim.annoHeat;
    } else {
      dim.numToColor = dim.annoReg;
    }
    dim.scaleAnnoColor.domain(values);
    dim.labelsAnno.updateScale(values);
    // visual updates
    dim.annoTitle.setText(undersToSpaces(dim.annoBy));
    dim.annoColors.setup(values); // clear previous rects and add new ones in
    dim.labelsAnno.updateNT();
    dim.sideColors.selection.transition().duration(animDuration).attr("fill", dim.sideColors.fill);
  }

  // sorts the rows/columns (depending on dim) of the 3 heatmaps according to the currently selected
  // sorting option for the given dimension
  function sortUpdate(dim, annotype) {
    if (annotype != "Clustered Order") { // sort the rows/columns by the chosen annotype
      var values = dim.annotations[annotype],
          valueToIndex = {}; // hashmap to determine priority for sorting
      for (var j = 0; j < values.length; j++) valueToIndex[values[j]] = j;
      dim.labelsAnnotated.sort(function(a, b) {
        var val1 = valueToIndex[a.annos[annotype]],
            val2 = valueToIndex[b.annos[annotype]];
        return val1 === val2 ? a.key.localeCompare(b.key) : val1 - val2;
      });
    }
    dim.names = annotype === "Clustered Order" ? dim.clustOrder : dim.labelsAnnotated.map(key);
    // update scales
    dim.scaleCell.domain(dim.names);
    dim.scaleCellSub.domain(dim.names);
    dim.brusher.inverter.range(dim.names);
    dim.labelsSub.updateScale(dim.names);
    // visual updates for the brushable heatmaps
    dim.labelsSub.update();
    dim.cellsSub.update([dim.pos]);
    dim.other.cellsSub.update([dim.pos]);
    renderScope(dim, true);
  }

  // updates the fill of the heatmap cells based on the currently selected scaling option
  function updateColorScaling(newScalingDim) {
    scalingDim = newScalingDim;
    if (scalingDim != "bucket") {
      mainColorScale.domain(scalingDim === "none" ? [dataset.stats.totalMin, dataset.stats.totalMax]
                              : [-dataset.stats.zMax[scalingDim], dataset.stats.zMax[scalingDim]]);
    }
    colorKey.change(scalingDim);
    cells.update(["fill"]);
    col.cellsSub.update(["fill"]);
    row.cellsSub.update(["fill"]);
  }

  function resizeSVG() {
    w = parent.clientWidth - margin.left - margin.right;
    h = height - margin.top - margin.bottom;
    svgSetup(w, h);
    marginsSetup(w, h);
    anchorsSetup(w, h);
    scalesSetup(w, h);
    col.brusher.extentsSetup();
    row.brusher.extentsSetup();
    positionAllElements();
    col.currentScope[0] != 0 || col.currentScope[1] != col.names.length ? col.brusher.brushToScope()
                                                                        : col.brusher.clearBrush();
    row.currentScope[0] != 0 || row.currentScope[1] != row.names.length ? row.brusher.brushToScope()
                                                                        : row.brusher.clearBrush();
  }

  //================================================================================================
  //                            TOOLTIP GENERATING/DISPLAYING FUNCTIONS + MISC
  // These handle the setup and displaying of various visual/interactive elements in the heatmap.
  //================================================================================================

  function settingsPanelSetup() {
      var isRawData = true;
      // This test assumes that the normalized data will contain decimal values. For now checking to see if there is at least 1 decimal value. If not then assuming data is raw data counts.
      for (var i = 0; i < dataset.matrix.length; i++) {
          if (dataset.matrix[i][0].value % 1 != 0) {
              i = dataset.matrix.length;
              isRawData = false;
          }
      }

    var panel = container.append("div").attr("id", "settings")
                  .attr("class", "tip").classed("hidden", true);
    panel.append("p").text("Settings");
    panel.append("button").text("Fit to Labels").on("click", resizeSVG);
    panel.append("button").text("Close Settings").on("click", function() {
      settingsHidden = !settingsHidden;
      panel.classed("hidden", settingsHidden);
    });
    var table;

    if (isRawData) {
        table = panel.append("table"),
      row1 = table.append("tr");
        row1.append("td").append("p").text("Scale by");
        scaleBy = row1.append("td").append("select")
                      .on("change", function () { updateColorScaling(this.value); });
        scaleBy.selectAll("option")
          .data([{ value: "bucket", text: "Buckets" },
                 { value: col.self, text: col.title },
                 { value: row.self, text: row.title },
                 { value: "none", text: "None" }])
          .enter()
          .append("option")
          .attr("value", function (d) { return d.value; })
          .text(function (d) { return d.text; });
        scalingDim = "bucket";
    } else {
        table = panel.append("table"),
        row1 = table.append("tr");
        row1.append("td").append("p").text("Scale by");
        scaleBy = row1.append("td").append("select")
                      .on("change", function () { updateColorScaling(this.value); });
        scaleBy.selectAll("option")
          .data([{ value: col.self, text: col.title },
                 { value: row.self, text: row.title },
                 { value: "none", text: "None" },
                 { value: "bucket", text: "Buckets" }])
          .enter()
          .append("option")
          .attr("value", function (d) { return d.value; })
          .text(function (d) { return d.text; });
        scalingDim = col.self;
    }
    if (row.annotated) controlsSetup(row);
    if (col.annotated) controlsSetup(col);

    function controlsSetup(dim) {
      var r1 = table.append("tr"),
          r2 = table.append("tr");
      r1.append("td").append("p").text(dim.title + "s: annotate by");
      r2.append("td").append("p").text(dim.title + "s: sort by");
      dim.annoByDropdown = selectorSetup(r1, dim, annoUpdate);
      dim.sortByDropdown = selectorSetup(r2, dim, sortUpdate);
      dim.annoByDropdown.selectAll("option")
        .data(dim.annotypes)
        .enter()
        .append("option")
        .attr("value", identity)
        .text(function(d) { return undersToSpaces(d); });
      dim.sortByDropdown.selectAll("option")
        .data(["Clustered Order"].concat(dim.annotypes))
        .enter()
        .append("option")
        .attr("value", identity)
        .text(function(d) { return undersToSpaces(d); });
    }

    function selectorSetup(s, dim, update) {
      return s.append("td").append("select").on("change", function() { update(dim, this.value); });
    }

    return panel;
  }

  // sets up the tooltip for hovering over the cells of the main heatmap
  function cellTooltipSetup() {
    var tooltip = container.append("div").attr("class", "tip").classed("hidden", true);
    tooltip.append("p").text("Cell Info");
    var table = tooltip.append("table"),
        row1 = table.append("tr"),
        row2 = table.append("tr"),
        row3 = table.append("tr");
    row1.append("td").append("p").text("Value");
    row1.append("td").append("p").attr("id", "value");
    row2.append("td").append("p").text("Row");
    row2.append("td").append("p").attr("id", "row");
    row3.append("td").append("p").text("Column");
    row3.append("td").append("p").attr("id", "col");
    return tooltip;
  }

  // sets up the tooltip for hovering over row/column (determined by dim) side colors
  function sideTooltipSetup(dim) {
    var tooltip = container.append("div").attr("class", "tip").classed("hidden", true);
    tooltip.append("p").text(dim.title + " Info");
    var table = tooltip.append("table"),
        rows = table.selectAll("tr")
                .data(Object.keys(dim.labelsAnnotated[0].annos))
                .enter()
                .append("tr");
    rows.append("td").append("p").text(function(d) { return undersToSpaces(d); });
    rows.append("td").append("p").attr("id", identity);
    return tooltip;
  }

  // sets up the tooltip for hovering over the annotation colors
  function annoTooltipSetup() {
    var tooltip = container.append("div").attr("class", "tip").classed("hidden", true);
    tooltip.append("p").text("Annotation Info");
    var row1 = tooltip.append("table").append("tr");
    row1.append("td").append("p").attr("id", "annotype");
    row1.append("td").append("p").attr("id", "value");
    return tooltip;
  }

  // positions the settings panel at the lower-right corner of the cell (clickedRect), with width
  // function widthOffset and height function heightOffset. Sets settingsHidden to !settingsHidden
  // and then hides the given tooltip if settingsHidden is false and hides the settings panel if
  // settingsHidden is true (else shows the settings panel)
  function toggleSettingsPanel(clickedRect, widthOffset, heightOffset, tooltip) {
    settingsHidden = !settingsHidden;
    if (!settingsHidden) tooltip.classed("hidden", true);
    var box = clickedRect.getBoundingClientRect(),
        anchor = [box.left + widthOffset() + window.pageXOffset,
                  box.top + heightOffset() + window.pageYOffset];
    toggleTooltip(settingsPanel, "left", "top", anchor, settingsHidden);
  }

  // displays the tooltip for the heatmap cell (mousedOverRect) with the given data d
  function displayCellTooltip(d, mousedOverRect) {
    var box = mousedOverRect.getBoundingClientRect(),
        anchor = [box.left + cells.width() + window.pageXOffset,
                  box.top + cells.height() + window.pageYOffset];
    toggleTooltip(cellTooltip, "left", "top", anchor, false);
    cellTooltip.select("#value").text(d.value);
    cellTooltip.select("#row").text(d.row);
    cellTooltip.select("#col").text(d.col);
  }

  // displays the tooltip for the side color cell (mousedOverRect) with data d of the given dim
  function displaySideTooltip(d, mousedOverRect, dim) {
    var box = mousedOverRect.getBoundingClientRect(),
        anchor = [box.left + dim.sideColors.width() + window.pageXOffset,
                  box.top + dim.sideColors.height() + window.pageYOffset];
    toggleTooltip(dim.tooltip, "left", "top", anchor, false);
    var annotypes = Object.keys(d.annos);
    for (var j = 0; j < annotypes.length; j++) {
      dim.tooltip.select("#" + annotypes[j]).text(d.annos[annotypes[j]]);
    }
  }

  // displays the tooltip for the annotation cell (mouserOverRect) with data d of the given dim
  // TODO: fix scroll bar weirdness for Windows (document.body.offsetHeight > window.innerHeight ?)
  function displayAnnoTooltip(d, mousedOverRect, dim) {
    var box = mousedOverRect.getBoundingClientRect(),
        anchor = [window.innerWidth - box.left - window.pageXOffset, box.top + window.pageYOffset];
    toggleTooltip(annoTooltip, "right", "top", anchor, false);
    annoTooltip.select("#annotype").text(undersToSpaces(dim.annoBy));
    annoTooltip.select("#value").text(d);
  }

  function toggleTooltip(tip, xPos, yPos, anchor, hidden) {
    tip.style(xPos, anchor[0] + "px").style(yPos, anchor[1] + "px").classed("hidden", hidden);
  }

  function positionAllElements() {
    cells.position();
    cells.update(["x", "y", "width", "height"]);
    colorKey.updateCells(["x", "y", "width", "height"]);
    colorKey.positionCells();
    colorKey.positionLabels();
    colorKey.updateNTLabels();
    colorKey.positionTitles();
    colorKey.change(scalingDim);
    positionElements(col);
    positionElements(row);

    function positionElements(dim) {
      dim.labels.position();
      dim.labels.updateNT();
      dim.labelsSub.position();
      dim.labelsSub.updateNT();
      dim.cellsSub.position();
      dim.cellsSub.update(["x", "y", "width", "height"]);
      dim.brusher.callBrush();
      if (dim.annotated) {
        dim.labelsAnno.position();
        dim.labelsAnno.updateNT();
        dim.sideColors.position();
        dim.sideColors.update(["x", "y", "width", "height"]);
        dim.annoColors.position();
        dim.annoColors.update(["x", "y", "width", "height"]);
        dim.annoTitle.position();
      }
    }
  }

  function svgSetup(w, h) {
    SVG.attr("width", w + margin.left + margin.right)
      .attr("height", h + margin.top + margin.bottom);
  }

  // places the given element (e) at the anchor point (a)
  function positionElement(e, a) { e.attr("transform", "translate(" + a[0] + "," + a[1] + ")"); }

  //================================================================================================
  //                                     OTHER HELPER FUNCTIONS
  //================================================================================================

  function colorsSetup(dim, cat, categoricalScheme, continuousScheme, annoHeatScheme) {
    var categoricalSchemes = {
          ns:       ["#7fff00", "#eead0e", "#00b2ee", "#ee2c2c", "#bf3eff",
                     "#d2b48c", "#6959cd", "#228b22", "#ff7f50", "#7a7a7a"],
          google:   ["#3366cc", "#dc3912", "#ff9900", "#109618", "#990099",
                     "#0099c6", "#dd4477", "#66aa00", "#b82e2e", "#316395",
                     "#994499", "#22aa99", "#aaaa11", "#6633cc", "#e67300",
                     "#8b0707", "#651067", "#329262", "#5574a6", "#3b3eac"],
          rainbow:  ["#843c39", "#ad494a", "#d6616b", "#e7969c", "#e6550d",
                     "#fd8d3c", "#fdae6b", "#fdd0a2", "#8c6d31", "#bd9e39",
                     "#e7ba52", "#e7cb94", "#637939", "#8ca252", "#b5cf6b",
                     "#cedb9c", "#31a354", "#74c476", "#a1d99b", "#c7e9c0",
                     "#3182bd", "#6baed6", "#9ecae1", "#c6dbef", "#393b79",
                     "#5254a3", "#6b6ecf", "#9c9ede", "#756bb1", "#9e9ac8",
                     "#bcbddc", "#dadaeb"," #7b4173", "#a55194", "#ce6dbd",
                     "#de9ed6", "#636363", "#969696", "#bdbdbd"," #d9d9d9"]
        },
        continuousSchemes = {
          cubehelix:  d3.interpolateCubehelixDefault,
          rainbow:    d3.interpolateRainbow
        },
        annoHeatSchemes = {
          viridis:  d3.interpolateViridis,
          inferno:  d3.interpolateInferno,
          magma:    d3.interpolateMagma,
          plasma:   d3.interpolatePlasma,
          warm:     d3.interpolateWarm,
          cool:     d3.interpolateCool
        };
    var catColors = categoricalSchemes[categoricalScheme],
        conColors = continuousSchemes[continuousScheme];
    dim.annoReg = cat ? function(index) { return catColors[index % catColors.length]; } : conColors;
    dim.annoHeat = annoHeatSchemes[annoHeatScheme];
  }

  // returns the size, in pixels, of the heatmap along the given dim (height - col, width - row),
  // but does not take into account margins used for color keys
  function sizeHeatmap(dim) {
    return dim.marginTotal - dim.marginSideColor - dim.marginLabel - dim.marginBrush
                                                                              - dim.marginLabelSub;
  }

  //================================================================================================
  //                                         PARSING FUNCTIONS
  // These take strings in CSV format and turn them into the data structures needed for the heatmap.
  //================================================================================================

  // parses the given string into the data structures used for generating the heatmap
  function parseDataMatrix(file) {
    var parsedRows = d3.csvParseRows(file); // parses the string into an array of arrays
    var colnames = parsedRows.shift(); // column names should be stored in the first row
    colnames.shift(); // removes whatever name was given to the column containing the row names
    var rownames = []; // the array of rownames will grow as we process each row
    // stats will hold all relevant statistics for the dataset
    var stats = {
                  col: {},
                  row: {},
                  zMax: { col: 0, row: 0 },
                  totalMin: Number.POSITIVE_INFINITY,
                  totalMax: Number.NEGATIVE_INFINITY
    };

    // traverse the parsed rows to create the matrix (a doubly-nested array) for the heatmap, adding
    // to the rownames array and updating the stats object as we go
    var matrix = d3.range(parsedRows.length).map(function(j) { // j = index of parsedRows
        // grab the row name out of the parsed row. This makes parsedRows[j] the same length as
        // colnames, with parsedRows[j][k] being the value in row 'rowname' and column 'colnames[k]'
        var rowname = parsedRows[j].shift();
        // add the new row name to the list of row names
        rownames.push(rowname);
        // traverse the parsed row, reformatting each element (a number) and updating stats
        return d3.range(colnames.length).map(function(k) { // k = index of colnames
          // the "+" converts parsedRows[j][k] to a number (since it was parsed as a string)
          var value = +parsedRows[j][k];
          // update the stats for the current column and the current row with this value
          updateStats(stats, "col", dotsToUnders(colnames[k]), value);
          updateStats(stats, "row", dotsToUnders(rowname), value);
          return {
            key: j + " " + k, // useful for d3 data joins
            row: rowname,     // determines cell attributes (position (y), size (height))
            col: colnames[k], // determines cell attributes (position (x), size (width))
            value: value      // determines cell attributes (fill)
          };
        });
    });

    // perform final calculations of the stats for each column, and find the totalMin and totalMax
    // of the dataset (this could also be done in the final calculations for the row stats)
    var cStatNames = Object.keys(stats.col);
    for (var j = 0; j < cStatNames.length; j++) {
      finalCalculations(stats, "col", cStatNames[j], rownames.length);
      stats.totalMin = Math.min(stats.totalMin, stats.col[cStatNames[j]].min); // reassign if needed
      stats.totalMax = Math.max(stats.totalMax, stats.col[cStatNames[j]].max); // reassign if needed
    }
    // perform final calculations of the stats for each row
    var rStatNames = Object.keys(stats.row);
    for (var j = 0; j < rStatNames.length; j++) {
      finalCalculations(stats, "row", rStatNames[j], colnames.length);
    }
    // find the z-score in the dataset with the largest magnitude
    for (var j = 0; j < matrix.length; j++) {
      for (var k = 0; k < matrix[j].length; k++) {
        var value = matrix[j][k].value, // grab the value and compute its z-score for to its row/col
            colname = dotsToUnders(colnames[k]),
            rowname = dotsToUnders(rownames[j]),
            colZ = (value - stats.col[colname].mean) / stats.col[colname].stdev,
            rowZ = (value - stats.row[rowname].mean) / stats.row[rowname].stdev;
        stats.zMax.col = Math.max(stats.zMax.col, Math.abs(colZ)); // reassign max if necessary
        stats.zMax.row = Math.max(stats.zMax.row, Math.abs(rowZ)); // reassign max if necessary
      }
    }

    // updates the stats object for the given dimension at the given name with the given value
    function updateStats(stats, dim, name, value) {
      if (stats[dim][name] === undefined) { // if unseen, give it a fresh new stats object
        // an stdev field will be added to this object during final calculations
        stats[dim][name] = {
          min: value,       // helps to find most negative z-score
          max: value,       // helps to find most positive z-score
          mean: 0,          // used in calculating standard deviation/z-scores for cell fills
          meanOfSquares: 0  // used in calculating standard deviation
        };
      }
      if (value < stats[dim][name].min) stats[dim][name].min = value; // reassign min if necessary
      if (value > stats[dim][name].max) stats[dim][name].max = value; // reassign max if necessary
      stats[dim][name].mean += value;                   // this will be averaged later
      stats[dim][name].meanOfSquares += value * value;  // this will be averaged later
    }

    // adds the stdev field to the stats object for the dimension at the given name
    function finalCalculations(stats, dim, name, numVals) {
      stats[dim][name].mean *= (1 / numVals);
      stats[dim][name].meanOfSquares *= (1 / numVals);
      stats[dim][name].stdev = Math.sqrt(stats[dim][name].meanOfSquares
                                                              - Math.pow(stats[dim][name].mean, 2));
    }

    return {
      matrix: matrix,     // array of arrays of objects (cells have value, row, col, key)
      rownames: rownames, // arrays of strings (list of all row names, assumed to be clustered)
      colnames: colnames, // arrays of strings (list of all column names, assumed to be clustered)
      stats: stats        // object with 5 fields: row and col (hashmaps from row/col name to object
                          // of statistics, zMax stores largest z-score (by magnitude) for both row
                          // and col, and totalMin/totalMax store min and max of the entire dataset
    };
  }

  // parses the given string into the data structures used for annotating/sorting the heatmap
  function parseAnnotations(file) {
    file = file.charAt(0) === "," ? "Name" + file : file; // add a nameKey if there isn't one
    var na = "{ no data }"; // this will be used as a readable name if a cell only holds ""
    var parsedRows = d3.csvParseRows(file); // parse the file into an array of arrays
    // the names of the different kinds of annotations should be stored in the first row of the file
    var annotypes = parsedRows.shift(); // pops off the first element (ACTUALLY modifies parsedRows)
    annotypes = annotypes.map(dotsToUnders); // periods in names of annotypes will mess up JS code

    var shouldShift = true;
    if (annotypes[0]) {
        var nameKey = annotypes[0]; // First position in the table contains a header, don't shift.
        shouldShift = false;
    } else {
        var nameKey = annotypes.shift(); // trims annotypes down to JUST the actual annotation types
    }

    var annotations = {}; // each type of annotation will map to a sorted array of its unique values
    for (var j = 0; j < annotypes.length; j++) annotations[annotypes[j]] = [];

    // in these nested loops, examine all values for each annotation type and add them to the
    // hashmap of annotation types -> array of unique values
    for (var j = 0; j < parsedRows.length; j++) {
      // toss out the first element in the row (the name of the dimension for this value); what's
      // left is an array of the same length as the annotypes array, with values[k] being a value
        // for the annotation type annotypes[k]
        if (shouldShift) {
            parsedRows[j].shift();
        }
      var values = parsedRows[j];
      // associate new unique values with their corresponding annotation types as necessary
      for (var k = 0; k < annotypes.length; k++) {
        var value = values[k] || na; // give a readable name if blank
        // add this value into the array of unique values for its corresponding annotation type
        if (annotations[annotypes[k]].indexOf(value) < 0) annotations[annotypes[k]].push(value);
      }
    }

    // sort each annotation type's values (numerically if both numbers, otherwise lexicographically)
    for (var j = 0; j < annotypes.length; j++) {
      annotations[annotypes[j]].sort(function(a, b) {
        if (!isNaN(a) && !isNaN(b)) return (+a) - (+b); // the "+" converts a and b to numbers
        return a.localeCompare(b);
      });
    }

    // parse the file into an array of objects (each dimension (row or column) is grouped with all
    // its values for each annotation type, with the column names for the file being the keys). Then
    // restructure and reformat the elements so that each object is now a nested object, with a key
    // field holding the original object's value for the nameKey and an annos holding the entire
    // original object (reformatted). This allows for easier lookup of the annotations for a given
    // row/column name, and makes d3 data joins easier
    var labels = d3.csvParse(file).map(function(obj) {
      // reformat so that keys contain no periods and values are renamed if blank
      var objClean = {}, keys = Object.keys(obj);
      for (var j = 0; j < keys.length; j++) objClean[dotsToUnders(keys[j])] = obj[keys[j]] || na;
      return {
        key: objClean[nameKey], // this corresponds to the name of the row/column
        annos: objClean
      };
    });

    return {
      annotations: annotations, // hashmap string to string[] (annotation types -> values)
      labels: labels            // array of objects (list of annotated dimension names)
    };
  }
}

//--------------------------------------------------------------------------------------------------
//                                        END HEATMAP FUNCTION
//--------------------------------------------------------------------------------------------------
```
