## Supplementary material for "Comparative pathway enrichment analysis in gastrointestinal cell lines Caco-2, HT-29, HEPG2, and colon fibroblasts using a custom expression panel for tight-junction and cytoskeletal regulatory genes": Sup. Results 1: helpers.js

```
// returns the given array if its length is less than or equal to the given max, otherwise returns
// a new array of length max whose elements are a subset of the given array such that their
// distance from each other in the given array is approximately even (surely there's a better way
// to word this...)
function sample(array, max) {
  if (array.length <= max) {
    return array;
  } else {
    var sampler = d3.scaleLinear().domain([0, max - 1]).range([0, array.length - 1]),
        sampledIndices = d3.range(max).map(function(i) { return Math.floor(sampler(i)); });
    return sampledIndices.map(function(i) { return array[i]; });
  }
}

// returns the given string with all period characters replaced with underscores
function dotsToUnders(str) {
  return str.replace(/\./g, "_");
}

// returns the given string will all underscore characters replaced with spaces (necessary???)
function undersToSpaces(str) {
  return str.replace(/_/g, " ");
}

// returns an array of the given length whose elements are hex colors representing an interpolated
// gradient from low to mid to high (if length is odd, then there will be one more color in the
// high end than in the low end)
function interpolateColors(low, mid, high, length) {
  var lowToMidF = d3.interpolateLab(low, mid),
      lowToMid = d3.range(Math.floor(length / 2))
                   .map(function(j) { return lowToMidF(j / Math.floor(length / 2)); }),
      midToHighF = d3.interpolateLab(mid, high),
      midToHigh = d3.range(Math.ceil(length / 2))
                    .map(function(j) { return midToHighF(j / Math.ceil(length / 2)); });
  return lowToMid.concat(midToHigh);
}

function Bucketizer(dividers, colors) {
  this.domain = dividers.map(function(d) { return d; }); // copy
  this.domain.push(Number.POSITIVE_INFINITY);
  this.range = colors;
  this.bucketize = function(value) {
    for (var j = 0; j < this.domain.length - 1; j++) {
      if (value < this.domain[j] && value < this.domain[j + 1]) return this.range[j];
    }
    return this.range[this.range.length - 1];
  }
}

// returns the key field of the given object
function key(d) { return d.key; }

// return the given object
function identity(d) { return d; }
```
