## Supplementary material for "Comparative pathway enrichment analysis in gastrointestinal cell lines Caco-2, HT-29, HEPG2, and colon fibroblasts using a custom expression panel for tight-junction and cytoskeletal regulatory genes": Sup. Results 1: jquery.cookie.js

```
/*!
 * jQuery Cookie Plugin v1.4.1
 * https://github.com/carhartl/jquery-cookie
 *
 * Copyright 2013 Klaus Hartl
 * Released under the MIT license
 */
!function(a){"function"==typeof define&&define.amd?define(["jquery"],a):a("object"==typeof exports?require("jquery"):jQuery)}(function(a){function b(a){return h.raw?a:encodeURIComponent(a)}function c(a){return h.raw?a:decodeURIComponent(a)}function d(a){return b(h.json?JSON.stringify(a):String(a))}function e(a){0===a.indexOf('"')&&(a=a.slice(1,-1).replace(/\\"/g,'"').replace(/\\\\/g,"\\"));try{return a=decodeURIComponent(a.replace(g," ")),h.json?JSON.parse(a):a}catch(b){}}function f(b,c){var d=h.raw?b:e(b);return a.isFunction(c)?c(d):d}var g=/\+/g,h=a.cookie=function(e,g,i){if(void 0!==g&&!a.isFunction(g)){if(i=a.extend({},h.defaults,i),"number"==typeof i.expires){var j=i.expires,k=i.expires=new Date;k.setTime(+k+864e5*j)}return document.cookie=[b(e),"=",d(g),i.expires?"; expires="+i.expires.toUTCString():"",i.path?"; path="+i.path:"",i.domain?"; domain="+i.domain:"",i.secure?"; secure":""].join("")}for(var l=e?void 0:{},m=document.cookie?document.cookie.split("; "):[],n=0,o=m.length;o>n;n++){var p=m[n].split("="),q=c(p.shift()),r=p.join("=");if(e&&e===q){l=f(r,g);break}e||void 0===(r=f(r))||(l[q]=r)}return l};h.defaults={},a.removeCookie=function(b,c){return void 0===a.cookie(b)?!1:(a.cookie(b,"",a.extend({},c,{expires:-1})),!a.cookie(b))}});
```
