## Supplementary material for "Comparative pathway enrichment analysis in gastrointestinal cell lines Caco-2, HT-29, HEPG2, and colon fibroblasts using a custom expression panel for tight-junction and cytoskeletal regulatory genes": Sup. Results 1: jquery.dataTables.min.js

```
/*! DataTables 1.10.6
 * ©2008-2015 SpryMedia Ltd - datatables.net/license
 */
(function(Ea,P,k){var O=function(h){function V(a){var b,c,e={};h.each(a,function(d){if((b=d.match(/^([^A-Z]+?)([A-Z])/))&&-1!=="a aa ai ao as b fn i m o s ".indexOf(b[1]+" "))c=d.replace(b[0],b[2].toLowerCase()),e[c]=d,"o"===b[1]&&V(a[d])});a._hungarianMap=e}function H(a,b,c){a._hungarianMap||V(a);var e;h.each(b,function(d){e=a._hungarianMap[d];if(e!==k&&(c||b[e]===k))"o"===e.charAt(0)?(b[e]||(b[e]={}),h.extend(!0,b[e],b[d]),H(a[e],b[e],c)):b[e]=b[d]})}function O(a){var b=m.defaults.oLanguage,c=a.sZeroRecords;
!a.sEmptyTable&&(c&&"No data available in table"===b.sEmptyTable)&&E(a,a,"sZeroRecords","sEmptyTable");!a.sLoadingRecords&&(c&&"Loading..."===b.sLoadingRecords)&&E(a,a,"sZeroRecords","sLoadingRecords");a.sInfoThousands&&(a.sThousands=a.sInfoThousands);(a=a.sDecimal)&&db(a)}function eb(a){A(a,"ordering","bSort");A(a,"orderMulti","bSortMulti");A(a,"orderClasses","bSortClasses");A(a,"orderCellsTop","bSortCellsTop");A(a,"order","aaSorting");A(a,"orderFixed","aaSortingFixed");A(a,"paging","bPaginate");
A(a,"pagingType","sPaginationType");A(a,"pageLength","iDisplayLength");A(a,"searching","bFilter");if(a=a.aoSearchCols)for(var b=0,c=a.length;b<c;b++)a[b]&&H(m.models.oSearch,a[b])}function fb(a){A(a,"orderable","bSortable");A(a,"orderData","aDataSort");A(a,"orderSequence","asSorting");A(a,"orderDataType","sortDataType");var b=a.aDataSort;b&&!h.isArray(b)&&(a.aDataSort=[b])}function gb(a){var a=a.oBrowser,b=h("<div/>").css({position:"absolute",top:0,left:0,height:1,width:1,overflow:"hidden"}).append(h("<div/>").css({position:"absolute",
top:1,left:1,width:100,overflow:"scroll"}).append(h('<div class="test"/>').css({width:"100%",height:10}))).appendTo("body"),c=b.find(".test");a.bScrollOversize=100===c[0].offsetWidth;a.bScrollbarLeft=1!==Math.round(c.offset().left);b.remove()}function hb(a,b,c,e,d,f){var g,i=!1;c!==k&&(g=c,i=!0);for(;e!==d;)a.hasOwnProperty(e)&&(g=i?b(g,a[e],e,a):a[e],i=!0,e+=f);return g}function Fa(a,b){var c=m.defaults.column,e=a.aoColumns.length,c=h.extend({},m.models.oColumn,c,{nTh:b?b:P.createElement("th"),sTitle:c.sTitle?
c.sTitle:b?b.innerHTML:"",aDataSort:c.aDataSort?c.aDataSort:[e],mData:c.mData?c.mData:e,idx:e});a.aoColumns.push(c);c=a.aoPreSearchCols;c[e]=h.extend({},m.models.oSearch,c[e]);ka(a,e,h(b).data())}function ka(a,b,c){var b=a.aoColumns[b],e=a.oClasses,d=h(b.nTh);if(!b.sWidthOrig){b.sWidthOrig=d.attr("width")||null;var f=(d.attr("style")||"").match(/width:\s*(\d+[pxem%]+)/);f&&(b.sWidthOrig=f[1])}c!==k&&null!==c&&(fb(c),H(m.defaults.column,c),c.mDataProp!==k&&!c.mData&&(c.mData=c.mDataProp),c.sType&&
(b._sManualType=c.sType),c.className&&!c.sClass&&(c.sClass=c.className),h.extend(b,c),E(b,c,"sWidth","sWidthOrig"),c.iDataSort!==k&&(b.aDataSort=[c.iDataSort]),E(b,c,"aDataSort"));var g=b.mData,i=W(g),j=b.mRender?W(b.mRender):null,c=function(a){return"string"===typeof a&&-1!==a.indexOf("@")};b._bAttrSrc=h.isPlainObject(g)&&(c(g.sort)||c(g.type)||c(g.filter));b.fnGetData=function(a,b,c){var e=i(a,b,k,c);return j&&b?j(e,b,a,c):e};b.fnSetData=function(a,b,c){return Q(g)(a,b,c)};"number"!==typeof g&&
(a._rowReadObject=!0);a.oFeatures.bSort||(b.bSortable=!1,d.addClass(e.sSortableNone));a=-1!==h.inArray("asc",b.asSorting);c=-1!==h.inArray("desc",b.asSorting);!b.bSortable||!a&&!c?(b.sSortingClass=e.sSortableNone,b.sSortingClassJUI=""):a&&!c?(b.sSortingClass=e.sSortableAsc,b.sSortingClassJUI=e.sSortJUIAscAllowed):!a&&c?(b.sSortingClass=e.sSortableDesc,b.sSortingClassJUI=e.sSortJUIDescAllowed):(b.sSortingClass=e.sSortable,b.sSortingClassJUI=e.sSortJUI)}function X(a){if(!1!==a.oFeatures.bAutoWidth){var b=
a.aoColumns;Ga(a);for(var c=0,e=b.length;c<e;c++)b[c].nTh.style.width=b[c].sWidth}b=a.oScroll;(""!==b.sY||""!==b.sX)&&Y(a);w(a,null,"column-sizing",[a])}function la(a,b){var c=Z(a,"bVisible");return"number"===typeof c[b]?c[b]:null}function $(a,b){var c=Z(a,"bVisible"),c=h.inArray(b,c);return-1!==c?c:null}function aa(a){return Z(a,"bVisible").length}function Z(a,b){var c=[];h.map(a.aoColumns,function(a,d){a[b]&&c.push(d)});return c}function Ha(a){var b=a.aoColumns,c=a.aoData,e=m.ext.type.detect,d,
f,g,i,j,h,l,q,o;d=0;for(f=b.length;d<f;d++)if(l=b[d],o=[],!l.sType&&l._sManualType)l.sType=l._sManualType;else if(!l.sType){g=0;for(i=e.length;g<i;g++){j=0;for(h=c.length;j<h;j++){o[j]===k&&(o[j]=y(a,j,d,"type"));q=e[g](o[j],a);if(!q&&g!==e.length-1)break;if("html"===q)break}if(q){l.sType=q;break}}l.sType||(l.sType="string")}}function ib(a,b,c,e){var d,f,g,i,j,n,l=a.aoColumns;if(b)for(d=b.length-1;0<=d;d--){n=b[d];var q=n.targets!==k?n.targets:n.aTargets;h.isArray(q)||(q=[q]);f=0;for(g=q.length;f<
g;f++)if("number"===typeof q[f]&&0<=q[f]){for(;l.length<=q[f];)Fa(a);e(q[f],n)}else if("number"===typeof q[f]&&0>q[f])e(l.length+q[f],n);else if("string"===typeof q[f]){i=0;for(j=l.length;i<j;i++)("_all"==q[f]||h(l[i].nTh).hasClass(q[f]))&&e(i,n)}}if(c){d=0;for(a=c.length;d<a;d++)e(d,c[d])}}function J(a,b,c,e){var d=a.aoData.length,f=h.extend(!0,{},m.models.oRow,{src:c?"dom":"data"});f._aData=b;a.aoData.push(f);for(var b=a.aoColumns,f=0,g=b.length;f<g;f++)c&&Ia(a,d,f,y(a,d,f)),b[f].sType=null;a.aiDisplayMaster.push(d);
(c||!a.oFeatures.bDeferRender)&&Ja(a,d,c,e);return d}function ma(a,b){var c;b instanceof h||(b=h(b));return b.map(function(b,d){c=na(a,d);return J(a,c.data,d,c.cells)})}function y(a,b,c,e){var d=a.iDraw,f=a.aoColumns[c],g=a.aoData[b]._aData,i=f.sDefaultContent,c=f.fnGetData(g,e,{settings:a,row:b,col:c});if(c===k)return a.iDrawError!=d&&null===i&&(R(a,0,"Requested unknown parameter "+("function"==typeof f.mData?"{function}":"'"+f.mData+"'")+" for row "+b,4),a.iDrawError=d),i;if((c===g||null===c)&&
null!==i)c=i;else if("function"===typeof c)return c.call(g);return null===c&&"display"==e?"":c}function Ia(a,b,c,e){a.aoColumns[c].fnSetData(a.aoData[b]._aData,e,{settings:a,row:b,col:c})}function Ka(a){return h.map(a.match(/(\\.|[^\.])+/g),function(a){return a.replace(/\\./g,".")})}function W(a){if(h.isPlainObject(a)){var b={};h.each(a,function(a,c){c&&(b[a]=W(c))});return function(a,c,f,g){var i=b[c]||b._;return i!==k?i(a,c,f,g):a}}if(null===a)return function(a){return a};if("function"===typeof a)return function(b,
c,f,g){return a(b,c,f,g)};if("string"===typeof a&&(-1!==a.indexOf(".")||-1!==a.indexOf("[")||-1!==a.indexOf("("))){var c=function(a,b,f){var g,i;if(""!==f){i=Ka(f);for(var j=0,h=i.length;j<h;j++){f=i[j].match(ba);g=i[j].match(S);if(f){i[j]=i[j].replace(ba,"");""!==i[j]&&(a=a[i[j]]);g=[];i.splice(0,j+1);i=i.join(".");j=0;for(h=a.length;j<h;j++)g.push(c(a[j],b,i));a=f[0].substring(1,f[0].length-1);a=""===a?g:g.join(a);break}else if(g){i[j]=i[j].replace(S,"");a=a[i[j]]();continue}if(null===a||a[i[j]]===
k)return k;a=a[i[j]]}}return a};return function(b,d){return c(b,d,a)}}return function(b){return b[a]}}function Q(a){if(h.isPlainObject(a))return Q(a._);if(null===a)return function(){};if("function"===typeof a)return function(b,e,d){a(b,"set",e,d)};if("string"===typeof a&&(-1!==a.indexOf(".")||-1!==a.indexOf("[")||-1!==a.indexOf("("))){var b=function(a,e,d){var d=Ka(d),f;f=d[d.length-1];for(var g,i,j=0,h=d.length-1;j<h;j++){g=d[j].match(ba);i=d[j].match(S);if(g){d[j]=d[j].replace(ba,"");a[d[j]]=[];
f=d.slice();f.splice(0,j+1);g=f.join(".");i=0;for(h=e.length;i<h;i++)f={},b(f,e[i],g),a[d[j]].push(f);return}i&&(d[j]=d[j].replace(S,""),a=a[d[j]](e));if(null===a[d[j]]||a[d[j]]===k)a[d[j]]={};a=a[d[j]]}if(f.match(S))a[f.replace(S,"")](e);else a[f.replace(ba,"")]=e};return function(c,e){return b(c,e,a)}}return function(b,e){b[a]=e}}function La(a){return D(a.aoData,"_aData")}function oa(a){a.aoData.length=0;a.aiDisplayMaster.length=0;a.aiDisplay.length=0}function pa(a,b,c){for(var e=-1,d=0,f=a.length;d<
f;d++)a[d]==b?e=d:a[d]>b&&a[d]--; -1!=e&&c===k&&a.splice(e,1)}function ca(a,b,c,e){var d=a.aoData[b],f,g=function(c,f){for(;c.childNodes.length;)c.removeChild(c.firstChild);c.innerHTML=y(a,b,f,"display")};if("dom"===c||(!c||"auto"===c)&&"dom"===d.src)d._aData=na(a,d,e,e===k?k:d._aData).data;else{var i=d.anCells;if(i)if(e!==k)g(i[e],e);else{c=0;for(f=i.length;c<f;c++)g(i[c],c)}}d._aSortData=null;d._aFilterData=null;g=a.aoColumns;if(e!==k)g[e].sType=null;else{c=0;for(f=g.length;c<f;c++)g[c].sType=null;
Ma(d)}}function na(a,b,c,e){var d=[],f=b.firstChild,g,i=0,j,n=a.aoColumns,l=a._rowReadObject,e=e||l?{}:[],q=function(a,b){if("string"===typeof a){var c=a.indexOf("@");-1!==c&&(c=a.substring(c+1),Q(a)(e,b.getAttribute(c)))}},a=function(a){if(c===k||c===i)g=n[i],j=h.trim(a.innerHTML),g&&g._bAttrSrc?(Q(g.mData._)(e,j),q(g.mData.sort,a),q(g.mData.type,a),q(g.mData.filter,a)):l?(g._setter||(g._setter=Q(g.mData)),g._setter(e,j)):e[i]=j;i++};if(f)for(;f;){b=f.nodeName.toUpperCase();if("TD"==b||"TH"==b)a(f),
d.push(f);f=f.nextSibling}else{d=b.anCells;f=0;for(b=d.length;f<b;f++)a(d[f])}return{data:e,cells:d}}function Ja(a,b,c,e){var d=a.aoData[b],f=d._aData,g=[],i,j,h,l,q;if(null===d.nTr){i=c||P.createElement("tr");d.nTr=i;d.anCells=g;i._DT_RowIndex=b;Ma(d);l=0;for(q=a.aoColumns.length;l<q;l++){h=a.aoColumns[l];j=c?e[l]:P.createElement(h.sCellType);g.push(j);if(!c||h.mRender||h.mData!==l)j.innerHTML=y(a,b,l,"display");h.sClass&&(j.className+=" "+h.sClass);h.bVisible&&!c?i.appendChild(j):!h.bVisible&&c&&
j.parentNode.removeChild(j);h.fnCreatedCell&&h.fnCreatedCell.call(a.oInstance,j,y(a,b,l),f,b,l)}w(a,"aoRowCreatedCallback",null,[i,f,b])}d.nTr.setAttribute("role","row")}function Ma(a){var b=a.nTr,c=a._aData;if(b){c.DT_RowId&&(b.id=c.DT_RowId);if(c.DT_RowClass){var e=c.DT_RowClass.split(" ");a.__rowc=a.__rowc?Na(a.__rowc.concat(e)):e;h(b).removeClass(a.__rowc.join(" ")).addClass(c.DT_RowClass)}c.DT_RowAttr&&h(b).attr(c.DT_RowAttr);c.DT_RowData&&h(b).data(c.DT_RowData)}}function jb(a){var b,c,e,d,
f,g=a.nTHead,i=a.nTFoot,j=0===h("th, td",g).length,n=a.oClasses,l=a.aoColumns;j&&(d=h("<tr/>").appendTo(g));b=0;for(c=l.length;b<c;b++)f=l[b],e=h(f.nTh).addClass(f.sClass),j&&e.appendTo(d),a.oFeatures.bSort&&(e.addClass(f.sSortingClass),!1!==f.bSortable&&(e.attr("tabindex",a.iTabIndex).attr("aria-controls",a.sTableId),Oa(a,f.nTh,b))),f.sTitle!=e.html()&&e.html(f.sTitle),Pa(a,"header")(a,e,f,n);j&&da(a.aoHeader,g);h(g).find(">tr").attr("role","row");h(g).find(">tr>th, >tr>td").addClass(n.sHeaderTH);
h(i).find(">tr>th, >tr>td").addClass(n.sFooterTH);if(null!==i){a=a.aoFooter[0];b=0;for(c=a.length;b<c;b++)f=l[b],f.nTf=a[b].cell,f.sClass&&h(f.nTf).addClass(f.sClass)}}function ea(a,b,c){var e,d,f,g=[],i=[],j=a.aoColumns.length,n;if(b){c===k&&(c=!1);e=0;for(d=b.length;e<d;e++){g[e]=b[e].slice();g[e].nTr=b[e].nTr;for(f=j-1;0<=f;f--)!a.aoColumns[f].bVisible&&!c&&g[e].splice(f,1);i.push([])}e=0;for(d=g.length;e<d;e++){if(a=g[e].nTr)for(;f=a.firstChild;)a.removeChild(f);f=0;for(b=g[e].length;f<b;f++)if(n=
j=1,i[e][f]===k){a.appendChild(g[e][f].cell);for(i[e][f]=1;g[e+j]!==k&&g[e][f].cell==g[e+j][f].cell;)i[e+j][f]=1,j++;for(;g[e][f+n]!==k&&g[e][f].cell==g[e][f+n].cell;){for(c=0;c<j;c++)i[e+c][f+n]=1;n++}h(g[e][f].cell).attr("rowspan",j).attr("colspan",n)}}}}function M(a){var b=w(a,"aoPreDrawCallback","preDraw",[a]);if(-1!==h.inArray(!1,b))C(a,!1);else{var b=[],c=0,e=a.asStripeClasses,d=e.length,f=a.oLanguage,g=a.iInitDisplayStart,i="ssp"==B(a),j=a.aiDisplay;a.bDrawing=!0;g!==k&&-1!==g&&(a._iDisplayStart=
i?g:g>=a.fnRecordsDisplay()?0:g,a.iInitDisplayStart=-1);var g=a._iDisplayStart,n=a.fnDisplayEnd();if(a.bDeferLoading)a.bDeferLoading=!1,a.iDraw++,C(a,!1);else if(i){if(!a.bDestroying&&!kb(a))return}else a.iDraw++;if(0!==j.length){f=i?a.aoData.length:n;for(i=i?0:g;i<f;i++){var l=j[i],q=a.aoData[l];null===q.nTr&&Ja(a,l);l=q.nTr;if(0!==d){var o=e[c%d];q._sRowStripe!=o&&(h(l).removeClass(q._sRowStripe).addClass(o),q._sRowStripe=o)}w(a,"aoRowCallback",null,[l,q._aData,c,i]);b.push(l);c++}}else c=f.sZeroRecords,
1==a.iDraw&&"ajax"==B(a)?c=f.sLoadingRecords:f.sEmptyTable&&0===a.fnRecordsTotal()&&(c=f.sEmptyTable),b[0]=h("<tr/>",{"class":d?e[0]:""}).append(h("<td />",{valign:"top",colSpan:aa(a),"class":a.oClasses.sRowEmpty}).html(c))[0];w(a,"aoHeaderCallback","header",[h(a.nTHead).children("tr")[0],La(a),g,n,j]);w(a,"aoFooterCallback","footer",[h(a.nTFoot).children("tr")[0],La(a),g,n,j]);e=h(a.nTBody);e.children().detach();e.append(h(b));w(a,"aoDrawCallback","draw",[a]);a.bSorted=!1;a.bFiltered=!1;a.bDrawing=
!1}}function N(a,b){var c=a.oFeatures,e=c.bFilter;c.bSort&&lb(a);e?fa(a,a.oPreviousSearch):a.aiDisplay=a.aiDisplayMaster.slice();!0!==b&&(a._iDisplayStart=0);a._drawHold=b;M(a);a._drawHold=!1}function mb(a){var b=a.oClasses,c=h(a.nTable),c=h("<div/>").insertBefore(c),e=a.oFeatures,d=h("<div/>",{id:a.sTableId+"_wrapper","class":b.sWrapper+(a.nTFoot?"":" "+b.sNoFooter)});a.nHolding=c[0];a.nTableWrapper=d[0];a.nTableReinsertBefore=a.nTable.nextSibling;for(var f=a.sDom.split(""),g,i,j,n,l,q,o=0;o<f.length;o++){g=
null;i=f[o];if("<"==i){j=h("<div/>")[0];n=f[o+1];if("'"==n||'"'==n){l="";for(q=2;f[o+q]!=n;)l+=f[o+q],q++;"H"==l?l=b.sJUIHeader:"F"==l&&(l=b.sJUIFooter);-1!=l.indexOf(".")?(n=l.split("."),j.id=n[0].substr(1,n[0].length-1),j.className=n[1]):"#"==l.charAt(0)?j.id=l.substr(1,l.length-1):j.className=l;o+=q}d.append(j);d=h(j)}else if(">"==i)d=d.parent();else if("l"==i&&e.bPaginate&&e.bLengthChange)g=nb(a);else if("f"==i&&e.bFilter)g=ob(a);else if("r"==i&&e.bProcessing)g=pb(a);else if("t"==i)g=qb(a);else if("i"==
i&&e.bInfo)g=rb(a);else if("p"==i&&e.bPaginate)g=sb(a);else if(0!==m.ext.feature.length){j=m.ext.feature;q=0;for(n=j.length;q<n;q++)if(i==j[q].cFeature){g=j[q].fnInit(a);break}}g&&(j=a.aanFeatures,j[i]||(j[i]=[]),j[i].push(g),d.append(g))}c.replaceWith(d)}function da(a,b){var c=h(b).children("tr"),e,d,f,g,i,j,n,l,q,o;a.splice(0,a.length);f=0;for(j=c.length;f<j;f++)a.push([]);f=0;for(j=c.length;f<j;f++){e=c[f];for(d=e.firstChild;d;){if("TD"==d.nodeName.toUpperCase()||"TH"==d.nodeName.toUpperCase()){l=
1*d.getAttribute("colspan");q=1*d.getAttribute("rowspan");l=!l||0===l||1===l?1:l;q=!q||0===q||1===q?1:q;g=0;for(i=a[f];i[g];)g++;n=g;o=1===l?!0:!1;for(i=0;i<l;i++)for(g=0;g<q;g++)a[f+g][n+i]={cell:d,unique:o},a[f+g].nTr=e}d=d.nextSibling}}}function qa(a,b,c){var e=[];c||(c=a.aoHeader,b&&(c=[],da(c,b)));for(var b=0,d=c.length;b<d;b++)for(var f=0,g=c[b].length;f<g;f++)if(c[b][f].unique&&(!e[f]||!a.bSortCellsTop))e[f]=c[b][f].cell;return e}function ra(a,b,c){w(a,"aoServerParams","serverParams",[b]);
if(b&&h.isArray(b)){var e={},d=/(.*?)\[\]$/;h.each(b,function(a,b){var c=b.name.match(d);c?(c=c[0],e[c]||(e[c]=[]),e[c].push(b.value)):e[b.name]=b.value});b=e}var f,g=a.ajax,i=a.oInstance,j=function(b){w(a,null,"xhr",[a,b]);c(b)};if(h.isPlainObject(g)&&g.data){f=g.data;var n=h.isFunction(f)?f(b,a):f,b=h.isFunction(f)&&n?n:h.extend(!0,b,n);delete g.data}n={data:b,success:function(b){var c=b.error||b.sError;c&&a.oApi._fnLog(a,0,c);a.json=b;j(b)},dataType:"json",cache:!1,type:a.sServerMethod,error:function(b,
c){var f=a.oApi._fnLog;"parsererror"==c?f(a,0,"Invalid JSON response",1):4===b.readyState&&f(a,0,"Ajax error",7);C(a,!1)}};a.oAjaxData=b;w(a,null,"preXhr",[a,b]);a.fnServerData?a.fnServerData.call(i,a.sAjaxSource,h.map(b,function(a,b){return{name:b,value:a}}),j,a):a.sAjaxSource||"string"===typeof g?a.jqXHR=h.ajax(h.extend(n,{url:g||a.sAjaxSource})):h.isFunction(g)?a.jqXHR=g.call(i,b,j,a):(a.jqXHR=h.ajax(h.extend(n,g)),g.data=f)}function kb(a){return a.bAjaxDataGet?(a.iDraw++,C(a,!0),ra(a,tb(a),function(b){ub(a,
b)}),!1):!0}function tb(a){var b=a.aoColumns,c=b.length,e=a.oFeatures,d=a.oPreviousSearch,f=a.aoPreSearchCols,g,i=[],j,n,l,q=T(a);g=a._iDisplayStart;j=!1!==e.bPaginate?a._iDisplayLength:-1;var o=function(a,b){i.push({name:a,value:b})};o("sEcho",a.iDraw);o("iColumns",c);o("sColumns",D(b,"sName").join(","));o("iDisplayStart",g);o("iDisplayLength",j);var k={draw:a.iDraw,columns:[],order:[],start:g,length:j,search:{value:d.sSearch,regex:d.bRegex}};for(g=0;g<c;g++)n=b[g],l=f[g],j="function"==typeof n.mData?
"function":n.mData,k.columns.push({data:j,name:n.sName,searchable:n.bSearchable,orderable:n.bSortable,search:{value:l.sSearch,regex:l.bRegex}}),o("mDataProp_"+g,j),e.bFilter&&(o("sSearch_"+g,l.sSearch),o("bRegex_"+g,l.bRegex),o("bSearchable_"+g,n.bSearchable)),e.bSort&&o("bSortable_"+g,n.bSortable);e.bFilter&&(o("sSearch",d.sSearch),o("bRegex",d.bRegex));e.bSort&&(h.each(q,function(a,b){k.order.push({column:b.col,dir:b.dir});o("iSortCol_"+a,b.col);o("sSortDir_"+a,b.dir)}),o("iSortingCols",q.length));
b=m.ext.legacy.ajax;return null===b?a.sAjaxSource?i:k:b?i:k}function ub(a,b){var c=sa(a,b),e=b.sEcho!==k?b.sEcho:b.draw,d=b.iTotalRecords!==k?b.iTotalRecords:b.recordsTotal,f=b.iTotalDisplayRecords!==k?b.iTotalDisplayRecords:b.recordsFiltered;if(e){if(1*e<a.iDraw)return;a.iDraw=1*e}oa(a);a._iRecordsTotal=parseInt(d,10);a._iRecordsDisplay=parseInt(f,10);e=0;for(d=c.length;e<d;e++)J(a,c[e]);a.aiDisplay=a.aiDisplayMaster.slice();a.bAjaxDataGet=!1;M(a);a._bInitComplete||ta(a,b);a.bAjaxDataGet=!0;C(a,
!1)}function sa(a,b){var c=h.isPlainObject(a.ajax)&&a.ajax.dataSrc!==k?a.ajax.dataSrc:a.sAjaxDataProp;return"data"===c?b.aaData||b[c]:""!==c?W(c)(b):b}function ob(a){var b=a.oClasses,c=a.sTableId,e=a.oLanguage,d=a.oPreviousSearch,f=a.aanFeatures,g='<input type="search" class="'+b.sFilterInput+'"/>',i=e.sSearch,i=i.match(/_INPUT_/)?i.replace("_INPUT_",g):i+g,b=h("<div/>",{id:!f.f?c+"_filter":null,"class":b.sFilter}).append(h("<label/>").append(i)),f=function(){var b=!this.value?"":this.value;b!=d.sSearch&&
(fa(a,{sSearch:b,bRegex:d.bRegex,bSmart:d.bSmart,bCaseInsensitive:d.bCaseInsensitive}),a._iDisplayStart=0,M(a))},g=null!==a.searchDelay?a.searchDelay:"ssp"===B(a)?400:0,j=h("input",b).val(d.sSearch).attr("placeholder",e.sSearchPlaceholder).bind("keyup.DT search.DT input.DT paste.DT cut.DT",g?ua(f,g):f).bind("keypress.DT",function(a){if(13==a.keyCode)return!1}).attr("aria-controls",c);h(a.nTable).on("search.dt.DT",function(b,c){if(a===c)try{j[0]!==P.activeElement&&j.val(d.sSearch)}catch(f){}});return b[0]}
function fa(a,b,c){var e=a.oPreviousSearch,d=a.aoPreSearchCols,f=function(a){e.sSearch=a.sSearch;e.bRegex=a.bRegex;e.bSmart=a.bSmart;e.bCaseInsensitive=a.bCaseInsensitive};Ha(a);if("ssp"!=B(a)){vb(a,b.sSearch,c,b.bEscapeRegex!==k?!b.bEscapeRegex:b.bRegex,b.bSmart,b.bCaseInsensitive);f(b);for(b=0;b<d.length;b++)wb(a,d[b].sSearch,b,d[b].bEscapeRegex!==k?!d[b].bEscapeRegex:d[b].bRegex,d[b].bSmart,d[b].bCaseInsensitive);xb(a)}else f(b);a.bFiltered=!0;w(a,null,"search",[a])}function xb(a){for(var b=m.ext.search,
c=a.aiDisplay,e,d,f=0,g=b.length;f<g;f++){for(var i=[],j=0,h=c.length;j<h;j++)d=c[j],e=a.aoData[d],b[f](a,e._aFilterData,d,e._aData,j)&&i.push(d);c.length=0;c.push.apply(c,i)}}function wb(a,b,c,e,d,f){if(""!==b)for(var g=a.aiDisplay,e=Qa(b,e,d,f),d=g.length-1;0<=d;d--)b=a.aoData[g[d]]._aFilterData[c],e.test(b)||g.splice(d,1)}function vb(a,b,c,e,d,f){var e=Qa(b,e,d,f),d=a.oPreviousSearch.sSearch,f=a.aiDisplayMaster,g;0!==m.ext.search.length&&(c=!0);g=yb(a);if(0>=b.length)a.aiDisplay=f.slice();else{if(g||
c||d.length>b.length||0!==b.indexOf(d)||a.bSorted)a.aiDisplay=f.slice();b=a.aiDisplay;for(c=b.length-1;0<=c;c--)e.test(a.aoData[b[c]]._sFilterRow)||b.splice(c,1)}}function Qa(a,b,c,e){a=b?a:va(a);c&&(a="^(?=.*?"+h.map(a.match(/"[^"]+"|[^ ]+/g)||"",function(a){if('"'===a.charAt(0))var b=a.match(/^"(.*)"$/),a=b?b[1]:a;return a.replace('"',"")}).join(")(?=.*?")+").*$");return RegExp(a,e?"i":"")}function va(a){return a.replace(Yb,"\\$1")}function yb(a){var b=a.aoColumns,c,e,d,f,g,i,j,h,l=m.ext.type.search;
c=!1;e=0;for(f=a.aoData.length;e<f;e++)if(h=a.aoData[e],!h._aFilterData){i=[];d=0;for(g=b.length;d<g;d++)c=b[d],c.bSearchable?(j=y(a,e,d,"filter"),l[c.sType]&&(j=l[c.sType](j)),null===j&&(j=""),"string"!==typeof j&&j.toString&&(j=j.toString())):j="",j.indexOf&&-1!==j.indexOf("&")&&(wa.innerHTML=j,j=Zb?wa.textContent:wa.innerText),j.replace&&(j=j.replace(/[\r\n]/g,"")),i.push(j);h._aFilterData=i;h._sFilterRow=i.join("  ");c=!0}return c}function zb(a){return{search:a.sSearch,smart:a.bSmart,regex:a.bRegex,
caseInsensitive:a.bCaseInsensitive}}function Ab(a){return{sSearch:a.search,bSmart:a.smart,bRegex:a.regex,bCaseInsensitive:a.caseInsensitive}}function rb(a){var b=a.sTableId,c=a.aanFeatures.i,e=h("<div/>",{"class":a.oClasses.sInfo,id:!c?b+"_info":null});c||(a.aoDrawCallback.push({fn:Bb,sName:"information"}),e.attr("role","status").attr("aria-live","polite"),h(a.nTable).attr("aria-describedby",b+"_info"));return e[0]}function Bb(a){var b=a.aanFeatures.i;if(0!==b.length){var c=a.oLanguage,e=a._iDisplayStart+
1,d=a.fnDisplayEnd(),f=a.fnRecordsTotal(),g=a.fnRecordsDisplay(),i=g?c.sInfo:c.sInfoEmpty;g!==f&&(i+=" "+c.sInfoFiltered);i+=c.sInfoPostFix;i=Cb(a,i);c=c.fnInfoCallback;null!==c&&(i=c.call(a.oInstance,a,e,d,f,g,i));h(b).html(i)}}function Cb(a,b){var c=a.fnFormatNumber,e=a._iDisplayStart+1,d=a._iDisplayLength,f=a.fnRecordsDisplay(),g=-1===d;return b.replace(/_START_/g,c.call(a,e)).replace(/_END_/g,c.call(a,a.fnDisplayEnd())).replace(/_MAX_/g,c.call(a,a.fnRecordsTotal())).replace(/_TOTAL_/g,c.call(a,
f)).replace(/_PAGE_/g,c.call(a,g?1:Math.ceil(e/d))).replace(/_PAGES_/g,c.call(a,g?1:Math.ceil(f/d)))}function ga(a){var b,c,e=a.iInitDisplayStart,d=a.aoColumns,f;c=a.oFeatures;if(a.bInitialised){mb(a);jb(a);ea(a,a.aoHeader);ea(a,a.aoFooter);C(a,!0);c.bAutoWidth&&Ga(a);b=0;for(c=d.length;b<c;b++)f=d[b],f.sWidth&&(f.nTh.style.width=s(f.sWidth));N(a);d=B(a);"ssp"!=d&&("ajax"==d?ra(a,[],function(c){var f=sa(a,c);for(b=0;b<f.length;b++)J(a,f[b]);a.iInitDisplayStart=e;N(a);C(a,!1);ta(a,c)},a):(C(a,!1),
ta(a)))}else setTimeout(function(){ga(a)},200)}function ta(a,b){a._bInitComplete=!0;b&&X(a);w(a,"aoInitComplete","init",[a,b])}function Ra(a,b){var c=parseInt(b,10);a._iDisplayLength=c;Sa(a);w(a,null,"length",[a,c])}function nb(a){for(var b=a.oClasses,c=a.sTableId,e=a.aLengthMenu,d=h.isArray(e[0]),f=d?e[0]:e,e=d?e[1]:e,d=h("<select/>",{name:c+"_length","aria-controls":c,"class":b.sLengthSelect}),g=0,i=f.length;g<i;g++)d[0][g]=new Option(e[g],f[g]);var j=h("<div><label/></div>").addClass(b.sLength);
a.aanFeatures.l||(j[0].id=c+"_length");j.children().append(a.oLanguage.sLengthMenu.replace("_MENU_",d[0].outerHTML));h("select",j).val(a._iDisplayLength).bind("change.DT",function(){Ra(a,h(this).val());M(a)});h(a.nTable).bind("length.dt.DT",function(b,c,f){a===c&&h("select",j).val(f)});return j[0]}function sb(a){var b=a.sPaginationType,c=m.ext.pager[b],e="function"===typeof c,d=function(a){M(a)},b=h("<div/>").addClass(a.oClasses.sPaging+b)[0],f=a.aanFeatures;e||c.fnInit(a,b,d);f.p||(b.id=a.sTableId+
"_paginate",a.aoDrawCallback.push({fn:function(a){if(e){var b=a._iDisplayStart,h=a._iDisplayLength,n=a.fnRecordsDisplay(),l=-1===h,b=l?0:Math.ceil(b/h),h=l?1:Math.ceil(n/h),n=c(b,h),q,l=0;for(q=f.p.length;l<q;l++)Pa(a,"pageButton")(a,f.p[l],l,n,b,h)}else c.fnUpdate(a,d)},sName:"pagination"}));return b}function Ta(a,b,c){var e=a._iDisplayStart,d=a._iDisplayLength,f=a.fnRecordsDisplay();0===f||-1===d?e=0:"number"===typeof b?(e=b*d,e>f&&(e=0)):"first"==b?e=0:"previous"==b?(e=0<=d?e-d:0,0>e&&(e=0)):"next"==
b?e+d<f&&(e+=d):"last"==b?e=Math.floor((f-1)/d)*d:R(a,0,"Unknown paging action: "+b,5);b=a._iDisplayStart!==e;a._iDisplayStart=e;b&&(w(a,null,"page",[a]),c&&M(a));return b}function pb(a){return h("<div/>",{id:!a.aanFeatures.r?a.sTableId+"_processing":null,"class":a.oClasses.sProcessing}).html(a.oLanguage.sProcessing).insertBefore(a.nTable)[0]}function C(a,b){a.oFeatures.bProcessing&&h(a.aanFeatures.r).css("display",b?"block":"none");w(a,null,"processing",[a,b])}function qb(a){var b=h(a.nTable);b.attr("role",
"grid");var c=a.oScroll;if(""===c.sX&&""===c.sY)return a.nTable;var e=c.sX,d=c.sY,f=a.oClasses,g=b.children("caption"),i=g.length?g[0]._captionSide:null,j=h(b[0].cloneNode(!1)),n=h(b[0].cloneNode(!1)),l=b.children("tfoot");c.sX&&"100%"===b.attr("width")&&b.removeAttr("width");l.length||(l=null);c=h("<div/>",{"class":f.sScrollWrapper}).append(h("<div/>",{"class":f.sScrollHead}).css({overflow:"hidden",position:"relative",border:0,width:e?!e?null:s(e):"100%"}).append(h("<div/>",{"class":f.sScrollHeadInner}).css({"box-sizing":"content-box",
width:c.sXInner||"100%"}).append(j.removeAttr("id").css("margin-left",0).append("top"===i?g:null).append(b.children("thead"))))).append(h("<div/>",{"class":f.sScrollBody}).css({overflow:"auto",height:!d?null:s(d),width:!e?null:s(e)}).append(b));l&&c.append(h("<div/>",{"class":f.sScrollFoot}).css({overflow:"hidden",border:0,width:e?!e?null:s(e):"100%"}).append(h("<div/>",{"class":f.sScrollFootInner}).append(n.removeAttr("id").css("margin-left",0).append("bottom"===i?g:null).append(b.children("tfoot")))));
var b=c.children(),q=b[0],f=b[1],o=l?b[2]:null;if(e)h(f).on("scroll.DT",function(){var a=this.scrollLeft;q.scrollLeft=a;l&&(o.scrollLeft=a)});a.nScrollHead=q;a.nScrollBody=f;a.nScrollFoot=o;a.aoDrawCallback.push({fn:Y,sName:"scrolling"});return c[0]}function Y(a){var b=a.oScroll,c=b.sX,e=b.sXInner,d=b.sY,f=b.iBarWidth,g=h(a.nScrollHead),i=g[0].style,j=g.children("div"),n=j[0].style,l=j.children("table"),j=a.nScrollBody,q=h(j),o=j.style,k=h(a.nScrollFoot).children("div"),p=k.children("table"),m=h(a.nTHead),
r=h(a.nTable),t=r[0],u=t.style,K=a.nTFoot?h(a.nTFoot):null,ha=a.oBrowser,w=ha.bScrollOversize,x,v,y,L,z,A=[],B=[],C=[],D,E=function(a){a=a.style;a.paddingTop="0";a.paddingBottom="0";a.borderTopWidth="0";a.borderBottomWidth="0";a.height=0};r.children("thead, tfoot").remove();z=m.clone().prependTo(r);x=m.find("tr");y=z.find("tr");z.find("th, td").removeAttr("tabindex");K&&(L=K.clone().prependTo(r),v=K.find("tr"),L=L.find("tr"));c||(o.width="100%",g[0].style.width="100%");h.each(qa(a,z),function(b,c){D=
la(a,b);c.style.width=a.aoColumns[D].sWidth});K&&G(function(a){a.style.width=""},L);b.bCollapse&&""!==d&&(o.height=q[0].offsetHeight+m[0].offsetHeight+"px");g=r.outerWidth();if(""===c){if(u.width="100%",w&&(r.find("tbody").height()>j.offsetHeight||"scroll"==q.css("overflow-y")))u.width=s(r.outerWidth()-f)}else""!==e?u.width=s(e):g==q.width()&&q.height()<r.height()?(u.width=s(g-f),r.outerWidth()>g-f&&(u.width=s(g))):u.width=s(g);g=r.outerWidth();G(E,y);G(function(a){C.push(a.innerHTML);A.push(s(h(a).css("width")))},
y);G(function(a,b){a.style.width=A[b]},x);h(y).height(0);K&&(G(E,L),G(function(a){B.push(s(h(a).css("width")))},L),G(function(a,b){a.style.width=B[b]},v),h(L).height(0));G(function(a,b){a.innerHTML='<div class="dataTables_sizing" style="height:0;overflow:hidden;">'+C[b]+"</div>";a.style.width=A[b]},y);K&&G(function(a,b){a.innerHTML="";a.style.width=B[b]},L);if(r.outerWidth()<g){v=j.scrollHeight>j.offsetHeight||"scroll"==q.css("overflow-y")?g+f:g;if(w&&(j.scrollHeight>j.offsetHeight||"scroll"==q.css("overflow-y")))u.width=
s(v-f);(""===c||""!==e)&&R(a,1,"Possible column misalignment",6)}else v="100%";o.width=s(v);i.width=s(v);K&&(a.nScrollFoot.style.width=s(v));!d&&w&&(o.height=s(t.offsetHeight+f));d&&b.bCollapse&&(o.height=s(d),b=c&&t.offsetWidth>j.offsetWidth?f:0,t.offsetHeight<j.offsetHeight&&(o.height=s(t.offsetHeight+b)));b=r.outerWidth();l[0].style.width=s(b);n.width=s(b);l=r.height()>j.clientHeight||"scroll"==q.css("overflow-y");ha="padding"+(ha.bScrollbarLeft?"Left":"Right");n[ha]=l?f+"px":"0px";K&&(p[0].style.width=
s(b),k[0].style.width=s(b),k[0].style[ha]=l?f+"px":"0px");q.scroll();if((a.bSorted||a.bFiltered)&&!a._drawHold)j.scrollTop=0}function G(a,b,c){for(var e=0,d=0,f=b.length,g,i;d<f;){g=b[d].firstChild;for(i=c?c[d].firstChild:null;g;)1===g.nodeType&&(c?a(g,i,e):a(g,e),e++),g=g.nextSibling,i=c?i.nextSibling:null;d++}}function Ga(a){var b=a.nTable,c=a.aoColumns,e=a.oScroll,d=e.sY,f=e.sX,g=e.sXInner,i=c.length,e=Z(a,"bVisible"),j=h("th",a.nTHead),n=b.getAttribute("width"),l=b.parentNode,k=!1,o,m;(o=b.style.width)&&
-1!==o.indexOf("%")&&(n=o);for(o=0;o<e.length;o++)m=c[e[o]],null!==m.sWidth&&(m.sWidth=Db(m.sWidthOrig,l),k=!0);if(!k&&!f&&!d&&i==aa(a)&&i==j.length)for(o=0;o<i;o++)c[o].sWidth=s(j.eq(o).width());else{i=h(b).clone().empty().css("visibility","hidden").removeAttr("id").append(h(a.nTHead).clone(!1)).append(h(a.nTFoot).clone(!1)).append(h("<tbody><tr/></tbody>"));i.find("tfoot th, tfoot td").css("width","");var p=i.find("tbody tr"),j=qa(a,i.find("thead")[0]);for(o=0;o<e.length;o++)m=c[e[o]],j[o].style.width=
null!==m.sWidthOrig&&""!==m.sWidthOrig?s(m.sWidthOrig):"";if(a.aoData.length)for(o=0;o<e.length;o++)k=e[o],m=c[k],h(Eb(a,k)).clone(!1).append(m.sContentPadding).appendTo(p);i.appendTo(l);f&&g?i.width(g):f?(i.css("width","auto"),i.width()<l.offsetWidth&&i.width(l.offsetWidth)):d?i.width(l.offsetWidth):n&&i.width(n);Fb(a,i[0]);if(f){for(o=g=0;o<e.length;o++)m=c[e[o]],d=h(j[o]).outerWidth(),g+=null===m.sWidthOrig?d:parseInt(m.sWidth,10)+d-h(j[o]).width();i.width(s(g));b.style.width=s(g)}for(o=0;o<e.length;o++)if(m=
c[e[o]],d=h(j[o]).width())m.sWidth=s(d);b.style.width=s(i.css("width"));i.remove()}n&&(b.style.width=s(n));if((n||f)&&!a._reszEvt)h(Ea).bind("resize.DT-"+a.sInstance,ua(function(){X(a)})),a._reszEvt=!0}function ua(a,b){var c=b!==k?b:200,e,d;return function(){var b=this,g=+new Date,i=arguments;e&&g<e+c?(clearTimeout(d),d=setTimeout(function(){e=k;a.apply(b,i)},c)):(e=g,a.apply(b,i))}}function Db(a,b){if(!a)return 0;var c=h("<div/>").css("width",s(a)).appendTo(b||P.body),e=c[0].offsetWidth;c.remove();
return e}function Fb(a,b){var c=a.oScroll;if(c.sX||c.sY)c=!c.sX?c.iBarWidth:0,b.style.width=s(h(b).outerWidth()-c)}function Eb(a,b){var c=Gb(a,b);if(0>c)return null;var e=a.aoData[c];return!e.nTr?h("<td/>").html(y(a,c,b,"display"))[0]:e.anCells[b]}function Gb(a,b){for(var c,e=-1,d=-1,f=0,g=a.aoData.length;f<g;f++)c=y(a,f,b,"display")+"",c=c.replace($b,""),c.length>e&&(e=c.length,d=f);return d}function s(a){return null===a?"0px":"number"==typeof a?0>a?"0px":a+"px":a.match(/\d$/)?a+"px":a}function Hb(){var a=
m.__scrollbarWidth;if(a===k){var b=h("<p/>").css({position:"absolute",top:0,left:0,width:"100%",height:150,padding:0,overflow:"scroll",visibility:"hidden"}).appendTo("body"),a=b[0].offsetWidth-b[0].clientWidth;m.__scrollbarWidth=a;b.remove()}return a}function T(a){var b,c,e=[],d=a.aoColumns,f,g,i,j;b=a.aaSortingFixed;c=h.isPlainObject(b);var n=[];f=function(a){a.length&&!h.isArray(a[0])?n.push(a):n.push.apply(n,a)};h.isArray(b)&&f(b);c&&b.pre&&f(b.pre);f(a.aaSorting);c&&b.post&&f(b.post);for(a=0;a<
n.length;a++){j=n[a][0];f=d[j].aDataSort;b=0;for(c=f.length;b<c;b++)g=f[b],i=d[g].sType||"string",n[a]._idx===k&&(n[a]._idx=h.inArray(n[a][1],d[g].asSorting)),e.push({src:j,col:g,dir:n[a][1],index:n[a]._idx,type:i,formatter:m.ext.type.order[i+"-pre"]})}return e}function lb(a){var b,c,e=[],d=m.ext.type.order,f=a.aoData,g=0,i,h=a.aiDisplayMaster,n;Ha(a);n=T(a);b=0;for(c=n.length;b<c;b++)i=n[b],i.formatter&&g++,Ib(a,i.col);if("ssp"!=B(a)&&0!==n.length){b=0;for(c=h.length;b<c;b++)e[h[b]]=b;g===n.length?
h.sort(function(a,b){var c,d,g,h,i=n.length,j=f[a]._aSortData,k=f[b]._aSortData;for(g=0;g<i;g++)if(h=n[g],c=j[h.col],d=k[h.col],c=c<d?-1:c>d?1:0,0!==c)return"asc"===h.dir?c:-c;c=e[a];d=e[b];return c<d?-1:c>d?1:0}):h.sort(function(a,b){var c,g,h,i,j=n.length,k=f[a]._aSortData,m=f[b]._aSortData;for(h=0;h<j;h++)if(i=n[h],c=k[i.col],g=m[i.col],i=d[i.type+"-"+i.dir]||d["string-"+i.dir],c=i(c,g),0!==c)return c;c=e[a];g=e[b];return c<g?-1:c>g?1:0})}a.bSorted=!0}function Jb(a){for(var b,c,e=a.aoColumns,d=
T(a),a=a.oLanguage.oAria,f=0,g=e.length;f<g;f++){c=e[f];var i=c.asSorting;b=c.sTitle.replace(/<.*?>/g,"");var h=c.nTh;h.removeAttribute("aria-sort");c.bSortable&&(0<d.length&&d[0].col==f?(h.setAttribute("aria-sort","asc"==d[0].dir?"ascending":"descending"),c=i[d[0].index+1]||i[0]):c=i[0],b+="asc"===c?a.sSortAscending:a.sSortDescending);h.setAttribute("aria-label",b)}}function Ua(a,b,c,e){var d=a.aaSorting,f=a.aoColumns[b].asSorting,g=function(a,b){var c=a._idx;c===k&&(c=h.inArray(a[1],f));return c+
1<f.length?c+1:b?null:0};"number"===typeof d[0]&&(d=a.aaSorting=[d]);c&&a.oFeatures.bSortMulti?(c=h.inArray(b,D(d,"0")),-1!==c?(b=g(d[c],!0),null===b&&1===d.length&&(b=0),null===b?d.splice(c,1):(d[c][1]=f[b],d[c]._idx=b)):(d.push([b,f[0],0]),d[d.length-1]._idx=0)):d.length&&d[0][0]==b?(b=g(d[0]),d.length=1,d[0][1]=f[b],d[0]._idx=b):(d.length=0,d.push([b,f[0]]),d[0]._idx=0);N(a);"function"==typeof e&&e(a)}function Oa(a,b,c,e){var d=a.aoColumns[c];Va(b,{},function(b){!1!==d.bSortable&&(a.oFeatures.bProcessing?
(C(a,!0),setTimeout(function(){Ua(a,c,b.shiftKey,e);"ssp"!==B(a)&&C(a,!1)},0)):Ua(a,c,b.shiftKey,e))})}function xa(a){var b=a.aLastSort,c=a.oClasses.sSortColumn,e=T(a),d=a.oFeatures,f,g;if(d.bSort&&d.bSortClasses){d=0;for(f=b.length;d<f;d++)g=b[d].src,h(D(a.aoData,"anCells",g)).removeClass(c+(2>d?d+1:3));d=0;for(f=e.length;d<f;d++)g=e[d].src,h(D(a.aoData,"anCells",g)).addClass(c+(2>d?d+1:3))}a.aLastSort=e}function Ib(a,b){var c=a.aoColumns[b],e=m.ext.order[c.sSortDataType],d;e&&(d=e.call(a.oInstance,
a,b,$(a,b)));for(var f,g=m.ext.type.order[c.sType+"-pre"],h=0,j=a.aoData.length;h<j;h++)if(c=a.aoData[h],c._aSortData||(c._aSortData=[]),!c._aSortData[b]||e)f=e?d[h]:y(a,h,b,"sort"),c._aSortData[b]=g?g(f):f}function ya(a){if(a.oFeatures.bStateSave&&!a.bDestroying){var b={time:+new Date,start:a._iDisplayStart,length:a._iDisplayLength,order:h.extend(!0,[],a.aaSorting),search:zb(a.oPreviousSearch),columns:h.map(a.aoColumns,function(b,e){return{visible:b.bVisible,search:zb(a.aoPreSearchCols[e])}})};w(a,
"aoStateSaveParams","stateSaveParams",[a,b]);a.oSavedState=b;a.fnStateSaveCallback.call(a.oInstance,a,b)}}function Kb(a){var b,c,e=a.aoColumns;if(a.oFeatures.bStateSave){var d=a.fnStateLoadCallback.call(a.oInstance,a);if(d&&d.time&&(b=w(a,"aoStateLoadParams","stateLoadParams",[a,d]),-1===h.inArray(!1,b)&&(b=a.iStateDuration,!(0<b&&d.time<+new Date-1E3*b)&&e.length===d.columns.length))){a.oLoadedState=h.extend(!0,{},d);d.start!==k&&(a._iDisplayStart=d.start,a.iInitDisplayStart=d.start);d.length!==
k&&(a._iDisplayLength=d.length);d.order!==k&&(a.aaSorting=[],h.each(d.order,function(b,c){a.aaSorting.push(c[0]>=e.length?[0,c[1]]:c)}));d.search!==k&&h.extend(a.oPreviousSearch,Ab(d.search));b=0;for(c=d.columns.length;b<c;b++){var f=d.columns[b];f.visible!==k&&(e[b].bVisible=f.visible);f.search!==k&&h.extend(a.aoPreSearchCols[b],Ab(f.search))}w(a,"aoStateLoaded","stateLoaded",[a,d])}}}function za(a){var b=m.settings,a=h.inArray(a,D(b,"nTable"));return-1!==a?b[a]:null}function R(a,b,c,e){c="DataTables warning: "+
(null!==a?"table id="+a.sTableId+" - ":"")+c;e&&(c+=". For more information about this error, please see http://datatables.net/tn/"+e);if(b)Ea.console&&console.log&&console.log(c);else if(b=m.ext,b=b.sErrMode||b.errMode,w(a,null,"error",[a,e,c]),"alert"==b)alert(c);else{if("throw"==b)throw Error(c);"function"==typeof b&&b(a,e,c)}}function E(a,b,c,e){h.isArray(c)?h.each(c,function(c,f){h.isArray(f)?E(a,b,f[0],f[1]):E(a,b,f)}):(e===k&&(e=c),b[c]!==k&&(a[e]=b[c]))}function Lb(a,b,c){var e,d;for(d in b)b.hasOwnProperty(d)&&
(e=b[d],h.isPlainObject(e)?(h.isPlainObject(a[d])||(a[d]={}),h.extend(!0,a[d],e)):a[d]=c&&"data"!==d&&"aaData"!==d&&h.isArray(e)?e.slice():e);return a}function Va(a,b,c){h(a).bind("click.DT",b,function(b){a.blur();c(b)}).bind("keypress.DT",b,function(a){13===a.which&&(a.preventDefault(),c(a))}).bind("selectstart.DT",function(){return!1})}function z(a,b,c,e){c&&a[b].push({fn:c,sName:e})}function w(a,b,c,e){var d=[];b&&(d=h.map(a[b].slice().reverse(),function(b){return b.fn.apply(a.oInstance,e)}));
null!==c&&h(a.nTable).trigger(c+".dt",e);return d}function Sa(a){var b=a._iDisplayStart,c=a.fnDisplayEnd(),e=a._iDisplayLength;b>=c&&(b=c-e);b-=b%e;if(-1===e||0>b)b=0;a._iDisplayStart=b}function Pa(a,b){var c=a.renderer,e=m.ext.renderer[b];return h.isPlainObject(c)&&c[b]?e[c[b]]||e._:"string"===typeof c?e[c]||e._:e._}function B(a){return a.oFeatures.bServerSide?"ssp":a.ajax||a.sAjaxSource?"ajax":"dom"}function Wa(a,b){var c=[],c=Mb.numbers_length,e=Math.floor(c/2);b<=c?c=U(0,b):a<=e?(c=U(0,c-2),c.push("ellipsis"),
c.push(b-1)):(a>=b-1-e?c=U(b-(c-2),b):(c=U(a-e+2,a+e-1),c.push("ellipsis"),c.push(b-1)),c.splice(0,0,"ellipsis"),c.splice(0,0,0));c.DT_el="span";return c}function db(a){h.each({num:function(b){return Aa(b,a)},"num-fmt":function(b){return Aa(b,a,Xa)},"html-num":function(b){return Aa(b,a,Ba)},"html-num-fmt":function(b){return Aa(b,a,Ba,Xa)}},function(b,c){x.type.order[b+a+"-pre"]=c;b.match(/^html\-/)&&(x.type.search[b+a]=x.type.search.html)})}function Nb(a){return function(){var b=[za(this[m.ext.iApiIndex])].concat(Array.prototype.slice.call(arguments));
return m.ext.internal[a].apply(this,b)}}var m,x,t,r,u,Ya={},Ob=/[\r\n]/g,Ba=/<.*?>/g,ac=/^[\w\+\-]/,bc=/[\w\+\-]$/,Yb=RegExp("(\\/|\\.|\\*|\\+|\\?|\\||\\(|\\)|\\[|\\]|\\{|\\}|\\\\|\\$|\\^|\\-)","g"),Xa=/[',$\u00a3\u20ac\u00a5%\u2009\u202F\u20BD\u20a9\u20BArfk]/gi,I=function(a){return!a||!0===a||"-"===a?!0:!1},Pb=function(a){var b=parseInt(a,10);return!isNaN(b)&&isFinite(a)?b:null},Qb=function(a,b){Ya[b]||(Ya[b]=RegExp(va(b),"g"));return"string"===typeof a&&"."!==b?a.replace(/\./g,"").replace(Ya[b],
"."):a},Za=function(a,b,c){var e="string"===typeof a;if(I(a))return!0;b&&e&&(a=Qb(a,b));c&&e&&(a=a.replace(Xa,""));return!isNaN(parseFloat(a))&&isFinite(a)},Rb=function(a,b,c){return I(a)?!0:!(I(a)||"string"===typeof a)?null:Za(a.replace(Ba,""),b,c)?!0:null},D=function(a,b,c){var e=[],d=0,f=a.length;if(c!==k)for(;d<f;d++)a[d]&&a[d][b]&&e.push(a[d][b][c]);else for(;d<f;d++)a[d]&&e.push(a[d][b]);return e},ia=function(a,b,c,e){var d=[],f=0,g=b.length;if(e!==k)for(;f<g;f++)a[b[f]][c]&&d.push(a[b[f]][c][e]);
else for(;f<g;f++)d.push(a[b[f]][c]);return d},U=function(a,b){var c=[],e;b===k?(b=0,e=a):(e=b,b=a);for(var d=b;d<e;d++)c.push(d);return c},Sb=function(a){for(var b=[],c=0,e=a.length;c<e;c++)a[c]&&b.push(a[c]);return b},Na=function(a){var b=[],c,e,d=a.length,f,g=0;e=0;a:for(;e<d;e++){c=a[e];for(f=0;f<g;f++)if(b[f]===c)continue a;b.push(c);g++}return b},A=function(a,b,c){a[b]!==k&&(a[c]=a[b])},ba=/\[.*?\]$/,S=/\(\)$/,wa=h("<div>")[0],Zb=wa.textContent!==k,$b=/<.*?>/g;m=function(a){this.$=function(a,
b){return this.api(!0).$(a,b)};this._=function(a,b){return this.api(!0).rows(a,b).data()};this.api=function(a){return a?new t(za(this[x.iApiIndex])):new t(this)};this.fnAddData=function(a,b){var c=this.api(!0),e=h.isArray(a)&&(h.isArray(a[0])||h.isPlainObject(a[0]))?c.rows.add(a):c.row.add(a);(b===k||b)&&c.draw();return e.flatten().toArray()};this.fnAdjustColumnSizing=function(a){var b=this.api(!0).columns.adjust(),c=b.settings()[0],e=c.oScroll;a===k||a?b.draw(!1):(""!==e.sX||""!==e.sY)&&Y(c)};this.fnClearTable=
function(a){var b=this.api(!0).clear();(a===k||a)&&b.draw()};this.fnClose=function(a){this.api(!0).row(a).child.hide()};this.fnDeleteRow=function(a,b,c){var e=this.api(!0),a=e.rows(a),d=a.settings()[0],h=d.aoData[a[0][0]];a.remove();b&&b.call(this,d,h);(c===k||c)&&e.draw();return h};this.fnDestroy=function(a){this.api(!0).destroy(a)};this.fnDraw=function(a){this.api(!0).draw(a)};this.fnFilter=function(a,b,c,e,d,h){d=this.api(!0);null===b||b===k?d.search(a,c,e,h):d.column(b).search(a,c,e,h);d.draw()};
this.fnGetData=function(a,b){var c=this.api(!0);if(a!==k){var e=a.nodeName?a.nodeName.toLowerCase():"";return b!==k||"td"==e||"th"==e?c.cell(a,b).data():c.row(a).data()||null}return c.data().toArray()};this.fnGetNodes=function(a){var b=this.api(!0);return a!==k?b.row(a).node():b.rows().nodes().flatten().toArray()};this.fnGetPosition=function(a){var b=this.api(!0),c=a.nodeName.toUpperCase();return"TR"==c?b.row(a).index():"TD"==c||"TH"==c?(a=b.cell(a).index(),[a.row,a.columnVisible,a.column]):null};
this.fnIsOpen=function(a){return this.api(!0).row(a).child.isShown()};this.fnOpen=function(a,b,c){return this.api(!0).row(a).child(b,c).show().child()[0]};this.fnPageChange=function(a,b){var c=this.api(!0).page(a);(b===k||b)&&c.draw(!1)};this.fnSetColumnVis=function(a,b,c){a=this.api(!0).column(a).visible(b);(c===k||c)&&a.columns.adjust().draw()};this.fnSettings=function(){return za(this[x.iApiIndex])};this.fnSort=function(a){this.api(!0).order(a).draw()};this.fnSortListener=function(a,b,c){this.api(!0).order.listener(a,
b,c)};this.fnUpdate=function(a,b,c,e,d){var h=this.api(!0);c===k||null===c?h.row(b).data(a):h.cell(b,c).data(a);(d===k||d)&&h.columns.adjust();(e===k||e)&&h.draw();return 0};this.fnVersionCheck=x.fnVersionCheck;var b=this,c=a===k,e=this.length;c&&(a={});this.oApi=this.internal=x.internal;for(var d in m.ext.internal)d&&(this[d]=Nb(d));this.each(function(){var d={},d=1<e?Lb(d,a,!0):a,g=0,i,j=this.getAttribute("id"),n=!1,l=m.defaults,q=h(this);if("table"!=this.nodeName.toLowerCase())R(null,0,"Non-table node initialisation ("+
this.nodeName+")",2);else{eb(l);fb(l.column);H(l,l,!0);H(l.column,l.column,!0);H(l,h.extend(d,q.data()));var o=m.settings,g=0;for(i=o.length;g<i;g++){var r=o[g];if(r.nTable==this||r.nTHead.parentNode==this||r.nTFoot&&r.nTFoot.parentNode==this){g=d.bRetrieve!==k?d.bRetrieve:l.bRetrieve;if(c||g)return r.oInstance;if(d.bDestroy!==k?d.bDestroy:l.bDestroy){r.oInstance.fnDestroy();break}else{R(r,0,"Cannot reinitialise DataTable",3);return}}if(r.sTableId==this.id){o.splice(g,1);break}}if(null===j||""===
j)this.id=j="DataTables_Table_"+m.ext._unique++;var p=h.extend(!0,{},m.models.oSettings,{sDestroyWidth:q[0].style.width,sInstance:j,sTableId:j});p.nTable=this;p.oApi=b.internal;p.oInit=d;o.push(p);p.oInstance=1===b.length?b:q.dataTable();eb(d);d.oLanguage&&O(d.oLanguage);d.aLengthMenu&&!d.iDisplayLength&&(d.iDisplayLength=h.isArray(d.aLengthMenu[0])?d.aLengthMenu[0][0]:d.aLengthMenu[0]);d=Lb(h.extend(!0,{},l),d);E(p.oFeatures,d,"bPaginate bLengthChange bFilter bSort bSortMulti bInfo bProcessing bAutoWidth bSortClasses bServerSide bDeferRender".split(" "));
E(p,d,["asStripeClasses","ajax","fnServerData","fnFormatNumber","sServerMethod","aaSorting","aaSortingFixed","aLengthMenu","sPaginationType","sAjaxSource","sAjaxDataProp","iStateDuration","sDom","bSortCellsTop","iTabIndex","fnStateLoadCallback","fnStateSaveCallback","renderer","searchDelay",["iCookieDuration","iStateDuration"],["oSearch","oPreviousSearch"],["aoSearchCols","aoPreSearchCols"],["iDisplayLength","_iDisplayLength"],["bJQueryUI","bJUI"]]);E(p.oScroll,d,[["sScrollX","sX"],["sScrollXInner",
"sXInner"],["sScrollY","sY"],["bScrollCollapse","bCollapse"]]);E(p.oLanguage,d,"fnInfoCallback");z(p,"aoDrawCallback",d.fnDrawCallback,"user");z(p,"aoServerParams",d.fnServerParams,"user");z(p,"aoStateSaveParams",d.fnStateSaveParams,"user");z(p,"aoStateLoadParams",d.fnStateLoadParams,"user");z(p,"aoStateLoaded",d.fnStateLoaded,"user");z(p,"aoRowCallback",d.fnRowCallback,"user");z(p,"aoRowCreatedCallback",d.fnCreatedRow,"user");z(p,"aoHeaderCallback",d.fnHeaderCallback,"user");z(p,"aoFooterCallback",
d.fnFooterCallback,"user");z(p,"aoInitComplete",d.fnInitComplete,"user");z(p,"aoPreDrawCallback",d.fnPreDrawCallback,"user");j=p.oClasses;d.bJQueryUI?(h.extend(j,m.ext.oJUIClasses,d.oClasses),d.sDom===l.sDom&&"lfrtip"===l.sDom&&(p.sDom='<"H"lfr>t<"F"ip>'),p.renderer)?h.isPlainObject(p.renderer)&&!p.renderer.header&&(p.renderer.header="jqueryui"):p.renderer="jqueryui":h.extend(j,m.ext.classes,d.oClasses);q.addClass(j.sTable);if(""!==p.oScroll.sX||""!==p.oScroll.sY)p.oScroll.iBarWidth=Hb();!0===p.oScroll.sX&&
(p.oScroll.sX="100%");p.iInitDisplayStart===k&&(p.iInitDisplayStart=d.iDisplayStart,p._iDisplayStart=d.iDisplayStart);null!==d.iDeferLoading&&(p.bDeferLoading=!0,g=h.isArray(d.iDeferLoading),p._iRecordsDisplay=g?d.iDeferLoading[0]:d.iDeferLoading,p._iRecordsTotal=g?d.iDeferLoading[1]:d.iDeferLoading);var t=p.oLanguage;h.extend(!0,t,d.oLanguage);""!==t.sUrl&&(h.ajax({dataType:"json",url:t.sUrl,success:function(a){O(a);H(l.oLanguage,a);h.extend(true,t,a);ga(p)},error:function(){ga(p)}}),n=!0);null===
d.asStripeClasses&&(p.asStripeClasses=[j.sStripeOdd,j.sStripeEven]);var g=p.asStripeClasses,s=q.children("tbody").find("tr").eq(0);-1!==h.inArray(!0,h.map(g,function(a){return s.hasClass(a)}))&&(h("tbody tr",this).removeClass(g.join(" ")),p.asDestroyStripes=g.slice());o=[];g=this.getElementsByTagName("thead");0!==g.length&&(da(p.aoHeader,g[0]),o=qa(p));if(null===d.aoColumns){r=[];g=0;for(i=o.length;g<i;g++)r.push(null)}else r=d.aoColumns;g=0;for(i=r.length;g<i;g++)Fa(p,o?o[g]:null);ib(p,d.aoColumnDefs,
r,function(a,b){ka(p,a,b)});if(s.length){var u=function(a,b){return a.getAttribute("data-"+b)!==null?b:null};h.each(na(p,s[0]).cells,function(a,b){var c=p.aoColumns[a];if(c.mData===a){var d=u(b,"sort")||u(b,"order"),e=u(b,"filter")||u(b,"search");if(d!==null||e!==null){c.mData={_:a+".display",sort:d!==null?a+".@data-"+d:k,type:d!==null?a+".@data-"+d:k,filter:e!==null?a+".@data-"+e:k};ka(p,a)}}})}var v=p.oFeatures;d.bStateSave&&(v.bStateSave=!0,Kb(p,d),z(p,"aoDrawCallback",ya,"state_save"));if(d.aaSorting===
k){o=p.aaSorting;g=0;for(i=o.length;g<i;g++)o[g][1]=p.aoColumns[g].asSorting[0]}xa(p);v.bSort&&z(p,"aoDrawCallback",function(){if(p.bSorted){var a=T(p),b={};h.each(a,function(a,c){b[c.src]=c.dir});w(p,null,"order",[p,a,b]);Jb(p)}});z(p,"aoDrawCallback",function(){(p.bSorted||B(p)==="ssp"||v.bDeferRender)&&xa(p)},"sc");gb(p);g=q.children("caption").each(function(){this._captionSide=q.css("caption-side")});i=q.children("thead");0===i.length&&(i=h("<thead/>").appendTo(this));p.nTHead=i[0];i=q.children("tbody");
0===i.length&&(i=h("<tbody/>").appendTo(this));p.nTBody=i[0];i=q.children("tfoot");if(0===i.length&&0<g.length&&(""!==p.oScroll.sX||""!==p.oScroll.sY))i=h("<tfoot/>").appendTo(this);0===i.length||0===i.children().length?q.addClass(j.sNoFooter):0<i.length&&(p.nTFoot=i[0],da(p.aoFooter,p.nTFoot));if(d.aaData)for(g=0;g<d.aaData.length;g++)J(p,d.aaData[g]);else(p.bDeferLoading||"dom"==B(p))&&ma(p,h(p.nTBody).children("tr"));p.aiDisplay=p.aiDisplayMaster.slice();p.bInitialised=!0;!1===n&&ga(p)}});b=null;
return this};var Tb=[],v=Array.prototype,cc=function(a){var b,c,e=m.settings,d=h.map(e,function(a){return a.nTable});if(a){if(a.nTable&&a.oApi)return[a];if(a.nodeName&&"table"===a.nodeName.toLowerCase())return b=h.inArray(a,d),-1!==b?[e[b]]:null;if(a&&"function"===typeof a.settings)return a.settings().toArray();"string"===typeof a?c=h(a):a instanceof h&&(c=a)}else return[];if(c)return c.map(function(){b=h.inArray(this,d);return-1!==b?e[b]:null}).toArray()};t=function(a,b){if(!this instanceof t)throw"DT API must be constructed as a new object";
var c=[],e=function(a){(a=cc(a))&&c.push.apply(c,a)};if(h.isArray(a))for(var d=0,f=a.length;d<f;d++)e(a[d]);else e(a);this.context=Na(c);b&&this.push.apply(this,b.toArray?b.toArray():b);this.selector={rows:null,cols:null,opts:null};t.extend(this,this,Tb)};m.Api=t;t.prototype={concat:v.concat,context:[],each:function(a){for(var b=0,c=this.length;b<c;b++)a.call(this,this[b],b,this);return this},eq:function(a){var b=this.context;return b.length>a?new t(b[a],this[a]):null},filter:function(a){var b=[];
if(v.filter)b=v.filter.call(this,a,this);else for(var c=0,e=this.length;c<e;c++)a.call(this,this[c],c,this)&&b.push(this[c]);return new t(this.context,b)},flatten:function(){var a=[];return new t(this.context,a.concat.apply(a,this.toArray()))},join:v.join,indexOf:v.indexOf||function(a,b){for(var c=b||0,e=this.length;c<e;c++)if(this[c]===a)return c;return-1},iterator:function(a,b,c,e){var d=[],f,g,h,j,n,l=this.context,q,o,m=this.selector;"string"===typeof a&&(e=c,c=b,b=a,a=!1);g=0;for(h=l.length;g<
h;g++){var p=new t(l[g]);if("table"===b)f=c.call(p,l[g],g),f!==k&&d.push(f);else if("columns"===b||"rows"===b)f=c.call(p,l[g],this[g],g),f!==k&&d.push(f);else if("column"===b||"column-rows"===b||"row"===b||"cell"===b){o=this[g];"column-rows"===b&&(q=Ca(l[g],m.opts));j=0;for(n=o.length;j<n;j++)f=o[j],f="cell"===b?c.call(p,l[g],f.row,f.column,g,j):c.call(p,l[g],f,g,j,q),f!==k&&d.push(f)}}return d.length||e?(a=new t(l,a?d.concat.apply([],d):d),b=a.selector,b.rows=m.rows,b.cols=m.cols,b.opts=m.opts,a):
this},lastIndexOf:v.lastIndexOf||function(a,b){return this.indexOf.apply(this.toArray.reverse(),arguments)},length:0,map:function(a){var b=[];if(v.map)b=v.map.call(this,a,this);else for(var c=0,e=this.length;c<e;c++)b.push(a.call(this,this[c],c));return new t(this.context,b)},pluck:function(a){return this.map(function(b){return b[a]})},pop:v.pop,push:v.push,reduce:v.reduce||function(a,b){return hb(this,a,b,0,this.length,1)},reduceRight:v.reduceRight||function(a,b){return hb(this,a,b,this.length-1,
-1,-1)},reverse:v.reverse,selector:null,shift:v.shift,sort:v.sort,splice:v.splice,toArray:function(){return v.slice.call(this)},to$:function(){return h(this)},toJQuery:function(){return h(this)},unique:function(){return new t(this.context,Na(this))},unshift:v.unshift};t.extend=function(a,b,c){if(c.length&&b&&(b instanceof t||b.__dt_wrapper)){var e,d,f,g=function(a,b,c){return function(){var d=b.apply(a,arguments);t.extend(d,d,c.methodExt);return d}};e=0;for(d=c.length;e<d;e++)f=c[e],b[f.name]="function"===
typeof f.val?g(a,f.val,f):h.isPlainObject(f.val)?{}:f.val,b[f.name].__dt_wrapper=!0,t.extend(a,b[f.name],f.propExt)}};t.register=r=function(a,b){if(h.isArray(a))for(var c=0,e=a.length;c<e;c++)t.register(a[c],b);else for(var d=a.split("."),f=Tb,g,i,c=0,e=d.length;c<e;c++){g=(i=-1!==d[c].indexOf("()"))?d[c].replace("()",""):d[c];var j;a:{j=0;for(var n=f.length;j<n;j++)if(f[j].name===g){j=f[j];break a}j=null}j||(j={name:g,val:{},methodExt:[],propExt:[]},f.push(j));c===e-1?j.val=b:f=i?j.methodExt:j.propExt}};
t.registerPlural=u=function(a,b,c){t.register(a,c);t.register(b,function(){var a=c.apply(this,arguments);return a===this?this:a instanceof t?a.length?h.isArray(a[0])?new t(a.context,a[0]):a[0]:k:a})};r("tables()",function(a){var b;if(a){b=t;var c=this.context;if("number"===typeof a)a=[c[a]];else var e=h.map(c,function(a){return a.nTable}),a=h(e).filter(a).map(function(){var a=h.inArray(this,e);return c[a]}).toArray();b=new b(a)}else b=this;return b});r("table()",function(a){var a=this.tables(a),b=
a.context;return b.length?new t(b[0]):a});u("tables().nodes()","table().node()",function(){return this.iterator("table",function(a){return a.nTable},1)});u("tables().body()","table().body()",function(){return this.iterator("table",function(a){return a.nTBody},1)});u("tables().header()","table().header()",function(){return this.iterator("table",function(a){return a.nTHead},1)});u("tables().footer()","table().footer()",function(){return this.iterator("table",function(a){return a.nTFoot},1)});u("tables().containers()",
"table().container()",function(){return this.iterator("table",function(a){return a.nTableWrapper},1)});r("draw()",function(a){return this.iterator("table",function(b){N(b,!1===a)})});r("page()",function(a){return a===k?this.page.info().page:this.iterator("table",function(b){Ta(b,a)})});r("page.info()",function(){if(0===this.context.length)return k;var a=this.context[0],b=a._iDisplayStart,c=a._iDisplayLength,e=a.fnRecordsDisplay(),d=-1===c;return{page:d?0:Math.floor(b/c),pages:d?1:Math.ceil(e/c),start:b,
end:a.fnDisplayEnd(),length:c,recordsTotal:a.fnRecordsTotal(),recordsDisplay:e}});r("page.len()",function(a){return a===k?0!==this.context.length?this.context[0]._iDisplayLength:k:this.iterator("table",function(b){Ra(b,a)})});var Ub=function(a,b,c){if(c){var e=new t(a);e.one("draw",function(){c(e.ajax.json())})}"ssp"==B(a)?N(a,b):(C(a,!0),ra(a,[],function(c){oa(a);for(var c=sa(a,c),e=0,g=c.length;e<g;e++)J(a,c[e]);N(a,b);C(a,!1)}))};r("ajax.json()",function(){var a=this.context;if(0<a.length)return a[0].json});
r("ajax.params()",function(){var a=this.context;if(0<a.length)return a[0].oAjaxData});r("ajax.reload()",function(a,b){return this.iterator("table",function(c){Ub(c,!1===b,a)})});r("ajax.url()",function(a){var b=this.context;if(a===k){if(0===b.length)return k;b=b[0];return b.ajax?h.isPlainObject(b.ajax)?b.ajax.url:b.ajax:b.sAjaxSource}return this.iterator("table",function(b){h.isPlainObject(b.ajax)?b.ajax.url=a:b.ajax=a})});r("ajax.url().load()",function(a,b){return this.iterator("table",function(c){Ub(c,
!1===b,a)})});var $a=function(a,b){var c=[],e,d,f,g,i,j;e=typeof a;if(!a||"string"===e||"function"===e||a.length===k)a=[a];f=0;for(g=a.length;f<g;f++){d=a[f]&&a[f].split?a[f].split(","):[a[f]];i=0;for(j=d.length;i<j;i++)(e=b("string"===typeof d[i]?h.trim(d[i]):d[i]))&&e.length&&c.push.apply(c,e)}return c},ab=function(a){a||(a={});a.filter&&!a.search&&(a.search=a.filter);return{search:a.search||"none",order:a.order||"current",page:a.page||"all"}},bb=function(a){for(var b=0,c=a.length;b<c;b++)if(0<
a[b].length)return a[0]=a[b],a.length=1,a.context=[a.context[b]],a;a.length=0;return a},Ca=function(a,b){var c,e,d,f=[],g=a.aiDisplay;c=a.aiDisplayMaster;var i=b.search;e=b.order;d=b.page;if("ssp"==B(a))return"removed"===i?[]:U(0,c.length);if("current"==d){c=a._iDisplayStart;for(e=a.fnDisplayEnd();c<e;c++)f.push(g[c])}else if("current"==e||"applied"==e)f="none"==i?c.slice():"applied"==i?g.slice():h.map(c,function(a){return-1===h.inArray(a,g)?a:null});else if("index"==e||"original"==e){c=0;for(e=a.aoData.length;c<
e;c++)"none"==i?f.push(c):(d=h.inArray(c,g),(-1===d&&"removed"==i||0<=d&&"applied"==i)&&f.push(c))}return f};r("rows()",function(a,b){a===k?a="":h.isPlainObject(a)&&(b=a,a="");var b=ab(b),c=this.iterator("table",function(c){var d=b;return $a(a,function(a){var b=Pb(a);if(b!==null&&!d)return[b];var i=Ca(c,d);if(b!==null&&h.inArray(b,i)!==-1)return[b];if(!a)return i;if(typeof a==="function")return h.map(i,function(b){var d=c.aoData[b];return a(b,d._aData,d.nTr)?b:null});b=Sb(ia(c.aoData,i,"nTr"));return a.nodeName&&
h.inArray(a,b)!==-1?[a._DT_RowIndex]:h(b).filter(a).map(function(){return this._DT_RowIndex}).toArray()})},1);c.selector.rows=a;c.selector.opts=b;return c});r("rows().nodes()",function(){return this.iterator("row",function(a,b){return a.aoData[b].nTr||k},1)});r("rows().data()",function(){return this.iterator(!0,"rows",function(a,b){return ia(a.aoData,b,"_aData")},1)});u("rows().cache()","row().cache()",function(a){return this.iterator("row",function(b,c){var e=b.aoData[c];return"search"===a?e._aFilterData:
e._aSortData},1)});u("rows().invalidate()","row().invalidate()",function(a){return this.iterator("row",function(b,c){ca(b,c,a)})});u("rows().indexes()","row().index()",function(){return this.iterator("row",function(a,b){return b},1)});u("rows().remove()","row().remove()",function(){var a=this;return this.iterator("row",function(b,c,e){var d=b.aoData;d.splice(c,1);for(var f=0,g=d.length;f<g;f++)null!==d[f].nTr&&(d[f].nTr._DT_RowIndex=f);h.inArray(c,b.aiDisplay);pa(b.aiDisplayMaster,c);pa(b.aiDisplay,
c);pa(a[e],c,!1);Sa(b)})});r("rows.add()",function(a){var b=this.iterator("table",function(b){var c,f,g,h=[];f=0;for(g=a.length;f<g;f++)c=a[f],c.nodeName&&"TR"===c.nodeName.toUpperCase()?h.push(ma(b,c)[0]):h.push(J(b,c));return h},1),c=this.rows(-1);c.pop();c.push.apply(c,b.toArray());return c});r("row()",function(a,b){return bb(this.rows(a,b))});r("row().data()",function(a){var b=this.context;if(a===k)return b.length&&this.length?b[0].aoData[this[0]]._aData:k;b[0].aoData[this[0]]._aData=a;ca(b[0],
this[0],"data");return this});r("row().node()",function(){var a=this.context;return a.length&&this.length?a[0].aoData[this[0]].nTr||null:null});r("row.add()",function(a){a instanceof h&&a.length&&(a=a[0]);var b=this.iterator("table",function(b){return a.nodeName&&"TR"===a.nodeName.toUpperCase()?ma(b,a)[0]:J(b,a)});return this.row(b[0])});var cb=function(a,b){var c=a.context;c.length&&(c=c[0].aoData[b!==k?b:a[0]],c._details&&(c._details.remove(),c._detailsShow=k,c._details=k))},Vb=function(a,b){var c=
a.context;if(c.length&&a.length){var e=c[0].aoData[a[0]];if(e._details){(e._detailsShow=b)?e._details.insertAfter(e.nTr):e._details.detach();var d=c[0],f=new t(d),g=d.aoData;f.off("draw.dt.DT_details column-visibility.dt.DT_details destroy.dt.DT_details");0<D(g,"_details").length&&(f.on("draw.dt.DT_details",function(a,b){d===b&&f.rows({page:"current"}).eq(0).each(function(a){a=g[a];a._detailsShow&&a._details.insertAfter(a.nTr)})}),f.on("column-visibility.dt.DT_details",function(a,b){if(d===b)for(var c,
e=aa(b),f=0,h=g.length;f<h;f++)c=g[f],c._details&&c._details.children("td[colspan]").attr("colspan",e)}),f.on("destroy.dt.DT_details",function(a,b){if(d===b)for(var c=0,e=g.length;c<e;c++)g[c]._details&&cb(f,c)}))}}};r("row().child()",function(a,b){var c=this.context;if(a===k)return c.length&&this.length?c[0].aoData[this[0]]._details:k;if(!0===a)this.child.show();else if(!1===a)cb(this);else if(c.length&&this.length){var e=c[0],c=c[0].aoData[this[0]],d=[],f=function(a,b){if(h.isArray(a)||a instanceof
h)for(var c=0,k=a.length;c<k;c++)f(a[c],b);else a.nodeName&&"tr"===a.nodeName.toLowerCase()?d.push(a):(c=h("<tr><td/></tr>").addClass(b),h("td",c).addClass(b).html(a)[0].colSpan=aa(e),d.push(c[0]))};f(a,b);c._details&&c._details.remove();c._details=h(d);c._detailsShow&&c._details.insertAfter(c.nTr)}return this});r(["row().child.show()","row().child().show()"],function(){Vb(this,!0);return this});r(["row().child.hide()","row().child().hide()"],function(){Vb(this,!1);return this});r(["row().child.remove()",
"row().child().remove()"],function(){cb(this);return this});r("row().child.isShown()",function(){var a=this.context;return a.length&&this.length?a[0].aoData[this[0]]._detailsShow||!1:!1});var dc=/^(.+):(name|visIdx|visible)$/,Wb=function(a,b,c,e,d){for(var c=[],e=0,f=d.length;e<f;e++)c.push(y(a,d[e],b));return c};r("columns()",function(a,b){a===k?a="":h.isPlainObject(a)&&(b=a,a="");var b=ab(b),c=this.iterator("table",function(c){var d=a,f=b,g=c.aoColumns,i=D(g,"sName"),j=D(g,"nTh");return $a(d,function(a){var b=
Pb(a);if(a==="")return U(g.length);if(b!==null)return[b>=0?b:g.length+b];if(typeof a==="function"){var d=Ca(c,f);return h.map(g,function(b,f){return a(f,Wb(c,f,0,0,d),j[f])?f:null})}var k=typeof a==="string"?a.match(dc):"";if(k)switch(k[2]){case "visIdx":case "visible":b=parseInt(k[1],10);if(b<0){var m=h.map(g,function(a,b){return a.bVisible?b:null});return[m[m.length+b]]}return[la(c,b)];case "name":return h.map(i,function(a,b){return a===k[1]?b:null})}else return h(j).filter(a).map(function(){return h.inArray(this,
j)}).toArray()})},1);c.selector.cols=a;c.selector.opts=b;return c});u("columns().header()","column().header()",function(){return this.iterator("column",function(a,b){return a.aoColumns[b].nTh},1)});u("columns().footer()","column().footer()",function(){return this.iterator("column",function(a,b){return a.aoColumns[b].nTf},1)});u("columns().data()","column().data()",function(){return this.iterator("column-rows",Wb,1)});u("columns().dataSrc()","column().dataSrc()",function(){return this.iterator("column",
function(a,b){return a.aoColumns[b].mData},1)});u("columns().cache()","column().cache()",function(a){return this.iterator("column-rows",function(b,c,e,d,f){return ia(b.aoData,f,"search"===a?"_aFilterData":"_aSortData",c)},1)});u("columns().nodes()","column().nodes()",function(){return this.iterator("column-rows",function(a,b,c,e,d){return ia(a.aoData,d,"anCells",b)},1)});u("columns().visible()","column().visible()",function(a,b){return this.iterator("column",function(c,e){if(a===k)return c.aoColumns[e].bVisible;
var d=c.aoColumns,f=d[e],g=c.aoData,i,j,n;if(a!==k&&f.bVisible!==a){if(a){var l=h.inArray(!0,D(d,"bVisible"),e+1);i=0;for(j=g.length;i<j;i++)n=g[i].nTr,d=g[i].anCells,n&&n.insertBefore(d[e],d[l]||null)}else h(D(c.aoData,"anCells",e)).detach();f.bVisible=a;ea(c,c.aoHeader);ea(c,c.aoFooter);if(b===k||b)X(c),(c.oScroll.sX||c.oScroll.sY)&&Y(c);w(c,null,"column-visibility",[c,e,a]);ya(c)}})});u("columns().indexes()","column().index()",function(a){return this.iterator("column",function(b,c){return"visible"===
a?$(b,c):c},1)});r("columns.adjust()",function(){return this.iterator("table",function(a){X(a)},1)});r("column.index()",function(a,b){if(0!==this.context.length){var c=this.context[0];if("fromVisible"===a||"toData"===a)return la(c,b);if("fromData"===a||"toVisible"===a)return $(c,b)}});r("column()",function(a,b){return bb(this.columns(a,b))});r("cells()",function(a,b,c){h.isPlainObject(a)&&(a.row===k?(c=a,a=null):(c=b,b=null));h.isPlainObject(b)&&(c=b,b=null);if(null===b||b===k)return this.iterator("table",
function(b){var d=a,e=ab(c),f=b.aoData,g=Ca(b,e),e=Sb(ia(f,g,"anCells")),i=h([].concat.apply([],e)),j,l=b.aoColumns.length,n,m,r,t,s,u;return $a(d,function(a){var c=typeof a==="function";if(a===null||a===k||c){n=[];m=0;for(r=g.length;m<r;m++){j=g[m];for(t=0;t<l;t++){s={row:j,column:t};if(c){u=b.aoData[j];a(s,y(b,j,t),u.anCells[t])&&n.push(s)}else n.push(s)}}return n}return h.isPlainObject(a)?[a]:i.filter(a).map(function(a,b){j=b.parentNode._DT_RowIndex;return{row:j,column:h.inArray(b,f[j].anCells)}}).toArray()})});
var e=this.columns(b,c),d=this.rows(a,c),f,g,i,j,n,l=this.iterator("table",function(a,b){f=[];g=0;for(i=d[b].length;g<i;g++){j=0;for(n=e[b].length;j<n;j++)f.push({row:d[b][g],column:e[b][j]})}return f},1);h.extend(l.selector,{cols:b,rows:a,opts:c});return l});u("cells().nodes()","cell().node()",function(){return this.iterator("cell",function(a,b,c){return(a=a.aoData[b].anCells)?a[c]:k},1)});r("cells().data()",function(){return this.iterator("cell",function(a,b,c){return y(a,b,c)},1)});u("cells().cache()",
"cell().cache()",function(a){a="search"===a?"_aFilterData":"_aSortData";return this.iterator("cell",function(b,c,e){return b.aoData[c][a][e]},1)});u("cells().render()","cell().render()",function(a){return this.iterator("cell",function(b,c,e){return y(b,c,e,a)},1)});u("cells().indexes()","cell().index()",function(){return this.iterator("cell",function(a,b,c){return{row:b,column:c,columnVisible:$(a,c)}},1)});u("cells().invalidate()","cell().invalidate()",function(a){return this.iterator("cell",function(b,
c,e){ca(b,c,a,e)})});r("cell()",function(a,b,c){return bb(this.cells(a,b,c))});r("cell().data()",function(a){var b=this.context,c=this[0];if(a===k)return b.length&&c.length?y(b[0],c[0].row,c[0].column):k;Ia(b[0],c[0].row,c[0].column,a);ca(b[0],c[0].row,"data",c[0].column);return this});r("order()",function(a,b){var c=this.context;if(a===k)return 0!==c.length?c[0].aaSorting:k;"number"===typeof a?a=[[a,b]]:h.isArray(a[0])||(a=Array.prototype.slice.call(arguments));return this.iterator("table",function(b){b.aaSorting=
a.slice()})});r("order.listener()",function(a,b,c){return this.iterator("table",function(e){Oa(e,a,b,c)})});r(["columns().order()","column().order()"],function(a){var b=this;return this.iterator("table",function(c,e){var d=[];h.each(b[e],function(b,c){d.push([c,a])});c.aaSorting=d})});r("search()",function(a,b,c,e){var d=this.context;return a===k?0!==d.length?d[0].oPreviousSearch.sSearch:k:this.iterator("table",function(d){d.oFeatures.bFilter&&fa(d,h.extend({},d.oPreviousSearch,{sSearch:a+"",bRegex:null===
b?!1:b,bSmart:null===c?!0:c,bCaseInsensitive:null===e?!0:e}),1)})});u("columns().search()","column().search()",function(a,b,c,e){return this.iterator("column",function(d,f){var g=d.aoPreSearchCols;if(a===k)return g[f].sSearch;d.oFeatures.bFilter&&(h.extend(g[f],{sSearch:a+"",bRegex:null===b?!1:b,bSmart:null===c?!0:c,bCaseInsensitive:null===e?!0:e}),fa(d,d.oPreviousSearch,1))})});r("state()",function(){return this.context.length?this.context[0].oSavedState:null});r("state.clear()",function(){return this.iterator("table",
function(a){a.fnStateSaveCallback.call(a.oInstance,a,{})})});r("state.loaded()",function(){return this.context.length?this.context[0].oLoadedState:null});r("state.save()",function(){return this.iterator("table",function(a){ya(a)})});m.versionCheck=m.fnVersionCheck=function(a){for(var b=m.version.split("."),a=a.split("."),c,e,d=0,f=a.length;d<f;d++)if(c=parseInt(b[d],10)||0,e=parseInt(a[d],10)||0,c!==e)return c>e;return!0};m.isDataTable=m.fnIsDataTable=function(a){var b=h(a).get(0),c=!1;h.each(m.settings,
function(a,d){var f=d.nScrollHead?h("table",d.nScrollHead)[0]:null,g=d.nScrollFoot?h("table",d.nScrollFoot)[0]:null;if(d.nTable===b||f===b||g===b)c=!0});return c};m.tables=m.fnTables=function(a){return h.map(m.settings,function(b){if(!a||a&&h(b.nTable).is(":visible"))return b.nTable})};m.util={throttle:ua,escapeRegex:va};m.camelToHungarian=H;r("$()",function(a,b){var c=this.rows(b).nodes(),c=h(c);return h([].concat(c.filter(a).toArray(),c.find(a).toArray()))});h.each(["on","one","off"],function(a,
b){r(b+"()",function(){var a=Array.prototype.slice.call(arguments);a[0].match(/\.dt\b/)||(a[0]+=".dt");var e=h(this.tables().nodes());e[b].apply(e,a);return this})});r("clear()",function(){return this.iterator("table",function(a){oa(a)})});r("settings()",function(){return new t(this.context,this.context)});r("init()",function(){var a=this.context;return a.length?a[0].oInit:null});r("data()",function(){return this.iterator("table",function(a){return D(a.aoData,"_aData")}).flatten()});r("destroy()",
function(a){a=a||!1;return this.iterator("table",function(b){var c=b.nTableWrapper.parentNode,e=b.oClasses,d=b.nTable,f=b.nTBody,g=b.nTHead,i=b.nTFoot,j=h(d),f=h(f),k=h(b.nTableWrapper),l=h.map(b.aoData,function(a){return a.nTr}),q;b.bDestroying=!0;w(b,"aoDestroyCallback","destroy",[b]);a||(new t(b)).columns().visible(!0);k.unbind(".DT").find(":not(tbody *)").unbind(".DT");h(Ea).unbind(".DT-"+b.sInstance);d!=g.parentNode&&(j.children("thead").detach(),j.append(g));i&&d!=i.parentNode&&(j.children("tfoot").detach(),
j.append(i));j.detach();k.detach();b.aaSorting=[];b.aaSortingFixed=[];xa(b);h(l).removeClass(b.asStripeClasses.join(" "));h("th, td",g).removeClass(e.sSortable+" "+e.sSortableAsc+" "+e.sSortableDesc+" "+e.sSortableNone);b.bJUI&&(h("th span."+e.sSortIcon+", td span."+e.sSortIcon,g).detach(),h("th, td",g).each(function(){var a=h("div."+e.sSortJUIWrapper,this);h(this).append(a.contents());a.detach()}));!a&&c&&c.insertBefore(d,b.nTableReinsertBefore);f.children().detach();f.append(l);j.css("width",b.sDestroyWidth).removeClass(e.sTable);
(q=b.asDestroyStripes.length)&&f.children().each(function(a){h(this).addClass(b.asDestroyStripes[a%q])});c=h.inArray(b,m.settings);-1!==c&&m.settings.splice(c,1)})});h.each(["column","row","cell"],function(a,b){r(b+"s().every()",function(a){return this.iterator(b,function(e,d,f){a.call((new t(e))[b](d,f))})})});m.version="1.10.6";m.settings=[];m.models={};m.models.oSearch={bCaseInsensitive:!0,sSearch:"",bRegex:!1,bSmart:!0};m.models.oRow={nTr:null,anCells:null,_aData:[],_aSortData:null,_aFilterData:null,
_sFilterRow:null,_sRowStripe:"",src:null};m.models.oColumn={idx:null,aDataSort:null,asSorting:null,bSearchable:null,bSortable:null,bVisible:null,_sManualType:null,_bAttrSrc:!1,fnCreatedCell:null,fnGetData:null,fnSetData:null,mData:null,mRender:null,nTh:null,nTf:null,sClass:null,sContentPadding:null,sDefaultContent:null,sName:null,sSortDataType:"std",sSortingClass:null,sSortingClassJUI:null,sTitle:null,sType:null,sWidth:null,sWidthOrig:null};m.defaults={aaData:null,aaSorting:[[0,"asc"]],aaSortingFixed:[],
ajax:null,aLengthMenu:[10,25,50,100],aoColumns:null,aoColumnDefs:null,aoSearchCols:[],asStripeClasses:null,bAutoWidth:!0,bDeferRender:!1,bDestroy:!1,bFilter:!0,bInfo:!0,bJQueryUI:!1,bLengthChange:!0,bPaginate:!0,bProcessing:!1,bRetrieve:!1,bScrollCollapse:!1,bServerSide:!1,bSort:!0,bSortMulti:!0,bSortCellsTop:!1,bSortClasses:!0,bStateSave:!1,fnCreatedRow:null,fnDrawCallback:null,fnFooterCallback:null,fnFormatNumber:function(a){return a.toString().replace(/\B(?=(\d{3})+(?!\d))/g,this.oLanguage.sThousands)},
fnHeaderCallback:null,fnInfoCallback:null,fnInitComplete:null,fnPreDrawCallback:null,fnRowCallback:null,fnServerData:null,fnServerParams:null,fnStateLoadCallback:function(a){try{return JSON.parse((-1===a.iStateDuration?sessionStorage:localStorage).getItem("DataTables_"+a.sInstance+"_"+location.pathname))}catch(b){}},fnStateLoadParams:null,fnStateLoaded:null,fnStateSaveCallback:function(a,b){try{(-1===a.iStateDuration?sessionStorage:localStorage).setItem("DataTables_"+a.sInstance+"_"+location.pathname,
JSON.stringify(b))}catch(c){}},fnStateSaveParams:null,iStateDuration:7200,iDeferLoading:null,iDisplayLength:10,iDisplayStart:0,iTabIndex:0,oClasses:{},oLanguage:{oAria:{sSortAscending:": activate to sort column ascending",sSortDescending:": activate to sort column descending"},oPaginate:{sFirst:"First",sLast:"Last",sNext:"Next",sPrevious:"Previous"},sEmptyTable:"No data available in table",sInfo:"Showing _START_ to _END_ of _TOTAL_ entries",sInfoEmpty:"Showing 0 to 0 of 0 entries",sInfoFiltered:"(filtered from _MAX_ total entries)",
sInfoPostFix:"",sDecimal:"",sThousands:",",sLengthMenu:"Show _MENU_ ",sLoadingRecords:"Loading...",sProcessing:"Processing...",sSearch:"Search:",sSearchPlaceholder:"",sUrl:"",sZeroRecords:"No matching records found"},oSearch:h.extend({},m.models.oSearch),sAjaxDataProp:"data",sAjaxSource:null,sDom:"lfrtip",searchDelay:null,sPaginationType:"simple_numbers",sScrollX:"",sScrollXInner:"",sScrollY:"",sServerMethod:"GET",renderer:null};V(m.defaults);m.defaults.column={aDataSort:null,iDataSort:-1,
asSorting:["asc","desc"],bSearchable:!0,bSortable:!0,bVisible:!0,fnCreatedCell:null,mData:null,mRender:null,sCellType:"td",sClass:"",sContentPadding:"",sDefaultContent:null,sName:"",sSortDataType:"std",sTitle:null,sType:null,sWidth:null};V(m.defaults.column);m.models.oSettings={oFeatures:{bAutoWidth:null,bDeferRender:null,bFilter:null,bInfo:null,bLengthChange:null,bPaginate:null,bProcessing:null,bServerSide:null,bSort:null,bSortMulti:null,bSortClasses:null,bStateSave:null},oScroll:{bCollapse:null,
iBarWidth:0,sX:null,sXInner:null,sY:null},oLanguage:{fnInfoCallback:null},oBrowser:{bScrollOversize:!1,bScrollbarLeft:!1},ajax:null,aanFeatures:[],aoData:[],aiDisplay:[],aiDisplayMaster:[],aoColumns:[],aoHeader:[],aoFooter:[],oPreviousSearch:{},aoPreSearchCols:[],aaSorting:null,aaSortingFixed:[],asStripeClasses:null,asDestroyStripes:[],sDestroyWidth:0,aoRowCallback:[],aoHeaderCallback:[],aoFooterCallback:[],aoDrawCallback:[],aoRowCreatedCallback:[],aoPreDrawCallback:[],aoInitComplete:[],aoStateSaveParams:[],
aoStateLoadParams:[],aoStateLoaded:[],sTableId:"",nTable:null,nTHead:null,nTFoot:null,nTBody:null,nTableWrapper:null,bDeferLoading:!1,bInitialised:!1,aoOpenRows:[],sDom:null,searchDelay:null,sPaginationType:"two_button",iStateDuration:0,aoStateSave:[],aoStateLoad:[],oSavedState:null,oLoadedState:null,sAjaxSource:null,sAjaxDataProp:null,bAjaxDataGet:!0,jqXHR:null,json:k,oAjaxData:k,fnServerData:null,aoServerParams:[],sServerMethod:null,fnFormatNumber:null,aLengthMenu:null,iDraw:0,bDrawing:!1,iDrawError:-1,
_iDisplayLength:10,_iDisplayStart:0,_iRecordsTotal:0,_iRecordsDisplay:0,bJUI:null,oClasses:{},bFiltered:!1,bSorted:!1,bSortCellsTop:null,oInit:null,aoDestroyCallback:[],fnRecordsTotal:function(){return"ssp"==B(this)?1*this._iRecordsTotal:this.aiDisplayMaster.length},fnRecordsDisplay:function(){return"ssp"==B(this)?1*this._iRecordsDisplay:this.aiDisplay.length},fnDisplayEnd:function(){var a=this._iDisplayLength,b=this._iDisplayStart,c=b+a,e=this.aiDisplay.length,d=this.oFeatures,f=d.bPaginate;return d.bServerSide?
!1===f||-1===a?b+e:Math.min(b+a,this._iRecordsDisplay):!f||c>e||-1===a?e:c},oInstance:null,sInstance:null,iTabIndex:0,nScrollHead:null,nScrollFoot:null,aLastSort:[],oPlugins:{}};m.ext=x={buttons:{},classes:{},errMode:"alert",feature:[],search:[],internal:{},legacy:{ajax:null},pager:{},renderer:{pageButton:{},header:{}},order:{},type:{detect:[],search:{},order:{}},_unique:0,fnVersionCheck:m.fnVersionCheck,iApiIndex:0,oJUIClasses:{},sVersion:m.version};h.extend(x,{afnFiltering:x.search,aTypes:x.type.detect,
ofnSearch:x.type.search,oSort:x.type.order,afnSortData:x.order,aoFeatures:x.feature,oApi:x.internal,oStdClasses:x.classes,oPagination:x.pager});h.extend(m.ext.classes,{sTable:"dataTable",sNoFooter:"no-footer",sPageButton:"paginate_button",sPageButtonActive:"current",sPageButtonDisabled:"disabled",sStripeOdd:"odd",sStripeEven:"even",sRowEmpty:"dataTables_empty",sWrapper:"dataTables_wrapper",sFilter:"dataTables_filter",sInfo:"dataTables_info",sPaging:"dataTables_paginate paging_",sLength:"dataTables_length",
sProcessing:"dataTables_processing",sSortAsc:"sorting_asc",sSortDesc:"sorting_desc",sSortable:"sorting",sSortableAsc:"sorting_asc_disabled",sSortableDesc:"sorting_desc_disabled",sSortableNone:"sorting_disabled",sSortColumn:"sorting_",sFilterInput:"",sLengthSelect:"",sScrollWrapper:"dataTables_scroll",sScrollHead:"dataTables_scrollHead",sScrollHeadInner:"dataTables_scrollHeadInner",sScrollBody:"dataTables_scrollBody",sScrollFoot:"dataTables_scrollFoot",sScrollFootInner:"dataTables_scrollFootInner",
sHeaderTH:"",sFooterTH:"",sSortJUIAsc:"",sSortJUIDesc:"",sSortJUI:"",sSortJUIAscAllowed:"",sSortJUIDescAllowed:"",sSortJUIWrapper:"",sSortIcon:"",sJUIHeader:"",sJUIFooter:""});var Da="",Da="",F=Da+"ui-state-default",ja=Da+"css_right ui-icon ui-icon-",Xb=Da+"fg-toolbar ui-toolbar ui-widget-header ui-helper-clearfix";h.extend(m.ext.oJUIClasses,m.ext.classes,{sPageButton:"fg-button ui-button "+F,sPageButtonActive:"ui-state-disabled",sPageButtonDisabled:"ui-state-disabled",sPaging:"dataTables_paginate fg-buttonset ui-buttonset fg-buttonset-multi ui-buttonset-multi paging_",
sSortAsc:F+" sorting_asc",sSortDesc:F+" sorting_desc",sSortable:F+" sorting",sSortableAsc:F+" sorting_asc_disabled",sSortableDesc:F+" sorting_desc_disabled",sSortableNone:F+" sorting_disabled",sSortJUIAsc:ja+"triangle-1-n",sSortJUIDesc:ja+"triangle-1-s",sSortJUI:ja+"carat-2-n-s",sSortJUIAscAllowed:ja+"carat-1-n",sSortJUIDescAllowed:ja+"carat-1-s",sSortJUIWrapper:"DataTables_sort_wrapper",sSortIcon:"DataTables_sort_icon",sScrollHead:"dataTables_scrollHead "+F,sScrollFoot:"dataTables_scrollFoot "+F,
sHeaderTH:F,sFooterTH:F,sJUIHeader:Xb+" ui-corner-tl ui-corner-tr",sJUIFooter:Xb+" ui-corner-bl ui-corner-br"});var Mb=m.ext.pager;h.extend(Mb,{simple:function(){return["previous","next"]},full:function(){return["first","previous","next","last"]},simple_numbers:function(a,b){return["previous",Wa(a,b),"next"]},full_numbers:function(a,b){return["first","previous",Wa(a,b),"next","last"]},_numbers:Wa,numbers_length:7});h.extend(!0,m.ext.renderer,{pageButton:{_:function(a,b,c,e,d,f){var g=a.oClasses,i=
a.oLanguage.oPaginate,j,k,l=0,m=function(b,e){var o,r,t,s,u=function(b){Ta(a,b.data.action,true)};o=0;for(r=e.length;o<r;o++){s=e[o];if(h.isArray(s)){t=h("<"+(s.DT_el||"div")+"/>").appendTo(b);m(t,s)}else{k=j="";switch(s){case "ellipsis":b.append('<span class="ellipsis">&#x2026;</span>');break;case "first":j=i.sFirst;k=s+(d>0?"":" "+g.sPageButtonDisabled);break;case "previous":j=i.sPrevious;k=s+(d>0?"":" "+g.sPageButtonDisabled);break;case "next":j=i.sNext;k=s+(d<f-1?"":" "+g.sPageButtonDisabled);
break;case "last":j=i.sLast;k=s+(d<f-1?"":" "+g.sPageButtonDisabled);break;default:j=s+1;k=d===s?g.sPageButtonActive:""}if(j){t=h("<a>",{"class":g.sPageButton+" "+k,"aria-controls":a.sTableId,"data-dt-idx":l,tabindex:a.iTabIndex,id:c===0&&typeof s==="string"?a.sTableId+"_"+s:null}).html(j).appendTo(b);Va(t,{action:s},u);l++}}}},o;try{o=h(P.activeElement).data("dt-idx")}catch(r){}m(h(b).empty(),e);o&&h(b).find("[data-dt-idx="+o+"]").focus()}}});h.extend(m.ext.type.detect,[function(a,b){var c=b.oLanguage.sDecimal;
return Za(a,c)?"num"+c:null},function(a){if(a&&!(a instanceof Date)&&(!ac.test(a)||!bc.test(a)))return null;var b=Date.parse(a);return null!==b&&!isNaN(b)||I(a)?"date":null},function(a,b){var c=b.oLanguage.sDecimal;return Za(a,c,!0)?"num-fmt"+c:null},function(a,b){var c=b.oLanguage.sDecimal;return Rb(a,c)?"html-num"+c:null},function(a,b){var c=b.oLanguage.sDecimal;return Rb(a,c,!0)?"html-num-fmt"+c:null},function(a){return I(a)||"string"===typeof a&&-1!==a.indexOf("<")?"html":null}]);h.extend(m.ext.type.search,
{html:function(a){return I(a)?a:"string"===typeof a?a.replace(Ob," ").replace(Ba,""):""},string:function(a){return I(a)?a:"string"===typeof a?a.replace(Ob," "):a}});var Aa=function(a,b,c,e){if(0!==a&&(!a||"-"===a))return-Infinity;b&&(a=Qb(a,b));a.replace&&(c&&(a=a.replace(c,"")),e&&(a=a.replace(e,"")));return 1*a};h.extend(x.type.order,{"date-pre":function(a){return Date.parse(a)||0},"html-pre":function(a){return I(a)?"":a.replace?a.replace(/<.*?>/g,"").toLowerCase():a+""},"string-pre":function(a){return I(a)?
"":"string"===typeof a?a.toLowerCase():!a.toString?"":a.toString()},"string-asc":function(a,b){return a<b?-1:a>b?1:0},"string-desc":function(a,b){return a<b?1:a>b?-1:0}});db("");h.extend(!0,m.ext.renderer,{header:{_:function(a,b,c,e){h(a.nTable).on("order.dt.DT",function(d,f,g,h){if(a===f){d=c.idx;b.removeClass(c.sSortingClass+" "+e.sSortAsc+" "+e.sSortDesc).addClass(h[d]=="asc"?e.sSortAsc:h[d]=="desc"?e.sSortDesc:c.sSortingClass)}})},jqueryui:function(a,b,c,e){h("<div/>").addClass(e.sSortJUIWrapper).append(b.contents()).append(h("<span/>").addClass(e.sSortIcon+
" "+c.sSortingClassJUI)).appendTo(b);h(a.nTable).on("order.dt.DT",function(d,f,g,h){if(a===f){d=c.idx;b.removeClass(e.sSortAsc+" "+e.sSortDesc).addClass(h[d]=="asc"?e.sSortAsc:h[d]=="desc"?e.sSortDesc:c.sSortingClass);b.find("span."+e.sSortIcon).removeClass(e.sSortJUIAsc+" "+e.sSortJUIDesc+" "+e.sSortJUI+" "+e.sSortJUIAscAllowed+" "+e.sSortJUIDescAllowed).addClass(h[d]=="asc"?e.sSortJUIAsc:h[d]=="desc"?e.sSortJUIDesc:c.sSortingClassJUI)}})}}});m.render={number:function(a,b,c,e){return{display:function(d){if("number"!==
typeof d&&"string"!==typeof d)return d;var f=0>d?"-":"",d=Math.abs(parseFloat(d)),g=parseInt(d,10),d=c?b+(d-g).toFixed(c).substring(2):"";return f+(e||"")+g.toString().replace(/\B(?=(\d{3})+(?!\d))/g,a)+d}}}};h.extend(m.ext.internal,{_fnExternApiFunc:Nb,_fnBuildAjax:ra,_fnAjaxUpdate:kb,_fnAjaxParameters:tb,_fnAjaxUpdateDraw:ub,_fnAjaxDataSrc:sa,_fnAddColumn:Fa,_fnColumnOptions:ka,_fnAdjustColumnSizing:X,_fnVisibleToColumnIndex:la,_fnColumnIndexToVisible:$,_fnVisbleColumns:aa,_fnGetColumns:Z,_fnColumnTypes:Ha,
_fnApplyColumnDefs:ib,_fnHungarianMap:V,_fnCamelToHungarian:H,_fnLanguageCompat:O,_fnBrowserDetect:gb,_fnAddData:J,_fnAddTr:ma,_fnNodeToDataIndex:function(a,b){return b._DT_RowIndex!==k?b._DT_RowIndex:null},_fnNodeToColumnIndex:function(a,b,c){return h.inArray(c,a.aoData[b].anCells)},_fnGetCellData:y,_fnSetCellData:Ia,_fnSplitObjNotation:Ka,_fnGetObjectDataFn:W,_fnSetObjectDataFn:Q,_fnGetDataMaster:La,_fnClearTable:oa,_fnDeleteIndex:pa,_fnInvalidate:ca,_fnGetRowElements:na,_fnCreateTr:Ja,_fnBuildHead:jb,
_fnDrawHead:ea,_fnDraw:M,_fnReDraw:N,_fnAddOptionsHtml:mb,_fnDetectHeader:da,_fnGetUniqueThs:qa,_fnFeatureHtmlFilter:ob,_fnFilterComplete:fa,_fnFilterCustom:xb,_fnFilterColumn:wb,_fnFilter:vb,_fnFilterCreateSearch:Qa,_fnEscapeRegex:va,_fnFilterData:yb,_fnFeatureHtmlInfo:rb,_fnUpdateInfo:Bb,_fnInfoMacros:Cb,_fnInitialise:ga,_fnInitComplete:ta,_fnLengthChange:Ra,_fnFeatureHtmlLength:nb,_fnFeatureHtmlPaginate:sb,_fnPageChange:Ta,_fnFeatureHtmlProcessing:pb,_fnProcessingDisplay:C,_fnFeatureHtmlTable:qb,
_fnScrollDraw:Y,_fnApplyToChildren:G,_fnCalculateColumnWidths:Ga,_fnThrottle:ua,_fnConvertToWidth:Db,_fnScrollingWidthAdjust:Fb,_fnGetWidestNode:Eb,_fnGetMaxLenString:Gb,_fnStringToCss:s,_fnScrollBarWidth:Hb,_fnSortFlatten:T,_fnSort:lb,_fnSortAria:Jb,_fnSortListener:Ua,_fnSortAttachListener:Oa,_fnSortingClasses:xa,_fnSortData:Ib,_fnSaveState:ya,_fnLoadState:Kb,_fnSettingsFromNode:za,_fnLog:R,_fnMap:E,_fnBindAction:Va,_fnCallbackReg:z,_fnCallbackFire:w,_fnLengthOverflow:Sa,_fnRenderer:Pa,_fnDataSource:B,
_fnRowAttributes:Ma,_fnCalculateEnd:function(){}});h.fn.dataTable=m;h.fn.dataTableSettings=m.settings;h.fn.dataTableExt=m.ext;h.fn.DataTable=function(a){return h(this).dataTable(a).api()};h.each(m,function(a,b){h.fn.DataTable[a]=b});return h.fn.dataTable};"function"===typeof define&&define.amd?define("datatables",["jquery"],O):"object"===typeof exports?module.exports=O(require("jquery")):jQuery&&!jQuery.fn.dataTable&&O(jQuery)})(window,document);
```
