## Supplementary material for "Comparative pathway enrichment analysis in gastrointestinal cell lines Caco-2, HT-29, HEPG2, and colon fibroblasts using a custom expression panel for tight-junction and cytoskeletal regulatory genes": Sup. Results 1: jquery.js

```
/*!
 * jQuery JavaScript Library v2.1.1
 * http://jquery.com/
 *
 * Includes Sizzle.js
 * http://sizzlejs.com/
 *
 * Copyright 2005, 2014 jQuery Foundation, Inc. and other contributors
 * Released under the MIT license
 * http://jquery.org/license
 *
 * Date: 2014-05-01T17:11Z
 */
!function(a,b){"object"==typeof module&&"object"==typeof module.exports?module.exports=a.document?b(a,!0):function(a){if(!a.document)throw new Error("jQuery requires a window with a document");return b(a)}:b(a)}("undefined"!=typeof window?window:this,function(a,b){function c(a){var b=a.length,c=_.type(a);return"function"===c||_.isWindow(a)?!1:1===a.nodeType&&b?!0:"array"===c||0===b||"number"==typeof b&&b>0&&b-1 in a}function d(a,b,c){if(_.isFunction(b))return _.grep(a,function(a,d){return!!b.call(a,d,a)!==c});if(b.nodeType)return _.grep(a,function(a){return a===b!==c});if("string"==typeof b){if(hb.test(b))return _.filter(b,a,c);b=_.filter(b,a)}return _.grep(a,function(a){return U.call(b,a)>=0!==c})}function e(a,b){for(;(a=a[b])&&1!==a.nodeType;);return a}function f(a){var b=ob[a]={};return _.each(a.match(nb)||[],function(a,c){b[c]=!0}),b}function g(){Z.removeEventListener("DOMContentLoaded",g,!1),a.removeEventListener("load",g,!1),_.ready()}function h(){Object.defineProperty(this.cache={},0,{get:function(){return{}}}),this.expando=_.expando+Math.random()}function i(a,b,c){var d;if(void 0===c&&1===a.nodeType)if(d="data-"+b.replace(ub,"-$1").toLowerCase(),c=a.getAttribute(d),"string"==typeof c){try{c="true"===c?!0:"false"===c?!1:"null"===c?null:+c+""===c?+c:tb.test(c)?_.parseJSON(c):c}catch(e){}sb.set(a,b,c)}else c=void 0;return c}function j(){return!0}function k(){return!1}function l(){try{return Z.activeElement}catch(a){}}function m(a,b){return _.nodeName(a,"table")&&_.nodeName(11!==b.nodeType?b:b.firstChild,"tr")?a.getElementsByTagName("tbody")[0]||a.appendChild(a.ownerDocument.createElement("tbody")):a}function n(a){return a.type=(null!==a.getAttribute("type"))+"/"+a.type,a}function o(a){var b=Kb.exec(a.type);return b?a.type=b[1]:a.removeAttribute("type"),a}function p(a,b){for(var c=0,d=a.length;d>c;c++)rb.set(a[c],"globalEval",!b||rb.get(b[c],"globalEval"))}function q(a,b){var c,d,e,f,g,h,i,j;if(1===b.nodeType){if(rb.hasData(a)&&(f=rb.access(a),g=rb.set(b,f),j=f.events)){delete g.handle,g.events={};for(e in j)for(c=0,d=j[e].length;d>c;c++)_.event.add(b,e,j[e][c])}sb.hasData(a)&&(h=sb.access(a),i=_.extend({},h),sb.set(b,i))}}function r(a,b){var c=a.getElementsByTagName?a.getElementsByTagName(b||"*"):a.querySelectorAll?a.querySelectorAll(b||"*"):[];return void 0===b||b&&_.nodeName(a,b)?_.merge([a],c):c}function s(a,b){var c=b.nodeName.toLowerCase();"input"===c&&yb.test(a.type)?b.checked=a.checked:("input"===c||"textarea"===c)&&(b.defaultValue=a.defaultValue)}function t(b,c){var d,e=_(c.createElement(b)).appendTo(c.body),f=a.getDefaultComputedStyle&&(d=a.getDefaultComputedStyle(e[0]))?d.display:_.css(e[0],"display");return e.detach(),f}function u(a){var b=Z,c=Ob[a];return c||(c=t(a,b),"none"!==c&&c||(Nb=(Nb||_("<iframe frameborder='0' width='0' height='0'/>")).appendTo(b.documentElement),b=Nb[0].contentDocument,b.write(),b.close(),c=t(a,b),Nb.detach()),Ob[a]=c),c}function v(a,b,c){var d,e,f,g,h=a.style;return c=c||Rb(a),c&&(g=c.getPropertyValue(b)||c[b]),c&&(""!==g||_.contains(a.ownerDocument,a)||(g=_.style(a,b)),Qb.test(g)&&Pb.test(b)&&(d=h.width,e=h.minWidth,f=h.maxWidth,h.minWidth=h.maxWidth=h.width=g,g=c.width,h.width=d,h.minWidth=e,h.maxWidth=f)),void 0!==g?g+"":g}function w(a,b){return{get:function(){return a()?void delete this.get:(this.get=b).apply(this,arguments)}}}function x(a,b){if(b in a)return b;for(var c=b[0].toUpperCase()+b.slice(1),d=b,e=Xb.length;e--;)if(b=Xb[e]+c,b in a)return b;return d}function y(a,b,c){var d=Tb.exec(b);return d?Math.max(0,d[1]-(c||0))+(d[2]||"px"):b}function z(a,b,c,d,e){for(var f=c===(d?"border":"content")?4:"width"===b?1:0,g=0;4>f;f+=2)"margin"===c&&(g+=_.css(a,c+wb[f],!0,e)),d?("content"===c&&(g-=_.css(a,"padding"+wb[f],!0,e)),"margin"!==c&&(g-=_.css(a,"border"+wb[f]+"Width",!0,e))):(g+=_.css(a,"padding"+wb[f],!0,e),"padding"!==c&&(g+=_.css(a,"border"+wb[f]+"Width",!0,e)));return g}function A(a,b,c){var d=!0,e="width"===b?a.offsetWidth:a.offsetHeight,f=Rb(a),g="border-box"===_.css(a,"boxSizing",!1,f);if(0>=e||null==e){if(e=v(a,b,f),(0>e||null==e)&&(e=a.style[b]),Qb.test(e))return e;d=g&&(Y.boxSizingReliable()||e===a.style[b]),e=parseFloat(e)||0}return e+z(a,b,c||(g?"border":"content"),d,f)+"px"}function B(a,b){for(var c,d,e,f=[],g=0,h=a.length;h>g;g++)d=a[g],d.style&&(f[g]=rb.get(d,"olddisplay"),c=d.style.display,b?(f[g]||"none"!==c||(d.style.display=""),""===d.style.display&&xb(d)&&(f[g]=rb.access(d,"olddisplay",u(d.nodeName)))):(e=xb(d),"none"===c&&e||rb.set(d,"olddisplay",e?c:_.css(d,"display"))));for(g=0;h>g;g++)d=a[g],d.style&&(b&&"none"!==d.style.display&&""!==d.style.display||(d.style.display=b?f[g]||"":"none"));return a}function C(a,b,c,d,e){return new C.prototype.init(a,b,c,d,e)}function D(){return setTimeout(function(){Yb=void 0}),Yb=_.now()}function E(a,b){var c,d=0,e={height:a};for(b=b?1:0;4>d;d+=2-b)c=wb[d],e["margin"+c]=e["padding"+c]=a;return b&&(e.opacity=e.width=a),e}function F(a,b,c){for(var d,e=(cc[b]||[]).concat(cc["*"]),f=0,g=e.length;g>f;f++)if(d=e[f].call(c,b,a))return d}function G(a,b,c){var d,e,f,g,h,i,j,k,l=this,m={},n=a.style,o=a.nodeType&&xb(a),p=rb.get(a,"fxshow");c.queue||(h=_._queueHooks(a,"fx"),null==h.unqueued&&(h.unqueued=0,i=h.empty.fire,h.empty.fire=function(){h.unqueued||i()}),h.unqueued++,l.always(function(){l.always(function(){h.unqueued--,_.queue(a,"fx").length||h.empty.fire()})})),1===a.nodeType&&("height"in b||"width"in b)&&(c.overflow=[n.overflow,n.overflowX,n.overflowY],j=_.css(a,"display"),k="none"===j?rb.get(a,"olddisplay")||u(a.nodeName):j,"inline"===k&&"none"===_.css(a,"float")&&(n.display="inline-block")),c.overflow&&(n.overflow="hidden",l.always(function(){n.overflow=c.overflow[0],n.overflowX=c.overflow[1],n.overflowY=c.overflow[2]}));for(d in b)if(e=b[d],$b.exec(e)){if(delete b[d],f=f||"toggle"===e,e===(o?"hide":"show")){if("show"!==e||!p||void 0===p[d])continue;o=!0}m[d]=p&&p[d]||_.style(a,d)}else j=void 0;if(_.isEmptyObject(m))"inline"===("none"===j?u(a.nodeName):j)&&(n.display=j);else{p?"hidden"in p&&(o=p.hidden):p=rb.access(a,"fxshow",{}),f&&(p.hidden=!o),o?_(a).show():l.done(function(){_(a).hide()}),l.done(function(){var b;rb.remove(a,"fxshow");for(b in m)_.style(a,b,m[b])});for(d in m)g=F(o?p[d]:0,d,l),d in p||(p[d]=g.start,o&&(g.end=g.start,g.start="width"===d||"height"===d?1:0))}}function H(a,b){var c,d,e,f,g;for(c in a)if(d=_.camelCase(c),e=b[d],f=a[c],_.isArray(f)&&(e=f[1],f=a[c]=f[0]),c!==d&&(a[d]=f,delete a[c]),g=_.cssHooks[d],g&&"expand"in g){f=g.expand(f),delete a[d];for(c in f)c in a||(a[c]=f[c],b[c]=e)}else b[d]=e}function I(a,b,c){var d,e,f=0,g=bc.length,h=_.Deferred().always(function(){delete i.elem}),i=function(){if(e)return!1;for(var b=Yb||D(),c=Math.max(0,j.startTime+j.duration-b),d=c/j.duration||0,f=1-d,g=0,i=j.tweens.length;i>g;g++)j.tweens[g].run(f);return h.notifyWith(a,[j,f,c]),1>f&&i?c:(h.resolveWith(a,[j]),!1)},j=h.promise({elem:a,props:_.extend({},b),opts:_.extend(!0,{specialEasing:{}},c),originalProperties:b,originalOptions:c,startTime:Yb||D(),duration:c.duration,tweens:[],createTween:function(b,c){var d=_.Tween(a,j.opts,b,c,j.opts.specialEasing[b]||j.opts.easing);return j.tweens.push(d),d},stop:function(b){var c=0,d=b?j.tweens.length:0;if(e)return this;for(e=!0;d>c;c++)j.tweens[c].run(1);return b?h.resolveWith(a,[j,b]):h.rejectWith(a,[j,b]),this}}),k=j.props;for(H(k,j.opts.specialEasing);g>f;f++)if(d=bc[f].call(j,a,k,j.opts))return d;return _.map(k,F,j),_.isFunction(j.opts.start)&&j.opts.start.call(a,j),_.fx.timer(_.extend(i,{elem:a,anim:j,queue:j.opts.queue})),j.progress(j.opts.progress).done(j.opts.done,j.opts.complete).fail(j.opts.fail).always(j.opts.always)}function J(a){return function(b,c){"string"!=typeof b&&(c=b,b="*");var d,e=0,f=b.toLowerCase().match(nb)||[];if(_.isFunction(c))for(;d=f[e++];)"+"===d[0]?(d=d.slice(1)||"*",(a[d]=a[d]||[]).unshift(c)):(a[d]=a[d]||[]).push(c)}}function K(a,b,c,d){function e(h){var i;return f[h]=!0,_.each(a[h]||[],function(a,h){var j=h(b,c,d);return"string"!=typeof j||g||f[j]?g?!(i=j):void 0:(b.dataTypes.unshift(j),e(j),!1)}),i}var f={},g=a===vc;return e(b.dataTypes[0])||!f["*"]&&e("*")}function L(a,b){var c,d,e=_.ajaxSettings.flatOptions||{};for(c in b)void 0!==b[c]&&((e[c]?a:d||(d={}))[c]=b[c]);return d&&_.extend(!0,a,d),a}function M(a,b,c){for(var d,e,f,g,h=a.contents,i=a.dataTypes;"*"===i[0];)i.shift(),void 0===d&&(d=a.mimeType||b.getResponseHeader("Content-Type"));if(d)for(e in h)if(h[e]&&h[e].test(d)){i.unshift(e);break}if(i[0]in c)f=i[0];else{for(e in c){if(!i[0]||a.converters[e+" "+i[0]]){f=e;break}g||(g=e)}f=f||g}return f?(f!==i[0]&&i.unshift(f),c[f]):void 0}function N(a,b,c,d){var e,f,g,h,i,j={},k=a.dataTypes.slice();if(k[1])for(g in a.converters)j[g.toLowerCase()]=a.converters[g];for(f=k.shift();f;)if(a.responseFields[f]&&(c[a.responseFields[f]]=b),!i&&d&&a.dataFilter&&(b=a.dataFilter(b,a.dataType)),i=f,f=k.shift())if("*"===f)f=i;else if("*"!==i&&i!==f){if(g=j[i+" "+f]||j["* "+f],!g)for(e in j)if(h=e.split(" "),h[1]===f&&(g=j[i+" "+h[0]]||j["* "+h[0]])){g===!0?g=j[e]:j[e]!==!0&&(f=h[0],k.unshift(h[1]));break}if(g!==!0)if(g&&a["throws"])b=g(b);else try{b=g(b)}catch(l){return{state:"parsererror",error:g?l:"No conversion from "+i+" to "+f}}}return{state:"success",data:b}}function O(a,b,c,d){var e;if(_.isArray(b))_.each(b,function(b,e){c||zc.test(a)?d(a,e):O(a+"["+("object"==typeof e?b:"")+"]",e,c,d)});else if(c||"object"!==_.type(b))d(a,b);else for(e in b)O(a+"["+e+"]",b[e],c,d)}function P(a){return _.isWindow(a)?a:9===a.nodeType&&a.defaultView}var Q=[],R=Q.slice,S=Q.concat,T=Q.push,U=Q.indexOf,V={},W=V.toString,X=V.hasOwnProperty,Y={},Z=a.document,$="2.1.1",_=function(a,b){return new _.fn.init(a,b)},ab=/^[\s\uFEFF\xA0]+|[\s\uFEFF\xA0]+$/g,bb=/^-ms-/,cb=/-([\da-z])/gi,db=function(a,b){return b.toUpperCase()};_.fn=_.prototype={jquery:$,constructor:_,selector:"",length:0,toArray:function(){return R.call(this)},get:function(a){return null!=a?0>a?this[a+this.length]:this[a]:R.call(this)},pushStack:function(a){var b=_.merge(this.constructor(),a);return b.prevObject=this,b.context=this.context,b},each:function(a,b){return _.each(this,a,b)},map:function(a){return this.pushStack(_.map(this,function(b,c){return a.call(b,c,b)}))},slice:function(){return this.pushStack(R.apply(this,arguments))},first:function(){return this.eq(0)},last:function(){return this.eq(-1)},eq:function(a){var b=this.length,c=+a+(0>a?b:0);return this.pushStack(c>=0&&b>c?[this[c]]:[])},end:function(){return this.prevObject||this.constructor(null)},push:T,sort:Q.sort,splice:Q.splice},_.extend=_.fn.extend=function(){var a,b,c,d,e,f,g=arguments[0]||{},h=1,i=arguments.length,j=!1;for("boolean"==typeof g&&(j=g,g=arguments[h]||{},h++),"object"==typeof g||_.isFunction(g)||(g={}),h===i&&(g=this,h--);i>h;h++)if(null!=(a=arguments[h]))for(b in a)c=g[b],d=a[b],g!==d&&(j&&d&&(_.isPlainObject(d)||(e=_.isArray(d)))?(e?(e=!1,f=c&&_.isArray(c)?c:[]):f=c&&_.isPlainObject(c)?c:{},g[b]=_.extend(j,f,d)):void 0!==d&&(g[b]=d));return g},_.extend({expando:"jQuery"+($+Math.random()).replace(/\D/g,""),isReady:!0,error:function(a){throw new Error(a)},noop:function(){},isFunction:function(a){return"function"===_.type(a)},isArray:Array.isArray,isWindow:function(a){return null!=a&&a===a.window},isNumeric:function(a){return!_.isArray(a)&&a-parseFloat(a)>=0},isPlainObject:function(a){return"object"!==_.type(a)||a.nodeType||_.isWindow(a)?!1:a.constructor&&!X.call(a.constructor.prototype,"isPrototypeOf")?!1:!0},isEmptyObject:function(a){var b;for(b in a)return!1;return!0},type:function(a){return null==a?a+"":"object"==typeof a||"function"==typeof a?V[W.call(a)]||"object":typeof a},globalEval:function(a){var b,c=eval;a=_.trim(a),a&&(1===a.indexOf("use strict")?(b=Z.createElement("script"),b.text=a,Z.head.appendChild(b).parentNode.removeChild(b)):c(a))},camelCase:function(a){return a.replace(bb,"ms-").replace(cb,db)},nodeName:function(a,b){return a.nodeName&&a.nodeName.toLowerCase()===b.toLowerCase()},each:function(a,b,d){var e,f=0,g=a.length,h=c(a);if(d){if(h)for(;g>f&&(e=b.apply(a[f],d),e!==!1);f++);else for(f in a)if(e=b.apply(a[f],d),e===!1)break}else if(h)for(;g>f&&(e=b.call(a[f],f,a[f]),e!==!1);f++);else for(f in a)if(e=b.call(a[f],f,a[f]),e===!1)break;return a},trim:function(a){return null==a?"":(a+"").replace(ab,"")},makeArray:function(a,b){var d=b||[];return null!=a&&(c(Object(a))?_.merge(d,"string"==typeof a?[a]:a):T.call(d,a)),d},inArray:function(a,b,c){return null==b?-1:U.call(b,a,c)},merge:function(a,b){for(var c=+b.length,d=0,e=a.length;c>d;d++)a[e++]=b[d];return a.length=e,a},grep:function(a,b,c){for(var d,e=[],f=0,g=a.length,h=!c;g>f;f++)d=!b(a[f],f),d!==h&&e.push(a[f]);return e},map:function(a,b,d){var e,f=0,g=a.length,h=c(a),i=[];if(h)for(;g>f;f++)e=b(a[f],f,d),null!=e&&i.push(e);else for(f in a)e=b(a[f],f,d),null!=e&&i.push(e);return S.apply([],i)},guid:1,proxy:function(a,b){var c,d,e;return"string"==typeof b&&(c=a[b],b=a,a=c),_.isFunction(a)?(d=R.call(arguments,2),e=function(){return a.apply(b||this,d.concat(R.call(arguments)))},e.guid=a.guid=a.guid||_.guid++,e):void 0},now:Date.now,support:Y}),_.each("Boolean Number String Function Array Date RegExp Object Error".split(" "),function(a,b){V["[object "+b+"]"]=b.toLowerCase()});var eb=/*!
 * Sizzle CSS Selector Engine v1.10.19
 * http://sizzlejs.com/
 *
 * Copyright 2013 jQuery Foundation, Inc. and other contributors
 * Released under the MIT license
 * http://jquery.org/license
 *
 * Date: 2014-04-18
 */
function(a){function b(a,b,c,d){var e,f,g,h,i,j,l,n,o,p;if((b?b.ownerDocument||b:O)!==G&&F(b),b=b||G,c=c||[],!a||"string"!=typeof a)return c;if(1!==(h=b.nodeType)&&9!==h)return[];if(I&&!d){if(e=sb.exec(a))if(g=e[1]){if(9===h){if(f=b.getElementById(g),!f||!f.parentNode)return c;if(f.id===g)return c.push(f),c}else if(b.ownerDocument&&(f=b.ownerDocument.getElementById(g))&&M(b,f)&&f.id===g)return c.push(f),c}else{if(e[2])return _.apply(c,b.getElementsByTagName(a)),c;if((g=e[3])&&v.getElementsByClassName&&b.getElementsByClassName)return _.apply(c,b.getElementsByClassName(g)),c}if(v.qsa&&(!J||!J.test(a))){if(n=l=N,o=b,p=9===h&&a,1===h&&"object"!==b.nodeName.toLowerCase()){for(j=z(a),(l=b.getAttribute("id"))?n=l.replace(ub,"\\$&"):b.setAttribute("id",n),n="[id='"+n+"'] ",i=j.length;i--;)j[i]=n+m(j[i]);o=tb.test(a)&&k(b.parentNode)||b,p=j.join(",")}if(p)try{return _.apply(c,o.querySelectorAll(p)),c}catch(q){}finally{l||b.removeAttribute("id")}}}return B(a.replace(ib,"$1"),b,c,d)}function c(){function a(c,d){return b.push(c+" ")>w.cacheLength&&delete a[b.shift()],a[c+" "]=d}var b=[];return a}function d(a){return a[N]=!0,a}function e(a){var b=G.createElement("div");try{return!!a(b)}catch(c){return!1}finally{b.parentNode&&b.parentNode.removeChild(b),b=null}}function f(a,b){for(var c=a.split("|"),d=a.length;d--;)w.attrHandle[c[d]]=b}function g(a,b){var c=b&&a,d=c&&1===a.nodeType&&1===b.nodeType&&(~b.sourceIndex||W)-(~a.sourceIndex||W);if(d)return d;if(c)for(;c=c.nextSibling;)if(c===b)return-1;return a?1:-1}function h(a){return function(b){var c=b.nodeName.toLowerCase();return"input"===c&&b.type===a}}function i(a){return function(b){var c=b.nodeName.toLowerCase();return("input"===c||"button"===c)&&b.type===a}}function j(a){return d(function(b){return b=+b,d(function(c,d){for(var e,f=a([],c.length,b),g=f.length;g--;)c[e=f[g]]&&(c[e]=!(d[e]=c[e]))})})}function k(a){return a&&typeof a.getElementsByTagName!==V&&a}function l(){}function m(a){for(var b=0,c=a.length,d="";c>b;b++)d+=a[b].value;return d}function n(a,b,c){var d=b.dir,e=c&&"parentNode"===d,f=Q++;return b.first?function(b,c,f){for(;b=b[d];)if(1===b.nodeType||e)return a(b,c,f)}:function(b,c,g){var h,i,j=[P,f];if(g){for(;b=b[d];)if((1===b.nodeType||e)&&a(b,c,g))return!0}else for(;b=b[d];)if(1===b.nodeType||e){if(i=b[N]||(b[N]={}),(h=i[d])&&h[0]===P&&h[1]===f)return j[2]=h[2];if(i[d]=j,j[2]=a(b,c,g))return!0}}}function o(a){return a.length>1?function(b,c,d){for(var e=a.length;e--;)if(!a[e](b,c,d))return!1;return!0}:a[0]}function p(a,c,d){for(var e=0,f=c.length;f>e;e++)b(a,c[e],d);return d}function q(a,b,c,d,e){for(var f,g=[],h=0,i=a.length,j=null!=b;i>h;h++)(f=a[h])&&(!c||c(f,d,e))&&(g.push(f),j&&b.push(h));return g}function r(a,b,c,e,f,g){return e&&!e[N]&&(e=r(e)),f&&!f[N]&&(f=r(f,g)),d(function(d,g,h,i){var j,k,l,m=[],n=[],o=g.length,r=d||p(b||"*",h.nodeType?[h]:h,[]),s=!a||!d&&b?r:q(r,m,a,h,i),t=c?f||(d?a:o||e)?[]:g:s;if(c&&c(s,t,h,i),e)for(j=q(t,n),e(j,[],h,i),k=j.length;k--;)(l=j[k])&&(t[n[k]]=!(s[n[k]]=l));if(d){if(f||a){if(f){for(j=[],k=t.length;k--;)(l=t[k])&&j.push(s[k]=l);f(null,t=[],j,i)}for(k=t.length;k--;)(l=t[k])&&(j=f?bb.call(d,l):m[k])>-1&&(d[j]=!(g[j]=l))}}else t=q(t===g?t.splice(o,t.length):t),f?f(null,g,t,i):_.apply(g,t)})}function s(a){for(var b,c,d,e=a.length,f=w.relative[a[0].type],g=f||w.relative[" "],h=f?1:0,i=n(function(a){return a===b},g,!0),j=n(function(a){return bb.call(b,a)>-1},g,!0),k=[function(a,c,d){return!f&&(d||c!==C)||((b=c).nodeType?i(a,c,d):j(a,c,d))}];e>h;h++)if(c=w.relative[a[h].type])k=[n(o(k),c)];else{if(c=w.filter[a[h].type].apply(null,a[h].matches),c[N]){for(d=++h;e>d&&!w.relative[a[d].type];d++);return r(h>1&&o(k),h>1&&m(a.slice(0,h-1).concat({value:" "===a[h-2].type?"*":""})).replace(ib,"$1"),c,d>h&&s(a.slice(h,d)),e>d&&s(a=a.slice(d)),e>d&&m(a))}k.push(c)}return o(k)}function t(a,c){var e=c.length>0,f=a.length>0,g=function(d,g,h,i,j){var k,l,m,n=0,o="0",p=d&&[],r=[],s=C,t=d||f&&w.find.TAG("*",j),u=P+=null==s?1:Math.random()||.1,v=t.length;for(j&&(C=g!==G&&g);o!==v&&null!=(k=t[o]);o++){if(f&&k){for(l=0;m=a[l++];)if(m(k,g,h)){i.push(k);break}j&&(P=u)}e&&((k=!m&&k)&&n--,d&&p.push(k))}if(n+=o,e&&o!==n){for(l=0;m=c[l++];)m(p,r,g,h);if(d){if(n>0)for(;o--;)p[o]||r[o]||(r[o]=Z.call(i));r=q(r)}_.apply(i,r),j&&!d&&r.length>0&&n+c.length>1&&b.uniqueSort(i)}return j&&(P=u,C=s),p};return e?d(g):g}var u,v,w,x,y,z,A,B,C,D,E,F,G,H,I,J,K,L,M,N="sizzle"+-new Date,O=a.document,P=0,Q=0,R=c(),S=c(),T=c(),U=function(a,b){return a===b&&(E=!0),0},V="undefined",W=1<<31,X={}.hasOwnProperty,Y=[],Z=Y.pop,$=Y.push,_=Y.push,ab=Y.slice,bb=Y.indexOf||function(a){for(var b=0,c=this.length;c>b;b++)if(this[b]===a)return b;return-1},cb="checked|selected|async|autofocus|autoplay|controls|defer|disabled|hidden|ismap|loop|multiple|open|readonly|required|scoped",db="[\\x20\\t\\r\\n\\f]",eb="(?:\\\\.|[\\w-]|[^\\x00-\\xa0])+",fb=eb.replace("w","w#"),gb="\\["+db+"*("+eb+")(?:"+db+"*([*^$|!~]?=)"+db+"*(?:'((?:\\\\.|[^\\\\'])*)'|\"((?:\\\\.|[^\\\\\"])*)\"|("+fb+"))|)"+db+"*\\]",hb=":("+eb+")(?:\\((('((?:\\\\.|[^\\\\'])*)'|\"((?:\\\\.|[^\\\\\"])*)\")|((?:\\\\.|[^\\\\()[\\]]|"+gb+")*)|.*)\\)|)",ib=new RegExp("^"+db+"+|((?:^|[^\\\\])(?:\\\\.)*)"+db+"+$","g"),jb=new RegExp("^"+db+"*,"+db+"*"),kb=new RegExp("^"+db+"*([>+~]|"+db+")"+db+"*"),lb=new RegExp("="+db+"*([^\\]'\"]*?)"+db+"*\\]","g"),mb=new RegExp(hb),nb=new RegExp("^"+fb+"$"),ob={ID:new RegExp("^#("+eb+")"),CLASS:new RegExp("^\\.("+eb+")"),TAG:new RegExp("^("+eb.replace("w","w*")+")"),ATTR:new RegExp("^"+gb),PSEUDO:new RegExp("^"+hb),CHILD:new RegExp("^:(only|first|last|nth|nth-last)-(child|of-type)(?:\\("+db+"*(even|odd|(([+-]|)(\\d*)n|)"+db+"*(?:([+-]|)"+db+"*(\\d+)|))"+db+"*\\)|)","i"),bool:new RegExp("^(?:"+cb+")$","i"),needsContext:new RegExp("^"+db+"*[>+~]|:(even|odd|eq|gt|lt|nth|first|last)(?:\\("+db+"*((?:-\\d)?\\d*)"+db+"*\\)|)(?=[^-]|$)","i")},pb=/^(?:input|select|textarea|button)$/i,qb=/^h\d$/i,rb=/^[^{]+\{\s*\[native \w/,sb=/^(?:#([\w-]+)|(\w+)|\.([\w-]+))$/,tb=/[+~]/,ub=/'|\\/g,vb=new RegExp("\\\\([\\da-f]{1,6}"+db+"?|("+db+")|.)","ig"),wb=function(a,b,c){var d="0x"+b-65536;return d!==d||c?b:0>d?String.fromCharCode(d+65536):String.fromCharCode(d>>10|55296,1023&d|56320)};try{_.apply(Y=ab.call(O.childNodes),O.childNodes),Y[O.childNodes.length].nodeType}catch(xb){_={apply:Y.length?function(a,b){$.apply(a,ab.call(b))}:function(a,b){for(var c=a.length,d=0;a[c++]=b[d++];);a.length=c-1}}}v=b.support={},y=b.isXML=function(a){var b=a&&(a.ownerDocument||a).documentElement;return b?"HTML"!==b.nodeName:!1},F=b.setDocument=function(a){var b,c=a?a.ownerDocument||a:O,d=c.defaultView;return c!==G&&9===c.nodeType&&c.documentElement?(G=c,H=c.documentElement,I=!y(c),d&&d!==d.top&&(d.addEventListener?d.addEventListener("unload",function(){F()},!1):d.attachEvent&&d.attachEvent("onunload",function(){F()})),v.attributes=e(function(a){return a.className="i",!a.getAttribute("className")}),v.getElementsByTagName=e(function(a){return a.appendChild(c.createComment("")),!a.getElementsByTagName("*").length}),v.getElementsByClassName=rb.test(c.getElementsByClassName)&&e(function(a){return a.innerHTML="<div class='a'></div><div class='a i'></div>",a.firstChild.className="i",2===a.getElementsByClassName("i").length}),v.getById=e(function(a){return H.appendChild(a).id=N,!c.getElementsByName||!c.getElementsByName(N).length}),v.getById?(w.find.ID=function(a,b){if(typeof b.getElementById!==V&&I){var c=b.getElementById(a);return c&&c.parentNode?[c]:[]}},w.filter.ID=function(a){var b=a.replace(vb,wb);return function(a){return a.getAttribute("id")===b}}):(delete w.find.ID,w.filter.ID=function(a){var b=a.replace(vb,wb);return function(a){var c=typeof a.getAttributeNode!==V&&a.getAttributeNode("id");return c&&c.value===b}}),w.find.TAG=v.getElementsByTagName?function(a,b){return typeof b.getElementsByTagName!==V?b.getElementsByTagName(a):void 0}:function(a,b){var c,d=[],e=0,f=b.getElementsByTagName(a);if("*"===a){for(;c=f[e++];)1===c.nodeType&&d.push(c);return d}return f},w.find.CLASS=v.getElementsByClassName&&function(a,b){return typeof b.getElementsByClassName!==V&&I?b.getElementsByClassName(a):void 0},K=[],J=[],(v.qsa=rb.test(c.querySelectorAll))&&(e(function(a){a.innerHTML="<select msallowclip=''><option selected=''></option></select>",a.querySelectorAll("[msallowclip^='']").length&&J.push("[*^$]="+db+"*(?:''|\"\")"),a.querySelectorAll("[selected]").length||J.push("\\["+db+"*(?:value|"+cb+")"),a.querySelectorAll(":checked").length||J.push(":checked")}),e(function(a){var b=c.createElement("input");b.setAttribute("type","hidden"),a.appendChild(b).setAttribute("name","D"),a.querySelectorAll("[name=d]").length&&J.push("name"+db+"*[*^$|!~]?="),a.querySelectorAll(":enabled").length||J.push(":enabled",":disabled"),a.querySelectorAll("*,:x"),J.push(",.*:")})),(v.matchesSelector=rb.test(L=H.matches||H.webkitMatchesSelector||H.mozMatchesSelector||H.oMatchesSelector||H.msMatchesSelector))&&e(function(a){v.disconnectedMatch=L.call(a,"div"),L.call(a,"[s!='']:x"),K.push("!=",hb)}),J=J.length&&new RegExp(J.join("|")),K=K.length&&new RegExp(K.join("|")),b=rb.test(H.compareDocumentPosition),M=b||rb.test(H.contains)?function(a,b){var c=9===a.nodeType?a.documentElement:a,d=b&&b.parentNode;return a===d||!(!d||1!==d.nodeType||!(c.contains?c.contains(d):a.compareDocumentPosition&&16&a.compareDocumentPosition(d)))}:function(a,b){if(b)for(;b=b.parentNode;)if(b===a)return!0;return!1},U=b?function(a,b){if(a===b)return E=!0,0;var d=!a.compareDocumentPosition-!b.compareDocumentPosition;return d?d:(d=(a.ownerDocument||a)===(b.ownerDocument||b)?a.compareDocumentPosition(b):1,1&d||!v.sortDetached&&b.compareDocumentPosition(a)===d?a===c||a.ownerDocument===O&&M(O,a)?-1:b===c||b.ownerDocument===O&&M(O,b)?1:D?bb.call(D,a)-bb.call(D,b):0:4&d?-1:1)}:function(a,b){if(a===b)return E=!0,0;var d,e=0,f=a.parentNode,h=b.parentNode,i=[a],j=[b];if(!f||!h)return a===c?-1:b===c?1:f?-1:h?1:D?bb.call(D,a)-bb.call(D,b):0;if(f===h)return g(a,b);for(d=a;d=d.parentNode;)i.unshift(d);for(d=b;d=d.parentNode;)j.unshift(d);for(;i[e]===j[e];)e++;return e?g(i[e],j[e]):i[e]===O?-1:j[e]===O?1:0},c):G},b.matches=function(a,c){return b(a,null,null,c)},b.matchesSelector=function(a,c){if((a.ownerDocument||a)!==G&&F(a),c=c.replace(lb,"='$1']"),!(!v.matchesSelector||!I||K&&K.test(c)||J&&J.test(c)))try{var d=L.call(a,c);if(d||v.disconnectedMatch||a.document&&11!==a.document.nodeType)return d}catch(e){}return b(c,G,null,[a]).length>0},b.contains=function(a,b){return(a.ownerDocument||a)!==G&&F(a),M(a,b)},b.attr=function(a,b){(a.ownerDocument||a)!==G&&F(a);var c=w.attrHandle[b.toLowerCase()],d=c&&X.call(w.attrHandle,b.toLowerCase())?c(a,b,!I):void 0;return void 0!==d?d:v.attributes||!I?a.getAttribute(b):(d=a.getAttributeNode(b))&&d.specified?d.value:null},b.error=function(a){throw new Error("Syntax error, unrecognized expression: "+a)},b.uniqueSort=function(a){var b,c=[],d=0,e=0;if(E=!v.detectDuplicates,D=!v.sortStable&&a.slice(0),a.sort(U),E){for(;b=a[e++];)b===a[e]&&(d=c.push(e));for(;d--;)a.splice(c[d],1)}return D=null,a},x=b.getText=function(a){var b,c="",d=0,e=a.nodeType;if(e){if(1===e||9===e||11===e){if("string"==typeof a.textContent)return a.textContent;for(a=a.firstChild;a;a=a.nextSibling)c+=x(a)}else if(3===e||4===e)return a.nodeValue}else for(;b=a[d++];)c+=x(b);return c},w=b.selectors={cacheLength:50,createPseudo:d,match:ob,attrHandle:{},find:{},relative:{">":{dir:"parentNode",first:!0}," ":{dir:"parentNode"},"+":{dir:"previousSibling",first:!0},"~":{dir:"previousSibling"}},preFilter:{ATTR:function(a){return a[1]=a[1].replace(vb,wb),a[3]=(a[3]||a[4]||a[5]||"").replace(vb,wb),"~="===a[2]&&(a[3]=" "+a[3]+" "),a.slice(0,4)},CHILD:function(a){return a[1]=a[1].toLowerCase(),"nth"===a[1].slice(0,3)?(a[3]||b.error(a[0]),a[4]=+(a[4]?a[5]+(a[6]||1):2*("even"===a[3]||"odd"===a[3])),a[5]=+(a[7]+a[8]||"odd"===a[3])):a[3]&&b.error(a[0]),a},PSEUDO:function(a){var b,c=!a[6]&&a[2];return ob.CHILD.test(a[0])?null:(a[3]?a[2]=a[4]||a[5]||"":c&&mb.test(c)&&(b=z(c,!0))&&(b=c.indexOf(")",c.length-b)-c.length)&&(a[0]=a[0].slice(0,b),a[2]=c.slice(0,b)),a.slice(0,3))}},filter:{TAG:function(a){var b=a.replace(vb,wb).toLowerCase();return"*"===a?function(){return!0}:function(a){return a.nodeName&&a.nodeName.toLowerCase()===b}},CLASS:function(a){var b=R[a+" "];return b||(b=new RegExp("(^|"+db+")"+a+"("+db+"|$)"))&&R(a,function(a){return b.test("string"==typeof a.className&&a.className||typeof a.getAttribute!==V&&a.getAttribute("class")||"")})},ATTR:function(a,c,d){return function(e){var f=b.attr(e,a);return null==f?"!="===c:c?(f+="","="===c?f===d:"!="===c?f!==d:"^="===c?d&&0===f.indexOf(d):"*="===c?d&&f.indexOf(d)>-1:"$="===c?d&&f.slice(-d.length)===d:"~="===c?(" "+f+" ").indexOf(d)>-1:"|="===c?f===d||f.slice(0,d.length+1)===d+"-":!1):!0}},CHILD:function(a,b,c,d,e){var f="nth"!==a.slice(0,3),g="last"!==a.slice(-4),h="of-type"===b;return 1===d&&0===e?function(a){return!!a.parentNode}:function(b,c,i){var j,k,l,m,n,o,p=f!==g?"nextSibling":"previousSibling",q=b.parentNode,r=h&&b.nodeName.toLowerCase(),s=!i&&!h;if(q){if(f){for(;p;){for(l=b;l=l[p];)if(h?l.nodeName.toLowerCase()===r:1===l.nodeType)return!1;o=p="only"===a&&!o&&"nextSibling"}return!0}if(o=[g?q.firstChild:q.lastChild],g&&s){for(k=q[N]||(q[N]={}),j=k[a]||[],n=j[0]===P&&j[1],m=j[0]===P&&j[2],l=n&&q.childNodes[n];l=++n&&l&&l[p]||(m=n=0)||o.pop();)if(1===l.nodeType&&++m&&l===b){k[a]=[P,n,m];break}}else if(s&&(j=(b[N]||(b[N]={}))[a])&&j[0]===P)m=j[1];else for(;(l=++n&&l&&l[p]||(m=n=0)||o.pop())&&((h?l.nodeName.toLowerCase()!==r:1!==l.nodeType)||!++m||(s&&((l[N]||(l[N]={}))[a]=[P,m]),l!==b)););return m-=e,m===d||m%d===0&&m/d>=0}}},PSEUDO:function(a,c){var e,f=w.pseudos[a]||w.setFilters[a.toLowerCase()]||b.error("unsupported pseudo: "+a);return f[N]?f(c):f.length>1?(e=[a,a,"",c],w.setFilters.hasOwnProperty(a.toLowerCase())?d(function(a,b){for(var d,e=f(a,c),g=e.length;g--;)d=bb.call(a,e[g]),a[d]=!(b[d]=e[g])}):function(a){return f(a,0,e)}):f}},pseudos:{not:d(function(a){var b=[],c=[],e=A(a.replace(ib,"$1"));return e[N]?d(function(a,b,c,d){for(var f,g=e(a,null,d,[]),h=a.length;h--;)(f=g[h])&&(a[h]=!(b[h]=f))}):function(a,d,f){return b[0]=a,e(b,null,f,c),!c.pop()}}),has:d(function(a){return function(c){return b(a,c).length>0}}),contains:d(function(a){return function(b){return(b.textContent||b.innerText||x(b)).indexOf(a)>-1}}),lang:d(function(a){return nb.test(a||"")||b.error("unsupported lang: "+a),a=a.replace(vb,wb).toLowerCase(),function(b){var c;do if(c=I?b.lang:b.getAttribute("xml:lang")||b.getAttribute("lang"))return c=c.toLowerCase(),c===a||0===c.indexOf(a+"-");while((b=b.parentNode)&&1===b.nodeType);return!1}}),target:function(b){var c=a.location&&a.location.hash;return c&&c.slice(1)===b.id},root:function(a){return a===H},focus:function(a){return a===G.activeElement&&(!G.hasFocus||G.hasFocus())&&!!(a.type||a.href||~a.tabIndex)},enabled:function(a){return a.disabled===!1},disabled:function(a){return a.disabled===!0},checked:function(a){var b=a.nodeName.toLowerCase();return"input"===b&&!!a.checked||"option"===b&&!!a.selected},selected:function(a){return a.parentNode&&a.parentNode.selectedIndex,a.selected===!0},empty:function(a){for(a=a.firstChild;a;a=a.nextSibling)if(a.nodeType<6)return!1;return!0},parent:function(a){return!w.pseudos.empty(a)},header:function(a){return qb.test(a.nodeName)},input:function(a){return pb.test(a.nodeName)},button:function(a){var b=a.nodeName.toLowerCase();return"input"===b&&"button"===a.type||"button"===b},text:function(a){var b;return"input"===a.nodeName.toLowerCase()&&"text"===a.type&&(null==(b=a.getAttribute("type"))||"text"===b.toLowerCase())},first:j(function(){return[0]}),last:j(function(a,b){return[b-1]}),eq:j(function(a,b,c){return[0>c?c+b:c]}),even:j(function(a,b){for(var c=0;b>c;c+=2)a.push(c);return a}),odd:j(function(a,b){for(var c=1;b>c;c+=2)a.push(c);return a}),lt:j(function(a,b,c){for(var d=0>c?c+b:c;--d>=0;)a.push(d);return a}),gt:j(function(a,b,c){for(var d=0>c?c+b:c;++d<b;)a.push(d);return a})}},w.pseudos.nth=w.pseudos.eq;for(u in{radio:!0,checkbox:!0,file:!0,password:!0,image:!0})w.pseudos[u]=h(u);for(u in{submit:!0,reset:!0})w.pseudos[u]=i(u);return l.prototype=w.filters=w.pseudos,w.setFilters=new l,z=b.tokenize=function(a,c){var d,e,f,g,h,i,j,k=S[a+" "];if(k)return c?0:k.slice(0);for(h=a,i=[],j=w.preFilter;h;){(!d||(e=jb.exec(h)))&&(e&&(h=h.slice(e[0].length)||h),i.push(f=[])),d=!1,(e=kb.exec(h))&&(d=e.shift(),f.push({value:d,type:e[0].replace(ib," ")}),h=h.slice(d.length));for(g in w.filter)!(e=ob[g].exec(h))||j[g]&&!(e=j[g](e))||(d=e.shift(),f.push({value:d,type:g,matches:e}),h=h.slice(d.length));if(!d)break}return c?h.length:h?b.error(a):S(a,i).slice(0)},A=b.compile=function(a,b){var c,d=[],e=[],f=T[a+" "];if(!f){for(b||(b=z(a)),c=b.length;c--;)f=s(b[c]),f[N]?d.push(f):e.push(f);f=T(a,t(e,d)),f.selector=a}return f},B=b.select=function(a,b,c,d){var e,f,g,h,i,j="function"==typeof a&&a,l=!d&&z(a=j.selector||a);if(c=c||[],1===l.length){if(f=l[0]=l[0].slice(0),f.length>2&&"ID"===(g=f[0]).type&&v.getById&&9===b.nodeType&&I&&w.relative[f[1].type]){if(b=(w.find.ID(g.matches[0].replace(vb,wb),b)||[])[0],!b)return c;j&&(b=b.parentNode),a=a.slice(f.shift().value.length)}for(e=ob.needsContext.test(a)?0:f.length;e--&&(g=f[e],!w.relative[h=g.type]);)if((i=w.find[h])&&(d=i(g.matches[0].replace(vb,wb),tb.test(f[0].type)&&k(b.parentNode)||b))){if(f.splice(e,1),a=d.length&&m(f),!a)return _.apply(c,d),c;break}}return(j||A(a,l))(d,b,!I,c,tb.test(a)&&k(b.parentNode)||b),c},v.sortStable=N.split("").sort(U).join("")===N,v.detectDuplicates=!!E,F(),v.sortDetached=e(function(a){return 1&a.compareDocumentPosition(G.createElement("div"))}),e(function(a){return a.innerHTML="<a href='#'></a>","#"===a.firstChild.getAttribute("href")})||f("type|href|height|width",function(a,b,c){return c?void 0:a.getAttribute(b,"type"===b.toLowerCase()?1:2)}),v.attributes&&e(function(a){return a.innerHTML="<input/>",a.firstChild.setAttribute("value",""),""===a.firstChild.getAttribute("value")})||f("value",function(a,b,c){return c||"input"!==a.nodeName.toLowerCase()?void 0:a.defaultValue}),e(function(a){return null==a.getAttribute("disabled")})||f(cb,function(a,b,c){var d;return c?void 0:a[b]===!0?b.toLowerCase():(d=a.getAttributeNode(b))&&d.specified?d.value:null}),b}(a);_.find=eb,_.expr=eb.selectors,_.expr[":"]=_.expr.pseudos,_.unique=eb.uniqueSort,_.text=eb.getText,_.isXMLDoc=eb.isXML,_.contains=eb.contains;var fb=_.expr.match.needsContext,gb=/^<(\w+)\s*\/?>(?:<\/\1>|)$/,hb=/^.[^:#\[\.,]*$/;_.filter=function(a,b,c){var d=b[0];return c&&(a=":not("+a+")"),1===b.length&&1===d.nodeType?_.find.matchesSelector(d,a)?[d]:[]:_.find.matches(a,_.grep(b,function(a){return 1===a.nodeType}))},_.fn.extend({find:function(a){var b,c=this.length,d=[],e=this;if("string"!=typeof a)return this.pushStack(_(a).filter(function(){for(b=0;c>b;b++)if(_.contains(e[b],this))return!0}));for(b=0;c>b;b++)_.find(a,e[b],d);return d=this.pushStack(c>1?_.unique(d):d),d.selector=this.selector?this.selector+" "+a:a,d},filter:function(a){return this.pushStack(d(this,a||[],!1))},not:function(a){return this.pushStack(d(this,a||[],!0))},is:function(a){return!!d(this,"string"==typeof a&&fb.test(a)?_(a):a||[],!1).length}});var ib,jb=/^(?:\s*(<[\w\W]+>)[^>]*|#([\w-]*))$/,kb=_.fn.init=function(a,b){var c,d;if(!a)return this;if("string"==typeof a){if(c="<"===a[0]&&">"===a[a.length-1]&&a.length>=3?[null,a,null]:jb.exec(a),!c||!c[1]&&b)return!b||b.jquery?(b||ib).find(a):this.constructor(b).find(a);if(c[1]){if(b=b instanceof _?b[0]:b,_.merge(this,_.parseHTML(c[1],b&&b.nodeType?b.ownerDocument||b:Z,!0)),gb.test(c[1])&&_.isPlainObject(b))for(c in b)_.isFunction(this[c])?this[c](b[c]):this.attr(c,b[c]);return this}return d=Z.getElementById(c[2]),d&&d.parentNode&&(this.length=1,this[0]=d),this.context=Z,this.selector=a,this}return a.nodeType?(this.context=this[0]=a,this.length=1,this):_.isFunction(a)?"undefined"!=typeof ib.ready?ib.ready(a):a(_):(void 0!==a.selector&&(this.selector=a.selector,this.context=a.context),_.makeArray(a,this))};kb.prototype=_.fn,ib=_(Z);var lb=/^(?:parents|prev(?:Until|All))/,mb={children:!0,contents:!0,next:!0,prev:!0};_.extend({dir:function(a,b,c){for(var d=[],e=void 0!==c;(a=a[b])&&9!==a.nodeType;)if(1===a.nodeType){if(e&&_(a).is(c))break;d.push(a)}return d},sibling:function(a,b){for(var c=[];a;a=a.nextSibling)1===a.nodeType&&a!==b&&c.push(a);return c}}),_.fn.extend({has:function(a){var b=_(a,this),c=b.length;return this.filter(function(){for(var a=0;c>a;a++)if(_.contains(this,b[a]))return!0})},closest:function(a,b){for(var c,d=0,e=this.length,f=[],g=fb.test(a)||"string"!=typeof a?_(a,b||this.context):0;e>d;d++)for(c=this[d];c&&c!==b;c=c.parentNode)if(c.nodeType<11&&(g?g.index(c)>-1:1===c.nodeType&&_.find.matchesSelector(c,a))){f.push(c);break}return this.pushStack(f.length>1?_.unique(f):f)},index:function(a){return a?"string"==typeof a?U.call(_(a),this[0]):U.call(this,a.jquery?a[0]:a):this[0]&&this[0].parentNode?this.first().prevAll().length:-1},add:function(a,b){return this.pushStack(_.unique(_.merge(this.get(),_(a,b))))},addBack:function(a){return this.add(null==a?this.prevObject:this.prevObject.filter(a))}}),_.each({parent:function(a){var b=a.parentNode;return b&&11!==b.nodeType?b:null},parents:function(a){return _.dir(a,"parentNode")},parentsUntil:function(a,b,c){return _.dir(a,"parentNode",c)},next:function(a){return e(a,"nextSibling")},prev:function(a){return e(a,"previousSibling")},nextAll:function(a){return _.dir(a,"nextSibling")},prevAll:function(a){return _.dir(a,"previousSibling")},nextUntil:function(a,b,c){return _.dir(a,"nextSibling",c)},prevUntil:function(a,b,c){return _.dir(a,"previousSibling",c)},siblings:function(a){return _.sibling((a.parentNode||{}).firstChild,a)},children:function(a){return _.sibling(a.firstChild)},contents:function(a){return a.contentDocument||_.merge([],a.childNodes)}},function(a,b){_.fn[a]=function(c,d){var e=_.map(this,b,c);return"Until"!==a.slice(-5)&&(d=c),d&&"string"==typeof d&&(e=_.filter(d,e)),this.length>1&&(mb[a]||_.unique(e),lb.test(a)&&e.reverse()),this.pushStack(e)}});var nb=/\S+/g,ob={};_.Callbacks=function(a){a="string"==typeof a?ob[a]||f(a):_.extend({},a);var b,c,d,e,g,h,i=[],j=!a.once&&[],k=function(f){for(b=a.memory&&f,c=!0,h=e||0,e=0,g=i.length,d=!0;i&&g>h;h++)if(i[h].apply(f[0],f[1])===!1&&a.stopOnFalse){b=!1;break}d=!1,i&&(j?j.length&&k(j.shift()):b?i=[]:l.disable())},l={add:function(){if(i){var c=i.length;!function f(b){_.each(b,function(b,c){var d=_.type(c);"function"===d?a.unique&&l.has(c)||i.push(c):c&&c.length&&"string"!==d&&f(c)})}(arguments),d?g=i.length:b&&(e=c,k(b))}return this},remove:function(){return i&&_.each(arguments,function(a,b){for(var c;(c=_.inArray(b,i,c))>-1;)i.splice(c,1),d&&(g>=c&&g--,h>=c&&h--)}),this},has:function(a){return a?_.inArray(a,i)>-1:!(!i||!i.length)},empty:function(){return i=[],g=0,this},disable:function(){return i=j=b=void 0,this},disabled:function(){return!i},lock:function(){return j=void 0,b||l.disable(),this},locked:function(){return!j},fireWith:function(a,b){return!i||c&&!j||(b=b||[],b=[a,b.slice?b.slice():b],d?j.push(b):k(b)),this},fire:function(){return l.fireWith(this,arguments),this},fired:function(){return!!c}};return l},_.extend({Deferred:function(a){var b=[["resolve","done",_.Callbacks("once memory"),"resolved"],["reject","fail",_.Callbacks("once memory"),"rejected"],["notify","progress",_.Callbacks("memory")]],c="pending",d={state:function(){return c},always:function(){return e.done(arguments).fail(arguments),this},then:function(){var a=arguments;return _.Deferred(function(c){_.each(b,function(b,f){var g=_.isFunction(a[b])&&a[b];e[f[1]](function(){var a=g&&g.apply(this,arguments);a&&_.isFunction(a.promise)?a.promise().done(c.resolve).fail(c.reject).progress(c.notify):c[f[0]+"With"](this===d?c.promise():this,g?[a]:arguments)})}),a=null}).promise()},promise:function(a){return null!=a?_.extend(a,d):d}},e={};return d.pipe=d.then,_.each(b,function(a,f){var g=f[2],h=f[3];d[f[1]]=g.add,h&&g.add(function(){c=h},b[1^a][2].disable,b[2][2].lock),e[f[0]]=function(){return e[f[0]+"With"](this===e?d:this,arguments),this},e[f[0]+"With"]=g.fireWith}),d.promise(e),a&&a.call(e,e),e},when:function(a){var b,c,d,e=0,f=R.call(arguments),g=f.length,h=1!==g||a&&_.isFunction(a.promise)?g:0,i=1===h?a:_.Deferred(),j=function(a,c,d){return function(e){c[a]=this,d[a]=arguments.length>1?R.call(arguments):e,d===b?i.notifyWith(c,d):--h||i.resolveWith(c,d)}};if(g>1)for(b=new Array(g),c=new Array(g),d=new Array(g);g>e;e++)f[e]&&_.isFunction(f[e].promise)?f[e].promise().done(j(e,d,f)).fail(i.reject).progress(j(e,c,b)):--h;return h||i.resolveWith(d,f),i.promise()}});var pb;_.fn.ready=function(a){return _.ready.promise().done(a),this},_.extend({isReady:!1,readyWait:1,holdReady:function(a){a?_.readyWait++:_.ready(!0)},ready:function(a){(a===!0?--_.readyWait:_.isReady)||(_.isReady=!0,a!==!0&&--_.readyWait>0||(pb.resolveWith(Z,[_]),_.fn.triggerHandler&&(_(Z).triggerHandler("ready"),_(Z).off("ready"))))}}),_.ready.promise=function(b){return pb||(pb=_.Deferred(),"complete"===Z.readyState?setTimeout(_.ready):(Z.addEventListener("DOMContentLoaded",g,!1),a.addEventListener("load",g,!1))),pb.promise(b)},_.ready.promise();var qb=_.access=function(a,b,c,d,e,f,g){var h=0,i=a.length,j=null==c;if("object"===_.type(c)){e=!0;for(h in c)_.access(a,b,h,c[h],!0,f,g)}else if(void 0!==d&&(e=!0,_.isFunction(d)||(g=!0),j&&(g?(b.call(a,d),b=null):(j=b,b=function(a,b,c){return j.call(_(a),c)})),b))for(;i>h;h++)b(a[h],c,g?d:d.call(a[h],h,b(a[h],c)));return e?a:j?b.call(a):i?b(a[0],c):f};_.acceptData=function(a){return 1===a.nodeType||9===a.nodeType||!+a.nodeType},h.uid=1,h.accepts=_.acceptData,h.prototype={key:function(a){if(!h.accepts(a))return 0;var b={},c=a[this.expando];if(!c){c=h.uid++;try{b[this.expando]={value:c},Object.defineProperties(a,b)}catch(d){b[this.expando]=c,_.extend(a,b)}}return this.cache[c]||(this.cache[c]={}),c},set:function(a,b,c){var d,e=this.key(a),f=this.cache[e];if("string"==typeof b)f[b]=c;else if(_.isEmptyObject(f))_.extend(this.cache[e],b);else for(d in b)f[d]=b[d];return f},get:function(a,b){var c=this.cache[this.key(a)];return void 0===b?c:c[b]},access:function(a,b,c){var d;return void 0===b||b&&"string"==typeof b&&void 0===c?(d=this.get(a,b),void 0!==d?d:this.get(a,_.camelCase(b))):(this.set(a,b,c),void 0!==c?c:b)},remove:function(a,b){var c,d,e,f=this.key(a),g=this.cache[f];if(void 0===b)this.cache[f]={};else{_.isArray(b)?d=b.concat(b.map(_.camelCase)):(e=_.camelCase(b),b in g?d=[b,e]:(d=e,d=d in g?[d]:d.match(nb)||[])),c=d.length;for(;c--;)delete g[d[c]]}},hasData:function(a){return!_.isEmptyObject(this.cache[a[this.expando]]||{})},discard:function(a){a[this.expando]&&delete this.cache[a[this.expando]]}};var rb=new h,sb=new h,tb=/^(?:\{[\w\W]*\}|\[[\w\W]*\])$/,ub=/([A-Z])/g;_.extend({hasData:function(a){return sb.hasData(a)||rb.hasData(a)},data:function(a,b,c){return sb.access(a,b,c)},removeData:function(a,b){sb.remove(a,b)},_data:function(a,b,c){return rb.access(a,b,c)},_removeData:function(a,b){rb.remove(a,b)}}),_.fn.extend({data:function(a,b){var c,d,e,f=this[0],g=f&&f.attributes;if(void 0===a){if(this.length&&(e=sb.get(f),1===f.nodeType&&!rb.get(f,"hasDataAttrs"))){for(c=g.length;c--;)g[c]&&(d=g[c].name,0===d.indexOf("data-")&&(d=_.camelCase(d.slice(5)),i(f,d,e[d])));rb.set(f,"hasDataAttrs",!0)}return e}return"object"==typeof a?this.each(function(){sb.set(this,a)}):qb(this,function(b){var c,d=_.camelCase(a);if(f&&void 0===b){if(c=sb.get(f,a),void 0!==c)return c;if(c=sb.get(f,d),void 0!==c)return c;if(c=i(f,d,void 0),void 0!==c)return c}else this.each(function(){var c=sb.get(this,d);sb.set(this,d,b),-1!==a.indexOf("-")&&void 0!==c&&sb.set(this,a,b)})},null,b,arguments.length>1,null,!0)},removeData:function(a){return this.each(function(){sb.remove(this,a)})}}),_.extend({queue:function(a,b,c){var d;return a?(b=(b||"fx")+"queue",d=rb.get(a,b),c&&(!d||_.isArray(c)?d=rb.access(a,b,_.makeArray(c)):d.push(c)),d||[]):void 0},dequeue:function(a,b){b=b||"fx";var c=_.queue(a,b),d=c.length,e=c.shift(),f=_._queueHooks(a,b),g=function(){_.dequeue(a,b)};"inprogress"===e&&(e=c.shift(),d--),e&&("fx"===b&&c.unshift("inprogress"),delete f.stop,e.call(a,g,f)),!d&&f&&f.empty.fire()},_queueHooks:function(a,b){var c=b+"queueHooks";return rb.get(a,c)||rb.access(a,c,{empty:_.Callbacks("once memory").add(function(){rb.remove(a,[b+"queue",c])})})}}),_.fn.extend({queue:function(a,b){var c=2;return"string"!=typeof a&&(b=a,a="fx",c--),arguments.length<c?_.queue(this[0],a):void 0===b?this:this.each(function(){var c=_.queue(this,a,b);_._queueHooks(this,a),"fx"===a&&"inprogress"!==c[0]&&_.dequeue(this,a)})},dequeue:function(a){return this.each(function(){_.dequeue(this,a)})},clearQueue:function(a){return this.queue(a||"fx",[])},promise:function(a,b){var c,d=1,e=_.Deferred(),f=this,g=this.length,h=function(){--d||e.resolveWith(f,[f])};for("string"!=typeof a&&(b=a,a=void 0),a=a||"fx";g--;)c=rb.get(f[g],a+"queueHooks"),c&&c.empty&&(d++,c.empty.add(h));return h(),e.promise(b)}});var vb=/[+-]?(?:\d*\.|)\d+(?:[eE][+-]?\d+|)/.source,wb=["Top","Right","Bottom","Left"],xb=function(a,b){return a=b||a,"none"===_.css(a,"display")||!_.contains(a.ownerDocument,a)},yb=/^(?:checkbox|radio)$/i;!function(){var a=Z.createDocumentFragment(),b=a.appendChild(Z.createElement("div")),c=Z.createElement("input");c.setAttribute("type","radio"),c.setAttribute("checked","checked"),c.setAttribute("name","t"),b.appendChild(c),Y.checkClone=b.cloneNode(!0).cloneNode(!0).lastChild.checked,b.innerHTML="<textarea>x</textarea>",Y.noCloneChecked=!!b.cloneNode(!0).lastChild.defaultValue}();var zb="undefined";Y.focusinBubbles="onfocusin"in a;var Ab=/^key/,Bb=/^(?:mouse|pointer|contextmenu)|click/,Cb=/^(?:focusinfocus|focusoutblur)$/,Db=/^([^.]*)(?:\.(.+)|)$/;_.event={global:{},add:function(a,b,c,d,e){var f,g,h,i,j,k,l,m,n,o,p,q=rb.get(a);if(q)for(c.handler&&(f=c,c=f.handler,e=f.selector),c.guid||(c.guid=_.guid++),(i=q.events)||(i=q.events={}),(g=q.handle)||(g=q.handle=function(b){return typeof _!==zb&&_.event.triggered!==b.type?_.event.dispatch.apply(a,arguments):void 0}),b=(b||"").match(nb)||[""],j=b.length;j--;)h=Db.exec(b[j])||[],n=p=h[1],o=(h[2]||"").split(".").sort(),n&&(l=_.event.special[n]||{},n=(e?l.delegateType:l.bindType)||n,l=_.event.special[n]||{},k=_.extend({type:n,origType:p,data:d,handler:c,guid:c.guid,selector:e,needsContext:e&&_.expr.match.needsContext.test(e),namespace:o.join(".")},f),(m=i[n])||(m=i[n]=[],m.delegateCount=0,l.setup&&l.setup.call(a,d,o,g)!==!1||a.addEventListener&&a.addEventListener(n,g,!1)),l.add&&(l.add.call(a,k),k.handler.guid||(k.handler.guid=c.guid)),e?m.splice(m.delegateCount++,0,k):m.push(k),_.event.global[n]=!0)},remove:function(a,b,c,d,e){var f,g,h,i,j,k,l,m,n,o,p,q=rb.hasData(a)&&rb.get(a);if(q&&(i=q.events)){for(b=(b||"").match(nb)||[""],j=b.length;j--;)if(h=Db.exec(b[j])||[],n=p=h[1],o=(h[2]||"").split(".").sort(),n){for(l=_.event.special[n]||{},n=(d?l.delegateType:l.bindType)||n,m=i[n]||[],h=h[2]&&new RegExp("(^|\\.)"+o.join("\\.(?:.*\\.|)")+"(\\.|$)"),g=f=m.length;f--;)k=m[f],!e&&p!==k.origType||c&&c.guid!==k.guid||h&&!h.test(k.namespace)||d&&d!==k.selector&&("**"!==d||!k.selector)||(m.splice(f,1),k.selector&&m.delegateCount--,l.remove&&l.remove.call(a,k));g&&!m.length&&(l.teardown&&l.teardown.call(a,o,q.handle)!==!1||_.removeEvent(a,n,q.handle),delete i[n])}else for(n in i)_.event.remove(a,n+b[j],c,d,!0);_.isEmptyObject(i)&&(delete q.handle,rb.remove(a,"events"))}},trigger:function(b,c,d,e){var f,g,h,i,j,k,l,m=[d||Z],n=X.call(b,"type")?b.type:b,o=X.call(b,"namespace")?b.namespace.split("."):[];if(g=h=d=d||Z,3!==d.nodeType&&8!==d.nodeType&&!Cb.test(n+_.event.triggered)&&(n.indexOf(".")>=0&&(o=n.split("."),n=o.shift(),o.sort()),j=n.indexOf(":")<0&&"on"+n,b=b[_.expando]?b:new _.Event(n,"object"==typeof b&&b),b.isTrigger=e?2:3,b.namespace=o.join("."),b.namespace_re=b.namespace?new RegExp("(^|\\.)"+o.join("\\.(?:.*\\.|)")+"(\\.|$)"):null,b.result=void 0,b.target||(b.target=d),c=null==c?[b]:_.makeArray(c,[b]),l=_.event.special[n]||{},e||!l.trigger||l.trigger.apply(d,c)!==!1)){if(!e&&!l.noBubble&&!_.isWindow(d)){for(i=l.delegateType||n,Cb.test(i+n)||(g=g.parentNode);g;g=g.parentNode)m.push(g),h=g;
h===(d.ownerDocument||Z)&&m.push(h.defaultView||h.parentWindow||a)}for(f=0;(g=m[f++])&&!b.isPropagationStopped();)b.type=f>1?i:l.bindType||n,k=(rb.get(g,"events")||{})[b.type]&&rb.get(g,"handle"),k&&k.apply(g,c),k=j&&g[j],k&&k.apply&&_.acceptData(g)&&(b.result=k.apply(g,c),b.result===!1&&b.preventDefault());return b.type=n,e||b.isDefaultPrevented()||l._default&&l._default.apply(m.pop(),c)!==!1||!_.acceptData(d)||j&&_.isFunction(d[n])&&!_.isWindow(d)&&(h=d[j],h&&(d[j]=null),_.event.triggered=n,d[n](),_.event.triggered=void 0,h&&(d[j]=h)),b.result}},dispatch:function(a){a=_.event.fix(a);var b,c,d,e,f,g=[],h=R.call(arguments),i=(rb.get(this,"events")||{})[a.type]||[],j=_.event.special[a.type]||{};if(h[0]=a,a.delegateTarget=this,!j.preDispatch||j.preDispatch.call(this,a)!==!1){for(g=_.event.handlers.call(this,a,i),b=0;(e=g[b++])&&!a.isPropagationStopped();)for(a.currentTarget=e.elem,c=0;(f=e.handlers[c++])&&!a.isImmediatePropagationStopped();)(!a.namespace_re||a.namespace_re.test(f.namespace))&&(a.handleObj=f,a.data=f.data,d=((_.event.special[f.origType]||{}).handle||f.handler).apply(e.elem,h),void 0!==d&&(a.result=d)===!1&&(a.preventDefault(),a.stopPropagation()));return j.postDispatch&&j.postDispatch.call(this,a),a.result}},handlers:function(a,b){var c,d,e,f,g=[],h=b.delegateCount,i=a.target;if(h&&i.nodeType&&(!a.button||"click"!==a.type))for(;i!==this;i=i.parentNode||this)if(i.disabled!==!0||"click"!==a.type){for(d=[],c=0;h>c;c++)f=b[c],e=f.selector+" ",void 0===d[e]&&(d[e]=f.needsContext?_(e,this).index(i)>=0:_.find(e,this,null,[i]).length),d[e]&&d.push(f);d.length&&g.push({elem:i,handlers:d})}return h<b.length&&g.push({elem:this,handlers:b.slice(h)}),g},props:"altKey bubbles cancelable ctrlKey currentTarget eventPhase metaKey relatedTarget shiftKey target timeStamp view which".split(" "),fixHooks:{},keyHooks:{props:"char charCode key keyCode".split(" "),filter:function(a,b){return null==a.which&&(a.which=null!=b.charCode?b.charCode:b.keyCode),a}},mouseHooks:{props:"button buttons clientX clientY offsetX offsetY pageX pageY screenX screenY toElement".split(" "),filter:function(a,b){var c,d,e,f=b.button;return null==a.pageX&&null!=b.clientX&&(c=a.target.ownerDocument||Z,d=c.documentElement,e=c.body,a.pageX=b.clientX+(d&&d.scrollLeft||e&&e.scrollLeft||0)-(d&&d.clientLeft||e&&e.clientLeft||0),a.pageY=b.clientY+(d&&d.scrollTop||e&&e.scrollTop||0)-(d&&d.clientTop||e&&e.clientTop||0)),a.which||void 0===f||(a.which=1&f?1:2&f?3:4&f?2:0),a}},fix:function(a){if(a[_.expando])return a;var b,c,d,e=a.type,f=a,g=this.fixHooks[e];for(g||(this.fixHooks[e]=g=Bb.test(e)?this.mouseHooks:Ab.test(e)?this.keyHooks:{}),d=g.props?this.props.concat(g.props):this.props,a=new _.Event(f),b=d.length;b--;)c=d[b],a[c]=f[c];return a.target||(a.target=Z),3===a.target.nodeType&&(a.target=a.target.parentNode),g.filter?g.filter(a,f):a},special:{load:{noBubble:!0},focus:{trigger:function(){return this!==l()&&this.focus?(this.focus(),!1):void 0},delegateType:"focusin"},blur:{trigger:function(){return this===l()&&this.blur?(this.blur(),!1):void 0},delegateType:"focusout"},click:{trigger:function(){return"checkbox"===this.type&&this.click&&_.nodeName(this,"input")?(this.click(),!1):void 0},_default:function(a){return _.nodeName(a.target,"a")}},beforeunload:{postDispatch:function(a){void 0!==a.result&&a.originalEvent&&(a.originalEvent.returnValue=a.result)}}},simulate:function(a,b,c,d){var e=_.extend(new _.Event,c,{type:a,isSimulated:!0,originalEvent:{}});d?_.event.trigger(e,null,b):_.event.dispatch.call(b,e),e.isDefaultPrevented()&&c.preventDefault()}},_.removeEvent=function(a,b,c){a.removeEventListener&&a.removeEventListener(b,c,!1)},_.Event=function(a,b){return this instanceof _.Event?(a&&a.type?(this.originalEvent=a,this.type=a.type,this.isDefaultPrevented=a.defaultPrevented||void 0===a.defaultPrevented&&a.returnValue===!1?j:k):this.type=a,b&&_.extend(this,b),this.timeStamp=a&&a.timeStamp||_.now(),void(this[_.expando]=!0)):new _.Event(a,b)},_.Event.prototype={isDefaultPrevented:k,isPropagationStopped:k,isImmediatePropagationStopped:k,preventDefault:function(){var a=this.originalEvent;this.isDefaultPrevented=j,a&&a.preventDefault&&a.preventDefault()},stopPropagation:function(){var a=this.originalEvent;this.isPropagationStopped=j,a&&a.stopPropagation&&a.stopPropagation()},stopImmediatePropagation:function(){var a=this.originalEvent;this.isImmediatePropagationStopped=j,a&&a.stopImmediatePropagation&&a.stopImmediatePropagation(),this.stopPropagation()}},_.each({mouseenter:"mouseover",mouseleave:"mouseout",pointerenter:"pointerover",pointerleave:"pointerout"},function(a,b){_.event.special[a]={delegateType:b,bindType:b,handle:function(a){var c,d=this,e=a.relatedTarget,f=a.handleObj;return(!e||e!==d&&!_.contains(d,e))&&(a.type=f.origType,c=f.handler.apply(this,arguments),a.type=b),c}}}),Y.focusinBubbles||_.each({focus:"focusin",blur:"focusout"},function(a,b){var c=function(a){_.event.simulate(b,a.target,_.event.fix(a),!0)};_.event.special[b]={setup:function(){var d=this.ownerDocument||this,e=rb.access(d,b);e||d.addEventListener(a,c,!0),rb.access(d,b,(e||0)+1)},teardown:function(){var d=this.ownerDocument||this,e=rb.access(d,b)-1;e?rb.access(d,b,e):(d.removeEventListener(a,c,!0),rb.remove(d,b))}}}),_.fn.extend({on:function(a,b,c,d,e){var f,g;if("object"==typeof a){"string"!=typeof b&&(c=c||b,b=void 0);for(g in a)this.on(g,b,c,a[g],e);return this}if(null==c&&null==d?(d=b,c=b=void 0):null==d&&("string"==typeof b?(d=c,c=void 0):(d=c,c=b,b=void 0)),d===!1)d=k;else if(!d)return this;return 1===e&&(f=d,d=function(a){return _().off(a),f.apply(this,arguments)},d.guid=f.guid||(f.guid=_.guid++)),this.each(function(){_.event.add(this,a,d,c,b)})},one:function(a,b,c,d){return this.on(a,b,c,d,1)},off:function(a,b,c){var d,e;if(a&&a.preventDefault&&a.handleObj)return d=a.handleObj,_(a.delegateTarget).off(d.namespace?d.origType+"."+d.namespace:d.origType,d.selector,d.handler),this;if("object"==typeof a){for(e in a)this.off(e,b,a[e]);return this}return(b===!1||"function"==typeof b)&&(c=b,b=void 0),c===!1&&(c=k),this.each(function(){_.event.remove(this,a,c,b)})},trigger:function(a,b){return this.each(function(){_.event.trigger(a,b,this)})},triggerHandler:function(a,b){var c=this[0];return c?_.event.trigger(a,b,c,!0):void 0}});var Eb=/<(?!area|br|col|embed|hr|img|input|link|meta|param)(([\w:]+)[^>]*)\/>/gi,Fb=/<([\w:]+)/,Gb=/<|&#?\w+;/,Hb=/<(?:script|style|link)/i,Ib=/checked\s*(?:[^=]|=\s*.checked.)/i,Jb=/^$|\/(?:java|ecma)script/i,Kb=/^true\/(.*)/,Lb=/^\s*<!(?:\[CDATA\[|--)|(?:\]\]|--)>\s*$/g,Mb={option:[1,"<select multiple='multiple'>","</select>"],thead:[1,"<table>","</table>"],col:[2,"<table><colgroup>","</colgroup></table>"],tr:[2,"<table><tbody>","</tbody></table>"],td:[3,"<table><tbody><tr>","</tr></tbody></table>"],_default:[0,"",""]};Mb.optgroup=Mb.option,Mb.tbody=Mb.tfoot=Mb.colgroup=Mb.caption=Mb.thead,Mb.th=Mb.td,_.extend({clone:function(a,b,c){var d,e,f,g,h=a.cloneNode(!0),i=_.contains(a.ownerDocument,a);if(!(Y.noCloneChecked||1!==a.nodeType&&11!==a.nodeType||_.isXMLDoc(a)))for(g=r(h),f=r(a),d=0,e=f.length;e>d;d++)s(f[d],g[d]);if(b)if(c)for(f=f||r(a),g=g||r(h),d=0,e=f.length;e>d;d++)q(f[d],g[d]);else q(a,h);return g=r(h,"script"),g.length>0&&p(g,!i&&r(a,"script")),h},buildFragment:function(a,b,c,d){for(var e,f,g,h,i,j,k=b.createDocumentFragment(),l=[],m=0,n=a.length;n>m;m++)if(e=a[m],e||0===e)if("object"===_.type(e))_.merge(l,e.nodeType?[e]:e);else if(Gb.test(e)){for(f=f||k.appendChild(b.createElement("div")),g=(Fb.exec(e)||["",""])[1].toLowerCase(),h=Mb[g]||Mb._default,f.innerHTML=h[1]+e.replace(Eb,"<$1></$2>")+h[2],j=h[0];j--;)f=f.lastChild;_.merge(l,f.childNodes),f=k.firstChild,f.textContent=""}else l.push(b.createTextNode(e));for(k.textContent="",m=0;e=l[m++];)if((!d||-1===_.inArray(e,d))&&(i=_.contains(e.ownerDocument,e),f=r(k.appendChild(e),"script"),i&&p(f),c))for(j=0;e=f[j++];)Jb.test(e.type||"")&&c.push(e);return k},cleanData:function(a){for(var b,c,d,e,f=_.event.special,g=0;void 0!==(c=a[g]);g++){if(_.acceptData(c)&&(e=c[rb.expando],e&&(b=rb.cache[e]))){if(b.events)for(d in b.events)f[d]?_.event.remove(c,d):_.removeEvent(c,d,b.handle);rb.cache[e]&&delete rb.cache[e]}delete sb.cache[c[sb.expando]]}}}),_.fn.extend({text:function(a){return qb(this,function(a){return void 0===a?_.text(this):this.empty().each(function(){(1===this.nodeType||11===this.nodeType||9===this.nodeType)&&(this.textContent=a)})},null,a,arguments.length)},append:function(){return this.domManip(arguments,function(a){if(1===this.nodeType||11===this.nodeType||9===this.nodeType){var b=m(this,a);b.appendChild(a)}})},prepend:function(){return this.domManip(arguments,function(a){if(1===this.nodeType||11===this.nodeType||9===this.nodeType){var b=m(this,a);b.insertBefore(a,b.firstChild)}})},before:function(){return this.domManip(arguments,function(a){this.parentNode&&this.parentNode.insertBefore(a,this)})},after:function(){return this.domManip(arguments,function(a){this.parentNode&&this.parentNode.insertBefore(a,this.nextSibling)})},remove:function(a,b){for(var c,d=a?_.filter(a,this):this,e=0;null!=(c=d[e]);e++)b||1!==c.nodeType||_.cleanData(r(c)),c.parentNode&&(b&&_.contains(c.ownerDocument,c)&&p(r(c,"script")),c.parentNode.removeChild(c));return this},empty:function(){for(var a,b=0;null!=(a=this[b]);b++)1===a.nodeType&&(_.cleanData(r(a,!1)),a.textContent="");return this},clone:function(a,b){return a=null==a?!1:a,b=null==b?a:b,this.map(function(){return _.clone(this,a,b)})},html:function(a){return qb(this,function(a){var b=this[0]||{},c=0,d=this.length;if(void 0===a&&1===b.nodeType)return b.innerHTML;if("string"==typeof a&&!Hb.test(a)&&!Mb[(Fb.exec(a)||["",""])[1].toLowerCase()]){a=a.replace(Eb,"<$1></$2>");try{for(;d>c;c++)b=this[c]||{},1===b.nodeType&&(_.cleanData(r(b,!1)),b.innerHTML=a);b=0}catch(e){}}b&&this.empty().append(a)},null,a,arguments.length)},replaceWith:function(){var a=arguments[0];return this.domManip(arguments,function(b){a=this.parentNode,_.cleanData(r(this)),a&&a.replaceChild(b,this)}),a&&(a.length||a.nodeType)?this:this.remove()},detach:function(a){return this.remove(a,!0)},domManip:function(a,b){a=S.apply([],a);var c,d,e,f,g,h,i=0,j=this.length,k=this,l=j-1,m=a[0],p=_.isFunction(m);if(p||j>1&&"string"==typeof m&&!Y.checkClone&&Ib.test(m))return this.each(function(c){var d=k.eq(c);p&&(a[0]=m.call(this,c,d.html())),d.domManip(a,b)});if(j&&(c=_.buildFragment(a,this[0].ownerDocument,!1,this),d=c.firstChild,1===c.childNodes.length&&(c=d),d)){for(e=_.map(r(c,"script"),n),f=e.length;j>i;i++)g=c,i!==l&&(g=_.clone(g,!0,!0),f&&_.merge(e,r(g,"script"))),b.call(this[i],g,i);if(f)for(h=e[e.length-1].ownerDocument,_.map(e,o),i=0;f>i;i++)g=e[i],Jb.test(g.type||"")&&!rb.access(g,"globalEval")&&_.contains(h,g)&&(g.src?_._evalUrl&&_._evalUrl(g.src):_.globalEval(g.textContent.replace(Lb,"")))}return this}}),_.each({appendTo:"append",prependTo:"prepend",insertBefore:"before",insertAfter:"after",replaceAll:"replaceWith"},function(a,b){_.fn[a]=function(a){for(var c,d=[],e=_(a),f=e.length-1,g=0;f>=g;g++)c=g===f?this:this.clone(!0),_(e[g])[b](c),T.apply(d,c.get());return this.pushStack(d)}});var Nb,Ob={},Pb=/^margin/,Qb=new RegExp("^("+vb+")(?!px)[a-z%]+$","i"),Rb=function(a){return a.ownerDocument.defaultView.getComputedStyle(a,null)};!function(){function b(){g.style.cssText="-webkit-box-sizing:border-box;-moz-box-sizing:border-box;box-sizing:border-box;display:block;margin-top:1%;top:1%;border:1px;padding:1px;width:4px;position:absolute",g.innerHTML="",e.appendChild(f);var b=a.getComputedStyle(g,null);c="1%"!==b.top,d="4px"===b.width,e.removeChild(f)}var c,d,e=Z.documentElement,f=Z.createElement("div"),g=Z.createElement("div");g.style&&(g.style.backgroundClip="content-box",g.cloneNode(!0).style.backgroundClip="",Y.clearCloneStyle="content-box"===g.style.backgroundClip,f.style.cssText="border:0;width:0;height:0;top:0;left:-9999px;margin-top:1px;position:absolute",f.appendChild(g),a.getComputedStyle&&_.extend(Y,{pixelPosition:function(){return b(),c},boxSizingReliable:function(){return null==d&&b(),d},reliableMarginRight:function(){var b,c=g.appendChild(Z.createElement("div"));return c.style.cssText=g.style.cssText="-webkit-box-sizing:content-box;-moz-box-sizing:content-box;box-sizing:content-box;display:block;margin:0;border:0;padding:0",c.style.marginRight=c.style.width="0",g.style.width="1px",e.appendChild(f),b=!parseFloat(a.getComputedStyle(c,null).marginRight),e.removeChild(f),b}}))}(),_.swap=function(a,b,c,d){var e,f,g={};for(f in b)g[f]=a.style[f],a.style[f]=b[f];e=c.apply(a,d||[]);for(f in b)a.style[f]=g[f];return e};var Sb=/^(none|table(?!-c[ea]).+)/,Tb=new RegExp("^("+vb+")(.*)$","i"),Ub=new RegExp("^([+-])=("+vb+")","i"),Vb={position:"absolute",visibility:"hidden",display:"block"},Wb={letterSpacing:"0",fontWeight:"400"},Xb=["Webkit","O","Moz","ms"];_.extend({cssHooks:{opacity:{get:function(a,b){if(b){var c=v(a,"opacity");return""===c?"1":c}}}},cssNumber:{columnCount:!0,fillOpacity:!0,flexGrow:!0,flexShrink:!0,fontWeight:!0,lineHeight:!0,opacity:!0,order:!0,orphans:!0,widows:!0,zIndex:!0,zoom:!0},cssProps:{"float":"cssFloat"},style:function(a,b,c,d){if(a&&3!==a.nodeType&&8!==a.nodeType&&a.style){var e,f,g,h=_.camelCase(b),i=a.style;return b=_.cssProps[h]||(_.cssProps[h]=x(i,h)),g=_.cssHooks[b]||_.cssHooks[h],void 0===c?g&&"get"in g&&void 0!==(e=g.get(a,!1,d))?e:i[b]:(f=typeof c,"string"===f&&(e=Ub.exec(c))&&(c=(e[1]+1)*e[2]+parseFloat(_.css(a,b)),f="number"),null!=c&&c===c&&("number"!==f||_.cssNumber[h]||(c+="px"),Y.clearCloneStyle||""!==c||0!==b.indexOf("background")||(i[b]="inherit"),g&&"set"in g&&void 0===(c=g.set(a,c,d))||(i[b]=c)),void 0)}},css:function(a,b,c,d){var e,f,g,h=_.camelCase(b);return b=_.cssProps[h]||(_.cssProps[h]=x(a.style,h)),g=_.cssHooks[b]||_.cssHooks[h],g&&"get"in g&&(e=g.get(a,!0,c)),void 0===e&&(e=v(a,b,d)),"normal"===e&&b in Wb&&(e=Wb[b]),""===c||c?(f=parseFloat(e),c===!0||_.isNumeric(f)?f||0:e):e}}),_.each(["height","width"],function(a,b){_.cssHooks[b]={get:function(a,c,d){return c?Sb.test(_.css(a,"display"))&&0===a.offsetWidth?_.swap(a,Vb,function(){return A(a,b,d)}):A(a,b,d):void 0},set:function(a,c,d){var e=d&&Rb(a);return y(a,c,d?z(a,b,d,"border-box"===_.css(a,"boxSizing",!1,e),e):0)}}}),_.cssHooks.marginRight=w(Y.reliableMarginRight,function(a,b){return b?_.swap(a,{display:"inline-block"},v,[a,"marginRight"]):void 0}),_.each({margin:"",padding:"",border:"Width"},function(a,b){_.cssHooks[a+b]={expand:function(c){for(var d=0,e={},f="string"==typeof c?c.split(" "):[c];4>d;d++)e[a+wb[d]+b]=f[d]||f[d-2]||f[0];return e}},Pb.test(a)||(_.cssHooks[a+b].set=y)}),_.fn.extend({css:function(a,b){return qb(this,function(a,b,c){var d,e,f={},g=0;if(_.isArray(b)){for(d=Rb(a),e=b.length;e>g;g++)f[b[g]]=_.css(a,b[g],!1,d);return f}return void 0!==c?_.style(a,b,c):_.css(a,b)},a,b,arguments.length>1)},show:function(){return B(this,!0)},hide:function(){return B(this)},toggle:function(a){return"boolean"==typeof a?a?this.show():this.hide():this.each(function(){xb(this)?_(this).show():_(this).hide()})}}),_.Tween=C,C.prototype={constructor:C,init:function(a,b,c,d,e,f){this.elem=a,this.prop=c,this.easing=e||"swing",this.options=b,this.start=this.now=this.cur(),this.end=d,this.unit=f||(_.cssNumber[c]?"":"px")},cur:function(){var a=C.propHooks[this.prop];return a&&a.get?a.get(this):C.propHooks._default.get(this)},run:function(a){var b,c=C.propHooks[this.prop];return this.pos=b=this.options.duration?_.easing[this.easing](a,this.options.duration*a,0,1,this.options.duration):a,this.now=(this.end-this.start)*b+this.start,this.options.step&&this.options.step.call(this.elem,this.now,this),c&&c.set?c.set(this):C.propHooks._default.set(this),this}},C.prototype.init.prototype=C.prototype,C.propHooks={_default:{get:function(a){var b;return null==a.elem[a.prop]||a.elem.style&&null!=a.elem.style[a.prop]?(b=_.css(a.elem,a.prop,""),b&&"auto"!==b?b:0):a.elem[a.prop]},set:function(a){_.fx.step[a.prop]?_.fx.step[a.prop](a):a.elem.style&&(null!=a.elem.style[_.cssProps[a.prop]]||_.cssHooks[a.prop])?_.style(a.elem,a.prop,a.now+a.unit):a.elem[a.prop]=a.now}}},C.propHooks.scrollTop=C.propHooks.scrollLeft={set:function(a){a.elem.nodeType&&a.elem.parentNode&&(a.elem[a.prop]=a.now)}},_.easing={linear:function(a){return a},swing:function(a){return.5-Math.cos(a*Math.PI)/2}},_.fx=C.prototype.init,_.fx.step={};var Yb,Zb,$b=/^(?:toggle|show|hide)$/,_b=new RegExp("^(?:([+-])=|)("+vb+")([a-z%]*)$","i"),ac=/queueHooks$/,bc=[G],cc={"*":[function(a,b){var c=this.createTween(a,b),d=c.cur(),e=_b.exec(b),f=e&&e[3]||(_.cssNumber[a]?"":"px"),g=(_.cssNumber[a]||"px"!==f&&+d)&&_b.exec(_.css(c.elem,a)),h=1,i=20;if(g&&g[3]!==f){f=f||g[3],e=e||[],g=+d||1;do h=h||".5",g/=h,_.style(c.elem,a,g+f);while(h!==(h=c.cur()/d)&&1!==h&&--i)}return e&&(g=c.start=+g||+d||0,c.unit=f,c.end=e[1]?g+(e[1]+1)*e[2]:+e[2]),c}]};_.Animation=_.extend(I,{tweener:function(a,b){_.isFunction(a)?(b=a,a=["*"]):a=a.split(" ");for(var c,d=0,e=a.length;e>d;d++)c=a[d],cc[c]=cc[c]||[],cc[c].unshift(b)},prefilter:function(a,b){b?bc.unshift(a):bc.push(a)}}),_.speed=function(a,b,c){var d=a&&"object"==typeof a?_.extend({},a):{complete:c||!c&&b||_.isFunction(a)&&a,duration:a,easing:c&&b||b&&!_.isFunction(b)&&b};return d.duration=_.fx.off?0:"number"==typeof d.duration?d.duration:d.duration in _.fx.speeds?_.fx.speeds[d.duration]:_.fx.speeds._default,(null==d.queue||d.queue===!0)&&(d.queue="fx"),d.old=d.complete,d.complete=function(){_.isFunction(d.old)&&d.old.call(this),d.queue&&_.dequeue(this,d.queue)},d},_.fn.extend({fadeTo:function(a,b,c,d){return this.filter(xb).css("opacity",0).show().end().animate({opacity:b},a,c,d)},animate:function(a,b,c,d){var e=_.isEmptyObject(a),f=_.speed(b,c,d),g=function(){var b=I(this,_.extend({},a),f);(e||rb.get(this,"finish"))&&b.stop(!0)};return g.finish=g,e||f.queue===!1?this.each(g):this.queue(f.queue,g)},stop:function(a,b,c){var d=function(a){var b=a.stop;delete a.stop,b(c)};return"string"!=typeof a&&(c=b,b=a,a=void 0),b&&a!==!1&&this.queue(a||"fx",[]),this.each(function(){var b=!0,e=null!=a&&a+"queueHooks",f=_.timers,g=rb.get(this);if(e)g[e]&&g[e].stop&&d(g[e]);else for(e in g)g[e]&&g[e].stop&&ac.test(e)&&d(g[e]);for(e=f.length;e--;)f[e].elem!==this||null!=a&&f[e].queue!==a||(f[e].anim.stop(c),b=!1,f.splice(e,1));(b||!c)&&_.dequeue(this,a)})},finish:function(a){return a!==!1&&(a=a||"fx"),this.each(function(){var b,c=rb.get(this),d=c[a+"queue"],e=c[a+"queueHooks"],f=_.timers,g=d?d.length:0;for(c.finish=!0,_.queue(this,a,[]),e&&e.stop&&e.stop.call(this,!0),b=f.length;b--;)f[b].elem===this&&f[b].queue===a&&(f[b].anim.stop(!0),f.splice(b,1));for(b=0;g>b;b++)d[b]&&d[b].finish&&d[b].finish.call(this);delete c.finish})}}),_.each(["toggle","show","hide"],function(a,b){var c=_.fn[b];_.fn[b]=function(a,d,e){return null==a||"boolean"==typeof a?c.apply(this,arguments):this.animate(E(b,!0),a,d,e)}}),_.each({slideDown:E("show"),slideUp:E("hide"),slideToggle:E("toggle"),fadeIn:{opacity:"show"},fadeOut:{opacity:"hide"},fadeToggle:{opacity:"toggle"}},function(a,b){_.fn[a]=function(a,c,d){return this.animate(b,a,c,d)}}),_.timers=[],_.fx.tick=function(){var a,b=0,c=_.timers;for(Yb=_.now();b<c.length;b++)a=c[b],a()||c[b]!==a||c.splice(b--,1);c.length||_.fx.stop(),Yb=void 0},_.fx.timer=function(a){_.timers.push(a),a()?_.fx.start():_.timers.pop()},_.fx.interval=13,_.fx.start=function(){Zb||(Zb=setInterval(_.fx.tick,_.fx.interval))},_.fx.stop=function(){clearInterval(Zb),Zb=null},_.fx.speeds={slow:600,fast:200,_default:400},_.fn.delay=function(a,b){return a=_.fx?_.fx.speeds[a]||a:a,b=b||"fx",this.queue(b,function(b,c){var d=setTimeout(b,a);c.stop=function(){clearTimeout(d)}})},function(){var a=Z.createElement("input"),b=Z.createElement("select"),c=b.appendChild(Z.createElement("option"));a.type="checkbox",Y.checkOn=""!==a.value,Y.optSelected=c.selected,b.disabled=!0,Y.optDisabled=!c.disabled,a=Z.createElement("input"),a.value="t",a.type="radio",Y.radioValue="t"===a.value}();var dc,ec,fc=_.expr.attrHandle;_.fn.extend({attr:function(a,b){return qb(this,_.attr,a,b,arguments.length>1)},removeAttr:function(a){return this.each(function(){_.removeAttr(this,a)})}}),_.extend({attr:function(a,b,c){var d,e,f=a.nodeType;if(a&&3!==f&&8!==f&&2!==f)return typeof a.getAttribute===zb?_.prop(a,b,c):(1===f&&_.isXMLDoc(a)||(b=b.toLowerCase(),d=_.attrHooks[b]||(_.expr.match.bool.test(b)?ec:dc)),void 0===c?d&&"get"in d&&null!==(e=d.get(a,b))?e:(e=_.find.attr(a,b),null==e?void 0:e):null!==c?d&&"set"in d&&void 0!==(e=d.set(a,c,b))?e:(a.setAttribute(b,c+""),c):void _.removeAttr(a,b))},removeAttr:function(a,b){var c,d,e=0,f=b&&b.match(nb);if(f&&1===a.nodeType)for(;c=f[e++];)d=_.propFix[c]||c,_.expr.match.bool.test(c)&&(a[d]=!1),a.removeAttribute(c)},attrHooks:{type:{set:function(a,b){if(!Y.radioValue&&"radio"===b&&_.nodeName(a,"input")){var c=a.value;return a.setAttribute("type",b),c&&(a.value=c),b}}}}}),ec={set:function(a,b,c){return b===!1?_.removeAttr(a,c):a.setAttribute(c,c),c}},_.each(_.expr.match.bool.source.match(/\w+/g),function(a,b){var c=fc[b]||_.find.attr;fc[b]=function(a,b,d){var e,f;return d||(f=fc[b],fc[b]=e,e=null!=c(a,b,d)?b.toLowerCase():null,fc[b]=f),e}});var gc=/^(?:input|select|textarea|button)$/i;_.fn.extend({prop:function(a,b){return qb(this,_.prop,a,b,arguments.length>1)},removeProp:function(a){return this.each(function(){delete this[_.propFix[a]||a]})}}),_.extend({propFix:{"for":"htmlFor","class":"className"},prop:function(a,b,c){var d,e,f,g=a.nodeType;if(a&&3!==g&&8!==g&&2!==g)return f=1!==g||!_.isXMLDoc(a),f&&(b=_.propFix[b]||b,e=_.propHooks[b]),void 0!==c?e&&"set"in e&&void 0!==(d=e.set(a,c,b))?d:a[b]=c:e&&"get"in e&&null!==(d=e.get(a,b))?d:a[b]},propHooks:{tabIndex:{get:function(a){return a.hasAttribute("tabindex")||gc.test(a.nodeName)||a.href?a.tabIndex:-1}}}}),Y.optSelected||(_.propHooks.selected={get:function(a){var b=a.parentNode;return b&&b.parentNode&&b.parentNode.selectedIndex,null}}),_.each(["tabIndex","readOnly","maxLength","cellSpacing","cellPadding","rowSpan","colSpan","useMap","frameBorder","contentEditable"],function(){_.propFix[this.toLowerCase()]=this});var hc=/[\t\r\n\f]/g;_.fn.extend({addClass:function(a){var b,c,d,e,f,g,h="string"==typeof a&&a,i=0,j=this.length;if(_.isFunction(a))return this.each(function(b){_(this).addClass(a.call(this,b,this.className))});if(h)for(b=(a||"").match(nb)||[];j>i;i++)if(c=this[i],d=1===c.nodeType&&(c.className?(" "+c.className+" ").replace(hc," "):" ")){for(f=0;e=b[f++];)d.indexOf(" "+e+" ")<0&&(d+=e+" ");g=_.trim(d),c.className!==g&&(c.className=g)}return this},removeClass:function(a){var b,c,d,e,f,g,h=0===arguments.length||"string"==typeof a&&a,i=0,j=this.length;if(_.isFunction(a))return this.each(function(b){_(this).removeClass(a.call(this,b,this.className))});if(h)for(b=(a||"").match(nb)||[];j>i;i++)if(c=this[i],d=1===c.nodeType&&(c.className?(" "+c.className+" ").replace(hc," "):"")){for(f=0;e=b[f++];)for(;d.indexOf(" "+e+" ")>=0;)d=d.replace(" "+e+" "," ");g=a?_.trim(d):"",c.className!==g&&(c.className=g)}return this},toggleClass:function(a,b){var c=typeof a;return"boolean"==typeof b&&"string"===c?b?this.addClass(a):this.removeClass(a):this.each(_.isFunction(a)?function(c){_(this).toggleClass(a.call(this,c,this.className,b),b)}:function(){if("string"===c)for(var b,d=0,e=_(this),f=a.match(nb)||[];b=f[d++];)e.hasClass(b)?e.removeClass(b):e.addClass(b);else(c===zb||"boolean"===c)&&(this.className&&rb.set(this,"__className__",this.className),this.className=this.className||a===!1?"":rb.get(this,"__className__")||"")})},hasClass:function(a){for(var b=" "+a+" ",c=0,d=this.length;d>c;c++)if(1===this[c].nodeType&&(" "+this[c].className+" ").replace(hc," ").indexOf(b)>=0)return!0;return!1}});var ic=/\r/g;_.fn.extend({val:function(a){var b,c,d,e=this[0];{if(arguments.length)return d=_.isFunction(a),this.each(function(c){var e;1===this.nodeType&&(e=d?a.call(this,c,_(this).val()):a,null==e?e="":"number"==typeof e?e+="":_.isArray(e)&&(e=_.map(e,function(a){return null==a?"":a+""})),b=_.valHooks[this.type]||_.valHooks[this.nodeName.toLowerCase()],b&&"set"in b&&void 0!==b.set(this,e,"value")||(this.value=e))});if(e)return b=_.valHooks[e.type]||_.valHooks[e.nodeName.toLowerCase()],b&&"get"in b&&void 0!==(c=b.get(e,"value"))?c:(c=e.value,"string"==typeof c?c.replace(ic,""):null==c?"":c)}}}),_.extend({valHooks:{option:{get:function(a){var b=_.find.attr(a,"value");return null!=b?b:_.trim(_.text(a))}},select:{get:function(a){for(var b,c,d=a.options,e=a.selectedIndex,f="select-one"===a.type||0>e,g=f?null:[],h=f?e+1:d.length,i=0>e?h:f?e:0;h>i;i++)if(c=d[i],!(!c.selected&&i!==e||(Y.optDisabled?c.disabled:null!==c.getAttribute("disabled"))||c.parentNode.disabled&&_.nodeName(c.parentNode,"optgroup"))){if(b=_(c).val(),f)return b;g.push(b)}return g},set:function(a,b){for(var c,d,e=a.options,f=_.makeArray(b),g=e.length;g--;)d=e[g],(d.selected=_.inArray(d.value,f)>=0)&&(c=!0);return c||(a.selectedIndex=-1),f}}}}),_.each(["radio","checkbox"],function(){_.valHooks[this]={set:function(a,b){return _.isArray(b)?a.checked=_.inArray(_(a).val(),b)>=0:void 0}},Y.checkOn||(_.valHooks[this].get=function(a){return null===a.getAttribute("value")?"on":a.value})}),_.each("blur focus focusin focusout load resize scroll unload click dblclick mousedown mouseup mousemove mouseover mouseout mouseenter mouseleave change select submit keydown keypress keyup error contextmenu".split(" "),function(a,b){_.fn[b]=function(a,c){return arguments.length>0?this.on(b,null,a,c):this.trigger(b)}}),_.fn.extend({hover:function(a,b){return this.mouseenter(a).mouseleave(b||a)},bind:function(a,b,c){return this.on(a,null,b,c)},unbind:function(a,b){return this.off(a,null,b)},delegate:function(a,b,c,d){return this.on(b,a,c,d)},undelegate:function(a,b,c){return 1===arguments.length?this.off(a,"**"):this.off(b,a||"**",c)}});var jc=_.now(),kc=/\?/;_.parseJSON=function(a){return JSON.parse(a+"")},_.parseXML=function(a){var b,c;if(!a||"string"!=typeof a)return null;try{c=new DOMParser,b=c.parseFromString(a,"text/xml")}catch(d){b=void 0}return(!b||b.getElementsByTagName("parsererror").length)&&_.error("Invalid XML: "+a),b};var lc,mc,nc=/#.*$/,oc=/([?&])_=[^&]*/,pc=/^(.*?):[ \t]*([^\r\n]*)$/gm,qc=/^(?:about|app|app-storage|.+-extension|file|res|widget):$/,rc=/^(?:GET|HEAD)$/,sc=/^\/\//,tc=/^([\w.+-]+:)(?:\/\/(?:[^\/?#]*@|)([^\/?#:]*)(?::(\d+)|)|)/,uc={},vc={},wc="*/".concat("*");try{mc=location.href}catch(xc){mc=Z.createElement("a"),mc.href="",mc=mc.href}lc=tc.exec(mc.toLowerCase())||[],_.extend({active:0,lastModified:{},etag:{},ajaxSettings:{url:mc,type:"GET",isLocal:qc.test(lc[1]),global:!0,processData:!0,async:!0,contentType:"application/x-www-form-urlencoded; charset=UTF-8",accepts:{"*":wc,text:"text/plain",html:"text/html",xml:"application/xml, text/xml",json:"application/json, text/javascript"},contents:{xml:/xml/,html:/html/,json:/json/},responseFields:{xml:"responseXML",text:"responseText",json:"responseJSON"},converters:{"* text":String,"text html":!0,"text json":_.parseJSON,"text xml":_.parseXML},flatOptions:{url:!0,context:!0}},ajaxSetup:function(a,b){return b?L(L(a,_.ajaxSettings),b):L(_.ajaxSettings,a)},ajaxPrefilter:J(uc),ajaxTransport:J(vc),ajax:function(a,b){function c(a,b,c,g){var i,k,r,s,u,w=b;2!==t&&(t=2,h&&clearTimeout(h),d=void 0,f=g||"",v.readyState=a>0?4:0,i=a>=200&&300>a||304===a,c&&(s=M(l,v,c)),s=N(l,s,v,i),i?(l.ifModified&&(u=v.getResponseHeader("Last-Modified"),u&&(_.lastModified[e]=u),u=v.getResponseHeader("etag"),u&&(_.etag[e]=u)),204===a||"HEAD"===l.type?w="nocontent":304===a?w="notmodified":(w=s.state,k=s.data,r=s.error,i=!r)):(r=w,(a||!w)&&(w="error",0>a&&(a=0))),v.status=a,v.statusText=(b||w)+"",i?o.resolveWith(m,[k,w,v]):o.rejectWith(m,[v,w,r]),v.statusCode(q),q=void 0,j&&n.trigger(i?"ajaxSuccess":"ajaxError",[v,l,i?k:r]),p.fireWith(m,[v,w]),j&&(n.trigger("ajaxComplete",[v,l]),--_.active||_.event.trigger("ajaxStop")))}"object"==typeof a&&(b=a,a=void 0),b=b||{};var d,e,f,g,h,i,j,k,l=_.ajaxSetup({},b),m=l.context||l,n=l.context&&(m.nodeType||m.jquery)?_(m):_.event,o=_.Deferred(),p=_.Callbacks("once memory"),q=l.statusCode||{},r={},s={},t=0,u="canceled",v={readyState:0,getResponseHeader:function(a){var b;if(2===t){if(!g)for(g={};b=pc.exec(f);)g[b[1].toLowerCase()]=b[2];b=g[a.toLowerCase()]}return null==b?null:b},getAllResponseHeaders:function(){return 2===t?f:null},setRequestHeader:function(a,b){var c=a.toLowerCase();return t||(a=s[c]=s[c]||a,r[a]=b),this},overrideMimeType:function(a){return t||(l.mimeType=a),this},statusCode:function(a){var b;if(a)if(2>t)for(b in a)q[b]=[q[b],a[b]];else v.always(a[v.status]);return this},abort:function(a){var b=a||u;return d&&d.abort(b),c(0,b),this}};if(o.promise(v).complete=p.add,v.success=v.done,v.error=v.fail,l.url=((a||l.url||mc)+"").replace(nc,"").replace(sc,lc[1]+"//"),l.type=b.method||b.type||l.method||l.type,l.dataTypes=_.trim(l.dataType||"*").toLowerCase().match(nb)||[""],null==l.crossDomain&&(i=tc.exec(l.url.toLowerCase()),l.crossDomain=!(!i||i[1]===lc[1]&&i[2]===lc[2]&&(i[3]||("http:"===i[1]?"80":"443"))===(lc[3]||("http:"===lc[1]?"80":"443")))),l.data&&l.processData&&"string"!=typeof l.data&&(l.data=_.param(l.data,l.traditional)),K(uc,l,b,v),2===t)return v;j=l.global,j&&0===_.active++&&_.event.trigger("ajaxStart"),l.type=l.type.toUpperCase(),l.hasContent=!rc.test(l.type),e=l.url,l.hasContent||(l.data&&(e=l.url+=(kc.test(e)?"&":"?")+l.data,delete l.data),l.cache===!1&&(l.url=oc.test(e)?e.replace(oc,"$1_="+jc++):e+(kc.test(e)?"&":"?")+"_="+jc++)),l.ifModified&&(_.lastModified[e]&&v.setRequestHeader("If-Modified-Since",_.lastModified[e]),_.etag[e]&&v.setRequestHeader("If-None-Match",_.etag[e])),(l.data&&l.hasContent&&l.contentType!==!1||b.contentType)&&v.setRequestHeader("Content-Type",l.contentType),v.setRequestHeader("Accept",l.dataTypes[0]&&l.accepts[l.dataTypes[0]]?l.accepts[l.dataTypes[0]]+("*"!==l.dataTypes[0]?", "+wc+"; q=0.01":""):l.accepts["*"]);for(k in l.headers)v.setRequestHeader(k,l.headers[k]);if(l.beforeSend&&(l.beforeSend.call(m,v,l)===!1||2===t))return v.abort();u="abort";for(k in{success:1,error:1,complete:1})v[k](l[k]);if(d=K(vc,l,b,v)){v.readyState=1,j&&n.trigger("ajaxSend",[v,l]),l.async&&l.timeout>0&&(h=setTimeout(function(){v.abort("timeout")},l.timeout));try{t=1,d.send(r,c)}catch(w){if(!(2>t))throw w;c(-1,w)}}else c(-1,"No Transport");return v},getJSON:function(a,b,c){return _.get(a,b,c,"json")},getScript:function(a,b){return _.get(a,void 0,b,"script")}}),_.each(["get","post"],function(a,b){_[b]=function(a,c,d,e){return _.isFunction(c)&&(e=e||d,d=c,c=void 0),_.ajax({url:a,type:b,dataType:e,data:c,success:d})}}),_.each(["ajaxStart","ajaxStop","ajaxComplete","ajaxError","ajaxSuccess","ajaxSend"],function(a,b){_.fn[b]=function(a){return this.on(b,a)}}),_._evalUrl=function(a){return _.ajax({url:a,type:"GET",dataType:"script",async:!1,global:!1,"throws":!0})},_.fn.extend({wrapAll:function(a){var b;return _.isFunction(a)?this.each(function(b){_(this).wrapAll(a.call(this,b))}):(this[0]&&(b=_(a,this[0].ownerDocument).eq(0).clone(!0),this[0].parentNode&&b.insertBefore(this[0]),b.map(function(){for(var a=this;a.firstElementChild;)a=a.firstElementChild;return a}).append(this)),this)},wrapInner:function(a){return this.each(_.isFunction(a)?function(b){_(this).wrapInner(a.call(this,b))}:function(){var b=_(this),c=b.contents();c.length?c.wrapAll(a):b.append(a)})},wrap:function(a){var b=_.isFunction(a);return this.each(function(c){_(this).wrapAll(b?a.call(this,c):a)})},unwrap:function(){return this.parent().each(function(){_.nodeName(this,"body")||_(this).replaceWith(this.childNodes)}).end()}}),_.expr.filters.hidden=function(a){return a.offsetWidth<=0&&a.offsetHeight<=0},_.expr.filters.visible=function(a){return!_.expr.filters.hidden(a)};var yc=/%20/g,zc=/\[\]$/,Ac=/\r?\n/g,Bc=/^(?:submit|button|image|reset|file)$/i,Cc=/^(?:input|select|textarea|keygen)/i;_.param=function(a,b){var c,d=[],e=function(a,b){b=_.isFunction(b)?b():null==b?"":b,d[d.length]=encodeURIComponent(a)+"="+encodeURIComponent(b)};if(void 0===b&&(b=_.ajaxSettings&&_.ajaxSettings.traditional),_.isArray(a)||a.jquery&&!_.isPlainObject(a))_.each(a,function(){e(this.name,this.value)
});else for(c in a)O(c,a[c],b,e);return d.join("&").replace(yc,"+")},_.fn.extend({serialize:function(){return _.param(this.serializeArray())},serializeArray:function(){return this.map(function(){var a=_.prop(this,"elements");return a?_.makeArray(a):this}).filter(function(){var a=this.type;return this.name&&!_(this).is(":disabled")&&Cc.test(this.nodeName)&&!Bc.test(a)&&(this.checked||!yb.test(a))}).map(function(a,b){var c=_(this).val();return null==c?null:_.isArray(c)?_.map(c,function(a){return{name:b.name,value:a.replace(Ac,"\r\n")}}):{name:b.name,value:c.replace(Ac,"\r\n")}}).get()}}),_.ajaxSettings.xhr=function(){try{return new XMLHttpRequest}catch(a){}};var Dc=0,Ec={},Fc={0:200,1223:204},Gc=_.ajaxSettings.xhr();a.ActiveXObject&&_(a).on("unload",function(){for(var a in Ec)Ec[a]()}),Y.cors=!!Gc&&"withCredentials"in Gc,Y.ajax=Gc=!!Gc,_.ajaxTransport(function(a){var b;return Y.cors||Gc&&!a.crossDomain?{send:function(c,d){var e,f=a.xhr(),g=++Dc;if(f.open(a.type,a.url,a.async,a.username,a.password),a.xhrFields)for(e in a.xhrFields)f[e]=a.xhrFields[e];a.mimeType&&f.overrideMimeType&&f.overrideMimeType(a.mimeType),a.crossDomain||c["X-Requested-With"]||(c["X-Requested-With"]="XMLHttpRequest");for(e in c)f.setRequestHeader(e,c[e]);b=function(a){return function(){b&&(delete Ec[g],b=f.onload=f.onerror=null,"abort"===a?f.abort():"error"===a?d(f.status,f.statusText):d(Fc[f.status]||f.status,f.statusText,"string"==typeof f.responseText?{text:f.responseText}:void 0,f.getAllResponseHeaders()))}},f.onload=b(),f.onerror=b("error"),b=Ec[g]=b("abort");try{f.send(a.hasContent&&a.data||null)}catch(h){if(b)throw h}},abort:function(){b&&b()}}:void 0}),_.ajaxSetup({accepts:{script:"text/javascript, application/javascript, application/ecmascript, application/x-ecmascript"},contents:{script:/(?:java|ecma)script/},converters:{"text script":function(a){return _.globalEval(a),a}}}),_.ajaxPrefilter("script",function(a){void 0===a.cache&&(a.cache=!1),a.crossDomain&&(a.type="GET")}),_.ajaxTransport("script",function(a){if(a.crossDomain){var b,c;return{send:function(d,e){b=_("<script>").prop({async:!0,charset:a.scriptCharset,src:a.url}).on("load error",c=function(a){b.remove(),c=null,a&&e("error"===a.type?404:200,a.type)}),Z.head.appendChild(b[0])},abort:function(){c&&c()}}}});var Hc=[],Ic=/(=)\?(?=&|$)|\?\?/;_.ajaxSetup({jsonp:"callback",jsonpCallback:function(){var a=Hc.pop()||_.expando+"_"+jc++;return this[a]=!0,a}}),_.ajaxPrefilter("json jsonp",function(b,c,d){var e,f,g,h=b.jsonp!==!1&&(Ic.test(b.url)?"url":"string"==typeof b.data&&!(b.contentType||"").indexOf("application/x-www-form-urlencoded")&&Ic.test(b.data)&&"data");return h||"jsonp"===b.dataTypes[0]?(e=b.jsonpCallback=_.isFunction(b.jsonpCallback)?b.jsonpCallback():b.jsonpCallback,h?b[h]=b[h].replace(Ic,"$1"+e):b.jsonp!==!1&&(b.url+=(kc.test(b.url)?"&":"?")+b.jsonp+"="+e),b.converters["script json"]=function(){return g||_.error(e+" was not called"),g[0]},b.dataTypes[0]="json",f=a[e],a[e]=function(){g=arguments},d.always(function(){a[e]=f,b[e]&&(b.jsonpCallback=c.jsonpCallback,Hc.push(e)),g&&_.isFunction(f)&&f(g[0]),g=f=void 0}),"script"):void 0}),_.parseHTML=function(a,b,c){if(!a||"string"!=typeof a)return null;"boolean"==typeof b&&(c=b,b=!1),b=b||Z;var d=gb.exec(a),e=!c&&[];return d?[b.createElement(d[1])]:(d=_.buildFragment([a],b,e),e&&e.length&&_(e).remove(),_.merge([],d.childNodes))};var Jc=_.fn.load;_.fn.load=function(a,b,c){if("string"!=typeof a&&Jc)return Jc.apply(this,arguments);var d,e,f,g=this,h=a.indexOf(" ");return h>=0&&(d=_.trim(a.slice(h)),a=a.slice(0,h)),_.isFunction(b)?(c=b,b=void 0):b&&"object"==typeof b&&(e="POST"),g.length>0&&_.ajax({url:a,type:e,dataType:"html",data:b}).done(function(a){f=arguments,g.html(d?_("<div>").append(_.parseHTML(a)).find(d):a)}).complete(c&&function(a,b){g.each(c,f||[a.responseText,b,a])}),this},_.expr.filters.animated=function(a){return _.grep(_.timers,function(b){return a===b.elem}).length};var Kc=a.document.documentElement;_.offset={setOffset:function(a,b,c){var d,e,f,g,h,i,j,k=_.css(a,"position"),l=_(a),m={};"static"===k&&(a.style.position="relative"),h=l.offset(),f=_.css(a,"top"),i=_.css(a,"left"),j=("absolute"===k||"fixed"===k)&&(f+i).indexOf("auto")>-1,j?(d=l.position(),g=d.top,e=d.left):(g=parseFloat(f)||0,e=parseFloat(i)||0),_.isFunction(b)&&(b=b.call(a,c,h)),null!=b.top&&(m.top=b.top-h.top+g),null!=b.left&&(m.left=b.left-h.left+e),"using"in b?b.using.call(a,m):l.css(m)}},_.fn.extend({offset:function(a){if(arguments.length)return void 0===a?this:this.each(function(b){_.offset.setOffset(this,a,b)});var b,c,d=this[0],e={top:0,left:0},f=d&&d.ownerDocument;if(f)return b=f.documentElement,_.contains(b,d)?(typeof d.getBoundingClientRect!==zb&&(e=d.getBoundingClientRect()),c=P(f),{top:e.top+c.pageYOffset-b.clientTop,left:e.left+c.pageXOffset-b.clientLeft}):e},position:function(){if(this[0]){var a,b,c=this[0],d={top:0,left:0};return"fixed"===_.css(c,"position")?b=c.getBoundingClientRect():(a=this.offsetParent(),b=this.offset(),_.nodeName(a[0],"html")||(d=a.offset()),d.top+=_.css(a[0],"borderTopWidth",!0),d.left+=_.css(a[0],"borderLeftWidth",!0)),{top:b.top-d.top-_.css(c,"marginTop",!0),left:b.left-d.left-_.css(c,"marginLeft",!0)}}},offsetParent:function(){return this.map(function(){for(var a=this.offsetParent||Kc;a&&!_.nodeName(a,"html")&&"static"===_.css(a,"position");)a=a.offsetParent;return a||Kc})}}),_.each({scrollLeft:"pageXOffset",scrollTop:"pageYOffset"},function(b,c){var d="pageYOffset"===c;_.fn[b]=function(e){return qb(this,function(b,e,f){var g=P(b);return void 0===f?g?g[c]:b[e]:void(g?g.scrollTo(d?a.pageXOffset:f,d?f:a.pageYOffset):b[e]=f)},b,e,arguments.length,null)}}),_.each(["top","left"],function(a,b){_.cssHooks[b]=w(Y.pixelPosition,function(a,c){return c?(c=v(a,b),Qb.test(c)?_(a).position()[b]+"px":c):void 0})}),_.each({Height:"height",Width:"width"},function(a,b){_.each({padding:"inner"+a,content:b,"":"outer"+a},function(c,d){_.fn[d]=function(d,e){var f=arguments.length&&(c||"boolean"!=typeof d),g=c||(d===!0||e===!0?"margin":"border");return qb(this,function(b,c,d){var e;return _.isWindow(b)?b.document.documentElement["client"+a]:9===b.nodeType?(e=b.documentElement,Math.max(b.body["scroll"+a],e["scroll"+a],b.body["offset"+a],e["offset"+a],e["client"+a])):void 0===d?_.css(b,c,g):_.style(b,c,d,g)},b,f?d:void 0,f,null)}})}),_.fn.size=function(){return this.length},_.fn.andSelf=_.fn.addBack,"function"==typeof define&&define.amd&&define("jquery",[],function(){return _});var Lc=a.jQuery,Mc=a.$;return _.noConflict=function(b){return a.$===_&&(a.$=Mc),b&&a.jQuery===_&&(a.jQuery=Lc),_},typeof b===zb&&(a.jQuery=a.$=_),_});
```
