## Supplementary material for "Comparative pathway enrichment analysis in gastrointestinal cell lines Caco-2, HT-29, HEPG2, and colon fibroblasts using a custom expression panel for tight-junction and cytoskeletal regulatory genes": Sup. Results 1: modernizr.js

```
/*!
 * Modernizr v2.8.2
 * www.modernizr.com
 *
 * Copyright (c) Faruk Ates, Paul Irish, Alex Sexton
 * Available under the BSD and MIT licenses: www.modernizr.com/license/
 */
window.Modernizr=function(a,b,c){function d(a){t.cssText=a}function e(a,b){return d(x.join(a+";")+(b||""))}function f(a,b){return typeof a===b}function g(a,b){return!!~(""+a).indexOf(b)}function h(a,b){for(var d in a){var e=a[d];if(!g(e,"-")&&t[e]!==c)return"pfx"==b?e:!0}return!1}function i(a,b,d){for(var e in a){var g=b[a[e]];if(g!==c)return d===!1?a[e]:f(g,"function")?g.bind(d||b):g}return!1}function j(a,b,c){var d=a.charAt(0).toUpperCase()+a.slice(1),e=(a+" "+z.join(d+" ")+d).split(" ");return f(b,"string")||f(b,"undefined")?h(e,b):(e=(a+" "+A.join(d+" ")+d).split(" "),i(e,b,c))}function k(){o.input=function(c){for(var d=0,e=c.length;e>d;d++)E[c[d]]=!!(c[d]in u);return E.list&&(E.list=!(!b.createElement("datalist")||!a.HTMLDataListElement)),E}("autocomplete autofocus list placeholder max min multiple pattern required step".split(" ")),o.inputtypes=function(a){for(var d,e,f,g=0,h=a.length;h>g;g++)u.setAttribute("type",e=a[g]),d="text"!==u.type,d&&(u.value=v,u.style.cssText="position:absolute;visibility:hidden;",/^range$/.test(e)&&u.style.WebkitAppearance!==c?(q.appendChild(u),f=b.defaultView,d=f.getComputedStyle&&"textfield"!==f.getComputedStyle(u,null).WebkitAppearance&&0!==u.offsetHeight,q.removeChild(u)):/^(search|tel)$/.test(e)||(d=/^(url|email)$/.test(e)?u.checkValidity&&u.checkValidity()===!1:u.value!=v)),D[a[g]]=!!d;return D}("search tel url email datetime date month week time datetime-local number range color".split(" "))}var l,m,n="2.8.2",o={},p=!0,q=b.documentElement,r="modernizr",s=b.createElement(r),t=s.style,u=b.createElement("input"),v=":)",w={}.toString,x=" -webkit- -moz- -o- -ms- ".split(" "),y="Webkit Moz O ms",z=y.split(" "),A=y.toLowerCase().split(" "),B={svg:"http://www.w3.org/2000/svg"},C={},D={},E={},F=[],G=F.slice,H=function(a,c,d,e){var f,g,h,i,j=b.createElement("div"),k=b.body,l=k||b.createElement("body");if(parseInt(d,10))for(;d--;)h=b.createElement("div"),h.id=e?e[d]:r+(d+1),j.appendChild(h);return f=["&#173;",'<style id="s',r,'">',a,"</style>"].join(""),j.id=r,(k?j:l).innerHTML+=f,l.appendChild(j),k||(l.style.background="",l.style.overflow="hidden",i=q.style.overflow,q.style.overflow="hidden",q.appendChild(l)),g=c(j,a),k?j.parentNode.removeChild(j):(l.parentNode.removeChild(l),q.style.overflow=i),!!g},I=function(b){var c=a.matchMedia||a.msMatchMedia;if(c)return c(b)&&c(b).matches||!1;var d;return H("@media "+b+" { #"+r+" { position: absolute; } }",function(b){d="absolute"==(a.getComputedStyle?getComputedStyle(b,null):b.currentStyle).position}),d},J=function(){function a(a,e){e=e||b.createElement(d[a]||"div"),a="on"+a;var g=a in e;return g||(e.setAttribute||(e=b.createElement("div")),e.setAttribute&&e.removeAttribute&&(e.setAttribute(a,""),g=f(e[a],"function"),f(e[a],"undefined")||(e[a]=c),e.removeAttribute(a))),e=null,g}var d={select:"input",change:"input",submit:"form",reset:"form",error:"img",load:"img",abort:"img"};return a}(),K={}.hasOwnProperty;m=f(K,"undefined")||f(K.call,"undefined")?function(a,b){return b in a&&f(a.constructor.prototype[b],"undefined")}:function(a,b){return K.call(a,b)},Function.prototype.bind||(Function.prototype.bind=function(a){var b=this;if("function"!=typeof b)throw new TypeError;var c=G.call(arguments,1),d=function(){if(this instanceof d){var e=function(){};e.prototype=b.prototype;var f=new e,g=b.apply(f,c.concat(G.call(arguments)));return Object(g)===g?g:f}return b.apply(a,c.concat(G.call(arguments)))};return d}),C.flexbox=function(){return j("flexWrap")},C.flexboxlegacy=function(){return j("boxDirection")},C.canvas=function(){var a=b.createElement("canvas");return!(!a.getContext||!a.getContext("2d"))},C.canvastext=function(){return!(!o.canvas||!f(b.createElement("canvas").getContext("2d").fillText,"function"))},C.webgl=function(){return!!a.WebGLRenderingContext},C.touch=function(){var c;return"ontouchstart"in a||a.DocumentTouch&&b instanceof DocumentTouch?c=!0:H(["@media (",x.join("touch-enabled),("),r,")","{#modernizr{top:9px;position:absolute}}"].join(""),function(a){c=9===a.offsetTop}),c},C.geolocation=function(){return"geolocation"in navigator},C.postmessage=function(){return!!a.postMessage},C.websqldatabase=function(){return!!a.openDatabase},C.indexedDB=function(){return!!j("indexedDB",a)},C.hashchange=function(){return J("hashchange",a)&&(b.documentMode===c||b.documentMode>7)},C.history=function(){return!(!a.history||!history.pushState)},C.draganddrop=function(){var a=b.createElement("div");return"draggable"in a||"ondragstart"in a&&"ondrop"in a},C.websockets=function(){return"WebSocket"in a||"MozWebSocket"in a},C.rgba=function(){return d("background-color:rgba(150,255,150,.5)"),g(t.backgroundColor,"rgba")},C.hsla=function(){return d("background-color:hsla(120,40%,100%,.5)"),g(t.backgroundColor,"rgba")||g(t.backgroundColor,"hsla")},C.multiplebgs=function(){return d("background:url(https://),url(https://),red url(https://)"),/(url\s*\(.*?){3}/.test(t.background)},C.backgroundsize=function(){return j("backgroundSize")},C.borderimage=function(){return j("borderImage")},C.borderradius=function(){return j("borderRadius")},C.boxshadow=function(){return j("boxShadow")},C.textshadow=function(){return""===b.createElement("div").style.textShadow},C.opacity=function(){return e("opacity:.55"),/^0.55$/.test(t.opacity)},C.cssanimations=function(){return j("animationName")},C.csscolumns=function(){return j("columnCount")},C.cssgradients=function(){var a="background-image:",b="gradient(linear,left top,right bottom,from(#9f9),to(white));",c="linear-gradient(left top,#9f9, white);";return d((a+"-webkit- ".split(" ").join(b+a)+x.join(c+a)).slice(0,-a.length)),g(t.backgroundImage,"gradient")},C.cssreflections=function(){return j("boxReflect")},C.csstransforms=function(){return!!j("transform")},C.csstransforms3d=function(){var a=!!j("perspective");return a&&"webkitPerspective"in q.style&&H("@media (transform-3d),(-webkit-transform-3d){#modernizr{left:9px;position:absolute;height:3px;}}",function(b){a=9===b.offsetLeft&&3===b.offsetHeight}),a},C.csstransitions=function(){return j("transition")},C.fontface=function(){var a;return H('@font-face {font-family:"font";src:url("https://")}',function(c,d){var e=b.getElementById("smodernizr"),f=e.sheet||e.styleSheet,g=f?f.cssRules&&f.cssRules[0]?f.cssRules[0].cssText:f.cssText||"":"";a=/src/i.test(g)&&0===g.indexOf(d.split(" ")[0])}),a},C.generatedcontent=function(){var a;return H(["#",r,"{font:0/0 a}#",r,':after{content:"',v,'";visibility:hidden;font:3px/1 a}'].join(""),function(b){a=b.offsetHeight>=3}),a},C.video=function(){var a=b.createElement("video"),c=!1;try{(c=!!a.canPlayType)&&(c=new Boolean(c),c.ogg=a.canPlayType('video/ogg; codecs="theora"').replace(/^no$/,""),c.h264=a.canPlayType('video/mp4; codecs="avc1.42E01E"').replace(/^no$/,""),c.webm=a.canPlayType('video/webm; codecs="vp8, vorbis"').replace(/^no$/,""))}catch(d){}return c},C.audio=function(){var a=b.createElement("audio"),c=!1;try{(c=!!a.canPlayType)&&(c=new Boolean(c),c.ogg=a.canPlayType('audio/ogg; codecs="vorbis"').replace(/^no$/,""),c.mp3=a.canPlayType("audio/mpeg;").replace(/^no$/,""),c.wav=a.canPlayType('audio/wav; codecs="1"').replace(/^no$/,""),c.m4a=(a.canPlayType("audio/x-m4a;")||a.canPlayType("audio/aac;")).replace(/^no$/,""))}catch(d){}return c},C.localstorage=function(){try{return localStorage.setItem(r,r),localStorage.removeItem(r),!0}catch(a){return!1}},C.sessionstorage=function(){try{return sessionStorage.setItem(r,r),sessionStorage.removeItem(r),!0}catch(a){return!1}},C.webworkers=function(){return!!a.Worker},C.applicationcache=function(){return!!a.applicationCache},C.svg=function(){return!!b.createElementNS&&!!b.createElementNS(B.svg,"svg").createSVGRect},C.inlinesvg=function(){var a=b.createElement("div");return a.innerHTML="<svg/>",(a.firstChild&&a.firstChild.namespaceURI)==B.svg},C.smil=function(){return!!b.createElementNS&&/SVGAnimate/.test(w.call(b.createElementNS(B.svg,"animate")))},C.svgclippaths=function(){return!!b.createElementNS&&/SVGClipPath/.test(w.call(b.createElementNS(B.svg,"clipPath")))};for(var L in C)m(C,L)&&(l=L.toLowerCase(),o[l]=C[L](),F.push((o[l]?"":"no-")+l));return o.input||k(),o.addTest=function(a,b){if("object"==typeof a)for(var d in a)m(a,d)&&o.addTest(d,a[d]);else{if(a=a.toLowerCase(),o[a]!==c)return o;b="function"==typeof b?b():b,"undefined"!=typeof p&&p&&(q.className+=" "+(b?"":"no-")+a),o[a]=b}return o},d(""),s=u=null,function(a,b){function c(a,b){var c=a.createElement("p"),d=a.getElementsByTagName("head")[0]||a.documentElement;return c.innerHTML="x<style>"+b+"</style>",d.insertBefore(c.lastChild,d.firstChild)}function d(){var a=s.elements;return"string"==typeof a?a.split(" "):a}function e(a){var b=r[a[p]];return b||(b={},q++,a[p]=q,r[q]=b),b}function f(a,c,d){if(c||(c=b),k)return c.createElement(a);d||(d=e(c));var f;return f=d.cache[a]?d.cache[a].cloneNode():o.test(a)?(d.cache[a]=d.createElem(a)).cloneNode():d.createElem(a),!f.canHaveChildren||n.test(a)||f.tagUrn?f:d.frag.appendChild(f)}function g(a,c){if(a||(a=b),k)return a.createDocumentFragment();c=c||e(a);for(var f=c.frag.cloneNode(),g=0,h=d(),i=h.length;i>g;g++)f.createElement(h[g]);return f}function h(a,b){b.cache||(b.cache={},b.createElem=a.createElement,b.createFrag=a.createDocumentFragment,b.frag=b.createFrag()),a.createElement=function(c){return s.shivMethods?f(c,a,b):b.createElem(c)},a.createDocumentFragment=Function("h,f","return function(){var n=f.cloneNode(),c=n.createElement;h.shivMethods&&("+d().join().replace(/[\w\-]+/g,function(a){return b.createElem(a),b.frag.createElement(a),'c("'+a+'")'})+");return n}")(s,b.frag)}function i(a){a||(a=b);var d=e(a);return!s.shivCSS||j||d.hasCSS||(d.hasCSS=!!c(a,"article,aside,dialog,figcaption,figure,footer,header,hgroup,main,nav,section{display:block}mark{background:#FF0;color:#000}template{display:none}")),k||h(a,d),a}var j,k,l="3.7.0",m=a.html5||{},n=/^<|^(?:button|map|select|textarea|object|iframe|option|optgroup)$/i,o=/^(?:a|b|code|div|fieldset|h1|h2|h3|h4|h5|h6|i|label|li|ol|p|q|span|strong|style|table|tbody|td|th|tr|ul)$/i,p="_html5shiv",q=0,r={};!function(){try{var a=b.createElement("a");a.innerHTML="<xyz></xyz>",j="hidden"in a,k=1==a.childNodes.length||function(){b.createElement("a");var a=b.createDocumentFragment();return"undefined"==typeof a.cloneNode||"undefined"==typeof a.createDocumentFragment||"undefined"==typeof a.createElement}()}catch(c){j=!0,k=!0}}();var s={elements:m.elements||"abbr article aside audio bdi canvas data datalist details dialog figcaption figure footer header hgroup main mark meter nav output progress section summary template time video",version:l,shivCSS:m.shivCSS!==!1,supportsUnknownElements:k,shivMethods:m.shivMethods!==!1,type:"default",shivDocument:i,createElement:f,createDocumentFragment:g};a.html5=s,i(b)}(this,b),o._version=n,o._prefixes=x,o._domPrefixes=A,o._cssomPrefixes=z,o.mq=I,o.hasEvent=J,o.testProp=function(a){return h([a])},o.testAllProps=j,o.testStyles=H,o.prefixed=function(a,b,c){return b?j(a,b,c):j(a,"pfx")},q.className=q.className.replace(/(^|\s)no-js(\s|$)/,"$1$2")+(p?" js "+F.join(" "):""),o}(this,this.document);
```
