## Supplementary material for "Comparative pathway enrichment analysis in gastrointestinal cell lines Caco-2, HT-29, HEPG2, and colon fibroblasts using a custom expression panel for tight-junction and cytoskeletal regulatory genes": Sup. Results 1: placeholder.js

```
/*! http://mths.be/placeholder v2.0.9 by @mathias */
!function(a){"function"==typeof define&&define.amd?define(["jquery"],a):a(jQuery)}(function(a){function b(b){var c={},d=/^jQuery\d+$/;return a.each(b.attributes,function(a,b){b.specified&&!d.test(b.name)&&(c[b.name]=b.value)}),c}function c(b,c){var d=this,f=a(d);if(d.value==f.attr("placeholder")&&f.hasClass("placeholder"))if(f.data("placeholder-password")){if(f=f.hide().nextAll('input[type="password"]:first').show().attr("id",f.removeAttr("id").data("placeholder-id")),b===!0)return f[0].value=c;f.focus()}else d.value="",f.removeClass("placeholder"),d==e()&&d.select()}function d(){var d,e=this,f=a(e),g=this.id;if(""===e.value){if("password"===e.type){if(!f.data("placeholder-textinput")){try{d=f.clone().attr({type:"text"})}catch(h){d=a("<input>").attr(a.extend(b(this),{type:"text"}))}d.removeAttr("name").data({"placeholder-password":f,"placeholder-id":g}).bind("focus.placeholder",c),f.data({"placeholder-textinput":d,"placeholder-id":g}).before(d)}f=f.removeAttr("id").hide().prevAll('input[type="text"]:first').attr("id",g).show()}f.addClass("placeholder"),f[0].value=f.attr("placeholder")}else f.removeClass("placeholder")}function e(){try{return document.activeElement}catch(a){}}var f,g,h="[object OperaMini]"==Object.prototype.toString.call(window.operamini),i="placeholder"in document.createElement("input")&&!h,j="placeholder"in document.createElement("textarea")&&!h,k=a.valHooks,l=a.propHooks;i&&j?(g=a.fn.placeholder=function(){return this},g.input=g.textarea=!0):(g=a.fn.placeholder=function(){var a=this;return a.filter((i?"textarea":":input")+"[placeholder]").not(".placeholder").bind({"focus.placeholder":c,"blur.placeholder":d}).data("placeholder-enabled",!0).trigger("blur.placeholder"),a},g.input=i,g.textarea=j,f={get:function(b){var c=a(b),d=c.data("placeholder-password");return d?d[0].value:c.data("placeholder-enabled")&&c.hasClass("placeholder")?"":b.value},set:function(b,f){var g=a(b),h=g.data("placeholder-password");return h?h[0].value=f:g.data("placeholder-enabled")?(""===f?(b.value=f,b!=e()&&d.call(b)):g.hasClass("placeholder")?c.call(b,!0,f)||(b.value=f):b.value=f,g):b.value=f}},i||(k.input=f,l.value=f),j||(k.textarea=f,l.value=f),a(function(){a(document).delegate("form","submit.placeholder",function(){var b=a(".placeholder",this).each(c);setTimeout(function(){b.each(d)},10)})}),a(window).bind("beforeunload.placeholder",function(){a(".placeholder").each(function(){this.value=""})}))});
```
