## Supplementary material for "Comparative pathway enrichment analysis in gastrointestinal cell lines Caco-2, HT-29, HEPG2, and colon fibroblasts using a custom expression panel for tight-junction and cytoskeletal regulatory genes": Sup. Results 1: report.js

```
	function HideContent(d) {
		document.getElementById(d).style.display = "none";
	}
	function ShowContent(d) {
	    var p = document.getElementsByClassName("panelContent");
		var i;
		for (i = 0; i < p.length; i++) {
			p[i].style.display = "none";
		}
		document.getElementById(d).style.display = "block";

	}
	function ReverseDisplay(d) {
		if(document.getElementById(d).style.display == "none") { document.getElementById(d).style.display = "block"; }
		else { document.getElementById(d).style.display = "none"; }
	}

	function dompath(element) {
	    var path = '';
	    for (; element && element.nodeType == 1; element = element.parentNode) {
	        var inner = $(element).children().length == 0 ? $(element).text() : '';
	        var eleSelector = element.tagName.toLowerCase() +
               ((inner.length > 0) ? ':contains(\'' + inner + '\')' : '');
	        path = ' ' + eleSelector + path;
	    }
	    return path;
	}

	function openImagePopup(popupName) {
	    $('#' + popupName).dialog({
	        modal: true,
	        autoOpen: true,
	        height: ($(window).height()),
	        width: ($(window).width()),
	        position: { my: "left top", at: "left top", of: window }
	    });
	};

	function openInteractiveHeatmapPopup(popupName) {
	    $("body").addClass("loading");

	    setTimeout(function () {
	        $('#' + popupName).dialog({
	            modal: true,
	            autoOpen: true,
	            height: ($(window).height()),
	            width: ($(window).width()),
	            position: { my: "left top", at: "left top", of: window }
	        });

            // Remove the SVG element so when heatmaps are redrawn, they are drawn correctly.
	        d3.select("svg").remove();
	            heatmap(
                    id = "ih-div-" + popupName,
                    datasetFile = this["heatmapData_" + popupName],
                    colAnnoFile = this["colAnnotation_" + popupName],
                    rowAnnoFile = this["rowAnnotation_" + popupName],
                    colClustOrder = null,
                    rowClustOrder = null,
                    height = 700,                                       // make dynamic
                    renderOnBrushEnd = true,				            // set to true for extremely large datasets / slow browsers
                    categorical = true,				                    // true = discrete colors for annotations false = continuous
                    categoricalScheme = "google", 	                    // "google", "rainbow" (ignored if categorical = false)
                    continuousScheme = "rainbow",	                    // "cubehelix", "rainbow" (ignored if categorical = true)
                    annoHeatScheme = "plasma" 	                        // "viridis", "inferno", "magma", "plasma", "warm", "cool"
                    );

	        $("body").removeClass("loading");
	    }, 50);
	};

	function openPopup(popupName) {
	    $('#' + popupName).dialog({
	        modal: true,
	        autoOpen: true,
            minWidth: 600
	    });
	};
```
