## Supplementary figures and images for "Comparative pathway enrichment analysis in gastrointestinal cell lines Caco-2, HT-29, HEPG2, and colon fibroblasts using a custom expression panel for tight-junction and cytoskeletal regulatory genes"

### color legend - Day.png

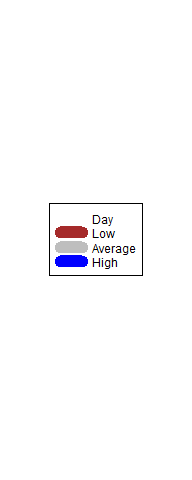

### color legend - Line.png

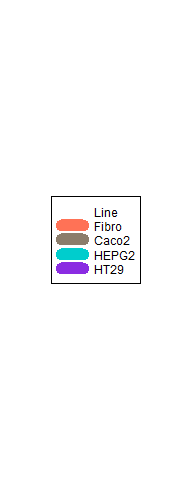

### color legend.png

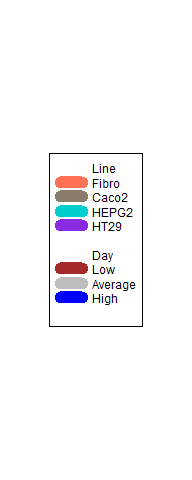

### logo_nanostring.png

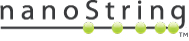

### logo_nanostring_Flat_189x40.png

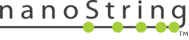

### logo_nanostring_white_Flat.png

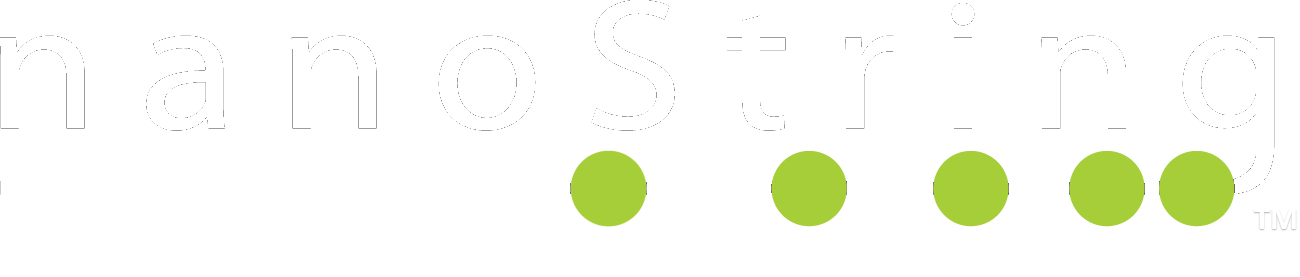

### logo_nanostring_white_Flat_189x40.png

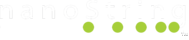

### nanostring_icon.png

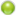

### ui-bg_flat_0_aaaaaa_40x100.png

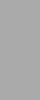

### ui-bg_flat_75_ffffff_40x100.png

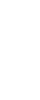

### ui-icons_2e83ff_256x240.png

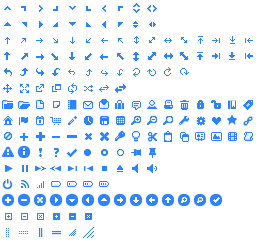

### ui-icons_222222_256x240.png

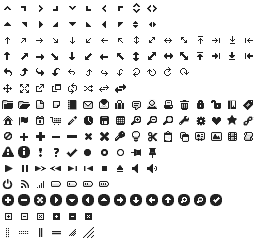

### ui-icons_454545_256x240.png

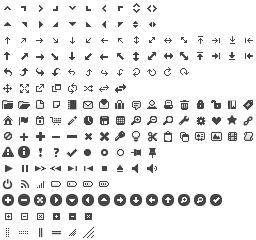

### ui-icons_888888_256x240.png

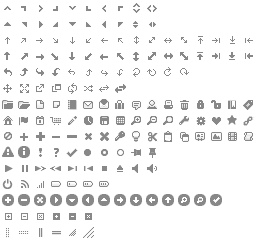

### ui-icons_cd0a0a_256x240.png

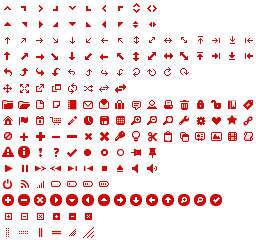

### volcano plotDay.png

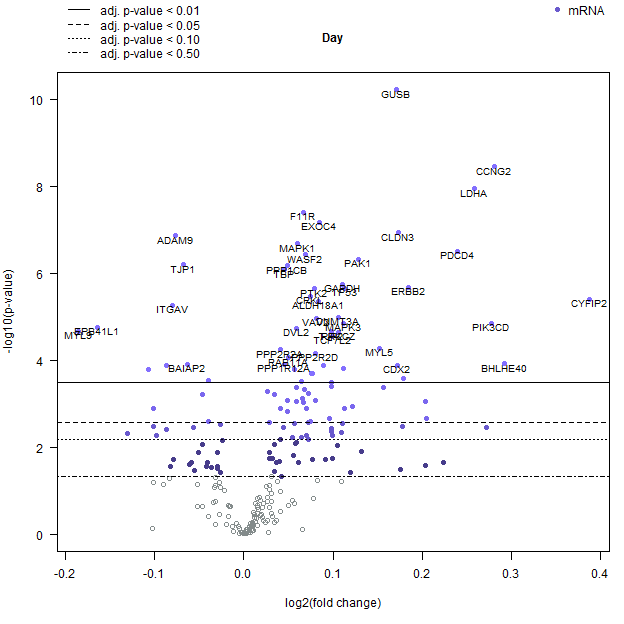

### volcano plotLineCaco2.png

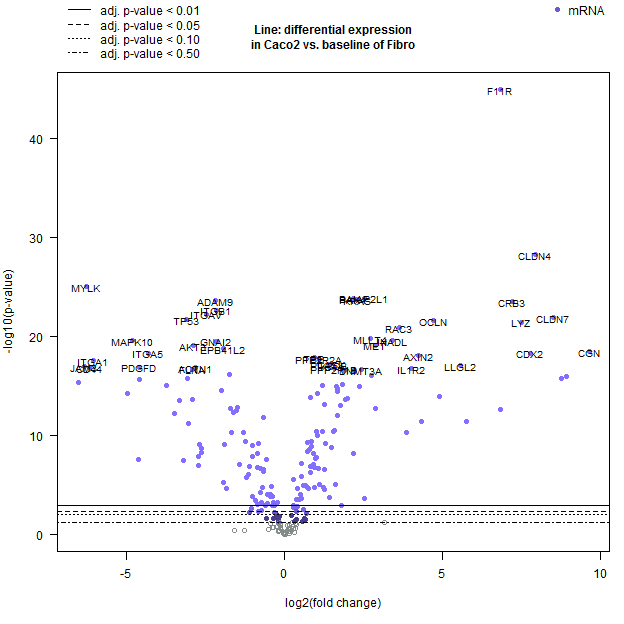

### volcano plotLineHEPG2.png

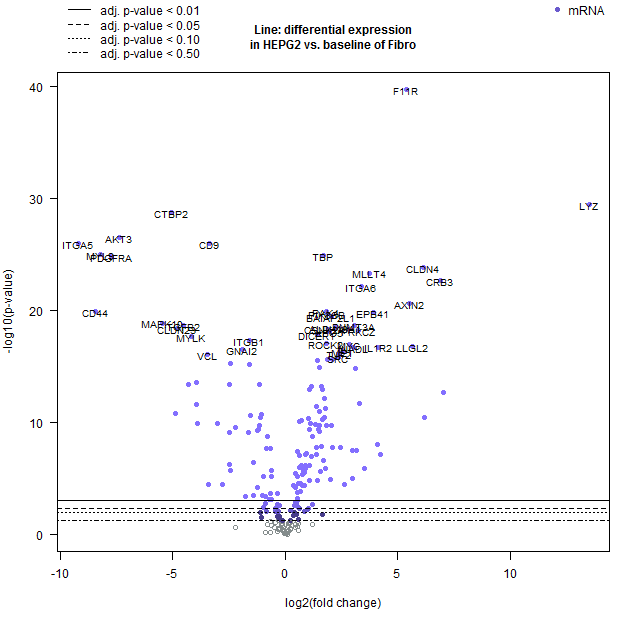

### volcano plotLineHT29.png

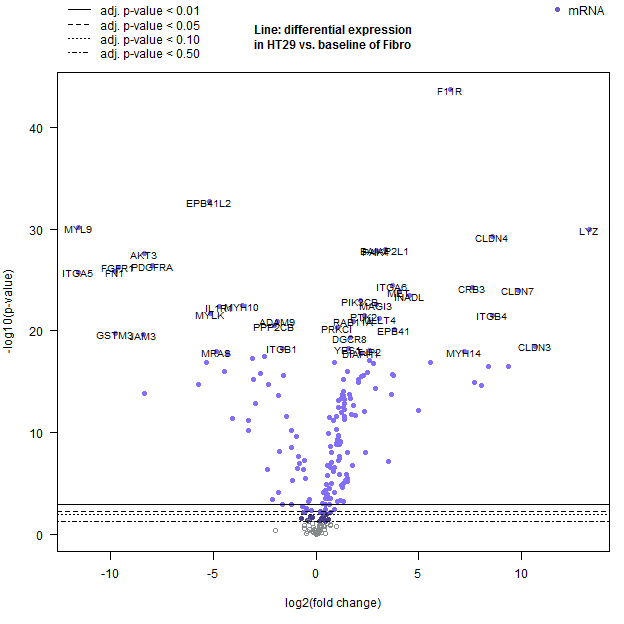
